# Characterizing and inferring quantitative cell cycle phase in single-cell RNA-seq data analysis

**DOI:** 10.1101/526848

**Authors:** Chiaowen Joyce Hsiao, PoYuan Tung, John D. Blischak, Jonathan E. Burnett, Kenneth A. Barr, Kushal K. Dey, Matthew Stephens, Yoav Gilad

## Abstract

Cellular heterogeneity in gene expression is driven by cellular processes such as cell cycle and cell-type identity, and cellular environment such as spatial location. The cell cycle, in particular, is thought to be a key driver of cell-to-cell heterogeneity in gene expression, even in otherwise homogeneous cell populations. Recent advances in single-cell RNA-sequencing (scRNA-seq) facilitate detailed characterization of gene expression heterogeneity, and can thus shed new light on the processes driving heterogeneity. Here, we combined fluorescence imaging with scRNA-seq to measure cell cycle phase and gene expression levels in human induced pluripotent stem cells (iPSCs). Using these data, we developed a novel approach to characterize cell cycle progression. While standard methods assign cells to discrete cell cycle stages, our method goes beyond this, and quantifies cell cycle progression on a continuum. We found that, on average, scRNA-seq data from only five genes predicted a cell’s position on the cell cycle continuum to within 14% of the entire cycle, and that using more genes did not improve this accuracy. Our data and predictor of cell cycle phase can directly help future studies to account for cell-cycle-related heterogeneity in iPSCs. Our results and methods also provide a foundation for future work to characterize the effects of the cell cycle on expression heterogeneity in other cell types.

## Introduction

Single-cell RNA-sequencing (scRNA-seq) can help characterize cellular heterogeneity in gene expression at unprecedented resolution (Kelsey et al. 2017; Macaulay et al. 2017; Tanay and Regev 2017; Papalexi and Satija 2018). By using scRNA-seq one can study not only the mean expression level of genes across an entire cell population, but also the variation in gene expression levels among cells (Kowalczyk et al. 2015; Lu et al. 2016; Stubbington et al. 2017; Velten et al. 2017; Nguyen et al. 2018; Skelly et al. 2018).

There are many reasons for differences in gene expression among cells, with arguably the most obvious candidates being differences in regulation among cell types, and differences in cell cycle phase among cells (Sanchez and Golding 2013; Keren et al. 2015; Soltani and Singh 2016). Cell type and cell cycle phase, while interesting to study directly, are often considered confounders in single cell studies that focus on other factors influencing gene expression (Buettner et al. 2015; Barron and Li 2016; Chen and Zhou 2017), such as genotype, treatment (Kolodziejczyk et al. 2015), or developmental time (Kowalczyk et al. 2015; Lauridsen et al. 2018). The ability to characterize, correctly classify, and correct for cell type and cell cycle phase are therefore important, even in studies that do not specifically aim to study either of these factors.

For these reasons, many studies have used single cell data to characterize the gene regulatory signatures of individual cells of different types and of cells at different cell cycle phases (e.g., Buettner et al. 2015; Leng et al. 2015; Povinelli et al. 2018). Often the ultimate goal of such studies is to be able to develop an effective approach to account for the variation associated with cell cycle or cell type. To characterize cell cycle phase, a common strategy in scRNA-seq studies is to first use flow cytometry to sort and pool cells that are in the same phase, followed by single-cell sequencing of the different pools (Buettner et al. 2015; Leng et al. 2015). In this common study design, cell cycle phase is completely confounded with the technical batch used to process single-cell RNA. This design flaw can inflate expression differences between the pools of cells in different cell cycle phase, resulting in inaccurate estimates of multi-gene signatures of cell cycle phase. When cells are not sorted before sequencing, cell cycle phase is typically accounted for by classifying the cells into discrete states based on the expression level of a few known markers (Butler et al. 2018).

Regardless of whether or not cells are sorted, all single-cell studies to date have accounted for cell cycle by using the standard classification of cell cycle phases, which is based on the notion that a cell passes through a consecutive series of distinct phases (G1, S, G2, M, and G0) marked by irreversible abrupt transitions. This standard definition of cell phases, however, is based on physiological observations and low-resolution data.

The traditional approach to classify and sort cells into distinct cell cycle states relies on a few known markers, and quite arbitrary gating cutoffs. Most cells of any given non-synchronized culture do not, in fact, show an unambiguous signature of being in one of the standard discrete cell cycle phases (Ingolia and Murray 2004; Pauklin and Vallier 2013; Kowalczyk et al. 2015). This makes intuitive sense: while from a physiological perspective, transitions between cell cycle states can be clearly defined (the DNA is either being replicated or not; the cell is either dividing or not), this is not the case when we try to define the cell states using molecular data. Indeed, we do not expect the gene expression signature of cell state transitions to occur in abrupt steps but rather to be a continuous process. High resolution single-cell data can provide a quantitative description of cell cycle progression and thus can allow us to move beyond the arbitrary classification of cells into discrete states.

From an analysis perspective, the ability to assign cells to a more precise point on the cell cycle continuum could capture fine-scale differences in the transcriptional profiles of single cells - differences that would be masked by grouping cells into discrete categories. Our goal here is therefore to study the relationship between cell cycle progression and gene expression at high resolution in single cells, without confounding cell cycle with batch effects as in (Buettner et al. 2015; Leng et al. 2015). To do so, we used fluorescent ubiquitination cell cycle indicators (FUCCI) (Sakaue-Sawano, Kurokawa, et al. 2008) to measure cell cycle progression, and scRNA-seq to measure gene expression in induced pluripotent stem cells (iPSCs) from six Yoruba individuals from Ibadan, Nigeria (abbreviation: YRI). To avoid the confounding of cell cycle with batch, we did not sort the cells by cell cycle phase before we collected the RNA-seq data. Instead, we measured FUCCI fluorescence intensities on intact single cells that were sorted into the C1 Fluidigm plate, prior to the preparation of the sequencing libraries. We also used a balanced incomplete block design to avoid confounding individual effects with batch effects. Using these data, we developed an analysis approach to characterize cell cycle progression on a continuous scale. We also developed a predictor of cell cycle progression in the iPSCs based on the scRNA-seq data. Our experimental and analytical strategies can help future scRNA-seq studies to explore the complex interplay between cell cycle progression, transcriptional heterogeneity, and other cellular phenotypes.

## Results

### Study design and data collection

We generated FUCCI-iPSCs using six YRI iPSC lines (see Methods for details) that we had characterized previously (Banovich et al. 2018). FUCCI-expressing iPSCs constitutively express two fluorescent reporter constructs transcribed from a shared promoter (Sakaue-Sawano, Kurokawa, et al. 2008; Sakaue-Sawano, Yo, et al. 2017). Reporters consist of either EGFP or mCherry fused to the degron domain of Geminin (geminin DNA replication inhibitor) or Cdt1 (Chromatin licensing and DNA replication factor 1). Due to their precisely-timed and specific regulation by the ubiquitin ligases APC/C and SCF, Geminin and Cdt1 are expressed in an inverse pattern throughout the cell cycle. Specifically, Geminin accumulates during S/G2/M and declines as the cell enters G1, whereas Cdt1 accumulates during G1 and declines after the onset of S phase. Thus, FUCCI reporters provide a way to assign cell cycle phase by tracking the degradation of Geminin-EGFP and Cdt1-mCherry through the enzymatic activity of their corresponding regulators, APC/C and SCF.

**Figure 1:**
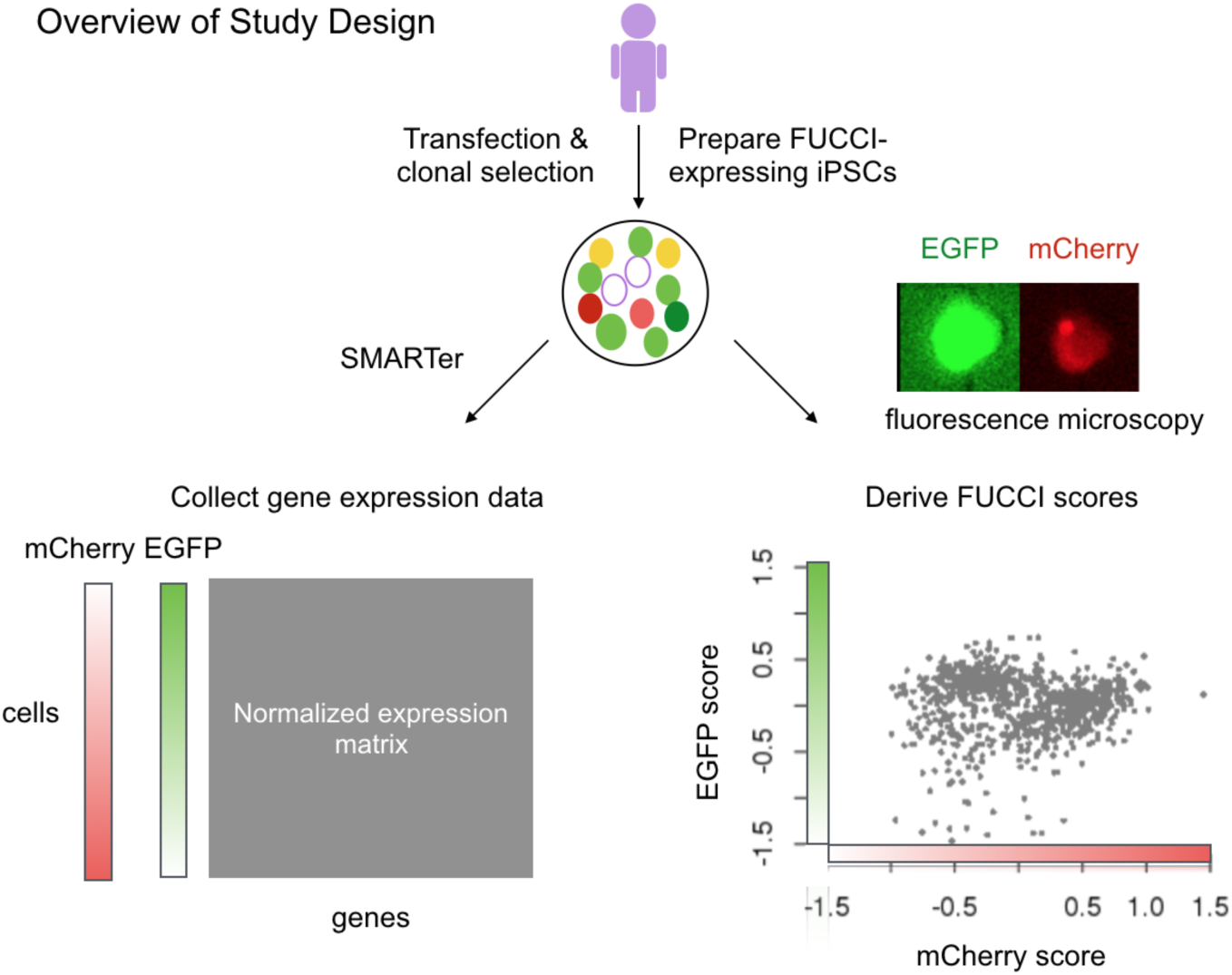
Overview of study design. We collected two types of data from the same single cells using FUCCI-expressing iPSCs: in situ fluorescence images and scRNA-seq. After quality control, we obtained 888 single cells for which we had high quality RNA-seq data. We computed two FUCCI scores for each cell by individually summing the EGFP (green) and mCherry (red) intensities in a fixed cell area (100 x 100 px), correcting for background noise outside the defined cell area, and then taking the log10 transformation of the sum of corrected intensities. In the bottom-right scatter plot, we show the FUCCI scores for the 888 high quality single-cell samples, i.e., mCherry and EGFP log10 sum intensities after background noise correction. Finally, we standardized the molecule counts to counts per million (CPM) and transformed the data per gene to a standard normal distribution.

We collected FUCCI fluorescence images (EGFP-Geminin and mCherry-Cdt1) and scRNA-seq data from the same single cells using an automated system designed for the Fluidigm C1 platform (Fig. 1; see Methods). After image capture, we prepared scRNA-seq libraries for sequencing using a SMARTer protocol adapted for iPSCs (Tung et al. 2017). To minimize bias caused by batch effects (Hicks et al. 2018; Tung et al. 2017), we used a balanced incomplete block design in which cells from unique pairs of iPSC lines were distributed across fifteen 96-well plates on the C1 platform (see Supplemental Fig. S1 for our C1 study design). We also included data from one additional plate (containing individuals NA18855 and NA18511), which we collected as part of a pilot study in which we optimized our protocols. In total, we collected data from 1,536 scRNA-seq samples distributed across 16 C1 plates.

### Single-cell RNA-sequencing

We applied quality metrics previously described in Tung et al. 2017 to determine criteria for including high-quality scRNA-seq samples (see Supplemental Fig. S2 and Methods for details). In addition, we used DAPI staining to help determine the number of single cells captured in each C1 well. This approach excludes any scRNA-seq samples containing cells undergoing mitosis, broken cells, or more than one cell. After quality control, we retained RNA-seq data from 888 single-cell samples, with a range of 103 to 206 cells from each of the six individuals (see Supplemental Fig. S3, Supplemental Fig. S4). We standardized the molecule counts to counts per million (CPM) and retained 11,040 genes with CPM 1 or higher in order to evaluate as many genes as possible. This resulted in a mean gene detection rate of 70 % across cells (standard deviation of 25 %, no significant difference between the six cell lines, see Supplemental Fig. S3). Finally, we quantile-normalized the expression levels across all single-cell samples to a standard normal distribution for each gene.

We used principal components analysis (PCA) to assess the global influence of technical factors on expression, including plate, individual, and read depth (see Supplemental Fig. S5). The primary source of sample variation in our data was the proportion of genes detected (>1 log_2_ CPM; adj. R-squared=0.39 for PC1; 0.25 for PC2), consistent with results from previous studies (Hicks et al. 2018). We found that the proportion of genes detected in our samples showed a stronger correlation with the number of reads mapped (adj. R-squared=0.32) than with plate (adj. R-squared=0.01) or individual (adj. R-squared=0.09). Thus, we confirmed that further statistical adjustment to account for batch effects will not yield noticeably different results. This demonstrates that our use of a balanced incomplete block design was an effective strategy to minimize the effects of confounding technical variables.

### Quantifying continuous cell cycle phase using FUCCI intensities

Proceeding with the 888 single cells for which we had high quality RNA-seq data, we turned our attention to the corresponding FUCCI data. For each cell, we defined a fixed cell area for all sample images (100 x 100 px) for the EGFP-Geminin and mCherry-Cdt1 images. This allowed us to account for differences in cell size. We computed two FUCCI scores for each cell to assign cell cycle phase. These scores sum up the EGFP/mCherry intensities in the fixed cell area after correcting for background noise outside the defined cell area (see Methods for more details).

Because images were captured one plate at a time, we scanned the data for evidence of batch effects. We found mean FUCCI scores to be significantly different between plates (F-test P-value < 2 × 10^−16^ for both EGFP and mCherry, see Supplemental Fig. S6 for comparisons between C1 plates, Supplemental Fig. S7 for comparisons between the six cell lines). We hence applied a linear model to account for plate effects on FUCCI scores without removing individual effects (FUCCI score ∼ plate + individual). Figure 1C shows the relationship between EGFP and mCherry scores after batch effect correction.

FUCCI intensities are commonly used to sort cells into discrete cell cycle phases. For example, cells expressing EGFP-Geminin in the absence of mCherry-Cdt1 would traditionally be assigned to G2/M, cells with the opposite pattern of expression would be assigned to G1, and cells expressing equal amounts of EGFP-Geminin and mCherry-Cdt1 would be assigned to the S/G2 transition (Sakaue-Sawano, Kurokawa, et al. 2008). As a representative of this approach, we applied Partition Around Medoids (PAM) from (Kaufman and Rousseeuw 1990) to FUCCI scores to assign single-cell samples to G1, S, and G2/M phase (G1 384 cells, S 172 cells, G2/M 332 cells, see Supplemental Fig S8). Henceforth, the classification obtained from PAM is referred to as PAM-based classification.

However, FUCCI intensities are known to be continuously distributed within each phase (Sakaue-Sawano, Kurokawa, et al. 2008), suggesting that they could also be used to quantify cell cycle progression through a continuum (conventionally represented using radians in the range [0, 2*π*]). With this in mind, we ordered the corrected FUCCI scores by phase and plotted them on a unit circle, using the co-oscillation of mCherry-Cdt1 and EGFP-Geminin to infer an angle, or ‘FUCCI phase’, for each cell (Fig. 2A; see Methods). For example, Fig. 2B shows that as a cell progresses through *π*/2 to *π* radians, mCherry-Ctd1 intensity decreases from its maximum, while EGFP-Geminin intensity changes from negative to positive, suggesting progression through G1/S transition. Overall, FUCCI phase explains 87% of variation in mCherry intensity and 70% of variation in EGFP intensity.

**Figure 2:**
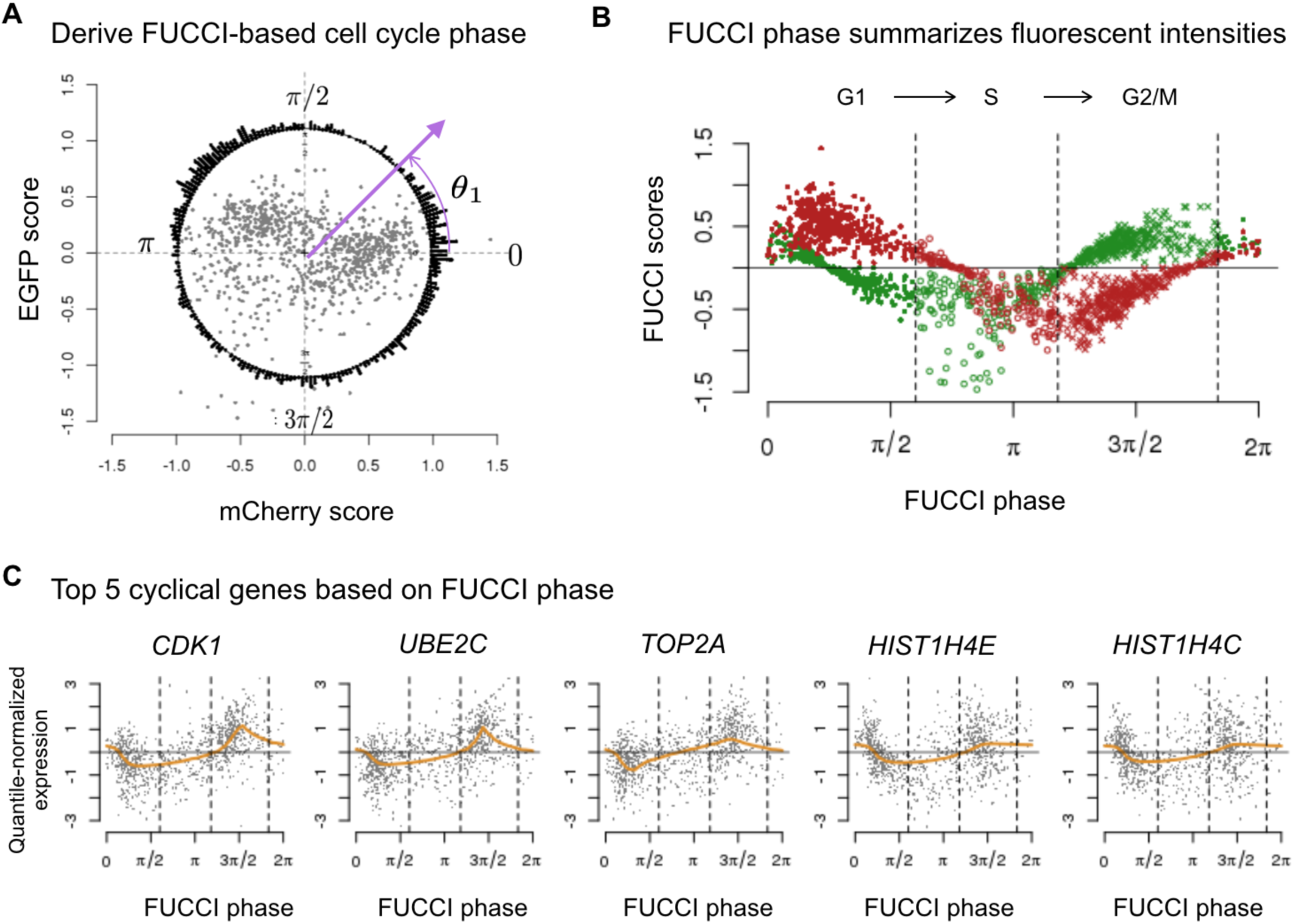
Characterizing cell cycle phase using FUCCI fluorescence intensities. (A) We inferred FUCCI phase (angles in a circle) based on FUCCI scores of EGFP and mCherry. The points in center correspond to PC scores based on EGFP and mCherry scores, and the circle histogram shows the corresponding FUCCI phase distribution. For example, we inferred *θ*_1_ based on the PC scores derived from the cell’s FUCCI scores. (B) We ordered FUCCI scores of EGFP and mCherry by FUCCI phase to visualize the co-oscillation of EGFP and mCherry along the cell cycle. Red and green points correspond to EGFP and mCherry scores, respectively. The vertical lines correspond to phase boundaries derived from the PAM-based classification (G1 384 cells, S 172 cells, G2/M 332 cells). (C) Given FUCCI phase, we ordered cells along the cell cycle to estimate the cyclic trend of gene expression levels for each gene. We identified these 5 genes as the top 5 cyclic genes in the data: *CDK1*, *UBE2C*, *TOP2A*, *H4C5*, and *H4C3*. Each plot shows the expression levels of 888 single-cell samples and the estimated cyclic trend (orange line). All 5 genes were previously identified as related to cell cycle regulation. The vertical lines correspond to phase boundaries derived from the PAM-based classification.

We next sought to identify genes whose expression levels vary in a cyclic way through the cell cycle, as captured by FUCCI phase. Specifically, we used a non-parametric smoothing method, trend filtering (Tibshirani 2014), to estimate the change in expression for each gene through the cell cycle. We refer to these estimates as the “cyclic trend” for each gene. We used a permutation-based test (see Methods) to assess the significance of each inferred cyclic trend, and ranked the genes by statistical significance. Results showed that genes with a significant cyclic trend were strongly enriched for known cell cycle genes. Using a curated set of 622 cell cycle genes in Macosko et al. 2015 (a subset of genes annotated in Whitfield et al. 2002, See Supplemental Table S1), we found Odds Ratio=25.79 for the 101 significant cyclic genes, Odds Ratio=31 for the top 5 significant cyclic genes, 30 for the top 50 genes, and 27 for the top 100 genes (Fisher’s exact test P-value < .001. See Supplemental Table S2 for the gene list and Supplemental Fig S9A,B, C for cyclic trends of known cell cycle genes). These results provide strong independent support that the inferred FUCCI phase is indeed meaningfully capturing cell cycle progression.

For illustration, Fig. 2C shows the cyclic trends for the top 5 significant cyclic genes: *CDK1*, *UBE2C*, *TOP2A*, *H4C5*, *H4C3*. These genes have all been previously identified as cell cycle genes in synchronization experiments of HeLa cells (Whitfield et al. 2002) and in scRNA-seq studies of FUCCI-sorted cells (Leng et al. 2015). *CDK1* (Cyclin Dependent Kinase 1, previously known as *CDC2*) promotes the transition to mitosis. *TOP2A* (DNA topoisomerase II-alpha) controls the topological state of DNA during cell state transitions. *UBE2C* (Ubiquitin Conjugating Enzyme E2 C) is required for the degradation of mitotic cyclins and the transition to G2 stage. Finally, *H4C3*, and *H4C5* (Histone gene cluster 1, H4 histone family) are replication-dependent histone genes expressed mainly during S phase.

### Predicting FUCCI phase from gene expression data

#### Our supervised approach

Building on these results, we developed a statistical method for predicting continuous cell cycle phase from gene expression data. The intuition behind our approach is that given a set of labeled training data – cells for which we have both FUCCI phase (*Y*) and scRNA-seq data (*X*) – our trend-filtering approach learns the cyclic trend for each gene (i.e., *p*(*X|Y*)). We combine this with a prior for the phase (*p*(*Y*)) using the idea of a “naive Bayes” predictor, to predict FUCCI phase from gene expression (i.e., *p*(*Y|X*)). Given scRNA-seq data, *X*, on any additional cell without FUCCI data, we can then apply this method to predict its FUCCI phase, *Y* (see Methods for more details). Henceforth, our continuous predictor is referred to as peco.

To assess the performance of our predictor, we applied six-fold cross-validation. In each fold, we trained our predictor on cells from five individuals and tested its performance on cells from the remaining individual. This allowed us to assess the ability of our predictor to generalize to individuals not seen in training. We measured the prediction error as the difference between the predicted phase and the measured FUCCI phase (as a percentage of the entire cycle, 2*π*). Note that since phases lie on a circle, the maximum possible error is 50% of the circle, and the expected error from random guessing would be 25% of the circle. Using our approach, on average, we were able to predict a cell’s position on the cell cycle continuum to within 14% of the entire cycle (i.e., .28*π* between inferred phase and FUCCI phase).

Supplemental Fig. S11A shows the performance of predictors built using between 2 and 50 genes. The genes were ranked and included in the predictors according to the significance of their cyclic trend. We observed that the mean prediction error was robust to the number of genes included in the predictor, and that the simplest predictor using only the top five genes (*CDK1*, *UBE2C*, *TOP2A*, *H4C5*, *H4C3*) performed just as well as the predictors with more genes.

We also checked the robustness of our predictors for data with lower effective sequencing depth compared to the C1 platform (e.g., Drop-seq and 10x Genomics). Specifically, we repeated the analysis above after thinning the test data (sample molecule count in the unthinned data was 56,724 ± 12,762) by a factor of 2.2 (sample molecule count 25,581 ± 15,220) and 4.4 (sample molecule count 13,651 ± 13,577). Results in Supplemental Fig. S10C,D show that the predictors based on fewer genes (e.g. 5-15) were relatively robust to this thinning; predictors based on more genes showed worse performance in the lower-count data. These results were somewhat expected, as we demonstrated in the unthinned data (Supplemental Fig. S10A) that adding genes with weak signals increased prediction error.

#### Comparisons with existing methods on our data

Several methods exist for making inferences on cell cycle from RNA-seq data. Here we consider two methods that attempt to infer a “cyclic ordering” of cells from RNA-seq data in an unsupervised way (Oscope by Leng et al. 2015, reCAT by Liu et al. 2017) and two methods that assign cells to discrete cell cycle states (Seurat by Butler et al. 2018, Cyclone by Scialdone et al. 2015). Coming to concrete conclusions that one analytic method is better than another is difficult in most settings, and is particularly difficult in settings where, as here, “gold standard” data are hard to come by. It is further complicated here by the fact that the methods differ in their precise goals (e.g. discrete vs continuous assignments, and supervised vs unsupervised assignments). Nonetheless, we compared the methods on both our data and on other data sets in an effort to provide some indication of their differences and commonalities.

First we ran the four other methods on our RNA-seq data from all 888 single cells, and compared their results with our FUCCI data on the same cells.

For the unsupervised methods Oscope and reCAT, we first applied each method to infer an ordering of cells from the RNA-seq data, and then assessed whether the inferred orderings produced cyclic patterns in the FUCCI scores (which one would expect if the inferred orderings accurately represented cell cycle). In both cases the ordering explained only very little variation in the FUCCI scores, with Oscope slightly higher than reCAT (Oscope: 13% EGFP, 16% mCherry; reCAT: 4% EGFP, 9% mCherry, see Supplemental Fig. S11B,D). In contrast, inferred phase from peco explained an average 29% of the variation in EGFP score and an average of 24% of the variation in mCherry score across six cell lines (see Supplemental Fig. S12).

For Seurat and Cyclone, we compared the discrete classifications (G1 vs S vs G2/M) they produced from RNA-seq data with the corresponding classifications obtained from FUCCI data using the PAM-based method from Kaufman and Rousseeuw 1990. Neither the Seurat nor Cyclone classifications agreed well with the FUCCI data (See Supplemental Fig. S14). Treating the FUCCI results as a gold standard, Seurat misclassification rates were 78% (G1), 74% (S), 43% (G2/M); and Cyclone misclassification rates were 34% (G1), 88% (S), 31% (G2/M). See Supplemental Fig. S13A,B).

To directly compare existing methods with our method requires translating results from existing methods into a continuous predictor of cell cycle phase that is comparable with our continuous predictor. For Oscope/reCAT, we did this by using their cyclic ordering to assign cells to equidistant points on the unit circle. For Seurat/Cyclone, we built a continuous predictor based on the Seurat/Cyclone phase-specific scores. Specifically, we applied the same approach used to derive FUCCI phase to transform the two Seurat scores and the three Cyclone scores to cell cycle angles (see Supplemental Fig. S14 for an example of Seurat score transformation).

This comparison will likely favor our predictor because existing methods were not optimized for continuous phase predictions (indeed, no method other than ours has been so optimized). Also our predictor was trained on the same cell types as are being used for assessment. With these caveats in mind, on these data our predictor outperformed predictors built from existing methods(Fig. 3B), with lower prediction error than all the other methods on all cell lines (and in most cases significantly lower at P-value < .05; see Supplemental Fig. S15, Supplemental Fig. S16). Overall, the mean prediction error of our predictor across the six cell lines was approximately 60% of the Seurat/Oscope/reCAT-based predictors and 80% of the Cyclone-based predictor.

Visual comparisons of the results from the different methods on the top 5 cyclic genes used by peco (see Fig. 3C, Supplemental Fig. S17A, B, C, D, E), suggest that on these data Oscope agrees most closely with peco than other methods; in particular, results from peco and Oscope show a clearer cyclic trend in the expression levels of *H4C5* and *H4C3* than do other methods.

**Figure 3:**
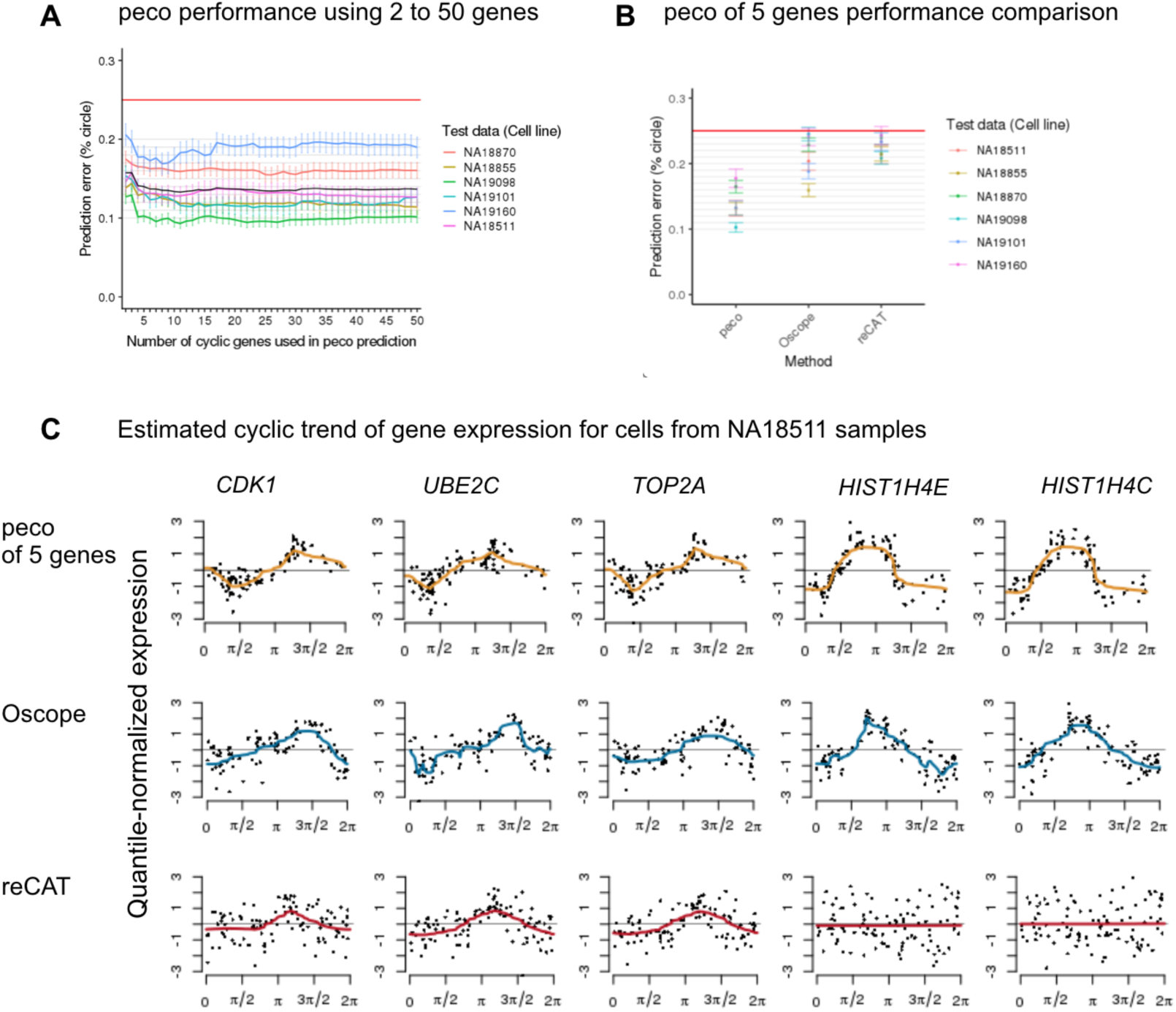
Inferring cell cycle phase from scRNA-seq data. (A) We applied six-fold cross-validation to test the performance of our predictor. In each fold, we trained our predictor on cells from five individuals and tested its performance on cells from the remaining individual. Y-axis corresponds to prediction error (between 0 to 25%, or *π*/4), and X-axis corresponds to the number of top cyclic genes used in the predictor. The six colored lines correspond to performances in the six folds, specifically average prediction error among cells in the test samples, and error bars correspond to standard errors. (B) Performance comparison of peco built from the top 5 cyclic genes (*CDK1*, *UBE2C*, *TOP2A*, *H4C5*, and *H4C3*) with Oscope (Leng et al. 2015) and recAT (Liu et al. 2017). (C) Estimated cyclic trend of top 5 cyclic genes for samples from cell line NA18511. The rows correspond to prediction results from peco of 5 genes, Oscope and reCAT. For the Oscope/reCAT results, we ordered the single-cell samples from NA18511 using the Oscope/reCAT-based predicted phase (based on 888 samples in the data) and used *trendfilter* to estimate cyclic trend of gene expression. For the peco results, We ordered the samples according to the predicted phase and used *trendfilter* to estimate cyclic trend of gene expression. The colored line corresponds to the estimated cyclic expression level along the predicted phase.

#### Comparisons on data from Leng et al

Leng et al. (2015) collected scRNA-seq and FUCCI data on Human embryonic cells (hESCs). The cells were transfected with the same FUCCI reporters used in our study (in fact, the co-author Dr. Chris Barry generously gifted us their plasmid). However, in contrast to our study, cells were first sorted into discrete cell cycle phases based on the FUCCI data (G1, S, and G2/M, henceforth referred to as “gating-based classification”), and then cells in each phase were prepared on different 96-well C1 plates prior to RNA-seq. In contrast to our data, this design means that plate effects are confounded with cell cycle phase, which is far from ideal. In addition, the sorted cells are not a random sample of all cells across all cell cycle states, but rather represent cells whose FUCCI data place them confidently into one of three discretely-defined cell cycle states. These issues were major motivations for our own data collection efforts. However, since this is one of the very few available single-cell datasets with RNA-seq and FUCCI data on the same cells, we nonetheless compared methods on these data.

We analyzed the 247 FUCCI-expressing hESC single-cell samples from Leng et al. (2015) that passed quality control: 91 G1 phase, 80 S phase, and 76 G2/M phase. Fig. 4A shows the average gene expression levels of top 4 cyclic genes in G1, S, and G2/M phase. We applied peco and the four existing methods to these data (for peco we used only the top 4 cyclic genes because *H4C5*, was not mapped in these data).

Comparing results from the three continuous assignment methods (peco, Oscope and reCAT), we found that the orderings from reCAT agree most closely with the gating-based classification (Fig. 4B). Results from peco also show strong agreement with gating-based classification, but the S-phase cells are spread out on either side of the G2/M cells, rather than only preceding them as in the reCAT results. In contrast, the ordering from Oscope shows less agreement with the gating-based classification. (Quantifying these qualitative statements is not straightforward because it is not obvious how to quantitatively compare a continuous cyclic ordering with a discrete classification; nonetheless we believe the qualitative patterns are clear in Fig. 4B.)

**Figure 4:**
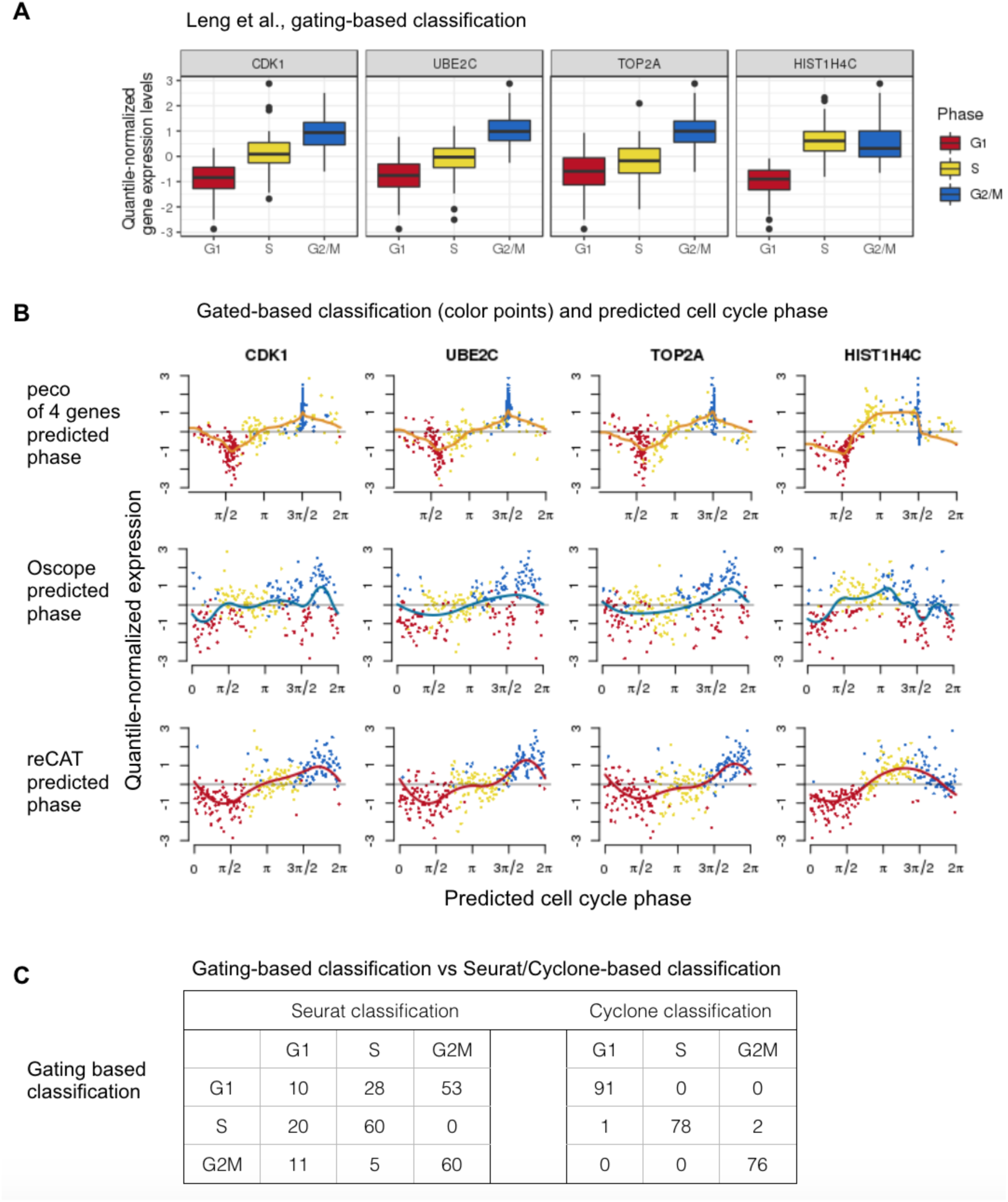
Applying peco and existing tools to data from Leng et al. 2015. The single-cell samples in this data were sorted into G1, S and G2M phase. (A) We plot the distribution of gene expression for the top 4 cyclic genes per cell cycle phase. (B) We compare predicted phases based on peco, Oscope and reCAT. Rows correspond to prediction results based on the three methods. Specifically, we sort the single-cell samples according to the predicted phase, and color the sample points according to the gated phase. For example, in the first row, we show that the peak expression profile of peco prediction is consistent with results based on gating. The orange line corresponds to the cyclic trend of expression levels. (C) We compare the phase assignment based on gating with Seurat/Cyclone-based classification.

Turning to the discrete classification methods (Seurat and Cyclone), in these data the Cyclone discrete assignments show much better agreement with the gating-based patterns than Seurat (see Fig. 4C, Cyclone misclassification rates 0% G1, 2.5% S, and 0% G2/M; Seurat misclassification rates 89% G1, 25% S, and 21% G2/M).

Overall our results suggest the need for more research and better data to quantify the accuracy and relative performance of the different available methods, including ours.

## Discussion

In this study we sought to characterize the effects of cell cycle progression on gene expression data from single cells (iPSCs), by jointly measuring both cell cycle phase (via FUCCI) and expression (via scRNA-seq) from the same cells. Our study differs in two key ways from previous similar studies. First, unlike the most commonly-cited previous studies (Leng et al. 2015; Buettner et al. 2015), our experimental design avoided confounding batch/plate effects with cell cycle phase. In these previous studies, cells were FACS-sorted by discrete cell cycle phase and loaded onto different C1 plates, making it difficult to decouple batch effects from cell cycle effects (Hicks et al. 2018). Second, our study focused on characterizing cell cycle progression in a continuum, rather than as abrupt transitions between discrete cell cycle phases.

We found that a simple predictor, based on 5 genes with a cyclic expression pattern (*CDK1*, *UBE2C*, *TOP2A*, *H4C3*, *H4C5*), was sufficient to predict cell cycle progression in our data, and that adding information from other genes did not improve prediction accuracy. That these particular genes should be helpful predictors of cell cycle is somewhat expected, as they have been reported as potential markers in previous studies, including synchronization experiments in HeLa cells (Whitfield et al. 2002) and yeast (Spellman et al. 1998), and in previous scRNA-seq studies of FUCCI-sorted hESCs (Leng et al. 2015). However, our finding that additional genes did not further improve prediction accuracy is unexpected, and contrasts with the common use of dozens of genes for cell cycle prediction (e.g., Seurat by Butler et al. 2018). Of course, our results do not imply that only these five genes are associated with cell cycle progression in iPSCs, only that additional genes provide redundant information in our data.

As noted in the Introduction, one reason to estimate cell cycle from RNA-seq data is to control for it when performing other downstream tasks. Although our methods provide a way to estimate cell cycle information, they do not dictate a specific way to control for cell cycle in downstream analyses. Indeed, how best to do this remains an interesting and open question, and the ease with which it can be achieved will depend on the downstream analyses being performed. For example, if the downstream analyses rely on Gaussian models for transformed single-cell data (e.g., Z Ji and H Ji 2016; Kiselev et al. 2017) then it may suffice to first regress out the effects of cell cycle from the transformed data (e.g. using a non-parametric regression method such as trend filtering, to allow for the non-linear trends that must occur in any cyclic phenomenon) before applying downstream analyses to the residuals. On the other hand, if the downstream methods rely on explicit models for count data (e.g., Dey et al. 2017) then controlling for cell cycle may be more complicated and require further methodological development. However, we note that these issues are not unique to our approach: controlling for cell cycle within count-based analyses poses additional methodological challenges for whatever method is used to estimate cell cycle.

One important question is how well our methods will generalize beyond the data collected here. We believe that our methods should be useful in other iPSC studies because we were able to effectively predict cell cycle progression in cells from one individual using scRNA-seq data from five other individuals (that is, our approach worked well in out-of-sample prediction assessment). However, further data are required to assess how well our methods generalize to studies involving different cell types than the iPSCs studied here.

Single-cell omics technology allows us to characterize cellular heterogeneity at an ever-increasing scale and high resolution. We argue that the standard way of classifying biological states in general, and cell cycle in particular, to discrete types, is no longer sufficient for capturing the complexities of expression variation at the cellular level. Our study provides a foundation for future work to characterize the effect of the cell cycle at single-cell resolution and to study cellular heterogeneity in single-cell expression studies.

## Methods

### FUCCI-iPSC cell lines and cell culture

Six previously characterized YRI iPSCs (Banovich et al. 2018), including three females (NA18855, NA18511, and NA18870) and three males (NA19098, NA19101, and NA19160), were used to generate FUCCI iPSC lines by the PiggyBAC insertion of a cassette encoding an EEF1A1 promoter-driven mCherryCDT1-IRES-EgfpGMNN double transgene (the plasmid was generously gifted by Dr. Chris Barry) (Sakaue-Sawano, Kurokawa, et al. 2008; Leng et al. 2015). Transfection of these iPSCs with the plasmid and Super piggyBacTM transposase mRNA (Transposagen) was done using the Human Stem Cell Nucleofector Kit 1 (VAPH-5012) by Nucleofector 2b Device (AAB-1001, Lonza) according to the manual. Single-cell suspension for the transfection was freshly prepared each time using TrypLETM Select Enzyme (1X) with no phenol red (Thermo Fisher Scientific) to maintain cell viability. For standard maintenance, cells were split every 3–4 days using cell release solution (0.5 mM EDTA and NaCl in PBS) at the confluence of roughly 80%.

After two regular passages on the 6-wells, the transfected cells were submitted to fluorescence activated cell sorting (FACS) for the selection of double positive (EGFP and mCherry) single cells. To increase the cell survival after FACS, Y27632 ROCK inhibitor (Sigma-Aldrich) was included in E8 medium (Life Technologies) for the first day. FACS was performed on the FACSAria IIIu instrument at University of Chicago Flow Cytometry Facility. Up to 12 individual clones from each of the six iPSC lines were maintained in E8 medium on Matrigel-coated tissue culture plates with daily media feeding at 37°C with 5% (vol/vol) CO2, same as regular iPSCs. After another ten passages of the FUCCI-iPSCs, a second round of FACS was performed to confirm the activation of the FUCCI transgene before single-cell collection on the C1 platform.

### Single-cell capture and image acquisition

Single-cell loading, capture, and library preparations were performed following the Fluidigm protocol (PN 100-7168) and as described in Tung et al. (2017). Specifically, the reverse transcription primer and the 1:50,000 Ambion® ERCC Spike-In Mix1 (Life Technologies) were added to the lysis buffer, and the template-switching RNA oligos which contain the UMI (6-bp random sequence) were included in the reverse transcription mix. A cell mixture of two different YRI FUCCI-iPSC lines was freshly prepared using TrypLE^TM^ at 37°C for three minutes. Cell viability and cell number were measured to have an equal number of live cells from the two FUCCI-iPSC lines. In addition, single-cell suspensions were stained with 5 uM Vybrant^TM^ DyeCycle^TM^ Violet Stain (Thermo Fisher Scientific) at 37°C for five minutes right before adding the C1 suspension buffer.

After the cell sorting step on the C1 machine, the C1 IFC microfluidic chip was immediately transferred to JuLI Stage (NanoEnTek) for imaging. The JuLI stage was specifically designed as an automated single-cell observation system for C1 IFC vessel. For each cell capture site, four images were captured, including bright field, DAPI, EGFP, and mCherry. The total imaging time, together with the setup time, was roughly 45 minutes for one 96-well C1 IFC. The JuLI Stage runs a series of standardized steps for each C1 IFC and for each fluorescence channel, separately. First, the camera scans the four corners of the C1 IFC and sets the exposure setting accordingly. Then, the camera proceeds to capture images of each C1 well.

### Library preparation and read mapping

For sequencing library preparation, tagmentation and isolation of 5 fragments were performed as described in our previous work (Tung et al. 2017). The sequencing libraries generated from the 96 single-cell samples of each C1 chip were pooled and then sequenced in two lanes on an Illumina HiSeq 2500 instrument using TruSeq SBS Kit v3-HS (FC-401-3002).

We mapped the reads with Subjunc (Liao et al. 2013) to a combined genome that included human genome GRCh37, ERCC RNA Spike-In Mix 1 (Invitrogen), and the mCherry and EGFP open reading frames from the FUCCI plasmid (we included the latter to ensure that the transgene was being transcribed). Next, we extracted the UMIs from the 5′ end of each read (pre-mapping) and deduplicated the UMIs (post-mapping) with UMI-tools (Smith et al. 2017). We counted the molecules per protein-coding gene (Ensembl 75, February 2014) with featureCounts (Liao et al. 2014). Note that we observed quantitatively similar results when using genome build GRCh38 and gene annotations from Ensembl 96 (April 2019) (Supplemental Fig. S18). Lastly, we matched each single cell to its individual of origin with verifyBamID (Jun et al. 2012) by comparing the genetic variation present in the RNA-seq reads to the known genotypes.

### Image analysis and FUCCI phase quantification

We analyzed images captured for each C1 well in the EGFP, mCherry, and DAPI channels. We used the DAPI images to identify individual nuclei location. This allowed us to identify the number of cells captured in each C1 well, and to align the EGFP and mCherry cell images based on the nucleus location. EBImage package in R/Bioconductor (Pau et al. 2010) was used for image processing and analysis. First, we normalized pixel intensities in each DAPI image and applied a ten-pixel median filter. Next, we generated 10 a nuclear mask using the EBImage adaptive thresholding algorithm. We filled holes in the resulting binary image and smoothed borders with a single round of erosion and dilation. Finally, we identified individual nuclei using the EBImage bwlabel function. The code that implements these methods is available at https://raw.githubusercontent.com/jdblischak/fucci-seq/master/code/create_mask.R.

To score the fluorescence intensity signals in each channel, we defined a 100 by 100 pixel cell area for all channel images centered on the nucleus centroid location. We estimated the background florescence in each channel image by taking the median intensity value of all pixels outside the defined cell area. We then subtracted this background intensity from intensity values of pixels located within the defined cell area. Finally, we summed and log-transformed the background-removed fluorescence intensities in the defined cell area. For each cell, this yielded a DAPI score and two FUCCI scores (mCherry and EGFP scores) summarizing fluorescence intensities of mCherry-Cdt1 and EGFP-Geminin.

We tested batch effects on FUCCI and DAPI scores using analysis of variance (score ∼ plate + individual). Type III sum of squares were computed to test for C1 plate effect while controlling for individual effect, and vice versa. To adjust for C1 plate effect, we subtracted the marginal means of plate effect from FUCCI and DAPI scores controlling for individual effect. Supplemental Fig. S19 shows the relationship between the corrected FUCCI and DAPI scores in the 888 single-cell samples.

For FUCCI phase quantification, we used the corrected EGFP and mCherry scores - log_10_ sum of fluorescence intensity in the 100 x 100 defined cell area after background and C1 plate effect correction - to infer an angle for each cell on a unit circle, where the angle is the inverse tangent function of (EGFP/mCherry). We refer to these angles as FUCCI phase, namely the estimated cell cycle phase based on FUCCI intensities.

Finally, we applied PAM from Kaufman and Rousseeuw 1990 to FUCCI scores to assign single-cell samples to G1, S and G2/M phases. DAPI is also commonly used to sort single cells into discrete cell cycle phases based on their relative quantification of celluar DNA content (Krishan 1975; Roukos et al. 2015). In our data, we observed substantial plate-to-plate variability in the range of DAPI scores both before and after batch correction (see Supplemental Fig. S20). To avoid batch bias in assigning discrete cell cycle phases, the DAPI results were not used in the cell cycle analysis in our data.

### Filtering and normalization of gene expression data

We used DAPI staining results to inform our RNA-seq quality control analysis in two steps. First, we used DAPI staining results to classify each C1 well into empty or non-empty wells. We then used data from the empty wells to determine filtering criteria for the non-empty wells (see Supplemental Fig. S2): number of mapped reads, percentage of unmapped reads, percentage of ERCC reads, and percentage of genes detected to have at least one read. Second, we determined the number of cells captured in each C1 well using linear discriminant analysis (LDA; see our previous work for the rationale, Tung et al. 2017). We fitted two LDA models: 1) number of cells per well ∼ gene molecule count + concentration of cDNA amplicons, and 2) number of cells per well ∼ read-to-molecule conversion efficiency of ERCC spike-in controls + read-to-molecule conversion efficiency of endogeneous genes. We used DAPI staining results to determine the number of cells captured in each well. Supplemental Fig. S21 shows the results of our LDA analysis. These scRNA-seq sample quality control steps have been described in details in Tung et al. (2017).

In summary, our quality control criteria include the following:

- Only one cell observed per well
- At least one molecule mapped to EGFP (to ensure the transgene is transcribed)
- The individual assigned by verifyBamID was included on the C1 chip
- At least 1,309,921 reads mapped to the genome
- Less than 44% unmapped reads
- Less than 18% ERCC reads
- At least 6,292 genes with at least one read

After sample filtering, we excluded genes based on the following criteria.

- Over-expressed genes with more than 6^4^ molecules across the samples.
- Lowly-expressed genes with sample average of CPM less than 2.

In total, we collected 20,327 genes from 1,536 scRNA-seq samples after read mapping. After the quality filtering steps described above, we were left with 888 samples and 11,040 genes. We standardized the molecule counts to CPM using per-sample total molecule count pre-filtering from the 20,327 genes.

### Estimating cyclic trends in gene expression data

To estimate the cyclic trend of gene expression, we ordered the single-cell samples by the measured FUCCI phase and applied nonparametric trend filtering. We quantile-normalized CPM values of each gene to a standard normal distribution. This way, the samples with zero molecule count were assigned the lowest level of gene expression. We applied quadratic (second order) trend filtering using the *trendfilter* function in the *genlasso* package (Tibshirani 2014). The *trendfilter* function implements a nonparametric smoothing method which chooses the smoothing parameter by cross-validation and fits a piecewise polynomial regression. In more specifics: The *trendfilter* method determines the folds in cross-validation in a nonrandom manner. Every k-th data point in the ordered sample is placed in the k-th fold, so the folds contain ordered subsamples. We applied five-fold cross-validation and chose the smoothing penalty using the option *lambda.1se*: among all possible values of the penalty term, the largest value such that the cross-validation standard error is within one standard error of the minimum. Furthermore, we desired that the estimated expression trend be cyclical. To encourage this, we concatenated the ordered gene expression data three times, with one added after another. The quadratic trend filtering was applied to the concatenated data series of each gene. The estimates from the middle series were extracted and taken as the estimated cyclic trend of each gene. Using this approach, we ensured that the estimated trend be continuous at the boundaries of the ordered data: the estimates at the beginning always meet the estimates at the end of the ordered data series.

We used a permutation-based test to assess the significance of each inferred cyclic trend. For each gene, we computed the proportion of variance explained (PVE) by the inferred cyclic trend in the expression levels. Then, we constructed an empirical null distribution of PVE. We randomly chose a gene with less than 10 % of the cells observed as undetected (CPM ≥ 1) and permuted the expression levels in the selected gene 1,000 times. Each time, we fit *trendfilter* and computed PVE of the cyclic trend. We found that the significance (p-value) of the inferred cyclic trend was more conservative when the empirical null was based on a gene with low proportion of undetected cells, compared to when the empirical null was based on a gene with high proportion of detected cells (> 80 %). Using these empirical p-values, we were able to assess significance of the cyclic trends for each gene.

### Predicting quantitative cell cycle phase of single cells: a supervised learning approach

Our goal was to build a statistical method to predict continuous cell cycle phase from gene expression data. We implemented the method in a two-step algorithm. In the first step, we trained our predictor on data from 5 individuals and learned the cyclic trend for each gene using *trendfilter*. In the second step, we applied the predictor and used the gene-specific trends to compute the likelihood of gene expression levels in the test data for each cell. We evaluated the likelihood on grid points selected along a circle (default to 100 equally-spaced cell cycle phases). Finally, we assigned each cell in the test data to a grid point (phase) at which its likelihood reaches the maximum. Because we independently assigned each cell based on its gene expression levels, prediction accuracy does not depend on the number of cells in the test data.

#### Notations

- 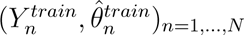: For each individual cell *n* in the training sample, we denote 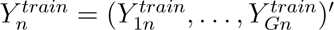 as the quantile-normalized gene expression vector, and 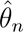 the FUCCI-based cell cycle phases. The single-cell samples are ordered in FUCCI time, where 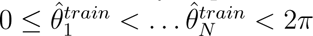.
- 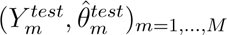: For each cell *m* in the test data, 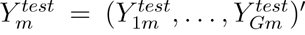 denotes the log_2_ normalized gene expression vector. The method estimates 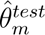 the cell cycle phase for each sample *m*.
- 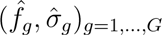: Using the training data *Y^train^*, we estimate a function 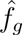 for each gene describing the cyclic trend of gene expression levels in FUCCI phase. *f* is a cyclic function assumed to be continuous at 0 and 2*π*.

Our approach to predicting quantitative cell cycle phase is related to methods that use gene expression data to predict circadian time in humans (e.g., Braun et al. 2018; Hughey et al. 2016). Among these methods, Hughey et al. 2016 is perhaps the most similar to ours, but uses smoothing splines instead of trend filtering to estimate cyclic trends, and a more complex method to combine information across genes.

#### Methods

1. Estimate 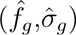 using 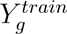 gene expression levels of gene *g*
  a. Sort the gene expression levels 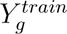 in ascending order according to the cell times 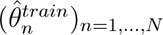.
  b. For each gene *g*, fit a piecewise polynomial function 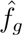 using *trendfilter*. (This function uses internal 5-fold cross-validation to determine an appropriate amount of smoothing for each *g*).
  c. Compute the gene-specific standard error 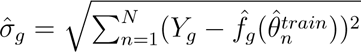
2. Predict 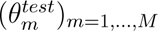 using the gene expression data 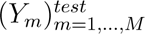
  a. Choose *K* discrete and equally-spaced cell times between 0 to 2*π*. For now, we choose *K* = 100, which is pretty large considering the size of 155 cells in the test sample.
  b. Compute the likelihood of 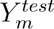 at each cell time *k*:

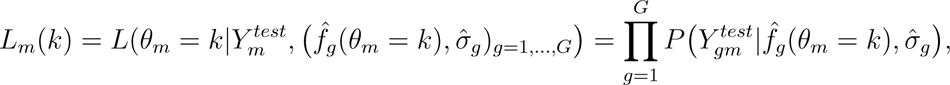

where 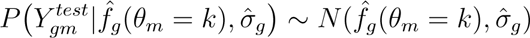
  c. Maximize *L_m_*(*k*) over *k* = 1, …, 100:

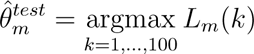

## Supporting information

Supplemental_Table_S1: A curated set of 622 cell cycle genes from Macosko et al. 2015 (a subset of genes annotated in Whitfield et al. 2002).

Supplemental_Table_S2: Complete list of genes analyzed in our study

Source code of our analysis, including scripts and data necessary to reproduce the work.

## Data access

All raw and processed sequencing data generated in this study have been submitted to the NCBI Gene Expression Omnibus (GEO; http://www.ncbi.nlm.nih.gov/geo/, Edgar et al. 2002) under accession number GSE121265. We also make the processed data available at https://github.com/jhsiao999/peco-paper and https://giladlab.uchicago.edu/wp-content/uploads/2019/02/Hsiao_et_al_2019.tar.gz. All analysis results, scripts and data required to reproduce this work are available at https://jhsiao999.github.io/peco-paper. as well as in Supplemental Material. The source code is available in an R/Bioconductor package peco (the development version of peco is available at https://github.com/jhsiao999/peco.

## Acknowledgements

We thank members of the Gilad and Stephens laboratories for valuable discussions during the preparation of this manuscript. We also thank Natalia Gonzales for constructive feedback on this manuscript. We are also grateful to the editors and referees for insightful comments and suggestions. This work was funded by NIH grant HG002585 to M.S. and NIH grant GM122930 to Y.G. The content is solely the responsibility of the authors and does not necessarily represent the official views of the National Institutes of Health.

## Author Contributions

CJH, PYT, YG, and MS conceived of the study, designed the experiments, and formulated the analysis framework. PYT performed the experiments with assistance from JEB. CJH and MS developed the statistical approach for predicting continuous cell cycle phase, and CJH implemented the algorithm. CJH wrote the R package with assistance from KAB and KKD. CJH analyzed the data, with assistance from KB, JDB, PYT, and MS. CJH, MS and YG wrote the original draft with input from PYT, JDB and KAB. All authors reviewed the final manuscript.

## References

Banovich NE et al. 2018. Impact of regulatory variation across human iPSCs and differentiated cells. Genome Res. 28: 122–131.

Barron M and Li J. 2016. Identifying and removing the cell-cycle effect from single-cell RNA-Sequencing data. Sci. Rep. 6: 33892. doi: 10.1038/srep33892.

Braun R, Kath WL, Iwanaszko M, Kula-Eversole E, Abbott SM, Reid KJ, Zee PC, and Allada R. 2018. Universal method for robust detection of circadian state from gene expression. Proc. Natl. Acad. Sci. U. S. A. 115: E9247–E9256.

Buettner F, Natarajan KN, Casale FP, Proserpio V, Scialdone A, Theis FJ, Teichmann SA, Marioni JC, and Stegle O. 2015. Computational analysis of cell-to-cell heterogeneity in single-cell RNA-sequencing data reveals hidden subpopulations of cells. Nat. Biotechnol. 33: 155–160.

Butler A, Hoffman P, Smibert P, Papalexi E, and Satija R. 2018. Integrating single-cell transcriptomic data across different conditions, technologies, and species. Nat. Biotechnol. 36: 411–420.

Chen M and Zhou X. 2017. Controlling for confounding effects in single cell RNA sequencing studies using both control and target genes. Sci. Rep. 7: 13587. doi:10.1038/s41598-017-13665-w.

Dey KK, Hsiao CJ, and Stephens M. 2017. Visualizing the structure of RNA-seq expression data using grade of membership models. PLoS Genet. 13: e1006599. doi: 10.1371/journal.pgen.1006599.

Edgar R, Domrachev M, and Lash AE. 2002. Gene Expression Omnibus: NCBI gene expression and hybridization array data repository. Nucleic acids research. 30: 207–210.

Hicks SC, Townes FW, Teng M, and Irizarry RA. 2018. Missing data and technical variability in single-cell RNA-sequencing experiments. Biostatistics. 19: 562–578.

Hughey JJ, Hastie T, and Butte AJ. 2016. ZeitZeiger: supervised learning for high-dimensional data from an oscillatory system. Nucleic Acids Research. 44: e80–e80. doi: 10.1093/nar/gkw030.

Ingolia NT and Murray AW. 2004. The ups and downs of modeling the cell cycle. Curr. Biol. 14: PR771–R777.

Ji Z and Ji H. 2016. TSCAN: Pseudo-time reconstruction and evaluation in single-cell RNA-seq analysis. Nucleic Acids Res. 44: e117. doi: 10.1093/nar/gkw430.

Jun G, Flickinger M, Hetrick KN, Romm JM, Doheny KF, Abecasis GR, Boehnke M, and Kang HM. 2012. Detecting and estimating contamination of human DNA samples in sequencing and array-based genotype data. Am. J. Hum. Genet. 91: 839–848.

Kaufman L and Rousseeuw PJ (1990). Innovation and Intellectual Property Rights. In: Finding Groups in Data: An Introduction to Cluster Analysis. Ed. by L Kaufman and PJ Rousseeuw. Hoboken: John Wiley Sons, Inc. Chap. 2, pp. 68–125.

Kelsey G, Stegle O, and Reik W. 2017. Single-cell epigenomics: Recording the past and predicting the future. Science. 358: 69–75.

Keren L, Dijk D van, Weingarten-Gabbay S, Davidi D, Jona G, Weinberger A, Milo R, and Segal E. 2015. Noise in gene expression is coupled to growth rate. Genome Research. 25: 1893–1902.

Kiselev VY et al. 2017. SC3: Consensus clustering of single-cell RNA-seq data. Nat. Methods. 14: 483–486.

Kolodziejczyk AA et al. 2015. Single Cell RNA-Sequencing of Pluripotent States Unlocks Modular Transcriptional Variation. Cell Stem Cell. 17: 471–485.

Kowalczyk MS, Tirosh I, Heckl D, Rao TN, Dixit A, Haas BJ, Schneider RK, Wagers AJ, Ebert BL, and Regev A. 2015. Single-cell RNA-seq reveals changes in cell cycle and differentiation programs upon aging of hematopoietic stem cells. Genome Res. 25: 1860–1872.

Krishan A. 1975. Rapid flow cytofluorometric analysis of mammalian cell cycle by propidium iodide staining. Journal of Cell Biology. 66: 188-193.

Lauridsen FKB et al. 2018. Differences in cell cycle status underlie transcriptional heterogeneity in the HSC Compartment. Cell Rep. 24: 766–780.

Leng N, Chu LF, Barry C, Li Y, Choi J, Li X, Jiang P, Stewart RM, Thomson JA, and Kendziorski C. 2015. Oscope identifies oscillatory genes in unsynchronized single-cell RNA-seq experiments. Nat. Methods. 12: 947–950.

Liao Y, Smyth GK, and Shi W. 2013. The Subread aligner: fast, accurate and scalable read mapping by seed-and-vote. Nucleic Acids Res. 41: e108. doi: 10.1093/nar/gkt214.

Liao Y, Smyth GK, and Shi W. 2014. featureCounts: an efficient general purpose program for assigning sequence reads to genomic features. Bioinformatics. 30: 923–930.

Liu Z, Lou H, Xie K, Wang H, Chen N, Aparicio OM, Zhang MQ, Jiang R, and Chen T. 2017. Reconstructing cell cycle pseudo time-series via single-cell transcriptome data. Nat. Commun. 8: doi: 10.1038/s41467-017-00039-z.

Lu Y, Biancotto A, Cheung F, Remmers E, Shah N, McCoy JP, and Tsang JS. 2016. Systematic analysis of cell-to-cell expression variation of T lymphocytes in a human cohort identifies aging and genetic associations. Immunity. 45: 1162–1175.

Macaulay IC, Ponting CP, and Voet T. 2017. Single-Cell Multiomics: Multiple Measurements from Single Cells. Trends Genet. 33: 155–168.

Macosko EZ et al. 2015. Highly Parallel Genome-wide Expression Profiling of Individual Cells Using Nanoliter Droplets. Cell. 161: 1202–1214.

Maechler M, Rousseeuw P, Struyf A, Hubert M, and Hornik K (2019). cluster: Cluster Analysis Basics and Extensions. R package version 2.1.0.

Nguyen QH et al. 2018. Profiling human breast epithelial cells using single cell RNA sequencing identifies cell diversity. Nat. Commun. 9: doi: 10.1038/s41467-018-04334-1.

Papalexi E and Satija R. 2018. Single-cell RNA sequencing to explore immune cell heterogeneity. Nat. Rev. Immunol. 18: 35–45.

Pau G, Fuchs F, Sklyar O, Boutros M, and Huber W. 2010. EBImage–an R package for image processing with applications to cellular phenotype\s. Bioinformatics. 26: 979–981.

Pauklin S and Vallier L. 2013. The cell-cycle state of stem cells determines cell fate propensity. Cell. 155: 135–47.

Povinelli B, Wills Q, Barkas N, Booth C, Campbell K, Rodriguez-Meira A, Jacobsen SE, Yau C, and Mead A. 2018. Integrated single cell analysis reveals cell cycle and ontogeny related transcriptional heterogeneity in HSCs. Exp. Hematol. 64: S95–S96.

Roukos V, Pegoraro G, Voss TC, and Misteli T. 2015. Cell cycle staging of individual cells by fluorescence microscopy. Nat. Protoc. 10: 334-348.

Sakaue-Sawano A, Kurokawa H, et al. 2008. Visualizing spatiotemporal dynamics of multicellular cell-cycle progression. Cell. 132: 487–498.

Sakaue-Sawano A, Yo M, Komatsu N, Hiratsuka T, Kogure T, Hoshida T, Goshima N, Matsuda M, Miyoshi H, and Miyawaki A. 2017. Genetically encoded tools for optical dissection of the mammalian cell cycle. Mol. Cell. 68: 626–640.E5.

Sanchez A and Golding I. 2013. Genetic determinants and cellular constraints in noisy gene expression. Science. 342: 1188–1193.

Scialdone A, Natarajan KN, Saraiva LR, Proserpio V, Teichmann SA, Stegle O, Marioni JC, and Buettner F. 2015. Computational assignment of cell-cycle stage from single-cell transcriptome data. Methods. 85: 54–61.

Skelly DA, Squiers GT, McLellan MA, Bolisetty MT, Robson P, Rosenthal NA, and Pinto AR. 2018. Single-cell transcriptional profiling reveals cellular diversity and intercommunication in the mouse heart. Cell Rep. 22: 600–610.

Smith T, Heger A, and Sudbery I. 2017. UMI-tools: modeling sequencing errors in Unique Molecular Identifiers to improve quantification accuracy. Genome Res. 27: 491–499.

Soltani M and Singh A. 2016. Effects of cell-cycle-dependent expression on random fluctuations in protein levels. R Soc Open Sci. 3: doi: 10.1098/rsos.160578.

Spellman PT, Sherlock G, Zhang MQ, Iyer VR, Anders K, Eisen MB, Brown PO, Botstein D, Futcher B, and Fink GR. 1998. Comprehensive identification of cell cycle–regulated genes of the yeast saccharomyces cerevisiae by microarray hybridization. MBoC. 9: 3273–3297.

Stubbington MJT, Rozenblatt-Rosen O, Regev A, and Teichmann SA. 2017. Single-cell transcriptomics to explore the immune system in health and disease. Science. 358: 58–63.

Tanay A and Regev A. 2017. Scaling single-cell genomics from phenomenology to mechanism. Nature. 541: 331–338.

Tibshirani RJ. 2014. Adaptive piecewise polynomial estimation via trend filtering. Ann. Stat. 42: 285–323.

Tung PY, Blischak JD, Hsiao CJ, Knowles DA, Burnett JE, Pritchard JK, and Gilad Y. 2017. Batch effects and the effective design of single-cell gene expression studies. Sci. Rep. 7: 39921. doi: 10.1038/srep39921.

Velten L et al. 2017. Human haematopoietic stem cell lineage commitment is a continuous process. Nat. Cell Biol. 19: 271–281.

Whitfield ML et al. 2002. Identification of genes periodically expressed in the human cell cycle and their expression in tumors. Mol. Biol. Cell. 13: 1977–2000.

