## Supplementary material for "Characterizing and inferring quantitative cell cycle phase in single-cell RNA-seq data analysis": Source code of our analysis, including scripts and data necessary to reproduce the work.: _workflowr.yml

```
# workflowr options
# Version 1.4.0

# The seed to use for random number generation. See ?set.seed for details.
seed: 20190814
# The working directory to build the R Markdown files. The path is relative to
# _workflowr.yml. See ?rmarkdown::render for details.
knit_root_dir: "."
```
