## Supplementary material for "Characterizing and inferring quantitative cell cycle phase in single-cell RNA-seq data analysis": Source code of our analysis, including scripts and data necessary to reproduce the work.: _site.yml

```
name: peco-paper
output_dir: ../docs
navbar:
  title: peco-paper
  left:
  - text: Home
    href: index.html
  - text: About
    href: about.html
  - text: License
    href: license.html
  right:
  - icon: fa-github
    text: Source code
    href: https://github.com/jhsiao999/peco-paper
output:
  workflowr::wflow_html:
    toc: yes
    toc_float: yes
    theme: cosmo
    highlight: textmate
```
