## Supplementary material for "Characterizing and inferring quantitative cell cycle phase in single-cell RNA-seq data analysis": Source code of our analysis, including scripts and data necessary to reproduce the work.: about.Rmd

```
---
title: "About"
output:
  workflowr::wflow_html:
    toc: false
editor_options:
  chunk_output_type: console
---


We combined fluorescence imaging with scRNA-seq to measure cell cycle phase and gene expression levels in human induced pluripotent stem cells (iPSCs). Using these data, we developed a novel approach to characterize cell cycle progression. While standard methods assign cells to discrete cell cycle stages, our method goes beyond this, and quantifies cell cycle progression on a continuum.

Find out more about our study in our paper.

* bioRxiv: [Characterizing and inferring quantitative cell-cycle phase in single-cell RNA-seq data analysis](https://www.biorxiv.org/content/10.1101/526848v2)
```
