## Supplementary material for "Characterizing and inferring quantitative cell cycle phase in single-cell RNA-seq data analysis": Source code of our analysis, including scripts and data necessary to reproduce the work.: access_data.Rmd

````
---
title: "Access data"
author: "John Blischak, PoYuan Tung, Joyce Hsiao"
output:
  workflowr::wflow_html:
    toc: TRUE
    toc_float: TRUE
---


```{r setup, include=FALSE}
knitr::opts_chunk$set(echo = F)
```


Here are some useful links for how we processed our data.

  * [Correct for C1 batch effects in FUCCI intensity data](https://jhsiao999.github.io/peco-paper/images_batchcorrect.html).

  * Assess potential C1 batch effects in [sequencing depth](https://jhsiao999.github.io/peco-paper/totals.html) and [read to molecule conversion rate](https://jhsiao999.github.io/peco-paper/reads-v-molecules.html).
  
  * Identify high quality samples using [sample quality metrics](https://jhsiao999.github.io/peco-paper/sampleqc.html), such as total number of mapped reads, ERCC percentages, etc.
    
  * [Filter out over-expressed and lowly-expressed genes](https://jhsiao999.github.io/peco-paper/gene_filtering.html). 
  
  * Use [Principal Component Analysis](https://jhsiao999.github.io/peco-paper/pca_tf.html) to examined potential sources of gene expression variation.
````
