## Supplementary material for "Characterizing and inferring quantitative cell cycle phase in single-cell RNA-seq data analysis": Source code of our analysis, including scripts and data necessary to reproduce the work.: gene_filtering.Rmd

````
---
title: "Gene filtering and sample filtering"
author: "Joyce Hsiao"
output: 
  workflowr::wflow_html:
    toc: TRUE
    toc_float: TRUE
---


```{r setup, include=FALSE}
knitr::opts_chunk$set(echo = TRUE)
```

---

## Summary

I performed gene filtering based on the criterion set forth in our previous paper. 

1. Remove outlier genes: molecule counts > 4,096 in any sample (x is the theoretical maximum of UMI count for 6-bp UMI)

*Results* There's one, and turns out this over-expressed gene is one of the mitochrondrial genes.

$~$

2. Remove lowly expressed genes: Lowly-expressed genes := gene mean < 2 CPM.

*Results*: * Of 20,421 genes, 7,637 genes are classifed as lowly-expressed. Of these, 34 are ERCC control genes and 7,603 are endogenoeus genes. 


---

## Set-up


```{r, message=FALSE}
library(knitr)
library(SingleCellExperiment)
library(dplyr)
library(heatmap3)
library(testit)
library(cowplot)
library(biomaRt)
library(knitr)
library(data.table)

sce_raw <- readRDS("data/sce-raw.rds")
anno = data.frame(colData(sce_raw))
```

Filter out low-quality single cell samples.

```{r}
anno_filter <- anno[anno$filter_all == TRUE,]
count_filter <- assay(sce_raw)[,anno$filter_all == TRUE]
dim(count_filter)
```


## Over-expressed genes

There's one, and turns out this over-expressed gene.

```{r}
which_over_expressed <- which(apply(count_filter, 1, function(x) any(x>(4^6)) ))
over_expressed_genes <- rownames(count_filter)[which_over_expressed]
over_expressed_genes
```

Get over-expressed gene info via `biomaRt`.

```{r, eval = FALSE}
over_expressed_genes_info <- getBM(
  attributes = c("ensembl_gene_id", "chromosome_name",
                 "external_gene_name", "transcript_count",
                 "description"),
  filters = "ensembl_gene_id",
  values = over_expressed_genes,
  mart = ensembl)
```


## Filter out lowly-expressed genes

* Of 20,421 genes, 7,864 genes are classifed as lowly-expressed. Of these, 34 are ERCC control genes and 7,830 are endogenoeus genes. 

Compute CPM

```{r}
cpm <- t(t(count_filter)/colSums(count_filter))*(10^6)
```

Lowly-expressed genes := gene mean < 2 CPM

```{r}
cpm <- t(t(count_filter)/anno_filter$molecules)*(10^6)
which_lowly_expressed <- which(rowMeans(cpm) < 2)

length(grep("ERCC", which_lowly_expressed))
length(grep("ENSG", which_lowly_expressed))
```

Get gene info via `biomaRt`.

```{r, eval = FALSE}
lowly_expressed_genes_info <- getBM(
  attributes = c("ensembl_gene_id", "chromosome_name",
                 "external_gene_name", "transcript_count",
                 "description"),
  filters = "ensembl_gene_id",
  values = which_lowly_expressed[grep("ENSG", which_lowly_expressed_genes)],
  mart = ensembl)
```


## Combine filters

Including 12784 genes.

```{r}
gene_filter <- unique(c(which_over_expressed, which_lowly_expressed))
genes_to_include <- setdiff(1:nrow(count_filter), gene_filter)
length(genes_to_include)
```


## Session information

```{r, echo = FALSE}
sessionInfo()
```
````
