## Supplementary material for "Characterizing and inferring quantitative cell cycle phase in single-cell RNA-seq data analysis": Source code of our analysis, including scripts and data necessary to reproduce the work.: images_batchcorrect.Rmd

````
---
title: "Normalize intensities across C1 batches"
author: "Joyce Hsiao"
output: workflowr::wflow_html
---


```{r setup, include=FALSE}
knitr::opts_chunk$set(echo = TRUE, message=FALSE, warning=FALSE)
```

---

## Introduction/summary

We fit the following model to estimate individual effect $\gamma_i$ and C1 plate effect $\beta_j$ in fluorescence intensities. In notation, 

$$
y_{ijk} = \mu + \tau_i + \beta_j + \epsilon_{ijk}
$$
where $i = 1,2,..., I$ and $j = 1,2,..., J$. The parameters are estimated under sum-to-zero constraints $\sum \tau_i = 0$ and $\sum \beta_j = 0$.

Note that in this model 1) not all $y_{ij.}$ exists due to the incompleteness of the design, 2) the effects of individual and block are nonorthogonal, 3) the effects are additive due to the block design.


We found significant C1 plate effect in fluorescence intensities and corrected for this effect in GFP, RFP and DAPI channels. The corrected estimates are computed as

$$
\hat{y}_{ijk} = y_{ijk} - \hat{\beta}_j
$$
for all channels.


## Data and packages

```{r, echo = T}
library(data.table)
library(dplyr)
library(ggplot2)
library(cowplot)
library(RColorBrewer)
library(scales)
library(car)
library(lsmeans)
library(SingleCellExperiment)
```

Read in filtered data.

```{r}
sce <- readRDS(file="data/sce-final.rds")
cdata <- data.frame(colData(sce))
rdata <- data.frame(rowData(sce))
```


## Estimate source of variation

Statistical tests show that for GFP, there's significant plate effect (corresponds to *experiment* variable, P-value < 2E-16) but not significant individual effect (corresponds to *chip_id* varialbe).


```{r}
lm.rfp <- lm(rfp.median.log10sum~factor(chip_id)+factor(experiment),
             data = cdata)
lm.gfp <- lm(gfp.median.log10sum~factor(chip_id)+factor(experiment),
             data = cdata)
lm.dapi <- lm(dapi.median.log10sum~factor(chip_id)+factor(experiment),
             data = cdata)

aov.lm.rfp <- Anova(lm.rfp, type = "III")
aov.lm.gfp <- Anova(lm.gfp, type = "III")
aov.lm.dapi <- Anova(lm.dapi, type = "III")
aov.lm.rfp
aov.lm.gfp
aov.lm.dapi
```

Visualize indivdual and plate variation

```{r, fig.width=12, fig.height=8, echo = FALSE}
gfp.plot <- plot_grid(
  ggplot(cdata, 
          aes(x=chip_id, y=gfp.median.log10sum,
              col=chip_id)) +
    geom_violin() + geom_boxplot(width=.1),
  ggplot(cdata, 
        aes(x=factor(experiment), y=gfp.median.log10sum,
            col=factor(experiment))) +
  geom_violin() + geom_boxplot(width=.1), 
  ncol=2, labels = c("Individual variation", "Plate variation"))
title <- ggdraw() + draw_label("Green fluorescence intensites", fontface='bold')
plot_grid(title, gfp.plot, 
          ncol=1, rel_heights=c(0.1, 1)) # rel_heights values control title margins

## RFP
rfp.plot <- plot_grid(
  ggplot(cdata, 
          aes(x=chip_id, y=rfp.median.log10sum,
              col=chip_id)) +
    geom_violin() + geom_boxplot(width=.1),
  ggplot(cdata, 
        aes(x=factor(experiment), y=rfp.median.log10sum,
            col=factor(experiment))) +
  geom_violin() + geom_boxplot(width=.1), 
  ncol=2, labels = c("Individual variation", "Plate variation"))
title <- ggdraw() + draw_label("Red fluorescence intensites", fontface='bold')
plot_grid(title, rfp.plot, 
          ncol=1, rel_heights=c(0.1, 1)) # rel_heights values control title margins


## DAPI
dapi.plot <- plot_grid(
  ggplot(cdata, 
          aes(x=chip_id, y=dapi.median.log10sum,
              col=chip_id)) +
    geom_violin() + geom_boxplot(width=.1),
  ggplot(cdata, 
        aes(x=factor(experiment), y=dapi.median.log10sum,
            col=factor(experiment))) +
  geom_violin() + geom_boxplot(width=.1), 
  ncol=2, labels = c("Individual variation", "Plate variation"))
title <- ggdraw() + draw_label("DAPI intensites", fontface='bold')
plot_grid(title, dapi.plot, 
          ncol=1, rel_heights=c(0.1, 1)) # rel_heights values control title margins


gfp.plot
rfp.plot
dapi.plot
```

## Estimate plate

Contrast test to estimate the effect of C1 plate.

```{r}
# make contrast matrix for plates
# each plate is compared to the average
n_plates <- uniqueN(cdata$experiment)
contrast_plates <- matrix(-1, nrow=n_plates, ncol=n_plates)
diag(contrast_plates) <- n_plates-1


gfp.plates <- summary(lsmeans(lm.gfp, specs="experiment", contrast=contrast_plates))
rfp.plates <- summary(lsmeans(lm.rfp, specs="experiment", contrast=contrast_plates))
dapi.plates <- summary(lsmeans(lm.dapi, specs="experiment", contrast=contrast_plates))
```


Substract plate effect from the raw estimates.

```{r compute-adjusted, eval = T}
## RFP
cdata$rfp.median.log10sum.adjust <- cdata$rfp.median.log10sum
rfp.plates$experiment <- as.character(rfp.plates$experiment)
cdata$experiment <- as.character(cdata$experiment)

exps <- unique(cdata$experiment)
for (i in 1:uniqueN(exps)) {
  exp <- exps[i]
  ii_exp <- which(cdata$experiment == exp)
  est_exp <- rfp.plates$lsmean[which(rfp.plates$experiment==exp)]
  cdata$rfp.median.log10sum.adjust[ii_exp] <- (cdata$rfp.median.log10sum[ii_exp] - est_exp)
}


## GFP
cdata$gfp.median.log10sum.adjust <- cdata$gfp.median.log10sum
gfp.plates$experiment <- as.character(gfp.plates$experiment)
cdata$experiment <- as.character(cdata$experiment)

exps <- unique(cdata$experiment)
for (i in 1:uniqueN(exps)) {
  exp <- exps[i]
  ii_exp <- which(cdata$experiment == exp)
  est_exp <- gfp.plates$lsmean[which(gfp.plates$experiment==exp)]
  cdata$gfp.median.log10sum.adjust[ii_exp] <- (cdata$gfp.median.log10sum[ii_exp] - est_exp)
}


## DAPI
cdata$dapi.median.log10sum.adjust <- cdata$dapi.median.log10sum
dapi.plates$experiment <- as.character(dapi.plates$experiment)
cdata$experiment <- as.character(cdata$experiment)

exps <- unique(cdata$experiment)
for (i in 1:uniqueN(exps)) {
  exp <- exps[i]
  ii_exp <- which(cdata$experiment == exp)
  est_exp <- dapi.plates$lsmean[which(dapi.plates$experiment==exp)]
  cdata$dapi.median.log10sum.adjust[ii_exp] <- (cdata$dapi.median.log10sum[ii_exp] - est_exp)
}
```


Visualize intensities after adjusting for C1 plate effect.

```{r boxplot-adjusted, fig.width=12, fig.height=7, echo = FALSE}
gfp.plot <- plot_grid(
  ggplot(cdata, 
          aes(x=chip_id, y=gfp.median.log10sum.adjust,
              col=chip_id)) +
    geom_violin() + geom_boxplot(width=.1),
  ggplot(cdata, 
        aes(x=factor(experiment), y=gfp.median.log10sum.adjust,
            col=factor(experiment))) +
  geom_violin() + geom_boxplot(width=.1), 
  ncol=2, labels = c("Individual variation", "Plate variation"))
title <- ggdraw() + draw_label("Adjusted Green fluorescence intensites", fontface='bold')
plot_grid(title, gfp.plot, 
          ncol=1, rel_heights=c(0.1, 1)) # rel_heights values control title margins

## RFP
rfp.plot <- plot_grid(
  ggplot(cdata, 
          aes(x=chip_id, y=rfp.median.log10sum.adjust,
              col=chip_id)) +
    geom_violin() + geom_boxplot(width=.1),
  ggplot(cdata, 
        aes(x=factor(experiment), y=rfp.median.log10sum.adjust,
            col=factor(experiment))) +
  geom_violin() + geom_boxplot(width=.1), 
  ncol=2, labels = c("Individual variation", "Plate variation"))
title <- ggdraw() + draw_label("Adjusted Red fluorescence intensites", fontface='bold')
plot_grid(title, rfp.plot, 
          ncol=1, rel_heights=c(0.1, 1)) # rel_heights values control title margins


## DAPI
dapi.plot <- plot_grid(
  ggplot(cdata, 
          aes(x=chip_id, y=dapi.median.log10sum.adjust,
              col=chip_id)) +
    geom_violin() + geom_boxplot(width=.1),
  ggplot(cdata, 
        aes(x=factor(experiment), y=dapi.median.log10sum.adjust,
            col=factor(experiment))) +
  geom_violin() + geom_boxplot(width=.1), 
  ncol=2, labels = c("Individual variation", "Plate variation"))
title <- ggdraw() + draw_label("Adjusted DAPI intensites", fontface='bold')
plot_grid(title, dapi.plot, 
          ncol=1, rel_heights=c(0.1, 1)) # rel_heights values control title margins

gfp.plot
rfp.plot
dapi.plot
```

Analysis of variation on data after correcting for plate effect shows that plate effect is no longer a significant contributor of variation in fluorescence intensity.


```{r}
lm.rfp.adjust <- lm(rfp.median.log10sum.adjust~factor(chip_id)+factor(experiment),
             data = cdata)
lm.gfp.adjust <- lm(gfp.median.log10sum.adjust~factor(chip_id)+factor(experiment),
             data = cdata)
lm.dapi.adjust <- lm(dapi.median.log10sum.adjust~factor(chip_id)+factor(experiment),
             data = cdata)

aov.lm.rfp <- Anova(lm.rfp, type = "III")
aov.lm.gfp <- Anova(lm.gfp, type = "III")
aov.lm.dapi <- Anova(lm.dapi, type = "III")
aov.lm.rfp
aov.lm.gfp
aov.lm.dapi
```


```{r, echo = FALSE}
sessionInfo()
```
````
