## Supplementary material for "Characterizing and inferring quantitative cell cycle phase in single-cell RNA-seq data analysis": Source code of our analysis, including scripts and data necessary to reproduce the work.: index.Rmd

````
---
title: "peco"
site: workflowr::wflow_site
output:
  workflowr::wflow_html:
    toc: TRUE
    toc_float: FALSE
editor_options:
  chunk_output_type: console
---

## Overview

__peco__ is a supervised approach for predicting _continuous_ cell cycle phase using single-cell RNA-seq (scRNA-seq) data. The approach is described in [our paper](doi.org/10.1101/526848).

We use this site to document and share the code used to produce our analysis. Please feel free to explore. Comments and feedbacks are welcome!


## The software peco

We quantified continous cell cycle phase using FUCCI fluorescence imaging and trained peco to predict this continuous cell cycle phase using scRNA-seq data collected from six human cell lines. 
[Our paper](doi.org/10.1101/526848) showed that peco produces robust cell cycle phase predictions using strong cyclic genes - genes which expression levels oscillate along the cell cycle. 

The software __peco__ will be released in [Biconductor 3.11](https://www.bioconductor.org/developers/release-schedule/). This release will use the latest [R3.6.1](https://cran.r-project.org/).

The development version is available on [GitHub](https://github.com/jhsiao999/peco). To install the development version,

```{r, eval=F}
devtools::install_github("jhsiao999/peco")
library(peco)
```

## The analysis 

* [Process scRNA-seq and imaging data](https://jhsiao999.github.io/peco-paper/access_data.html)

* [Estimate the cyclic trends of gene expression levels](https://jhsiao999.github.io/peco-paper/npreg_trendfilter_quantile.html)

* [Compare the performance of peco with other methods on our data](https://jhsiao999.github.io/peco-paper/eval_on_our_data.html)

* [Compare the performance of peco with other methods using Leng et al. 2015 data](https://jhsiao999.github.io/peco-paper/eval_on_leng2015_data.html)

* [Assess the performance of peco in thinned data](https://jhsiao999.github.io/peco-paper/predict_thinned_data.html)

* [Produce figures shown in our paper](https://github.com/jhsiao999/peco-paper/tree/master/code)


## Citation

[Characterizing and inferring quantitative cell-cycle phase in single-cell RNA-seq data analysis](doi.org/10.1101/526848).


## Downloading the data files

You have two main options for downloading the data files. First, you can
manually download the individual files by clicking on the links on this page or
navigating to the files in the
[peco-paper](https://github.com/jhsiao999/peco-paper) GitHub repository. This is
the recommended strategy if you only need a few data files.

Second, you can install [git-lfs](https://git-lfs.github.com/). To handle large
files, we used Git Large File Storage (LFS). This means that the files that you
download with `git clone` are only plain text files that contain identifiers for
the files saved on GitHub's servers. If you want to download all of the data
files at once, you can do this with after you install git-lfs.

To install git-lfs, follow their instructions to download, install, and setup
(`git lfs install`). Alternatively, if you use conda, you can install git-lfs
with `conda install -c conda-forge git-lfs`. Once installed, you can download
the latest version of the data files with `git lfs pull`.


## Other information

* GEO record [GSE121265](http://www.ncbi.nlm.nih.gov/ geo/query/acc.cgi?acc=GSE121265) 
for all raw and processed sequencing data.

* We also provide the processed data in TXT format as a gzip compressed 
tarball on the [Gilad lab website](https://giladlab.uchicago.edu/wp-content/uploads/2019/02/Hsiao_et_al_2019.tar.gz).
````
