## Supplementary material for "Characterizing and inferring quantitative cell cycle phase in single-cell RNA-seq data analysis": Source code of our analysis, including scripts and data necessary to reproduce the work.: license.Rmd

```
---
title: "License"
output:
  workflowr::wflow_html:
    toc: false
editor_options:
  chunk_output_type: console
---

The code is available under the [MIT license][mit]. The text, data, and figures
are available under the [CC-BY license][cc-by]. Please see the file
[LICENSE][license] for the full license text.

Both of these allow you to reuse, adapt, or redistribute the content as long as
you attribute us. If you use our data, please attribute us as follows:

The single-cell RNA-seq data and FUCCI imaging data were generated by the Gilad lab at the University of Chicago and the Pritchard lab at Stanford University (see https://github.com/jdblischak/singlecell-qtl for details).

If you use our code, please attribute us as follows:

The code used to process the single-cell RNA-seq data was adapted from the code written by the [Gilad lab](gilad-lab) and the [Stephens lab](stephens-lab) at the University of Chicago (see https://github.com/jdblischak/fucci-seq for details).

[cc-by]: https://creativecommons.org/licenses/by/4.0/
[license]: https://github.com/jdblischak/fucci-seq/LICENSE
[mit]: https://choosealicense.com/licenses/mit/
[gilad-lab]: http://giladlab.uchicago.edu/
[stephens-lab]: http://stephenslab.uchicago.edu/
```
