## Supplementary material for "Characterizing and inferring quantitative cell cycle phase in single-cell RNA-seq data analysis": Source code of our analysis, including scripts and data necessary to reproduce the work.: totals.Rmd

````
---
title: "Sequencing depth per C1 chip"
author: "John Blischak"
date: 2017-11-28
output: workflowr::wflow_html
---

## Setup

```{r packages, message=FALSE}
library("dplyr")
library("DT")
library("ggplot2")
library("reshape2")
library("SingleCellExperiment")
theme_set(cowplot::theme_cowplot())
```


```{r data}
sce_raw = readRDS("data/sce-raw.rds")
anno = data.frame(colData(sce_raw))
anno$experiment = factor(anno$experiment)
```

## Total sequencing depth

```{r distribution-sequencing-depth}
ggplot(anno, aes(x = raw, color = experiment)) +
  geom_density() +
  labs(x = "Number of raw sequences per single cell", y = "Number of cells",
       title = "Distribution of total raw sequences per single cell") +
  scale_color_discrete(name = "C1 chip")
```

Mapped reads per cell

```{r mapped}
ggplot(anno, aes(x = experiment, y = mapped, color = experiment)) +
  geom_violin() + 
  geom_boxplot(alpha = .01, width = .2, position = position_dodge(width = .9)) +
  labs(x = "C1 chip", y = "Number of reads",
       title = "Number of mapped sequences per single cell") +
  theme(legend.title = element_blank(),
        axis.text.x = element_text(angle = 45, hjust = 1, vjust = 1))
```

Sum of sequences across the 96 single cells per C1 chip.

```{r datatable}
total_per_experiment <- anno %>%
  group_by(experiment) %>%
  summarize(raw = sum(raw) / 10^6,
            mapped = sum(mapped) / 10^6,
            molecules = sum(molecules) / 10^6)
datatable(total_per_experiment,
          options = list(pageLength = nrow(total_per_experiment)),
          colnames = c("C1 chip", "Number of raw sequences",
                       "Number of mapped",
                       "Number of molecules"))
```


```{r}
ggplot(melt(total_per_experiment, id.vars = "experiment",
            variable.name = "type", value.name = "count"),
       aes(x = experiment, y = count, color = type)) +
  geom_point() +
  labs(title = "Sequencing depth per C1 chip",
       x = "C1 chip", y = "Number of sequences") +
  theme(legend.title = element_blank(),
        axis.text.x = element_text(angle = 45, hjust = 1, vjust = 1))
```
````
