## Supplementary material for "Characterizing and inferring quantitative cell cycle phase in single-cell RNA-seq data analysis": Source code of our analysis, including scripts and data necessary to reproduce the work.: fig2.R

```
# Figure 2
#   Characterizing cell cycle phase using FUCCI fluorescence intensities.


sce <- readRDS("data/sce-final.rds")
sce <- sce[grep("ENSG", rownames(sce)),]
pdata <- data.frame(colData(sce))
fdata <- data.frame(rowData(sce))

sce <- data_transform_quantile(sce)
log2cpm_quantNormed <- assay(sce, "cpm_quantNormed")

# Fig 2A. Derive FUCCI-based cell cycle phase
ints <- data.frame(rfp=colData(sce)$rfp.median.log10sum.adjust,
                   gfp=colData(sce)$gfp.median.log10sum.adjust)
plot(x=ints$rfp, y=ints$gfp, pch=16, cex=.5, xlim=c(-1.5, 1.5), ylim=c(-1.5,1.5),
     xlab="eCherry score", ylab="EGFP score",
     main = "FUCCI scores", col="gray50", axes=F)
axis(1);axis(2)
abline(h=0,v=0, col="gray50", lty=2)
par(new=TRUE)
theta <- coord2rad(scale(ints))
plot(circular(theta), stack=T, shrink=1.3, cex=.5, bins=200)


# Fig 2B. FUCCI phase summarizes fluorescent intensities

# Compute FUCCI phase, do a PCA rotation
ints <- data.frame(rfp=colData(sce)$rfp.median.log10sum.adjust,
                   gfp=colData(sce)$gfp.median.log10sum.adjust)
pca <- prcomp(ints)
theta <- coord2rad(pca$x)
theta_final <- shift_origin(as.numeric(theta), origin = 3*pi/4)

# get PAM-based clusters
pam_res <- pam(ints, k=3)
clust <- data.frame(clust=pam_res$clustering,
                    sample_id=rownames(colData(sce)))
plot(theta_final, clust$clust)
b1 <- mean(max(range(theta_final[clust$clust==2])), min(range(theta_final[clust$clust==3])))
b2 <- mean(max(range(theta_final[clust$clust==3])), max(range(theta_final[clust$clust==1])))
b3 <- mean(min(range(theta_final[clust$clust==2])), min(range(theta_final[clust$clust==1])))
abline(v=c(b1,b2,b3), lty=2)
# clust 1 = G1
# clust 2 = S
# clust 3 = G2/M

par(mfrow=c(1,1))
plot(x=theta_final,
     y=ints$gfp, col="forestgreen",
     pch=c(16,1,4)[clust$clust],
     ylim=c(-1.5, 1.5), cex=.5,
     xlab="FUCCI phase", ylab="FUCCI scores",
     main="FUCCI scores", axes=F)
axis(2); axis(1,at=c(0,pi/2, pi, 3*pi/2, 2*pi),
              labels=c(0,expression(pi/2), expression(pi), expression(3*pi/2),
                       expression(2*pi)))
abline(h=0, lty=1, col="black", lwd=.7)
points(x=theta_final,
       y=ints$rfp, col="firebrick",
       ylim=c(-1.5, 1.5),
       pch=c(16,1,4)[clust$clust],
       cex=.5)
abline(v=c(b1,b2,b3), lty=2)


# Fig 2C. FUCCI phase summarizes fluorescent intensities
# Note:
#   - For details of how we estimated cyclic trend of gene expression levels for each gene.
#     See https://jhsiao999.github.io/peco-paper/npreg_trendfilter_quantile.html. This file
#     makes "data/fit.quant.rds"
fits_all <- readRDS("data/fit.quant.rds")

# get top 5 cyclic genes
pve_all <- sapply(fits_all, "[[", 3)
pve_all_ord <- pve_all[order(pve_all, decreasing = T)]
gene_ensg <- names(pve_all_ord)[1:5]
gene_symbol <- c("CDK1", "UBE2C", "TOP2A", "HIST1H4E", "HIST1H4C")

sample_ord <- rownames(colData(sce))[order(theta_final)]
data_quant_ord <- log2cpm_quantNormed[,match(sample_ord,colnames(log2cpm_quantNormed))]
fits_tmp <- lapply(1:5, function(i) {
  ii <- which(rownames(data_quant_ord)==genes[i])
  fit_g <- fit_trendfilter_generic(data_quant_ord[ii,])
  fun_g <- approxfun(x=as.numeric(theta_final[order(theta_final)]),
                     y=as.numeric(fit_g$trend.yy), rule=2)
  return(fun_g)
})
names(fits_tmp) <- gene_ensg

par(mfrow=c(1,5), mar=c(2,2,2,1))
for (i in 1:5) {
  plot(x=theta_final[order(theta_final)],
       y=data_quant_ord[rownames(data_quant_ord)==gene_ensg[i],], col="gray50",
       xlab="FUCCI phase",
       ylab="", axes=F, cex=.3, pch=16,
       main = gene_symbol[i])
  lines(x = seq(0, 2*pi, length.out=200),
        y = fits_tmp[[which(names(fits_tmp) == gene_ensg[i])]](seq(0, 2*pi, length.out=200)),
        col=wesanderson::wes_palette("FantasticFox1")[1], lty=1, lwd=2)
  axis(2);
  axis(1,at=c(0,pi/2,pi,3*pi/2,2*pi), labels=c(0,expression(pi/2), expression(pi),
                                               expression(3*pi/2), expression(2*pi)))
  abline(h=0, col="black", lty=1, lwd=.7)
  abline(v=c(b1,b2,b3), lty=2)
}
```
