## Supplementary material for "Characterizing and inferring quantitative cell cycle phase in single-cell RNA-seq data analysis": Source code of our analysis, including scripts and data necessary to reproduce the work.: fig3.R

```
# Figure 3: Inferring cell cycle phase form scRNA-seq data

# Notes
#   - For details of how we applied each method, estimated cyclic trends and
#     computed prediction error, see
#     https://jhsiao999.github.io/peco-paper/eval_on_our_data.html. This
#     file produced "data/fit_diff*.rds".

library(SingleCellExperiment)
library(circular)
library(peco)
library(tidyverse)
library(wesanderson)


sce <- readRDS("data/sce-final.rds")
sce <- sce[grep("ENSG", rownames(sce)),]
fdata <- data.frame(colData(sce))
pdata <- data.frame(rowData(sce))
counts <- data.frame(assay(sce, "counts"))

sce_normed <- data_transform_quantile(sce)
log2cpm_quant <- assay(sce_normed, "cpm_quantNormed")


# Figure 3A. Performance of peco using 2 to 50 genes
# See predict_thinned_data.html for more details

# function to make training/testing data
makedata_supervised <- function(sce, log2cpm_quant,
                                theta) {
  message("Create data/data_training_test folder \n")
  if (!file.exists("data/data_training_test")) { dir.create("data/data_training_test") }
  library(SingleCellExperiment)
  library(peco)
  pdata <- data.frame(colData(sce))
  fdata <- data.frame(rowData(sce))
  counts <- data.frame(assay(sce))
  counts <- counts[grep("ENSG", rownames(counts)), ]
  log2cpm <- t(log2(1+(10^6)*(t(counts)/pdata$molecules)))

  for (ind in unique(pdata$chip_id)) {
    ii_test <- c(1:nrow(pdata))[which(pdata$chip_id == ind)]
    ii_train <- c(1:nrow(pdata))[which(pdata$chip_id != ind)]

    pdata_test <- pdata[ii_test,]
    pdata_train <- pdata[ii_train,]

    log2cpm_quant_test <- log2cpm_quant[,ii_test]
    log2cpm_quant_train <- log2cpm_quant[,ii_train]
    theta <- pdata$theta
    names(theta) <- rownames(pdata)

    log2cpm_test <- log2cpm[,ii_test]
    log2cpm_train <- log2cpm[,ii_train]

    counts_test <- counts[,ii_test]
    counts_train <- counts[,ii_train]

    theta_test <- theta[ii_test]
    theta_train <- theta[ii_train]

    #sig.genes <- readRDS("output/npreg-trendfilter-quantile.Rmd/out.stats.ordered.sig.476.rds")
    data_training <- list(theta_train=theta_train,
                          log2cpm_quant_train=log2cpm_quant_train,
                          log2cpm_train=log2cpm_train,
                          counts_train=counts_train,
                          pdata_train=pdata_train,
                          fdata=fdata)

    data_test <- list(theta_test=theta_test,
                      log2cpm_quant_test=log2cpm_quant_test,
                      log2cpm_test=log2cpm_test,
                      counts_test = counts_test,
                      pdata_test=pdata_test,
                      fdata=fdata)

    saveRDS(data_training,
            file=file.path(paste0("data/data_training_test/ind_",ind,"_data_training.rds")))
    saveRDS(data_test,
            file=file.path(paste0("data/data_training_test/ind_",ind,"_data_test.rds")))
  }
}
makedata_supervised(sce, log2cpm_quant, theta)


# compute prediction error based on 2 to 50 genes

# Note:
#   - For details of how we estimated cyclic trend of gene expression levels for each gene.
#     See https://jhsiao999.github.io/peco-paper/npreg_trendfilter_quantile.html. This file
#     makes "data/fit.quant.rds"
fits_all <- readRDS("data/fit.quant.rds")
genes_all <- names(fits_all)[order(sapply(fits_all,"[[",3), decreasing=T)]

res_unthinned <- do.call(rbind, lapply(seq_along(unique(pdata$chip_id)), function(ind) {
  ind <- unique(pdata$chip_id)[i]
  res_thin_each <- do.call(rbind, lapply(2:50, function(ngenes) {
    data_test <- readRDS(paste0("data/data_training_test/ind_",ind,"_data_test.rds"))
    data_train <- readRDS(paste0("data/data_training_test/ind_",ind,"_data_training.rds"))

    which_genes <- genes_all[1:ngenes]
    fit_train <- cycle_npreg_insample(
      Y = with(data_train,
               log2cpm_quant_train[which(rownames(log2cpm_quant_train) %in% which_genes), ]),
      theta = with(data_train, theta_train),
      method.trend="trendfilter")
    fit_test <- cycle_npreg_outsample(
      Y_test=with(data_test,
                  log2cpm_quant_test[which(rownames(log2cpm_quant_test) %in% which_genes), ]),
      sigma_est=with(fit_train, sigma_est),
      funs_est=with(fit_train, funs_est))

    diff_time <- circ_dist(data_test$theta_test,
                           rotation(data_test$theta_test, fit_test$cell_times_est))
    out <- data.frame(phase_pred_rot=rotation(data_test$theta_test, fit_test$cell_times_est),
                      phase_ref=data_test$theta_test,
                      diff_time=diff_time,
                      ind = ind,
                      ngenes = ngenes)
    return(out)
  }) )
}) )


res_unthinned %>% group_by(ind, ngenes) %>%
  #  filter(ngenes <= 20) %>%
  summarise(diff_mean = mean(diff_time/2/pi),
            diff_se = sd(diff_time/2/pi)/sqrt(length(diff_time/2/pi))) %>%
  ggplot(., aes(x=factor(ngenes), y=diff_mean, group = ind)) +
  #  geom_vline(xintercept=seq(5, 50,5)-1, col="gray90", lty=1) +
  geom_hline(yintercept=seq(.1, .2, .01), col="gray90", lty=1) +
  geom_line(aes(col=ind), lwd=.7) + # ggtitle("thinlog2 = .80") +
  scale_fill_brewer(palette="Dark2") +
  geom_errorbar(aes(ymin=diff_mean-diff_se,
                    ymax=diff_mean+diff_se, col=ind), width=.2, alpha=.5) +
  stat_summary(fun.y=mean,geom="line",lwd=.5, group=1) +
  ylim(0,.3) + geom_hline(yintercept=.25, col="red") +
  labs(color="Test data (Cell line)") +
  xlab("Number of cyclic genes used in peco prediction") +
  ylab("Prediction error (% circle)") +
  scale_x_discrete(breaks=c(2:50),
                   labels=c(rep("",3),5, rep("",4), 10, rep("",4), 15,
                            rep("",4), 20, rep("",4), 25,
                            rep("",4), 30, rep("",4), 35,
                            rep("",4), 40, rep("",4), 45, rep("",4), 50)) +
  ggtitle("Performance in unthinned data")


# Figure 3B. Prediction error in the test data
# For details of our computations of prediction error of each method, see
# https://jhsiao999.github.io/peco-paper/eval_on_our_data.html
diff_cyclone <- readRDS("data/fit_diff_cyclone.rds")
diff_recat <- readRDS("data/fit_diff_recat.rds")
diff_peco <- readRDS("data/fit_diff_peco.rds")
diff_seurat <- readRDS("data/fit_diff_seurat.rds")
diff_oscope <- readRDS("data/fit_diff_oscope.rds")

diff_peco %>% group_by(ind) %>% summarise(mn=mean(diff_time)/2/pi,
                                          sd=sd(diff_time/2/pi)/sqrt(sum(phase_pred_rot>0)))
diff_cyclone %>% group_by(ind) %>% summarise(mn=mean(diff_time)/2/pi,
                                             sd=sd(diff_time/2/pi)/sqrt(sum(phase_pred_rot>0)))
diff_seurat %>% group_by(ind) %>% summarise(mn=mean(diff_time)/2/pi,
                                            sd=sd(diff_time/2/pi)/sqrt(sum(phase_pred_rot>0)))
diff_oscope %>% group_by(ind) %>% summarise(mn=mean(diff_time)/2/pi,
                                            sd=sd(diff_time/2/pi)/sqrt(sum(phase_pred_rot>0)))
diff_recat %>% group_by(ind) %>% summarise(mn=mean(diff_time)/2/pi,
                                           sd=sd(diff_time/2/pi)/sqrt(sum(phase_pred_rot>0)))

diff_all <- rbind(diff_recat,
                  diff_cyclone,
                  diff_peco,
                  diff_seurat,
                  diff_oscope)

# only 3 continuous methods
cols_method <- c(wes_palette("FantasticFox1")[1],
                 wes_palette("Darjeeling1")[2],
                 wes_palette("FantasticFox1")[5])
diff_all$method <- factor(diff_all$method, levels=c("peco", "Oscope", "reCAT"))
diff_all %>% dplyr::filter(method %in% c("peco", "Oscope", "reCAT")) %>%
  group_by(ind, method) %>%
  summarise(diff_mean = mean(diff_time/2/pi),
            diff_se = sd(diff_time/2/pi)/sqrt(length(diff_time/2/pi))) %>%
  ggplot(., aes(x=method,y=diff_mean, col=ind, group=ind)) +
  geom_point(aes(x=method, y=diff_mean), size = .5) +
  geom_hline(yintercept=seq(.1, .25, .01), col="gray90", alpha=.6, lty=1) +
  geom_errorbar(aes(ymin=diff_mean-diff_se,
                    ymax=diff_mean+diff_se, col=ind), width=.2, alpha=.5) +
  #  scale_colour_manual(values = c(as.vector(cols_method))) +
  ylab("Prediction error (% circle)") + xlab("Method") +
  labs(color="Test data (Cell line)") + ylim(0,.3) +
  geom_hline(yintercept = .25, col = "red") + theme_classic() +
  #  facet_wrap(~method) +
  theme(axis.text.x  = element_text(angle=45, vjust=1, hjust=1,size=10))


# Supp Fig 3C

sce <- readRDS("data/sce-final.rds")
sce <- sce[grep("ENSG", rownames(sce)),]
fdata <- data.frame(colData(sce))
fdata <- data.frame(rowData(sce))
counts <- data.frame(assay(sce, "counts"))
sce_normed <- data_transform_quantile(sce)
log2cpm_quant <- assay(sce_normed, "cpm_quantNormed")

labs <- c("CDK1", "UBE2C", "TOP2A", "HIST1H4E", "HIST1H4C")
genes <- c("ENSG00000170312","ENSG00000175063",
           "ENSG00000131747", "ENSG00000198518", "ENSG00000197061")
inds <- "NA18511"

# peco
pred_peco <- lapply(1:length(inds), function(i) {
  ind <- inds[i]
  out <- readRDS(paste0("data/ourdata_peco_",
                        ind, "_top",sprintf("%03d", 5), "genes.rds"))
  fits <- lapply(1:5, function(g) {
    gexp <- with(out$fit_test, Y[g,order(cell_times_est)])
    phase <- shift_origin(with(out$fit_test, cell_times_est[order(cell_times_est)]),
                          origin = 3*pi/4)
    fit_g <- with(out$fit_test, fit_trendfilter_generic(gexp))
    fun_g <- approxfun(x=as.numeric(phase),
                       y=as.numeric(fit_g$trend.yy), rule=2)
    fit_g$phase <- phase
    fit_g$gexp <- gexp
    fit_g$fun_g <- fun_g
    return(fit_g)
  })
  names(fits) <- labs
  return(fits)
})
names(pred_peco) <- inds


# Oscope
# For details of how we applied Oscope and estimated cyclic trends based
# on these two methods, see https://jhsiao999.github.io/peco-paper/eval_on_our_data.html
load("data/ourdata_oscope_366genes.rda")

pred_oscope <- lapply(1:length(inds), function(i) {
  out_oscope_sub <- out_oscope[out_oscope$ind == inds[i],]
  fits <- lapply(1:5, function(g) {
    gindex <- which(rownames(log2cpm_quant) == genes[g])
    fit_g <- data.frame(
      gexp=log2cpm_quant[gindex, match(rownames(out_oscope_sub), colnames(log2cpm_quant))],
      phase=shift_origin(2*pi-out_oscope_sub$phase_pred_rot, origin=3*pi/4))
    fit_g <- fit_g[order(fit_g$phase),]

    fit_trend <- fit_trendfilter_generic(fit_g$gexp)
    fit_g$trend.yy <- fit_trend$trend.yy
    fun_g <- approxfun(x=as.numeric(fit_g$phase),
                       y=as.numeric(fit_g$trend.yy), rule=2)
    fit_out <- list(fit_g=fit_g,
                    fun_g = fun_g)
    return(fit_out)
  })
  names(fits) <- labs
  return(fits)
})
names(pred_oscope) <- inds

# recat
# For details of how we applied Oscope and estimated cyclic trends based
# on these two methods, see https://jhsiao999.github.io/peco-paper/eval_on_our_data.html
load("data/ourdata_recat.rda")
samples_ordered <- rownames(test_exp)[ordIndex]
phase_pred <- seq(0, 2*pi, length.out= length(ordIndex))
names(phase_pred) <- samples_ordered

out_recat <- data.frame(phase_ref=pdata$theta[match(samples_ordered, colnames(sce))])
out_recat$ind <- pdata$chip_id[match(samples_ordered, colnames(sce))]
out_recat$phase_pred_rot <- rotation(ref_var= out_recat$phase_ref, shift_var = phase_pred)
rownames(out_recat) <- samples_ordered

pred_recat <- lapply(1:length(inds), function(i) {
  out_recat_sub <- out_recat[out_recat$ind == inds[i],]
  fits <- lapply(1:5, function(g) {
    gindex <- which(rownames(log2cpm_quant) == genes[g])
    fit_g <- data.frame(
      gexp=log2cpm_quant[gindex, rownames(out_recat_sub)],
      phase=shift_origin(2*pi-out_recat_sub$phase_pred_rot, origin = 2*pi/3))
    fit_g <- fit_g[order(fit_g$phase),]

    fit_trend <- fit_trendfilter_generic(fit_g$gexp)
    fit_g$trend.yy <- fit_trend$trend.yy
    #    fit_g$pve <- fit$pve
    fun_g <- approxfun(x=as.numeric(fit_g$phase),
                       y=as.numeric(fit_g$trend.yy), rule=2)
    fit_out <- list(fit_g=fit_g,
                    pve = fit_trend$pve,
                    fun_g = fun_g)
    return(fit_out)
  })
  names(fits) <- labs
  return(fits)
})
names(pred_recat) <- inds


par(mfrow=c(3,5), mar=c(2,2,2,1))
# peco
res <- pred_peco[[1]]
for (g in 1:5) {
  if (g>1) { ylab <- ""} else { ylab <- "Quantile-normalized expression level"}
  #    if (i==1) { xlab <- "Predicted phase"} else { xlab <- "Inferred phase"}
  res_g <- res[[g]]
  plot(x= res_g$phase,
       y = res_g$gexp, pch = 16, cex=.5, ylab=ylab,
       xlab="Predicted phase",
       main = names(res)[g], axes=F, ylim=c(-3,3))
  abline(h=0, lwd=.5)
  lines(x = seq(0, 2*pi, length.out=200),
        y = res_g$fun_g(seq(0, 2*pi, length.out=200)),
        col = wesanderson::wes_palette("FantasticFox1")[1],
        lty=1, lwd=2)
  axis(2); axis(1,at=c(0,pi/2, pi, 3*pi/2, 2*pi),
                labels=c(0,expression(pi/2), expression(pi), expression(3*pi/2),
                         expression(2*pi)))
}

res <- pred_oscope[[1]]
for (g in 1:5) {
  if (g>1) { ylab <- ""} else { ylab <- "Quantile-normalized expression level"}
  res_g <- res[[g]]
  plot(x= res_g$fit_g$phase,
       y = res_g$fit_g$gexp, pch = 16, cex=.5, ylab=ylab,
       xlab="Predicted phase",
       main = names(res)[g], axes=F, ylim=c(-3,3))
  abline(h=0, lwd=.5)
  lines(x = seq(0, 2*pi, length.out=200),
        y = res_g$fun_g(seq(0, 2*pi, length.out=200)),
        col = wesanderson::wes_palette("Darjeeling2")[2], lty=1, lwd=2)
  axis(2); axis(1,at=c(0,pi/2, pi, 3*pi/2, 2*pi),
                labels=c(0,expression(pi/2), expression(pi), expression(3*pi/2),
                         expression(2*pi)))
}

res <- pred_recat[[1]]
for (g in 1:5) {
  if (g>1) { ylab <- ""} else { ylab <- "Quantile-normalized expression level"}
  res_g <- res[[g]]
  plot(x= res_g$fit_g$phase,
       y = res_g$fit_g$gexp, pch = 16, cex=.5, ylab=ylab,
       xlab="Predicted phase",
       main = names(res)[g], axes=F, ylim=c(-3,3))
  abline(h=0, lwd=.5)
  lines(x = seq(0, 2*pi, length.out=200),
        y = res_g$fun_g(seq(0, 2*pi, length.out=200)),
        col = wesanderson::wes_palette("FantasticFox1")[5], lty=1, lwd=2)
  axis(2); axis(1,at=c(0,pi/2, pi, 3*pi/2, 2*pi),
                labels=c(0,expression(pi/2), expression(pi), expression(3*pi/2),
                         expression(2*pi)))
}
```
