## Supplementary material for "Characterizing and inferring quantitative cell cycle phase in single-cell RNA-seq data analysis": Source code of our analysis, including scripts and data necessary to reproduce the work.: suppfig02.R

```
# Supplemental Figure S2
# Authored by PoYuan Tung

library("cowplot")
library("dplyr")
library("ggplot2")
library("MASS")
library("SingleCellExperiment")

# The palette with grey:
cbPalette <- c("#999999", "#E69F00", "#56B4E9", "#009E73", "#F0E442", "#0072B2", "#D55E00", "#CC79A7")
indi_palette <- c("tomato4", "chocolate1", "rosybrown", "purple4", "plum3", "slateblue")


sce_raw <- readRDS("data/sce-raw.rds")
anno <- data.frame(colData(sce_raw))


## Total mapped reads
## Note: Using the 20 % cutoff of samples with no cells excludes all the samples

## calculate the cut-off
cut_off_reads <- quantile(anno[anno$cell_number == 0,"mapped"], 0.82)
cut_off_reads

anno$cut_off_reads <- anno$mapped > cut_off_reads

## numbers of cells
sum(anno[anno$cell_number == 1, "mapped"] > cut_off_reads)
sum(anno[anno$cell_number == 1, "mapped"] <= cut_off_reads)

## density plots
plot_reads <- ggplot(anno[anno$cell_number == 0 |
                            anno$cell_number == 1 , ],
                     aes(x = mapped, fill = as.factor(cell_number))) +
  geom_density(alpha = 0.5) +
  geom_vline(xintercept = cut_off_reads, colour="grey", linetype = "longdash") +
  labs(x = "Total mapped reads", title = "Number of total mapped reads", fill = "Cell number")


## Unmapped ratios
# Using the 40 % cutoff of samples with no cells excludes all the samples

## calculate unmapped ratios
anno$unmapped_ratios <- anno$unmapped/anno$umi

## cut off
cut_off_unmapped <- quantile(anno[anno$cell_number == 0,"unmapped_ratios"], 0.40)
cut_off_unmapped

anno$cut_off_unmapped <- anno$unmapped_ratios < cut_off_unmapped

## numbers of cells
sum(anno[anno$cell_number == 1, "unmapped_ratios"] >= cut_off_unmapped)

sum(anno[anno$cell_number == 1, "unmapped_ratios"] < cut_off_unmapped)

## density plots
plot_unmapped <- ggplot(anno[anno$cell_number == 0 |
                               anno$cell_number == 1 , ],
                        aes(x = unmapped_ratios *100, fill = as.factor(cell_number))) +
  geom_density(alpha = 0.5) +
  geom_vline(xintercept = cut_off_unmapped *100, colour="grey", linetype = "longdash") +
  labs(x = "Unmapped reads/ total reads", title = "Unmapped reads percentage")


## ERCC percentage

## calculate ercc reads percentage
anno$ercc_percentage <- anno$reads_ercc / anno$mapped

## cut off
cut_off_ercc <- quantile(anno[anno$cell_number == 0,"ercc_percentage"], 0.20)
cut_off_ercc

anno$cut_off_ercc <- anno$ercc_percentage < cut_off_ercc

## numbers of cells
sum(anno[anno$cell_number == 1, "ercc_percentage"] >= cut_off_ercc)

sum(anno[anno$cell_number == 1, "ercc_percentage"] < cut_off_ercc)

## density plots
plot_ercc <- ggplot(anno[anno$cell_number == 0 |
                           anno$cell_number == 1 , ],
                    aes(x = ercc_percentage *100, fill = as.factor(cell_number))) +
  geom_density(alpha = 0.5) +
  geom_vline(xintercept = cut_off_ercc *100, colour="grey", linetype = "longdash") +
  labs(x = "ERCC reads / total mapped reads", title = "ERCC reads percentage")


## Number of genes detected
## cut off
cut_off_genes <- quantile(anno[anno$cell_number == 0,"detect_hs"], 0.80)
cut_off_genes

anno$cut_off_genes <- anno$detect_hs > cut_off_genes

## numbers of cells
sum(anno[anno$cell_number == 1, "detect_hs"] > cut_off_genes)
sum(anno[anno$cell_number == 1, "detect_hs"] <= cut_off_genes)

## density plots
plot_gene <- ggplot(anno[anno$cell_number == 0 |
                           anno$cell_number == 1 , ],
                    aes(x = detect_hs, fill = as.factor(cell_number))) +
  geom_density(alpha = 0.5) +
  geom_vline(xintercept = cut_off_genes, colour="grey", linetype = "longdash") +
  labs(x = "Gene numbers", title = "Numbers of detected genes")


plot_grid(plot_reads, plot_unmapped,
          plot_ercc, plot_gene, labels = LETTERS[1:4], ncol = 2)
```
