## Supplementary material for "Characterizing and inferring quantitative cell cycle phase in single-cell RNA-seq data analysis": Source code of our analysis, including scripts and data necessary to reproduce the work.: suppfig03.R

```
# Supplemental Figure 3
# RNA-seq data quality, PCA, etc ---------------------------------------------------------

library(SingleCellExperiment)
library(peco)
sce_final <- readRDS("data/sce-final.rds")
sce_final <- sce_final[grep("ENSG", rownames(sce_final)),]

counts <- assay(sce_final)
pdata <- data.frame(colData(sce_final))


# drop-out rate, number of singletons
log2cpm.all <- t(log2(1+(10^6)*(t(counts)/pdata$molecules)))
mean(rowMeans(log2cpm.all>0))
sd(rowMeans(log2cpm.all>0))

inds <- unique(pdata$chip_id)
cbind(inds, sapply(1:length(inds), function(i) {
  tmp <- log2cpm.all[,pdata$chip_id == inds[i]]
  mean(rowMeans(tmp>0)) }))

pdata$dropout <- colMeans(log2cpm.all >0)
pdata$experiment <- factor(pdata$experiment)
pdata$chip_id <- factor(pdata$chip_id)

# compare dropout rate between individuals
kruskal.test(pdata$dropout ~ pdata$chip_id)
kruskal.test(log10(pdata$molecules) ~ pdata$chip_id)

# compare dropout rate between C1 plates
kruskal.test(pdata$dropout ~ pdata$experiment)
kruskal.test(log10(pdata$molecules) ~ pdata$experiment)

dd <- table(pdata$experiment)/96
dd <- as.numeric(dd)
mean(dd)
sd(dd)
range(dd)
table(pdata$chip_id)

# Result section on sequencing reads
# prior to filtering
sce_raw = readRDS("data/sce-raw.rds")
pdata = data.frame(colData(sce_raw))
mean(pdata$mapped)
sd(pdata$mapped)
range(pdata$mapped)


# sequencing results by individual, including
# all data before filtering of samples or genes
# prior to filtering
sce_final <- readRDS("data/sce-final.rds")
pdata <- data.frame(colData(sce_final))
inds <- unique(pdata$chip_id)

# number of reads with a valid UMI and also mapped to the index
# prior to filtering
do.call(rbind, lapply(1:length(inds), function(i) {
  tmp <- pdata$mapped[pdata$chip_id == inds[i]]
  data.frame(ind=inds[i],
             mn=mean(tmp),
             std=sd(tmp), stringsAsFactors = F) }))
cbind(mean(pdata$mapped), sd(pdata$mapped))


# proportin of unmapped reads with a valid UMI
# prior to filtering
percents_unmapped <- pdata$unmapped/pdata$umi
do.call(rbind, lapply(1:length(inds), function(i) {
  tmp <- percents_unmapped[pdata$chip_id == inds[i]]
  data.frame(ind=inds[i],
             mn=mean(tmp)*100,
             std=sd(tmp)*100, stringsAsFactors = F) }))
cbind(mean(percents_unmapped)*100, sd(percents_unmapped)*100)


# number of singletons
table(pdata$chip_id)
length(pdata$chip_id)


# fraction of ERCC reads: reads_ercc/mapped
percentage_ercc <- pdata$ercc_percentage
do.call(rbind, lapply(1:length(inds), function(i) {
  tmp <- percentage_ercc[pdata$chip_id == inds[i]]
  data.frame(ind=inds[i],
             mn=mean(tmp)*100,
             std=sd(tmp)*100, stringsAsFactors = F) }))
cbind(mean(percentage_ercc)*100, sd(percentage_ercc)*100)


# total molecular count
molecules <- pdata$molecules
do.call(rbind, lapply(1:length(inds), function(i) {
  tmp <- molecules[pdata$chip_id == inds[i]]
  data.frame(ind=inds[i],
             mn=mean(tmp)/(10^6),
             std=sd(tmp)/(10^6), stringsAsFactors = F) }))
cbind(mean(molecules)/(10^6), sd(molecules)/(10^6))
```
