## Supplementary material for "Characterizing and inferring quantitative cell cycle phase in single-cell RNA-seq data analysis": Source code of our analysis, including scripts and data necessary to reproduce the work.: suppfig04.R

```
# Supplemental Figure 4
# Quality metrics in individual cell lines

library(ggplot2)
library(cowplot)

sce <- readRDS("data/sce-final.rds")
sce <- sce[grep("ENSG", rownames(sce)), ]

counts <- assay(sce)
pdata <- data.frame(colData(sce))

log2cpm.all <- t(log2(1+(10^6)*(t(counts)/pdata$molecules)))

pdata$dropout <- colMeans(log2cpm.all >0)
pdata$experiment <- factor(pdata$experiment)
pdata$chip_id <- factor(pdata$chip_id)

plot_grid(ggplot(pdata, aes(x=chip_id, y=log10(molecules),
                  group=chip_id, col=chip_id)) +
  geom_boxplot() + coord_flip() +
  ylab("log10 molecule count") + xlab("Individual") +
  scale_color_brewer(palette="Dark2") +
  ggtitle("Library size and individual") + theme(legend.position="none"),
ggplot(pdata, aes(x=chip_id, y=dropout,
                group=chip_id, col=chip_id)) +
  geom_boxplot() + coord_flip() +
  ylab("Sample detection rate") + xlab("Individual") +
  scale_color_brewer(palette="Dark2") +
  ggtitle("Detection rate and individual") + theme(legend.position="none"),
ggplot(pdata, aes(x=chip_id, fill=chip_id)) +
  geom_bar() + coord_flip() +
  ylab("Singletons") + xlab("Individual") +
  scale_fill_brewer("Individual",palette="Dark2") +
  ggtitle("Singletons and individual") + theme(legend.position="none"),
ncol = 2, labels = LETTERS[1:3])
```
