## Supplementary material for "Characterizing and inferring quantitative cell cycle phase in single-cell RNA-seq data analysis": Source code of our analysis, including scripts and data necessary to reproduce the work.: suppfig05.R

```
# Supplemental Figure S5
#   Major sources of variation in our gene expression data of 888 quality samples and 11,040 genes.

library(cowplot)
library(dplyr)
library(edgeR)
library(ggplot2)
library(heatmap3)
library(reshape2)
library(SingleCellExperiment)
library(heatmap3)
library(RColorBrewer)

# load code to apply PCA to gene expression data
source("code/utility.R")


sce_filtered <- readRDS("data/sce-filtered.rds")
counts <- assay(sce_filtered)
pdata <- data.frame(colData(sce_filtered))
pdata$experiment <- as.factor(pdata$experiment)

# Compute log2 CPM based on the library size before filtering.
log2cpm <- edgeR::cpm(counts, log = TRUE)
pca_log2cpm <- run_pca(log2cpm)


# selection of technical factor
covariates <- pdata %>% dplyr::select(experiment, well, chip_id,
                                            concentration, raw:unmapped,
                                            starts_with("detect"),  molecules)


# Supp Fig S5A: Association between Principal Components of gene e
#   expression data and technical factors.

# R-square between PCs and the covariates
get_r2 <- function(x, y) {
  stopifnot(length(x) == length(y))
  model <- lm(y ~ x)
  stats <- summary(model)
  return(stats$adj.r.squared)
}

# look at the first 6 PCs
pca_log2cpm <- prcomp(t(log2cpm), scale. = TRUE, center = TRUE)
pcs <- pca_log2cpm$x[, 1:6]

r2 <- matrix(NA, nrow = ncol(covariates), ncol = ncol(pcs),
             dimnames = list(colnames(covariates), colnames(pcs)))
for (cov in colnames(covariates)) {
  for (pc in colnames(pcs)) {
    r2[cov, pc] <- get_r2(covariates[, cov], pcs[, pc])
  }
}

heatmap3(r2, cexRow=1, cexCol=1, margins=c(4,15), scale="none",
         ylab="", main = "", symm=F,
         Colv=NA, showColDendro = F,
         labRow = c("C1: C1 batch",
                    "Well: C1 capture site (well) ID",
                    "Individual",
                    "cDNA: cDNA concentration",
                    "Reads raw: Raw reads",
                    "Reads UMI: Reads with valid UMI",
                    "Reads mapped: Reads with valid UMI mapped to genome",
                    "Reads unmapped: Reads with valid UMI not mapped to genome",
                    "ERCC: ERCC gene proportion detected",
                    "ENSG: Endogeneous gene proportion detected",
                    "Molecules: Total sample molecule count"
         ))

# r-squared between PC1 and individual
f1 <- lm(pcs[,1] ~ covariates$chip_id)
f2 <- lm(pcs[,1] ~ covariates$molecules)
summary(f1)

summary(f1)$adj.r.squared
summary(f2)$adj.r.squared


# PC proportions
100*((pca_log2cpm$sdev^2)/sum(pca_log2cpm$sdev^2))[1:6]


# Supp Fig S5B: Correlation between Technical factors
cor_tech <- cor(as.matrix(covariates[,4:11]),use="pairwise.complete.obs")
heatmap3(cor_tech, symm = TRUE, margins=c(7,15),
         col=brewer.pal (9, "Blues" ), cexRow=1, cexCol=1, scale="none",
         labRow = c("cDNA: cDNA concentration",
                    "Reads raw: Raw reads",
                    "Reads UMI: Reads with valid UMI",
                    "Reads mapped: Reads with valid UMI mapped to genome",
                    "Reads unmapped: Reads with valid UMI not mapped to genome",
                    "ERCC: ERCC gene proportion detected",
                    "ENSG: Endogeneous gene proportion detected",
                    "Molecules: Total sample molecule count"),
         labCol = c("cDNA",
                    "Reads raw",
                    "Reads UMI",
                    "Reads mapped",
                    "Reads unmapped",
                    "ERCC",
                    "ENSG",
                    "Molecules"))

summary(lm(covariates$detect_hs ~ covariates$experiment))

summary(lm(covariates$detect_hs ~ covariates$mapped))

summary(lm(covariates$detect_hs ~ covariates$chip_id))

summary(lm(covariates$detect_hs ~ covariates$molecules))

summary(lm(covariates$detect_hs ~ covariates$concentration))
```
