## Supplementary material for "Characterizing and inferring quantitative cell cycle phase in single-cell RNA-seq data analysis": Source code of our analysis, including scripts and data necessary to reproduce the work.: suppfig06.R

```
# Supplemental Figure 6
#   FUCCI scores for the sixteen C1 plates before and after correcting for C1 plate effect.

library(ggplot2)
library(RColorBrewer)
library(cowplot)


sce <- readRDS("data/sce-final.rds")
counts <- assay(sce)
counts <- counts[grep("ENSG", rownames(counts)), ]
pdata <- data.frame(colData(sce))
pdata$experiment <- factor(pdata$experiment)


# Intensity batch effect ---------------------------------------------------------
lm.rfp <- lm(rfp.median.log10sum~factor(chip_id)+factor(experiment),
             data = pdata)
lm.gfp <- lm(gfp.median.log10sum~factor(chip_id)+factor(experiment),
             data = pdata)
lm.dapi <- lm(dapi.median.log10sum~factor(chip_id)+factor(experiment),
              data = pdata)

library(car)
aov.lm.rfp <- Anova(lm.rfp, type = "III")
aov.lm.gfp <- Anova(lm.gfp, type = "III")
aov.lm.dapi <- Anova(lm.dapi, type = "III")

aov.lm.rfp
aov.lm.gfp

plot_grid(
  ggplot(pdata,
         aes(x=experiment, y=gfp.median.log10sum,
             fill=experiment)) +
    geom_violin(alpha=.5) + geom_boxplot(width=.1, fill="gray90") +
    ylab("FUCC score") + xlab("Individual") +
    ggtitle("GFP before batch-correction") + labs(fill="experiment") +
    theme(axis.text.x  = element_text(angle=90, vjust=1, hjust=1,size=10),
          axis.text.y  = element_text(angle=0, vjust=0.2, size=10)) +
    theme(legend.position="none"),
  ggplot(pdata,
         aes(x=experiment, y=rfp.median.log10sum, fill=experiment)) +
    geom_violin(alpha=.5) + geom_boxplot(width=.1, fill="gray90") +
    ylab("FUCC score") + xlab("Individual") +
    ggtitle("RFP before batch-correction") + labs(fill="experiment") +
    theme(axis.text.x  = element_text(angle=90, vjust=1, hjust=1,size=10),
          axis.text.y  = element_text(angle=0, vjust=0.2, size=10)) +
    theme(legend.position="none"),
  ggplot(pdata,
         aes(x=experiment, y=gfp.median.log10sum.adjust, fill=experiment)) +
    geom_violin(alpha=.5) + geom_boxplot(width=.1, fill="gray90") +
    ylab("FUCC score") + xlab("Individual") +
    ggtitle("GFP after batch-correction") + labs(fill="experiment") +
    theme(axis.text.x  = element_text(angle=90, vjust=1, hjust=1,size=10),
          axis.text.y  = element_text(angle=0, vjust=0.2, size=10)) +
    theme(legend.position="none"),
  ggplot(pdata,
         aes(x=experiment, y=rfp.median.log10sum.adjust, fill=experiment)) +
    geom_violin(alpha=.5) + geom_boxplot(width=.1, fill="gray90") +
    ylab("FUCC score") + xlab("Individual") +
    ggtitle("RFP after batch-correction") + labs(fill="experiment") +
    theme(axis.text.x  = element_text(angle=90, vjust=1, hjust=1,size=10),
          axis.text.y  = element_text(angle=0, vjust=0.2, size=10)) +
    theme(legend.position="none"),
  labels=LETTERS[1:4])
```
