## Supplementary material for "Characterizing and inferring quantitative cell cycle phase in single-cell RNA-seq data analysis": Source code of our analysis, including scripts and data necessary to reproduce the work.: suppfig08.R

```
# Supplemental Figure 8

library(SingleCellExperiment)
library(circular)
library(peco)
library(cluster)
library(ggplot2)

sce <- readRDS("data/sce-final.rds")
sce <- sce[grep("ENSG", rownames(sce)),]
pdata <- data.frame(colData(sce))
fdata <- data.frame(rowData(sce))

sce <- data_transform_quantile(sce)
log2cpm_quantNormed <- assay(sce, "cpm_quantNormed")

# Figure S8A: PAM-based classification and FUCCI scores

# derive cell cycle phase
# use PCA to rotate the axes
pca <- prcomp(cbind(pdata$rfp.median.log10sum.adjust,
                    pdata$gfp.median.log10sum.adjust))
theta <- coord2rad(pca$x)
theta_final <- shift_origin(as.numeric(theta), 3*pi/4)

# get PAM-based clusters
pam_res <- pam(cbind(pdata$rfp.median.log10sum.adjust,
                     pdata$gfp.median.log10sum.adjust), k=3)
clust <- data.frame(clust=pam_res$clustering,
                    sample_id=rownames(pdata))

# check cluster boundaries
plot(theta_final, clust$clust)

ggplot(data.frame(gfp=pdata$gfp.median.log10sum.adjust,
                  rfp=pdata$rfp.median.log10sum.adjust,
                  clust=factor(clust$clust)), aes(x=gfp, y=rfp, shape=clust)) +
  geom_point() + theme_classic() +
  xlab("EGFP score") +
  ylab("mCherry score") +
  scale_shape_manual(values=c(16,1,4),
                     labels = c("G1", "S", "G2M"),
                     name = "") +
  ggtitle("")


# Figure S8B:
# PAM-based gene classification vs Whitfield et al. 2012 classification

# check all cell cycle phase markers in Whitfield et al.
sample_ord <- rownames(pdata)[order(theta_final)]
data_quant_ord <- log2cpm_quantNormed[,match(sample_ord,colnames(log2cpm_quantNormed))]

# import the list of cell cycle genes in Whitfeld et al. 2012 (10.1091/mbc.02-02-0030)
# the Whitfeld et al. 2012 list is annotated and reported in Macosko et al. 2015
# data was downloaded from Macosko et al. 2015 (10.1016/j.cell.2015.05.002)
# https://www.ncbi.nlm.nih.gov/pmc/articles/PMC4481139/bin/NIHMS687993-supplement-supp_data_2.xlsx
macosko <- readRDS("data/macosko-2015.rds")

subdata_plot = do.call(rbind, lapply(1:nrow(macosko), function(g) {
  gindex = which(rownames(data_quant_ord) == macosko$ensembl[g])
  if (length(gindex) != 0 ) {
    gexp = data_quant_ord[gindex,]
    tmp = data.frame(gexp=gexp,
                     ensembl=macosko$ensembl[g], hgnc = macosko$hgnc[g],
                     cell_state=as.character(clust$clust)[match(colnames(data_quant_ord),
                                                                as.character(clust$sample_id))])
    tmp2 = data.frame(max_state = which.max(sapply(1:3, function(i) mean(tmp$gexp[tmp$cell_state==i]))),
                      ensembl=macosko$ensembl[g], hgnc = macosko$hgnc[g])
    return(tmp2)
  }
}))
subdata_plot$max_state = factor(subdata_plot$max_state,
                                labels=c("G1", "S", "G2M"))
macosko_sub = macosko[match(subdata_plot$ensembl, macosko$ensembl),]
table(subdata_plot$max_state, macosko_sub$phase)

table(subdata_plot$max_state,
      macosko_sub$phase)

# compute mis-classfication rate
# G1
(39+13+27)/(53+39+13+27+22)
# S
(14+24+25)/(8+14+14+24+25)
# G2M
(26+51)/(26+51+99+91+53)
```
