## Supplementary material for "Characterizing and inferring quantitative cell cycle phase in single-cell RNA-seq data analysis": Source code of our analysis, including scripts and data necessary to reproduce the work.: suppfig09.R

```
# Supplemental Figure 9A-C
#   Cyclic trends of gene expression based on FUCCI phase in 54 genes,
# which is a subset of the top 91 cyclic genes that is also known to be cell
# cycle genes in Macosko et al. 2015.

# Notes
#   - For details of how we estimate cycic trends, see
#     https://jhsiao999.github.io/peco-paper/npreg_trendfilter_quantile.html

library(SingleCellExperiment)
library(circular)
library(peco)
library(cluster)
library(ggplot2)

sce <- readRDS("data/sce-final.rds")
sce <- sce[grep("ENSG", rownames(sce)),]
pdata <- data.frame(colData(sce))
fdata <- data.frame(rowData(sce))
counts <- data.frame(assay(sce, "counts"))

sce_normed <- data_transform_quantile(sce)
log2cpm_quant <- assay(sce_normed, "cpm_quantNormed")

# derive cell cycle phase
# use PCA to rotate the axes
pca <- prcomp(cbind(pdata$rfp.median.log10sum.adjust,
                    pdata$gfp.median.log10sum.adjust))

plot(pca$x[,1], pca$x[,2], pch=16, cex=.5, xlim=c(-4, 4), ylim=c(-4,4),
     xlab="PC1 (67%)", ylab="PC2 (33%)",
     main = "fucci intensities PC1 vs PC2", col="gray50", axes=F)
axis(1);axis(2)
abline(h=0,v=0, col="gray50", lty=2)
par(new=TRUE)
theta <- coord2rad(pca$x)
plot(circular(theta), stack=T, shrink=1.3, cex=.5, bins=200)
theta_final <- shift_origin(as.numeric(theta), 3*pi/4)

# get PAM-based clusters
pam_res <- pam(cbind(pdata$rfp.median.log10sum.adjust,
                     pdata$gfp.median.log10sum.adjust), k=3)
clust <- data.frame(clust=pam_res$clustering,
                    sample_id=rownames(pdata))
plot(theta_final, clust$clust)
b1 <- mean(max(range(theta_final[clust$clust==2])), min(range(theta_final[clust$clust==3])))
b2 <- mean(max(range(theta_final[clust$clust==3])), max(range(theta_final[clust$clust==1])))
b3 <- mean(min(range(theta_final[clust$clust==2])), min(range(theta_final[clust$clust==1])))
abline(v=c(b1,b2,b3), lty=2)
# clust 1 = G1
# clust 2 = S
# clust 3 = G2/M


# check all cell cycle phase markers in Whitfield et al.

# import the list of cell cycle genes in Whitfeld et al. 2012 (10.1091/mbc.02-02-0030)
# the Whitfeld et al. 2012 list is annotated and reported in Macosko et al. 2015
# data was downloaded from Macosko et al. 2015 (10.1016/j.cell.2015.05.002)
# https://www.ncbi.nlm.nih.gov/pmc/articles/PMC4481139/bin/NIHMS687993-supplement-supp_data_2.xlsx
macosko <- readRDS("data/macosko-2015.rds")

# import quantile-normalized data
data_quant <- readRDS("data/log2cpm.quant.rds")
sample_ord <- rownames(pdata)[order(theta_final)]
data_quant_ord <- log2cpm_quant[,match(sample_ord,colnames(log2cpm_quant))]

# Get all significant cyclic genes assigend as cell cycle genes.
# Specifically, get significant cyclic genes that are also classifed as cell cycle genes
# permutation-based PVE.
#   - For details of how we estimate cycic trends, see
#     https://jhsiao999.github.io/peco-paper/npreg_trendfilter_quantile.html

perm.lowmiss <- readRDS("data/fit.trend.perm.lowmiss.rds")
pve.perm.lowmiss <- sapply(perm.lowmiss, "[[", "trend.pve")
fit.quant <- readRDS("data/fit.quant.rds")
pval_peco <- data.frame(pve=sapply(fit.quant, "[[", "trend.pve"),
                        pval_perm = sapply(fit.quant, function(x) (1+sum(pve.perm.lowmiss > x$trend.pve))/(1+ length(pve.perm.lowmiss)) ),
                        row.names = fdata$name[match(names(fit.quant),
                                                     rownames(fdata))])
pval_peco_ordered <- pval_peco[order(pval_peco$pve, decreasing = T),]


# choose cyclic genes that are also cell cycle genes in Whitfield et al. 2012
metrics <- pval_peco_ordered$pval_perm
cutoff <- .001
cyclegenes <- toupper(macosko$hgnc)
allgenes <- toupper(rownames(pval_peco_ordered))

which_sig_cycle <- allgenes[metrics < cutoff][allgenes[metrics < cutoff] %in% cyclegenes]
which_sig_cycle_genes <- data.frame(names=which_sig_cycle,
                                    ensg=rownames(fdata)[match(which_sig_cycle, fdata$name)])

fits_tmp <- lapply(seq_len(nrow(which_sig_cycle_genes)), function(i) {
  ii <- which(rownames(data_quant_ord)==which_sig_cycle_genes$ensg[i])
  fit_g <- fit_trendfilter_generic(data_quant_ord[ii,])
  fun_g <- approxfun(x=as.numeric(theta_final[order(theta_final)]),
                     y=as.numeric(fit_g$trend.yy), rule=2)
  return(fun_g)
})
names(fits_tmp) <- which_sig_cycle_genes$names


# Supp Fig 9A
par(mfrow=c(4,5), mar=c(2,2,2,1))
for (i in 1:20) {
  gene_name = which_sig_cycle_genes$names[i]
  plot(x=theta_final[order(theta_final)],
       y=data_quant_ord[rownames(data_quant_ord)==which_sig_cycle_genes$ensg[i],],
       col="gray50",
       xlab="FUCCI phase",
       ylab="", axes=F, cex=.3, pch=16,
       main = paste(which_sig_cycle_genes$names[i], paste0("(", i ,")")))
  lines(x = seq(0, 2*pi, length.out=200),
        y = fits_tmp[[which(names(fits_tmp) == which_sig_cycle_genes$names[i])]](seq(0, 2*pi, length.out=200)),
        col=wesanderson::wes_palette("FantasticFox1")[1], lty=1, lwd=2)
  axis(2);
  axis(1,at=c(0,pi/2,pi,3*pi/2,2*pi), labels=c(0,expression(pi/2), expression(pi),
                                               expression(3*pi/2), expression(2*pi)))
  abline(h=0, col="black", lty=1, lwd=.7)
  abline(v=c(b1,b2,b3), lty=2)
}

# Supp Fig 9B
par(mfrow=c(4,5), mar=c(2,2,2,1))
for (i in 21:40) {
  plot(x=theta_final[order(theta_final)],
       y=data_quant_ord[rownames(data_quant_ord)==which_sig_cycle_genes$ensg[i],],
       col="gray50",
       xlab="FUCCI phase",
       ylab="", axes=F, cex=.3, pch=16,
       main = paste(which_sig_cycle_genes$names[i], paste0("(", i ,")")))
  lines(x = seq(0, 2*pi, length.out=200),
        y = fits_tmp[[which(names(fits_tmp) == which_sig_cycle_genes$names[i])]](seq(0, 2*pi, length.out=200)),
        col=wesanderson::wes_palette("FantasticFox1")[1], lty=1, lwd=2)
  axis(2);
  axis(1,at=c(0,pi/2,pi,3*pi/2,2*pi), labels=c(0,expression(pi/2), expression(pi),
                                               expression(3*pi/2), expression(2*pi)))
  abline(h=0, col="black", lty=1, lwd=.7)
  abline(v=c(b1,b2,b3), lty=2)
}

# Supp Fig 9C
par(mfrow=c(4,5), mar=c(2,2,2,1))
for (i in 41:54) {
  plot(x=theta_final[order(theta_final)],
       y=data_quant_ord[rownames(data_quant_ord)==which_sig_cycle_genes$ensg[i],],
       col="gray50",
       xlab="FUCCI phase",
       ylab="", axes=F, cex=.3, pch=16,
       main = paste(which_sig_cycle_genes$names[i], paste0("(", i ,")")))
  lines(x = seq(0, 2*pi, length.out=200),
        y = fits_tmp[[which(names(fits_tmp) == which_sig_cycle_genes$names[i])]](seq(0, 2*pi, length.out=200)),
        col=wesanderson::wes_palette("FantasticFox1")[1], lty=1, lwd=2)
  axis(2);
  axis(1,at=c(0,pi/2,pi,3*pi/2,2*pi), labels=c(0,expression(pi/2), expression(pi),
                                               expression(3*pi/2), expression(2*pi)))
  abline(h=0, col="black", lty=1, lwd=.7)
  abline(v=c(b1,b2,b3), lty=2)
}
```
