## Supplementary material for "Characterizing and inferring quantitative cell cycle phase in single-cell RNA-seq data analysis": Source code of our analysis, including scripts and data necessary to reproduce the work.: suppfig10.R

```
# Supplemental Fig S10
#   Performance of peco in unthinned and thinned data.

# Note:
#   - For more details of how we thinned the data and computed prediction error in
#     thinned and unthinned data, see https://jhsiao999.github.io/peco-paper/predict_thinned_data.html.

# First, prepare test/training data
sce <- readRDS("data/sce-final.rds")
sce <- sce[grep("ENSG", rownames(sce)),]
fdata <- data.frame(colData(sce))
fdata <- data.frame(rowData(sce))
counts <- data.frame(assay(sce, "counts"))

sce_normed <- data_transform_quantile(sce)
log2cpm_quant <- assay(sce_normed, "cpm_quantNormed")

inds <- c("NA18511", "NA18855", "NA18870", "NA19098", "NA19101", "NA19160")
theta <- pdata$theta

# function to make training/testing data
makedata_supervised <- function(sce, log2cpm_quant,
                                theta) {
  message("Create data/data_training_test folder \n")
  if (!file.exists("data/data_training_test")) { dir.create("data/data_training_test") }
  library(SingleCellExperiment)
  library(peco)
  pdata <- data.frame(colData(sce))
  fdata <- data.frame(rowData(sce))
  counts <- data.frame(assay(sce))
  counts <- counts[grep("ENSG", rownames(counts)), ]
  log2cpm <- t(log2(1+(10^6)*(t(counts)/pdata$molecules)))

  for (ind in unique(pdata$chip_id)) {
    ii_test <- c(1:nrow(pdata))[which(pdata$chip_id == ind)]
    ii_train <- c(1:nrow(pdata))[which(pdata$chip_id != ind)]

    pdata_test <- pdata[ii_test,]
    pdata_train <- pdata[ii_train,]

    log2cpm_quant_test <- log2cpm_quant[,ii_test]
    log2cpm_quant_train <- log2cpm_quant[,ii_train]
    theta <- pdata$theta
    names(theta) <- rownames(pdata)

    log2cpm_test <- log2cpm[,ii_test]
    log2cpm_train <- log2cpm[,ii_train]

    counts_test <- counts[,ii_test]
    counts_train <- counts[,ii_train]

    theta_test <- theta[ii_test]
    theta_train <- theta[ii_train]

    #sig.genes <- readRDS("output/npreg-trendfilter-quantile.Rmd/out.stats.ordered.sig.476.rds")
    data_training <- list(theta_train=theta_train,
                          log2cpm_quant_train=log2cpm_quant_train,
                          log2cpm_train=log2cpm_train,
                          counts_train=counts_train,
                          pdata_train=pdata_train,
                          fdata=fdata)

    data_test <- list(theta_test=theta_test,
                      log2cpm_quant_test=log2cpm_quant_test,
                      log2cpm_test=log2cpm_test,
                      counts_test = counts_test,
                      pdata_test=pdata_test,
                      fdata=fdata)

    saveRDS(data_training,
            file=file.path(paste0("data/data_training_test/ind_",ind,"_data_training.rds")))
    saveRDS(data_test,
            file=file.path(paste0("data/data_training_test/ind_",ind,"_data_test.rds")))
  }
}
makedata_supervised(sce, log2cpm_quant, theta)


# Supp Fig 11A

# Note:
#   - For details of how we estimated cyclic trend of gene expression levels for each gene.
#     See https://jhsiao999.github.io/peco-paper/npreg_trendfilter_quantile.html. This file
#     makes "data/fit.quant.rds"
fits_all <- readRDS("data/fit.quant.rds")
genes_all <- names(fits_all)[order(sapply(fits_all,"[[",3), decreasing=T)]

res_unthinned <- do.call(rbind, lapply(seq_along(unique(pdata$chip_id)), function(ind) {
  ind <- unique(pdata$chip_id)[i]
  res_thin_each <- do.call(rbind, lapply(2:50, function(ngenes) {
    data_test <- readRDS(paste0("data/data_training_test/ind_",ind,"_data_test.rds"))
    data_train <- readRDS(paste0("data/data_training_test/ind_",ind,"_data_training.rds"))

    which_genes <- genes_all[1:ngenes]
    fit_train <- cycle_npreg_insample(
      Y = with(data_train,
               log2cpm_quant_train[which(rownames(log2cpm_quant_train) %in% which_genes), ]),
      theta = with(data_train, theta_train),
      method.trend="trendfilter")
    fit_test <- cycle_npreg_outsample(
      Y_test=with(data_test,
                  log2cpm_quant_test[which(rownames(log2cpm_quant_test) %in% which_genes), ]),
      sigma_est=with(fit_train, sigma_est),
      funs_est=with(fit_train, funs_est))

    diff_time <- circ_dist(data_test$theta_test,
                           rotation(data_test$theta_test, fit_test$cell_times_est))
    out <- data.frame(phase_pred_rot=rotation(data_test$theta_test, fit_test$cell_times_est),
                      phase_ref=data_test$theta_test,
                      diff_time=diff_time,
                      ind = ind,
                      ngenes = ngenes)
    return(out)
  }) )
}) )


res_unthinned %>% group_by(ind, ngenes) %>%
  #  filter(ngenes <= 20) %>%
  summarise(diff_mean = mean(diff_time/2/pi),
            diff_se = sd(diff_time/2/pi)/sqrt(length(diff_time/2/pi))) %>%
  ggplot(., aes(x=factor(ngenes), y=diff_mean, group = ind)) +
  #  geom_vline(xintercept=seq(5, 50,5)-1, col="gray90", lty=1) +
  geom_hline(yintercept=seq(.1, .2, .01), col="gray90", lty=1) +
  geom_line(aes(col=ind), lwd=.7) + # ggtitle("thinlog2 = .80") +
  scale_fill_brewer(palette="Dark2") +
  geom_errorbar(aes(ymin=diff_mean-diff_se,
                    ymax=diff_mean+diff_se, col=ind), width=.2, alpha=.5) +
  stat_summary(fun.y=mean,geom="line",lwd=.5, group=1) +
  ylim(0,.3) + geom_hline(yintercept=.25, col="red") +
  labs(color="Test data (Cell line)") +
  xlab("Number of cyclic genes used in peco prediction") +
  ylab("Prediction error (% circle)") +
  scale_x_discrete(breaks=c(2:50),
                   labels=c(rep("",3),5, rep("",4), 10, rep("",4), 15,
                            rep("",4), 20, rep("",4), 25,
                            rep("",4), 30, rep("",4), 35,
                            rep("",4), 40, rep("",4), 45, rep("",4), 50)) +
  ggtitle("Performance in unthinned data")


# Supp Fig s10B
inds <- c("NA19098","NA18511","NA18870","NA19101","NA18855","NA19160")
ngene <- 5
eval_fit <- res_unthinned %>%
  filter(ngenes ==5)

# sample size and prediction error
ns_error <- do.call(rbind, lapply(seq_along(inds), function(i) {
  data.frame(ind=inds[i],
             error=mean(eval_fit[eval_fit$ind == inds[i],]$diff_time),
             ns=sum(eval_fit$ind == inds[i]),
             stringsAsFactors = F)
}))
cols <- RColorBrewer::brewer.pal(9,"Dark2")
par(mfrow=c(1,1))
plot(y=(ns_error$error/2/pi),x=ns_error$ns, pch=16,
     ylab="Prediction error",
     xlab="Sample size in test sample", axes=F,
     xlim=c(80, 220), col=cols[1:6])
axis(1); axis(2)
text(labels=ns_error$ind[1],
     y=(ns_error$error[1]/2/pi),
     x=ns_error$ns[1], pos=4, offset=1, col=cols[1])
text(labels=ns_error$ind[2],
     y=(ns_error$error[2]/2/pi),
     x=ns_error$ns[2], pos=3, offset=1, col=cols[2])
text(labels=ns_error$ind[3],
     y=(ns_error$error[3]/2/pi),
     x=ns_error$ns[3], pos=3, offset=1, col=cols[3])
text(labels=ns_error$ind[4],
     y=(ns_error$error[4]/2/pi),
     x=ns_error$ns[4], pos=1, offset=1, col=cols[4])
text(labels=ns_error$ind[5],
     y=(ns_error$error[5]/2/pi),
     x=ns_error$ns[5], pos=3, offset=1, col=cols[5])
text(labels=ns_error$ind[6],
     y=(ns_error$error[6]/2/pi),
     x=ns_error$ns[6], pos=1, offset=1, col=cols[6])
cor.test(ns_error$error, ns_error$ns, method = "spearman")
title("Supp Fig S10B")


# Supp Fig 11C,D
#
# Prepare thinned data
makedata_thinned <- function(sce, thinlog2_rate) {

  library(SingleCellExperiment)
  library(peco)
  pdata <- data.frame(colData(sce))
  fdata <- data.frame(rowData(sce))
  counts <- assay(sce)[grep("ENSG", rownames(sce)), ]
  log2cpm <- t(log2(1+(10^6)*(t(counts)/pdata$molecules)))

  # ---- thinning!
  library(seqgendiff)
  nsamp <- ncol(counts)
  inputmat <- counts
  thinlog2 <- rexp(nsamp, rate = thinlog2_rate)
  outmat <- thin_lib(inputmat, thinlog2 = thinlog2)

  counts_thin <- outmat$mat
  dimnames(counts_thin) <- dimnames(counts)
  libsize <- colSums(counts_thin)

  keep_samples <- which(colMeans(counts_thin==0)>.01)
  keep_genes <- which(rowMeans(counts_thin == 0 ) < .5)
  counts_thin <- counts_thin[keep_genes, keep_samples]
  libsize <- libsize[keep_samples]

  log2cpm_thin <- t(log2(1+(10^6)*t(counts_thin)/libsize))
  dimnames(log2cpm_thin) <- dimnames(counts_thin)

  log2cpm_thin_quant <- do.call(rbind,
                                lapply(1:nrow(log2cpm_thin), function(i) {
                                  qqnorm(log2cpm_thin[i,], plot.it = F )$x }) )
  dimnames(log2cpm_thin_quant) <- dimnames(counts_thin)

  saveRDS(list(counts_thin=counts_thin,
               log2cpm_thin= log2cpm_thin,
               log2cpm_thin_quant=log2cpm_thin_quant),
          file=file.path(paste0("data/data_training_test/data_thinlog2_",
                                sprintf("%03d", 100*thinlog2_rate),".rds")))

  pdata_thin <- pdata[keep_samples,]

  for (i in 1:length(unique(pdata$chip_id))) {
    ind <- unique(pdata_thin$chip_id)[i]
    ii_test <- c(1:nrow(pdata_thin))[which(pdata_thin$chip_id == ind)]
    counts_test <- counts[,ii_test]
    counts_thin_test <- counts_thin[,ii_test]
    log2cpm_thin_test <- log2cpm_thin[,ii_test]
    log2cpm_thin_quant_test <- log2cpm_thin_quant[,ii_test]

    data_thin_test <- list(counts_test=counts_test,
                           counts_thin_test=counts_thin_test,
                           log2cpm_thin_test=log2cpm_thin_test,
                           log2cpm_thin_quant_test=log2cpm_thin_quant_test)

    saveRDS(data_thin_test,
            file=file.path(paste0("data/data_training_test/ind_",ind,"_data_test_thinlog2_",
                                  sprintf("%03d", 100*thinlog2_rate),".rds")))
  }
}

makedata_thinned(sce, thinlog2_rate = .8)
makedata_thinned(sce, thinlog2_rate = .33)
```
