## Supplementary material for "Characterizing and inferring quantitative cell cycle phase in single-cell RNA-seq data analysis": Source code of our analysis, including scripts and data necessary to reproduce the work.: suppfig11.R

```
# Supplemental Figure S11
#     Comparison of FUCCI phase with phase assignment by Oscope and reCAT
# Notes
#   - For details of how we applied each method, estimated cyclic trends and
#     computed prediction error, see https://jhsiao999.github.io/peco-paper/eval_on_our_data.html

library(SingleCellExperiment)
sce <- readRDS("data/sce-final.rds")
pdata <- data.frame(colData(sce))


# Supp Fig S11A, 11B
# Oscope and FUCCI score
load("data/ourdata_oscope_366genes.rda")
samples_ordered <- colnames(DataInput)[ENIRes[["cluster2"]]]
phase_oscope <- seq(0, 2*pi, length.out= length(ENIRes[[2]]))
names(phase_oscope) <- samples_ordered
ii_match <- match(samples_ordered, rownames(pdata) )

theta_final <- shift_origin(pdata$theta, origin=3*pi/4)
names(theta_final) <- rownames(pdata)

map_oscope <- data.frame(theta_final=theta_final,
                         phase_oscope=phase_oscope[match(names(theta_final), names(phase_oscope))],
                         sample_id=names(theta_final))
map_oscope$phase_oscope_rot <- rotation(ref_var=map_oscope$theta_final,
                                        shift_var=map_oscope$phase_oscope)

# Supp Fig S11A
par(mfrow=c(1,1))
plot(x=map_oscope$theta_final,
     y=map_oscope$phase_oscope_rot%%(2*pi),
     xlab="FUCCI phase", ylab="Oscope predicted phase",
     pch=16, cex=.6, axes=F, col="gray50")
abline(0,1, lty=1)
axis(1, at=c(0,pi/2, pi, 3*pi/2, 2*pi),
     labels=c(0,expression(pi/2), expression(pi), expression(3*pi/2),
              expression(2*pi)))
axis(2, at=c(0,pi/2, pi, 3*pi/2, 2*pi),
     labels=c(0,expression(pi/2), expression(pi), expression(3*pi/2),
              expression(2*pi)))
abline(a=pi/2, b=1, lty=2, col="black", lwd=.7)
abline(a=-pi/2, b=1, lty=2, col="black", lwd=.7)
title("Supp Fig S11A")


# Supp Fig S11B
par(mfrow=c(1,1))
plot(x=phase_oscope,
     y=pdata$gfp.median.log10sum.adjust[ii_match], col="forestgreen",
     ylim=c(-1.5, 1.5), cex=.5, pch=16,
     xlab="Oscope phase", ylab="FUCCI intensities",
     main="Oscope phase and FUCCI intensities", axes=F)
axis(2); axis(1,at=c(0,pi/2, pi, 3*pi/2, 2*pi),
              labels=c(0,expression(pi/2), expression(pi), expression(3*pi/2),
                       expression(2*pi)))
abline(h=0, lty=1, col="black", lwd=.7)
points(x=phase_oscope,
       y=pdata$rfp.median.log10sum.adjust[ii_match], col="firebrick",
       ylim=c(-1.5, 1.5), cex=.5, pch=16)
gfp <- pdata$gfp.median.log10sum.adjust[ii_match]
gfp_ordered <- gfp[order(phase_oscope)]
rfp <- pdata$rfp.median.log10sum.adjust[ii_match]
rfp_ordered <- rfp[order(phase_oscope)]
fit_gfp_oscope <- fit_trendfilter_generic(gfp_ordered)
fit_rfp_oscope <- fit_trendfilter_generic(rfp_ordered)
fit_gfp_oscope$pve
fit_rfp_oscope$pve
title("Supp Fig S11B")


# Supp Fig S11C, S11D
# reCAT phase and FUCCI score
load("data/ourdata_recat.rda")
samples_ordered <- rownames(test_exp)[ordIndex]
phase_recat <- seq(0, 2*pi, length.out= length(ordIndex))
names(phase_recat) <- samples_ordered
ii_match <- match(samples_ordered, rownames(pdata) )

theta_final <- shift_origin(pdata$theta, origin=3*pi/4)
names(theta_final) <- rownames(pdata)

map_recat <- data.frame(theta_final=theta_final,
                        phase_recat=phase_recat[match(names(theta_final), names(phase_recat))],
                        sample_id=names(theta_final))
map_recat$phase_recat_rot <- rotation(ref_var=map_recat$theta_final,
                                      shift_var=map_recat$phase_recat)

par(mfrow=c(1,1))
plot(x=map_recat$theta_final,
     y=map_recat$phase_recat_rot %% (2*pi),
     xlab="FUCCI phase", ylab="reCAT predicted phase",
     pch=16, cex=.6, axes=F, col="gray50")
abline(0,1, lty=1)
abline(a=pi/2, b=1, lty=2, col="black", lwd=.7)
abline(a=-pi/2, b=1, lty=2, col="black", lwd=.7)
axis(1, at=c(0,pi/2, pi, 3*pi/2, 2*pi),
     labels=c(0,expression(pi/2), expression(pi), expression(3*pi/2),
              expression(2*pi)))
axis(2, at=c(0,pi/2, pi, 3*pi/2, 2*pi),
     labels=c(0,expression(pi/2), expression(pi), expression(3*pi/2),
              expression(2*pi)))
title("Supp Fig S11C")


par(mfrow=c(1,1))
plot(x=phase_recat,
     y=pdata$gfp.median.log10sum.adjust[ii_match], col="forestgreen",
     ylim=c(-1.5, 1.5), cex=.5, pch=16,
     xlab="reCAT phase", ylab="FUCCI intensities",
     main="reCAT phase and FUCCI intensities", axes=F)
axis(2); axis(1,at=c(0,pi/2, pi, 3*pi/2, 2*pi),
              labels=c(0,expression(pi/2), expression(pi), expression(3*pi/2),
                       expression(2*pi)))
abline(h=0, lty=1, col="black", lwd=.7)
points(x=phase_recat,
       y=pdata$rfp.median.log10sum.adjust[ii_match], col="firebrick",
       ylim=c(-1.5, 1.5), cex=.5, pch=16)

gfp <- pdata$gfp.median.log10sum.adjust[ii_match]
gfp_ordered <- gfp[order(phase_recat)]
rfp <- pdata$rfp.median.log10sum.adjust[ii_match]
rfp_ordered <- rfp[order(phase_recat)]
fit_gfp_recat <- fit_trendfilter_generic(gfp_ordered)
fit_rfp_recat <- fit_trendfilter_generic(rfp_ordered)
fit_gfp_recat$pve
fit_rfp_recat$pve
title("Supp Fig S11D")
```
