## Supplementary material for "Characterizing and inferring quantitative cell cycle phase in single-cell RNA-seq data analysis": Source code of our analysis, including scripts and data necessary to reproduce the work.: suppfig12.R

```
# Supplemental Figure S12
#   FUCCI scores and inferred phase from peco in individual cell lines.
#
# Notes
#   - For details of how we used peco to estimate continuous cell cycle phase in
#     individual cell lines, see https://jhsiao999.github.io/peco-paper/predict_thinned_data.html


library(peco)
library(SingleCellExperiment)

inds <- c("NA19098","NA18511","NA18870","NA19101","NA18855","NA19160")
sce <- readRDS("data/sce-final.rds")
pdata <- data.frame(colData(sce))

pred_peco <- lapply(1:length(inds), function(i) {
  ind <- inds[i]
  out <- readRDS(paste0("data/ourdata_peco_",
                        ind, "_top",sprintf("%03d", 5), "genes.rds"))
  gfp <- pdata$gfp.median.log10sum.adjust
  rfp <- pdata$rfp.median.log10sum.adjust
  gfp_ind <- gfp[match(names(out$fit_test$cell_times_est)[order(out$fit_test$cell_times_est)],
                       rownames(pdata))]
  phase <- shift_origin(with(out$fit_test, cell_times_est[order(cell_times_est)]), origin = 3*pi/4)
  fit_gfp <- with(out$fit_test, fit_trendfilter_generic(gfp_ind))
  fun_gfp <- approxfun(x=as.numeric(phase),
                       y=as.numeric(fit_gfp$trend.yy), rule=2)
  fit_gfp$phase <- phase
  fit_gfp$gfp_ind <- gfp_ind
  fit_gfp$fun <- fun_gfp

  rfp_ind <- rfp[match(names(out$fit_test$cell_times_est)[order(out$fit_test$cell_times_est)],
                       rownames(pdata))]
  fit_rfp <- with(out$fit_test, fit_trendfilter_generic(rfp_ind))
  fun_rfp <- approxfun(x=as.numeric(phase),
                       y=as.numeric(fit_rfp$trend.yy), rule=2)
  fit_rfp$phase <- phase
  fit_rfp$rfp_ind <- rfp_ind
  fit_rfp$fun <- fun_rfp
  return(list(fit_rfp=fit_rfp, fit_gfp=fit_gfp))
})
names(pred_peco) <- inds

mean(sapply(pred_peco, function(x) x$fit_gfp$pve))
mean(sapply(pred_peco, function(x) x$fit_rfp$pve))


par(mfrow=c(3,2), mar=c(4,2,4,1))
for (i in 1:6) {
  plot(x=pred_peco[[i]]$fit_gfp$phase,
       y=pred_peco[[i]]$fit_gfp$gfp_ind, col="forestgreen",
       ylim=c(-1.5, 1.5), cex=.5, pch=16,
       xlab="peco phase", ylab="FUCCI score",
       main=paste0(names(pred_peco)[i], "\n",
                   "EGFP ", round(100*pred_peco[[i]]$fit_gfp$pve,2), "%,",
                   " mCherry ", round(100*round(pred_peco[[i]]$fit_rfp$pve,2)), "%"),
       axes=F)
  axis(2); axis(1,at=c(0,pi/2, pi, 3*pi/2, 2*pi),
                labels=c(0,expression(pi/2), expression(pi), expression(3*pi/2),
                         expression(2*pi)))
  abline(h=0, lty=1, col="black", lwd=.7)
  points(x=pred_peco[[i]]$fit_rfp$phase,
         y=pred_peco[[i]]$fit_rfp$rfp_ind, col="firebrick",
         ylim=c(-1.5, 1.5), cex=.5, pch=16)
}
```
