## Supplementary material for "Characterizing and inferring quantitative cell cycle phase in single-cell RNA-seq data analysis": Source code of our analysis, including scripts and data necessary to reproduce the work.: suppfig13.R

```
# Supplemental Figure S13
#   Comparison of FUCCI phase with phase assignment by Seurat and Cyclone

library(SingleCellExperiment)
library(peco)
library(cluster)

sce <- readRDS("data/sce-final.rds")
sce <- sce[grep("ENSG", rownames(sce)),]
pdata <- data.frame(colData(sce))
fdata <- data.frame(rowData(sce))
counts <- assay(sce, "counts")

sce <- data_transform_quantile(sce)
log2cpm_quantNormed <- assay(sce, "cpm_quantNormed")


# derive cell cycle phase
# use PCA to rotate the axes
pca <- prcomp(cbind(pdata$rfp.median.log10sum.adjust,
                    pdata$gfp.median.log10sum.adjust))
theta <- coord2rad(pca$x)
theta_final <- shift_origin(as.numeric(theta), 3*pi/4)
names(theta_final) = rownames(pdata)

# get PAM-based clusters
pam_res <- pam(cbind(pdata$rfp.median.log10sum.adjust,
                     pdata$gfp.median.log10sum.adjust), k=3)
clust <- data.frame(clust=pam_res$clustering,
                    sample_id=rownames(pdata))
plot(theta_final, clust$clust)

# find boundaries
b1 <- mean(max(range(theta_final[clust$clust==2])), min(range(theta_final[clust$clust==3])))
b2 <- mean(max(range(theta_final[clust$clust==3])), max(range(theta_final[clust$clust==1])))
b3 <- mean(min(range(theta_final[clust$clust==2])), min(range(theta_final[clust$clust==1])))
abline(v=c(b1,b2,b3), lty=2)


# Supp Fig 13A
# Seurat and PAM-based classification obtained from FUCCI scores
library(Seurat)
cc.genes <- readLines(con = "data/regev_lab_cell_cycle_genes.txt")
s.genes <- cc.genes[1:43]; g2m.genes <- cc.genes[44:97]
rownames_hgnc <- fdata[rownames(counts),]$name

obj <- CreateSeuratObject(counts = counts)
obj <- NormalizeData(obj)
obj <- FindVariableFeatures(obj, selection.method = "vst")
obj <- ScaleData(obj, features = rownames(obj))
obj <- CellCycleScoring(obj, s.features = s.genes, g2m.features = g2m.genes, set.ident = TRUE)

out_seurat <- obj[[]]
all.equal(rownames(out_seurat), colnames(log2cpm_quantNormed))
out_seurat <- out_seurat[match(colnames(log2cpm_quantNormed), rownames(out_seurat)),]


out_seurat$Phase <- factor(out_seurat$Phase,
                           levels = c("G1", "S", "G2M"))
out_seurat_ordered <- out_seurat[match(rownames(pdata), rownames(out_seurat)),]

all.equal(as.character(clust$sample_id), rownames(out_seurat_ordered))
table(out_seurat_ordered$Phase, clust$clust)


# Supp Fig S13B
# Cyclone and PAM-based classification obtained from FUCCI scores
#
# run Cyclone
Y_cyclone <- log2cpm_quantNormed

library(scran)
hs.pairs <- readRDS(system.file("exdata", "human_cycle_markers.rds", package="scran"))

out_cyclone <- cyclone(Y_cyclone, pairs = hs.pairs,
                       gene.names=rownames(Y_cyclone),
                       iter=1000, min.iter=100, min.pairs=50,
                       BPPARAM=SerialParam(), verbose=T, subset.row=NULL)
names(out_cyclone$phases) <- colnames(Y_cyclone)
all.equal(names(out_cyclone$phases), colnames(log2cpm_quantNormed))

out_cyclone$phases <- factor(out_cyclone$phases,
                             levels = c("G1", "S", "G2M"))
out_cyclone_ordered_phases <- out_cyclone$phases[match(rownames(pdata), names(out_cyclone$phases))]

all.equal(as.character(clust$sample_id), names(out_cyclone_ordered_phases))
table(out_cyclone_ordered_phases, clust$clust)


# Supp Fig S13C
# Seurat-based classification and FUCCI phase
# seurat discrete classes vs FUCCI phase
out_seurat2 = out_seurat$Phase[match(names(theta_final), rownames(out_seurat))]
all.equal(rownames(out_seurat2), names(theta_final))

phases = c("G1", "S", "G2M")
cols = as.character(yarrr::piratepal("espresso")[c(3,2,1)])
par(mfrow = c(1,3))
for (i in seq_along(phases)) {
  hist(theta_final[out_seurat2==phases[i]],
       breaks = 80, ylim = c(0,40), axes = F, ann = F, lty = "blank",
       col = cols[i])
  title(main = paste(sum(out_seurat2==phases[i]), "cells"))
  axis(2); axis(1,at=c(0,pi/2, pi, 3*pi/2, 2*pi),
                labels=c(0,expression(pi/2), expression(pi), expression(3*pi/2),
                         expression(2*pi)))
  abline(v=c(b1,b2,b3), lty=2)
}


# Supp Fig S13D
# Cyclone-based classification and FUCCI phase
out_cyclone2 = out_cyclone$phases[match(names(theta_final),
                                        names(out_cyclone$phases))]
all.equal(names(out_cyclone2), names(theta_final))

phases = c("G1", "S", "G2M")
cols = as.character(yarrr::piratepal("espresso")[c(3,2,1)])
par(mfrow = c(1,3))
for (i in seq_along(phases)) {
  hist(theta_final[out_cyclone2==phases[i]],
       breaks = 80, ylim = c(0,40), axes = F, ann = F, lty = "blank",
       col = cols[i])
  title(main = paste(sum(out_cyclone2==phases[i]), "cells"))
  axis(2); axis(1,at=c(0,pi/2, pi, 3*pi/2, 2*pi),
                labels=c(0,expression(pi/2), expression(pi), expression(3*pi/2),
                         expression(2*pi)))
  abline(v=c(b1,b2,b3), lty=2)
}
```
