## Supplementary material for "Characterizing and inferring quantitative cell cycle phase in single-cell RNA-seq data analysis": Source code of our analysis, including scripts and data necessary to reproduce the work.: suppfig14.R

```
# Supplemental Figure S14
#   Continuous cell cycle phase assignmetn based on the two Seurat phase-specific
# scores for samples from cell line NA19098.

library(SingleCellExperiment)
sce <- readRDS("data/sce-final.rds")

# import Seurat results
inds <- c("NA19098","NA18511","NA18870","NA19101","NA18855","NA19160")
eval_seurat <- lapply(1:length(inds), function(j) {
  ind <- inds[j]
  fl_name <- file.path(paste0("data/ind_",ind,
                           "_results_overallcyclical.top5.rds"))
  df <- readRDS(fl_name)
  return(df$fit.test)
})
names(eval_seurat) <- inds

# Seurat scores before PCA transformation
par(mfrow=c(1,1), mar=c(5,4,3,1))
plot(eval_seurat[[1]]$fit.seurat$S,
     eval_seurat[[1]]$fit.seurat$G2M,
     xlab="S score",
     ylab="G2M score",
     pch=16, cex=.5, axes=F, xlim=c(-1.5,1.5), ylim=c(-1.5,1.5),
     col=c("darkgoldenrod1", "coral", "darkred")[eval_seurat[[1]]$fit.seurat$assignments])
axis(1); axis(2)
abline(v=0,a=0,b=1,h=0, lty=2, lwd=.5)

# Seurat scores after PCA transformation
par(mfrow=c(1,1), mar=c(5,4,3,1))
seurat.pca <- prcomp(cbind(eval_seurat[[1]]$fit.seurat$G2M,
                           eval_seurat[[1]]$fit.seurat$S), scale=TRUE)
seurat.cell_times_est <- as.numeric(coord2rad(cbind(seurat.pca$x[,1],seurat.pca$x[,2])))
(seurat.pca$sdev^2)/sum(seurat.pca$sdev^2)
plot(seurat.pca$x[,1],seurat.pca$x[,2], pch=16, cex=.5,
     xlim=c(-4,4),ylim=c(-4,4),
     col=c("darkgoldenrod1", "coral", "darkred")[eval_seurat[[1]]$fit.seurat$assignments],
     axes=F, xlab="PC1 (83%)", ylab="PC2 (17%)")
axis(1); axis(2)
abline(v=0,h=0, lty=2, lwd=.5)
par(new=TRUE)
library(circular)
plot(as.circular(eval_seurat[[1]]$fit.seurat$cell_times_est[
  eval_seurat[[1]]$fit.seurat$assignments=="G1"]),
  stack=T, shrink=1, cex=.5, col="darkgoldenrod1")
points(as.circular(eval_seurat[[1]]$fit.seurat$cell_times_est[
  eval_seurat[[1]]$fit.seurat$assignments=="S"]),
  stack=T, shrink=1, cex=.5, col="coral")
points(as.circular(eval_seurat[[1]]$fit.seurat$cell_times_est[
  eval_seurat[[1]]$fit.seurat$assignments=="G2M"]),
  stack=T, shrink=1, cex=.5, col="darkred", bins=200)
```
