## Supplementary material for "Characterizing and inferring quantitative cell cycle phase in single-cell RNA-seq data analysis": Source code of our analysis, including scripts and data necessary to reproduce the work.: suppfig15.R

```
# Supplemental Figure S15
# Comparison of prediction error on our data in cross-validation.

# Notes
#   - For details of how we applied each method, estimated cyclic trends and
#     computed prediction error, see https://jhsiao999.github.io/peco-paper/eval_on_our_data.html


diff_cyclone <- readRDS("data/fit_diff_cyclone.rds")
diff_recat <- readRDS("data/fit_diff_recat.rds")
diff_peco <- readRDS("data/fit_diff_peco.rds")
diff_seurat <- readRDS("data/fit_diff_seurat.rds")
diff_oscope <- readRDS("data/fit_diff_oscope.rds")

diff_peco %>% group_by(ind) %>% summarise(mn=mean(diff_time)/2/pi,
                                          sd=sd(diff_time/2/pi)/sqrt(sum(phase_pred_rot>0)))

diff_cyclone %>% group_by(ind) %>% summarise(mn=mean(diff_time)/2/pi,
                                             sd=sd(diff_time/2/pi)/sqrt(sum(phase_pred_rot>0)))

diff_seurat %>% group_by(ind) %>% summarise(mn=mean(diff_time)/2/pi,
                                            sd=sd(diff_time/2/pi)/sqrt(sum(phase_pred_rot>0)))

diff_oscope %>% group_by(ind) %>% summarise(mn=mean(diff_time)/2/pi,
                                            sd=sd(diff_time/2/pi)/sqrt(sum(phase_pred_rot>0)))

diff_recat %>% group_by(ind) %>% summarise(mn=mean(diff_time)/2/pi,
                                           sd=sd(diff_time/2/pi)/sqrt(sum(phase_pred_rot>0)))

diff_all <- rbind(diff_recat,
                  diff_cyclone,
                  diff_peco,
                  diff_seurat,
                  diff_oscope)

pval_difftime <- do.call(rbind, lapply(inds, function(ind) {
  data.frame(ind=ind,
             p_cyclone=wilcox.test(diff_peco[diff_peco$ind==ind,]$diff_time/2/pi,
                                   diff_cyclone[diff_cyclone$ind==ind,]$diff_time/2/pi)$p.value,
             p_seurat=wilcox.test(diff_peco[diff_peco$ind==ind,]$diff_time/2/pi,
                                  diff_seurat[diff_peco$ind==ind,]$diff_time/2/pi)$p.value,
             p_oscope=wilcox.test(diff_peco[diff_peco$ind==ind,]$diff_time/2/pi,
                                  diff_oscope[diff_peco$ind==ind,]$diff_time/2/pi)$p.value,
             p_recat=wilcox.test(diff_peco[diff_peco$ind==ind,]$diff_time/2/pi,
                                 diff_recat[diff_peco$ind==ind,]$diff_time/2/pi)$p.value,
             ratio_cyclone=mean(diff_peco[diff_peco$ind==ind,]$diff_time)/mean(diff_cyclone[diff_cyclone$ind==ind,]$diff_time),
             ratio_seurat=mean(diff_peco[diff_peco$ind==ind,]$diff_time)/mean(diff_seurat[diff_seurat$ind==ind,]$diff_time),
             ratio_oscope=mean(diff_peco[diff_peco$ind==ind,]$diff_time)/mean(diff_oscope[diff_oscope$ind==ind,]$diff_time),
             ratio_recat=mean(diff_peco[diff_peco$ind==ind,]$diff_time)/mean(diff_recat[diff_recat$ind==ind,]$diff_time))
}))
cbind(inds, round(pval_difftime[,-1],4))

mean(pval_difftime$ratio_cyclone)
mean(pval_difftime$ratio_seurat)
mean(pval_difftime$ratio_oscope)
mean(pval_difftime$ratio_recat)
```
