## Supplementary material for "Characterizing and inferring quantitative cell cycle phase in single-cell RNA-seq data analysis": Source code of our analysis, including scripts and data necessary to reproduce the work.: suppfig17.R

```
# Supplemental Figure S17
# Cyclic expression predicted results of each method
#
# Notes
#   - For details of how we applied each method, estimated cyclic trends and
#     computed prediction error, see
#     https://jhsiao999.github.io/peco-paper/eval_on_our_data.html. This
#     file produced "data/fit_diff*.rds" files and also the results of cell cycle
#     prediction in each method.

library(SingleCellExperiment)
library(peco)
library(wesanderson)

sce <- readRDS("data/sce-final.rds")
sce <- sce[grep("ENSG", rownames(sce)),]
fdata <- data.frame(colData(sce))
fdata <- data.frame(rowData(sce))
counts <- data.frame(assay(sce, "counts"))
sce_normed <- data_transform_quantile(sce)
log2cpm_quant <- assay(sce_normed, "cpm_quantNormed")


# peco
labs <- c("CDK1", "UBE2C", "TOP2A", "HIST1H4E", "HIST1H4C")
genes <- c("ENSG00000170312","ENSG00000175063",
           "ENSG00000131747", "ENSG00000198518", "ENSG00000197061")
pred_peco <- lapply(1:length(inds), function(i) {
  ind <- inds[i]
  out <- readRDS(paste0("data/ourdata_peco_",
                                  ind, "_top",sprintf("%03d", 5), "genes.rds"))
  fits <- lapply(1:5, function(g) {
    gexp <- with(out$fit_test, Y[g,order(cell_times_est)])
    phase <- shift_origin(with(out$fit_test, cell_times_est[order(cell_times_est)]),
                          origin = 3*pi/4)
    fit_g <- with(out$fit_test, fit_trendfilter_generic(gexp))
    fun_g <- approxfun(x=as.numeric(phase),
                       y=as.numeric(fit_g$trend.yy), rule=2)
    fit_g$phase <- phase
    fit_g$gexp <- gexp
    fit_g$fun_g <- fun_g
    return(fit_g)
  })
  names(fits) <- labs
  return(fits)
})
names(pred_peco) <- inds

par(mfrow=c(6,5), mar=c(2,2,2,1))
for (i in 1:6) {
  res <- pred_peco[[i]]
  ind <- names(pred_peco)[i]
  for (g in 1:5) {
    if (g>1) { ylab <- ""} else { ylab <- "Quantile-normalized expression level"}
    #    if (i==1) { xlab <- "Predicted phase"} else { xlab <- "Inferred phase"}
    res_g <- res[[g]]
    plot(x= res_g$phase,
         y = res_g$gexp, pch = 16, cex=.5, ylab=ylab,
         xlab="Predicted phase",
         main = names(res)[g], axes=F, ylim=c(-3,3))
    abline(h=0, lwd=.5)
    lines(x = seq(0, 2*pi, length.out=200),
          y = res_g$fun_g(seq(0, 2*pi, length.out=200)),
          col = wesanderson::wes_palette("FantasticFox1")[1],
          lty=1, lwd=2)
    axis(2); axis(1,at=c(0,pi/2, pi, 3*pi/2, 2*pi),
                  labels=c(0,expression(pi/2), expression(pi), expression(3*pi/2),
                           expression(2*pi)))
  }
}


# Cyclone
inds <- c("NA19098","NA18511","NA18870","NA19101","NA18855","NA19160")
labs <- c("CDK1", "UBE2C", "TOP2A", "HIST1H4E", "HIST1H4C")
genes <- c("ENSG00000170312","ENSG00000175063",
           "ENSG00000131747", "ENSG00000198518", "ENSG00000197061")
pred_cyclone <- lapply(1:length(inds), function(i) {
  res <- readRDS(paste0("data/ourdata_cyclone_", inds[i], ".rds"))
  phase_pred <- res$phase_pred
  names(phase_pred) <- rownames(res)
  phase_ref <- pdata$theta[match(names(phase_pred), rownames(pdata))]
  phase_pred_rot <- rotation(ref_var= phase_ref, shift_var = phase_pred)
  fits <- lapply(1:5, function(g) {
    gindex <- which(rownames(log2cpm_quant) == genes[g])
    fit_g <- data.frame(
      gexp=log2cpm_quant[gindex, match(names(phase_pred_rot), colnames(log2cpm_quant))],
      phase=shift_origin(phase_pred_rot, origin = pi/2))
    fit_g <- fit_g[order(fit_g$phase),]

    fit_trend <- fit_trendfilter_generic(fit_g$gexp)
    fit_g$trend.yy <- fit_trend$trend.yy
    fun_g <- approxfun(x=as.numeric(fit_g$phase),
                       y=as.numeric(fit_g$trend.yy), rule=2)
    fit_out <- list(fit_g=fit_g,
                    fun_g = fun_g)
    return(fit_out)
  })
  names(fits) <- labs
  return(fits)
})
names(pred_cyclone) <- inds

par(mfrow=c(6,5), mar=c(2,2,2,1))
for (i in 1:6) {
  res <- pred_cyclone[[i]]
  ind <- names(pred_cyclone)[i]
  for (g in 1:5) {
    if (g>1) { ylab <- ""} else { ylab <- "Quantile-normalized expression level"}
    res_g <- res[[g]]
    plot(x= res_g$fit_g$phase,
         y = res_g$fit_g$gexp, pch = 16, cex=.5, ylab=ylab,
         xlab="Predicted phase",
         main = names(res)[g], axes=F, ylim=c(-3,3))
    abline(h=0, lwd=.5)
    lines(x = seq(0, 2*pi, length.out=200),
          y = res_g$fun_g(seq(0, 2*pi, length.out=200)),
          col = wesanderson::wes_palette("FantasticFox1")[2], lty=1, lwd=2)
    axis(2); axis(1,at=c(0,pi/2, pi, 3*pi/2, 2*pi),
                  labels=c(0,expression(pi/2), expression(pi), expression(3*pi/2),
                           expression(2*pi)))
  }
}


# Seurat
inds <- c("NA19098","NA18511","NA18870","NA19101","NA18855","NA19160")
labs <- c("CDK1", "UBE2C", "TOP2A", "HIST1H4E", "HIST1H4C")
genes <- c("ENSG00000170312","ENSG00000175063",
           "ENSG00000131747", "ENSG00000198518", "ENSG00000197061")
diff_seurat <- readRDS(file="data/fit_diff_seurat.rds")

pred_seurat <- lapply(1:length(inds), function(i) {
  out <- diff_seurat[diff_seurat$ind==inds[i],c("phase_ref", "phase_pred_rot")]
  fits <- lapply(1:5, function(g) {
    gindex <- which(rownames(log2cpm_quant) == genes[g])
    fit_g <- data.frame(
      gexp=log2cpm_quant[gindex, match(rownames(out), colnames(log2cpm_quant))],
      phase=shift_origin(2*pi-out$phase_pred_rot, pi/3))
    fit_g <- fit_g[order(fit_g$phase),]

    fit_trend <- fit_trendfilter_generic(fit_g$gexp)
    fit_g$trend.yy <- fit_trend$trend.yy
    fun_g <- approxfun(x=as.numeric(fit_g$phase),
                       y=as.numeric(fit_g$trend.yy), rule=2)
    fit_out <- list(fit_g=fit_g,
                    fun_g = fun_g)
    return(fit_out)
  })
  names(fits) <- labs
  return(fits)
})
names(pred_seurat) <- inds

par(mfrow=c(6,5), mar=c(2,2,2,1))
for (i in 1:6) {
  res <- pred_seurat[[i]]
  ind <- names(pred_seurat)[i]
  for (g in 1:5) {
    if (g>1) { ylab <- ""} else { ylab <- "Quantile-normalized expression level"}
    res_g <- res[[g]]
    plot(x= res_g$fit_g$phase,
         y = res_g$fit_g$gexp, pch = 16, cex=.5, ylab=ylab,
         xlab="Predicted phase",
         main = names(res)[g], axes=F, ylim=c(-3,3))
    lines(x = seq(0, 2*pi, length.out=200),
          y = res_g$fun_g(seq(0, 2*pi, length.out=200)),
          col = wesanderson::wes_palette("FantasticFox1")[3], lty=1, lwd=2)
    axis(2); axis(1,at=c(0,pi/2, pi, 3*pi/2, 2*pi),
                  labels=c(0,expression(pi/2), expression(pi), expression(3*pi/2),
                           expression(2*pi)))
  }
}


# Oscope cyclic expression patterns
inds <- c("NA19098","NA18511","NA18870","NA19101","NA18855","NA19160")
labs <- c("CDK1", "UBE2C", "TOP2A", "HIST1H4E", "HIST1H4C")
genes <- c("ENSG00000170312","ENSG00000175063",
           "ENSG00000131747", "ENSG00000198518", "ENSG00000197061")

load("data/ourdata_oscope_366genes.rda")

pred_oscope <- lapply(1:length(inds), function(i) {
  out_oscope_sub <- out_oscope[out_oscope$ind == inds[i],]
  fits <- lapply(1:5, function(g) {
    gindex <- which(rownames(log2cpm_quant) == genes[g])
    fit_g <- data.frame(
      gexp=log2cpm_quant[gindex, match(rownames(out_oscope_sub), colnames(log2cpm_quant))],
      phase=shift_origin(2*pi-out_oscope_sub$phase_pred_rot, origin=3*pi/4))
    fit_g <- fit_g[order(fit_g$phase),]

    fit_trend <- fit_trendfilter_generic(fit_g$gexp)
    fit_g$trend.yy <- fit_trend$trend.yy
    fun_g <- approxfun(x=as.numeric(fit_g$phase),
                       y=as.numeric(fit_g$trend.yy), rule=2)
    fit_out <- list(fit_g=fit_g,
                    fun_g = fun_g)
    return(fit_out)
  })
  names(fits) <- labs
  return(fits)
})
names(pred_oscope) <- inds

par(mfrow=c(6,5), mar=c(2,2,2,1))
for (i in 1:6) {
  res <- pred_oscope[[i]]
  print(inds[i])
  for (g in 1:5) {
    if (g>1) { ylab <- ""} else { ylab <- "Quantile-normalized expression level"}
    res_g <- res[[g]]
    plot(x= res_g$fit_g$phase,
         y = res_g$fit_g$gexp, pch = 16, cex=.5, ylab=ylab,
         xlab="Predicted phase",
         main = names(res)[g], axes=F, ylim=c(-3,3))
    abline(h=0, lwd=.5)
    lines(x = seq(0, 2*pi, length.out=200),
          y = res_g$fun_g(seq(0, 2*pi, length.out=200)),
          col = wesanderson::wes_palette("Darjeeling2")[2], lty=1, lwd=2)
    axis(2); axis(1,at=c(0,pi/2, pi, 3*pi/2, 2*pi),
                  labels=c(0,expression(pi/2), expression(pi), expression(3*pi/2),
                           expression(2*pi)))
  }
}


# recat
labs <- c("CDK1", "UBE2C", "TOP2A", "HIST1H4E", "HIST1H4C")
genes <- c("ENSG00000170312","ENSG00000175063",
           "ENSG00000131747", "ENSG00000198518", "ENSG00000197061")
# load pre-computed predicte results
load("data/ourdata_recat.rda")
samples_ordered <- rownames(test_exp)[ordIndex]
phase_pred <- seq(0, 2*pi, length.out= length(ordIndex))
names(phase_pred) <- samples_ordered

out_recat <- data.frame(phase_ref=pdata$theta[match(samples_ordered, colnames(sce))])
out_recat$ind <- pdata$chip_id[match(samples_ordered, colnames(sce))]

out_recat$phase_pred_rot <- rotation(ref_var= out_recat$phase_ref, shift_var = phase_pred)
rownames(out_recat) <- samples_ordered

pred_recat <- lapply(1:length(inds), function(i) {
  out_recat_sub <- out_recat[out_recat$ind == inds[i],]
  fits <- lapply(1:5, function(g) {
    gindex <- which(rownames(log2cpm_quant) == genes[g])
    fit_g <- data.frame(
      gexp=log2cpm_quant[gindex, rownames(out_recat_sub)],
      phase=shift_origin(2*pi-out_recat_sub$phase_pred_rot, origin = 2*pi/3))
    fit_g <- fit_g[order(fit_g$phase),]

    fit_trend <- fit_trendfilter_generic(fit_g$gexp)
    fit_g$trend.yy <- fit_trend$trend.yy
    #    fit_g$pve <- fit$pve
    fun_g <- approxfun(x=as.numeric(fit_g$phase),
                       y=as.numeric(fit_g$trend.yy), rule=2)
    fit_out <- list(fit_g=fit_g,
                    pve = fit_trend$pve,
                    fun_g = fun_g)
    return(fit_out)
  })
  names(fits) <- labs
  return(fits)
})
names(pred_recat) <- inds


par(mfrow=c(6,5), mar=c(2,2,2,1))
for (i in 1:6) {
  res <- pred_recat[[i]]
  print(inds[i])
  for (g in 1:5) {
    if (g>1) { ylab <- ""} else { ylab <- "Quantile-normalized expression level"}
    res_g <- res[[g]]
    plot(x= res_g$fit_g$phase,
         y = res_g$fit_g$gexp, pch = 16, cex=.5, ylab=ylab,
         xlab="Predicted phase",
         main = names(res)[g], axes=F, ylim=c(-3,3))
    abline(h=0, lwd=.5)
    lines(x = seq(0, 2*pi, length.out=200),
          y = res_g$fun_g(seq(0, 2*pi, length.out=200)),
          col = wesanderson::wes_palette("FantasticFox1")[5], lty=1, lwd=2)
    axis(2); axis(1,at=c(0,pi/2, pi, 3*pi/2, 2*pi),
                  labels=c(0,expression(pi/2), expression(pi), expression(3*pi/2),
                           expression(2*pi)))
  }
}
```
