## Supplementary material for "Characterizing and inferring quantitative cell cycle phase in single-cell RNA-seq data analysis": Source code of our analysis, including scripts and data necessary to reproduce the work.: suppfig18.R

```
# Supplemental Figure S18
# Authored by John Blischak
# Please see https://github.com/jdblischak/fucci-seq/blob/hg38/code/hg38.R


#!/usr/bin/env Rscript

# Hacky script to compare hg19 molecule counts in ~/fucci-seq/data/molecules and
# hg38 molecules counts in ~/fucci-seq-hg38/data/molecules.
#
# Script executed in ~/fucci-seq-hg38/

library(SingleCellExperiment)
library(cowplot)
library(data.table)
library(ggplot2)
library(stringr)
theme_set(theme_cowplot())

# Input ------------------------------------------------------------------------
# the following txt.gz files are available upon request

mol_hg19 <- fread("zcat ../fucci-seq/data/molecules/*.txt.gz | grep -v sample")
colnames(mol_hg19) <- colnames(fread("zcat ../fucci-seq/data/molecules/20170905.txt.gz",
                                     nrows = 1))
dim(mol_hg19)
mol_hg19[, 1:5]

mol_hg38 <- fread("zcat data/molecules/*.txt.gz | grep -v sample")
colnames(mol_hg38) <- colnames(fread("zcat data/molecules/20170905.txt.gz",
                                     nrows = 1))
dim(mol_hg38)
mol_hg38[, 1:5]

# The sample should be identical
stopifnot(mol_hg19$sample == mol_hg38$sample,
          mol_hg19$experiment == mol_hg38$experiment,
          mol_hg19$well == mol_hg38$well)

# There are slight differences in annotation between releases 75 and 96
genes_hg19 <- str_subset(colnames(mol_hg19), "^ENSG")
genes_hg38 <- str_subset(colnames(mol_hg38), "^ENSG")
genes_all <- union(genes_hg19, genes_hg38)
genes_shared <- intersect(genes_hg19, genes_hg38)
length(genes_shared) / length(genes_all)
## [1] 0.8929394

# Limiting to protein-coding genes used in the actual study
sce <- readRDS("data/sce-final.rds")
genes_study <- rownames(sce)
genes_hg19 <- genes_hg19[genes_hg19 %in% genes_study]
genes_hg38 <- genes_hg38[genes_hg38 %in% genes_study]
genes_all <- union(genes_hg19, genes_hg38)
genes_shared <- intersect(genes_hg19, genes_hg38)
length(genes_shared) / length(genes_all)
## [1] 0.9897645

# Filter -----------------------------------------------------------------------

cols_shared <- c("sample", "experiment", "well", genes_shared)

mol_hg19_shared <- mol_hg19[, ..cols_shared]
dim(mol_hg19_shared)
mol_hg38_shared <- mol_hg38[, ..cols_shared]
dim(mol_hg38_shared)
stopifnot(dim(mol_hg19_shared) == dim(mol_hg38_shared),
          colnames(mol_hg19_shared) == colnames(mol_hg38_shared))


# Confirm no genes with zero counts across all cells
sums_hg19 <- mol_hg19_shared[, vapply(.SD, sum, integer(1)), .SDcols = -(sample:well)]
sums_hg38 <- mol_hg38_shared[, vapply(.SD, sum, integer(1)), .SDcols = -(sample:well)]
cor(sums_hg19, sums_hg38)
plot(sums_hg19, sums_hg38)
sums_total <- sums_hg19 + sums_hg38
zeros <- sums_total == 0
stopifnot(sum(zeros) == 0)

# Correlation per gene ---------------------------------------------------------

corrs <- numeric(length = length(genes_shared))
names(corrs) <- genes_shared
for (i in seq_along(genes_shared)) {
  counts_hg19 <- unlist(mol_hg19_shared[, genes_shared[i], with = FALSE])
  counts_hg38 <- unlist(mol_hg38_shared[, genes_shared[i], with = FALSE])
  if (sum(counts_hg19) == 0 || sum(counts_hg38) == 0) {
    corrs[i] <- 0
  } else {
    corrs[i] <- cor(counts_hg19, counts_hg38)
  }
}
summary(corrs)
hist(corrs)
mean(corrs < 0.9, na.rm = TRUE)
## [1] 0.09792258
sum(corrs < 0.9)
## [1] 1070
sum(corrs < 0)
## [1] 2

negative <- names(corrs)[corrs < 0]
plot(unlist(mol_hg19_shared[, negative[1], with = FALSE]),
     unlist(mol_hg38_shared[, negative[1], with = FALSE]))
plot(unlist(mol_hg19_shared[, negative[2], with = FALSE]),
     unlist(mol_hg38_shared[, negative[2], with = FALSE]))

five <- c("CDK1", "UBE2C", "TOP2A", "HIST1H4E", "HIST1H4C")
five_ensg <- rownames(sce)[data.frame(rowData(sce))$name %in% five]
five_ensg %in% genes_shared
cycling <- five[five_ensg %in% genes_shared]
cycling_ensg <- five_ensg[five_ensg %in% genes_shared]

plot(unlist(mol_hg19_shared[, 5]), unlist(mol_hg38_shared[, 5]))
cor(unlist(mol_hg19_shared[, 5]), unlist(mol_hg38_shared[, 5]))

for (i in seq_along(cycling)) {
  cycling_hg19 <- unlist(mol_hg19_shared[, cycling_ensg[i], with = FALSE])
  cycling_hg38 <- unlist(mol_hg38_shared[, cycling_ensg[i], with = FALSE])
  plot(cycling_hg19, cycling_hg38, main = paste(cycling[i], cycling_ensg[i]),
       sub = cor(cycling_hg19, cycling_hg38))
}

# Correlation per sample -------------------------------------------------------

# Remove columns with sample metadata
mol_hg19_shared_mat <- as.matrix(mol_hg19_shared[, -(sample:well)])
mol_hg38_shared_mat <- as.matrix(mol_hg38_shared[, -(sample:well)])

cor_sample <- numeric(length = nrow(mol_hg19_shared_mat))

for (i in seq_len(nrow(mol_hg19_shared_mat))) {
  cor_sample[i] <- cor(mol_hg19_shared_mat[i, ],
                       mol_hg38_shared_mat[i, ])
}

summary(cor_sample)
hist(cor_sample)
mean(cor_sample < 0.9, na.rm = TRUE)
## [1] 0.1992188
sum(cor_sample < 0.9)
## [1] 306
sum(cor_sample < 0)
## [1] 0

sums_per_sample <- rowSums(mol_hg19_shared_mat) + rowSums(mol_hg38_shared_mat)
plot(sums_per_sample, cor_sample)

# Plot -------------------------------------------------------------------------

(p_gene <- ggplot(data.frame(cor = corrs), aes(x = cor)) +
   geom_histogram(binwidth = 0.025) +
   labs(x = "Pearson correlation", y = "Number of genes",
        title = "Gene-gene correlation"))
(p_samp <- ggplot(data.frame(cor = cor_sample), aes(x = cor)) +
    geom_histogram(binwidth = 0.025) +
    labs(x = "Pearson correlation", y = "Number of cells",
         title = "Sample-sample correlation"))
(p_cdk1 <- ggplot(data.frame(hg19 = unlist(mol_hg19_shared[, cycling_ensg[1], with = FALSE]),
                             hg38 = unlist(mol_hg38_shared[, cycling_ensg[1], with = FALSE])),
                  aes(x = hg19, y = hg38)) +
    geom_point(shape = "circle open") +
    geom_abline(intercept = 0, slope = 1, color = "red", linetype = "dashed") +
    labs(x = "hg19 gene counts", y = "hg38 gene counts",
         title = sprintf("%s (%s)", cycling[1], cycling_ensg[1]),
         subtitle = sprintf("Pearson correlation: %.5f",
                            cor(unlist(mol_hg19_shared[, cycling_ensg[1], with = FALSE]),
                                unlist(mol_hg38_shared[, cycling_ensg[1], with = FALSE])))))

(p_ube2c <- ggplot(data.frame(hg19 = unlist(mol_hg19_shared[, cycling_ensg[2], with = FALSE]),
                              hg38 = unlist(mol_hg38_shared[, cycling_ensg[2], with = FALSE])),
                   aes(x = hg19, y = hg38)) +
    geom_point(shape = "circle open") +
    geom_abline(intercept = 0, slope = 1, color = "red", linetype = "dashed") +
    labs(x = "hg19 gene counts", y = "hg38 gene counts",
         title = sprintf("%s (%s)", cycling[2], cycling_ensg[2]),
         subtitle = sprintf("Pearson correlation: %.5f",
                            cor(unlist(mol_hg19_shared[, cycling_ensg[2], with = FALSE]),
                                unlist(mol_hg38_shared[, cycling_ensg[2], with = FALSE])))))

(p_top2a <- ggplot(data.frame(hg19 = unlist(mol_hg19_shared[, cycling_ensg[3], with = FALSE]),
                              hg38 = unlist(mol_hg38_shared[, cycling_ensg[3], with = FALSE])),
                   aes(x = hg19, y = hg38)) +
    geom_point(shape = "circle open") +
    geom_abline(intercept = 0, slope = 1, color = "red", linetype = "dashed") +
    labs(x = "hg19 gene counts", y = "hg38 gene counts",
         title = sprintf("%s (%s)", cycling[3], cycling_ensg[3]),
         subtitle = sprintf("Pearson correlation: %.5f",
                            cor(unlist(mol_hg19_shared[, cycling_ensg[3], with = FALSE]),
                                unlist(mol_hg38_shared[, cycling_ensg[3], with = FALSE])))))

(p_hist1h4e <- ggplot(data.frame(hg19 = unlist(mol_hg19_shared[, cycling_ensg[4], with = FALSE]),
                                 hg38 = unlist(mol_hg38_shared[, cycling_ensg[4], with = FALSE])),
                      aes(x = hg19, y = hg38)) +
    geom_point(shape = "circle open") +
    geom_abline(intercept = 0, slope = 1, color = "red", linetype = "dashed") +
    labs(x = "hg19 gene counts", y = "hg38 gene counts",
         title = sprintf("%s (%s)", cycling[4], cycling_ensg[4]),
         subtitle = sprintf("Pearson correlation: %.5f",
                            cor(unlist(mol_hg19_shared[, cycling_ensg[4], with = FALSE]),
                                unlist(mol_hg38_shared[, cycling_ensg[4], with = FALSE])))))

plot_grid(p_gene, p_samp, p_cdk1, p_ube2c, p_top2a, p_hist1h4e,
          nrow = 3, labels = "AUTO")
ggsave(filename = "hg38.png", width = 8, height = 12)

# (A) The Pearson correlation for each gene comparing its counts across single
# cells when mapping to genome build hg19 versus hg38. The median correlation
# was 0.983919, and 1,070 genes (9.8%) had a correlation less than 0.9. (B) The
# Pearson correlation for each sample comparing its gene counts when mapped to
# genome build hg19 versus hg38.  The median correlation was 0.9849, and 306
# single cells (19.9%) had a correlation less than 0.9. (C) A scatterplot of
# cell cycle gene \textit{CDK1} (ENSG00000131747) gene counts for hg19 (x-axis)
# versus hg38 (y-axis). The Pearson correlation across single cells was 0.99542.
# (D) A scatterplot of cell cycle gene \textit{UBE2C} (ENSG00000170312) gene
# counts for hg19 (x-axis) versus hg38 (y-axis). The Pearson correlation across
# single cells was 0.99814. (E) A scatterplot of cell cycle gene \textit{TOP2A}
# (ENSG00000175063) gene counts for hg19 (x-axis) versus hg38 (y-axis). The
# Pearson correlation across single cells was 0.99948. (F) A scatterplot of cell
# cycle gene \textit{HIST1H4E} (ENSG00000197061) gene counts for hg19 (x-axis)
# versus hg38 (y-axis). The Pearson correlation across single cells was 0.99998.
# The dashed red line represents the 1-1 line. All plots used data for the
# 10,297 protein-coding genes that were shared between genome builds hg19 and
# hg38. Note that cell cycle gene \textit{HIST1H4C} (ENSG00000198518) was
# deprecated in the annotation for genome build hg38.
```
