## Supplementary material for "Characterizing and inferring quantitative cell cycle phase in single-cell RNA-seq data analysis": Source code of our analysis, including scripts and data necessary to reproduce the work.: suppfig19.R

```
# Supplemental Figure 19
#   FUCCI and DAPI intensity before C1 plate effect correction of the 888 samples in the data.

library(SingleCellExperiment)
library(circular)
library(peco)
library(cluster)
library(ggplot2)

sce <- readRDS("data/sce-final.rds")
pdata = data.frame(colData(sce))


# split samples into 40 bins based on DAPI intensity distribution
# then plot GFP/RFP distribution in the 40 bins
pdata %>%
  mutate(dapi_bins=cut((pdata$dapi.median.log10sum), breaks=40)) %>%
  group_by(dapi_bins) %>%
  ggplot(., aes(x = dapi_bins, y = (pdata$gfp.median.log10sum))) +
    geom_boxplot() +
    ylab("EGFP score") +
    scale_x_discrete("DAPI score (40 equally-spaced bins)",
        breaks = levels(cut((pdata$dapi.median.log10sum), breaks=40))[c(1,5,10,15,20,25,30,35,40)],
        labels = levels(cut((pdata$dapi.median.log10sum), breaks=40))[c(1,5,10,15,20,25,30,35,40)]) +
  theme_gray() + theme(axis.text.x = element_text(size = 10, angle = 90,
                                     vjust=.5, hjust=1))

# split samples into 40 bins based on DAPI intensity distribution
# then plot GFP/RFP distribution in the 40 bins
pdata %>%
  mutate(dapi_bins=cut((pdata$dapi.median.log10sum), breaks=40)) %>%
  group_by(dapi_bins) %>%
  ggplot(., aes(x = dapi_bins, y = (pdata$rfp.median.log10sum))) +
  geom_boxplot() +
  ylab("mCherry score") +
  scale_x_discrete("DAPI score (40 equally-spaced bins)",
                   breaks = levels(cut((pdata$dapi.median.log10sum), breaks=40))[c(1,5,10,15,20,25,30,35,40)],
                   labels = levels(cut((pdata$dapi.median.log10sum), breaks=40))[c(1,5,10,15,20,25,30,35,40)]) +
  theme_gray() + theme(axis.text.x = element_text(size = 10, angle = 90,
                                   vjust=.5, hjust=1))


pdata %>%
  mutate(dapi_bins=cut((pdata$dapi.median.log10sum.adjust), breaks=40)) %>%
  group_by(dapi_bins) %>%
  ggplot(., aes(x = dapi_bins, y = (pdata$gfp.median.log10sum.adjust))) +
  geom_boxplot() +
  ylab("EGFP score") +
  scale_x_discrete("DAPI score (40 equally-spaced bins)",
                   breaks = levels(cut((pdata$dapi.median.log10sum.adjust), breaks=40))[c(1,5,10,15,20,25,30,35,40)],
                   labels = levels(cut((pdata$dapi.median.log10sum.adjust), breaks=40))[c(1,5,10,15,20,25,30,35,40)]) +
  theme_gray() + theme(axis.text.x = element_text(size = 10, angle = 90,
                                   vjust=.5, hjust=1)) #+ theme_gray(base_size = 12)

# split samples into 40 bins based on DAPI intensity distribution
# then plot GFP/RFP distribution in the 40 bins
pdata %>%
  mutate(dapi_bins=cut((pdata$dapi.median.log10sum.adjust), breaks=40)) %>%
  group_by(dapi_bins) %>%
  ggplot(., aes(x = dapi_bins, y = (pdata$rfp.median.log10sum.adjust))) +
  geom_boxplot() +
  ylab("mCherry score") +
  scale_x_discrete("DAPI score (40 equally-spaced bins)",
                   breaks = levels(cut((pdata$dapi.median.log10sum.adjust), breaks=40))[c(1,5,10,15,20,25,30,35,40)],
                   labels = levels(cut((pdata$dapi.median.log10sum.adjust), breaks=40))[c(1,5,10,15,20,25,30,35,40)]) +
  theme_gray() + theme(axis.text.x = element_text(size = 10, angle = 90,
                                   vjust=.5, hjust=1))

#
# pdata %>%
#   mutate(dapi_bins=cut(10^(pdata$dapi.median.log10sum), breaks=40)) %>%
#   group_by(dapi_bins) %>%
#   ggplot(., aes(x = dapi_bins, y = (10^pdata$gfp.median.log10sum))) +
#   geom_boxplot() +
#   ylab("EGFP intensity") +
#   scale_x_discrete("DAPI intensity (40 equally-spaced bins)",
#                    breaks = levels(cut((10^pdata$dapi.median.log10sum.adjust), breaks=40))[c(1,5,10,15,20,25,30,35,40)],
#                    labels = levels(cut((10^pdata$dapi.median.log10sum), breaks=40))[c(1,5,10,15,20,25,30,35,40)]) +
#   theme(axis.text.x = element_text(size = 10, angle = 90,
#                                    vjust=.5, hjust=1))
#
# # split samples into 40 bins based on DAPI intensity distribution
# # then plot GFP/RFP distribution in the 40 bins
# pdata %>%
#   mutate(dapi_bins=cut(10^pdata$dapi.median.log10sum, breaks=40)) %>%
#   group_by(dapi_bins) %>%
#   ggplot(., aes(x = dapi_bins, y = 10^pdata$rfp.median.log10sum)) +
#   geom_boxplot() +
#   ylab("mCherry intensity") +
#   scale_x_discrete("DAPI intensity (40 equally-spaced bins)",
#                    breaks = levels(cut((10^pdata$dapi.median.log10sum), breaks=40))[c(1,5,10,15,20,25,30,35,40)],
#                    labels = levels(cut((10^pdata$dapi.median.log10sum), breaks=40))[c(1,5,10,15,20,25,30,35,40)]) +
#   theme(axis.text.x = element_text(size = 10, angle = 90,
#                                    vjust=.5, hjust=1))


# par(mfrow=c(1,2))
# # histogram displaying distribution of DAPI intensities
# hist(10^pdata$dapi.median.log10sum, breaks = 40,
#      xlab = "DAPI intensity", xlim = c(0, 2600),
#      main = "")
# # histogram displaying distribution of DAPI log10 sum corrected intensities
# hist((10^pdata$dapi.median.log10sum.adjust)*800, breaks = 40,
#      xlab = "DAPI intensity",
#      main = "DAPI intensities \n after background and batch correction")


# library(cowplot)
# pdata$experiment = factor(pdata$experiment)
# plot_grid(
#   ggplot(pdata,
#          aes(x=experiment, y=dapi.median.log10sum,
#              fill=experiment)) +
#     geom_violin(alpha=.5) + geom_boxplot(width=.1, fill="gray90") +
#     ylab("") + xlab("Individual") +
#     ggtitle("DAPI before batch-correction") + labs(fill="experiment") +
#     theme(axis.text.x  = element_text(angle=90, vjust=1, hjust=1,size=10),
#           axis.text.y  = element_text(angle=0, vjust=0.2, size=10)) +
#     theme(legend.position="none"),
#   ggplot(pdata,
#          aes(x=experiment, y=dapi.median.log10sum.adjust, fill=experiment)) +
#     geom_violin(alpha=.5) + geom_boxplot(width=.1, fill="gray90") +
#     ylab("") + xlab("Individual") +
#     ggtitle("DAPI after batch-correction") + labs(fill="experiment") +
#     theme(axis.text.x  = element_text(angle=90, vjust=1, hjust=1,size=10),
#           axis.text.y  = element_text(angle=0, vjust=0.2, size=10)) +
#     theme(legend.position="none"))


# compute spearman correlation between DAPI intensities and GFP/RFP intensities
cor.test(pdata$dapi.median.log10sum.adjust,
         pdata$gfp.median.log10sum.adjust, method = "spearman")
cor.test(pdata$dapi.median.log10sum.adjust,
         pdata$rfp.median.log10sum.adjust, method = "spearman")
cor.test(pdata$dapi.median.log10sum.adjust,
         pdata$gfp.median.log10sum.adjust, method = "pearson")
cor.test(pdata$dapi.median.log10sum.adjust,
         pdata$rfp.median.log10sum.adjust, method = "pearson")

cor.test(pdata$gfp.median.log10sum.adjust,
         pdata$rfp.median.log10sum.adjust, method = "pearson")


# Compute FUCCI phase, do a PCA rotation
ints <- data.frame(rfp=colData(sce)$rfp.median.log10sum.adjust,
                   gfp=colData(sce)$gfp.median.log10sum.adjust)
pca <- prcomp(ints)
theta <- coord2rad(pca$x)
theta_final <- shift_origin(as.numeric(theta), origin = 3*pi/4)
```
