## Supplementary material for "Characterizing and inferring quantitative cell cycle phase in single-cell RNA-seq data analysis": Source code of our analysis, including scripts and data necessary to reproduce the work.: suppfig20.R

```
# Supplemental Figure 20
#   FUCCI scores for the six cell lines before and after correcting for C1 plate effect.

library(ggplot2)
library(RColorBrewer)
library(cowplot)
library(car)

sce <- readRDS("data/sce-final.rds")
counts <- assay(sce)
counts <- counts[grep("ENSG", rownames(counts)), ]
pdata <- data.frame(colData(sce))
pdata$chip_id <- factor(pdata$chip_id)

# Intensity batch effect ---------------------------------------------------------
lm.dapi <- lm(dapi.median.log10sum~factor(chip_id)+factor(experiment),
              data = pdata)

aov.lm.dapi <- Anova(lm.dapi, type = "III")
aov.lm.dapi

# before and after plate effect correction
plot_grid(
  ggplot(cdata,
         aes(x=chip_id, y=dapi.median.log10sum,
             col=chip_id)) +
    geom_violin() + geom_boxplot(width=.1) +
    theme(axis.text.x  = element_text(angle=90, vjust=1, hjust=1,size=10),
          axis.text.y  = element_text(angle=0, vjust=0.2, size=10)),
  ggplot(cdata,
         aes(x=factor(experiment), y=dapi.median.log10sum,
             col=factor(experiment))) +
    geom_violin() + geom_boxplot(width=.1) +
    theme(axis.text.x  = element_text(angle=90, vjust=1, hjust=1,size=10),
          axis.text.y  = element_text(angle=0, vjust=0.2, size=10)),
  ggplot(cdata,
         aes(x=chip_id, y=dapi.median.log10sum.adjust,
             col=chip_id)) +
    geom_violin() + geom_boxplot(width=.1) +
    theme(axis.text.x  = element_text(angle=90, vjust=1, hjust=1,size=10),
          axis.text.y  = element_text(angle=0, vjust=0.2, size=10)),
  ggplot(cdata,
         aes(x=factor(experiment), y=dapi.median.log10sum.adjust,
             col=factor(experiment))) +
    geom_violin() + geom_boxplot(width=.1) +
    theme(axis.text.x  = element_text(angle=90, vjust=1, hjust=1,size=10),
          axis.text.y  = element_text(angle=0, vjust=0.2, size=10)),
  labels = LETTERS[1:4])
```
