## Supplementary material for "Characterizing and inferring quantitative cell cycle phase in single-cell RNA-seq data analysis": Source code of our analysis, including scripts and data necessary to reproduce the work.: suppfig21.R

```
# Supplemental Figure S21
# Author: PoYuan Tung

library(SingleCellExperiment)
library(MASS)
library(ggplot2)
library(cowplot)


sce_raw <- readRDS("data/sce-raw.rds")
pdata <- data.frame(colData(sce_raw))

group_3 <- rep("2",dim(pdata)[1])
group_3[grep("0", pdata$cell_number)] <- "0"
group_3[grep("1", pdata$cell_number)] <- "1"


# Supp Fig S20A-C: Total molecule vs concentration
# LDA model: no. cells per well ~ gene molecules + cDNA concentration

data <- pdata %>% dplyr::select(experiment:concentration, mapped, molecules)
data <- data.frame(data, group = group_3)
data_lda <- lda(group ~ concentration + molecules, data = data)
data_lda_p <- predict(data_lda, newdata = data[,c("concentration", "molecules")])$class

outliers_lda <- data %>% rownames_to_column("sample_id") %>%
  filter(cell_number == 1, data_lda_p == "two")
pdata$molecule_outlier <- row.names(pdata) %in% outliers_lda$sample_id

## determine how well the model fix
table(data_lda_p, data[, "group"])
data$data_lda_p <- data_lda_p
data$data_lda_p <- factor(data_lda_p, levels = c(0, 1, 2))

## Metrics by the inferred number of cells
p1 <- ggplot(data, aes(y = molecules / 10^3, x = concentration, color=data_lda_p)) +
  geom_point(aes(color=data_lda_p), alpha=.6, cex=.6) +
  scale_color_manual(values = RColorBrewer::brewer.pal(3, "Set2"),
                     name = "Est. number of cells per well") +
  labs(x = "cDNA concentration", y = "Gene molecules (thousands)") +
  theme_bw()

p2 <- ggplot(data, aes(y = molecules / 10^3, x = data_lda_p)) +
  geom_violin(aes(fill = data_lda_p)) +
  geom_boxplot(width=.1) +
  scale_fill_manual(values = RColorBrewer::brewer.pal(3, "Set2"),
                    name = "Number of cells per well") +
  labs(x = "Estimated number of cells", y = "Total molecules (thousands)") +
  theme_bw()

p3 <- ggplot(data, aes(y = concentration, x = as.factor(data_lda_p))) +
  geom_violin(aes(fill = as.factor(data_lda_p))) +
  geom_boxplot(width=.1) +
  scale_fill_manual(values = RColorBrewer::brewer.pal(3, "Set2"), name = "Number of cells per well") +
  labs(x = "Estimated number of cells", y = "cDNA Concentration (ng/ul)") +
  theme_bw()
plot_grid(p1 + theme(legend.position = "none"),
          p2 + theme(legend.position = "none"),
          p3 + theme(legend.position = "none"), ncol = 3, labels = LETTERS[1:3])


# Supp Fig S12D-F: Reads to molecule conversion
# LDA model: no. cell per well ~ ERCC conversion efficiency + endogeneous gene conversion efficiency

## calculate convertion
pdata$ercc_conversion <- pdata$mol_ercc / pdata$reads_ercc
pdata$conversion <- pdata$mol_hs / pdata$reads_hs

data$conversion <- pdata$conversion
data$ercc_conversion <- pdata$ercc_conversion

data_ercc_lda <- lda(group ~ ercc_conversion + conversion, data = data)
data_ercc_lda_p <- predict(data_ercc_lda,  newdata = data[,c("ercc_conversion", "conversion")])$class

table(data_ercc_lda_p, data[, "group"])
data$data_ercc_lda_p <- data_ercc_lda_p
data$data_ercc_lda_p <- factor(data_ercc_lda_p, levels = c(0, 1, 2))

p1 <- ggplot(data, aes(y = conversion, x = ercc_conversion, color=data_ercc_lda_p)) +
  geom_point(aes(color=data_ercc_lda_p), alpha=.6, cex=.6) +
  scale_color_manual(values = RColorBrewer::brewer.pal(3, "Set2"),
                     name = "Est. number of cells per well") +
  labs(x = "ERCC conversion efficiency", y = "Endogeneous conversion efficiency") +
  theme_bw() + xlim(0, .38) + ylim(0, .38)

p2 <- ggplot(data, aes(y = conversion, x = data_ercc_lda_p)) +
  geom_violin(aes(fill = data_ercc_lda_p)) +
  geom_boxplot(width=.1) +
  scale_fill_manual(values = RColorBrewer::brewer.pal(3, "Set2"), name = "Number of cells per well") + ylim(0, .38) +
  labs(x = "Estimated number of cells", y = "Endogeneous gene \n read-to-molecule conversion efficiency") +
  theme_bw()

p3 <- ggplot(data, aes(y = ercc_conversion, x = data_ercc_lda_p)) +
  geom_violin(aes(fill = data_ercc_lda_p)) +
  geom_boxplot(width=.1) +
  scale_fill_manual(values = RColorBrewer::brewer.pal(3, "Set2"), name = "Number of cells per well") + ylim(0, .38) +
  labs(x = "Estimated number of cells", y = "ERCC \n read-to-molecule conversion efficiency") +
  theme_bw()

plot_grid(p1 + theme(legend.position="none"),
          p2 + theme(legend.position="none"),
          p3 + theme(legend.position="none"), ncol=3, labels = LETTERS[4:6])
```
