## Supplementary material for "Characterizing and inferring quantitative cell cycle phase in single-cell RNA-seq data analysis": Source code of our analysis, including scripts and data necessary to reproduce the work.: utility.R

```
#' @title Perform Principal Components Analysis (PCA).
#
#' @param x gene-by-sample matrix
#' @param retx center, scale - see ?prcomp
#
#' @return a list with the following elements:
#'   PCs - sample-by-PC matrix of principal components
#'   explained - proportion of variance explained by each PC
#'
#' @author John Blischak
#'
#' @references Zhang et al. 2009 (http://www.ncbi.nlm.nih.gov/pubmed/19763933)
#'
#' @export
run_pca <- function(x, retx = TRUE, center = TRUE, scale = TRUE) {
  library("testit")

  pca <- prcomp(t(x), retx = retx, center = center, scale. = scale)
  variances <- pca$sdev^2
  explained <- variances / sum(variances)
  assert("Variance explained is calculated correctly.",
         explained[1:2] - summary(pca)$importance[2, 1:2] < 0.0001)
  return(list(PCs = pca$x, explained = explained))
}

#' @title Plot PCA results.
#
#' @param x numeric matrix of PCs
#' @param pcx PC to plot on x-axis (default: 1)
#' @param pcy PC to plot on y-axis (default: 2)
#' @param explained numeric vector of fractions of variance explained by each PC
#' @param metadata data frame or matrix that contains the metadata used to annotate
#             the plot
#' @param color, shape, size: column name of metadata used to pass column to ggplot
#                       aesthetic
#' @param factors character vector which contains the column names of metadata that
#            need to be explicitly converted to a factor
#' @param ...  Additional arguments passed to geom_point
#'
#' @author John Blischak
#'
#' @export
plot_pca <- function(x, pcx = 1, pcy = 2, explained = NULL, metadata = NULL,
                     color = NULL, shape = NULL, factors = NULL,
                     ...) {
  library("ggplot2")
  library("testit")

  # Prepare data frame to pass to ggplot
  if (!is.null(metadata)) {
    assert("PC and metadata have same number of rows.",
           nrow(x) == nrow(metadata))
    plot_data <- cbind(x, metadata)
    plot_data <- as.data.frame(plot_data)
    # Convert numeric factors to class "factor"
    for (f in factors) {
      plot_data[, f] <- as.factor(plot_data[, f])
    }
  } else {
    plot_data <- as.data.frame(x)
  }
  # Prepare axis labels
  if (!is.null(explained)) {
    assert("Number of PCs differs between x and explained.",
           length(explained) == ncol(x))
    xaxis <- sprintf("PC%d (%.2f%%)", pcx, round(explained[pcx] * 100, 2))
    yaxis <- sprintf("PC%d (%.2f%%)", pcy, round(explained[pcy] * 100, 2))
  } else {
    xaxis <- paste0("PC", pcx)
    yaxis <- paste0("PC", pcy)
  }
  # Plot
  p <- ggplot(plot_data, aes_string(x = paste0("PC", pcx),
                                    y = paste0("PC", pcy),
                                    color = color,
                                    shape = shape)) +
    geom_point(...) +
    labs(x = xaxis, y = yaxis)
  p
}
```
