## Supplementary material for "Characterizing and inferring quantitative cell cycle phase in single-cell RNA-seq data analysis": Source code of our analysis, including scripts and data necessary to reproduce the work.: about.html


peco-paper

- Home
- About
- License

- Source code


### About

workflowr

- Summary
- Checks
- Past versions

**Last updated:** 2019-09-13

**Checks:**  2  0

**Knit directory:** `peco-paper/`

This reproducible R Markdown analysis was created with workflowr (version 1.4.0). The *Checks* tab describes the reproducibility checks that were applied when the results were created. The *Past versions* tab lists the development history.

---

**R Markdown file:** up-to-date

Great! Since the R Markdown file has been committed to the Git repository, you know the exact version of the code that produced these results.

**Repository version:** fc6d3f2

Great! You are using Git for version control. Tracking code development and connecting the code version to the results is critical for reproducibility. The version displayed above was the version of the Git repository at the time these results were generated.   
  
 Note that you need to be careful to ensure that all relevant files for the analysis have been committed to Git prior to generating the results (you can use `wflow_publish` or `wflow_git_commit`). workflowr only checks the R Markdown file, but you know if there are other scripts or data files that it depends on. Below is the status of the Git repository when the results were generated:

```
Ignored files:
    Ignored:    .Rhistory
    Ignored:    .Rproj.user/
```

Note that any generated files, e.g. HTML, png, CSS, etc., are not included in this status report because it is ok for generated content to have uncommitted changes.

---

These are the previous versions of the R Markdown and HTML files. If you’ve configured a remote Git repository (see `?wflow_git_remote`), click on the hyperlinks in the table below to view them.

| File | Version | Author | Date | Message |
| --- | --- | --- | --- | --- |
| Rmd | fc6d3f2 | jhsiao999 | 2019-09-13 | fix biorxiv link |
| html | bf95f68 | jhsiao999 | 2019-09-13 | Build site. |
| Rmd | 33b9a34 | jhsiao999 | 2019-09-13 | updates |
| html | 2d3a990 | jhsiao999 | 2019-09-13 | Build site. |
| html | ba2d647 | Joyce Hsiao | 2019-08-14 | Build site. |
| Rmd | 63517ff | Joyce Hsiao | 2019-08-14 | Start workflowr project. |

---

We combined fluorescence imaging with scRNA-seq to measure cell cycle phase and gene expression levels in human induced pluripotent stem cells (iPSCs). Using these data, we developed a novel approach to characterize cell cycle progression. While standard methods assign cells to discrete cell cycle stages, our method goes beyond this, and quantifies cell cycle progression on a continuum.

Find out more about our study in our paper.

- bioRxiv: Characterizing and inferring quantitative cell-cycle phase in single-cell RNA-seq data analysis
