## Supplementary material for "Characterizing and inferring quantitative cell cycle phase in single-cell RNA-seq data analysis": Source code of our analysis, including scripts and data necessary to reproduce the work.: about_data.html


peco-paper

- Home
- About
- License

- Source code


### About data

workflowr

- Summary
- Checks
- Past versions

**Last updated:** 2019-09-04

**Checks:**  6  1

**Knit directory:** `peco-paper/`

This reproducible R Markdown analysis was created with workflowr (version 1.4.0). The *Checks* tab describes the reproducibility checks that were applied when the results were created. The *Past versions* tab lists the development history.

---

**R Markdown file:** uncommitted changes

The R Markdown is untracked by Git. To know which version of the R Markdown file created these results, you’ll want to first commit it to the Git repo. If you’re still working on the analysis, you can ignore this warning. When you’re finished, you can run `wflow_publish` to commit the R Markdown file and build the HTML.

**Environment:** empty

Great job! The global environment was empty. Objects defined in the global environment can affect the analysis in your R Markdown file in unknown ways. For reproduciblity it’s best to always run the code in an empty environment.

**Seed:** `set.seed(20190814)`

The command `set.seed(20190814)` was run prior to running the code in the R Markdown file. Setting a seed ensures that any results that rely on randomness, e.g. subsampling or permutations, are reproducible.

**Session information:** recorded

Great job! Recording the operating system, R version, and package versions is critical for reproducibility.

**Cache:** none

Nice! There were no cached chunks for this analysis, so you can be confident that you successfully produced the results during this run.

**File paths:** relative

Great job! Using relative paths to the files within your workflowr project makes it easier to run your code on other machines.

**Repository version:** 9d74622

Great! You are using Git for version control. Tracking code development and connecting the code version to the results is critical for reproducibility. The version displayed above was the version of the Git repository at the time these results were generated.   
  
 Note that you need to be careful to ensure that all relevant files for the analysis have been committed to Git prior to generating the results (you can use `wflow_publish` or `wflow_git_commit`). workflowr only checks the R Markdown file, but you know if there are other scripts or data files that it depends on. Below is the status of the Git repository when the results were generated:

```
Ignored files:
    Ignored:    .Rhistory
    Ignored:    .Rproj.user/

Untracked files:
    Untracked:  analysis/about_data.Rmd
    Untracked:  analysis/rnaseq_processing.Rmd
    Untracked:  data/peco_ordered_cyclic_genes.txt

Unstaged changes:
    Modified:   analysis/index.Rmd
```

Note that any generated files, e.g. HTML, png, CSS, etc., are not included in this status report because it is ok for generated content to have uncommitted changes.

---

There are no past versions. Publish this analysis with `wflow_publish()` to start tracking its development.

---

On this page, you can find out how we processed imaging data and RNA-seq data.

- processing our FUCCI and DAPI images here and corrected for C1 batch effects in FUCCI intensity data
- quality control of scRNA-seq data
- find all the files used in the paper here

  

Session information

```
sessionInfo()
```

```
R version 3.5.1 (2018-07-02)
Platform: x86_64-pc-linux-gnu (64-bit)
Running under: Scientific Linux 7.4 (Nitrogen)

Matrix products: default
BLAS/LAPACK: /software/openblas-0.2.19-el7-x86_64/lib/libopenblas_haswellp-r0.2.19.so

locale:
 [1] LC_CTYPE=en_US.UTF-8       LC_NUMERIC=C              
 [3] LC_TIME=en_US.UTF-8        LC_COLLATE=en_US.UTF-8    
 [5] LC_MONETARY=en_US.UTF-8    LC_MESSAGES=en_US.UTF-8   
 [7] LC_PAPER=en_US.UTF-8       LC_NAME=C                 
 [9] LC_ADDRESS=C               LC_TELEPHONE=C            
[11] LC_MEASUREMENT=en_US.UTF-8 LC_IDENTIFICATION=C       

attached base packages:
[1] stats     graphics  grDevices utils     datasets  methods   base     

loaded via a namespace (and not attached):
 [1] workflowr_1.4.0 Rcpp_1.0.1      digest_0.6.19   rprojroot_1.3-2
 [5] backports_1.1.2 git2r_0.25.2    magrittr_1.5    evaluate_0.12  
 [9] stringi_1.2.4   fs_1.3.1        rmarkdown_1.10  tools_3.5.1    
[13] stringr_1.3.1   glue_1.3.0      yaml_2.2.0      compiler_3.5.1 
[17] htmltools_0.3.6 knitr_1.20
```
