## Supplementary material for "Characterizing and inferring quantitative cell cycle phase in single-cell RNA-seq data analysis": Source code of our analysis, including scripts and data necessary to reproduce the work.: access_data.html


peco-paper

- Home
- About
- License

- Source code


### Access data

###### *John Blischak, PoYuan Tung, Joyce Hsiao*

workflowr

- Summary
- Checks
- Past versions

**Last updated:** 2020-01-26

**Checks:**  7  0

**Knit directory:** `peco-paper/`

This reproducible R Markdown analysis was created with workflowr (version 1.6.0). The *Checks* tab describes the reproducibility checks that were applied when the results were created. The *Past versions* tab lists the development history.

---

**R Markdown file:** up-to-date

Great! Since the R Markdown file has been committed to the Git repository, you know the exact version of the code that produced these results.

**Environment:** empty

Great job! The global environment was empty. Objects defined in the global environment can affect the analysis in your R Markdown file in unknown ways. For reproduciblity it’s best to always run the code in an empty environment.

**Seed:** `set.seed(20190814)`

The command `set.seed(20190814)` was run prior to running the code in the R Markdown file. Setting a seed ensures that any results that rely on randomness, e.g. subsampling or permutations, are reproducible.

**Session information:** recorded

Great job! Recording the operating system, R version, and package versions is critical for reproducibility.

**Cache:** none

Nice! There were no cached chunks for this analysis, so you can be confident that you successfully produced the results during this run.

**File paths:** relative

Great job! Using relative paths to the files within your workflowr project makes it easier to run your code on other machines.

**Repository version:** 7ab2cab

Great! You are using Git for version control. Tracking code development and connecting the code version to the results is critical for reproducibility. The version displayed above was the version of the Git repository at the time these results were generated.   
  
 Note that you need to be careful to ensure that all relevant files for the analysis have been committed to Git prior to generating the results (you can use `wflow_publish` or `wflow_git_commit`). workflowr only checks the R Markdown file, but you know if there are other scripts or data files that it depends on. Below is the status of the Git repository when the results were generated:

```
Ignored files:
    Ignored:    .Rhistory
    Ignored:    .Rproj.user/

Untracked files:
    Untracked:  code/note_wo_w_pca.R
    Untracked:  data/intensity.rds
    Untracked:  data/log2cpm.quant.rds
    Untracked:  data/ourdata_phase_cyclone.rds
    Untracked:  data/ourdata_phase_seurat.rds

Unstaged changes:
    Modified:   analysis/index.Rmd
    Modified:   code/run_seurat.R
```

Note that any generated files, e.g. HTML, png, CSS, etc., are not included in this status report because it is ok for generated content to have uncommitted changes.

---

These are the previous versions of the R Markdown and HTML files. If you’ve configured a remote Git repository (see `?wflow_git_remote`), click on the hyperlinks in the table below to view them.

| File | Version | Author | Date | Message |
| --- | --- | --- | --- | --- |
| Rmd | 7ab2cab | jhsiao999 | 2020-01-26 | updates |
| html | f468a43 | jhsiao999 | 2020-01-23 | Build site. |
| Rmd | 86d7297 | jhsiao999 | 2020-01-23 | update link to output-raw-2-final.R |
| html | 99d0556 | jhsiao999 | 2020-01-23 | Build site. |
| Rmd | 39f5a01 | jhsiao999 | 2020-01-23 | data processing steps |
| html | bf95f68 | jhsiao999 | 2019-09-13 | Build site. |
| html | 2d3a990 | jhsiao999 | 2019-09-13 | Build site. |
| Rmd | 60e3281 | jhsiao999 | 2019-09-13 | wflow\_publish(c(“analysis/index.Rmd”, “analysis/access\_data.Rmd”, “analysis/license.Rmd”, |

---

Here are some useful links for how we processed our data.

- Correct for C1 batch effects in FUCCI intensity data.
- Assess potential C1 batch effects in sequencing depth and read to molecule conversion rate.
- Identify high quality samples using sample quality metrics, such as total number of mapped reads, ERCC percentages, etc.
- Filter out over-expressed and lowly-expressed genes.
- Use Principal Component Analysis to examined potential sources of gene expression variation.

  

Session information

```
R version 3.5.1 (2018-07-02)
Platform: x86_64-pc-linux-gnu (64-bit)
Running under: Scientific Linux 7.4 (Nitrogen)

Matrix products: default
BLAS/LAPACK: /software/openblas-0.2.19-el7-x86_64/lib/libopenblas_haswellp-r0.2.19.so

locale:
 [1] LC_CTYPE=en_US.UTF-8       LC_NUMERIC=C              
 [3] LC_TIME=en_US.UTF-8        LC_COLLATE=en_US.UTF-8    
 [5] LC_MONETARY=en_US.UTF-8    LC_MESSAGES=en_US.UTF-8   
 [7] LC_PAPER=en_US.UTF-8       LC_NAME=C                 
 [9] LC_ADDRESS=C               LC_TELEPHONE=C            
[11] LC_MEASUREMENT=en_US.UTF-8 LC_IDENTIFICATION=C       

attached base packages:
[1] stats     graphics  grDevices utils     datasets  methods   base     

loaded via a namespace (and not attached):
 [1] workflowr_1.6.0 Rcpp_1.0.3      digest_0.6.20   later_0.7.5    
 [5] rprojroot_1.3-2 R6_2.4.0        backports_1.1.2 git2r_0.26.1   
 [9] magrittr_1.5    evaluate_0.12   stringi_1.2.4   fs_1.3.1       
[13] promises_1.0.1  whisker_0.3-2   rmarkdown_1.10  tools_3.5.1    
[17] stringr_1.3.1   glue_1.3.0      httpuv_1.4.5    yaml_2.2.0     
[21] compiler_3.5.1  htmltools_0.3.6 knitr_1.20
```
