## Supplementary material for "Characterizing and inferring quantitative cell cycle phase in single-cell RNA-seq data analysis": Source code of our analysis, including scripts and data necessary to reproduce the work.: eval_on_leng2015_data.html

Compare the performance of peco with other methods using Leng et al. 2015 data


peco-paper

- Home
- About
- License

- Source code


### Compare the performance of peco with other methods using Leng et al. 2015 data

workflowr

- Summary
- Checks
- Past versions

**Last updated:** 2020-01-26

**Checks:**  7  0

**Knit directory:** `peco-paper/`

This reproducible R Markdown analysis was created with workflowr (version 1.6.0). The *Checks* tab describes the reproducibility checks that were applied when the results were created. The *Past versions* tab lists the development history.

---

**R Markdown file:** up-to-date

Great! Since the R Markdown file has been committed to the Git repository, you know the exact version of the code that produced these results.

**Environment:** empty

Great job! The global environment was empty. Objects defined in the global environment can affect the analysis in your R Markdown file in unknown ways. For reproduciblity it’s best to always run the code in an empty environment.

**Seed:** `set.seed(20190814)`

The command `set.seed(20190814)` was run prior to running the code in the R Markdown file. Setting a seed ensures that any results that rely on randomness, e.g. subsampling or permutations, are reproducible.

**Session information:** recorded

Great job! Recording the operating system, R version, and package versions is critical for reproducibility.

**Cache:** none

Nice! There were no cached chunks for this analysis, so you can be confident that you successfully produced the results during this run.

**File paths:** relative

Great job! Using relative paths to the files within your workflowr project makes it easier to run your code on other machines.

**Repository version:** cfbce75

Great! You are using Git for version control. Tracking code development and connecting the code version to the results is critical for reproducibility. The version displayed above was the version of the Git repository at the time these results were generated.   
  
 Note that you need to be careful to ensure that all relevant files for the analysis have been committed to Git prior to generating the results (you can use `wflow_publish` or `wflow_git_commit`). workflowr only checks the R Markdown file, but you know if there are other scripts or data files that it depends on. Below is the status of the Git repository when the results were generated:

```
Ignored files:
    Ignored:    .Rhistory
    Ignored:    .Rproj.user/
    Ignored:    analysis/figure/

Untracked files:
    Untracked:  analysis/npreg_trendfilter_quantile.Rmd
    Untracked:  data/HumanLengESC.rds
    Untracked:  data/data_training_test/
    Untracked:  data/eset-filtered.rds
    Untracked:  data/fit.quant.rds
    Untracked:  data/fit.trend.perm.lowmiss.rds
    Untracked:  data/fit_diff_cyclone.rds
    Untracked:  data/fit_diff_oscope.rds
    Untracked:  data/fit_diff_peco.rds
    Untracked:  data/fit_diff_recat.rds
    Untracked:  data/fit_diff_seurat.rds
    Untracked:  data/intensity.rds
    Untracked:  data/leng2015_data.rds
    Untracked:  data/leng_fucci_oscope_29genes.rda
    Untracked:  data/leng_fucci_recat.rda
    Untracked:  data/leng_geneinfo.txt
    Untracked:  data/log2cpm.quant.rds
    Untracked:  data/macosko-2015.rds
    Untracked:  data/nmeth.3549-S2.xlsx
    Untracked:  data/ourdata_cyclone_NA18511.rds
    Untracked:  data/ourdata_cyclone_NA18855.rds
    Untracked:  data/ourdata_cyclone_NA18870.rds
    Untracked:  data/ourdata_cyclone_NA19098.rds
    Untracked:  data/ourdata_cyclone_NA19101.rds
    Untracked:  data/ourdata_cyclone_NA19160.rds
    Untracked:  data/ourdata_oscope_366genes.rda
    Untracked:  data/ourdata_peco_NA18511_top005genes.rds
    Untracked:  data/ourdata_peco_NA18855_top005genes.rds
    Untracked:  data/ourdata_peco_NA18870_top005genes.rds
    Untracked:  data/ourdata_peco_NA19098_top005genes.rds
    Untracked:  data/ourdata_peco_NA19101_top005genes.rds
    Untracked:  data/ourdata_peco_NA19160_top005genes.rds
    Untracked:  data/ourdata_phase_cyclone.rds
    Untracked:  data/ourdata_phase_seurat.rds
    Untracked:  data/ourdata_recat.rda
    Untracked:  data/sce-filtered.rds

Unstaged changes:
    Modified:   analysis/index.Rmd
    Modified:   code/run_seurat.R
```

Note that any generated files, e.g. HTML, png, CSS, etc., are not included in this status report because it is ok for generated content to have uncommitted changes.

---

These are the previous versions of the R Markdown and HTML files. If you’ve configured a remote Git repository (see `?wflow_git_remote`), click on the hyperlinks in the table below to view them.

| File | Version | Author | Date | Message |
| --- | --- | --- | --- | --- |
| Rmd | cfbce75 | jhsiao999 | 2020-01-26 | Leng et al. 2015 data results |

---

#### Introduction

In Leng et al. 2015, FUCCI reporters were used to sort single-cell samples (Human Embryonic Stem Cells) into G1, S and G2M phases. The single-cell samples in each phase were then captured separated in 3 Fluidigm C1 plates.

We used this datat to assess the ability of peco to predict cell cycle phase against a “ground truth”. Specifically, we compared the performance of peco with Oscope and reCAT in predicting continuous cell cycle phase. We also compared the reuslts of Seurat and Oscope with the Leng et al. gating-based classification and our PAM-based discrete classification.

#### Set up

Load packages

```
library(SingleCellExperiment)
library(peco)
library(yarrr)
library(tidyverse)
library(SingleCellExperiment)
library(peco)
```

Getting data

```
# ------ getting data
sce <- readRDS("data/leng2015_data.rds")
pdata <- data.frame(colData(sce))
fdata <- data.frame(rowData(sce))
counts <- assay(sce)

sce_fucci <- sce[,pdata$cell_state != "H1"]

# compute quantile-normalized gene expression levels
sce_fucci <- data_transform_quantile(sce_fucci)
```

```
computing on 2 cores
```

```
counts_fucci <- assay(sce_fucci, "counts")
cpm_quantNormed_fucci <- assay(sce_fucci, "cpm_quantNormed")

pdata_fucci <- data.frame(colData(sce_fucci))
pdata_fucci$cell_state <- droplevels(pdata_fucci$cell_state)
pdata_fucci$cell_state <- factor(pdata_fucci$cell_state,
                                      levels = c("G1", "S", "G2"),
                                      labels = c("G1", "S", "G2M"))
```

#### peco prediction

```
# load training data
library(peco)
data("sce_top101genes")
data("training_human")


# Use top 4 genes for HIST1H4E was deprecated in genome build hg38
gene_symbols <- c("CDK1", "UBE2C", "TOP2A", "HIST1H4C")
gene_ensg <- c("ENSG00000170312","ENSG00000175063",
           "ENSG00000131747", "ENSG00000197061")
# use top 101 genes to predict
training_fun = training_human$cellcycle_function[gene_ensg]
training_sigma = training_human$sigma[gene_ensg,]


# use get_trend_estimateds=TRUE to estimate cyclic expression levels of 
# the four genes in Leng et al. 2015 data
out_peco <- cycle_npreg_outsample(
  Y_test=sce_fucci[rownames(sce_fucci) %in% gene_symbols],
  funs_est=training_fun,
  sigma_est=training_sigma,
  method.trend="trendfilter",
  ncores=4,
  get_trend_estimates=TRUE)
```

```
computing on 4 cores
```

```
subdata_plot <- do.call(rbind, lapply(1:4, function(g) {
  gindex <- which(rownames(out_peco$Y_reordered) == gene_symbols[g])
  gexp <- out_peco$Y_reordered[gindex,]
  data.frame(gexp=gexp, gene=gene_symbols[g], cell_state=pdata_fucci$cell_state)
}))
ggplot(subdata_plot, aes(x=cell_state, y = gexp, fill=cell_state)) +
  geom_boxplot() + facet_wrap(~gene, ncol=4) +
  ylab("Quantile-normalized \n gene expression levels") + xlab("") +
  scale_fill_manual(values=as.character(yarrr::piratepal("espresso")[3:1]),
                    name="Phase") + theme_bw()
```

```
par(mfrow=c(1,4), mar=c(2,2,2,1), oma = c(0,0,2,0))

all.equal(colnames(out_peco$Y_reordered), rownames(pdata_fucci))
```

```
[1] "246 string mismatches"
```

```
cell_state_col <- data.frame(cell_state = pdata_fucci$cell_state[match(colnames(out_peco$Y_reordered),
                                                                            rownames(pdata_fucci))],
                             sample_id = pdata_fucci$sample_id[match(colnames(out_peco$Y_reordered),
                                                                            rownames(pdata_fucci))])
cell_state_col$cols <- as.character(yarrr::piratepal("espresso")[3:1])[cell_state_col$cell_state]

for (g in 1:4) {
  if (g>1) { ylab <- ""} else { ylab <- "Quantile-normalized expression value"}
  #    if (i==1) { xlab <- "Predicted phase"} else { xlab <- "Inferred phase"}
  gindex <- which(rownames(out_peco$Y_reordered) == gene_symbols[g])

  plot(x= out_peco$cell_times_reordered,
       y = out_peco$Y_reordered[gindex,], pch = 16, cex=.5, ylab=ylab,
       xlab="Predicted phase", col = cell_state_col$cols,
       main = gene_symbols[g], axes=F, ylim=c(-3,3))
  abline(h=0, lwd=.5)
  lines(x = seq(0, 2*pi, length.out=200),
        y = out_peco$funs_reordered[[g]](seq(0, 2*pi, length.out=200)),
        col=wesanderson::wes_palette("FantasticFox1")[1], lty=1, lwd=2)
  axis(2); axis(1,at=c(0,pi/2, pi, 3*pi/2, 2*pi),
                labels=c(0,expression(pi/2), expression(pi), expression(3*pi/2),
                         expression(2*pi)))
}
title("peco prediction results", outer = TRUE, line = .5)
```

#### Oscope prediction

We applied Oscope to estimate cell cycle ordering across the 888 single-cell samples. The analysis used the 29 genes that were identified as oscillating cell cycle genes in Leng et al. 2015 (doi:10.1038/nmeth.3549). The data file was downloaed from nmeth.3549-S2.xlsx.

```
# Oscope
library(openxlsx)
oscope29genes <- unlist(read.xlsx("data/nmeth.3549-S2.xlsx", colNames = F))

# run Oscope on FUCCI cells ------------------------------
library(Oscope)
Sizes <- MedianNorm(counts_fucci)
DataNorm <- GetNormalizedMat(counts_fucci, Sizes)
DataNorm_sub <- DataNorm[which(rownames(DataNorm) %in% oscope29genes),]
DataInput <- NormForSine(DataNorm_sub)
SineRes <- OscopeSine(DataInput, parallel=F)
KMResUse <- list(cluster1=oscope29genes)
ENIRes <- OscopeENI(KMRes = KMResUse, Data = DataInput,
                    NCThre = 100, parallel=T)

save(DataInput,
     KMResUse, ENIRes, fits_leng_oscope, pdata_fucci,
     file = "data/leng_fucci_oscope_29genes.rda")
```

```
# load pre-computed results,
load("data/leng_fucci_oscope_29genes.rda")

# lot predicted results
gene_symbols <- c("CDK1", "UBE2C", "TOP2A", "HIST1H4C")
gene_ensg <- c("ENSG00000170312","ENSG00000175063",
           "ENSG00000131747", "ENSG00000197061")

phase_pred <- seq(0, 2*pi, length.out= length(ENIRes$cluster1))
names(phase_pred) <- colnames(DataInput)[ENIRes[["cluster1"]]]

fits_leng_oscope <- lapply(1:4, function(g) {
  gindex <- which(rownames(cpm_quantNormed_fucci) == gene_symbols[g])
  fit_g <- data.frame(
    gexp=cpm_quantNormed_fucci[gindex, match(names(phase_pred), colnames(cpm_quantNormed_fucci))],
    phase=shift_origin((-phase_pred+2*pi), origin = pi/2))
  fit_g <- fit_g[order(fit_g$phase),]

  fit_trend <- fit_trendfilter_generic(fit_g$gexp)
  fit_g$trend.yy <- fit_trend$trend.yy
  fun_g <- approxfun(x=as.numeric(fit_g$phase),
                     y=as.numeric(fit_g$trend.yy), rule=2)
  fit_out <- list(fit_g=fit_g,
                  pve = fit_trend$pve,
                  fun_g = fun_g)
  return(fit_out)
})
```

```
Fold 1 ... Fold 2 ... Fold 3 ... Fold 4 ... Fold 5 ... 
Fold 1 ... Fold 2 ... Fold 3 ... Fold 4 ... Fold 5 ... 
Fold 1 ... Fold 2 ... Fold 3 ... Fold 4 ... Fold 5 ... 
Fold 1 ... Fold 2 ... Fold 3 ... Fold 4 ... Fold 5 ...
```

```
names(fits_leng_oscope) <- gene_symbols


all.equal(rownames(fits_leng_oscope[[1]]$fit_g), rownames(pdata_fucci))
```

```
[1] "247 string mismatches"
```

```
cell_state_oscope_col <- 
    data.frame(cell_state = pdata_fucci$cell_state[match(rownames(fits_leng_oscope[[1]]$fit_g),
                                               rownames(pdata_fucci))],
              sample_id = rownames(fits_leng_oscope[[1]]$fit_g) )
cell_state_oscope_col$cols <-
    as.character(yarrr::piratepal("espresso")[3:1])[cell_state_oscope_col$cell_state]


par(mfrow=c(1,4), mar=c(2,2,2,1), oma = c(0,0,2,1))
for (g in 1:4) {
  if (g>1) { ylab <- ""} else { ylab <- "Normalized log2CPM"}
  #    if (i==1) { xlab <- "Predicted phase"} else { xlab <- "Inferred phase"}
  res_g <- fits_leng_oscope[[g]]
  plot(x= res_g$fit_g$phase,
       y = res_g$fit_g$gexp, pch = 16, cex=.5, ylab=ylab,
       xlab="Predicted phase", col=cell_state_oscope_col$cols,
       main = names(fits_leng_oscope)[g], axes=F, ylim=c(-3,3))
  abline(h=0, lwd=.5)
  lines(x = seq(0, 2*pi, length.out=200),
        y = res_g$fun_g(seq(0, 2*pi, length.out=200)),
        col = wesanderson::wes_palette("Darjeeling2")[2], lty=1, lwd=2)
  axis(2); axis(1,at=c(0,pi/2, pi, 3*pi/2, 2*pi),
                labels=c(0,expression(pi/2), expression(pi), expression(3*pi/2),
                         expression(2*pi)))
}
title("Oscope prediction results", outer = TRUE, line = .5)
```

#### reCAT prediction

We applied reCAT to estimate cell cycle ordering across the 888 single-cell samples. The analysis used all the 11,040 genes.

To run recAT, please clone the reCAT GitHUb repository and then cd to the directory.

`git clone https://github.com/tinglab/reCAT`

`cd "reCAT/R"`

```
# recat
input <- t(cpm_quantNormed_fucci)
input <- input[sample(1:nrow(input)),]
source("get_test_exp.R")
test_exp <- get_test_exp(t(input))
source("get_ordIndex.R")
res_ord <- get_ordIndex(test_exp, 10)
ordIndex <- res_ord$ordIndex

save(test_exp, res_ord, ordIndex, "data/leng_fucci_recat.rda")
```

```
# load pre-computed results
load("data/leng_fucci_recat.rda")

sample_ordered <- rownames(test_exp)[ordIndex]
phase_pred <- seq(0, 2*pi, length.out= length(ordIndex))
names(phase_pred) <- sample_ordered

fits_recat <- lapply(1:4, function(g) {
  gindex <- which(rownames(cpm_quantNormed_fucci) == gene_symbols[g])
  fit_g <- data.frame(
    gexp=cpm_quantNormed_fucci[gindex, match(sample_ordered, colnames(cpm_quantNormed_fucci))],
    phase=(-phase_pred)+2*pi )
  fit_g <- fit_g[order(fit_g$phase),]

  fit_trend <- fit_trendfilter_generic(fit_g$gexp, polyorder = 2)
  fit_g$trend.yy <- fit_trend$trend.yy
  #    fit_g$pve <- fit$pve
  fun_g <- approxfun(x=as.numeric(fit_g$phase),
                     y=as.numeric(fit_g$trend.yy), rule=2)
  fit_out <- list(fit_g=fit_g,
                  #                  pve = fit$pve,
                  fun_g = fun_g)
  return(fit_out)
})
```

```
Fold 1 ... Fold 2 ... Fold 3 ... Fold 4 ... Fold 5 ... 
Fold 1 ... Fold 2 ... Fold 3 ... Fold 4 ... Fold 5 ... 
Fold 1 ... Fold 2 ... Fold 3 ... Fold 4 ... Fold 5 ... 
Fold 1 ... Fold 2 ... Fold 3 ... Fold 4 ... Fold 5 ...
```

```
names(fits_recat) <- gene_symbols

all.equal(rownames(fits_recat[[1]]$fit_g), rownames(pdata_fucci))
```

```
[1] "246 string mismatches"
```

```
cell_state_recat_col <- 
    data.frame(cell_state = pdata_fucci$cell_state[match(rownames(fits_recat[[1]]$fit_g),
                                                       rownames(pdata_fucci))],
               sample_id = rownames(fits_recat[[1]]$fit_g) )
cell_state_recat_col$cols <- 
    as.character(yarrr::piratepal("espresso")[3:1])[cell_state_recat_col$cell_state]

par(mfrow=c(1,4), mar=c(2,2,2,1), oma = c(0,0,2,0) )
for (g in 1:4) {
  if (g>1) { ylab <- ""} else { ylab <- "Quantile-normalized expression value"}
  res_g <- fits_recat[[g]]
  plot(x= res_g$fit_g$phase,
       y = res_g$fit_g$gexp, pch = 16, cex=.5, ylab=ylab,
       xlab="Predicted phase", col = cell_state_recat_col$cols,
       main = names(fits_recat)[g], axes=F, ylim=c(-3,3))
  abline(h=0, lwd=.5)
  lines(x = seq(0, 2*pi, length.out=200),
        y = res_g$fun_g(seq(0, 2*pi, length.out=200)),
        col=wesanderson::wes_palette("FantasticFox1")[5], lty=1, lwd=2)
  axis(2); axis(1,at=c(0,pi/2, pi, 3*pi/2, 2*pi),
                labels=c(0,expression(pi/2), expression(pi), expression(3*pi/2),
                         expression(2*pi)))
}
title("reCAT prediction results", outer = TRUE, line = .5)
```

#### Seurat

```
library(Seurat)
```

```
Attaching package: 'Seurat'
```

```
The following object is masked from 'package:SummarizedExperiment':

    Assays
```

```
cc.genes <- readLines(con = "data//regev_lab_cell_cycle_genes.txt")

obj <- CreateSeuratObject(counts = counts_fucci)
obj <- NormalizeData(obj)
obj <- FindVariableFeatures(obj, selection.method = "vst")
obj <- ScaleData(obj, features = rownames(obj))
```

```
Centering and scaling data matrix
```

```
obj <- CellCycleScoring(obj, s.features = cc.genes[1:43], 
                        g2m.features = cc.genes[44:97], set.ident = TRUE)
out_seurat <- obj[[]]

all.equal(rownames(out_seurat), colnames(counts_fucci))
```

```
[1] TRUE
```

```
out_seurat <- out_seurat[match(colnames(counts_fucci), rownames(out_seurat)),]

out_seurat$Phase <- factor(out_seurat$Phase,
                           levels = c("G1", "S", "G2M"))
out_seurat$gates <- pdata_fucci$cell_state[match(rownames(out_seurat),
                                                       rownames(pdata_fucci))]
table(out_seurat$gates, out_seurat$Phase)
```

```
      G1  S G2M
  G1  10 28  53
  S   20 60   0
  G2M 11  5  60
```

#### Cyclone

```
library(scran)
hs.pairs <- readRDS(system.file("exdata", "human_cycle_markers.rds", package="scran"))

## get ENSG ID
# library(biomaRt)
# mart <- useMart(biomart = "ensembl", dataset = "hsapiens_gene_ensembl")
# geneinfo <- getBM(attributes = c("hgnc_symbol", "ensembl_gene_id"),
#                filters = "hgnc_symbol",
#                values = rownames(cpm_quantNormed_fucci), bmHeader = T, mart = mart)
#write.table(geneinfo, file = "data/leng_geneinfo.txt"))

geneinfo <- read.table("data/leng_geneinfo.txt")


indata <- which(geneinfo$ensg %in% unlist(hs.pairs))
geneinfo_indata <- geneinfo[indata,]

Y_cyclone <- cpm_quantNormed_fucci[rownames(cpm_quantNormed_fucci) %in% as.character(geneinfo_indata$symbol),]
rownames(Y_cyclone) <- geneinfo_indata$ensg[match(rownames(Y_cyclone),
                                                  geneinfo_indata$symbol)]

out_cyclone <- cyclone(Y_cyclone, pairs = hs.pairs,
                       gene.names=rownames(Y_cyclone),
                       iter=1000, min.iter=100, min.pairs=50,
                       BPPARAM=SerialParam(), verbose=T, subset.row=NULL)
```

```
Number of G1 pairs: 21666
```

```
Number of S pairs: 26446
```

```
Number of G2M pairs: 18738
```

```
names(out_cyclone$phases) <- colnames(Y_cyclone)
all.equal(names(out_cyclone$phases), colnames(cpm_quantNormed_fucci))
```

```
[1] TRUE
```

```
all.equal(names(out_cyclone$phases), rownames(pdata_fucci))
```

```
[1] TRUE
```

```
out_cyclone$cell_state <- pdata_fucci$cell_state[match(names(out_cyclone$phases),
                                                       rownames(pdata_fucci))]
out_cyclone$phases <- factor(out_cyclone$phases,
                             levels = c("G1", "S", "G2M"))
table(pdata_fucci$cell_state, out_cyclone$phases)
```

```
      G1  S G2M
  G1  91  0   0
  S    0 78   2
  G2M  0  0  76
```

#### Session information

```
sessionInfo()
```

```
R version 3.5.1 (2018-07-02)
Platform: x86_64-pc-linux-gnu (64-bit)
Running under: Scientific Linux 7.4 (Nitrogen)

Matrix products: default
BLAS/LAPACK: /software/openblas-0.2.19-el7-x86_64/lib/libopenblas_haswellp-r0.2.19.so

locale:
 [1] LC_CTYPE=en_US.UTF-8       LC_NUMERIC=C              
 [3] LC_TIME=en_US.UTF-8        LC_COLLATE=en_US.UTF-8    
 [5] LC_MONETARY=en_US.UTF-8    LC_MESSAGES=en_US.UTF-8   
 [7] LC_PAPER=en_US.UTF-8       LC_NAME=C                 
 [9] LC_ADDRESS=C               LC_TELEPHONE=C            
[11] LC_MEASUREMENT=en_US.UTF-8 LC_IDENTIFICATION=C       

attached base packages:
[1] parallel  stats4    stats     graphics  grDevices utils     datasets 
[8] methods   base     

other attached packages:
 [1] scran_1.10.2                Seurat_3.1.0               
 [3] forcats_0.3.0               stringr_1.3.1              
 [5] dplyr_0.8.0.1               purrr_0.3.2                
 [7] readr_1.3.1                 tidyr_0.8.3                
 [9] tibble_2.1.1                ggplot2_3.2.1              
[11] tidyverse_1.2.1             yarrr_0.1.5                
[13] circlize_0.4.8              BayesFactor_0.9.12-4.2     
[15] Matrix_1.2-17               coda_0.19-2                
[17] jpeg_0.1-8                  peco_0.99.10               
[19] SingleCellExperiment_1.4.1  SummarizedExperiment_1.12.0
[21] DelayedArray_0.8.0          BiocParallel_1.16.0        
[23] matrixStats_0.55.0          Biobase_2.42.0             
[25] GenomicRanges_1.34.0        GenomeInfoDb_1.18.1        
[27] IRanges_2.16.0              S4Vectors_0.20.1           
[29] BiocGenerics_0.28.0        

loaded via a namespace (and not attached):
  [1] reticulate_1.10          R.utils_2.7.0           
  [3] tidyselect_0.2.5         htmlwidgets_1.3         
  [5] grid_3.5.1               Rtsne_0.15              
  [7] munsell_0.5.0            codetools_0.2-15        
  [9] ica_1.0-2                statmod_1.4.30          
 [11] future_1.14.0            withr_2.1.2             
 [13] colorspace_1.3-2         knitr_1.20              
 [15] rstudioapi_0.10          ROCR_1.0-7              
 [17] gbRd_0.4-11              listenv_0.7.0           
 [19] Rdpack_0.11-0            labeling_0.3            
 [21] git2r_0.26.1             GenomeInfoDbData_1.2.0  
 [23] rhdf5_2.26.2             rprojroot_1.3-2         
 [25] generics_0.0.2           circular_0.4-93         
 [27] R6_2.4.0                 doParallel_1.0.14       
 [29] ggbeeswarm_0.6.0         rsvd_1.0.0              
 [31] conicfit_1.0.4           locfit_1.5-9.1          
 [33] bitops_1.0-6             assertthat_0.2.1        
 [35] promises_1.0.1           SDMTools_1.1-221.1      
 [37] scales_1.0.0             beeswarm_0.2.3          
 [39] gtable_0.2.0             npsurv_0.4-0            
 [41] globals_0.12.4           workflowr_1.6.0         
 [43] rlang_0.4.0              MatrixModels_0.4-1      
 [45] GlobalOptions_0.1.0      splines_3.5.1           
 [47] lazyeval_0.2.1           genlasso_1.4            
 [49] broom_0.5.1              yaml_2.2.0              
 [51] reshape2_1.4.3           modelr_0.1.2            
 [53] backports_1.1.2          httpuv_1.4.5            
 [55] tools_3.5.1              gplots_3.0.1            
 [57] RColorBrewer_1.1-2       dynamicTreeCut_1.63-1   
 [59] ggridges_0.5.1           Rcpp_1.0.3              
 [61] plyr_1.8.4               zlibbioc_1.28.0         
 [63] RCurl_1.95-4.11          pbapply_1.3-4           
 [65] viridis_0.5.1            cowplot_0.9.4           
 [67] zoo_1.8-4                haven_1.1.2             
 [69] ggrepel_0.8.0            cluster_2.0.7-1         
 [71] fs_1.3.1                 magrittr_1.5            
 [73] data.table_1.12.0        lmtest_0.9-36           
 [75] RANN_2.6.1               mvtnorm_1.0-11          
 [77] whisker_0.3-2            fitdistrplus_1.0-14     
 [79] hms_0.4.2                lsei_1.2-0              
 [81] evaluate_0.12            readxl_1.1.0            
 [83] gridExtra_2.3            shape_1.4.4             
 [85] compiler_3.5.1           scater_1.10.1           
 [87] KernSmooth_2.23-15       crayon_1.3.4            
 [89] R.oo_1.22.0              htmltools_0.3.6         
 [91] later_0.7.5              RcppParallel_4.4.3      
 [93] lubridate_1.7.4          MASS_7.3-51.1           
 [95] boot_1.3-20              wesanderson_0.3.6       
 [97] cli_1.1.0                R.methodsS3_1.7.1       
 [99] gdata_2.18.0             metap_1.1               
[101] igraph_1.2.2             pkgconfig_2.0.3         
[103] plotly_4.8.0             xml2_1.2.0              
[105] foreach_1.4.4            vipor_0.4.5             
[107] XVector_0.22.0           bibtex_0.4.2            
[109] rvest_0.3.2              digest_0.6.20           
[111] sctransform_0.2.0        RcppAnnoy_0.0.11        
[113] pracma_2.2.9             tsne_0.1-3              
[115] rmarkdown_1.10           cellranger_1.1.0        
[117] leiden_0.3.1             edgeR_3.24.0            
[119] uwot_0.1.3               geigen_2.3              
[121] DelayedMatrixStats_1.4.0 gtools_3.8.1            
[123] nlme_3.1-137             jsonlite_1.6            
[125] Rhdf5lib_1.4.3           BiocNeighbors_1.0.0     
[127] limma_3.38.3             viridisLite_0.3.0       
[129] pillar_1.3.1             lattice_0.20-38         
[131] httr_1.3.1               survival_2.43-1         
[133] glue_1.3.0               png_0.1-7               
[135] iterators_1.0.12         stringi_1.2.4           
[137] HDF5Array_1.10.1         caTools_1.17.1.1        
[139] irlba_2.3.3              future.apply_1.3.0      
[141] ape_5.2
```

  

Session information

```
sessionInfo()
```

```
R version 3.5.1 (2018-07-02)
Platform: x86_64-pc-linux-gnu (64-bit)
Running under: Scientific Linux 7.4 (Nitrogen)

Matrix products: default
BLAS/LAPACK: /software/openblas-0.2.19-el7-x86_64/lib/libopenblas_haswellp-r0.2.19.so

locale:
 [1] LC_CTYPE=en_US.UTF-8       LC_NUMERIC=C              
 [3] LC_TIME=en_US.UTF-8        LC_COLLATE=en_US.UTF-8    
 [5] LC_MONETARY=en_US.UTF-8    LC_MESSAGES=en_US.UTF-8   
 [7] LC_PAPER=en_US.UTF-8       LC_NAME=C                 
 [9] LC_ADDRESS=C               LC_TELEPHONE=C            
[11] LC_MEASUREMENT=en_US.UTF-8 LC_IDENTIFICATION=C       

attached base packages:
[1] parallel  stats4    stats     graphics  grDevices utils     datasets 
[8] methods   base     

other attached packages:
 [1] scran_1.10.2                Seurat_3.1.0               
 [3] forcats_0.3.0               stringr_1.3.1              
 [5] dplyr_0.8.0.1               purrr_0.3.2                
 [7] readr_1.3.1                 tidyr_0.8.3                
 [9] tibble_2.1.1                ggplot2_3.2.1              
[11] tidyverse_1.2.1             yarrr_0.1.5                
[13] circlize_0.4.8              BayesFactor_0.9.12-4.2     
[15] Matrix_1.2-17               coda_0.19-2                
[17] jpeg_0.1-8                  peco_0.99.10               
[19] SingleCellExperiment_1.4.1  SummarizedExperiment_1.12.0
[21] DelayedArray_0.8.0          BiocParallel_1.16.0        
[23] matrixStats_0.55.0          Biobase_2.42.0             
[25] GenomicRanges_1.34.0        GenomeInfoDb_1.18.1        
[27] IRanges_2.16.0              S4Vectors_0.20.1           
[29] BiocGenerics_0.28.0        

loaded via a namespace (and not attached):
  [1] reticulate_1.10          R.utils_2.7.0           
  [3] tidyselect_0.2.5         htmlwidgets_1.3         
  [5] grid_3.5.1               Rtsne_0.15              
  [7] munsell_0.5.0            codetools_0.2-15        
  [9] ica_1.0-2                statmod_1.4.30          
 [11] future_1.14.0            withr_2.1.2             
 [13] colorspace_1.3-2         knitr_1.20              
 [15] rstudioapi_0.10          ROCR_1.0-7              
 [17] gbRd_0.4-11              listenv_0.7.0           
 [19] Rdpack_0.11-0            labeling_0.3            
 [21] git2r_0.26.1             GenomeInfoDbData_1.2.0  
 [23] rhdf5_2.26.2             rprojroot_1.3-2         
 [25] generics_0.0.2           circular_0.4-93         
 [27] R6_2.4.0                 doParallel_1.0.14       
 [29] ggbeeswarm_0.6.0         rsvd_1.0.0              
 [31] conicfit_1.0.4           locfit_1.5-9.1          
 [33] bitops_1.0-6             assertthat_0.2.1        
 [35] promises_1.0.1           SDMTools_1.1-221.1      
 [37] scales_1.0.0             beeswarm_0.2.3          
 [39] gtable_0.2.0             npsurv_0.4-0            
 [41] globals_0.12.4           workflowr_1.6.0         
 [43] rlang_0.4.0              MatrixModels_0.4-1      
 [45] GlobalOptions_0.1.0      splines_3.5.1           
 [47] lazyeval_0.2.1           genlasso_1.4            
 [49] broom_0.5.1              yaml_2.2.0              
 [51] reshape2_1.4.3           modelr_0.1.2            
 [53] backports_1.1.2          httpuv_1.4.5            
 [55] tools_3.5.1              gplots_3.0.1            
 [57] RColorBrewer_1.1-2       dynamicTreeCut_1.63-1   
 [59] ggridges_0.5.1           Rcpp_1.0.3              
 [61] plyr_1.8.4               zlibbioc_1.28.0         
 [63] RCurl_1.95-4.11          pbapply_1.3-4           
 [65] viridis_0.5.1            cowplot_0.9.4           
 [67] zoo_1.8-4                haven_1.1.2             
 [69] ggrepel_0.8.0            cluster_2.0.7-1         
 [71] fs_1.3.1                 magrittr_1.5            
 [73] data.table_1.12.0        lmtest_0.9-36           
 [75] RANN_2.6.1               mvtnorm_1.0-11          
 [77] whisker_0.3-2            fitdistrplus_1.0-14     
 [79] hms_0.4.2                lsei_1.2-0              
 [81] evaluate_0.12            readxl_1.1.0            
 [83] gridExtra_2.3            shape_1.4.4             
 [85] compiler_3.5.1           scater_1.10.1           
 [87] KernSmooth_2.23-15       crayon_1.3.4            
 [89] R.oo_1.22.0              htmltools_0.3.6         
 [91] later_0.7.5              RcppParallel_4.4.3      
 [93] lubridate_1.7.4          MASS_7.3-51.1           
 [95] boot_1.3-20              wesanderson_0.3.6       
 [97] cli_1.1.0                R.methodsS3_1.7.1       
 [99] gdata_2.18.0             metap_1.1               
[101] igraph_1.2.2             pkgconfig_2.0.3         
[103] plotly_4.8.0             xml2_1.2.0              
[105] foreach_1.4.4            vipor_0.4.5             
[107] XVector_0.22.0           bibtex_0.4.2            
[109] rvest_0.3.2              digest_0.6.20           
[111] sctransform_0.2.0        RcppAnnoy_0.0.11        
[113] pracma_2.2.9             tsne_0.1-3              
[115] rmarkdown_1.10           cellranger_1.1.0        
[117] leiden_0.3.1             edgeR_3.24.0            
[119] uwot_0.1.3               geigen_2.3              
[121] DelayedMatrixStats_1.4.0 gtools_3.8.1            
[123] nlme_3.1-137             jsonlite_1.6            
[125] Rhdf5lib_1.4.3           BiocNeighbors_1.0.0     
[127] limma_3.38.3             viridisLite_0.3.0       
[129] pillar_1.3.1             lattice_0.20-38         
[131] httr_1.3.1               survival_2.43-1         
[133] glue_1.3.0               png_0.1-7               
[135] iterators_1.0.12         stringi_1.2.4           
[137] HDF5Array_1.10.1         caTools_1.17.1.1        
[139] irlba_2.3.3              future.apply_1.3.0      
[141] ape_5.2
```
