## Supplementary material for "Characterizing and inferring quantitative cell cycle phase in single-cell RNA-seq data analysis": Source code of our analysis, including scripts and data necessary to reproduce the work.: eval_on_our_data.html

Comparison of performances on our data


peco-paper

- Home
- About
- License

- Source code


### Comparison of performances on our data

workflowr

- Summary
- Checks
- Past versions

**Last updated:** 2020-01-26

**Checks:**  7  0

**Knit directory:** `peco-paper/`

```
Ignored files:
    Ignored:    .Rhistory
    Ignored:    .Rproj.user/

Untracked files:
    Untracked:  analysis/eval_on_leng2015_data.Rmd
    Untracked:  analysis/npreg_trendfilter_quantile.Rmd
    Untracked:  data/HumanLengESC.rds
    Untracked:  data/data_training_test/
    Untracked:  data/eset-filtered.rds
    Untracked:  data/fit.quant.rds
    Untracked:  data/fit.trend.perm.lowmiss.rds
    Untracked:  data/fit_diff_cyclone.rds
    Untracked:  data/fit_diff_oscope.rds
    Untracked:  data/fit_diff_peco.rds
    Untracked:  data/fit_diff_recat.rds
    Untracked:  data/fit_diff_seurat.rds
    Untracked:  data/intensity.rds
    Untracked:  data/leng2015_data.rds
    Untracked:  data/leng_fucci_oscope_29genes.rda
    Untracked:  data/leng_fucci_recat.rda
    Untracked:  data/leng_geneinfo.txt
    Untracked:  data/log2cpm.quant.rds
    Untracked:  data/macosko-2015.rds
    Untracked:  data/nmeth.3549-S2.xlsx
    Untracked:  data/ourdata_cyclone_NA18511.rds
    Untracked:  data/ourdata_cyclone_NA18855.rds
    Untracked:  data/ourdata_cyclone_NA18870.rds
    Untracked:  data/ourdata_cyclone_NA19098.rds
    Untracked:  data/ourdata_cyclone_NA19101.rds
    Untracked:  data/ourdata_cyclone_NA19160.rds
    Untracked:  data/ourdata_oscope_366genes.rda
    Untracked:  data/ourdata_peco_NA18511_top005genes.rds
    Untracked:  data/ourdata_peco_NA18855_top005genes.rds
    Untracked:  data/ourdata_peco_NA18870_top005genes.rds
    Untracked:  data/ourdata_peco_NA19098_top005genes.rds
    Untracked:  data/ourdata_peco_NA19101_top005genes.rds
    Untracked:  data/ourdata_peco_NA19160_top005genes.rds
    Untracked:  data/ourdata_phase_cyclone.rds
    Untracked:  data/ourdata_phase_seurat.rds
    Untracked:  data/ourdata_recat.rda
    Untracked:  data/sce-filtered.rds

Unstaged changes:
    Modified:   analysis/index.Rmd
    Modified:   code/run_seurat.R
```

Note that any generated files, e.g. HTML, png, CSS, etc., are not included in this status report because it is ok for generated content to have uncommitted changes.

---

These are the previous versions of the R Markdown and HTML files. If you’ve configured a remote Git repository (see `?wflow_git_remote`), click on the hyperlinks in the table below to view them.

| File | Version | Author | Date | Message |
| --- | --- | --- | --- | --- |
| Rmd | c9afbb3 | jhsiao999 | 2020-01-26 | minor edits |
| html | 958a49a | jhsiao999 | 2020-01-25 | Build site. |
| Rmd | 2a13ee4 | jhsiao999 | 2020-01-25 | Comparisons of predictions on our data |

---

#### Introduction

We compare the performance of peco with Seurat, Cyclone, Oscope and reCAT using our data in six-fold cross-validation. Here we provide the code that we used to predict continuous cel cycle phase in each method and to compute prediction error between FUCCI phase and predicted cell cycle phase.

#### Set up

Load packages

```
library(SingleCellExperiment)
library(peco)
```

Prepare training/testing data

```
sce <- readRDS("data/sce-final.rds")
sce <- sce[grep("ENSG", rownames(sce)),]
fdata <- data.frame(colData(sce))
fdata <- data.frame(rowData(sce))
counts <- data.frame(assay(sce, "counts"))

sce_normed <- data_transform_quantile(sce)
log2cpm_quant <- assay(sce_normed, "cpm_quantNormed")

inds <- c("NA18511", "NA18855", "NA18870", "NA19098", "NA19101", "NA19160")
theta <- pdata$theta

# function to make training/testing data
makedata_supervised <- function(sce, log2cpm_quant,
                                theta) {
  message("Create data/data_training_test folder \n")
  if (!file.exists("data/data_training_test")) { dir.create("data/data_training_test") }
  library(SingleCellExperiment)
  library(peco)
  pdata <- data.frame(colData(sce))
  fdata <- data.frame(rowData(sce))
  counts <- data.frame(assay(sce))
  counts <- counts[grep("ENSG", rownames(counts)), ]
  log2cpm <- t(log2(1+(10^6)*(t(counts)/pdata$molecules)))

  for (ind in unique(pdata$chip_id)) {
    ii_test <- c(1:nrow(pdata))[which(pdata$chip_id == ind)]
    ii_train <- c(1:nrow(pdata))[which(pdata$chip_id != ind)]

    pdata_test <- pdata[ii_test,]
    pdata_train <- pdata[ii_train,]

    log2cpm_quant_test <- log2cpm_quant[,ii_test]
    log2cpm_quant_train <- log2cpm_quant[,ii_train]
    theta <- pdata$theta
    names(theta) <- rownames(pdata)

    log2cpm_test <- log2cpm[,ii_test]
    log2cpm_train <- log2cpm[,ii_train]

    counts_test <- counts[,ii_test]
    counts_train <- counts[,ii_train]

    theta_test <- theta[ii_test]
    theta_train <- theta[ii_train]

    #sig.genes <- readRDS("output/npreg-trendfilter-quantile.Rmd/out.stats.ordered.sig.476.rds")
    data_training <- list(theta_train=theta_train,
                          log2cpm_quant_train=log2cpm_quant_train,
                          log2cpm_train=log2cpm_train,
                          counts_train=counts_train,
                          pdata_train=pdata_train,
                          fdata=fdata)

    data_test <- list(theta_test=theta_test,
                      log2cpm_quant_test=log2cpm_quant_test,
                      log2cpm_test=log2cpm_test,
                      counts_test = counts_test,
                      pdata_test=pdata_test,
                      fdata=fdata)

    saveRDS(data_training,
            file=file.path(paste0("data/data_training_test/ind_",ind,"_data_training.rds")))
    saveRDS(data_test,
            file=file.path(paste0("data/data_training_test/ind_",ind,"_data_test.rds")))
  }
}
makedata_supervised(sce, log2cpm_quant, theta)
```

#### peco

Use peco predictor of 5 genes to predict cell cycle phase.

```
# --- prediction for samples from one individual cell ine
ngenes = 5

for (i in seq_along(inds)) {
  ind = inds[i]
  data_train <- readRDS(paste0("data/data_training_test/ind_", ind, "_data_training.rds"))
  data_test <- readRDS(paste0("data/data_training_test/ind_",ind, "_data_test.rds"))

  fits_all <- readRDS("data/fit.quant.rds")
  genes_all <- names(fits_all)[order(sapply(fits_all,"[[",3), decreasing=T)]

  which_genes <- genes_all[1:ngenes]
  fit_train <- cycle_npreg_insample(
    Y = with(data_train, log2cpm_quant_train[which(rownames(log2cpm_quant_train) %in% which_genes), ]),
    theta = with(data_train, theta_train),
    polyorder=2,
    ncores=4,
    method.trend="trendfilter")

  fit_test <- cycle_npreg_outsample(
    Y_test=with(data_test, log2cpm_quant_test[which(rownames(log2cpm_quant_test) %in% which_genes), ]),
    sigma_est=with(fit_train, sigma_est),
    funs_est=with(fit_train, funs_est),
    method.grid = "uniform",
    method.trend="trendfilter",
    polyorder=2,
    ncores=2)

  out_peco <- list(fit_train=fit_train,
            fit_test=fit_test)
  saveRDS(out, file.path(paste0("data/ourdata_peco_",
                                ind, "_top", sprintf("%03d", ngenes),"genes.rds")))
}


# compute prediction error
inds <- c("NA19098","NA18511","NA18870","NA19101","NA18855","NA19160")
ngenes <- 5
diff_peco <- do.call(rbind, lapply(1:length(inds), function(i) {
  ind <- inds[i]
  fl_name <- list.files(file.path("data"),
                        pattern=paste0("ourdata_peco_", ind, "_top", sprintf("%03d", ngenes),"genes.rds"),
                        full.names = TRUE)
  res_pred <- readRDS(fl_name)$fit_test
  data_test <- readRDS(paste0("data/data_training_test/ind_",ind, "_data_test.rds"))
  all.equal(rownames(data_test$pdata_test), names(res_pred$cell_times_est))

  df_pred <- readRDS(fl_name)$fit_test
  phase_ref = data_test$theta_test
  phase_pred_rot = rotation(ref_var=data_test$theta_test, 
                          shift_var=res_pred$cell_times_est)

  return(data.frame(phase_ref = phase_ref,
                    phase_pred_rot = phase_pred_rot,
                    diff_time = circ_dist(phase_ref, phase_pred_rot),
                    ind = as.character(ind),
                    method = "peco"))
}))
diff_peco$ind = factor(diff_peco$ind, 
                       levels = c("NA18511", "NA18855", "NA18870", 
                                  "NA19098", "NA19101", "NA19160"))
saveRDS(diff_peco, file=("data/fit_diff_peco.rds"))

diff_peco %>% group_by(ind) %>% summarise(mn=mean(diff_time)/2/pi,
                                          sd=sd(diff_time/2/pi)/sqrt(sum(phase_pred_rot>0)))
```

#### Seurat

We computed Seurat cell cycle scores and transformed these two scores into continuous cell cycle phase. Specifically, we applied PCA to the two cell cycle scores and then assigend each cell an angle on a unit circle. We did this for samples of each individual cell lines separately.

```
# use Seurat to assign phase for each individual
# and then compute predictione error for samples in individual cell lines
diff_seurat <- do.call(rbind, lapply(1:length(inds), function(i) {
  source("code/run_seurat.R")
  seurat.genes <- readLines(con = "data/regev_lab_cell_cycle_genes.txt")
  seurat.genes <- list(s.genes=seurat.genes[1:43],
                       g2m.genes=seurat.genes[44:97])

  #  print(i)
  data <- readRDS(paste0("data/data_training_test/ind_",inds[i], "_data_test.rds"))
  Y_fit <- data$log2cpm_test
  rownames(Y_fit) <- fdata$name[match(rownames(Y_fit), rownames(fdata))]
  fit <- run_seurat(Y_fit,
                    s.genes=seurat.genes$s.genes,
                    g2m.genes=seurat.genes$g2m.genes)
  fit <- as.list(fit)
  seurat.pca <- prcomp(cbind(fit$G2M, fit$S), scale=TRUE)
  phase_pred <- intensity2circle(seurat.pca$x, plot.it = F, method = "trig")
  names(phase_pred) <- colnames(Y_fit)
  phase_ref <- pdata$theta[match(names(phase_pred), rownames(pdata))]
  phase_pred_rot <- rotation(ref_var=phase_ref, shift_var=phase_pred)
  return(data.frame(phase_ref = phase_ref,
                    phase_pred_rot = phase_pred_rot,
                    diff_time = circ_dist(phase_ref, phase_pred_rot),
                    ind = inds[i],
                    method = "Seurat"))
}) )
saveRDS(diff_seurat, file="data/fit_diff_seurat.rds")

diff_seurat %>% group_by(ind) %>% summarise(mn=mean(diff_time)/2/pi,
                                            sd=sd(diff_time/2/pi)/sqrt(sum(phase_pred_rot>0)))
```

#### Cyclone

We computed Cyclone cell cycle scores and transformed these two scores into continuous cell cycle phase. Specifically, we applied PCA to the two cell cycle scores and then assigend each cell an angle on a unit circle. We did this for samples of each individual cell lines separately.

```
library(scran)
hs.pairs <- readRDS(system.file("exdata", "human_cycle_markers.rds", package="scran"))

for (ind in unique(pData(eset)$chip_id)) {
   data <- readRDS(file.path(paste0("data/data_training_test/ind_",
                                    ind, "_data_test.rds")))
   input <- data.frame(data$log2cpm_quant_test)
   ii <- which(colSums(is.na(input))==0)
   input <- input[,ii]

   # --- begin cyclone
   hs.pairs <- readRDS(system.file("exdata", "human_cycle_markers.rds", package="scran"))
#   message("run cyclone", "\n")
   out <- cyclone(input, pairs = hs.pairs,
                  gene.names=rownames(input),
                  iter=1000, min.iter=100, min.pairs=50,
                  BPPARAM=SerialParam(), verbose=T, subset.row=NULL)

   out_cyclone <- data.frame(phase_cyclone = out$phases,
                             normalized.scores_G1 = out$normalized.scores$G1,
                             normalized.scores_S = out$normalized.scores$S,
                             normalized.scores_G2M = out$normalized.scores$G2M)
   rownames(out_cyclone) <- colnames(input)

   # Infer phase using Seurat
   pc_cyclone <- prcomp(out$normalized.scores, scale. = T, center = T)

   library(peco)
   phase_peco <- intensity2circle(pc_cyclone$x[,1:2], plot.it = F, method = "trig")
   out_cyclone$phase_pred <- rotation(data$theta_test,
                                          phase_peco)
   out_cyclone$phase_ref <- data$theta_test
   out_cyclone$ind <- ind

   saveRDS(out_cyclone, file = paste0("data/ourdata_cyclone_",
                                   ind,".rds")) 
}


# compute prediction error of cyclone results
inds <- c("NA19098","NA18511","NA18870","NA19101","NA18855","NA19160")
diff_cyclone <- do.call(rbind, lapply(1:length(inds), function(i) {
  out_cyclone <- readRDS(paste0("data/ourdata_cyclone_",inds[i],".rds"))
  phase_peco <- pdata$theta[match(rownames(out_cyclone), rownames(pdata))]
  phase_cyclone_rot <- rotation(ref_var= phase_peco, shift_var = out_cyclone$phase_pred)
  return(data.frame(phase_ref = phase_peco,
                    phase_pred_rot = phase_cyclone_rot,
                    diff_time = circ_dist(phase_peco, phase_cyclone_rot),
                    ind = inds[i],
                    method = "Cyclone"))
}) )
saveRDS(diff_cyclone, file="data/fit_diff_cyclone.rds")

diff_cyclone %>% group_by(ind) %>% summarise(mn=mean(diff_time)/2/pi,
                                             sd=sd(diff_time/2/pi)/sqrt(sum(phase_pred_rot>0)))
```

#### Oscope

We applied Oscope to estimate cell cycle ordering across the 888 single-cell samples. The analysis used 366 genes that were selected using Oscope’s criterion of high variabilty genes. Specifically, we changed the default ’MeanCutlow` option of 100 to 10. We also found that the results were similar when adding more genes in the analysis.

```
library(Oscope)

# to run Oscope
# use gene symbols as labels 
rownames(counts) <- fdata$name

Sizes <- MedianNorm(counts)
DataNorm <- GetNormalizedMat(counts, Sizes)
MV <- CalcMV(Data = as.matrix(counts), Sizes = Sizes, MeanCutLow = 10)
DataNorm_high_var <- DataNorm[MV$GeneToUse,]
DataInput <- NormForSine(DataNorm_high_var)
dim(DataInput)

# select samples from all individuals
SineRes <- OscopeSine(DataInput, parallel=T)
KMRes <- OscopeKM(SineRes, maxK = NULL)

# to flag clusters...
# no need to flag clusters
KMResUse <- KMRes
ENIRes <- OscopeENI(KMRes = KMResUse, Data = DataInput,
            NCThre = 100, parallel=T)

save(DataInput,
     KMResUse, ENIRes,
     file = file.path("data/ourdata_oscope_366genes.rda")


# Oscope results
load("data/ourdata_oscope_366genes.rda")
samples_ordered <- colnames(DataInput)[ENIRes[["cluster2"]]]

phase_pred <- seq(0, 2*pi, length.out= length(ENIRes[["cluster2"]]))
names(phase_pred) <- samples_ordered

diff_oscope <- data.frame(phase_ref=pdata$theta[match(samples_ordered, colnames(eset))])
diff_oscope$phase_pred_rot <- rotation(ref_var= diff_oscope$phase_ref, shift_var = phase_pred)
diff_oscope$diff_time <- circ_dist(diff_oscope$phase_ref, diff_oscope$phase_pred_rot)
diff_oscope$ind <- pdata$chip_id[match(names(phase_pred), colnames(counts))]
diff_oscope$method <- "Oscope"

saveRDS(diff_oscope, file="data/fit_diff_oscope.rds")

diff_oscope %>% group_by(ind) %>% summarise(mn=mean(diff_time)/2/pi,
                                            sd=sd(diff_time/2/pi)/sqrt(sum(phase_pred_rot>0)))
```

#### recAT

We applied reCAT to estimate cell cycle ordering across the 888 single-cell samples. The analysis used all the 11,040 genes.

To run recAT, please clone the reCAT GitHUb repository and then cd to the directory.

`git clone https://github.com/tinglab/reCAT`

`cd "reCAT/R"`

```
# recat
input <- t(log2cpm_quant)
input <- input[sample(1:nrow(input)),]
source("get_test_exp.R")
test_exp <- get_test_exp(t(input))
source("get_ordIndex.R")
res_ord <- get_ordIndex(test_exp, 10)
ordIndex <- res_ord$ordIndex

save(test_exp, ordIndex,
     file = file.path("data/ourdata_recat_rda"))

# compute predictor error of reCAT predicted time
sample_ordered <- rownames(test_exp)[ordIndex]
phase_pred <- seq(0, 2*pi, length.out= length(ordIndex))
names(phase_pred) <- sample_ordered

diff_recat <- data.frame(phase_ref=pdata$theta[match(samples_ordered, colnames(eset))])
diff_recat$ind <- pdata$chip_id[match(names(phase_pred), colnames(eset))]

diff_recat$phase_pred_rot <- rotation(ref_var= diff_recat$phase_ref, shift_var = phase_pred)
rownames(diff_recat) <- samples_ordered

diff_recat$diff_time <- circ_dist(diff_recat$phase_ref, diff_recat$phase_pred_rot)
diff_recat$method <- "reCAT"
diff_recat <- diff_recat[,c(1,3,4,2,5)]

saveRDS(diff_recat, file="data/fit_diff_recat.rds")

diff_recat %>% group_by(ind) %>% summarise(mn=mean(diff_time)/2/pi,
                                           sd=sd(diff_time/2/pi)/sqrt(sum(phase_pred_rot>0)))
```

#### Session information

```
sessionInfo()
```

```
R version 3.5.1 (2018-07-02)
Platform: x86_64-pc-linux-gnu (64-bit)
Running under: Scientific Linux 7.4 (Nitrogen)

Matrix products: default
BLAS/LAPACK: /software/openblas-0.2.19-el7-x86_64/lib/libopenblas_haswellp-r0.2.19.so

locale:
 [1] LC_CTYPE=en_US.UTF-8       LC_NUMERIC=C              
 [3] LC_TIME=en_US.UTF-8        LC_COLLATE=en_US.UTF-8    
 [5] LC_MONETARY=en_US.UTF-8    LC_MESSAGES=en_US.UTF-8   
 [7] LC_PAPER=en_US.UTF-8       LC_NAME=C                 
 [9] LC_ADDRESS=C               LC_TELEPHONE=C            
[11] LC_MEASUREMENT=en_US.UTF-8 LC_IDENTIFICATION=C       

attached base packages:
[1] parallel  stats4    stats     graphics  grDevices utils     datasets 
[8] methods   base     

other attached packages:
 [1] peco_0.99.6                 SingleCellExperiment_1.4.1 
 [3] SummarizedExperiment_1.12.0 DelayedArray_0.8.0         
 [5] BiocParallel_1.16.0         matrixStats_0.55.0         
 [7] Biobase_2.42.0              GenomicRanges_1.34.0       
 [9] GenomeInfoDb_1.18.1         IRanges_2.16.0             
[11] S4Vectors_0.20.1            BiocGenerics_0.28.0        

loaded via a namespace (and not attached):
 [1] viridis_0.5.1            genlasso_1.4            
 [3] viridisLite_0.3.0        foreach_1.4.4           
 [5] DelayedMatrixStats_1.4.0 assertthat_0.2.1        
 [7] vipor_0.4.5              GenomeInfoDbData_1.2.0  
 [9] yaml_2.2.0               pillar_1.3.1            
[11] backports_1.1.2          lattice_0.20-38         
[13] glue_1.3.0               digest_0.6.20           
[15] promises_1.0.1           XVector_0.22.0          
[17] colorspace_1.3-2         plyr_1.8.4              
[19] htmltools_0.3.6          httpuv_1.4.5            
[21] Matrix_1.2-17            pkgconfig_2.0.3         
[23] zlibbioc_1.28.0          purrr_0.3.2             
[25] mvtnorm_1.0-11           scales_1.0.0            
[27] HDF5Array_1.10.1         whisker_0.3-2           
[29] later_0.7.5              pracma_2.2.9            
[31] git2r_0.26.1             tibble_2.1.1            
[33] ggplot2_3.2.1            conicfit_1.0.4          
[35] lazyeval_0.2.1           magrittr_1.5            
[37] crayon_1.3.4             evaluate_0.12           
[39] fs_1.3.1                 doParallel_1.0.14       
[41] MASS_7.3-51.1            beeswarm_0.2.3          
[43] geigen_2.3               tools_3.5.1             
[45] scater_1.10.1            stringr_1.3.1           
[47] Rhdf5lib_1.4.3           munsell_0.5.0           
[49] compiler_3.5.1           rlang_0.4.0             
[51] rhdf5_2.26.2             grid_3.5.1              
[53] RCurl_1.95-4.11          iterators_1.0.12        
[55] circular_0.4-93          igraph_1.2.2            
[57] bitops_1.0-6             rmarkdown_1.10          
[59] boot_1.3-20              gtable_0.2.0            
[61] codetools_0.2-15         reshape2_1.4.3          
[63] R6_2.4.0                 gridExtra_2.3           
[65] knitr_1.20               dplyr_0.8.0.1           
[67] workflowr_1.6.0          rprojroot_1.3-2         
[69] stringi_1.2.4            ggbeeswarm_0.6.0        
[71] Rcpp_1.0.3               tidyselect_0.2.5
```

  

Session information

```
sessionInfo()
```

```
R version 3.5.1 (2018-07-02)
Platform: x86_64-pc-linux-gnu (64-bit)
Running under: Scientific Linux 7.4 (Nitrogen)

Matrix products: default
BLAS/LAPACK: /software/openblas-0.2.19-el7-x86_64/lib/libopenblas_haswellp-r0.2.19.so

locale:
 [1] LC_CTYPE=en_US.UTF-8       LC_NUMERIC=C              
 [3] LC_TIME=en_US.UTF-8        LC_COLLATE=en_US.UTF-8    
 [5] LC_MONETARY=en_US.UTF-8    LC_MESSAGES=en_US.UTF-8   
 [7] LC_PAPER=en_US.UTF-8       LC_NAME=C                 
 [9] LC_ADDRESS=C               LC_TELEPHONE=C            
[11] LC_MEASUREMENT=en_US.UTF-8 LC_IDENTIFICATION=C       

attached base packages:
[1] parallel  stats4    stats     graphics  grDevices utils     datasets 
[8] methods   base     

other attached packages:
 [1] peco_0.99.6                 SingleCellExperiment_1.4.1 
 [3] SummarizedExperiment_1.12.0 DelayedArray_0.8.0         
 [5] BiocParallel_1.16.0         matrixStats_0.55.0         
 [7] Biobase_2.42.0              GenomicRanges_1.34.0       
 [9] GenomeInfoDb_1.18.1         IRanges_2.16.0             
[11] S4Vectors_0.20.1            BiocGenerics_0.28.0        

loaded via a namespace (and not attached):
 [1] viridis_0.5.1            genlasso_1.4            
 [3] viridisLite_0.3.0        foreach_1.4.4           
 [5] DelayedMatrixStats_1.4.0 assertthat_0.2.1        
 [7] vipor_0.4.5              GenomeInfoDbData_1.2.0  
 [9] yaml_2.2.0               pillar_1.3.1            
[11] backports_1.1.2          lattice_0.20-38         
[13] glue_1.3.0               digest_0.6.20           
[15] promises_1.0.1           XVector_0.22.0          
[17] colorspace_1.3-2         plyr_1.8.4              
[19] htmltools_0.3.6          httpuv_1.4.5            
[21] Matrix_1.2-17            pkgconfig_2.0.3         
[23] zlibbioc_1.28.0          purrr_0.3.2             
[25] mvtnorm_1.0-11           scales_1.0.0            
[27] HDF5Array_1.10.1         whisker_0.3-2           
[29] later_0.7.5              pracma_2.2.9            
[31] git2r_0.26.1             tibble_2.1.1            
[33] ggplot2_3.2.1            conicfit_1.0.4          
[35] lazyeval_0.2.1           magrittr_1.5            
[37] crayon_1.3.4             evaluate_0.12           
[39] fs_1.3.1                 doParallel_1.0.14       
[41] MASS_7.3-51.1            beeswarm_0.2.3          
[43] geigen_2.3               tools_3.5.1             
[45] scater_1.10.1            stringr_1.3.1           
[47] Rhdf5lib_1.4.3           munsell_0.5.0           
[49] compiler_3.5.1           rlang_0.4.0             
[51] rhdf5_2.26.2             grid_3.5.1              
[53] RCurl_1.95-4.11          iterators_1.0.12        
[55] circular_0.4-93          igraph_1.2.2            
[57] bitops_1.0-6             rmarkdown_1.10          
[59] boot_1.3-20              gtable_0.2.0            
[61] codetools_0.2-15         reshape2_1.4.3          
[63] R6_2.4.0                 gridExtra_2.3           
[65] knitr_1.20               dplyr_0.8.0.1           
[67] workflowr_1.6.0          rprojroot_1.3-2         
[69] stringi_1.2.4            ggbeeswarm_0.6.0        
[71] Rcpp_1.0.3               tidyselect_0.2.5
```
