## Supplementary material for "Characterizing and inferring quantitative cell cycle phase in single-cell RNA-seq data analysis": Source code of our analysis, including scripts and data necessary to reproduce the work.: gene_filtering.html

Gene filtering and sample filtering


peco-paper

- Home
- About
- License

- Source code


### Gene filtering and sample filtering

###### *Joyce Hsiao*

workflowr

- Summary
- Checks
- Past versions

**Last updated:** 2020-01-23

**Checks:**  7  0

**Knit directory:** `peco-paper/`

```
Ignored files:
    Ignored:    .Rhistory
    Ignored:    .Rproj.user/

Untracked files:
    Untracked:  analysis/npreg_trendfilter_quantile.Rmd
    Untracked:  analysis/pca-tf.Rmd
    Untracked:  code/fig2_rev.R
    Untracked:  data/fit.quant.rds
    Untracked:  data/intensity.rds
    Untracked:  data/log2cpm.quant.rds

Unstaged changes:
    Modified:   analysis/access_data.Rmd
    Modified:   analysis/index.Rmd
    Modified:   code/fig2.R
```

Note that any generated files, e.g. HTML, png, CSS, etc., are not included in this status report because it is ok for generated content to have uncommitted changes.

---

These are the previous versions of the R Markdown and HTML files. If you’ve configured a remote Git repository (see `?wflow_git_remote`), click on the hyperlinks in the table below to view them.

| File | Version | Author | Date | Message |
| --- | --- | --- | --- | --- |
| Rmd | 66c350a | jhsiao999 | 2020-01-23 | move gene\_filtering.Rmd and change eset to sce |
| html | 6107391 | jhsiao999 | 2020-01-23 | Build site. |
| Rmd | 2d495fb | jhsiao999 | 2020-01-23 | move gene\_filtering.Rmd and change eset to sce |

---

---
