## Supplementary material for "Characterizing and inferring quantitative cell cycle phase in single-cell RNA-seq data analysis": Source code of our analysis, including scripts and data necessary to reproduce the work.: images_batchcorrect.html

Normalize intensities across C1 batches


peco-paper

- Home
- About
- License

- Source code


### Normalize intensities across C1 batches

###### *Joyce Hsiao*

workflowr

- Summary
- Checks
- Past versions

**Last updated:** 2020-03-14

**Checks:**  7  0

**Knit directory:** `peco-paper/`

```
Ignored files:
    Ignored:    .Rhistory
    Ignored:    .Rproj.user/

Untracked files:
    Untracked:  code/note_wo_w_pca.R
    Untracked:  code/run_seurat.R
    Untracked:  data/intensity.rds
    Untracked:  data/log2cpm.quant.rds
    Untracked:  data/ourdata_phase_cyclone.rds
    Untracked:  data/ourdata_phase_seurat.rds

Unstaged changes:
    Modified:   code/suppfig06.R
```

Note that any generated files, e.g. HTML, png, CSS, etc., are not included in this status report because it is ok for generated content to have uncommitted changes.

---

These are the previous versions of the R Markdown and HTML files. If you’ve configured a remote Git repository (see `?wflow_git_remote`), click on the hyperlinks in the table below to view them.

| File | Version | Author | Date | Message |
| --- | --- | --- | --- | --- |
| Rmd | 909764a | jhsiao999 | 2020-03-14 | add anova results post batch correction |
| html | 72c318c | jhsiao999 | 2020-01-12 | Build site. |
| Rmd | 060d627 | jhsiao999 | 2020-01-12 | correct for C1 batch effect in fluorescence intensities |

---

---
