## Supplementary material for "Characterizing and inferring quantitative cell cycle phase in single-cell RNA-seq data analysis": Source code of our analysis, including scripts and data necessary to reproduce the work.: index.html

peco


peco-paper

- Home
- About
- License

- Source code


### peco

- Overview
- The software peco
- The analysis
- Citation
- Downloading the data files
- Other information

workflowr

- Summary
- Checks
- Past versions

**Last updated:** 2020-01-26

```
Ignored files:
    Ignored:    .Rhistory
    Ignored:    .Rproj.user/

Untracked files:
    Untracked:  code/note_wo_w_pca.R
    Untracked:  data/intensity.rds
    Untracked:  data/log2cpm.quant.rds
    Untracked:  data/ourdata_phase_cyclone.rds
    Untracked:  data/ourdata_phase_seurat.rds

Unstaged changes:
    Modified:   code/run_seurat.R
```

Note that any generated files, e.g. HTML, png, CSS, etc., are not included in this status report because it is ok for generated content to have uncommitted changes.

---

These are the previous versions of the R Markdown and HTML files. If you’ve configured a remote Git repository (see `?wflow_git_remote`), click on the hyperlinks in the table below to view them.

| File | Version | Author | Date | Message |
| --- | --- | --- | --- | --- |
| Rmd | ea63cb6 | jhsiao999 | 2020-01-26 | homepage |
| Rmd | 206dbf5 | jhsiao999 | 2020-01-12 | updates |
| html | 35b41c2 | jhsiao999 | 2019-09-16 | Build site. |
| Rmd | 944d881 | jhsiao999 | 2019-09-16 | fix code/ hyperlink |
| html | bf95f68 | jhsiao999 | 2019-09-13 | Build site. |
| Rmd | 33b9a34 | jhsiao999 | 2019-09-13 | updates |
| html | 2d3a990 | jhsiao999 | 2019-09-13 | Build site. |
| Rmd | 60e3281 | jhsiao999 | 2019-09-13 | wflow\_publish(c(“analysis/index.Rmd”, “analysis/access\_data.Rmd”, “analysis/license.Rmd”, |
| html | 60a309f | jhsiao999 | 2019-09-06 | Build site. |
| Rmd | 37ffff6 | jhsiao999 | 2019-09-06 | update table of content |
| html | ba2d647 | Joyce Hsiao | 2019-08-14 | Build site. |
| Rmd | 63517ff | Joyce Hsiao | 2019-08-14 | Start workflowr project. |

---

#### Overview

**peco** is a supervised approach for predicting *continuous* cell cycle phase using single-cell RNA-seq (scRNA-seq) data. The approach is described in our paper.

We use this site to document and share the code used to produce our analysis. Please feel free to explore. Comments and feedbacks are welcome!

#### The software peco

We quantified continous cell cycle phase using FUCCI fluorescence imaging and trained peco to predict this continuous cell cycle phase using scRNA-seq data collected from six human cell lines. Our paper showed that peco produces robust cell cycle phase predictions using strong cyclic genes - genes which expression levels oscillate along the cell cycle.

The software **peco** will be released in Biconductor 3.11. This release will use the latest R3.6.1.

The development version is available on GitHub. To install the development version,

```
devtools::install_github("jhsiao999/peco")
library(peco)
```

#### The analysis

- Process scRNA-seq and imaging data
- Estimate the cyclic trends of gene expression levels
- Compare the performance of peco with other methods on our data
- Compare the performance of peco with other methods using Leng et al. 2015 data
- Assess the performance of peco in thinned data
- Produce figures shown in our paper

#### Citation

Characterizing and inferring quantitative cell-cycle phase in single-cell RNA-seq data analysis.

#### Downloading the data files

You have two main options for downloading the data files. First, you can manually download the individual files by clicking on the links on this page or navigating to the files in the peco-paper GitHub repository. This is the recommended strategy if you only need a few data files.

Second, you can install git-lfs. To handle large files, we used Git Large File Storage (LFS). This means that the files that you download with `git clone` are only plain text files that contain identifiers for the files saved on GitHub’s servers. If you want to download all of the data files at once, you can do this with after you install git-lfs.

To install git-lfs, follow their instructions to download, install, and setup (`git lfs install`). Alternatively, if you use conda, you can install git-lfs with `conda install -c conda-forge git-lfs`. Once installed, you can download the latest version of the data files with `git lfs pull`.

#### Other information

- GEO record GSE121265 for all raw and processed sequencing data.
- We also provide the processed data in TXT format as a gzip compressed tarball on the Gilad lab website.

  

Session information

```
sessionInfo()
```

```
R version 3.5.1 (2018-07-02)
Platform: x86_64-pc-linux-gnu (64-bit)
Running under: Scientific Linux 7.4 (Nitrogen)

Matrix products: default
BLAS/LAPACK: /software/openblas-0.2.19-el7-x86_64/lib/libopenblas_haswellp-r0.2.19.so

locale:
 [1] LC_CTYPE=en_US.UTF-8       LC_NUMERIC=C              
 [3] LC_TIME=en_US.UTF-8        LC_COLLATE=en_US.UTF-8    
 [5] LC_MONETARY=en_US.UTF-8    LC_MESSAGES=en_US.UTF-8   
 [7] LC_PAPER=en_US.UTF-8       LC_NAME=C                 
 [9] LC_ADDRESS=C               LC_TELEPHONE=C            
[11] LC_MEASUREMENT=en_US.UTF-8 LC_IDENTIFICATION=C       

attached base packages:
[1] stats     graphics  grDevices utils     datasets  methods   base     

loaded via a namespace (and not attached):
 [1] workflowr_1.6.0 Rcpp_1.0.3      digest_0.6.20   later_0.7.5    
 [5] rprojroot_1.3-2 R6_2.4.0        backports_1.1.2 git2r_0.26.1   
 [9] magrittr_1.5    evaluate_0.12   stringi_1.2.4   fs_1.3.1       
[13] promises_1.0.1  whisker_0.3-2   rmarkdown_1.10  tools_3.5.1    
[17] stringr_1.3.1   glue_1.3.0      httpuv_1.4.5    yaml_2.2.0     
[21] compiler_3.5.1  htmltools_0.3.6 knitr_1.20
```
