## Supplementary material for "Characterizing and inferring quantitative cell cycle phase in single-cell RNA-seq data analysis": Source code of our analysis, including scripts and data necessary to reproduce the work.: npreg_trendfilter_quantile.html

Estimate cyclic trend of gene expression levels


peco-paper

- Home
- About
- License

- Source code


### Estimate cyclic trend of gene expression levels

###### *Joyce Hsiao*

workflowr

- Summary
- Checks
- Past versions

**Last updated:** 2020-01-26

```
Ignored files:
    Ignored:    .Rhistory
    Ignored:    .Rproj.user/

Untracked files:
    Untracked:  code/note_wo_w_pca.R
    Untracked:  data/data_training_test/
    Untracked:  data/eset-filtered.rds
    Untracked:  data/fit.trend.perm.highmiss.rds
    Untracked:  data/fit.trend.perm.lowmiss.rds
    Untracked:  data/fit_diff_cyclone.rds
    Untracked:  data/fit_diff_oscope.rds
    Untracked:  data/fit_diff_peco.rds
    Untracked:  data/fit_diff_recat.rds
    Untracked:  data/fit_diff_seurat.rds
    Untracked:  data/intensity.rds
    Untracked:  data/leng2015_data.rds
    Untracked:  data/leng_fucci_oscope_29genes.rda
    Untracked:  data/leng_fucci_recat.rda
    Untracked:  data/leng_geneinfo.txt
    Untracked:  data/log2cpm.quant.rds
    Untracked:  data/macosko-2015.rds
    Untracked:  data/nmeth.3549-S2.xlsx
    Untracked:  data/ourdata_cyclone_NA18511.rds
    Untracked:  data/ourdata_cyclone_NA18855.rds
    Untracked:  data/ourdata_cyclone_NA18870.rds
    Untracked:  data/ourdata_cyclone_NA19098.rds
    Untracked:  data/ourdata_cyclone_NA19101.rds
    Untracked:  data/ourdata_cyclone_NA19160.rds
    Untracked:  data/ourdata_oscope_366genes.rda
    Untracked:  data/ourdata_peco_NA18511_top005genes.rds
    Untracked:  data/ourdata_peco_NA18855_top005genes.rds
    Untracked:  data/ourdata_peco_NA18870_top005genes.rds
    Untracked:  data/ourdata_peco_NA19098_top005genes.rds
    Untracked:  data/ourdata_peco_NA19101_top005genes.rds
    Untracked:  data/ourdata_peco_NA19160_top005genes.rds
    Untracked:  data/ourdata_phase_cyclone.rds
    Untracked:  data/ourdata_phase_seurat.rds
    Untracked:  data/ourdata_recat.rda
    Untracked:  data/sce-filtered.rds

Unstaged changes:
    Modified:   analysis/index.Rmd
    Modified:   code/run_seurat.R
```

Note that any generated files, e.g. HTML, png, CSS, etc., are not included in this status report because it is ok for generated content to have uncommitted changes.

---

These are the previous versions of the R Markdown and HTML files. If you’ve configured a remote Git repository (see `?wflow_git_remote`), click on the hyperlinks in the table below to view them.

| File | Version | Author | Date | Message |
| --- | --- | --- | --- | --- |
| Rmd | 12d976a | jhsiao999 | 2020-01-26 | estimate cyclic trends of gene expression levels |

---

#### Introduction

We used trendfilter to estimate cyclic trend of gene expression levels for each gene. We computed proportion of variance explained (PVE) by the cyclic trend for each gene and quantify the signficance of cycliic trend by permutation-based p-values.

#### Set-up

Load packages

```
library(SingleCellExperiment)
library(dplyr)
library(matrixStats)
library(edgeR)
library(circular)
library(peco)
```

Load data

```
sce <- readRDS("data/sce-final.rds")
sce <- sce[grep("ENSG", rownames(sce)),]
pdata <- data.frame(colData(sce))
fdata <- data.frame(rowData(sce))

sce <- data_transform_quantile(sce)
log2cpm_quantNormed <- assay(sce, "cpm_quantNormed")
log2cpm_beforequant <- assay(sce, "cpm")

# derive and rotate cell cycle phase
pca <- prcomp(cbind(pdata$rfp.median.log10sum.adjust,
                    pdata$gfp.median.log10sum.adjust))
theta <- coord2rad(pca$x)
theta_final <- shift_origin(as.numeric(theta), 3*pi/4)

log2cpm_quantNormed <- log2cpm_quantNormed[,order(theta_final)]
log2cpm_beforequant <- log2cpm_beforequant[,order(theta_final)]

plot(log2cpm_quantNormed["ENSG00000170312",],
     main = "CDK1", ylab = "Normalized gene expression values", 
     xlab = "FUCCI phase")
```

#### Evaluate data after standardizing expression

Map log2cpm expression to standard normal distribution. The transformation is non-linear. Sort N expression values from the largest to the smalles. General N standard normal random variable. For the non-zero expression value, find the correspnoding standard normal random variable that has the same quantile and subsitute the value with the corresponding normal random variable value. We then find the stanadard normal random variable values correspond to non-zero expression values and randomly assign these non-zero expression values to a standard normal random variable value.

- For genes with relatively low fraction of deteted cells, this method allows to move the zero expression values closer to the non-zero expression value.
- For genes with high fraction of undetected cells, this method creates a

Check genes with low/high fraction of undetected cells.

```
ii.high <- order(rowMeans(log2cpm_beforequant > 0), decreasing = F)[1:5]
par(mfcol=c(3,5))
for (i in 1:5) {
  plot(log2cpm_beforequant[ii.high[i],], ylab = "log2 CPM expression values")
  plot(log2cpm_quantNormed[ii.high[i],], ylab = "quantile-normalized log2CPM")
  plot(x=log2cpm_beforequant[ii.high[i],], y=log2cpm_quantNormed[ii.high[i],],
       ylab = "quantile-normalized log2CPM",
       xlab = "log2 CPM expression values")
}
```

```
ii.low <- order(rowMeans(log2cpm_beforequant > 0), decreasing = T)[1:5]
par(mfcol=c(3,5))
for (i in 1:5) {
  plot(log2cpm_beforequant[ii.low[i],], ylab = "log2 CPM expression values")
  plot(log2cpm_quantNormed[ii.low[i],], ylab = "quantile-normalized log2CPM")
  plot(x=log2cpm_beforequant[ii.low[i],], y=log2cpm_quantNormed[ii.low[i],],
       ylab = "quantile-normalized log2CPM",
       xlab = "log2 CPM expression values")
}
```

Check genes that we previously found to have cyclical patterns in Whitfield et al 2002.

The Whitfeld et al. 2002 list was downloaded from Macosko et al. 2015 (10.1016/j.cell.2015.05.002). Link to the file is https://www.ncbi.nlm.nih.gov/pmc/articles/PMC4481139/bin/NIHMS687993-supplement-supp\_data\_2.xlsx.

```
macosko <- readRDS("data/macosko-2015.rds")

log2cpm_quantNormed_macosko <- log2cpm_quantNormed[rownames(log2cpm_quantNormed) %in%macosko$ensembl,]
macosko_present <- macosko[macosko$ensembl %in% rownames(log2cpm_quantNormed),]

par(mfrow=c(8,5), mar = c(2,2,2,1))
for (g in seq_along(macosko_present$ensembl[1:25])) {
  ind <- rownames(log2cpm_quantNormed_macosko) == macosko_present$ensembl[g]
  plot(log2cpm_quantNormed_macosko[ind,], ylab = "Quantile-normalized log2CPM expression values",
       main = paste(macosko_present$hgnc[g], ",", macosko_present$phase[g]),
       pch = 16, cex=.6, ylim = c(-3,3))
}
```

#### Fit trendfilter to the data after quantile normalization

```
fit.trend <- mclapply(1:10, function(g) {
  fit_trendfilter_generic(log2cpm_quantNormed[g,],
                              polyorder = 2)
}, mc.cores=25)
names(fit.trend) <- rownames(log2cpm_quantNormed)

saveRDS(fit.trend, "data/fit.quant.rds"))
```

```
# load pre-computed resutls
fit.quant <- readRDS("data/fit.quant.rds")

pve <- sapply(fit.quant, "[[", "trend.pve")
summary(pve)
```

```
      Min.    1st Qu.     Median       Mean    3rd Qu.       Max. 
-0.0043839  0.0000798  0.0002166  0.0008961  0.0004973  0.3266288
```

Plot top 10 genes in PVE.

```
pve.genes <- names(pve)[order(c(pve), decreasing = T)[1:10]]
par(mfrow=c(2,5))
for (g in 1:length(pve.genes)) {
  ii.g <- which(names(fit.quant)==pve.genes[g])
  plot(log2cpm_quantNormed[rownames(log2cpm_quantNormed)==pve.genes[g],],
       main = fdata[pve.genes[g],]$name, ylab = "Normalized expression")
  points(fit.quant[[ii.g]]$trend.yy, pch=16, col = "blue", cex=.7)
}
```

quickily check top 100 enrichment for cell cycle genes.

```
enrich.order <- function(cutoffs, metrics, cyclegenes, allgenes) {
  #  out <- order(mad.ratio$smash.mad.ratio)
  # cutoffs <- c(100, 200, 300)
  cycle.rich <- sapply(cutoffs, function(x) {
    which_top <- order(metrics, decreasing = T)[1:x]
    sig.cycle <- sum(allgenes[which_top] %in% cyclegenes)/x
    non.cycle <- sum(allgenes[-which_top] %in% cyclegenes)/(length(allgenes)-x)
    cbind(as.numeric(sum(allgenes[which_top] %in% cyclegenes)), 
          sig.cycle/non.cycle)
  })
  colnames(cycle.rich) <- cutoffs
  rownames(cycle.rich) <- c("nsig.genes.cycle", "fold.sig.vs.nonsig.cycle")
  cycle.rich
}

macosko <- readRDS("data/macosko-2015.rds")
enrich.order(cutoffs = c(100, 200, 300), 
             metrics = pve, cyclegenes = macosko$ensembl,
             allgenes = rownames(log2cpm_quantNormed))
```

```
                              100       200   300
nsig.genes.cycle         54.00000 73.000000 86.00
fold.sig.vs.nonsig.cycle 12.78701  8.931377  7.16
```

#### Compute permutation-based p-values

Consider two genes, one with large fraction of undetected cells and one with small fraction of undeteted cells. See if the null distribution is similar.

```
nperm <- 1000

# choose a gene with high fraction of missing and permute data
set.seed(17)
ii.frac.miss.high <- names(sample(which(rowMeans(log2cpm_beforequant==0) > .8),1))

fit.trend.highmiss <- mclapply(1:nperm, function(g) {
  fit.trendfilter.generic(sample(log2cpm_quantNormed[ii.frac.miss.high,]),
                              polyorder = 2)
}, mc.cores=25)
saveRDS(fit.trend.highmiss, "data/fit.trend.perm.highmiss.rds")


# choose a gene with low fraction of missing and permute data
set.seed(31)
ii.frac.miss.low <- names(sample(which(rowMeans(log2cpm_beforequant==0) < .1),1))

fit.trend.lowmiss <- mclapply(1:nperm, function(g) {
  fit.trendfilter.generic(sample(log2cpm_quantNormed[ii.frac.miss.low,]),
                  polyorder = 2)
}, mc.cores=25)
saveRDS(fit.trend.lowmiss, "data/fit.trend.perm.lowmiss.rds")
```

Turns out the p-value based on permuted data is not the same for genes with low and high fraction of undetected cells.

```
# load pre-computed results
perm.lowmiss <- readRDS("data/fit.trend.perm.lowmiss.rds")

perm.highmiss <- readRDS("data/fit.trend.perm.highmiss.rds")

pve.perm.lowmiss <- sapply(perm.lowmiss, "[[", "trend.pve")
pve.perm.highmiss <- sapply(perm.highmiss, "[[", "trend.pve")

summary(pve.perm.lowmiss)
```

```
     Min.   1st Qu.    Median      Mean   3rd Qu.      Max. 
3.700e-08 4.998e-05 1.388e-04 2.821e-04 3.382e-04 1.617e-02
```

```
summary(pve.perm.highmiss)
```

```
     Min.   1st Qu.    Median      Mean   3rd Qu.      Max. 
1.340e-07 5.089e-05 1.389e-04 2.664e-04 3.403e-04 4.214e-03
```

```
par(mfrow=c(1,2))
hist(pve.perm.lowmiss, nclass=30,
     main = "Fraction undetected < 10%", xlab = "p-value")
hist(pve.perm.highmiss, nclass=30,
     main = "Fraction undetected > 80%", xlab = "p-value")
```

Compute p-value based on two different distributions. High consistency between the two.

Use permutated distribution based data with low missing value, which turns out to be more conservative.

```
B <- length(pve.perm.lowmiss)
pval.perm.low <- sapply(fit.quant, function(x) (1+sum(pve.perm.lowmiss > as.numeric(x$trend.pve)))/(1+B))
pval.perm.high <- sapply(fit.quant, function(x) (1+sum(pve.perm.highmiss > as.numeric(x$trend.pve)))/(1+B))

summary(pval.perm.low)
```

```
    Min.  1st Qu.   Median     Mean  3rd Qu.     Max. 
0.000999 0.163836 0.365634 0.415336 0.648352 1.000000
```

```
summary(pval.perm.high)
```

```
    Min.  1st Qu.   Median     Mean  3rd Qu.     Max. 
0.000999 0.150849 0.360639 0.411520 0.630370 1.000000
```

```
plot(x=pval.perm.low, y=pval.perm.high,
     main = "permutation-based p-values",
     xlab = "Based on data with low zero fractions", 
     yalb = "Based on data with high zero fractions")
```

```
sum(pval.perm.high < .001)
```

```
[1] 278
```

```
sum(pval.perm.low < .001)
```

```
[1] 101
```

Cell cycle signals in signficant cyclic genes.

```
which.sig <- pval.perm.low < .001
enrich.sigval <- function(cutoffs, metrics, cyclegenes, allgenes) {
  #  out <- order(mad.ratio$smash.mad.ratio)
  # cutoffs <- c(100, 200, 300)
  cycle.rich <- sapply(cutoffs, function(x) {
    #which_top <- order(metrics, decreasing = T)[1:x]
    sig.cycle <- sum(allgenes[metrics < x] %in% cyclegenes)/sum(metrics < x)
    non.cycle <- sum(allgenes[metrics > x] %in% cyclegenes)/sum(metrics > x)
    cbind(sum(metrics < x), as.numeric(sum(allgenes[metrics < x] %in% cyclegenes)), 
          sig.cycle/non.cycle)
  })
  colnames(cycle.rich) <- cutoffs
  rownames(cycle.rich) <- c("nsig.genes", "nsig.genes.cycle", "fold.sig.vs.nonsig.cycle")
  cycle.rich
}

enrich.sigval(cutoffs = c(.001, .005, .01), metrics=pval.perm.low,
              cyclegenes = macosko$ensembl,
              allgenes = rownames(log2cpm_quantNormed))
```

```
                             0.001      0.005       0.01
nsig.genes               101.00000 476.000000 553.000000
nsig.genes.cycle          54.00000  99.000000  99.000000
fold.sig.vs.nonsig.cycle  12.65925   5.268908   4.502205
```

```
sessionInfo()
```

```
R version 3.5.1 (2018-07-02)
Platform: x86_64-pc-linux-gnu (64-bit)
Running under: Scientific Linux 7.4 (Nitrogen)

Matrix products: default
BLAS/LAPACK: /software/openblas-0.2.19-el7-x86_64/lib/libopenblas_haswellp-r0.2.19.so

locale:
 [1] LC_CTYPE=en_US.UTF-8       LC_NUMERIC=C              
 [3] LC_TIME=en_US.UTF-8        LC_COLLATE=en_US.UTF-8    
 [5] LC_MONETARY=en_US.UTF-8    LC_MESSAGES=en_US.UTF-8   
 [7] LC_PAPER=en_US.UTF-8       LC_NAME=C                 
 [9] LC_ADDRESS=C               LC_TELEPHONE=C            
[11] LC_MEASUREMENT=en_US.UTF-8 LC_IDENTIFICATION=C       

attached base packages:
[1] parallel  stats4    stats     graphics  grDevices utils     datasets 
[8] methods   base     

other attached packages:
 [1] peco_0.99.10                circular_0.4-93            
 [3] edgeR_3.24.0                limma_3.38.3               
 [5] dplyr_0.8.0.1               SingleCellExperiment_1.4.1 
 [7] SummarizedExperiment_1.12.0 DelayedArray_0.8.0         
 [9] BiocParallel_1.16.0         matrixStats_0.55.0         
[11] Biobase_2.42.0              GenomicRanges_1.34.0       
[13] GenomeInfoDb_1.18.1         IRanges_2.16.0             
[15] S4Vectors_0.20.1            BiocGenerics_0.28.0        

loaded via a namespace (and not attached):
 [1] viridis_0.5.1            genlasso_1.4            
 [3] viridisLite_0.3.0        foreach_1.4.4           
 [5] DelayedMatrixStats_1.4.0 assertthat_0.2.1        
 [7] vipor_0.4.5              GenomeInfoDbData_1.2.0  
 [9] yaml_2.2.0               pillar_1.3.1            
[11] backports_1.1.2          lattice_0.20-38         
[13] glue_1.3.0               digest_0.6.20           
[15] promises_1.0.1           XVector_0.22.0          
[17] colorspace_1.3-2         plyr_1.8.4              
[19] htmltools_0.3.6          httpuv_1.4.5            
[21] Matrix_1.2-17            pkgconfig_2.0.3         
[23] zlibbioc_1.28.0          purrr_0.3.2             
[25] mvtnorm_1.0-11           scales_1.0.0            
[27] HDF5Array_1.10.1         whisker_0.3-2           
[29] later_0.7.5              pracma_2.2.9            
[31] git2r_0.26.1             tibble_2.1.1            
[33] ggplot2_3.2.1            conicfit_1.0.4          
[35] lazyeval_0.2.1           magrittr_1.5            
[37] crayon_1.3.4             evaluate_0.12           
[39] fs_1.3.1                 doParallel_1.0.14       
[41] MASS_7.3-51.1            beeswarm_0.2.3          
[43] geigen_2.3               tools_3.5.1             
[45] scater_1.10.1            stringr_1.3.1           
[47] Rhdf5lib_1.4.3           munsell_0.5.0           
[49] locfit_1.5-9.1           compiler_3.5.1          
[51] rlang_0.4.0              rhdf5_2.26.2            
[53] grid_3.5.1               RCurl_1.95-4.11         
[55] iterators_1.0.12         igraph_1.2.2            
[57] bitops_1.0-6             rmarkdown_1.10          
[59] boot_1.3-20              gtable_0.2.0            
[61] codetools_0.2-15         reshape2_1.4.3          
[63] R6_2.4.0                 gridExtra_2.3           
[65] knitr_1.20               workflowr_1.6.0         
[67] rprojroot_1.3-2          ggbeeswarm_0.6.0        
[69] stringi_1.2.4            Rcpp_1.0.3              
[71] tidyselect_0.2.5
```

  

Session information

```
sessionInfo()
```

```
R version 3.5.1 (2018-07-02)
Platform: x86_64-pc-linux-gnu (64-bit)
Running under: Scientific Linux 7.4 (Nitrogen)

Matrix products: default
BLAS/LAPACK: /software/openblas-0.2.19-el7-x86_64/lib/libopenblas_haswellp-r0.2.19.so

locale:
 [1] LC_CTYPE=en_US.UTF-8       LC_NUMERIC=C              
 [3] LC_TIME=en_US.UTF-8        LC_COLLATE=en_US.UTF-8    
 [5] LC_MONETARY=en_US.UTF-8    LC_MESSAGES=en_US.UTF-8   
 [7] LC_PAPER=en_US.UTF-8       LC_NAME=C                 
 [9] LC_ADDRESS=C               LC_TELEPHONE=C            
[11] LC_MEASUREMENT=en_US.UTF-8 LC_IDENTIFICATION=C       

attached base packages:
[1] parallel  stats4    stats     graphics  grDevices utils     datasets 
[8] methods   base     

other attached packages:
 [1] peco_0.99.10                circular_0.4-93            
 [3] edgeR_3.24.0                limma_3.38.3               
 [5] dplyr_0.8.0.1               SingleCellExperiment_1.4.1 
 [7] SummarizedExperiment_1.12.0 DelayedArray_0.8.0         
 [9] BiocParallel_1.16.0         matrixStats_0.55.0         
[11] Biobase_2.42.0              GenomicRanges_1.34.0       
[13] GenomeInfoDb_1.18.1         IRanges_2.16.0             
[15] S4Vectors_0.20.1            BiocGenerics_0.28.0        

loaded via a namespace (and not attached):
 [1] viridis_0.5.1            genlasso_1.4            
 [3] viridisLite_0.3.0        foreach_1.4.4           
 [5] DelayedMatrixStats_1.4.0 assertthat_0.2.1        
 [7] vipor_0.4.5              GenomeInfoDbData_1.2.0  
 [9] yaml_2.2.0               pillar_1.3.1            
[11] backports_1.1.2          lattice_0.20-38         
[13] glue_1.3.0               digest_0.6.20           
[15] promises_1.0.1           XVector_0.22.0          
[17] colorspace_1.3-2         plyr_1.8.4              
[19] htmltools_0.3.6          httpuv_1.4.5            
[21] Matrix_1.2-17            pkgconfig_2.0.3         
[23] zlibbioc_1.28.0          purrr_0.3.2             
[25] mvtnorm_1.0-11           scales_1.0.0            
[27] HDF5Array_1.10.1         whisker_0.3-2           
[29] later_0.7.5              pracma_2.2.9            
[31] git2r_0.26.1             tibble_2.1.1            
[33] ggplot2_3.2.1            conicfit_1.0.4          
[35] lazyeval_0.2.1           magrittr_1.5            
[37] crayon_1.3.4             evaluate_0.12           
[39] fs_1.3.1                 doParallel_1.0.14       
[41] MASS_7.3-51.1            beeswarm_0.2.3          
[43] geigen_2.3               tools_3.5.1             
[45] scater_1.10.1            stringr_1.3.1           
[47] Rhdf5lib_1.4.3           munsell_0.5.0           
[49] locfit_1.5-9.1           compiler_3.5.1          
[51] rlang_0.4.0              rhdf5_2.26.2            
[53] grid_3.5.1               RCurl_1.95-4.11         
[55] iterators_1.0.12         igraph_1.2.2            
[57] bitops_1.0-6             rmarkdown_1.10          
[59] boot_1.3-20              gtable_0.2.0            
[61] codetools_0.2-15         reshape2_1.4.3          
[63] R6_2.4.0                 gridExtra_2.3           
[65] knitr_1.20               workflowr_1.6.0         
[67] rprojroot_1.3-2          ggbeeswarm_0.6.0        
[69] stringi_1.2.4            Rcpp_1.0.3              
[71] tidyselect_0.2.5
```
