## Supplementary material for "Characterizing and inferring quantitative cell cycle phase in single-cell RNA-seq data analysis": Source code of our analysis, including scripts and data necessary to reproduce the work.: pca_tf.html

PCA vs Technical Variables


peco-paper

- Home
- About
- License

- Source code


### PCA vs Technical Variables

###### *Po-Yuan Tung*

#### *2018-01-31*

workflowr

- Summary
- Checks
- Past versions

**Last updated:** 2020-01-23

**Checks:**  7  0

**Knit directory:** `peco-paper/`

| File | Version | Author | Date | Message |
| --- | --- | --- | --- | --- |
| Rmd | 5205df7 | jhsiao999 | 2020-01-23 | move pca\_tf.Rmd and change eset to sce |

---

#### Setup

```
library("cowplot")
library("dplyr")
library("edgeR")
library("ggplot2")
library("heatmap3")
library("reshape2")
library("SingleCellExperiment")
source("code/utility.R")
```

#### PCA

##### Before fileter

```
sce_raw <- readRDS("data/sce-raw.rds")

## look at human genes
sce_raw_hs <- sce_raw[rowData(sce_raw)$source == "H. sapiens", ]
head(colData(sce_raw_hs))
```

```
DataFrame with 6 rows and 44 columns
             experiment        well cell_number concentration         ERCC
              <integer> <character>   <integer>     <numeric>  <character>
20170905-A01   20170905         A01           1   1.726404375 50x dilution
20170905-A02   20170905         A02           1   1.445692561 50x dilution
20170905-A03   20170905         A03           1   1.889617025 50x dilution
20170905-A04   20170905         A04           1    0.47537227 50x dilution
20170905-A05   20170905         A05           1   0.559682703 50x dilution
20170905-A06   20170905         A06           1   2.135351839 50x dilution
             individual.1 individual.2 image_individual image_label
              <character>  <character>      <character>   <integer>
20170905-A01      NA18855      NA18870      18870_18855           3
20170905-A02      NA18855      NA18870      18870_18855           2
20170905-A03      NA18855      NA18870      18870_18855           1
20170905-A04      NA18855      NA18870      18870_18855          49
20170905-A05      NA18855      NA18870      18870_18855          50
20170905-A06      NA18855      NA18870      18870_18855          51
                   raw       umi    mapped  unmapped reads_ercc  reads_hs
             <integer> <integer> <integer> <integer>  <integer> <integer>
20170905-A01   5746265   3709414   2597589   1111825     161686   2435427
20170905-A02   3997709   2642317   1799823    842494     253670   1545769
20170905-A03   4765829   3301270   2274259   1027011     261145   2011561
20170905-A04   1926305   1286653    806647    480006     247028    559614
20170905-A05   2626155   1740464   1036933    703531     247831    788955
20170905-A06   5249443   3662342   2631138   1031204     210518   2419498
             reads_egfp reads_mcherry molecules  mol_ercc    mol_hs
              <integer>     <integer> <integer> <integer> <integer>
20170905-A01        475             1    158546      3692    154830
20170905-A02        379             5     88552      3648     84887
20170905-A03       1550             3    107984      3820    104119
20170905-A04          5             0     44772      3505     41262
20170905-A05        145             2     64926      3505     61413
20170905-A06       1118             4    134989      3664    131293
              mol_egfp mol_mcherry detect_ercc detect_hs     chip_id
             <integer>   <integer>   <numeric> <numeric> <character>
20170905-A01        23           1          41      9018     NA18870
20170905-A02        12           5          45      6848     NA18870
20170905-A03        44           1          44      7228     NA18855
20170905-A04         5           0          43      3653     NA18870
20170905-A05         6           2          44      4704     NA18870
20170905-A06        31           1          43      8198     NA18870
               chipmix   freemix      snps     reads    avg_dp    min_dp
             <numeric> <numeric> <integer> <integer> <numeric> <integer>
20170905-A01   0.16693   0.05395    311848      9356      0.03         1
20170905-A02   0.26917   0.13813    311848      4678      0.02         1
20170905-A03   0.36964   0.07778    311848      6201      0.02         1
20170905-A04    0.5132    0.2615    311848      1356         0         1
20170905-A05   0.54431   0.21419    311848      1906      0.01         1
20170905-A06   0.22935   0.08126    311848      7929      0.03         1
             snps_w_min  valid_id cut_off_reads   unmapped_ratios
              <integer> <logical>     <logical>         <numeric>
20170905-A01       3961      TRUE          TRUE 0.299730631307263
20170905-A02       2201      TRUE          TRUE 0.318846678880694
20170905-A03       2550      TRUE          TRUE 0.311095729825188
20170905-A04        857      TRUE         FALSE 0.373065620645193
20170905-A05       1139      TRUE         FALSE 0.404220368821188
20170905-A06       3190      TRUE          TRUE 0.281569553034643
             cut_off_unmapped    ercc_percentage cut_off_ercc
                    <logical>          <numeric>    <logical>
20170905-A01             TRUE 0.0622446430131942         TRUE
20170905-A02             TRUE  0.140941637038753         TRUE
20170905-A03             TRUE  0.114826411591644         TRUE
20170905-A04             TRUE  0.306240524045834        FALSE
20170905-A05             TRUE  0.239003870066822        FALSE
20170905-A06             TRUE 0.0800102465169064         TRUE
             cut_off_genes    ercc_conversion         conversion
                 <logical>          <numeric>          <numeric>
20170905-A01          TRUE 0.0228343826923791 0.0635740672990814
20170905-A02          TRUE  0.014380888555998 0.0549157086214046
20170905-A03          TRUE 0.0146278887208256 0.0517602995882302
20170905-A04         FALSE 0.0141886749680198 0.0737329659372353
20170905-A05         FALSE 0.0141427020832745 0.0778409414985646
20170905-A06          TRUE 0.0174046874851557 0.0542645623182991
             conversion_outlier molecule_outlier filter_all
                      <logical>        <logical>  <logical>
20170905-A01              FALSE            FALSE       TRUE
20170905-A02              FALSE            FALSE       TRUE
20170905-A03              FALSE            FALSE       TRUE
20170905-A04              FALSE            FALSE      FALSE
20170905-A05              FALSE            FALSE      FALSE
20170905-A06              FALSE            FALSE       TRUE
```

```
## remove genes of all 0s
sce_raw_hs_clean <- sce_raw_hs[rowSums(assay(sce_raw_hs)) != 0, ]
dim(sce_raw_hs_clean)
```

```
[1] 19348  1536
```

```
## convert to log2 cpm
mol_raw_hs_cpm <- edgeR::cpm(assay(sce_raw_hs_clean), log = TRUE)
mol_raw_hs_cpm_means <- rowMeans(mol_raw_hs_cpm)
summary(mol_raw_hs_cpm_means)
```

```
   Min. 1st Qu.  Median    Mean 3rd Qu.    Max. 
  5.412   5.431   5.634   5.987   6.210  13.007
```

```
## keep genes with reasonable expression levels 
mol_raw_hs_cpm <- mol_raw_hs_cpm[mol_raw_hs_cpm_means > median(mol_raw_hs_cpm_means), ]
dim(mol_raw_hs_cpm)
```

```
[1] 9674 1536
```

```
anno_raw = data.frame(colData(sce_raw))
anno_raw_hs = data.frame(colData(sce_raw_hs))
```

```
## pca of genes with reasonable expression levels
pca_raw_hs <- run_pca(mol_raw_hs_cpm)

## a function of pca vs technical factors
get_r2 <- function(x, y) {
  stopifnot(length(x) == length(y))
  model <- lm(y ~ x)
  stats <- summary(model)
  return(stats$adj.r.squared)
}

## selection of technical factor
covariates <- anno_raw %>% dplyr::select(experiment, well, concentration, raw:unmapped,
                                                     starts_with("detect"), chip_id, molecules)
## look at the first 6 PCs
pcs <- pca_raw_hs$PCs[, 1:6]

## generate the data
r2_before <- matrix(NA, nrow = ncol(covariates), ncol = ncol(pcs),
             dimnames = list(colnames(covariates), colnames(pcs)))
for (cov in colnames(covariates)) {
  for (pc in colnames(pcs)) {
    r2_before[cov, pc] <- get_r2(covariates[, cov], pcs[, pc])
  }
}

## plot
heatmap3(r2_before, cexRow=1, cexCol=1, margins=c(8,8), scale = "none",
                       ylab="technical factor", main = "Before filter")
```

```
plot_pca(pca_raw_hs$PCs, pcx = 1, pcy = 2, explained = pca_raw_hs$explained,
         metadata = anno_raw_hs, color="chip_id")
```

##### After filter

```
sce_filtered = sce_raw[,sce_raw$filter_all == TRUE]
```

Compute log2 CPM based on the library size before filtering.

```
log2cpm <- edgeR::cpm(assay(sce_filtered), log = TRUE)
dim(log2cpm)
```

```
[1] 20421   923
```

```
pca_log2cpm <- run_pca(log2cpm)

anno = data.frame(colData(sce_filtered))
anno$experiment <- as.factor(anno$experiment)

plot_pca(x=pca_log2cpm$PCs, explained=pca_log2cpm$explained,
         metadata=anno, color="chip_id")
```

```
plot_pca(x=pca_log2cpm$PCs, explained=pca_log2cpm$explained,
         metadata=anno, color="experiment")
```

```
## selection of technical factor
covariates <- anno %>% dplyr::select(experiment, well, chip_id, 
                                                     concentration, raw:unmapped,
                                                     starts_with("detect"),  molecules)
## look at the first 6 PCs
pcs <- pca_log2cpm$PCs[, 1:6]

## generate the data
r2 <- matrix(NA, nrow = ncol(covariates), ncol = ncol(pcs),
             dimnames = list(colnames(covariates), colnames(pcs)))
for (cov in colnames(covariates)) {
  for (pc in colnames(pcs)) {
    r2[cov, pc] <- get_r2(covariates[, cov], pcs[, pc])
  }
}

## plot heatmap
heatmap3(r2, cexRow=1, cexCol=1, margins=c(8,8), scale = "none", 
         ylab="technical factor", main = "After filter")
```

PC1 correlated with number of genes detected, which is described in Hicks et al 2017

Number of genes detected also highly correlated with sequencing metrics, especially total molecule number per sample.

```
cor_tech <- cor(as.matrix(covariates[,4:11]),use="pairwise.complete.obs")
heatmap(cor_tech, symm = TRUE)
```

Look at the top 10% expression genes to see if the correlation of PC1 and number of detected gene would go away. However, the PC1 is still not individual (chip\_id).

```
## look at top 10% of genes
log2cpm_mean <- rowMeans(log2cpm)
summary(log2cpm_mean)
```

```
   Min. 1st Qu.  Median    Mean 3rd Qu.    Max. 
  5.124   5.138   5.337   5.738   5.982  13.416
```

```
log2cpm_top <- log2cpm[rank(log2cpm_mean) / length(log2cpm_mean) > 1 - 0.1, ]
dim(log2cpm_top)
```

```
[1] 2043  923
```

```
pca_top <- run_pca(log2cpm_top)

## look at the first 6 PCs
pcs <- pca_top$PCs[, 1:6]

## generate the data
r2_top <- matrix(NA, nrow = ncol(covariates), ncol = ncol(pcs),
             dimnames = list(colnames(covariates), colnames(pcs)))
for (cov in colnames(covariates)) {
  for (pc in colnames(pcs)) {
    r2_top[cov, pc] <- get_r2(covariates[, cov], pcs[, pc])
  }
}

## plot heatmap
heatmap3(r2_top, cexRow=1, cexCol=1, margins=c(8,8), scale = "none", 
         ylab="technical factor", main = "Top 10 % gene")
```

  

Session information

```
sessionInfo()
```

```
R version 3.5.1 (2018-07-02)
Platform: x86_64-pc-linux-gnu (64-bit)
Running under: Scientific Linux 7.4 (Nitrogen)

Matrix products: default
BLAS/LAPACK: /software/openblas-0.2.19-el7-x86_64/lib/libopenblas_haswellp-r0.2.19.so

locale:
 [1] LC_CTYPE=en_US.UTF-8       LC_NUMERIC=C              
 [3] LC_TIME=en_US.UTF-8        LC_COLLATE=en_US.UTF-8    
 [5] LC_MONETARY=en_US.UTF-8    LC_MESSAGES=en_US.UTF-8   
 [7] LC_PAPER=en_US.UTF-8       LC_NAME=C                 
 [9] LC_ADDRESS=C               LC_TELEPHONE=C            
[11] LC_MEASUREMENT=en_US.UTF-8 LC_IDENTIFICATION=C       

attached base packages:
[1] parallel  stats4    stats     graphics  grDevices utils     datasets 
[8] methods   base     

other attached packages:
 [1] testit_0.9                  SingleCellExperiment_1.4.1 
 [3] SummarizedExperiment_1.12.0 DelayedArray_0.8.0         
 [5] BiocParallel_1.16.0         matrixStats_0.55.0         
 [7] Biobase_2.42.0              GenomicRanges_1.34.0       
 [9] GenomeInfoDb_1.18.1         IRanges_2.16.0             
[11] S4Vectors_0.20.1            BiocGenerics_0.28.0        
[13] reshape2_1.4.3              heatmap3_1.1.6             
[15] edgeR_3.24.0                limma_3.38.3               
[17] dplyr_0.8.0.1               cowplot_0.9.4              
[19] ggplot2_3.2.1              

loaded via a namespace (and not attached):
 [1] fastcluster_1.1.25     tidyselect_0.2.5       locfit_1.5-9.1        
 [4] purrr_0.3.2            lattice_0.20-38        colorspace_1.3-2      
 [7] htmltools_0.3.6        yaml_2.2.0             rlang_0.4.0           
[10] later_0.7.5            pillar_1.3.1           glue_1.3.0            
[13] withr_2.1.2            GenomeInfoDbData_1.2.0 plyr_1.8.4            
[16] stringr_1.3.1          zlibbioc_1.28.0        munsell_0.5.0         
[19] gtable_0.2.0           workflowr_1.6.0        evaluate_0.12         
[22] labeling_0.3           knitr_1.20             httpuv_1.4.5          
[25] Rcpp_1.0.3             promises_1.0.1         scales_1.0.0          
[28] backports_1.1.2        XVector_0.22.0         fs_1.3.1              
[31] digest_0.6.20          stringi_1.2.4          grid_3.5.1            
[34] rprojroot_1.3-2        tools_3.5.1            bitops_1.0-6          
[37] magrittr_1.5           lazyeval_0.2.1         RCurl_1.95-4.11       
[40] tibble_2.1.1           crayon_1.3.4           whisker_0.3-2         
[43] pkgconfig_2.0.3        Matrix_1.2-17          assertthat_0.2.1      
[46] rmarkdown_1.10         R6_2.4.0               git2r_0.26.1          
[49] compiler_3.5.1
```
