## Supplementary material for "Characterizing and inferring quantitative cell cycle phase in single-cell RNA-seq data analysis": Source code of our analysis, including scripts and data necessary to reproduce the work.: predict_thinned_data.html

```
Ignored files:
    Ignored:    .Rhistory
    Ignored:    .Rproj.user/

Untracked files:
    Untracked:  analysis/npreg_trendfilter_quantile.Rmd
    Untracked:  code/suppfig09_?.R
    Untracked:  data/HumanLengESC.rds
    Untracked:  data/data_training_test/
    Untracked:  data/eset-filtered.rds
    Untracked:  data/fit.quant.rds
    Untracked:  data/fit.trend.perm.lowmiss.rds
    Untracked:  data/fit_diff_cyclone.rds
    Untracked:  data/fit_diff_oscope.rds
    Untracked:  data/fit_diff_peco.rds
    Untracked:  data/fit_diff_recat.rds
    Untracked:  data/fit_diff_seurat.rds
    Untracked:  data/intensity.rds
    Untracked:  data/log2cpm.quant.rds
    Untracked:  data/macosko-2015.rds
    Untracked:  data/ourdata_cyclone_NA18511.rds
    Untracked:  data/ourdata_cyclone_NA18855.rds
    Untracked:  data/ourdata_cyclone_NA18870.rds
    Untracked:  data/ourdata_cyclone_NA19098.rds
    Untracked:  data/ourdata_cyclone_NA19101.rds
    Untracked:  data/ourdata_cyclone_NA19160.rds
    Untracked:  data/ourdata_oscope_366genes.rda
    Untracked:  data/ourdata_peco_NA18511_top005genes.rds
    Untracked:  data/ourdata_peco_NA18855_top005genes.rds
    Untracked:  data/ourdata_peco_NA18870_top005genes.rds
    Untracked:  data/ourdata_peco_NA19098_top005genes.rds
    Untracked:  data/ourdata_peco_NA19101_top005genes.rds
    Untracked:  data/ourdata_peco_NA19160_top005genes.rds
    Untracked:  data/ourdata_phase_cyclone.rds
    Untracked:  data/ourdata_phase_seurat.rds
    Untracked:  data/ourdata_recat.rda
    Untracked:  data/sce-filtered.rds

Unstaged changes:
    Modified:   analysis/eval_on_our_data.Rmd
    Modified:   analysis/index.Rmd
    Modified:   code/fig2.R
    Modified:   code/fig3.R
    Modified:   code/fig4.R
```

Note that any generated files, e.g. HTML, png, CSS, etc., are not included in this status report because it is ok for generated content to have uncommitted changes.

---

These are the previous versions of the R Markdown and HTML files. If you’ve configured a remote Git repository (see `?wflow_git_remote`), click on the hyperlinks in the table below to view them.

| File | Version | Author | Date | Message |
| --- | --- | --- | --- | --- |
| Rmd | fdebe2b | jhsiao999 | 2020-01-25 | code to compute prediction error in unthinned and thinned data |

---

### Introduction

### Set up

Load packages

```
library(SingleCellExperiment)
library(peco)
library(ggplot2)
library(dplyr)
```

Prepare thinned data. We used R package `seqgendiff` to thin the expression.

```
# function to make thinned data
makedata_thinned <- function(sce, thinlog2_rate) {

  library(SingleCellExperiment)
  library(peco)
  pdata <- data.frame(colData(sce))
  fdata <- data.frame(rowData(sce))
  counts <- assay(sce)[grep("ENSG", rownames(sce)), ]
  log2cpm <- t(log2(1+(10^6)*(t(counts)/pdata$molecules)))

  # ---- thinning!
  library(seqgendiff)
  nsamp <- ncol(counts)
  inputmat <- counts
  thinlog2 <- rexp(nsamp, rate = thinlog2_rate)
  outmat <- thin_lib(inputmat, thinlog2 = thinlog2)

  counts_thin <- outmat$mat
  dimnames(counts_thin) <- dimnames(counts)
  libsize <- colSums(counts_thin)

  keep_samples <- which(colMeans(counts_thin==0)>.01)
  keep_genes <- which(rowMeans(counts_thin == 0 ) < .5)
  counts_thin <- counts_thin[keep_genes, keep_samples]
  libsize <- libsize[keep_samples]

  log2cpm_thin <- t(log2(1+(10^6)*t(counts_thin)/libsize))
  dimnames(log2cpm_thin) <- dimnames(counts_thin)

  log2cpm_thin_quant <- do.call(rbind,
                                lapply(1:nrow(log2cpm_thin), function(i) {
                                  qqnorm(log2cpm_thin[i,], plot.it = F )$x }) )
  dimnames(log2cpm_thin_quant) <- dimnames(counts_thin)

  saveRDS(list(counts_thin=counts_thin,
               log2cpm_thin= log2cpm_thin,
               log2cpm_thin_quant=log2cpm_thin_quant),
          file=file.path(paste0("data/data_training_test/data_thinlog2_",
                                     sprintf("%03d", 100*thinlog2_rate),".rds")))

  pdata_thin <- pdata[keep_samples,]

  for (i in 1:length(unique(pdata$chip_id))) {
    ind <- unique(pdata_thin$chip_id)[i]
    ii_test <- c(1:nrow(pdata_thin))[which(pdata_thin$chip_id == ind)]
    counts_test <- counts[,ii_test]
    counts_thin_test <- counts_thin[,ii_test]
    log2cpm_thin_test <- log2cpm_thin[,ii_test]
    log2cpm_thin_quant_test <- log2cpm_thin_quant[,ii_test]

    data_thin_test <- list(counts_test=counts_test,
                           counts_thin_test=counts_thin_test,
                           log2cpm_thin_test=log2cpm_thin_test,
                           log2cpm_thin_quant_test=log2cpm_thin_quant_test)

    saveRDS(data_thin_test,
            file=file.path(paste0("data/data_training_test/ind_",ind,"_data_test_thinlog2_",
                                       sprintf("%03d", 100*thinlog2_rate),".rds")))
  }
}

makedata_thinned(sce, thinlog2_rate = .8)
makedata_thinned(sce, thinlog2_rate = .33)
```

### Properties of the thinned datasets

Library size

```
data_thinlog2_033 <- readRDS("data/data_training_test/data_thinlog2_033.rds")
data_thinlog2_080 <- readRDS("data/data_training_test/data_thinlog2_080.rds")


# thin library size by a factor of 4.24
mean(colSums(data_thinlog2_033$counts_thin))/mean(colSums(counts))
mean(colSums(counts)/(1/.23))
mean(colSums(data_thinlog2_033$counts_thin))
1/.23

# thin library size by a factor of 2.2
mean(colSums(data_thinlog2_080$counts_thin))/mean(colSums(counts))
mean(colSums(counts)/(1/.45))
mean(colSums(data_thinlog2_080$counts_thin))
1/.45
```

Number of genes detected

```
top5genes <- c("ENSG00000170312","ENSG00000175063",
           "ENSG00000131747", "ENSG00000198518", "ENSG00000197061")
inds <- unique(pdata$chip_id)

do.call(rbind, lapply(1:length(inds), function(i) {
  ind <- inds[i]
  samps <- rownames(pdata)[which(pdata$chip_id == ind)]
  exp_tmp <- data_thinlog2_080$counts_thin[,colnames(data_thinlog2_080$counts_thin) %in% samps]
  data.frame(ind = ind,
             g_detect = min(rowMeans(exp_tmp > 0)),
             g_detect_top50 = mean(rowMeans(exp_tmp[which(rownames(exp_tmp) %in% top5genes),] > 0)))
}))


do.call(rbind, lapply(1:length(inds), function(i) {
  ind <- inds[i]
  samps <- rownames(pdata)[which(pdata$chip_id == ind)]
  exp_tmp <- data_thinlog2_033$counts_thin[,colnames(data_thinlog2_033$counts_thin) %in% samps]
  data.frame(ind = ind,
             g_detect = min(rowMeans(exp_tmp > 0)))
}))

do.call(rbind, lapply(1:length(inds), function(i) {
  ind <- inds[i]
  samps <- rownames(pdata)[which(pdata$chip_id == ind)]
  exp_tmp <- counts[,colnames(counts) %in% samps]
  data.frame(ind = ind,
             g_detect = min(rowMeans(exp_tmp > 0)))
}))
```

Plot out properties of thinned data

```
# checking...
thing_080 <- readRDS("data/data_training_test/data_thinlog2_080.rds")
thing_033 <- readRDS("data/data_training_test/data_thinlog2_033.rds")

subsam <- do.call(rbind, list(data.frame(libsize=colSums(counts), thinlog2="1"),
                              data.frame(libsize=colSums(thing_080$counts_thin), thinlog2="080"),
                              data.frame(libsize=colSums(thing_033$counts_thin), thinlog2="033")))
subsam$thinlog2 <- factor(subsam$thinlog2,
                          levels=c("033", "080", "1"),
                          labels=c("4.4", "2.2", "1 (none)"))
ggplot(subsam, aes(x=thinlog2, y=libsize, group=thinlog2, fill=thinlog2)) +
  geom_violin() +
  geom_boxplot(width=.2, col="black") +
  labs(fill="Thinning factor") +
  ylab("Sample molecule count") +
  xlab("Thinning factor")
subsam %>% group_by(thinlog2) %>% summarize(mn = mean(libsize), sd = sd(libsize))
```

### Compute prediction error on unthinned data

```
fits_all <- readRDS("data/fit.quant.rds")
genes_all <- names(fits_all)[order(sapply(fits_all,"[[",3), decreasing=T)]

res_unthinned <- do.call(rbind, lapply(seq_along(unique(pdata$chip_id)), function(ind) {
    ind <- unique(pdata$chip_id)[i]
    res_thin_each <- do.call(rbind, lapply(2:50, function(ngenes) {
        data_test <- readRDS(paste0("data/data_training_test/ind_",ind,"_data_test.rds"))
        data_train <- readRDS(paste0("data/data_training_test/ind_",ind,"_data_training.rds"))

        which_genes <- genes_all[1:ngenes]
        fit_train <- cycle_npreg_insample(
          Y = with(data_train, 
                   log2cpm_quant_train[which(rownames(log2cpm_quant_train) %in% which_genes), ]),
          theta = with(data_train, theta_train))
        fit_test <- cycle_npreg_outsample(
            Y_test=with(data_test, 
                        log2cpm_quant_test[which(rownames(log2cpm_quant_test) %in% which_genes), ]),
            sigma_est=with(fit_train, sigma_est),
            funs_est=with(fit_train, funs_est))
    
        diff_time <- circ_dist(data_test$theta_test,
                               rotation(data_test$theta_test, fit_test$cell_times_est))
        out <- data.frame(phase_pred_rot=rotation(data_test$theta_test, fit_test$cell_times_est),
                   phase_ref=data_test$theta_test,
                   diff_time=diff_time,
                   ind = ind,
                   ngenes = ngenes)
        return(out)
    }) )
}) )


res_unthinned %>% group_by(ind, ngenes) %>%
  #  filter(ngenes <= 20) %>%
  summarise(diff_mean = mean(diff_time/2/pi),
            diff_se = sd(diff_time/2/pi)/sqrt(length(diff_time/2/pi))) %>%
  ggplot(., aes(x=factor(ngenes), y=diff_mean, group = ind)) +
  #  geom_vline(xintercept=seq(5, 50,5)-1, col="gray90", lty=1) +
  geom_hline(yintercept=seq(.1, .2, .01), col="gray90", lty=1) +
  geom_line(aes(col=ind), lwd=.7) + # ggtitle("thinlog2 = .80") +
  scale_fill_brewer(palette="Dark2") +
  geom_errorbar(aes(ymin=diff_mean-diff_se,
                    ymax=diff_mean+diff_se, col=ind), width=.2, alpha=.5) +
  stat_summary(fun.y=mean,geom="line",lwd=.5, group=1) +
  ylim(0,.3) + geom_hline(yintercept=.25, col="red") +
  labs(color="Test data (Cell line)") +
  xlab("Number of cyclic genes used in peco prediction") +
  ylab("Prediction error (% circle)") +
  scale_x_discrete(breaks=c(2:50),
                   labels=c(rep("",3),5, rep("",4), 10, rep("",4), 15,
                            rep("",4), 20, rep("",4), 25,
                            rep("",4), 30, rep("",4), 35,
                            rep("",4), 40, rep("",4), 45, rep("",4), 50)) +
  ggtitle("Performance in unthinned data")
```

### Compute prediction error on thinned data

Use peco predictors trained on unthinned data of samples from 5 individual cell lines to predict cell cycle phase in thinned data of samples from an individual not seen in training.

```
fits_all <- readRDS("data/fit.quant.rds")
genes_all <- names(fits_all)[order(sapply(fits_all,"[[",3), decreasing=T)]

res_thin_080 <- do.call(rbind, lapply(seq_along(unique(pdata$chip_id)), function(ind) {
    ind <- unique(pdata$chip_id)[i]
    res_thin_each <- do.call(rbind, lapply(2:50, function(ngenes) {
        data_test <- readRDS(paste0("data/data_training_test/ind_",ind,"_data_test.rds"))
        data_train <- readRDS(paste0("data/data_training_test/ind_",ind,"_data_training.rds"))

        which_genes <- genes_all[1:ngenes]
        fit_train <- cycle_npreg_insample(
          Y = with(data_train, 
                   log2cpm_quant_train[which(rownames(log2cpm_quant_train) %in% which_genes), ]),
          theta = with(data_train, theta_train))
        
        data_thin_test_080 <- readRDS(paste0("data/data_training_test/ind_", 
                                           ind, "_data_test_thinlog2_080.rds"))
        fit_test_080 <- cycle_npreg_outsample(
            Y_test=with(data_thin_test_080, 
                        log2cpm_thin_quant_test[which(rownames(log2cpm_thin_quant_test) %in% which_genes), ]),
            sigma_est=with(fit_train, sigma_est),
            funs_est=with(fit_train, funs_est))
    
        diff_time <- circ_dist(data_test$theta_test,
                               rotation(data_test$theta_test, fit_test_080$cell_times_est))
        out <- data.frame(phase_pred_rot=rotation(data_test$theta_test, fit_test$cell_times_est),
                   phase_ref=data_test$theta_test,
                   diff_time=diff_time,
                   ind = ind,
                   ngenes = ngenes)
        return(out)
    }) )
}) )

res_thin_033 %>% group_by(ind, ngenes) %>%
  #  filter(ngenes <= 20) %>%
  summarise(diff_mean = mean(diff_time/2/pi),
            diff_se = sd(diff_time/2/pi)/sqrt(length(diff_time/2/pi))) %>%
  ggplot(., aes(x=factor(ngenes), y=diff_mean, group = ind)) +
  #  geom_vline(xintercept=seq(5, 50,5)-1, col="gray90", lty=1) +
  geom_hline(yintercept=seq(.1, .2, .01), col="gray90", lty=1) +
  geom_line(aes(col=ind), lwd=.7) + # ggtitle("thinlog2 = .80") +
  scale_fill_brewer(palette="Dark2") +
  geom_errorbar(aes(ymin=diff_mean-diff_se,
                    ymax=diff_mean+diff_se, col=ind), width=.2, alpha=.5) +
  stat_summary(fun.y=mean,geom="line",lwd=.5, group=1) +
  ylim(0,.3) + geom_hline(yintercept=.25, col="red") +
  labs(color="Test data (Cell line)") +
  xlab("Number of cyclic genes used in peco prediction") +
  ylab("Prediction error (% circle)") +
  scale_x_discrete(breaks=c(2:50),
                   labels=c(rep("",3),5, rep("",4), 10, rep("",4), 15,
                            rep("",4), 20, rep("",4), 25,
                            rep("",4), 30, rep("",4), 35,
                            rep("",4), 40, rep("",4), 45, rep("",4), 50)) +
  ggtitle("Performance in data thinned by a factor of 2.2")


res_thin_033 <- do.call(rbind, lapply(seq_along(unique(pdata$chip_id)), function(ind) {
    ind <- unique(pdata$chip_id)[i]
    res_thin_each <- do.call(rbind, lapply(2:50, function(ngenes) {
        data_test <- readRDS(paste0("data/data_training_test/ind_",ind,"_data_test.rds"))
        data_train <- readRDS(paste0("data/data_training_test/ind_",ind,"_data_training.rds"))

        which_genes <- genes_all[1:ngenes]
        fit_train <- cycle_npreg_insample(
          Y = with(data_train, 
                   log2cpm_quant_train[which(rownames(log2cpm_quant_train) %in% which_genes), ]),
          theta = with(data_train, theta_train))
        
        data_thin_test_033 <- readRDS(paste0("data/data_training_test/ind_", 
                                           ind, "_data_test_thinlog2_033.rds"))
        fit_test_033 <- cycle_npreg_outsample(
            Y_test=with(data_thin_test_033, 
                        log2cpm_thin_quant_test[which(rownames(log2cpm_thin_quant_test) %in% which_genes), ]),
            sigma_est=with(fit_train, sigma_est),
            funs_est=with(fit_train, funs_est))
    
        diff_time <- circ_dist(data_test$theta_test,
                               rotation(data_test$theta_test, fit_test_033$cell_times_est))
        out <- data.frame(phase_pred_rot=rotation(data_test$theta_test, fit_test$cell_times_est),
                   phase_ref=data_test$theta_test,
                   diff_time=diff_time,
                   ind = ind,
                   ngenes = ngenes)
        return(out)
    }) )
}) )


res_thin_033 %>% group_by(ind, ngenes) %>%
  #  filter(ngenes <= 20) %>%
  summarise(diff_mean = mean(diff_time/2/pi),
            diff_se = sd(diff_time/2/pi)/sqrt(length(diff_time/2/pi))) %>%
  ggplot(., aes(x=factor(ngenes), y=diff_mean, group = ind)) +
  #  geom_vline(xintercept=seq(5, 50,5)-1, col="gray90", lty=1) +
  geom_hline(yintercept=seq(.1, .2, .01), col="gray90", lty=1) +
  geom_line(aes(col=ind), lwd=.7) + # ggtitle("thinlog2 = .80") +
  scale_fill_brewer(palette="Dark2") +
  geom_errorbar(aes(ymin=diff_mean-diff_se,
                    ymax=diff_mean+diff_se, col=ind), width=.2, alpha=.5) +
  stat_summary(fun.y=mean,geom="line",lwd=.5, group=1) +
  ylim(0,.3) + geom_hline(yintercept=.25, col="red") +
  labs(color="Test data (Cell line)") +
  xlab("Number of cyclic genes used in peco prediction") +
  ylab("Prediction error (% circle)") +
  scale_x_discrete(breaks=c(2:50),
                   labels=c(rep("",3),5, rep("",4), 10, rep("",4), 15,
                            rep("",4), 20, rep("",4), 25,
                            rep("",4), 30, rep("",4), 35,
                            rep("",4), 40, rep("",4), 45, rep("",4), 50)) +
  ggtitle("Performance in data thinned by a factor of 4.3")
```

### Session information

```
sessionInfo()
```

```
R version 3.5.1 (2018-07-02)
Platform: x86_64-pc-linux-gnu (64-bit)
Running under: Scientific Linux 7.4 (Nitrogen)

Matrix products: default
BLAS/LAPACK: /software/openblas-0.2.19-el7-x86_64/lib/libopenblas_haswellp-r0.2.19.so

locale:
 [1] LC_CTYPE=en_US.UTF-8       LC_NUMERIC=C              
 [3] LC_TIME=en_US.UTF-8        LC_COLLATE=en_US.UTF-8    
 [5] LC_MONETARY=en_US.UTF-8    LC_MESSAGES=en_US.UTF-8   
 [7] LC_PAPER=en_US.UTF-8       LC_NAME=C                 
 [9] LC_ADDRESS=C               LC_TELEPHONE=C            
[11] LC_MEASUREMENT=en_US.UTF-8 LC_IDENTIFICATION=C       

attached base packages:
[1] parallel  stats4    stats     graphics  grDevices utils     datasets 
[8] methods   base     

other attached packages:
 [1] dplyr_0.8.0.1               ggplot2_3.2.1              
 [3] peco_0.99.6                 SingleCellExperiment_1.4.1 
 [5] SummarizedExperiment_1.12.0 DelayedArray_0.8.0         
 [7] BiocParallel_1.16.0         matrixStats_0.55.0         
 [9] Biobase_2.42.0              GenomicRanges_1.34.0       
[11] GenomeInfoDb_1.18.1         IRanges_2.16.0             
[13] S4Vectors_0.20.1            BiocGenerics_0.28.0        

loaded via a namespace (and not attached):
 [1] viridis_0.5.1            genlasso_1.4            
 [3] viridisLite_0.3.0        foreach_1.4.4           
 [5] DelayedMatrixStats_1.4.0 assertthat_0.2.1        
 [7] vipor_0.4.5              GenomeInfoDbData_1.2.0  
 [9] yaml_2.2.0               pillar_1.3.1            
[11] backports_1.1.2          lattice_0.20-38         
[13] glue_1.3.0               digest_0.6.20           
[15] promises_1.0.1           XVector_0.22.0          
[17] colorspace_1.3-2         plyr_1.8.4              
[19] htmltools_0.3.6          httpuv_1.4.5            
[21] Matrix_1.2-17            pkgconfig_2.0.3         
[23] zlibbioc_1.28.0          purrr_0.3.2             
[25] mvtnorm_1.0-11           scales_1.0.0            
[27] HDF5Array_1.10.1         whisker_0.3-2           
[29] later_0.7.5              pracma_2.2.9            
[31] git2r_0.26.1             tibble_2.1.1            
[33] conicfit_1.0.4           withr_2.1.2             
[35] lazyeval_0.2.1           magrittr_1.5            
[37] crayon_1.3.4             evaluate_0.12           
[39] fs_1.3.1                 doParallel_1.0.14       
[41] MASS_7.3-51.1            beeswarm_0.2.3          
[43] geigen_2.3               tools_3.5.1             
[45] scater_1.10.1            stringr_1.3.1           
[47] Rhdf5lib_1.4.3           munsell_0.5.0           
[49] compiler_3.5.1           rlang_0.4.0             
[51] rhdf5_2.26.2             grid_3.5.1              
[53] RCurl_1.95-4.11          iterators_1.0.12        
[55] circular_0.4-93          igraph_1.2.2            
[57] bitops_1.0-6             rmarkdown_1.10          
[59] boot_1.3-20              gtable_0.2.0            
[61] codetools_0.2-15         reshape2_1.4.3          
[63] R6_2.4.0                 gridExtra_2.3           
[65] knitr_1.20               workflowr_1.6.0         
[67] rprojroot_1.3-2          stringi_1.2.4           
[69] ggbeeswarm_0.6.0         Rcpp_1.0.3              
[71] tidyselect_0.2.5
```

  

Session information

```
sessionInfo()
```

```
R version 3.5.1 (2018-07-02)
Platform: x86_64-pc-linux-gnu (64-bit)
Running under: Scientific Linux 7.4 (Nitrogen)

Matrix products: default
BLAS/LAPACK: /software/openblas-0.2.19-el7-x86_64/lib/libopenblas_haswellp-r0.2.19.so

locale:
 [1] LC_CTYPE=en_US.UTF-8       LC_NUMERIC=C              
 [3] LC_TIME=en_US.UTF-8        LC_COLLATE=en_US.UTF-8    
 [5] LC_MONETARY=en_US.UTF-8    LC_MESSAGES=en_US.UTF-8   
 [7] LC_PAPER=en_US.UTF-8       LC_NAME=C                 
 [9] LC_ADDRESS=C               LC_TELEPHONE=C            
[11] LC_MEASUREMENT=en_US.UTF-8 LC_IDENTIFICATION=C       

attached base packages:
[1] parallel  stats4    stats     graphics  grDevices utils     datasets 
[8] methods   base     

other attached packages:
 [1] dplyr_0.8.0.1               ggplot2_3.2.1              
 [3] peco_0.99.6                 SingleCellExperiment_1.4.1 
 [5] SummarizedExperiment_1.12.0 DelayedArray_0.8.0         
 [7] BiocParallel_1.16.0         matrixStats_0.55.0         
 [9] Biobase_2.42.0              GenomicRanges_1.34.0       
[11] GenomeInfoDb_1.18.1         IRanges_2.16.0             
[13] S4Vectors_0.20.1            BiocGenerics_0.28.0        

loaded via a namespace (and not attached):
 [1] viridis_0.5.1            genlasso_1.4            
 [3] viridisLite_0.3.0        foreach_1.4.4           
 [5] DelayedMatrixStats_1.4.0 assertthat_0.2.1        
 [7] vipor_0.4.5              GenomeInfoDbData_1.2.0  
 [9] yaml_2.2.0               pillar_1.3.1            
[11] backports_1.1.2          lattice_0.20-38         
[13] glue_1.3.0               digest_0.6.20           
[15] promises_1.0.1           XVector_0.22.0          
[17] colorspace_1.3-2         plyr_1.8.4              
[19] htmltools_0.3.6          httpuv_1.4.5            
[21] Matrix_1.2-17            pkgconfig_2.0.3         
[23] zlibbioc_1.28.0          purrr_0.3.2             
[25] mvtnorm_1.0-11           scales_1.0.0            
[27] HDF5Array_1.10.1         whisker_0.3-2           
[29] later_0.7.5              pracma_2.2.9            
[31] git2r_0.26.1             tibble_2.1.1            
[33] conicfit_1.0.4           withr_2.1.2             
[35] lazyeval_0.2.1           magrittr_1.5            
[37] crayon_1.3.4             evaluate_0.12           
[39] fs_1.3.1                 doParallel_1.0.14       
[41] MASS_7.3-51.1            beeswarm_0.2.3          
[43] geigen_2.3               tools_3.5.1             
[45] scater_1.10.1            stringr_1.3.1           
[47] Rhdf5lib_1.4.3           munsell_0.5.0           
[49] compiler_3.5.1           rlang_0.4.0             
[51] rhdf5_2.26.2             grid_3.5.1              
[53] RCurl_1.95-4.11          iterators_1.0.12        
[55] circular_0.4-93          igraph_1.2.2            
[57] bitops_1.0-6             rmarkdown_1.10          
[59] boot_1.3-20              gtable_0.2.0            
[61] codetools_0.2-15         reshape2_1.4.3          
[63] R6_2.4.0                 gridExtra_2.3           
[65] knitr_1.20               workflowr_1.6.0         
[67] rprojroot_1.3-2          stringi_1.2.4           
[69] ggbeeswarm_0.6.0         Rcpp_1.0.3              
[71] tidyselect_0.2.5
```
