## Supplementary material for "Characterizing and inferring quantitative cell cycle phase in single-cell RNA-seq data analysis": Source code of our analysis, including scripts and data necessary to reproduce the work.: reads-v-molecules.html

Read to molecule conversion


peco-paper

- Home
- About
- License

- Source code


### Read to molecule conversion

###### *John Blischak*

#### *2017-11-29*

workflowr

- Summary
- Checks
- Past versions

**Last updated:** 2020-01-23

**Checks:**  7  0

**Knit directory:** `peco-paper/`

| File | Version | Author | Date | Message |
| --- | --- | --- | --- | --- |
| Rmd | fdc569c | jhsiao999 | 2020-01-23 | move reads-v-molecules.Rmd and change eset to sce |

---

#### Setup

```
library("cowplot")
library("dplyr")
library("DT")
library("ggplot2")
library("reshape2")
library("SingleCellExperiment")
theme_set(theme_cowplot())
```

```
sce_raw = readRDS("data/sce-raw.rds")
anno = data.frame(colData(sce_raw))
anno$experiment = factor(anno$experiment)
```

#### Reads versus molecules

```
conv_hs_c1 <- ggplot(anno, aes(x = reads_hs, y = mol_hs,
                                     color = experiment)) +
  geom_point(alpha = 1/2) +
  labs(x = "Total read count",
       y = "Total molecule count",
       title = "Endogenous genes by C1 chip") +
  theme(legend.position = "none")

conv_hs_ind <- ggplot(anno, aes(x = reads_hs, y = mol_hs,
                                      color = chip_id)) +
  geom_point(alpha = 1/2) +
  scale_color_brewer(palette = "Dark2") +
  labs(x = "Total read count",
       y = "Total molecule count",
       title = "Endogenous genes by individual") +
  theme(legend.position = "none")

conv_ercc_c1 <- ggplot(anno, aes(x = reads_ercc, y = mol_ercc,
                                        color = experiment)) +
  geom_point(alpha = 1/2) +
  labs(x = "Total read count",
       y = "Total molecule count",
       title = "ERCC genes by C1 chip") +
  theme(legend.position = "none")

conv_ercc_ind <- ggplot(anno, aes(x = reads_ercc, y = mol_ercc,
                                         color = chip_id)) +
  geom_point(alpha = 1/2) +
  scale_color_brewer(palette = "Dark2") +
  labs(x = "Total read count",
       y = "Total molecule count",
       title = "ERCC genes by individual") +
  theme(legend.position = "none")

plot_grid(conv_hs_c1, conv_hs_ind, conv_ercc_c1, conv_ercc_ind,
        labels = letters[1:4])
```

#### Read to molecule conversion

```
anno$conv_hs <- anno$mol_hs / anno$reads_hs
anno$conv_ercc <- anno$mol_ercc / anno$reads_ercc

r2_hs_c1 <- summary(lm(conv_hs ~ experiment, data = anno))$r.squared
box_hs_c1 <- ggplot(anno, aes(x = experiment, y = conv_hs,
                                     fill = experiment)) +
  geom_boxplot() +
  labs(x = "C1 chip", y = "Conversion efficiency",
       title = sprintf("Endogenous genes R-squared: %.2f", r2_hs_c1)) +
  theme(legend.position = "none",
        axis.text.x = element_text(angle = 45, hjust = 1, vjust = 1))

r2_hs_ind <- summary(lm(conv_hs ~ chip_id, data = anno))$r.squared
box_hs_ind <- ggplot(anno, aes(x = chip_id, y = conv_hs,
                                      fill = chip_id)) +
  geom_boxplot() +
  scale_fill_brewer(palette = "Dark2") +
  labs(x = "Individual", y = "Conversion efficiency",
       title = sprintf("Endogenous genes R-squared: %.2f", r2_hs_ind)) +
  theme(legend.position = "none",
        axis.text.x = element_text(angle = 45, hjust = 1, vjust = 1))

r2_ercc_c1 <- summary(lm(conv_ercc ~ experiment, data = anno))$r.squared
box_ercc_c1 <- ggplot(anno, aes(x = experiment, y = conv_ercc,
                                       fill = experiment)) +
  geom_boxplot() +
  labs(x = "C1 chip", y = "Conversion efficiency",
       title = sprintf("ERCC genes R-squared: %.2f", r2_ercc_c1)) +
  theme(legend.position = "none",
        axis.text.x = element_text(angle = 45, hjust = 1, vjust = 1))

r2_ercc_ind <- summary(lm(conv_ercc ~ chip_id, data = anno))$r.squared
box_ercc_ind <- ggplot(anno, aes(x = chip_id, y = conv_ercc,
                                        fill = chip_id)) +
  geom_boxplot() +
  scale_fill_brewer(palette = "Dark2") +
  labs(x = "Individual", y = "Conversion efficiency",
       title = sprintf("ERCC genes R-squared: %.2f", r2_ercc_ind)) +
  theme(legend.position = "none",
        axis.text.x = element_text(angle = 45, hjust = 1, vjust = 1))

plot_grid(box_hs_c1, box_hs_ind, box_ercc_c1, box_ercc_ind,
          labels = letters[1:4])
```

#### Total ERCC versus total endogenous molecules

Recreating Tung et al., 2017 Figure 3b:

Tung et al., 2017 Figure 3b

```
gene_v_ercc_c1 <- ggplot(anno, aes(x = mol_hs, y = mol_ercc,
                        color = experiment)) +
  geom_point(alpha = 1/2) +
  labs(x = "Total gene molecule-counts per sample",
       y = "Total ERCC molecule-counts per sample",
       title = "C1 chip") +
  theme(legend.position = "none")

gene_v_ercc_ind <- ggplot(anno, aes(x = mol_hs, y = mol_ercc,
                                           color = chip_id)) +
  geom_point(alpha = 1/2) +
  scale_color_brewer(palette = "Dark2") +
  labs(x = "Total gene molecule-counts per sample",
       y = "Total ERCC molecule-counts per sample",
       title = "Individual") +
  theme(legend.position = "none")

plot_grid(gene_v_ercc_c1, gene_v_ercc_ind, labels = letters[1:2])
```

  

Session information

```
sessionInfo()
```

```
R version 3.5.1 (2018-07-02)
Platform: x86_64-pc-linux-gnu (64-bit)
Running under: Scientific Linux 7.4 (Nitrogen)

Matrix products: default
BLAS/LAPACK: /software/openblas-0.2.19-el7-x86_64/lib/libopenblas_haswellp-r0.2.19.so

locale:
 [1] LC_CTYPE=en_US.UTF-8       LC_NUMERIC=C              
 [3] LC_TIME=en_US.UTF-8        LC_COLLATE=en_US.UTF-8    
 [5] LC_MONETARY=en_US.UTF-8    LC_MESSAGES=en_US.UTF-8   
 [7] LC_PAPER=en_US.UTF-8       LC_NAME=C                 
 [9] LC_ADDRESS=C               LC_TELEPHONE=C            
[11] LC_MEASUREMENT=en_US.UTF-8 LC_IDENTIFICATION=C       

attached base packages:
[1] parallel  stats4    stats     graphics  grDevices utils     datasets 
[8] methods   base     

other attached packages:
 [1] SingleCellExperiment_1.4.1  SummarizedExperiment_1.12.0
 [3] DelayedArray_0.8.0          BiocParallel_1.16.0        
 [5] matrixStats_0.55.0          Biobase_2.42.0             
 [7] GenomicRanges_1.34.0        GenomeInfoDb_1.18.1        
 [9] IRanges_2.16.0              S4Vectors_0.20.1           
[11] BiocGenerics_0.28.0         reshape2_1.4.3             
[13] DT_0.5                      dplyr_0.8.0.1              
[15] cowplot_0.9.4               ggplot2_3.2.1              

loaded via a namespace (and not attached):
 [1] tidyselect_0.2.5       purrr_0.3.2            lattice_0.20-38       
 [4] colorspace_1.3-2       htmltools_0.3.6        yaml_2.2.0            
 [7] rlang_0.4.0            later_0.7.5            pillar_1.3.1          
[10] glue_1.3.0             withr_2.1.2            RColorBrewer_1.1-2    
[13] GenomeInfoDbData_1.2.0 plyr_1.8.4             stringr_1.3.1         
[16] zlibbioc_1.28.0        munsell_0.5.0          gtable_0.2.0          
[19] workflowr_1.6.0        htmlwidgets_1.3        evaluate_0.12         
[22] labeling_0.3           knitr_1.20             httpuv_1.4.5          
[25] Rcpp_1.0.3             promises_1.0.1         scales_1.0.0          
[28] backports_1.1.2        XVector_0.22.0         fs_1.3.1              
[31] digest_0.6.20          stringi_1.2.4          grid_3.5.1            
[34] rprojroot_1.3-2        tools_3.5.1            bitops_1.0-6          
[37] magrittr_1.5           lazyeval_0.2.1         RCurl_1.95-4.11       
[40] tibble_2.1.1           crayon_1.3.4           whisker_0.3-2         
[43] pkgconfig_2.0.3        Matrix_1.2-17          assertthat_0.2.1      
[46] rmarkdown_1.10         R6_2.4.0               git2r_0.26.1          
[49] compiler_3.5.1
```
