## Supplementary material for "Characterizing and inferring quantitative cell cycle phase in single-cell RNA-seq data analysis": Source code of our analysis, including scripts and data necessary to reproduce the work.: sampleqc.html

```
Ignored files:
    Ignored:    .Rhistory
    Ignored:    .Rproj.user/

Untracked files:
    Untracked:  analysis/gene-filtering.Rmd
    Untracked:  analysis/npreg_trendfilter_quantile.Rmd
    Untracked:  analysis/pca-tf.Rmd
    Untracked:  code/fig2_rev.R
    Untracked:  data/fit.quant.rds
    Untracked:  data/intensity.rds
    Untracked:  data/log2cpm.quant.rds

Unstaged changes:
    Modified:   analysis/access_data.Rmd
    Modified:   analysis/index.Rmd
    Modified:   code/fig2.R
```

Note that any generated files, e.g. HTML, png, CSS, etc., are not included in this status report because it is ok for generated content to have uncommitted changes.

---

These are the previous versions of the R Markdown and HTML files. If you’ve configured a remote Git repository (see `?wflow_git_remote`), click on the hyperlinks in the table below to view them.

| File | Version | Author | Date | Message |
| --- | --- | --- | --- | --- |
| Rmd | 3f2b8a8 | jhsiao999 | 2020-01-23 | move sampleqc.Rmd and change eset to sce |
| html | 179ec00 | jhsiao999 | 2020-01-23 | Build site. |
| Rmd | 0f6f533 | jhsiao999 | 2020-01-23 | copy and update eset to sce in sampleqc.Rmd |

---

### Setup

```
library("cowplot")
library("dplyr")
library("edgeR")
library("ggplot2")
library("MASS")
library("tibble")
library("SingleCellExperiment")
theme_set(cowplot::theme_cowplot())

# The palette with grey:
cbPalette <- c("#999999", "#E69F00", "#56B4E9", "#009E73", "#F0E442", "#0072B2", "#D55E00", "#CC79A7")
indi_palette <- c("tomato4", "chocolate1", "rosybrown", "purple4", "plum3", "slateblue")
```

```
sce_raw <- readRDS("data/sce-raw.rds")
anno <- data.frame(colData(sce_raw))
```

### Total mapped reads

Note: Using the 20 % cutoff of samples with no cells excludes all the samples

```
## calculate the cut-off  
cut_off_reads <- quantile(anno[anno$cell_number == 0,"mapped"], 0.82)

cut_off_reads
```

```
    82% 
1309921
```

```
anno$cut_off_reads <- anno$mapped > cut_off_reads

## numbers of cells 
sum(anno[anno$cell_number == 1, "mapped"] > cut_off_reads)
```

```
[1] 978
```

```
sum(anno[anno$cell_number == 1, "mapped"] <= cut_off_reads)
```

```
[1] 349
```

```
## density plots
plot_reads <- ggplot(anno[anno$cell_number == 0 |
                          anno$cell_number == 1 , ],
       aes(x = mapped, fill = as.factor(cell_number))) + 
       geom_density(alpha = 0.5) +
       geom_vline(xintercept = cut_off_reads, colour="grey", linetype = "longdash") +
       labs(x = "Total mapped reads", title = "Number of total mapped reads", fill = "Cell number")

plot_reads
```

Past versions of total-reads-1.png

| Version | Author | Date |
| --- | --- | --- |
| 179ec00 | jhsiao999 | 2020-01-23 |

### Unmapped ratios

Note: Using the 40 % cutoff of samples with no cells excludes all the samples

```
## calculate unmapped ratios
anno$unmapped_ratios <- anno$unmapped/anno$umi

## cut off 
cut_off_unmapped <- quantile(anno[anno$cell_number == 0,"unmapped_ratios"], 0.40)

cut_off_unmapped
```

```
      40% 
0.4362152
```

```
anno$cut_off_unmapped <- anno$unmapped_ratios < cut_off_unmapped

## numbers of cells 
sum(anno[anno$cell_number == 1, "unmapped_ratios"] >= cut_off_unmapped)
```

```
[1] 221
```

```
sum(anno[anno$cell_number == 1, "unmapped_ratios"] < cut_off_unmapped)
```

```
[1] 1106
```

```
## density plots
plot_unmapped <- ggplot(anno[anno$cell_number == 0 |
                             anno$cell_number == 1 , ],
       aes(x = unmapped_ratios *100, fill = as.factor(cell_number))) + 
       geom_density(alpha = 0.5) +
       geom_vline(xintercept = cut_off_unmapped *100, colour="grey", linetype = "longdash") +
       labs(x = "Unmapped reads/ total reads", title = "Unmapped reads percentage")

plot_unmapped
```

Past versions of unmapped-ratios-1.png

| Version | Author | Date |
| --- | --- | --- |
| 179ec00 | jhsiao999 | 2020-01-23 |

Look at the unmapped percentage per sample by C1 experimnet and by individual.

```
unmapped_exp <- ggplot(anno, aes(x = as.factor(experiment), y = unmapped_ratios, color = as.factor(experiment))) +
  geom_violin() + 
  geom_boxplot(alpha = .01, width = .2, position = position_dodge(width = .9)) +
  labs(x = "C1 chip", y = "Unmapped reads/ total reads",
       title = "Unmapped reads percentage") +
  theme(legend.title = element_blank(),
        axis.text.x = element_text(angle = 45, hjust = 1, vjust = 1))

unmapped_indi <- ggplot(anno, aes(x = chip_id, y = unmapped_ratios, color = as.factor(chip_id))) +
  geom_violin() + 
  geom_boxplot(alpha = .01, width = .2, position = position_dodge(width = .9)) +
  scale_color_manual(values = indi_palette) +
  labs(x = "C1 chip", y = "Unmapped reads/ total reads",
       title = "Unmapped reads percentage") +
  theme(legend.title = element_blank(),
        axis.text.x = element_text(angle = 45, hjust = 1, vjust = 1))

plot_grid(unmapped_exp + theme(legend.position = "none"),
          unmapped_indi + theme(legend.position = "none"),
          labels = letters[1:2])
```

Past versions of unmapped\_exp-1.png

| Version | Author | Date |
| --- | --- | --- |
| 179ec00 | jhsiao999 | 2020-01-23 |

### ERCC percentage

```
## calculate ercc reads percentage
anno$ercc_percentage <- anno$reads_ercc / anno$mapped

## cut off 
cut_off_ercc <- quantile(anno[anno$cell_number == 0,"ercc_percentage"], 0.20)

cut_off_ercc
```

```
     20% 
0.179423
```

```
anno$cut_off_ercc <- anno$ercc_percentage < cut_off_ercc

## numbers of cells 
sum(anno[anno$cell_number == 1, "ercc_percentage"] >= cut_off_ercc)
```

```
[1] 223
```

```
sum(anno[anno$cell_number == 1, "ercc_percentage"] < cut_off_ercc)
```

```
[1] 1104
```

```
## density plots
plot_ercc <- ggplot(anno[anno$cell_number == 0 |
                                anno$cell_number == 1 , ],
       aes(x = ercc_percentage *100, fill = as.factor(cell_number))) + 
       geom_density(alpha = 0.5) +
       geom_vline(xintercept = cut_off_ercc *100, colour="grey", linetype = "longdash") +
       labs(x = "ERCC reads / total mapped reads", title = "ERCC reads percentage")

plot_ercc
```

Past versions of ercc-percentage-1.png

| Version | Author | Date |
| --- | --- | --- |
| 179ec00 | jhsiao999 | 2020-01-23 |

Look at the ERCC spike-in percentage per sample by C1 experimnet and by individual.

```
ercc_exp <- ggplot(anno, aes(x = as.factor(experiment), y = ercc_percentage, color = as.factor(experiment))) +
  geom_violin() + 
  geom_boxplot(alpha = .01, width = .2, position = position_dodge(width = .9)) +
  labs(x = "C1 chip", y = "ERCC percentage",
       title = "ERCC percentage per sample") +
  theme(legend.title = element_blank(),
        axis.text.x = element_text(angle = 45, hjust = 1, vjust = 1))

ercc_indi <- ggplot(anno, aes(x = chip_id, y = ercc_percentage, color = as.factor(chip_id))) +
  geom_violin() + 
  geom_boxplot(alpha = .01, width = .2, position = position_dodge(width = .9)) +
  scale_colour_manual(values = indi_palette) +
  labs(x = "C1 chip", y = "ERCC percentage",
       title = "ERCC percentage per sample") +
  theme(legend.title = element_blank(),
        axis.text.x = element_text(angle = 45, hjust = 1, vjust = 1))

plot_grid(ercc_exp + theme(legend.position = "none"),
          ercc_indi + theme(legend.position = "none"),
          labels = letters[1:2])
```

Past versions of ercc\_exp-1.png

| Version | Author | Date |
| --- | --- | --- |
| 179ec00 | jhsiao999 | 2020-01-23 |

### Number of genes detected

```
## cut off 
cut_off_genes <- quantile(anno[anno$cell_number == 0,"detect_hs"], 0.80)

cut_off_genes
```

```
   80% 
6291.8
```

```
anno$cut_off_genes <- anno$detect_hs > cut_off_genes

## numbers of cells 
sum(anno[anno$cell_number == 1, "detect_hs"] > cut_off_genes)
```

```
[1] 1040
```

```
sum(anno[anno$cell_number == 1, "detect_hs"] <= cut_off_genes)
```

```
[1] 287
```

```
## density plots
plot_gene <- ggplot(anno[anno$cell_number == 0 |
                         anno$cell_number == 1 , ],
       aes(x = detect_hs, fill = as.factor(cell_number))) + 
       geom_density(alpha = 0.5) +
       geom_vline(xintercept = cut_off_genes, colour="grey", linetype = "longdash") +
       labs(x = "Gene numbers", title = "Numbers of detected genes")

plot_gene
```

Past versions of gene-number-1.png

| Version | Author | Date |
| --- | --- | --- |
| 179ec00 | jhsiao999 | 2020-01-23 |

```
number_exp <- ggplot(anno, aes(x = as.factor(experiment), y = detect_hs, color = as.factor(experiment))) +
  geom_violin() + 
  geom_boxplot(alpha = .01, width = .2, position = position_dodge(width = .9)) +
  labs(x = "C1 chip", y = "Number of genes detected",
       title = "Number of genes per sample") +
  theme(legend.title = element_blank(),
        axis.text.x = element_text(angle = 45, hjust = 1, vjust = 1))

number_indi <- ggplot(anno, aes(x = chip_id, y = detect_hs, color = as.factor(chip_id))) +
  geom_violin() + 
  geom_boxplot(alpha = .01, width = .2, position = position_dodge(width = .9)) +
  scale_colour_manual(values = indi_palette) +
  labs(x = "C1 chip", y = "Number of genes detected",
       title = "Number of genes per sample") +
  theme(legend.title = element_blank(),
        axis.text.x = element_text(angle = 45, hjust = 1, vjust = 1))

plot_grid(number_exp + theme(legend.position = "none"),
          number_indi + theme(legend.position = "none"),
          labels = letters[1:2])
```

Past versions of gene-number-exp-1.png

| Version | Author | Date |
| --- | --- | --- |
| 179ec00 | jhsiao999 | 2020-01-23 |

### FUCCI transgene

```
## plot molecule number of egfp and mCherry 
egfp_mol <- ggplot(anno[anno$cell_number == 0 |
            anno$cell_number == 1 , ],
       aes(x = mol_egfp, fill = as.factor(cell_number))) + 
       geom_density(alpha = 0.5) +
       labs(x = "EGFP molecule numbers", title = "Numbers of EGFP molecules")

mcherry_mol <- ggplot(anno[anno$cell_number == 0 |
            anno$cell_number == 1 , ],
       aes(x = mol_mcherry, fill = as.factor(cell_number))) + 
       geom_density(alpha = 0.5) +
       labs(x = "mCherry molecule numbers", title = "Numbers of mCherry molecules")

plot_grid(egfp_mol + theme(legend.position = c(.5,.9)), 
          mcherry_mol + theme(legend.position = "none"),
          labels = letters[1:2])
```

Past versions of fucci-1.png

| Version | Author | Date |
| --- | --- | --- |
| 179ec00 | jhsiao999 | 2020-01-23 |

### Linear Discriminat Analysis

#### Total molecule vs concentration

```
## create 3 groups according to cell number
group_3 <- rep("two",dim(anno)[1])
         group_3[grep("0", anno$cell_number)] <- "no"
         group_3[grep("1", anno$cell_number)] <- "one"

## create data frame
data <- anno %>% dplyr::select(experiment:concentration, mapped, molecules)
data <- data.frame(data, group = group_3)

## perform lda
data_lda <- lda(group ~ concentration + molecules, data = data)
data_lda_p <- predict(data_lda, newdata = data[,c("concentration", "molecules")])$class

## determine how well the model fix
table(data_lda_p, data[, "group"])
```

```
data_lda_p   no  one  two
       no     0    0    0
       one   16 1317  147
       two    0   11   45
```

```
data$data_lda_p <- data_lda_p

## identify the outlier
outliers_lda <- data %>% rownames_to_column("sample_id") %>% filter(cell_number == 1, data_lda_p == "two")
outliers_lda
```

```
      sample_id experiment well cell_number concentration  mapped
1  20170910-H09   20170910  H09           1     1.6540139 3013565
2  20170914-B04   20170914  B04           1     1.3324572 2284753
3  20170914-C04   20170914  C04           1     1.2389752 2489489
4  20170916-A08   20170916  A08           1     0.4900687 2276922
5  20170921-A12   20170921  A12           1     1.7967793 1868421
6  20170921-C01   20170921  C01           1     1.4171401 1543484
7  20170921-D04   20170921  D04           1     1.0853582 1753336
8  20170921-H09   20170921  H09           1     0.5802048 1541776
9  20170924-A04   20170924  A04           1     0.9856825 1414428
10 20170924-E01   20170924  E01           1     0.4581647 1817801
11 20170924-E03   20170924  E03           1     0.4484874 1924140
   molecules group data_lda_p
1     281516   one        two
2     190473   one        two
3     205586   one        two
4     169065   one        two
5     205868   one        two
6     202756   one        two
7     255148   one        two
8     415411   one        two
9     253058   one        two
10    187137   one        two
11    346383   one        two
```

```
## create filter
anno$molecule_outlier <- row.names(anno) %in% outliers_lda$sample_id

## plot before and after
plot_before <- ggplot(data, aes(x = concentration, y = molecules / 10^3,
               color = as.factor(group))) +
               geom_text(aes(label = cell_number, alpha = 0.5)) +
               labs(x = "Concentration", y = "Gene molecules (thousands)", title = "Before") +
               scale_color_brewer(palette = "Dark2") +
               theme(legend.position = "none")


plot_after <- ggplot(data, aes(x = concentration, y = molecules / 10^3,
               color = as.factor(data_lda_p))) +
               geom_text(aes(label = cell_number, alpha = 0.5)) +
               labs(x = "Concentration", y = "Gene molecules (thousands)", title = "After") +
               scale_color_brewer(palette = "Dark2") +
               theme(legend.position = "none")

plot_grid(plot_before + theme(legend.position=c(.8,.85)), 
          plot_after + theme(legend.position = "none"),
          labels = LETTERS[1:2])
```

Past versions of lda-1.png

| Version | Author | Date |
| --- | --- | --- |
| 179ec00 | jhsiao999 | 2020-01-23 |

#### Reads to molecule conversion

```
## calculate convertion
anno$ercc_conversion <- anno$mol_ercc / anno$reads_ercc

anno$conversion <- anno$mol_hs / anno$reads_hs

## try lda
data$conversion <- anno$conversion
data$ercc_conversion <- anno$ercc_conversion

data_ercc_lda <- lda(group ~ ercc_conversion + conversion, data = data)

data_ercc_lda_p <- predict(data_ercc_lda,  newdata = data[,c("ercc_conversion", "conversion")])$class

## determine how well the model fix
table(data_ercc_lda_p, data[, "group"])
```

```
data_ercc_lda_p   no  one  two
            no     5   20    0
            one   11 1303  166
            two    0    5   26
```

```
data$data_ercc_lda_p <- data_ercc_lda_p

## identify the outlier
outliers_conversion <- data %>% rownames_to_column("sample_id") %>% filter(cell_number == 1, data_ercc_lda_p == "two")
outliers_conversion
```

```
     sample_id experiment well cell_number concentration  mapped molecules
1 20170908-C07   20170908  C07           1     2.6936993  850577     95163
2 20170920-A10   20170920  A10           1     0.1369852   81371     28680
3 20170921-H09   20170921  H09           1     0.5802048 1541776    415411
4 20170924-A04   20170924  A04           1     0.9856825 1414428    253058
5 20170924-E03   20170924  E03           1     0.4484874 1924140    346383
  group data_lda_p conversion ercc_conversion data_ercc_lda_p
1   one        one  0.1126820      0.08186312             two
2   one        one  0.3585794      0.24727354             two
3   one        two  0.2716902      0.13085675             two
4   one        two  0.1809263      0.09389137             two
5   one        two  0.1813526      0.10260360             two
```

```
## create filter
anno$conversion_outlier <- row.names(anno) %in% outliers_conversion$sample_id

## plot before and after
plot_ercc_before <- ggplot(data, aes(x = ercc_conversion, y = conversion,
               color = as.factor(group))) +
               geom_text(aes(label = cell_number, alpha = 0.5)) +
               labs(x = "Convertion of ERCC spike-ins", y = "Conversion of genes", title = "Before") +
               scale_color_brewer(palette = "Dark2") +
               theme(legend.position = "none")

plot_ercc_after <- ggplot(data, aes(x = ercc_conversion, y = conversion,
               color = as.factor(data_ercc_lda_p))) +
               geom_text(aes(label = cell_number, alpha = 0.5)) +
               labs(x = "Convertion of ERCC spike-ins", y = "Conversion of genes", title = "After") +
               scale_color_brewer(palette = "Dark2") +
               theme(legend.position = "none")

plot_grid(plot_ercc_before, 
          plot_ercc_after,
          labels = LETTERS[3:4])
```

Past versions of convertion-1.png

| Version | Author | Date |
| --- | --- | --- |
| 179ec00 | jhsiao999 | 2020-01-23 |

### PCA

```
## look at human genes
sce_hs <- sce_raw[rowData(sce_raw)$source == "H. sapiens", ]
head(colData(sce_hs))
```

```
DataFrame with 6 rows and 44 columns
             experiment        well cell_number concentration         ERCC
              <integer> <character>   <integer>     <numeric>  <character>
20170905-A01   20170905         A01           1   1.726404375 50x dilution
20170905-A02   20170905         A02           1   1.445692561 50x dilution
20170905-A03   20170905         A03           1   1.889617025 50x dilution
20170905-A04   20170905         A04           1    0.47537227 50x dilution
20170905-A05   20170905         A05           1   0.559682703 50x dilution
20170905-A06   20170905         A06           1   2.135351839 50x dilution
             individual.1 individual.2 image_individual image_label
              <character>  <character>      <character>   <integer>
20170905-A01      NA18855      NA18870      18870_18855           3
20170905-A02      NA18855      NA18870      18870_18855           2
20170905-A03      NA18855      NA18870      18870_18855           1
20170905-A04      NA18855      NA18870      18870_18855          49
20170905-A05      NA18855      NA18870      18870_18855          50
20170905-A06      NA18855      NA18870      18870_18855          51
                   raw       umi    mapped  unmapped reads_ercc  reads_hs
             <integer> <integer> <integer> <integer>  <integer> <integer>
20170905-A01   5746265   3709414   2597589   1111825     161686   2435427
20170905-A02   3997709   2642317   1799823    842494     253670   1545769
20170905-A03   4765829   3301270   2274259   1027011     261145   2011561
20170905-A04   1926305   1286653    806647    480006     247028    559614
20170905-A05   2626155   1740464   1036933    703531     247831    788955
20170905-A06   5249443   3662342   2631138   1031204     210518   2419498
             reads_egfp reads_mcherry molecules  mol_ercc    mol_hs
              <integer>     <integer> <integer> <integer> <integer>
20170905-A01        475             1    158546      3692    154830
20170905-A02        379             5     88552      3648     84887
20170905-A03       1550             3    107984      3820    104119
20170905-A04          5             0     44772      3505     41262
20170905-A05        145             2     64926      3505     61413
20170905-A06       1118             4    134989      3664    131293
              mol_egfp mol_mcherry detect_ercc detect_hs     chip_id
             <integer>   <integer>   <numeric> <numeric> <character>
20170905-A01        23           1          41      9018     NA18870
20170905-A02        12           5          45      6848     NA18870
20170905-A03        44           1          44      7228     NA18855
20170905-A04         5           0          43      3653     NA18870
20170905-A05         6           2          44      4704     NA18870
20170905-A06        31           1          43      8198     NA18870
               chipmix   freemix      snps     reads    avg_dp    min_dp
             <numeric> <numeric> <integer> <integer> <numeric> <integer>
20170905-A01   0.16693   0.05395    311848      9356      0.03         1
20170905-A02   0.26917   0.13813    311848      4678      0.02         1
20170905-A03   0.36964   0.07778    311848      6201      0.02         1
20170905-A04    0.5132    0.2615    311848      1356         0         1
20170905-A05   0.54431   0.21419    311848      1906      0.01         1
20170905-A06   0.22935   0.08126    311848      7929      0.03         1
             snps_w_min  valid_id cut_off_reads   unmapped_ratios
              <integer> <logical>     <logical>         <numeric>
20170905-A01       3961      TRUE          TRUE 0.299730631307263
20170905-A02       2201      TRUE          TRUE 0.318846678880694
20170905-A03       2550      TRUE          TRUE 0.311095729825188
20170905-A04        857      TRUE         FALSE 0.373065620645193
20170905-A05       1139      TRUE         FALSE 0.404220368821188
20170905-A06       3190      TRUE          TRUE 0.281569553034643
             cut_off_unmapped    ercc_percentage cut_off_ercc
                    <logical>          <numeric>    <logical>
20170905-A01             TRUE 0.0622446430131942         TRUE
20170905-A02             TRUE  0.140941637038753         TRUE
20170905-A03             TRUE  0.114826411591644         TRUE
20170905-A04             TRUE  0.306240524045834        FALSE
20170905-A05             TRUE  0.239003870066822        FALSE
20170905-A06             TRUE 0.0800102465169064         TRUE
             cut_off_genes    ercc_conversion         conversion
                 <logical>          <numeric>          <numeric>
20170905-A01          TRUE 0.0228343826923791 0.0635740672990814
20170905-A02          TRUE  0.014380888555998 0.0549157086214046
20170905-A03          TRUE 0.0146278887208256 0.0517602995882302
20170905-A04         FALSE 0.0141886749680198 0.0737329659372353
20170905-A05         FALSE 0.0141427020832745 0.0778409414985646
20170905-A06          TRUE 0.0174046874851557 0.0542645623182991
             conversion_outlier molecule_outlier filter_all
                      <logical>        <logical>  <logical>
20170905-A01              FALSE            FALSE       TRUE
20170905-A02              FALSE            FALSE       TRUE
20170905-A03              FALSE            FALSE       TRUE
20170905-A04              FALSE            FALSE      FALSE
20170905-A05              FALSE            FALSE      FALSE
20170905-A06              FALSE            FALSE       TRUE
```

```
## remove genes of all 0s
sce_hs_clean <- sce_hs[rowSums(assay(sce_hs)) != 0, ]
dim(sce_hs_clean)
```

```
[1] 19348  1536
```

```
## convert to log2 cpm
mol_hs_cpm <- edgeR::cpm(assay(sce_hs_clean), log = TRUE)
mol_hs_cpm_means <- rowMeans(mol_hs_cpm)
summary(mol_hs_cpm_means)
```

```
   Min. 1st Qu.  Median    Mean 3rd Qu.    Max. 
  5.412   5.431   5.634   5.987   6.210  13.007
```

```
mol_hs_cpm <- mol_hs_cpm[mol_hs_cpm_means > median(mol_hs_cpm_means), ]
dim(mol_hs_cpm)
```

```
[1] 9674 1536
```

```
## pca of genes with reasonable expression levels
source("code/utility.R")
pca_hs <- run_pca(mol_hs_cpm)

plot_pca_id <- plot_pca(pca_hs$PCs, pcx = 1, pcy = 2, explained = pca_hs$explained,
         metadata = data.frame(colData(sce_hs_clean)), color = "chip_id")
plot_pca_id
```

Past versions of pca-1.png

| Version | Author | Date |
| --- | --- | --- |
| 179ec00 | jhsiao999 | 2020-01-23 |

### Filter

#### Final list

```
## all filter
anno$filter_all <- anno$cell_number == 1 &
                   anno$mol_egfp > 0 &
                   anno$valid_id &
                   anno$cut_off_reads &
                   anno$cut_off_unmapped &
                   anno$cut_off_ercc &
                   anno$cut_off_genes &
                   anno$molecule_outlier == "FALSE" &
                   anno$conversion_outlier == "FALSE"
sort(table(anno[anno$filter_all, "chip_id"]))
```

```
NA18511 NA19160 NA19101 NA18855 NA19098 NA18870 
    104     123     124     171     187     214
```

```
table(anno[anno$filter_all, c("experiment","chip_id")])
```

```
          chip_id
experiment NA18511 NA18855 NA18870 NA19098 NA19101 NA19160
  20170905       0      37      32       0       0       0
  20170906       0       0       0      41      22       0
  20170907       0      29       0      23       0       0
  20170908       0       0      32       0      31       0
  20170910       0      30       0       0      24       0
  20170912       0       0      42      37       0       0
  20170913       0      41       0       0       0       9
  20170914       0       0       0       0      24      34
  20170915      24      34       0       0       0       0
  20170916      13       0       0       0      23       0
  20170917       0       0       0      40       0      12
  20170919      11       0       0      46       0       0
  20170920      39       0       0       0       0      18
  20170921       0       0      45       0       0      22
  20170922      17       0      25       0       0       0
  20170924       0       0      38       0       0      28
```

#### Plots

```
genes_unmapped <-  ggplot(anno,
                   aes(x = detect_hs, y = unmapped_ratios * 100,
                       col = as.factor(chip_id), 
                       label = as.character(cell_number),
                       height = 600, width = 2000)) +
                   scale_colour_manual(values = indi_palette) +
                   geom_text(fontface = 3, alpha = 0.5) + 
                   geom_vline(xintercept = cut_off_genes, 
                              colour="grey", linetype = "longdash") +
                   geom_hline(yintercept = cut_off_unmapped * 100, 
                              colour="grey", linetype = "longdash") +
                   labs(x = "Number of detected genes / sample", 
                        y = "Percentage of unmapped reads (%)") 

genes_spike <- ggplot(anno,
               aes(x = detect_hs, y = ercc_percentage * 100,
                   col = as.factor(chip_id), 
                   label = as.character(cell_number), 
                   height = 600, width = 2000)) +
               scale_colour_manual(values = indi_palette) +
               scale_shape_manual(values=c(1:10)) +
               geom_text(fontface = 3, alpha = 0.5) + 
               geom_vline(xintercept = cut_off_genes, 
                          colour="grey", linetype = "longdash") +
               geom_hline(yintercept = cut_off_ercc * 100, 
                          colour="grey", linetype = "longdash") +
               labs(x = "Number of detected genes / samlpe", 
                    y = "Percentage of ERCC spike-in reads (%)") 

reads_unmapped_num <-  ggplot(anno,
                       aes(x = mapped, y = unmapped_ratios * 100,
                           col = as.factor(experiment), 
                           label = as.character(cell_number), 
                           height = 600, width = 2000)) +
                       geom_text(fontface = 3, alpha = 0.5) + 
                       geom_vline(xintercept = cut_off_reads, 
                                  colour="grey", linetype = "longdash") +
                       geom_hline(yintercept = cut_off_unmapped * 100,
                                  colour="grey", linetype = "longdash") +
                       labs(x = "Total mapped reads / sample", 
                            y = "Percentage of unmapped reads (%)") 

reads_spike_num <- ggplot(anno,
                   aes(x = mapped, y = ercc_percentage * 100,
                       col = as.factor(experiment), 
                       label = as.character(cell_number), 
                       height = 600, width = 2000)) +
                   geom_text(fontface = 3, alpha = 0.5) + 
                   geom_vline(xintercept = cut_off_reads, 
                              colour="grey", linetype = "longdash") +
                   geom_hline(yintercept = cut_off_ercc * 100, 
                              colour="grey", linetype = "longdash") +
                   labs(x = "Total mapped reads / sample",
                        y = "Percentage of ERCC spike-in reads (%)") 

plot_grid(genes_unmapped + theme(legend.position = c(.7,.9)), 
          genes_spike + theme(legend.position = "none"),
          labels = letters[1:2])
```

Past versions of plots-1.png

| Version | Author | Date |
| --- | --- | --- |
| 179ec00 | jhsiao999 | 2020-01-23 |

```
plot_grid(reads_unmapped_num + theme(legend.position = c(.7,.9)), 
          reads_spike_num + theme(legend.position = "none"),
          labels = letters[3:4])
```

Past versions of plots-2.png

| Version | Author | Date |
| --- | --- | --- |
| 179ec00 | jhsiao999 | 2020-01-23 |

```
ggplot(anno, aes(x = mol_hs, y = detect_hs,
                   col = as.factor(chip_id), 
                   height = 600, width = 2000)) +
               scale_colour_manual(values = indi_palette) +
               scale_shape_manual(values=c(1:10)) +
               geom_point(alpha = 0.6) + 
               labs(col = "Individual") +
               theme(legend.position = c(.7,.5)) +
               labs(x = "Total molecule counts / samlpe", 
                    y = "Number of detected genes / sample")
```

Past versions of unnamed-chunk-1-1.png

| Version | Author | Date |
| --- | --- | --- |
| 179ec00 | jhsiao999 | 2020-01-23 |

### Session information

```
R version 3.5.1 (2018-07-02)
Platform: x86_64-pc-linux-gnu (64-bit)
Running under: Scientific Linux 7.4 (Nitrogen)

Matrix products: default
BLAS/LAPACK: /software/openblas-0.2.19-el7-x86_64/lib/libopenblas_haswellp-r0.2.19.so

locale:
 [1] LC_CTYPE=en_US.UTF-8       LC_NUMERIC=C              
 [3] LC_TIME=en_US.UTF-8        LC_COLLATE=en_US.UTF-8    
 [5] LC_MONETARY=en_US.UTF-8    LC_MESSAGES=en_US.UTF-8   
 [7] LC_PAPER=en_US.UTF-8       LC_NAME=C                 
 [9] LC_ADDRESS=C               LC_TELEPHONE=C            
[11] LC_MEASUREMENT=en_US.UTF-8 LC_IDENTIFICATION=C       

attached base packages:
[1] parallel  stats4    stats     graphics  grDevices utils     datasets 
[8] methods   base     

other attached packages:
 [1] testit_0.9                  SingleCellExperiment_1.4.1 
 [3] SummarizedExperiment_1.12.0 DelayedArray_0.8.0         
 [5] BiocParallel_1.16.0         matrixStats_0.55.0         
 [7] Biobase_2.42.0              GenomicRanges_1.34.0       
 [9] GenomeInfoDb_1.18.1         IRanges_2.16.0             
[11] S4Vectors_0.20.1            BiocGenerics_0.28.0        
[13] tibble_2.1.1                MASS_7.3-51.1              
[15] edgeR_3.24.0                limma_3.38.3               
[17] dplyr_0.8.0.1               cowplot_0.9.4              
[19] ggplot2_3.2.1              

loaded via a namespace (and not attached):
 [1] tidyselect_0.2.5       locfit_1.5-9.1         purrr_0.3.2           
 [4] lattice_0.20-38        colorspace_1.3-2       htmltools_0.3.6       
 [7] yaml_2.2.0             rlang_0.4.0            later_0.7.5           
[10] pillar_1.3.1           glue_1.3.0             withr_2.1.2           
[13] RColorBrewer_1.1-2     GenomeInfoDbData_1.2.0 stringr_1.3.1         
[16] zlibbioc_1.28.0        munsell_0.5.0          gtable_0.2.0          
[19] workflowr_1.6.0        evaluate_0.12          labeling_0.3          
[22] knitr_1.20             httpuv_1.4.5           Rcpp_1.0.3            
[25] promises_1.0.1         scales_1.0.0           backports_1.1.2       
[28] XVector_0.22.0         fs_1.3.1               digest_0.6.20         
[31] stringi_1.2.4          grid_3.5.1             rprojroot_1.3-2       
[34] tools_3.5.1            bitops_1.0-6           magrittr_1.5          
[37] lazyeval_0.2.1         RCurl_1.95-4.11        crayon_1.3.4          
[40] whisker_0.3-2          pkgconfig_2.0.3        Matrix_1.2-17         
[43] assertthat_0.2.1       rmarkdown_1.10         R6_2.4.0              
[46] git2r_0.26.1           compiler_3.5.1
```

  

Session information

```
sessionInfo()
```

```
R version 3.5.1 (2018-07-02)
Platform: x86_64-pc-linux-gnu (64-bit)
Running under: Scientific Linux 7.4 (Nitrogen)

Matrix products: default
BLAS/LAPACK: /software/openblas-0.2.19-el7-x86_64/lib/libopenblas_haswellp-r0.2.19.so

locale:
 [1] LC_CTYPE=en_US.UTF-8       LC_NUMERIC=C              
 [3] LC_TIME=en_US.UTF-8        LC_COLLATE=en_US.UTF-8    
 [5] LC_MONETARY=en_US.UTF-8    LC_MESSAGES=en_US.UTF-8   
 [7] LC_PAPER=en_US.UTF-8       LC_NAME=C                 
 [9] LC_ADDRESS=C               LC_TELEPHONE=C            
[11] LC_MEASUREMENT=en_US.UTF-8 LC_IDENTIFICATION=C       

attached base packages:
[1] parallel  stats4    stats     graphics  grDevices utils     datasets 
[8] methods   base     

other attached packages:
 [1] testit_0.9                  SingleCellExperiment_1.4.1 
 [3] SummarizedExperiment_1.12.0 DelayedArray_0.8.0         
 [5] BiocParallel_1.16.0         matrixStats_0.55.0         
 [7] Biobase_2.42.0              GenomicRanges_1.34.0       
 [9] GenomeInfoDb_1.18.1         IRanges_2.16.0             
[11] S4Vectors_0.20.1            BiocGenerics_0.28.0        
[13] tibble_2.1.1                MASS_7.3-51.1              
[15] edgeR_3.24.0                limma_3.38.3               
[17] dplyr_0.8.0.1               cowplot_0.9.4              
[19] ggplot2_3.2.1              

loaded via a namespace (and not attached):
 [1] tidyselect_0.2.5       locfit_1.5-9.1         purrr_0.3.2           
 [4] lattice_0.20-38        colorspace_1.3-2       htmltools_0.3.6       
 [7] yaml_2.2.0             rlang_0.4.0            later_0.7.5           
[10] pillar_1.3.1           glue_1.3.0             withr_2.1.2           
[13] RColorBrewer_1.1-2     GenomeInfoDbData_1.2.0 stringr_1.3.1         
[16] zlibbioc_1.28.0        munsell_0.5.0          gtable_0.2.0          
[19] workflowr_1.6.0        evaluate_0.12          labeling_0.3          
[22] knitr_1.20             httpuv_1.4.5           Rcpp_1.0.3            
[25] promises_1.0.1         scales_1.0.0           backports_1.1.2       
[28] XVector_0.22.0         fs_1.3.1               digest_0.6.20         
[31] stringi_1.2.4          grid_3.5.1             rprojroot_1.3-2       
[34] tools_3.5.1            bitops_1.0-6           magrittr_1.5          
[37] lazyeval_0.2.1         RCurl_1.95-4.11        crayon_1.3.4          
[40] whisker_0.3-2          pkgconfig_2.0.3        Matrix_1.2-17         
[43] assertthat_0.2.1       rmarkdown_1.10         R6_2.4.0              
[46] git2r_0.26.1           compiler_3.5.1
```
