## Supplementary material for "Characterizing and inferring quantitative cell cycle phase in single-cell RNA-seq data analysis": Source code of our analysis, including scripts and data necessary to reproduce the work.: bootstrap.js

```
/*!
 * Bootstrap v3.3.5 (http://getbootstrap.com)
 * Copyright 2011-2015 Twitter, Inc.
 * Licensed under the MIT license
 */

if (typeof jQuery === 'undefined') {
  throw new Error('Bootstrap\'s JavaScript requires jQuery')
}

+function ($) {
  'use strict';
  var version = $.fn.jquery.split(' ')[0].split('.')
  if ((version[0] < 2 && version[1] < 9) || (version[0] == 1 && version[1] == 9 && version[2] < 1)) {
    throw new Error('Bootstrap\'s JavaScript requires jQuery version 1.9.1 or higher')
  }
}(jQuery);

/* ========================================================================
 * Bootstrap: transition.js v3.3.5
 * http://getbootstrap.com/javascript/#transitions
 * ========================================================================
 * Copyright 2011-2015 Twitter, Inc.
 * Licensed under MIT (https://github.com/twbs/bootstrap/blob/master/LICENSE)
 * ======================================================================== */


+function ($) {
  'use strict';

  // CSS TRANSITION SUPPORT (Shoutout: http://www.modernizr.com/)
  // ============================================================

  function transitionEnd() {
    var el = document.createElement('bootstrap')

    var transEndEventNames = {
      WebkitTransition : 'webkitTransitionEnd',
      MozTransition    : 'transitionend',
      OTransition      : 'oTransitionEnd otransitionend',
      transition       : 'transitionend'
    }

    for (var name in transEndEventNames) {
      if (el.style[name] !== undefined) {
        return { end: transEndEventNames[name] }
      }
    }

    return false // explicit for ie8 (  ._.)
  }

  // http://blog.alexmaccaw.com/css-transitions
  $.fn.emulateTransitionEnd = function (duration) {
    var called = false
    var $el = this
    $(this).one('bsTransitionEnd', function () { called = true })
    var callback = function () { if (!called) $($el).trigger($.support.transition.end) }
    setTimeout(callback, duration)
    return this
  }

  $(function () {
    $.support.transition = transitionEnd()

    if (!$.support.transition) return

    $.event.special.bsTransitionEnd = {
      bindType: $.support.transition.end,
      delegateType: $.support.transition.end,
      handle: function (e) {
        if ($(e.target).is(this)) return e.handleObj.handler.apply(this, arguments)
      }
    }
  })

}(jQuery);

/* ========================================================================
 * Bootstrap: alert.js v3.3.5
 * http://getbootstrap.com/javascript/#alerts
 * ========================================================================
 * Copyright 2011-2015 Twitter, Inc.
 * Licensed under MIT (https://github.com/twbs/bootstrap/blob/master/LICENSE)
 * ======================================================================== */


+function ($) {
  'use strict';

  // ALERT CLASS DEFINITION
  // ======================

  var dismiss = '[data-dismiss="alert"]'
  var Alert   = function (el) {
    $(el).on('click', dismiss, this.close)
  }

  Alert.VERSION = '3.3.5'

  Alert.TRANSITION_DURATION = 150

  Alert.prototype.close = function (e) {
    var $this    = $(this)
    var selector = $this.attr('data-target')

    if (!selector) {
      selector = $this.attr('href')
      selector = selector && selector.replace(/.*(?=#[^\s]*$)/, '') // strip for ie7
    }

    var $parent = $(selector)

    if (e) e.preventDefault()

    if (!$parent.length) {
      $parent = $this.closest('.alert')
    }

    $parent.trigger(e = $.Event('close.bs.alert'))

    if (e.isDefaultPrevented()) return

    $parent.removeClass('in')

    function removeElement() {
      // detach from parent, fire event then clean up data
      $parent.detach().trigger('closed.bs.alert').remove()
    }

    $.support.transition && $parent.hasClass('fade') ?
      $parent
        .one('bsTransitionEnd', removeElement)
        .emulateTransitionEnd(Alert.TRANSITION_DURATION) :
      removeElement()
  }


  // ALERT PLUGIN DEFINITION
  // =======================

  function Plugin(option) {
    return this.each(function () {
      var $this = $(this)
      var data  = $this.data('bs.alert')

      if (!data) $this.data('bs.alert', (data = new Alert(this)))
      if (typeof option == 'string') data[option].call($this)
    })
  }

  var old = $.fn.alert

  $.fn.alert             = Plugin
  $.fn.alert.Constructor = Alert


  // ALERT NO CONFLICT
  // =================

  $.fn.alert.noConflict = function () {
    $.fn.alert = old
    return this
  }


  // ALERT DATA-API
  // ==============

  $(document).on('click.bs.alert.data-api', dismiss, Alert.prototype.close)

}(jQuery);

/* ========================================================================
 * Bootstrap: button.js v3.3.5
 * http://getbootstrap.com/javascript/#buttons
 * ========================================================================
 * Copyright 2011-2015 Twitter, Inc.
 * Licensed under MIT (https://github.com/twbs/bootstrap/blob/master/LICENSE)
 * ======================================================================== */


+function ($) {
  'use strict';

  // BUTTON PUBLIC CLASS DEFINITION
  // ==============================

  var Button = function (element, options) {
    this.$element  = $(element)
    this.options   = $.extend({}, Button.DEFAULTS, options)
    this.isLoading = false
  }

  Button.VERSION  = '3.3.5'

  Button.DEFAULTS = {
    loadingText: 'loading...'
  }

  Button.prototype.setState = function (state) {
    var d    = 'disabled'
    var $el  = this.$element
    var val  = $el.is('input') ? 'val' : 'html'
    var data = $el.data()

    state += 'Text'

    if (data.resetText == null) $el.data('resetText', $el[val]())

    // push to event loop to allow forms to submit
    setTimeout($.proxy(function () {
      $el[val](data[state] == null ? this.options[state] : data[state])

      if (state == 'loadingText') {
        this.isLoading = true
        $el.addClass(d).attr(d, d)
      } else if (this.isLoading) {
        this.isLoading = false
        $el.removeClass(d).removeAttr(d)
      }
    }, this), 0)
  }

  Button.prototype.toggle = function () {
    var changed = true
    var $parent = this.$element.closest('[data-toggle="buttons"]')

    if ($parent.length) {
      var $input = this.$element.find('input')
      if ($input.prop('type') == 'radio') {
        if ($input.prop('checked')) changed = false
        $parent.find('.active').removeClass('active')
        this.$element.addClass('active')
      } else if ($input.prop('type') == 'checkbox') {
        if (($input.prop('checked')) !== this.$element.hasClass('active')) changed = false
        this.$element.toggleClass('active')
      }
      $input.prop('checked', this.$element.hasClass('active'))
      if (changed) $input.trigger('change')
    } else {
      this.$element.attr('aria-pressed', !this.$element.hasClass('active'))
      this.$element.toggleClass('active')
    }
  }


  // BUTTON PLUGIN DEFINITION
  // ========================

  function Plugin(option) {
    return this.each(function () {
      var $this   = $(this)
      var data    = $this.data('bs.button')
      var options = typeof option == 'object' && option

      if (!data) $this.data('bs.button', (data = new Button(this, options)))

      if (option == 'toggle') data.toggle()
      else if (option) data.setState(option)
    })
  }

  var old = $.fn.button

  $.fn.button             = Plugin
  $.fn.button.Constructor = Button


  // BUTTON NO CONFLICT
  // ==================

  $.fn.button.noConflict = function () {
    $.fn.button = old
    return this
  }


  // BUTTON DATA-API
  // ===============

  $(document)
    .on('click.bs.button.data-api', '[data-toggle^="button"]', function (e) {
      var $btn = $(e.target)
      if (!$btn.hasClass('btn')) $btn = $btn.closest('.btn')
      Plugin.call($btn, 'toggle')
      if (!($(e.target).is('input[type="radio"]') || $(e.target).is('input[type="checkbox"]'))) e.preventDefault()
    })
    .on('focus.bs.button.data-api blur.bs.button.data-api', '[data-toggle^="button"]', function (e) {
      $(e.target).closest('.btn').toggleClass('focus', /^focus(in)?$/.test(e.type))
    })

}(jQuery);

/* ========================================================================
 * Bootstrap: carousel.js v3.3.5
 * http://getbootstrap.com/javascript/#carousel
 * ========================================================================
 * Copyright 2011-2015 Twitter, Inc.
 * Licensed under MIT (https://github.com/twbs/bootstrap/blob/master/LICENSE)
 * ======================================================================== */


+function ($) {
  'use strict';

  // CAROUSEL CLASS DEFINITION
  // =========================

  var Carousel = function (element, options) {
    this.$element    = $(element)
    this.$indicators = this.$element.find('.carousel-indicators')
    this.options     = options
    this.paused      = null
    this.sliding     = null
    this.interval    = null
    this.$active     = null
    this.$items      = null

    this.options.keyboard && this.$element.on('keydown.bs.carousel', $.proxy(this.keydown, this))

    this.options.pause == 'hover' && !('ontouchstart' in document.documentElement) && this.$element
      .on('mouseenter.bs.carousel', $.proxy(this.pause, this))
      .on('mouseleave.bs.carousel', $.proxy(this.cycle, this))
  }

  Carousel.VERSION  = '3.3.5'

  Carousel.TRANSITION_DURATION = 600

  Carousel.DEFAULTS = {
    interval: 5000,
    pause: 'hover',
    wrap: true,
    keyboard: true
  }

  Carousel.prototype.keydown = function (e) {
    if (/input|textarea/i.test(e.target.tagName)) return
    switch (e.which) {
      case 37: this.prev(); break
      case 39: this.next(); break
      default: return
    }

    e.preventDefault()
  }

  Carousel.prototype.cycle = function (e) {
    e || (this.paused = false)

    this.interval && clearInterval(this.interval)

    this.options.interval
      && !this.paused
      && (this.interval = setInterval($.proxy(this.next, this), this.options.interval))

    return this
  }

  Carousel.prototype.getItemIndex = function (item) {
    this.$items = item.parent().children('.item')
    return this.$items.index(item || this.$active)
  }

  Carousel.prototype.getItemForDirection = function (direction, active) {
    var activeIndex = this.getItemIndex(active)
    var willWrap = (direction == 'prev' && activeIndex === 0)
                || (direction == 'next' && activeIndex == (this.$items.length - 1))
    if (willWrap && !this.options.wrap) return active
    var delta = direction == 'prev' ? -1 : 1
    var itemIndex = (activeIndex + delta) % this.$items.length
    return this.$items.eq(itemIndex)
  }

  Carousel.prototype.to = function (pos) {
    var that        = this
    var activeIndex = this.getItemIndex(this.$active = this.$element.find('.item.active'))

    if (pos > (this.$items.length - 1) || pos < 0) return

    if (this.sliding)       return this.$element.one('slid.bs.carousel', function () { that.to(pos) }) // yes, "slid"
    if (activeIndex == pos) return this.pause().cycle()

    return this.slide(pos > activeIndex ? 'next' : 'prev', this.$items.eq(pos))
  }

  Carousel.prototype.pause = function (e) {
    e || (this.paused = true)

    if (this.$element.find('.next, .prev').length && $.support.transition) {
      this.$element.trigger($.support.transition.end)
      this.cycle(true)
    }

    this.interval = clearInterval(this.interval)

    return this
  }

  Carousel.prototype.next = function () {
    if (this.sliding) return
    return this.slide('next')
  }

  Carousel.prototype.prev = function () {
    if (this.sliding) return
    return this.slide('prev')
  }

  Carousel.prototype.slide = function (type, next) {
    var $active   = this.$element.find('.item.active')
    var $next     = next || this.getItemForDirection(type, $active)
    var isCycling = this.interval
    var direction = type == 'next' ? 'left' : 'right'
    var that      = this

    if ($next.hasClass('active')) return (this.sliding = false)

    var relatedTarget = $next[0]
    var slideEvent = $.Event('slide.bs.carousel', {
      relatedTarget: relatedTarget,
      direction: direction
    })
    this.$element.trigger(slideEvent)
    if (slideEvent.isDefaultPrevented()) return

    this.sliding = true

    isCycling && this.pause()

    if (this.$indicators.length) {
      this.$indicators.find('.active').removeClass('active')
      var $nextIndicator = $(this.$indicators.children()[this.getItemIndex($next)])
      $nextIndicator && $nextIndicator.addClass('active')
    }

    var slidEvent = $.Event('slid.bs.carousel', { relatedTarget: relatedTarget, direction: direction }) // yes, "slid"
    if ($.support.transition && this.$element.hasClass('slide')) {
      $next.addClass(type)
      $next[0].offsetWidth // force reflow
      $active.addClass(direction)
      $next.addClass(direction)
      $active
        .one('bsTransitionEnd', function () {
          $next.removeClass([type, direction].join(' ')).addClass('active')
          $active.removeClass(['active', direction].join(' '))
          that.sliding = false
          setTimeout(function () {
            that.$element.trigger(slidEvent)
          }, 0)
        })
        .emulateTransitionEnd(Carousel.TRANSITION_DURATION)
    } else {
      $active.removeClass('active')
      $next.addClass('active')
      this.sliding = false
      this.$element.trigger(slidEvent)
    }

    isCycling && this.cycle()

    return this
  }


  // CAROUSEL PLUGIN DEFINITION
  // ==========================

  function Plugin(option) {
    return this.each(function () {
      var $this   = $(this)
      var data    = $this.data('bs.carousel')
      var options = $.extend({}, Carousel.DEFAULTS, $this.data(), typeof option == 'object' && option)
      var action  = typeof option == 'string' ? option : options.slide

      if (!data) $this.data('bs.carousel', (data = new Carousel(this, options)))
      if (typeof option == 'number') data.to(option)
      else if (action) data[action]()
      else if (options.interval) data.pause().cycle()
    })
  }

  var old = $.fn.carousel

  $.fn.carousel             = Plugin
  $.fn.carousel.Constructor = Carousel


  // CAROUSEL NO CONFLICT
  // ====================

  $.fn.carousel.noConflict = function () {
    $.fn.carousel = old
    return this
  }


  // CAROUSEL DATA-API
  // =================

  var clickHandler = function (e) {
    var href
    var $this   = $(this)
    var $target = $($this.attr('data-target') || (href = $this.attr('href')) && href.replace(/.*(?=#[^\s]+$)/, '')) // strip for ie7
    if (!$target.hasClass('carousel')) return
    var options = $.extend({}, $target.data(), $this.data())
    var slideIndex = $this.attr('data-slide-to')
    if (slideIndex) options.interval = false

    Plugin.call($target, options)

    if (slideIndex) {
      $target.data('bs.carousel').to(slideIndex)
    }

    e.preventDefault()
  }

  $(document)
    .on('click.bs.carousel.data-api', '[data-slide]', clickHandler)
    .on('click.bs.carousel.data-api', '[data-slide-to]', clickHandler)

  $(window).on('load', function () {
    $('[data-ride="carousel"]').each(function () {
      var $carousel = $(this)
      Plugin.call($carousel, $carousel.data())
    })
  })

}(jQuery);

/* ========================================================================
 * Bootstrap: collapse.js v3.3.5
 * http://getbootstrap.com/javascript/#collapse
 * ========================================================================
 * Copyright 2011-2015 Twitter, Inc.
 * Licensed under MIT (https://github.com/twbs/bootstrap/blob/master/LICENSE)
 * ======================================================================== */


+function ($) {
  'use strict';

  // COLLAPSE PUBLIC CLASS DEFINITION
  // ================================

  var Collapse = function (element, options) {
    this.$element      = $(element)
    this.options       = $.extend({}, Collapse.DEFAULTS, options)
    this.$trigger      = $('[data-toggle="collapse"][href="#' + element.id + '"],' +
                           '[data-toggle="collapse"][data-target="#' + element.id + '"]')
    this.transitioning = null

    if (this.options.parent) {
      this.$parent = this.getParent()
    } else {
      this.addAriaAndCollapsedClass(this.$element, this.$trigger)
    }

    if (this.options.toggle) this.toggle()
  }

  Collapse.VERSION  = '3.3.5'

  Collapse.TRANSITION_DURATION = 350

  Collapse.DEFAULTS = {
    toggle: true
  }

  Collapse.prototype.dimension = function () {
    var hasWidth = this.$element.hasClass('width')
    return hasWidth ? 'width' : 'height'
  }

  Collapse.prototype.show = function () {
    if (this.transitioning || this.$element.hasClass('in')) return

    var activesData
    var actives = this.$parent && this.$parent.children('.panel').children('.in, .collapsing')

    if (actives && actives.length) {
      activesData = actives.data('bs.collapse')
      if (activesData && activesData.transitioning) return
    }

    var startEvent = $.Event('show.bs.collapse')
    this.$element.trigger(startEvent)
    if (startEvent.isDefaultPrevented()) return

    if (actives && actives.length) {
      Plugin.call(actives, 'hide')
      activesData || actives.data('bs.collapse', null)
    }

    var dimension = this.dimension()

    this.$element
      .removeClass('collapse')
      .addClass('collapsing')[dimension](0)
      .attr('aria-expanded', true)

    this.$trigger
      .removeClass('collapsed')
      .attr('aria-expanded', true)

    this.transitioning = 1

    var complete = function () {
      this.$element
        .removeClass('collapsing')
        .addClass('collapse in')[dimension]('')
      this.transitioning = 0
      this.$element
        .trigger('shown.bs.collapse')
    }

    if (!$.support.transition) return complete.call(this)

    var scrollSize = $.camelCase(['scroll', dimension].join('-'))

    this.$element
      .one('bsTransitionEnd', $.proxy(complete, this))
      .emulateTransitionEnd(Collapse.TRANSITION_DURATION)[dimension](this.$element[0][scrollSize])
  }

  Collapse.prototype.hide = function () {
    if (this.transitioning || !this.$element.hasClass('in')) return

    var startEvent = $.Event('hide.bs.collapse')
    this.$element.trigger(startEvent)
    if (startEvent.isDefaultPrevented()) return

    var dimension = this.dimension()

    this.$element[dimension](this.$element[dimension]())[0].offsetHeight

    this.$element
      .addClass('collapsing')
      .removeClass('collapse in')
      .attr('aria-expanded', false)

    this.$trigger
      .addClass('collapsed')
      .attr('aria-expanded', false)

    this.transitioning = 1

    var complete = function () {
      this.transitioning = 0
      this.$element
        .removeClass('collapsing')
        .addClass('collapse')
        .trigger('hidden.bs.collapse')
    }

    if (!$.support.transition) return complete.call(this)

    this.$element
      [dimension](0)
      .one('bsTransitionEnd', $.proxy(complete, this))
      .emulateTransitionEnd(Collapse.TRANSITION_DURATION)
  }

  Collapse.prototype.toggle = function () {
    this[this.$element.hasClass('in') ? 'hide' : 'show']()
  }

  Collapse.prototype.getParent = function () {
    return $(this.options.parent)
      .find('[data-toggle="collapse"][data-parent="' + this.options.parent + '"]')
      .each($.proxy(function (i, element) {
        var $element = $(element)
        this.addAriaAndCollapsedClass(getTargetFromTrigger($element), $element)
      }, this))
      .end()
  }

  Collapse.prototype.addAriaAndCollapsedClass = function ($element, $trigger) {
    var isOpen = $element.hasClass('in')

    $element.attr('aria-expanded', isOpen)
    $trigger
      .toggleClass('collapsed', !isOpen)
      .attr('aria-expanded', isOpen)
  }

  function getTargetFromTrigger($trigger) {
    var href
    var target = $trigger.attr('data-target')
      || (href = $trigger.attr('href')) && href.replace(/.*(?=#[^\s]+$)/, '') // strip for ie7

    return $(target)
  }


  // COLLAPSE PLUGIN DEFINITION
  // ==========================

  function Plugin(option) {
    return this.each(function () {
      var $this   = $(this)
      var data    = $this.data('bs.collapse')
      var options = $.extend({}, Collapse.DEFAULTS, $this.data(), typeof option == 'object' && option)

      if (!data && options.toggle && /show|hide/.test(option)) options.toggle = false
      if (!data) $this.data('bs.collapse', (data = new Collapse(this, options)))
      if (typeof option == 'string') data[option]()
    })
  }

  var old = $.fn.collapse

  $.fn.collapse             = Plugin
  $.fn.collapse.Constructor = Collapse


  // COLLAPSE NO CONFLICT
  // ====================

  $.fn.collapse.noConflict = function () {
    $.fn.collapse = old
    return this
  }


  // COLLAPSE DATA-API
  // =================

  $(document).on('click.bs.collapse.data-api', '[data-toggle="collapse"]', function (e) {
    var $this   = $(this)

    if (!$this.attr('data-target')) e.preventDefault()

    var $target = getTargetFromTrigger($this)
    var data    = $target.data('bs.collapse')
    var option  = data ? 'toggle' : $this.data()

    Plugin.call($target, option)
  })

}(jQuery);

/* ========================================================================
 * Bootstrap: dropdown.js v3.3.5
 * http://getbootstrap.com/javascript/#dropdowns
 * ========================================================================
 * Copyright 2011-2015 Twitter, Inc.
 * Licensed under MIT (https://github.com/twbs/bootstrap/blob/master/LICENSE)
 * ======================================================================== */


+function ($) {
  'use strict';

  // DROPDOWN CLASS DEFINITION
  // =========================

  var backdrop = '.dropdown-backdrop'
  var toggle   = '[data-toggle="dropdown"]'
  var Dropdown = function (element) {
    $(element).on('click.bs.dropdown', this.toggle)
  }

  Dropdown.VERSION = '3.3.5'

  function getParent($this) {
    var selector = $this.attr('data-target')

    if (!selector) {
      selector = $this.attr('href')
      selector = selector && /#[A-Za-z]/.test(selector) && selector.replace(/.*(?=#[^\s]*$)/, '') // strip for ie7
    }

    var $parent = selector && $(selector)

    return $parent && $parent.length ? $parent : $this.parent()
  }

  function clearMenus(e) {
    if (e && e.which === 3) return
    $(backdrop).remove()
    $(toggle).each(function () {
      var $this         = $(this)
      var $parent       = getParent($this)
      var relatedTarget = { relatedTarget: this }

      if (!$parent.hasClass('open')) return

      if (e && e.type == 'click' && /input|textarea/i.test(e.target.tagName) && $.contains($parent[0], e.target)) return

      $parent.trigger(e = $.Event('hide.bs.dropdown', relatedTarget))

      if (e.isDefaultPrevented()) return

      $this.attr('aria-expanded', 'false')
      $parent.removeClass('open').trigger('hidden.bs.dropdown', relatedTarget)
    })
  }

  Dropdown.prototype.toggle = function (e) {
    var $this = $(this)

    if ($this.is('.disabled, :disabled')) return

    var $parent  = getParent($this)
    var isActive = $parent.hasClass('open')

    clearMenus()

    if (!isActive) {
      if ('ontouchstart' in document.documentElement && !$parent.closest('.navbar-nav').length) {
        // if mobile we use a backdrop because click events don't delegate
        $(document.createElement('div'))
          .addClass('dropdown-backdrop')
          .insertAfter($(this))
          .on('click', clearMenus)
      }

      var relatedTarget = { relatedTarget: this }
      $parent.trigger(e = $.Event('show.bs.dropdown', relatedTarget))

      if (e.isDefaultPrevented()) return

      $this
        .trigger('focus')
        .attr('aria-expanded', 'true')

      $parent
        .toggleClass('open')
        .trigger('shown.bs.dropdown', relatedTarget)
    }

    return false
  }

  Dropdown.prototype.keydown = function (e) {
    if (!/(38|40|27|32)/.test(e.which) || /input|textarea/i.test(e.target.tagName)) return

    var $this = $(this)

    e.preventDefault()
    e.stopPropagation()

    if ($this.is('.disabled, :disabled')) return

    var $parent  = getParent($this)
    var isActive = $parent.hasClass('open')

    if (!isActive && e.which != 27 || isActive && e.which == 27) {
      if (e.which == 27) $parent.find(toggle).trigger('focus')
      return $this.trigger('click')
    }

    var desc = ' li:not(.disabled):visible a'
    var $items = $parent.find('.dropdown-menu' + desc)

    if (!$items.length) return

    var index = $items.index(e.target)

    if (e.which == 38 && index > 0)                 index--         // up
    if (e.which == 40 && index < $items.length - 1) index++         // down
    if (!~index)                                    index = 0

    $items.eq(index).trigger('focus')
  }


  // DROPDOWN PLUGIN DEFINITION
  // ==========================

  function Plugin(option) {
    return this.each(function () {
      var $this = $(this)
      var data  = $this.data('bs.dropdown')

      if (!data) $this.data('bs.dropdown', (data = new Dropdown(this)))
      if (typeof option == 'string') data[option].call($this)
    })
  }

  var old = $.fn.dropdown

  $.fn.dropdown             = Plugin
  $.fn.dropdown.Constructor = Dropdown


  // DROPDOWN NO CONFLICT
  // ====================

  $.fn.dropdown.noConflict = function () {
    $.fn.dropdown = old
    return this
  }


  // APPLY TO STANDARD DROPDOWN ELEMENTS
  // ===================================

  $(document)
    .on('click.bs.dropdown.data-api', clearMenus)
    .on('click.bs.dropdown.data-api', '.dropdown form', function (e) { e.stopPropagation() })
    .on('click.bs.dropdown.data-api', toggle, Dropdown.prototype.toggle)
    .on('keydown.bs.dropdown.data-api', toggle, Dropdown.prototype.keydown)
    .on('keydown.bs.dropdown.data-api', '.dropdown-menu', Dropdown.prototype.keydown)

}(jQuery);

/* ========================================================================
 * Bootstrap: modal.js v3.3.5
 * http://getbootstrap.com/javascript/#modals
 * ========================================================================
 * Copyright 2011-2015 Twitter, Inc.
 * Licensed under MIT (https://github.com/twbs/bootstrap/blob/master/LICENSE)
 * ======================================================================== */


+function ($) {
  'use strict';

  // MODAL CLASS DEFINITION
  // ======================

  var Modal = function (element, options) {
    this.options             = options
    this.$body               = $(document.body)
    this.$element            = $(element)
    this.$dialog             = this.$element.find('.modal-dialog')
    this.$backdrop           = null
    this.isShown             = null
    this.originalBodyPad     = null
    this.scrollbarWidth      = 0
    this.ignoreBackdropClick = false

    if (this.options.remote) {
      this.$element
        .find('.modal-content')
        .load(this.options.remote, $.proxy(function () {
          this.$element.trigger('loaded.bs.modal')
        }, this))
    }
  }

  Modal.VERSION  = '3.3.5'

  Modal.TRANSITION_DURATION = 300
  Modal.BACKDROP_TRANSITION_DURATION = 150

  Modal.DEFAULTS = {
    backdrop: true,
    keyboard: true,
    show: true
  }

  Modal.prototype.toggle = function (_relatedTarget) {
    return this.isShown ? this.hide() : this.show(_relatedTarget)
  }

  Modal.prototype.show = function (_relatedTarget) {
    var that = this
    var e    = $.Event('show.bs.modal', { relatedTarget: _relatedTarget })

    this.$element.trigger(e)

    if (this.isShown || e.isDefaultPrevented()) return

    this.isShown = true

    this.checkScrollbar()
    this.setScrollbar()
    this.$body.addClass('modal-open')

    this.escape()
    this.resize()

    this.$element.on('click.dismiss.bs.modal', '[data-dismiss="modal"]', $.proxy(this.hide, this))

    this.$dialog.on('mousedown.dismiss.bs.modal', function () {
      that.$element.one('mouseup.dismiss.bs.modal', function (e) {
        if ($(e.target).is(that.$element)) that.ignoreBackdropClick = true
      })
    })

    this.backdrop(function () {
      var transition = $.support.transition && that.$element.hasClass('fade')

      if (!that.$element.parent().length) {
        that.$element.appendTo(that.$body) // don't move modals dom position
      }

      that.$element
        .show()
        .scrollTop(0)

      that.adjustDialog()

      if (transition) {
        that.$element[0].offsetWidth // force reflow
      }

      that.$element.addClass('in')

      that.enforceFocus()

      var e = $.Event('shown.bs.modal', { relatedTarget: _relatedTarget })

      transition ?
        that.$dialog // wait for modal to slide in
          .one('bsTransitionEnd', function () {
            that.$element.trigger('focus').trigger(e)
          })
          .emulateTransitionEnd(Modal.TRANSITION_DURATION) :
        that.$element.trigger('focus').trigger(e)
    })
  }

  Modal.prototype.hide = function (e) {
    if (e) e.preventDefault()

    e = $.Event('hide.bs.modal')

    this.$element.trigger(e)

    if (!this.isShown || e.isDefaultPrevented()) return

    this.isShown = false

    this.escape()
    this.resize()

    $(document).off('focusin.bs.modal')

    this.$element
      .removeClass('in')
      .off('click.dismiss.bs.modal')
      .off('mouseup.dismiss.bs.modal')

    this.$dialog.off('mousedown.dismiss.bs.modal')

    $.support.transition && this.$element.hasClass('fade') ?
      this.$element
        .one('bsTransitionEnd', $.proxy(this.hideModal, this))
        .emulateTransitionEnd(Modal.TRANSITION_DURATION) :
      this.hideModal()
  }

  Modal.prototype.enforceFocus = function () {
    $(document)
      .off('focusin.bs.modal') // guard against infinite focus loop
      .on('focusin.bs.modal', $.proxy(function (e) {
        if (this.$element[0] !== e.target && !this.$element.has(e.target).length) {
          this.$element.trigger('focus')
        }
      }, this))
  }

  Modal.prototype.escape = function () {
    if (this.isShown && this.options.keyboard) {
      this.$element.on('keydown.dismiss.bs.modal', $.proxy(function (e) {
        e.which == 27 && this.hide()
      }, this))
    } else if (!this.isShown) {
      this.$element.off('keydown.dismiss.bs.modal')
    }
  }

  Modal.prototype.resize = function () {
    if (this.isShown) {
      $(window).on('resize.bs.modal', $.proxy(this.handleUpdate, this))
    } else {
      $(window).off('resize.bs.modal')
    }
  }

  Modal.prototype.hideModal = function () {
    var that = this
    this.$element.hide()
    this.backdrop(function () {
      that.$body.removeClass('modal-open')
      that.resetAdjustments()
      that.resetScrollbar()
      that.$element.trigger('hidden.bs.modal')
    })
  }

  Modal.prototype.removeBackdrop = function () {
    this.$backdrop && this.$backdrop.remove()
    this.$backdrop = null
  }

  Modal.prototype.backdrop = function (callback) {
    var that = this
    var animate = this.$element.hasClass('fade') ? 'fade' : ''

    if (this.isShown && this.options.backdrop) {
      var doAnimate = $.support.transition && animate

      this.$backdrop = $(document.createElement('div'))
        .addClass('modal-backdrop ' + animate)
        .appendTo(this.$body)

      this.$element.on('click.dismiss.bs.modal', $.proxy(function (e) {
        if (this.ignoreBackdropClick) {
          this.ignoreBackdropClick = false
          return
        }
        if (e.target !== e.currentTarget) return
        this.options.backdrop == 'static'
          ? this.$element[0].focus()
          : this.hide()
      }, this))

      if (doAnimate) this.$backdrop[0].offsetWidth // force reflow

      this.$backdrop.addClass('in')

      if (!callback) return

      doAnimate ?
        this.$backdrop
          .one('bsTransitionEnd', callback)
          .emulateTransitionEnd(Modal.BACKDROP_TRANSITION_DURATION) :
        callback()

    } else if (!this.isShown && this.$backdrop) {
      this.$backdrop.removeClass('in')

      var callbackRemove = function () {
        that.removeBackdrop()
        callback && callback()
      }
      $.support.transition && this.$element.hasClass('fade') ?
        this.$backdrop
          .one('bsTransitionEnd', callbackRemove)
          .emulateTransitionEnd(Modal.BACKDROP_TRANSITION_DURATION) :
        callbackRemove()

    } else if (callback) {
      callback()
    }
  }

  // these following methods are used to handle overflowing modals

  Modal.prototype.handleUpdate = function () {
    this.adjustDialog()
  }

  Modal.prototype.adjustDialog = function () {
    var modalIsOverflowing = this.$element[0].scrollHeight > document.documentElement.clientHeight

    this.$element.css({
      paddingLeft:  !this.bodyIsOverflowing && modalIsOverflowing ? this.scrollbarWidth : '',
      paddingRight: this.bodyIsOverflowing && !modalIsOverflowing ? this.scrollbarWidth : ''
    })
  }

  Modal.prototype.resetAdjustments = function () {
    this.$element.css({
      paddingLeft: '',
      paddingRight: ''
    })
  }

  Modal.prototype.checkScrollbar = function () {
    var fullWindowWidth = window.innerWidth
    if (!fullWindowWidth) { // workaround for missing window.innerWidth in IE8
      var documentElementRect = document.documentElement.getBoundingClientRect()
      fullWindowWidth = documentElementRect.right - Math.abs(documentElementRect.left)
    }
    this.bodyIsOverflowing = document.body.clientWidth < fullWindowWidth
    this.scrollbarWidth = this.measureScrollbar()
  }

  Modal.prototype.setScrollbar = function () {
    var bodyPad = parseInt((this.$body.css('padding-right') || 0), 10)
    this.originalBodyPad = document.body.style.paddingRight || ''
    if (this.bodyIsOverflowing) this.$body.css('padding-right', bodyPad + this.scrollbarWidth)
  }

  Modal.prototype.resetScrollbar = function () {
    this.$body.css('padding-right', this.originalBodyPad)
  }

  Modal.prototype.measureScrollbar = function () { // thx walsh
    var scrollDiv = document.createElement('div')
    scrollDiv.className = 'modal-scrollbar-measure'
    this.$body.append(scrollDiv)
    var scrollbarWidth = scrollDiv.offsetWidth - scrollDiv.clientWidth
    this.$body[0].removeChild(scrollDiv)
    return scrollbarWidth
  }


  // MODAL PLUGIN DEFINITION
  // =======================

  function Plugin(option, _relatedTarget) {
    return this.each(function () {
      var $this   = $(this)
      var data    = $this.data('bs.modal')
      var options = $.extend({}, Modal.DEFAULTS, $this.data(), typeof option == 'object' && option)

      if (!data) $this.data('bs.modal', (data = new Modal(this, options)))
      if (typeof option == 'string') data[option](_relatedTarget)
      else if (options.show) data.show(_relatedTarget)
    })
  }

  var old = $.fn.modal

  $.fn.modal             = Plugin
  $.fn.modal.Constructor = Modal


  // MODAL NO CONFLICT
  // =================

  $.fn.modal.noConflict = function () {
    $.fn.modal = old
    return this
  }


  // MODAL DATA-API
  // ==============

  $(document).on('click.bs.modal.data-api', '[data-toggle="modal"]', function (e) {
    var $this   = $(this)
    var href    = $this.attr('href')
    var $target = $($this.attr('data-target') || (href && href.replace(/.*(?=#[^\s]+$)/, ''))) // strip for ie7
    var option  = $target.data('bs.modal') ? 'toggle' : $.extend({ remote: !/#/.test(href) && href }, $target.data(), $this.data())

    if ($this.is('a')) e.preventDefault()

    $target.one('show.bs.modal', function (showEvent) {
      if (showEvent.isDefaultPrevented()) return // only register focus restorer if modal will actually get shown
      $target.one('hidden.bs.modal', function () {
        $this.is(':visible') && $this.trigger('focus')
      })
    })
    Plugin.call($target, option, this)
  })

}(jQuery);

/* ========================================================================
 * Bootstrap: tooltip.js v3.3.5
 * http://getbootstrap.com/javascript/#tooltip
 * Inspired by the original jQuery.tipsy by Jason Frame
 * ========================================================================
 * Copyright 2011-2015 Twitter, Inc.
 * Licensed under MIT (https://github.com/twbs/bootstrap/blob/master/LICENSE)
 * ======================================================================== */


+function ($) {
  'use strict';

  // TOOLTIP PUBLIC CLASS DEFINITION
  // ===============================

  var Tooltip = function (element, options) {
    this.type       = null
    this.options    = null
    this.enabled    = null
    this.timeout    = null
    this.hoverState = null
    this.$element   = null
    this.inState    = null

    this.init('tooltip', element, options)
  }

  Tooltip.VERSION  = '3.3.5'

  Tooltip.TRANSITION_DURATION = 150

  Tooltip.DEFAULTS = {
    animation: true,
    placement: 'top',
    selector: false,
    template: '<div class="tooltip" role="tooltip"><div class="tooltip-arrow"></div><div class="tooltip-inner"></div></div>',
    trigger: 'hover focus',
    title: '',
    delay: 0,
    html: false,
    container: false,
    viewport: {
      selector: 'body',
      padding: 0
    }
  }

  Tooltip.prototype.init = function (type, element, options) {
    this.enabled   = true
    this.type      = type
    this.$element  = $(element)
    this.options   = this.getOptions(options)
    this.$viewport = this.options.viewport && $($.isFunction(this.options.viewport) ? this.options.viewport.call(this, this.$element) : (this.options.viewport.selector || this.options.viewport))
    this.inState   = { click: false, hover: false, focus: false }

    if (this.$element[0] instanceof document.constructor && !this.options.selector) {
      throw new Error('`selector` option must be specified when initializing ' + this.type + ' on the window.document object!')
    }

    var triggers = this.options.trigger.split(' ')

    for (var i = triggers.length; i--;) {
      var trigger = triggers[i]

      if (trigger == 'click') {
        this.$element.on('click.' + this.type, this.options.selector, $.proxy(this.toggle, this))
      } else if (trigger != 'manual') {
        var eventIn  = trigger == 'hover' ? 'mouseenter' : 'focusin'
        var eventOut = trigger == 'hover' ? 'mouseleave' : 'focusout'

        this.$element.on(eventIn  + '.' + this.type, this.options.selector, $.proxy(this.enter, this))
        this.$element.on(eventOut + '.' + this.type, this.options.selector, $.proxy(this.leave, this))
      }
    }

    this.options.selector ?
      (this._options = $.extend({}, this.options, { trigger: 'manual', selector: '' })) :
      this.fixTitle()
  }

  Tooltip.prototype.getDefaults = function () {
    return Tooltip.DEFAULTS
  }

  Tooltip.prototype.getOptions = function (options) {
    options = $.extend({}, this.getDefaults(), this.$element.data(), options)

    if (options.delay && typeof options.delay == 'number') {
      options.delay = {
        show: options.delay,
        hide: options.delay
      }
    }

    return options
  }

  Tooltip.prototype.getDelegateOptions = function () {
    var options  = {}
    var defaults = this.getDefaults()

    this._options && $.each(this._options, function (key, value) {
      if (defaults[key] != value) options[key] = value
    })

    return options
  }

  Tooltip.prototype.enter = function (obj) {
    var self = obj instanceof this.constructor ?
      obj : $(obj.currentTarget).data('bs.' + this.type)

    if (!self) {
      self = new this.constructor(obj.currentTarget, this.getDelegateOptions())
      $(obj.currentTarget).data('bs.' + this.type, self)
    }

    if (obj instanceof $.Event) {
      self.inState[obj.type == 'focusin' ? 'focus' : 'hover'] = true
    }

    if (self.tip().hasClass('in') || self.hoverState == 'in') {
      self.hoverState = 'in'
      return
    }

    clearTimeout(self.timeout)

    self.hoverState = 'in'

    if (!self.options.delay || !self.options.delay.show) return self.show()

    self.timeout = setTimeout(function () {
      if (self.hoverState == 'in') self.show()
    }, self.options.delay.show)
  }

  Tooltip.prototype.isInStateTrue = function () {
    for (var key in this.inState) {
      if (this.inState[key]) return true
    }

    return false
  }

  Tooltip.prototype.leave = function (obj) {
    var self = obj instanceof this.constructor ?
      obj : $(obj.currentTarget).data('bs.' + this.type)

    if (!self) {
      self = new this.constructor(obj.currentTarget, this.getDelegateOptions())
      $(obj.currentTarget).data('bs.' + this.type, self)
    }

    if (obj instanceof $.Event) {
      self.inState[obj.type == 'focusout' ? 'focus' : 'hover'] = false
    }

    if (self.isInStateTrue()) return

    clearTimeout(self.timeout)

    self.hoverState = 'out'

    if (!self.options.delay || !self.options.delay.hide) return self.hide()

    self.timeout = setTimeout(function () {
      if (self.hoverState == 'out') self.hide()
    }, self.options.delay.hide)
  }

  Tooltip.prototype.show = function () {
    var e = $.Event('show.bs.' + this.type)

    if (this.hasContent() && this.enabled) {
      this.$element.trigger(e)

      var inDom = $.contains(this.$element[0].ownerDocument.documentElement, this.$element[0])
      if (e.isDefaultPrevented() || !inDom) return
      var that = this

      var $tip = this.tip()

      var tipId = this.getUID(this.type)

      this.setContent()
      $tip.attr('id', tipId)
      this.$element.attr('aria-describedby', tipId)

      if (this.options.animation) $tip.addClass('fade')

      var placement = typeof this.options.placement == 'function' ?
        this.options.placement.call(this, $tip[0], this.$element[0]) :
        this.options.placement

      var autoToken = /\s?auto?\s?/i
      var autoPlace = autoToken.test(placement)
      if (autoPlace) placement = placement.replace(autoToken, '') || 'top'

      $tip
        .detach()
        .css({ top: 0, left: 0, display: 'block' })
        .addClass(placement)
        .data('bs.' + this.type, this)

      this.options.container ? $tip.appendTo(this.options.container) : $tip.insertAfter(this.$element)
      this.$element.trigger('inserted.bs.' + this.type)

      var pos          = this.getPosition()
      var actualWidth  = $tip[0].offsetWidth
      var actualHeight = $tip[0].offsetHeight

      if (autoPlace) {
        var orgPlacement = placement
        var viewportDim = this.getPosition(this.$viewport)

        placement = placement == 'bottom' && pos.bottom + actualHeight > viewportDim.bottom ? 'top'    :
                    placement == 'top'    && pos.top    - actualHeight < viewportDim.top    ? 'bottom' :
                    placement == 'right'  && pos.right  + actualWidth  > viewportDim.width  ? 'left'   :
                    placement == 'left'   && pos.left   - actualWidth  < viewportDim.left   ? 'right'  :
                    placement

        $tip
          .removeClass(orgPlacement)
          .addClass(placement)
      }

      var calculatedOffset = this.getCalculatedOffset(placement, pos, actualWidth, actualHeight)

      this.applyPlacement(calculatedOffset, placement)

      var complete = function () {
        var prevHoverState = that.hoverState
        that.$element.trigger('shown.bs.' + that.type)
        that.hoverState = null

        if (prevHoverState == 'out') that.leave(that)
      }

      $.support.transition && this.$tip.hasClass('fade') ?
        $tip
          .one('bsTransitionEnd', complete)
          .emulateTransitionEnd(Tooltip.TRANSITION_DURATION) :
        complete()
    }
  }

  Tooltip.prototype.applyPlacement = function (offset, placement) {
    var $tip   = this.tip()
    var width  = $tip[0].offsetWidth
    var height = $tip[0].offsetHeight

    // manually read margins because getBoundingClientRect includes difference
    var marginTop = parseInt($tip.css('margin-top'), 10)
    var marginLeft = parseInt($tip.css('margin-left'), 10)

    // we must check for NaN for ie 8/9
    if (isNaN(marginTop))  marginTop  = 0
    if (isNaN(marginLeft)) marginLeft = 0

    offset.top  += marginTop
    offset.left += marginLeft

    // $.fn.offset doesn't round pixel values
    // so we use setOffset directly with our own function B-0
    $.offset.setOffset($tip[0], $.extend({
      using: function (props) {
        $tip.css({
          top: Math.round(props.top),
          left: Math.round(props.left)
        })
      }
    }, offset), 0)

    $tip.addClass('in')

    // check to see if placing tip in new offset caused the tip to resize itself
    var actualWidth  = $tip[0].offsetWidth
    var actualHeight = $tip[0].offsetHeight

    if (placement == 'top' && actualHeight != height) {
      offset.top = offset.top + height - actualHeight
    }

    var delta = this.getViewportAdjustedDelta(placement, offset, actualWidth, actualHeight)

    if (delta.left) offset.left += delta.left
    else offset.top += delta.top

    var isVertical          = /top|bottom/.test(placement)
    var arrowDelta          = isVertical ? delta.left * 2 - width + actualWidth : delta.top * 2 - height + actualHeight
    var arrowOffsetPosition = isVertical ? 'offsetWidth' : 'offsetHeight'

    $tip.offset(offset)
    this.replaceArrow(arrowDelta, $tip[0][arrowOffsetPosition], isVertical)
  }

  Tooltip.prototype.replaceArrow = function (delta, dimension, isVertical) {
    this.arrow()
      .css(isVertical ? 'left' : 'top', 50 * (1 - delta / dimension) + '%')
      .css(isVertical ? 'top' : 'left', '')
  }

  Tooltip.prototype.setContent = function () {
    var $tip  = this.tip()
    var title = this.getTitle()

    $tip.find('.tooltip-inner')[this.options.html ? 'html' : 'text'](title)
    $tip.removeClass('fade in top bottom left right')
  }

  Tooltip.prototype.hide = function (callback) {
    var that = this
    var $tip = $(this.$tip)
    var e    = $.Event('hide.bs.' + this.type)

    function complete() {
      if (that.hoverState != 'in') $tip.detach()
      that.$element
        .removeAttr('aria-describedby')
        .trigger('hidden.bs.' + that.type)
      callback && callback()
    }

    this.$element.trigger(e)

    if (e.isDefaultPrevented()) return

    $tip.removeClass('in')

    $.support.transition && $tip.hasClass('fade') ?
      $tip
        .one('bsTransitionEnd', complete)
        .emulateTransitionEnd(Tooltip.TRANSITION_DURATION) :
      complete()

    this.hoverState = null

    return this
  }

  Tooltip.prototype.fixTitle = function () {
    var $e = this.$element
    if ($e.attr('title') || typeof $e.attr('data-original-title') != 'string') {
      $e.attr('data-original-title', $e.attr('title') || '').attr('title', '')
    }
  }

  Tooltip.prototype.hasContent = function () {
    return this.getTitle()
  }

  Tooltip.prototype.getPosition = function ($element) {
    $element   = $element || this.$element

    var el     = $element[0]
    var isBody = el.tagName == 'BODY'

    var elRect    = el.getBoundingClientRect()
    if (elRect.width == null) {
      // width and height are missing in IE8, so compute them manually; see https://github.com/twbs/bootstrap/issues/14093
      elRect = $.extend({}, elRect, { width: elRect.right - elRect.left, height: elRect.bottom - elRect.top })
    }
    var elOffset  = isBody ? { top: 0, left: 0 } : $element.offset()
    var scroll    = { scroll: isBody ? document.documentElement.scrollTop || document.body.scrollTop : $element.scrollTop() }
    var outerDims = isBody ? { width: $(window).width(), height: $(window).height() } : null

    return $.extend({}, elRect, scroll, outerDims, elOffset)
  }

  Tooltip.prototype.getCalculatedOffset = function (placement, pos, actualWidth, actualHeight) {
    return placement == 'bottom' ? { top: pos.top + pos.height,   left: pos.left + pos.width / 2 - actualWidth / 2 } :
           placement == 'top'    ? { top: pos.top - actualHeight, left: pos.left + pos.width / 2 - actualWidth / 2 } :
           placement == 'left'   ? { top: pos.top + pos.height / 2 - actualHeight / 2, left: pos.left - actualWidth } :
        /* placement == 'right' */ { top: pos.top + pos.height / 2 - actualHeight / 2, left: pos.left + pos.width }

  }

  Tooltip.prototype.getViewportAdjustedDelta = function (placement, pos, actualWidth, actualHeight) {
    var delta = { top: 0, left: 0 }
    if (!this.$viewport) return delta

    var viewportPadding = this.options.viewport && this.options.viewport.padding || 0
    var viewportDimensions = this.getPosition(this.$viewport)

    if (/right|left/.test(placement)) {
      var topEdgeOffset    = pos.top - viewportPadding - viewportDimensions.scroll
      var bottomEdgeOffset = pos.top + viewportPadding - viewportDimensions.scroll + actualHeight
      if (topEdgeOffset < viewportDimensions.top) { // top overflow
        delta.top = viewportDimensions.top - topEdgeOffset
      } else if (bottomEdgeOffset > viewportDimensions.top + viewportDimensions.height) { // bottom overflow
        delta.top = viewportDimensions.top + viewportDimensions.height - bottomEdgeOffset
      }
    } else {
      var leftEdgeOffset  = pos.left - viewportPadding
      var rightEdgeOffset = pos.left + viewportPadding + actualWidth
      if (leftEdgeOffset < viewportDimensions.left) { // left overflow
        delta.left = viewportDimensions.left - leftEdgeOffset
      } else if (rightEdgeOffset > viewportDimensions.right) { // right overflow
        delta.left = viewportDimensions.left + viewportDimensions.width - rightEdgeOffset
      }
    }

    return delta
  }

  Tooltip.prototype.getTitle = function () {
    var title
    var $e = this.$element
    var o  = this.options

    title = $e.attr('data-original-title')
      || (typeof o.title == 'function' ? o.title.call($e[0]) :  o.title)

    return title
  }

  Tooltip.prototype.getUID = function (prefix) {
    do prefix += ~~(Math.random() * 1000000)
    while (document.getElementById(prefix))
    return prefix
  }

  Tooltip.prototype.tip = function () {
    if (!this.$tip) {
      this.$tip = $(this.options.template)
      if (this.$tip.length != 1) {
        throw new Error(this.type + ' `template` option must consist of exactly 1 top-level element!')
      }
    }
    return this.$tip
  }

  Tooltip.prototype.arrow = function () {
    return (this.$arrow = this.$arrow || this.tip().find('.tooltip-arrow'))
  }

  Tooltip.prototype.enable = function () {
    this.enabled = true
  }

  Tooltip.prototype.disable = function () {
    this.enabled = false
  }

  Tooltip.prototype.toggleEnabled = function () {
    this.enabled = !this.enabled
  }

  Tooltip.prototype.toggle = function (e) {
    var self = this
    if (e) {
      self = $(e.currentTarget).data('bs.' + this.type)
      if (!self) {
        self = new this.constructor(e.currentTarget, this.getDelegateOptions())
        $(e.currentTarget).data('bs.' + this.type, self)
      }
    }

    if (e) {
      self.inState.click = !self.inState.click
      if (self.isInStateTrue()) self.enter(self)
      else self.leave(self)
    } else {
      self.tip().hasClass('in') ? self.leave(self) : self.enter(self)
    }
  }

  Tooltip.prototype.destroy = function () {
    var that = this
    clearTimeout(this.timeout)
    this.hide(function () {
      that.$element.off('.' + that.type).removeData('bs.' + that.type)
      if (that.$tip) {
        that.$tip.detach()
      }
      that.$tip = null
      that.$arrow = null
      that.$viewport = null
    })
  }


  // TOOLTIP PLUGIN DEFINITION
  // =========================

  function Plugin(option) {
    return this.each(function () {
      var $this   = $(this)
      var data    = $this.data('bs.tooltip')
      var options = typeof option == 'object' && option

      if (!data && /destroy|hide/.test(option)) return
      if (!data) $this.data('bs.tooltip', (data = new Tooltip(this, options)))
      if (typeof option == 'string') data[option]()
    })
  }

  var old = $.fn.tooltip

  $.fn.tooltip             = Plugin
  $.fn.tooltip.Constructor = Tooltip


  // TOOLTIP NO CONFLICT
  // ===================

  $.fn.tooltip.noConflict = function () {
    $.fn.tooltip = old
    return this
  }

}(jQuery);

/* ========================================================================
 * Bootstrap: popover.js v3.3.5
 * http://getbootstrap.com/javascript/#popovers
 * ========================================================================
 * Copyright 2011-2015 Twitter, Inc.
 * Licensed under MIT (https://github.com/twbs/bootstrap/blob/master/LICENSE)
 * ======================================================================== */


+function ($) {
  'use strict';

  // POPOVER PUBLIC CLASS DEFINITION
  // ===============================

  var Popover = function (element, options) {
    this.init('popover', element, options)
  }

  if (!$.fn.tooltip) throw new Error('Popover requires tooltip.js')

  Popover.VERSION  = '3.3.5'

  Popover.DEFAULTS = $.extend({}, $.fn.tooltip.Constructor.DEFAULTS, {
    placement: 'right',
    trigger: 'click',
    content: '',
    template: '<div class="popover" role="tooltip"><div class="arrow"></div><h3 class="popover-title"></h3><div class="popover-content"></div></div>'
  })


  // NOTE: POPOVER EXTENDS tooltip.js
  // ================================

  Popover.prototype = $.extend({}, $.fn.tooltip.Constructor.prototype)

  Popover.prototype.constructor = Popover

  Popover.prototype.getDefaults = function () {
    return Popover.DEFAULTS
  }

  Popover.prototype.setContent = function () {
    var $tip    = this.tip()
    var title   = this.getTitle()
    var content = this.getContent()

    $tip.find('.popover-title')[this.options.html ? 'html' : 'text'](title)
    $tip.find('.popover-content').children().detach().end()[ // we use append for html objects to maintain js events
      this.options.html ? (typeof content == 'string' ? 'html' : 'append') : 'text'
    ](content)

    $tip.removeClass('fade top bottom left right in')

    // IE8 doesn't accept hiding via the `:empty` pseudo selector, we have to do
    // this manually by checking the contents.
    if (!$tip.find('.popover-title').html()) $tip.find('.popover-title').hide()
  }

  Popover.prototype.hasContent = function () {
    return this.getTitle() || this.getContent()
  }

  Popover.prototype.getContent = function () {
    var $e = this.$element
    var o  = this.options

    return $e.attr('data-content')
      || (typeof o.content == 'function' ?
            o.content.call($e[0]) :
            o.content)
  }

  Popover.prototype.arrow = function () {
    return (this.$arrow = this.$arrow || this.tip().find('.arrow'))
  }


  // POPOVER PLUGIN DEFINITION
  // =========================

  function Plugin(option) {
    return this.each(function () {
      var $this   = $(this)
      var data    = $this.data('bs.popover')
      var options = typeof option == 'object' && option

      if (!data && /destroy|hide/.test(option)) return
      if (!data) $this.data('bs.popover', (data = new Popover(this, options)))
      if (typeof option == 'string') data[option]()
    })
  }

  var old = $.fn.popover

  $.fn.popover             = Plugin
  $.fn.popover.Constructor = Popover


  // POPOVER NO CONFLICT
  // ===================

  $.fn.popover.noConflict = function () {
    $.fn.popover = old
    return this
  }

}(jQuery);

/* ========================================================================
 * Bootstrap: scrollspy.js v3.3.5
 * http://getbootstrap.com/javascript/#scrollspy
 * ========================================================================
 * Copyright 2011-2015 Twitter, Inc.
 * Licensed under MIT (https://github.com/twbs/bootstrap/blob/master/LICENSE)
 * ======================================================================== */


+function ($) {
  'use strict';

  // SCROLLSPY CLASS DEFINITION
  // ==========================

  function ScrollSpy(element, options) {
    this.$body          = $(document.body)
    this.$scrollElement = $(element).is(document.body) ? $(window) : $(element)
    this.options        = $.extend({}, ScrollSpy.DEFAULTS, options)
    this.selector       = (this.options.target || '') + ' .nav li > a'
    this.offsets        = []
    this.targets        = []
    this.activeTarget   = null
    this.scrollHeight   = 0

    this.$scrollElement.on('scroll.bs.scrollspy', $.proxy(this.process, this))
    this.refresh()
    this.process()
  }

  ScrollSpy.VERSION  = '3.3.5'

  ScrollSpy.DEFAULTS = {
    offset: 10
  }

  ScrollSpy.prototype.getScrollHeight = function () {
    return this.$scrollElement[0].scrollHeight || Math.max(this.$body[0].scrollHeight, document.documentElement.scrollHeight)
  }

  ScrollSpy.prototype.refresh = function () {
    var that          = this
    var offsetMethod  = 'offset'
    var offsetBase    = 0

    this.offsets      = []
    this.targets      = []
    this.scrollHeight = this.getScrollHeight()

    if (!$.isWindow(this.$scrollElement[0])) {
      offsetMethod = 'position'
      offsetBase   = this.$scrollElement.scrollTop()
    }

    this.$body
      .find(this.selector)
      .map(function () {
        var $el   = $(this)
        var href  = $el.data('target') || $el.attr('href')
        var $href = /^#./.test(href) && $(href)

        return ($href
          && $href.length
          && $href.is(':visible')
          && [[$href[offsetMethod]().top + offsetBase, href]]) || null
      })
      .sort(function (a, b) { return a[0] - b[0] })
      .each(function () {
        that.offsets.push(this[0])
        that.targets.push(this[1])
      })
  }

  ScrollSpy.prototype.process = function () {
    var scrollTop    = this.$scrollElement.scrollTop() + this.options.offset
    var scrollHeight = this.getScrollHeight()
    var maxScroll    = this.options.offset + scrollHeight - this.$scrollElement.height()
    var offsets      = this.offsets
    var targets      = this.targets
    var activeTarget = this.activeTarget
    var i

    if (this.scrollHeight != scrollHeight) {
      this.refresh()
    }

    if (scrollTop >= maxScroll) {
      return activeTarget != (i = targets[targets.length - 1]) && this.activate(i)
    }

    if (activeTarget && scrollTop < offsets[0]) {
      this.activeTarget = null
      return this.clear()
    }

    for (i = offsets.length; i--;) {
      activeTarget != targets[i]
        && scrollTop >= offsets[i]
        && (offsets[i + 1] === undefined || scrollTop < offsets[i + 1])
        && this.activate(targets[i])
    }
  }

  ScrollSpy.prototype.activate = function (target) {
    this.activeTarget = target

    this.clear()

    var selector = this.selector +
      '[data-target="' + target + '"],' +
      this.selector + '[href="' + target + '"]'

    var active = $(selector)
      .parents('li')
      .addClass('active')

    if (active.parent('.dropdown-menu').length) {
      active = active
        .closest('li.dropdown')
        .addClass('active')
    }

    active.trigger('activate.bs.scrollspy')
  }

  ScrollSpy.prototype.clear = function () {
    $(this.selector)
      .parentsUntil(this.options.target, '.active')
      .removeClass('active')
  }


  // SCROLLSPY PLUGIN DEFINITION
  // ===========================

  function Plugin(option) {
    return this.each(function () {
      var $this   = $(this)
      var data    = $this.data('bs.scrollspy')
      var options = typeof option == 'object' && option

      if (!data) $this.data('bs.scrollspy', (data = new ScrollSpy(this, options)))
      if (typeof option == 'string') data[option]()
    })
  }

  var old = $.fn.scrollspy

  $.fn.scrollspy             = Plugin
  $.fn.scrollspy.Constructor = ScrollSpy


  // SCROLLSPY NO CONFLICT
  // =====================

  $.fn.scrollspy.noConflict = function () {
    $.fn.scrollspy = old
    return this
  }


  // SCROLLSPY DATA-API
  // ==================

  $(window).on('load.bs.scrollspy.data-api', function () {
    $('[data-spy="scroll"]').each(function () {
      var $spy = $(this)
      Plugin.call($spy, $spy.data())
    })
  })

}(jQuery);

/* ========================================================================
 * Bootstrap: tab.js v3.3.5
 * http://getbootstrap.com/javascript/#tabs
 * ========================================================================
 * Copyright 2011-2015 Twitter, Inc.
 * Licensed under MIT (https://github.com/twbs/bootstrap/blob/master/LICENSE)
 * ======================================================================== */


+function ($) {
  'use strict';

  // TAB CLASS DEFINITION
  // ====================

  var Tab = function (element) {
    // jscs:disable requireDollarBeforejQueryAssignment
    this.element = $(element)
    // jscs:enable requireDollarBeforejQueryAssignment
  }

  Tab.VERSION = '3.3.5'

  Tab.TRANSITION_DURATION = 150

  Tab.prototype.show = function () {
    var $this    = this.element
    var $ul      = $this.closest('ul:not(.dropdown-menu)')
    var selector = $this.data('target')

    if (!selector) {
      selector = $this.attr('href')
      selector = selector && selector.replace(/.*(?=#[^\s]*$)/, '') // strip for ie7
    }

    if ($this.parent('li').hasClass('active')) return

    var $previous = $ul.find('.active:last a')
    var hideEvent = $.Event('hide.bs.tab', {
      relatedTarget: $this[0]
    })
    var showEvent = $.Event('show.bs.tab', {
      relatedTarget: $previous[0]
    })

    $previous.trigger(hideEvent)
    $this.trigger(showEvent)

    if (showEvent.isDefaultPrevented() || hideEvent.isDefaultPrevented()) return

    var $target = $(selector)

    this.activate($this.closest('li'), $ul)
    this.activate($target, $target.parent(), function () {
      $previous.trigger({
        type: 'hidden.bs.tab',
        relatedTarget: $this[0]
      })
      $this.trigger({
        type: 'shown.bs.tab',
        relatedTarget: $previous[0]
      })
    })
  }

  Tab.prototype.activate = function (element, container, callback) {
    var $active    = container.find('> .active')
    var transition = callback
      && $.support.transition
      && ($active.length && $active.hasClass('fade') || !!container.find('> .fade').length)

    function next() {
      $active
        .removeClass('active')
        .find('> .dropdown-menu > .active')
          .removeClass('active')
        .end()
        .find('[data-toggle="tab"]')
          .attr('aria-expanded', false)

      element
        .addClass('active')
        .find('[data-toggle="tab"]')
          .attr('aria-expanded', true)

      if (transition) {
        element[0].offsetWidth // reflow for transition
        element.addClass('in')
      } else {
        element.removeClass('fade')
      }

      if (element.parent('.dropdown-menu').length) {
        element
          .closest('li.dropdown')
            .addClass('active')
          .end()
          .find('[data-toggle="tab"]')
            .attr('aria-expanded', true)
      }

      callback && callback()
    }

    $active.length && transition ?
      $active
        .one('bsTransitionEnd', next)
        .emulateTransitionEnd(Tab.TRANSITION_DURATION) :
      next()

    $active.removeClass('in')
  }


  // TAB PLUGIN DEFINITION
  // =====================

  function Plugin(option) {
    return this.each(function () {
      var $this = $(this)
      var data  = $this.data('bs.tab')

      if (!data) $this.data('bs.tab', (data = new Tab(this)))
      if (typeof option == 'string') data[option]()
    })
  }

  var old = $.fn.tab

  $.fn.tab             = Plugin
  $.fn.tab.Constructor = Tab


  // TAB NO CONFLICT
  // ===============

  $.fn.tab.noConflict = function () {
    $.fn.tab = old
    return this
  }


  // TAB DATA-API
  // ============

  var clickHandler = function (e) {
    e.preventDefault()
    Plugin.call($(this), 'show')
  }

  $(document)
    .on('click.bs.tab.data-api', '[data-toggle="tab"]', clickHandler)
    .on('click.bs.tab.data-api', '[data-toggle="pill"]', clickHandler)

}(jQuery);

/* ========================================================================
 * Bootstrap: affix.js v3.3.5
 * http://getbootstrap.com/javascript/#affix
 * ========================================================================
 * Copyright 2011-2015 Twitter, Inc.
 * Licensed under MIT (https://github.com/twbs/bootstrap/blob/master/LICENSE)
 * ======================================================================== */


+function ($) {
  'use strict';

  // AFFIX CLASS DEFINITION
  // ======================

  var Affix = function (element, options) {
    this.options = $.extend({}, Affix.DEFAULTS, options)

    this.$target = $(this.options.target)
      .on('scroll.bs.affix.data-api', $.proxy(this.checkPosition, this))
      .on('click.bs.affix.data-api',  $.proxy(this.checkPositionWithEventLoop, this))

    this.$element     = $(element)
    this.affixed      = null
    this.unpin        = null
    this.pinnedOffset = null

    this.checkPosition()
  }

  Affix.VERSION  = '3.3.5'

  Affix.RESET    = 'affix affix-top affix-bottom'

  Affix.DEFAULTS = {
    offset: 0,
    target: window
  }

  Affix.prototype.getState = function (scrollHeight, height, offsetTop, offsetBottom) {
    var scrollTop    = this.$target.scrollTop()
    var position     = this.$element.offset()
    var targetHeight = this.$target.height()

    if (offsetTop != null && this.affixed == 'top') return scrollTop < offsetTop ? 'top' : false

    if (this.affixed == 'bottom') {
      if (offsetTop != null) return (scrollTop + this.unpin <= position.top) ? false : 'bottom'
      return (scrollTop + targetHeight <= scrollHeight - offsetBottom) ? false : 'bottom'
    }

    var initializing   = this.affixed == null
    var colliderTop    = initializing ? scrollTop : position.top
    var colliderHeight = initializing ? targetHeight : height

    if (offsetTop != null && scrollTop <= offsetTop) return 'top'
    if (offsetBottom != null && (colliderTop + colliderHeight >= scrollHeight - offsetBottom)) return 'bottom'

    return false
  }

  Affix.prototype.getPinnedOffset = function () {
    if (this.pinnedOffset) return this.pinnedOffset
    this.$element.removeClass(Affix.RESET).addClass('affix')
    var scrollTop = this.$target.scrollTop()
    var position  = this.$element.offset()
    return (this.pinnedOffset = position.top - scrollTop)
  }

  Affix.prototype.checkPositionWithEventLoop = function () {
    setTimeout($.proxy(this.checkPosition, this), 1)
  }

  Affix.prototype.checkPosition = function () {
    if (!this.$element.is(':visible')) return

    var height       = this.$element.height()
    var offset       = this.options.offset
    var offsetTop    = offset.top
    var offsetBottom = offset.bottom
    var scrollHeight = Math.max($(document).height(), $(document.body).height())

    if (typeof offset != 'object')         offsetBottom = offsetTop = offset
    if (typeof offsetTop == 'function')    offsetTop    = offset.top(this.$element)
    if (typeof offsetBottom == 'function') offsetBottom = offset.bottom(this.$element)

    var affix = this.getState(scrollHeight, height, offsetTop, offsetBottom)

    if (this.affixed != affix) {
      if (this.unpin != null) this.$element.css('top', '')

      var affixType = 'affix' + (affix ? '-' + affix : '')
      var e         = $.Event(affixType + '.bs.affix')

      this.$element.trigger(e)

      if (e.isDefaultPrevented()) return

      this.affixed = affix
      this.unpin = affix == 'bottom' ? this.getPinnedOffset() : null

      this.$element
        .removeClass(Affix.RESET)
        .addClass(affixType)
        .trigger(affixType.replace('affix', 'affixed') + '.bs.affix')
    }

    if (affix == 'bottom') {
      this.$element.offset({
        top: scrollHeight - height - offsetBottom
      })
    }
  }


  // AFFIX PLUGIN DEFINITION
  // =======================

  function Plugin(option) {
    return this.each(function () {
      var $this   = $(this)
      var data    = $this.data('bs.affix')
      var options = typeof option == 'object' && option

      if (!data) $this.data('bs.affix', (data = new Affix(this, options)))
      if (typeof option == 'string') data[option]()
    })
  }

  var old = $.fn.affix

  $.fn.affix             = Plugin
  $.fn.affix.Constructor = Affix


  // AFFIX NO CONFLICT
  // =================

  $.fn.affix.noConflict = function () {
    $.fn.affix = old
    return this
  }


  // AFFIX DATA-API
  // ==============

  $(window).on('load', function () {
    $('[data-spy="affix"]').each(function () {
      var $spy = $(this)
      var data = $spy.data()

      data.offset = data.offset || {}

      if (data.offsetBottom != null) data.offset.bottom = data.offsetBottom
      if (data.offsetTop    != null) data.offset.top    = data.offsetTop

      Plugin.call($spy, data)
    })
  })

}(jQuery);
```
