## Supplementary material for "Characterizing and inferring quantitative cell cycle phase in single-cell RNA-seq data analysis": Source code of our analysis, including scripts and data necessary to reproduce the work.: bootstrap.min.js

```
/*!
 * Bootstrap v3.3.5 (http://getbootstrap.com)
 * Copyright 2011-2015 Twitter, Inc.
 * Licensed under the MIT license
 */
if("undefined"==typeof jQuery)throw new Error("Bootstrap's JavaScript requires jQuery");+function(a){"use strict";var b=a.fn.jquery.split(" ")[0].split(".");if(b[0]<2&&b[1]<9||1==b[0]&&9==b[1]&&b[2]<1)throw new Error("Bootstrap's JavaScript requires jQuery version 1.9.1 or higher")}(jQuery),+function(a){"use strict";function b(){var a=document.createElement("bootstrap"),b={WebkitTransition:"webkitTransitionEnd",MozTransition:"transitionend",OTransition:"oTransitionEnd otransitionend",transition:"transitionend"};for(var c in b)if(void 0!==a.style[c])return{end:b[c]};return!1}a.fn.emulateTransitionEnd=function(b){var c=!1,d=this;a(this).one("bsTransitionEnd",function(){c=!0});var e=function(){c||a(d).trigger(a.support.transition.end)};return setTimeout(e,b),this},a(function(){a.support.transition=b(),a.support.transition&&(a.event.special.bsTransitionEnd={bindType:a.support.transition.end,delegateType:a.support.transition.end,handle:function(b){return a(b.target).is(this)?b.handleObj.handler.apply(this,arguments):void 0}})})}(jQuery),+function(a){"use strict";function b(b){return this.each(function(){var c=a(this),e=c.data("bs.alert");e||c.data("bs.alert",e=new d(this)),"string"==typeof b&&e[b].call(c)})}var c='[data-dismiss="alert"]',d=function(b){a(b).on("click",c,this.close)};d.VERSION="3.3.5",d.TRANSITION_DURATION=150,d.prototype.close=function(b){function c(){g.detach().trigger("closed.bs.alert").remove()}var e=a(this),f=e.attr("data-target");f||(f=e.attr("href"),f=f&&f.replace(/.*(?=#[^\s]*$)/,""));var g=a(f);b&&b.preventDefault(),g.length||(g=e.closest(".alert")),g.trigger(b=a.Event("close.bs.alert")),b.isDefaultPrevented()||(g.removeClass("in"),a.support.transition&&g.hasClass("fade")?g.one("bsTransitionEnd",c).emulateTransitionEnd(d.TRANSITION_DURATION):c())};var e=a.fn.alert;a.fn.alert=b,a.fn.alert.Constructor=d,a.fn.alert.noConflict=function(){return a.fn.alert=e,this},a(document).on("click.bs.alert.data-api",c,d.prototype.close)}(jQuery),+function(a){"use strict";function b(b){return this.each(function(){var d=a(this),e=d.data("bs.button"),f="object"==typeof b&&b;e||d.data("bs.button",e=new c(this,f)),"toggle"==b?e.toggle():b&&e.setState(b)})}var c=function(b,d){this.$element=a(b),this.options=a.extend({},c.DEFAULTS,d),this.isLoading=!1};c.VERSION="3.3.5",c.DEFAULTS={loadingText:"loading..."},c.prototype.setState=function(b){var c="disabled",d=this.$element,e=d.is("input")?"val":"html",f=d.data();b+="Text",null==f.resetText&&d.data("resetText",d[e]()),setTimeout(a.proxy(function(){d[e](null==f[b]?this.options[b]:f[b]),"loadingText"==b?(this.isLoading=!0,d.addClass(c).attr(c,c)):this.isLoading&&(this.isLoading=!1,d.removeClass(c).removeAttr(c))},this),0)},c.prototype.toggle=function(){var a=!0,b=this.$element.closest('[data-toggle="buttons"]');if(b.length){var c=this.$element.find("input");"radio"==c.prop("type")?(c.prop("checked")&&(a=!1),b.find(".active").removeClass("active"),this.$element.addClass("active")):"checkbox"==c.prop("type")&&(c.prop("checked")!==this.$element.hasClass("active")&&(a=!1),this.$element.toggleClass("active")),c.prop("checked",this.$element.hasClass("active")),a&&c.trigger("change")}else this.$element.attr("aria-pressed",!this.$element.hasClass("active")),this.$element.toggleClass("active")};var d=a.fn.button;a.fn.button=b,a.fn.button.Constructor=c,a.fn.button.noConflict=function(){return a.fn.button=d,this},a(document).on("click.bs.button.data-api",'[data-toggle^="button"]',function(c){var d=a(c.target);d.hasClass("btn")||(d=d.closest(".btn")),b.call(d,"toggle"),a(c.target).is('input[type="radio"]')||a(c.target).is('input[type="checkbox"]')||c.preventDefault()}).on("focus.bs.button.data-api blur.bs.button.data-api",'[data-toggle^="button"]',function(b){a(b.target).closest(".btn").toggleClass("focus",/^focus(in)?$/.test(b.type))})}(jQuery),+function(a){"use strict";function b(b){return this.each(function(){var d=a(this),e=d.data("bs.carousel"),f=a.extend({},c.DEFAULTS,d.data(),"object"==typeof b&&b),g="string"==typeof b?b:f.slide;e||d.data("bs.carousel",e=new c(this,f)),"number"==typeof b?e.to(b):g?e[g]():f.interval&&e.pause().cycle()})}var c=function(b,c){this.$element=a(b),this.$indicators=this.$element.find(".carousel-indicators"),this.options=c,this.paused=null,this.sliding=null,this.interval=null,this.$active=null,this.$items=null,this.options.keyboard&&this.$element.on("keydown.bs.carousel",a.proxy(this.keydown,this)),"hover"==this.options.pause&&!("ontouchstart"in document.documentElement)&&this.$element.on("mouseenter.bs.carousel",a.proxy(this.pause,this)).on("mouseleave.bs.carousel",a.proxy(this.cycle,this))};c.VERSION="3.3.5",c.TRANSITION_DURATION=600,c.DEFAULTS={interval:5e3,pause:"hover",wrap:!0,keyboard:!0},c.prototype.keydown=function(a){if(!/input|textarea/i.test(a.target.tagName)){switch(a.which){case 37:this.prev();break;case 39:this.next();break;default:return}a.preventDefault()}},c.prototype.cycle=function(b){return b||(this.paused=!1),this.interval&&clearInterval(this.interval),this.options.interval&&!this.paused&&(this.interval=setInterval(a.proxy(this.next,this),this.options.interval)),this},c.prototype.getItemIndex=function(a){return this.$items=a.parent().children(".item"),this.$items.index(a||this.$active)},c.prototype.getItemForDirection=function(a,b){var c=this.getItemIndex(b),d="prev"==a&&0===c||"next"==a&&c==this.$items.length-1;if(d&&!this.options.wrap)return b;var e="prev"==a?-1:1,f=(c+e)%this.$items.length;return this.$items.eq(f)},c.prototype.to=function(a){var b=this,c=this.getItemIndex(this.$active=this.$element.find(".item.active"));return a>this.$items.length-1||0>a?void 0:this.sliding?this.$element.one("slid.bs.carousel",function(){b.to(a)}):c==a?this.pause().cycle():this.slide(a>c?"next":"prev",this.$items.eq(a))},c.prototype.pause=function(b){return b||(this.paused=!0),this.$element.find(".next, .prev").length&&a.support.transition&&(this.$element.trigger(a.support.transition.end),this.cycle(!0)),this.interval=clearInterval(this.interval),this},c.prototype.next=function(){return this.sliding?void 0:this.slide("next")},c.prototype.prev=function(){return this.sliding?void 0:this.slide("prev")},c.prototype.slide=function(b,d){var e=this.$element.find(".item.active"),f=d||this.getItemForDirection(b,e),g=this.interval,h="next"==b?"left":"right",i=this;if(f.hasClass("active"))return this.sliding=!1;var j=f[0],k=a.Event("slide.bs.carousel",{relatedTarget:j,direction:h});if(this.$element.trigger(k),!k.isDefaultPrevented()){if(this.sliding=!0,g&&this.pause(),this.$indicators.length){this.$indicators.find(".active").removeClass("active");var l=a(this.$indicators.children()[this.getItemIndex(f)]);l&&l.addClass("active")}var m=a.Event("slid.bs.carousel",{relatedTarget:j,direction:h});return a.support.transition&&this.$element.hasClass("slide")?(f.addClass(b),f[0].offsetWidth,e.addClass(h),f.addClass(h),e.one("bsTransitionEnd",function(){f.removeClass([b,h].join(" ")).addClass("active"),e.removeClass(["active",h].join(" ")),i.sliding=!1,setTimeout(function(){i.$element.trigger(m)},0)}).emulateTransitionEnd(c.TRANSITION_DURATION)):(e.removeClass("active"),f.addClass("active"),this.sliding=!1,this.$element.trigger(m)),g&&this.cycle(),this}};var d=a.fn.carousel;a.fn.carousel=b,a.fn.carousel.Constructor=c,a.fn.carousel.noConflict=function(){return a.fn.carousel=d,this};var e=function(c){var d,e=a(this),f=a(e.attr("data-target")||(d=e.attr("href"))&&d.replace(/.*(?=#[^\s]+$)/,""));if(f.hasClass("carousel")){var g=a.extend({},f.data(),e.data()),h=e.attr("data-slide-to");h&&(g.interval=!1),b.call(f,g),h&&f.data("bs.carousel").to(h),c.preventDefault()}};a(document).on("click.bs.carousel.data-api","[data-slide]",e).on("click.bs.carousel.data-api","[data-slide-to]",e),a(window).on("load",function(){a('[data-ride="carousel"]').each(function(){var c=a(this);b.call(c,c.data())})})}(jQuery),+function(a){"use strict";function b(b){var c,d=b.attr("data-target")||(c=b.attr("href"))&&c.replace(/.*(?=#[^\s]+$)/,"");return a(d)}function c(b){return this.each(function(){var c=a(this),e=c.data("bs.collapse"),f=a.extend({},d.DEFAULTS,c.data(),"object"==typeof b&&b);!e&&f.toggle&&/show|hide/.test(b)&&(f.toggle=!1),e||c.data("bs.collapse",e=new d(this,f)),"string"==typeof b&&e[b]()})}var d=function(b,c){this.$element=a(b),this.options=a.extend({},d.DEFAULTS,c),this.$trigger=a('[data-toggle="collapse"][href="#'+b.id+'"],[data-toggle="collapse"][data-target="#'+b.id+'"]'),this.transitioning=null,this.options.parent?this.$parent=this.getParent():this.addAriaAndCollapsedClass(this.$element,this.$trigger),this.options.toggle&&this.toggle()};d.VERSION="3.3.5",d.TRANSITION_DURATION=350,d.DEFAULTS={toggle:!0},d.prototype.dimension=function(){var a=this.$element.hasClass("width");return a?"width":"height"},d.prototype.show=function(){if(!this.transitioning&&!this.$element.hasClass("in")){var b,e=this.$parent&&this.$parent.children(".panel").children(".in, .collapsing");if(!(e&&e.length&&(b=e.data("bs.collapse"),b&&b.transitioning))){var f=a.Event("show.bs.collapse");if(this.$element.trigger(f),!f.isDefaultPrevented()){e&&e.length&&(c.call(e,"hide"),b||e.data("bs.collapse",null));var g=this.dimension();this.$element.removeClass("collapse").addClass("collapsing")[g](0).attr("aria-expanded",!0),this.$trigger.removeClass("collapsed").attr("aria-expanded",!0),this.transitioning=1;var h=function(){this.$element.removeClass("collapsing").addClass("collapse in")[g](""),this.transitioning=0,this.$element.trigger("shown.bs.collapse")};if(!a.support.transition)return h.call(this);var i=a.camelCase(["scroll",g].join("-"));this.$element.one("bsTransitionEnd",a.proxy(h,this)).emulateTransitionEnd(d.TRANSITION_DURATION)[g](this.$element[0][i])}}}},d.prototype.hide=function(){if(!this.transitioning&&this.$element.hasClass("in")){var b=a.Event("hide.bs.collapse");if(this.$element.trigger(b),!b.isDefaultPrevented()){var c=this.dimension();this.$element[c](this.$element[c]())[0].offsetHeight,this.$element.addClass("collapsing").removeClass("collapse in").attr("aria-expanded",!1),this.$trigger.addClass("collapsed").attr("aria-expanded",!1),this.transitioning=1;var e=function(){this.transitioning=0,this.$element.removeClass("collapsing").addClass("collapse").trigger("hidden.bs.collapse")};return a.support.transition?void this.$element[c](0).one("bsTransitionEnd",a.proxy(e,this)).emulateTransitionEnd(d.TRANSITION_DURATION):e.call(this)}}},d.prototype.toggle=function(){this[this.$element.hasClass("in")?"hide":"show"]()},d.prototype.getParent=function(){return a(this.options.parent).find('[data-toggle="collapse"][data-parent="'+this.options.parent+'"]').each(a.proxy(function(c,d){var e=a(d);this.addAriaAndCollapsedClass(b(e),e)},this)).end()},d.prototype.addAriaAndCollapsedClass=function(a,b){var c=a.hasClass("in");a.attr("aria-expanded",c),b.toggleClass("collapsed",!c).attr("aria-expanded",c)};var e=a.fn.collapse;a.fn.collapse=c,a.fn.collapse.Constructor=d,a.fn.collapse.noConflict=function(){return a.fn.collapse=e,this},a(document).on("click.bs.collapse.data-api",'[data-toggle="collapse"]',function(d){var e=a(this);e.attr("data-target")||d.preventDefault();var f=b(e),g=f.data("bs.collapse"),h=g?"toggle":e.data();c.call(f,h)})}(jQuery),+function(a){"use strict";function b(b){var c=b.attr("data-target");c||(c=b.attr("href"),c=c&&/#[A-Za-z]/.test(c)&&c.replace(/.*(?=#[^\s]*$)/,""));var d=c&&a(c);return d&&d.length?d:b.parent()}function c(c){c&&3===c.which||(a(e).remove(),a(f).each(function(){var d=a(this),e=b(d),f={relatedTarget:this};e.hasClass("open")&&(c&&"click"==c.type&&/input|textarea/i.test(c.target.tagName)&&a.contains(e[0],c.target)||(e.trigger(c=a.Event("hide.bs.dropdown",f)),c.isDefaultPrevented()||(d.attr("aria-expanded","false"),e.removeClass("open").trigger("hidden.bs.dropdown",f))))}))}function d(b){return this.each(function(){var c=a(this),d=c.data("bs.dropdown");d||c.data("bs.dropdown",d=new g(this)),"string"==typeof b&&d[b].call(c)})}var e=".dropdown-backdrop",f='[data-toggle="dropdown"]',g=function(b){a(b).on("click.bs.dropdown",this.toggle)};g.VERSION="3.3.5",g.prototype.toggle=function(d){var e=a(this);if(!e.is(".disabled, :disabled")){var f=b(e),g=f.hasClass("open");if(c(),!g){"ontouchstart"in document.documentElement&&!f.closest(".navbar-nav").length&&a(document.createElement("div")).addClass("dropdown-backdrop").insertAfter(a(this)).on("click",c);var h={relatedTarget:this};if(f.trigger(d=a.Event("show.bs.dropdown",h)),d.isDefaultPrevented())return;e.trigger("focus").attr("aria-expanded","true"),f.toggleClass("open").trigger("shown.bs.dropdown",h)}return!1}},g.prototype.keydown=function(c){if(/(38|40|27|32)/.test(c.which)&&!/input|textarea/i.test(c.target.tagName)){var d=a(this);if(c.preventDefault(),c.stopPropagation(),!d.is(".disabled, :disabled")){var e=b(d),g=e.hasClass("open");if(!g&&27!=c.which||g&&27==c.which)return 27==c.which&&e.find(f).trigger("focus"),d.trigger("click");var h=" li:not(.disabled):visible a",i=e.find(".dropdown-menu"+h);if(i.length){var j=i.index(c.target);38==c.which&&j>0&&j--,40==c.which&&j<i.length-1&&j++,~j||(j=0),i.eq(j).trigger("focus")}}}};var h=a.fn.dropdown;a.fn.dropdown=d,a.fn.dropdown.Constructor=g,a.fn.dropdown.noConflict=function(){return a.fn.dropdown=h,this},a(document).on("click.bs.dropdown.data-api",c).on("click.bs.dropdown.data-api",".dropdown form",function(a){a.stopPropagation()}).on("click.bs.dropdown.data-api",f,g.prototype.toggle).on("keydown.bs.dropdown.data-api",f,g.prototype.keydown).on("keydown.bs.dropdown.data-api",".dropdown-menu",g.prototype.keydown)}(jQuery),+function(a){"use strict";function b(b,d){return this.each(function(){var e=a(this),f=e.data("bs.modal"),g=a.extend({},c.DEFAULTS,e.data(),"object"==typeof b&&b);f||e.data("bs.modal",f=new c(this,g)),"string"==typeof b?f[b](d):g.show&&f.show(d)})}var c=function(b,c){this.options=c,this.$body=a(document.body),this.$element=a(b),this.$dialog=this.$element.find(".modal-dialog"),this.$backdrop=null,this.isShown=null,this.originalBodyPad=null,this.scrollbarWidth=0,this.ignoreBackdropClick=!1,this.options.remote&&this.$element.find(".modal-content").load(this.options.remote,a.proxy(function(){this.$element.trigger("loaded.bs.modal")},this))};c.VERSION="3.3.5",c.TRANSITION_DURATION=300,c.BACKDROP_TRANSITION_DURATION=150,c.DEFAULTS={backdrop:!0,keyboard:!0,show:!0},c.prototype.toggle=function(a){return this.isShown?this.hide():this.show(a)},c.prototype.show=function(b){var d=this,e=a.Event("show.bs.modal",{relatedTarget:b});this.$element.trigger(e),this.isShown||e.isDefaultPrevented()||(this.isShown=!0,this.checkScrollbar(),this.setScrollbar(),this.$body.addClass("modal-open"),this.escape(),this.resize(),this.$element.on("click.dismiss.bs.modal",'[data-dismiss="modal"]',a.proxy(this.hide,this)),this.$dialog.on("mousedown.dismiss.bs.modal",function(){d.$element.one("mouseup.dismiss.bs.modal",function(b){a(b.target).is(d.$element)&&(d.ignoreBackdropClick=!0)})}),this.backdrop(function(){var e=a.support.transition&&d.$element.hasClass("fade");d.$element.parent().length||d.$element.appendTo(d.$body),d.$element.show().scrollTop(0),d.adjustDialog(),e&&d.$element[0].offsetWidth,d.$element.addClass("in"),d.enforceFocus();var f=a.Event("shown.bs.modal",{relatedTarget:b});e?d.$dialog.one("bsTransitionEnd",function(){d.$element.trigger("focus").trigger(f)}).emulateTransitionEnd(c.TRANSITION_DURATION):d.$element.trigger("focus").trigger(f)}))},c.prototype.hide=function(b){b&&b.preventDefault(),b=a.Event("hide.bs.modal"),this.$element.trigger(b),this.isShown&&!b.isDefaultPrevented()&&(this.isShown=!1,this.escape(),this.resize(),a(document).off("focusin.bs.modal"),this.$element.removeClass("in").off("click.dismiss.bs.modal").off("mouseup.dismiss.bs.modal"),this.$dialog.off("mousedown.dismiss.bs.modal"),a.support.transition&&this.$element.hasClass("fade")?this.$element.one("bsTransitionEnd",a.proxy(this.hideModal,this)).emulateTransitionEnd(c.TRANSITION_DURATION):this.hideModal())},c.prototype.enforceFocus=function(){a(document).off("focusin.bs.modal").on("focusin.bs.modal",a.proxy(function(a){this.$element[0]===a.target||this.$element.has(a.target).length||this.$element.trigger("focus")},this))},c.prototype.escape=function(){this.isShown&&this.options.keyboard?this.$element.on("keydown.dismiss.bs.modal",a.proxy(function(a){27==a.which&&this.hide()},this)):this.isShown||this.$element.off("keydown.dismiss.bs.modal")},c.prototype.resize=function(){this.isShown?a(window).on("resize.bs.modal",a.proxy(this.handleUpdate,this)):a(window).off("resize.bs.modal")},c.prototype.hideModal=function(){var a=this;this.$element.hide(),this.backdrop(function(){a.$body.removeClass("modal-open"),a.resetAdjustments(),a.resetScrollbar(),a.$element.trigger("hidden.bs.modal")})},c.prototype.removeBackdrop=function(){this.$backdrop&&this.$backdrop.remove(),this.$backdrop=null},c.prototype.backdrop=function(b){var d=this,e=this.$element.hasClass("fade")?"fade":"";if(this.isShown&&this.options.backdrop){var f=a.support.transition&&e;if(this.$backdrop=a(document.createElement("div")).addClass("modal-backdrop "+e).appendTo(this.$body),this.$element.on("click.dismiss.bs.modal",a.proxy(function(a){return this.ignoreBackdropClick?void(this.ignoreBackdropClick=!1):void(a.target===a.currentTarget&&("static"==this.options.backdrop?this.$element[0].focus():this.hide()))},this)),f&&this.$backdrop[0].offsetWidth,this.$backdrop.addClass("in"),!b)return;f?this.$backdrop.one("bsTransitionEnd",b).emulateTransitionEnd(c.BACKDROP_TRANSITION_DURATION):b()}else if(!this.isShown&&this.$backdrop){this.$backdrop.removeClass("in");var g=function(){d.removeBackdrop(),b&&b()};a.support.transition&&this.$element.hasClass("fade")?this.$backdrop.one("bsTransitionEnd",g).emulateTransitionEnd(c.BACKDROP_TRANSITION_DURATION):g()}else b&&b()},c.prototype.handleUpdate=function(){this.adjustDialog()},c.prototype.adjustDialog=function(){var a=this.$element[0].scrollHeight>document.documentElement.clientHeight;this.$element.css({paddingLeft:!this.bodyIsOverflowing&&a?this.scrollbarWidth:"",paddingRight:this.bodyIsOverflowing&&!a?this.scrollbarWidth:""})},c.prototype.resetAdjustments=function(){this.$element.css({paddingLeft:"",paddingRight:""})},c.prototype.checkScrollbar=function(){var a=window.innerWidth;if(!a){var b=document.documentElement.getBoundingClientRect();a=b.right-Math.abs(b.left)}this.bodyIsOverflowing=document.body.clientWidth<a,this.scrollbarWidth=this.measureScrollbar()},c.prototype.setScrollbar=function(){var a=parseInt(this.$body.css("padding-right")||0,10);this.originalBodyPad=document.body.style.paddingRight||"",this.bodyIsOverflowing&&this.$body.css("padding-right",a+this.scrollbarWidth)},c.prototype.resetScrollbar=function(){this.$body.css("padding-right",this.originalBodyPad)},c.prototype.measureScrollbar=function(){var a=document.createElement("div");a.className="modal-scrollbar-measure",this.$body.append(a);var b=a.offsetWidth-a.clientWidth;return this.$body[0].removeChild(a),b};var d=a.fn.modal;a.fn.modal=b,a.fn.modal.Constructor=c,a.fn.modal.noConflict=function(){return a.fn.modal=d,this},a(document).on("click.bs.modal.data-api",'[data-toggle="modal"]',function(c){var d=a(this),e=d.attr("href"),f=a(d.attr("data-target")||e&&e.replace(/.*(?=#[^\s]+$)/,"")),g=f.data("bs.modal")?"toggle":a.extend({remote:!/#/.test(e)&&e},f.data(),d.data());d.is("a")&&c.preventDefault(),f.one("show.bs.modal",function(a){a.isDefaultPrevented()||f.one("hidden.bs.modal",function(){d.is(":visible")&&d.trigger("focus")})}),b.call(f,g,this)})}(jQuery),+function(a){"use strict";function b(b){return this.each(function(){var d=a(this),e=d.data("bs.tooltip"),f="object"==typeof b&&b;(e||!/destroy|hide/.test(b))&&(e||d.data("bs.tooltip",e=new c(this,f)),"string"==typeof b&&e[b]())})}var c=function(a,b){this.type=null,this.options=null,this.enabled=null,this.timeout=null,this.hoverState=null,this.$element=null,this.inState=null,this.init("tooltip",a,b)};c.VERSION="3.3.5",c.TRANSITION_DURATION=150,c.DEFAULTS={animation:!0,placement:"top",selector:!1,template:'<div class="tooltip" role="tooltip"><div class="tooltip-arrow"></div><div class="tooltip-inner"></div></div>',trigger:"hover focus",title:"",delay:0,html:!1,container:!1,viewport:{selector:"body",padding:0}},c.prototype.init=function(b,c,d){if(this.enabled=!0,this.type=b,this.$element=a(c),this.options=this.getOptions(d),this.$viewport=this.options.viewport&&a(a.isFunction(this.options.viewport)?this.options.viewport.call(this,this.$element):this.options.viewport.selector||this.options.viewport),this.inState={click:!1,hover:!1,focus:!1},this.$element[0]instanceof document.constructor&&!this.options.selector)throw new Error("`selector` option must be specified when initializing "+this.type+" on the window.document object!");for(var e=this.options.trigger.split(" "),f=e.length;f--;){var g=e[f];if("click"==g)this.$element.on("click."+this.type,this.options.selector,a.proxy(this.toggle,this));else if("manual"!=g){var h="hover"==g?"mouseenter":"focusin",i="hover"==g?"mouseleave":"focusout";this.$element.on(h+"."+this.type,this.options.selector,a.proxy(this.enter,this)),this.$element.on(i+"."+this.type,this.options.selector,a.proxy(this.leave,this))}}this.options.selector?this._options=a.extend({},this.options,{trigger:"manual",selector:""}):this.fixTitle()},c.prototype.getDefaults=function(){return c.DEFAULTS},c.prototype.getOptions=function(b){return b=a.extend({},this.getDefaults(),this.$element.data(),b),b.delay&&"number"==typeof b.delay&&(b.delay={show:b.delay,hide:b.delay}),b},c.prototype.getDelegateOptions=function(){var b={},c=this.getDefaults();return this._options&&a.each(this._options,function(a,d){c[a]!=d&&(b[a]=d)}),b},c.prototype.enter=function(b){var c=b instanceof this.constructor?b:a(b.currentTarget).data("bs."+this.type);return c||(c=new this.constructor(b.currentTarget,this.getDelegateOptions()),a(b.currentTarget).data("bs."+this.type,c)),b instanceof a.Event&&(c.inState["focusin"==b.type?"focus":"hover"]=!0),c.tip().hasClass("in")||"in"==c.hoverState?void(c.hoverState="in"):(clearTimeout(c.timeout),c.hoverState="in",c.options.delay&&c.options.delay.show?void(c.timeout=setTimeout(function(){"in"==c.hoverState&&c.show()},c.options.delay.show)):c.show())},c.prototype.isInStateTrue=function(){for(var a in this.inState)if(this.inState[a])return!0;return!1},c.prototype.leave=function(b){var c=b instanceof this.constructor?b:a(b.currentTarget).data("bs."+this.type);return c||(c=new this.constructor(b.currentTarget,this.getDelegateOptions()),a(b.currentTarget).data("bs."+this.type,c)),b instanceof a.Event&&(c.inState["focusout"==b.type?"focus":"hover"]=!1),c.isInStateTrue()?void 0:(clearTimeout(c.timeout),c.hoverState="out",c.options.delay&&c.options.delay.hide?void(c.timeout=setTimeout(function(){"out"==c.hoverState&&c.hide()},c.options.delay.hide)):c.hide())},c.prototype.show=function(){var b=a.Event("show.bs."+this.type);if(this.hasContent()&&this.enabled){this.$element.trigger(b);var d=a.contains(this.$element[0].ownerDocument.documentElement,this.$element[0]);if(b.isDefaultPrevented()||!d)return;var e=this,f=this.tip(),g=this.getUID(this.type);this.setContent(),f.attr("id",g),this.$element.attr("aria-describedby",g),this.options.animation&&f.addClass("fade");var h="function"==typeof this.options.placement?this.options.placement.call(this,f[0],this.$element[0]):this.options.placement,i=/\s?auto?\s?/i,j=i.test(h);j&&(h=h.replace(i,"")||"top"),f.detach().css({top:0,left:0,display:"block"}).addClass(h).data("bs."+this.type,this),this.options.container?f.appendTo(this.options.container):f.insertAfter(this.$element),this.$element.trigger("inserted.bs."+this.type);var k=this.getPosition(),l=f[0].offsetWidth,m=f[0].offsetHeight;if(j){var n=h,o=this.getPosition(this.$viewport);h="bottom"==h&&k.bottom+m>o.bottom?"top":"top"==h&&k.top-m<o.top?"bottom":"right"==h&&k.right+l>o.width?"left":"left"==h&&k.left-l<o.left?"right":h,f.removeClass(n).addClass(h)}var p=this.getCalculatedOffset(h,k,l,m);this.applyPlacement(p,h);var q=function(){var a=e.hoverState;e.$element.trigger("shown.bs."+e.type),e.hoverState=null,"out"==a&&e.leave(e)};a.support.transition&&this.$tip.hasClass("fade")?f.one("bsTransitionEnd",q).emulateTransitionEnd(c.TRANSITION_DURATION):q()}},c.prototype.applyPlacement=function(b,c){var d=this.tip(),e=d[0].offsetWidth,f=d[0].offsetHeight,g=parseInt(d.css("margin-top"),10),h=parseInt(d.css("margin-left"),10);isNaN(g)&&(g=0),isNaN(h)&&(h=0),b.top+=g,b.left+=h,a.offset.setOffset(d[0],a.extend({using:function(a){d.css({top:Math.round(a.top),left:Math.round(a.left)})}},b),0),d.addClass("in");var i=d[0].offsetWidth,j=d[0].offsetHeight;"top"==c&&j!=f&&(b.top=b.top+f-j);var k=this.getViewportAdjustedDelta(c,b,i,j);k.left?b.left+=k.left:b.top+=k.top;var l=/top|bottom/.test(c),m=l?2*k.left-e+i:2*k.top-f+j,n=l?"offsetWidth":"offsetHeight";d.offset(b),this.replaceArrow(m,d[0][n],l)},c.prototype.replaceArrow=function(a,b,c){this.arrow().css(c?"left":"top",50*(1-a/b)+"%").css(c?"top":"left","")},c.prototype.setContent=function(){var a=this.tip(),b=this.getTitle();a.find(".tooltip-inner")[this.options.html?"html":"text"](b),a.removeClass("fade in top bottom left right")},c.prototype.hide=function(b){function d(){"in"!=e.hoverState&&f.detach(),e.$element.removeAttr("aria-describedby").trigger("hidden.bs."+e.type),b&&b()}var e=this,f=a(this.$tip),g=a.Event("hide.bs."+this.type);return this.$element.trigger(g),g.isDefaultPrevented()?void 0:(f.removeClass("in"),a.support.transition&&f.hasClass("fade")?f.one("bsTransitionEnd",d).emulateTransitionEnd(c.TRANSITION_DURATION):d(),this.hoverState=null,this)},c.prototype.fixTitle=function(){var a=this.$element;(a.attr("title")||"string"!=typeof a.attr("data-original-title"))&&a.attr("data-original-title",a.attr("title")||"").attr("title","")},c.prototype.hasContent=function(){return this.getTitle()},c.prototype.getPosition=function(b){b=b||this.$element;var c=b[0],d="BODY"==c.tagName,e=c.getBoundingClientRect();null==e.width&&(e=a.extend({},e,{width:e.right-e.left,height:e.bottom-e.top}));var f=d?{top:0,left:0}:b.offset(),g={scroll:d?document.documentElement.scrollTop||document.body.scrollTop:b.scrollTop()},h=d?{width:a(window).width(),height:a(window).height()}:null;return a.extend({},e,g,h,f)},c.prototype.getCalculatedOffset=function(a,b,c,d){return"bottom"==a?{top:b.top+b.height,left:b.left+b.width/2-c/2}:"top"==a?{top:b.top-d,left:b.left+b.width/2-c/2}:"left"==a?{top:b.top+b.height/2-d/2,left:b.left-c}:{top:b.top+b.height/2-d/2,left:b.left+b.width}},c.prototype.getViewportAdjustedDelta=function(a,b,c,d){var e={top:0,left:0};if(!this.$viewport)return e;var f=this.options.viewport&&this.options.viewport.padding||0,g=this.getPosition(this.$viewport);if(/right|left/.test(a)){var h=b.top-f-g.scroll,i=b.top+f-g.scroll+d;h<g.top?e.top=g.top-h:i>g.top+g.height&&(e.top=g.top+g.height-i)}else{var j=b.left-f,k=b.left+f+c;j<g.left?e.left=g.left-j:k>g.right&&(e.left=g.left+g.width-k)}return e},c.prototype.getTitle=function(){var a,b=this.$element,c=this.options;return a=b.attr("data-original-title")||("function"==typeof c.title?c.title.call(b[0]):c.title)},c.prototype.getUID=function(a){do a+=~~(1e6*Math.random());while(document.getElementById(a));return a},c.prototype.tip=function(){if(!this.$tip&&(this.$tip=a(this.options.template),1!=this.$tip.length))throw new Error(this.type+" `template` option must consist of exactly 1 top-level element!");return this.$tip},c.prototype.arrow=function(){return this.$arrow=this.$arrow||this.tip().find(".tooltip-arrow")},c.prototype.enable=function(){this.enabled=!0},c.prototype.disable=function(){this.enabled=!1},c.prototype.toggleEnabled=function(){this.enabled=!this.enabled},c.prototype.toggle=function(b){var c=this;b&&(c=a(b.currentTarget).data("bs."+this.type),c||(c=new this.constructor(b.currentTarget,this.getDelegateOptions()),a(b.currentTarget).data("bs."+this.type,c))),b?(c.inState.click=!c.inState.click,c.isInStateTrue()?c.enter(c):c.leave(c)):c.tip().hasClass("in")?c.leave(c):c.enter(c)},c.prototype.destroy=function(){var a=this;clearTimeout(this.timeout),this.hide(function(){a.$element.off("."+a.type).removeData("bs."+a.type),a.$tip&&a.$tip.detach(),a.$tip=null,a.$arrow=null,a.$viewport=null})};var d=a.fn.tooltip;a.fn.tooltip=b,a.fn.tooltip.Constructor=c,a.fn.tooltip.noConflict=function(){return a.fn.tooltip=d,this}}(jQuery),+function(a){"use strict";function b(b){return this.each(function(){var d=a(this),e=d.data("bs.popover"),f="object"==typeof b&&b;(e||!/destroy|hide/.test(b))&&(e||d.data("bs.popover",e=new c(this,f)),"string"==typeof b&&e[b]())})}var c=function(a,b){this.init("popover",a,b)};if(!a.fn.tooltip)throw new Error("Popover requires tooltip.js");c.VERSION="3.3.5",c.DEFAULTS=a.extend({},a.fn.tooltip.Constructor.DEFAULTS,{placement:"right",trigger:"click",content:"",template:'<div class="popover" role="tooltip"><div class="arrow"></div><h3 class="popover-title"></h3><div class="popover-content"></div></div>'}),c.prototype=a.extend({},a.fn.tooltip.Constructor.prototype),c.prototype.constructor=c,c.prototype.getDefaults=function(){return c.DEFAULTS},c.prototype.setContent=function(){var a=this.tip(),b=this.getTitle(),c=this.getContent();a.find(".popover-title")[this.options.html?"html":"text"](b),a.find(".popover-content").children().detach().end()[this.options.html?"string"==typeof c?"html":"append":"text"](c),a.removeClass("fade top bottom left right in"),a.find(".popover-title").html()||a.find(".popover-title").hide()},c.prototype.hasContent=function(){return this.getTitle()||this.getContent()},c.prototype.getContent=function(){var a=this.$element,b=this.options;return a.attr("data-content")||("function"==typeof b.content?b.content.call(a[0]):b.content)},c.prototype.arrow=function(){return this.$arrow=this.$arrow||this.tip().find(".arrow")};var d=a.fn.popover;a.fn.popover=b,a.fn.popover.Constructor=c,a.fn.popover.noConflict=function(){return a.fn.popover=d,this}}(jQuery),+function(a){"use strict";function b(c,d){this.$body=a(document.body),this.$scrollElement=a(a(c).is(document.body)?window:c),this.options=a.extend({},b.DEFAULTS,d),this.selector=(this.options.target||"")+" .nav li > a",this.offsets=[],this.targets=[],this.activeTarget=null,this.scrollHeight=0,this.$scrollElement.on("scroll.bs.scrollspy",a.proxy(this.process,this)),this.refresh(),this.process()}function c(c){return this.each(function(){var d=a(this),e=d.data("bs.scrollspy"),f="object"==typeof c&&c;e||d.data("bs.scrollspy",e=new b(this,f)),"string"==typeof c&&e[c]()})}b.VERSION="3.3.5",b.DEFAULTS={offset:10},b.prototype.getScrollHeight=function(){return this.$scrollElement[0].scrollHeight||Math.max(this.$body[0].scrollHeight,document.documentElement.scrollHeight)},b.prototype.refresh=function(){var b=this,c="offset",d=0;this.offsets=[],this.targets=[],this.scrollHeight=this.getScrollHeight(),a.isWindow(this.$scrollElement[0])||(c="position",d=this.$scrollElement.scrollTop()),this.$body.find(this.selector).map(function(){var b=a(this),e=b.data("target")||b.attr("href"),f=/^#./.test(e)&&a(e);return f&&f.length&&f.is(":visible")&&[[f[c]().top+d,e]]||null}).sort(function(a,b){return a[0]-b[0]}).each(function(){b.offsets.push(this[0]),b.targets.push(this[1])})},b.prototype.process=function(){var a,b=this.$scrollElement.scrollTop()+this.options.offset,c=this.getScrollHeight(),d=this.options.offset+c-this.$scrollElement.height(),e=this.offsets,f=this.targets,g=this.activeTarget;if(this.scrollHeight!=c&&this.refresh(),b>=d)return g!=(a=f[f.length-1])&&this.activate(a);if(g&&b<e[0])return this.activeTarget=null,this.clear();for(a=e.length;a--;)g!=f[a]&&b>=e[a]&&(void 0===e[a+1]||b<e[a+1])&&this.activate(f[a])},b.prototype.activate=function(b){this.activeTarget=b,this.clear();var c=this.selector+'[data-target="'+b+'"],'+this.selector+'[href="'+b+'"]',d=a(c).parents("li").addClass("active");d.parent(".dropdown-menu").length&&(d=d.closest("li.dropdown").addClass("active")),
d.trigger("activate.bs.scrollspy")},b.prototype.clear=function(){a(this.selector).parentsUntil(this.options.target,".active").removeClass("active")};var d=a.fn.scrollspy;a.fn.scrollspy=c,a.fn.scrollspy.Constructor=b,a.fn.scrollspy.noConflict=function(){return a.fn.scrollspy=d,this},a(window).on("load.bs.scrollspy.data-api",function(){a('[data-spy="scroll"]').each(function(){var b=a(this);c.call(b,b.data())})})}(jQuery),+function(a){"use strict";function b(b){return this.each(function(){var d=a(this),e=d.data("bs.tab");e||d.data("bs.tab",e=new c(this)),"string"==typeof b&&e[b]()})}var c=function(b){this.element=a(b)};c.VERSION="3.3.5",c.TRANSITION_DURATION=150,c.prototype.show=function(){var b=this.element,c=b.closest("ul:not(.dropdown-menu)"),d=b.data("target");if(d||(d=b.attr("href"),d=d&&d.replace(/.*(?=#[^\s]*$)/,"")),!b.parent("li").hasClass("active")){var e=c.find(".active:last a"),f=a.Event("hide.bs.tab",{relatedTarget:b[0]}),g=a.Event("show.bs.tab",{relatedTarget:e[0]});if(e.trigger(f),b.trigger(g),!g.isDefaultPrevented()&&!f.isDefaultPrevented()){var h=a(d);this.activate(b.closest("li"),c),this.activate(h,h.parent(),function(){e.trigger({type:"hidden.bs.tab",relatedTarget:b[0]}),b.trigger({type:"shown.bs.tab",relatedTarget:e[0]})})}}},c.prototype.activate=function(b,d,e){function f(){g.removeClass("active").find("> .dropdown-menu > .active").removeClass("active").end().find('[data-toggle="tab"]').attr("aria-expanded",!1),b.addClass("active").find('[data-toggle="tab"]').attr("aria-expanded",!0),h?(b[0].offsetWidth,b.addClass("in")):b.removeClass("fade"),b.parent(".dropdown-menu").length&&b.closest("li.dropdown").addClass("active").end().find('[data-toggle="tab"]').attr("aria-expanded",!0),e&&e()}var g=d.find("> .active"),h=e&&a.support.transition&&(g.length&&g.hasClass("fade")||!!d.find("> .fade").length);g.length&&h?g.one("bsTransitionEnd",f).emulateTransitionEnd(c.TRANSITION_DURATION):f(),g.removeClass("in")};var d=a.fn.tab;a.fn.tab=b,a.fn.tab.Constructor=c,a.fn.tab.noConflict=function(){return a.fn.tab=d,this};var e=function(c){c.preventDefault(),b.call(a(this),"show")};a(document).on("click.bs.tab.data-api",'[data-toggle="tab"]',e).on("click.bs.tab.data-api",'[data-toggle="pill"]',e)}(jQuery),+function(a){"use strict";function b(b){return this.each(function(){var d=a(this),e=d.data("bs.affix"),f="object"==typeof b&&b;e||d.data("bs.affix",e=new c(this,f)),"string"==typeof b&&e[b]()})}var c=function(b,d){this.options=a.extend({},c.DEFAULTS,d),this.$target=a(this.options.target).on("scroll.bs.affix.data-api",a.proxy(this.checkPosition,this)).on("click.bs.affix.data-api",a.proxy(this.checkPositionWithEventLoop,this)),this.$element=a(b),this.affixed=null,this.unpin=null,this.pinnedOffset=null,this.checkPosition()};c.VERSION="3.3.5",c.RESET="affix affix-top affix-bottom",c.DEFAULTS={offset:0,target:window},c.prototype.getState=function(a,b,c,d){var e=this.$target.scrollTop(),f=this.$element.offset(),g=this.$target.height();if(null!=c&&"top"==this.affixed)return c>e?"top":!1;if("bottom"==this.affixed)return null!=c?e+this.unpin<=f.top?!1:"bottom":a-d>=e+g?!1:"bottom";var h=null==this.affixed,i=h?e:f.top,j=h?g:b;return null!=c&&c>=e?"top":null!=d&&i+j>=a-d?"bottom":!1},c.prototype.getPinnedOffset=function(){if(this.pinnedOffset)return this.pinnedOffset;this.$element.removeClass(c.RESET).addClass("affix");var a=this.$target.scrollTop(),b=this.$element.offset();return this.pinnedOffset=b.top-a},c.prototype.checkPositionWithEventLoop=function(){setTimeout(a.proxy(this.checkPosition,this),1)},c.prototype.checkPosition=function(){if(this.$element.is(":visible")){var b=this.$element.height(),d=this.options.offset,e=d.top,f=d.bottom,g=Math.max(a(document).height(),a(document.body).height());"object"!=typeof d&&(f=e=d),"function"==typeof e&&(e=d.top(this.$element)),"function"==typeof f&&(f=d.bottom(this.$element));var h=this.getState(g,b,e,f);if(this.affixed!=h){null!=this.unpin&&this.$element.css("top","");var i="affix"+(h?"-"+h:""),j=a.Event(i+".bs.affix");if(this.$element.trigger(j),j.isDefaultPrevented())return;this.affixed=h,this.unpin="bottom"==h?this.getPinnedOffset():null,this.$element.removeClass(c.RESET).addClass(i).trigger(i.replace("affix","affixed")+".bs.affix")}"bottom"==h&&this.$element.offset({top:g-b-f})}};var d=a.fn.affix;a.fn.affix=b,a.fn.affix.Constructor=c,a.fn.affix.noConflict=function(){return a.fn.affix=d,this},a(window).on("load",function(){a('[data-spy="affix"]').each(function(){var c=a(this),d=c.data();d.offset=d.offset||{},null!=d.offsetBottom&&(d.offset.bottom=d.offsetBottom),null!=d.offsetTop&&(d.offset.top=d.offsetTop),b.call(c,d)})})}(jQuery);
```
