## Supplementary material for "Characterizing and inferring quantitative cell cycle phase in single-cell RNA-seq data analysis": Source code of our analysis, including scripts and data necessary to reproduce the work.: npm.js

```
// This file is autogenerated via the `commonjs` Grunt task. You can require() this file in a CommonJS environment.
require('../../js/transition.js')
require('../../js/alert.js')
require('../../js/button.js')
require('../../js/carousel.js')
require('../../js/collapse.js')
require('../../js/dropdown.js')
require('../../js/modal.js')
require('../../js/tooltip.js')
require('../../js/popover.js')
require('../../js/scrollspy.js')
require('../../js/tab.js')
require('../../js/affix.js')
```
