## Supplementary material for "Characterizing and inferring quantitative cell cycle phase in single-cell RNA-seq data analysis": Source code of our analysis, including scripts and data necessary to reproduce the work.: html5shiv.min.js

```
/**
* @preserve HTML5 Shiv 3.7.2 | @afarkas @jdalton @jon_neal @rem | MIT/GPL2 Licensed
*/
// Only run this code in IE 8
if (!!window.navigator.userAgent.match("MSIE 8")) {
!function(a,b){function c(a,b){var c=a.createElement("p"),d=a.getElementsByTagName("head")[0]||a.documentElement;return c.innerHTML="x<style>"+b+"</style>",d.insertBefore(c.lastChild,d.firstChild)}function d(){var a=t.elements;return"string"==typeof a?a.split(" "):a}function e(a,b){var c=t.elements;"string"!=typeof c&&(c=c.join(" ")),"string"!=typeof a&&(a=a.join(" ")),t.elements=c+" "+a,j(b)}function f(a){var b=s[a[q]];return b||(b={},r++,a[q]=r,s[r]=b),b}function g(a,c,d){if(c||(c=b),l)return c.createElement(a);d||(d=f(c));var e;return e=d.cache[a]?d.cache[a].cloneNode():p.test(a)?(d.cache[a]=d.createElem(a)).cloneNode():d.createElem(a),!e.canHaveChildren||o.test(a)||e.tagUrn?e:d.frag.appendChild(e)}function h(a,c){if(a||(a=b),l)return a.createDocumentFragment();c=c||f(a);for(var e=c.frag.cloneNode(),g=0,h=d(),i=h.length;i>g;g++)e.createElement(h[g]);return e}function i(a,b){b.cache||(b.cache={},b.createElem=a.createElement,b.createFrag=a.createDocumentFragment,b.frag=b.createFrag()),a.createElement=function(c){return t.shivMethods?g(c,a,b):b.createElem(c)},a.createDocumentFragment=Function("h,f","return function(){var n=f.cloneNode(),c=n.createElement;h.shivMethods&&("+d().join().replace(/[\w\-:]+/g,function(a){return b.createElem(a),b.frag.createElement(a),'c("'+a+'")'})+");return n}")(t,b.frag)}function j(a){a||(a=b);var d=f(a);return!t.shivCSS||k||d.hasCSS||(d.hasCSS=!!c(a,"article,aside,dialog,figcaption,figure,footer,header,hgroup,main,nav,section{display:block}mark{background:#FF0;color:#000}template{display:none}")),l||i(a,d),a}var k,l,m="3.7.2",n=a.html5||{},o=/^<|^(?:button|map|select|textarea|object|iframe|option|optgroup)$/i,p=/^(?:a|b|code|div|fieldset|h1|h2|h3|h4|h5|h6|i|label|li|ol|p|q|span|strong|style|table|tbody|td|th|tr|ul)$/i,q="_html5shiv",r=0,s={};!function(){try{var a=b.createElement("a");a.innerHTML="<xyz></xyz>",k="hidden"in a,l=1==a.childNodes.length||function(){b.createElement("a");var a=b.createDocumentFragment();return"undefined"==typeof a.cloneNode||"undefined"==typeof a.createDocumentFragment||"undefined"==typeof a.createElement}()}catch(c){k=!0,l=!0}}();var t={elements:n.elements||"abbr article aside audio bdi canvas data datalist details dialog figcaption figure footer header hgroup main mark meter nav output picture progress section summary template time video",version:m,shivCSS:n.shivCSS!==!1,supportsUnknownElements:l,shivMethods:n.shivMethods!==!1,type:"default",shivDocument:j,createElement:g,createDocumentFragment:h,addElements:e};a.html5=t,j(b)}(this,document);
};
```
