## Supplementary material for "Characterizing and inferring quantitative cell cycle phase in single-cell RNA-seq data analysis": Source code of our analysis, including scripts and data necessary to reproduce the work.: respond.min.js

```
/*! Respond.js v1.4.2: min/max-width media query polyfill * Copyright 2013 Scott Jehl
 * Licensed under https://github.com/scottjehl/Respond/blob/master/LICENSE-MIT
 *  */

// Only run this code in IE 8
if (!!window.navigator.userAgent.match("MSIE 8")) {
!function(a){"use strict";a.matchMedia=a.matchMedia||function(a){var b,c=a.documentElement,d=c.firstElementChild||c.firstChild,e=a.createElement("body"),f=a.createElement("div");return f.id="mq-test-1",f.style.cssText="position:absolute;top:-100em",e.style.background="none",e.appendChild(f),function(a){return f.innerHTML='&shy;<style media="'+a+'"> #mq-test-1 { width: 42px; }</style>',c.insertBefore(e,d),b=42===f.offsetWidth,c.removeChild(e),{matches:b,media:a}}}(a.document)}(this),function(a){"use strict";function b(){u(!0)}var c={};a.respond=c,c.update=function(){};var d=[],e=function(){var b=!1;try{b=new a.XMLHttpRequest}catch(c){b=new a.ActiveXObject("Microsoft.XMLHTTP")}return function(){return b}}(),f=function(a,b){var c=e();c&&(c.open("GET",a,!0),c.onreadystatechange=function(){4!==c.readyState||200!==c.status&&304!==c.status||b(c.responseText)},4!==c.readyState&&c.send(null))};if(c.ajax=f,c.queue=d,c.regex={media:/@media[^\{]+\{([^\{\}]*\{[^\}\{]*\})+/gi,keyframes:/@(?:\-(?:o|moz|webkit)\-)?keyframes[^\{]+\{(?:[^\{\}]*\{[^\}\{]*\})+[^\}]*\}/gi,urls:/(url\()['"]?([^\/\)'"][^:\)'"]+)['"]?(\))/g,findStyles:/@media *([^\{]+)\{([\S\s]+?)$/,only:/(only\s+)?([a-zA-Z]+)\s?/,minw:/\([\s]*min\-width\s*:[\s]*([\s]*[0-9\.]+)(px|em)[\s]*\)/,maxw:/\([\s]*max\-width\s*:[\s]*([\s]*[0-9\.]+)(px|em)[\s]*\)/},c.mediaQueriesSupported=a.matchMedia&&null!==a.matchMedia("only all")&&a.matchMedia("only all").matches,!c.mediaQueriesSupported){var g,h,i,j=a.document,k=j.documentElement,l=[],m=[],n=[],o={},p=30,q=j.getElementsByTagName("head")[0]||k,r=j.getElementsByTagName("base")[0],s=q.getElementsByTagName("link"),t=function(){var a,b=j.createElement("div"),c=j.body,d=k.style.fontSize,e=c&&c.style.fontSize,f=!1;return b.style.cssText="position:absolute;font-size:1em;width:1em",c||(c=f=j.createElement("body"),c.style.background="none"),k.style.fontSize="100%",c.style.fontSize="100%",c.appendChild(b),f&&k.insertBefore(c,k.firstChild),a=b.offsetWidth,f?k.removeChild(c):c.removeChild(b),k.style.fontSize=d,e&&(c.style.fontSize=e),a=i=parseFloat(a)},u=function(b){var c="clientWidth",d=k[c],e="CSS1Compat"===j.compatMode&&d||j.body[c]||d,f={},o=s[s.length-1],r=(new Date).getTime();if(b&&g&&p>r-g)return a.clearTimeout(h),h=a.setTimeout(u,p),void 0;g=r;for(var v in l)if(l.hasOwnProperty(v)){var w=l[v],x=w.minw,y=w.maxw,z=null===x,A=null===y,B="em";x&&(x=parseFloat(x)*(x.indexOf(B)>-1?i||t():1)),y&&(y=parseFloat(y)*(y.indexOf(B)>-1?i||t():1)),w.hasquery&&(z&&A||!(z||e>=x)||!(A||y>=e))||(f[w.media]||(f[w.media]=[]),f[w.media].push(m[w.rules]))}for(var C in n)n.hasOwnProperty(C)&&n[C]&&n[C].parentNode===q&&q.removeChild(n[C]);n.length=0;for(var D in f)if(f.hasOwnProperty(D)){var E=j.createElement("style"),F=f[D].join("\n");E.type="text/css",E.media=D,q.insertBefore(E,o.nextSibling),E.styleSheet?E.styleSheet.cssText=F:E.appendChild(j.createTextNode(F)),n.push(E)}},v=function(a,b,d){var e=a.replace(c.regex.keyframes,"").match(c.regex.media),f=e&&e.length||0;b=b.substring(0,b.lastIndexOf("/"));var g=function(a){return a.replace(c.regex.urls,"$1"+b+"$2$3")},h=!f&&d;b.length&&(b+="/"),h&&(f=1);for(var i=0;f>i;i++){var j,k,n,o;h?(j=d,m.push(g(a))):(j=e[i].match(c.regex.findStyles)&&RegExp.$1,m.push(RegExp.$2&&g(RegExp.$2))),n=j.split(","),o=n.length;for(var p=0;o>p;p++)k=n[p],l.push({media:k.split("(")[0].match(c.regex.only)&&RegExp.$2||"all",rules:m.length-1,hasquery:k.indexOf("(")>-1,minw:k.match(c.regex.minw)&&parseFloat(RegExp.$1)+(RegExp.$2||""),maxw:k.match(c.regex.maxw)&&parseFloat(RegExp.$1)+(RegExp.$2||"")})}u()},w=function(){if(d.length){var b=d.shift();f(b.href,function(c){v(c,b.href,b.media),o[b.href]=!0,a.setTimeout(function(){w()},0)})}},x=function(){for(var b=0;b<s.length;b++){var c=s[b],e=c.href,f=c.media,g=c.rel&&"stylesheet"===c.rel.toLowerCase();e&&g&&!o[e]&&(c.styleSheet&&c.styleSheet.rawCssText?(v(c.styleSheet.rawCssText,e,f),o[e]=!0):(!/^([a-zA-Z:]*\/\/)/.test(e)&&!r||e.replace(RegExp.$1,"").split("/")[0]===a.location.host)&&("//"===e.substring(0,2)&&(e=a.location.protocol+e),d.push({href:e,media:f})))}w()};x(),c.update=x,c.getEmValue=t,a.addEventListener?a.addEventListener("resize",b,!1):a.attachEvent&&a.attachEvent("onresize",b)}}(this);
};
```
