## Supplementary material for "Characterizing and inferring quantitative cell cycle phase in single-cell RNA-seq data analysis": Source code of our analysis, including scripts and data necessary to reproduce the work.: crosstalk.js

```
(function e(t,n,r){function s(o,u){if(!n[o]){if(!t[o]){var a=typeof require=="function"&&require;if(!u&&a)return a(o,!0);if(i)return i(o,!0);var f=new Error("Cannot find module '"+o+"'");throw f.code="MODULE_NOT_FOUND",f}var l=n[o]={exports:{}};t[o][0].call(l.exports,function(e){var n=t[o][1][e];return s(n?n:e)},l,l.exports,e,t,n,r)}return n[o].exports}var i=typeof require=="function"&&require;for(var o=0;o<r.length;o++)s(r[o]);return s})({1:[function(require,module,exports){
"use strict";

Object.defineProperty(exports, "__esModule", {
  value: true
});

var _createClass = function () { function defineProperties(target, props) { for (var i = 0; i < props.length; i++) { var descriptor = props[i]; descriptor.enumerable = descriptor.enumerable || false; descriptor.configurable = true; if ("value" in descriptor) descriptor.writable = true; Object.defineProperty(target, descriptor.key, descriptor); } } return function (Constructor, protoProps, staticProps) { if (protoProps) defineProperties(Constructor.prototype, protoProps); if (staticProps) defineProperties(Constructor, staticProps); return Constructor; }; }();

function _classCallCheck(instance, Constructor) { if (!(instance instanceof Constructor)) { throw new TypeError("Cannot call a class as a function"); } }

var Events = function () {
  function Events() {
    _classCallCheck(this, Events);

    this._types = {};
    this._seq = 0;
  }

  _createClass(Events, [{
    key: "on",
    value: function on(eventType, listener) {
      var subs = this._types[eventType];
      if (!subs) {
        subs = this._types[eventType] = {};
      }
      var sub = "sub" + this._seq++;
      subs[sub] = listener;
      return sub;
    }

    // Returns false if no match, or string for sub name if matched

  }, {
    key: "off",
    value: function off(eventType, listener) {
      var subs = this._types[eventType];
      if (typeof listener === "function") {
        for (var key in subs) {
          if (subs.hasOwnProperty(key)) {
            if (subs[key] === listener) {
              delete subs[key];
              return key;
            }
          }
        }
        return false;
      } else if (typeof listener === "string") {
        if (subs && subs[listener]) {
          delete subs[listener];
          return listener;
        }
        return false;
      } else {
        throw new Error("Unexpected type for listener");
      }
    }
  }, {
    key: "trigger",
    value: function trigger(eventType, arg, thisObj) {
      var subs = this._types[eventType];
      for (var key in subs) {
        if (subs.hasOwnProperty(key)) {
          subs[key].call(thisObj, arg);
        }
      }
    }
  }]);

  return Events;
}();

exports.default = Events;

},{}],2:[function(require,module,exports){
"use strict";

Object.defineProperty(exports, "__esModule", {
  value: true
});
exports.FilterHandle = undefined;

var _createClass = function () { function defineProperties(target, props) { for (var i = 0; i < props.length; i++) { var descriptor = props[i]; descriptor.enumerable = descriptor.enumerable || false; descriptor.configurable = true; if ("value" in descriptor) descriptor.writable = true; Object.defineProperty(target, descriptor.key, descriptor); } } return function (Constructor, protoProps, staticProps) { if (protoProps) defineProperties(Constructor.prototype, protoProps); if (staticProps) defineProperties(Constructor, staticProps); return Constructor; }; }();

var _events = require("./events");

var _events2 = _interopRequireDefault(_events);

var _filterset = require("./filterset");

var _filterset2 = _interopRequireDefault(_filterset);

var _group = require("./group");

var _group2 = _interopRequireDefault(_group);

var _util = require("./util");

var util = _interopRequireWildcard(_util);

function _interopRequireWildcard(obj) { if (obj && obj.__esModule) { return obj; } else { var newObj = {}; if (obj != null) { for (var key in obj) { if (Object.prototype.hasOwnProperty.call(obj, key)) newObj[key] = obj[key]; } } newObj.default = obj; return newObj; } }

function _interopRequireDefault(obj) { return obj && obj.__esModule ? obj : { default: obj }; }

function _classCallCheck(instance, Constructor) { if (!(instance instanceof Constructor)) { throw new TypeError("Cannot call a class as a function"); } }

function getFilterSet(group) {
  var fsVar = group.var("filterset");
  var result = fsVar.get();
  if (!result) {
    result = new _filterset2.default();
    fsVar.set(result);
  }
  return result;
}

var id = 1;
function nextId() {
  return id++;
}

var FilterHandle = exports.FilterHandle = function () {
  /**
   * @classdesc
   * Use this class to contribute to, and listen for changes to, the filter set
   * for the given group of widgets. Filter input controls should create one
   * `FilterHandle` and only call {@link FilterHandle#set}. Output widgets that
   * wish to displayed filtered data should create one `FilterHandle` and use
   * the {@link FilterHandle#filteredKeys} property and listen for change
   * events.
   *
   * If two (or more) `FilterHandle` instances in the same webpage share the
   * same group name, they will contribute to a single "filter set". Each
   * `FilterHandle` starts out with a `null` value, which means they take
   * nothing away from the set of data that should be shown. To make a
   * `FilterHandle` actually remove data from the filter set, set its value to
   * an array of keys which should be displayed. Crosstalk will aggregate the
   * various key arrays by finding their intersection; only keys that are
   * present in all non-null filter handles are considered part of the filter
   * set.
   *
   * @param {string} [group] - The name of the Crosstalk group, or if none,
   *   null or undefined (or any other falsy value). This can be changed later
   *   via the @{link FilterHandle#setGroup} method.
   * @param {Object} [extraInfo] - An object whose properties will be copied to
   *   the event object whenever an event is emitted.
   */
  function FilterHandle(group, extraInfo) {
    _classCallCheck(this, FilterHandle);

    this._eventRelay = new _events2.default();
    this._emitter = new util.SubscriptionTracker(this._eventRelay);

    // Name of the group we're currently tracking, if any. Can change over time.
    this._group = null;
    // The filterSet that we're tracking, if any. Can change over time.
    this._filterSet = null;
    // The Var we're currently tracking, if any. Can change over time.
    this._filterVar = null;
    // The event handler subscription we currently have on var.on("change").
    this._varOnChangeSub = null;

    this._extraInfo = util.extend({ sender: this }, extraInfo);

    this._id = "filter" + nextId();

    this.setGroup(group);
  }

  /**
   * Changes the Crosstalk group membership of this FilterHandle. If `set()` was
   * previously called on this handle, switching groups will clear those keys
   * from the old group's filter set. These keys will not be applied to the new
   * group's filter set either. In other words, `setGroup()` effectively calls
   * `clear()` before switching groups.
   *
   * @param {string} group - The name of the Crosstalk group, or null (or
   *   undefined) to clear the group.
   */


  _createClass(FilterHandle, [{
    key: "setGroup",
    value: function setGroup(group) {
      var _this = this;

      // If group is unchanged, do nothing
      if (this._group === group) return;
      // Treat null, undefined, and other falsy values the same
      if (!this._group && !group) return;

      if (this._filterVar) {
        this._filterVar.off("change", this._varOnChangeSub);
        this.clear();
        this._varOnChangeSub = null;
        this._filterVar = null;
        this._filterSet = null;
      }

      this._group = group;

      if (group) {
        group = (0, _group2.default)(group);
        this._filterSet = getFilterSet(group);
        this._filterVar = (0, _group2.default)(group).var("filter");
        var sub = this._filterVar.on("change", function (e) {
          _this._eventRelay.trigger("change", e, _this);
        });
        this._varOnChangeSub = sub;
      }
    }

    /**
     * Combine the given `extraInfo` (if any) with the handle's default
     * `_extraInfo` (if any).
     * @private
     */

  }, {
    key: "_mergeExtraInfo",
    value: function _mergeExtraInfo(extraInfo) {
      return util.extend({}, this._extraInfo ? this._extraInfo : null, extraInfo ? extraInfo : null);
    }

    /**
     * Close the handle. This clears this handle's contribution to the filter set,
     * and unsubscribes all event listeners.
     */

  }, {
    key: "close",
    value: function close() {
      this._emitter.removeAllListeners();
      this.clear();
      this.setGroup(null);
    }

    /**
     * Clear this handle's contribution to the filter set.
     *
     * @param {Object} [extraInfo] - Extra properties to be included on the event
     *   object that's passed to listeners (in addition to any options that were
     *   passed into the `FilterHandle` constructor).
     */

  }, {
    key: "clear",
    value: function clear(extraInfo) {
      if (!this._filterSet) return;
      this._filterSet.clear(this._id);
      this._onChange(extraInfo);
    }

    /**
     * Set this handle's contribution to the filter set. This array should consist
     * of the keys of the rows that _should_ be displayed; any keys that are not
     * present in the array will be considered _filtered out_. Note that multiple
     * `FilterHandle` instances in the group may each contribute an array of keys,
     * and only those keys that appear in _all_ of the arrays make it through the
     * filter.
     *
     * @param {string[]} keys - Empty array, or array of keys. To clear the
     *   filter, don't pass an empty array; instead, use the
     *   {@link FilterHandle#clear} method.
     * @param {Object} [extraInfo] - Extra properties to be included on the event
     *   object that's passed to listeners (in addition to any options that were
     *   passed into the `FilterHandle` constructor).
     */

  }, {
    key: "set",
    value: function set(keys, extraInfo) {
      if (!this._filterSet) return;
      this._filterSet.update(this._id, keys);
      this._onChange(extraInfo);
    }

    /**
     * @return {string[]|null} - Either: 1) an array of keys that made it through
     *   all of the `FilterHandle` instances, or, 2) `null`, which means no filter
     *   is being applied (all data should be displayed).
     */

  }, {
    key: "on",


    /**
     * Subscribe to events on this `FilterHandle`.
     *
     * @param {string} eventType - Indicates the type of events to listen to.
     *   Currently, only `"change"` is supported.
     * @param {FilterHandle~listener} listener - The callback function that
     *   will be invoked when the event occurs.
     * @return {string} - A token to pass to {@link FilterHandle#off} to cancel
     *   this subscription.
     */
    value: function on(eventType, listener) {
      return this._emitter.on(eventType, listener);
    }

    /**
     * Cancel event subscriptions created by {@link FilterHandle#on}.
     *
     * @param {string} eventType - The type of event to unsubscribe.
     * @param {string|FilterHandle~listener} listener - Either the callback
     *   function previously passed into {@link FilterHandle#on}, or the
     *   string that was returned from {@link FilterHandle#on}.
     */

  }, {
    key: "off",
    value: function off(eventType, listener) {
      return this._emitter.off(eventType, listener);
    }
  }, {
    key: "_onChange",
    value: function _onChange(extraInfo) {
      if (!this._filterSet) return;
      this._filterVar.set(this._filterSet.value, this._mergeExtraInfo(extraInfo));
    }

    /**
     * @callback FilterHandle~listener
     * @param {Object} event - An object containing details of the event. For
     *   `"change"` events, this includes the properties `value` (the new
     *   value of the filter set, or `null` if no filter set is active),
     *   `oldValue` (the previous value of the filter set), and `sender` (the
     *   `FilterHandle` instance that made the change).
     */

    /**
     * @event FilterHandle#change
     * @type {object}
     * @property {object} value - The new value of the filter set, or `null`
     *   if no filter set is active.
     * @property {object} oldValue - The previous value of the filter set.
     * @property {FilterHandle} sender - The `FilterHandle` instance that
     *   changed the value.
     */

  }, {
    key: "filteredKeys",
    get: function get() {
      return this._filterSet ? this._filterSet.value : null;
    }
  }]);

  return FilterHandle;
}();

},{"./events":1,"./filterset":3,"./group":4,"./util":11}],3:[function(require,module,exports){
"use strict";

Object.defineProperty(exports, "__esModule", {
  value: true
});

var _createClass = function () { function defineProperties(target, props) { for (var i = 0; i < props.length; i++) { var descriptor = props[i]; descriptor.enumerable = descriptor.enumerable || false; descriptor.configurable = true; if ("value" in descriptor) descriptor.writable = true; Object.defineProperty(target, descriptor.key, descriptor); } } return function (Constructor, protoProps, staticProps) { if (protoProps) defineProperties(Constructor.prototype, protoProps); if (staticProps) defineProperties(Constructor, staticProps); return Constructor; }; }();

var _util = require("./util");

function _classCallCheck(instance, Constructor) { if (!(instance instanceof Constructor)) { throw new TypeError("Cannot call a class as a function"); } }

function naturalComparator(a, b) {
  if (a === b) {
    return 0;
  } else if (a < b) {
    return -1;
  } else if (a > b) {
    return 1;
  }
}

/**
 * @private
 */

var FilterSet = function () {
  function FilterSet() {
    _classCallCheck(this, FilterSet);

    this.reset();
  }

  _createClass(FilterSet, [{
    key: "reset",
    value: function reset() {
      // Key: handle ID, Value: array of selected keys, or null
      this._handles = {};
      // Key: key string, Value: count of handles that include it
      this._keys = {};
      this._value = null;
      this._activeHandles = 0;
    }
  }, {
    key: "update",
    value: function update(handleId, keys) {
      if (keys !== null) {
        keys = keys.slice(0); // clone before sorting
        keys.sort(naturalComparator);
      }

      var _diffSortedLists = (0, _util.diffSortedLists)(this._handles[handleId], keys),
          added = _diffSortedLists.added,
          removed = _diffSortedLists.removed;

      this._handles[handleId] = keys;

      for (var i = 0; i < added.length; i++) {
        this._keys[added[i]] = (this._keys[added[i]] || 0) + 1;
      }
      for (var _i = 0; _i < removed.length; _i++) {
        this._keys[removed[_i]]--;
      }

      this._updateValue(keys);
    }

    /**
     * @param {string[]} keys Sorted array of strings that indicate
     * a superset of possible keys.
     * @private
     */

  }, {
    key: "_updateValue",
    value: function _updateValue() {
      var keys = arguments.length > 0 && arguments[0] !== undefined ? arguments[0] : this._allKeys;

      var handleCount = Object.keys(this._handles).length;
      if (handleCount === 0) {
        this._value = null;
      } else {
        this._value = [];
        for (var i = 0; i < keys.length; i++) {
          var count = this._keys[keys[i]];
          if (count === handleCount) {
            this._value.push(keys[i]);
          }
        }
      }
    }
  }, {
    key: "clear",
    value: function clear(handleId) {
      if (typeof this._handles[handleId] === "undefined") {
        return;
      }

      var keys = this._handles[handleId];
      if (!keys) {
        keys = [];
      }

      for (var i = 0; i < keys.length; i++) {
        this._keys[keys[i]]--;
      }
      delete this._handles[handleId];

      this._updateValue();
    }
  }, {
    key: "value",
    get: function get() {
      return this._value;
    }
  }, {
    key: "_allKeys",
    get: function get() {
      var allKeys = Object.keys(this._keys);
      allKeys.sort(naturalComparator);
      return allKeys;
    }
  }]);

  return FilterSet;
}();

exports.default = FilterSet;

},{"./util":11}],4:[function(require,module,exports){
(function (global){
"use strict";

Object.defineProperty(exports, "__esModule", {
  value: true
});

var _createClass = function () { function defineProperties(target, props) { for (var i = 0; i < props.length; i++) { var descriptor = props[i]; descriptor.enumerable = descriptor.enumerable || false; descriptor.configurable = true; if ("value" in descriptor) descriptor.writable = true; Object.defineProperty(target, descriptor.key, descriptor); } } return function (Constructor, protoProps, staticProps) { if (protoProps) defineProperties(Constructor.prototype, protoProps); if (staticProps) defineProperties(Constructor, staticProps); return Constructor; }; }();

var _typeof = typeof Symbol === "function" && typeof Symbol.iterator === "symbol" ? function (obj) { return typeof obj; } : function (obj) { return obj && typeof Symbol === "function" && obj.constructor === Symbol && obj !== Symbol.prototype ? "symbol" : typeof obj; };

exports.default = group;

var _var2 = require("./var");

var _var3 = _interopRequireDefault(_var2);

function _interopRequireDefault(obj) { return obj && obj.__esModule ? obj : { default: obj }; }

function _classCallCheck(instance, Constructor) { if (!(instance instanceof Constructor)) { throw new TypeError("Cannot call a class as a function"); } }

// Use a global so that multiple copies of crosstalk.js can be loaded and still
// have groups behave as singletons across all copies.
global.__crosstalk_groups = global.__crosstalk_groups || {};
var groups = global.__crosstalk_groups;

function group(groupName) {
  if (groupName && typeof groupName === "string") {
    if (!groups.hasOwnProperty(groupName)) {
      groups[groupName] = new Group(groupName);
    }
    return groups[groupName];
  } else if ((typeof groupName === "undefined" ? "undefined" : _typeof(groupName)) === "object" && groupName._vars && groupName.var) {
    // Appears to already be a group object
    return groupName;
  } else if (Array.isArray(groupName) && groupName.length == 1 && typeof groupName[0] === "string") {
    return group(groupName[0]);
  } else {
    throw new Error("Invalid groupName argument");
  }
}

var Group = function () {
  function Group(name) {
    _classCallCheck(this, Group);

    this.name = name;
    this._vars = {};
  }

  _createClass(Group, [{
    key: "var",
    value: function _var(name) {
      if (!name || typeof name !== "string") {
        throw new Error("Invalid var name");
      }

      if (!this._vars.hasOwnProperty(name)) this._vars[name] = new _var3.default(this, name);
      return this._vars[name];
    }
  }, {
    key: "has",
    value: function has(name) {
      if (!name || typeof name !== "string") {
        throw new Error("Invalid var name");
      }

      return this._vars.hasOwnProperty(name);
    }
  }]);

  return Group;
}();

}).call(this,typeof global !== "undefined" ? global : typeof self !== "undefined" ? self : typeof window !== "undefined" ? window : {})

},{"./var":12}],5:[function(require,module,exports){
(function (global){
"use strict";

Object.defineProperty(exports, "__esModule", {
  value: true
});

var _group = require("./group");

var _group2 = _interopRequireDefault(_group);

var _selection = require("./selection");

var _filter = require("./filter");

require("./input");

require("./input_selectize");

require("./input_checkboxgroup");

require("./input_slider");

function _interopRequireDefault(obj) { return obj && obj.__esModule ? obj : { default: obj }; }

var defaultGroup = (0, _group2.default)("default");

function var_(name) {
  return defaultGroup.var(name);
}

function has(name) {
  return defaultGroup.has(name);
}

if (global.Shiny) {
  global.Shiny.addCustomMessageHandler("update-client-value", function (message) {
    if (typeof message.group === "string") {
      (0, _group2.default)(message.group).var(message.name).set(message.value);
    } else {
      var_(message.name).set(message.value);
    }
  });
}

var crosstalk = {
  group: _group2.default,
  var: var_,
  has: has,
  SelectionHandle: _selection.SelectionHandle,
  FilterHandle: _filter.FilterHandle
};

/**
 * @namespace crosstalk
 */
exports.default = crosstalk;

global.crosstalk = crosstalk;

}).call(this,typeof global !== "undefined" ? global : typeof self !== "undefined" ? self : typeof window !== "undefined" ? window : {})

},{"./filter":2,"./group":4,"./input":6,"./input_checkboxgroup":7,"./input_selectize":8,"./input_slider":9,"./selection":10}],6:[function(require,module,exports){
(function (global){
"use strict";

Object.defineProperty(exports, "__esModule", {
  value: true
});
exports.register = register;
var $ = global.jQuery;

var bindings = {};

function register(reg) {
  bindings[reg.className] = reg;
  if (global.document && global.document.readyState !== "complete") {
    $(function () {
      bind();
    });
  } else if (global.document) {
    setTimeout(bind, 100);
  }
}

function bind() {
  Object.keys(bindings).forEach(function (className) {
    var binding = bindings[className];
    $("." + binding.className).not(".crosstalk-input-bound").each(function (i, el) {
      bindInstance(binding, el);
    });
  });
}

// Escape jQuery identifier
function $escape(val) {
  return val.replace(/([!"#$%&'()*+,.\/:;<=>?@\[\\\]^`{|}~])/g, "\\$1");
}

function bindEl(el) {
  var $el = $(el);
  Object.keys(bindings).forEach(function (className) {
    if ($el.hasClass(className) && !$el.hasClass("crosstalk-input-bound")) {
      var binding = bindings[className];
      bindInstance(binding, el);
    }
  });
}

function bindInstance(binding, el) {
  var jsonEl = $(el).find("script[type='application/json'][data-for='" + $escape(el.id) + "']");
  var data = JSON.parse(jsonEl[0].innerText);

  var instance = binding.factory(el, data);
  $(el).data("crosstalk-instance", instance);
  $(el).addClass("crosstalk-input-bound");
}

if (global.Shiny) {
  (function () {
    var inputBinding = new global.Shiny.InputBinding();
    var $ = global.jQuery;
    $.extend(inputBinding, {
      find: function find(scope) {
        return $(scope).find(".crosstalk-input");
      },
      initialize: function initialize(el) {
        if (!$(el).hasClass("crosstalk-input-bound")) {
          bindEl(el);
        }
      },
      getId: function getId(el) {
        return el.id;
      },
      getValue: function getValue(el) {},
      setValue: function setValue(el, value) {},
      receiveMessage: function receiveMessage(el, data) {},
      subscribe: function subscribe(el, callback) {
        $(el).data("crosstalk-instance").resume();
      },
      unsubscribe: function unsubscribe(el) {
        $(el).data("crosstalk-instance").suspend();
      }
    });
    global.Shiny.inputBindings.register(inputBinding, "crosstalk.inputBinding");
  })();
}

}).call(this,typeof global !== "undefined" ? global : typeof self !== "undefined" ? self : typeof window !== "undefined" ? window : {})

},{}],7:[function(require,module,exports){
(function (global){
"use strict";

var _input = require("./input");

var input = _interopRequireWildcard(_input);

var _filter = require("./filter");

function _interopRequireWildcard(obj) { if (obj && obj.__esModule) { return obj; } else { var newObj = {}; if (obj != null) { for (var key in obj) { if (Object.prototype.hasOwnProperty.call(obj, key)) newObj[key] = obj[key]; } } newObj.default = obj; return newObj; } }

var $ = global.jQuery;

input.register({
  className: "crosstalk-input-checkboxgroup",

  factory: function factory(el, data) {
    /*
     * map: {"groupA": ["keyA", "keyB", ...], ...}
     * group: "ct-groupname"
     */
    var ctHandle = new _filter.FilterHandle(data.group);

    var lastKnownKeys = void 0;
    var $el = $(el);
    $el.on("change", "input[type='checkbox']", function () {
      var checked = $el.find("input[type='checkbox']:checked");
      if (checked.length === 0) {
        lastKnownKeys = null;
        ctHandle.clear();
      } else {
        (function () {
          var keys = {};
          checked.each(function () {
            data.map[this.value].forEach(function (key) {
              keys[key] = true;
            });
          });
          var keyArray = Object.keys(keys);
          keyArray.sort();
          lastKnownKeys = keyArray;
          ctHandle.set(keyArray);
        })();
      }
    });

    return {
      suspend: function suspend() {
        ctHandle.clear();
      },
      resume: function resume() {
        if (lastKnownKeys) ctHandle.set(lastKnownKeys);
      }
    };
  }
});

}).call(this,typeof global !== "undefined" ? global : typeof self !== "undefined" ? self : typeof window !== "undefined" ? window : {})

},{"./filter":2,"./input":6}],8:[function(require,module,exports){
(function (global){
"use strict";

var _input = require("./input");

var input = _interopRequireWildcard(_input);

var _util = require("./util");

var util = _interopRequireWildcard(_util);

var _filter = require("./filter");

function _interopRequireWildcard(obj) { if (obj && obj.__esModule) { return obj; } else { var newObj = {}; if (obj != null) { for (var key in obj) { if (Object.prototype.hasOwnProperty.call(obj, key)) newObj[key] = obj[key]; } } newObj.default = obj; return newObj; } }

var $ = global.jQuery;

input.register({
  className: "crosstalk-input-select",

  factory: function factory(el, data) {
    /*
     * items: {value: [...], label: [...]}
     * map: {"groupA": ["keyA", "keyB", ...], ...}
     * group: "ct-groupname"
     */

    var first = [{ value: "", label: "(All)" }];
    var items = util.dataframeToD3(data.items);
    var opts = {
      options: first.concat(items),
      valueField: "value",
      labelField: "label",
      searchField: "label"
    };

    var select = $(el).find("select")[0];

    var selectize = $(select).selectize(opts)[0].selectize;

    var ctHandle = new _filter.FilterHandle(data.group);

    var lastKnownKeys = void 0;
    selectize.on("change", function () {
      if (selectize.items.length === 0) {
        lastKnownKeys = null;
        ctHandle.clear();
      } else {
        (function () {
          var keys = {};
          selectize.items.forEach(function (group) {
            data.map[group].forEach(function (key) {
              keys[key] = true;
            });
          });
          var keyArray = Object.keys(keys);
          keyArray.sort();
          lastKnownKeys = keyArray;
          ctHandle.set(keyArray);
        })();
      }
    });

    return {
      suspend: function suspend() {
        ctHandle.clear();
      },
      resume: function resume() {
        if (lastKnownKeys) ctHandle.set(lastKnownKeys);
      }
    };
  }
});

}).call(this,typeof global !== "undefined" ? global : typeof self !== "undefined" ? self : typeof window !== "undefined" ? window : {})

},{"./filter":2,"./input":6,"./util":11}],9:[function(require,module,exports){
(function (global){
"use strict";

var _slicedToArray = function () { function sliceIterator(arr, i) { var _arr = []; var _n = true; var _d = false; var _e = undefined; try { for (var _i = arr[Symbol.iterator](), _s; !(_n = (_s = _i.next()).done); _n = true) { _arr.push(_s.value); if (i && _arr.length === i) break; } } catch (err) { _d = true; _e = err; } finally { try { if (!_n && _i["return"]) _i["return"](); } finally { if (_d) throw _e; } } return _arr; } return function (arr, i) { if (Array.isArray(arr)) { return arr; } else if (Symbol.iterator in Object(arr)) { return sliceIterator(arr, i); } else { throw new TypeError("Invalid attempt to destructure non-iterable instance"); } }; }();

var _input = require("./input");

var input = _interopRequireWildcard(_input);

var _filter = require("./filter");

function _interopRequireWildcard(obj) { if (obj && obj.__esModule) { return obj; } else { var newObj = {}; if (obj != null) { for (var key in obj) { if (Object.prototype.hasOwnProperty.call(obj, key)) newObj[key] = obj[key]; } } newObj.default = obj; return newObj; } }

var $ = global.jQuery;
var strftime = global.strftime;

input.register({
  className: "crosstalk-input-slider",

  factory: function factory(el, data) {
    /*
     * map: {"groupA": ["keyA", "keyB", ...], ...}
     * group: "ct-groupname"
     */
    var ctHandle = new _filter.FilterHandle(data.group);

    var opts = {};
    var $el = $(el).find("input");
    var dataType = $el.data("data-type");
    var timeFormat = $el.data("time-format");
    var timeFormatter = void 0;

    // Set up formatting functions
    if (dataType === "date") {
      timeFormatter = strftime.utc();
      opts.prettify = function (num) {
        return timeFormatter(timeFormat, new Date(num));
      };
    } else if (dataType === "datetime") {
      var timezone = $el.data("timezone");
      if (timezone) timeFormatter = strftime.timezone(timezone);else timeFormatter = strftime;

      opts.prettify = function (num) {
        return timeFormatter(timeFormat, new Date(num));
      };
    }

    $el.ionRangeSlider(opts);

    function getValue() {
      var result = $el.data("ionRangeSlider").result;

      // Function for converting numeric value from slider to appropriate type.
      var convert = void 0;
      var dataType = $el.data("data-type");
      if (dataType === "date") {
        convert = function convert(val) {
          return formatDateUTC(new Date(+val));
        };
      } else if (dataType === "datetime") {
        convert = function convert(val) {
          // Convert ms to s
          return +val / 1000;
        };
      } else {
        convert = function convert(val) {
          return +val;
        };
      }

      if ($el.data("ionRangeSlider").options.type === "double") {
        return [convert(result.from), convert(result.to)];
      } else {
        return convert(result.from);
      }
    }

    var lastKnownKeys = null;

    $el.on("change.crosstalkSliderInput", function (event) {
      if (!$el.data("updating") && !$el.data("animating")) {
        var _getValue = getValue(),
            _getValue2 = _slicedToArray(_getValue, 2),
            from = _getValue2[0],
            to = _getValue2[1];

        var keys = [];
        for (var i = 0; i < data.values.length; i++) {
          var val = data.values[i];
          if (val >= from && val <= to) {
            keys.push(data.keys[i]);
          }
        }
        keys.sort();
        ctHandle.set(keys);
        lastKnownKeys = keys;
      }
    });

    // let $el = $(el);
    // $el.on("change", "input[type="checkbox"]", function() {
    //   let checked = $el.find("input[type="checkbox"]:checked");
    //   if (checked.length === 0) {
    //     ctHandle.clear();
    //   } else {
    //     let keys = {};
    //     checked.each(function() {
    //       data.map[this.value].forEach(function(key) {
    //         keys[key] = true;
    //       });
    //     });
    //     let keyArray = Object.keys(keys);
    //     keyArray.sort();
    //     ctHandle.set(keyArray);
    //   }
    // });

    return {
      suspend: function suspend() {
        ctHandle.clear();
      },
      resume: function resume() {
        if (lastKnownKeys) ctHandle.set(lastKnownKeys);
      }
    };
  }
});

// Convert a number to a string with leading zeros
function padZeros(n, digits) {
  var str = n.toString();
  while (str.length < digits) {
    str = "0" + str;
  }return str;
}

// Given a Date object, return a string in yyyy-mm-dd format, using the
// UTC date. This may be a day off from the date in the local time zone.
function formatDateUTC(date) {
  if (date instanceof Date) {
    return date.getUTCFullYear() + "-" + padZeros(date.getUTCMonth() + 1, 2) + "-" + padZeros(date.getUTCDate(), 2);
  } else {
    return null;
  }
}

}).call(this,typeof global !== "undefined" ? global : typeof self !== "undefined" ? self : typeof window !== "undefined" ? window : {})

},{"./filter":2,"./input":6}],10:[function(require,module,exports){
"use strict";

Object.defineProperty(exports, "__esModule", {
  value: true
});
exports.SelectionHandle = undefined;

var _createClass = function () { function defineProperties(target, props) { for (var i = 0; i < props.length; i++) { var descriptor = props[i]; descriptor.enumerable = descriptor.enumerable || false; descriptor.configurable = true; if ("value" in descriptor) descriptor.writable = true; Object.defineProperty(target, descriptor.key, descriptor); } } return function (Constructor, protoProps, staticProps) { if (protoProps) defineProperties(Constructor.prototype, protoProps); if (staticProps) defineProperties(Constructor, staticProps); return Constructor; }; }();

var _events = require("./events");

var _events2 = _interopRequireDefault(_events);

var _group = require("./group");

var _group2 = _interopRequireDefault(_group);

var _util = require("./util");

var util = _interopRequireWildcard(_util);

function _interopRequireWildcard(obj) { if (obj && obj.__esModule) { return obj; } else { var newObj = {}; if (obj != null) { for (var key in obj) { if (Object.prototype.hasOwnProperty.call(obj, key)) newObj[key] = obj[key]; } } newObj.default = obj; return newObj; } }

function _interopRequireDefault(obj) { return obj && obj.__esModule ? obj : { default: obj }; }

function _classCallCheck(instance, Constructor) { if (!(instance instanceof Constructor)) { throw new TypeError("Cannot call a class as a function"); } }

var SelectionHandle = exports.SelectionHandle = function () {

  /**
   * @classdesc
   * Use this class to read and write (and listen for changes to) the selection
   * for a Crosstalk group. This is intended to be used for linked brushing.
   *
   * If two (or more) `SelectionHandle` instances in the same webpage share the
   * same group name, they will share the same state. Setting the selection using
   * one `SelectionHandle` instance will result in the `value` property instantly
   * changing across the others, and `"change"` event listeners on all instances
   * (including the one that initiated the sending) will fire.
   *
   * @param {string} [group] - The name of the Crosstalk group, or if none,
   *   null or undefined (or any other falsy value). This can be changed later
   *   via the [SelectionHandle#setGroup](#setGroup) method.
   * @param {Object} [extraInfo] - An object whose properties will be copied to
   *   the event object whenever an event is emitted.
   */
  function SelectionHandle() {
    var group = arguments.length > 0 && arguments[0] !== undefined ? arguments[0] : null;
    var extraInfo = arguments.length > 1 && arguments[1] !== undefined ? arguments[1] : null;

    _classCallCheck(this, SelectionHandle);

    this._eventRelay = new _events2.default();
    this._emitter = new util.SubscriptionTracker(this._eventRelay);

    // Name of the group we're currently tracking, if any. Can change over time.
    this._group = null;
    // The Var we're currently tracking, if any. Can change over time.
    this._var = null;
    // The event handler subscription we currently have on var.on("change").
    this._varOnChangeSub = null;

    this._extraInfo = util.extend({ sender: this }, extraInfo);

    this.setGroup(group);
  }

  /**
   * Changes the Crosstalk group membership of this SelectionHandle. The group
   * being switched away from (if any) will not have its selection value
   * modified as a result of calling `setGroup`, even if this handle was the
   * most recent handle to set the selection of the group.
   *
   * The group being switched to (if any) will also not have its selection value
   * modified as a result of calling `setGroup`. If you want to set the
   * selection value of the new group, call `set` explicitly.
   *
   * @param {string} group - The name of the Crosstalk group, or null (or
   *   undefined) to clear the group.
   */


  _createClass(SelectionHandle, [{
    key: "setGroup",
    value: function setGroup(group) {
      var _this = this;

      // If group is unchanged, do nothing
      if (this._group === group) return;
      // Treat null, undefined, and other falsy values the same
      if (!this._group && !group) return;

      if (this._var) {
        this._var.off("change", this._varOnChangeSub);
        this._var = null;
        this._varOnChangeSub = null;
      }

      this._group = group;

      if (group) {
        this._var = (0, _group2.default)(group).var("selection");
        var sub = this._var.on("change", function (e) {
          _this._eventRelay.trigger("change", e, _this);
        });
        this._varOnChangeSub = sub;
      }
    }

    /**
     * Retrieves the current selection for the group represented by this
     * `SelectionHandle`.
     *
     * - If no selection is active, then this value will be falsy.
     * - If a selection is active, but no data points are selected, then this
     *   value will be an empty array.
     * - If a selection is active, and data points are selected, then the keys
     *   of the selected data points will be present in the array.
     */

  }, {
    key: "_mergeExtraInfo",


    /**
     * Combines the given `extraInfo` (if any) with the handle's default
     * `_extraInfo` (if any).
     * @private
     */
    value: function _mergeExtraInfo(extraInfo) {
      // Important incidental effect: shallow clone is returned
      return util.extend({}, this._extraInfo ? this._extraInfo : null, extraInfo ? extraInfo : null);
    }

    /**
     * Overwrites the current selection for the group, and raises the `"change"`
     * event among all of the group's '`SelectionHandle` instances (including
     * this one).
     *
     * @fires SelectionHandle#change
     * @param {string[]} selectedKeys - Falsy, empty array, or array of keys (see
     *   {@link SelectionHandle#value}).
     * @param {Object} [extraInfo] - Extra properties to be included on the event
     *   object that's passed to listeners (in addition to any options that were
     *   passed into the `SelectionHandle` constructor).
     */

  }, {
    key: "set",
    value: function set(selectedKeys, extraInfo) {
      if (this._var) this._var.set(selectedKeys, this._mergeExtraInfo(extraInfo));
    }

    /**
     * Overwrites the current selection for the group, and raises the `"change"`
     * event among all of the group's '`SelectionHandle` instances (including
     * this one).
     *
     * @fires SelectionHandle#change
     * @param {Object} [extraInfo] - Extra properties to be included on the event
     *   object that's passed to listeners (in addition to any that were passed
     *   into the `SelectionHandle` constructor).
     */

  }, {
    key: "clear",
    value: function clear(extraInfo) {
      if (this._var) this.set(void 0, this._mergeExtraInfo(extraInfo));
    }

    /**
     * Subscribes to events on this `SelectionHandle`.
     *
     * @param {string} eventType - Indicates the type of events to listen to.
     *   Currently, only `"change"` is supported.
     * @param {SelectionHandle~listener} listener - The callback function that
     *   will be invoked when the event occurs.
     * @return {string} - A token to pass to {@link SelectionHandle#off} to cancel
     *   this subscription.
     */

  }, {
    key: "on",
    value: function on(eventType, listener) {
      return this._emitter.on(eventType, listener);
    }

    /**
     * Cancels event subscriptions created by {@link SelectionHandle#on}.
     *
     * @param {string} eventType - The type of event to unsubscribe.
     * @param {string|SelectionHandle~listener} listener - Either the callback
     *   function previously passed into {@link SelectionHandle#on}, or the
     *   string that was returned from {@link SelectionHandle#on}.
     */

  }, {
    key: "off",
    value: function off(eventType, listener) {
      return this._emitter.off(eventType, listener);
    }

    /**
     * Shuts down the `SelectionHandle` object.
     *
     * Removes all event listeners that were added through this handle.
     */

  }, {
    key: "close",
    value: function close() {
      this._emitter.removeAllListeners();
      this.setGroup(null);
    }

    /**
     * @callback SelectionHandle~listener
     * @param {Object} event - An object containing details of the event. For
     *   `"change"` events, this includes the properties `value` (the new
     *   value of the selection, or `undefined` if no selection is active),
     *   `oldValue` (the previous value of the selection), and `sender` (the
     *   `SelectionHandle` instance that made the change).
     */

    /**
     * @event SelectionHandle#change
     * @type {object}
     * @property {object} value - The new value of the selection, or `undefined`
     *   if no selection is active.
     * @property {object} oldValue - The previous value of the selection.
     * @property {SelectionHandle} sender - The `SelectionHandle` instance that
     *   changed the value.
     */

  }, {
    key: "value",
    get: function get() {
      return this._var ? this._var.get() : null;
    }
  }]);

  return SelectionHandle;
}();

},{"./events":1,"./group":4,"./util":11}],11:[function(require,module,exports){
"use strict";

Object.defineProperty(exports, "__esModule", {
  value: true
});

var _createClass = function () { function defineProperties(target, props) { for (var i = 0; i < props.length; i++) { var descriptor = props[i]; descriptor.enumerable = descriptor.enumerable || false; descriptor.configurable = true; if ("value" in descriptor) descriptor.writable = true; Object.defineProperty(target, descriptor.key, descriptor); } } return function (Constructor, protoProps, staticProps) { if (protoProps) defineProperties(Constructor.prototype, protoProps); if (staticProps) defineProperties(Constructor, staticProps); return Constructor; }; }();

var _typeof = typeof Symbol === "function" && typeof Symbol.iterator === "symbol" ? function (obj) { return typeof obj; } : function (obj) { return obj && typeof Symbol === "function" && obj.constructor === Symbol && obj !== Symbol.prototype ? "symbol" : typeof obj; };

exports.extend = extend;
exports.checkSorted = checkSorted;
exports.diffSortedLists = diffSortedLists;
exports.dataframeToD3 = dataframeToD3;

function _classCallCheck(instance, Constructor) { if (!(instance instanceof Constructor)) { throw new TypeError("Cannot call a class as a function"); } }

function extend(target) {
  for (var _len = arguments.length, sources = Array(_len > 1 ? _len - 1 : 0), _key = 1; _key < _len; _key++) {
    sources[_key - 1] = arguments[_key];
  }

  for (var i = 0; i < sources.length; i++) {
    var src = sources[i];
    if (typeof src === "undefined" || src === null) continue;

    for (var key in src) {
      if (src.hasOwnProperty(key)) {
        target[key] = src[key];
      }
    }
  }
  return target;
}

function checkSorted(list) {
  for (var i = 1; i < list.length; i++) {
    if (list[i] <= list[i - 1]) {
      throw new Error("List is not sorted or contains duplicate");
    }
  }
}

function diffSortedLists(a, b) {
  var i_a = 0;
  var i_b = 0;

  if (!a) a = [];
  if (!b) b = [];

  var a_only = [];
  var b_only = [];

  checkSorted(a);
  checkSorted(b);

  while (i_a < a.length && i_b < b.length) {
    if (a[i_a] === b[i_b]) {
      i_a++;
      i_b++;
    } else if (a[i_a] < b[i_b]) {
      a_only.push(a[i_a++]);
    } else {
      b_only.push(b[i_b++]);
    }
  }

  if (i_a < a.length) a_only = a_only.concat(a.slice(i_a));
  if (i_b < b.length) b_only = b_only.concat(b.slice(i_b));
  return {
    removed: a_only,
    added: b_only
  };
}

// Convert from wide: { colA: [1,2,3], colB: [4,5,6], ... }
// to long: [ {colA: 1, colB: 4}, {colA: 2, colB: 5}, ... ]
function dataframeToD3(df) {
  var names = [];
  var length = void 0;
  for (var name in df) {
    if (df.hasOwnProperty(name)) names.push(name);
    if (_typeof(df[name]) !== "object" || typeof df[name].length === "undefined") {
      throw new Error("All fields must be arrays");
    } else if (typeof length !== "undefined" && length !== df[name].length) {
      throw new Error("All fields must be arrays of the same length");
    }
    length = df[name].length;
  }
  var results = [];
  var item = void 0;
  for (var row = 0; row < length; row++) {
    item = {};
    for (var col = 0; col < names.length; col++) {
      item[names[col]] = df[names[col]][row];
    }
    results.push(item);
  }
  return results;
}

/**
 * Keeps track of all event listener additions/removals and lets all active
 * listeners be removed with a single operation.
 *
 * @private
 */

var SubscriptionTracker = exports.SubscriptionTracker = function () {
  function SubscriptionTracker(emitter) {
    _classCallCheck(this, SubscriptionTracker);

    this._emitter = emitter;
    this._subs = {};
  }

  _createClass(SubscriptionTracker, [{
    key: "on",
    value: function on(eventType, listener) {
      var sub = this._emitter.on(eventType, listener);
      this._subs[sub] = eventType;
      return sub;
    }
  }, {
    key: "off",
    value: function off(eventType, listener) {
      var sub = this._emitter.off(eventType, listener);
      if (sub) {
        delete this._subs[sub];
      }
      return sub;
    }
  }, {
    key: "removeAllListeners",
    value: function removeAllListeners() {
      var _this = this;

      var current_subs = this._subs;
      this._subs = {};
      Object.keys(current_subs).forEach(function (sub) {
        _this._emitter.off(current_subs[sub], sub);
      });
    }
  }]);

  return SubscriptionTracker;
}();

},{}],12:[function(require,module,exports){
(function (global){
"use strict";

Object.defineProperty(exports, "__esModule", {
  value: true
});

var _typeof = typeof Symbol === "function" && typeof Symbol.iterator === "symbol" ? function (obj) { return typeof obj; } : function (obj) { return obj && typeof Symbol === "function" && obj.constructor === Symbol && obj !== Symbol.prototype ? "symbol" : typeof obj; };

var _createClass = function () { function defineProperties(target, props) { for (var i = 0; i < props.length; i++) { var descriptor = props[i]; descriptor.enumerable = descriptor.enumerable || false; descriptor.configurable = true; if ("value" in descriptor) descriptor.writable = true; Object.defineProperty(target, descriptor.key, descriptor); } } return function (Constructor, protoProps, staticProps) { if (protoProps) defineProperties(Constructor.prototype, protoProps); if (staticProps) defineProperties(Constructor, staticProps); return Constructor; }; }();

var _events = require("./events");

var _events2 = _interopRequireDefault(_events);

function _interopRequireDefault(obj) { return obj && obj.__esModule ? obj : { default: obj }; }

function _classCallCheck(instance, Constructor) { if (!(instance instanceof Constructor)) { throw new TypeError("Cannot call a class as a function"); } }

var Var = function () {
  function Var(group, name, /*optional*/value) {
    _classCallCheck(this, Var);

    this._group = group;
    this._name = name;
    this._value = value;
    this._events = new _events2.default();
  }

  _createClass(Var, [{
    key: "get",
    value: function get() {
      return this._value;
    }
  }, {
    key: "set",
    value: function set(value, /*optional*/event) {
      if (this._value === value) {
        // Do nothing; the value hasn't changed
        return;
      }
      var oldValue = this._value;
      this._value = value;
      // Alert JavaScript listeners that the value has changed
      var evt = {};
      if (event && (typeof event === "undefined" ? "undefined" : _typeof(event)) === "object") {
        for (var k in event) {
          if (event.hasOwnProperty(k)) evt[k] = event[k];
        }
      }
      evt.oldValue = oldValue;
      evt.value = value;
      this._events.trigger("change", evt, this);

      // TODO: Make this extensible, to let arbitrary back-ends know that
      // something has changed
      if (global.Shiny && global.Shiny.onInputChange) {
        global.Shiny.onInputChange(".clientValue-" + (this._group.name !== null ? this._group.name + "-" : "") + this._name, typeof value === "undefined" ? null : value);
      }
    }
  }, {
    key: "on",
    value: function on(eventType, listener) {
      return this._events.on(eventType, listener);
    }
  }, {
    key: "off",
    value: function off(eventType, listener) {
      return this._events.off(eventType, listener);
    }
  }]);

  return Var;
}();

exports.default = Var;

}).call(this,typeof global !== "undefined" ? global : typeof self !== "undefined" ? self : typeof window !== "undefined" ? window : {})

},{"./events":1}]},{},[5])
//# sourceMappingURL=crosstalk.js.map
```
