## Supplementary material for "Characterizing and inferring quantitative cell cycle phase in single-cell RNA-seq data analysis": Source code of our analysis, including scripts and data necessary to reproduce the work.: crosstalk.min.js

```
!function a(b,c,d){function e(g,h){if(!c[g]){if(!b[g]){var i="function"==typeof require&&require;if(!h&&i)return i(g,!0);if(f)return f(g,!0);var j=new Error("Cannot find module '"+g+"'");throw j.code="MODULE_NOT_FOUND",j}var k=c[g]={exports:{}};b[g][0].call(k.exports,function(a){var c=b[g][1][a];return e(c?c:a)},k,k.exports,a,b,c,d)}return c[g].exports}for(var f="function"==typeof require&&require,g=0;g<d.length;g++)e(d[g]);return e}({1:[function(a,b,c){"use strict";function d(a,b){if(!(a instanceof b))throw new TypeError("Cannot call a class as a function")}Object.defineProperty(c,"__esModule",{value:!0});var e=function(){function a(a,b){for(var c=0;c<b.length;c++){var d=b[c];d.enumerable=d.enumerable||!1,d.configurable=!0,"value"in d&&(d.writable=!0),Object.defineProperty(a,d.key,d)}}return function(b,c,d){return c&&a(b.prototype,c),d&&a(b,d),b}}(),f=function(){function a(){d(this,a),this._types={},this._seq=0}return e(a,[{key:"on",value:function(a,b){var c=this._types[a];c||(c=this._types[a]={});var d="sub"+this._seq++;return c[d]=b,d}},{key:"off",value:function(a,b){var c=this._types[a];if("function"==typeof b){for(var d in c)if(c.hasOwnProperty(d)&&c[d]===b)return delete c[d],d;return!1}if("string"==typeof b)return!(!c||!c[b])&&(delete c[b],b);throw new Error("Unexpected type for listener")}},{key:"trigger",value:function(a,b,c){var d=this._types[a];for(var e in d)d.hasOwnProperty(e)&&d[e].call(c,b)}}]),a}();c.default=f},{}],2:[function(a,b,c){"use strict";function d(a){if(a&&a.__esModule)return a;var b={};if(null!=a)for(var c in a)Object.prototype.hasOwnProperty.call(a,c)&&(b[c]=a[c]);return b.default=a,b}function e(a){return a&&a.__esModule?a:{default:a}}function f(a,b){if(!(a instanceof b))throw new TypeError("Cannot call a class as a function")}function g(a){var b=a.var("filterset"),c=b.get();return c||(c=new m.default,b.set(c)),c}function h(){return r++}Object.defineProperty(c,"__esModule",{value:!0}),c.FilterHandle=void 0;var i=function(){function a(a,b){for(var c=0;c<b.length;c++){var d=b[c];d.enumerable=d.enumerable||!1,d.configurable=!0,"value"in d&&(d.writable=!0),Object.defineProperty(a,d.key,d)}}return function(b,c,d){return c&&a(b.prototype,c),d&&a(b,d),b}}(),j=a("./events"),k=e(j),l=a("./filterset"),m=e(l),n=a("./group"),o=e(n),p=a("./util"),q=d(p),r=1;c.FilterHandle=function(){function a(b,c){f(this,a),this._eventRelay=new k.default,this._emitter=new q.SubscriptionTracker(this._eventRelay),this._group=null,this._filterSet=null,this._filterVar=null,this._varOnChangeSub=null,this._extraInfo=q.extend({sender:this},c),this._id="filter"+h(),this.setGroup(b)}return i(a,[{key:"setGroup",value:function(a){var b=this;if(this._group!==a&&(this._group||a)&&(this._filterVar&&(this._filterVar.off("change",this._varOnChangeSub),this.clear(),this._varOnChangeSub=null,this._filterVar=null,this._filterSet=null),this._group=a,a)){a=(0,o.default)(a),this._filterSet=g(a),this._filterVar=(0,o.default)(a).var("filter");var c=this._filterVar.on("change",function(a){b._eventRelay.trigger("change",a,b)});this._varOnChangeSub=c}}},{key:"_mergeExtraInfo",value:function(a){return q.extend({},this._extraInfo?this._extraInfo:null,a?a:null)}},{key:"close",value:function(){this._emitter.removeAllListeners(),this.clear(),this.setGroup(null)}},{key:"clear",value:function(a){this._filterSet&&(this._filterSet.clear(this._id),this._onChange(a))}},{key:"set",value:function(a,b){this._filterSet&&(this._filterSet.update(this._id,a),this._onChange(b))}},{key:"on",value:function(a,b){return this._emitter.on(a,b)}},{key:"off",value:function(a,b){return this._emitter.off(a,b)}},{key:"_onChange",value:function(a){this._filterSet&&this._filterVar.set(this._filterSet.value,this._mergeExtraInfo(a))}},{key:"filteredKeys",get:function(){return this._filterSet?this._filterSet.value:null}}]),a}()},{"./events":1,"./filterset":3,"./group":4,"./util":11}],3:[function(a,b,c){"use strict";function d(a,b){if(!(a instanceof b))throw new TypeError("Cannot call a class as a function")}function e(a,b){return a===b?0:a<b?-1:a>b?1:void 0}Object.defineProperty(c,"__esModule",{value:!0});var f=function(){function a(a,b){for(var c=0;c<b.length;c++){var d=b[c];d.enumerable=d.enumerable||!1,d.configurable=!0,"value"in d&&(d.writable=!0),Object.defineProperty(a,d.key,d)}}return function(b,c,d){return c&&a(b.prototype,c),d&&a(b,d),b}}(),g=a("./util"),h=function(){function a(){d(this,a),this.reset()}return f(a,[{key:"reset",value:function(){this._handles={},this._keys={},this._value=null,this._activeHandles=0}},{key:"update",value:function(a,b){null!==b&&(b=b.slice(0),b.sort(e));var c=(0,g.diffSortedLists)(this._handles[a],b),d=c.added,f=c.removed;this._handles[a]=b;for(var h=0;h<d.length;h++)this._keys[d[h]]=(this._keys[d[h]]||0)+1;for(var i=0;i<f.length;i++)this._keys[f[i]]--;this._updateValue(b)}},{key:"_updateValue",value:function(){var a=arguments.length>0&&void 0!==arguments[0]?arguments[0]:this._allKeys,b=Object.keys(this._handles).length;if(0===b)this._value=null;else{this._value=[];for(var c=0;c<a.length;c++){var d=this._keys[a[c]];d===b&&this._value.push(a[c])}}}},{key:"clear",value:function(a){if("undefined"!=typeof this._handles[a]){var b=this._handles[a];b||(b=[]);for(var c=0;c<b.length;c++)this._keys[b[c]]--;delete this._handles[a],this._updateValue()}}},{key:"value",get:function(){return this._value}},{key:"_allKeys",get:function(){var a=Object.keys(this._keys);return a.sort(e),a}}]),a}();c.default=h},{"./util":11}],4:[function(a,b,c){(function(b){"use strict";function d(a){return a&&a.__esModule?a:{default:a}}function e(a,b){if(!(a instanceof b))throw new TypeError("Cannot call a class as a function")}function f(a){if(a&&"string"==typeof a)return k.hasOwnProperty(a)||(k[a]=new l(a)),k[a];if("object"===("undefined"==typeof a?"undefined":h(a))&&a._vars&&a.var)return a;if(Array.isArray(a)&&1==a.length&&"string"==typeof a[0])return f(a[0]);throw new Error("Invalid groupName argument")}Object.defineProperty(c,"__esModule",{value:!0});var g=function(){function a(a,b){for(var c=0;c<b.length;c++){var d=b[c];d.enumerable=d.enumerable||!1,d.configurable=!0,"value"in d&&(d.writable=!0),Object.defineProperty(a,d.key,d)}}return function(b,c,d){return c&&a(b.prototype,c),d&&a(b,d),b}}(),h="function"==typeof Symbol&&"symbol"==typeof Symbol.iterator?function(a){return typeof a}:function(a){return a&&"function"==typeof Symbol&&a.constructor===Symbol&&a!==Symbol.prototype?"symbol":typeof a};c.default=f;var i=a("./var"),j=d(i);b.__crosstalk_groups=b.__crosstalk_groups||{};var k=b.__crosstalk_groups,l=function(){function a(b){e(this,a),this.name=b,this._vars={}}return g(a,[{key:"var",value:function(a){if(!a||"string"!=typeof a)throw new Error("Invalid var name");return this._vars.hasOwnProperty(a)||(this._vars[a]=new j.default(this,a)),this._vars[a]}},{key:"has",value:function(a){if(!a||"string"!=typeof a)throw new Error("Invalid var name");return this._vars.hasOwnProperty(a)}}]),a}()}).call(this,"undefined"!=typeof global?global:"undefined"!=typeof self?self:"undefined"!=typeof window?window:{})},{"./var":12}],5:[function(a,b,c){(function(b){"use strict";function d(a){return a&&a.__esModule?a:{default:a}}function e(a){return k.var(a)}function f(a){return k.has(a)}Object.defineProperty(c,"__esModule",{value:!0});var g=a("./group"),h=d(g),i=a("./selection"),j=a("./filter");a("./input"),a("./input_selectize"),a("./input_checkboxgroup"),a("./input_slider");var k=(0,h.default)("default");b.Shiny&&b.Shiny.addCustomMessageHandler("update-client-value",function(a){"string"==typeof a.group?(0,h.default)(a.group).var(a.name).set(a.value):e(a.name).set(a.value)});var l={group:h.default,var:e,has:f,SelectionHandle:i.SelectionHandle,FilterHandle:j.FilterHandle};c.default=l,b.crosstalk=l}).call(this,"undefined"!=typeof global?global:"undefined"!=typeof self?self:"undefined"!=typeof window?window:{})},{"./filter":2,"./group":4,"./input":6,"./input_checkboxgroup":7,"./input_selectize":8,"./input_slider":9,"./selection":10}],6:[function(a,b,c){(function(a){"use strict";function b(b){i[b.className]=b,a.document&&"complete"!==a.document.readyState?h(function(){d()}):a.document&&setTimeout(d,100)}function d(){Object.keys(i).forEach(function(a){var b=i[a];h("."+b.className).not(".crosstalk-input-bound").each(function(a,c){g(b,c)})})}function e(a){return a.replace(/([!"#$%&'()*+,.\/:;<=>?@\[\\\]^`{|}~])/g,"\\$1")}function f(a){var b=h(a);Object.keys(i).forEach(function(c){if(b.hasClass(c)&&!b.hasClass("crosstalk-input-bound")){var d=i[c];g(d,a)}})}function g(a,b){var c=h(b).find("script[type='application/json'][data-for='"+e(b.id)+"']"),d=JSON.parse(c[0].innerText),f=a.factory(b,d);h(b).data("crosstalk-instance",f),h(b).addClass("crosstalk-input-bound")}Object.defineProperty(c,"__esModule",{value:!0}),c.register=b;var h=a.jQuery,i={};a.Shiny&&!function(){var b=new a.Shiny.InputBinding,c=a.jQuery;c.extend(b,{find:function(a){return c(a).find(".crosstalk-input")},initialize:function(a){c(a).hasClass("crosstalk-input-bound")||f(a)},getId:function(a){return a.id},getValue:function(a){},setValue:function(a,b){},receiveMessage:function(a,b){},subscribe:function(a,b){c(a).data("crosstalk-instance").resume()},unsubscribe:function(a){c(a).data("crosstalk-instance").suspend()}}),a.Shiny.inputBindings.register(b,"crosstalk.inputBinding")}()}).call(this,"undefined"!=typeof global?global:"undefined"!=typeof self?self:"undefined"!=typeof window?window:{})},{}],7:[function(a,b,c){(function(b){"use strict";function c(a){if(a&&a.__esModule)return a;var b={};if(null!=a)for(var c in a)Object.prototype.hasOwnProperty.call(a,c)&&(b[c]=a[c]);return b.default=a,b}var d=a("./input"),e=c(d),f=a("./filter"),g=b.jQuery;e.register({className:"crosstalk-input-checkboxgroup",factory:function(a,b){var c=new f.FilterHandle(b.group),d=void 0,e=g(a);return e.on("change","input[type='checkbox']",function(){var a=e.find("input[type='checkbox']:checked");0===a.length?(d=null,c.clear()):!function(){var e={};a.each(function(){b.map[this.value].forEach(function(a){e[a]=!0})});var f=Object.keys(e);f.sort(),d=f,c.set(f)}()}),{suspend:function(){c.clear()},resume:function(){d&&c.set(d)}}}})}).call(this,"undefined"!=typeof global?global:"undefined"!=typeof self?self:"undefined"!=typeof window?window:{})},{"./filter":2,"./input":6}],8:[function(a,b,c){(function(b){"use strict";function c(a){if(a&&a.__esModule)return a;var b={};if(null!=a)for(var c in a)Object.prototype.hasOwnProperty.call(a,c)&&(b[c]=a[c]);return b.default=a,b}var d=a("./input"),e=c(d),f=a("./util"),g=c(f),h=a("./filter"),i=b.jQuery;e.register({className:"crosstalk-input-select",factory:function(a,b){var c=[{value:"",label:"(All)"}],d=g.dataframeToD3(b.items),e={options:c.concat(d),valueField:"value",labelField:"label",searchField:"label"},f=i(a).find("select")[0],j=i(f).selectize(e)[0].selectize,k=new h.FilterHandle(b.group),l=void 0;return j.on("change",function(){0===j.items.length?(l=null,k.clear()):!function(){var a={};j.items.forEach(function(c){b.map[c].forEach(function(b){a[b]=!0})});var c=Object.keys(a);c.sort(),l=c,k.set(c)}()}),{suspend:function(){k.clear()},resume:function(){l&&k.set(l)}}}})}).call(this,"undefined"!=typeof global?global:"undefined"!=typeof self?self:"undefined"!=typeof window?window:{})},{"./filter":2,"./input":6,"./util":11}],9:[function(a,b,c){(function(b){"use strict";function c(a){if(a&&a.__esModule)return a;var b={};if(null!=a)for(var c in a)Object.prototype.hasOwnProperty.call(a,c)&&(b[c]=a[c]);return b.default=a,b}function d(a,b){for(var c=a.toString();c.length<b;)c="0"+c;return c}function e(a){return a instanceof Date?a.getUTCFullYear()+"-"+d(a.getUTCMonth()+1,2)+"-"+d(a.getUTCDate(),2):null}var f=function(){function a(a,b){var c=[],d=!0,e=!1,f=void 0;try{for(var g,h=a[Symbol.iterator]();!(d=(g=h.next()).done)&&(c.push(g.value),!b||c.length!==b);d=!0);}catch(a){e=!0,f=a}finally{try{!d&&h.return&&h.return()}finally{if(e)throw f}}return c}return function(b,c){if(Array.isArray(b))return b;if(Symbol.iterator in Object(b))return a(b,c);throw new TypeError("Invalid attempt to destructure non-iterable instance")}}(),g=a("./input"),h=c(g),i=a("./filter"),j=b.jQuery,k=b.strftime;h.register({className:"crosstalk-input-slider",factory:function(a,b){function c(){var a=h.data("ionRangeSlider").result,b=void 0,c=h.data("data-type");return b="date"===c?function(a){return e(new Date(+a))}:"datetime"===c?function(a){return+a/1e3}:function(a){return+a},"double"===h.data("ionRangeSlider").options.type?[b(a.from),b(a.to)]:b(a.from)}var d=new i.FilterHandle(b.group),g={},h=j(a).find("input"),l=h.data("data-type"),m=h.data("time-format"),n=void 0;if("date"===l)n=k.utc(),g.prettify=function(a){return n(m,new Date(a))};else if("datetime"===l){var o=h.data("timezone");n=o?k.timezone(o):k,g.prettify=function(a){return n(m,new Date(a))}}h.ionRangeSlider(g);var p=null;return h.on("change.crosstalkSliderInput",function(a){if(!h.data("updating")&&!h.data("animating")){for(var e=c(),g=f(e,2),i=g[0],j=g[1],k=[],l=0;l<b.values.length;l++){var m=b.values[l];m>=i&&m<=j&&k.push(b.keys[l])}k.sort(),d.set(k),p=k}}),{suspend:function(){d.clear()},resume:function(){p&&d.set(p)}}}})}).call(this,"undefined"!=typeof global?global:"undefined"!=typeof self?self:"undefined"!=typeof window?window:{})},{"./filter":2,"./input":6}],10:[function(a,b,c){"use strict";function d(a){if(a&&a.__esModule)return a;var b={};if(null!=a)for(var c in a)Object.prototype.hasOwnProperty.call(a,c)&&(b[c]=a[c]);return b.default=a,b}function e(a){return a&&a.__esModule?a:{default:a}}function f(a,b){if(!(a instanceof b))throw new TypeError("Cannot call a class as a function")}Object.defineProperty(c,"__esModule",{value:!0}),c.SelectionHandle=void 0;var g=function(){function a(a,b){for(var c=0;c<b.length;c++){var d=b[c];d.enumerable=d.enumerable||!1,d.configurable=!0,"value"in d&&(d.writable=!0),Object.defineProperty(a,d.key,d)}}return function(b,c,d){return c&&a(b.prototype,c),d&&a(b,d),b}}(),h=a("./events"),i=e(h),j=a("./group"),k=e(j),l=a("./util"),m=d(l);c.SelectionHandle=function(){function a(){var b=arguments.length>0&&void 0!==arguments[0]?arguments[0]:null,c=arguments.length>1&&void 0!==arguments[1]?arguments[1]:null;f(this,a),this._eventRelay=new i.default,this._emitter=new m.SubscriptionTracker(this._eventRelay),this._group=null,this._var=null,this._varOnChangeSub=null,this._extraInfo=m.extend({sender:this},c),this.setGroup(b)}return g(a,[{key:"setGroup",value:function(a){var b=this;if(this._group!==a&&(this._group||a)&&(this._var&&(this._var.off("change",this._varOnChangeSub),this._var=null,this._varOnChangeSub=null),this._group=a,a)){this._var=(0,k.default)(a).var("selection");var c=this._var.on("change",function(a){b._eventRelay.trigger("change",a,b)});this._varOnChangeSub=c}}},{key:"_mergeExtraInfo",value:function(a){return m.extend({},this._extraInfo?this._extraInfo:null,a?a:null)}},{key:"set",value:function(a,b){this._var&&this._var.set(a,this._mergeExtraInfo(b))}},{key:"clear",value:function(a){this._var&&this.set(void 0,this._mergeExtraInfo(a))}},{key:"on",value:function(a,b){return this._emitter.on(a,b)}},{key:"off",value:function(a,b){return this._emitter.off(a,b)}},{key:"close",value:function(){this._emitter.removeAllListeners(),this.setGroup(null)}},{key:"value",get:function(){return this._var?this._var.get():null}}]),a}()},{"./events":1,"./group":4,"./util":11}],11:[function(a,b,c){"use strict";function d(a,b){if(!(a instanceof b))throw new TypeError("Cannot call a class as a function")}function e(a){for(var b=arguments.length,c=Array(b>1?b-1:0),d=1;d<b;d++)c[d-1]=arguments[d];for(var e=0;e<c.length;e++){var f=c[e];if("undefined"!=typeof f&&null!==f)for(var g in f)f.hasOwnProperty(g)&&(a[g]=f[g])}return a}function f(a){for(var b=1;b<a.length;b++)if(a[b]<=a[b-1])throw new Error("List is not sorted or contains duplicate")}function g(a,b){var c=0,d=0;a||(a=[]),b||(b=[]);var e=[],g=[];for(f(a),f(b);c<a.length&&d<b.length;)a[c]===b[d]?(c++,d++):a[c]<b[d]?e.push(a[c++]):g.push(b[d++]);return c<a.length&&(e=e.concat(a.slice(c))),d<b.length&&(g=g.concat(b.slice(d))),{removed:e,added:g}}function h(a){var b=[],c=void 0;for(var d in a){if(a.hasOwnProperty(d)&&b.push(d),"object"!==j(a[d])||"undefined"==typeof a[d].length)throw new Error("All fields must be arrays");if("undefined"!=typeof c&&c!==a[d].length)throw new Error("All fields must be arrays of the same length");c=a[d].length}for(var e=[],f=void 0,g=0;g<c;g++){f={};for(var h=0;h<b.length;h++)f[b[h]]=a[b[h]][g];e.push(f)}return e}Object.defineProperty(c,"__esModule",{value:!0});var i=function(){function a(a,b){for(var c=0;c<b.length;c++){var d=b[c];d.enumerable=d.enumerable||!1,d.configurable=!0,"value"in d&&(d.writable=!0),Object.defineProperty(a,d.key,d)}}return function(b,c,d){return c&&a(b.prototype,c),d&&a(b,d),b}}(),j="function"==typeof Symbol&&"symbol"==typeof Symbol.iterator?function(a){return typeof a}:function(a){return a&&"function"==typeof Symbol&&a.constructor===Symbol&&a!==Symbol.prototype?"symbol":typeof a};c.extend=e,c.checkSorted=f,c.diffSortedLists=g,c.dataframeToD3=h;c.SubscriptionTracker=function(){function a(b){d(this,a),this._emitter=b,this._subs={}}return i(a,[{key:"on",value:function(a,b){var c=this._emitter.on(a,b);return this._subs[c]=a,c}},{key:"off",value:function(a,b){var c=this._emitter.off(a,b);return c&&delete this._subs[c],c}},{key:"removeAllListeners",value:function(){var a=this,b=this._subs;this._subs={},Object.keys(b).forEach(function(c){a._emitter.off(b[c],c)})}}]),a}()},{}],12:[function(a,b,c){(function(b){"use strict";function d(a){return a&&a.__esModule?a:{default:a}}function e(a,b){if(!(a instanceof b))throw new TypeError("Cannot call a class as a function")}Object.defineProperty(c,"__esModule",{value:!0});var f="function"==typeof Symbol&&"symbol"==typeof Symbol.iterator?function(a){return typeof a}:function(a){return a&&"function"==typeof Symbol&&a.constructor===Symbol&&a!==Symbol.prototype?"symbol":typeof a},g=function(){function a(a,b){for(var c=0;c<b.length;c++){var d=b[c];d.enumerable=d.enumerable||!1,d.configurable=!0,"value"in d&&(d.writable=!0),Object.defineProperty(a,d.key,d)}}return function(b,c,d){return c&&a(b.prototype,c),d&&a(b,d),b}}(),h=a("./events"),i=d(h),j=function(){function a(b,c,d){e(this,a),this._group=b,this._name=c,this._value=d,this._events=new i.default}return g(a,[{key:"get",value:function(){return this._value}},{key:"set",value:function(a,c){if(this._value!==a){var d=this._value;this._value=a;var e={};if(c&&"object"===("undefined"==typeof c?"undefined":f(c)))for(var g in c)c.hasOwnProperty(g)&&(e[g]=c[g]);e.oldValue=d,e.value=a,this._events.trigger("change",e,this),b.Shiny&&b.Shiny.onInputChange&&b.Shiny.onInputChange(".clientValue-"+(null!==this._group.name?this._group.name+"-":"")+this._name,"undefined"==typeof a?null:a)}}},{key:"on",value:function(a,b){return this._events.on(a,b)}},{key:"off",value:function(a,b){return this._events.off(a,b)}}]),a}();c.default=j}).call(this,"undefined"!=typeof global?global:"undefined"!=typeof self?self:"undefined"!=typeof window?window:{})},{"./events":1}]},{},[5]);
//# sourceMappingURL=crosstalk.min.js.map
```
