## Supplementary material for "Characterizing and inferring quantitative cell cycle phase in single-cell RNA-seq data analysis": Source code of our analysis, including scripts and data necessary to reproduce the work.: datatables.js

```
(function() {

// some helper functions: using a global object DTWidget so that it can be used
// in JS() code, e.g. datatable(options = list(foo = JS('code'))); unlike R's
// dynamic scoping, when 'code' is eval()'ed, JavaScript does not know objects
// from the "parent frame", e.g. JS('DTWidget') will not work unless it was made
// a global object
var DTWidget = {};

// 123456666.7890 -> 123,456,666.7890
var markInterval = function(d, digits, interval, mark, decMark, precision) {
  x = precision ? d.toPrecision(digits) : d.toFixed(digits);
  if (!/^-?[\d.]+$/.test(x)) return x;
  var xv = x.split('.');
  if (xv.length > 2) return x;  // should have at most one decimal point
  xv[0] = xv[0].replace(new RegExp('\\B(?=(\\d{' + interval + '})+(?!\\d))', 'g'), mark);
  return xv.join(decMark);
};

DTWidget.formatCurrency = function(thiz, row, data, col, currency, digits, interval, mark, decMark, before) {
  var d = parseFloat(data[col]);
  if (isNaN(d)) return;
  var res = markInterval(d, digits, interval, mark, decMark);
  res = before ? (/^-/.test(res) ? '-' + currency + res.replace(/^-/, '') : currency + res) :
    res + currency;
  $(thiz.api().cell(row, col).node()).html(res);
};

DTWidget.formatString = function(thiz, row, data, col, prefix, suffix) {
  var d = data[col];
  if (d === null) return;
  var cell = $(thiz.api().cell(row, col).node());
  cell.html(prefix + cell.html() + suffix);
};

DTWidget.formatPercentage = function(thiz, row, data, col, digits, interval, mark, decMark) {
  var d = parseFloat(data[col]);
  if (isNaN(d)) return;
  $(thiz.api().cell(row, col).node())
  .html(markInterval(d * 100, digits, interval, mark, decMark) + '%');
};

DTWidget.formatRound = function(thiz, row, data, col, digits, interval, mark, decMark) {
  var d = parseFloat(data[col]);
  if (isNaN(d)) return;
  $(thiz.api().cell(row, col).node()).html(markInterval(d, digits, interval, mark, decMark));
};

DTWidget.formatSignif = function(thiz, row, data, col, digits, interval, mark, decMark) {
  var d = parseFloat(data[col]);
  if (isNaN(d)) return;
  $(thiz.api().cell(row, col).node())
    .html(markInterval(d, digits, interval, mark, decMark, true));
};

DTWidget.formatDate = function(thiz, row, data, col, method, params) {
  var d = data[col];
  if (d === null) return;
  // (new Date('2015-10-28')).toDateString() may return 2015-10-27 because the
  // actual time created could be like 'Tue Oct 27 2015 19:00:00 GMT-0500 (CDT)',
  // i.e. the date-only string is treated as UTC time instead of local time
  if ((method === 'toDateString' || method === 'toLocaleDateString') && /^\d{4,}\D\d{2}\D\d{2}$/.test(d)) {
    d = d.split(/\D/);
    d = new Date(d[0], d[1] - 1, d[2]);
  } else {
    d = new Date(d);
  }
  $(thiz.api().cell(row, col).node()).html(d[method].apply(d, params));
};

window.DTWidget = DTWidget;

var transposeArray2D = function(a) {
  return a.length === 0 ? a : HTMLWidgets.transposeArray2D(a);
};

var crosstalkPluginsInstalled = false;

function maybeInstallCrosstalkPlugins() {
  if (crosstalkPluginsInstalled)
    return;
  crosstalkPluginsInstalled = true;

  $.fn.dataTable.ext.afnFiltering.push(
    function(oSettings, aData, iDataIndex) {
      var ctfilter = oSettings.nTable.ctfilter;
      if (ctfilter && !ctfilter[iDataIndex])
        return false;

      var ctselect = oSettings.nTable.ctselect;
      if (ctselect && !ctselect[iDataIndex])
        return false;

      return true;
    }
  );
}

HTMLWidgets.widget({
  name: "datatables",
  type: "output",
  renderOnNullValue: true,
  initialize: function(el, width, height) {
    $(el).html('&nbsp;');
    return {
      data: null,
      ctfilterHandle: new crosstalk.FilterHandle(),
      ctfilterSubscription: null,
      ctselectHandle: new crosstalk.SelectionHandle(),
      ctselectSubscription: null
    };
  },
  renderValue: function(el, data, instance) {
    if (el.offsetWidth === 0 || el.offsetHeight === 0) {
      instance.data = data;
      return;
    }
    instance.data = null;
    var $el = $(el);
    $el.empty();

    if (data === null) {
      $el.append('&nbsp;');
      // clear previous Shiny inputs (if any)
      for (var i in instance.clearInputs) instance.clearInputs[i]();
      instance.clearInputs = {};
      return;
    }

    var crosstalkOptions = data.crosstalkOptions;
    if (!crosstalkOptions) crosstalkOptions = {
      'key': null, 'group': null
    };
    if (crosstalkOptions.group) {
      maybeInstallCrosstalkPlugins();
      instance.ctfilterHandle.setGroup(crosstalkOptions.group);
      instance.ctselectHandle.setGroup(crosstalkOptions.group);
    }

    // If we are in a flexdashboard scroll layout then we:
    //  (a) Always want to use pagination (otherwise we'll have
    //      a "double scroll bar" effect on the phone); and
    //  (b) Never want to fill the container (we want the pagination
    //      level to determine the size of the container)
    if (window.FlexDashboard && !window.FlexDashboard.isFillPage()) {
      data.options.bPaginate = true;
      data.fillContainer = false;
    }

    // if we are in the viewer then we always want to fillContainer and
    // and autoHideNavigation (unless the user has explicitly set these)
    if (window.HTMLWidgets.viewerMode) {
      if (!data.hasOwnProperty("fillContainer"))
        data.fillContainer = true;
      if (!data.hasOwnProperty("autoHideNavigation"))
        data.autoHideNavigation = true;
    }

    // propagate fillContainer to instance (so we have it in resize)
    instance.fillContainer = data.fillContainer;

    var cells = data.data;

    if (cells instanceof Array) cells = transposeArray2D(cells);

    $el.append(data.container);
    var $table = $el.find('table');
    if (data.class) $table.addClass(data.class);
    if (data.caption) $table.prepend(data.caption);

    if (!data.selection) data.selection = {
      mode: 'none', selected: null, target: 'row'
    };
    if (HTMLWidgets.shinyMode && data.selection.mode !== 'none' &&
        data.selection.target === 'row+column') {
      if ($table.children('tfoot').length === 0) {
        $table.append($('<tfoot>'));
        $table.find('thead tr').clone().appendTo($table.find('tfoot'));
      }
    }

    // column filters
    var filterRow;
    switch (data.filter) {
      case 'top':
        $table.children('thead').append(data.filterHTML);
        filterRow = $table.find('thead tr:last td');
        break;
      case 'bottom':
        if ($table.children('tfoot').length === 0) {
          $table.append($('<tfoot>'));
        }
        $table.children('tfoot').prepend(data.filterHTML);
        filterRow = $table.find('tfoot tr:first td');
        break;
    }

    var options = { searchDelay: 1000 };
    if (cells !== null) $.extend(options, {
      data: cells
    });

    // options for fillContainer
    var bootstrapActive = typeof($.fn.popover) != 'undefined';
    if (instance.fillContainer) {

      // force scrollX/scrollY and turn off autoWidth
      options.scrollX = true;
      options.scrollY = "100px"; // can be any value, we'll adjust below

      // if we aren't paginating then move around the info/filter controls
      // to save space at the bottom and rephrase the info callback
      if (data.options.bPaginate === false) {

        // we know how to do this cleanly for bootstrap, not so much
        // for other themes/layouts
        if (bootstrapActive) {
          options.dom = "<'row'<'col-sm-4'i><'col-sm-8'f>>" +
                        "<'row'<'col-sm-12'tr>>";
        }

        options.fnInfoCallback = function(oSettings, iStart, iEnd,
                                           iMax, iTotal, sPre) {
          return Number(iTotal).toLocaleString() + " records";
        };
      }
    }

    // auto hide navigation if requested
    if (data.autoHideNavigation === true) {
      if (bootstrapActive && data.options.bPaginate !== false) {
        // strip all nav if length >= cells
        if ((cells instanceof Array) && data.options.iDisplayLength >= cells.length)
          options.dom = "<'row'<'col-sm-12'tr>>";
        // alternatively lean things out for flexdashboard mobile portrait
        else if (window.FlexDashboard && window.FlexDashboard.isMobilePhone())
          options.dom = "<'row'<'col-sm-12'f>>" +
                        "<'row'<'col-sm-12'tr>>"  +
                        "<'row'<'col-sm-12'p>>";
      }
    }

    $.extend(true, options, data.options || {});

    var searchCols = options.searchCols;
    if (searchCols) {
      searchCols = searchCols.map(function(x) {
        return x === null ? '' : x.search;
      });
      // FIXME: this means I don't respect the escapeRegex setting
      delete options.searchCols;
    }

    // server-side processing?
    var server = options.serverSide === true;

    // use the dataSrc function to pre-process JSON data returned from R
    var DT_rows_all = [], DT_rows_current = [];
    if (server && HTMLWidgets.shinyMode && typeof options.ajax === 'object' &&
        /^session\/[\da-z]+\/dataobj/.test(options.ajax.url) && !options.ajax.dataSrc) {
      options.ajax.dataSrc = function(json) {
        DT_rows_all = $.makeArray(json.DT_rows_all);
        DT_rows_current = $.makeArray(json.DT_rows_current);
        var data = json.data;
        if (!colReorderEnabled()) return data;
        var table = $table.DataTable(), order = table.colReorder.order(), flag = true, i, j, row;
        for (i = 0; i < order.length; ++i) if (order[i] !== i) flag = false;
        if (flag) return data;
        for (i = 0; i < data.length; ++i) {
          row = data[i].slice();
          for (j = 0; j < order.length; ++j) data[i][j] = row[order[j]];
        }
        return data;
      };
    }

    var thiz = this;
    if (instance.fillContainer) $table.on('init.dt', function(e) {
      thiz.fillAvailableHeight(el, $(el).innerHeight());
    });
    // If the page contains serveral datatables and one of which enables colReorder,
    // the table.colReorder.order() function will exist but throws error when called.
    // So it seems like the only way to know if colReorder is enabled or not is to
    // check the options.
    var colReorderEnabled = function() { return "colReorder" in options; };
    var table = $table.DataTable(options);
    $el.data('datatable', table);

    // Unregister previous Crosstalk event subscriptions, if they exist
    if (instance.ctfilterSubscription) {
      instance.ctfilterHandle.off("change", instance.ctfilterSubscription);
      instance.ctfilterSubscription = null;
    }
    if (instance.ctselectSubscription) {
      instance.ctselectHandle.off("change", instance.ctselectSubscription);
      instance.ctselectSubscription = null;
    }

    if (!crosstalkOptions.group) {
      $table[0].ctfilter = null;
      $table[0].ctselect = null;
    } else {
      var key = crosstalkOptions.key;
      function keysToMatches(keys) {
        if (!keys) {
          return null;
        } else {
          var selectedKeys = {};
          for (var i = 0; i < keys.length; i++) {
            selectedKeys[keys[i]] = true;
          }
          var matches = {};
          for (var j = 0; j < key.length; j++) {
            if (selectedKeys[key[j]])
              matches[j] = true;
          }
          return matches;
        }
      }

      function applyCrosstalkFilter(e) {
        $table[0].ctfilter = keysToMatches(e.value);
        table.draw();
      }
      instance.ctfilterSubscription = instance.ctfilterHandle.on("change", applyCrosstalkFilter);
      applyCrosstalkFilter({value: instance.ctfilterHandle.filteredKeys});

      function applyCrosstalkSelection(e) {
        if (e.sender !== instance.ctselectHandle) {
          table
            .rows('.' + selClass, {search: 'applied'})
            .nodes()
            .to$()
            .removeClass(selClass);
          if (selectedRows)
            changeInput('rows_selected', selectedRows(), void 0, true);
        }

        if (e.sender !== instance.ctselectHandle && e.value && e.value.length) {
          var matches = keysToMatches(e.value);

          // persistent selection with plotly (& leaflet)
          var ctOpts = crosstalk.var("plotlyCrosstalkOpts").get() || {};
          if (ctOpts.persistent === true) {
            var matches = $.extend(matches, $table[0].ctselect);
          }

          $table[0].ctselect = matches;
          table.draw();
        } else {
          if ($table[0].ctselect) {
            $table[0].ctselect = null;
            table.draw();
          }
        }
      }
      instance.ctselectSubscription = instance.ctselectHandle.on("change", applyCrosstalkSelection);
      // TODO: This next line doesn't seem to work when renderDataTable is used
      applyCrosstalkSelection({value: instance.ctselectHandle.value});
    }

    var inArray = function(val, array) {
      return $.inArray(val, $.makeArray(array)) > -1;
    };

    // encode + to %2B when searching in the table on server side, because
    // shiny::parseQueryString() treats + as spaces, and DataTables does not
    // encode + to %2B (or % to %25) when sending the request
    var encode_plus = function(x) {
      return server ? x.replace(/%/g, '%25').replace(/\+/g, '%2B') : x;
    };

    // search the i-th column
    var searchColumn = function(i, value) {
      var regex = false, ci = true;
      if (options.search) {
        regex = options.search.regex,
        ci = options.search.caseInsensitive !== false;
      }
      return table.column(i).search(encode_plus(value), regex, !regex, ci);
    };

    if (data.filter !== 'none') {

      filterRow.each(function(i, td) {

        var $td = $(td), type = $td.data('type'), filter;
        var $input = $td.children('div').first().children('input');
        $input.prop('disabled', !table.settings()[0].aoColumns[i].bSearchable || type === 'disabled');
        $input.on('input blur', function() {
          $input.next('span').toggle(Boolean($input.val()));
        });
        // Bootstrap sets pointer-events to none and we won't be able to click
        // the clear button
        $input.next('span').css('pointer-events', 'auto').hide().click(function() {
          $(this).hide().prev('input').val('').trigger('input').focus();
        });
        var searchCol;  // search string for this column
        if (searchCols && searchCols[i]) {
          searchCol = searchCols[i];
          $input.val(searchCol).trigger('input');
        }
        var $x = $td.children('div').last();

        // remove the overflow: hidden attribute of the scrollHead
        // (otherwise the scrolling table body obscures the filters)
        var scrollHead = $(el).find('.dataTables_scrollHead,.dataTables_scrollFoot');
        var cssOverflow = scrollHead.css('overflow');
        if (cssOverflow === 'hidden') {
          $x.on('show hide', function(e) {
            scrollHead.css('overflow', e.type === 'show' ? '' : cssOverflow);
          });
          $x.css('z-index', 25);
        }

        if (inArray(type, ['factor', 'logical'])) {
          $input.on({
            click: function() {
              $input.parent().hide(); $x.show().trigger('show'); filter[0].selectize.focus();
            },
            input: function() {
              if ($input.val() === '') filter[0].selectize.setValue([]);
            }
          });
          var $input2 = $x.children('select');
          filter = $input2.selectize({
            options: $input2.data('options').map(function(v, i) {
              return ({text: v, value: v});
            }),
            plugins: ['remove_button'],
            hideSelected: true,
            onChange: function(value) {
              if (value === null) value = []; // compatibility with jQuery 3.0
              $input.val(value.length ? JSON.stringify(value) : '');
              if (value.length) $input.trigger('input');
              $input.attr('title', $input.val());
              if (server) {
                table.column(i).search(value.length ? encode_plus(JSON.stringify(value)) : '').draw();
                return;
              }
              // turn off filter if nothing selected
              $td.data('filter', value.length > 0);
              table.draw();  // redraw table, and filters will be applied
            }
          });
          if (searchCol) filter[0].selectize.setValue(JSON.parse(searchCol));
          filter[0].selectize.on('blur', function() {
            $x.hide().trigger('hide'); $input.parent().show(); $input.trigger('blur');
          });
          filter.next('div').css('margin-bottom', 'auto');
        } else if (type === 'character') {
          var fun = function() {
            searchColumn(i, $input.val()).draw();
          };
          if (server) {
            fun = $.fn.dataTable.util.throttle(fun, options.searchDelay);
          }
          $input.on('input', fun);
        } else if (inArray(type, ['number', 'integer', 'date', 'time'])) {
          var $x0 = $x;
          $x = $x0.children('div').first();
          $x0.css({
            'background-color': '#fff',
            'border': '1px #ddd solid',
            'border-radius': '4px',
            'padding': '20px 20px 10px 20px'
          });
          var $spans = $x0.children('span').css({
            'margin-top': '10px',
            'white-space': 'nowrap'
          });
          var $span1 = $spans.first(), $span2 = $spans.last();
          var r1 = +$x.data('min'), r2 = +$x.data('max');
          // when the numbers are too small or have many decimal places, the
          // slider may have numeric precision problems (#150)
          var scale = Math.pow(10, Math.max(0, +$x.data('scale') || 0));
          r1 = Math.round(r1 * scale); r2 = Math.round(r2 * scale);
          var scaleBack = function(x, scale) {
            if (scale === 1) return x;
            var d = Math.round(Math.log(scale) / Math.log(10));
            // to avoid problems like 3.423/100 -> 0.034230000000000003
            return (x / scale).toFixed(d);
          };
          $input.on({
            focus: function() {
              $x0.show().trigger('show');
              // first, make sure the slider div leaves at least 20px between
              // the two (slider value) span's
              $x0.width(Math.max(160, $span1.outerWidth() + $span2.outerWidth() + 20));
              // then, if the input is really wide, make the slider the same
              // width as the input
              if ($x0.outerWidth() < $input.outerWidth()) {
                $x0.outerWidth($input.outerWidth());
              }
              // make sure the slider div does not reach beyond the right margin
              if ($(window).width() < $x0.offset().left + $x0.width()) {
                $x0.offset({
                  'left': $input.offset().left + $input.outerWidth() - $x0.outerWidth()
                });
              }
            },
            blur: function() {
              $x0.hide().trigger('hide');
            },
            input: function() {
              if ($input.val() === '') filter.val([r1, r2]);
            },
            change: function() {
              var v = $input.val().replace(/\s/g, '');
              if (v === '') return;
              v = v.split('...');
              if (v.length !== 2) {
                $input.parent().addClass('has-error');
                return;
              }
              if (v[0] === '') v[0] = r1;
              if (v[1] === '') v[1] = r2;
              $input.parent().removeClass('has-error');
              // treat date as UTC time at midnight
              var strTime = function(x) {
                var s = type === 'date' ? 'T00:00:00Z' : '';
                var t = new Date(x + s).getTime();
                // add 10 minutes to date since it does not hurt the date, and
                // it helps avoid the tricky floating point arithmetic problems,
                // e.g. sometimes the date may be a few milliseconds earlier
                // than the midnight due to precision problems in noUiSlider
                return type === 'date' ? t + 3600000 : t;
              };
              if (inArray(type, ['date', 'time'])) {
                v[0] = strTime(v[0]);
                v[1] = strTime(v[1]);
              }
              if (v[0] != r1) v[0] *= scale;
              if (v[1] != r2) v[1] *= scale;
              filter.val(v);
            }
          });
          var formatDate = function(d, isoFmt) {
            d = scaleBack(d, scale);
            if (type === 'number') return d;
            if (type === 'integer') return parseInt(d);
            var x = new Date(+d);
            var fmt = ('filterDateFmt' in data) ? data.filterDateFmt[i] : undefined;
            if (fmt !== undefined && isoFmt === false) return x[fmt.method].apply(x, fmt.params);
            if (type === 'date') {
              var pad0 = function(x) {
                return ('0' + x).substr(-2, 2);
              };
              return x.getUTCFullYear() + '-' + pad0(1 + x.getUTCMonth())
                      + '-' + pad0(x.getUTCDate());
            } else {
              return x.toISOString();
            }
          };
          var opts = type === 'date' ? { step: 60 * 60 * 1000 } :
                     type === 'integer' ? { step: 1 } : {};
          filter = $x.noUiSlider($.extend({
            start: [r1, r2],
            range: {min: r1, max: r2},
            connect: true
          }, opts));
          if (scale > 1) (function() {
            var t1 = r1, t2 = r2;
            var val = filter.val();
            while (val[0] > r1 || val[1] < r2) {
              if (val[0] > r1) {
                t1 -= val[0] - r1;
              }
              if (val[1] < r2) {
                t2 += r2 - val[1];
              }
              filter = $x.noUiSlider($.extend({
                start: [t1, t2],
                range: {min: t1, max: t2},
                connect: true
              }, opts), true);
              val = filter.val();
            }
            r1  = t1; r2 = t2;
          })();
          var updateSliderText = function(v1, v2) {
            $span1.text(formatDate(v1, false)); $span2.text(formatDate(v2, false));
          };
          updateSliderText(r1, r2);
          var updateSlider = function(e) {
            var val = filter.val();
            // turn off filter if in full range
            $td.data('filter', val[0] > r1 || val[1] < r2);
            var v1 = formatDate(val[0]), v2 = formatDate(val[1]), ival;
            if ($td.data('filter')) {
              ival = v1 + ' ... ' + v2;
              $input.attr('title', ival).val(ival).trigger('input');
            } else {
              $input.attr('title', '').val('');
            }
            updateSliderText(val[0], val[1]);
            if (e.type === 'slide') return;  // no searching when sliding only
            if (server) {
              table.column(i).search($td.data('filter') ? ival : '').draw();
              return;
            }
            table.draw();
          };
          filter.on({
            set: updateSlider,
            slide: updateSlider
          });
        }

        // server-side processing will be handled by R (or whatever server
        // language you use); the following code is only needed for client-side
        // processing
        if (server) {
          // if a search string has been pre-set, search now
          if (searchCol) searchColumn(i, searchCol).draw();
          return;
        }

        var customFilter = function(settings, data, dataIndex) {
          // there is no way to attach a search function to a specific table,
          // and we need to make sure a global search function is not applied to
          // all tables (i.e. a range filter in a previous table should not be
          // applied to the current table); we use the settings object to
          // determine if we want to perform searching on the current table,
          // since settings.sTableId will be different to different tables
          if (table.settings()[0] !== settings) return true;
          // no filter on this column or no need to filter this column
          if (typeof filter === 'undefined' || !$td.data('filter')) return true;

          var r = filter.val(), v, r0, r1;
          var i_data = function(i) {
            if (!colReorderEnabled()) return i;
            var order = table.colReorder.order(), k;
            for (k = 0; k < order.length; ++k) if (order[k] === i) return k;
            return i; // in theory it will never be here...
          }
          v = data[i_data(i)];
          if (type === 'number' || type === 'integer') {
            v = parseFloat(v);
            // how to handle NaN? currently exclude these rows
            if (isNaN(v)) return(false);
            r0 = parseFloat(scaleBack(r[0], scale))
            r1 = parseFloat(scaleBack(r[1], scale));
            if (v >= r0 && v <= r1) return true;
          } else if (type === 'date' || type === 'time') {
            v = new Date(v);
            r0 = new Date(r[0] / scale); r1 = new Date(r[1] / scale);
            if (v >= r0 && v <= r1) return true;
          } else if (type === 'factor') {
            if (r.length === 0 || inArray(v, r)) return true;
          } else if (type === 'logical') {
            if (r.length === 0) return true;
            if (inArray(v === '' ? 'na' : v, r)) return true;
          }
          return false;
        };

        $.fn.dataTable.ext.search.push(customFilter);

        // search for the preset search strings if it is non-empty
        if (searchCol) {
          if (inArray(type, ['factor', 'logical'])) {
            filter[0].selectize.setValue(JSON.parse(searchCol));
          } else if (type === 'character') {
            $input.trigger('input');
          } else if (inArray(type, ['number', 'integer', 'date', 'time'])) {
            $input.trigger('change');
          }
        }

      });

    }

    // highlight search keywords
    var highlight = function() {
      var body = $(table.table().body());
      // removing the old highlighting first
      body.unhighlight();

      // don't highlight the "not found" row, so we get the rows using the api
      if (table.rows({ filter: 'applied' }).data().length === 0) return;
      // highlight gloal search keywords
      body.highlight($.trim(table.search()).split(/\s+/));
      // then highlight keywords from individual column filters
      if (filterRow) filterRow.each(function(i, td) {
        var $td = $(td), type = $td.data('type');
        if (type !== 'character') return;
        var $input = $td.children('div').first().children('input');
        var column = table.column(i).nodes().to$(),
            val = $.trim($input.val());
        if (type !== 'character' || val === '') return;
        column.highlight(val.split(/\s+/));
      });
    };

    if (options.searchHighlight) {
      table
      .on('draw.dt.dth column-visibility.dt.dth column-reorder.dt.dth', highlight)
      .on('destroy', function() {
        // remove event handler
        table.off('draw.dt.dth column-visibility.dt.dth column-reorder.dt.dth');
      });

      // initial highlight for state saved conditions and initial states
      highlight();
    }

    // run the callback function on the table instance
    if (typeof data.callback === 'function') data.callback(table);

    // double click to edit the cell
    if (data.editable) table.on('dblclick.dt', 'tbody td', function() {
      var $input = $('<input type="text">');
      var $this = $(this), value = table.cell(this).data(), html = $this.html();
      var changed = false;
      $input.val(value);
      $this.empty().append($input);
      $input.css('width', '100%').focus().on('change', function() {
        changed = true;
        var valueNew = $input.val();
        if (valueNew != value) {
          table.cell($this).data(valueNew);
          if (HTMLWidgets.shinyMode) changeInput('cell_edit', cellInfo($this));
          // for server-side processing, users have to call replaceData() to update the table
          if (!server) table.draw(false);
        } else {
          $this.html(html);
        }
        $input.remove();
      }).on('blur', function() {
        if (!changed) $input.trigger('change');
      });
    });

    // interaction with shiny
    if (!HTMLWidgets.shinyMode && !crosstalkOptions.group) return;

    var methods = {};
    var shinyData = {};

    methods.updateCaption = function(caption) {
      if (!caption) return;
      $table.children('caption').replaceWith(caption);
    }

    // register clear functions to remove input values when the table is removed
    instance.clearInputs = {};

    var changeInput = function(id, value, type, noCrosstalk) {
      var event = id;
      id = el.id + '_' + id;
      if (type) id = id + ':' + type;
      // do not update if the new value is the same as old value
      if (shinyData.hasOwnProperty(id) && shinyData[id] === JSON.stringify(value))
        return;
      shinyData[id] = JSON.stringify(value);
      if (HTMLWidgets.shinyMode) {
        Shiny.onInputChange(id, value);
        if (!instance.clearInputs[id]) instance.clearInputs[id] = function() {
          Shiny.onInputChange(id, null);
        }
      }

      // HACK
      if (event === "rows_selected" && !noCrosstalk) {
        if (crosstalkOptions.group) {
          var keys = crosstalkOptions.key;
          var selectedKeys = null;
          if (value) {
            selectedKeys = [];
            for (var i = 0; i < value.length; i++) {
              // The value array's contents use 1-based row numbers, so we must
              // convert to 0-based before indexing into the keys array.
              selectedKeys.push(keys[value[i] - 1]);
            }
          }
          instance.ctselectHandle.set(selectedKeys);
        }
      }
    };

    var addOne = function(x) {
      return x.map(function(i) { return 1 + i; });
    };

    var unique = function(x) {
      var ux = [];
      $.each(x, function(i, el){
        if ($.inArray(el, ux) === -1) ux.push(el);
      });
      return ux;
    }

    // change the row index of a cell
    var tweakCellIndex = function(cell) {
      var info = cell.index();
      if (server) {
        info.row = DT_rows_current[info.row];
      } else {
        info.row += 1;
      }
      return {row: info.row, col: info.column};
    }

    var selMode = data.selection.mode, selTarget = data.selection.target;
    if (inArray(selMode, ['single', 'multiple'])) {
      var selClass = data.style === 'bootstrap' ? 'active' : 'selected';
      var selected = data.selection.selected, selected1, selected2;
      // selected1: row indices; selected2: column indices
      if (selected === null) {
        selected1 = selected2 = [];
      } else if (selTarget === 'row') {
        selected1 = $.makeArray(selected);
      } else if (selTarget === 'column') {
        selected2 = $.makeArray(selected);
      } else if (selTarget === 'row+column') {
        selected1 = $.makeArray(selected.rows);
        selected2 = $.makeArray(selected.cols);
      }

      // After users reorder the rows or filter the table, we cannot use the table index
      // directly. Instead, we need this function to find out the rows between the two clicks.
      // If user filter the table again between the start click and the end click, the behavior
      // would be undefined, but it should not be a problem.
      var shiftSelRowsIndex = function(start, end) {
        var indexes = server ? DT_rows_all : table.rows({ search: 'applied' }).indexes().toArray();
        start = indexes.indexOf(start); end = indexes.indexOf(end);
        // if start is larger than end, we need to swap
        if (start > end) {
          var tmp = end; end = start; start = tmp;
        }
        return indexes.slice(start, end + 1);
      }

      var serverRowIndex = function(clientRowIndex) {
        return server ? DT_rows_current[clientRowIndex] : clientRowIndex + 1;
      }

      // row, column, or cell selection
      var lastClickedRow;
      if (inArray(selTarget, ['row', 'row+column'])) {
        var selectedRows = function() {
          var rows = table.rows('.' + selClass);
          var idx = rows.indexes().toArray();
          if (!server) return addOne(idx);
          idx = idx.map(function(i) {
            return DT_rows_current[i];
          });
          selected1 = selMode === 'multiple' ? unique(selected1.concat(idx)) : idx;
          return selected1;
        }
        table.on('mousedown.dt', 'tbody tr', function(e) {
          var $this = $(this), thisRow = table.row(this);
          if (selMode === 'multiple') {
            if (e.shiftKey && lastClickedRow !== undefined) {
              // select or de-select depends on the last clicked row's status
              var flagSel = !$this.hasClass(selClass);
              var crtClickedRow = serverRowIndex(thisRow.index());
              if (server) {
                var rowsIndex = shiftSelRowsIndex(lastClickedRow, crtClickedRow);
                // update current page's selClass
                rowsIndex.map(function(i) {
                  var rowIndex = DT_rows_current.indexOf(i);
                  if (rowIndex >= 0) {
                    var row = table.row(rowIndex).nodes().to$();
                    var flagRowSel = !row.hasClass(selClass);
                    if (flagSel === flagRowSel) row.toggleClass(selClass);
                  }
                });
                // update selected1
                if (flagSel) {
                  selected1 = unique(selected1.concat(rowsIndex));
                } else {
                  selected1 = selected1.filter(function(index) {
                    return !inArray(index, rowsIndex);
                  });
                }
              } else {
                // js starts from 0
                shiftSelRowsIndex(lastClickedRow - 1, crtClickedRow - 1).map(function(value) {
                  var row = table.row(value).nodes().to$();
                  var flagRowSel = !row.hasClass(selClass);
                  if (flagSel === flagRowSel) row.toggleClass(selClass);
                });
              }
              e.preventDefault();
            } else {
              $this.toggleClass(selClass);
            }
          } else {
            if ($this.hasClass(selClass)) {
              $this.removeClass(selClass);
            } else {
              table.$('tr.' + selClass).removeClass(selClass);
              $this.addClass(selClass);
            }
          }
          if (server && !$this.hasClass(selClass)) {
            var id = DT_rows_current[thisRow.index()];
            // remove id from selected1 since its class .selected has been removed
            if (inArray(id, selected1)) selected1.splice($.inArray(id, selected1), 1);
          }
          changeInput('rows_selected', selectedRows());
          changeInput('row_last_clicked', serverRowIndex(thisRow.index()));
          lastClickedRow = serverRowIndex(thisRow.index());
        });
        changeInput('rows_selected', selected1);
        var selectRows = function() {
          table.$('tr.' + selClass).removeClass(selClass);
          if (selected1.length === 0) return;
          if (server) {
            table.rows({page: 'current'}).every(function() {
              if (inArray(DT_rows_current[this.index()], selected1)) {
                $(this.node()).addClass(selClass);
              }
            });
          } else {
            var selected0 = selected1.map(function(i) { return i - 1; });
            $(table.rows(selected0).nodes()).addClass(selClass);
          }
        }
        selectRows();  // in case users have specified pre-selected rows
        // restore selected rows after the table is redrawn (e.g. sort/search/page);
        // client-side tables will preserve the selections automatically; for
        // server-side tables, we have to *real* row indices are in `selected1`
        if (server) table.on('draw.dt', selectRows);
        methods.selectRows = function(selected) {
          selected1 = $.makeArray(selected);
          selectRows();
          changeInput('rows_selected', selected1);
        }
      }

      if (inArray(selTarget, ['column', 'row+column'])) {
        if (selTarget === 'row+column') {
          $(table.columns().footer()).css('cursor', 'pointer');
        }
        table.on('click.dt', selTarget === 'column' ? 'tbody td' : 'tfoot tr th', function() {
          var colIdx = selTarget === 'column' ? table.cell(this).index().column :
              $.inArray(this, table.columns().footer()),
              thisCol = $(table.column(colIdx).nodes());
          if (colIdx === -1) return;
          if (thisCol.hasClass(selClass)) {
            thisCol.removeClass(selClass);
            selected2.splice($.inArray(colIdx, selected2), 1);
          } else {
            if (selMode === 'single') $(table.cells().nodes()).removeClass(selClass);
            thisCol.addClass(selClass);
            selected2 = selMode === 'single' ? [colIdx] : unique(selected2.concat([colIdx]));
          }
          changeInput('columns_selected', selected2);
        });
        changeInput('columns_selected', selected2);
        var selectCols = function() {
          table.columns().nodes().flatten().to$().removeClass(selClass);
          if (selected2.length > 0)
            table.columns(selected2).nodes().flatten().to$().addClass(selClass);
        }
        selectCols();  // in case users have specified pre-selected columns
        if (server) table.on('draw.dt', selectCols);
        methods.selectColumns = function(selected) {
          selected2 = $.makeArray(selected);
          selectCols();
          changeInput('columns_selected', selected2);
        }
      }

      if (selTarget === 'cell') {
        var selected3;
        if (selected === null) {
          selected3 = [];
        } else {
          selected3 = selected;
        }
        var findIndex = function(ij) {
          for (var i = 0; i < selected3.length; i++) {
            if (ij[0] === selected3[i][0] && ij[1] === selected3[i][1]) return i;
          }
          return -1;
        }
        table.on('click.dt', 'tbody td', function() {
          var $this = $(this), info = tweakCellIndex(table.cell(this));
          if ($this.hasClass(selClass)) {
            $this.removeClass(selClass);
            selected3.splice(findIndex([info.row, info.col]), 1);
          } else {
            if (selMode === 'single') $(table.cells().nodes()).removeClass(selClass);
            $this.addClass(selClass);
            selected3 = selMode === 'single' ? [[info.row, info.col]] :
              unique(selected3.concat([[info.row, info.col]]));
          }
          changeInput('cells_selected', transposeArray2D(selected3), 'shiny.matrix');
        });
        changeInput('cells_selected', transposeArray2D(selected3), 'shiny.matrix');
        var selectCells = function() {
          table.$('td.' + selClass).removeClass(selClass);
          if (selected3.length === 0) return;
          if (server) {
            table.cells({page: 'current'}).every(function() {
              var info = tweakCellIndex(this);
              if (findIndex([info.row, info.col], selected3) > -1)
                $(this.node()).addClass(selClass);
            });
          } else {
            selected3.map(function(ij) {
              $(table.cell(ij[0] - 1, ij[1]).node()).addClass(selClass);
            });
          }
        };
        selectCells();  // in case users have specified pre-selected columns
        if (server) table.on('draw.dt', selectCells);
        methods.selectCells = function(selected) {
          selected3 = selected ? selected : [];
          selectCells();
          changeInput('cells_selected', transposeArray2D(selected3), 'shiny.matrix');
        }
      }
    }

    // expose some table info to Shiny
    var updateTableInfo = function(e, settings) {
      // TODO: is anyone interested in the page info?
      // changeInput('page_info', table.page.info());
      var updateRowInfo = function(id, modifier) {
        var idx;
        if (server) {
          idx = modifier.page === 'current' ? DT_rows_current : DT_rows_all;
        } else {
          var rows = table.rows($.extend({
            search: 'applied',
            page: 'all'
          }, modifier));
          idx = addOne(rows.indexes().toArray());
        }
        changeInput('rows' + '_' + id, idx);
      };
      updateRowInfo('current', {page: 'current'});
      updateRowInfo('all', {});
    }
    table.on('draw.dt', updateTableInfo);
    updateTableInfo();

    // state info
    table.on('draw.dt column-visibility.dt', function() {
      changeInput('state', table.state());
    });
    changeInput('state', table.state());

    // search info
    var updateSearchInfo = function() {
      changeInput('search', table.search());
      if (filterRow) changeInput('search_columns', filterRow.toArray().map(function(td) {
        return $(td).find('input').first().val();
      }));
    }
    table.on('draw.dt', updateSearchInfo);
    updateSearchInfo();

    var cellInfo = function(thiz) {
      var info = tweakCellIndex(table.cell(thiz));
      info.value = table.cell(thiz).data();
      return info;
    }
    // the current cell clicked on
    table.on('click.dt', 'tbody td', function() {
      changeInput('cell_clicked', cellInfo(this));
    })
    changeInput('cell_clicked', {});

    // do not trigger table selection when clicking on links unless they have classes
    table.on('click.dt', 'tbody td a', function(e) {
      if (this.className === '') e.stopPropagation();
    });

    methods.addRow = function(data, rowname) {
      var data0 = table.row(0).data(), n = data0.length, d = n - data.length;
      if (d === 1) {
        data = rowname.concat(data)
      } else if (d !== 0) {
        console.log(data);
        console.log(data0);
        throw 'New data must be of the same length as current data (' + n + ')';
      };
      table.row.add(data).draw();
    }

    methods.updateSearch = function(keywords) {
      if (keywords.global !== null)
        $(table.table().container()).find('input[type=search]').first()
             .val(keywords.global).trigger('input');
      var columns = keywords.columns;
      if (!filterRow || columns === null) return;
      filterRow.toArray().map(function(td, i) {
        var v = typeof columns === 'string' ? columns : columns[i];
        if (typeof v === 'undefined') {
          console.log('The search keyword for column ' + i + ' is undefined')
          return;
        }
        $(td).find('input').first().val(v);
        searchColumn(i, v);
      });
      table.draw();
    }

    methods.hideCols = function(hide, reset) {
      if (reset) table.columns().visible(true, false);
      table.columns(hide).visible(false);
    }

    methods.showCols = function(show, reset) {
      if (reset) table.columns().visible(false, false);
      table.columns(show).visible(true);
    }

    methods.colReorder = function(order, origOrder) {
      table.colReorder.order(order, origOrder);
    }

    methods.selectPage = function(page) {
      if (table.page.info().pages < page || page < 1) {
        throw 'Selected page is out of range';
      };
      table.page(page - 1).draw(false);
    }

    methods.reloadData = function(resetPaging, clearSelection) {
      // empty selections first if necessary
      if (methods.selectRows && inArray('row', clearSelection)) methods.selectRows([]);
      if (methods.selectColumns && inArray('column', clearSelection)) methods.selectColumns([]);
      if (methods.selectCells && inArray('cell', clearSelection)) methods.selectCells([]);
      table.ajax.reload(null, resetPaging);
    }

    table.shinyMethods = methods;
  },
  resize: function(el, width, height, instance) {
    if (instance.data) this.renderValue(el, instance.data, instance);

    // dynamically adjust height if fillContainer = TRUE
    if (instance.fillContainer)
      this.fillAvailableHeight(el, height);

    this.adjustWidth(el);
  },

  // dynamically set the scroll body to fill available height
  // (used with fillContainer = TRUE)
  fillAvailableHeight: function(el, availableHeight) {

    // see how much of the table is occupied by header/footer elements
    // and use that to compute a target scroll body height
    var dtWrapper = $(el).find('div.dataTables_wrapper');
    var dtScrollBody = $(el).find($('div.dataTables_scrollBody'));
    var framingHeight = dtWrapper.innerHeight() - dtScrollBody.innerHeight();
    var scrollBodyHeight = availableHeight - framingHeight;

    // set the height
    dtScrollBody.height(scrollBodyHeight + 'px');
  },

  // adjust the width of columns; remove the hard-coded widths on table and the
  // scroll header when scrollX/Y are enabled
  adjustWidth: function(el) {
    var $el = $(el), table = $el.data('datatable');
    if (table) table.columns.adjust();
    $el.find('.dataTables_scrollHeadInner').css('width', '')
        .children('table').css('margin-left', '');
  }
});

  if (!HTMLWidgets.shinyMode) return;

  Shiny.addCustomMessageHandler('datatable-calls', function(data) {
    var id = data.id;
    var el = document.getElementById(id);
    var table = el ? $(el).data('datatable') : null;
    if (!table) {
      console.log("Couldn't find table with id " + id);
      return;
    }

    var methods = table.shinyMethods, call = data.call;
    if (methods[call.method]) {
      methods[call.method].apply(table, call.args);
    } else {
      console.log("Unknown method " + call.method);
    }
  });

})();
```
