## Supplementary material for "Characterizing and inferring quantitative cell cycle phase in single-cell RNA-seq data analysis": Source code of our analysis, including scripts and data necessary to reproduce the work.: jquery.dataTables.min.js

```
/*!
 DataTables 1.10.16
 ©2008-2017 SpryMedia Ltd - datatables.net/license
*/
(function(h){"function"===typeof define&&define.amd?define(["jquery"],function(E){return h(E,window,document)}):"object"===typeof exports?module.exports=function(E,G){E||(E=window);G||(G="undefined"!==typeof window?require("jquery"):require("jquery")(E));return h(G,E,E.document)}:h(jQuery,window,document)})(function(h,E,G,k){function X(a){var b,c,d={};h.each(a,function(e){if((b=e.match(/^([^A-Z]+?)([A-Z])/))&&-1!=="a aa ai ao as b fn i m o s ".indexOf(b[1]+" "))c=e.replace(b[0],b[2].toLowerCase()),
d[c]=e,"o"===b[1]&&X(a[e])});a._hungarianMap=d}function I(a,b,c){a._hungarianMap||X(a);var d;h.each(b,function(e){d=a._hungarianMap[e];if(d!==k&&(c||b[d]===k))"o"===d.charAt(0)?(b[d]||(b[d]={}),h.extend(!0,b[d],b[e]),I(a[d],b[d],c)):b[d]=b[e]})}function Ca(a){var b=m.defaults.oLanguage,c=a.sZeroRecords;!a.sEmptyTable&&(c&&"No data available in table"===b.sEmptyTable)&&F(a,a,"sZeroRecords","sEmptyTable");!a.sLoadingRecords&&(c&&"Loading..."===b.sLoadingRecords)&&F(a,a,"sZeroRecords","sLoadingRecords");
a.sInfoThousands&&(a.sThousands=a.sInfoThousands);(a=a.sDecimal)&&cb(a)}function db(a){A(a,"ordering","bSort");A(a,"orderMulti","bSortMulti");A(a,"orderClasses","bSortClasses");A(a,"orderCellsTop","bSortCellsTop");A(a,"order","aaSorting");A(a,"orderFixed","aaSortingFixed");A(a,"paging","bPaginate");A(a,"pagingType","sPaginationType");A(a,"pageLength","iDisplayLength");A(a,"searching","bFilter");"boolean"===typeof a.sScrollX&&(a.sScrollX=a.sScrollX?"100%":"");"boolean"===typeof a.scrollX&&(a.scrollX=
a.scrollX?"100%":"");if(a=a.aoSearchCols)for(var b=0,c=a.length;b<c;b++)a[b]&&I(m.models.oSearch,a[b])}function eb(a){A(a,"orderable","bSortable");A(a,"orderData","aDataSort");A(a,"orderSequence","asSorting");A(a,"orderDataType","sortDataType");var b=a.aDataSort;"number"===typeof b&&!h.isArray(b)&&(a.aDataSort=[b])}function fb(a){if(!m.__browser){var b={};m.__browser=b;var c=h("<div/>").css({position:"fixed",top:0,left:-1*h(E).scrollLeft(),height:1,width:1,overflow:"hidden"}).append(h("<div/>").css({position:"absolute",
top:1,left:1,width:100,overflow:"scroll"}).append(h("<div/>").css({width:"100%",height:10}))).appendTo("body"),d=c.children(),e=d.children();b.barWidth=d[0].offsetWidth-d[0].clientWidth;b.bScrollOversize=100===e[0].offsetWidth&&100!==d[0].clientWidth;b.bScrollbarLeft=1!==Math.round(e.offset().left);b.bBounding=c[0].getBoundingClientRect().width?!0:!1;c.remove()}h.extend(a.oBrowser,m.__browser);a.oScroll.iBarWidth=m.__browser.barWidth}function gb(a,b,c,d,e,f){var g,j=!1;c!==k&&(g=c,j=!0);for(;d!==
e;)a.hasOwnProperty(d)&&(g=j?b(g,a[d],d,a):a[d],j=!0,d+=f);return g}function Da(a,b){var c=m.defaults.column,d=a.aoColumns.length,c=h.extend({},m.models.oColumn,c,{nTh:b?b:G.createElement("th"),sTitle:c.sTitle?c.sTitle:b?b.innerHTML:"",aDataSort:c.aDataSort?c.aDataSort:[d],mData:c.mData?c.mData:d,idx:d});a.aoColumns.push(c);c=a.aoPreSearchCols;c[d]=h.extend({},m.models.oSearch,c[d]);ja(a,d,h(b).data())}function ja(a,b,c){var b=a.aoColumns[b],d=a.oClasses,e=h(b.nTh);if(!b.sWidthOrig){b.sWidthOrig=
e.attr("width")||null;var f=(e.attr("style")||"").match(/width:\s*(\d+[pxem%]+)/);f&&(b.sWidthOrig=f[1])}c!==k&&null!==c&&(eb(c),I(m.defaults.column,c),c.mDataProp!==k&&!c.mData&&(c.mData=c.mDataProp),c.sType&&(b._sManualType=c.sType),c.className&&!c.sClass&&(c.sClass=c.className),c.sClass&&e.addClass(c.sClass),h.extend(b,c),F(b,c,"sWidth","sWidthOrig"),c.iDataSort!==k&&(b.aDataSort=[c.iDataSort]),F(b,c,"aDataSort"));var g=b.mData,j=Q(g),i=b.mRender?Q(b.mRender):null,c=function(a){return"string"===
typeof a&&-1!==a.indexOf("@")};b._bAttrSrc=h.isPlainObject(g)&&(c(g.sort)||c(g.type)||c(g.filter));b._setter=null;b.fnGetData=function(a,b,c){var d=j(a,b,k,c);return i&&b?i(d,b,a,c):d};b.fnSetData=function(a,b,c){return R(g)(a,b,c)};"number"!==typeof g&&(a._rowReadObject=!0);a.oFeatures.bSort||(b.bSortable=!1,e.addClass(d.sSortableNone));a=-1!==h.inArray("asc",b.asSorting);c=-1!==h.inArray("desc",b.asSorting);!b.bSortable||!a&&!c?(b.sSortingClass=d.sSortableNone,b.sSortingClassJUI=""):a&&!c?(b.sSortingClass=
d.sSortableAsc,b.sSortingClassJUI=d.sSortJUIAscAllowed):!a&&c?(b.sSortingClass=d.sSortableDesc,b.sSortingClassJUI=d.sSortJUIDescAllowed):(b.sSortingClass=d.sSortable,b.sSortingClassJUI=d.sSortJUI)}function Y(a){if(!1!==a.oFeatures.bAutoWidth){var b=a.aoColumns;Ea(a);for(var c=0,d=b.length;c<d;c++)b[c].nTh.style.width=b[c].sWidth}b=a.oScroll;(""!==b.sY||""!==b.sX)&&ka(a);r(a,null,"column-sizing",[a])}function Z(a,b){var c=la(a,"bVisible");return"number"===typeof c[b]?c[b]:null}function $(a,b){var c=
la(a,"bVisible"),c=h.inArray(b,c);return-1!==c?c:null}function aa(a){var b=0;h.each(a.aoColumns,function(a,d){d.bVisible&&"none"!==h(d.nTh).css("display")&&b++});return b}function la(a,b){var c=[];h.map(a.aoColumns,function(a,e){a[b]&&c.push(e)});return c}function Fa(a){var b=a.aoColumns,c=a.aoData,d=m.ext.type.detect,e,f,g,j,i,h,l,q,t;e=0;for(f=b.length;e<f;e++)if(l=b[e],t=[],!l.sType&&l._sManualType)l.sType=l._sManualType;else if(!l.sType){g=0;for(j=d.length;g<j;g++){i=0;for(h=c.length;i<h;i++){t[i]===
k&&(t[i]=B(a,i,e,"type"));q=d[g](t[i],a);if(!q&&g!==d.length-1)break;if("html"===q)break}if(q){l.sType=q;break}}l.sType||(l.sType="string")}}function hb(a,b,c,d){var e,f,g,j,i,n,l=a.aoColumns;if(b)for(e=b.length-1;0<=e;e--){n=b[e];var q=n.targets!==k?n.targets:n.aTargets;h.isArray(q)||(q=[q]);f=0;for(g=q.length;f<g;f++)if("number"===typeof q[f]&&0<=q[f]){for(;l.length<=q[f];)Da(a);d(q[f],n)}else if("number"===typeof q[f]&&0>q[f])d(l.length+q[f],n);else if("string"===typeof q[f]){j=0;for(i=l.length;j<
i;j++)("_all"==q[f]||h(l[j].nTh).hasClass(q[f]))&&d(j,n)}}if(c){e=0;for(a=c.length;e<a;e++)d(e,c[e])}}function M(a,b,c,d){var e=a.aoData.length,f=h.extend(!0,{},m.models.oRow,{src:c?"dom":"data",idx:e});f._aData=b;a.aoData.push(f);for(var g=a.aoColumns,j=0,i=g.length;j<i;j++)g[j].sType=null;a.aiDisplayMaster.push(e);b=a.rowIdFn(b);b!==k&&(a.aIds[b]=f);(c||!a.oFeatures.bDeferRender)&&Ga(a,e,c,d);return e}function ma(a,b){var c;b instanceof h||(b=h(b));return b.map(function(b,e){c=Ha(a,e);return M(a,
c.data,e,c.cells)})}function B(a,b,c,d){var e=a.iDraw,f=a.aoColumns[c],g=a.aoData[b]._aData,j=f.sDefaultContent,i=f.fnGetData(g,d,{settings:a,row:b,col:c});if(i===k)return a.iDrawError!=e&&null===j&&(J(a,0,"Requested unknown parameter "+("function"==typeof f.mData?"{function}":"'"+f.mData+"'")+" for row "+b+", column "+c,4),a.iDrawError=e),j;if((i===g||null===i)&&null!==j&&d!==k)i=j;else if("function"===typeof i)return i.call(g);return null===i&&"display"==d?"":i}function ib(a,b,c,d){a.aoColumns[c].fnSetData(a.aoData[b]._aData,
d,{settings:a,row:b,col:c})}function Ia(a){return h.map(a.match(/(\\.|[^\.])+/g)||[""],function(a){return a.replace(/\\\./g,".")})}function Q(a){if(h.isPlainObject(a)){var b={};h.each(a,function(a,c){c&&(b[a]=Q(c))});return function(a,c,f,g){var j=b[c]||b._;return j!==k?j(a,c,f,g):a}}if(null===a)return function(a){return a};if("function"===typeof a)return function(b,c,f,g){return a(b,c,f,g)};if("string"===typeof a&&(-1!==a.indexOf(".")||-1!==a.indexOf("[")||-1!==a.indexOf("("))){var c=function(a,
b,f){var g,j;if(""!==f){j=Ia(f);for(var i=0,n=j.length;i<n;i++){f=j[i].match(ba);g=j[i].match(U);if(f){j[i]=j[i].replace(ba,"");""!==j[i]&&(a=a[j[i]]);g=[];j.splice(0,i+1);j=j.join(".");if(h.isArray(a)){i=0;for(n=a.length;i<n;i++)g.push(c(a[i],b,j))}a=f[0].substring(1,f[0].length-1);a=""===a?g:g.join(a);break}else if(g){j[i]=j[i].replace(U,"");a=a[j[i]]();continue}if(null===a||a[j[i]]===k)return k;a=a[j[i]]}}return a};return function(b,e){return c(b,e,a)}}return function(b){return b[a]}}function R(a){if(h.isPlainObject(a))return R(a._);
if(null===a)return function(){};if("function"===typeof a)return function(b,d,e){a(b,"set",d,e)};if("string"===typeof a&&(-1!==a.indexOf(".")||-1!==a.indexOf("[")||-1!==a.indexOf("("))){var b=function(a,d,e){var e=Ia(e),f;f=e[e.length-1];for(var g,j,i=0,n=e.length-1;i<n;i++){g=e[i].match(ba);j=e[i].match(U);if(g){e[i]=e[i].replace(ba,"");a[e[i]]=[];f=e.slice();f.splice(0,i+1);g=f.join(".");if(h.isArray(d)){j=0;for(n=d.length;j<n;j++)f={},b(f,d[j],g),a[e[i]].push(f)}else a[e[i]]=d;return}j&&(e[i]=e[i].replace(U,
""),a=a[e[i]](d));if(null===a[e[i]]||a[e[i]]===k)a[e[i]]={};a=a[e[i]]}if(f.match(U))a[f.replace(U,"")](d);else a[f.replace(ba,"")]=d};return function(c,d){return b(c,d,a)}}return function(b,d){b[a]=d}}function Ja(a){return D(a.aoData,"_aData")}function na(a){a.aoData.length=0;a.aiDisplayMaster.length=0;a.aiDisplay.length=0;a.aIds={}}function oa(a,b,c){for(var d=-1,e=0,f=a.length;e<f;e++)a[e]==b?d=e:a[e]>b&&a[e]--; -1!=d&&c===k&&a.splice(d,1)}function ca(a,b,c,d){var e=a.aoData[b],f,g=function(c,d){for(;c.childNodes.length;)c.removeChild(c.firstChild);
c.innerHTML=B(a,b,d,"display")};if("dom"===c||(!c||"auto"===c)&&"dom"===e.src)e._aData=Ha(a,e,d,d===k?k:e._aData).data;else{var j=e.anCells;if(j)if(d!==k)g(j[d],d);else{c=0;for(f=j.length;c<f;c++)g(j[c],c)}}e._aSortData=null;e._aFilterData=null;g=a.aoColumns;if(d!==k)g[d].sType=null;else{c=0;for(f=g.length;c<f;c++)g[c].sType=null;Ka(a,e)}}function Ha(a,b,c,d){var e=[],f=b.firstChild,g,j,i=0,n,l=a.aoColumns,q=a._rowReadObject,d=d!==k?d:q?{}:[],t=function(a,b){if("string"===typeof a){var c=a.indexOf("@");
-1!==c&&(c=a.substring(c+1),R(a)(d,b.getAttribute(c)))}},m=function(a){if(c===k||c===i)j=l[i],n=h.trim(a.innerHTML),j&&j._bAttrSrc?(R(j.mData._)(d,n),t(j.mData.sort,a),t(j.mData.type,a),t(j.mData.filter,a)):q?(j._setter||(j._setter=R(j.mData)),j._setter(d,n)):d[i]=n;i++};if(f)for(;f;){g=f.nodeName.toUpperCase();if("TD"==g||"TH"==g)m(f),e.push(f);f=f.nextSibling}else{e=b.anCells;f=0;for(g=e.length;f<g;f++)m(e[f])}if(b=b.firstChild?b:b.nTr)(b=b.getAttribute("id"))&&R(a.rowId)(d,b);return{data:d,cells:e}}
function Ga(a,b,c,d){var e=a.aoData[b],f=e._aData,g=[],j,i,n,l,q;if(null===e.nTr){j=c||G.createElement("tr");e.nTr=j;e.anCells=g;j._DT_RowIndex=b;Ka(a,e);l=0;for(q=a.aoColumns.length;l<q;l++){n=a.aoColumns[l];i=c?d[l]:G.createElement(n.sCellType);i._DT_CellIndex={row:b,column:l};g.push(i);if((!c||n.mRender||n.mData!==l)&&(!h.isPlainObject(n.mData)||n.mData._!==l+".display"))i.innerHTML=B(a,b,l,"display");n.sClass&&(i.className+=" "+n.sClass);n.bVisible&&!c?j.appendChild(i):!n.bVisible&&c&&i.parentNode.removeChild(i);
n.fnCreatedCell&&n.fnCreatedCell.call(a.oInstance,i,B(a,b,l),f,b,l)}r(a,"aoRowCreatedCallback",null,[j,f,b])}e.nTr.setAttribute("role","row")}function Ka(a,b){var c=b.nTr,d=b._aData;if(c){var e=a.rowIdFn(d);e&&(c.id=e);d.DT_RowClass&&(e=d.DT_RowClass.split(" "),b.__rowc=b.__rowc?qa(b.__rowc.concat(e)):e,h(c).removeClass(b.__rowc.join(" ")).addClass(d.DT_RowClass));d.DT_RowAttr&&h(c).attr(d.DT_RowAttr);d.DT_RowData&&h(c).data(d.DT_RowData)}}function jb(a){var b,c,d,e,f,g=a.nTHead,j=a.nTFoot,i=0===
h("th, td",g).length,n=a.oClasses,l=a.aoColumns;i&&(e=h("<tr/>").appendTo(g));b=0;for(c=l.length;b<c;b++)f=l[b],d=h(f.nTh).addClass(f.sClass),i&&d.appendTo(e),a.oFeatures.bSort&&(d.addClass(f.sSortingClass),!1!==f.bSortable&&(d.attr("tabindex",a.iTabIndex).attr("aria-controls",a.sTableId),La(a,f.nTh,b))),f.sTitle!=d[0].innerHTML&&d.html(f.sTitle),Ma(a,"header")(a,d,f,n);i&&da(a.aoHeader,g);h(g).find(">tr").attr("role","row");h(g).find(">tr>th, >tr>td").addClass(n.sHeaderTH);h(j).find(">tr>th, >tr>td").addClass(n.sFooterTH);
if(null!==j){a=a.aoFooter[0];b=0;for(c=a.length;b<c;b++)f=l[b],f.nTf=a[b].cell,f.sClass&&h(f.nTf).addClass(f.sClass)}}function ea(a,b,c){var d,e,f,g=[],j=[],i=a.aoColumns.length,n;if(b){c===k&&(c=!1);d=0;for(e=b.length;d<e;d++){g[d]=b[d].slice();g[d].nTr=b[d].nTr;for(f=i-1;0<=f;f--)!a.aoColumns[f].bVisible&&!c&&g[d].splice(f,1);j.push([])}d=0;for(e=g.length;d<e;d++){if(a=g[d].nTr)for(;f=a.firstChild;)a.removeChild(f);f=0;for(b=g[d].length;f<b;f++)if(n=i=1,j[d][f]===k){a.appendChild(g[d][f].cell);
for(j[d][f]=1;g[d+i]!==k&&g[d][f].cell==g[d+i][f].cell;)j[d+i][f]=1,i++;for(;g[d][f+n]!==k&&g[d][f].cell==g[d][f+n].cell;){for(c=0;c<i;c++)j[d+c][f+n]=1;n++}h(g[d][f].cell).attr("rowspan",i).attr("colspan",n)}}}}function N(a){var b=r(a,"aoPreDrawCallback","preDraw",[a]);if(-1!==h.inArray(!1,b))C(a,!1);else{var b=[],c=0,d=a.asStripeClasses,e=d.length,f=a.oLanguage,g=a.iInitDisplayStart,j="ssp"==y(a),i=a.aiDisplay;a.bDrawing=!0;g!==k&&-1!==g&&(a._iDisplayStart=j?g:g>=a.fnRecordsDisplay()?0:g,a.iInitDisplayStart=
-1);var g=a._iDisplayStart,n=a.fnDisplayEnd();if(a.bDeferLoading)a.bDeferLoading=!1,a.iDraw++,C(a,!1);else if(j){if(!a.bDestroying&&!kb(a))return}else a.iDraw++;if(0!==i.length){f=j?a.aoData.length:n;for(j=j?0:g;j<f;j++){var l=i[j],q=a.aoData[l];null===q.nTr&&Ga(a,l);l=q.nTr;if(0!==e){var t=d[c%e];q._sRowStripe!=t&&(h(l).removeClass(q._sRowStripe).addClass(t),q._sRowStripe=t)}r(a,"aoRowCallback",null,[l,q._aData,c,j]);b.push(l);c++}}else c=f.sZeroRecords,1==a.iDraw&&"ajax"==y(a)?c=f.sLoadingRecords:
f.sEmptyTable&&0===a.fnRecordsTotal()&&(c=f.sEmptyTable),b[0]=h("<tr/>",{"class":e?d[0]:""}).append(h("<td />",{valign:"top",colSpan:aa(a),"class":a.oClasses.sRowEmpty}).html(c))[0];r(a,"aoHeaderCallback","header",[h(a.nTHead).children("tr")[0],Ja(a),g,n,i]);r(a,"aoFooterCallback","footer",[h(a.nTFoot).children("tr")[0],Ja(a),g,n,i]);d=h(a.nTBody);d.children().detach();d.append(h(b));r(a,"aoDrawCallback","draw",[a]);a.bSorted=!1;a.bFiltered=!1;a.bDrawing=!1}}function S(a,b){var c=a.oFeatures,d=c.bFilter;
c.bSort&&lb(a);d?fa(a,a.oPreviousSearch):a.aiDisplay=a.aiDisplayMaster.slice();!0!==b&&(a._iDisplayStart=0);a._drawHold=b;N(a);a._drawHold=!1}function mb(a){var b=a.oClasses,c=h(a.nTable),c=h("<div/>").insertBefore(c),d=a.oFeatures,e=h("<div/>",{id:a.sTableId+"_wrapper","class":b.sWrapper+(a.nTFoot?"":" "+b.sNoFooter)});a.nHolding=c[0];a.nTableWrapper=e[0];a.nTableReinsertBefore=a.nTable.nextSibling;for(var f=a.sDom.split(""),g,j,i,n,l,q,k=0;k<f.length;k++){g=null;j=f[k];if("<"==j){i=h("<div/>")[0];
n=f[k+1];if("'"==n||'"'==n){l="";for(q=2;f[k+q]!=n;)l+=f[k+q],q++;"H"==l?l=b.sJUIHeader:"F"==l&&(l=b.sJUIFooter);-1!=l.indexOf(".")?(n=l.split("."),i.id=n[0].substr(1,n[0].length-1),i.className=n[1]):"#"==l.charAt(0)?i.id=l.substr(1,l.length-1):i.className=l;k+=q}e.append(i);e=h(i)}else if(">"==j)e=e.parent();else if("l"==j&&d.bPaginate&&d.bLengthChange)g=nb(a);else if("f"==j&&d.bFilter)g=ob(a);else if("r"==j&&d.bProcessing)g=pb(a);else if("t"==j)g=qb(a);else if("i"==j&&d.bInfo)g=rb(a);else if("p"==
j&&d.bPaginate)g=sb(a);else if(0!==m.ext.feature.length){i=m.ext.feature;q=0;for(n=i.length;q<n;q++)if(j==i[q].cFeature){g=i[q].fnInit(a);break}}g&&(i=a.aanFeatures,i[j]||(i[j]=[]),i[j].push(g),e.append(g))}c.replaceWith(e);a.nHolding=null}function da(a,b){var c=h(b).children("tr"),d,e,f,g,j,i,n,l,q,k;a.splice(0,a.length);f=0;for(i=c.length;f<i;f++)a.push([]);f=0;for(i=c.length;f<i;f++){d=c[f];for(e=d.firstChild;e;){if("TD"==e.nodeName.toUpperCase()||"TH"==e.nodeName.toUpperCase()){l=1*e.getAttribute("colspan");
q=1*e.getAttribute("rowspan");l=!l||0===l||1===l?1:l;q=!q||0===q||1===q?1:q;g=0;for(j=a[f];j[g];)g++;n=g;k=1===l?!0:!1;for(j=0;j<l;j++)for(g=0;g<q;g++)a[f+g][n+j]={cell:e,unique:k},a[f+g].nTr=d}e=e.nextSibling}}}function ra(a,b,c){var d=[];c||(c=a.aoHeader,b&&(c=[],da(c,b)));for(var b=0,e=c.length;b<e;b++)for(var f=0,g=c[b].length;f<g;f++)if(c[b][f].unique&&(!d[f]||!a.bSortCellsTop))d[f]=c[b][f].cell;return d}function sa(a,b,c){r(a,"aoServerParams","serverParams",[b]);if(b&&h.isArray(b)){var d={},
e=/(.*?)\[\]$/;h.each(b,function(a,b){var c=b.name.match(e);c?(c=c[0],d[c]||(d[c]=[]),d[c].push(b.value)):d[b.name]=b.value});b=d}var f,g=a.ajax,j=a.oInstance,i=function(b){r(a,null,"xhr",[a,b,a.jqXHR]);c(b)};if(h.isPlainObject(g)&&g.data){f=g.data;var n=h.isFunction(f)?f(b,a):f,b=h.isFunction(f)&&n?n:h.extend(!0,b,n);delete g.data}n={data:b,success:function(b){var c=b.error||b.sError;c&&J(a,0,c);a.json=b;i(b)},dataType:"json",cache:!1,type:a.sServerMethod,error:function(b,c){var d=r(a,null,"xhr",
[a,null,a.jqXHR]);-1===h.inArray(!0,d)&&("parsererror"==c?J(a,0,"Invalid JSON response",1):4===b.readyState&&J(a,0,"Ajax error",7));C(a,!1)}};a.oAjaxData=b;r(a,null,"preXhr",[a,b]);a.fnServerData?a.fnServerData.call(j,a.sAjaxSource,h.map(b,function(a,b){return{name:b,value:a}}),i,a):a.sAjaxSource||"string"===typeof g?a.jqXHR=h.ajax(h.extend(n,{url:g||a.sAjaxSource})):h.isFunction(g)?a.jqXHR=g.call(j,b,i,a):(a.jqXHR=h.ajax(h.extend(n,g)),g.data=f)}function kb(a){return a.bAjaxDataGet?(a.iDraw++,C(a,
!0),sa(a,tb(a),function(b){ub(a,b)}),!1):!0}function tb(a){var b=a.aoColumns,c=b.length,d=a.oFeatures,e=a.oPreviousSearch,f=a.aoPreSearchCols,g,j=[],i,n,l,k=V(a);g=a._iDisplayStart;i=!1!==d.bPaginate?a._iDisplayLength:-1;var t=function(a,b){j.push({name:a,value:b})};t("sEcho",a.iDraw);t("iColumns",c);t("sColumns",D(b,"sName").join(","));t("iDisplayStart",g);t("iDisplayLength",i);var pa={draw:a.iDraw,columns:[],order:[],start:g,length:i,search:{value:e.sSearch,regex:e.bRegex}};for(g=0;g<c;g++)n=b[g],
l=f[g],i="function"==typeof n.mData?"function":n.mData,pa.columns.push({data:i,name:n.sName,searchable:n.bSearchable,orderable:n.bSortable,search:{value:l.sSearch,regex:l.bRegex}}),t("mDataProp_"+g,i),d.bFilter&&(t("sSearch_"+g,l.sSearch),t("bRegex_"+g,l.bRegex),t("bSearchable_"+g,n.bSearchable)),d.bSort&&t("bSortable_"+g,n.bSortable);d.bFilter&&(t("sSearch",e.sSearch),t("bRegex",e.bRegex));d.bSort&&(h.each(k,function(a,b){pa.order.push({column:b.col,dir:b.dir});t("iSortCol_"+a,b.col);t("sSortDir_"+
a,b.dir)}),t("iSortingCols",k.length));b=m.ext.legacy.ajax;return null===b?a.sAjaxSource?j:pa:b?j:pa}function ub(a,b){var c=ta(a,b),d=b.sEcho!==k?b.sEcho:b.draw,e=b.iTotalRecords!==k?b.iTotalRecords:b.recordsTotal,f=b.iTotalDisplayRecords!==k?b.iTotalDisplayRecords:b.recordsFiltered;if(d){if(1*d<a.iDraw)return;a.iDraw=1*d}na(a);a._iRecordsTotal=parseInt(e,10);a._iRecordsDisplay=parseInt(f,10);d=0;for(e=c.length;d<e;d++)M(a,c[d]);a.aiDisplay=a.aiDisplayMaster.slice();a.bAjaxDataGet=!1;N(a);a._bInitComplete||
ua(a,b);a.bAjaxDataGet=!0;C(a,!1)}function ta(a,b){var c=h.isPlainObject(a.ajax)&&a.ajax.dataSrc!==k?a.ajax.dataSrc:a.sAjaxDataProp;return"data"===c?b.aaData||b[c]:""!==c?Q(c)(b):b}function ob(a){var b=a.oClasses,c=a.sTableId,d=a.oLanguage,e=a.oPreviousSearch,f=a.aanFeatures,g='<input type="search" class="'+b.sFilterInput+'"/>',j=d.sSearch,j=j.match(/_INPUT_/)?j.replace("_INPUT_",g):j+g,b=h("<div/>",{id:!f.f?c+"_filter":null,"class":b.sFilter}).append(h("<label/>").append(j)),f=function(){var b=!this.value?
"":this.value;b!=e.sSearch&&(fa(a,{sSearch:b,bRegex:e.bRegex,bSmart:e.bSmart,bCaseInsensitive:e.bCaseInsensitive}),a._iDisplayStart=0,N(a))},g=null!==a.searchDelay?a.searchDelay:"ssp"===y(a)?400:0,i=h("input",b).val(e.sSearch).attr("placeholder",d.sSearchPlaceholder).on("keyup.DT search.DT input.DT paste.DT cut.DT",g?Na(f,g):f).on("keypress.DT",function(a){if(13==a.keyCode)return!1}).attr("aria-controls",c);h(a.nTable).on("search.dt.DT",function(b,c){if(a===c)try{i[0]!==G.activeElement&&i.val(e.sSearch)}catch(d){}});
return b[0]}function fa(a,b,c){var d=a.oPreviousSearch,e=a.aoPreSearchCols,f=function(a){d.sSearch=a.sSearch;d.bRegex=a.bRegex;d.bSmart=a.bSmart;d.bCaseInsensitive=a.bCaseInsensitive};Fa(a);if("ssp"!=y(a)){vb(a,b.sSearch,c,b.bEscapeRegex!==k?!b.bEscapeRegex:b.bRegex,b.bSmart,b.bCaseInsensitive);f(b);for(b=0;b<e.length;b++)wb(a,e[b].sSearch,b,e[b].bEscapeRegex!==k?!e[b].bEscapeRegex:e[b].bRegex,e[b].bSmart,e[b].bCaseInsensitive);xb(a)}else f(b);a.bFiltered=!0;r(a,null,"search",[a])}function xb(a){for(var b=
m.ext.search,c=a.aiDisplay,d,e,f=0,g=b.length;f<g;f++){for(var j=[],i=0,n=c.length;i<n;i++)e=c[i],d=a.aoData[e],b[f](a,d._aFilterData,e,d._aData,i)&&j.push(e);c.length=0;h.merge(c,j)}}function wb(a,b,c,d,e,f){if(""!==b){for(var g=[],j=a.aiDisplay,d=Oa(b,d,e,f),e=0;e<j.length;e++)b=a.aoData[j[e]]._aFilterData[c],d.test(b)&&g.push(j[e]);a.aiDisplay=g}}function vb(a,b,c,d,e,f){var d=Oa(b,d,e,f),f=a.oPreviousSearch.sSearch,g=a.aiDisplayMaster,j,e=[];0!==m.ext.search.length&&(c=!0);j=yb(a);if(0>=b.length)a.aiDisplay=
g.slice();else{if(j||c||f.length>b.length||0!==b.indexOf(f)||a.bSorted)a.aiDisplay=g.slice();b=a.aiDisplay;for(c=0;c<b.length;c++)d.test(a.aoData[b[c]]._sFilterRow)&&e.push(b[c]);a.aiDisplay=e}}function Oa(a,b,c,d){a=b?a:Pa(a);c&&(a="^(?=.*?"+h.map(a.match(/"[^"]+"|[^ ]+/g)||[""],function(a){if('"'===a.charAt(0))var b=a.match(/^"(.*)"$/),a=b?b[1]:a;return a.replace('"',"")}).join(")(?=.*?")+").*$");return RegExp(a,d?"i":"")}function yb(a){var b=a.aoColumns,c,d,e,f,g,j,i,h,l=m.ext.type.search;c=!1;
d=0;for(f=a.aoData.length;d<f;d++)if(h=a.aoData[d],!h._aFilterData){j=[];e=0;for(g=b.length;e<g;e++)c=b[e],c.bSearchable?(i=B(a,d,e,"filter"),l[c.sType]&&(i=l[c.sType](i)),null===i&&(i=""),"string"!==typeof i&&i.toString&&(i=i.toString())):i="",i.indexOf&&-1!==i.indexOf("&")&&(va.innerHTML=i,i=Wb?va.textContent:va.innerText),i.replace&&(i=i.replace(/[\r\n]/g,"")),j.push(i);h._aFilterData=j;h._sFilterRow=j.join("  ");c=!0}return c}function zb(a){return{search:a.sSearch,smart:a.bSmart,regex:a.bRegex,
caseInsensitive:a.bCaseInsensitive}}function Ab(a){return{sSearch:a.search,bSmart:a.smart,bRegex:a.regex,bCaseInsensitive:a.caseInsensitive}}function rb(a){var b=a.sTableId,c=a.aanFeatures.i,d=h("<div/>",{"class":a.oClasses.sInfo,id:!c?b+"_info":null});c||(a.aoDrawCallback.push({fn:Bb,sName:"information"}),d.attr("role","status").attr("aria-live","polite"),h(a.nTable).attr("aria-describedby",b+"_info"));return d[0]}function Bb(a){var b=a.aanFeatures.i;if(0!==b.length){var c=a.oLanguage,d=a._iDisplayStart+
1,e=a.fnDisplayEnd(),f=a.fnRecordsTotal(),g=a.fnRecordsDisplay(),j=g?c.sInfo:c.sInfoEmpty;g!==f&&(j+=" "+c.sInfoFiltered);j+=c.sInfoPostFix;j=Cb(a,j);c=c.fnInfoCallback;null!==c&&(j=c.call(a.oInstance,a,d,e,f,g,j));h(b).html(j)}}function Cb(a,b){var c=a.fnFormatNumber,d=a._iDisplayStart+1,e=a._iDisplayLength,f=a.fnRecordsDisplay(),g=-1===e;return b.replace(/_START_/g,c.call(a,d)).replace(/_END_/g,c.call(a,a.fnDisplayEnd())).replace(/_MAX_/g,c.call(a,a.fnRecordsTotal())).replace(/_TOTAL_/g,c.call(a,
f)).replace(/_PAGE_/g,c.call(a,g?1:Math.ceil(d/e))).replace(/_PAGES_/g,c.call(a,g?1:Math.ceil(f/e)))}function ga(a){var b,c,d=a.iInitDisplayStart,e=a.aoColumns,f;c=a.oFeatures;var g=a.bDeferLoading;if(a.bInitialised){mb(a);jb(a);ea(a,a.aoHeader);ea(a,a.aoFooter);C(a,!0);c.bAutoWidth&&Ea(a);b=0;for(c=e.length;b<c;b++)f=e[b],f.sWidth&&(f.nTh.style.width=v(f.sWidth));r(a,null,"preInit",[a]);S(a);e=y(a);if("ssp"!=e||g)"ajax"==e?sa(a,[],function(c){var f=ta(a,c);for(b=0;b<f.length;b++)M(a,f[b]);a.iInitDisplayStart=
d;S(a);C(a,!1);ua(a,c)},a):(C(a,!1),ua(a))}else setTimeout(function(){ga(a)},200)}function ua(a,b){a._bInitComplete=!0;(b||a.oInit.aaData)&&Y(a);r(a,null,"plugin-init",[a,b]);r(a,"aoInitComplete","init",[a,b])}function Qa(a,b){var c=parseInt(b,10);a._iDisplayLength=c;Ra(a);r(a,null,"length",[a,c])}function nb(a){for(var b=a.oClasses,c=a.sTableId,d=a.aLengthMenu,e=h.isArray(d[0]),f=e?d[0]:d,d=e?d[1]:d,e=h("<select/>",{name:c+"_length","aria-controls":c,"class":b.sLengthSelect}),g=0,j=f.length;g<j;g++)e[0][g]=
new Option("number"===typeof d[g]?a.fnFormatNumber(d[g]):d[g],f[g]);var i=h("<div><label/></div>").addClass(b.sLength);a.aanFeatures.l||(i[0].id=c+"_length");i.children().append(a.oLanguage.sLengthMenu.replace("_MENU_",e[0].outerHTML));h("select",i).val(a._iDisplayLength).on("change.DT",function(){Qa(a,h(this).val());N(a)});h(a.nTable).on("length.dt.DT",function(b,c,d){a===c&&h("select",i).val(d)});return i[0]}function sb(a){var b=a.sPaginationType,c=m.ext.pager[b],d="function"===typeof c,e=function(a){N(a)},
b=h("<div/>").addClass(a.oClasses.sPaging+b)[0],f=a.aanFeatures;d||c.fnInit(a,b,e);f.p||(b.id=a.sTableId+"_paginate",a.aoDrawCallback.push({fn:function(a){if(d){var b=a._iDisplayStart,i=a._iDisplayLength,h=a.fnRecordsDisplay(),l=-1===i,b=l?0:Math.ceil(b/i),i=l?1:Math.ceil(h/i),h=c(b,i),k,l=0;for(k=f.p.length;l<k;l++)Ma(a,"pageButton")(a,f.p[l],l,h,b,i)}else c.fnUpdate(a,e)},sName:"pagination"}));return b}function Sa(a,b,c){var d=a._iDisplayStart,e=a._iDisplayLength,f=a.fnRecordsDisplay();0===f||-1===
e?d=0:"number"===typeof b?(d=b*e,d>f&&(d=0)):"first"==b?d=0:"previous"==b?(d=0<=e?d-e:0,0>d&&(d=0)):"next"==b?d+e<f&&(d+=e):"last"==b?d=Math.floor((f-1)/e)*e:J(a,0,"Unknown paging action: "+b,5);b=a._iDisplayStart!==d;a._iDisplayStart=d;b&&(r(a,null,"page",[a]),c&&N(a));return b}function pb(a){return h("<div/>",{id:!a.aanFeatures.r?a.sTableId+"_processing":null,"class":a.oClasses.sProcessing}).html(a.oLanguage.sProcessing).insertBefore(a.nTable)[0]}function C(a,b){a.oFeatures.bProcessing&&h(a.aanFeatures.r).css("display",
b?"block":"none");r(a,null,"processing",[a,b])}function qb(a){var b=h(a.nTable);b.attr("role","grid");var c=a.oScroll;if(""===c.sX&&""===c.sY)return a.nTable;var d=c.sX,e=c.sY,f=a.oClasses,g=b.children("caption"),j=g.length?g[0]._captionSide:null,i=h(b[0].cloneNode(!1)),n=h(b[0].cloneNode(!1)),l=b.children("tfoot");l.length||(l=null);i=h("<div/>",{"class":f.sScrollWrapper}).append(h("<div/>",{"class":f.sScrollHead}).css({overflow:"hidden",position:"relative",border:0,width:d?!d?null:v(d):"100%"}).append(h("<div/>",
{"class":f.sScrollHeadInner}).css({"box-sizing":"content-box",width:c.sXInner||"100%"}).append(i.removeAttr("id").css("margin-left",0).append("top"===j?g:null).append(b.children("thead"))))).append(h("<div/>",{"class":f.sScrollBody}).css({position:"relative",overflow:"auto",width:!d?null:v(d)}).append(b));l&&i.append(h("<div/>",{"class":f.sScrollFoot}).css({overflow:"hidden",border:0,width:d?!d?null:v(d):"100%"}).append(h("<div/>",{"class":f.sScrollFootInner}).append(n.removeAttr("id").css("margin-left",
0).append("bottom"===j?g:null).append(b.children("tfoot")))));var b=i.children(),k=b[0],f=b[1],t=l?b[2]:null;if(d)h(f).on("scroll.DT",function(){var a=this.scrollLeft;k.scrollLeft=a;l&&(t.scrollLeft=a)});h(f).css(e&&c.bCollapse?"max-height":"height",e);a.nScrollHead=k;a.nScrollBody=f;a.nScrollFoot=t;a.aoDrawCallback.push({fn:ka,sName:"scrolling"});return i[0]}function ka(a){var b=a.oScroll,c=b.sX,d=b.sXInner,e=b.sY,b=b.iBarWidth,f=h(a.nScrollHead),g=f[0].style,j=f.children("div"),i=j[0].style,n=j.children("table"),
j=a.nScrollBody,l=h(j),q=j.style,t=h(a.nScrollFoot).children("div"),m=t.children("table"),o=h(a.nTHead),p=h(a.nTable),s=p[0],r=s.style,u=a.nTFoot?h(a.nTFoot):null,x=a.oBrowser,T=x.bScrollOversize,Xb=D(a.aoColumns,"nTh"),O,K,P,w,Ta=[],y=[],z=[],A=[],B,C=function(a){a=a.style;a.paddingTop="0";a.paddingBottom="0";a.borderTopWidth="0";a.borderBottomWidth="0";a.height=0};K=j.scrollHeight>j.clientHeight;if(a.scrollBarVis!==K&&a.scrollBarVis!==k)a.scrollBarVis=K,Y(a);else{a.scrollBarVis=K;p.children("thead, tfoot").remove();
u&&(P=u.clone().prependTo(p),O=u.find("tr"),P=P.find("tr"));w=o.clone().prependTo(p);o=o.find("tr");K=w.find("tr");w.find("th, td").removeAttr("tabindex");c||(q.width="100%",f[0].style.width="100%");h.each(ra(a,w),function(b,c){B=Z(a,b);c.style.width=a.aoColumns[B].sWidth});u&&H(function(a){a.style.width=""},P);f=p.outerWidth();if(""===c){r.width="100%";if(T&&(p.find("tbody").height()>j.offsetHeight||"scroll"==l.css("overflow-y")))r.width=v(p.outerWidth()-b);f=p.outerWidth()}else""!==d&&(r.width=
v(d),f=p.outerWidth());H(C,K);H(function(a){z.push(a.innerHTML);Ta.push(v(h(a).css("width")))},K);H(function(a,b){if(h.inArray(a,Xb)!==-1)a.style.width=Ta[b]},o);h(K).height(0);u&&(H(C,P),H(function(a){A.push(a.innerHTML);y.push(v(h(a).css("width")))},P),H(function(a,b){a.style.width=y[b]},O),h(P).height(0));H(function(a,b){a.innerHTML='<div class="dataTables_sizing" style="height:0;overflow:hidden;">'+z[b]+"</div>";a.style.width=Ta[b]},K);u&&H(function(a,b){a.innerHTML='<div class="dataTables_sizing" style="height:0;overflow:hidden;">'+
A[b]+"</div>";a.style.width=y[b]},P);if(p.outerWidth()<f){O=j.scrollHeight>j.offsetHeight||"scroll"==l.css("overflow-y")?f+b:f;if(T&&(j.scrollHeight>j.offsetHeight||"scroll"==l.css("overflow-y")))r.width=v(O-b);(""===c||""!==d)&&J(a,1,"Possible column misalignment",6)}else O="100%";q.width=v(O);g.width=v(O);u&&(a.nScrollFoot.style.width=v(O));!e&&T&&(q.height=v(s.offsetHeight+b));c=p.outerWidth();n[0].style.width=v(c);i.width=v(c);d=p.height()>j.clientHeight||"scroll"==l.css("overflow-y");e="padding"+
(x.bScrollbarLeft?"Left":"Right");i[e]=d?b+"px":"0px";u&&(m[0].style.width=v(c),t[0].style.width=v(c),t[0].style[e]=d?b+"px":"0px");p.children("colgroup").insertBefore(p.children("thead"));l.scroll();if((a.bSorted||a.bFiltered)&&!a._drawHold)j.scrollTop=0}}function H(a,b,c){for(var d=0,e=0,f=b.length,g,j;e<f;){g=b[e].firstChild;for(j=c?c[e].firstChild:null;g;)1===g.nodeType&&(c?a(g,j,d):a(g,d),d++),g=g.nextSibling,j=c?j.nextSibling:null;e++}}function Ea(a){var b=a.nTable,c=a.aoColumns,d=a.oScroll,
e=d.sY,f=d.sX,g=d.sXInner,j=c.length,i=la(a,"bVisible"),n=h("th",a.nTHead),l=b.getAttribute("width"),k=b.parentNode,t=!1,m,o,p=a.oBrowser,d=p.bScrollOversize;(m=b.style.width)&&-1!==m.indexOf("%")&&(l=m);for(m=0;m<i.length;m++)o=c[i[m]],null!==o.sWidth&&(o.sWidth=Db(o.sWidthOrig,k),t=!0);if(d||!t&&!f&&!e&&j==aa(a)&&j==n.length)for(m=0;m<j;m++)i=Z(a,m),null!==i&&(c[i].sWidth=v(n.eq(m).width()));else{j=h(b).clone().css("visibility","hidden").removeAttr("id");j.find("tbody tr").remove();var s=h("<tr/>").appendTo(j.find("tbody"));
j.find("thead, tfoot").remove();j.append(h(a.nTHead).clone()).append(h(a.nTFoot).clone());j.find("tfoot th, tfoot td").css("width","");n=ra(a,j.find("thead")[0]);for(m=0;m<i.length;m++)o=c[i[m]],n[m].style.width=null!==o.sWidthOrig&&""!==o.sWidthOrig?v(o.sWidthOrig):"",o.sWidthOrig&&f&&h(n[m]).append(h("<div/>").css({width:o.sWidthOrig,margin:0,padding:0,border:0,height:1}));if(a.aoData.length)for(m=0;m<i.length;m++)t=i[m],o=c[t],h(Eb(a,t)).clone(!1).append(o.sContentPadding).appendTo(s);h("[name]",
j).removeAttr("name");o=h("<div/>").css(f||e?{position:"absolute",top:0,left:0,height:1,right:0,overflow:"hidden"}:{}).append(j).appendTo(k);f&&g?j.width(g):f?(j.css("width","auto"),j.removeAttr("width"),j.width()<k.clientWidth&&l&&j.width(k.clientWidth)):e?j.width(k.clientWidth):l&&j.width(l);for(m=e=0;m<i.length;m++)k=h(n[m]),g=k.outerWidth()-k.width(),k=p.bBounding?Math.ceil(n[m].getBoundingClientRect().width):k.outerWidth(),e+=k,c[i[m]].sWidth=v(k-g);b.style.width=v(e);o.remove()}l&&(b.style.width=
v(l));if((l||f)&&!a._reszEvt)b=function(){h(E).on("resize.DT-"+a.sInstance,Na(function(){Y(a)}))},d?setTimeout(b,1E3):b(),a._reszEvt=!0}function Db(a,b){if(!a)return 0;var c=h("<div/>").css("width",v(a)).appendTo(b||G.body),d=c[0].offsetWidth;c.remove();return d}function Eb(a,b){var c=Fb(a,b);if(0>c)return null;var d=a.aoData[c];return!d.nTr?h("<td/>").html(B(a,c,b,"display"))[0]:d.anCells[b]}function Fb(a,b){for(var c,d=-1,e=-1,f=0,g=a.aoData.length;f<g;f++)c=B(a,f,b,"display")+"",c=c.replace(Yb,
""),c=c.replace(/&nbsp;/g," "),c.length>d&&(d=c.length,e=f);return e}function v(a){return null===a?"0px":"number"==typeof a?0>a?"0px":a+"px":a.match(/\d$/)?a+"px":a}function V(a){var b,c,d=[],e=a.aoColumns,f,g,j,i;b=a.aaSortingFixed;c=h.isPlainObject(b);var n=[];f=function(a){a.length&&!h.isArray(a[0])?n.push(a):h.merge(n,a)};h.isArray(b)&&f(b);c&&b.pre&&f(b.pre);f(a.aaSorting);c&&b.post&&f(b.post);for(a=0;a<n.length;a++){i=n[a][0];f=e[i].aDataSort;b=0;for(c=f.length;b<c;b++)g=f[b],j=e[g].sType||
"string",n[a]._idx===k&&(n[a]._idx=h.inArray(n[a][1],e[g].asSorting)),d.push({src:i,col:g,dir:n[a][1],index:n[a]._idx,type:j,formatter:m.ext.type.order[j+"-pre"]})}return d}function lb(a){var b,c,d=[],e=m.ext.type.order,f=a.aoData,g=0,j,i=a.aiDisplayMaster,h;Fa(a);h=V(a);b=0;for(c=h.length;b<c;b++)j=h[b],j.formatter&&g++,Gb(a,j.col);if("ssp"!=y(a)&&0!==h.length){b=0;for(c=i.length;b<c;b++)d[i[b]]=b;g===h.length?i.sort(function(a,b){var c,e,g,j,i=h.length,k=f[a]._aSortData,m=f[b]._aSortData;for(g=
0;g<i;g++)if(j=h[g],c=k[j.col],e=m[j.col],c=c<e?-1:c>e?1:0,0!==c)return"asc"===j.dir?c:-c;c=d[a];e=d[b];return c<e?-1:c>e?1:0}):i.sort(function(a,b){var c,g,j,i,k=h.length,m=f[a]._aSortData,o=f[b]._aSortData;for(j=0;j<k;j++)if(i=h[j],c=m[i.col],g=o[i.col],i=e[i.type+"-"+i.dir]||e["string-"+i.dir],c=i(c,g),0!==c)return c;c=d[a];g=d[b];return c<g?-1:c>g?1:0})}a.bSorted=!0}function Hb(a){for(var b,c,d=a.aoColumns,e=V(a),a=a.oLanguage.oAria,f=0,g=d.length;f<g;f++){c=d[f];var j=c.asSorting;b=c.sTitle.replace(/<.*?>/g,
"");var i=c.nTh;i.removeAttribute("aria-sort");c.bSortable&&(0<e.length&&e[0].col==f?(i.setAttribute("aria-sort","asc"==e[0].dir?"ascending":"descending"),c=j[e[0].index+1]||j[0]):c=j[0],b+="asc"===c?a.sSortAscending:a.sSortDescending);i.setAttribute("aria-label",b)}}function Ua(a,b,c,d){var e=a.aaSorting,f=a.aoColumns[b].asSorting,g=function(a,b){var c=a._idx;c===k&&(c=h.inArray(a[1],f));return c+1<f.length?c+1:b?null:0};"number"===typeof e[0]&&(e=a.aaSorting=[e]);c&&a.oFeatures.bSortMulti?(c=h.inArray(b,
D(e,"0")),-1!==c?(b=g(e[c],!0),null===b&&1===e.length&&(b=0),null===b?e.splice(c,1):(e[c][1]=f[b],e[c]._idx=b)):(e.push([b,f[0],0]),e[e.length-1]._idx=0)):e.length&&e[0][0]==b?(b=g(e[0]),e.length=1,e[0][1]=f[b],e[0]._idx=b):(e.length=0,e.push([b,f[0]]),e[0]._idx=0);S(a);"function"==typeof d&&d(a)}function La(a,b,c,d){var e=a.aoColumns[c];Va(b,{},function(b){!1!==e.bSortable&&(a.oFeatures.bProcessing?(C(a,!0),setTimeout(function(){Ua(a,c,b.shiftKey,d);"ssp"!==y(a)&&C(a,!1)},0)):Ua(a,c,b.shiftKey,d))})}
function wa(a){var b=a.aLastSort,c=a.oClasses.sSortColumn,d=V(a),e=a.oFeatures,f,g;if(e.bSort&&e.bSortClasses){e=0;for(f=b.length;e<f;e++)g=b[e].src,h(D(a.aoData,"anCells",g)).removeClass(c+(2>e?e+1:3));e=0;for(f=d.length;e<f;e++)g=d[e].src,h(D(a.aoData,"anCells",g)).addClass(c+(2>e?e+1:3))}a.aLastSort=d}function Gb(a,b){var c=a.aoColumns[b],d=m.ext.order[c.sSortDataType],e;d&&(e=d.call(a.oInstance,a,b,$(a,b)));for(var f,g=m.ext.type.order[c.sType+"-pre"],j=0,i=a.aoData.length;j<i;j++)if(c=a.aoData[j],
c._aSortData||(c._aSortData=[]),!c._aSortData[b]||d)f=d?e[j]:B(a,j,b,"sort"),c._aSortData[b]=g?g(f):f}function xa(a){if(a.oFeatures.bStateSave&&!a.bDestroying){var b={time:+new Date,start:a._iDisplayStart,length:a._iDisplayLength,order:h.extend(!0,[],a.aaSorting),search:zb(a.oPreviousSearch),columns:h.map(a.aoColumns,function(b,d){return{visible:b.bVisible,search:zb(a.aoPreSearchCols[d])}})};r(a,"aoStateSaveParams","stateSaveParams",[a,b]);a.oSavedState=b;a.fnStateSaveCallback.call(a.oInstance,a,
b)}}function Ib(a,b,c){var d,e,f=a.aoColumns,b=function(b){if(b&&b.time){var g=r(a,"aoStateLoadParams","stateLoadParams",[a,b]);if(-1===h.inArray(!1,g)&&(g=a.iStateDuration,!(0<g&&b.time<+new Date-1E3*g)&&!(b.columns&&f.length!==b.columns.length))){a.oLoadedState=h.extend(!0,{},b);b.start!==k&&(a._iDisplayStart=b.start,a.iInitDisplayStart=b.start);b.length!==k&&(a._iDisplayLength=b.length);b.order!==k&&(a.aaSorting=[],h.each(b.order,function(b,c){a.aaSorting.push(c[0]>=f.length?[0,c[1]]:c)}));b.search!==
k&&h.extend(a.oPreviousSearch,Ab(b.search));if(b.columns){d=0;for(e=b.columns.length;d<e;d++)g=b.columns[d],g.visible!==k&&(f[d].bVisible=g.visible),g.search!==k&&h.extend(a.aoPreSearchCols[d],Ab(g.search))}r(a,"aoStateLoaded","stateLoaded",[a,b])}}c()};if(a.oFeatures.bStateSave){var g=a.fnStateLoadCallback.call(a.oInstance,a,b);g!==k&&b(g)}else c()}function ya(a){var b=m.settings,a=h.inArray(a,D(b,"nTable"));return-1!==a?b[a]:null}function J(a,b,c,d){c="DataTables warning: "+(a?"table id="+a.sTableId+
" - ":"")+c;d&&(c+=". For more information about this error, please see http://datatables.net/tn/"+d);if(b)E.console&&console.log&&console.log(c);else if(b=m.ext,b=b.sErrMode||b.errMode,a&&r(a,null,"error",[a,d,c]),"alert"==b)alert(c);else{if("throw"==b)throw Error(c);"function"==typeof b&&b(a,d,c)}}function F(a,b,c,d){h.isArray(c)?h.each(c,function(c,d){h.isArray(d)?F(a,b,d[0],d[1]):F(a,b,d)}):(d===k&&(d=c),b[c]!==k&&(a[d]=b[c]))}function Jb(a,b,c){var d,e;for(e in b)b.hasOwnProperty(e)&&(d=b[e],
h.isPlainObject(d)?(h.isPlainObject(a[e])||(a[e]={}),h.extend(!0,a[e],d)):a[e]=c&&"data"!==e&&"aaData"!==e&&h.isArray(d)?d.slice():d);return a}function Va(a,b,c){h(a).on("click.DT",b,function(b){a.blur();c(b)}).on("keypress.DT",b,function(a){13===a.which&&(a.preventDefault(),c(a))}).on("selectstart.DT",function(){return!1})}function z(a,b,c,d){c&&a[b].push({fn:c,sName:d})}function r(a,b,c,d){var e=[];b&&(e=h.map(a[b].slice().reverse(),function(b){return b.fn.apply(a.oInstance,d)}));null!==c&&(b=h.Event(c+
".dt"),h(a.nTable).trigger(b,d),e.push(b.result));return e}function Ra(a){var b=a._iDisplayStart,c=a.fnDisplayEnd(),d=a._iDisplayLength;b>=c&&(b=c-d);b-=b%d;if(-1===d||0>b)b=0;a._iDisplayStart=b}function Ma(a,b){var c=a.renderer,d=m.ext.renderer[b];return h.isPlainObject(c)&&c[b]?d[c[b]]||d._:"string"===typeof c?d[c]||d._:d._}function y(a){return a.oFeatures.bServerSide?"ssp":a.ajax||a.sAjaxSource?"ajax":"dom"}function ha(a,b){var c=[],c=Kb.numbers_length,d=Math.floor(c/2);b<=c?c=W(0,b):a<=d?(c=W(0,
c-2),c.push("ellipsis"),c.push(b-1)):(a>=b-1-d?c=W(b-(c-2),b):(c=W(a-d+2,a+d-1),c.push("ellipsis"),c.push(b-1)),c.splice(0,0,"ellipsis"),c.splice(0,0,0));c.DT_el="span";return c}function cb(a){h.each({num:function(b){return za(b,a)},"num-fmt":function(b){return za(b,a,Wa)},"html-num":function(b){return za(b,a,Aa)},"html-num-fmt":function(b){return za(b,a,Aa,Wa)}},function(b,c){x.type.order[b+a+"-pre"]=c;b.match(/^html\-/)&&(x.type.search[b+a]=x.type.search.html)})}function Lb(a){return function(){var b=
[ya(this[m.ext.iApiIndex])].concat(Array.prototype.slice.call(arguments));return m.ext.internal[a].apply(this,b)}}var m=function(a){this.$=function(a,b){return this.api(!0).$(a,b)};this._=function(a,b){return this.api(!0).rows(a,b).data()};this.api=function(a){return a?new s(ya(this[x.iApiIndex])):new s(this)};this.fnAddData=function(a,b){var c=this.api(!0),d=h.isArray(a)&&(h.isArray(a[0])||h.isPlainObject(a[0]))?c.rows.add(a):c.row.add(a);(b===k||b)&&c.draw();return d.flatten().toArray()};this.fnAdjustColumnSizing=
function(a){var b=this.api(!0).columns.adjust(),c=b.settings()[0],d=c.oScroll;a===k||a?b.draw(!1):(""!==d.sX||""!==d.sY)&&ka(c)};this.fnClearTable=function(a){var b=this.api(!0).clear();(a===k||a)&&b.draw()};this.fnClose=function(a){this.api(!0).row(a).child.hide()};this.fnDeleteRow=function(a,b,c){var d=this.api(!0),a=d.rows(a),e=a.settings()[0],h=e.aoData[a[0][0]];a.remove();b&&b.call(this,e,h);(c===k||c)&&d.draw();return h};this.fnDestroy=function(a){this.api(!0).destroy(a)};this.fnDraw=function(a){this.api(!0).draw(a)};
this.fnFilter=function(a,b,c,d,e,h){e=this.api(!0);null===b||b===k?e.search(a,c,d,h):e.column(b).search(a,c,d,h);e.draw()};this.fnGetData=function(a,b){var c=this.api(!0);if(a!==k){var d=a.nodeName?a.nodeName.toLowerCase():"";return b!==k||"td"==d||"th"==d?c.cell(a,b).data():c.row(a).data()||null}return c.data().toArray()};this.fnGetNodes=function(a){var b=this.api(!0);return a!==k?b.row(a).node():b.rows().nodes().flatten().toArray()};this.fnGetPosition=function(a){var b=this.api(!0),c=a.nodeName.toUpperCase();
return"TR"==c?b.row(a).index():"TD"==c||"TH"==c?(a=b.cell(a).index(),[a.row,a.columnVisible,a.column]):null};this.fnIsOpen=function(a){return this.api(!0).row(a).child.isShown()};this.fnOpen=function(a,b,c){return this.api(!0).row(a).child(b,c).show().child()[0]};this.fnPageChange=function(a,b){var c=this.api(!0).page(a);(b===k||b)&&c.draw(!1)};this.fnSetColumnVis=function(a,b,c){a=this.api(!0).column(a).visible(b);(c===k||c)&&a.columns.adjust().draw()};this.fnSettings=function(){return ya(this[x.iApiIndex])};
this.fnSort=function(a){this.api(!0).order(a).draw()};this.fnSortListener=function(a,b,c){this.api(!0).order.listener(a,b,c)};this.fnUpdate=function(a,b,c,d,e){var h=this.api(!0);c===k||null===c?h.row(b).data(a):h.cell(b,c).data(a);(e===k||e)&&h.columns.adjust();(d===k||d)&&h.draw();return 0};this.fnVersionCheck=x.fnVersionCheck;var b=this,c=a===k,d=this.length;c&&(a={});this.oApi=this.internal=x.internal;for(var e in m.ext.internal)e&&(this[e]=Lb(e));this.each(function(){var e={},g=1<d?Jb(e,a,!0):
a,j=0,i,e=this.getAttribute("id"),n=!1,l=m.defaults,q=h(this);if("table"!=this.nodeName.toLowerCase())J(null,0,"Non-table node initialisation ("+this.nodeName+")",2);else{db(l);eb(l.column);I(l,l,!0);I(l.column,l.column,!0);I(l,h.extend(g,q.data()));var t=m.settings,j=0;for(i=t.length;j<i;j++){var o=t[j];if(o.nTable==this||o.nTHead.parentNode==this||o.nTFoot&&o.nTFoot.parentNode==this){var s=g.bRetrieve!==k?g.bRetrieve:l.bRetrieve;if(c||s)return o.oInstance;if(g.bDestroy!==k?g.bDestroy:l.bDestroy){o.oInstance.fnDestroy();
break}else{J(o,0,"Cannot reinitialise DataTable",3);return}}if(o.sTableId==this.id){t.splice(j,1);break}}if(null===e||""===e)this.id=e="DataTables_Table_"+m.ext._unique++;var p=h.extend(!0,{},m.models.oSettings,{sDestroyWidth:q[0].style.width,sInstance:e,sTableId:e});p.nTable=this;p.oApi=b.internal;p.oInit=g;t.push(p);p.oInstance=1===b.length?b:q.dataTable();db(g);g.oLanguage&&Ca(g.oLanguage);g.aLengthMenu&&!g.iDisplayLength&&(g.iDisplayLength=h.isArray(g.aLengthMenu[0])?g.aLengthMenu[0][0]:g.aLengthMenu[0]);
g=Jb(h.extend(!0,{},l),g);F(p.oFeatures,g,"bPaginate bLengthChange bFilter bSort bSortMulti bInfo bProcessing bAutoWidth bSortClasses bServerSide bDeferRender".split(" "));F(p,g,["asStripeClasses","ajax","fnServerData","fnFormatNumber","sServerMethod","aaSorting","aaSortingFixed","aLengthMenu","sPaginationType","sAjaxSource","sAjaxDataProp","iStateDuration","sDom","bSortCellsTop","iTabIndex","fnStateLoadCallback","fnStateSaveCallback","renderer","searchDelay","rowId",["iCookieDuration","iStateDuration"],
["oSearch","oPreviousSearch"],["aoSearchCols","aoPreSearchCols"],["iDisplayLength","_iDisplayLength"]]);F(p.oScroll,g,[["sScrollX","sX"],["sScrollXInner","sXInner"],["sScrollY","sY"],["bScrollCollapse","bCollapse"]]);F(p.oLanguage,g,"fnInfoCallback");z(p,"aoDrawCallback",g.fnDrawCallback,"user");z(p,"aoServerParams",g.fnServerParams,"user");z(p,"aoStateSaveParams",g.fnStateSaveParams,"user");z(p,"aoStateLoadParams",g.fnStateLoadParams,"user");z(p,"aoStateLoaded",g.fnStateLoaded,"user");z(p,"aoRowCallback",
g.fnRowCallback,"user");z(p,"aoRowCreatedCallback",g.fnCreatedRow,"user");z(p,"aoHeaderCallback",g.fnHeaderCallback,"user");z(p,"aoFooterCallback",g.fnFooterCallback,"user");z(p,"aoInitComplete",g.fnInitComplete,"user");z(p,"aoPreDrawCallback",g.fnPreDrawCallback,"user");p.rowIdFn=Q(g.rowId);fb(p);var u=p.oClasses;h.extend(u,m.ext.classes,g.oClasses);q.addClass(u.sTable);p.iInitDisplayStart===k&&(p.iInitDisplayStart=g.iDisplayStart,p._iDisplayStart=g.iDisplayStart);null!==g.iDeferLoading&&(p.bDeferLoading=
!0,e=h.isArray(g.iDeferLoading),p._iRecordsDisplay=e?g.iDeferLoading[0]:g.iDeferLoading,p._iRecordsTotal=e?g.iDeferLoading[1]:g.iDeferLoading);var v=p.oLanguage;h.extend(!0,v,g.oLanguage);v.sUrl&&(h.ajax({dataType:"json",url:v.sUrl,success:function(a){Ca(a);I(l.oLanguage,a);h.extend(true,v,a);ga(p)},error:function(){ga(p)}}),n=!0);null===g.asStripeClasses&&(p.asStripeClasses=[u.sStripeOdd,u.sStripeEven]);var e=p.asStripeClasses,x=q.children("tbody").find("tr").eq(0);-1!==h.inArray(!0,h.map(e,function(a){return x.hasClass(a)}))&&
(h("tbody tr",this).removeClass(e.join(" ")),p.asDestroyStripes=e.slice());e=[];t=this.getElementsByTagName("thead");0!==t.length&&(da(p.aoHeader,t[0]),e=ra(p));if(null===g.aoColumns){t=[];j=0;for(i=e.length;j<i;j++)t.push(null)}else t=g.aoColumns;j=0;for(i=t.length;j<i;j++)Da(p,e?e[j]:null);hb(p,g.aoColumnDefs,t,function(a,b){ja(p,a,b)});if(x.length){var w=function(a,b){return a.getAttribute("data-"+b)!==null?b:null};h(x[0]).children("th, td").each(function(a,b){var c=p.aoColumns[a];if(c.mData===
a){var d=w(b,"sort")||w(b,"order"),e=w(b,"filter")||w(b,"search");if(d!==null||e!==null){c.mData={_:a+".display",sort:d!==null?a+".@data-"+d:k,type:d!==null?a+".@data-"+d:k,filter:e!==null?a+".@data-"+e:k};ja(p,a)}}})}var T=p.oFeatures,e=function(){if(g.aaSorting===k){var a=p.aaSorting;j=0;for(i=a.length;j<i;j++)a[j][1]=p.aoColumns[j].asSorting[0]}wa(p);T.bSort&&z(p,"aoDrawCallback",function(){if(p.bSorted){var a=V(p),b={};h.each(a,function(a,c){b[c.src]=c.dir});r(p,null,"order",[p,a,b]);Hb(p)}});
z(p,"aoDrawCallback",function(){(p.bSorted||y(p)==="ssp"||T.bDeferRender)&&wa(p)},"sc");var a=q.children("caption").each(function(){this._captionSide=h(this).css("caption-side")}),b=q.children("thead");b.length===0&&(b=h("<thead/>").appendTo(q));p.nTHead=b[0];b=q.children("tbody");b.length===0&&(b=h("<tbody/>").appendTo(q));p.nTBody=b[0];b=q.children("tfoot");if(b.length===0&&a.length>0&&(p.oScroll.sX!==""||p.oScroll.sY!==""))b=h("<tfoot/>").appendTo(q);if(b.length===0||b.children().length===0)q.addClass(u.sNoFooter);
else if(b.length>0){p.nTFoot=b[0];da(p.aoFooter,p.nTFoot)}if(g.aaData)for(j=0;j<g.aaData.length;j++)M(p,g.aaData[j]);else(p.bDeferLoading||y(p)=="dom")&&ma(p,h(p.nTBody).children("tr"));p.aiDisplay=p.aiDisplayMaster.slice();p.bInitialised=true;n===false&&ga(p)};g.bStateSave?(T.bStateSave=!0,z(p,"aoDrawCallback",xa,"state_save"),Ib(p,g,e)):e()}});b=null;return this},x,s,o,u,Xa={},Mb=/[\r\n]/g,Aa=/<.*?>/g,Zb=/^\d{2,4}[\.\/\-]\d{1,2}[\.\/\-]\d{1,2}([T ]{1}\d{1,2}[:\.]\d{2}([\.:]\d{2})?)?$/,$b=RegExp("(\\/|\\.|\\*|\\+|\\?|\\||\\(|\\)|\\[|\\]|\\{|\\}|\\\\|\\$|\\^|\\-)",
"g"),Wa=/[',$£€¥%\u2009\u202F\u20BD\u20a9\u20BArfk]/gi,L=function(a){return!a||!0===a||"-"===a?!0:!1},Nb=function(a){var b=parseInt(a,10);return!isNaN(b)&&isFinite(a)?b:null},Ob=function(a,b){Xa[b]||(Xa[b]=RegExp(Pa(b),"g"));return"string"===typeof a&&"."!==b?a.replace(/\./g,"").replace(Xa[b],"."):a},Ya=function(a,b,c){var d="string"===typeof a;if(L(a))return!0;b&&d&&(a=Ob(a,b));c&&d&&(a=a.replace(Wa,""));return!isNaN(parseFloat(a))&&isFinite(a)},Pb=function(a,b,c){return L(a)?!0:!(L(a)||"string"===
typeof a)?null:Ya(a.replace(Aa,""),b,c)?!0:null},D=function(a,b,c){var d=[],e=0,f=a.length;if(c!==k)for(;e<f;e++)a[e]&&a[e][b]&&d.push(a[e][b][c]);else for(;e<f;e++)a[e]&&d.push(a[e][b]);return d},ia=function(a,b,c,d){var e=[],f=0,g=b.length;if(d!==k)for(;f<g;f++)a[b[f]][c]&&e.push(a[b[f]][c][d]);else for(;f<g;f++)e.push(a[b[f]][c]);return e},W=function(a,b){var c=[],d;b===k?(b=0,d=a):(d=b,b=a);for(var e=b;e<d;e++)c.push(e);return c},Qb=function(a){for(var b=[],c=0,d=a.length;c<d;c++)a[c]&&b.push(a[c]);
return b},qa=function(a){var b;a:{if(!(2>a.length)){b=a.slice().sort();for(var c=b[0],d=1,e=b.length;d<e;d++){if(b[d]===c){b=!1;break a}c=b[d]}}b=!0}if(b)return a.slice();b=[];var e=a.length,f,g=0,d=0;a:for(;d<e;d++){c=a[d];for(f=0;f<g;f++)if(b[f]===c)continue a;b.push(c);g++}return b};m.util={throttle:function(a,b){var c=b!==k?b:200,d,e;return function(){var b=this,g=+new Date,j=arguments;d&&g<d+c?(clearTimeout(e),e=setTimeout(function(){d=k;a.apply(b,j)},c)):(d=g,a.apply(b,j))}},escapeRegex:function(a){return a.replace($b,
"\\$1")}};var A=function(a,b,c){a[b]!==k&&(a[c]=a[b])},ba=/\[.*?\]$/,U=/\(\)$/,Pa=m.util.escapeRegex,va=h("<div>")[0],Wb=va.textContent!==k,Yb=/<.*?>/g,Na=m.util.throttle,Rb=[],w=Array.prototype,ac=function(a){var b,c,d=m.settings,e=h.map(d,function(a){return a.nTable});if(a){if(a.nTable&&a.oApi)return[a];if(a.nodeName&&"table"===a.nodeName.toLowerCase())return b=h.inArray(a,e),-1!==b?[d[b]]:null;if(a&&"function"===typeof a.settings)return a.settings().toArray();"string"===typeof a?c=h(a):a instanceof
h&&(c=a)}else return[];if(c)return c.map(function(){b=h.inArray(this,e);return-1!==b?d[b]:null}).toArray()};s=function(a,b){if(!(this instanceof s))return new s(a,b);var c=[],d=function(a){(a=ac(a))&&(c=c.concat(a))};if(h.isArray(a))for(var e=0,f=a.length;e<f;e++)d(a[e]);else d(a);this.context=qa(c);b&&h.merge(this,b);this.selector={rows:null,cols:null,opts:null};s.extend(this,this,Rb)};m.Api=s;h.extend(s.prototype,{any:function(){return 0!==this.count()},concat:w.concat,context:[],count:function(){return this.flatten().length},
each:function(a){for(var b=0,c=this.length;b<c;b++)a.call(this,this[b],b,this);return this},eq:function(a){var b=this.context;return b.length>a?new s(b[a],this[a]):null},filter:function(a){var b=[];if(w.filter)b=w.filter.call(this,a,this);else for(var c=0,d=this.length;c<d;c++)a.call(this,this[c],c,this)&&b.push(this[c]);return new s(this.context,b)},flatten:function(){var a=[];return new s(this.context,a.concat.apply(a,this.toArray()))},join:w.join,indexOf:w.indexOf||function(a,b){for(var c=b||0,
d=this.length;c<d;c++)if(this[c]===a)return c;return-1},iterator:function(a,b,c,d){var e=[],f,g,j,h,n,l=this.context,m,o,u=this.selector;"string"===typeof a&&(d=c,c=b,b=a,a=!1);g=0;for(j=l.length;g<j;g++){var r=new s(l[g]);if("table"===b)f=c.call(r,l[g],g),f!==k&&e.push(f);else if("columns"===b||"rows"===b)f=c.call(r,l[g],this[g],g),f!==k&&e.push(f);else if("column"===b||"column-rows"===b||"row"===b||"cell"===b){o=this[g];"column-rows"===b&&(m=Ba(l[g],u.opts));h=0;for(n=o.length;h<n;h++)f=o[h],f=
"cell"===b?c.call(r,l[g],f.row,f.column,g,h):c.call(r,l[g],f,g,h,m),f!==k&&e.push(f)}}return e.length||d?(a=new s(l,a?e.concat.apply([],e):e),b=a.selector,b.rows=u.rows,b.cols=u.cols,b.opts=u.opts,a):this},lastIndexOf:w.lastIndexOf||function(a,b){return this.indexOf.apply(this.toArray.reverse(),arguments)},length:0,map:function(a){var b=[];if(w.map)b=w.map.call(this,a,this);else for(var c=0,d=this.length;c<d;c++)b.push(a.call(this,this[c],c));return new s(this.context,b)},pluck:function(a){return this.map(function(b){return b[a]})},
pop:w.pop,push:w.push,reduce:w.reduce||function(a,b){return gb(this,a,b,0,this.length,1)},reduceRight:w.reduceRight||function(a,b){return gb(this,a,b,this.length-1,-1,-1)},reverse:w.reverse,selector:null,shift:w.shift,slice:function(){return new s(this.context,this)},sort:w.sort,splice:w.splice,toArray:function(){return w.slice.call(this)},to$:function(){return h(this)},toJQuery:function(){return h(this)},unique:function(){return new s(this.context,qa(this))},unshift:w.unshift});s.extend=function(a,
b,c){if(c.length&&b&&(b instanceof s||b.__dt_wrapper)){var d,e,f,g=function(a,b,c){return function(){var d=b.apply(a,arguments);s.extend(d,d,c.methodExt);return d}};d=0;for(e=c.length;d<e;d++)f=c[d],b[f.name]="function"===typeof f.val?g(a,f.val,f):h.isPlainObject(f.val)?{}:f.val,b[f.name].__dt_wrapper=!0,s.extend(a,b[f.name],f.propExt)}};s.register=o=function(a,b){if(h.isArray(a))for(var c=0,d=a.length;c<d;c++)s.register(a[c],b);else for(var e=a.split("."),f=Rb,g,j,c=0,d=e.length;c<d;c++){g=(j=-1!==
e[c].indexOf("()"))?e[c].replace("()",""):e[c];var i;a:{i=0;for(var n=f.length;i<n;i++)if(f[i].name===g){i=f[i];break a}i=null}i||(i={name:g,val:{},methodExt:[],propExt:[]},f.push(i));c===d-1?i.val=b:f=j?i.methodExt:i.propExt}};s.registerPlural=u=function(a,b,c){s.register(a,c);s.register(b,function(){var a=c.apply(this,arguments);return a===this?this:a instanceof s?a.length?h.isArray(a[0])?new s(a.context,a[0]):a[0]:k:a})};o("tables()",function(a){var b;if(a){b=s;var c=this.context;if("number"===
typeof a)a=[c[a]];else var d=h.map(c,function(a){return a.nTable}),a=h(d).filter(a).map(function(){var a=h.inArray(this,d);return c[a]}).toArray();b=new b(a)}else b=this;return b});o("table()",function(a){var a=this.tables(a),b=a.context;return b.length?new s(b[0]):a});u("tables().nodes()","table().node()",function(){return this.iterator("table",function(a){return a.nTable},1)});u("tables().body()","table().body()",function(){return this.iterator("table",function(a){return a.nTBody},1)});u("tables().header()",
"table().header()",function(){return this.iterator("table",function(a){return a.nTHead},1)});u("tables().footer()","table().footer()",function(){return this.iterator("table",function(a){return a.nTFoot},1)});u("tables().containers()","table().container()",function(){return this.iterator("table",function(a){return a.nTableWrapper},1)});o("draw()",function(a){return this.iterator("table",function(b){"page"===a?N(b):("string"===typeof a&&(a="full-hold"===a?!1:!0),S(b,!1===a))})});o("page()",function(a){return a===
k?this.page.info().page:this.iterator("table",function(b){Sa(b,a)})});o("page.info()",function(){if(0===this.context.length)return k;var a=this.context[0],b=a._iDisplayStart,c=a.oFeatures.bPaginate?a._iDisplayLength:-1,d=a.fnRecordsDisplay(),e=-1===c;return{page:e?0:Math.floor(b/c),pages:e?1:Math.ceil(d/c),start:b,end:a.fnDisplayEnd(),length:c,recordsTotal:a.fnRecordsTotal(),recordsDisplay:d,serverSide:"ssp"===y(a)}});o("page.len()",function(a){return a===k?0!==this.context.length?this.context[0]._iDisplayLength:
k:this.iterator("table",function(b){Qa(b,a)})});var Sb=function(a,b,c){if(c){var d=new s(a);d.one("draw",function(){c(d.ajax.json())})}if("ssp"==y(a))S(a,b);else{C(a,!0);var e=a.jqXHR;e&&4!==e.readyState&&e.abort();sa(a,[],function(c){na(a);for(var c=ta(a,c),d=0,e=c.length;d<e;d++)M(a,c[d]);S(a,b);C(a,!1)})}};o("ajax.json()",function(){var a=this.context;if(0<a.length)return a[0].json});o("ajax.params()",function(){var a=this.context;if(0<a.length)return a[0].oAjaxData});o("ajax.reload()",function(a,
b){return this.iterator("table",function(c){Sb(c,!1===b,a)})});o("ajax.url()",function(a){var b=this.context;if(a===k){if(0===b.length)return k;b=b[0];return b.ajax?h.isPlainObject(b.ajax)?b.ajax.url:b.ajax:b.sAjaxSource}return this.iterator("table",function(b){h.isPlainObject(b.ajax)?b.ajax.url=a:b.ajax=a})});o("ajax.url().load()",function(a,b){return this.iterator("table",function(c){Sb(c,!1===b,a)})});var Za=function(a,b,c,d,e){var f=[],g,j,i,n,l,m;i=typeof b;if(!b||"string"===i||"function"===
i||b.length===k)b=[b];i=0;for(n=b.length;i<n;i++){j=b[i]&&b[i].split&&!b[i].match(/[\[\(:]/)?b[i].split(","):[b[i]];l=0;for(m=j.length;l<m;l++)(g=c("string"===typeof j[l]?h.trim(j[l]):j[l]))&&g.length&&(f=f.concat(g))}a=x.selector[a];if(a.length){i=0;for(n=a.length;i<n;i++)f=a[i](d,e,f)}return qa(f)},$a=function(a){a||(a={});a.filter&&a.search===k&&(a.search=a.filter);return h.extend({search:"none",order:"current",page:"all"},a)},ab=function(a){for(var b=0,c=a.length;b<c;b++)if(0<a[b].length)return a[0]=
a[b],a[0].length=1,a.length=1,a.context=[a.context[b]],a;a.length=0;return a},Ba=function(a,b){var c,d,e,f=[],g=a.aiDisplay;c=a.aiDisplayMaster;var j=b.search;d=b.order;e=b.page;if("ssp"==y(a))return"removed"===j?[]:W(0,c.length);if("current"==e){c=a._iDisplayStart;for(d=a.fnDisplayEnd();c<d;c++)f.push(g[c])}else if("current"==d||"applied"==d)f="none"==j?c.slice():"applied"==j?g.slice():h.map(c,function(a){return-1===h.inArray(a,g)?a:null});else if("index"==d||"original"==d){c=0;for(d=a.aoData.length;c<
d;c++)"none"==j?f.push(c):(e=h.inArray(c,g),(-1===e&&"removed"==j||0<=e&&"applied"==j)&&f.push(c))}return f};o("rows()",function(a,b){a===k?a="":h.isPlainObject(a)&&(b=a,a="");var b=$a(b),c=this.iterator("table",function(c){var e=b,f;return Za("row",a,function(a){var b=Nb(a);if(b!==null&&!e)return[b];f||(f=Ba(c,e));if(b!==null&&h.inArray(b,f)!==-1)return[b];if(a===null||a===k||a==="")return f;if(typeof a==="function")return h.map(f,function(b){var e=c.aoData[b];return a(b,e._aData,e.nTr)?b:null});
b=Qb(ia(c.aoData,f,"nTr"));if(a.nodeName){if(a._DT_RowIndex!==k)return[a._DT_RowIndex];if(a._DT_CellIndex)return[a._DT_CellIndex.row];b=h(a).closest("*[data-dt-row]");return b.length?[b.data("dt-row")]:[]}if(typeof a==="string"&&a.charAt(0)==="#"){var i=c.aIds[a.replace(/^#/,"")];if(i!==k)return[i.idx]}return h(b).filter(a).map(function(){return this._DT_RowIndex}).toArray()},c,e)},1);c.selector.rows=a;c.selector.opts=b;return c});o("rows().nodes()",function(){return this.iterator("row",function(a,
b){return a.aoData[b].nTr||k},1)});o("rows().data()",function(){return this.iterator(!0,"rows",function(a,b){return ia(a.aoData,b,"_aData")},1)});u("rows().cache()","row().cache()",function(a){return this.iterator("row",function(b,c){var d=b.aoData[c];return"search"===a?d._aFilterData:d._aSortData},1)});u("rows().invalidate()","row().invalidate()",function(a){return this.iterator("row",function(b,c){ca(b,c,a)})});u("rows().indexes()","row().index()",function(){return this.iterator("row",function(a,
b){return b},1)});u("rows().ids()","row().id()",function(a){for(var b=[],c=this.context,d=0,e=c.length;d<e;d++)for(var f=0,g=this[d].length;f<g;f++){var h=c[d].rowIdFn(c[d].aoData[this[d][f]]._aData);b.push((!0===a?"#":"")+h)}return new s(c,b)});u("rows().remove()","row().remove()",function(){var a=this;this.iterator("row",function(b,c,d){var e=b.aoData,f=e[c],g,h,i,n,l;e.splice(c,1);g=0;for(h=e.length;g<h;g++)if(i=e[g],l=i.anCells,null!==i.nTr&&(i.nTr._DT_RowIndex=g),null!==l){i=0;for(n=l.length;i<
n;i++)l[i]._DT_CellIndex.row=g}oa(b.aiDisplayMaster,c);oa(b.aiDisplay,c);oa(a[d],c,!1);0<b._iRecordsDisplay&&b._iRecordsDisplay--;Ra(b);c=b.rowIdFn(f._aData);c!==k&&delete b.aIds[c]});this.iterator("table",function(a){for(var c=0,d=a.aoData.length;c<d;c++)a.aoData[c].idx=c});return this});o("rows.add()",function(a){var b=this.iterator("table",function(b){var c,f,g,h=[];f=0;for(g=a.length;f<g;f++)c=a[f],c.nodeName&&"TR"===c.nodeName.toUpperCase()?h.push(ma(b,c)[0]):h.push(M(b,c));return h},1),c=this.rows(-1);
c.pop();h.merge(c,b);return c});o("row()",function(a,b){return ab(this.rows(a,b))});o("row().data()",function(a){var b=this.context;if(a===k)return b.length&&this.length?b[0].aoData[this[0]]._aData:k;b[0].aoData[this[0]]._aData=a;ca(b[0],this[0],"data");return this});o("row().node()",function(){var a=this.context;return a.length&&this.length?a[0].aoData[this[0]].nTr||null:null});o("row.add()",function(a){a instanceof h&&a.length&&(a=a[0]);var b=this.iterator("table",function(b){return a.nodeName&&
"TR"===a.nodeName.toUpperCase()?ma(b,a)[0]:M(b,a)});return this.row(b[0])});var bb=function(a,b){var c=a.context;if(c.length&&(c=c[0].aoData[b!==k?b:a[0]])&&c._details)c._details.remove(),c._detailsShow=k,c._details=k},Tb=function(a,b){var c=a.context;if(c.length&&a.length){var d=c[0].aoData[a[0]];if(d._details){(d._detailsShow=b)?d._details.insertAfter(d.nTr):d._details.detach();var e=c[0],f=new s(e),g=e.aoData;f.off("draw.dt.DT_details column-visibility.dt.DT_details destroy.dt.DT_details");0<D(g,
"_details").length&&(f.on("draw.dt.DT_details",function(a,b){e===b&&f.rows({page:"current"}).eq(0).each(function(a){a=g[a];a._detailsShow&&a._details.insertAfter(a.nTr)})}),f.on("column-visibility.dt.DT_details",function(a,b){if(e===b)for(var c,d=aa(b),f=0,h=g.length;f<h;f++)c=g[f],c._details&&c._details.children("td[colspan]").attr("colspan",d)}),f.on("destroy.dt.DT_details",function(a,b){if(e===b)for(var c=0,d=g.length;c<d;c++)g[c]._details&&bb(f,c)}))}}};o("row().child()",function(a,b){var c=this.context;
if(a===k)return c.length&&this.length?c[0].aoData[this[0]]._details:k;if(!0===a)this.child.show();else if(!1===a)bb(this);else if(c.length&&this.length){var d=c[0],c=c[0].aoData[this[0]],e=[],f=function(a,b){if(h.isArray(a)||a instanceof h)for(var c=0,k=a.length;c<k;c++)f(a[c],b);else a.nodeName&&"tr"===a.nodeName.toLowerCase()?e.push(a):(c=h("<tr><td/></tr>").addClass(b),h("td",c).addClass(b).html(a)[0].colSpan=aa(d),e.push(c[0]))};f(a,b);c._details&&c._details.detach();c._details=h(e);c._detailsShow&&
c._details.insertAfter(c.nTr)}return this});o(["row().child.show()","row().child().show()"],function(){Tb(this,!0);return this});o(["row().child.hide()","row().child().hide()"],function(){Tb(this,!1);return this});o(["row().child.remove()","row().child().remove()"],function(){bb(this);return this});o("row().child.isShown()",function(){var a=this.context;return a.length&&this.length?a[0].aoData[this[0]]._detailsShow||!1:!1});var bc=/^([^:]+):(name|visIdx|visible)$/,Ub=function(a,b,c,d,e){for(var c=
[],d=0,f=e.length;d<f;d++)c.push(B(a,e[d],b));return c};o("columns()",function(a,b){a===k?a="":h.isPlainObject(a)&&(b=a,a="");var b=$a(b),c=this.iterator("table",function(c){var e=a,f=b,g=c.aoColumns,j=D(g,"sName"),i=D(g,"nTh");return Za("column",e,function(a){var b=Nb(a);if(a==="")return W(g.length);if(b!==null)return[b>=0?b:g.length+b];if(typeof a==="function"){var e=Ba(c,f);return h.map(g,function(b,f){return a(f,Ub(c,f,0,0,e),i[f])?f:null})}var k=typeof a==="string"?a.match(bc):"";if(k)switch(k[2]){case "visIdx":case "visible":b=
parseInt(k[1],10);if(b<0){var m=h.map(g,function(a,b){return a.bVisible?b:null});return[m[m.length+b]]}return[Z(c,b)];case "name":return h.map(j,function(a,b){return a===k[1]?b:null});default:return[]}if(a.nodeName&&a._DT_CellIndex)return[a._DT_CellIndex.column];b=h(i).filter(a).map(function(){return h.inArray(this,i)}).toArray();if(b.length||!a.nodeName)return b;b=h(a).closest("*[data-dt-column]");return b.length?[b.data("dt-column")]:[]},c,f)},1);c.selector.cols=a;c.selector.opts=b;return c});u("columns().header()",
"column().header()",function(){return this.iterator("column",function(a,b){return a.aoColumns[b].nTh},1)});u("columns().footer()","column().footer()",function(){return this.iterator("column",function(a,b){return a.aoColumns[b].nTf},1)});u("columns().data()","column().data()",function(){return this.iterator("column-rows",Ub,1)});u("columns().dataSrc()","column().dataSrc()",function(){return this.iterator("column",function(a,b){return a.aoColumns[b].mData},1)});u("columns().cache()","column().cache()",
function(a){return this.iterator("column-rows",function(b,c,d,e,f){return ia(b.aoData,f,"search"===a?"_aFilterData":"_aSortData",c)},1)});u("columns().nodes()","column().nodes()",function(){return this.iterator("column-rows",function(a,b,c,d,e){return ia(a.aoData,e,"anCells",b)},1)});u("columns().visible()","column().visible()",function(a,b){var c=this.iterator("column",function(b,c){if(a===k)return b.aoColumns[c].bVisible;var f=b.aoColumns,g=f[c],j=b.aoData,i,n,l;if(a!==k&&g.bVisible!==a){if(a){var m=
h.inArray(!0,D(f,"bVisible"),c+1);i=0;for(n=j.length;i<n;i++)l=j[i].nTr,f=j[i].anCells,l&&l.insertBefore(f[c],f[m]||null)}else h(D(b.aoData,"anCells",c)).detach();g.bVisible=a;ea(b,b.aoHeader);ea(b,b.aoFooter);xa(b)}});a!==k&&(this.iterator("column",function(c,e){r(c,null,"column-visibility",[c,e,a,b])}),(b===k||b)&&this.columns.adjust());return c});u("columns().indexes()","column().index()",function(a){return this.iterator("column",function(b,c){return"visible"===a?$(b,c):c},1)});o("columns.adjust()",
function(){return this.iterator("table",function(a){Y(a)},1)});o("column.index()",function(a,b){if(0!==this.context.length){var c=this.context[0];if("fromVisible"===a||"toData"===a)return Z(c,b);if("fromData"===a||"toVisible"===a)return $(c,b)}});o("column()",function(a,b){return ab(this.columns(a,b))});o("cells()",function(a,b,c){h.isPlainObject(a)&&(a.row===k?(c=a,a=null):(c=b,b=null));h.isPlainObject(b)&&(c=b,b=null);if(null===b||b===k)return this.iterator("table",function(b){var d=a,e=$a(c),f=
b.aoData,g=Ba(b,e),j=Qb(ia(f,g,"anCells")),i=h([].concat.apply([],j)),l,n=b.aoColumns.length,m,o,u,s,r,v;return Za("cell",d,function(a){var c=typeof a==="function";if(a===null||a===k||c){m=[];o=0;for(u=g.length;o<u;o++){l=g[o];for(s=0;s<n;s++){r={row:l,column:s};if(c){v=f[l];a(r,B(b,l,s),v.anCells?v.anCells[s]:null)&&m.push(r)}else m.push(r)}}return m}if(h.isPlainObject(a))return[a];c=i.filter(a).map(function(a,b){return{row:b._DT_CellIndex.row,column:b._DT_CellIndex.column}}).toArray();if(c.length||
!a.nodeName)return c;v=h(a).closest("*[data-dt-row]");return v.length?[{row:v.data("dt-row"),column:v.data("dt-column")}]:[]},b,e)});var d=this.columns(b,c),e=this.rows(a,c),f,g,j,i,n,l=this.iterator("table",function(a,b){f=[];g=0;for(j=e[b].length;g<j;g++){i=0;for(n=d[b].length;i<n;i++)f.push({row:e[b][g],column:d[b][i]})}return f},1);h.extend(l.selector,{cols:b,rows:a,opts:c});return l});u("cells().nodes()","cell().node()",function(){return this.iterator("cell",function(a,b,c){return(a=a.aoData[b])&&
a.anCells?a.anCells[c]:k},1)});o("cells().data()",function(){return this.iterator("cell",function(a,b,c){return B(a,b,c)},1)});u("cells().cache()","cell().cache()",function(a){a="search"===a?"_aFilterData":"_aSortData";return this.iterator("cell",function(b,c,d){return b.aoData[c][a][d]},1)});u("cells().render()","cell().render()",function(a){return this.iterator("cell",function(b,c,d){return B(b,c,d,a)},1)});u("cells().indexes()","cell().index()",function(){return this.iterator("cell",function(a,
b,c){return{row:b,column:c,columnVisible:$(a,c)}},1)});u("cells().invalidate()","cell().invalidate()",function(a){return this.iterator("cell",function(b,c,d){ca(b,c,a,d)})});o("cell()",function(a,b,c){return ab(this.cells(a,b,c))});o("cell().data()",function(a){var b=this.context,c=this[0];if(a===k)return b.length&&c.length?B(b[0],c[0].row,c[0].column):k;ib(b[0],c[0].row,c[0].column,a);ca(b[0],c[0].row,"data",c[0].column);return this});o("order()",function(a,b){var c=this.context;if(a===k)return 0!==
c.length?c[0].aaSorting:k;"number"===typeof a?a=[[a,b]]:a.length&&!h.isArray(a[0])&&(a=Array.prototype.slice.call(arguments));return this.iterator("table",function(b){b.aaSorting=a.slice()})});o("order.listener()",function(a,b,c){return this.iterator("table",function(d){La(d,a,b,c)})});o("order.fixed()",function(a){if(!a){var b=this.context,b=b.length?b[0].aaSortingFixed:k;return h.isArray(b)?{pre:b}:b}return this.iterator("table",function(b){b.aaSortingFixed=h.extend(!0,{},a)})});o(["columns().order()",
"column().order()"],function(a){var b=this;return this.iterator("table",function(c,d){var e=[];h.each(b[d],function(b,c){e.push([c,a])});c.aaSorting=e})});o("search()",function(a,b,c,d){var e=this.context;return a===k?0!==e.length?e[0].oPreviousSearch.sSearch:k:this.iterator("table",function(e){e.oFeatures.bFilter&&fa(e,h.extend({},e.oPreviousSearch,{sSearch:a+"",bRegex:null===b?!1:b,bSmart:null===c?!0:c,bCaseInsensitive:null===d?!0:d}),1)})});u("columns().search()","column().search()",function(a,
b,c,d){return this.iterator("column",function(e,f){var g=e.aoPreSearchCols;if(a===k)return g[f].sSearch;e.oFeatures.bFilter&&(h.extend(g[f],{sSearch:a+"",bRegex:null===b?!1:b,bSmart:null===c?!0:c,bCaseInsensitive:null===d?!0:d}),fa(e,e.oPreviousSearch,1))})});o("state()",function(){return this.context.length?this.context[0].oSavedState:null});o("state.clear()",function(){return this.iterator("table",function(a){a.fnStateSaveCallback.call(a.oInstance,a,{})})});o("state.loaded()",function(){return this.context.length?
this.context[0].oLoadedState:null});o("state.save()",function(){return this.iterator("table",function(a){xa(a)})});m.versionCheck=m.fnVersionCheck=function(a){for(var b=m.version.split("."),a=a.split("."),c,d,e=0,f=a.length;e<f;e++)if(c=parseInt(b[e],10)||0,d=parseInt(a[e],10)||0,c!==d)return c>d;return!0};m.isDataTable=m.fnIsDataTable=function(a){var b=h(a).get(0),c=!1;if(a instanceof m.Api)return!0;h.each(m.settings,function(a,e){var f=e.nScrollHead?h("table",e.nScrollHead)[0]:null,g=e.nScrollFoot?
h("table",e.nScrollFoot)[0]:null;if(e.nTable===b||f===b||g===b)c=!0});return c};m.tables=m.fnTables=function(a){var b=!1;h.isPlainObject(a)&&(b=a.api,a=a.visible);var c=h.map(m.settings,function(b){if(!a||a&&h(b.nTable).is(":visible"))return b.nTable});return b?new s(c):c};m.camelToHungarian=I;o("$()",function(a,b){var c=this.rows(b).nodes(),c=h(c);return h([].concat(c.filter(a).toArray(),c.find(a).toArray()))});h.each(["on","one","off"],function(a,b){o(b+"()",function(){var a=Array.prototype.slice.call(arguments);
a[0]=h.map(a[0].split(/\s/),function(a){return!a.match(/\.dt\b/)?a+".dt":a}).join(" ");var d=h(this.tables().nodes());d[b].apply(d,a);return this})});o("clear()",function(){return this.iterator("table",function(a){na(a)})});o("settings()",function(){return new s(this.context,this.context)});o("init()",function(){var a=this.context;return a.length?a[0].oInit:null});o("data()",function(){return this.iterator("table",function(a){return D(a.aoData,"_aData")}).flatten()});o("destroy()",function(a){a=a||
!1;return this.iterator("table",function(b){var c=b.nTableWrapper.parentNode,d=b.oClasses,e=b.nTable,f=b.nTBody,g=b.nTHead,j=b.nTFoot,i=h(e),f=h(f),k=h(b.nTableWrapper),l=h.map(b.aoData,function(a){return a.nTr}),o;b.bDestroying=!0;r(b,"aoDestroyCallback","destroy",[b]);a||(new s(b)).columns().visible(!0);k.off(".DT").find(":not(tbody *)").off(".DT");h(E).off(".DT-"+b.sInstance);e!=g.parentNode&&(i.children("thead").detach(),i.append(g));j&&e!=j.parentNode&&(i.children("tfoot").detach(),i.append(j));
b.aaSorting=[];b.aaSortingFixed=[];wa(b);h(l).removeClass(b.asStripeClasses.join(" "));h("th, td",g).removeClass(d.sSortable+" "+d.sSortableAsc+" "+d.sSortableDesc+" "+d.sSortableNone);f.children().detach();f.append(l);g=a?"remove":"detach";i[g]();k[g]();!a&&c&&(c.insertBefore(e,b.nTableReinsertBefore),i.css("width",b.sDestroyWidth).removeClass(d.sTable),(o=b.asDestroyStripes.length)&&f.children().each(function(a){h(this).addClass(b.asDestroyStripes[a%o])}));c=h.inArray(b,m.settings);-1!==c&&m.settings.splice(c,
1)})});h.each(["column","row","cell"],function(a,b){o(b+"s().every()",function(a){var d=this.selector.opts,e=this;return this.iterator(b,function(f,g,h,i,n){a.call(e[b](g,"cell"===b?h:d,"cell"===b?d:k),g,h,i,n)})})});o("i18n()",function(a,b,c){var d=this.context[0],a=Q(a)(d.oLanguage);a===k&&(a=b);c!==k&&h.isPlainObject(a)&&(a=a[c]!==k?a[c]:a._);return a.replace("%d",c)});m.version="1.10.16";m.settings=[];m.models={};m.models.oSearch={bCaseInsensitive:!0,sSearch:"",bRegex:!1,bSmart:!0};m.models.oRow=
{nTr:null,anCells:null,_aData:[],_aSortData:null,_aFilterData:null,_sFilterRow:null,_sRowStripe:"",src:null,idx:-1};m.models.oColumn={idx:null,aDataSort:null,asSorting:null,bSearchable:null,bSortable:null,bVisible:null,_sManualType:null,_bAttrSrc:!1,fnCreatedCell:null,fnGetData:null,fnSetData:null,mData:null,mRender:null,nTh:null,nTf:null,sClass:null,sContentPadding:null,sDefaultContent:null,sName:null,sSortDataType:"std",sSortingClass:null,sSortingClassJUI:null,sTitle:null,sType:null,sWidth:null,
sWidthOrig:null};m.defaults={aaData:null,aaSorting:[[0,"asc"]],aaSortingFixed:[],ajax:null,aLengthMenu:[10,25,50,100],aoColumns:null,aoColumnDefs:null,aoSearchCols:[],asStripeClasses:null,bAutoWidth:!0,bDeferRender:!1,bDestroy:!1,bFilter:!0,bInfo:!0,bLengthChange:!0,bPaginate:!0,bProcessing:!1,bRetrieve:!1,bScrollCollapse:!1,bServerSide:!1,bSort:!0,bSortMulti:!0,bSortCellsTop:!1,bSortClasses:!0,bStateSave:!1,fnCreatedRow:null,fnDrawCallback:null,fnFooterCallback:null,fnFormatNumber:function(a){return a.toString().replace(/\B(?=(\d{3})+(?!\d))/g,
this.oLanguage.sThousands)},fnHeaderCallback:null,fnInfoCallback:null,fnInitComplete:null,fnPreDrawCallback:null,fnRowCallback:null,fnServerData:null,fnServerParams:null,fnStateLoadCallback:function(a){try{return JSON.parse((-1===a.iStateDuration?sessionStorage:localStorage).getItem("DataTables_"+a.sInstance+"_"+location.pathname))}catch(b){}},fnStateLoadParams:null,fnStateLoaded:null,fnStateSaveCallback:function(a,b){try{(-1===a.iStateDuration?sessionStorage:localStorage).setItem("DataTables_"+a.sInstance+
"_"+location.pathname,JSON.stringify(b))}catch(c){}},fnStateSaveParams:null,iStateDuration:7200,iDeferLoading:null,iDisplayLength:10,iDisplayStart:0,iTabIndex:0,oClasses:{},oLanguage:{oAria:{sSortAscending:": activate to sort column ascending",sSortDescending:": activate to sort column descending"},oPaginate:{sFirst:"First",sLast:"Last",sNext:"Next",sPrevious:"Previous"},sEmptyTable:"No data available in table",sInfo:"Showing _START_ to _END_ of _TOTAL_ entries",sInfoEmpty:"Showing 0 to 0 of 0 entries",
sInfoFiltered:"(filtered from _MAX_ total entries)",sInfoPostFix:"",sDecimal:"",sThousands:",",sLengthMenu:"Show _MENU_ entries",sLoadingRecords:"Loading...",sProcessing:"Processing...",sSearch:"Search:",sSearchPlaceholder:"",sUrl:"",sZeroRecords:"No matching records found"},oSearch:h.extend({},m.models.oSearch),sAjaxDataProp:"data",sAjaxSource:null,sDom:"lfrtip",searchDelay:null,sPaginationType:"simple_numbers",sScrollX:"",sScrollXInner:"",sScrollY:"",sServerMethod:"GET",renderer:null,rowId:"DT_RowId"};
X(m.defaults);m.defaults.column={aDataSort:null,iDataSort:-1,asSorting:["asc","desc"],bSearchable:!0,bSortable:!0,bVisible:!0,fnCreatedCell:null,mData:null,mRender:null,sCellType:"td",sClass:"",sContentPadding:"",sDefaultContent:null,sName:"",sSortDataType:"std",sTitle:null,sType:null,sWidth:null};X(m.defaults.column);m.models.oSettings={oFeatures:{bAutoWidth:null,bDeferRender:null,bFilter:null,bInfo:null,bLengthChange:null,bPaginate:null,bProcessing:null,bServerSide:null,bSort:null,bSortMulti:null,
bSortClasses:null,bStateSave:null},oScroll:{bCollapse:null,iBarWidth:0,sX:null,sXInner:null,sY:null},oLanguage:{fnInfoCallback:null},oBrowser:{bScrollOversize:!1,bScrollbarLeft:!1,bBounding:!1,barWidth:0},ajax:null,aanFeatures:[],aoData:[],aiDisplay:[],aiDisplayMaster:[],aIds:{},aoColumns:[],aoHeader:[],aoFooter:[],oPreviousSearch:{},aoPreSearchCols:[],aaSorting:null,aaSortingFixed:[],asStripeClasses:null,asDestroyStripes:[],sDestroyWidth:0,aoRowCallback:[],aoHeaderCallback:[],aoFooterCallback:[],
aoDrawCallback:[],aoRowCreatedCallback:[],aoPreDrawCallback:[],aoInitComplete:[],aoStateSaveParams:[],aoStateLoadParams:[],aoStateLoaded:[],sTableId:"",nTable:null,nTHead:null,nTFoot:null,nTBody:null,nTableWrapper:null,bDeferLoading:!1,bInitialised:!1,aoOpenRows:[],sDom:null,searchDelay:null,sPaginationType:"two_button",iStateDuration:0,aoStateSave:[],aoStateLoad:[],oSavedState:null,oLoadedState:null,sAjaxSource:null,sAjaxDataProp:null,bAjaxDataGet:!0,jqXHR:null,json:k,oAjaxData:k,fnServerData:null,
aoServerParams:[],sServerMethod:null,fnFormatNumber:null,aLengthMenu:null,iDraw:0,bDrawing:!1,iDrawError:-1,_iDisplayLength:10,_iDisplayStart:0,_iRecordsTotal:0,_iRecordsDisplay:0,oClasses:{},bFiltered:!1,bSorted:!1,bSortCellsTop:null,oInit:null,aoDestroyCallback:[],fnRecordsTotal:function(){return"ssp"==y(this)?1*this._iRecordsTotal:this.aiDisplayMaster.length},fnRecordsDisplay:function(){return"ssp"==y(this)?1*this._iRecordsDisplay:this.aiDisplay.length},fnDisplayEnd:function(){var a=this._iDisplayLength,
b=this._iDisplayStart,c=b+a,d=this.aiDisplay.length,e=this.oFeatures,f=e.bPaginate;return e.bServerSide?!1===f||-1===a?b+d:Math.min(b+a,this._iRecordsDisplay):!f||c>d||-1===a?d:c},oInstance:null,sInstance:null,iTabIndex:0,nScrollHead:null,nScrollFoot:null,aLastSort:[],oPlugins:{},rowIdFn:null,rowId:null};m.ext=x={buttons:{},classes:{},builder:"-source-",errMode:"alert",feature:[],search:[],selector:{cell:[],column:[],row:[]},internal:{},legacy:{ajax:null},pager:{},renderer:{pageButton:{},header:{}},
order:{},type:{detect:[],search:{},order:{}},_unique:0,fnVersionCheck:m.fnVersionCheck,iApiIndex:0,oJUIClasses:{},sVersion:m.version};h.extend(x,{afnFiltering:x.search,aTypes:x.type.detect,ofnSearch:x.type.search,oSort:x.type.order,afnSortData:x.order,aoFeatures:x.feature,oApi:x.internal,oStdClasses:x.classes,oPagination:x.pager});h.extend(m.ext.classes,{sTable:"dataTable",sNoFooter:"no-footer",sPageButton:"paginate_button",sPageButtonActive:"current",sPageButtonDisabled:"disabled",sStripeOdd:"odd",
sStripeEven:"even",sRowEmpty:"dataTables_empty",sWrapper:"dataTables_wrapper",sFilter:"dataTables_filter",sInfo:"dataTables_info",sPaging:"dataTables_paginate paging_",sLength:"dataTables_length",sProcessing:"dataTables_processing",sSortAsc:"sorting_asc",sSortDesc:"sorting_desc",sSortable:"sorting",sSortableAsc:"sorting_asc_disabled",sSortableDesc:"sorting_desc_disabled",sSortableNone:"sorting_disabled",sSortColumn:"sorting_",sFilterInput:"",sLengthSelect:"",sScrollWrapper:"dataTables_scroll",sScrollHead:"dataTables_scrollHead",
sScrollHeadInner:"dataTables_scrollHeadInner",sScrollBody:"dataTables_scrollBody",sScrollFoot:"dataTables_scrollFoot",sScrollFootInner:"dataTables_scrollFootInner",sHeaderTH:"",sFooterTH:"",sSortJUIAsc:"",sSortJUIDesc:"",sSortJUI:"",sSortJUIAscAllowed:"",sSortJUIDescAllowed:"",sSortJUIWrapper:"",sSortIcon:"",sJUIHeader:"",sJUIFooter:""});var Kb=m.ext.pager;h.extend(Kb,{simple:function(){return["previous","next"]},full:function(){return["first","previous","next","last"]},numbers:function(a,b){return[ha(a,
b)]},simple_numbers:function(a,b){return["previous",ha(a,b),"next"]},full_numbers:function(a,b){return["first","previous",ha(a,b),"next","last"]},first_last_numbers:function(a,b){return["first",ha(a,b),"last"]},_numbers:ha,numbers_length:7});h.extend(!0,m.ext.renderer,{pageButton:{_:function(a,b,c,d,e,f){var g=a.oClasses,j=a.oLanguage.oPaginate,i=a.oLanguage.oAria.paginate||{},n,l,m=0,o=function(b,d){var k,s,u,r,v=function(b){Sa(a,b.data.action,true)};k=0;for(s=d.length;k<s;k++){r=d[k];if(h.isArray(r)){u=
h("<"+(r.DT_el||"div")+"/>").appendTo(b);o(u,r)}else{n=null;l="";switch(r){case "ellipsis":b.append('<span class="ellipsis">&#x2026;</span>');break;case "first":n=j.sFirst;l=r+(e>0?"":" "+g.sPageButtonDisabled);break;case "previous":n=j.sPrevious;l=r+(e>0?"":" "+g.sPageButtonDisabled);break;case "next":n=j.sNext;l=r+(e<f-1?"":" "+g.sPageButtonDisabled);break;case "last":n=j.sLast;l=r+(e<f-1?"":" "+g.sPageButtonDisabled);break;default:n=r+1;l=e===r?g.sPageButtonActive:""}if(n!==null){u=h("<a>",{"class":g.sPageButton+
" "+l,"aria-controls":a.sTableId,"aria-label":i[r],"data-dt-idx":m,tabindex:a.iTabIndex,id:c===0&&typeof r==="string"?a.sTableId+"_"+r:null}).html(n).appendTo(b);Va(u,{action:r},v);m++}}}},s;try{s=h(b).find(G.activeElement).data("dt-idx")}catch(u){}o(h(b).empty(),d);s!==k&&h(b).find("[data-dt-idx="+s+"]").focus()}}});h.extend(m.ext.type.detect,[function(a,b){var c=b.oLanguage.sDecimal;return Ya(a,c)?"num"+c:null},function(a){if(a&&!(a instanceof Date)&&!Zb.test(a))return null;var b=Date.parse(a);
return null!==b&&!isNaN(b)||L(a)?"date":null},function(a,b){var c=b.oLanguage.sDecimal;return Ya(a,c,!0)?"num-fmt"+c:null},function(a,b){var c=b.oLanguage.sDecimal;return Pb(a,c)?"html-num"+c:null},function(a,b){var c=b.oLanguage.sDecimal;return Pb(a,c,!0)?"html-num-fmt"+c:null},function(a){return L(a)||"string"===typeof a&&-1!==a.indexOf("<")?"html":null}]);h.extend(m.ext.type.search,{html:function(a){return L(a)?a:"string"===typeof a?a.replace(Mb," ").replace(Aa,""):""},string:function(a){return L(a)?
a:"string"===typeof a?a.replace(Mb," "):a}});var za=function(a,b,c,d){if(0!==a&&(!a||"-"===a))return-Infinity;b&&(a=Ob(a,b));a.replace&&(c&&(a=a.replace(c,"")),d&&(a=a.replace(d,"")));return 1*a};h.extend(x.type.order,{"date-pre":function(a){return Date.parse(a)||-Infinity},"html-pre":function(a){return L(a)?"":a.replace?a.replace(/<.*?>/g,"").toLowerCase():a+""},"string-pre":function(a){return L(a)?"":"string"===typeof a?a.toLowerCase():!a.toString?"":a.toString()},"string-asc":function(a,b){return a<
b?-1:a>b?1:0},"string-desc":function(a,b){return a<b?1:a>b?-1:0}});cb("");h.extend(!0,m.ext.renderer,{header:{_:function(a,b,c,d){h(a.nTable).on("order.dt.DT",function(e,f,g,h){if(a===f){e=c.idx;b.removeClass(c.sSortingClass+" "+d.sSortAsc+" "+d.sSortDesc).addClass(h[e]=="asc"?d.sSortAsc:h[e]=="desc"?d.sSortDesc:c.sSortingClass)}})},jqueryui:function(a,b,c,d){h("<div/>").addClass(d.sSortJUIWrapper).append(b.contents()).append(h("<span/>").addClass(d.sSortIcon+" "+c.sSortingClassJUI)).appendTo(b);
h(a.nTable).on("order.dt.DT",function(e,f,g,h){if(a===f){e=c.idx;b.removeClass(d.sSortAsc+" "+d.sSortDesc).addClass(h[e]=="asc"?d.sSortAsc:h[e]=="desc"?d.sSortDesc:c.sSortingClass);b.find("span."+d.sSortIcon).removeClass(d.sSortJUIAsc+" "+d.sSortJUIDesc+" "+d.sSortJUI+" "+d.sSortJUIAscAllowed+" "+d.sSortJUIDescAllowed).addClass(h[e]=="asc"?d.sSortJUIAsc:h[e]=="desc"?d.sSortJUIDesc:c.sSortingClassJUI)}})}}});var Vb=function(a){return"string"===typeof a?a.replace(/</g,"&lt;").replace(/>/g,"&gt;").replace(/"/g,
"&quot;"):a};m.render={number:function(a,b,c,d,e){return{display:function(f){if("number"!==typeof f&&"string"!==typeof f)return f;var g=0>f?"-":"",h=parseFloat(f);if(isNaN(h))return Vb(f);h=h.toFixed(c);f=Math.abs(h);h=parseInt(f,10);f=c?b+(f-h).toFixed(c).substring(2):"";return g+(d||"")+h.toString().replace(/\B(?=(\d{3})+(?!\d))/g,a)+f+(e||"")}}},text:function(){return{display:Vb}}};h.extend(m.ext.internal,{_fnExternApiFunc:Lb,_fnBuildAjax:sa,_fnAjaxUpdate:kb,_fnAjaxParameters:tb,_fnAjaxUpdateDraw:ub,
_fnAjaxDataSrc:ta,_fnAddColumn:Da,_fnColumnOptions:ja,_fnAdjustColumnSizing:Y,_fnVisibleToColumnIndex:Z,_fnColumnIndexToVisible:$,_fnVisbleColumns:aa,_fnGetColumns:la,_fnColumnTypes:Fa,_fnApplyColumnDefs:hb,_fnHungarianMap:X,_fnCamelToHungarian:I,_fnLanguageCompat:Ca,_fnBrowserDetect:fb,_fnAddData:M,_fnAddTr:ma,_fnNodeToDataIndex:function(a,b){return b._DT_RowIndex!==k?b._DT_RowIndex:null},_fnNodeToColumnIndex:function(a,b,c){return h.inArray(c,a.aoData[b].anCells)},_fnGetCellData:B,_fnSetCellData:ib,
_fnSplitObjNotation:Ia,_fnGetObjectDataFn:Q,_fnSetObjectDataFn:R,_fnGetDataMaster:Ja,_fnClearTable:na,_fnDeleteIndex:oa,_fnInvalidate:ca,_fnGetRowElements:Ha,_fnCreateTr:Ga,_fnBuildHead:jb,_fnDrawHead:ea,_fnDraw:N,_fnReDraw:S,_fnAddOptionsHtml:mb,_fnDetectHeader:da,_fnGetUniqueThs:ra,_fnFeatureHtmlFilter:ob,_fnFilterComplete:fa,_fnFilterCustom:xb,_fnFilterColumn:wb,_fnFilter:vb,_fnFilterCreateSearch:Oa,_fnEscapeRegex:Pa,_fnFilterData:yb,_fnFeatureHtmlInfo:rb,_fnUpdateInfo:Bb,_fnInfoMacros:Cb,_fnInitialise:ga,
_fnInitComplete:ua,_fnLengthChange:Qa,_fnFeatureHtmlLength:nb,_fnFeatureHtmlPaginate:sb,_fnPageChange:Sa,_fnFeatureHtmlProcessing:pb,_fnProcessingDisplay:C,_fnFeatureHtmlTable:qb,_fnScrollDraw:ka,_fnApplyToChildren:H,_fnCalculateColumnWidths:Ea,_fnThrottle:Na,_fnConvertToWidth:Db,_fnGetWidestNode:Eb,_fnGetMaxLenString:Fb,_fnStringToCss:v,_fnSortFlatten:V,_fnSort:lb,_fnSortAria:Hb,_fnSortListener:Ua,_fnSortAttachListener:La,_fnSortingClasses:wa,_fnSortData:Gb,_fnSaveState:xa,_fnLoadState:Ib,_fnSettingsFromNode:ya,
_fnLog:J,_fnMap:F,_fnBindAction:Va,_fnCallbackReg:z,_fnCallbackFire:r,_fnLengthOverflow:Ra,_fnRenderer:Ma,_fnDataSource:y,_fnRowAttributes:Ka,_fnCalculateEnd:function(){}});h.fn.dataTable=m;m.$=h;h.fn.dataTableSettings=m.settings;h.fn.dataTableExt=m.ext;h.fn.DataTable=function(a){return h(this).dataTable(a).api()};h.each(m,function(a,b){h.fn.DataTable[a]=b});return h.fn.dataTable});
```
