## Supplementary material for "Characterizing and inferring quantitative cell cycle phase in single-cell RNA-seq data analysis": Source code of our analysis, including scripts and data necessary to reproduce the work.: fa-v4-shims.min.js

```
/*!
 * Font Awesome Free 5.0.13 by @fontawesome - https://fontawesome.com
 * License - https://fontawesome.com/license (Icons: CC BY 4.0, Fonts: SIL OFL 1.1, Code: MIT License)
 */
var l,a;l=this,a=function(){"use strict";var l={};try{"undefined"!=typeof window&&(l=window)}catch(l){}var a=(l.navigator||{}).userAgent,e=void 0===a?"":a,r=l,n=(~e.indexOf("MSIE")||e.indexOf("Trident/"),"___FONT_AWESOME___"),o=function(){try{return"production"===process.env.NODE_ENV}catch(l){return!1}}(),u=[1,2,3,4,5,6,7,8,9,10],t=u.concat([11,12,13,14,15,16,17,18,19,20]);["xs","sm","lg","fw","ul","li","border","pull-left","pull-right","spin","pulse","rotate-90","rotate-180","rotate-270","flip-horizontal","flip-vertical","stack","stack-1x","stack-2x","inverse","layers","layers-text","layers-counter"].concat(u.map(function(l){return l+"x"})).concat(t.map(function(l){return"w-"+l}));var f=r||{};f[n]||(f[n]={}),f[n].styles||(f[n].styles={}),f[n].hooks||(f[n].hooks={}),f[n].shims||(f[n].shims=[]);var i=f[n],s=[["glass",null,"glass-martini"],["meetup","fab",null],["star-o","far","star"],["remove",null,"times"],["close",null,"times"],["gear",null,"cog"],["trash-o","far","trash-alt"],["file-o","far","file"],["clock-o","far","clock"],["arrow-circle-o-down","far","arrow-alt-circle-down"],["arrow-circle-o-up","far","arrow-alt-circle-up"],["play-circle-o","far","play-circle"],["repeat",null,"redo"],["rotate-right",null,"redo"],["refresh",null,"sync"],["list-alt","far",null],["dedent",null,"outdent"],["video-camera",null,"video"],["picture-o","far","image"],["photo","far","image"],["image","far","image"],["pencil",null,"pencil-alt"],["map-marker",null,"map-marker-alt"],["pencil-square-o","far","edit"],["share-square-o","far","share-square"],["check-square-o","far","check-square"],["arrows",null,"arrows-alt"],["times-circle-o","far","times-circle"],["check-circle-o","far","check-circle"],["mail-forward",null,"share"],["eye","far",null],["eye-slash","far",null],["warning",null,"exclamation-triangle"],["calendar",null,"calendar-alt"],["arrows-v",null,"arrows-alt-v"],["arrows-h",null,"arrows-alt-h"],["bar-chart","far","chart-bar"],["bar-chart-o","far","chart-bar"],["twitter-square","fab",null],["facebook-square","fab",null],["gears",null,"cogs"],["thumbs-o-up","far","thumbs-up"],["thumbs-o-down","far","thumbs-down"],["heart-o","far","heart"],["sign-out",null,"sign-out-alt"],["linkedin-square","fab","linkedin"],["thumb-tack",null,"thumbtack"],["external-link",null,"external-link-alt"],["sign-in",null,"sign-in-alt"],["github-square","fab",null],["lemon-o","far","lemon"],["square-o","far","square"],["bookmark-o","far","bookmark"],["twitter","fab",null],["facebook","fab","facebook-f"],["facebook-f","fab","facebook-f"],["github","fab",null],["credit-card","far",null],["feed",null,"rss"],["hdd-o","far","hdd"],["hand-o-right","far","hand-point-right"],["hand-o-left","far","hand-point-left"],["hand-o-up","far","hand-point-up"],["hand-o-down","far","hand-point-down"],["arrows-alt",null,"expand-arrows-alt"],["group",null,"users"],["chain",null,"link"],["scissors",null,"cut"],["files-o","far","copy"],["floppy-o","far","save"],["navicon",null,"bars"],["reorder",null,"bars"],["pinterest","fab",null],["pinterest-square","fab",null],["google-plus-square","fab",null],["google-plus","fab","google-plus-g"],["money","far","money-bill-alt"],["unsorted",null,"sort"],["sort-desc",null,"sort-down"],["sort-asc",null,"sort-up"],["linkedin","fab","linkedin-in"],["rotate-left",null,"undo"],["legal",null,"gavel"],["tachometer",null,"tachometer-alt"],["dashboard",null,"tachometer-alt"],["comment-o","far","comment"],["comments-o","far","comments"],["flash",null,"bolt"],["clipboard","far",null],["paste","far","clipboard"],["lightbulb-o","far","lightbulb"],["exchange",null,"exchange-alt"],["cloud-download",null,"cloud-download-alt"],["cloud-upload",null,"cloud-upload-alt"],["bell-o","far","bell"],["cutlery",null,"utensils"],["file-text-o","far","file-alt"],["building-o","far","building"],["hospital-o","far","hospital"],["tablet",null,"tablet-alt"],["mobile",null,"mobile-alt"],["mobile-phone",null,"mobile-alt"],["circle-o","far","circle"],["mail-reply",null,"reply"],["github-alt","fab",null],["folder-o","far","folder"],["folder-open-o","far","folder-open"],["smile-o","far","smile"],["frown-o","far","frown"],["meh-o","far","meh"],["keyboard-o","far","keyboard"],["flag-o","far","flag"],["mail-reply-all",null,"reply-all"],["star-half-o","far","star-half"],["star-half-empty","far","star-half"],["star-half-full","far","star-half"],["code-fork",null,"code-branch"],["chain-broken",null,"unlink"],["shield",null,"shield-alt"],["calendar-o","far","calendar"],["maxcdn","fab",null],["html5","fab",null],["css3","fab",null],["ticket",null,"ticket-alt"],["minus-square-o","far","minus-square"],["level-up",null,"level-up-alt"],["level-down",null,"level-down-alt"],["pencil-square",null,"pen-square"],["external-link-square",null,"external-link-square-alt"],["compass","far",null],["caret-square-o-down","far","caret-square-down"],["toggle-down","far","caret-square-down"],["caret-square-o-up","far","caret-square-up"],["toggle-up","far","caret-square-up"],["caret-square-o-right","far","caret-square-right"],["toggle-right","far","caret-square-right"],["eur",null,"euro-sign"],["euro",null,"euro-sign"],["gbp",null,"pound-sign"],["usd",null,"dollar-sign"],["dollar",null,"dollar-sign"],["inr",null,"rupee-sign"],["rupee",null,"rupee-sign"],["jpy",null,"yen-sign"],["cny",null,"yen-sign"],["rmb",null,"yen-sign"],["yen",null,"yen-sign"],["rub",null,"ruble-sign"],["ruble",null,"ruble-sign"],["rouble",null,"ruble-sign"],["krw",null,"won-sign"],["won",null,"won-sign"],["btc","fab",null],["bitcoin","fab","btc"],["file-text",null,"file-alt"],["sort-alpha-asc",null,"sort-alpha-down"],["sort-alpha-desc",null,"sort-alpha-up"],["sort-amount-asc",null,"sort-amount-down"],["sort-amount-desc",null,"sort-amount-up"],["sort-numeric-asc",null,"sort-numeric-down"],["sort-numeric-desc",null,"sort-numeric-up"],["youtube-square","fab",null],["youtube","fab",null],["xing","fab",null],["xing-square","fab",null],["youtube-play","fab","youtube"],["dropbox","fab",null],["stack-overflow","fab",null],["instagram","fab",null],["flickr","fab",null],["adn","fab",null],["bitbucket","fab",null],["bitbucket-square","fab","bitbucket"],["tumblr","fab",null],["tumblr-square","fab",null],["long-arrow-down",null,"long-arrow-alt-down"],["long-arrow-up",null,"long-arrow-alt-up"],["long-arrow-left",null,"long-arrow-alt-left"],["long-arrow-right",null,"long-arrow-alt-right"],["apple","fab",null],["windows","fab",null],["android","fab",null],["linux","fab",null],["dribbble","fab",null],["skype","fab",null],["foursquare","fab",null],["trello","fab",null],["gratipay","fab",null],["gittip","fab","gratipay"],["sun-o","far","sun"],["moon-o","far","moon"],["vk","fab",null],["weibo","fab",null],["renren","fab",null],["pagelines","fab",null],["stack-exchange","fab",null],["arrow-circle-o-right","far","arrow-alt-circle-right"],["arrow-circle-o-left","far","arrow-alt-circle-left"],["caret-square-o-left","far","caret-square-left"],["toggle-left","far","caret-square-left"],["dot-circle-o","far","dot-circle"],["vimeo-square","fab",null],["try",null,"lira-sign"],["turkish-lira",null,"lira-sign"],["plus-square-o","far","plus-square"],["slack","fab",null],["wordpress","fab",null],["openid","fab",null],["institution",null,"university"],["bank",null,"university"],["mortar-board",null,"graduation-cap"],["yahoo","fab",null],["google","fab",null],["reddit","fab",null],["reddit-square","fab",null],["stumbleupon-circle","fab",null],["stumbleupon","fab",null],["delicious","fab",null],["digg","fab",null],["pied-piper-pp","fab",null],["pied-piper-alt","fab",null],["drupal","fab",null],["joomla","fab",null],["spoon",null,"utensil-spoon"],["behance","fab",null],["behance-square","fab",null],["steam","fab",null],["steam-square","fab",null],["automobile",null,"car"],["cab",null,"taxi"],["envelope-o","far","envelope"],["deviantart","fab",null],["soundcloud","fab",null],["file-pdf-o","far","file-pdf"],["file-word-o","far","file-word"],["file-excel-o","far","file-excel"],["file-powerpoint-o","far","file-powerpoint"],["file-image-o","far","file-image"],["file-photo-o","far","file-image"],["file-picture-o","far","file-image"],["file-archive-o","far","file-archive"],["file-zip-o","far","file-archive"],["file-audio-o","far","file-audio"],["file-sound-o","far","file-audio"],["file-video-o","far","file-video"],["file-movie-o","far","file-video"],["file-code-o","far","file-code"],["vine","fab",null],["codepen","fab",null],["jsfiddle","fab",null],["life-ring","far",null],["life-bouy","far","life-ring"],["life-buoy","far","life-ring"],["life-saver","far","life-ring"],["support","far","life-ring"],["circle-o-notch",null,"circle-notch"],["rebel","fab",null],["ra","fab","rebel"],["resistance","fab","rebel"],["empire","fab",null],["ge","fab","empire"],["git-square","fab",null],["git","fab",null],["hacker-news","fab",null],["y-combinator-square","fab","hacker-news"],["yc-square","fab","hacker-news"],["tencent-weibo","fab",null],["qq","fab",null],["weixin","fab",null],["wechat","fab","weixin"],["send",null,"paper-plane"],["paper-plane-o","far","paper-plane"],["send-o","far","paper-plane"],["circle-thin","far","circle"],["header",null,"heading"],["sliders",null,"sliders-h"],["futbol-o","far","futbol"],["soccer-ball-o","far","futbol"],["slideshare","fab",null],["twitch","fab",null],["yelp","fab",null],["newspaper-o","far","newspaper"],["paypal","fab",null],["google-wallet","fab",null],["cc-visa","fab",null],["cc-mastercard","fab",null],["cc-discover","fab",null],["cc-amex","fab",null],["cc-paypal","fab",null],["cc-stripe","fab",null],["bell-slash-o","far","bell-slash"],["trash",null,"trash-alt"],["copyright","far",null],["eyedropper",null,"eye-dropper"],["area-chart",null,"chart-area"],["pie-chart",null,"chart-pie"],["line-chart",null,"chart-line"],["lastfm","fab",null],["lastfm-square","fab",null],["ioxhost","fab",null],["angellist","fab",null],["cc","far","closed-captioning"],["ils",null,"shekel-sign"],["shekel",null,"shekel-sign"],["sheqel",null,"shekel-sign"],["meanpath","fab","font-awesome"],["buysellads","fab",null],["connectdevelop","fab",null],["dashcube","fab",null],["forumbee","fab",null],["leanpub","fab",null],["sellsy","fab",null],["shirtsinbulk","fab",null],["simplybuilt","fab",null],["skyatlas","fab",null],["diamond","far","gem"],["intersex",null,"transgender"],["facebook-official","fab","facebook"],["pinterest-p","fab",null],["whatsapp","fab",null],["hotel",null,"bed"],["viacoin","fab",null],["medium","fab",null],["y-combinator","fab",null],["yc","fab","y-combinator"],["optin-monster","fab",null],["opencart","fab",null],["expeditedssl","fab",null],["battery-4",null,"battery-full"],["battery",null,"battery-full"],["battery-3",null,"battery-three-quarters"],["battery-2",null,"battery-half"],["battery-1",null,"battery-quarter"],["battery-0",null,"battery-empty"],["object-group","far",null],["object-ungroup","far",null],["sticky-note-o","far","sticky-note"],["cc-jcb","fab",null],["cc-diners-club","fab",null],["clone","far",null],["hourglass-o","far","hourglass"],["hourglass-1",null,"hourglass-start"],["hourglass-2",null,"hourglass-half"],["hourglass-3",null,"hourglass-end"],["hand-rock-o","far","hand-rock"],["hand-grab-o","far","hand-rock"],["hand-paper-o","far","hand-paper"],["hand-stop-o","far","hand-paper"],["hand-scissors-o","far","hand-scissors"],["hand-lizard-o","far","hand-lizard"],["hand-spock-o","far","hand-spock"],["hand-pointer-o","far","hand-pointer"],["hand-peace-o","far","hand-peace"],["registered","far",null],["creative-commons","fab",null],["gg","fab",null],["gg-circle","fab",null],["tripadvisor","fab",null],["odnoklassniki","fab",null],["odnoklassniki-square","fab",null],["get-pocket","fab",null],["wikipedia-w","fab",null],["safari","fab",null],["chrome","fab",null],["firefox","fab",null],["opera","fab",null],["internet-explorer","fab",null],["television",null,"tv"],["contao","fab",null],["500px","fab",null],["amazon","fab",null],["calendar-plus-o","far","calendar-plus"],["calendar-minus-o","far","calendar-minus"],["calendar-times-o","far","calendar-times"],["calendar-check-o","far","calendar-check"],["map-o","far","map"],["commenting","far","comment-dots"],["commenting-o","far","comment-dots"],["houzz","fab",null],["vimeo","fab","vimeo-v"],["black-tie","fab",null],["fonticons","fab",null],["reddit-alien","fab",null],["edge","fab",null],["credit-card-alt",null,"credit-card"],["codiepie","fab",null],["modx","fab",null],["fort-awesome","fab",null],["usb","fab",null],["product-hunt","fab",null],["mixcloud","fab",null],["scribd","fab",null],["pause-circle-o","far","pause-circle"],["stop-circle-o","far","stop-circle"],["bluetooth","fab",null],["bluetooth-b","fab",null],["gitlab","fab",null],["wpbeginner","fab",null],["wpforms","fab",null],["envira","fab",null],["wheelchair-alt","fab","accessible-icon"],["question-circle-o","far","question-circle"],["volume-control-phone",null,"phone-volume"],["asl-interpreting",null,"american-sign-language-interpreting"],["deafness",null,"deaf"],["hard-of-hearing",null,"deaf"],["glide","fab",null],["glide-g","fab",null],["signing",null,"sign-language"],["viadeo","fab",null],["viadeo-square","fab",null],["snapchat","fab",null],["snapchat-ghost","fab",null],["snapchat-square","fab",null],["pied-piper","fab",null],["first-order","fab",null],["yoast","fab",null],["themeisle","fab",null],["google-plus-official","fab","google-plus"],["google-plus-circle","fab","google-plus"],["font-awesome","fab",null],["fa","fab","font-awesome"],["handshake-o","far","handshake"],["envelope-open-o","far","envelope-open"],["linode","fab",null],["address-book-o","far","address-book"],["vcard",null,"address-card"],["address-card-o","far","address-card"],["vcard-o","far","address-card"],["user-circle-o","far","user-circle"],["user-o","far","user"],["id-badge","far",null],["drivers-license",null,"id-card"],["id-card-o","far","id-card"],["drivers-license-o","far","id-card"],["quora","fab",null],["free-code-camp","fab",null],["telegram","fab",null],["thermometer-4",null,"thermometer-full"],["thermometer",null,"thermometer-full"],["thermometer-3",null,"thermometer-three-quarters"],["thermometer-2",null,"thermometer-half"],["thermometer-1",null,"thermometer-quarter"],["thermometer-0",null,"thermometer-empty"],["bathtub",null,"bath"],["s15",null,"bath"],["window-maximize","far",null],["window-restore","far",null],["times-rectangle",null,"window-close"],["window-close-o","far","window-close"],["times-rectangle-o","far","window-close"],["bandcamp","fab",null],["grav","fab",null],["etsy","fab",null],["imdb","fab",null],["ravelry","fab",null],["eercast","fab","sellcast"],["snowflake-o","far","snowflake"],["superpowers","fab",null],["wpexplorer","fab",null],["spotify","fab",null]];return function(l){try{l()}catch(l){if(!o)throw l}}(function(){var l;"function"==typeof i.hooks.addShims?i.hooks.addShims(s):(l=i.shims).push.apply(l,s)}),s},"object"==typeof exports&&"undefined"!=typeof module?module.exports=a():"function"==typeof define&&define.amd?define(a):l["fontawesome-free-shims"]=a();
```
