## Supplementary material for "Characterizing and inferring quantitative cell cycle phase in single-cell RNA-seq data analysis": Source code of our analysis, including scripts and data necessary to reproduce the work.: fontawesome-all.min.js

```
/*!
 * Font Awesome Free 5.0.13 by @fontawesome - https://fontawesome.com
 * License - https://fontawesome.com/license (Icons: CC BY 4.0, Fonts: SIL OFL 1.1, Code: MIT License)
 */
!function(){"use strict";var c={};try{"undefined"!=typeof window&&(c=window)}catch(c){}var l=(c.navigator||{}).userAgent,h=void 0===l?"":l,v=c,z=(~h.indexOf("MSIE")||h.indexOf("Trident/"),"___FONT_AWESOME___"),e=function(){try{return!0}catch(c){return!1}}(),m=[1,2,3,4,5,6,7,8,9,10],a=m.concat([11,12,13,14,15,16,17,18,19,20]);["xs","sm","lg","fw","ul","li","border","pull-left","pull-right","spin","pulse","rotate-90","rotate-180","rotate-270","flip-horizontal","flip-vertical","stack","stack-1x","stack-2x","inverse","layers","layers-text","layers-counter"].concat(m.map(function(c){return c+"x"})).concat(a.map(function(c){return"w-"+c}));var s=v||{};s[z]||(s[z]={}),s[z].styles||(s[z].styles={}),s[z].hooks||(s[z].hooks={}),s[z].shims||(s[z].shims=[]);var t=s[z],f=Object.assign||function(c){for(var l=1;l<arguments.length;l++){var h=arguments[l];for(var v in h)Object.prototype.hasOwnProperty.call(h,v)&&(c[v]=h[v])}return c};var M={"500px":[448,512,[],"f26e","M103.3 344.3c-6.5-14.2-6.9-18.3 7.4-23.1 25.6-8 8 9.2 43.2 49.2h.3v-93.9c1.2-50.2 44-92.2 97.7-92.2 53.9 0 97.7 43.5 97.7 96.8 0 63.4-60.8 113.2-128.5 93.3-10.5-4.2-2.1-31.7 8.5-28.6 53 0 89.4-10.1 89.4-64.4 0-61-77.1-89.6-116.9-44.6-23.5 26.4-17.6 42.1-17.6 157.6 50.7 31 118.3 22 160.4-20.1 24.8-24.8 38.5-58 38.5-93 0-35.2-13.8-68.2-38.8-93.3-24.8-24.8-57.8-38.5-93.3-38.5s-68.8 13.8-93.5 38.5c-.3.3-16 16.5-21.2 23.9l-.5.6c-3.3 4.7-6.3 9.1-20.1 6.1-6.9-1.7-14.3-5.8-14.3-11.8V20c0-5 3.9-10.5 10.5-10.5h241.3c8.3 0 8.3 11.6 8.3 15.1 0 3.9 0 15.1-8.3 15.1H130.3v132.9h.3c104.2-109.8 282.8-36 282.8 108.9 0 178.1-244.8 220.3-310.1 62.8zm63.3-260.8c-.5 4.2 4.6 24.5 14.6 20.6C306 56.6 384 144.5 390.6 144.5c4.8 0 22.8-15.3 14.3-22.8-93.2-89-234.5-57-238.3-38.2zM393 414.7C283 524.6 94 475.5 61 310.5c0-12.2-30.4-7.4-28.9 3.3 24 173.4 246 256.9 381.6 121.3 6.9-7.8-12.6-28.4-20.7-20.4zM213.6 306.6c0 4 4.3 7.3 5.5 8.5 3 3 6.1 4.4 8.5 4.4 3.8 0 2.6.2 22.3-19.5 19.6 19.3 19.1 19.5 22.3 19.5 5.4 0 18.5-10.4 10.7-18.2L265.6 284l18.2-18.2c6.3-6.8-10.1-21.8-16.2-15.7L249.7 268c-18.6-18.8-18.4-19.5-21.5-19.5-5 0-18 11.7-12.4 17.3L234 284c-18.1 17.9-20.4 19.2-20.4 22.6z"],"accessible-icon":[448,512,[],"f368","M423.9 255.8L411 413.1c-3.3 40.7-63.9 35.1-60.6-4.9l10-122.5-41.1 2.3c10.1 20.7 15.8 43.9 15.8 68.5 0 41.2-16.1 78.7-42.3 106.5l-39.3-39.3c57.9-63.7 13.1-167.2-74-167.2-25.9 0-49.5 9.9-67.2 26L73 243.2c22-20.7 50.1-35.1 81.4-40.2l75.3-85.7-42.6-24.8-51.6 46c-30 26.8-70.6-18.5-40.5-45.4l68-60.7c9.8-8.8 24.1-10.2 35.5-3.6 0 0 139.3 80.9 139.5 81.1 16.2 10.1 20.7 36 6.1 52.6L285.7 229l106.1-5.9c18.5-1.1 33.6 14.4 32.1 32.7zm-64.9-154c28.1 0 50.9-22.8 50.9-50.9C409.9 22.8 387.1 0 359 0c-28.1 0-50.9 22.8-50.9 50.9 0 28.1 22.8 50.9 50.9 50.9zM179.6 456.5c-80.6 0-127.4-90.6-82.7-156.1l-39.7-39.7C36.4 287 24 320.3 24 356.4c0 130.7 150.7 201.4 251.4 122.5l-39.7-39.7c-16 10.9-35.3 17.3-56.1 17.3z"],accusoft:[640,512,[],"f369","M482.2 372.1C476.5 365.2 250 75 242.3 65.5c-13.7-17.2 0-16.8 19.2-16.9 9.7-.1 106.3-.6 116.5-.6 24.1-.1 28.7.6 38.4 12.8 2.1 2.7 205.1 245.8 207.2 248.3 5.5 6.7 15.2 19.1 7.2 23.4-2.4 1.3-114.6 47.7-117.8 48.9-10.1 4-17.5 6.8-30.8-9.3m114.7-5.6s-115 50.4-117.5 51.6c-16 7.3-26.9-3.2-36.7-14.6l-57.1-74c-5.4-.9-60.4-9.6-65.3-9.3-3.1.2-9.6.8-14.4 2.9-4.9 2.1-145.2 52.8-150.2 54.7-5.1 2-11.4 3.6-11.1 7.6.2 2.5 2 2.6 4.6 3.5 2.7.8 300.9 67.6 308 69.1 15.6 3.3 38.5 10.5 53.6 1.7 2.1-1.2 123.8-76.4 125.8-77.8 5.4-4 4.3-6.8-1.7-8.2-2.3-.3-24.6-4.7-38-7.2m-326-181.3s-12 1.6-25 15.1c-9 9.3-242.1 239.1-243.4 240.9-7 10 1.6 6.8 15.7 1.7.8 0 114.5-36.6 114.5-36.6.5-.6-.1-.1.6-.6-.4-5.1-.8-26.2-1-27.7-.6-5.2 2.2-6.9 7-8.9l92.6-33.8c.6-.8 88.5-81.7 90.2-83.3v-1l-51.2-65.8"],adn:[496,512,[],"f170","M248 167.5l64.9 98.8H183.1l64.9-98.8zM496 256c0 136.9-111.1 248-248 248S0 392.9 0 256 111.1 8 248 8s248 111.1 248 248zm-99.8 82.7L248 115.5 99.8 338.7h30.4l33.6-51.7h168.6l33.6 51.7h30.2z"],adversal:[512,512,[],"f36a","M482.1 32H28.7C5.8 32 0 37.9 0 60.9v390.2C0 474.4 5.8 480 28.7 480h453.4c24.4 0 29.9-5.2 29.9-29.7V62.2c0-24.6-5.4-30.2-29.9-30.2zM178.4 220.3c-27.5-20.2-72.1-8.7-84.2 23.4-4.3 11.1-9.3 9.5-17.5 8.3-9.7-1.5-17.2-3.2-22.5-5.5-28.8-11.4 8.6-55.3 24.9-64.3 41.1-21.4 83.4-22.2 125.3-4.8 40.9 16.8 34.5 59.2 34.5 128.5 2.7 25.8-4.3 58.3 9.3 88.8 1.9 4.4.4 7.9-2.7 10.7-8.4 6.7-39.3 2.2-46.6-7.4-1.9-2.2-1.8-3.6-3.9-6.2-3.6-3.9-7.3-2.2-11.9 1-57.4 36.4-140.3 21.4-147-43.3-3.1-29.3 12.4-57.1 39.6-71 38.2-19.5 112.2-11.8 114-30.9 1.1-10.2-1.9-20.1-11.3-27.3zm286.7 222c0 15.1-11.1 9.9-17.8 9.9H52.4c-7.4 0-18.2 4.8-17.8-10.7.4-13.9 10.5-9.1 17.1-9.1 132.3-.4 264.5-.4 396.8 0 6.8 0 16.6-4.4 16.6 9.9zm3.8-340.5v291c0 5.7-.7 13.9-8.1 13.9-12.4-.4-27.5 7.1-36.1-5.6-5.8-8.7-7.8-4-12.4-1.2-53.4 29.7-128.1 7.1-144.4-85.2-6.1-33.4-.7-67.1 15.7-100 11.8-23.9 56.9-76.1 136.1-30.5v-71c0-26.2-.1-26.2 26-26.2 3.1 0 6.6.4 9.7 0 10.1-.8 13.6 4.4 13.6 14.3-.1.2-.1.3-.1.5zm-51.5 232.3c-19.5 47.6-72.9 43.3-90 5.2-15.1-33.3-15.5-68.2.4-101.5 16.3-34.1 59.7-35.7 81.5-4.8 20.6 28.8 14.9 84.6 8.1 101.1zm-294.8 35.3c-7.5-1.3-33-3.3-33.7-27.8-.4-13.9 7.8-23 19.8-25.8 24.4-5.9 49.3-9.9 73.7-14.7 8.9-2 7.4 4.4 7.8 9.5 1.4 33-26.1 59.2-67.6 58.8z"],affiliatetheme:[512,512,[],"f36b","M159.7 237.4C108.4 308.3 43.1 348.2 14 326.6-15.2 304.9 2.8 230 54.2 159.1c51.3-70.9 116.6-110.8 145.7-89.2 29.1 21.6 11.1 96.6-40.2 167.5zm351.2-57.3C437.1 303.5 319 367.8 246.4 323.7c-25-15.2-41.3-41.2-49-73.8-33.6 64.8-92.8 113.8-164.1 133.2 49.8 59.3 124.1 96.9 207 96.9 150 0 271.6-123.1 271.6-274.9.1-8.5-.3-16.8-1-25z"],algolia:[448,512,[],"f36c","M229.3 182.6c-49.3 0-89.2 39.9-89.2 89.2 0 49.3 39.9 89.2 89.2 89.2s89.2-39.9 89.2-89.2c0-49.3-40-89.2-89.2-89.2zm62.7 56.6l-58.9 30.6c-1.8.9-3.8-.4-3.8-2.3V201c0-1.5 1.3-2.7 2.7-2.6 26.2 1 48.9 15.7 61.1 37.1.7 1.3.2 3-1.1 3.7zM389.1 32H58.9C26.4 32 0 58.4 0 90.9V421c0 32.6 26.4 59 58.9 59H389c32.6 0 58.9-26.4 58.9-58.9V90.9C448 58.4 421.6 32 389.1 32zm-202.6 84.7c0-10.8 8.7-19.5 19.5-19.5h45.3c10.8 0 19.5 8.7 19.5 19.5v15.4c0 1.8-1.7 3-3.3 2.5-12.3-3.4-25.1-5.1-38.1-5.1-13.5 0-26.7 1.8-39.4 5.5-1.7.5-3.4-.8-3.4-2.5v-15.8zm-84.4 37l9.2-9.2c7.6-7.6 19.9-7.6 27.5 0l7.7 7.7c1.1 1.1 1 3-.3 4-6.2 4.5-12.1 9.4-17.6 14.9-5.4 5.4-10.4 11.3-14.8 17.4-1 1.3-2.9 1.5-4 .3l-7.7-7.7c-7.6-7.5-7.6-19.8 0-27.4zm127.2 244.8c-70 0-126.6-56.7-126.6-126.6s56.7-126.6 126.6-126.6c70 0 126.6 56.6 126.6 126.6 0 69.8-56.7 126.6-126.6 126.6z"],amazon:[448,512,[],"f270","M257.2 162.7c-48.7 1.8-169.5 15.5-169.5 117.5 0 109.5 138.3 114 183.5 43.2 6.5 10.2 35.4 37.5 45.3 46.8l56.8-56S341 288.9 341 261.4V114.3C341 89 316.5 32 228.7 32 140.7 32 94 87 94 136.3l73.5 6.8c16.3-49.5 54.2-49.5 54.2-49.5 40.7-.1 35.5 29.8 35.5 69.1zm0 86.8c0 80-84.2 68-84.2 17.2 0-47.2 50.5-56.7 84.2-57.8v40.6zm136 163.5c-7.7 10-70 67-174.5 67S34.2 408.5 9.7 379c-6.8-7.7 1-11.3 5.5-8.3C88.5 415.2 203 488.5 387.7 401c7.5-3.7 13.3 2 5.5 12zm39.8 2.2c-6.5 15.8-16 26.8-21.2 31-5.5 4.5-9.5 2.7-6.5-3.8s19.3-46.5 12.7-55c-6.5-8.3-37-4.3-48-3.2-10.8 1-13 2-14-.3-2.3-5.7 21.7-15.5 37.5-17.5 15.7-1.8 41-.8 46 5.7 3.7 5.1 0 27.1-6.5 43.1z"],"amazon-pay":[611,512,[],"f42c","M0 325.2c2.3-4.2 5.2-4.9 9.7-2.5 10.4 5.6 20.6 11.4 31.2 16.7 40.7 20.4 83.2 35.6 127.4 46.3 20.9 5 41.9 9 63.2 11.8 31.5 4.2 63.2 6 95 5.2 17.4-.4 34.8-1.8 52.1-3.8 56.4-6.7 110.9-20.8 163.3-42.8 2.9-1.2 5.9-2 9.1-1.2 6.7 1.8 9 9 4.1 13.9-2.8 2.8-6.3 5.1-9.6 7.4-30.7 21.1-64.2 36.4-99.6 47.9-24.6 7.9-49.6 13.8-75.1 17.6-17.6 2.6-35.4 4.4-53.2 4.8-.8 0-1.7.2-2.5.3H294c-.8-.1-1.7-.3-2.5-.3-3.6-.2-7.2-.3-10.7-.4-16.9-.7-33.7-2.6-50.4-5.3-27.4-4.5-54.2-11.4-80.4-20.9-54.1-19.6-102.6-48.6-145.6-87-1.8-1.6-3-3.8-4.4-5.7v-2zM158 65c-1.4.2-2.9.4-4.3.6-14 1.7-26.6 6.9-38 15.1-2.4 1.7-4.6 3.5-7.1 5.4-.2-.5-.4-1-.4-1.4-.4-2.7-.8-5.5-1.3-8.2-.7-4.6-3-6.6-7.6-6.6H87.8c-6.9 0-8.2 1.3-8.2 8.2v209.3c0 1 0 2 .1 3 .2 3 2 4.9 4.9 5 7 .1 14.1.1 21.1 0 2.9 0 4.7-2 5-5 .1-1 .1-2 .1-3V215c1.1.9 1.7 1.4 2.2 1.9 17.9 14.9 38.5 19.8 61 15.4 20.4-4 34.6-16.5 43.8-34.9 7-13.9 9.9-28.7 10.3-44.1.5-17.1-1.2-33.9-8.1-49.8-8.5-19.6-22.6-32.5-43.9-36.9-3.2-.7-6.5-1-9.8-1.5-2.8-.1-5.5-.1-8.3-.1zm-47.4 41.9c0-1.5.4-2.4 1.7-3.3 13.7-9.5 28.8-14.5 45.6-13.2 14.9 1.1 27.1 8.4 33.5 25.9 3.9 10.7 4.9 21.8 4.9 33 0 10.4-.8 20.6-4 30.6-6.8 21.3-22.4 29.4-42.6 28.5-14-.6-26.2-6-37.4-13.9-1.2-.9-1.7-1.7-1.7-3.3.1-14.1 0-28.1 0-42.2 0-14 .1-28 0-42.1zM316.3 65c-1 .1-2 .3-2.9.4-9.8.5-19.4 1.7-28.9 4.1-6.1 1.6-12 3.8-17.9 5.8-3.6 1.2-5.4 3.8-5.3 7.7.1 3.3-.1 6.6 0 9.9.1 4.8 2.1 6.1 6.8 4.9 7.8-2 15.6-4.2 23.5-5.7 12.3-2.3 24.7-3.3 37.2-1.4 6.5 1 12.6 2.9 16.8 8.4 3.7 4.8 5.1 10.5 5.3 16.4.3 8.3.2 16.6.3 24.9 0 .4-.1.9-.2 1.4-.5-.1-.9 0-1.3-.1-10.5-2.5-21.1-4.3-32-4.9-11.3-.6-22.5.1-33.3 3.9-12.9 4.5-23.3 12.3-29.4 24.9-4.7 9.8-5.4 20.2-3.9 30.7 2 14 9 24.8 21.4 31.7 11.9 6.6 24.8 7.4 37.9 5.4 15.1-2.3 28.5-8.7 40.3-18.4.4-.4.9-.7 1.6-1.1.6 3.8 1.1 7.4 1.8 11 .6 3.1 2.5 5.1 5.4 5.2 5.4.1 10.9.1 16.3 0 2.7-.1 4.5-1.9 4.8-4.7.1-.9.1-1.9.1-2.8v-106c0-4.3-.2-8.6-.9-12.9-1.9-12.9-7.4-23.5-19-30.4-6.7-4-14.1-6-21.8-7.1-3.6-.5-7.2-.8-10.8-1.3-3.9.1-7.9.1-11.9.1zm35 127.7c0 1.3-.4 2.2-1.5 3-11.2 8.1-23.5 13.5-37.4 14.9-5.7.6-11.4.4-16.8-1.8-6.3-2.5-10.4-6.9-12.4-13.3s-2-13-.1-19.4c2.5-8.3 8.4-13 16.4-15.6 8.1-2.6 16.5-3 24.8-2.2 8.4.7 16.6 2.3 25 3.4 1.6.2 2.1 1 2.1 2.6-.1 4.8 0 9.5 0 14.3-.1 4.7-.2 9.4-.1 14.1zm259.9 129.4c-1-5-4.8-6.9-9.1-8.3-6.8-2.3-13.9-3.3-21-3.9-13.1-1.1-26.2-.5-39.2 1.9-14.3 2.7-27.9 7.3-40 15.6-1.4 1-2.8 2.1-3.7 3.5-.7 1.1-.9 2.8-.5 4 .4 1.5 2.1 1.9 3.6 1.8.7 0 1.5 0 2.2-.1 7.8-.8 15.5-1.7 23.3-2.5 11.4-1.1 22.9-1.8 34.3-.9 4.8.3 9.7 1.4 14.4 2.7 5.1 1.4 7.4 5.2 7.6 10.4.4 8-1.4 15.7-3.5 23.3-4.1 15.4-10 30.3-15.8 45.1-.4 1-.8 2-1 3-.5 2.9 1.2 4.8 4.1 4.1 1.7-.4 3.6-1.3 4.8-2.5 4.4-4.3 8.9-8.6 12.7-13.4 12.8-16.4 20.3-35.3 24.7-55.6.8-3.6 1.4-7.3 2.1-10.9v-17.3zM479.1 198.9c-12.9-35.7-25.8-71.5-38.7-107.2-2-5.7-4.2-11.3-6.3-16.9-1.1-2.9-3.2-4.8-6.4-4.8-7.6-.1-15.2-.2-22.9-.1-2.5 0-3.7 2-3.2 4.5.5 2.1 1.1 4.1 1.9 6.1 19.6 48.5 39.3 97.1 59.1 145.5 1.7 4.1 2.1 7.6.2 11.8-3.3 7.3-5.9 15-9.3 22.3-3 6.5-8 11.4-15.2 13.3-5.1 1.4-10.2 1.6-15.4 1.1-2.5-.2-5-.8-7.5-1-3.4-.2-5.1 1.3-5.2 4.8-.1 3.3-.1 6.6 0 9.9.1 5.5 2 8 7.4 8.9 5.6 1 11.3 1.9 16.9 2 17.1.4 30.7-6.5 39.5-21.4 3.5-5.9 6.7-12.1 9.2-18.4 23.7-59.8 47.1-119.7 70.6-179.6.7-1.8 1.3-3.6 1.6-5.5.4-2.8-.9-4.4-3.7-4.4-6.6-.1-13.3 0-19.9 0-3.7 0-6.3 1.6-7.7 5.2-.5 1.4-1.1 2.7-1.6 4.1-11.6 33.3-23.2 66.6-34.8 100-2.5 7.2-5.1 14.5-7.7 22.2-.4-1.1-.6-1.7-.9-2.4z"],amilia:[448,512,[],"f36d","M240.1 32c-61.9 0-131.5 16.9-184.2 55.4-5.1 3.1-9.1 9.2-7.2 19.4 1.1 5.1 5.1 27.4 10.2 39.6 4.1 10.2 14.2 10.2 20.3 6.1 32.5-22.3 96.5-47.7 152.3-47.7 57.9 0 58.9 28.4 58.9 73.1v38.5C203 227.7 78.2 251 46.7 264.2 11.2 280.5 16.3 357.7 16.3 376s15.2 104 124.9 104c47.8 0 113.7-20.7 153.3-42.1v25.4c0 3 2.1 8.2 6.1 9.1 3.1 1 50.7 2 59.9 2s62.5.3 66.5-.7c4.1-1 5.1-6.1 5.1-9.1V168c-.1-80.3-57.9-136-192-136zm-87.9 327.7c0-12.2-3-42.7 18.3-52.9 24.3-13.2 75.1-29.4 119.8-33.5V380c-21.4 13.2-48.7 24.4-79.1 24.4-52.8 0-58.9-33.5-59-44.7"],android:[448,512,[],"f17b","M89.6 204.5v115.8c0 15.4-12.1 27.7-27.5 27.7-15.3 0-30.1-12.4-30.1-27.7V204.5c0-15.1 14.8-27.5 30.1-27.5 15.1 0 27.5 12.4 27.5 27.5zm10.8 157c0 16.4 13.2 29.6 29.6 29.6h19.9l.3 61.1c0 36.9 55.2 36.6 55.2 0v-61.1h37.2v61.1c0 36.7 55.5 36.8 55.5 0v-61.1h20.2c16.2 0 29.4-13.2 29.4-29.6V182.1H100.4v179.4zm248-189.1H99.3c0-42.8 25.6-80 63.6-99.4l-19.1-35.3c-2.8-4.9 4.3-8 6.7-3.8l19.4 35.6c34.9-15.5 75-14.7 108.3 0L297.5 34c2.5-4.3 9.5-1.1 6.7 3.8L285.1 73c37.7 19.4 63.3 56.6 63.3 99.4zm-170.7-55.5c0-5.7-4.6-10.5-10.5-10.5-5.7 0-10.2 4.8-10.2 10.5s4.6 10.5 10.2 10.5c5.9 0 10.5-4.8 10.5-10.5zm113.4 0c0-5.7-4.6-10.5-10.2-10.5-5.9 0-10.5 4.8-10.5 10.5s4.6 10.5 10.5 10.5c5.6 0 10.2-4.8 10.2-10.5zm94.8 60.1c-15.1 0-27.5 12.1-27.5 27.5v115.8c0 15.4 12.4 27.7 27.5 27.7 15.4 0 30.1-12.4 30.1-27.7V204.5c0-15.4-14.8-27.5-30.1-27.5z"],angellist:[448,512,[],"f209","M347.1 215.4c11.7-32.6 45.4-126.9 45.4-157.1 0-26.6-15.7-48.9-43.7-48.9-44.6 0-84.6 131.7-97.1 163.1C242 144 196.6 0 156.6 0c-31.1 0-45.7 22.9-45.7 51.7 0 35.3 34.2 126.8 46.6 162-6.3-2.3-13.1-4.3-20-4.3-23.4 0-48.3 29.1-48.3 52.6 0 8.9 4.9 21.4 8 29.7-36.9 10-51.1 34.6-51.1 71.7C46 435.6 114.4 512 210.6 512c118 0 191.4-88.6 191.4-202.9 0-43.1-6.9-82-54.9-93.7zM311.7 108c4-12.3 21.1-64.3 37.1-64.3 8.6 0 10.9 8.9 10.9 16 0 19.1-38.6 124.6-47.1 148l-34-6 33.1-93.7zM142.3 48.3c0-11.9 14.5-45.7 46.3 47.1l34.6 100.3c-15.6-1.3-27.7-3-35.4 1.4-10.9-28.8-45.5-119.7-45.5-148.8zM140 244c29.3 0 67.1 94.6 67.1 107.4 0 5.1-4.9 11.4-10.6 11.4-20.9 0-76.9-76.9-76.9-97.7.1-7.7 12.7-21.1 20.4-21.1zm184.3 186.3c-29.1 32-66.3 48.6-109.7 48.6-59.4 0-106.3-32.6-128.9-88.3-17.1-43.4 3.8-68.3 20.6-68.3 11.4 0 54.3 60.3 54.3 73.1 0 4.9-7.7 8.3-11.7 8.3-16.1 0-22.4-15.5-51.1-51.4-29.7 29.7 20.5 86.9 58.3 86.9 26.1 0 43.1-24.2 38-42 3.7 0 8.3.3 11.7-.6 1.1 27.1 9.1 59.4 41.7 61.7 0-.9 2-7.1 2-7.4 0-17.4-10.6-32.6-10.6-50.3 0-28.3 21.7-55.7 43.7-71.7 8-6 17.7-9.7 27.1-13.1 9.7-3.7 20-8 27.4-15.4-1.1-11.2-5.7-21.1-16.9-21.1-27.7 0-120.6 4-120.6-39.7 0-6.7.1-13.1 17.4-13.1 32.3 0 114.3 8 138.3 29.1 18.1 16.1 24.3 113.2-31 174.7zm-98.6-126c9.7 3.1 19.7 4 29.7 6-7.4 5.4-14 12-20.3 19.1-2.8-8.5-6.2-16.8-9.4-25.1z"],angrycreative:[640,512,[],"f36e","M640 238.2l-3.2 28.2-34.5 2.3-2 18.1 34.5-2.3-3.2 28.2-34.4 2.2-2.3 20.1 34.4-2.2-3 26.1-64.7 4.1 12.7-113.2L527 365.2l-31.9 2-23.8-117.8 30.3-2 13.6 79.4 31.7-82.4 93.1-6.2zM426.8 371.5l28.3-1.8L468 249.6l-28.4 1.9-12.8 120zM162 388.1l-19.4-36-3.5 37.4-28.2 1.7 2.7-29.1c-11 18-32 34.3-56.9 35.8C23.9 399.9-3 377 .3 339.7c2.6-29.3 26.7-62.8 67.5-65.4 37.7-2.4 47.6 23.2 51.3 28.8l2.8-30.8 38.9-2.5c20.1-1.3 38.7 3.7 42.5 23.7l2.6-26.6 64.8-4.2-2.7 27.9-36.4 2.4-1.7 17.9 36.4-2.3-2.7 27.9-36.4 2.3-1.9 19.9 36.3-2.3-2.1 20.8 55-117.2 23.8-1.6L370.4 369l8.9-85.6-22.3 1.4 2.9-27.9 75-4.9-3 28-24.3 1.6-9.7 91.9-58 3.7-4.3-15.6-39.4 2.5-8 16.3-126.2 7.7zm-44.3-70.2l-26.4 1.7C84.6 307.2 76.9 303 65 303.8c-19 1.2-33.3 17.5-34.6 33.3-1.4 16 7.3 32.5 28.7 31.2 12.8-.8 21.3-8.6 28.9-18.9l27-1.7 2.7-29.8zm56.1-7.7c1.2-12.9-7.6-13.6-26.1-12.4l-2.7 28.5c14.2-.9 27.5-2.1 28.8-16.1zm21.1 70.8l5.8-60c-5 13.5-14.7 21.1-27.9 26.6l22.1 33.4zm135.4-45l-7.9-37.8-15.8 39.3 23.7-1.5zm-170.1-74.6l-4.3-17.5-39.6 2.6-8.1 18.2-31.9 2.1 57-121.9 23.9-1.6 30.7 102 9.9-104.7 27-1.8 37.8 63.6 6.5-66.6 28.5-1.9-4 41.2c7.4-13.5 22.9-44.7 63.6-47.5 40.5-2.8 52.4 29.3 53.4 30.3l3.3-32 39.3-2.7c12.7-.9 27.8.3 36.3 9.7l-4.4-11.9 32.2-2.2 12.9 43.2 23-45.7 31-2.2-43.6 78.4-4.8 44.3-28.4 1.9 4.8-44.3-15.8-43c1 22.3-9.2 40.1-32 49.6l25.2 38.8-36.4 2.4-19.2-36.8-4 38.3-28.4 1.9 3.3-31.5c-6.7 9.3-19.7 35.4-59.6 38-26.2 1.7-45.6-10.3-55.4-39.2l-4 40.3-25 1.6-37.6-63.3-6.3 66.2-56.8 3.7zm276.6-82.1c10.2-.7 17.5-2.1 21.6-4.3 4.5-2.4 7-6.4 7.6-12.1.6-5.3-.6-8.8-3.4-10.4-3.6-2.1-10.6-2.8-22.9-2l-2.9 28.8zM327.7 214c5.6 5.9 12.7 8.5 21.3 7.9 4.7-.3 9.1-1.8 13.3-4.1 5.5-3 10.6-8 15.1-14.3l-34.2 2.3 2.4-23.9 63.1-4.3 1.2-12-31.2 2.1c-4.1-3.7-7.8-6.6-11.1-8.1-4-1.7-8.1-2.8-12.2-2.5-8 .5-15.3 3.6-22 9.2-7.7 6.4-12 14.5-12.9 24.4-1.1 9.6 1.4 17.3 7.2 23.3zm-201.3 8.2l23.8-1.6-8.3-37.6-15.5 39.2z"],angular:[415,512,[],"f420","M169.7 268.1h76.2l-38.1-91.6-38.1 91.6zM207.8 32L0 106.4l31.8 275.7 176 97.9 176-97.9 31.8-275.7L207.8 32zM338 373.8h-48.6l-26.2-65.4H152.6l-26.2 65.4H77.7L207.8 81.5 338 373.8z"],"app-store":[512,512,[],"f36f","M255.9 120.9l9.1-15.7c5.6-9.8 18.1-13.1 27.9-7.5 9.8 5.6 13.1 18.1 7.5 27.9l-87.5 151.5h63.3c20.5 0 32 24.1 23.1 40.8H113.8c-11.3 0-20.4-9.1-20.4-20.4 0-11.3 9.1-20.4 20.4-20.4h52l66.6-115.4-20.8-36.1c-5.6-9.8-2.3-22.2 7.5-27.9 9.8-5.6 22.2-2.3 27.9 7.5l8.9 15.7zm-78.7 218l-19.6 34c-5.6 9.8-18.1 13.1-27.9 7.5-9.8-5.6-13.1-18.1-7.5-27.9l14.6-25.2c16.4-5.1 29.8-1.2 40.4 11.6zm168.9-61.7h53.1c11.3 0 20.4 9.1 20.4 20.4 0 11.3-9.1 20.4-20.4 20.4h-29.5l19.9 34.5c5.6 9.8 2.3 22.2-7.5 27.9-9.8 5.6-22.2 2.3-27.9-7.5-33.5-58.1-58.7-101.6-75.4-130.6-17.1-29.5-4.9-59.1 7.2-69.1 13.4 23 33.4 57.7 60.1 104zM256 8C119 8 8 119 8 256s111 248 248 248 248-111 248-248S393 8 256 8zm216 248c0 118.7-96.1 216-216 216-118.7 0-216-96.1-216-216 0-118.7 96.1-216 216-216 118.7 0 216 96.1 216 216z"],"app-store-ios":[448,512,[],"f370","M400 32H48C21.5 32 0 53.5 0 80v352c0 26.5 21.5 48 48 48h352c26.5 0 48-21.5 48-48V80c0-26.5-21.5-48-48-48zM127 384.5c-5.5 9.6-17.8 12.8-27.3 7.3-9.6-5.5-12.8-17.8-7.3-27.3l14.3-24.7c16.1-4.9 29.3-1.1 39.6 11.4L127 384.5zm138.9-53.9H84c-11 0-20-9-20-20s9-20 20-20h51l65.4-113.2-20.5-35.4c-5.5-9.6-2.2-21.8 7.3-27.3 9.6-5.5 21.8-2.2 27.3 7.3l8.9 15.4 8.9-15.4c5.5-9.6 17.8-12.8 27.3-7.3 9.6 5.5 12.8 17.8 7.3 27.3l-85.8 148.6h62.1c20.2 0 31.5 23.7 22.7 40zm98.1 0h-29l19.6 33.9c5.5 9.6 2.2 21.8-7.3 27.3-9.6 5.5-21.8 2.2-27.3-7.3-32.9-56.9-57.5-99.7-74-128.1-16.7-29-4.8-58 7.1-67.8 13.1 22.7 32.7 56.7 58.9 102h52c11 0 20 9 20 20 0 11.1-9 20-20 20z"],apper:[640,512,[],"f371","M42.1 239.1c22.2 0 29 2.8 33.5 14.6h.8v-22.9c0-11.3-4.8-15.4-17.9-15.4-11.3 0-14.4 2.5-15.1 12.8H4.8c.3-13.9 1.5-19.1 5.8-24.4C17.9 195 29.5 192 56.7 192c33 0 47.1 5 53.9 18.9 2 4.3 4 15.6 4 23.7v76.3H76.3l1.3-19.1h-1c-5.3 15.6-13.6 20.4-35.5 20.4-30.3 0-41.1-10.1-41.1-37.3 0-25.2 12.3-35.8 42.1-35.8zm17.1 48.1c13.1 0 16.9-3 16.9-13.4 0-9.1-4.3-11.6-19.6-11.6-13.1 0-17.9 3-17.9 12.1-.1 10.4 3.7 12.9 20.6 12.9zm77.8-94.9h38.3l-1.5 20.6h.8c9.1-17.1 15.9-20.9 37.5-20.9 14.4 0 24.7 3 31.5 9.1 9.8 8.6 12.8 20.4 12.8 48.1 0 30-3 43.1-12.1 52.9-6.8 7.3-16.4 10.1-33.2 10.1-20.4 0-29.2-5.5-33.8-21.2h-.8v70.3H137v-169zm80.9 60.7c0-27.5-3.3-32.5-20.7-32.5-16.9 0-20.7 5-20.7 28.7 0 28 3.5 33.5 21.2 33.5 16.4 0 20.2-5.6 20.2-29.7zm57.9-60.7h38.3l-1.5 20.6h.8c9.1-17.1 15.9-20.9 37.5-20.9 14.4 0 24.7 3 31.5 9.1 9.8 8.6 12.8 20.4 12.8 48.1 0 30-3 43.1-12.1 52.9-6.8 7.3-16.4 10.1-33.3 10.1-20.4 0-29.2-5.5-33.8-21.2h-.8v70.3h-39.5v-169zm80.9 60.7c0-27.5-3.3-32.5-20.7-32.5-16.9 0-20.7 5-20.7 28.7 0 28 3.5 33.5 21.2 33.5 16.4 0 20.2-5.6 20.2-29.7zm53.8-3.8c0-25.4 3.3-37.8 12.3-45.8 8.8-8.1 22.2-11.3 45.1-11.3 42.8 0 55.7 12.8 55.7 55.7v11.1h-75.3c-.3 2-.3 4-.3 4.8 0 16.9 4.5 21.9 20.1 21.9 13.9 0 17.9-3 17.9-13.9h37.5v2.3c0 9.8-2.5 18.9-6.8 24.7-7.3 9.8-19.6 13.6-44.3 13.6-27.5 0-41.6-3.3-50.6-12.3-8.5-8.5-11.3-21.3-11.3-50.8zm76.4-11.6c-.3-1.8-.3-3.3-.3-3.8 0-12.3-3.3-14.6-19.6-14.6-14.4 0-17.1 3-18.1 15.1l-.3 3.3h38.3zm55.6-45.3h38.3l-1.8 19.9h.7c6.8-14.9 14.4-20.2 29.7-20.2 10.8 0 19.1 3.3 23.4 9.3 5.3 7.3 6.8 14.4 6.8 34 0 1.5 0 5 .2 9.3h-35c.3-1.8.3-3.3.3-4 0-15.4-2-19.4-10.3-19.4-6.3 0-10.8 3.3-13.1 9.3-1 3-1 4.3-1 12.3v68h-38.3V192.3z"],apple:[376,512,[],"f179","M314.7 268.7c-.2-36.7 16.4-64.4 50-84.8-18.8-26.9-47.2-41.7-84.7-44.6-35.5-2.8-74.3 20.7-88.5 20.7-15 0-49.4-19.7-76.4-19.7C59.3 141.2 0 184.8 0 273.5c0 26.2 4.8 53.3 14.4 81.2 12.8 36.7 59 126.7 107.2 125.2 25.2-.6 43-17.9 75.8-17.9 31.8 0 48.3 17.9 76.4 17.9 48.6-.7 90.4-82.5 102.6-119.3-65.2-30.7-61.7-90-61.7-91.9zm-56.6-164.2c27.3-32.4 24.8-61.9 24-72.5-24.1 1.4-52 16.4-67.9 34.9-17.5 19.8-27.8 44.3-25.6 71.9 26.1 2 49.9-11.4 69.5-34.3z"],"apple-pay":[640,512,[],"f415","M116.9 158.5c-7.5 8.9-19.5 15.9-31.5 14.9-1.5-12 4.4-24.8 11.3-32.6 7.5-9.1 20.6-15.6 31.3-16.1 1.2 12.4-3.7 24.7-11.1 33.8m10.9 17.2c-17.4-1-32.3 9.9-40.5 9.9-8.4 0-21-9.4-34.8-9.1-17.9.3-34.5 10.4-43.6 26.5-18.8 32.3-4.9 80 13.3 106.3 8.9 13 19.5 27.3 33.5 26.8 13.3-.5 18.5-8.6 34.5-8.6 16.1 0 20.8 8.6 34.8 8.4 14.5-.3 23.6-13 32.5-26 10.1-14.8 14.3-29.1 14.5-29.9-.3-.3-28-10.9-28.3-42.9-.3-26.8 21.9-39.5 22.9-40.3-12.5-18.6-32-20.6-38.8-21.1m100.4-36.2v194.9h30.3v-66.6h41.9c38.3 0 65.1-26.3 65.1-64.3s-26.4-64-64.1-64h-73.2zm30.3 25.5h34.9c26.3 0 41.3 14 41.3 38.6s-15 38.8-41.4 38.8h-34.8V165zm162.2 170.9c19 0 36.6-9.6 44.6-24.9h.6v23.4h28v-97c0-28.1-22.5-46.3-57.1-46.3-32.1 0-55.9 18.4-56.8 43.6h27.3c2.3-12 13.4-19.9 28.6-19.9 18.5 0 28.9 8.6 28.9 24.5v10.8l-37.8 2.3c-35.1 2.1-54.1 16.5-54.1 41.5.1 25.2 19.7 42 47.8 42zm8.2-23.1c-16.1 0-26.4-7.8-26.4-19.6 0-12.3 9.9-19.4 28.8-20.5l33.6-2.1v11c0 18.2-15.5 31.2-36 31.2zm102.5 74.6c29.5 0 43.4-11.3 55.5-45.4L640 193h-30.8l-35.6 115.1h-.6L537.4 193h-31.6L557 334.9l-2.8 8.6c-4.6 14.6-12.1 20.3-25.5 20.3-2.4 0-7-.3-8.9-.5v23.4c1.8.4 9.3.7 11.6.7z"],asymmetrik:[576,512,[],"f372","M517.5 309.2c38.8-40 58.1-80 58.5-116.1.8-65.5-59.4-118.2-169.4-135C277.9 38.4 118.1 73.6 0 140.5 52 114 110.6 92.3 170.7 82.3c74.5-20.5 153-25.4 221.3-14.8C544.5 91.3 588.8 195 490.8 299.2c-10.2 10.8-22 21.1-35 30.6L304.9 103.4 114.7 388.9c-65.6-29.4-76.5-90.2-19.1-151.2 20.8-22.2 48.3-41.9 79.5-58.1 20-12.2 39.7-22.6 62-30.7-65.1 20.3-122.7 52.9-161.6 92.9-27.7 28.6-41.4 57.1-41.7 82.9-.5 35.1 23.4 65.1 68.4 83l-34.5 51.7h101.6l22-34.4c22.2 1 45.3 0 68.6-2.7l-22.8 37.1h135.5L340 406.3c18.6-5.3 36.9-11.5 54.5-18.7l45.9 71.8H542L468.6 349c18.5-12.1 35-25.5 48.9-39.8zm-187.6 80.5l-25-40.6-32.7 53.3c-23.4 3.5-46.7 5.1-69.2 4.4l101.9-159.3 78.7 123c-17.2 7.4-35.3 13.9-53.7 19.2z"],audible:[640,512,[],"f373","M640 199.9v54l-320 200L0 254v-54l320 200 320-200.1zm-194.5 72l47.1-29.4c-37.2-55.8-100.7-92.6-172.7-92.6-72 0-135.5 36.7-172.6 92.4h.3c2.5-2.3 5.1-4.5 7.7-6.7 89.7-74.4 219.4-58.1 290.2 36.3zm-220.1 18.8c16.9-11.9 36.5-18.7 57.4-18.7 34.4 0 65.2 18.4 86.4 47.6l45.4-28.4c-20.9-29.9-55.6-49.5-94.8-49.5-38.9 0-73.4 19.4-94.4 49zM103.6 161.1c131.8-104.3 318.2-76.4 417.5 62.1l.7 1 48.8-30.4C517.1 112.1 424.8 58.1 319.9 58.1c-103.5 0-196.6 53.5-250.5 135.6 9.9-10.5 22.7-23.5 34.2-32.6zm467 32.7z"],autoprefixer:[640,512,[],"f41c","M318.4 16l-161 480h77.5l25.4-81.4h119.5L405 496h77.5L318.4 16zm-40.3 341.9l41.2-130.4h1.5l40.9 130.4h-83.6zM640 405l-10-31.4L462.1 358l19.4 56.5L640 405zm-462.1-47L10 373.7 0 405l158.5 9.4 19.4-56.4z"],avianex:[512,512,[],"f374","M453.1 32h-312c-38.9 0-76.2 31.2-83.3 69.7L1.2 410.3C-5.9 448.8 19.9 480 58.9 480h312c38.9 0 76.2-31.2 83.3-69.7l56.7-308.5c7-38.6-18.8-69.8-57.8-69.8zm-58.2 347.3l-32 13.5-115.4-110c-14.7 10-29.2 19.5-41.7 27.1l22.1 64.2-17.9 12.7-40.6-61-52.4-48.1 15.7-15.4 58 31.1c9.3-10.5 20.8-22.6 32.8-34.9L203 228.9l-68.8-99.8 18.8-28.9 8.9-4.8L265 207.8l4.9 4.5c19.4-18.8 33.8-32.4 33.8-32.4 7.7-6.5 21.5-2.9 30.7 7.9 9 10.5 10.6 24.7 2.7 31.3-1.8 1.3-15.5 11.4-35.3 25.6l4.5 7.3 94.9 119.4-6.3 7.9z"],aviato:[640,512,[],"f421","M107.2 283.5l-19-41.8H36.1l-19 41.8H0l62.2-131.4 62.2 131.4h-17.2zm-45-98.1l-19.6 42.5h39.2l-19.6-42.5zm112.7 102.4l-62.2-131.4h17.1l45.1 96 45.1-96h17l-62.1 131.4zm80.6-4.3V156.4H271v127.1h-15.5zm209.1-115.6v115.6h-17.3V167.9h-41.2v-11.5h99.6v11.5h-41.1zM640 218.8c0 9.2-1.7 17.8-5.1 25.8-3.4 8-8.2 15.1-14.2 21.1-6 6-13.1 10.8-21.1 14.2-8 3.4-16.6 5.1-25.8 5.1s-17.8-1.7-25.8-5.1c-8-3.4-15.1-8.2-21.1-14.2-6-6-10.8-13-14.2-21.1-3.4-8-5.1-16.6-5.1-25.8s1.7-17.8 5.1-25.8c3.4-8 8.2-15.1 14.2-21.1 6-6 13-8.4 21.1-11.9 8-3.4 16.6-5.1 25.8-5.1s17.8 1.7 25.8 5.1c8 3.4 15.1 5.8 21.1 11.9 6 6 10.7 13.1 14.2 21.1 3.4 8 5.1 16.6 5.1 25.8zm-15.5 0c0-7.3-1.3-14-3.9-20.3-2.6-6.3-6.2-11.7-10.8-16.3-4.6-4.6-10-8.2-16.2-10.9-6.2-2.7-12.8-4-19.8-4s-13.6 1.3-19.8 4c-6.2 2.7-11.6 6.3-16.2 10.9-4.6 4.6-8.2 10-10.8 16.3-2.6 6.3-3.9 13.1-3.9 20.3 0 7.3 1.3 14 3.9 20.3 2.6 6.3 6.2 11.7 10.8 16.3 4.6 4.6 10 8.2 16.2 10.9 6.2 2.7 12.8 4 19.8 4s13.6-1.3 19.8-4c6.2-2.7 11.6-6.3 16.2-10.9 4.6-4.6 8.2-10 10.8-16.3 2.6-6.3 3.9-13.1 3.9-20.3zm-94.8 96.7v-6.3l88.9-10-242.9 13.4c.6-2.2 1.1-4.6 1.4-7.2.3-2 .5-4.2.6-6.5l64.8-8.1-64.9 1.9c0-.4-.1-.7-.1-1.1-2.8-17.2-25.5-23.7-25.5-23.7l-1.1-26.3h23.8l19 41.8h17.1L348.6 152l-62.2 131.4h17.1l19-41.8h23.6L345 268s-22.7 6.5-25.5 23.7c-.1.3-.1.7-.1 1.1l-64.9-1.9 64.8 8.1c.1 2.3.3 4.4.6 6.5.3 2.6.8 5 1.4 7.2L78.4 299.2l88.9 10v6.3c-5.9.9-10.5 6-10.5 12.2 0 6.8 5.6 12.4 12.4 12.4 6.8 0 12.4-5.6 12.4-12.4 0-6.2-4.6-11.3-10.5-12.2v-5.8l80.3 9v5.4c-5.7 1.1-9.9 6.2-9.9 12.1 0 6.8 5.6 10.2 12.4 10.2 6.8 0 12.4-3.4 12.4-10.2 0-6-4.3-11-9.9-12.1v-4.9l28.4 3.2v23.7h-5.9V360h5.9v-6.6h5v6.6h5.9v-13.8h-5.9V323l38.3 4.3c8.1 11.4 19 13.6 19 13.6l-.1 6.7-5.1.2-.1 12.1h4.1l.1-5h5.2l.1 5h4.1l-.1-12.1-5.1-.2-.1-6.7s10.9-2.1 19-13.6l38.3-4.3v23.2h-5.9V360h5.9v-6.6h5v6.6h5.9v-13.8h-5.9v-23.7l28.4-3.2v4.9c-5.7 1.1-9.9 6.2-9.9 12.1 0 6.8 5.6 10.2 12.4 10.2 6.8 0 12.4-3.4 12.4-10.2 0-6-4.3-11-9.9-12.1v-5.4l80.3-9v5.8c-5.9.9-10.5 6-10.5 12.2 0 6.8 5.6 12.4 12.4 12.4 6.8 0 12.4-5.6 12.4-12.4-.2-6.3-4.7-11.4-10.7-12.3zm-200.8-87.6l19.6-42.5 19.6 42.5h-17.9l-1.7-40.3-1.7 40.3h-17.9z"],aws:[512,512,[],"f375","M261.2 136.1c-14 57.5-13.1 54.4-25.8 107-1.6 6.5-4.1 8.4-10.7 8.5h-14.4c-5.8-.1-8.2-1.6-9.9-7.3-12.3-39.4-28.8-94.1-39.9-130.7-4.1-13.5-1.4-13.2 9.3-12.9 3.7.1 7.3 0 11 0 5.1.1 7.7 2 9.1 7.1 3.6 12.9 6 22.8 26.6 104.1.4 1.6.9 3.1 1.4 4.6h1.1c.5-2 1.1-3.9 1.6-5.9 7.8-32.9 15.5-65.9 23.3-98.8 2.4-10.2 6.7-11.2 17-11.2h7.6c6.9.1 9 1.5 10.7 8.3 6 23.4 23.5 101.8 26.7 110.4 5.1-18.3-1.8 7.9 28.5-109 2.1-8.1 4.1-9.7 12.3-9.7h12.7c5.4.1 7 1.8 5.7 7.1-2.4 9.5-2.9 9.9-41.3 132.9-3.1 9.9-4.2 10.8-14.6 10.8h-10.6c-7.3 0-9.2-1.3-11-8.4-4.3-16.2-23.3-95.7-26.4-106.9zM125.4 247.3c4.2 5.8 8.1 6.3 14.1 2.4l6.3-4.2c6.8-4.5 7.3-6.3 3.6-13.5-4.3-8.4-6.4-17.3-6.3-26.9 0-3.1.6-55.7-.9-66.8-2.7-19.3-12.5-32.8-31.7-38.7-10.7-3.4-21.7-3.3-32.7-3-15.1.4-29.4 4.6-42.8 11.4-1.8.9-3.7 3.1-4.1 4.9-.8 3.9-1.1 8.1-.7 12.1.6 5.9 2.6 7 8.2 5.1 5.1-1.7 10-3.9 15.1-5.4 14.5-4.4 29.2-6.4 44.1-1.7 7.1 2.2 11.7 6.9 14.3 13.8 3 7.9 2.4 16.1 2.4 24.2 0 5.5-.1 5.5-5.5 4.5-13.9-2.6-27.7-5-41.9-3.1-15.2 2.1-28.6 7.3-38.2 20-9.1 12-10 25.6-7.4 39.5 2.8 15 11.8 25.7 26.4 30.4 20.6 6.7 40.1 3.3 57.7-9.5 3.8-2.8 7.2-6.2 11.1-9.5 3.1 5 5.8 9.7 8.9 14zm-15.3-61.6c3 .4 4.5 1.9 4.3 5.1-.2 3.8.1 7.6-.3 11.4-1.2 11.7-7.7 19.7-17.9 24.9-8.2 4.2-16.9 5.8-26.1 5-15.2-1.3-21-13.1-19.6-26.3C51.8 193.2 59 186.2 72 184c13.8-2.4 16-1.1 38.1 1.7zm348.8 65.1c21.3-8.6 32.9-26.2 29.2-50-2.2-14.6-11.8-24.2-25.2-29.5-14.7-5.9-33.8-10.3-48.1-18.2-4.4-2.4-7.4-6.3-7.6-11.9-.4-11.1 4.2-17.2 15.4-19.8 9.3-2.1 18.8-2.2 28.1-.4 7.3 1.4 14.3 4.2 21.4 6.3 2.8.9 5.9 2.1 7.8-1.6 3.8-7.3.4-18.7-7.3-21.8-22.5-9-45.5-11.6-68.2-1.6-14.6 6.4-24.6 17.4-26 34.2-1.6 19.3 6.9 33.4 24.1 41.7 7.7 3.7 16.1 5.9 24.2 8.9 8.1 3 16.2 5.8 24.1 9.1 12.3 5.3 11.6 24.2 1.2 30-27.7 15.3-64.9-2.4-69.2-3.8-3.3-1.1-5.3.2-6.3 3.7-3 11.3.7 18.8 11.6 22.7 21.7 7.9 49.6 10.5 70.8 2zM296 413.5c50.8-5.8 98.7-20.8 142.7-47 8-4.7 15.5-10.3 23.1-15.7 7.3-5.2 3.2-18.4-11.3-12.2-54.4 23.2-111.2 36.1-170.2 38.9-30.5 1.5-60.8-.3-91.1-4.7-63.1-9.2-122.4-29.2-177.6-61.2-2.1-1.2-4.2-2.5-6.5-3-4.9-1.1-7.7 4.7-2.4 9.7 24 22.1 50.3 40.8 79.1 55.7 53.7 27.7 110.5 42.7 171.2 42 14.4-.8 28.8-.9 43-2.5zm174.7-92.2c14.8.8 19.4 5.9 15.7 20.2-3.8 14.8-9.3 29.2-13.9 43.8-.9 2.9-4.2 6.3-.8 8.8 3.7 2.6 6.5-1 9-3.3 10.2-9.5 17.4-21 22.5-33.8 5.4-13.4 9.3-27.2 8.7-41.9-.2-6.2-1.8-8.8-7.8-10.5-5.4-1.5-11-2.8-16.5-3.2-21.6-1.8-42.5.5-62 10.6-3.1 1.6-6 3.7-8.7 5.9-1.1.9-3.2 5.3 2.4 6.1 1.9.3 3.9-.1 5.9-.3 16.9-1.6 28.6-3.3 45.5-2.4z"],bandcamp:[496,512,[],"f2d5","M248 8C111 8 0 119 0 256s111 248 248 248 248-111 248-248S385 8 248 8zm48.2 326.1h-181L199.9 178h181l-84.7 156.1z"],behance:[576,512,[],"f1b4","M232 237.2c31.8-15.2 48.4-38.2 48.4-74 0-70.6-52.6-87.8-113.3-87.8H0v354.4h171.8c64.4 0 124.9-30.9 124.9-102.9 0-44.5-21.1-77.4-64.7-89.7zM77.9 135.9H151c28.1 0 53.4 7.9 53.4 40.5 0 30.1-19.7 42.2-47.5 42.2h-79v-82.7zm83.3 233.7H77.9V272h84.9c34.3 0 56 14.3 56 50.6 0 35.8-25.9 47-57.6 47zm358.5-240.7H376V94h143.7v34.9zM576 305.2c0-75.9-44.4-139.2-124.9-139.2-78.2 0-131.3 58.8-131.3 135.8 0 79.9 50.3 134.7 131.3 134.7 61.3 0 101-27.6 120.1-86.3H509c-6.7 21.9-34.3 33.5-55.7 33.5-41.3 0-63-24.2-63-65.3h185.1c.3-4.2.6-8.7.6-13.2zM390.4 274c2.3-33.7 24.7-54.8 58.5-54.8 35.4 0 53.2 20.8 56.2 54.8H390.4z"],"behance-square":[448,512,[],"f1b5","M186.5 293c0 19.3-14 25.4-31.2 25.4h-45.1v-52.9h46c18.6.1 30.3 7.8 30.3 27.5zm-7.7-82.3c0-17.7-13.7-21.9-28.9-21.9h-39.6v44.8H153c15.1 0 25.8-6.6 25.8-22.9zm132.3 23.2c-18.3 0-30.5 11.4-31.7 29.7h62.2c-1.7-18.5-11.3-29.7-30.5-29.7zM448 80v352c0 26.5-21.5 48-48 48H48c-26.5 0-48-21.5-48-48V80c0-26.5 21.5-48 48-48h352c26.5 0 48 21.5 48 48zM271.7 185h77.8v-18.9h-77.8V185zm-43 110.3c0-24.1-11.4-44.9-35-51.6 17.2-8.2 26.2-17.7 26.2-37 0-38.2-28.5-47.5-61.4-47.5H68v192h93.1c34.9-.2 67.6-16.9 67.6-55.9zM380 280.5c0-41.1-24.1-75.4-67.6-75.4-42.4 0-71.1 31.8-71.1 73.6 0 43.3 27.3 73 71.1 73 33.2 0 54.7-14.9 65.1-46.8h-33.7c-3.7 11.9-18.6 18.1-30.2 18.1-22.4 0-34.1-13.1-34.1-35.3h100.2c.1-2.3.3-4.8.3-7.2z"],bimobject:[448,512,[],"f378","M416 32H32C14.4 32 0 46.4 0 64v384c0 17.6 14.4 32 32 32h384c17.6 0 32-14.4 32-32V64c0-17.6-14.4-32-32-32zm-64 257.4c0 49.4-11.4 82.6-103.8 82.6h-16.9c-44.1 0-62.4-14.9-70.4-38.8h-.9V368H96V136h64v74.7h1.1c4.6-30.5 39.7-38.8 69.7-38.8h17.3c92.4 0 103.8 33.1 103.8 82.5v35zm-64-28.9v22.9c0 21.7-3.4 33.8-38.4 33.8h-45.3c-28.9 0-44.1-6.5-44.1-35.7v-19c0-29.3 15.2-35.7 44.1-35.7h45.3c35-.2 38.4 12 38.4 33.7z"],bitbucket:[512,512,[],"f171","M23.1 32C14.2 31.9 7 38.9 6.9 47.8c0 .9.1 1.8.2 2.8L74.9 462c1.7 10.4 10.7 18 21.2 18.1h325.1c7.9.1 14.7-5.6 16-13.4l67.8-416c1.4-8.7-4.5-16.9-13.2-18.3-.9-.1-1.8-.2-2.8-.2L23.1 32zm285.3 297.3H204.6l-28.1-146.8h157l-25.1 146.8z"],bitcoin:[512,512,[],"f379","M504 256c0 136.967-111.033 248-248 248S8 392.967 8 256 119.033 8 256 8s248 111.033 248 248zm-141.651-35.33c4.937-32.999-20.191-50.739-54.55-62.573l11.146-44.702-27.213-6.781-10.851 43.524c-7.154-1.783-14.502-3.464-21.803-5.13l10.929-43.81-27.198-6.781-11.153 44.686c-5.922-1.349-11.735-2.682-17.377-4.084l.031-.14-37.53-9.37-7.239 29.062s20.191 4.627 19.765 4.913c11.022 2.751 13.014 10.044 12.68 15.825l-12.696 50.925c.76.194 1.744.473 2.829.907-.907-.225-1.876-.473-2.876-.713l-17.796 71.338c-1.349 3.348-4.767 8.37-12.471 6.464.271.395-19.78-4.937-19.78-4.937l-13.51 31.147 35.414 8.827c6.588 1.651 13.045 3.379 19.4 5.006l-11.262 45.213 27.182 6.781 11.153-44.733a1038.209 1038.209 0 0 0 21.687 5.627l-11.115 44.523 27.213 6.781 11.262-45.128c46.404 8.781 81.299 5.239 95.986-36.727 11.836-33.79-.589-53.281-25.004-65.991 17.78-4.098 31.174-15.792 34.747-39.949zm-62.177 87.179c-8.41 33.79-65.308 15.523-83.755 10.943l14.944-59.899c18.446 4.603 77.6 13.717 68.811 48.956zm8.417-87.667c-7.673 30.736-55.031 15.12-70.393 11.292l13.548-54.327c15.363 3.828 64.836 10.973 56.845 43.035z"],bity:[496,512,[],"f37a","M78.4 67.2C173.8-22 324.5-24 421.5 71c14.3 14.1-6.4 37.1-22.4 21.5-84.8-82.4-215.8-80.3-298.9-3.2-16.3 15.1-36.5-8.3-21.8-22.1zm98.9 418.6c19.3 5.7 29.3-23.6 7.9-30C73 421.9 9.4 306.1 37.7 194.8c5-19.6-24.9-28.1-30.2-7.1-32.1 127.4 41.1 259.8 169.8 298.1zm148.1-2c121.9-40.2 192.9-166.9 164.4-291-4.5-19.7-34.9-13.8-30 7.9 24.2 107.7-37.1 217.9-143.2 253.4-21.2 7-10.4 36 8.8 29.7zm-62.9-79l.2-71.8c0-8.2-6.6-14.8-14.8-14.8-8.2 0-14.8 6.7-14.8 14.8l-.2 71.8c0 8.2 6.6 14.8 14.8 14.8s14.8-6.6 14.8-14.8zm71-269c2.1 90.9 4.7 131.9-85.5 132.5-92.5-.7-86.9-44.3-85.5-132.5 0-21.8-32.5-19.6-32.5 0v71.6c0 69.3 60.7 90.9 118 90.1 57.3.8 118-20.8 118-90.1v-71.6c0-19.6-32.5-21.8-32.5 0z"],"black-tie":[448,512,[],"f27e","M0 32v448h448V32H0zm316.5 325.2L224 445.9l-92.5-88.7 64.5-184-64.5-86.6h184.9L252 173.2l64.5 184z"],blackberry:[512,512,[],"f37b","M166 116.9c0 23.4-16.4 49.1-72.5 49.1H23.4l21-88.8h67.8c42.1 0 53.8 23.3 53.8 39.7zm126.2-39.7h-67.8L205.7 166h70.1c53.8 0 70.1-25.7 70.1-49.1.1-16.4-11.6-39.7-53.7-39.7zM88.8 208.1H21L0 296.9h70.1c56.1 0 72.5-23.4 72.5-49.1 0-16.3-11.7-39.7-53.8-39.7zm180.1 0h-67.8l-18.7 88.8h70.1c53.8 0 70.1-23.4 70.1-49.1 0-16.3-11.7-39.7-53.7-39.7zm189.3-53.8h-67.8l-18.7 88.8h70.1c53.8 0 70.1-23.4 70.1-49.1.1-16.3-11.6-39.7-53.7-39.7zm-28 137.9h-67.8L343.7 381h70.1c56.1 0 70.1-23.4 70.1-49.1 0-16.3-11.6-39.7-53.7-39.7zM240.8 346H173l-18.7 88.8h70.1c56.1 0 70.1-25.7 70.1-49.1.1-16.3-11.6-39.7-53.7-39.7z"],blogger:[448,512,[],"f37c","M162.4 196c4.8-4.9 6.2-5.1 36.4-5.1 27.2 0 28.1.1 32.1 2.1 5.8 2.9 8.3 7 8.3 13.6 0 5.9-2.4 10-7.6 13.4-2.8 1.8-4.5 1.9-31.1 2.1-16.4.1-29.5-.2-31.5-.8-10.3-2.9-14.1-17.7-6.6-25.3zm61.4 94.5c-53.9 0-55.8.2-60.2 4.1-3.5 3.1-5.7 9.4-5.1 13.9.7 4.7 4.8 10.1 9.2 12 2.2 1 14.1 1.7 56.3 1.2l47.9-.6 9.2-1.5c9-5.1 10.5-17.4 3.1-24.4-5.3-4.7-5-4.7-60.4-4.7zm223.4 130.1c-3.5 28.4-23 50.4-51.1 57.5-7.2 1.8-9.7 1.9-172.9 1.8-157.8 0-165.9-.1-172-1.8-8.4-2.2-15.6-5.5-22.3-10-5.6-3.8-13.9-11.8-17-16.4-3.8-5.6-8.2-15.3-10-22C.1 423 0 420.3 0 256.3 0 93.2 0 89.7 1.8 82.6 8.1 57.9 27.7 39 53 33.4c7.3-1.6 332.1-1.9 340-.3 21.2 4.3 37.9 17.1 47.6 36.4 7.7 15.3 7-1.5 7.3 180.6.2 115.8 0 164.5-.7 170.5zm-85.4-185.2c-1.1-5-4.2-9.6-7.7-11.5-1.1-.6-8-1.3-15.5-1.7-12.4-.6-13.8-.8-17.8-3.1-6.2-3.6-7.9-7.6-8-18.3 0-20.4-8.5-39.4-25.3-56.5-12-12.2-25.3-20.5-40.6-25.1-3.6-1.1-11.8-1.5-39.2-1.8-42.9-.5-52.5.4-67.1 6.2-27 10.7-46.3 33.4-53.4 62.4-1.3 5.4-1.6 14.2-1.9 64.3-.4 62.8 0 72.1 4 84.5 9.7 30.7 37.1 53.4 64.6 58.4 9.2 1.7 122.2 2.1 133.7.5 20.1-2.7 35.9-10.8 50.7-25.9 10.7-10.9 17.4-22.8 21.8-38.5 3.2-10.9 2.9-88.4 1.7-93.9z"],"blogger-b":[448,512,[],"f37d","M446.6 222.7c-1.8-8-6.8-15.4-12.5-18.5-1.8-1-13-2.2-25-2.7-20.1-.9-22.3-1.3-28.7-5-10.1-5.9-12.8-12.3-12.9-29.5-.1-33-13.8-63.7-40.9-91.3-19.3-19.7-40.9-33-65.5-40.5-5.9-1.8-19.1-2.4-63.3-2.9-69.4-.8-84.8.6-108.4 10C45.9 59.5 14.7 96.1 3.3 142.9 1.2 151.7.7 165.8.2 246.8c-.6 101.5.1 116.4 6.4 136.5 15.6 49.6 59.9 86.3 104.4 94.3 14.8 2.7 197.3 3.3 216 .8 32.5-4.4 58-17.5 81.9-41.9 17.3-17.7 28.1-36.8 35.2-62.1 4.9-17.6 4.5-142.8 2.5-151.7zm-322.1-63.6c7.8-7.9 10-8.2 58.8-8.2 43.9 0 45.4.1 51.8 3.4 9.3 4.7 13.4 11.3 13.4 21.9 0 9.5-3.8 16.2-12.3 21.6-4.6 2.9-7.3 3.1-50.3 3.3-26.5.2-47.7-.4-50.8-1.2-16.6-4.7-22.8-28.5-10.6-40.8zm191.8 199.8l-14.9 2.4-77.5.9c-68.1.8-87.3-.4-90.9-2-7.1-3.1-13.8-11.7-14.9-19.4-1.1-7.3 2.6-17.3 8.2-22.4 7.1-6.4 10.2-6.6 97.3-6.7 89.6-.1 89.1-.1 97.6 7.8 12.1 11.3 9.5 31.2-4.9 39.4z"],bluetooth:[448,512,[],"f293","M292.6 171.1L249.7 214l-.3-86 43.2 43.1m-43.2 219.8l43.1-43.1-42.9-42.9-.2 86zM416 259.4C416 465 344.1 512 230.9 512S32 465 32 259.4 115.4 0 228.6 0 416 53.9 416 259.4zm-158.5 0l79.4-88.6L211.8 36.5v176.9L138 139.6l-27 26.9 92.7 93-92.7 93 26.9 26.9 73.8-73.8 2.3 170 127.4-127.5-83.9-88.7z"],"bluetooth-b":[320,512,[],"f294","M196.48 260.023l92.626-103.333L143.125 0v206.33l-86.111-86.111-31.406 31.405 108.061 108.399L25.608 368.422l31.406 31.405 86.111-86.111L145.84 512l148.552-148.644-97.912-103.333zm40.86-102.996l-49.977 49.978-.338-100.295 50.315 50.317zM187.363 313.04l49.977 49.978-50.315 50.316.338-100.294z"],btc:[384,512,[],"f15a","M310.204 242.638c27.73-14.18 45.377-39.39 41.28-81.3-5.358-57.351-52.458-76.573-114.85-81.929V0h-48.528v77.203c-12.605 0-25.525.315-38.444.63V0h-48.528v79.409c-17.842.539-38.622.276-97.37 0v51.678c38.314-.678 58.417-3.14 63.023 21.427v217.429c-2.925 19.492-18.524 16.685-53.255 16.071L3.765 443.68c88.481 0 97.37.315 97.37.315V512h48.528v-67.06c13.234.315 26.154.315 38.444.315V512h48.528v-68.005c81.299-4.412 135.647-24.894 142.895-101.467 5.671-61.446-23.32-88.862-69.326-99.89zM150.608 134.553c27.415 0 113.126-8.507 113.126 48.528 0 54.515-85.71 48.212-113.126 48.212v-96.74zm0 251.776V279.821c32.772 0 133.127-9.138 133.127 53.255-.001 60.186-100.355 53.253-133.127 53.253z"],buromobelexperte:[448,512,[],"f37f","M0 32v128h128V32H0zm120 120H8V40h112v112zm40-120v128h128V32H160zm120 120H168V40h112v112zm40-120v128h128V32H320zm120 120H328V40h112v112zM0 192v128h128V192H0zm120 120H8V200h112v112zm40-120v128h128V192H160zm120 120H168V200h112v112zm40-120v128h128V192H320zm120 120H328V200h112v112zM0 352v128h128V352H0zm120 120H8V360h112v112zm40-120v128h128V352H160zm120 120H168V360h112v112zm40-120v128h128V352H320z"],buysellads:[448,512,[],"f20d","M224 150.7l42.9 160.7h-85.8L224 150.7zM448 80v352c0 26.5-21.5 48-48 48H48c-26.5 0-48-21.5-48-48V80c0-26.5 21.5-48 48-48h352c26.5 0 48 21.5 48 48zm-65.3 325.3l-94.5-298.7H159.8L65.3 405.3H156l111.7-91.6 24.2 91.6h90.8z"],"cc-amazon-pay":[576,512,[],"f42d","M124.7 201.8c.1-11.8 0-23.5 0-35.3v-35.3c0-1.3.4-2 1.4-2.7 11.5-8 24.1-12.1 38.2-11.1 12.5.9 22.7 7 28.1 21.7 3.3 8.9 4.1 18.2 4.1 27.7 0 8.7-.7 17.3-3.4 25.6-5.7 17.8-18.7 24.7-35.7 23.9-11.7-.5-21.9-5-31.4-11.7-.9-.8-1.4-1.6-1.3-2.8zm154.9 14.6c4.6 1.8 9.3 2 14.1 1.5 11.6-1.2 21.9-5.7 31.3-12.5.9-.6 1.3-1.3 1.3-2.5-.1-3.9 0-7.9 0-11.8 0-4-.1-8 0-12 0-1.4-.4-2-1.8-2.2-7-.9-13.9-2.2-20.9-2.9-7-.6-14-.3-20.8 1.9-6.7 2.2-11.7 6.2-13.7 13.1-1.6 5.4-1.6 10.8.1 16.2 1.6 5.5 5.2 9.2 10.4 11.2zM576 80v352c0 26.5-21.5 48-48 48H48c-26.5 0-48-21.5-48-48V80c0-26.5 21.5-48 48-48h480c26.5 0 48 21.5 48 48zm-207.5 23.9c.4 1.7.9 3.4 1.6 5.1 16.5 40.6 32.9 81.3 49.5 121.9 1.4 3.5 1.7 6.4.2 9.9-2.8 6.2-4.9 12.6-7.8 18.7-2.6 5.5-6.7 9.5-12.7 11.2-4.2 1.1-8.5 1.3-12.9.9-2.1-.2-4.2-.7-6.3-.8-2.8-.2-4.2 1.1-4.3 4-.1 2.8-.1 5.6 0 8.3.1 4.6 1.6 6.7 6.2 7.5 4.7.8 9.4 1.6 14.2 1.7 14.3.3 25.7-5.4 33.1-17.9 2.9-4.9 5.6-10.1 7.7-15.4 19.8-50.1 39.5-100.3 59.2-150.5.6-1.5 1.1-3 1.3-4.6.4-2.4-.7-3.6-3.1-3.7-5.6-.1-11.1 0-16.7 0-3.1 0-5.3 1.4-6.4 4.3-.4 1.1-.9 2.3-1.3 3.4l-29.1 83.7c-2.1 6.1-4.2 12.1-6.5 18.6-.4-.9-.6-1.4-.8-1.9-10.8-29.9-21.6-59.9-32.4-89.8-1.7-4.7-3.5-9.5-5.3-14.2-.9-2.5-2.7-4-5.4-4-6.4-.1-12.8-.2-19.2-.1-2.2 0-3.3 1.6-2.8 3.7zM242.4 206c1.7 11.7 7.6 20.8 18 26.6 9.9 5.5 20.7 6.2 31.7 4.6 12.7-1.9 23.9-7.3 33.8-15.5.4-.3.8-.6 1.4-1 .5 3.2.9 6.2 1.5 9.2.5 2.6 2.1 4.3 4.5 4.4 4.6.1 9.1.1 13.7 0 2.3-.1 3.8-1.6 4-3.9.1-.8.1-1.6.1-2.3v-88.8c0-3.6-.2-7.2-.7-10.8-1.6-10.8-6.2-19.7-15.9-25.4-5.6-3.3-11.8-5-18.2-5.9-3-.4-6-.7-9.1-1.1h-10c-.8.1-1.6.3-2.5.3-8.2.4-16.3 1.4-24.2 3.5-5.1 1.3-10 3.2-15 4.9-3 1-4.5 3.2-4.4 6.5.1 2.8-.1 5.6 0 8.3.1 4.1 1.8 5.2 5.7 4.1 6.5-1.7 13.1-3.5 19.7-4.8 10.3-1.9 20.7-2.7 31.1-1.2 5.4.8 10.5 2.4 14.1 7 3.1 4 4.2 8.8 4.4 13.7.3 6.9.2 13.9.3 20.8 0 .4-.1.7-.2 1.2-.4 0-.8 0-1.1-.1-8.8-2.1-17.7-3.6-26.8-4.1-9.5-.5-18.9.1-27.9 3.2-10.8 3.8-19.5 10.3-24.6 20.8-4.1 8.3-4.6 17-3.4 25.8zM98.7 106.9v175.3c0 .8 0 1.7.1 2.5.2 2.5 1.7 4.1 4.1 4.2 5.9.1 11.8.1 17.7 0 2.5 0 4-1.7 4.1-4.1.1-.8.1-1.7.1-2.5v-60.7c.9.7 1.4 1.2 1.9 1.6 15 12.5 32.2 16.6 51.1 12.9 17.1-3.4 28.9-13.9 36.7-29.2 5.8-11.6 8.3-24.1 8.7-37 .5-14.3-1-28.4-6.8-41.7-7.1-16.4-18.9-27.3-36.7-30.9-2.7-.6-5.5-.8-8.2-1.2h-7c-1.2.2-2.4.3-3.6.5-11.7 1.4-22.3 5.8-31.8 12.7-2 1.4-3.9 3-5.9 4.5-.1-.5-.3-.8-.4-1.2-.4-2.3-.7-4.6-1.1-6.9-.6-3.9-2.5-5.5-6.4-5.6h-9.7c-5.9-.1-6.9 1-6.9 6.8zM493.6 339c-2.7-.7-5.1 0-7.6 1-43.9 18.4-89.5 30.2-136.8 35.8-14.5 1.7-29.1 2.8-43.7 3.2-26.6.7-53.2-.8-79.6-4.3-17.8-2.4-35.5-5.7-53-9.9-37-8.9-72.7-21.7-106.7-38.8-8.8-4.4-17.4-9.3-26.1-14-3.8-2.1-6.2-1.5-8.2 2.1v1.7c1.2 1.6 2.2 3.4 3.7 4.8 36 32.2 76.6 56.5 122 72.9 21.9 7.9 44.4 13.7 67.3 17.5 14 2.3 28 3.8 42.2 4.5 3 .1 6 .2 9 .4.7 0 1.4.2 2.1.3h17.7c.7-.1 1.4-.3 2.1-.3 14.9-.4 29.8-1.8 44.6-4 21.4-3.2 42.4-8.1 62.9-14.7 29.6-9.6 57.7-22.4 83.4-40.1 2.8-1.9 5.7-3.8 8-6.2 4.3-4.4 2.3-10.4-3.3-11.9zm50.4-27.7c-.8-4.2-4-5.8-7.6-7-5.7-1.9-11.6-2.8-17.6-3.3-11-.9-22-.4-32.8 1.6-12 2.2-23.4 6.1-33.5 13.1-1.2.8-2.4 1.8-3.1 3-.6.9-.7 2.3-.5 3.4.3 1.3 1.7 1.6 3 1.5.6 0 1.2 0 1.8-.1l19.5-2.1c9.6-.9 19.2-1.5 28.8-.8 4.1.3 8.1 1.2 12 2.2 4.3 1.1 6.2 4.4 6.4 8.7.3 6.7-1.2 13.1-2.9 19.5-3.5 12.9-8.3 25.4-13.3 37.8-.3.8-.7 1.7-.8 2.5-.4 2.5 1 4 3.4 3.5 1.4-.3 3-1.1 4-2.1 3.7-3.6 7.5-7.2 10.6-11.2 10.7-13.8 17-29.6 20.7-46.6.7-3 1.2-6.1 1.7-9.1.2-4.7.2-9.6.2-14.5z"],"cc-amex":[576,512,[],"f1f3","M576 255.4c-37.9-.2-44.2-.9-54.5 5v-5c-45.3 0-53.5-1.7-64.9 5.2v-5.2h-78.2v5.1c-11.4-6.5-21.4-5.1-75.7-5.1v5.6c-6.3-3.7-14.5-5.6-24.3-5.6h-58c-3.5 3.8-12.5 13.7-15.7 17.2-12.7-14.1-10.5-11.6-15.5-17.2h-83.1v92.3h82c3.3-3.5 12.9-13.9 16.1-17.4 12.7 14.3 10.3 11.7 15.4 17.4h48.9c0-14.7.1-8.3.1-23 11.5.2 24.3-.2 34.3-6.2 0 13.9-.1 17.1-.1 29.2h39.6c0-18.5.1-7.4.1-25.3 6.2 0 7.7 0 9.4.1.1 1.3 0 0 0 25.2 152.8 0 145.9 1.1 156.7-4.5v4.5c34.8 0 54.8 2.2 67.5-6.1V432c0 26.5-21.5 48-48 48H48c-26.5 0-48-21.5-48-48V228.3h26.6c4.2-10.1 2.2-5.3 6.4-15.3h19.2c4.2 10 2.2 5.2 6.4 15.3h52.9v-11.4c2.2 5 1.1 2.5 5.1 11.4h29.5c2.4-5.5 2.6-5.8 5.1-11.4v11.4h135.5v-25.1c6.4 0 8-.1 9.8.2 0 0-.2 10.9.1 24.8h66.5v-8.9c7.4 5.9 17.4 8.9 29.7 8.9h26.8c4.2-10.1 2.2-5.3 6.4-15.3h19c6.5 15 .2.5 6.6 15.3h52.8v-21.9c11.8 19.7 7.8 12.9 13.2 21.9h41.6v-92h-39.9v18.4c-12.2-20.2-6.3-10.4-11.2-18.4h-43.3v20.6c-6.2-14.6-4.6-10.8-8.8-20.6h-32.4c-.4 0-2.3.2-2.3-.3h-27.6c-12.8 0-23.1 3.2-30.7 9.3v-9.3h-39.9v5.3c-10.8-6.1-20.7-5.1-64.4-5.3-.1 0-11.6-.1-11.6 0h-103c-2.5 6.1-6.8 16.4-12.6 30-2.8-6-11-23.8-13.9-30h-46V157c-7.4-17.4-4.7-11-9-21.1H22.9c-3.4 7.9-13.7 32-23.1 53.9V80c0-26.5 21.5-48 48-48h480c26.5 0 48 21.5 48 48v175.4zm-186.6-80.6c-.3.2-1.4 2.2-1.4 7.6 0 6 .9 7.7 1.1 7.9.2.1 1.1.5 3.4.5l7.3-16.9c-1.1 0-2.1-.1-3.1-.1-5.6 0-7 .7-7.3 1zm-19.9 130.9c9.2 3.3 11 9.5 11 18.4l-.1 13.8h-16.6l.1-11.5c0-11.8-3.8-13.8-14.8-13.8h-17.6l-.1 25.3h-16.6l.1-69.3h39.4c13 0 27.1 2.3 27.1 18.7-.1 7.6-4.2 15.3-11.9 18.4zm-6.3-15.4c0-6.4-5.6-7.4-10.7-7.4h-21v15.6h20.7c5.6 0 11-1.3 11-8.2zm181.7-7.1H575v-14.6h-32.9c-12.8 0-23.8 6.6-23.8 20.7 0 33 42.7 12.8 42.7 27.4 0 5.1-4.3 6.4-8.4 6.4h-32l-.1 14.8h32c8.4 0 17.6-1.8 22.5-8.9v-25.8c-10.5-13.8-39.3-1.3-39.3-13.5 0-5.8 4.6-6.5 9.2-6.5zm-99.2-.3v-14.3h-55.2l-.1 69.3h55.2l.1-14.3-38.6-.3v-13.8H445v-14.1h-37.8v-12.5h38.5zm42.2 40.1h-32.2l-.1 14.8h32.2c14.8 0 26.2-5.6 26.2-22 0-33.2-42.9-11.2-42.9-26.3 0-5.6 4.9-6.4 9.2-6.4h30.4v-14.6h-33.2c-12.8 0-23.5 6.6-23.5 20.7 0 33 42.7 12.5 42.7 27.4-.1 5.4-4.7 6.4-8.8 6.4zm-78.1-158.7c-17.4-.3-33.2-4.1-33.2 19.7 0 11.8 2.8 19.9 16.1 19.9h7.4l23.5-54.5h24.8l27.9 65.4v-65.4h25.3l29.1 48.1v-48.1h16.9v69H524l-31.2-51.9v51.9h-33.7l-6.6-15.3h-34.3l-6.4 15.3h-19.2c-22.8 0-33-11.8-33-34 0-23.3 10.5-35.3 34-35.3h16.1v15.2zm14.3 24.5h22.8l-11.2-27.6-11.6 27.6zm-72.6-39.6h-16.9v69.3h16.9v-69.3zm-38.1 37.3c9.5 3.3 11 9.2 11 18.4v13.5h-16.6c-.3-14.8 3.6-25.1-14.8-25.1h-18v25.1h-16.4v-69.3l39.1.3c13.3 0 27.4 2 27.4 18.4.1 8-4.3 15.7-11.7 18.7zm-6.7-15.3c0-6.4-5.6-7.4-10.7-7.4h-21v15.3h20.7c5.7 0 11-1.3 11-7.9zm-59.5-7.4v-14.6h-55.5v69.3h55.5v-14.3h-38.9v-13.8h37.8v-14.1h-37.8v-12.5h38.9zm-84.6 54.7v-54.2l-24 54.2H124l-24-54.2v54.2H66.2l-6.4-15.3H25.3l-6.4 15.3H1l29.7-69.3h24.5l28.1 65.7v-65.7h27.1l21.7 47 19.7-47h27.6v69.3h-16.8zM53.9 188.8l-11.5-27.6-11.2 27.6h22.7zm253 102.5c0 27.9-30.4 23.3-49.3 23.3l-.1 23.3h-32.2l-20.4-23-21.3 23h-65.4l.1-69.3h66.5l20.5 22.8 21-22.8H279c15.6 0 27.9 5.4 27.9 22.7zm-112.7 11.8l-17.9-20.2h-41.7v12.5h36.3v14.1h-36.3v13.8h40.6l19-20.2zM241 276l-25.3 27.4 25.3 28.1V276zm48.3 15.3c0-6.1-4.6-8.4-10.2-8.4h-21.5v17.6h21.2c5.9 0 10.5-2.8 10.5-9.2z"],"cc-apple-pay":[576,512,[],"f416","M302.2 218.4c0 17.2-10.5 27.1-29 27.1h-24.3v-54.2h24.4c18.4 0 28.9 9.8 28.9 27.1zm47.5 62.6c0 8.3 7.2 13.7 18.5 13.7 14.4 0 25.2-9.1 25.2-21.9v-7.7l-23.5 1.5c-13.3.9-20.2 5.8-20.2 14.4zM576 79v352c0 26.5-21.5 48-48 48H48c-26.5 0-48-21.5-48-48V79c0-26.5 21.5-48 48-48h480c26.5 0 48 21.5 48 48zM127.8 197.2c8.4.7 16.8-4.2 22.1-10.4 5.2-6.4 8.6-15 7.7-23.7-7.4.3-16.6 4.9-21.9 11.3-4.8 5.5-8.9 14.4-7.9 22.8zm60.6 74.5c-.2-.2-19.6-7.6-19.8-30-.2-18.7 15.3-27.7 16-28.2-8.8-13-22.4-14.4-27.1-14.7-12.2-.7-22.6 6.9-28.4 6.9-5.9 0-14.7-6.6-24.3-6.4-12.5.2-24.2 7.3-30.5 18.6-13.1 22.6-3.4 56 9.3 74.4 6.2 9.1 13.7 19.1 23.5 18.7 9.3-.4 13-6 24.2-6 11.3 0 14.5 6 24.3 5.9 10.2-.2 16.5-9.1 22.8-18.2 6.9-10.4 9.8-20.4 10-21zm135.4-53.4c0-26.6-18.5-44.8-44.9-44.8h-51.2v136.4h21.2v-46.6h29.3c26.8 0 45.6-18.4 45.6-45zm90 23.7c0-19.7-15.8-32.4-40-32.4-22.5 0-39.1 12.9-39.7 30.5h19.1c1.6-8.4 9.4-13.9 20-13.9 13 0 20.2 6 20.2 17.2v7.5l-26.4 1.6c-24.6 1.5-37.9 11.6-37.9 29.1 0 17.7 13.7 29.4 33.4 29.4 13.3 0 25.6-6.7 31.2-17.4h.4V310h19.6v-68zM516 210.9h-21.5l-24.9 80.6h-.4l-24.9-80.6H422l35.9 99.3-1.9 6c-3.2 10.2-8.5 14.2-17.9 14.2-1.7 0-4.9-.2-6.2-.3v16.4c1.2.4 6.5.5 8.1.5 20.7 0 30.4-7.9 38.9-31.8L516 210.9z"],"cc-diners-club":[576,512,[],"f24c","M239.7 79.9c-96.9 0-175.8 78.6-175.8 175.8 0 96.9 78.9 175.8 175.8 175.8 97.2 0 175.8-78.9 175.8-175.8 0-97.2-78.6-175.8-175.8-175.8zm-39.9 279.6c-41.7-15.9-71.4-56.4-71.4-103.8s29.7-87.9 71.4-104.1v207.9zm79.8.3V151.6c41.7 16.2 71.4 56.7 71.4 104.1s-29.7 87.9-71.4 104.1zM528 32H48C21.5 32 0 53.5 0 80v352c0 26.5 21.5 48 48 48h480c26.5 0 48-21.5 48-48V80c0-26.5-21.5-48-48-48zM329.7 448h-90.3c-106.2 0-193.8-85.5-193.8-190.2C45.6 143.2 133.2 64 239.4 64h90.3c105 0 200.7 79.2 200.7 193.8 0 104.7-95.7 190.2-200.7 190.2z"],"cc-discover":[576,512,[],"f1f2","M83 212.1c0 7.9-3.2 15.5-8.9 20.7-4.9 4.4-11.6 6.4-21.9 6.4H48V185h4.2c10.3 0 16.7 1.7 21.9 6.6 5.7 5 8.9 12.6 8.9 20.5zM504.8 184h-4.9v24.9h4.7c10.3 0 15.8-4.4 15.8-12.8 0-7.9-5.5-12.1-15.6-12.1zM576 80v352c0 26.5-21.5 48-48 48H48c-26.5 0-48-21.5-48-48V80c0-26.5 21.5-48 48-48h480c26.5 0 48 21.5 48 48zM428 253h45.3v-13.8H444V217h28.3v-13.8H444V185h29.3v-14H428v82zm-86.2-82l35 84.2h8.6l35.5-84.2h-17.5l-22.2 55.2-21.9-55.2h-17.5zm-83 41.6c0 24.6 19.9 44.6 44.6 44.6 24.6 0 44.6-19.9 44.6-44.6 0-24.6-19.9-44.6-44.6-44.6-24.6 0-44.6 19.9-44.6 44.6zm-68-.5c0 32.5 33.6 52.5 63.3 38.2v-19c-19.3 19.3-46.8 5.8-46.8-19.2 0-23.7 26.7-39.1 46.8-19v-19c-30.2-15-63.3 6.8-63.3 38zm-33.9 28.3c-7.6 0-13.8-3.7-17.5-10.8l-10.3 9.9c17.8 26.1 56.6 18.2 56.6-11.3 0-13.1-5.4-19-23.6-25.6-9.6-3.4-12.3-5.9-12.3-10.3 0-8.7 14.5-14.1 24.9-2.5l8.4-10.8c-19.1-17.1-49.7-8.9-49.7 14.3 0 11.3 5.2 17.2 20.2 22.7 25.7 9.1 14.7 24.4 3.3 24.4zm-57.4-28.3c0-24.1-18-41.1-44.1-41.1H32v82h23.4c30.9 0 44.1-22.4 44.1-40.9zm23.4-41.1h-16v82h16v-82zM544 288c-33.3 20.8-226.4 124.4-416 160h401c8.2 0 15-6.8 15-15V288zm0-35l-25.9-34.5c12.1-2.5 18.7-10.6 18.7-23.2 0-28.5-30.3-24.4-52.9-24.4v82h16v-32.8h2.2l22.2 32.8H544z"],"cc-jcb":[576,512,[],"f24b","M431.5 244.3V212c41.2 0 38.5.2 38.5.2 7.3 1.3 13.3 7.3 13.3 16 0 8.8-6 14.5-13.3 15.8-1.2.4-3.3.3-38.5.3zm42.8 20.2c-2.8-.7-3.3-.5-42.8-.5v35c39.6 0 40 .2 42.8-.5 7.5-1.5 13.5-8 13.5-17 0-8.7-6-15.5-13.5-17zM576 80v352c0 26.5-21.5 48-48 48H48c-26.5 0-48-21.5-48-48V80c0-26.5 21.5-48 48-48h480c26.5 0 48 21.5 48 48zM182 192.3h-57c0 67.1 10.7 109.7-35.8 109.7-19.5 0-38.8-5.7-57.2-14.8v28c30 8.3 68 8.3 68 8.3 97.9 0 82-47.7 82-131.2zm178.5 4.5c-63.4-16-165-14.9-165 59.3 0 77.1 108.2 73.6 165 59.2V287C312.9 311.7 253 309 253 256s59.8-55.6 107.5-31.2v-28zM544 286.5c0-18.5-16.5-30.5-38-32v-.8c19.5-2.7 30.3-15.5 30.3-30.2 0-19-15.7-30-37-31 0 0 6.3-.3-120.3-.3v127.5h122.7c24.3.1 42.3-12.9 42.3-33.2z"],"cc-mastercard":[576,512,[],"f1f1","M482.9 410.3c0 6.8-4.6 11.7-11.2 11.7-6.8 0-11.2-5.2-11.2-11.7 0-6.5 4.4-11.7 11.2-11.7 6.6 0 11.2 5.2 11.2 11.7zm-310.8-11.7c-7.1 0-11.2 5.2-11.2 11.7 0 6.5 4.1 11.7 11.2 11.7 6.5 0 10.9-4.9 10.9-11.7-.1-6.5-4.4-11.7-10.9-11.7zm117.5-.3c-5.4 0-8.7 3.5-9.5 8.7h19.1c-.9-5.7-4.4-8.7-9.6-8.7zm107.8.3c-6.8 0-10.9 5.2-10.9 11.7 0 6.5 4.1 11.7 10.9 11.7 6.8 0 11.2-4.9 11.2-11.7 0-6.5-4.4-11.7-11.2-11.7zm105.9 26.1c0 .3.3.5.3 1.1 0 .3-.3.5-.3 1.1-.3.3-.3.5-.5.8-.3.3-.5.5-1.1.5-.3.3-.5.3-1.1.3-.3 0-.5 0-1.1-.3-.3 0-.5-.3-.8-.5-.3-.3-.5-.5-.5-.8-.3-.5-.3-.8-.3-1.1 0-.5 0-.8.3-1.1 0-.5.3-.8.5-1.1.3-.3.5-.3.8-.5.5-.3.8-.3 1.1-.3.5 0 .8 0 1.1.3.5.3.8.3 1.1.5s.2.6.5 1.1zm-2.2 1.4c.5 0 .5-.3.8-.3.3-.3.3-.5.3-.8 0-.3 0-.5-.3-.8-.3 0-.5-.3-1.1-.3h-1.6v3.5h.8V426h.3l1.1 1.4h.8l-1.1-1.3zM576 81v352c0 26.5-21.5 48-48 48H48c-26.5 0-48-21.5-48-48V81c0-26.5 21.5-48 48-48h480c26.5 0 48 21.5 48 48zM64 220.6c0 76.5 62.1 138.5 138.5 138.5 27.2 0 53.9-8.2 76.5-23.1-72.9-59.3-72.4-171.2 0-230.5-22.6-15-49.3-23.1-76.5-23.1-76.4-.1-138.5 62-138.5 138.2zm224 108.8c70.5-55 70.2-162.2 0-217.5-70.2 55.3-70.5 162.6 0 217.5zm-142.3 76.3c0-8.7-5.7-14.4-14.7-14.7-4.6 0-9.5 1.4-12.8 6.5-2.4-4.1-6.5-6.5-12.2-6.5-3.8 0-7.6 1.4-10.6 5.4V392h-8.2v36.7h8.2c0-18.9-2.5-30.2 9-30.2 10.2 0 8.2 10.2 8.2 30.2h7.9c0-18.3-2.5-30.2 9-30.2 10.2 0 8.2 10 8.2 30.2h8.2v-23zm44.9-13.7h-7.9v4.4c-2.7-3.3-6.5-5.4-11.7-5.4-10.3 0-18.2 8.2-18.2 19.3 0 11.2 7.9 19.3 18.2 19.3 5.2 0 9-1.9 11.7-5.4v4.6h7.9V392zm40.5 25.6c0-15-22.9-8.2-22.9-15.2 0-5.7 11.9-4.8 18.5-1.1l3.3-6.5c-9.4-6.1-30.2-6-30.2 8.2 0 14.3 22.9 8.3 22.9 15 0 6.3-13.5 5.8-20.7.8l-3.5 6.3c11.2 7.6 32.6 6 32.6-7.5zm35.4 9.3l-2.2-6.8c-3.8 2.1-12.2 4.4-12.2-4.1v-16.6h13.1V392h-13.1v-11.2h-8.2V392h-7.6v7.3h7.6V416c0 17.6 17.3 14.4 22.6 10.9zm13.3-13.4h27.5c0-16.2-7.4-22.6-17.4-22.6-10.6 0-18.2 7.9-18.2 19.3 0 20.5 22.6 23.9 33.8 14.2l-3.8-6c-7.8 6.4-19.6 5.8-21.9-4.9zm59.1-21.5c-4.6-2-11.6-1.8-15.2 4.4V392h-8.2v36.7h8.2V408c0-11.6 9.5-10.1 12.8-8.4l2.4-7.6zm10.6 18.3c0-11.4 11.6-15.1 20.7-8.4l3.8-6.5c-11.6-9.1-32.7-4.1-32.7 15 0 19.8 22.4 23.8 32.7 15l-3.8-6.5c-9.2 6.5-20.7 2.6-20.7-8.6zm66.7-18.3H408v4.4c-8.3-11-29.9-4.8-29.9 13.9 0 19.2 22.4 24.7 29.9 13.9v4.6h8.2V392zm33.7 0c-2.4-1.2-11-2.9-15.2 4.4V392h-7.9v36.7h7.9V408c0-11 9-10.3 12.8-8.4l2.4-7.6zm40.3-14.9h-7.9v19.3c-8.2-10.9-29.9-5.1-29.9 13.9 0 19.4 22.5 24.6 29.9 13.9v4.6h7.9v-51.7zm7.6-75.1v4.6h.8V302h1.9v-.8h-4.6v.8h1.9zm6.6 123.8c0-.5 0-1.1-.3-1.6-.3-.3-.5-.8-.8-1.1-.3-.3-.8-.5-1.1-.8-.5 0-1.1-.3-1.6-.3-.3 0-.8.3-1.4.3-.5.3-.8.5-1.1.8-.5.3-.8.8-.8 1.1-.3.5-.3 1.1-.3 1.6 0 .3 0 .8.3 1.4 0 .3.3.8.8 1.1.3.3.5.5 1.1.8.5.3 1.1.3 1.4.3.5 0 1.1 0 1.6-.3.3-.3.8-.5 1.1-.8.3-.3.5-.8.8-1.1.3-.6.3-1.1.3-1.4zm3.2-124.7h-1.4l-1.6 3.5-1.6-3.5h-1.4v5.4h.8v-4.1l1.6 3.5h1.1l1.4-3.5v4.1h1.1v-5.4zm4.4-80.5c0-76.2-62.1-138.3-138.5-138.3-27.2 0-53.9 8.2-76.5 23.1 72.1 59.3 73.2 171.5 0 230.5 22.6 15 49.5 23.1 76.5 23.1 76.4.1 138.5-61.9 138.5-138.4z"],"cc-paypal":[576,512,[],"f1f4","M186.3 258.2c0 12.2-9.7 21.5-22 21.5-9.2 0-16-5.2-16-15 0-12.2 9.5-22 21.7-22 9.3 0 16.3 5.7 16.3 15.5zM80.5 209.7h-4.7c-1.5 0-3 1-3.2 2.7l-4.3 26.7 8.2-.3c11 0 19.5-1.5 21.5-14.2 2.3-13.4-6.2-14.9-17.5-14.9zm284 0H360c-1.8 0-3 1-3.2 2.7l-4.2 26.7 8-.3c13 0 22-3 22-18-.1-10.6-9.6-11.1-18.1-11.1zM576 80v352c0 26.5-21.5 48-48 48H48c-26.5 0-48-21.5-48-48V80c0-26.5 21.5-48 48-48h480c26.5 0 48 21.5 48 48zM128.3 215.4c0-21-16.2-28-34.7-28h-40c-2.5 0-5 2-5.2 4.7L32 294.2c-.3 2 1.2 4 3.2 4h19c2.7 0 5.2-2.9 5.5-5.7l4.5-26.6c1-7.2 13.2-4.7 18-4.7 28.6 0 46.1-17 46.1-45.8zm84.2 8.8h-19c-3.8 0-4 5.5-4.2 8.2-5.8-8.5-14.2-10-23.7-10-24.5 0-43.2 21.5-43.2 45.2 0 19.5 12.2 32.2 31.7 32.2 9 0 20.2-4.9 26.5-11.9-.5 1.5-1 4.7-1 6.2 0 2.3 1 4 3.2 4H200c2.7 0 5-2.9 5.5-5.7l10.2-64.3c.3-1.9-1.2-3.9-3.2-3.9zm40.5 97.9l63.7-92.6c.5-.5.5-1 .5-1.7 0-1.7-1.5-3.5-3.2-3.5h-19.2c-1.7 0-3.5 1-4.5 2.5l-26.5 39-11-37.5c-.8-2.2-3-4-5.5-4h-18.7c-1.7 0-3.2 1.8-3.2 3.5 0 1.2 19.5 56.8 21.2 62.1-2.7 3.8-20.5 28.6-20.5 31.6 0 1.8 1.5 3.2 3.2 3.2h19.2c1.8-.1 3.5-1.1 4.5-2.6zm159.3-106.7c0-21-16.2-28-34.7-28h-39.7c-2.7 0-5.2 2-5.5 4.7l-16.2 102c-.2 2 1.3 4 3.2 4h20.5c2 0 3.5-1.5 4-3.2l4.5-29c1-7.2 13.2-4.7 18-4.7 28.4 0 45.9-17 45.9-45.8zm84.2 8.8h-19c-3.8 0-4 5.5-4.3 8.2-5.5-8.5-14-10-23.7-10-24.5 0-43.2 21.5-43.2 45.2 0 19.5 12.2 32.2 31.7 32.2 9.3 0 20.5-4.9 26.5-11.9-.3 1.5-1 4.7-1 6.2 0 2.3 1 4 3.2 4H484c2.7 0 5-2.9 5.5-5.7l10.2-64.3c.3-1.9-1.2-3.9-3.2-3.9zm47.5-33.3c0-2-1.5-3.5-3.2-3.5h-18.5c-1.5 0-3 1.2-3.2 2.7l-16.2 104-.3.5c0 1.8 1.5 3.5 3.5 3.5h16.5c2.5 0 5-2.9 5.2-5.7L544 191.2v-.3zm-90 51.8c-12.2 0-21.7 9.7-21.7 22 0 9.7 7 15 16.2 15 12 0 21.7-9.2 21.7-21.5.1-9.8-6.9-15.5-16.2-15.5z"],"cc-stripe":[576,512,[],"f1f5","M396.9 256.5c0 19.1-8.8 33.4-21.9 33.4-8.3 0-13.3-3-16.8-6.7l-.2-52.8c3.7-4.1 8.8-7 17-7 12.9-.1 21.9 14.5 21.9 33.1zM576 80v352c0 26.5-21.5 48-48 48H48c-26.5 0-48-21.5-48-48V80c0-26.5 21.5-48 48-48h480c26.5 0 48 21.5 48 48zM122.2 281.1c0-42.3-54.3-34.7-54.3-50.7 0-5.5 4.6-7.7 12.1-7.7 10.8 0 24.5 3.3 35.3 9.1v-33.4c-11.8-4.7-23.5-6.5-35.3-6.5-28.8 0-48 15-48 40.2 0 39.3 54 32.9 54 49.9 0 6.6-5.7 8.7-13.6 8.7-11.8 0-26.9-4.9-38.9-11.3v33.9c13.2 5.7 26.6 8.1 38.8 8.1 29.6-.2 49.9-14.7 49.9-40.3zm68.9-86.9h-27v-30.8l-34.7 7.4-.2 113.9c0 21 15.8 36.5 36.9 36.5 11.6 0 20.2-2.1 24.9-4.7v-28.9c-4.5 1.8-27 8.3-27-12.6v-50.5h27v-30.3zm73.8 0c-4.7-1.7-21.3-4.8-29.6 10.5l-2.2-10.5h-30.7v124.5h35.5v-84.4c8.4-11 22.6-8.9 27.1-7.4v-32.7zm44.2 0h-35.7v124.5h35.7V194.2zm0-47.3l-35.7 7.6v28.9l35.7-7.6v-28.9zm122.7 108.8c0-41.3-23.5-63.8-48.4-63.8-13.9 0-22.9 6.6-27.8 11.1l-1.8-8.8h-31.3V360l35.5-7.5.1-40.2c5.1 3.7 12.7 9 25.1 9 25.4-.1 48.6-20.5 48.6-65.6zm112.2 1.2c0-36.4-17.6-65.1-51.3-65.1-33.8 0-54.3 28.7-54.3 64.9 0 42.8 24.2 64.5 58.8 64.5 17 0 29.7-3.9 39.4-9.2v-28.6c-9.7 4.9-20.8 7.9-34.9 7.9-13.8 0-26-4.9-27.6-21.5h69.5c.1-2 .4-9.4.4-12.9zm-51.6-36.1c-8.9 0-18.7 6.7-18.7 22.7h36.7c0-16-9.3-22.7-18-22.7z"],"cc-visa":[576,512,[],"f1f0","M470.1 231.3s7.6 37.2 9.3 45H446c3.3-8.9 16-43.5 16-43.5-.2.3 3.3-9.1 5.3-14.9l2.8 13.4zM576 80v352c0 26.5-21.5 48-48 48H48c-26.5 0-48-21.5-48-48V80c0-26.5 21.5-48 48-48h480c26.5 0 48 21.5 48 48zM152.5 331.2L215.7 176h-42.5l-39.3 106-4.3-21.5-14-71.4c-2.3-9.9-9.4-12.7-18.2-13.1H32.7l-.7 3.1c15.8 4 29.9 9.8 42.2 17.1l35.8 135h42.5zm94.4.2L272.1 176h-40.2l-25.1 155.4h40.1zm139.9-50.8c.2-17.7-10.6-31.2-33.7-42.3-14.1-7.1-22.7-11.9-22.7-19.2.2-6.6 7.3-13.4 23.1-13.4 13.1-.3 22.7 2.8 29.9 5.9l3.6 1.7 5.5-33.6c-7.9-3.1-20.5-6.6-36-6.6-39.7 0-67.6 21.2-67.8 51.4-.3 22.3 20 34.7 35.2 42.2 15.5 7.6 20.8 12.6 20.8 19.3-.2 10.4-12.6 15.2-24.1 15.2-16 0-24.6-2.5-37.7-8.3l-5.3-2.5-5.6 34.9c9.4 4.3 26.8 8.1 44.8 8.3 42.2.1 69.7-20.8 70-53zM528 331.4L495.6 176h-31.1c-9.6 0-16.9 2.8-21 12.9l-59.7 142.5H426s6.9-19.2 8.4-23.3H486c1.2 5.5 4.8 23.3 4.8 23.3H528z"],centercode:[512,512,[],"f380","M329.2 268.6c-3.8 35.2-35.4 60.6-70.6 56.8-35.2-3.8-60.6-35.4-56.8-70.6 3.8-35.2 35.4-60.6 70.6-56.8 35.1 3.8 60.6 35.4 56.8 70.6zm-85.8 235.1C96.7 496-8.2 365.5 10.1 224.3c11.2-86.6 65.8-156.9 139.1-192 161-77.1 349.7 37.4 354.7 216.6 4.1 147-118.4 262.2-260.5 254.8zm179.9-180c27.9-118-160.5-205.9-237.2-234.2-57.5 56.3-69.1 188.6-33.8 344.4 68.8 15.8 169.1-26.4 271-110.2z"],chrome:[496,512,[],"f268","M131.5 217.5L55.1 100.1c47.6-59.2 119-91.8 192-92.1 42.3-.3 85.5 10.5 124.8 33.2 43.4 25.2 76.4 61.4 97.4 103L264 133.4c-58.1-3.4-113.4 29.3-132.5 84.1zm32.9 38.5c0 46.2 37.4 83.6 83.6 83.6s83.6-37.4 83.6-83.6-37.4-83.6-83.6-83.6-83.6 37.3-83.6 83.6zm314.9-89.2L339.6 174c37.9 44.3 38.5 108.2 6.6 157.2L234.1 503.6c46.5 2.5 94.4-7.7 137.8-32.9 107.4-62 150.9-192 107.4-303.9zM133.7 303.6L40.4 120.1C14.9 159.1 0 205.9 0 256c0 124 90.8 226.7 209.5 244.9l63.7-124.8c-57.6 10.8-113.2-20.8-139.5-72.5z"],cloudscale:[448,512,[],"f383","M318.1 154l-9.4 7.6c-22.5-19.3-51.5-33.6-83.3-33.6C153.8 128 96 188.8 96 260.3c0 6.6.4 13.1 1.4 19.4-2-56 41.8-97.4 92.6-97.4 24.2 0 46.2 9.4 62.6 24.7l-25.2 20.4c-8.3-.9-16.8 1.8-23.1 8.1-11.1 11-11.1 28.9 0 40 11.1 11 28.9 11 40 0 6.3-6.3 9-14.9 8.1-23.1l75.2-88.8c6.3-6.5-3.3-15.9-9.5-9.6zm-83.8 111.5c-5.6 5.5-14.6 5.5-20.2 0-5.6-5.6-5.6-14.6 0-20.2s14.6-5.6 20.2 0 5.6 14.7 0 20.2zM224 32C100.5 32 0 132.5 0 256s100.5 224 224 224 224-100.5 224-224S347.5 32 224 32zm0 384c-88.2 0-160-71.8-160-160S135.8 96 224 96s160 71.8 160 160-71.8 160-160 160z"],cloudsmith:[332,512,[],"f384","M332.5 419.9c0 46.4-37.6 84.1-84 84.1s-84-37.7-84-84.1 37.6-84 84-84 84 37.6 84 84zm-84-243.9c46.4 0 80-37.6 80-84s-33.6-84-80-84-88 37.6-88 84-29.6 76-76 76-84 41.6-84 88 37.6 80 84 80 84-33.6 84-80 33.6-80 80-80z"],cloudversify:[616,512,[],"f385","M148.6 304c8.2 68.5 67.4 115.5 146 111.3 51.2 43.3 136.8 45.8 186.4-5.6 69.2 1.1 118.5-44.6 131.5-99.5 14.8-62.5-18.2-132.5-92.1-155.1-33-88.1-131.4-101.5-186.5-85-57.3 17.3-84.3 53.2-99.3 109.7-7.8 2.7-26.5 8.9-45 24.1 11.7 0 15.2 8.9 15.2 19.5v20.4c0 10.7-8.7 19.5-19.5 19.5h-20.2c-10.7 0-19.5-6-19.5-16.7V240H98.8C95 240 88 244.3 88 251.9v40.4c0 6.4 5.3 11.8 11.7 11.8h48.9zm227.4 8c-10.7 46.3 21.7 72.4 55.3 86.8C324.1 432.6 259.7 348 296 288c-33.2 21.6-33.7 71.2-29.2 92.9-17.9-12.4-53.8-32.4-57.4-79.8-3-39.9 21.5-75.7 57-93.9C297 191.4 369.9 198.7 400 248c-14.1-48-53.8-70.1-101.8-74.8 30.9-30.7 64.4-50.3 114.2-43.7 69.8 9.3 133.2 82.8 67.7 150.5 35-16.3 48.7-54.4 47.5-76.9l10.5 19.6c11.8 22 15.2 47.6 9.4 72-9.2 39-40.6 68.8-79.7 76.5-32.1 6.3-83.1-5.1-91.8-59.2zM128 208H88.2c-8.9 0-16.2-7.3-16.2-16.2v-39.6c0-8.9 7.3-16.2 16.2-16.2H128c8.9 0 16.2 7.3 16.2 16.2v39.6c0 8.9-7.3 16.2-16.2 16.2zM10.1 168C4.5 168 0 163.5 0 157.9v-27.8c0-5.6 4.5-10.1 10.1-10.1h27.7c5.5 0 10.1 4.5 10.1 10.1v27.8c0 5.6-4.5 10.1-10.1 10.1H10.1zM168 142.7v-21.4c0-5.1 4.2-9.3 9.3-9.3h21.4c5.1 0 9.3 4.2 9.3 9.3v21.4c0 5.1-4.2 9.3-9.3 9.3h-21.4c-5.1 0-9.3-4.2-9.3-9.3zM56 235.5v25c0 6.3-5.1 11.5-11.4 11.5H19.4C13.1 272 8 266.8 8 260.5v-25c0-6.3 5.1-11.5 11.4-11.5h25.1c6.4 0 11.5 5.2 11.5 11.5z"],codepen:[512,512,[],"f1cb","M502.285 159.704l-234-156c-7.987-4.915-16.511-4.96-24.571 0l-234 156C3.714 163.703 0 170.847 0 177.989v155.999c0 7.143 3.714 14.286 9.715 18.286l234 156.022c7.987 4.915 16.511 4.96 24.571 0l234-156.022c6-3.999 9.715-11.143 9.715-18.286V177.989c-.001-7.142-3.715-14.286-9.716-18.285zM278 63.131l172.286 114.858-76.857 51.429L278 165.703V63.131zm-44 0v102.572l-95.429 63.715-76.857-51.429L234 63.131zM44 219.132l55.143 36.857L44 292.846v-73.714zm190 229.715L61.714 333.989l76.857-51.429L234 346.275v102.572zm22-140.858l-77.715-52 77.715-52 77.715 52-77.715 52zm22 140.858V346.275l95.429-63.715 76.857 51.429L278 448.847zm190-156.001l-55.143-36.857L468 219.132v73.714z"],codiepie:[472,512,[],"f284","M422.5 202.9c30.7 0 33.5 53.1-.3 53.1h-10.8v44.3h-26.6v-97.4h37.7zM472 352.6C429.9 444.5 350.4 504 248 504 111 504 0 393 0 256S111 8 248 8c97.4 0 172.8 53.7 218.2 138.4l-186 108.8L472 352.6zm-38.5 12.5l-60.3-30.7c-27.1 44.3-70.4 71.4-122.4 71.4-82.5 0-149.2-66.7-149.2-148.9 0-82.5 66.7-149.2 149.2-149.2 48.4 0 88.9 23.5 116.9 63.4l59.5-34.6c-40.7-62.6-104.7-100-179.2-100-121.2 0-219.5 98.3-219.5 219.5S126.8 475.5 248 475.5c78.6 0 146.5-42.1 185.5-110.4z"],connectdevelop:[576,512,[],"f20e","M550.5 241l-50.089-86.786c1.071-2.142 1.875-4.553 1.875-7.232 0-8.036-6.696-14.733-14.732-15.001l-55.447-95.893c.536-1.607 1.071-3.214 1.071-4.821 0-8.571-6.964-15.268-15.268-15.268-4.821 0-8.839 2.143-11.786 5.625H299.518C296.839 18.143 292.821 16 288 16s-8.839 2.143-11.518 5.625H170.411C167.464 18.143 163.447 16 158.625 16c-8.303 0-15.268 6.696-15.268 15.268 0 1.607.536 3.482 1.072 4.821l-55.983 97.233c-5.356 2.41-9.107 7.5-9.107 13.661 0 .535.268 1.071.268 1.607l-53.304 92.143c-7.232 1.339-12.59 7.5-12.59 15 0 7.232 5.089 13.393 12.054 15l55.179 95.358c-.536 1.607-.804 2.946-.804 4.821 0 7.232 5.089 13.393 12.054 14.732l51.697 89.732c-.536 1.607-1.071 3.482-1.071 5.357 0 8.571 6.964 15.268 15.268 15.268 4.821 0 8.839-2.143 11.518-5.357h106.875C279.161 493.857 283.447 496 288 496s8.839-2.143 11.518-5.357h107.143c2.678 2.946 6.696 4.821 10.982 4.821 8.571 0 15.268-6.964 15.268-15.268 0-1.607-.267-2.946-.803-4.285l51.697-90.268c6.964-1.339 12.054-7.5 12.054-14.732 0-1.607-.268-3.214-.804-4.821l54.911-95.358c6.964-1.339 12.322-7.5 12.322-15-.002-7.232-5.092-13.393-11.788-14.732zM153.535 450.732l-43.66-75.803h43.66v75.803zm0-83.839h-43.66c-.268-1.071-.804-2.142-1.339-3.214l44.999-47.41v50.624zm0-62.411l-50.357 53.304c-1.339-.536-2.679-1.34-4.018-1.607L43.447 259.75c.535-1.339.535-2.679.535-4.018s0-2.41-.268-3.482l51.965-90c2.679-.268 5.357-1.072 7.768-2.679l50.089 51.965v92.946zm0-102.322l-45.803-47.41c1.339-2.143 2.143-4.821 2.143-7.767 0-.268-.268-.804-.268-1.072l43.928-15.804v72.053zm0-80.625l-43.66 15.804 43.66-75.536v59.732zm326.519 39.108l.804 1.339L445.5 329.125l-63.75-67.232 98.036-101.518.268.268zM291.75 355.107l11.518 11.786H280.5l11.25-11.786zm-.268-11.25l-83.303-85.446 79.553-84.375 83.036 87.589-79.286 82.232zm5.357 5.893l79.286-82.232 67.5 71.25-5.892 28.125H313.714l-16.875-17.143zM410.411 44.393c1.071.536 2.142 1.072 3.482 1.34l57.857 100.714v.536c0 2.946.803 5.624 2.143 7.767L376.393 256l-83.035-87.589L410.411 44.393zm-9.107-2.143L287.732 162.518l-57.054-60.268 166.339-60h4.287zm-123.483 0c2.678 2.678 6.16 4.285 10.179 4.285s7.5-1.607 10.179-4.285h75L224.786 95.821 173.893 42.25h103.928zm-116.249 5.625l1.071-2.142a33.834 33.834 0 0 0 2.679-.804l51.161 53.84-54.911 19.821V47.875zm0 79.286l60.803-21.964 59.732 63.214-79.553 84.107-40.982-42.053v-83.304zm0 92.678L198 257.607l-36.428 38.304v-76.072zm0 87.858l42.053-44.464 82.768 85.982-17.143 17.678H161.572v-59.196zm6.964 162.053c-1.607-1.607-3.482-2.678-5.893-3.482l-1.071-1.607v-89.732h99.91l-91.607 94.821h-1.339zm129.911 0c-2.679-2.41-6.428-4.285-10.447-4.285s-7.767 1.875-10.447 4.285h-96.429l91.607-94.821h38.304l91.607 94.821H298.447zm120-11.786l-4.286 7.5c-1.339.268-2.41.803-3.482 1.339l-89.196-91.875h114.376l-17.412 83.036zm12.856-22.232l12.858-60.803h21.964l-34.822 60.803zm34.822-68.839h-20.357l4.553-21.16 17.143 18.214c-.535.803-1.071 1.874-1.339 2.946zm66.161-107.411l-55.447 96.697c-1.339.535-2.679 1.071-4.018 1.874l-20.625-21.964 34.554-163.928 45.803 79.286c-.267 1.339-.803 2.678-.803 4.285 0 1.339.268 2.411.536 3.75z"],contao:[512,512,[],"f26d","M45.4 305c14.4 67.1 26.4 129 68.2 175H34c-18.7 0-34-15.2-34-34V66c0-18.7 15.2-34 34-34h57.7C77.9 44.6 65.6 59.2 54.8 75.6c-45.4 70-27 146.8-9.4 229.4zM478 32h-90.2c21.4 21.4 39.2 49.5 52.7 84.1l-137.1 29.3c-14.9-29-37.8-53.3-82.6-43.9-24.6 5.3-41 19.3-48.3 34.6-8.8 18.7-13.2 39.8 8.2 140.3 21.1 100.2 33.7 117.7 49.5 131.2 12.9 11.1 33.4 17 58.3 11.7 44.5-9.4 55.7-40.7 57.4-73.2l137.4-29.6c3.2 71.5-18.7 125.2-57.4 163.6H478c18.7 0 34-15.2 34-34V66c0-18.8-15.2-34-34-34z"],cpanel:[640,512,[],"f388","M52.9 213.7h40l-6.2 23.6c-1.9 6.5-7.4 10.9-14.3 10.9H53.8c-24.9 0-24.7 37.4 0 37.4h11.3c4.2 0 7.6 3.9 6.4 8.3L64.4 320H52c-33.5 0-59-31.4-50.3-65.2 7.3-27 28.3-41.1 51.2-41.1M73.1 320L108 189.9c1.8-6.4 7.2-10.9 14.3-10.9h37c24.1 0 45.4 16.4 51 41.2 6.6 29.1-14.5 65.3-51.7 65.3h-32l6.4-23.8c1.8-6.2 7.3-10.8 14.3-10.8h10.3c12.4 0 20.8-11.7 18.3-22.6-2.1-9.2-9.9-14.8-18.3-14.8h-19.8L112 309.2c-1.9 6.2-7.4 10.7-14.2 10.7l-24.7.1m220.6-69.4c.3-1 1.9-5.3-2.1-5.3h-57.5c-9.7 0-16.6-8.9-14.2-18.5l3.5-13.4h77.9c18.8 0 33.3 17.6 28.5 36.8l-14 51.8c-2.8 10.6-12.2 17.8-23.4 17.8l-57.5-.2c-42.9 0-38.5-63.8.7-63.8H284l-3.5 13.2c-1.9 6.2-7.4 10.8-14.2 10.8h-21.6c-5.3 0-5.3 7.9 0 7.9h34.9c4.6 0 5.1-3.9 5.5-5.3l8.6-31.8m103.1-36.9c34.4 0 59.3 32.3 50.3 65.4l-8.8 33.1c-1.2 4.9-5.7 7.8-10.3 7.8h-19.1c-4.5 0-7.6-4-6.4-8.3l10.6-40c3.3-11.6-5.6-23.4-18.1-23.4h-19.8l-17.2 64c-1.2 4.8-5.6 7.8-10.4 7.8h-18.9c-4.2 0-7.6-3.9-6.4-8.3l26.2-98h48.3M498 251.6l-8 30c-.9 3.3 1.5 6.7 5.1 6.7h73.3l-5.7 21c-1.9 6.2-7.4 10.7-14.2 10.7h-66.7c-20 0-33.3-19-28.3-36.7l10.8-40c4.8-17.6 20.7-29.6 38.6-29.6h47.3c19 0 33.2 17.7 28.3 36.8l-3.2 12c-2.9 11-12.7 17.6-23.2 17.6h-53.4l3.5-13c1.6-6.2 7.2-10.8 14.2-10.8H538c2 0 3.3-1 3.9-3l.7-2.6c.7-2.7-1.3-5.1-3.9-5.1h-32.9c-4.1 0-6.9 2.1-7.8 6zm70.2 68.4l35.6-133.1c1.2-4.7 5.5-7.9 10.4-7.9h18.9c4.5 0 7.7 4 6.5 8.3l-26.5 98.2c-5.1 20.7-24.2 34.5-44.9 34.5"],"creative-commons":[496,512,[],"f25e","M254.8 214.8l-33.2 17.3c-9.4-19.6-25.2-19.9-27.5-19.9-22.1 0-33.2 14.6-33.2 43.8 0 23.6 9.2 43.8 33.2 43.8 14.5 0 24.7-7.1 30.6-21.3l30.6 15.5c-6.2 11.5-25.7 39-65.1 39-22.6 0-74-10.3-74-77.1 0-58.7 43-77.1 72.6-77.1 30.8.2 52.7 12.1 66 36zm143.1 0l-32.8 17.3c-9.5-19.8-25.7-19.9-27.9-19.9-22.1 0-33.2 14.6-33.2 43.8 0 23.6 9.2 43.8 33.2 43.8 14.5 0 24.6-7.1 30.5-21.3l31 15.5c-2.1 3.7-21.4 39-65.1 39-22.7 0-74-9.9-74-77.1 0-58.7 43-77.1 72.6-77.1 30.8.2 52.7 12.1 65.7 36zM247.6 8C389.4 8 496 118.1 496 256c0 147.1-118.5 248-248.4 248C113.6 504 0 394.5 0 256 0 123.1 104.7 8 247.6 8zm.8 44.7C130.2 52.7 44.7 150.6 44.7 256c0 109.8 91.2 202.8 203.7 202.8 103.2 0 202.8-81.1 202.8-202.8.1-113.8-90.2-203.3-202.8-203.3z"],"creative-commons-by":[496,512,[],"f4e7","M314.9 194.4v101.4h-28.3v120.5h-77.1V295.9h-28.3V194.4c0-4.4 1.6-8.2 4.6-11.3 3.1-3.1 6.9-4.7 11.3-4.7H299c4.1 0 7.8 1.6 11.1 4.7 3.1 3.2 4.8 6.9 4.8 11.3zm-101.5-63.7c0-23.3 11.5-35 34.5-35s34.5 11.7 34.5 35c0 23-11.5 34.5-34.5 34.5s-34.5-11.5-34.5-34.5zM247.6 8C389.4 8 496 118.1 496 256c0 147.1-118.5 248-248.4 248C113.6 504 0 394.5 0 256 0 123.1 104.7 8 247.6 8zm.8 44.7C130.2 52.7 44.7 150.6 44.7 256c0 109.8 91.2 202.8 203.7 202.8 103.2 0 202.8-81.1 202.8-202.8.1-113.8-90.2-203.3-202.8-203.3z"],"creative-commons-nc":[496,512,[],"f4e8","M247.6 8C387.4 8 496 115.9 496 256c0 147.2-118.5 248-248.4 248C113.1 504 0 393.2 0 256 0 123.1 104.7 8 247.6 8zM55.8 189.1c-7.4 20.4-11.1 42.7-11.1 66.9 0 110.9 92.1 202.4 203.7 202.4 122.4 0 177.2-101.8 178.5-104.1l-93.4-41.6c-7.7 37.1-41.2 53-68.2 55.4v38.1h-28.8V368c-27.5-.3-52.6-10.2-75.3-29.7l34.1-34.5c31.7 29.4 86.4 31.8 86.4-2.2 0-6.2-2.2-11.2-6.6-15.1-14.2-6-1.8-.1-219.3-97.4zM248.4 52.3c-38.4 0-112.4 8.7-170.5 93l94.8 42.5c10-31.3 40.4-42.9 63.8-44.3v-38.1h28.8v38.1c22.7 1.2 43.4 8.9 62 23L295 199.7c-42.7-29.9-83.5-8-70 11.1 53.4 24.1 43.8 19.8 93 41.6l127.1 56.7c4.1-17.4 6.2-35.1 6.2-53.1 0-57-19.8-105-59.3-143.9-39.3-39.9-87.2-59.8-143.6-59.8z"],"creative-commons-nc-eu":[496,512,[],"f4e9","M247.7 8C103.6 8 0 124.8 0 256c0 136.3 111.7 248 247.7 248C377.9 504 496 403.1 496 256 496 117 388.4 8 247.7 8zm.6 450.7c-112 0-203.6-92.5-203.6-202.7 0-23.2 3.7-45.2 10.9-66l65.7 29.1h-4.7v29.5h23.3c0 6.2-.4 3.2-.4 19.5h-22.8v29.5h27c11.4 67 67.2 101.3 124.6 101.3 26.6 0 50.6-7.9 64.8-15.8l-10-46.1c-8.7 4.6-28.2 10.8-47.3 10.8-28.2 0-58.1-10.9-67.3-50.2h90.3l128.3 56.8c-1.5 2.1-56.2 104.3-178.8 104.3zm-16.7-190.6l-.5-.4.9.4h-.4zm77.2-19.5h3.7v-29.5h-70.3l-28.6-12.6c2.5-5.5 5.4-10.5 8.8-14.3 12.9-15.8 31.1-22.4 51.1-22.4 18.3 0 35.3 5.4 46.1 10l11.6-47.3c-15-6.6-37-12.4-62.3-12.4-39 0-72.2 15.8-95.9 42.3-5.3 6.1-9.8 12.9-13.9 20.1l-81.6-36.1c64.6-96.8 157.7-93.6 170.7-93.6 113 0 203 90.2 203 203.4 0 18.7-2.1 36.3-6.3 52.9l-136.1-60.5z"],"creative-commons-nc-jp":[496,512,[],"f4ea","M247.7 8C103.6 8 0 124.8 0 256c0 136.4 111.8 248 247.7 248C377.9 504 496 403.2 496 256 496 117.2 388.5 8 247.7 8zm.6 450.7c-112 0-203.6-92.5-203.6-202.7 0-21.1 3-41.2 9-60.3l127 56.5h-27.9v38.6h58.1l5.7 11.8v18.7h-63.8V360h63.8v56h61.7v-56h64.2v-35.7l81 36.1c-1.5 2.2-57.1 98.3-175.2 98.3zm87.6-137.3h-57.6v-18.7l2.9-5.6 54.7 24.3zm6.5-51.4v-17.8h-38.6l63-116H301l-43.4 96-23-10.2-39.6-85.7h-65.8l27.3 51-81.9-36.5c27.8-44.1 82.6-98.1 173.7-98.1 112.8 0 203 90 203 203.4 0 21-2.7 40.6-7.9 59l-101-45.1z"],"creative-commons-nd":[496,512,[],"f4eb","M247.6 8C389.4 8 496 118.1 496 256c0 147.1-118.5 248-248.4 248C113.6 504 0 394.5 0 256 0 123.1 104.7 8 247.6 8zm.8 44.7C130.2 52.7 44.7 150.6 44.7 256c0 109.8 91.2 202.8 203.7 202.8 103.2 0 202.8-81.1 202.8-202.8.1-113.8-90.2-203.3-202.8-203.3zm94 144.3v42.5H162.1V197h180.3zm0 79.8v42.5H162.1v-42.5h180.3z"],"creative-commons-pd":[496,512,[],"f4ec","M248 8C111 8 0 119.1 0 256c0 137 111 248 248 248s248-111 248-248C496 119.1 385 8 248 8zm0 449.5c-139.2 0-235.8-138-190.2-267.9l78.8 35.1c-2.1 10.5-3.3 21.5-3.3 32.9 0 99 73.9 126.9 120.4 126.9 22.9 0 53.5-6.7 79.4-29.5L297 311.1c-5.5 6.3-17.6 16.7-36.3 16.7-37.8 0-53.7-39.9-53.9-71.9 230.4 102.6 216.5 96.5 217.9 96.8-34.3 62.4-100.6 104.8-176.7 104.8zm194.2-150l-224-100c18.8-34 54.9-30.7 74.7-11l40.4-41.6c-27.1-23.3-58-27.5-78.1-27.5-47.4 0-80.9 20.5-100.7 51.6l-74.9-33.4c36.1-54.9 98.1-91.2 168.5-91.2 111.1 0 201.5 90.4 201.5 201.5 0 18-2.4 35.4-6.8 52-.3-.1-.4-.2-.6-.4z"],"creative-commons-pd-alt":[496,512,[],"f4ed","M247.6 8C389.4 8 496 118.1 496 256c0 147.1-118.5 248-248.4 248C113.6 504 0 394.5 0 256 0 123.1 104.7 8 247.6 8zm.8 44.7C130.2 52.7 44.7 150.6 44.7 256c0 109.8 91.2 202.8 203.7 202.8 103.2 0 202.8-81.1 202.8-202.8.1-113.8-90.2-203.3-202.8-203.3zM135.5 323.3V186h68.2c33.7 0 50.5 15.5 50.5 46.5 0 9-3 46.5-57.1 46.5h-27v44.3h-34.6zm34.1-111.6v41.6h29.2c27.9 0 30-41.6-.9-41.6h-28.3zm93.9 111.6V186h53.2c21.4 0 70 5.2 70 68.6 0 63.5-48.6 68.6-70 68.6h-53.2zm34.1-108.5v79.7h19.9c24 0 34.5-15.3 34.5-39.9 0-42-31.2-39.9-35-39.9l-19.4.1z"],"creative-commons-remix":[496,512,[],"f4ee","M247.6 8C389.4 8 496 118.1 496 256c0 147.1-118.5 248-248.4 248C113.6 504 0 394.5 0 256 0 123.1 104.7 8 247.6 8zm.8 44.7C130.2 52.7 44.7 150.6 44.7 256c0 109.8 91.2 202.8 203.7 202.8 103.2 0 202.8-81.1 202.8-202.8.1-113.8-90.2-203.3-202.8-203.3zm161.7 207.7l4.9 2.2v70c-7.2 3.6-63.4 27.5-67.3 28.8-6.5-1.8-113.7-46.8-137.3-56.2l-64.2 26.6-63.3-27.5v-63.8l59.3-24.8c-.7-.7-.4 5-.4-70.4l67.3-29.7L361 178.5v61.6l49.1 20.3zm-70.4 81.5v-43.8h-.4v-1.8l-113.8-46.5V295l113.8 46.9v-.4l.4.4zm7.5-57.6l39.9-16.4-36.8-15.5-39 16.4 35.9 15.5zm52.3 38.1v-43L355.2 298v43.4l44.3-19z"],"creative-commons-sa":[496,512,[],"f4ef","M247.6 8C389.4 8 496 118.1 496 256c0 147.1-118.5 248-248.4 248C113.6 504 0 394.5 0 256 0 123.1 104.7 8 247.6 8zm.8 44.7C130.2 52.7 44.7 150.6 44.7 256c0 109.8 91.2 202.8 203.7 202.8 103.2 0 202.8-81.1 202.8-202.8.1-113.8-90.2-203.3-202.8-203.3zM137.7 221c13-83.9 80.5-95.7 108.9-95.7 99.8 0 127.5 82.5 127.5 134.2 0 63.6-41 132.9-128.9 132.9-38.9 0-99.1-20-109.4-97h62.5c1.5 30.1 19.6 45.2 54.5 45.2 23.3 0 58-18.2 58-82.8 0-82.5-49.1-80.6-56.7-80.6-33.1 0-51.7 14.6-55.8 43.8h18.2l-49.2 49.2-49-49.2h19.4z"],"creative-commons-sampling":[496,512,[],"f4f0","M247.6 8C389.4 8 496 118.1 496 256c0 147.1-118.5 248-248.4 248C113.6 504 0 394.5 0 256 0 123.1 104.7 8 247.6 8zm.8 44.7C130.2 52.7 44.7 150.6 44.7 256c0 109.8 91.2 202.8 203.7 202.8 103.2 0 202.8-81.1 202.8-202.8.1-113.8-90.2-203.3-202.8-203.3zm3.6 53.2c2.8-.3 11.5 1 11.5 11.5l6.6 107.2 4.9-59.3c0-6 4.7-10.6 10.6-10.6 5.9 0 10.6 4.7 10.6 10.6 0 2.5-.5-5.7 5.7 81.5l5.8-64.2c.3-2.9 2.9-9.3 10.2-9.3 3.8 0 9.9 2.3 10.6 8.9l11.5 96.5 5.3-12.8c1.8-4.4 5.2-6.6 10.2-6.6h58v21.3h-50.9l-18.2 44.3c-3.9 9.9-19.5 9.1-20.8-3.1l-4-31.9-7.5 92.6c-.3 3-3 9.3-10.2 9.3-3 0-9.8-2.1-10.6-9.3 0-1.9.6 5.8-6.2-77.9l-5.3 72.2c-1.1 4.8-4.8 9.3-10.6 9.3-2.9 0-9.8-2-10.6-9.3 0-1.9.5 6.7-5.8-87.7l-5.8 94.8c0 6.3-3.6 12.4-10.6 12.4-5.2 0-10.6-4.1-10.6-12l-5.8-87.7c-5.8 92.5-5.3 84-5.3 85.9-1.1 4.8-4.8 9.3-10.6 9.3-3 0-9.8-2.1-10.6-9.3 0-.7-.4-1.1-.4-2.6l-6.2-88.6L182 348c-.7 6.5-6.7 9.3-10.6 9.3-5.8 0-9.6-4.1-10.6-8.9L149.7 272c-2 4-3.5 8.4-11.1 8.4H87.2v-21.3H132l13.7-27.9c4.4-9.9 18.2-7.2 19.9 2.7l3.1 20.4 8.4-97.9c0-6 4.8-10.6 10.6-10.6.5 0 10.6-.2 10.6 12.4l4.9 69.1 6.6-92.6c0-10.1 9.5-10.6 10.2-10.6.6 0 10.6.7 10.6 10.6l5.3 80.6 6.2-97.9c.1-1.1-.6-10.3 9.9-11.5z"],"creative-commons-sampling-plus":[496,512,[],"f4f1","M247.6 8C389.4 8 496 118.1 496 256c0 147.1-118.5 248-248.4 248C113.6 504 0 394.5 0 256 0 123.1 104.7 8 247.6 8zm.8 44.7C130.2 52.7 44.7 150.6 44.7 256c0 109.8 91.2 202.8 203.7 202.8 103.2 0 202.8-81.1 202.8-202.8.1-113.8-90.2-203.3-202.8-203.3zm107 205.6c-4.7 0-9 2.8-10.7 7.2l-4 9.5-11-92.8c-1.7-13.9-22-13.4-23.1.4l-4.3 51.4-5.2-68.8c-1.1-14.3-22.1-14.2-23.2 0l-3.5 44.9-5.9-94.3c-.9-14.5-22.3-14.4-23.2 0l-5.1 83.7-4.3-66.3c-.9-14.4-22.2-14.4-23.2 0l-5.3 80.2-4.1-57c-1.1-14.3-22-14.3-23.2-.2l-7.7 89.8-1.8-12.2c-1.7-11.4-17.1-13.6-22-3.3l-13.2 27.7H87.5v23.2h51.3c4.4 0 8.4-2.5 10.4-6.4l10.7 73.1c2 13.5 21.9 13 23.1-.7l3.8-43.6 5.7 78.3c1.1 14.4 22.3 14.2 23.2-.1l4.6-70.4 4.8 73.3c.9 14.4 22.3 14.4 23.2-.1l4.9-80.5 4.5 71.8c.9 14.3 22.1 14.5 23.2.2l4.6-58.6 4.9 64.4c1.1 14.3 22 14.2 23.1.1l6.8-83 2.7 22.3c1.4 11.8 17.7 14.1 22.3 3.1l18-43.4h50.5V258l-58.4.3zm-78 5.2h-21.9v21.9c0 4.1-3.3 7.5-7.5 7.5-4.1 0-7.5-3.3-7.5-7.5v-21.9h-21.9c-4.1 0-7.5-3.3-7.5-7.5 0-4.1 3.4-7.5 7.5-7.5h21.9v-21.9c0-4.1 3.4-7.5 7.5-7.5s7.5 3.3 7.5 7.5v21.9h21.9c4.1 0 7.5 3.3 7.5 7.5 0 4.1-3.4 7.5-7.5 7.5z"],"creative-commons-share":[496,512,[],"f4f2","M247.6 8C389.4 8 496 118.1 496 256c0 147.1-118.5 248-248.4 248C113.6 504 0 394.5 0 256 0 123.1 104.7 8 247.6 8zm.8 44.7C130.2 52.7 44.7 150.6 44.7 256c0 109.8 91.2 202.8 203.7 202.8 103.2 0 202.8-81.1 202.8-202.8.1-113.8-90.2-203.3-202.8-203.3zm101 132.4c7.8 0 13.7 6.1 13.7 13.7v182.5c0 7.7-6.1 13.7-13.7 13.7H214.3c-7.7 0-13.7-6-13.7-13.7v-54h-54c-7.8 0-13.7-6-13.7-13.7V131.1c0-8.2 6.6-12.7 12.4-13.7h136.4c7.7 0 13.7 6 13.7 13.7v54h54zM159.9 300.3h40.7V198.9c0-7.4 5.8-12.6 12-13.7h55.8v-40.3H159.9v155.4zm176.2-88.1H227.6v155.4h108.5V212.2z"],css3:[512,512,[],"f13c","M480 32l-64 368-223.3 80L0 400l19.6-94.8h82l-8 40.6L210 390.2l134.1-44.4 18.8-97.1H29.5l16-82h333.7l10.5-52.7H56.3l16.3-82H480z"],"css3-alt":[384,512,[],"f38b","M0 32l34.9 395.8L192 480l157.1-52.2L384 32H0zm313.1 80l-4.8 47.3L193 208.6l-.3.1h111.5l-12.8 146.6-98.2 28.7-98.8-29.2-6.4-73.9h48.9l3.2 38.3 52.6 13.3 54.7-15.4 3.7-61.6-166.3-.5v-.1l-.2.1-3.6-46.3L193.1 162l6.5-2.7H76.7L70.9 112h242.2z"],cuttlefish:[440,512,[],"f38c","M344 305.5c-17.5 31.6-57.4 54.5-96 54.5-56.6 0-104-47.4-104-104s47.4-104 104-104c38.6 0 78.5 22.9 96 54.5 13.7-50.9 41.7-93.3 87-117.8C385.7 39.1 320.5 8 248 8 111 8 0 119 0 256s111 248 248 248c72.5 0 137.7-31.1 183-80.7-45.3-24.5-73.3-66.9-87-117.8z"],"d-and-d":[576,512,[],"f38d","M82.5 98.9c-.6-17.2 2-33.8 12.7-48.2.3 7.4 1.2 14.5 4.2 21.6 5.9-27.5 19.7-49.3 42.3-65.5-1.9 5.9-3.5 11.8-3 17.7 8.7-7.4 18.8-17.8 44.4-22.7 14.7-2.8 29.7-2 42.1 1 38.5 9.3 61 34.3 69.7 72.3 5.3 23.1.7 45-8.3 66.4-5.2 12.4-12 24.4-20.7 35.1-2-1.9-3.9-3.8-5.8-5.6-42.8-40.8-26.8-25.2-37.4-37.4-1.1-1.2-1-2.2-.1-3.6 8.3-13.5 11.8-28.2 10-44-1.1-9.8-4.3-18.9-11.3-26.2-14.5-15.3-39.2-15-53.5.6-11.4 12.5-14.1 27.4-10.9 43.6.2 1.3.4 2.7 0 3.9-3.4 13.7-4.6 27.6-2.5 41.6.1.5.1 1.1.1 1.6 0 .3-.1.5-.2 1.1-21.8-11-36-28.3-43.2-52.2-8.3 17.8-11.1 35.5-6.6 54.1-15.6-15.2-21.3-34.3-22-55.2zm469.6 123.2c-11.6-11.6-25-20.4-40.1-26.6-12.8-5.2-26-7.9-39.9-7.1-10 .6-19.6 3.1-29 6.4-2.5.9-5.1 1.6-7.7 2.2-4.9 1.2-7.3-3.1-4.7-6.8 3.2-4.6 3.4-4.2 15-12 .6-.4 1.2-.8 2.2-1.5h-2.5c-.6 0-1.2.2-1.9.3-19.3 3.3-30.7 15.5-48.9 29.6-10.4 8.1-13.8 3.8-12-.5 1.4-3.5 3.3-6.7 5.1-10 1-1.8 2.3-3.4 3.5-5.1-.2-.2-.5-.3-.7-.5-27 18.3-46.7 42.4-57.7 73.3.3.3.7.6 1 .9.3-.6.5-1.2.9-1.7 10.4-12.1 22.8-21.8 36.6-29.8 18.2-10.6 37.5-18.3 58.7-20.2 4.3-.4 8.7-.1 13.1-.1-1.8.7-3.5.9-5.3 1.1-18.5 2.4-35.5 9-51.5 18.5-30.2 17.9-54.5 42.2-75.1 70.4-.3.4-.4.9-.7 1.3 14.5 5.3 24 17.3 36.1 25.6.2-.1.3-.2.4-.4l1.2-2.7c12.2-26.9 27-52.3 46.7-74.5 16.7-18.8 38-25.3 62.5-20 5.9 1.3 11.4 4.4 17.2 6.8 2.3-1.4 5.1-3.2 8-4.7 8.4-4.3 17.4-7 26.7-9 14.7-3.1 29.5-4.9 44.5-1.3v-.5c-.5-.4-1.2-.8-1.7-1.4zM316.7 397.6c-39.4-33-22.8-19.5-42.7-35.6-.8.9 0-.2-1.9 3-11.2 19.1-25.5 35.3-44 47.6-10.3 6.8-21.5 11.8-34.1 11.8-21.6 0-38.2-9.5-49.4-27.8-12-19.5-13.3-40.7-8.2-62.6 7.8-33.8 30.1-55.2 38.6-64.3-18.7-6.2-33 1.7-46.4 13.9.8-13.9 4.3-26.2 11.8-37.3-24.3 10.6-45.9 25-64.8 43.9-.3-5.8 5.4-43.7 5.6-44.7.3-2.7-.6-5.3-3-7.4-24.2 24.7-44.5 51.8-56.1 84.6 7.4-5.9 14.9-11.4 23.6-16.2-8.3 22.3-19.6 52.8-7.8 101.1 4.6 19 11.9 36.8 24.1 52.3 2.9 3.7 6.3 6.9 9.5 10.3.2-.2.4-.3.6-.5-1.4-7-2.2-14.1-1.5-21.9 2.2 3.2 3.9 6 5.9 8.6 12.6 16 28.7 27.4 47.2 35.6 25 11.3 51.1 13.3 77.9 8.6 54.9-9.7 90.7-48.6 116-98.8 1-1.8.6-2.9-.9-4.2zm172-46.4c-9.5-3.1-22.2-4.2-28.7-2.9 9.9 4 14.1 6.6 18.8 12 12.6 14.4 10.4 34.7-5.4 45.6-11.7 8.1-24.9 10.5-38.9 9.1-1.2-.1-2.3-.4-3-.6 2.8-3.7 6-7 8.1-10.8 9.4-16.8 5.4-42.1-8.7-56.1-2.1-2.1-4.6-3.9-7-5.9-.3 1.3-.1 2.1.1 2.8 4.2 16.6-8.1 32.4-24.8 31.8-7.6-.3-13.9-3.8-19.6-8.5-19.5-16.1-39.1-32.1-58.5-48.3-5.9-4.9-12.5-8.1-20.1-8.7-4.6-.4-9.3-.6-13.9-.9-5.9-.4-8.8-2.8-10.4-8.4-.9-3.4-1.5-6.8-2.2-10.2-1.5-8.1-6.2-13-14.3-14.2-4.4-.7-8.9-1-13.3-1.5-13-1.4-19.8-7.4-22.6-20.3-5 11-1.6 22.4 7.3 29.9 4.5 3.8 9.3 7.3 13.8 11.2 4.6 3.8 7.4 8.7 7.9 14.8.4 4.7.8 9.5 1.8 14.1 2.2 10.6 8.9 18.4 17 25.1 16.5 13.7 33 27.3 49.5 41.1 17.9 15 13.9 32.8 13 56-.9 22.9 12.2 42.9 33.5 51.2 1 .4 2 .6 3.6 1.1-15.7-18.2-10.1-44.1.7-52.3.3 2.2.4 4.3.9 6.4 9.4 44.1 45.4 64.2 85 56.9 16-2.9 30.6-8.9 42.9-19.8 2-1.8 3.7-4.1 5.9-6.5-19.3 4.6-35.8.1-50.9-10.6.7-.3 1.3-.3 1.9-.3 21.3 1.8 40.6-3.4 57-17.4 19.5-16.6 26.6-42.9 17.4-66-8.3-20.1-23.6-32.3-43.8-38.9zM99.4 179.3c-5.3-9.2-13.2-15.6-22.1-21.3 13.7-.5 26.6.2 39.6 3.7-7-12.2-8.5-24.7-5-38.7 5.3 11.9 13.7 20.1 23.6 26.8 19.7 13.2 35.7 19.6 46.7 30.2 3.4 3.3 6.3 7.1 9.6 10.9-.8-2.1-1.4-4.1-2.2-6-5-10.6-13-18.6-22.6-25-1.8-1.2-2.8-2.5-3.4-4.5-3.3-12.5-3-25.1-.7-37.6 1-5.5 2.8-10.9 4.5-16.3.8-2.4 2.3-4.6 4-6.6.6 6.9 0 25.5 19.6 46 10.8 11.3 22.4 21.9 33.9 32.7 9 8.5 18.3 16.7 25.5 26.8 1.1 1.6 2.2 3.3 3.8 4.7-5-13-14.2-24.1-24.2-33.8-9.6-9.3-19.4-18.4-29.2-27.4-3.3-3-4.6-6.7-5.1-10.9-1.2-10.4 0-20.6 4.3-30.2.5-1 1.1-2 1.9-3.3.5 4.2.6 7.9 1.4 11.6 4.8 23.1 20.4 36.3 49.3 63.5 10 9.4 19.3 19.2 25.6 31.6 4.8 9.3 7.3 19 5.7 29.6-.1.6.5 1.7 1.1 2 6.2 2.6 10 6.9 9.7 14.3 7.7-2.6 12.5-8 16.4-14.5 4.2 20.2-9.1 50.3-27.2 58.7.4-4.5 5-23.4-16.5-27.7-6.8-1.3-12.8-1.3-22.9-2.1 4.7-9 10.4-20.6.5-22.4-24.9-4.6-52.8 1.9-57.8 4.6 8.2.4 16.3 1 23.5 3.3-2 6.5-4 12.7-5.8 18.9-1.9 6.5 2.1 14.6 9.3 9.6 1.2-.9 2.3-1.9 3.3-2.7-3.1 17.9-2.9 15.9-2.8 18.3.3 10.2 9.5 7.8 15.7 7.3-2.5 11.8-29.5 27.3-45.4 25.8 7-4.7 12.7-10.3 15.9-17.9-6.5.8-12.9 1.6-19.2 2.4l-.3-.9c4.7-3.4 8-7.8 10.2-13.1 8.7-21.1-3.6-38-25-39.9-9.1-.8-17.8.8-25.9 5.5 6.2-15.6 17.2-26.6 32.6-34.5-15.2-4.3-8.9-2.7-24.6-6.3 14.6-9.3 30.2-13.2 46.5-14.6-5.2-3.2-48.1-3.6-70.2 20.9 7.9 1.4 15.5 2.8 23.2 4.2-23.8 7-44 19.7-62.4 35.6 1.1-4.8 2.7-9.5 3.3-14.3.6-4.5.8-9.2.1-13.6-1.5-9.4-8.9-15.1-19.7-16.3-7.9-.9-15.6.1-23.3 1.3-.9.1-1.7.3-2.9 0 15.8-14.8 36-21.7 53.1-33.5 6-4.5 6.8-8.2 3-14.9zm128.4 26.8c3.3 16 12.6 25.5 23.8 24.3-4.6-11.3-12.1-19.5-23.8-24.3z"],dashcube:[448,512,[],"f210","M326.6 104H110.4c-51.1 0-91.2 43.3-91.2 93.5V427c0 50.5 40.1 85 91.2 85h227.2c51.1 0 91.2-34.5 91.2-85V0L326.6 104zM153.9 416.5c-17.7 0-32.4-15.1-32.4-32.8V240.8c0-17.7 14.7-32.5 32.4-32.5h140.7c17.7 0 32 14.8 32 32.5v123.5l51.1 52.3H153.9z"],delicious:[448,512,[],"f1a5","M446.5 68c-.4-1.5-.9-3-1.4-4.5-.9-2.5-2-4.8-3.3-7.1-1.4-2.4-3-4.8-4.7-6.9-2.1-2.5-4.4-4.8-6.9-6.8-1.1-.9-2.2-1.7-3.3-2.5-1.3-.9-2.6-1.7-4-2.4-1.8-1-3.6-1.8-5.5-2.5-1.7-.7-3.5-1.3-5.4-1.7-3.8-1-7.9-1.5-12-1.5H48C21.5 32 0 53.5 0 80v352c0 4.1.5 8.2 1.5 12 2 7.7 5.8 14.6 11 20.3 1 1.1 2.1 2.2 3.3 3.3 5.7 5.2 12.6 9 20.3 11 3.8 1 7.9 1.5 12 1.5h352c26.5 0 48-21.5 48-48V80c-.1-4.1-.6-8.2-1.6-12zM416 432c0 8.8-7.2 16-16 16H224V256H32V80c0-8.8 7.2-16 16-16h176v192h192v176z"],deploydog:[512,512,[],"f38e","M382.2 136h51.7v239.6h-51.7v-20.7c-19.8 24.8-52.8 24.1-73.8 14.7-26.2-11.7-44.3-38.1-44.3-71.8 0-29.8 14.8-57.9 43.3-70.8 20.2-9.1 52.7-10.6 74.8 12.9V136zm-64.7 161.8c0 18.2 13.6 33.5 33.2 33.5 19.8 0 33.2-16.4 33.2-32.9 0-17.1-13.7-33.2-33.2-33.2-19.6 0-33.2 16.4-33.2 32.6zM188.5 136h51.7v239.6h-51.7v-20.7c-19.8 24.8-52.8 24.1-73.8 14.7-26.2-11.7-44.3-38.1-44.3-71.8 0-29.8 14.8-57.9 43.3-70.8 20.2-9.1 52.7-10.6 74.8 12.9V136zm-64.7 161.8c0 18.2 13.6 33.5 33.2 33.5 19.8 0 33.2-16.4 33.2-32.9 0-17.1-13.7-33.2-33.2-33.2-19.7 0-33.2 16.4-33.2 32.6zM448 96c17.5 0 32 14.4 32 32v256c0 17.5-14.4 32-32 32H64c-17.5 0-32-14.4-32-32V128c0-17.5 14.4-32 32-32h384m0-32H64C28.8 64 0 92.8 0 128v256c0 35.2 28.8 64 64 64h384c35.2 0 64-28.8 64-64V128c0-35.2-28.8-64-64-64z"],deskpro:[480,512,[],"f38f","M205.9 512l31.1-38.4c12.3-.2 25.6-1.4 36.5-6.6 38.9-18.6 38.4-61.9 38.3-63.8-.1-5-.8-4.4-28.9-37.4H362c-.2 50.1-7.3 68.5-10.2 75.7-9.4 23.7-43.9 62.8-95.2 69.4-8.7 1.1-32.8 1.2-50.7 1.1zm200.4-167.7c38.6 0 58.5-13.6 73.7-30.9l-175.5-.3-17.4 31.3 119.2-.1zm-43.6-223.9v168.3h-73.5l-32.7 55.5H250c-52.3 0-58.1-56.5-58.3-58.9-1.2-13.2-21.3-11.6-20.1 1.8 1.4 15.8 8.8 40 26.4 57.1h-91c-25.5 0-110.8-26.8-107-114V16.9C0 .9 9.7.3 15 .1h82c.2 0 .3.1.5.1 4.3-.4 50.1-2.1 50.1 43.7 0 13.3 20.2 13.4 20.2 0 0-18.2-5.5-32.8-15.8-43.7h84.2c108.7-.4 126.5 79.4 126.5 120.2zm-132.5 56l64 29.3c13.3-45.5-42.2-71.7-64-29.3z"],deviantart:[320,512,[],"f1bd","M320 93.2l-98.2 179.1 7.4 9.5H320v127.7H159.1l-13.5 9.2-43.7 84c-.3 0-8.6 8.6-9.2 9.2H0v-93.2l93.2-179.4-7.4-9.2H0V102.5h156l13.5-9.2 43.7-84c.3 0 8.6-8.6 9.2-9.2H320v93.1z"],digg:[512,512,[],"f1a6","M81.7 172.3H0v174.4h132.7V96h-51v76.3zm0 133.4H50.9v-92.3h30.8v92.3zm297.2-133.4v174.4h81.8v28.5h-81.8V416H512V172.3H378.9zm81.8 133.4h-30.8v-92.3h30.8v92.3zm-235.6 41h82.1v28.5h-82.1V416h133.3V172.3H225.1v174.4zm51.2-133.3h30.8v92.3h-30.8v-92.3zM153.3 96h51.3v51h-51.3V96zm0 76.3h51.3v174.4h-51.3V172.3z"],"digital-ocean":[512,512,[],"f391","M256 504v-96.1c101.8 0 180.8-100.9 141.7-208-14.3-39.6-46.1-71.4-85.8-85.7-107.1-38.8-208.1 39.9-208.1 141.7H8C8 93.7 164.9-32.8 335 20.3c74.2 23.3 133.6 82.4 156.6 156.6C544.8 347.2 418.6 504 256 504zm.3-191.4h-95.6v95.6h95.6v-95.6zm-95.6 95.6H87v73.6h73.7v-73.6zM87 346.6H25.4v61.6H87v-61.6z"],discord:[448,512,[],"f392","M297.216 243.2c0 15.616-11.52 28.416-26.112 28.416-14.336 0-26.112-12.8-26.112-28.416s11.52-28.416 26.112-28.416c14.592 0 26.112 12.8 26.112 28.416zm-119.552-28.416c-14.592 0-26.112 12.8-26.112 28.416s11.776 28.416 26.112 28.416c14.592 0 26.112-12.8 26.112-28.416.256-15.616-11.52-28.416-26.112-28.416zM448 52.736V512c-64.494-56.994-43.868-38.128-118.784-107.776l13.568 47.36H52.48C23.552 451.584 0 428.032 0 398.848V52.736C0 23.552 23.552 0 52.48 0h343.04C424.448 0 448 23.552 448 52.736zm-72.96 242.688c0-82.432-36.864-149.248-36.864-149.248-36.864-27.648-71.936-26.88-71.936-26.88l-3.584 4.096c43.52 13.312 63.744 32.512 63.744 32.512-60.811-33.329-132.244-33.335-191.232-7.424-9.472 4.352-15.104 7.424-15.104 7.424s21.248-20.224 67.328-33.536l-2.56-3.072s-35.072-.768-71.936 26.88c0 0-36.864 66.816-36.864 149.248 0 0 21.504 37.12 78.08 38.912 0 0 9.472-11.52 17.152-21.248-32.512-9.728-44.8-30.208-44.8-30.208 3.766 2.636 9.976 6.053 10.496 6.4 43.21 24.198 104.588 32.126 159.744 8.96 8.96-3.328 18.944-8.192 29.44-15.104 0 0-12.8 20.992-46.336 30.464 7.68 9.728 16.896 20.736 16.896 20.736 56.576-1.792 78.336-38.912 78.336-38.912z"],discourse:[448,512,[],"f393","M225.9 32C103.3 32 0 130.5 0 252.1 0 256 .1 480 .1 480l225.8-.2c122.7 0 222.1-102.3 222.1-223.9C448 134.3 348.6 32 225.9 32zM224 384c-19.4 0-37.9-4.3-54.4-12.1L88.5 392l22.9-75c-9.8-18.1-15.4-38.9-15.4-61 0-70.7 57.3-128 128-128s128 57.3 128 128-57.3 128-128 128z"],dochub:[416,512,[],"f394","M397.9 160H256V19.6L397.9 160zM304 192v130c0 66.8-36.5 100.1-113.3 100.1H96V84.8h94.7c12 0 23.1.8 33.1 2.5v-84C212.9 1.1 201.4 0 189.2 0H0v512h189.2C329.7 512 400 447.4 400 318.1V192h-96z"],docker:[640,512,[],"f395","M349.9 236.3h-66.1v-59.4h66.1v59.4zm0-204.3h-66.1v60.7h66.1V32zm78.2 144.8H362v59.4h66.1v-59.4zm-156.3-72.1h-66.1v60.1h66.1v-60.1zm78.1 0h-66.1v60.1h66.1v-60.1zm276.8 100c-14.4-9.7-47.6-13.2-73.1-8.4-3.3-24-16.7-44.9-41.1-63.7l-14-9.3-9.3 14c-18.4 27.8-23.4 73.6-3.7 103.8-8.7 4.7-25.8 11.1-48.4 10.7H2.4c-8.7 50.8 5.8 116.8 44 162.1 37.1 43.9 92.7 66.2 165.4 66.2 157.4 0 273.9-72.5 328.4-204.2 21.4.4 67.6.1 91.3-45.2 1.5-2.5 6.6-13.2 8.5-17.1l-13.3-8.9zm-511.1-27.9h-66v59.4h66.1v-59.4zm78.1 0h-66.1v59.4h66.1v-59.4zm78.1 0h-66.1v59.4h66.1v-59.4zm-78.1-72.1h-66.1v60.1h66.1v-60.1z"],draft2digital:[480,512,[],"f396","M369.9 425.4V371l47.1 27.2-47.1 27.2zM82.4 380.6c25.5-27.3 97.7-104.7 150.9-170 35.1-43.1 40.3-82.4 28.4-112.7-7.4-18.8-17.5-30.2-24.3-35.7 45.3 2.1 68 23.4 82.2 38.3 0 0 42.4 48.2 5.8 113.3-37 65.9-110.9 147.5-128.5 166.7H82.4zm51.8-219.2c0 12.4-10 22.4-22.4 22.4-12.4 0-22.4-10-22.4-22.4 0-12.4 10-22.4 22.4-22.4 12.4 0 22.4 10.1 22.4 22.4M336 315.9v64.7h-91.3c30.8-35 81.8-95.9 111.8-149.3 35.2-62.6 16.1-123.4-12.8-153.3-4.4-4.6-62.2-62.9-166-41.2-59.1 12.4-89.4 43.4-104.3 67.3-13.1 20.9-17 39.8-18.2 47.7-5.5 33 19.4 67.1 56.7 67.1 31.7 0 57.3-25.7 57.3-57.4 0-27.1-19.7-52.1-48-56.8 1.8-7.3 17.7-21.1 26.3-24.7 41.1-17.3 78 5.2 83.3 33.5 8.3 44.3-37.1 90.4-69.7 127.6C84.5 328.1 18.3 396.8 0 415.9l336-.1V480l144-81.9-144-82.2z"],dribbble:[512,512,[],"f17d","M256 8C119.252 8 8 119.252 8 256s111.252 248 248 248 248-111.252 248-248S392.748 8 256 8zm163.97 114.366c29.503 36.046 47.369 81.957 47.835 131.955-6.984-1.477-77.018-15.682-147.502-6.818-5.752-14.041-11.181-26.393-18.617-41.614 78.321-31.977 113.818-77.482 118.284-83.523zM396.421 97.87c-3.81 5.427-35.697 48.286-111.021 76.519-34.712-63.776-73.185-116.168-79.04-124.008 67.176-16.193 137.966 1.27 190.061 47.489zm-230.48-33.25c5.585 7.659 43.438 60.116 78.537 122.509-99.087 26.313-186.36 25.934-195.834 25.809C62.38 147.205 106.678 92.573 165.941 64.62zM44.17 256.323c0-2.166.043-4.322.108-6.473 9.268.19 111.92 1.513 217.706-30.146 6.064 11.868 11.857 23.915 17.174 35.949-76.599 21.575-146.194 83.527-180.531 142.306C64.794 360.405 44.17 310.73 44.17 256.323zm81.807 167.113c22.127-45.233 82.178-103.622 167.579-132.756 29.74 77.283 42.039 142.053 45.189 160.638-68.112 29.013-150.015 21.053-212.768-27.882zm248.38 8.489c-2.171-12.886-13.446-74.897-41.152-151.033 66.38-10.626 124.7 6.768 131.947 9.055-9.442 58.941-43.273 109.844-90.795 141.978z"],"dribbble-square":[448,512,[],"f397","M90.2 228.2c8.9-42.4 37.4-77.7 75.7-95.7 3.6 4.9 28 38.8 50.7 79-64 17-120.3 16.8-126.4 16.7zM314.6 154c-33.6-29.8-79.3-41.1-122.6-30.6 3.8 5.1 28.6 38.9 51 80 48.6-18.3 69.1-45.9 71.6-49.4zM140.1 364c40.5 31.6 93.3 36.7 137.3 18-2-12-10-53.8-29.2-103.6-55.1 18.8-93.8 56.4-108.1 85.6zm98.8-108.2c-3.4-7.8-7.2-15.5-11.1-23.2C159.6 253 93.4 252.2 87.4 252c0 1.4-.1 2.8-.1 4.2 0 35.1 13.3 67.1 35.1 91.4 22.2-37.9 67.1-77.9 116.5-91.8zm34.9 16.3c17.9 49.1 25.1 89.1 26.5 97.4 30.7-20.7 52.5-53.6 58.6-91.6-4.6-1.5-42.3-12.7-85.1-5.8zm-20.3-48.4c4.8 9.8 8.3 17.8 12 26.8 45.5-5.7 90.7 3.4 95.2 4.4-.3-32.3-11.8-61.9-30.9-85.1-2.9 3.9-25.8 33.2-76.3 53.9zM448 80v352c0 26.5-21.5 48-48 48H48c-26.5 0-48-21.5-48-48V80c0-26.5 21.5-48 48-48h352c26.5 0 48 21.5 48 48zm-64 176c0-88.2-71.8-160-160-160S64 167.8 64 256s71.8 160 160 160 160-71.8 160-160z"],dropbox:[528,512,[],"f16b","M264.4 116.3l-132 84.3 132 84.3-132 84.3L0 284.1l132.3-84.3L0 116.3 132.3 32l132.1 84.3zM131.6 395.7l132-84.3 132 84.3-132 84.3-132-84.3zm132.8-111.6l132-84.3-132-83.6L395.7 32 528 116.3l-132.3 84.3L528 284.8l-132.3 84.3-131.3-85z"],drupal:[448,512,[],"f1a9","M319.5 114.7c-22.2-14-43.5-19.5-64.7-33.5-13-8.8-31.3-30-46.5-48.3-2.7 29.3-11.5 41.2-22 49.5-21.3 17-34.8 22.2-53.5 32.3C117 123 32 181.5 32 290.5 32 399.7 123.8 480 225.8 480 327.5 480 416 406 416 294c0-112.3-83-171-96.5-179.3zm2.5 325.6c-20.1 20.1-90.1 28.7-116.7 4.2-4.8-4.8.3-12 6.5-12 0 0 17 13.3 51.5 13.3 27 0 46-7.7 54.5-14 6.1-4.6 8.4 4.3 4.2 8.5zm-54.5-52.6c8.7-3.6 29-3.8 36.8 1.3 4.1 2.8 16.1 18.8 6.2 23.7-8.4 4.2-1.2-15.7-26.5-15.7-14.7 0-19.5 5.2-26.7 11-7 6-9.8 8-12.2 4.7-6-8.2 15.9-22.3 22.4-25zM360 405c-15.2-1-45.5-48.8-65-49.5-30.9-.9-104.1 80.7-161.3 42-38.8-26.6-14.6-104.8 51.8-105.2 49.5-.5 83.8 49 108.5 48.5 21.3-.3 61.8-41.8 81.8-41.8 48.7 0 23.3 109.3-15.8 106z"],dyalog:[416,512,[],"f399","M0 32v119.2h64V96h107.2C284.6 96 352 176.2 352 255.9 352 332 293.4 416 171.2 416H0v64h171.2C331.9 480 416 367.3 416 255.9c0-58.7-22.1-113.4-62.3-154.3C308.9 56 245.7 32 171.2 32H0z"],earlybirds:[480,512,[],"f39a","M313.2 47.5c1.2-13 21.3-14 36.6-8.7.9.3 26.2 9.7 19 15.2-27.9-7.4-56.4 18.2-55.6-6.5zm-201 6.9c30.7-8.1 62 20 61.1-7.1-1.3-14.2-23.4-15.3-40.2-9.6-1 .3-28.7 10.5-20.9 16.7zM319.4 160c-8.8 0-16 7.2-16 16s7.2 16 16 16 16-7.2 16-16-7.2-16-16-16zm-159.7 0c-8.8 0-16 7.2-16 16s7.2 16 16 16 16-7.2 16-16-7.2-16-16-16zm318.5 163.2c-9.9 24-40.7 11-63.9-1.2-13.5 69.1-58.1 111.4-126.3 124.2.3.9-2-.1 24 1 33.6 1.4 63.8-3.1 97.4-8-19.8-13.8-11.4-37.1-9.8-38.1 1.4-.9 14.7 1.7 21.6 11.5 8.6-12.5 28.4-14.8 30.2-13.6 1.6 1.1 6.6 20.9-6.9 34.6 4.7-.9 8.2-1.6 9.8-2.1 2.6-.8 17.7 11.3 3.1 13.3-14.3 2.3-22.6 5.1-47.1 10.8-45.9 10.7-85.9 11.8-117.7 12.8l1 11.6c3.8 18.1-23.4 24.3-27.6 6.2.8 17.9-27.1 21.8-28.4-1l-.5 5.3c-.7 18.4-28.4 17.9-28.3-.6-7.5 13.5-28.1 6.8-26.4-8.5l1.2-12.4c-36.7.9-59.7 3.1-61.8 3.1-20.9 0-20.9-31.6 0-31.6 2.4 0 27.7 1.3 63.2 2.8-61.1-15.5-103.7-55-114.9-118.2-25 12.8-57.5 26.8-68.2.8-10.5-25.4 21.5-42.6 66.8-73.4.7-6.6 1.6-13.3 2.7-19.8-14.4-19.6-11.6-36.3-16.1-60.4-16.8 2.4-23.2-9.1-23.6-23.1.3-7.3 2.1-14.9 2.4-15.4 1.1-1.8 10.1-2 12.7-2.6 6-31.7 50.6-33.2 90.9-34.5 19.7-21.8 45.2-41.5 80.9-48.3C203.3 29 215.2 8.5 216.2 8c1.7-.8 21.2 4.3 26.3 23.2 5.2-8.8 18.3-11.4 19.6-10.7 1.1.6 6.4 15-4.9 25.9 40.3 3.5 72.2 24.7 96 50.7 36.1 1.5 71.8 5.9 77.1 34 2.7.6 11.6.8 12.7 2.6.3.5 2.1 8.1 2.4 15.4-.5 13.9-6.8 25.4-23.6 23.1-3.2 17.3-2.7 32.9-8.7 47.7 2.4 11.7 4 23.8 4.8 36.4 37 25.4 70.3 42.5 60.3 66.9zM207.4 159.9c.9-44-37.9-42.2-78.6-40.3-21.7 1-38.9 1.9-45.5 13.9-11.4 20.9 5.9 92.9 23.2 101.2 9.8 4.7 73.4 7.9 86.3-7.1 8.2-9.4 15-49.4 14.6-67.7zm52 58.3c-4.3-12.4-6-30.1-15.3-32.7-2-.5-9-.5-11 0-10 2.8-10.8 22.1-17 37.2 15.4 0 19.3 9.7 23.7 9.7 4.3 0 6.3-11.3 19.6-14.2zm135.7-84.7c-6.6-12.1-24.8-12.9-46.5-13.9-40.2-1.9-78.2-3.8-77.3 40.3-.5 18.3 5 58.3 13.2 67.8 13 14.9 76.6 11.8 86.3 7.1 15.8-7.6 36.5-78.9 24.3-101.3z"],ebay:[640,512,[],"f4f4","M405.2 263.8c-29.1.9-47.2 6.2-47.2 25.3 0 12.4 9.9 25.8 35 25.8 33.7 0 51.6-18.4 51.6-48.4v-3.3c-11.8 0-26.3.1-39.4.6m71.5 39.7c0 9.3.3 18.6 1 26.8h-29.8c-.8-6.9-1.1-13.6-1.1-20.2-16.1 19.8-35.3 25.5-61.9 25.5-39.5 0-60.6-20.9-60.6-45 0-35 28.8-47.3 78.6-48.4 13.7-.3 29-.4 41.7-.4v-3.4c0-23.4-15-33-41-33-19.3 0-33.6 8-35 21.8h-33.7c3.6-34.4 39.7-43.1 71.5-43.1 38.1 0 70.3 13.5 70.3 53.8v65.6zm-349-56.8c-2.3-54.7-87.5-56.6-94.4 0h94.4zm-95 21.4c3.5 58.3 79.2 57.4 91.2 21.6H157c-6.4 34.4-43 46.1-74.4 46.1-57.2 0-82.5-31.5-82.5-74 0-46.8 26.2-77.6 83-77.6 45.3 0 78.4 23.7 78.4 75.4v8.5H32.7zm211 45.7c29.8 0 50.2-21.5 50.2-53.8 0-32.4-20.4-53.8-50.2-53.8-29.6 0-50.2 21.4-50.2 53.8 0 32.3 20.6 53.8 50.2 53.8m-82.2-186h32.1v80.6c15.7-18.7 37.4-24.2 58.7-24.2 35.7 0 75.4 24.1 75.4 76.2 0 43.6-31.5 75.4-76 75.4-23.3 0-45.1-8.3-58.7-24.9 0 6.6-.4 13.2-1.1 19.5h-31.5c.5-10.2 1.1-22.8 1.1-33.1V127.8zM640 189.5l-99.2 194.8h-35.9l28.5-54.1-74.6-140.7h37.5l54.9 109.9L606 189.5h34z"],edge:[512,512,[],"f282","M25.714 228.163c.111-.162.23-.323.342-.485-.021.162-.045.323-.065.485h-.277zm460.572 15.508c0-44.032-7.754-84.465-28.801-122.405C416.498 47.879 343.912 8.001 258.893 8.001 118.962 7.724 40.617 113.214 26.056 227.679c42.429-61.312 117.073-121.376 220.375-124.966 0 0 109.666 0 99.419 104.957H169.997c6.369-37.386 18.554-58.986 34.339-78.926-75.048 34.893-121.85 96.096-120.742 188.315.83 71.448 50.124 144.836 120.743 171.976 83.357 31.847 192.776 7.2 240.132-21.324V363.307c-80.864 56.494-270.871 60.925-272.255-67.572h314.073v-52.064z"],elementor:[448,512,[],"f430","M425.6 32H22.4C10 32 0 42 0 54.4v403.2C0 470 10 480 22.4 480h403.2c12.4 0 22.4-10 22.4-22.4V54.4C448 42 438 32 425.6 32M164.3 355.5h-39.8v-199h39.8v199zm159.3 0H204.1v-39.8h119.5v39.8zm0-79.6H204.1v-39.8h119.5v39.8zm0-79.7H204.1v-39.8h119.5v39.8z"],ember:[640,512,[],"f423","M639.9 254.6c-1.1-10.7-10.7-6.8-10.7-6.8s-15.6 12.1-29.3 10.7c-13.7-1.3-9.4-32-9.4-32s3-28.1-5.1-30.4c-8.1-2.4-18 7.3-18 7.3s-12.4 13.7-18.3 31.2l-1.6.5s1.9-30.6-.3-37.6c-1.6-3.5-16.4-3.2-18.8 3s-14.2 49.2-15 67.2c0 0-23.1 19.6-43.3 22.8s-25-9.4-25-9.4 54.8-15.3 52.9-59.1c-1.9-43.8-44.2-27.6-49-24-4.6 3.5-29.4 18.4-36.6 59.7-.2 1.4-.7 7.5-.7 7.5s-21.2 14.2-33 18c0 0 33-55.6-7.3-80.9-18.3-11-32.8 12.1-32.8 12.1s54.5-60.7 42.5-112c-5.8-24.4-18-27.1-29.2-23.1-17 6.7-23.5 16.7-23.5 16.7s-22 32-27.1 79.5-12.6 105.1-12.6 105.1-10.5 10.2-20.2 10.7-5.4-28.7-5.4-28.7 7.5-44.6 7-52.1-1.1-11.6-9.9-14.2c-8.9-2.7-18.5 8.6-18.5 8.6s-25.5 38.7-27.7 44.6l-1.3 2.4-1.3-1.6s18-52.7.8-53.5c-17.2-.8-28.5 18.8-28.5 18.8s-19.6 32.8-20.4 36.5l-1.3-1.6s8.1-38.2 6.4-47.6c-1.6-9.4-10.5-7.5-10.5-7.5s-11.3-1.3-14.2 5.9-13.7 55.3-15 70.7c0 0-28.2 20.2-46.8 20.4-18.5.3-16.7-11.8-16.7-11.8s68-23.3 49.4-69.2c-8.3-11.8-18-15.5-31.7-15.3-13.7.3-30.3 8.6-41.3 33.3-5.3 11.8-6.8 23-7.8 31.5 0 0-12.3 2.4-18.8-2.9s-10 0-10 0-11.2 14-.1 18.3 28.1 6.1 28.1 6.1c1.6 7.5 6.2 19.5 19.6 29.7 20.2 15.3 58.8-1.3 58.8-1.3l15.9-8.8s.5 14.6 12.1 16.7c11.6 2.1 16.4 1 36.5-47.9 11.8-25 12.6-23.6 12.6-23.6l1.3-.3s-9.1 46.8-5.6 59.7C187.7 319.4 203 318 203 318s8.3 2.4 15-21.2c6.7-23.6 19.6-49.9 19.6-49.9h1.6s-5.6 48.1 3 63.7c8.6 15.6 30.9 5.3 30.9 5.3s15.6-7.8 18-10.2c0 0 18.5 15.8 44.6 12.9 58.3-11.5 79.1-25.9 79.1-25.9s10 24.4 41.1 26.7c35.5 2.7 54.8-18.6 54.8-18.6s-.3 13.5 12.1 18.6c12.4 5.1 20.7-22.8 20.7-22.8l20.7-57.2h1.9s1.1 37.3 21.5 43.2 47-13.7 47-13.7 6.4-3.5 5.3-14.3zm-578 5.3c.8-32 21.8-45.9 29-39 7.3 7 4.6 22-9.1 31.4-13.7 9.5-19.9 7.6-19.9 7.6zm272.8-123.8s19.1-49.7 23.6-25.5-40 96.2-40 96.2c.5-16.2 16.4-70.7 16.4-70.7zm22.8 138.4c-12.6 33-43.3 19.6-43.3 19.6s-3.5-11.8 6.4-44.9 33.3-20.2 33.3-20.2 16.2 12.4 3.6 45.5zm84.6-14.6s-3-10.5 8.1-30.6c11-20.2 19.6-9.1 19.6-9.1s9.4 10.2-1.3 25.5-26.4 14.2-26.4 14.2z"],empire:[496,512,[],"f1d1","M287.6 54.2c-10.8-2.2-22.1-3.3-33.5-3.6V32.4c78.1 2.2 146.1 44 184.6 106.6l-15.8 9.1c-6.1-9.7-12.7-18.8-20.2-27.1l-18 15.5c-26-29.6-61.4-50.7-101.9-58.4l4.8-23.9zM53.4 322.4l23-7.7c-6.4-18.3-10-38.2-10-58.7s3.3-40.4 9.7-58.7l-22.7-7.7c3.6-10.8 8.3-21.3 13.6-31l-15.8-9.1C34 181 24.1 217.5 24.1 256s10 75 27.1 106.6l15.8-9.1c-5.3-10-9.7-20.3-13.6-31.1zM213.1 434c-40.4-8-75.8-29.1-101.9-58.7l-18 15.8c-7.5-8.6-14.4-17.7-20.2-27.4l-16 9.4c38.5 62.3 106.8 104.3 184.9 106.6v-18.3c-11.3-.3-22.7-1.7-33.5-3.6l4.7-23.8zM93.3 120.9l18 15.5c26-29.6 61.4-50.7 101.9-58.4l-4.7-23.8c10.8-2.2 22.1-3.3 33.5-3.6V32.4C163.9 34.6 95.9 76.4 57.4 139l15.8 9.1c6-9.7 12.6-18.9 20.1-27.2zm309.4 270.2l-18-15.8c-26 29.6-61.4 50.7-101.9 58.7l4.7 23.8c-10.8 1.9-22.1 3.3-33.5 3.6v18.3c78.1-2.2 146.4-44.3 184.9-106.6l-16.1-9.4c-5.7 9.7-12.6 18.8-20.1 27.4zM496 256c0 137-111 248-248 248S0 393 0 256 111 8 248 8s248 111 248 248zm-12.2 0c0-130.1-105.7-235.8-235.8-235.8S12.2 125.9 12.2 256 117.9 491.8 248 491.8 483.8 386.1 483.8 256zm-39-106.6l-15.8 9.1c5.3 9.7 10 20.2 13.6 31l-22.7 7.7c6.4 18.3 9.7 38.2 9.7 58.7s-3.6 40.4-10 58.7l23 7.7c-3.9 10.8-8.3 21-13.6 31l15.8 9.1C462 331 471.9 294.5 471.9 256s-9.9-75-27.1-106.6zm-183 177.7c16.3-3.3 30.4-11.6 40.7-23.5l51.2 44.8c11.9-13.6 21.3-29.3 27.1-46.8l-64.2-22.1c2.5-7.5 3.9-15.2 3.9-23.5s-1.4-16.1-3.9-23.5l64.5-22.1c-6.1-17.4-15.5-33.2-27.4-46.8l-51.2 44.8c-10.2-11.9-24.4-20.5-40.7-23.8l13.3-66.4c-8.6-1.9-17.7-2.8-27.1-2.8-9.4 0-18.5.8-27.1 2.8l13.3 66.4c-16.3 3.3-30.4 11.9-40.7 23.8l-51.2-44.8c-11.9 13.6-21.3 29.3-27.4 46.8l64.5 22.1c-2.5 7.5-3.9 15.2-3.9 23.5s1.4 16.1 3.9 23.5l-64.2 22.1c5.8 17.4 15.2 33.2 27.1 46.8l51.2-44.8c10.2 11.9 24.4 20.2 40.7 23.5l-13.3 66.7c8.6 1.7 17.7 2.8 27.1 2.8 9.4 0 18.5-1.1 27.1-2.8l-13.3-66.7z"],envira:[448,512,[],"f299","M0 32c477.6 0 366.6 317.3 367.1 366.3L448 480h-26l-70.4-71.2c-39 4.2-124.4 34.5-214.4-37C47 300.3 52 214.7 0 32zm79.7 46c-49.7-23.5-5.2 9.2-5.2 9.2 45.2 31.2 66 73.7 90.2 119.9 31.5 60.2 79 139.7 144.2 167.7 65 28 34.2 12.5 6-8.5-28.2-21.2-68.2-87-91-130.2-31.7-60-61-118.6-144.2-158.1z"],erlang:[640,512,[],"f39d","M21.7 246.4c-.1 86.8 29 159.5 78.7 212.1H0v-405h87.2c-41.5 50.2-65.6 116.2-65.5 192.9zM640 53.6h-83.6c31.4 42.7 48.7 97.5 46.2 162.7.5 6 .5 11.7 0 24.1H230.2c-.2 109.7 38.9 194.9 138.6 195.3 68.5-.3 118-51 151.9-106.1l96.4 48.2c-17.4 30.9-36.5 57.8-57.9 80.8H640v-405zm-80.8 405s0-.1 0 0h-.2.2zm-3.1-405h.3l-.1-.1-.2.1zm-230.7 9.6c-45.9.1-85.1 33.5-89.2 83.2h169.9c-1.1-49.7-34.5-83.1-80.7-83.2z"],ethereum:[320,512,[],"f42e","M311.9 260.8L160 353.6 8 260.8 160 0l151.9 260.8zM160 383.4L8 290.6 160 512l152-221.4-152 92.8z"],etsy:[384,512,[],"f2d7","M384 348c-1.75 10.75-13.75 110-15.5 132-117.879-4.299-219.895-4.743-368.5 0v-25.5c45.457-8.948 60.627-8.019 61-35.25 1.793-72.322 3.524-244.143 0-322-1.029-28.46-12.13-26.765-61-36v-25.5c73.886 2.358 255.933 8.551 362.999-3.75-3.5 38.25-7.75 126.5-7.75 126.5H332C320.947 115.665 313.241 68 277.25 68h-137c-10.25 0-10.75 3.5-10.75 9.75V241.5c58 .5 88.5-2.5 88.5-2.5 29.77-.951 27.56-8.502 40.75-65.251h25.75c-4.407 101.351-3.91 61.829-1.75 160.25H257c-9.155-40.086-9.065-61.045-39.501-61.5 0 0-21.5-2-88-2v139c0 26 14.25 38.25 44.25 38.25H263c63.636 0 66.564-24.996 98.751-99.75H384z"],expeditedssl:[496,512,[],"f23e","M248 43.4C130.6 43.4 35.4 138.6 35.4 256S130.6 468.6 248 468.6 460.6 373.4 460.6 256 365.4 43.4 248 43.4zm-97.4 132.9c0-53.7 43.7-97.4 97.4-97.4s97.4 43.7 97.4 97.4v26.6c0 5-3.9 8.9-8.9 8.9h-17.7c-5 0-8.9-3.9-8.9-8.9v-26.6c0-82.1-124-82.1-124 0v26.6c0 5-3.9 8.9-8.9 8.9h-17.7c-5 0-8.9-3.9-8.9-8.9v-26.6zM389.7 380c0 9.7-8 17.7-17.7 17.7H124c-9.7 0-17.7-8-17.7-17.7V238.3c0-9.7 8-17.7 17.7-17.7h248c9.7 0 17.7 8 17.7 17.7V380zm-248-137.3v132.9c0 2.5-1.9 4.4-4.4 4.4h-8.9c-2.5 0-4.4-1.9-4.4-4.4V242.7c0-2.5 1.9-4.4 4.4-4.4h8.9c2.5 0 4.4 1.9 4.4 4.4zm141.7 48.7c0 13-7.2 24.4-17.7 30.4v31.6c0 5-3.9 8.9-8.9 8.9h-17.7c-5 0-8.9-3.9-8.9-8.9v-31.6c-10.5-6.1-17.7-17.4-17.7-30.4 0-19.7 15.8-35.4 35.4-35.4s35.5 15.8 35.5 35.4zM248 8C111 8 0 119 0 256s111 248 248 248 248-111 248-248S385 8 248 8zm0 478.3C121 486.3 17.7 383 17.7 256S121 25.7 248 25.7 478.3 129 478.3 256 375 486.3 248 486.3z"],facebook:[448,512,[],"f09a","M448 56.7v398.5c0 13.7-11.1 24.7-24.7 24.7H309.1V306.5h58.2l8.7-67.6h-67v-43.2c0-19.6 5.4-32.9 33.5-32.9h35.8v-60.5c-6.2-.8-27.4-2.7-52.2-2.7-51.6 0-87 31.5-87 89.4v49.9h-58.4v67.6h58.4V480H24.7C11.1 480 0 468.9 0 455.3V56.7C0 43.1 11.1 32 24.7 32h398.5c13.7 0 24.8 11.1 24.8 24.7z"],"facebook-f":[264,512,[],"f39e","M76.7 512V283H0v-91h76.7v-71.7C76.7 42.4 124.3 0 193.8 0c33.3 0 61.9 2.5 70.2 3.6V85h-48.2c-37.8 0-45.1 18-45.1 44.3V192H256l-11.7 91h-73.6v229"],"facebook-messenger":[448,512,[],"f39f","M224 32C15.9 32-77.5 278 84.6 400.6V480l75.7-42c142.2 39.8 285.4-59.9 285.4-198.7C445.8 124.8 346.5 32 224 32zm23.4 278.1L190 250.5 79.6 311.6l121.1-128.5 57.4 59.6 110.4-61.1-121.1 128.5z"],"facebook-square":[448,512,[],"f082","M448 80v352c0 26.5-21.5 48-48 48h-85.3V302.8h60.6l8.7-67.6h-69.3V192c0-19.6 5.4-32.9 33.5-32.9H384V98.7c-6.2-.8-27.4-2.7-52.2-2.7-51.6 0-87 31.5-87 89.4v49.9H184v67.6h60.9V480H48c-26.5 0-48-21.5-48-48V80c0-26.5 21.5-48 48-48h352c26.5 0 48 21.5 48 48z"],firefox:[480,512,[],"f269","M478.1 235.3c-.7-4.5-1.4-7.1-1.4-7.1s-1.8 2-4.7 5.9c-.9-10.7-2.8-21.2-5.8-31.6-3.7-12.9-8.5-25.4-14.5-37.4-3.8-8-8.2-15.6-13.3-22.8-1.8-2.7-3.7-5.4-5.6-7.9-8.8-14.4-19-23.3-30.7-40-7.6-12.8-12.9-26.9-15.4-41.6-3.2 8.9-5.7 18-7.4 27.3-12.1-12.2-22.5-20.8-28.9-26.7C319.4 24.2 323 9.1 323 9.1S264.7 74.2 289.9 142c8.7 23 23.8 43.1 43.4 57.9 24.4 20.2 50.8 36 64.7 76.6-11.2-21.3-28.1-39.2-48.8-51.5 6.2 14.7 9.4 30.6 9.3 46.5 0 61-49.6 110.5-110.6 110.4-8.3 0-16.5-.9-24.5-2.8-9.5-1.8-18.7-4.9-27.4-9.3-12.9-7.8-24-18.1-32.8-30.3l-.2-.3 2 .7c4.6 1.6 9.2 2.8 14 3.7 18.7 4 38.3 1.7 55.6-6.6 17.5-9.7 28-16.9 36.6-14h.2c8.4 2.7 15-5.5 9-14-10.4-13.4-27.4-20-44.2-17-17.5 2.5-33.5 15-56.4 2.9-1.5-.8-2.9-1.6-4.3-2.5-1.6-.9 4.9 1.3 3.4.3-5-2.5-9.8-5.4-14.4-8.6-.3-.3 3.5 1.1 3.1.8-5.9-4-11-9.2-15-15.2-4.1-7.4-4.5-16.4-1-24.1 2.1-3.8 5.4-6.9 9.3-8.7 3 1.5 4.8 2.6 4.8 2.6s-1.3-2.5-2.1-3.8c.3-.1.5 0 .8-.2 2.6 1.1 8.3 4 11.4 5.8 2.1 1.1 3.8 2.7 5.2 4.7 0 0 1-.5.3-2.7-1.1-2.7-2.9-5-5.4-6.6h.2c2.3 1.2 4.5 2.6 6.6 4.1 1.9-4.4 2.8-9.2 2.6-14 .2-2.6-.2-5.3-1.1-7.8-.8-1.6.5-2.2 1.9-.5-.2-1.3-.7-2.5-1.2-3.7v-.1s.8-1.1 1.2-1.5c1-1 2.1-1.9 3.4-2.7 7.2-4.5 14.8-8.4 22.7-11.6 6.4-2.8 11.7-4.9 12.8-5.6 1.6-1 3.1-2.2 4.5-3.5 5.3-4.5 9-10.8 10.2-17.7.1-.9.2-1.8.3-2.8v-1.5c-.9-3.5-6.9-6.1-38.4-9.1-11.1-1.8-20-10.1-22.5-21.1v.1c-.4 1.1-.9 2.3-1.3 3.5.4-1.2.8-2.3 1.3-3.5v-.2c6-15.7 16.8-29.1 30.8-38.3.8-.7-3.2.2-2.4-.5 2.7-1.3 5.4-2.5 8.2-3.5 1.4-.6-6-3.4-12.6-2.7-4 .2-8 1.2-11.7 2.8 1.6-1.3 6.2-3.1 5.1-3.1-8.4 1.6-16.5 4.7-23.9 9 0-.8.1-1.5.5-2.2-5.9 2.5-11 6.5-15 11.5.1-.9.2-1.8.2-2.7-2.7 2-5.2 4.3-7.3 6.9l-.1.1c-17.4-6.7-36.3-8.3-54.6-4.7l-.2-.1h.2c-3.8-3.1-7.1-6.7-9.7-10.9l-.2.1-.4-.2c-1.2-1.8-2.4-3.8-3.7-6-.9-1.6-1.8-3.4-2.7-5.2 0-.1-.1-.2-.2-.2-.4 0-.6 1.7-.9 1.3v-.1c-3.2-8.3-4.7-17.2-4.4-26.2l-.2.1c-5.1 3.5-9 8.6-11.1 14.5-.9 2.1-1.6 3.3-2.2 4.5v-.5c.1-1.1.6-3.3.5-3.1-.1.2-.2.3-.3.4-1.5 1.7-2.9 3.7-3.9 5.8-.9 1.9-1.7 3.9-2.3 5.9-.1.3 0-.3 0-1s.1-2 0-1.7l-.3.7c-6.7 14.9-10.9 30.8-12.4 47.1-.4 2.8-.6 5.6-.5 8.3v.2c-4.8 5.2-9 11-12.7 17.1-12.1 20.4-21.1 42.5-26.8 65.6 4-8.8 8.8-17.2 14.3-25.1C5.5 228.5 0 257.4 0 286.6c1.8-8.6 4.2-17 7-25.3-1.7 34.5 4.9 68.9 19.4 100.3 19.4 43.5 51.6 80 92.3 104.7 16.6 11.2 34.7 19.9 53.8 25.8 2.5.9 5.1 1.8 7.7 2.7-.8-.3-1.6-.7-2.4-1 22.6 6.8 46.2 10.3 69.8 10.3 83.7 0 111.3-31.9 113.8-35 4.1-3.7 7.5-8.2 9.9-13.3 1.6-.7 3.2-1.4 4.9-2.1l1-.5 1.9-.9c12.6-5.9 24.5-13.4 35.3-22.1 16.3-11.7 27.9-28.7 32.9-48.1 3-7.1 3.1-15 .4-22.2.9-1.4 1.7-2.8 2.7-4.3 18-28.9 28.2-61.9 29.6-95.9v-2.8c0-7.3-.6-14.5-1.9-21.6z"],"first-order":[448,512,[],"f2b0","M12.9 229.2c.1-.1.2-.3.3-.4 0 .1 0 .3-.1.4h-.2zM224 96.6c-7.1 0-14.6.6-21.4 1.7l3.7 67.4-22-64c-14.3 3.7-27.7 9.4-40 16.6l29.4 61.4-45.1-50.9c-11.4 8.9-21.7 19.1-30.6 30.9l50.6 45.4-61.1-29.7c-7.1 12.3-12.9 25.7-16.6 40l64.3 22.6-68-4c-.9 7.1-1.4 14.6-1.4 22s.6 14.6 1.4 21.7l67.7-4-64 22.6c3.7 14.3 9.4 27.7 16.6 40.3l61.1-29.7L97.7 352c8.9 11.7 19.1 22.3 30.9 30.9l44.9-50.9-29.5 61.4c12.3 7.4 25.7 13.1 40 16.9l22.3-64.6-4 68c7.1 1.1 14.6 1.7 21.7 1.7 7.4 0 14.6-.6 21.7-1.7l-4-68.6 22.6 65.1c14.3-4 27.7-9.4 40-16.9L274.9 332l44.9 50.9c11.7-8.9 22-19.1 30.6-30.9l-50.6-45.1 61.1 29.4c7.1-12.3 12.9-25.7 16.6-40.3l-64-22.3 67.4 4c1.1-7.1 1.4-14.3 1.4-21.7s-.3-14.9-1.4-22l-67.7 4 64-22.3c-3.7-14.3-9.1-28-16.6-40.3l-60.9 29.7 50.6-45.4c-8.9-11.7-19.1-22-30.6-30.9l-45.1 50.9 29.4-61.1c-12.3-7.4-25.7-13.1-40-16.9L241.7 166l4-67.7c-7.1-1.2-14.3-1.7-21.7-1.7zM443.4 128v256L224 512 4.6 384V128L224 0l219.4 128zm-17.1 10.3L224 20.9 21.7 138.3v235.1L224 491.1l202.3-117.7V138.3zM224 37.1l187.7 109.4v218.9L224 474.9 36.3 365.4V146.6L224 37.1zm0 50.9c-92.3 0-166.9 75.1-166.9 168 0 92.6 74.6 167.7 166.9 167.7 92 0 166.9-75.1 166.9-167.7 0-92.9-74.9-168-166.9-168z"],"first-order-alt":[496,512,[],"f50a","M248 8C111.03 8 0 119.03 0 256s111.03 248 248 248 248-111.03 248-248S384.97 8 248 8zm0 488.21C115.34 496.21 7.79 388.66 7.79 256S115.34 15.79 248 15.79 488.21 123.34 488.21 256 380.66 496.21 248 496.21zm0-459.92C126.66 36.29 28.29 134.66 28.29 256S126.66 475.71 248 475.71 467.71 377.34 467.71 256 369.34 36.29 248 36.29zm0 431.22c-116.81 0-211.51-94.69-211.51-211.51S131.19 44.49 248 44.49 459.51 139.19 459.51 256 364.81 467.51 248 467.51zm186.23-162.98a191.613 191.613 0 0 1-20.13 48.69l-74.13-35.88 61.48 54.82a193.515 193.515 0 0 1-37.2 37.29l-54.8-61.57 35.88 74.27a190.944 190.944 0 0 1-48.63 20.23l-27.29-78.47 4.79 82.93c-8.61 1.18-17.4 1.8-26.33 1.8s-17.72-.62-26.33-1.8l4.76-82.46-27.15 78.03a191.365 191.365 0 0 1-48.65-20.2l35.93-74.34-54.87 61.64a193.85 193.85 0 0 1-37.22-37.28l61.59-54.9-74.26 35.93a191.638 191.638 0 0 1-20.14-48.69l77.84-27.11-82.23 4.76c-1.16-8.57-1.78-17.32-1.78-26.21 0-9 .63-17.84 1.82-26.51l82.38 4.77-77.94-27.16a191.726 191.726 0 0 1 20.23-48.67l74.22 35.92-61.52-54.86a193.85 193.85 0 0 1 37.28-37.22l54.76 61.53-35.83-74.17a191.49 191.49 0 0 1 48.65-20.13l26.87 77.25-4.71-81.61c8.61-1.18 17.39-1.8 26.32-1.8s17.71.62 26.32 1.8l-4.74 82.16 27.05-77.76c17.27 4.5 33.6 11.35 48.63 20.17l-35.82 74.12 54.72-61.47a193.13 193.13 0 0 1 37.24 37.23l-61.45 54.77 74.12-35.86a191.515 191.515 0 0 1 20.2 48.65l-77.81 27.1 82.24-4.75c1.19 8.66 1.82 17.5 1.82 26.49 0 8.88-.61 17.63-1.78 26.19l-82.12-4.75 77.72 27.09z"],firstdraft:[384,512,[],"f3a1","M384 192h-64v128H192v128H0v-25.6h166.4v-128h128v-128H384V192zm-25.6 38.4v128h-128v128H64V512h192V384h128V230.4h-25.6zm25.6 192h-89.6V512H320v-64h64v-25.6zM0 0v384h128V256h128V128h128V0H0z"],flickr:[448,512,[],"f16e","M400 32H48C21.5 32 0 53.5 0 80v352c0 26.5 21.5 48 48 48h352c26.5 0 48-21.5 48-48V80c0-26.5-21.5-48-48-48zM144.5 319c-35.1 0-63.5-28.4-63.5-63.5s28.4-63.5 63.5-63.5 63.5 28.4 63.5 63.5-28.4 63.5-63.5 63.5zm159 0c-35.1 0-63.5-28.4-63.5-63.5s28.4-63.5 63.5-63.5 63.5 28.4 63.5 63.5-28.4 63.5-63.5 63.5z"],flipboard:[448,512,[],"f44d","M0 32v448h448V32H0zm358.4 179.2h-89.6v89.6h-89.6v89.6H89.6V121.6h268.8v89.6z"],fly:[384,512,[],"f417","M197.8 427.8c12.9 11.7 33.7 33.3 33.2 50.7 0 .8-.1 1.6-.1 2.5-1.8 19.8-18.8 31.1-39.1 31-25-.1-39.9-16.8-38.7-35.8 1-16.2 20.5-36.7 32.4-47.6 2.3-2.1 2.7-2.7 5.6-3.6 3.4 0 3.9.3 6.7 2.8zM331.9 67.3c-16.3-25.7-38.6-40.6-63.3-52.1C243.1 4.5 214-.2 192 0c-44.1 0-71.2 13.2-81.1 17.3C57.3 45.2 26.5 87.2 28 158.6c7.1 82.2 97 176 155.8 233.8 1.7 1.6 4.5 4.5 6.2 5.1l3.3.1c2.1-.7 1.8-.5 3.5-2.1 52.3-49.2 140.7-145.8 155.9-215.7 7-39.2 3.1-72.5-20.8-112.5zM186.8 351.9c-28-51.1-65.2-130.7-69.3-189-3.4-47.5 11.4-131.2 69.3-136.7v325.7zM328.7 180c-16.4 56.8-77.3 128-118.9 170.3C237.6 298.4 275 217 277 158.4c1.6-45.9-9.8-105.8-48-131.4 88.8 18.3 115.5 98.1 99.7 153z"],"font-awesome":[448,512,[],"f2b4","M397.8 32H50.2C22.7 32 0 54.7 0 82.2v347.6C0 457.3 22.7 480 50.2 480h347.6c27.5 0 50.2-22.7 50.2-50.2V82.2c0-27.5-22.7-50.2-50.2-50.2zm-45.4 284.3c0 4.2-3.6 6-7.8 7.8-16.7 7.2-34.6 13.7-53.8 13.7-26.9 0-39.4-16.7-71.7-16.7-23.3 0-47.8 8.4-67.5 17.3-1.2.6-2.4.6-3.6 1.2V385c0 1.8 0 3.6-.6 4.8v1.2c-2.4 8.4-10.2 14.3-19.1 14.3-11.3 0-20.3-9-20.3-20.3V166.4c-7.8-6-13.1-15.5-13.1-26.3 0-18.5 14.9-33.5 33.5-33.5 18.5 0 33.5 14.9 33.5 33.5 0 10.8-4.8 20.3-13.1 26.3v18.5c1.8-.6 3.6-1.2 5.4-2.4 18.5-7.8 40.6-14.3 61.5-14.3 22.7 0 40.6 6 60.9 13.7 4.2 1.8 8.4 2.4 13.1 2.4 22.7 0 47.8-16.1 53.8-16.1 4.8 0 9 3.6 9 7.8v140.3z"],"font-awesome-alt":[448,512,[],"f35c","M397.8 67.8c7.8 0 14.3 6.6 14.3 14.3v347.6c0 7.8-6.6 14.3-14.3 14.3H50.2c-7.8 0-14.3-6.6-14.3-14.3V82.2c0-7.8 6.6-14.3 14.3-14.3h347.6m0-35.9H50.2C22.7 32 0 54.7 0 82.2v347.6C0 457.3 22.7 480 50.2 480h347.6c27.5 0 50.2-22.7 50.2-50.2V82.2c0-27.5-22.7-50.2-50.2-50.2zm-58.5 139.2c-6 0-29.9 15.5-52.6 15.5-4.2 0-8.4-.6-12.5-2.4-19.7-7.8-37-13.7-59.1-13.7-20.3 0-41.8 6.6-59.7 13.7-1.8.6-3.6 1.2-4.8 1.8v-17.9c7.8-6 12.5-14.9 12.5-25.7 0-17.9-14.3-32.3-32.3-32.3s-32.3 14.3-32.3 32.3c0 10.2 4.8 19.7 12.5 25.7v212.1c0 10.8 9 19.7 19.7 19.7 9 0 16.1-6 18.5-13.7V385c.6-1.8.6-3 .6-4.8V336c1.2 0 2.4-.6 3-1.2 19.7-8.4 43-16.7 65.7-16.7 31.1 0 43 16.1 69.3 16.1 18.5 0 36.4-6.6 52-13.7 4.2-1.8 7.2-3.6 7.2-7.8V178.3c1.8-4.1-2.3-7.1-7.7-7.1z"],"font-awesome-flag":[448,512,[],"f425","M444.373 359.424c0 7.168-6.144 10.24-13.312 13.312-28.672 12.288-59.392 23.552-92.16 23.552-46.08 0-67.584-28.672-122.88-28.672-39.936 0-81.92 14.336-115.712 29.696-2.048 1.024-4.096 1.024-6.144 2.048v77.824c0 21.405-16.122 34.816-33.792 34.816-19.456 0-34.816-15.36-34.816-34.816V102.4C12.245 92.16 3.029 75.776 3.029 57.344 3.029 25.6 28.629 0 60.373 0s57.344 25.6 57.344 57.344c0 18.432-8.192 34.816-22.528 45.056v31.744c4.124-1.374 58.768-28.672 114.688-28.672 65.27 0 97.676 27.648 126.976 27.648 38.912 0 81.92-27.648 92.16-27.648 8.192 0 15.36 6.144 15.36 13.312v240.64z"],"font-awesome-logo-full":[3992,512,["Font Awesome"],"f4e6","M454.6 0H57.4C25.9 0 0 25.9 0 57.4v397.3C0 486.1 25.9 512 57.4 512h397.3c31.4 0 57.4-25.9 57.4-57.4V57.4C512 25.9 486.1 0 454.6 0zm-58.9 324.9c0 4.8-4.1 6.9-8.9 8.9-19.2 8.1-39.7 15.7-61.5 15.7-40.5 0-68.7-44.8-163.2 2.5v51.8c0 30.3-45.7 30.2-45.7 0v-250c-9-7-15-17.9-15-30.3 0-21 17.1-38.2 38.2-38.2 21 0 38.2 17.1 38.2 38.2 0 12.2-5.8 23.2-14.9 30.2v21c37.1-12 65.5-34.4 146.1-3.4 26.6 11.4 68.7-15.7 76.5-15.7 5.5 0 10.3 4.1 10.3 8.9v160.4zm432.9-174.2h-137v70.1H825c39.8 0 40.4 62.2 0 62.2H691.6v105.6c0 45.5-70.7 46.4-70.7 0V128.3c0-22 18-39.8 39.8-39.8h167.8c39.6 0 40.5 62.2.1 62.2zm191.1 23.4c-169.3 0-169.1 252.4 0 252.4 169.9 0 169.9-252.4 0-252.4zm0 196.1c-81.6 0-82.1-139.8 0-139.8 82.5 0 82.4 139.8 0 139.8zm372.4 53.4c-17.5 0-31.4-13.9-31.4-31.4v-117c0-62.4-72.6-52.5-99.1-16.4v133.4c0 41.5-63.3 41.8-63.3 0V208c0-40 63.1-41.6 63.1 0v3.4c43.3-51.6 162.4-60.4 162.4 39.3v141.5c.3 30.4-31.5 31.4-31.7 31.4zm179.7 2.9c-44.3 0-68.3-22.9-68.3-65.8V235.2H1488c-35.6 0-36.7-55.3 0-55.3h15.5v-37.3c0-41.3 63.8-42.1 63.8 0v37.5h24.9c35.4 0 35.7 55.3 0 55.3h-24.9v108.5c0 29.6 26.1 26.3 27.4 26.3 31.4 0 52.6 56.3-22.9 56.3zM1992 123c-19.5-50.2-95.5-50-114.5 0-107.3 275.7-99.5 252.7-99.5 262.8 0 42.8 58.3 51.2 72.1 14.4l13.5-35.9H2006l13 35.9c14.2 37.7 72.1 27.2 72.1-14.4 0-10.1 5.3 6.8-99.1-262.8zm-108.9 179.1l51.7-142.9 51.8 142.9h-103.5zm591.3-85.6l-53.7 176.3c-12.4 41.2-72 41-84 0l-42.3-135.9-42.3 135.9c-12.4 40.9-72 41.2-84.5 0l-54.2-176.3c-12.5-39.4 49.8-56.1 60.2-16.9L2213 342l45.3-139.5c10.9-32.7 59.6-34.7 71.2 0l45.3 139.5 39.3-142.4c10.3-38.3 72.6-23.8 60.3 16.9zm275.4 75.1c0-42.4-33.9-117.5-119.5-117.5-73.2 0-124.4 56.3-124.4 126 0 77.2 55.3 126.4 128.5 126.4 31.7 0 93-11.5 93-39.8 0-18.3-21.1-31.5-39.3-22.4-49.4 26.2-109 8.4-115.9-43.8h148.3c16.3 0 29.3-13.4 29.3-28.9zM2571 277.7c9.5-73.4 113.9-68.6 118.6 0H2571zm316.7 148.8c-31.4 0-81.6-10.5-96.6-31.9-12.4-17 2.5-39.8 21.8-39.8 16.3 0 36.8 22.9 77.7 22.9 27.4 0 40.4-11 40.4-25.8 0-39.8-142.9-7.4-142.9-102 0-40.4 35.3-75.7 98.6-75.7 31.4 0 74.1 9.9 87.6 29.4 10.8 14.8-1.4 36.2-20.9 36.2-15.1 0-26.7-17.3-66.2-17.3-22.9 0-37.8 10.5-37.8 23.8 0 35.9 142.4 6 142.4 103.1-.1 43.7-37.4 77.1-104.1 77.1zm266.8-252.4c-169.3 0-169.1 252.4 0 252.4 170.1 0 169.6-252.4 0-252.4zm0 196.1c-81.8 0-82-139.8 0-139.8 82.5 0 82.4 139.8 0 139.8zm476.9 22V268.7c0-53.8-61.4-45.8-85.7-10.5v134c0 41.3-63.8 42.1-63.8 0V268.7c0-52.1-59.5-47.4-85.7-10.1v133.6c0 41.5-63.3 41.8-63.3 0V208c0-40 63.1-41.6 63.1 0v3.4c9.9-14.4 41.8-37.3 78.6-37.3 35.3 0 57.7 16.4 66.7 43.8 13.9-21.8 45.8-43.8 82.6-43.8 44.3 0 70.7 23.4 70.7 72.7v145.3c.5 17.3-13.5 31.4-31.9 31.4 3.5.1-31.3 1.1-31.3-31.3zM3992 291.6c0-42.4-32.4-117.5-117.9-117.5-73.2 0-127.5 56.3-127.5 126 0 77.2 58.3 126.4 131.6 126.4 31.7 0 91.5-11.5 91.5-39.8 0-18.3-21.1-31.5-39.3-22.4-49.4 26.2-110.5 8.4-117.5-43.8h149.8c16.3 0 29.1-13.4 29.3-28.9zm-180.5-13.9c9.7-74.4 115.9-68.3 120.1 0h-120.1z"],fonticons:[448,512,[],"f280","M0 32v448h448V32H0zm167.4 196h67.4l-11.1 37.3H168v112.9c0 5.8-2 6.7 3.2 7.3l43.5 4.1v25.1H84V389l21.3-2c5.2-.6 6.7-2.3 6.7-7.9V267.7c0-2.3-2.9-2.3-5.8-2.3H84V228h28v-21c0-49.6 26.5-70 77.3-70 34.1 0 64.7 8.2 64.7 52.8l-50.7 6.1c.3-18.7-4.4-23-16.3-23-18.4 0-19 9.9-19 27.4v23.3c0 2.4-3.5 4.4-.6 4.4zM364 414.7H261.3v-25.1l20.4-2.6c5.2-.6 7.6-1.7 7.6-7.3V271.8c0-4.1-2.9-6.7-6.7-7.9l-24.2-6.4 6.7-29.5h80.2v151.7c0 5.8-2.6 6.4 2.9 7.3l15.7 2.6v25.1zm-21.9-255.5l9 33.2-7.3 7.3-31.2-16.6-31.2 16.6-7.3-7.3 9-33.2-21.8-24.2 3.5-9.6h27.7l15.5-28h9.3l15.5 28h27.7l3.5 9.6-21.9 24.2z"],"fonticons-fi":[384,512,[],"f3a2","M114.4 224h92.4l-15.2 51.2h-76.4V433c0 8-2.8 9.2 4.4 10l59.6 5.6V483H0v-35.2l29.2-2.8c7.2-.8 9.2-3.2 9.2-10.8V278.4c0-3.2-4-3.2-8-3.2H0V224h38.4v-28.8c0-68 36.4-96 106-96 46.8 0 88.8 11.2 88.8 72.4l-69.6 8.4c.4-25.6-6-31.6-22.4-31.6-25.2 0-26 13.6-26 37.6v32c0 3.2-4.8 6-.8 6zM384 483H243.2v-34.4l28-3.6c7.2-.8 10.4-2.4 10.4-10V287c0-5.6-4-9.2-9.2-10.8l-33.2-8.8 9.2-40.4h110v208c0 8-3.6 8.8 4 10l21.6 3.6V483zm-30-347.2l12.4 45.6-10 10-42.8-22.8-42.8 22.8-10-10 12.4-45.6-30-36.4 4.8-10h38L307.2 51H320l21.2 38.4h38l4.8 13.2-30 33.2z"],"fort-awesome":[512,512,[],"f286","M489.2 287.9h-27.4c-2.6 0-4.6 2-4.6 4.6v32h-36.6V146.2c0-2.6-2-4.6-4.6-4.6h-27.4c-2.6 0-4.6 2-4.6 4.6v32h-36.6v-32c0-2.6-2-4.6-4.6-4.6h-27.4c-2.6 0-4.6 2-4.6 4.6v32h-36.6v-32c0-6-8-4.6-11.7-4.6v-38c8.3-2 17.1-3.4 25.7-3.4 10.9 0 20.9 4.3 31.4 4.3 4.6 0 27.7-1.1 27.7-8v-60c0-2.6-2-4.6-4.6-4.6-5.1 0-15.1 4.3-24 4.3-9.7 0-20.9-4.3-32.6-4.3-8 0-16 1.1-23.7 2.9v-4.9c5.4-2.6 9.1-8.3 9.1-14.3 0-20.7-31.4-20.8-31.4 0 0 6 3.7 11.7 9.1 14.3v111.7c-3.7 0-11.7-1.4-11.7 4.6v32h-36.6v-32c0-2.6-2-4.6-4.6-4.6h-27.4c-2.6 0-4.6 2-4.6 4.6v32H128v-32c0-2.6-2-4.6-4.6-4.6H96c-2.6 0-4.6 2-4.6 4.6v178.3H54.8v-32c0-2.6-2-4.6-4.6-4.6H22.8c-2.6 0-4.6 2-4.6 4.6V512h182.9v-96c0-72.6 109.7-72.6 109.7 0v96h182.9V292.5c.1-2.6-1.9-4.6-4.5-4.6zm-288.1-4.5c0 2.6-2 4.6-4.6 4.6h-27.4c-2.6 0-4.6-2-4.6-4.6v-64c0-2.6 2-4.6 4.6-4.6h27.4c2.6 0 4.6 2 4.6 4.6v64zm146.4 0c0 2.6-2 4.6-4.6 4.6h-27.4c-2.6 0-4.6-2-4.6-4.6v-64c0-2.6 2-4.6 4.6-4.6h27.4c2.6 0 4.6 2 4.6 4.6v64z"],"fort-awesome-alt":[512,512,[],"f3a3","M211.7 241.1v51.7c0 2.1-1.6 3.7-3.7 3.7h-22.2c-2.1 0-3.7-1.6-3.7-3.7v-51.7c0-2.1 1.6-3.7 3.7-3.7H208c2.1 0 3.7 1.6 3.7 3.7zm114.5-3.7H304c-2.1 0-3.7 1.6-3.7 3.7v51.7c0 2.1 1.6 3.7 3.7 3.7h22.2c2.1 0 3.7-1.6 3.7-3.7v-51.7c-.1-2.1-1.7-3.7-3.7-3.7zm-29.1 263.2c-.9.1-1.7.3-2.6.4-1 .2-2.1.3-3.1.5-.9.1-1.8.3-2.8.4-1 .1-2 .3-3 .4-1 .1-2 .2-2.9.3-1 .1-1.9.2-2.9.3-1 .1-2.1.2-3.1.3-.9.1-1.8.2-2.7.2-1.1.1-2.3.1-3.4.2-.8 0-1.7.1-2.5.1-1.3.1-2.6.1-3.9.1-.7 0-1.4.1-2.1.1-2 0-4 .1-6 .1s-4 0-6-.1c-.7 0-1.4 0-2.1-.1-1.3 0-2.6-.1-3.9-.1-.8 0-1.7-.1-2.5-.1-1.1-.1-2.3-.1-3.4-.2-.9-.1-1.8-.1-2.7-.2-1-.1-2.1-.2-3.1-.3-1-.1-1.9-.2-2.9-.3-1-.1-2-.2-2.9-.3-1-.1-2-.2-3-.4-.9-.1-1.8-.3-2.8-.4-1-.1-2.1-.3-3.1-.5-.9-.1-1.7-.3-2.6-.4-65.6-10.9-122.5-47.7-160-99.4-.2-.2-.3-.5-.5-.7-.8-1.1-1.6-2.2-2.3-3.3-.3-.4-.6-.8-.8-1.2-.7-1.1-1.4-2.1-2.1-3.2-.3-.5-.6-.9-.9-1.4-.7-1.1-1.4-2.1-2-3.2-.3-.5-.6-.9-.9-1.4-.7-1.1-1.3-2.2-2-3.3-.2-.4-.5-.8-.7-1.2-2.4-4-4.6-8.1-6.8-12.2-.1-.2-.2-.3-.3-.5-.6-1.1-1.1-2.2-1.7-3.3-.3-.6-.6-1.1-.8-1.7-.5-1-1-2.1-1.5-3.1-.3-.7-.6-1.3-.9-2-.5-1-.9-2-1.4-3l-.9-2.1c-.4-1-.9-2-1.3-3-.3-.7-.6-1.5-.9-2.2l-1.2-3c-.3-.8-.6-1.5-.9-2.3-.4-1-.8-2-1.1-3-.3-.9-.6-1.8-1-2.8-.6-1.6-1.1-3.3-1.7-4.9-.3-.9-.6-1.8-.9-2.8-.3-.9-.5-1.8-.8-2.7-.3-.9-.6-1.9-.8-2.8-.3-.9-.5-1.8-.8-2.7-.3-1-.5-1.9-.8-2.9-.2-.9-.5-1.8-.7-2.7-.3-1-.5-2-.7-3-.2-.9-.4-1.7-.6-2.6-.2-1.1-.5-2.2-.7-3.2-.2-.8-.3-1.6-.5-2.4-.3-1.3-.5-2.7-.8-4-.1-.6-.2-1.1-.3-1.7l-.9-5.7c-.1-.6-.2-1.3-.3-1.9-.2-1.3-.4-2.6-.5-3.9-.1-.8-.2-1.5-.3-2.3-.1-1.2-.3-2.4-.4-3.6-.1-.8-.2-1.6-.2-2.4-.1-1.2-.2-2.4-.3-3.5-.1-.8-.1-1.6-.2-2.4-.1-1.2-.2-2.4-.2-3.7 0-.8-.1-1.5-.1-2.3-.1-1.3-.1-2.7-.2-4 0-.7 0-1.3-.1-2 0-2-.1-4-.1-6 0-53.5 16.9-103 45.8-143.6 2.3-3.2 4.7-6.4 7.1-9.5 4.9-6.2 10.1-12.3 15.6-18 2.7-2.9 5.5-5.7 8.4-8.4 2.9-2.7 5.8-5.4 8.8-8 4.5-3.9 9.1-7.6 13.9-11.2 1.6-1.2 3.2-2.4 4.8-3.5C140 34.2 171.7 20.1 206 13c16.1-3.3 32.9-5 50-5s33.8 1.7 50 5c34.3 7 66 21.1 93.6 40.7 1.6 1.2 3.2 2.3 4.8 3.5 4.8 3.6 9.4 7.3 13.9 11.2 12 10.4 23 21.9 32.8 34.4 2.5 3.1 4.8 6.3 7.1 9.5C487.1 153 504 202.5 504 256c0 2 0 4-.1 6 0 .7 0 1.3-.1 2 0 1.3-.1 2.7-.2 4 0 .8-.1 1.5-.1 2.3-.1 1.2-.1 2.4-.2.7-.1.8-.1 1.6-.2 2.4-.1 1.2-.2 2.4-.3 3.5-.1.8-.2 1.6-.2 2.4-.1 1.2-.3 2.4-.4 3.6-.1.8-.2 1.5-.3 2.3-.2 1.3-.4 2.6-.5 3.9-.1.6-.2 1.3-.3 1.9l-.9 5.7c-.1.6-.2 1.1-.3 1.7-.2 1.3-.5 2.7-.8 4-.2.8-.3 1.6-.5 2.4-.2 1.1-.5 2.2-.7 3.2-.2.9-.4 1.7-.6 2.6-.2 1-.5 2-.7 3-.2.9-.5 1.8-.7 2.7-.3 1-.5 1.9-.8 2.9-.2.9-.5 1.8-.8 2.7-.3.9-.6 1.9-.8 2.8-.3.9-.5 1.8-.8 2.7-.3.9-.6 1.8-.9 2.8-.5 1.6-1.1 3.3-1.7 4.9-.3.9-.6 1.8-1 2.8-.4 1-.7 2-1.1 3-.3.8-.6 1.5-.9 2.3l-1.2 3c-.3.7-.6 1.5-.9 2.2-.4 1-.8 2-1.3 3l-.9 2.1c-.4 1-.9 2-1.4 3-.3.7-.6 1.3-.9 2-.5 1-1 2.1-1.5 3.1-.3.6-.6 1.1-.8 1.7-.6 1.1-1.1 2.2-1.7 3.3-.1.2-.2.3-.3.5-2.2 4.1-4.4 8.2-6.8 12.2-.2.4-.5.8-.7 1.2-.7 1.1-1.3 2.2-2 3.3-.3.5-.6.9-.9 1.4-.7 1.1-1.4 2.1-2 3.2-.3.5-.6.9-.9 1.4-.7 1.1-1.4 2.1-2.1 3.2-.3.4-.6.8-.8 1.2-.8 1.1-1.5 2.2-2.3 3.3-.2.2-.3.5-.5.7-37.6 54.7-94.5 91.4-160.1 102.4zm117.3-86.2c13-13 24.2-27.4 33.6-42.9v-71.3c0-2.1-1.6-3.7-3.7-3.7h-22.2c-2.1 0-3.7 1.6-3.7 3.7V326h-29.5V182c0-2.1-1.6-3.7-3.7-3.7h-22.1c-2.1 0-3.7 1.6-3.7 3.7v25.9h-29.5V182c0-2.1-1.6-3.7-3.7-3.7H304c-2.1 0-3.7 1.6-3.7 3.7v25.9h-29.5V182c0-4.8-6.5-3.7-9.5-3.7v-30.7c6.7-1.6 13.8-2.8 20.8-2.8 8.8 0 16.8 3.5 25.4 3.5 3.7 0 22.4-.9 22.4-6.5V93.4c0-2.1-1.6-3.7-3.7-3.7-4.2 0-12.2 3.5-19.4 3.5-7.9 0-16.9-3.5-26.3-3.5-6.5 0-12.9.9-19.2 2.3v-3.9c4.4-2.1 7.4-6.7 7.4-11.5 0-16.8-25.4-16.8-25.4 0 0 4.8 3 9.5 7.4 11.5v90.2c-3 0-9.5-1.1-9.5 3.7v25.9h-29.5V182c0-2.1-1.6-3.7-3.7-3.7h-22.2c-2.1 0-3.7 1.6-3.7 3.7v25.9h-29.5V182c0-2.1-1.6-3.7-3.7-3.7h-22.1c-2.1 0-3.7 1.6-3.7 3.7v144H93.5v-25.8c0-2.1-1.6-3.7-3.7-3.7H67.7c-2.1 0-3.7 1.6-3.7 3.7v71.3c9.4 15.5 20.6 29.9 33.6 42.9 20.6 20.6 44.5 36.7 71.2 48 13.9 5.9 28.2 10.3 42.9 13.2v-75.8c0-58.6 88.6-58.6 88.6 0v75.8c14.7-2.9 29-7.4 42.9-13.2 26.7-11.3 50.6-27.4 71.2-48"],forumbee:[448,512,[],"f211","M5.8 309.7C2 292.7 0 275.5 0 258.3 0 135 99.8 35 223.1 35c16.6 0 33.3 2 49.3 5.5C149 87.5 51.9 186 5.8 309.7zm392.9-189.2C385 103 369 87.8 350.9 75.2c-149.6 44.3-266.3 162.1-309.7 312 12.5 18.1 28 35.6 45.2 49 43.1-151.3 161.2-271.7 312.3-315.7zm15.8 252.7c15.2-25.1 25.4-53.7 29.5-82.8-79.4 42.9-145 110.6-187.6 190.3 30-4.4 58.9-15.3 84.6-31.3 35 13.1 70.9 24.3 107 33.6-9.3-36.5-20.4-74.5-33.5-109.8zm29.7-145.5c-2.6-19.5-7.9-38.7-15.8-56.8C290.5 216.7 182 327.5 137.1 466c18.1 7.6 37 12.5 56.6 15.2C240 367.1 330.5 274.4 444.2 227.7z"],foursquare:[368,512,[],"f180","M323.1 3H49.9C12.4 3 0 31.3 0 49.1v433.8c0 20.3 12.1 27.7 18.2 30.1 6.2 2.5 22.8 4.6 32.9-7.1C180 356.5 182.2 354 182.2 354c3.1-3.4 3.4-3.1 6.8-3.1h83.4c35.1 0 40.6-25.2 44.3-39.7l48.6-243C373.8 25.8 363.1 3 323.1 3zm-16.3 73.8l-11.4 59.7c-1.2 6.5-9.5 13.2-16.9 13.2H172.1c-12 0-20.6 8.3-20.6 20.3v13c0 12 8.6 20.6 20.6 20.6h90.4c8.3 0 16.6 9.2 14.8 18.2-1.8 8.9-10.5 53.8-11.4 58.8-.9 4.9-6.8 13.5-16.9 13.5h-73.5c-13.5 0-17.2 1.8-26.5 12.6 0 0-8.9 11.4-89.5 108.3-.9.9-1.8.6-1.8-.3V75.9c0-7.7 6.8-16.6 16.6-16.6h219c8.2 0 15.6 7.7 13.5 17.5z"],"free-code-camp":[576,512,[],"f2c5","M69.3 144.5c-41 68.5-36.4 163 1 227C92.5 409.7 120 423.9 120 438c0 6.8-6 13-12.8 13C87.7 451 8 375.5 8 253.2c0-111.5 78-186 97.1-186 6 0 14.9 4.8 14.9 11.1 0 12.7-28.3 28.6-50.7 66.2zm195.8 213.8c4.5 1.8 12.3 5.2 12.3-1.2 0-2.7-2.2-2.9-4.3-3.6-8.5-3.4-14-7.7-19.1-15.2-8.2-12.1-10.1-24.2-10.1-38.6 0-32.1 44.2-37.9 44.2-70 0-12.3-7.7-15.9-7.7-19.3 0-2.2.7-2.2 2.9-2.2 8 0 19.1 13.3 22.5 19.8 2.2 4.6 2.4 6 2.4 11.1 0 7-.7 14.2-.7 21.3 0 27 31.9 19.8 31.9 6.8 0-6-3.6-11.6-3.6-17.4 0-.7 0-1.2.7-1.2 3.4 0 9.4 7.7 11.1 10.1 5.8 8.9 8.5 20.8 8.5 31.4 0 32.4-29.5 49-29.5 56 0 1 2.9 7.7 12.1 1.9 29.7-15.1 53.1-47.6 53.1-89.8 0-33.6-8.7-57.7-32.1-82.6-3.9-4.1-16.4-16.9-22.5-16.9-8.2 0 7.2 18.6 7.2 31.2 0 7.2-4.8 12.3-12.3 12.3-11.6 0-14.5-25.4-15.9-33.3-5.8-33.8-12.8-58.2-46.4-74.1-10.4-5-36.5-11.8-36.5-2.2 0 2.4 2.7 4.1 4.6 5.1 9.2 5.6 19.6 21.4 19.6 38.2 0 46.1-57.7 88.2-57.7 136.2-.2 40.3 28.1 72.6 65.3 86.2zM470.4 67c-6 0-14.4 6.5-14.4 12.6 0 8.7 12.1 19.6 17.6 25.4 81.6 85.1 78.6 214.3 17.6 291-7 8.9-35.3 35.3-35.3 43.5 0 5.1 8.2 11.4 13.2 11.4 25.4 0 98.8-80.8 98.8-185.7C568 145.9 491.8 67 470.4 67zm-42.3 323.1H167c-9.4 0-15.5 7.5-15.5 16.4 0 8.5 7 15.5 15.5 15.5h261.1c9.4 0 11.9-7.5 11.9-16.4 0-8.5-3.5-15.5-11.9-15.5z"],freebsd:[448,512,[],"f3a4","M303.7 96.2c11.1-11.1 115.5-77 139.2-53.2 23.7 23.7-42.1 128.1-53.2 139.2-11.1 11.1-39.4.9-63.1-22.9-23.8-23.7-34.1-52-22.9-63.1zM109.9 68.1C73.6 47.5 22 24.6 5.6 41.1c-16.6 16.6 7.1 69.4 27.9 105.7 18.5-32.2 44.8-59.3 76.4-78.7zM406.7 174c3.3 11.3 2.7 20.7-2.7 26.1-20.3 20.3-87.5-27-109.3-70.1-18-32.3-11.1-53.4 14.9-48.7 5.7-3.6 12.3-7.6 19.6-11.6-29.8-15.5-63.6-24.3-99.5-24.3-119.1 0-215.6 96.5-215.6 215.6 0 119 96.5 215.6 215.6 215.6S445.3 380.1 445.3 261c0-38.4-10.1-74.5-27.7-105.8-3.9 7-7.6 13.3-10.9 18.8z"],fulcrum:[269,512,[],"f50b","M70.75 164.14l-35.38 43.55L0 164.14l35.38-43.55 35.37 43.55zM119.23 0L98.69 198.18 47.72 256l50.98 57.82L119.23 512V300.89L78.15 256l41.08-44.89V0zm79.67 164.14l35.38 43.55 35.38-43.55-35.38-43.55-35.38 43.55zm-48.48 46.97L191.5 256l-41.08 44.89V512l20.54-198.18L221.94 256l-50.98-57.82L150.42 0v211.11z"],"galactic-republic":[496,512,[],"f50c","M248 504C111.25 504 0 392.75 0 256S111.25 8 248 8s248 111.25 248 248-111.25 248-248 248zm0-479.47C120.37 24.53 16.53 128.37 16.53 256S120.37 487.47 248 487.47 479.47 383.63 479.47 256 375.63 24.53 248 24.53zm27.62 21.81v24.62a185.933 185.933 0 0 1 83.57 34.54l17.39-17.36c-28.75-22.06-63.3-36.89-100.96-41.8zm-55.37.07c-37.64 4.94-72.16 19.8-100.88 41.85l17.28 17.36h.08c24.07-17.84 52.55-30.06 83.52-34.67V46.41zm12.25 50.17v82.87c-10.04 2.03-19.42 5.94-27.67 11.42l-58.62-58.59-21.93 21.93 58.67 58.67c-5.47 8.23-9.45 17.59-11.47 27.62h-82.9v31h82.9c2.02 10.02 6.01 19.31 11.47 27.54l-58.67 58.69 21.93 21.93 58.62-58.62a77.873 77.873 0 0 0 27.67 11.47v82.9h31v-82.9c10.05-2.03 19.37-6.06 27.62-11.55l58.67 58.69 21.93-21.93-58.67-58.69c5.46-8.23 9.47-17.52 11.5-27.54h82.87v-31h-82.87c-2.02-10.02-6.03-19.38-11.5-27.62l58.67-58.67-21.93-21.93-58.67 58.67c-8.25-5.49-17.57-9.47-27.62-11.5V96.58h-31zm183.24 30.72l-17.36 17.36a186.337 186.337 0 0 1 34.67 83.67h24.62c-4.95-37.69-19.83-72.29-41.93-101.03zm-335.55.13c-22.06 28.72-36.91 63.26-41.85 100.91h24.65c4.6-30.96 16.76-59.45 34.59-83.52l-17.39-17.39zM38.34 283.67c4.92 37.64 19.75 72.18 41.8 100.9l17.36-17.39c-17.81-24.07-29.92-52.57-34.51-83.52H38.34zm394.7 0c-4.61 30.99-16.8 59.5-34.67 83.6l17.36 17.36c22.08-28.74 36.98-63.29 41.93-100.96h-24.62zM136.66 406.38l-17.36 17.36c28.73 22.09 63.3 36.98 100.96 41.93v-24.64c-30.99-4.63-59.53-16.79-83.6-34.65zm222.53.05c-24.09 17.84-52.58 30.08-83.57 34.67v24.57c37.67-4.92 72.21-19.79 100.96-41.85l-17.31-17.39h-.08z"],"galactic-senate":[512,512,[],"f50d","M249.86 33.48v26.07C236.28 80.17 226 168.14 225.39 274.9c11.74-15.62 19.13-33.33 19.13-48.24v-16.88c-.03-5.32.75-10.53 2.19-15.65.65-2.14 1.39-4.08 2.62-5.82 1.23-1.75 3.43-3.79 6.68-3.79 3.24 0 5.45 2.05 6.68 3.79 1.23 1.75 1.97 3.68 2.62 5.82 1.44 5.12 2.22 10.33 2.19 15.65v16.88c0 14.91 7.39 32.62 19.13 48.24-.63-106.76-10.91-194.73-24.49-215.35V33.48h-12.28zm-26.34 147.77c-9.52 2.15-18.7 5.19-27.46 9.08 8.9 16.12 9.76 32.64 1.71 37.29-8 4.62-21.85-4.23-31.36-19.82-11.58 8.79-21.88 19.32-30.56 31.09 14.73 9.62 22.89 22.92 18.32 30.66-4.54 7.7-20.03 7.14-35.47-.96-5.78 13.25-9.75 27.51-11.65 42.42 9.68.18 18.67 2.38 26.18 6.04 17.78-.3 32.77-1.96 40.49-4.22 5.55-26.35 23.02-48.23 46.32-59.51.73-25.55 1.88-49.67 3.48-72.07zm64.96 0c1.59 22.4 2.75 46.52 3.47 72.07 23.29 11.28 40.77 33.16 46.32 59.51 7.72 2.26 22.71 3.92 40.49 4.22 7.51-3.66 16.5-5.85 26.18-6.04-1.9-14.91-5.86-29.17-11.65-42.42-15.44 8.1-30.93 8.66-35.47.96-4.57-7.74 3.6-21.05 18.32-30.66-8.68-11.77-18.98-22.3-30.56-31.09-9.51 15.59-23.36 24.44-31.36 19.82-8.05-4.65-7.19-21.16 1.71-37.29a147.49 147.49 0 0 0-27.45-9.08zm-32.48 8.6c-3.23 0-5.86 8.81-6.09 19.93h-.05v16.88c0 41.42-49.01 95.04-93.49 95.04-52 0-122.75-1.45-156.37 29.17v2.51c9.42 17.12 20.58 33.17 33.18 47.97C45.7 380.26 84.77 360.4 141.2 360c45.68 1.02 79.03 20.33 90.76 40.87.01.01-.01.04 0 .05 7.67 2.14 15.85 3.23 24.04 3.21 8.19.02 16.37-1.07 24.04-3.21.01-.01-.01-.04 0-.05 11.74-20.54 45.08-39.85 90.76-40.87 56.43.39 95.49 20.26 108.02 41.35 12.6-14.8 23.76-30.86 33.18-47.97v-2.51c-33.61-30.62-104.37-29.17-156.37-29.17-44.48 0-93.49-53.62-93.49-95.04v-16.88h-.05c-.23-11.12-2.86-19.93-6.09-19.93zm0 96.59c22.42 0 40.6 18.18 40.6 40.6s-18.18 40.65-40.6 40.65-40.6-18.23-40.6-40.65c0-22.42 18.18-40.6 40.6-40.6zm0 7.64c-18.19 0-32.96 14.77-32.96 32.96S237.81 360 256 360s32.96-14.77 32.96-32.96-14.77-32.96-32.96-32.96zm0 6.14c14.81 0 26.82 12.01 26.82 26.82s-12.01 26.82-26.82 26.82-26.82-12.01-26.82-26.82 12.01-26.82 26.82-26.82zm-114.8 66.67c-10.19.07-21.6.36-30.5 1.66.43 4.42 1.51 18.63 7.11 29.76 9.11-2.56 18.36-3.9 27.62-3.9 41.28.94 71.48 34.35 78.26 74.47l.11 4.7c10.4 1.91 21.19 2.94 32.21 2.94 11.03 0 21.81-1.02 32.21-2.94l.11-4.7c6.78-40.12 36.98-73.53 78.26-74.47 9.26 0 18.51 1.34 27.62 3.9 5.6-11.13 6.68-25.34 7.11-29.76-8.9-1.3-20.32-1.58-30.5-1.66-18.76.42-35.19 4.17-48.61 9.67-12.54 16.03-29.16 30.03-49.58 33.07-.09.02-.17.04-.27.05-.05.01-.11.04-.16.05-5.24 1.07-10.63 1.6-16.19 1.6-5.55 0-10.95-.53-16.19-1.6-.05-.01-.11-.04-.16-.05-.1-.02-.17-.04-.27-.05-20.42-3.03-37.03-17.04-49.58-33.07-13.42-5.49-29.86-9.25-48.61-9.67z"],"get-pocket":[448,512,[],"f265","M407.6 64h-367C18.5 64 0 82.5 0 104.6v135.2C0 364.5 99.7 464 224.2 464c124 0 223.8-99.5 223.8-224.2V104.6c0-22.4-17.7-40.6-40.4-40.6zm-162 268.5c-12.4 11.8-31.4 11.1-42.4 0C89.5 223.6 88.3 227.4 88.3 209.3c0-16.9 13.8-30.7 30.7-30.7 17 0 16.1 3.8 105.2 89.3 90.6-86.9 88.6-89.3 105.5-89.3 16.9 0 30.7 13.8 30.7 30.7 0 17.8-2.9 15.7-114.8 123.2z"],gg:[512,512,[],"f260","M179.2 230.4l102.4 102.4-102.4 102.4L0 256 179.2 76.8l44.8 44.8-25.6 25.6-19.2-19.2-128 128 128 128 51.5-51.5-77.1-76.5 25.6-25.6zM332.8 76.8L230.4 179.2l102.4 102.4 25.6-25.6-77.1-76.5 51.5-51.5 128 128-128 128-19.2-19.2-25.6 25.6 44.8 44.8L512 256 332.8 76.8z"],"gg-circle":[512,512,[],"f261","M257 8C120 8 9 119 9 256s111 248 248 248 248-111 248-248S394 8 257 8zm-49.5 374.8L81.8 257.1l125.7-125.7 35.2 35.4-24.2 24.2-11.1-11.1-77.2 77.2 77.2 77.2 26.6-26.6-53.1-52.9 24.4-24.4 77.2 77.2-75 75.2zm99-2.2l-35.2-35.2 24.1-24.4 11.1 11.1 77.2-77.2-77.2-77.2-26.5 26.5 53.1 52.9-24.4 24.4-77.2-77.2 75-75L432.2 255 306.5 380.6z"],git:[448,512,[],"f1d3","M18.8 221.7c0 25.3 16.2 60 41.5 68.5v1c-18.8 8.3-24 50.6 1 65.8v1C34 367 16 384.3 16 414.2c0 51.5 48.8 65.8 91.5 65.8 52 0 90.7-18.7 90.7-76 0-70.5-101-44.5-101-82.8 0-13.5 7.2-18.7 19.7-21.3 41.5-7.7 67.5-40 67.5-82.2 0-7.3-1.5-14.2-4-21 6.7-1.5 13.2-3.3 19.7-5.5v-50.5c-17.2 6.8-35.7 11.8-54.5 11.8-53.8-31-126.8 1.3-126.8 69.2zm87.7 163.8c17 0 41.2 3 41.2 25 0 21.8-19.5 26.3-37.7 26.3-17.3 0-43.3-2.7-43.3-25.2.1-22.3 22.1-26.1 39.8-26.1zM103.3 256c-22 0-31.3-13-31.3-33.8 0-49.3 61-48.8 61-.5 0 20.3-8 34.3-29.7 34.3zM432 305.5v49c-13.3 7.3-30.5 9.8-45.5 9.8-53.5 0-59.8-42.2-59.8-85.7v-87.7h.5v-1c-7 0-7.3-1.6-24 1v-47.5h24c0-22.3.3-31-1.5-41.2h56.7c-2 13.8-1.5 27.5-1.5 41.2h51v47.5s-19.3-1-51-1V281c0 14.8 3.3 32.8 21.8 32.8 9.8 0 21.3-2.8 29.3-8.3zM286 68.7c0 18.7-14.5 36.2-33.8 36.2-19.8 0-34.5-17.2-34.5-36.2 0-19.3 14.5-36.7 34.5-36.7C272 32 286 50 286 68.7zm-6.2 74.5c-1.8 14.6-1.6 199.8 0 217.8h-55.5c1.6-18.1 1.8-203 0-217.8h55.5z"],"git-square":[448,512,[],"f1d2","M140.1 348.5c12.1 0 29.5 2.1 29.5 17.9 0 15.5-13.9 18.8-27 18.8-12.3 0-30.9-2-30.9-18s15.7-18.7 28.4-18.7zm-24.7-116.6c0 14.8 6.6 24.1 22.3 24.1 15.5 0 21.2-10 21.2-24.5.1-34.4-43.5-34.8-43.5.4zM448 80v352c0 26.5-21.5 48-48 48H48c-26.5 0-48-21.5-48-48V80c0-26.5 21.5-48 48-48h352c26.5 0 48 21.5 48 48zm-241 93.7c-12.3 4.8-25.5 8.4-38.9 8.4-38.5-22.1-90.7.9-90.7 49.5 0 18 11.6 42.9 29.6 48.9v.7c-13.4 5.9-17.1 36.1.7 47v.7c-19.5 6.4-32.3 18.8-32.3 40.2 0 36.8 34.8 47 65.4 47 37.1 0 64.8-13.4 64.8-54.3 0-50.4-72.1-31.8-72.1-59.1 0-9.6 5.2-13.4 14.1-15.2 29.6-5.5 48.2-28.6 48.2-58.7 0-5.2-1.1-10.2-2.9-15 4.8-1.1 9.5-2.3 14.1-3.9v-36.2zm56.8 1.8h-39.6c1.3 10.6 1.1 142.6 0 155.5h39.6c-1.1-12.8-1.2-145.1 0-155.5zm4.5-53.3c0-13.4-10-26.2-24.1-26.2-14.3 0-24.6 12.5-24.6 26.2 0 13.6 10.5 25.9 24.6 25.9 13.7 0 24.1-12.5 24.1-25.9zm104.3 53.3h-36.4c0-9.8-.4-19.6 1.1-29.5h-40.5c1.3 7.3 1.1 13.6 1.1 29.5h-17.1v33.9c11.9-1.9 12.1-.7 17.1-.7v.7h-.4v62.7c0 31.1 4.5 61.2 42.7 61.2 10.7 0 23-1.8 32.5-7v-35c-5.7 3.9-13.9 5.9-20.9 5.9-13.2 0-15.5-12.9-15.5-23.4v-65.2c22.7 0 36.4.7 36.4.7v-33.8z"],github:[496,512,[],"f09b","M165.9 397.4c0 2-2.3 3.6-5.2 3.6-3.3.3-5.6-1.3-5.6-3.6 0-2 2.3-3.6 5.2-3.6 3-.3 5.6 1.3 5.6 3.6zm-31.1-4.5c-.7 2 1.3 4.3 4.3 4.9 2.6 1 5.6 0 6.2-2s-1.3-4.3-4.3-5.2c-2.6-.7-5.5.3-6.2 2.3zm44.2-1.7c-2.9.7-4.9 2.6-4.6 4.9.3 2 2.9 3.3 5.9 2.6 2.9-.7 4.9-2.6 4.6-4.6-.3-1.9-3-3.2-5.9-2.9zM244.8 8C106.1 8 0 113.3 0 252c0 110.9 69.8 205.8 169.5 239.2 12.8 2.3 17.3-5.6 17.3-12.1 0-6.2-.3-40.4-.3-61.4 0 0-70 15-84.7-29.8 0 0-11.4-29.1-27.8-36.6 0 0-22.9-15.7 1.6-15.4 0 0 24.9 2 38.6 25.8 21.9 38.6 58.6 27.5 72.9 20.9 2.3-16 8.8-27.1 16-33.7-55.9-6.2-112.3-14.3-112.3-110.5 0-27.5 7.6-41.3 23.6-58.9-2.6-6.5-11.1-33.3 2.6-67.9 20.9-6.5 69 27 69 27 20-5.6 41.5-8.5 62.8-8.5s42.8 2.9 62.8 8.5c0 0 48.1-33.6 69-27 13.7 34.7 5.2 61.4 2.6 67.9 16 17.7 25.8 31.5 25.8 58.9 0 96.5-58.9 104.2-114.8 110.5 9.2 7.9 17 22.9 17 46.4 0 33.7-.3 75.4-.3 83.6 0 6.5 4.6 14.4 17.3 12.1C428.2 457.8 496 362.9 496 252 496 113.3 383.5 8 244.8 8zM97.2 352.9c-1.3 1-1 3.3.7 5.2 1.6 1.6 3.9 2.3 5.2 1 1.3-1 1-3.3-.7-5.2-1.6-1.6-3.9-2.3-5.2-1zm-10.8-8.1c-.7 1.3.3 2.9 2.3 3.9 1.6 1 3.6.7 4.3-.7.7-1.3-.3-2.9-2.3-3.9-2-.6-3.6-.3-4.3.7zm32.4 35.6c-1.6 1.3-1 4.3 1.3 6.2 2.3 2.3 5.2 2.6 6.5 1 1.3-1.3.7-4.3-1.3-6.2-2.2-2.3-5.2-2.6-6.5-1zm-11.4-14.7c-1.6 1-1.6 3.6 0 5.9 1.6 2.3 4.3 3.3 5.6 2.3 1.6-1.3 1.6-3.9 0-6.2-1.4-2.3-4-3.3-5.6-2z"],"github-alt":[480,512,[],"f113","M186.1 328.7c0 20.9-10.9 55.1-36.7 55.1s-36.7-34.2-36.7-55.1 10.9-55.1 36.7-55.1 36.7 34.2 36.7 55.1zM480 278.2c0 31.9-3.2 65.7-17.5 95-37.9 76.6-142.1 74.8-216.7 74.8-75.8 0-186.2 2.7-225.6-74.8-14.6-29-20.2-63.1-20.2-95 0-41.9 13.9-81.5 41.5-113.6-5.2-15.8-7.7-32.4-7.7-48.8 0-21.5 4.9-32.3 14.6-51.8 45.3 0 74.3 9 108.8 36 29-6.9 58.8-10 88.7-10 27 0 54.2 2.9 80.4 9.2 34-26.7 63-35.2 107.8-35.2 9.8 19.5 14.6 30.3 14.6 51.8 0 16.4-2.6 32.7-7.7 48.2 27.5 32.4 39 72.3 39 114.2zm-64.3 50.5c0-43.9-26.7-82.6-73.5-82.6-18.9 0-37 3.4-56 6-14.9 2.3-29.8 3.2-45.1 3.2-15.2 0-30.1-.9-45.1-3.2-18.7-2.6-37-6-56-6-46.8 0-73.5 38.7-73.5 82.6 0 87.8 80.4 101.3 150.4 101.3h48.2c70.3 0 150.6-13.4 150.6-101.3zm-82.6-55.1c-25.8 0-36.7 34.2-36.7 55.1s10.9 55.1 36.7 55.1 36.7-34.2 36.7-55.1-10.9-55.1-36.7-55.1z"],"github-square":[448,512,[],"f092","M400 32H48C21.5 32 0 53.5 0 80v352c0 26.5 21.5 48 48 48h352c26.5 0 48-21.5 48-48V80c0-26.5-21.5-48-48-48zM277.3 415.7c-8.4 1.5-11.5-3.7-11.5-8 0-5.4.2-33 .2-55.3 0-15.6-5.2-25.5-11.3-30.7 37-4.1 76-9.2 76-73.1 0-18.2-6.5-27.3-17.1-39 1.7-4.3 7.4-22-1.7-45-13.9-4.3-45.7 17.9-45.7 17.9-13.2-3.7-27.5-5.6-41.6-5.6-14.1 0-28.4 1.9-41.6 5.6 0 0-31.8-22.2-45.7-17.9-9.1 22.9-3.5 40.6-1.7 45-10.6 11.7-15.6 20.8-15.6 39 0 63.6 37.3 69 74.3 73.1-4.8 4.3-9.1 11.7-10.6 22.3-9.5 4.3-33.8 11.7-48.3-13.9-9.1-15.8-25.5-17.1-25.5-17.1-16.2-.2-1.1 10.2-1.1 10.2 10.8 5 18.4 24.2 18.4 24.2 9.7 29.7 56.1 19.7 56.1 19.7 0 13.9.2 36.5.2 40.6 0 4.3-3 9.5-11.5 8-66-22.1-112.2-84.9-112.2-158.3 0-91.8 70.2-161.5 162-161.5S388 165.6 388 257.4c.1 73.4-44.7 136.3-110.7 158.3zm-98.1-61.1c-1.9.4-3.7-.4-3.9-1.7-.2-1.5 1.1-2.8 3-3.2 1.9-.2 3.7.6 3.9 1.9.3 1.3-1 2.6-3 3zm-9.5-.9c0 1.3-1.5 2.4-3.5 2.4-2.2.2-3.7-.9-3.7-2.4 0-1.3 1.5-2.4 3.5-2.4 1.9-.2 3.7.9 3.7 2.4zm-13.7-1.1c-.4 1.3-2.4 1.9-4.1 1.3-1.9-.4-3.2-1.9-2.8-3.2.4-1.3 2.4-1.9 4.1-1.5 2 .6 3.3 2.1 2.8 3.4zm-12.3-5.4c-.9 1.1-2.8.9-4.3-.6-1.5-1.3-1.9-3.2-.9-4.1.9-1.1 2.8-.9 4.3.6 1.3 1.3 1.8 3.3.9 4.1zm-9.1-9.1c-.9.6-2.6 0-3.7-1.5s-1.1-3.2 0-3.9c1.1-.9 2.8-.2 3.7 1.3 1.1 1.5 1.1 3.3 0 4.1zm-6.5-9.7c-.9.9-2.4.4-3.5-.6-1.1-1.3-1.3-2.8-.4-3.5.9-.9 2.4-.4 3.5.6 1.1 1.3 1.3 2.8.4 3.5zm-6.7-7.4c-.4.9-1.7 1.1-2.8.4-1.3-.6-1.9-1.7-1.5-2.6.4-.6 1.5-.9 2.8-.4 1.3.7 1.9 1.8 1.5 2.6z"],gitkraken:[592,512,[],"f3a6","M565.7 118.1c-2.3-6.1-9.3-9.2-15.3-6.6-5.7 2.4-8.5 8.9-6.3 14.6 10.9 29 16.9 60.5 16.9 93.3 0 134.6-100.3 245.7-230.2 262.7V358.4c7.9-1.5 15.5-3.6 23-6.2v104c106.7-25.9 185.9-122.1 185.9-236.8 0-91.8-50.8-171.8-125.8-213.3-5.7-3.2-13-.9-15.9 5-2.7 5.5-.6 12.2 4.7 15.1 67.9 37.6 113.9 110 113.9 193.2 0 93.3-57.9 173.1-139.8 205.4v-92.2c14.2-4.5 24.9-17.7 24.9-33.5 0-13.1-6.8-24.4-17.3-30.5 8.3-79.5 44.5-58.6 44.5-83.9V170c0-38-87.9-161.8-129-164.7-2.5-.2-5-.2-7.6 0C251.1 8.3 163.2 132 163.2 170v14.8c0 25.3 36.3 4.3 44.5 83.9-10.6 6.1-17.3 17.4-17.3 30.5 0 15.8 10.6 29 24.8 33.5v92.2c-81.9-32.2-139.8-112-139.8-205.4 0-83.1 46-155.5 113.9-193.2 5.4-3 7.4-9.6 4.7-15.1-2.9-5.9-10.1-8.2-15.9-5-75 41.5-125.8 121.5-125.8 213.3 0 114.7 79.2 210.8 185.9 236.8v-104c7.6 2.5 15.1 4.6 23 6.2v123.7C131.4 465.2 31 354.1 31 219.5c0-32.8 6-64.3 16.9-93.3 2.2-5.8-.6-12.2-6.3-14.6-6-2.6-13 .4-15.3 6.6C14.5 149.7 8 183.8 8 219.5c0 155.1 122.6 281.6 276.3 287.8V361.4c6.8.4 15 .5 23.4 0v145.8C461.4 501.1 584 374.6 584 219.5c0-35.7-6.5-69.8-18.3-101.4zM365.9 275.5c13 0 23.7 10.5 23.7 23.7 0 13.1-10.6 23.7-23.7 23.7-13 0-23.7-10.5-23.7-23.7 0-13.1 10.6-23.7 23.7-23.7zm-139.8 47.3c-13.2 0-23.7-10.7-23.7-23.7s10.5-23.7 23.7-23.7c13.1 0 23.7 10.6 23.7 23.7 0 13-10.5 23.7-23.7 23.7z"],gitlab:[512,512,[],"f296","M29.782 199.732L256 493.714 8.074 309.699c-6.856-5.142-9.712-13.996-7.141-21.993l28.849-87.974zm75.405-174.806c-3.142-8.854-15.709-8.854-18.851 0L29.782 199.732h131.961L105.187 24.926zm56.556 174.806L256 493.714l94.257-293.982H161.743zm349.324 87.974l-28.849-87.974L256 493.714l247.926-184.015c6.855-5.142 9.711-13.996 7.141-21.993zm-85.404-262.78c-3.142-8.854-15.709-8.854-18.851 0l-56.555 174.806h131.961L425.663 24.926z"],gitter:[384,512,[],"f426","M66.4 322.5H16V0h50.4v322.5zM166.9 76.1h-50.4V512h50.4V76.1zm100.6 0h-50.4V512h50.4V76.1zM368 76h-50.4v247H368V76z"],glide:[448,512,[],"f2a5","M252.8 148.6c0 8.8-1.6 17.7-3.4 26.4-5.8 27.8-11.6 55.8-17.3 83.6-1.4 6.3-8.3 4.9-13.7 4.9-23.8 0-30.5-26-30.5-45.5 0-29.3 11.2-68.1 38.5-83.1 4.3-2.5 9.2-4.2 14.1-4.2 11.4 0 12.3 8.3 12.3 17.9zM448 80v352c0 26.5-21.5 48-48 48H48c-26.5 0-48-21.5-48-48V80c0-26.5 21.5-48 48-48h352c26.5 0 48 21.5 48 48zm-64 187c0-5.1-20.8-37.7-25.5-39.5-2.2-.9-7.2-2.3-9.6-2.3-23.1 0-38.7 10.5-58.2 21.5l-.5-.5c4.3-29.4 14.6-57.2 14.6-87.4 0-44.6-23.8-62.7-67.5-62.7-71.7 0-108 70.8-108 123.5 0 54.7 32 85 86.3 85 7.5 0 6.9-.6 6.9 2.3-10.5 80.3-56.5 82.9-56.5 58.9 0-24.4 28-36.5 28.3-38-.2-7.6-29.3-17.2-36.7-17.2-21.1 0-32.7 33-32.7 50.6 0 32.3 20.4 54.7 53.3 54.7 48.2 0 83.4-49.7 94.3-91.7 9.4-37.7 7-39.4 12.3-42.1 20-10.1 35.8-16.8 58.4-16.8 11.1 0 19 2.3 36.7 5.2 1.8.1 4.1-1.7 4.1-3.5z"],"glide-g":[448,512,[],"f2a6","M407.1 211.2c-3.5-1.4-11.6-3.8-15.4-3.8-37.1 0-62.2 16.8-93.5 34.5l-.9-.9c7-47.3 23.5-91.9 23.5-140.4C320.8 29.1 282.6 0 212.4 0 97.3 0 39 113.7 39 198.4 39 286.3 90.3 335 177.6 335c12 0 11-1 11 3.8-16.9 128.9-90.8 133.1-90.8 94.6 0-39.2 45-58.6 45.5-61-.3-12.2-47-27.6-58.9-27.6-33.9.1-52.4 51.2-52.4 79.3C32 476 64.8 512 117.5 512c77.4 0 134-77.8 151.4-145.4 15.1-60.5 11.2-63.3 19.7-67.6 32.2-16.2 57.5-27 93.8-27 17.8 0 30.5 3.7 58.9 8.4 2.9 0 6.7-2.9 6.7-5.8 0-8-33.4-60.5-40.9-63.4zm-175.3-84.4c-9.3 44.7-18.6 89.6-27.8 134.3-2.3 10.2-13.3 7.8-22 7.8-38.3 0-49-41.8-49-73.1 0-47 18-109.3 61.8-133.4 7-4.1 14.8-6.7 22.6-6.7 18.6 0 20 13.3 20 28.7-.1 14.3-2.7 28.5-5.6 42.4z"],gofore:[400,512,[],"f3a7","M324 319.8h-13.2v34.7c-24.5 23.1-56.3 35.8-89.9 35.8-73.2 0-132.4-60.2-132.4-134.4 0-74.1 59.2-134.4 132.4-134.4 35.3 0 68.6 14 93.6 39.4l62.3-63.3C335 55.3 279.7 32 220.7 32 98 32 0 132.6 0 256c0 122.5 97 224 220.7 224 63.2 0 124.5-26.2 171-82.5-2-27.6-13.4-77.7-67.7-77.7zm-12.1-112.5H205.6v89H324c33.5 0 60.5 15.1 76 41.8v-30.6c0-65.2-40.4-100.2-88.1-100.2z"],goodreads:[448,512,[],"f3a8","M299.9 191.2c5.1 37.3-4.7 79-35.9 100.7-22.3 15.5-52.8 14.1-70.8 5.7-37.1-17.3-49.5-58.6-46.8-97.2 4.3-60.9 40.9-87.9 75.3-87.5 46.9-.2 71.8 31.8 78.2 78.3zM448 88v336c0 30.9-25.1 56-56 56H56c-30.9 0-56-25.1-56-56V88c0-30.9 25.1-56 56-56h336c30.9 0 56 25.1 56 56zM330 313.2s-.1-34-.1-217.3h-29v40.3c-.8.3-1.2-.5-1.6-1.2-9.6-20.7-35.9-46.3-76-46-51.9.4-87.2 31.2-100.6 77.8-4.3 14.9-5.8 30.1-5.5 45.6 1.7 77.9 45.1 117.8 112.4 115.2 28.9-1.1 54.5-17 69-45.2.5-1 1.1-1.9 1.7-2.9.2.1.4.1.6.2.3 3.8.2 30.7.1 34.5-.2 14.8-2 29.5-7.2 43.5-7.8 21-22.3 34.7-44.5 39.5-17.8 3.9-35.6 3.8-53.2-1.2-21.5-6.1-36.5-19-41.1-41.8-.3-1.6-1.3-1.3-2.3-1.3h-26.8c.8 10.6 3.2 20.3 8.5 29.2 24.2 40.5 82.7 48.5 128.2 37.4 49.9-12.3 67.3-54.9 67.4-106.3z"],"goodreads-g":[384,512,[],"f3a9","M42.6 403.3h2.8c12.7 0 25.5 0 38.2.1 1.6 0 3.1-.4 3.6 2.1 7.1 34.9 30 54.6 62.9 63.9 26.9 7.6 54.1 7.8 81.3 1.8 33.8-7.4 56-28.3 68-60.4 8-21.5 10.7-43.8 11-66.5.1-5.8.3-47-.2-52.8l-.9-.3c-.8 1.5-1.7 2.9-2.5 4.4-22.1 43.1-61.3 67.4-105.4 69.1-103 4-169.4-57-172-176.2-.5-23.7 1.8-46.9 8.3-69.7C58.3 47.7 112.3.6 191.6 0c61.3-.4 101.5 38.7 116.2 70.3.5 1.1 1.3 2.3 2.4 1.9V10.6h44.3c0 280.3.1 332.2.1 332.2-.1 78.5-26.7 143.7-103 162.2-69.5 16.9-159 4.8-196-57.2-8-13.5-11.8-28.3-13-44.5zM188.9 36.5c-52.5-.5-108.5 40.7-115 133.8-4.1 59 14.8 122.2 71.5 148.6 27.6 12.9 74.3 15 108.3-8.7 47.6-33.2 62.7-97 54.8-154-9.7-71.1-47.8-120-119.6-119.7z"],google:[488,512,[],"f1a0","M488 261.8C488 403.3 391.1 504 248 504 110.8 504 0 393.2 0 256S110.8 8 248 8c66.8 0 123 24.5 166.3 64.9l-67.5 64.9C258.5 52.6 94.3 116.6 94.3 256c0 86.5 69.1 156.6 153.7 156.6 98.2 0 135-70.4 140.8-106.9H248v-85.3h236.1c2.3 12.7 3.9 24.9 3.9 41.4z"],"google-drive":[512,512,[],"f3aa","M339 314.9L175.4 32h161.2l163.6 282.9H339zm-137.5 23.6L120.9 480h310.5L512 338.5H201.5zM154.1 67.4L0 338.5 80.6 480 237 208.8 154.1 67.4z"],"google-play":[512,512,[],"f3ab","M325.3 234.3L104.6 13l280.8 161.2-60.1 60.1zM47 0C34 6.8 25.3 19.2 25.3 35.3v441.3c0 16.1 8.7 28.5 21.7 35.3l256.6-256L47 0zm425.2 225.6l-58.9-34.1-65.7 64.5 65.7 64.5 60.1-34.1c18-14.3 18-46.5-1.2-60.8zM104.6 499l280.8-161.2-60.1-60.1L104.6 499z"],"google-plus":[496,512,[],"f2b3","M248 8C111.1 8 0 119.1 0 256s111.1 248 248 248 248-111.1 248-248S384.9 8 248 8zm-70.7 372c-68.8 0-124-55.5-124-124s55.2-124 124-124c31.3 0 60.1 11 83 32.3l-33.6 32.6c-13.2-12.9-31.3-19.1-49.4-19.1-42.9 0-77.2 35.5-77.2 78.1s34.2 78.1 77.2 78.1c32.6 0 64.9-19.1 70.1-53.3h-70.1v-42.6h116.9c1.3 6.8 1.9 13.6 1.9 20.7 0 70.8-47.5 121.2-118.8 121.2zm230.2-106.2v35.5H372v-35.5h-35.5v-35.5H372v-35.5h35.5v35.5h35.2v35.5h-35.2z"],"google-plus-g":[640,512,[],"f0d5","M386.061 228.496c1.834 9.692 3.143 19.384 3.143 31.956C389.204 370.205 315.599 448 204.8 448c-106.084 0-192-85.915-192-192s85.916-192 192-192c51.864 0 95.083 18.859 128.611 50.292l-52.126 50.03c-14.145-13.621-39.028-29.599-76.485-29.599-65.484 0-118.92 54.221-118.92 121.277 0 67.056 53.436 121.277 118.92 121.277 75.961 0 104.513-54.745 108.965-82.773H204.8v-66.009h181.261zm185.406 6.437V179.2h-56.001v55.733h-55.733v56.001h55.733v55.733h56.001v-55.733H627.2v-56.001h-55.733z"],"google-plus-square":[448,512,[],"f0d4","M400 32H48C21.5 32 0 53.5 0 80v352c0 26.5 21.5 48 48 48h352c26.5 0 48-21.5 48-48V80c0-26.5-21.5-48-48-48zM164 356c-55.3 0-100-44.7-100-100s44.7-100 100-100c27 0 49.5 9.8 67 26.2l-27.1 26.1c-7.4-7.1-20.3-15.4-39.8-15.4-34.1 0-61.9 28.2-61.9 63.2 0 34.9 27.8 63.2 61.9 63.2 39.6 0 54.4-28.5 56.8-43.1H164v-34.4h94.4c1 5 1.6 10.1 1.6 16.6 0 57.1-38.3 97.6-96 97.6zm220-81.8h-29v29h-29.2v-29h-29V245h29v-29H355v29h29v29.2z"],"google-wallet":[448,512,[],"f1ee","M156.8 126.8c37.6 60.6 64.2 113.1 84.3 162.5-8.3 33.8-18.8 66.5-31.3 98.3-13.2-52.3-26.5-101.3-56-148.5 6.5-36.4 2.3-73.6 3-112.3zM109.3 200H16.1c-6.5 0-10.5 7.5-6.5 12.7C51.8 267 81.3 330.5 101.3 400h103.5c-16.2-69.7-38.7-133.7-82.5-193.5-3-4-8-6.5-13-6.5zm47.8-88c68.5 108 130 234.5 138.2 368H409c-12-138-68.4-265-143.2-368H157.1zm251.8-68.5c-1.8-6.8-8.2-11.5-15.2-11.5h-88.3c-5.3 0-9 5-7.8 10.3 13.2 46.5 22.3 95.5 26.5 146 48.2 86.2 79.7 178.3 90.6 270.8 15.8-60.5 25.3-133.5 25.3-203 0-73.6-12.1-145.1-31.1-212.6z"],gratipay:[496,512,[],"f184","M248 8C111.1 8 0 119.1 0 256s111.1 248 248 248 248-111.1 248-248S384.9 8 248 8zm114.6 226.4l-113 152.7-112.7-152.7c-8.7-11.9-19.1-50.4 13.6-72 28.1-18.1 54.6-4.2 68.5 11.9 15.9 17.9 46.6 16.9 61.7 0 13.9-16.1 40.4-30 68.1-11.9 32.9 21.6 22.6 60 13.8 72z"],grav:[512,512,[],"f2d6","M301.1 212c4.4 4.4 4.4 11.9 0 16.3l-9.7 9.7c-4.4 4.7-11.9 4.7-16.6 0l-10.5-10.5c-4.4-4.7-4.4-11.9 0-16.6l9.7-9.7c4.4-4.4 11.9-4.4 16.6 0l10.5 10.8zm-30.2-19.7c3-3 3-7.8 0-10.5-2.8-3-7.5-3-10.5 0-2.8 2.8-2.8 7.5 0 10.5 3.1 2.8 7.8 2.8 10.5 0zm-26 5.3c-3 2.8-3 7.5 0 10.2 2.8 3 7.5 3 10.5 0 2.8-2.8 2.8-7.5 0-10.2-3-3-7.7-3-10.5 0zm72.5-13.3c-19.9-14.4-33.8-43.2-11.9-68.1 21.6-24.9 40.7-17.2 59.8.8 11.9 11.3 29.3 24.9 17.2 48.2-12.5 23.5-45.1 33.2-65.1 19.1zm47.7-44.5c-8.9-10-23.3 6.9-15.5 16.1 7.4 9 32.1 2.4 15.5-16.1zM504 256c0 137-111 248-248 248S8 393 8 256 119 8 256 8s248 111 248 248zm-66.2 42.6c2.5-16.1-20.2-16.6-25.2-25.7-13.6-24.1-27.7-36.8-54.5-30.4 11.6-8 23.5-6.1 23.5-6.1.3-6.4 0-13-9.4-24.9 3.9-12.5.3-22.4.3-22.4 15.5-8.6 26.8-24.4 29.1-43.2 3.6-31-18.8-59.2-49.8-62.8-22.1-2.5-43.7 7.7-54.3 25.7-23.2 40.1 1.4 70.9 22.4 81.4-14.4-1.4-34.3-11.9-40.1-34.3-6.6-25.7 2.8-49.8 8.9-61.4 0 0-4.4-5.8-8-8.9 0 0-13.8 0-24.6 5.3 11.9-15.2 25.2-14.4 25.2-14.4 0-6.4-.6-14.9-3.6-21.6-5.4-11-23.8-12.9-31.7 2.8.1-.2.3-.4.4-.5-5 11.9-1.1 55.9 16.9 87.2-2.5 1.4-9.1 6.1-13 10-21.6 9.7-56.2 60.3-56.2 60.3-28.2 10.8-77.2 50.9-70.6 79.7.3 3 1.4 5.5 3 7.5-2.8 2.2-5.5 5-8.3 8.3-11.9 13.8-5.3 35.2 17.7 24.4 15.8-7.2 29.6-20.2 36.3-30.4 0 0-5.5-5-16.3-4.4 27.7-6.6 34.3-9.4 46.2-9.1 8 3.9 8-34.3 8-34.3 0-14.7-2.2-31-11.1-41.5 12.5 12.2 29.1 32.7 28 60.6-.8 18.3-15.2 23-15.2 23-9.1 16.6-43.2 65.9-30.4 106 0 0-9.7-14.9-10.2-22.1-17.4 19.4-46.5 52.3-24.6 64.5 26.6 14.7 108.8-88.6 126.2-142.3 34.6-20.8 55.4-47.3 63.9-65 22 43.5 95.3 94.5 101.1 59z"],gripfire:[384,512,[],"f3ac","M171.8 503.8c0-5.3 4.8-12.2 4.8-22.3 0-15.2-13-39.9-78.1-86.6C64.2 365.8 32 336.4 32 286.6 32 171.9 179.1 110.1 179.1 18c0-3.3-.2-6.7-.6-10 5.1 2.4 39.1 43.3 39.1 90.4 0 80.5-105.1 129.2-105.1 203 0 26.9 16.6 47.2 32.6 69.5 22.5 30.2 44.2 56.9 44.2 86.5-.1 14.5-4.4 29.7-17.5 46.4zm146-241.4c1.5 8.4 2.2 16.6 2.2 24.6 0 51.8-29.4 97.5-67.3 136.8-1 1-2.2 2.4-3.2 2.4-3.6 0-35.5-41.6-35.5-53.2 0 0 41.8-55.7 41.8-96.9 0-10.8-2.7-21.7-9.1-33.4-1.5 32.3-55.7 87.7-58.1 87.7-2.7 0-17.9-22-17.9-42.1 0-5.3 1-10.7 3.2-15.8 2.4-5.5 56.6-72 56.6-116.7 0-6.2-1-12-3.4-17.1l-4-7.2c16.7 6.5 82.6 64.1 94.7 130.9"],grunt:[384,512,[],"f3ad","M61.3 189.3c-1.1 10 5.2 19.1 5.2 19.1.7-7.5 2.2-12.8 4-16.6.4 10.3 3.2 23.5 12.8 34.1 6.9 7.6 35.6 23.3 54.9 6.1 1 2.4 2.1 5.3 3 8.5 2.9 10.3-2.7 25.3-2.7 25.3s15.1-17.1 13.9-32.5c10.8-.5 21.4-8.4 21.1-19.5 0 0-18.9 10.4-35.5-8.8-9.7-11.2-40.9-42-83.1-31.8 4.3 1 8.9 2.4 13.5 4.1h-.1c-4.2 2-6.5 7.1-7 12zm28.3-1.8c19.5 11 37.4 25.7 44.9 37-5.7 3.3-21.7 10.4-38-1.7-10.3-7.6-9.8-26.2-6.9-35.3zm79.2 233.7c2.2 2.3 1.5 5.3.9 6.8-1.1 2.7-5.5 11.6-13 19.8-2.7 2.9-6.6 4.6-11 4.6-4.3 0-8.7-1.6-11.8-4.3-2.3-2.1-10.2-9.5-13.7-18.6-1.3-3.4-1-6.1.9-8.1 1.3-1.3 4-2.9 9.5-2.9H160c4.1 0 7 .9 8.8 2.7zm62.9-187.9c-1.2 15.5 13.9 32.5 13.9 32.5s-5.6-15-2.7-25.3c.9-3.2 2-6 3-8.5 19.3 17.3 48 1.5 54.8-6.1 9.6-10.6 12.3-23.8 12.8-34.1 1.8 3.8 3.4 9.1 4 16.6 0 0 6.4-9.1 5.2-19.1-.6-5-2.9-10-7-11.8h-.1c4.6-1.8 9.2-3.2 13.5-4.1-42.3-10.2-73.4 20.6-83.1 31.8-16.7 19.2-35.5 8.8-35.5 8.8-.2 10.9 10.4 18.9 21.2 19.3zm17.8-8.8c7.5-11.4 25.4-26 44.9-37 3 9.1 3.4 27.7-7 35.4-16.3 12.1-32.2 5-37.9 1.6-.1.1 0 0 0 0zM263 421.4c1.9 1.9 2.2 4.6.9 7.9-3.5 8.9-11.4 16.1-13.7 18.1-3.1 2.6-7.4 4.2-11.8 4.2s-8.3-1.6-11-4.5c-7.5-8-12-16.7-13-19.3-.6-1.5-1.3-4.4.9-6.7 1.7-1.8 4.7-2.7 8.9-2.7h29.4c5.4.1 8.1 1.7 9.4 3zm-98.3-251.5c9.9 6 18.8 8.1 27.3 8.3 8.5-.2 17.4-2.3 27.3-8.3 0 0-14.5 17.7-27.2 17.8h-.2c-12.7-.2-27.2-17.8-27.2-17.8zm184.5 147.4c-2.4 17.9-13 33.8-24.6 43.7-3.1-22.7-3.7-55.5-3.7-62.4 0-14.7 9.5-24.5 12.2-26.1 2.5-1.5 5.4-3 8.3-4.6 18-9.6 40.4-21.6 40.4-43.7 0-16.2-9.3-23.2-15.4-27.8-.8-.6-1.5-1.1-2.2-1.7-2.1-1.7-3.7-3-4.3-4.4-4.4-9.8-3.6-34.2-1.7-37.6.6-.6 16.7-20.9 11.8-39.2-2-7.4-6.9-13.3-14.1-17-5.3-2.7-11.9-4.2-19.5-4.5-.1-2-.5-3.9-.9-5.9-.6-2.6-1.1-5.3-.9-8.1.4-4.7.8-9 2.2-11.3 8.4-13.3 28.8-17.6 29-17.6l12.3-2.4-8.1-9.5c-.1-.2-17.3-17.5-46.3-17.5-7.9 0-16 1.3-24.1 3.9-24.2 7.8-42.9 30.5-49.4 39.3-3.1-1-6.3-1.9-9.6-2.7-4.2-15.8 9-38.5 9-38.5s-13.6-3-33.7 15.2c-2.6-6.5-8.1-20.5-1.8-37.2C184.6 10.1 177.2 26 175 40.4c-7.6-5.4-6.7-23.1-7.2-27.6-7.5.9-29.2 21.9-28.2 48.3-2 .5-3.9 1.1-5.9 1.7-6.5-8.8-25.1-31.5-49.4-39.3-7.9-2.2-16-3.5-23.9-3.5-29 0-46.1 17.3-46.3 17.5L6 46.9l12.3 2.4c.2 0 20.6 4.3 29 17.6 1.4 2.2 1.8 6.6 2.2 11.3.2 2.8-.4 5.5-.9 8.1-.4 1.9-.8 3.9-.9 5.9-7.7.3-14.2 1.8-19.5 4.5-7.2 3.7-12.1 9.6-14.1 17-5 18.2 11.2 38.5 11.8 39.2 1.9 3.4 2.7 27.8-1.7 37.6-.6 1.4-2.2 2.7-4.3 4.4-.7.5-1.4 1.1-2.2 1.7-6.1 4.6-15.4 11.7-15.4 27.8 0 22.1 22.4 34.1 40.4 43.7 3 1.6 5.8 3.1 8.3 4.6 2.7 1.6 12.2 11.4 12.2 26.1 0 6.9-.6 39.7-3.7 62.4-11.6-9.9-22.2-25.9-24.6-43.8 0 0-29.2 22.6-20.6 70.8 5.2 29.5 23.2 46.1 47 54.7 8.8 19.1 29.4 45.7 67.3 49.6C143 504.3 163 512 192.2 512h.2c29.1 0 49.1-7.7 63.6-19.5 37.9-3.9 58.5-30.5 67.3-49.6 23.8-8.7 41.7-25.2 47-54.7 8.2-48.4-21.1-70.9-21.1-70.9zM305.7 37.7c5.6-1.8 11.6-2.7 17.7-2.7 11 0 19.9 3 24.7 5-3.1 1.4-6.4 3.2-9.7 5.3-2.4-.4-5.6-.8-9.2-.8-10.5 0-20.5 3.1-28.7 8.9-12.3 8.7-18 16.9-20.7 22.4-2.2-1.3-4.5-2.5-7.1-3.7-1.6-.8-3.1-1.5-4.7-2.2 6.1-9.1 19.9-26.5 37.7-32.2zm21 18.2c-.8 1-1.6 2.1-2.3 3.2-3.3 5.2-3.9 11.6-4.4 17.8-.5 6.4-1.1 12.5-4.4 17-4.2.8-8.1 1.7-11.5 2.7-2.3-3.1-5.6-7-10.5-11.2 1.4-4.8 5.5-16.1 13.5-22.5 5.6-4.3 12.2-6.7 19.6-7zM45.6 45.3c-3.3-2.2-6.6-4-9.7-5.3 4.8-2 13.7-5 24.7-5 6.1 0 12 .9 17.7 2.7 17.8 5.8 31.6 23.2 37.7 32.1-1.6.7-3.2 1.4-4.8 2.2-2.5 1.2-4.9 2.5-7.1 3.7-2.6-5.4-8.3-13.7-20.7-22.4-8.3-5.8-18.2-8.9-28.8-8.9-3.4.1-6.6.5-9 .9zm44.7 40.1c-4.9 4.2-8.3 8-10.5 11.2-3.4-.9-7.3-1.9-11.5-2.7C65 89.5 64.5 83.4 64 77c-.5-6.2-1.1-12.6-4.4-17.8-.7-1.1-1.5-2.2-2.3-3.2 7.4.3 14 2.6 19.5 7 8 6.3 12.1 17.6 13.5 22.4zM58.1 259.9c-2.7-1.6-5.6-3.1-8.4-4.6-14.9-8-30.2-16.3-30.2-30.5 0-11.1 4.3-14.6 8.9-18.2l.5-.4c.7-.6 1.4-1.2 2.2-1.8-.9 7.2-1.9 13.3-2.7 14.9 0 0 12.1-15 15.7-44.3 1.4-11.5-1.1-34.3-5.1-43 .2 4.9 0 9.8-.3 14.4-.4-.8-.8-1.6-1.3-2.2-3.2-4-11.8-17.5-9.4-26.6.9-3.5 3.1-6 6.7-7.8 3.8-1.9 8.8-2.9 15.1-2.9 12.3 0 25.9 3.7 32.9 6 25.1 8 55.4 30.9 64.1 37.7.2.2.4.3.4.3l5.6 3.9-3.5-5.8c-.2-.3-19.1-31.4-53.2-46.5 2-2.9 7.4-8.1 21.6-15.1 21.4-10.5 46.5-15.8 74.3-15.8 27.9 0 52.9 5.3 74.3 15.8 14.2 6.9 19.6 12.2 21.6 15.1-34 15.1-52.9 46.2-53.1 46.5l-3.5 5.8 5.6-3.9s.2-.1.4-.3c8.7-6.8 39-29.8 64.1-37.7 7-2.2 20.6-6 32.9-6 6.3 0 11.3 1 15.1 2.9 3.5 1.8 5.7 4.4 6.7 7.8 2.5 9.1-6.1 22.6-9.4 26.6-.5.6-.9 1.3-1.3 2.2-.3-4.6-.5-9.5-.3-14.4-4 8.8-6.5 31.5-5.1 43 3.6 29.3 15.7 44.3 15.7 44.3-.8-1.6-1.8-7.7-2.7-14.9.7.6 1.5 1.2 2.2 1.8l.5.4c4.6 3.7 8.9 7.1 8.9 18.2 0 14.2-15.4 22.5-30.2 30.5-2.9 1.5-5.7 3.1-8.4 4.6-8.7 5-18 16.7-19.1 34.2-.9 14.6.9 49.9 3.4 75.9-12.4 4.8-26.7 6.4-39.7 6.8-2-4.1-3.9-8.5-5.5-13.1-.7-2-19.6-51.1-26.4-62.2 5.5 39 17.5 73.7 23.5 89.6-3.5-.5-7.3-.7-11.7-.7h-117c-4.4 0-8.3.3-11.7.7 6-15.9 18.1-50.6 23.5-89.6-6.8 11.2-25.7 60.3-26.4 62.2-1.6 4.6-3.5 9-5.5 13.1-13-.4-27.2-2-39.7-6.8 2.5-26 4.3-61.2 3.4-75.9-.9-17.4-10.3-29.2-19-34.2zM34.8 404.6c-12.1-20-8.7-54.1-3.7-59.1 10.9 34.4 47.2 44.3 74.4 45.4-2.7 4.2-5.2 7.6-7 10l-1.4 1.4c-7.2 7.8-8.6 18.5-4.1 31.8-22.7-.1-46.3-9.8-58.2-29.5zm45.7 43.5c6 1.1 12.2 1.9 18.6 2.4 3.5 8 7.4 15.9 12.3 23.1-14.4-5.9-24.4-16-30.9-25.5zM192 498.2c-60.6-.1-78.3-45.8-84.9-64.7-3.7-10.5-3.4-18.2.9-23.1 2.9-3.3 9.5-7.2 24.6-7.2h118.8c15.1 0 21.8 3.9 24.6 7.2 4.2 4.8 4.5 12.6.9 23.1-6.6 18.8-24.3 64.6-84.9 64.7zm80.6-24.6c4.9-7.2 8.8-15.1 12.3-23.1 6.4-.5 12.6-1.3 18.6-2.4-6.5 9.5-16.5 19.6-30.9 25.5zm76.6-69c-12 19.7-35.6 29.3-58.1 29.7 4.5-13.3 3.1-24.1-4.1-31.8-.4-.5-.9-1-1.4-1.5-1.8-2.4-4.3-5.8-7-10 27.2-1.2 63.5-11 74.4-45.4 5 5 8.4 39.1-3.8 59z"],gulp:[256,512,[],"f3ae","M209.8 391.1l-14.1 24.6-4.6 80.2c0 8.9-28.3 16.1-63.1 16.1s-63.1-7.2-63.1-16.1l-5.8-79.4-14.9-25.4c41.2 17.3 126 16.7 165.6 0zm-196-253.3l13.6 125.5c5.9-20 20.8-47 40-55.2 6.3-2.7 12.7-2.7 18.7.9 5.2 3 9.6 9.3 10.1 11.8 1.2 6.5-2 9.1-4.5 9.1-3 0-5.3-4.6-6.8-7.3-4.1-7.3-10.3-7.6-16.9-2.8-6.9 5-12.9 13.4-17.1 20.7-5.1 8.8-9.4 18.5-12 28.2-1.5 5.6-2.9 14.6-.6 19.9 1 2.2 2.5 3.6 4.9 3.6 5 0 12.3-6.6 15.8-10.1 4.5-4.5 10.3-11.5 12.5-16l5.2-15.5c2.6-6.8 9.9-5.6 9.9 0 0 10.2-3.7 13.6-10 34.7-5.8 19.5-7.6 25.8-7.6 25.8-.7 2.8-3.4 7.5-6.3 7.5-1.2 0-2.1-.4-2.6-1.2-1-1.4-.9-5.3-.8-6.3.2-3.2 6.3-22.2 7.3-25.2-2 2.2-4.1 4.4-6.4 6.6-5.4 5.1-14.1 11.8-21.5 11.8-3.4 0-5.6-.9-7.7-2.4l7.6 79.6c2 5 39.2 17.1 88.2 17.1 49.1 0 86.3-12.2 88.2-17.1l10.9-94.6c-5.7 5.2-12.3 11.6-19.6 14.8-5.4 2.3-17.4 3.8-17.4-5.7 0-5.2 9.1-14.8 14.4-21.5 1.4-1.7 4.7-5.9 4.7-8.1 0-2.9-6-2.2-11.7 2.5-3.2 2.7-6.2 6.3-8.7 9.7-4.3 6-6.6 11.2-8.5 15.5-6.2 14.2-4.1 8.6-9.1 22-5 13.3-4.2 11.8-5.2 14-.9 1.9-2.2 3.5-4 4.5-1.9 1-4.5.9-6.1-.3-.9-.6-1.3-1.9-1.3-3.7 0-.9.1-1.8.3-2.7 1.5-6.1 7.8-18.1 15-34.3 1.6-3.7 1-2.6.8-2.3-6.2 6-10.9 8.9-14.4 10.5-5.8 2.6-13 2.6-14.5-4.1-.1-.4-.1-.8-.2-1.2-11.8 9.2-24.3 11.7-20-8.1-4.6 8.2-12.6 14.9-22.4 14.9-4.1 0-7.1-1.4-8.6-5.1-2.3-5.5 1.3-14.9 4.6-23.8 1.7-4.5 4-9.9 7.1-16.2 1.6-3.4 4.2-5.4 7.6-4.5.6.2 1.1.4 1.6.7 2.6 1.8 1.6 4.5.3 7.2-3.8 7.5-7.1 13-9.3 20.8-.9 3.3-2 9 1.5 9 2.4 0 4.7-.8 6.9-2.4 4.6-3.4 8.3-8.5 11.1-13.5 2-3.6 4.4-8.3 5.6-12.3.5-1.7 1.1-3.3 1.8-4.8 1.1-2.5 2.6-5.1 5.2-5.1 1.3 0 2.4.5 3.2 1.5 1.7 2.2 1.3 4.5.4 6.9-2 5.6-4.7 10.6-6.9 16.7-1.3 3.5-2.7 8-2.7 11.7 0 3.4 3.7 2.6 6.8 1.2 2.4-1.1 4.8-2.8 6.8-4.5 1.2-4.9.9-3.8 26.4-68.2 1.3-3.3 3.7-4.7 6.1-4.7 1.2 0 2.2.4 3.2 1.1 1.7 1.3 1.7 4.1 1 6.2-.7 1.9-.6 1.3-4.5 10.5-5.2 12.1-8.6 20.8-13.2 31.9-1.9 4.6-7.7 18.9-8.7 22.3-.6 2.2-1.3 5.8 1 5.8 5.4 0 19.3-13.1 23.1-17 .2-.3.5-.4.9-.6.6-1.9 1.2-3.7 1.7-5.5 1.4-3.8 2.7-8.2 5.3-11.3.8-1 1.7-1.6 2.7-1.6 2.8 0 4.2 1.2 4.2 4 0 1.1-.7 5.1-1.1 6.2 1.4-1.5 2.9-3 4.5-4.5 15-13.9 25.7-6.8 25.7.2 0 7.4-8.9 17.7-13.8 23.4-1.6 1.9-4.9 5.4-5 6.4 0 1.3.9 1.8 2.2 1.8 2 0 6.4-3.5 8-4.7 5-3.9 11.8-9.9 16.6-14.1l14.8-136.8c-30.5 17.1-197.6 17.2-228.3.2zm229.7-8.5c0 21-231.2 21-231.2 0 0-8.8 51.8-15.9 115.6-15.9 9 0 17.8.1 26.3.4l12.6-48.7L228.1.6c1.4-1.4 5.8-.2 9.9 3.5s6.6 7.9 5.3 9.3l-.1.1L185.9 74l-10 40.7c39.9 2.6 67.6 8.1 67.6 14.6zm-69.4 4.6c0-.8-.9-1.5-2.5-2.1l-.2.8c0 1.3-5 2.4-11.1 2.4s-11.1-1.1-11.1-2.4c0-.1 0-.2.1-.3l.2-.7c-1.8.6-3 1.4-3 2.3 0 2.1 6.2 3.7 13.7 3.7 7.7.1 13.9-1.6 13.9-3.7z"],"hacker-news":[448,512,[],"f1d4","M0 32v448h448V32H0zm21.2 197.2H21c.1-.1.2-.3.3-.4 0 .1 0 .3-.1.4zm218 53.9V384h-31.4V281.3L128 128h37.3c52.5 98.3 49.2 101.2 59.3 125.6 12.3-27 5.8-24.4 60.6-125.6H320l-80.8 155.1z"],"hacker-news-square":[448,512,[],"f3af","M400 32H48C21.5 32 0 53.5 0 80v352c0 26.5 21.5 48 48 48h352c26.5 0 48-21.5 48-48V80c0-26.5-21.5-48-48-48zM21.2 229.2H21c.1-.1.2-.3.3-.4 0 .1 0 .3-.1.4zm218 53.9V384h-31.4V281.3L128 128h37.3c52.5 98.3 49.2 101.2 59.3 125.6 12.3-27 5.8-24.4 60.6-125.6H320l-80.8 155.1z"],hips:[640,512,[],"f452","M0 80.2c0-1.8.9-2.7 2.7-2.7h40.9c1.9 0 2.8.9 2.8 2.7v81.2c15.2-7.7 31.7-11.5 49.8-11.4 24 .1 44.2 6.2 60.3 18 18.7 13.5 28 31.9 28 55.3v136.1c0 1.9-.9 2.8-2.7 2.8h-27.3c-9.1 0-16.4-7.3-16.4-16.3V223.3c0-.9 2.7-27-45.8-27-48.6 0-45.8 26.2-45.8 27v136.1c0 1.9-.9 2.8-2.8 2.8h-41c-1.8 0-2.7-.9-2.7-2.8V80.2zm497.7 221.5c13.7 10.2 34.1 19.1 58.4 19.1 23.3 0 32.8-4.5 36.5-13.6 3-7.9-.6-16.1-12.2-21.2l-53.6-23.5c-21.4-9.4-33.8-24-37.2-43.6-5.7-33.7 22.2-53.3 22.7-53.7 13.2-9.6 32-15.4 58.5-15.4 19 0 37.4 3.3 55.1 9.9 1.3.5 1.9 1.3 1.9 2.6V207c0 2.1-2.3 3.4-4 2.4-39.7-20.7-76.6-12.3-84-6.8-6.6 4.9-6 12.5 2.6 16.1L600 244c16.5 7.1 28.1 18.4 34.9 34.1 5.5 12.6 6.6 25.6 3.1 39.1-9.6 36.9-44.9 45.5-45.6 45.8-10.5 3.1-23.6 4.3-36.3 4.3-16.6 0-32.6-2.7-48.2-8.2-9.7-3.4-14.6-10.3-14.6-20.7V304c0-2.1 2.3-3.7 4.4-2.3zM376.2 149.8c-31.7 0-104.2 20.1-104.2 103.5v183.5c0 .8.6 2.7 2.7 2.7h40.9c1.9 0 2.8-.9 2.8-2.7V348c16.5 12.7 35.8 19.1 57.7 19.1 60.5 0 108.7-48.5 108.7-108.7.1-60.3-48.2-108.6-108.6-108.6zm0 170.9c-17.2 0-31.9-6.1-44-18.2-12.2-12.2-18.2-26.8-18.2-44 0-34.5 27.6-62.2 62.2-62.2 34.5 0 62.2 27.6 62.2 62.2.1 34.3-27.3 62.2-62.2 62.2zm-124.6 38.7c0 1.9-.9 2.8-2.8 2.8h-40.9c-1.6 0-2.7-1.4-2.7-2.8V157.6c0-1.4 1.1-2.8 2.7-2.8h40.9c1.9 0 2.8.9 2.8 2.8v201.8M228.3 72.5c15.9 0 28.9 12.7 28.9 28.9 0 15.8-12.7 28.9-28.9 28.9s-28.9-13.3-28.9-28.9c.1-16 13-28.9 28.9-28.9"],"hire-a-helper":[512,512,[],"f3b0","M443.1 0H71.9C67.9 37.3 37.4 67.8 0 71.7v371.5c37.4 4.9 66 32.4 71.9 68.8h372.2c3-36.4 32.5-65.8 67.9-69.8V71.7c-36.4-5.9-65-35.3-68.9-71.7zm-37 404.9c-36.3 0-18.8-2-55.1-2-35.8 0-21 2-56.1 2-5.9 0-4.9-8.2 0-9.8 22.8-7.6 22.9-10.2 24.6-12.8 10.4-15.6 5.9-83 5.9-113 0-5.3-6.4-12.8-13.8-12.8H200.4c-7.4 0-13.8 7.5-13.8 12.8 0 30-4.5 97.4 5.9 113 1.7 2.5 1.8 5.2 24.6 12.8 4.9 1.6 6 9.8 0 9.8-35.1 0-20.3-2-56.1-2-36.3 0-18.8 2-55.1 2-7.9 0-5.8-10.8 0-10.8 10.2-3.4 13.5-3.5 21.7-13.8 7.7-12.9 7.9-44.4 7.9-127.8V151.3c0-22.2-12.2-28.3-28.6-32.4-8.8-2.2-4-11.8 1-11.8 36.5 0 20.6 2 57.1 2 32.7 0 16.5-2 49.2-2 3.3 0 8.5 8.3 1 10.8-4.9 1.6-27.6 3.7-27.6 39.3 0 45.6-.2 55.8 1 68.8 0 1.3 2.3 12.8 12.8 12.8h109.2c10.5 0 12.8-11.5 12.8-12.8 1.2-13 1-23.2 1-68.8 0-35.6-22.7-37.7-27.6-39.3-7.5-2.5-2.3-10.8 1-10.8 32.7 0 16.5 2 49.2 2 36.5 0 20.6-2 57.1-2 4.9 0 9.9 9.6 1 11.8-16.4 4.1-28.6 10.3-28.6 32.4v101.2c0 83.4.1 114.9 7.9 127.8 8.2 10.2 11.4 10.4 21.7 13.8 5.8 0 7.8 10.8 0 10.8z"],hooli:[640,512,[],"f427","M508.4 352h57.9V156.7L508.4 184v168zm73.7-110.5V352H640V241.5h-57.9zm-250.7-8.9c-18.2-18.2-50.4-17.1-50.4-17.1s-32.2-1.1-50.4 17.1c-1.9 1.9-3.7 3.9-5.3 6-38.2-29.6-72.5-46.5-102.1-61.1v-20.7l-22.5 10.6c-54.4-22.1-89-18.2-97.3.1 0 0-24.9 32.8 61.9 110.9v-31c-48.8-54.6-39-76.1-35.3-79.2 13.5-11.4 37.5-8 64.4 2.1L65.2 184v63.3c13.1 14.7 30.5 31.5 53.5 50.4l4.5 3.6v-29.8c0-6.9 1.7-18.2 10.8-18.2s10.6 6.9 10.6 15V317c18 12.2 37.3 22.1 57.7 29.6v-93.9c0-18.7-13.4-37.4-40.6-37.4-15.8-.1-30.5 8.2-38.5 21.9v-54.3c41.9 20.9 83.9 46.5 99.9 58.3-10.2 14.6-9.3 28.1-9.3 43.7 0 18.7-1.4 34.3 16.8 52.5 18.2 18.2 50.4 17.1 50.4 17.1s32.3 1.1 50.4-17.1c18.2-18.2 16.7-33.8 16.7-52.5 0-18.5 1.5-34.2-16.7-52.3zm-39.7 71.9c0 3.6-1.8 12.5-10.7 12.5-8.9 0-10.7-8.9-10.7-12.5v-40.4c0-8.7 7.3-10.9 10.7-10.9 3.4 0 10.7 2.1 10.7 10.9v40.4zm185.7-71.9c-18.2-18.2-50.4-17.1-50.4-17.1s-32.3-1.1-50.4 17.1c-18.2 18.2-16.8 33.9-16.8 52.6 0 18.7-1.4 34.3 16.8 52.5 18.2 18.2 50.4 17.1 50.4 17.1s32.3 1.1 50.4-17.1c18.2-18.2 16.8-33.8 16.8-52.5-.1-18.8 1.3-34.5-16.8-52.6zm-39.8 71.9c0 3.6-1.8 12.5-10.7 12.5-8.9 0-10.7-8.9-10.7-12.5v-40.4c0-8.7 7.3-10.9 10.7-10.9 3.4 0 10.7 2.1 10.7 10.9v40.4zm173.5-73c15.9 0 28.9-12.9 28.9-28.9s-12.9-24.5-28.9-24.5c-15.9 0-28.9 8.6-28.9 24.5s12.9 28.9 28.9 28.9zM144.5 352l38.3.8c-13.2-4.6-26-10.2-38.3-16.8v16zm-21.4 0v-28.6c-6.5-4.2-13-8.7-19.4-13.6-14.8-11.2-27.5-21.7-38.5-31.5V352h57.9zm59.7.8c36.5 12.5 69.9 14.2 94.7 7.2-19.9.2-45.8-2.6-75.3-13.3v5.3l-19.4.8z"],hotjar:[448,512,[],"f3b1","M414.9 161.5C340.2 29 121.1 0 121.1 0S222.2 110.4 93 197.7C11.3 252.8-21 324.4 14 402.6c26.8 59.9 83.5 84.3 144.6 93.4-29.2-55.1-6.6-122.4-4.1-129.6 57.1 86.4 165 0 110.8-93.9 71 15.4 81.6 138.6 27.1 215.5 80.5-25.3 134.1-88.9 148.8-145.6 15.5-59.3 3.7-127.9-26.3-180.9z"],houzz:[414,512,[],"f27c","M258.9 330.7H154.3V480H0V32h109.5v104.5l305.1 85.6V480H258.9V330.7z"],html5:[384,512,[],"f13b","M0 32l34.9 395.8L191.5 480l157.6-52.2L384 32H0zm308.2 127.9H124.4l4.1 49.4h175.6l-13.6 148.4-97.9 27v.3h-1.1l-98.7-27.3-6-75.8h47.7L138 320l53.5 14.5 53.7-14.5 6-62.2H84.3L71.5 112.2h241.1l-4.4 47.7z"],hubspot:[512,512,[],"f3b2","M267.4 211.6c-25.1 23.7-40.8 57.3-40.8 94.6 0 29.3 9.7 56.3 26 78L203.1 434c-4.4-1.6-9.1-2.5-14-2.5-10.8 0-20.9 4.2-28.5 11.8-7.6 7.6-11.8 17.8-11.8 28.6s4.2 20.9 11.8 28.5c7.6 7.6 17.8 11.6 28.5 11.6 10.8 0 20.9-3.9 28.6-11.6 7.6-7.6 11.8-17.8 11.8-28.5 0-4.2-.6-8.2-1.9-12.1l50-50.2c22 16.9 49.4 26.9 79.3 26.9 71.9 0 130-58.3 130-130.2 0-65.2-47.7-119.2-110.2-128.7V116c17.5-7.4 28.2-23.8 28.2-42.9 0-26.1-20.9-47.9-47-47.9S311.2 47 311.2 73.1c0 19.1 10.7 35.5 28.2 42.9v61.2c-15.2 2.1-29.6 6.7-42.7 13.6-27.6-20.9-117.5-85.7-168.9-124.8 1.2-4.4 2-9 2-13.8C129.8 23.4 106.3 0 77.4 0 48.6 0 25.2 23.4 25.2 52.2c0 28.9 23.4 52.3 52.2 52.3 9.8 0 18.9-2.9 26.8-7.6l163.2 114.7zm89.5 163.6c-38.1 0-69-30.9-69-69s30.9-69 69-69 69 30.9 69 69-30.9 69-69 69z"],imdb:[448,512,[],"f2d8","M350.5 288.7c0 5.4 1.6 14.4-6.2 14.4-1.6 0-3-.8-3.8-2.4-2.2-5.1-1.1-44.1-1.1-44.7 0-3.8-1.1-12.7 4.9-12.7 7.3 0 6.2 7.3 6.2 12.7v32.7zM265 229.9c0-9.7 1.6-16-10.3-16v83.7c12.2.3 10.3-8.7 10.3-18.4v-49.3zM448 80v352c0 26.5-21.5 48-48 48H48c-26.5 0-48-21.5-48-48V80c0-26.5 21.5-48 48-48h352c26.5 0 48 21.5 48 48zM21.3 228.8c-.1.1-.2.3-.3.4h.3v-.4zM97 192H64v127.8h33V192zm113.3 0h-43.1l-7.6 59.9c-2.7-20-5.4-40.1-8.7-59.9h-42.8v127.8h29v-84.5l12.2 84.5h20.6l11.6-86.4v86.4h28.7V192zm86.3 45.3c0-8.1.3-16.8-1.4-24.4-4.3-22.5-31.4-20.9-49-20.9h-24.6v127.8c86.1.1 75 6 75-82.5zm85.9 17.3c0-17.3-.8-30.1-22.2-30.1-8.9 0-14.9 2.7-20.9 9.2V192h-31.7v127.8h29.8l1.9-8.1c5.7 6.8 11.9 9.8 20.9 9.8 19.8 0 22.2-15.2 22.2-30.9v-36z"],instagram:[448,512,[],"f16d","M224.1 141c-63.6 0-114.9 51.3-114.9 114.9s51.3 114.9 114.9 114.9S339 319.5 339 255.9 287.7 141 224.1 141zm0 189.6c-41.1 0-74.7-33.5-74.7-74.7s33.5-74.7 74.7-74.7 74.7 33.5 74.7 74.7-33.6 74.7-74.7 74.7zm146.4-194.3c0 14.9-12 26.8-26.8 26.8-14.9 0-26.8-12-26.8-26.8s12-26.8 26.8-26.8 26.8 12 26.8 26.8zm76.1 27.2c-1.7-35.9-9.9-67.7-36.2-93.9-26.2-26.2-58-34.4-93.9-36.2-37-2.1-147.9-2.1-184.9 0-35.8 1.7-67.6 9.9-93.9 36.1s-34.4 58-36.2 93.9c-2.1 37-2.1 147.9 0 184.9 1.7 35.9 9.9 67.7 36.2 93.9s58 34.4 93.9 36.2c37 2.1 147.9 2.1 184.9 0 35.9-1.7 67.7-9.9 93.9-36.2 26.2-26.2 34.4-58 36.2-93.9 2.1-37 2.1-147.8 0-184.8zM398.8 388c-7.8 19.6-22.9 34.7-42.6 42.6-29.5 11.7-99.5 9-132.1 9s-102.7 2.6-132.1-9c-19.6-7.8-34.7-22.9-42.6-42.6-11.7-29.5-9-99.5-9-132.1s-2.6-102.7 9-132.1c7.8-19.6 22.9-34.7 42.6-42.6 29.5-11.7 99.5-9 132.1-9s102.7-2.6 132.1 9c19.6 7.8 34.7 22.9 42.6 42.6 11.7 29.5 9 99.5 9 132.1s2.7 102.7-9 132.1z"],"internet-explorer":[512,512,[],"f26b","M483.049 159.706c10.855-24.575 21.424-60.438 21.424-87.871 0-72.722-79.641-98.371-209.673-38.577-107.632-7.181-211.221 73.67-237.098 186.457 30.852-34.862 78.271-82.298 121.977-101.158C125.404 166.85 79.128 228.002 43.992 291.725 23.246 329.651 0 390.94 0 436.747c0 98.575 92.854 86.5 180.251 42.006 31.423 15.43 66.559 15.573 101.695 15.573 97.124 0 184.249-54.294 216.814-146.022H377.927c-52.509 88.593-196.819 52.996-196.819-47.436H509.9c6.407-43.581-1.655-95.715-26.851-141.162zM64.559 346.877c17.711 51.15 53.703 95.871 100.266 123.304-88.741 48.94-173.267 29.096-100.266-123.304zm115.977-108.873c2-55.151 50.276-94.871 103.98-94.871 53.418 0 101.981 39.72 103.981 94.871H180.536zm184.536-187.6c21.425-10.287 48.563-22.003 72.558-22.003 31.422 0 54.274 21.717 54.274 53.722 0 20.003-7.427 49.007-14.569 67.867-26.28-42.292-65.986-81.584-112.263-99.586z"],ioxhost:[640,512,[],"f208","M616 160h-67.3C511.2 70.7 422.9 8 320 8 183 8 72 119 72 256c0 16.4 1.6 32.5 4.7 48H24c-13.3 0-24 10.8-24 24 0 13.3 10.7 24 24 24h67.3c37.5 89.3 125.8 152 228.7 152 137 0 248-111 248-248 0-16.4-1.6-32.5-4.7-48H616c13.3 0 24-10.8 24-24 0-13.3-10.7-24-24-24zm-96 96c0 110.5-89.5 200-200 200-75.7 0-141.6-42-175.5-104H424c13.3 0 24-10.8 24-24 0-13.3-10.7-24-24-24H125.8c-3.8-15.4-5.8-31.4-5.8-48 0-110.5 89.5-200 200-200 75.7 0 141.6 42 175.5 104H216c-13.3 0-24 10.8-24 24 0 13.3 10.7 24 24 24h298.2c3.8 15.4 5.8 31.4 5.8 48zm-304-24h208c13.3 0 24 10.7 24 24 0 13.2-10.7 24-24 24H216c-13.3 0-24-10.7-24-24 0-13.2 10.7-24 24-24z"],itunes:[448,512,[],"f3b4","M223.6 80.3C129 80.3 52.5 157 52.5 251.5S129 422.8 223.6 422.8s171.2-76.7 171.2-171.2c0-94.6-76.7-171.3-171.2-171.3zm79.4 240c-3.2 13.6-13.5 21.2-27.3 23.8-12.1 2.2-22.2 2.8-31.9-5-11.8-10-12-26.4-1.4-36.8 8.4-8 20.3-9.6 38-12.8 3-.5 5.6-1.2 7.7-3.7 3.2-3.6 2.2-2 2.2-80.8 0-5.6-2.7-7.1-8.4-6.1-4 .7-91.9 17.1-91.9 17.1-5 1.1-6.7 2.6-6.7 8.3 0 116.1.5 110.8-1.2 118.5-2.1 9-7.6 15.8-14.9 19.6-8.3 4.6-23.4 6.6-31.4 5.2-21.4-4-28.9-28.7-14.4-42.9 8.4-8 20.3-9.6 38-12.8 3-.5 5.6-1.2 7.7-3.7 5-5.7.9-127 2.6-133.7.4-2.6 1.5-4.8 3.5-6.4 2.1-1.7 5.8-2.7 6.7-2.7 101-19 113.3-21.4 115.1-21.4 5.7-.4 9 3 9 8.7-.1 170.6.4 161.4-1 167.6zM345.2 32H102.8C45.9 32 0 77.9 0 134.8v242.4C0 434.1 45.9 480 102.8 480h242.4c57 0 102.8-45.9 102.8-102.8V134.8C448 77.9 402.1 32 345.2 32zM223.6 444c-106.3 0-192.5-86.2-192.5-192.5S117.3 59 223.6 59s192.5 86.2 192.5 192.5S329.9 444 223.6 444z"],"itunes-note":[384,512,[],"f3b5","M381.9 388.2c-6.4 27.4-27.2 42.8-55.1 48-24.5 4.5-44.9 5.6-64.5-10.2-23.9-20.1-24.2-53.4-2.7-74.4 17-16.2 40.9-19.5 76.8-25.8 6-1.1 11.2-2.5 15.6-7.4 6.4-7.2 4.4-4.1 4.4-163.2 0-11.2-5.5-14.3-17-12.3-8.2 1.4-185.7 34.6-185.7 34.6-10.2 2.2-13.4 5.2-13.4 16.7 0 234.7 1.1 223.9-2.5 239.5-4.2 18.2-15.4 31.9-30.2 39.5-16.8 9.3-47.2 13.4-63.4 10.4-43.2-8.1-58.4-58-29.1-86.6 17-16.2 40.9-19.5 76.8-25.8 6-1.1 11.2-2.5 15.6-7.4 10.1-11.5 1.8-256.6 5.2-270.2.8-5.2 3-9.6 7.1-12.9 4.2-3.5 11.8-5.5 13.4-5.5 204-38.2 228.9-43.1 232.4-43.1 11.5-.8 18.1 6 18.1 17.6.2 344.5 1.1 326-1.8 338.5z"],java:[377,512,[],"f4e4","M121.6 396s-19.6 11.4 13.9 15.2c40.6 4.6 61.3 4 106-4.5 0 0 11.8 7.4 28.2 13.8C169.5 463.4 42.9 418 121.6 396m-12.2-56.1s-21.9 16.2 11.6 19.7c43.3 4.5 77.6 4.8 136.8-6.6 0 0 8.2 8.3 21.1 12.8-121.3 35.5-256.3 2.9-169.5-25.9m103.2-95.1c24.7 28.4-6.5 54-6.5 54s62.7-32.4 33.9-72.9c-26.9-37.8-47.5-56.6 64.1-121.3.1 0-175.2 43.8-91.5 140.2m132.6 192.6s14.5 11.9-15.9 21.2c-57.9 17.5-240.8 22.8-291.6.7-18.3-7.9 16-19 26.8-21.3 11.2-2.4 17.7-2 17.7-2-20.3-14.3-131.3 28.1-56.4 40.2 204.2 33.2 372.4-14.9 319.4-38.8M131 281.8s-93.1 22.1-33 30.1c25.4 3.4 76 2.6 123.1-1.3 38.5-3.2 77.2-10.2 77.2-10.2s-13.6 5.8-23.4 12.5c-94.5 24.9-277 13.3-224.5-12.1 44.5-21.4 80.6-19 80.6-19m167 93.3c96.1-49.9 51.6-97.9 20.6-91.4-7.6 1.6-11 3-11 3s2.8-4.4 8.2-6.3c61.3-21.6 108.5 63.6-19.8 97.3 0-.1 1.5-1.4 2-2.6M240 0s53.2 53.2-50.5 135c-83.1 65.6-19 103.1 0 145.8-48.5-43.8-84.1-82.3-60.2-118.2C164.4 110.1 261.5 84.5 240 0m-99.5 510.4c92.2 5.9 233.8-3.3 237.1-46.9 0 0-6.4 16.5-76.2 29.7-78.7 14.8-175.8 13.1-233.3 3.6 0-.1 11.8 9.7 72.4 13.6"],"jedi-order":[448,512,[],"f50e","M231.89 335.72l31.44-45.89-20.18 55.5 62.56 9.09-62.56 9.08 24.22 47.43s-30.22-31.77-34.49-36.25c1.76 68.78 2.19 85.7 2.19 85.7s154.4-71.65 68.62-230.09c0 0 106.97-118.07 10.09-190.73 0 0 165.5 99.91 60.55 271.46 0 0 86.78-84.77 41.37-170.54 0 0 78.71 111.01-17.16 233.11 0 0 26.24-16.15 49.45-77.71 0 0-16.93 183.26-221.96 185.66v.02h-4.08v-.02C16.93 479.14 0 295.88 0 295.88c23.21 61.56 49.44 77.71 49.44 77.71-95.87-122.11-17.15-233.11-17.15-233.11-45.41 85.78 41.38 170.54 41.38 170.54-104.95-171.56 60.54-271.46 60.54-271.46-96.88 72.66 10.09 190.73 10.09 190.73-85.78 158.44 68.62 230.09 68.62 230.09s.43-16.93 2.19-85.7l-34.48 36.25 24.22-47.43-62.56-9.08 62.56-9.09-20.18-55.5 31.44 45.89c2.25-87.85 7.82-305.82 7.85-306.85l.01-2.43.02 1 .03-1 .01 2.43c.05 1.72 5.61 219.2 7.86 306.85z"],jenkins:[512,512,[],"f3b6","M487.1 425c-1.4-11.2-19-23.1-28.2-31.9-5.1-5-29-23.1-30.4-29.9-1.4-6.6 9.7-21.5 13.3-28.9 5.1-10.7 8.8-23.7 11.3-32.6 18.8-66.1 20.7-156.9-6.2-211.2-10.2-20.6-38.6-49-56.4-62.5-42-31.7-119.6-35.3-170.1-16.6-14.1 5.2-27.8 9.8-40.1 17.1-33.1 19.4-68.3 32.5-78.1 71.6-24.2 10.8-31.5 41.8-30.3 77.8.2 7 4.1 15.8 2.7 22.4-.7 3.3-5.2 7.6-6.1 9.8-11.6 27.7-2.3 64 11.1 83.7 8.1 11.9 21.5 22.4 39.2 25.2.7 10.6 3.3 19.7 8.2 30.4 3.1 6.8 14.7 19 10.4 27.7-2.2 4.4-21 13.8-27.3 17.6C89 407.2 73.7 415 54.2 429c-12.6 9-32.3 10.2-29.2 31.1 2.1 14.1 10.1 31.6 14.7 45.8.7 2 1.4 4.1 2.1 6h422c4.9-15.3 9.7-30.9 14.6-47.2 3.4-11.4 10.2-27.8 8.7-39.7zM205.9 33.7c1.8-.5 3.4.7 4.9 2.4-.2 5.2-5.4 5.1-8.9 6.8-5.4 6.7-13.4 9.8-20 17.2-6.8 7.5-14.4 27.7-23.4 30-4.5 1.1-9.7-.8-13.6-.5-10.4.7-17.7 6-28.3 7.5 13.6-29.9 56.1-54 89.3-63.4zm-104.8 93.6c13.5-14.9 32.1-24.1 54.8-25.9 11.7 29.7-8.4 65-.9 97.6 2.3 9.9 10.2 25.4-2.4 25.7.3-28.3-34.8-46.3-61.3-29.6-1.8-21.5-4.9-51.7 9.8-67.8zm36.7 200.2c-1-4.1-2.7-12.9-2.3-15.1 1.6-8.7 17.1-12.5 11-24.7-11.3-.1-13.8 10.2-24.1 11.3-26.7 2.6-45.6-35.4-44.4-58.4 1-19.5 17.6-38.2 40.1-35.8 16 1.8 21.4 19.2 24.5 34.7 9.2.5 22.5-.4 26.9-7.6-.6-17.5-8.8-31.6-8.2-47.7 1-30.3 17.5-57.6 4.8-87.4 13.6-30.9 53.5-55.3 83.1-70 36.6-18.3 94.9-3.7 129.3 15.8 19.7 11.1 34.4 32.7 48.3 50.7-19.5-5.8-36.1 4.2-33.1 20.3 16.3-14.9 44.2-.2 52.5 16.4 7.9 15.8 7.8 39.3 9 62.8 2.9 57-10.4 115.9-39.1 157.1-7.7 11-14.1 23-24.9 30.6-26 18.2-65.4 34.7-99.2 23.4-44.7-15-65-44.8-89.5-78.8.7 18.7 13.8 34.1 26.8 48.4 11.3 12.5 25 26.6 39.7 32.4-12.3-2.9-31.1-3.8-36.2 7.2-28.6-1.9-55.1-4.8-68.7-24.2-10.6-15.4-21.4-41.4-26.3-61.4zm222 124.1c4.1-3 11.1-2.9 17.4-3.6-5.4-2.7-13-3.7-19.3-2.2-.1-4.2-2-6.8-3.2-10.2 10.6-3.8 35.5-28.5 49.6-20.3 6.7 3.9 9.5 26.2 10.1 37 .4 9-.8 18-4.5 22.8-18.8-.6-35.8-2.8-50.7-7 .9-6.1-1-12.1.6-16.5zm-17.2-20c-16.8.8-26-1.2-38.3-10.8.2-.8 1.4-.5 1.5-1.4 18 8 40.8-3.3 59-4.9-7.9 5.1-14.6 11.6-22.2 17.1zm-12.1 33.2c-1.6-9.4-3.5-12-2.8-20.2 25-16.6 29.7 28.6 2.8 20.2zM226 438.6c-11.6-.7-48.1-14-38.5-23.7 9.4 6.5 27.5 4.9 41.3 7.3.8 4.4-2.8 10.2-2.8 16.4zM57.7 497.1c-4.3-12.7-9.2-25.1-14.8-36.9 30.8-23.8 65.3-48.9 102.2-63.5 2.8-1.1 23.2 25.4 26.2 27.6 16.5 11.7 37 21 56.2 30.2 1.2 8.8 3.9 20.2 8.7 35.5.7 2.3 1.4 4.7 2.2 7.2H57.7zm240.6 5.7h-.8c.3-.2.5-.4.8-.5v.5zm7.5-5.7c2.1-1.4 4.3-2.8 6.4-4.3 1.1 1.4 2.2 2.8 3.2 4.3h-9.6zm15.1-24.7c-10.8 7.3-20.6 18.3-33.3 25.2-6 3.3-27 11.7-33.4 10.2-3.6-.8-3.9-5.3-5.4-9.5-3.1-9-10.1-23.4-10.8-37-.8-17.2-2.5-46 16-42.4 14.9 2.9 32.3 9.7 43.9 16.1 7.1 3.9 11.1 8.6 21.9 9.5-.1 1.4-.1 2.8-.2 4.3-5.9 3.9-15.3 3.8-21.8 7.1 9.5.4 17 2.7 23.5 5.9-.1 3.4-.3 7-.4 10.6zm53.4 24.7h-14c-.1-3.2-2.8-5.8-6.1-5.8s-5.9 2.6-6.1 5.8h-17.4c-2.8-4.4-5.7-8.6-8.9-12.5 2.1-2.2 4-4.7 6-6.9 9 3.7 14.8-4.9 21.7-4.2 7.9.8 14.2 11.7 25.4 11l-.6 12.6zm8.7 0c.2-4 .4-7.8.6-11.5 15.6-7.3 29 1.3 35.7 11.5H383zm83.4-37c-2.3 11.2-5.8 24-9.9 37.1-.2-.1-.4-.1-.6-.1H428c.6-1.1 1.2-2.2 1.9-3.3-2.6-6.1-9-8.7-10.9-15.5 12.1-22.7 6.5-93.4-24.2-78.5 4.3-6.3 15.6-11.5 20.8-19.3 13 10.4 20.8 20.3 33.2 31.4 6.8 6 20 13.3 21.4 23.1.8 5.5-2.6 18.9-3.8 25.1zM222.2 130.5c5.4-14.9 27.2-34.7 45-32 7.7 1.2 18 8.2 12.2 17.7-30.2-7-45.2 12.6-54.4 33.1-8.1-2-4.9-13.1-2.8-18.8zm184.1 63.1c8.2-3.6 22.4-.7 29.6-5.3-4.2-11.5-10.3-21.4-9.3-37.7.5 0 1 0 1.4.1 6.8 14.2 12.7 29.2 21.4 41.7-5.7 13.5-43.6 25.4-43.1 1.2zm20.4-43zm-117.2 45.7c-6.8-10.9-19-32.5-14.5-45.3 6.5 11.9 8.6 24.4 17.8 33.3 4.1 4 12.2 9 8.2 20.2-.9 2.7-7.8 8.6-11.7 9.7-14.4 4.3-47.9.9-36.6-17.1 11.9.7 27.9 7.8 36.8-.8zm27.3 70c3.8 6.6 1.4 18.7 12.1 20.6 20.2 3.4 43.6-12.3 58.1-17.8 9-15.2-.8-20.7-8.9-30.5-16.6-20-38.8-44.8-38-74.7 6.7-4.9 7.3 7.4 8.2 9.7 8.7 20.3 30.4 46.2 46.3 63.5 3.9 4.3 10.3 8.4 11 11.2 2.1 8.2-5.4 18-4.5 23.5-21.7 13.9-45.8 29.1-81.4 25.6-7.4-6.7-10.3-21.4-2.9-31.1zm-201.3-9.2c-6.8-3.9-8.4-21-16.4-21.4-11.4-.7-9.3 22.2-9.3 35.5-7.8-7.1-9.2-29.1-3.5-40.3-6.6-3.2-9.5 3.6-13.1 5.9 4.7-34.1 49.8-15.8 42.3 20.3zm299.6 28.8c-10.1 19.2-24.4 40.4-54 41-.6-6.2-1.1-15.6 0-19.4 22.7-2.2 36.6-13.7 54-21.6zm-141.9 12.4c18.9 9.9 53.6 11 79.3 10.2 1.4 5.6 1.3 12.6 1.4 19.4-33 1.8-72-6.4-80.7-29.6zm92.2 46.7c-1.7 4.3-5.3 9.3-9.8 11.1-12.1 4.9-45.6 8.7-62.4-.3-10.7-5.7-17.5-18.5-23.4-26-2.8-3.6-16.9-12.9-.2-12.9 13.1 32.7 58 29 95.8 28.1z"],joget:[496,512,[],"f3b7","M227.5 468.7c-9-13.6-19.9-33.3-23.7-42.4-5.7-13.7-27.2-45.6 31.2-67.1 51.7-19.1 176.7-16.5 208.8-17.6-4 9-8.6 17.9-13.9 26.6-40.4 65.5-110.4 101.5-182 101.5-6.8 0-13.6-.4-20.4-1M66.1 143.9C128 43.4 259.6 12.2 360.1 74.1c74.8 46.1 111.2 130.9 99.3 212.7-24.9-.5-179.3-3.6-230.3-4.9-55.5-1.4-81.7-20.8-58.5-48.2 23.2-27.4 51.1-40.7 68.9-51.2 17.9-10.5 27.3-33.7-23.6-29.7C87.3 161.5 48.6 252.1 37.6 293c-8.8-49.7-.1-102.7 28.5-149.1m-29.2-18c-71.9 116.6-35.6 269.3 81 341.2 116.6 71.9 269.3 35.6 341.2-80.9 71.9-116.6 35.6-269.4-81-341.2-40.5-25.1-85.5-37-129.9-37C165 8 83.8 49.9 36.9 125.9m244.4 110.4c-31.5 20.5-65.3 31.3-65.3 31.3l169.5-1.6 46.5-23.4s3.6-9.5-19.1-15.5c-22.7-6-57 11.3-86.7 27.2-29.7 15.8-31.1 8.2-31.1 8.2s40.2-28.1 50.7-34.5c10.5-6.4 31.9-14 13.4-24.6-3.2-1.8-6.7-2.7-10.4-2.7-17.8 0-41.5 18.7-67.5 35.6"],joomla:[448,512,[],"f1aa","M.6 92.1C.6 58.8 27.4 32 60.4 32c30 0 54.5 21.9 59.2 50.2 32.6-7.6 67.1.6 96.5 30l-44.3 44.3c-20.5-20.5-42.6-16.3-55.4-3.5-14.3 14.3-14.3 37.9 0 52.2l99.5 99.5-44 44.3c-87.7-87.2-49.7-49.7-99.8-99.7-26.8-26.5-35-64.8-24.8-98.9C20.4 144.6.6 120.7.6 92.1zm129.5 116.4l44.3 44.3c10-10 89.7-89.7 99.7-99.8 14.3-14.3 37.6-14.3 51.9 0 12.8 12.8 17 35-3.5 55.4l44 44.3c31.2-31.2 38.5-67.6 28.9-101.2 29.2-4.1 51.9-29.2 51.9-59.5 0-33.2-26.8-60.1-59.8-60.1-30.3 0-55.4 22.5-59.5 51.6-33.8-9.9-71.7-1.5-98.3 25.1-18.3 19.1-71.1 71.5-99.6 99.9zm266.3 152.2c8.2-32.7-.9-68.5-26.3-93.9-11.8-12.2 5 4.7-99.5-99.7l-44.3 44.3 99.7 99.7c14.3 14.3 14.3 37.6 0 51.9-12.8 12.8-35 17-55.4-3.5l-44 44.3c27.6 30.2 68 38.8 102.7 28 5.5 27.4 29.7 48.1 58.9 48.1 33 0 59.8-26.8 59.8-60.1 0-30.2-22.5-55-51.6-59.1zm-84.3-53.1l-44-44.3c-87 86.4-50.4 50.4-99.7 99.8-14.3 14.3-37.6 14.3-51.9 0-13.1-13.4-16.9-35.3 3.2-55.4l-44-44.3c-30.2 30.2-38 65.2-29.5 98.3-26.7 6-46.2 29.9-46.2 58.2C0 453.2 26.8 480 59.8 480c28.6 0 52.5-19.8 58.6-46.7 32.7 8.2 68.5-.6 94.2-26 32.1-32 12.2-12.4 99.5-99.7z"],js:[448,512,[],"f3b8","M0 32v448h448V32H0zm243.8 349.4c0 43.6-25.6 63.5-62.9 63.5-33.7 0-53.2-17.4-63.2-38.5l34.3-20.7c6.6 11.7 12.6 21.6 27.1 21.6 13.8 0 22.6-5.4 22.6-26.5V237.7h42.1v143.7zm99.6 63.5c-39.1 0-64.4-18.6-76.7-43l34.3-19.8c9 14.7 20.8 25.6 41.5 25.6 17.4 0 28.6-8.7 28.6-20.8 0-14.4-11.4-19.5-30.7-28l-10.5-4.5c-30.4-12.9-50.5-29.2-50.5-63.5 0-31.6 24.1-55.6 61.6-55.6 26.8 0 46 9.3 59.8 33.7L368 290c-7.2-12.9-15-18-27.1-18-12.3 0-20.1 7.8-20.1 18 0 12.6 7.8 17.7 25.9 25.6l10.5 4.5c35.8 15.3 55.9 31 55.9 66.2 0 37.8-29.8 58.6-69.7 58.6z"],"js-square":[448,512,[],"f3b9","M400 32H48C21.5 32 0 53.5 0 80v352c0 26.5 21.5 48 48 48h352c26.5 0 48-21.5 48-48V80c0-26.5-21.5-48-48-48zM243.8 381.4c0 43.6-25.6 63.5-62.9 63.5-33.7 0-53.2-17.4-63.2-38.5l34.3-20.7c6.6 11.7 12.6 21.6 27.1 21.6 13.8 0 22.6-5.4 22.6-26.5V237.7h42.1v143.7zm99.6 63.5c-39.1 0-64.4-18.6-76.7-43l34.3-19.8c9 14.7 20.8 25.6 41.5 25.6 17.4 0 28.6-8.7 28.6-20.8 0-14.4-11.4-19.5-30.7-28l-10.5-4.5c-30.4-12.9-50.5-29.2-50.5-63.5 0-31.6 24.1-55.6 61.6-55.6 26.8 0 46 9.3 59.8 33.7L368 290c-7.2-12.9-15-18-27.1-18-12.3 0-20.1 7.8-20.1 18 0 12.6 7.8 17.7 25.9 25.6l10.5 4.5c35.8 15.3 55.9 31 55.9 66.2 0 37.8-29.8 58.6-69.7 58.6z"],jsfiddle:[576,512,[],"f1cc","M510.634 237.462c-4.727-2.621-5.664-5.748-6.381-10.776-2.352-16.488-3.539-33.619-9.097-49.095-35.895-99.957-153.99-143.386-246.849-91.646-27.37 15.25-48.971 36.369-65.493 63.903-3.184-1.508-5.458-2.71-7.824-3.686-30.102-12.421-59.049-10.121-85.331 9.167-25.531 18.737-36.422 44.548-32.676 76.408.355 3.025-1.967 7.621-4.514 9.545-39.712 29.992-56.031 78.065-41.902 124.615 13.831 45.569 57.514 79.796 105.608 81.433 30.291 1.031 60.637.546 90.959.539 84.041-.021 168.09.531 252.12-.48 52.664-.634 96.108-36.873 108.212-87.293 11.54-48.074-11.144-97.3-56.832-122.634zm21.107 156.88c-18.23 22.432-42.343 35.253-71.28 35.65-56.874.781-113.767.23-170.652.23 0 .7-163.028.159-163.728.154-43.861-.332-76.739-19.766-95.175-59.995-18.902-41.245-4.004-90.848 34.186-116.106 9.182-6.073 12.505-11.566 10.096-23.136-5.49-26.361 4.453-47.956 26.42-62.981 22.987-15.723 47.422-16.146 72.034-3.083 10.269 5.45 14.607 11.564 22.198-2.527 14.222-26.399 34.557-46.727 60.671-61.294 97.46-54.366 228.37 7.568 230.24 132.697.122 8.15 2.412 12.428 9.848 15.894 57.56 26.829 74.456 96.122 35.142 144.497zm-87.789-80.499c-5.848 31.157-34.622 55.096-66.666 55.095-16.953-.001-32.058-6.545-44.079-17.705-27.697-25.713-71.141-74.98-95.937-93.387-20.056-14.888-41.99-12.333-60.272 3.782-49.996 44.071 15.859 121.775 67.063 77.188 4.548-3.96 7.84-9.543 12.744-12.844 8.184-5.509 20.766-.884 13.168 10.622-17.358 26.284-49.33 38.197-78.863 29.301-28.897-8.704-48.84-35.968-48.626-70.179 1.225-22.485 12.364-43.06 35.414-55.965 22.575-12.638 46.369-13.146 66.991 2.474C295.68 280.7 320.467 323.97 352.185 343.47c24.558 15.099 54.254 7.363 68.823-17.506 28.83-49.209-34.592-105.016-78.868-63.46-3.989 3.744-6.917 8.932-11.41 11.72-10.975 6.811-17.333-4.113-12.809-10.353 20.703-28.554 50.464-40.44 83.271-28.214 31.429 11.714 49.108 44.366 42.76 78.186z"],keybase:[412,512,[],"f4f5","M177.2 430.9c0 9.8-8 17.8-17.8 17.8s-17.8-8-17.8-17.8 8-17.8 17.8-17.8c9.8-.1 17.8 7.9 17.8 17.8zM270 413c-9.8 0-17.8 8-17.8 17.8s8 17.8 17.8 17.8 17.8-8 17.8-17.8-8-17.8-17.8-17.8zm142.3-36c0 38.9-7.6 73.9-22.2 103h-27.3c23.5-38.7 30.5-94.8 22.4-134.3-16.1 29.5-52.1 38.6-85.9 28.8-127.8-37.5-192.5 19.7-234.6 50.3l18.9-59.3-39.9 42.3c4.8 26.7 15.7 51.3 31.2 72.3H46.1c-9.7-15.8-17.2-33-22.2-51.3L.1 454c0-74.9-5.5-147.6 61.5-215.2 20.2-20.4 43.7-36.2 69.1-46.7-6.8-13.5-9.5-29.2-7.8-46l-19.9-1.2c-17.9-1.1-31.6-16.5-30.6-34.4v-.1L74 84.2c1.1-17.1 15.4-30.6 32.5-30.6 1.3 0-.3-.1 28.2 1.7 13.9.8 21.5 9.8 22.8 11.4 7.1-10.4 14.5-20.5 24.6-34.5l20.6 12.1c-13.6 29-9.1 36.2-9 36.3 3.9 0 13.9-.5 32.4 5.7C246 92.9 262 107 271 126c.4.9 15.5 29 1.2 62.6 19 6.1 51.3 19.9 82.4 51.8 36.6 37.6 57.7 87.4 57.7 136.6zM128 122.3c3.2-10 7.7-19.7 13.1-29.4.1-2 2.2-13.1-7.8-13.8-28.5-1.8-26.3-1.6-26.7-1.6-4.6 0-8.3 3.5-8.6 8.1l-1.6 26.2c-.3 4.7 3.4 8.8 8.1 9.1l23.5 1.4zm25.8 61.8c5.6 9.4 14.1 16.1 22.3 20 0-21.2 28.5-41.9 52.8-17.5l8.4 10.3c20.8-18.8 19.4-45.3 12.1-60.9-13.8-29.1-46.9-32-54.3-31.7-10.3.4-19.7-5.4-23.7-15.3-13.7 21.2-37.2 62.5-17.6 95.1zm82.9 68.4L217 268.6c-1.9 1.6-2.2 4.4-.6 6.3l8.9 10.9c1 1.2 3.8 2.7 6.3.6l19.6-16 5.5 6.8c4.9 6 13.8-1.4 9-7.3-63.6-78.3-41.5-51.1-55.3-68.1-4.7-6-13.9 1.4-9 7.3 1.9 2.3 18.4 22.6 19.8 24.3l-9.6 7.9c-4.6 3.8 2.6 13.3 7.4 9.4l9.7-8 8 9.8zm118.4 25.7c-16.9-23.7-42.6-46.7-73.4-60.4-7.9-3.5-15-6.1-22.9-8.6-2 2.2-4.1 4.3-6.4 6.2l31.9 39.2c10.4 12.7 8.5 31.5-4.2 41.9-1.3 1.1-13.1 10.7-29 4.9-2.9 2.3-10.1 9.9-22.2 9.9-8.6 0-16.6-3.8-22.1-10.5l-8.9-10.9c-6.3-7.8-7.9-17.9-5-26.8-8.2-9.9-8.3-21.3-4.6-30-7.2-1.3-26.7-6.2-42.7-21.4-55.8 20.7-88 64.4-101.3 91.2-14.9 30.2-18.8 60.9-19.9 90.2 8.2-8.7-3.9 4.1 114-120.9l-29.9 93.6c57.8-31.1 124-36 197.4-14.4 23.6 6.9 45.1 1.6 56-13.9 11.1-15.6 8.5-37.7-6.8-59.3zM110.6 107.3l15.6 1 1-15.6-15.6-1-1 15.6z"],keycdn:[512,512,[],"f3ba","M63.8 409.3l60.5-59c32.1 42.8 71.1 66 126.6 67.4 30.5.7 60.3-7 86.4-22.4 5.1 5.3 18.5 19.5 20.9 22-32.2 20.7-69.6 31.1-108.1 30.2-43.3-1.1-84.6-16.7-117.7-44.4.3-.6-38.2 37.5-38.6 37.9 9.5 29.8-13.1 62.4-46.3 62.4C20.7 503.3 0 481.7 0 454.9c0-34.3 33.1-56.6 63.8-45.6zm354.9-252.4c19.1 31.3 29.6 67.4 28.7 104-1.1 44.8-19 87.5-48.6 121 .3.3 23.8 25.2 24.1 25.5 9.6-1.3 19.2 2 25.9 9.1 11.3 12 10.9 30.9-1.1 42.4-12 11.3-30.9 10.9-42.4-1.1-6.7-7-9.4-16.8-7.6-26.3-24.9-26.6-44.4-47.2-44.4-47.2 42.7-34.1 63.3-79.6 64.4-124.2.7-28.9-7.2-57.2-21.1-82.2l22.1-21zM104 53.1c6.7 7 9.4 16.8 7.6 26.3l45.9 48.1c-4.7 3.8-13.3 10.4-22.8 21.3-25.4 28.5-39.6 64.8-40.7 102.9-.7 28.9 6.1 57.2 20 82.4l-22 21.5C72.7 324 63.1 287.9 64.2 250.9c1-44.6 18.3-87.6 47.5-121.1l-25.3-26.4c-9.6 1.3-19.2-2-25.9-9.1-11.3-12-10.9-30.9 1.1-42.4C73.5 40.7 92.2 41 104 53.1zM464.9 8c26 0 47.1 22.4 47.1 48.3S490.9 104 464.9 104c-6.3.1-14-1.1-15.9-1.8l-62.9 59.7c-32.7-43.6-76.7-65.9-126.9-67.2-30.5-.7-60.3 6.8-86.2 22.4l-21.1-22C184.1 74.3 221.5 64 260 64.9c43.3 1.1 84.6 16.7 117.7 44.6l41.1-38.6c-1.5-4.7-2.2-9.6-2.2-14.5C416.5 29.7 438.9 8 464.9 8zM256.7 113.4c5.5 0 10.9.4 16.4 1.1 78.1 9.8 133.4 81.1 123.8 159.1-9.8 78.1-81.1 133.4-159.1 123.8-78.1-9.8-133.4-81.1-123.8-159.2 9.3-72.4 70.1-124.6 142.7-124.8zm-59 119.4c.6 22.7 12.2 41.8 32.4 52.2l-11 51.7h73.7l-11-51.7c20.1-10.9 32.1-29 32.4-52.2-.4-32.8-25.8-57.5-58.3-58.3-32.1.8-57.3 24.8-58.2 58.3zM256 160"],kickstarter:[448,512,[],"f3bb","M400 480H48c-26.4 0-48-21.6-48-48V80c0-26.4 21.6-48 48-48h352c26.4 0 48 21.6 48 48v352c0 26.4-21.6 48-48 48zM199.6 178.5c0-30.7-17.6-45.1-39.7-45.1-25.8 0-40 19.8-40 44.5v154.8c0 25.8 13.7 45.6 40.5 45.6 21.5 0 39.2-14 39.2-45.6v-41.8l60.6 75.7c12.3 14.9 39 16.8 55.8 0 14.6-15.1 14.8-36.8 4-50.4l-49.1-62.8 40.5-58.7c9.4-13.5 9.5-34.5-5.6-49.1-16.4-15.9-44.6-17.3-61.4 7l-44.8 64.7v-38.8z"],"kickstarter-k":[384,512,[],"f3bc","M147.3 114.4c0-56.2-32.5-82.4-73.4-82.4C26.2 32 0 68.2 0 113.4v283c0 47.3 25.3 83.4 74.9 83.4 39.8 0 72.4-25.6 72.4-83.4v-76.5l112.1 138.3c22.7 27.2 72.1 30.7 103.2 0 27-27.6 27.3-67.4 7.4-92.2l-90.8-114.8 74.9-107.4c17.4-24.7 17.5-63.1-10.4-89.8-30.3-29-82.4-31.6-113.6 12.8L147.3 185v-70.6z"],korvue:[446,512,[],"f42f","M386.5 34h-327C26.8 34 0 60.8 0 93.5v327.1C0 453.2 26.8 480 59.5 480h327.1c33 0 59.5-26.8 59.5-59.5v-327C446 60.8 419.2 34 386.5 34zM87.1 120.8h96v116l61.8-116h110.9l-81.2 132H87.1v-132zm161.8 272.1l-65.7-113.6v113.6h-96V262.1h191.5l88.6 130.8H248.9z"],laravel:[640,512,[],"f3bd","M637.5 241.6c-4.2-4.8-62.8-78.1-73.1-90.5-10.3-12.4-15.4-10.2-21.7-9.3-6.4.9-80.5 13.4-89.1 14.8-8.6 1.5-14 4.9-8.7 12.3 4.7 6.6 53.4 75.7 64.2 90.9l-193.7 46.4L161.2 48.7c-6.1-9.1-7.4-12.3-21.4-11.6-14 .6-120.9 9.5-128.5 10.2-7.6.6-16 4-8.4 22s129 279.6 132.4 287.2c3.4 7.6 12.2 20 32.8 15 21.1-5.1 94.3-24.2 134.3-34.7 21.1 38.3 64.2 115.9 72.2 127 10.6 14.9 18 12.4 34.3 7.4 12.8-3.9 199.6-71.1 208-74.5 8.4-3.5 13.6-5.9 7.9-14.4-4.2-6.2-53.5-72.2-79.3-106.8 17.7-4.7 80.6-21.4 87.3-23.3 7.9-2 9-5.8 4.7-10.6zm-352.2 72c-2.3.5-110.8 26.5-116.6 27.8-5.8 1.3-5.8.7-6.5-1.3-.7-2-129-266.7-130.8-270-1.8-3.3-1.7-5.9 0-5.9s102.5-9 106-9.2c3.6-.2 3.2.6 4.5 2.8 0 0 142.2 245.4 144.6 249.7 2.6 4.3 1.1 5.6-1.2 6.1zm306 57.4c1.7 2.7 3.5 4.5-2 6.4-5.4 2-183.7 62.1-187.1 63.6-3.5 1.5-6.2 2-10.6-4.5s-62.4-106.8-62.4-106.8L518 280.6c4.7-1.5 6.2-2.5 9.2 2.2 2.9 4.8 62.4 85.5 64.1 88.2zm12.1-134.1c-4.2.9-73.6 18.1-73.6 18.1l-56.7-77.8c-1.6-2.3-2.9-4.5 1.1-5s68.4-12.2 71.3-12.8c2.9-.7 5.4-1.5 9 3.4 3.6 4.9 52.6 67 54.5 69.4 1.8 2.3-1.4 3.7-5.6 4.7z"],lastfm:[512,512,[],"f202","M225.8 367.1l-18.8-51s-30.5 34-76.2 34c-40.5 0-69.2-35.2-69.2-91.5 0-72.1 36.4-97.9 72.1-97.9 66.5 0 74.8 53.3 100.9 134.9 18.8 56.9 54 102.6 155.4 102.6 72.7 0 122-22.3 122-80.9 0-72.9-62.7-80.6-115-92.1-25.8-5.9-33.4-16.4-33.4-34 0-19.9 15.8-31.7 41.6-31.7 28.2 0 43.4 10.6 45.7 35.8l58.6-7c-4.7-52.8-41.1-74.5-100.9-74.5-52.8 0-104.4 19.9-104.4 83.9 0 39.9 19.4 65.1 68 76.8 44.9 10.6 79.8 13.8 79.8 45.7 0 21.7-21.1 30.5-61 30.5-59.2 0-83.9-31.1-97.9-73.9-32-96.8-43.6-163-161.3-163C45.7 113.8 0 168.3 0 261c0 89.1 45.7 137.2 127.9 137.2 66.2 0 97.9-31.1 97.9-31.1z"],"lastfm-square":[448,512,[],"f203","M400 32H48C21.5 32 0 53.5 0 80v352c0 26.5 21.5 48 48 48h352c26.5 0 48-21.5 48-48V80c0-26.5-21.5-48-48-48zm-92.2 312.9c-63.4 0-85.4-28.6-97.1-64.1-16.3-51-21.5-84.3-63-84.3-22.4 0-45.1 16.1-45.1 61.2 0 35.2 18 57.2 43.3 57.2 28.6 0 47.6-21.3 47.6-21.3l11.7 31.9s-19.8 19.4-61.2 19.4c-51.3 0-79.9-30.1-79.9-85.8 0-57.9 28.6-92 82.5-92 73.5 0 80.8 41.4 100.8 101.9 8.8 26.8 24.2 46.2 61.2 46.2 24.9 0 38.1-5.5 38.1-19.1 0-19.9-21.8-22-49.9-28.6-30.4-7.3-42.5-23.1-42.5-48 0-40 32.3-52.4 65.2-52.4 37.4 0 60.1 13.6 63 46.6l-36.7 4.4c-1.5-15.8-11-22.4-28.6-22.4-16.1 0-26 7.3-26 19.8 0 11 4.8 17.6 20.9 21.3 32.7 7.1 71.8 12 71.8 57.5.1 36.7-30.7 50.6-76.1 50.6z"],leanpub:[576,512,[],"f212","M386.539 111.485l15.096 248.955-10.979-.275c-36.232-.824-71.64 8.783-102.657 27.997-31.016-19.214-66.424-27.997-102.657-27.997-45.564 0-82.07 10.705-123.516 27.723L93.117 129.6c28.546-11.803 61.484-18.115 92.226-18.115 41.173 0 73.836 13.175 102.657 42.544 27.723-28.271 59.013-41.721 98.539-42.544zM569.07 448c-25.526 0-47.485-5.215-70.542-15.645-34.31-15.645-69.993-24.978-107.871-24.978-38.977 0-74.934 12.901-102.657 40.623-27.723-27.723-63.68-40.623-102.657-40.623-37.878 0-73.561 9.333-107.871 24.978C55.239 442.236 32.731 448 8.303 448H6.93L49.475 98.859C88.726 76.626 136.486 64 181.775 64 218.83 64 256.984 71.685 288 93.095 319.016 71.685 357.17 64 394.225 64c45.289 0 93.049 12.626 132.3 34.859L569.07 448zm-43.368-44.741l-34.036-280.246c-30.742-13.999-67.248-21.41-101.009-21.41-38.428 0-74.385 12.077-102.657 38.702-28.272-26.625-64.228-38.702-102.657-38.702-33.761 0-70.267 7.411-101.009 21.41L50.298 403.259c47.211-19.487 82.894-33.486 135.045-33.486 37.604 0 70.817 9.606 102.657 29.644 31.84-20.038 65.052-29.644 102.657-29.644 52.151 0 87.834 13.999 135.045 33.486z"],less:[640,512,[],"f41d","M612.7 219c0-20.5 3.2-32.6 3.2-54.6 0-34.2-12.6-45.2-40.5-45.2h-20.5v24.2h6.3c14.2 0 17.3 4.7 17.3 22.1 0 16.3-1.6 32.6-1.6 51.5 0 24.2 7.9 33.6 23.6 37.3v1.6c-15.8 3.7-23.6 13.1-23.6 37.3 0 18.9 1.6 34.2 1.6 51.5 0 17.9-3.7 22.6-17.3 22.6v.5h-6.3V393h20.5c27.8 0 40.5-11 40.5-45.2 0-22.6-3.2-34.2-3.2-54.6 0-11 6.8-22.6 27.3-23.6v-27.3c-20.5-.7-27.3-12.3-27.3-23.3zm-105.6 32c-15.8-6.3-30.5-10-30.5-20.5 0-7.9 6.3-12.6 17.9-12.6s22.1 4.7 33.6 13.1l21-27.8c-13.1-10-31-20.5-55.2-20.5-35.7 0-59.9 20.5-59.9 49.4 0 25.7 22.6 38.9 41.5 46.2 16.3 6.3 32.1 11.6 32.1 22.1 0 7.9-6.3 13.1-20.5 13.1-13.1 0-26.3-5.3-40.5-16.3l-21 30.5c15.8 13.1 39.9 22.1 59.9 22.1 42 0 64.6-22.1 64.6-51s-22.5-41-43-47.8zm-358.9 59.4c-3.7 0-8.4-3.2-8.4-13.1V119.1H65.2c-28.4 0-41 11-41 45.2 0 22.6 3.2 35.2 3.2 54.6 0 11-6.8 22.6-27.3 23.6v27.3c20.5.5 27.3 12.1 27.3 23.1 0 19.4-3.2 31-3.2 53.6 0 34.2 12.6 45.2 40.5 45.2h20.5v-24.2h-6.3c-13.1 0-17.3-5.3-17.3-22.6s1.6-32.1 1.6-51.5c0-24.2-7.9-33.6-23.6-37.3v-1.6c15.8-3.7 23.6-13.1 23.6-37.3 0-18.9-1.6-34.2-1.6-51.5s3.7-22.1 17.3-22.1H93v150.8c0 32.1 11 53.1 43.1 53.1 10 0 17.9-1.6 23.6-3.7l-5.3-34.2c-3.1.8-4.6.8-6.2.8zM379.9 251c-16.3-6.3-31-10-31-20.5 0-7.9 6.3-12.6 17.9-12.6 11.6 0 22.1 4.7 33.6 13.1l21-27.8c-13.1-10-31-20.5-55.2-20.5-35.7 0-59.9 20.5-59.9 49.4 0 25.7 22.6 38.9 41.5 46.2 16.3 6.3 32.1 11.6 32.1 22.1 0 7.9-6.3 13.1-20.5 13.1-13.1 0-26.3-5.3-40.5-16.3l-20.5 30.5c15.8 13.1 39.9 22.1 59.9 22.1 42 0 64.6-22.1 64.6-51 .1-28.9-22.5-41-43-47.8zm-155-68.8c-38.4 0-75.1 32.1-74.1 82.5 0 52 34.2 82.5 79.3 82.5 18.9 0 39.9-6.8 56.2-17.9l-15.8-27.8c-11.6 6.8-22.6 10-34.2 10-21 0-37.3-10-41.5-34.2H290c.5-3.7 1.6-11 1.6-19.4.6-42.6-22.6-75.7-66.7-75.7zm-30 66.2c3.2-21 15.8-31 30.5-31 18.9 0 26.3 13.1 26.3 31h-56.8z"],line:[448,512,[],"f3c0","M272.1 204.2v71.1c0 1.8-1.4 3.2-3.2 3.2h-11.4c-1.1 0-2.1-.6-2.6-1.3l-32.6-44v42.2c0 1.8-1.4 3.2-3.2 3.2h-11.4c-1.8 0-3.2-1.4-3.2-3.2v-71.1c0-1.8 1.4-3.2 3.2-3.2H219c1 0 2.1.5 2.6 1.4l32.6 44v-42.2c0-1.8 1.4-3.2 3.2-3.2h11.4c1.8-.1 3.3 1.4 3.3 3.1zm-82-3.2h-11.4c-1.8 0-3.2 1.4-3.2 3.2v71.1c0 1.8 1.4 3.2 3.2 3.2h11.4c1.8 0 3.2-1.4 3.2-3.2v-71.1c0-1.7-1.4-3.2-3.2-3.2zm-27.5 59.6h-31.1v-56.4c0-1.8-1.4-3.2-3.2-3.2h-11.4c-1.8 0-3.2 1.4-3.2 3.2v71.1c0 .9.3 1.6.9 2.2.6.5 1.3.9 2.2.9h45.7c1.8 0 3.2-1.4 3.2-3.2v-11.4c0-1.7-1.4-3.2-3.1-3.2zM332.1 201h-45.7c-1.7 0-3.2 1.4-3.2 3.2v71.1c0 1.7 1.4 3.2 3.2 3.2h45.7c1.8 0 3.2-1.4 3.2-3.2v-11.4c0-1.8-1.4-3.2-3.2-3.2H301v-12h31.1c1.8 0 3.2-1.4 3.2-3.2V234c0-1.8-1.4-3.2-3.2-3.2H301v-12h31.1c1.8 0 3.2-1.4 3.2-3.2v-11.4c-.1-1.7-1.5-3.2-3.2-3.2zM448 113.7V399c-.1 44.8-36.8 81.1-81.7 81H81c-44.8-.1-81.1-36.9-81-81.7V113c.1-44.8 36.9-81.1 81.7-81H367c44.8.1 81.1 36.8 81 81.7zm-61.6 122.6c0-73-73.2-132.4-163.1-132.4-89.9 0-163.1 59.4-163.1 132.4 0 65.4 58 120.2 136.4 130.6 19.1 4.1 16.9 11.1 12.6 36.8-.7 4.1-3.3 16.1 14.1 8.8 17.4-7.3 93.9-55.3 128.2-94.7 23.6-26 34.9-52.3 34.9-81.5z"],linkedin:[448,512,[],"f08c","M416 32H31.9C14.3 32 0 46.5 0 64.3v383.4C0 465.5 14.3 480 31.9 480H416c17.6 0 32-14.5 32-32.3V64.3c0-17.8-14.4-32.3-32-32.3zM135.4 416H69V202.2h66.5V416zm-33.2-243c-21.3 0-38.5-17.3-38.5-38.5S80.9 96 102.2 96c21.2 0 38.5 17.3 38.5 38.5 0 21.3-17.2 38.5-38.5 38.5zm282.1 243h-66.4V312c0-24.8-.5-56.7-34.5-56.7-34.6 0-39.9 27-39.9 54.9V416h-66.4V202.2h63.7v29.2h.9c8.9-16.8 30.6-34.5 62.9-34.5 67.2 0 79.7 44.3 79.7 101.9V416z"],"linkedin-in":[448,512,[],"f0e1","M100.3 480H7.4V180.9h92.9V480zM53.8 140.1C24.1 140.1 0 115.5 0 85.8 0 56.1 24.1 32 53.8 32c29.7 0 53.8 24.1 53.8 53.8 0 29.7-24.1 54.3-53.8 54.3zM448 480h-92.7V334.4c0-34.7-.7-79.2-48.3-79.2-48.3 0-55.7 37.7-55.7 76.7V480h-92.8V180.9h89.1v40.8h1.3c12.4-23.5 42.7-48.3 87.9-48.3 94 0 111.3 61.9 111.3 142.3V480z"],linode:[448,512,[],"f2b8","M437.4 226.3c-.3-.9-.9-1.4-1.4-2l-70-38.6c-.9-.6-2-.6-3.1 0l-58.9 36c-.9.6-1.4 1.7-1.4 2.6l-.9 31.4-24-16c-.9-.6-2.3-.6-3.1 0L240 260.9l-1.4-35.1c0-.9-.6-2-1.4-2.3l-36-24.3 33.7-17.4c1.1-.6 1.7-1.7 1.7-2.9l-5.7-132.3c0-.9-.9-2-1.7-2.6L138.6.3c-.9-.3-1.7-.3-2.3-.3L12.6 38.6c-1.4.6-2.3 2-2 3.7L38 175.4c.9 3.4 34 27.4 38.6 30.9l-26.9 12.9c-1.4.9-2 2.3-1.7 3.4l20.6 100.3c.6 2.9 23.7 23.1 27.1 26.3l-17.4 10.6c-.9.6-1.7 2-1.4 3.1 1.4 7.1 15.4 77.7 16.9 79.1l65.1 69.1c.6.6 1.4.6 2.3.9.6 0 1.1-.3 1.7-.6l83.7-66.9c.9-.6 1.1-1.4 1.1-2.3l-2-46 28 23.7c1.1.9 2.9.9 4 0l66.9-53.4c.9-.6 1.1-1.4 1.1-2.3l2.3-33.4 20.3 14c1.1.9 2.6.9 3.7 0l54.6-43.7c.6-.3 1.1-1.1 1.1-2 .9-6.5 10.3-70.8 9.7-72.8zm-204.8 4.8l4 92.6-90.6 61.2-14-96.6 100.6-57.2zm-7.7-180l5.4 126-106.6 55.4L104 97.7l120.9-46.6zM44 173.1L18 48l79.7 49.4 19.4 132.9L44 173.1zm30.6 147.8L55.7 230l70 58.3 13.7 93.4-64.8-60.8zm24.3 117.7l-13.7-67.1 61.7 60.9 9.7 67.4-57.7-61.2zm64.5 64.5l-10.6-70.9 85.7-61.4 3.1 70-78.2 62.3zm82-115.1c0-3.4.9-22.9-2-25.1l-24.3-20 22.3-14.9c2.3-1.7 1.1-5.7 1.1-8l29.4 22.6.6 68.3-27.1-22.9zm94.3-25.4l-60.9 48.6-.6-68.6 65.7-46.9-4.2 66.9zm27.7-25.7l-19.1-13.4 2-34c.3-.9-.3-2-1.1-2.6L308 259.7l.6-30 64.6 40.6-5.8 66.6zm54.6-39.8l-48.3 38.3 5.7-65.1 51.1-36.6-8.5 63.4z"],linux:[448,512,[],"f17c","M196.1 123.6c-.2-1.4 1.9-2.3 3.2-2.9 1.7-.7 3.9-1 5.5-.1.4.2.8.7.6 1.1-.4 1.2-2.4 1-3.5 1.6-1 .5-1.8 1.7-3 1.7-1 .1-2.7-.4-2.8-1.4zm24.7-.3c1 .5 1.8 1.7 3 1.7 1.1 0 2.8-.4 2.9-1.5.2-1.4-1.9-2.3-3.2-2.9-1.7-.7-3.9-1-5.5-.1-.4.2-.8.7-.6 1.1.3 1.3 2.3 1.1 3.4 1.7zm214.7 310.2c-.5 8.2-6.5 13.8-13.9 18.3-14.9 9-37.3 15.8-50.9 32.2l-2.6-2.2 2.6 2.2c-14.2 16.9-31.7 26.6-48.3 27.9-16.5 1.3-32-6.3-40.3-23v-.1c-1.1-2.1-1.9-4.4-2.5-6.7-21.5 1.2-40.2-5.3-55.1-4.1-22 1.2-35.8 6.5-48.3 6.6-4.8 10.6-14.3 17.6-25.9 20.2-16 3.7-36.1 0-55.9-10.4l1.6-3-1.6 3c-18.5-9.8-42-8.9-59.3-12.5-8.7-1.8-16.3-5-20.1-12.3-3.7-7.3-3-17.3 2.2-31.7 1.7-5.1.4-12.7-.8-20.8-.6-3.9-1.2-7.9-1.2-11.8 0-4.3.7-8.5 2.8-12.4 4.5-8.5 11.8-12.1 18.5-14.5 6.7-2.4 12.8-4 17-8.3 5.2-5.5 10.1-14.4 16.6-20.2-2.6-17.2.2-35.4 6.2-53.3 12.6-37.9 39.2-74.2 58.1-96.7 16.1-22.9 20.8-41.3 22.5-64.7C158 103.4 132.4-.2 234.8 0c80.9.1 76.3 85.4 75.8 131.3-.3 30.1 16.3 50.5 33.4 72 15.2 18 35.1 44.3 46.5 74.4 9.3 24.6 12.9 51.8 3.7 79.1 1.4.5 2.8 1.2 4.1 2 1.4.8 2.7 1.8 4 2.9 6.6 5.6 8.7 14.3 10.5 22.4 1.9 8.1 3.6 15.7 7.2 19.7 11.1 12.4 15.9 21.5 15.5 29.7zM220.8 109.1c3.6.9 8.9 2.4 13 4.4-2.1-12.2 4.5-23.5 11.8-23 8.9.3 13.9 15.5 9.1 27.3-.8 1.9-2.8 3.4-3.9 4.6 6.7 2.3 11 4.1 12.6 4.9 7.9-9.5 10.8-26.2 4.3-40.4-9.8-21.4-34.2-21.8-44 .4-3.2 7.2-3.9 14.9-2.9 21.8zm-46.2 18.8c7.8-5.7 6.9-4.7 5.9-5.5-8-6.9-6.6-27.4 1.8-28.1 6.3-.5 10.8 10.7 9.6 19.6 3.1-2.1 6.7-3.6 10.2-4.6 1.7-19.3-9-33.5-19.1-33.5-18.9 0-24 37.5-8.4 52.1zm-9.4 20.9c1.5 4.9 6.1 10.5 14.7 15.3 7.8 4.6 12 11.5 20 15 2.6 1.1 5.7 1.9 9.6 2.1 18.4 1.1 27.1-11.3 38.2-14.9 11.7-3.7 20.1-11 22.7-18.1 3.2-8.5-2.1-14.7-10.5-18.2-11.3-4.9-16.3-5.2-22.6-9.3-10.3-6.6-18.8-8.9-25.9-8.9-14.4 0-23.2 9.8-27.9 14.2-.5.5-7.9 5.9-14.1 10.5-4.2 3.3-5.6 7.4-4.2 12.3zm-33.5 252.8L112.1 366c-6.8-9.2-13.8-14.8-21.9-16-7.7-1.2-12.6 1.4-17.7 6.9-4.8 5.1-8.8 12.3-14.3 18-7.8 6.5-9.3 6.2-19.6 9.9-6.3 2.2-11.3 4.6-14.8 11.3-2.7 5-2.1 12.2-.9 20 1.2 7.9 3 16.3.6 23.9v.2c-5 13.7-5 21.7-2.6 26.4 7.9 15.4 46.6 6.1 76.5 21.9 31.4 16.4 72.6 17.1 75.3-18 2.1-20.5-31.5-49-41-68.9zm153.9 35.8c3.2-11 6.3-21.3 6.8-29 .8-15.2 1.6-28.7 4.4-39.9 3.1-12.6 9.3-23.1 21.4-27.3 2.3-21.1 18.7-21.1 38.3-12.5 18.9 8.5 26 16 22.8 26.1 1 0 2-.1 4.2 0 5.2-16.9-14.3-28-30.7-34.8 2.9-12 2.4-24.1-.4-35.7-6-25.3-22.6-47.8-35.2-59-2.3-.1-2.1 1.9 2.6 6.5 11.6 10.7 37.1 49.2 23.3 84.9-3.9-1-7.6-1.5-10.9-1.4-5.3-29.1-17.5-53.2-23.6-64.6-11.5-21.4-29.5-65.3-37.2-95.7-4.5 6.4-12.4 11.9-22.3 15-4.7 1.5-9.7 5.5-15.9 9-13.9 8-30 8.8-42.4-1.2-4.5-3.6-8-7.6-12.6-10.3-1.6-.9-5.1-3.3-6.2-4.1-2 37.8-27.3 85.3-39.3 112.7-8.3 19.7-13.2 40.8-13.8 61.5-21.8-29.1-5.9-66.3 2.6-82.4 9.5-17.6 11-22.5 8.7-20.8-8.6 14-22 36.3-27.2 59.2-2.7 11.9-3.2 24 .3 35.2 3.5 11.2 11.1 21.5 24.6 29.9 0 0 24.8 14.3 38.3 32.5 7.4 10 9.7 18.7 7.4 24.9-2.5 6.7-9.6 8.9-16.7 8.9 4.8 6 10.3 13 14.4 19.6 37.6 25.7 82.2 15.7 114.3-7.2zM415 408.5c-10-11.3-7.2-33.1-17.1-41.6-6.9-6-13.6-5.4-22.6-5.1-7.7 8.8-25.8 19.6-38.4 16.3-11.5-2.9-18-16.3-18.8-29.5-.3.2-.7.3-1 .5-7.1 3.9-11.1 10.8-13.7 21.1-2.5 10.2-3.4 23.5-4.2 38.7-.7 11.8-6.2 26.4-9.9 40.6-3.5 13.2-5.8 25.2-1.1 36.3 7.2 14.5 19.5 20.4 33.7 19.3 14.2-1.1 30.4-9.8 43.6-25.5 22-26.6 62.3-29.7 63.2-46.5.3-5.1-3.1-13-13.7-24.6zM173.3 148.7c2 1.9 4.7 4.5 8 7.1 6.6 5.2 15.8 10.6 27.3 10.6 11.6 0 22.5-5.9 31.8-10.8 4.9-2.6 10.9-7 14.8-10.4 3.9-3.4 5.9-6.3 3.1-6.6-2.8-.3-2.6 2.6-6 5.1-4.4 3.2-9.7 7.4-13.9 9.8-7.4 4.2-19.5 10.2-29.9 10.2-10.4 0-18.7-4.8-24.9-9.7-3.1-2.5-5.7-5-7.7-6.9-1.5-1.4-1.9-4.6-4.3-4.9-1.4-.1-1.8 3.7 1.7 6.5z"],lyft:[512,512,[],"f3c3","M0 81.1h77.8v208.7c0 33.1 15 52.8 27.2 61-12.7 11.1-51.2 20.9-80.2-2.8C7.8 334 0 310.7 0 289V81.1zm485.9 173.5v-22h23.8v-76.8h-26.1c-10.1-46.3-51.2-80.7-100.3-80.7-56.6 0-102.7 46-102.7 102.7V357c16 2.3 35.4-.3 51.7-14 17.1-14 24.8-37.2 24.8-59v-6.7h38.8v-76.8h-38.8v-23.3c0-34.6 52.2-34.6 52.2 0v77.1c0 56.6 46 102.7 102.7 102.7v-76.5c-14.5 0-26.1-11.7-26.1-25.9zm-294.3-99v113c0 15.4-23.8 15.4-23.8 0v-113H91v132.7c0 23.8 8 54 45 63.9 37 9.8 58.2-10.6 58.2-10.6-2.1 13.4-14.5 23.3-34.9 25.3-15.5 1.6-35.2-3.6-45-7.8v70.3c25.1 7.5 51.5 9.8 77.6 4.7 47.1-9.1 76.8-48.4 76.8-100.8V155.1h-77.1v.5z"],magento:[448,512,[],"f3c4","M445.7 127.9V384l-63.4 36.5V164.7L223.8 73.1 65.2 164.7l.4 255.9L2.3 384V128.1L224.2 0l221.5 127.9zM255.6 420.5L224 438.9l-31.8-18.2v-256l-63.3 36.6.1 255.9 94.9 54.9 95.1-54.9v-256l-63.4-36.6v255.9z"],mandalorian:[390,512,[],"f50f","M203.28 511.89c-.98-3.26-1.69-15.83-1.39-24.58.55-15.89.98-24.72 1.4-28.76.64-6.2 2.87-20.72 3.28-21.38.6-.96.4-27.87-.24-33.13-.31-2.58-.63-11.9-.69-20.73-.13-16.47-.53-20.12-2.73-24.76-1.1-2.32-1.23-3.84-.99-11.43.16-4.81 0-10.53-.34-12.71-2.05-12.97-3.46-27.7-3.25-33.9.21-6.12.43-7.15 2.06-9.67 3.05-4.71 6.51-14.04 8.62-23.27 2.26-9.86 3.88-17.18 4.59-20.74.89-4.42 2.43-9.72 4.36-15.05 2.27-6.25 2.49-15.39.37-15.39-.3 0-1.38 1.22-2.41 2.71-1.03 1.49-4.76 4.8-8.29 7.36-8.37 6.08-11.7 9.39-12.66 12.58-.93 3.11-1.02 7.23-.16 7.76.34.21 1.29 2.4 2.11 4.88 1.62 4.88 1.87 10.12.72 15.36-.39 1.77-1.05 5.47-1.46 8.23-.41 2.76-.98 6.46-1.25 8.22-.28 1.76-.97 3.68-1.55 4.26-.96.96-1.14.91-2.05-.53-.55-.87-1.2-3.01-1.44-4.75-.25-1.74-1.63-7.11-3.08-11.93-3.28-10.9-3.52-16.15-.96-20.96.92-1.73 1.67-3.81 1.67-4.61 0-2.39-2.2-5.32-7.41-9.89-7.05-6.18-8.63-7.92-10.23-11.3-1.71-3.6-3.06-4.06-4.54-1.54-1.78 3.01-2.6 9.11-2.97 22.02l-.35 12.13 1.95 2.25c3.21 3.7 12.07 16.45 13.78 19.83 3.41 6.74 4.34 11.69 4.41 23.56.07 11.84.95 22.75 2 24.71.36.66.51 1.35.34 1.52-.17.17.41 2.09 1.29 4.27.88 2.18 1.81 6.22 2.06 8.98.25 2.76 1.02 7.43 1.71 10.37 2.23 9.56 2.77 14.08 2.39 20.14-.2 3.27-.53 11.07-.73 17.32-1.31 41.76-1.85 57.98-2.04 61.21-.12 2.02-.39 11.51-.6 21.07-.36 16.3-1.3 27.37-2.42 28.65-.64.73-8.07-4.91-12.52-9.49-3.75-3.87-4.02-4.79-2.83-9.95.7-3.01 2.26-18.29 3.33-32.62.36-4.78.81-10.5 1.01-12.71.83-9.37 1.66-20.35 2.61-34.78.56-8.46 1.33-16.44 1.72-17.73.38-1.29.89-9.89 1.13-19.11l.43-16.77-2.26-4.3c-1.72-3.28-4.87-6.94-13.22-15.34-6.03-6.07-11.84-12.3-12.91-13.85l-1.95-2.81.75-10.9c1.09-15.71 1.1-48.57.02-59.06l-.89-8.7-3.28-4.52c-5.86-8.08-5.8-7.75-6.22-33.27-.1-6.07-.38-11.5-.63-12.06-.83-1.87-3.05-2.66-8.54-3.05-8.86-.62-10.96-1.9-23.85-14.55-6.15-6.04-12.34-11.97-13.75-13.19-2.81-2.42-2.79-1.99-.56-9.63l1.35-4.65-1.69-3.04c-.93-1.67-2.09-3.51-2.59-4.07-1.33-1.51-5.5-10.89-5.99-13.49-.31-1.66-.09-2.67.87-3.9 2.23-2.86 3.4-5.68 4.45-10.73 2.33-11.19 7.74-26.09 10.6-29.22 3.18-3.47 7.7-1.05 9.41 5.03 1.34 4.79 1.37 9.79.1 18.55-.53 3.68-.98 8.68-.99 11.11-.02 4.01.19 4.69 2.25 7.39 3.33 4.37 7.73 7.41 15.2 10.52 1.7.71 3.82 1.99 4.72 2.85 11.17 10.72 18.62 16.18 22.95 16.85 5.18.8 7.98 4.54 10.04 13.39 1.31 5.65 4 11.14 5.46 11.14.59 0 2.09-.63 3.33-1.39 1.98-1.22 2.25-1.73 2.25-4.18-.01-3.71-1.17-14.08-2-17.84-.37-1.66-.78-4.06-.93-5.35-.14-1.29-.61-3.85-1.03-5.69-2.55-11.16-3.65-15.46-4.1-16.05-1.55-2.02-4.08-10.2-4.93-15.92-1.64-11.11-3.96-14.23-12.91-17.39-4.64-1.64-8.89-4.12-13.32-7.78-1.15-.95-4.01-3.22-6.35-5.06-2.35-1.83-4.41-3.53-4.6-3.76-.18-.23-1.39-1.14-2.69-2.02-6.24-4.22-8.84-6.98-11.26-11.96l-2.44-5.02-.22-12.98-.22-12.98 6.91-6.55c3.95-3.75 8.48-7.35 10.59-8.43 3.31-1.69 4.45-1.89 11.37-2.05 8.53-.19 10.12.02 11.66 1.56 1.53 1.53 1.36 6.4-.29 8.5-.74.94-1.34 1.98-1.34 2.32 0 .58-2.61 4.91-5.42 8.99-.68.99-2.13 5.35-2.37 6.82 20.44 13.39 21.55 3.77 14.07 28.98l11.4 2.54c3.11-8.66 6.47-17.26 8.61-26.22.29-7.63-11.98-4.19-15.4-8.68-2.33-5.93 3.13-14.18 6.06-19.2 1.6-2.34 6.62-4.7 8.82-4.15.88.22 4.16-.35 7.37-1.28 3.18-.92 6.58-1.68 7.55-1.68.97 0 3.66-.58 5.98-1.29 3.65-1.11 4.5-1.17 6.35-.4 1.17.48 3.79 1.09 5.82 1.36 2.02.26 4.72 1.12 6 1.91 1.28.79 3.53 1.77 5.02 2.17 2.51.68 3 .57 7.05-1.67l4.35-2.4 10.7-.41c10.44-.4 10.81-.47 15.26-2.68l4.58-2.3 2.46 1.43c1.76 1.02 3.14 2.73 4.85 5.98 2.36 4.51 2.38 4.58 1.37 7.37-.88 2.44-.89 3.3-.1 6.39.5 1.96 1.45 4.62 2.1 5.91.65 1.29 1.24 3.09 1.31 4.01.31 4.33-.03 5.3-2.41 6.92-2.17 1.47-6.98 7.91-6.98 9.34 0 .32-.48 1.69-1.07 3.03-5.04 11.51-6.76 13.56-14.26 16.98-9.2 4.2-12.3 5.19-16.21 5.19-3.1 0-4 .25-4.54 1.26-.37.69-2.21 2.37-4.09 3.71-2.04 1.47-3.8 3.38-4.38 4.78-.54 1.28-1.66 2.59-2.49 2.91-.83.32-1.94 1.08-2.45 1.71-.52.62-3.66 3.04-7 5.38-3.33 2.34-6.87 5.02-7.87 5.96-1 .94-2.07 1.71-2.39 1.71s-1.28.74-2.13 1.65c-1.31 1.39-1.49 2.11-1.14 4.6.22 1.63.86 4.27 1.42 5.88 1.32 3.8 1.31 7.86-.05 10.57-1.43 2.86-.89 6.65 1.35 9.59 2.01 2.63 2.16 4.56.71 8.84-.61 1.8-1.05 5.45-1.06 8.91-.02 4.88.22 6.28 1.46 8.38 1.2 2.04 1.82 2.48 3.24 2.32 1.98-.23 2.3-1.05 4.71-12.12 2.18-10.03 3.71-11.92 13.76-17.08 2.94-1.51 7.46-3.96 10.03-5.44 2.58-1.48 6.79-3.69 9.37-4.91 6.67-3.16 11.05-6.52 15.22-11.67 7.11-8.79 9.98-16.22 12.85-33.3.55-3.28 1.43-5.65 2.86-7.73 1.29-1.87 2.37-4.62 2.89-7.31 1.02-5.3 2.85-9.08 5.58-11.51 4.7-4.18 6-1.09 4.59 10.87-.46 3.86-1.1 10.33-1.44 14.38l-.61 7.36 4.45 4.09 4.45 4.09.11 8.42c.06 4.63.47 9.53.92 10.89l.82 2.47-6.43 6.28c-8.54 8.33-12.88 13.93-16.76 21.61-1.77 3.49-3.74 7.11-4.38 8.03-2.18 3.11-6.46 13.01-8.76 20.26l-2.29 7.22-6.97 6.49c-3.83 3.57-7.96 7.25-9.17 8.17-3.05 2.32-4.26 5.15-4.26 9.99 0 2.98.43 4.96 1.59 7.26.87 1.74 1.81 3.91 2.09 4.83.28.92.98 2.22 1.57 2.89 1.4 1.59 1.92 16.12.83 23.22-.68 4.48-3.63 12.02-4.7 12.02-1.79 0-4.06 9.27-5.07 20.74-.18 2.02-.62 5.94-.98 8.7-.36 2.76-.96 9.98-1.35 16.05-.77 12.22-.19 18.77 2.05 23.15 3.41 6.69.52 12.69-11.03 22.84l-3.97 3.49.07 5.19c.04 2.86.55 6.85 1.14 8.87 4.61 15.98 4.73 16.92 4.38 37.13-.46 26.4-.26 40.27.63 44.15.42 1.84.91 5 1.08 7.02.17 2.02.66 5.33 1.08 7.36.47 2.26.78 11.02.79 22.74l.02 19.06-1.81 2.63c-2.71 3.91-15.11 13.54-15.49 12.29zm29.53-45.11c-.18-.3-.33-6.87-.33-14.59 0-14.06-.89-27.54-2.26-34.45-.4-2.02-.81-9.7-.9-17.06-.15-11.93-1.4-24.37-2.64-26.38-.66-1.07-3.02-17.66-3.03-21.3-.01-4.23 1.02-6 5.28-9.13 4.14-3.04 4.86-3.14 5.48-.72.28 1.1 1.45 5.62 2.6 10.03 3.93 15.12 4.14 16.27 4.05 21.74-.1 5.78-.13 6.13-1.74 17.73-.98 7.07-1.17 12.39-1.04 28.43.17 19.4-.64 35.73-2.04 41.27-.71 2.78-2.8 5.48-3.43 4.43zm-70.99-37.58c-.24-.38-1.01-5.24-1.73-10.79-.72-5.56-1.49-10.41-1.73-10.79-.23-.38-.68-3.3-.99-6.49-.31-3.19-.91-7.46-1.33-9.48-.99-4.79-3.35-19.35-3.42-21.07-.03-.74-.34-4.05-.7-7.36-.67-6.21-.84-27.67-.22-28.29.96-.96 6.63 2.76 11.33 7.43l5.28 5.25-.45 6.47c-.25 3.56-.6 10.23-.78 14.83-.18 4.6-.49 9.87-.67 11.71-.18 1.84-.61 9.36-.94 16.72-.79 17.41-1.94 31.29-2.65 32-.32.3-.76.24-1-.14zM74.63 162.61c21.07 12.79 17.84 14.15 28.49 17.66 13.01 4.29 18.87 7.13 23.15 16.87-43.66 36.14-69.01 57.9-76.71 70.82-31.02 52.01-5.99 101.59 62.75 87.21-14.18 29.23-77.97 28.63-98.68-4.9-24.68-39.95-22.09-118.3 61-187.66zm210.79 179.02c56.66 6.88 82.32-37.74 46.54-89.23 0 0-26.87-29.34-64.28-67.96 2.98-15.45 9.49-32.12 30.57-53.82 89.2 63.51 92 141.61 92.46 149.36 4.27 70.58-78.66 91.12-105.29 61.65z"],mastodon:[417,512,[],"f4f6","M417.8 179.1c0-97.2-63.7-125.7-63.7-125.7-62.5-28.7-228.5-28.4-290.4 0 0 0-63.7 28.5-63.7 125.7 0 115.7-6.6 259.4 105.6 289.1 40.5 10.7 75.3 13 103.3 11.4 50.8-2.8 79.3-18.1 79.3-18.1l-1.7-36.9s-36.3 11.4-77.1 10.1c-40.4-1.4-83-4.4-89.6-54-.6-4.4-.9-9-.9-13.9 85.6 20.9 158.6 9.1 178.7 6.7 56.1-6.7 105-41.3 111.2-72.9 9.8-49.8 9-121.5 9-121.5zm-75.1 125.2h-46.6V190.1c0-49.7-64-51.6-64 6.9v62.5h-46.3V197c0-58.5-64-56.6-64-6.9v114.2H75.1c0-122.1-5.2-147.9 18.4-175 25.9-28.9 79.8-30.8 103.8 6.1l11.6 19.5 11.6-19.5c24.1-37.1 78.1-34.8 103.8-6.1 23.7 27.3 18.4 53 18.4 175z"],maxcdn:[512,512,[],"f136","M461.1 442.7h-97.4L415.6 200c2.3-10.2.9-19.5-4.4-25.7-5-6.1-13.7-9.6-24.2-9.6h-49.3l-59.5 278h-97.4l59.5-278h-83.4l-59.5 278H0l59.5-278-44.6-95.4H387c39.4 0 75.3 16.3 98.3 44.9 23.3 28.6 31.8 67.4 23.6 105.9l-47.8 222.6z"],medapps:[320,512,[],"f3c6","M118.3 238.4c3.5-12.5 6.9-33.6 13.2-33.6 8.3 1.8 9.6 23.4 18.6 36.6 4.6-23.5 5.3-85.1 14.1-86.7 9-.7 19.7 66.5 22 77.5 9.9 4.1 48.9 6.6 48.9 6.6 1.9 7.3-24 7.6-40 7.8-4.6 14.8-5.4 27.7-11.4 28-4.7.2-8.2-28.8-17.5-49.6l-9.4 65.5c-4.4 13-15.5-22.5-21.9-39.3-3.3-.1-62.4-1.6-47.6-7.8l31-5zM228 448c21.2 0 21.2-32 0-32H92c-21.2 0-21.2 32 0 32h136zm-24 64c21.2 0 21.2-32 0-32h-88c-21.2 0-21.2 32 0 32h88zm34.2-141.5c3.2-18.9 5.2-36.4 11.9-48.8 7.9-14.7 16.1-28.1 24-41 24.6-40.4 45.9-75.2 45.9-125.5C320 69.6 248.2 0 160 0S0 69.6 0 155.2c0 50.2 21.3 85.1 45.9 125.5 7.9 12.9 16 26.3 24 41 6.7 12.5 8.7 29.8 11.9 48.9 3.5 21 36.1 15.7 32.6-5.1-3.6-21.7-5.6-40.7-15.3-58.6C66.5 246.5 33 211.3 33 155.2 33 87.3 90 32 160 32s127 55.3 127 123.2c0 56.1-33.5 91.3-66.1 151.6-9.7 18-11.7 37.4-15.3 58.6-3.4 20.6 29 26.4 32.6 5.1z"],medium:[448,512,[],"f23a","M0 32v448h448V32H0zm372.2 106.1l-24 23c-2.1 1.6-3.1 4.2-2.7 6.7v169.3c-.4 2.6.6 5.2 2.7 6.7l23.5 23v5.1h-118V367l24.3-23.6c2.4-2.4 2.4-3.1 2.4-6.7V199.8l-67.6 171.6h-9.1L125 199.8v115c-.7 4.8 1 9.7 4.4 13.2l31.6 38.3v5.1H71.2v-5.1l31.6-38.3c3.4-3.5 4.9-8.4 4.1-13.2v-133c.4-3.7-1-7.3-3.8-9.8L75 138.1V133h87.3l67.4 148L289 133.1h83.2v5z"],"medium-m":[512,512,[],"f3c7","M71.5 142.3c.6-5.9-1.7-11.8-6.1-15.8L20.3 72.1V64h140.2l108.4 237.7L364.2 64h133.7v8.1l-38.6 37c-3.3 2.5-5 6.7-4.3 10.8v272c-.7 4.1 1 8.3 4.3 10.8l37.7 37v8.1H307.3v-8.1l39.1-37.9c3.8-3.8 3.8-5 3.8-10.8V171.2L241.5 447.1h-14.7L100.4 171.2v184.9c-1.1 7.8 1.5 15.6 7 21.2l50.8 61.6v8.1h-144v-8L65 377.3c5.4-5.6 7.9-13.5 6.5-21.2V142.3z"],medrt:[544,512,[],"f3c8","M113.7 256c0 121.8 83.9 222.8 193.5 241.1-18.7 4.5-38.2 6.9-58.2 6.9C111.4 504 0 393 0 256S111.4 8 248.9 8c20.1 0 39.6 2.4 58.2 6.9C197.5 33.2 113.7 134.2 113.7 256m297.4 100.3c-77.7 55.4-179.6 47.5-240.4-14.6 5.5 14.1 12.7 27.7 21.7 40.5 61.6 88.2 182.4 109.3 269.7 47 87.3-62.3 108.1-184.3 46.5-272.6-9-12.9-19.3-24.3-30.5-34.2 37.4 78.8 10.7 178.5-67 233.9m-218.8-244c-1.4 1-2.7 2.1-4 3.1 64.3-17.8 135.9 4 178.9 60.5 35.7 47 42.9 106.6 24.4 158 56.7-56.2 67.6-142.1 22.3-201.8-50-65.5-149.1-74.4-221.6-19.8M296 224c-4.4 0-8-3.6-8-8v-40c0-4.4-3.6-8-8-8h-48c-4.4 0-8 3.6-8 8v40c0 4.4-3.6 8-8 8h-40c-4.4 0-8 3.6-8 8v48c0 4.4 3.6 8 8 8h40c4.4 0 8 3.6 8 8v40c0 4.4 3.6 8 8 8h48c4.4 0 8-3.6 8-8v-40c0-4.4 3.6-8 8-8h40c4.4 0 8-3.6 8-8v-48c0-4.4-3.6-8-8-8h-40z"],meetup:[512,512,[],"f2e0","M99 414.3c1.1 5.7-2.3 11.1-8 12.3-5.4 1.1-10.9-2.3-12-8-1.1-5.4 2.3-11.1 7.7-12.3 5.4-1.2 11.1 2.3 12.3 8zm143.1 71.4c-6.3 4.6-8 13.4-3.7 20 4.6 6.6 13.4 8.3 20 3.7 6.3-4.6 8-13.4 3.4-20-4.2-6.5-13.1-8.3-19.7-3.7zm-86-462.3c6.3-1.4 10.3-7.7 8.9-14-1.1-6.6-7.4-10.6-13.7-9.1-6.3 1.4-10.3 7.7-9.1 14 1.4 6.6 7.6 10.6 13.9 9.1zM34.4 226.3c-10-6.9-23.7-4.3-30.6 6-6.9 10-4.3 24 5.7 30.9 10 7.1 23.7 4.6 30.6-5.7 6.9-10.4 4.3-24.1-5.7-31.2zm272-170.9c10.6-6.3 13.7-20 7.7-30.3-6.3-10.6-19.7-14-30-7.7s-13.7 20-7.4 30.6c6 10.3 19.4 13.7 29.7 7.4zm-191.1 58c7.7-5.4 9.4-16 4.3-23.7s-15.7-9.4-23.1-4.3c-7.7 5.4-9.4 16-4.3 23.7 5.1 7.8 15.6 9.5 23.1 4.3zm372.3 156c-7.4 1.7-12.3 9.1-10.6 16.9 1.4 7.4 8.9 12.3 16.3 10.6 7.4-1.4 12.3-8.9 10.6-16.6-1.5-7.4-8.9-12.3-16.3-10.9zm39.7-56.8c-1.1-5.7-6.6-9.1-12-8-5.7 1.1-9.1 6.9-8 12.6 1.1 5.4 6.6 9.1 12.3 8 5.4-1.5 9.1-6.9 7.7-12.6zM447 138.9c-8.6 6-10.6 17.7-4.9 26.3 5.7 8.6 17.4 10.6 26 4.9 8.3-6 10.3-17.7 4.6-26.3-5.7-8.7-17.4-10.9-25.7-4.9zm-6.3 139.4c26.3 43.1 15.1 100-26.3 129.1-17.4 12.3-37.1 17.7-56.9 17.1-12 47.1-69.4 64.6-105.1 32.6-1.1.9-2.6 1.7-3.7 2.9-39.1 27.1-92.3 17.4-119.4-22.3-9.7-14.3-14.6-30.6-15.1-46.9-65.4-10.9-90-94-41.1-139.7-28.3-46.9.6-107.4 53.4-114.9C151.6 70 234.1 38.6 290.1 82c67.4-22.3 136.3 29.4 130.9 101.1 41.1 12.6 52.8 66.9 19.7 95.2zm-70 74.3c-3.1-20.6-40.9-4.6-43.1-27.1-3.1-32 43.7-101.1 40-128-3.4-24-19.4-29.1-33.4-29.4-13.4-.3-16.9 2-21.4 4.6-2.9 1.7-6.6 4.9-11.7-.3-6.3-6-11.1-11.7-19.4-12.9-12.3-2-17.7 2-26.6 9.7-3.4 2.9-12 12.9-20 9.1-3.4-1.7-15.4-7.7-24-11.4-16.3-7.1-40 4.6-48.6 20-12.9 22.9-38 113.1-41.7 125.1-8.6 26.6 10.9 48.6 36.9 47.1 11.1-.6 18.3-4.6 25.4-17.4 4-7.4 41.7-107.7 44.6-112.6 2-3.4 8.9-8 14.6-5.1 5.7 3.1 6.9 9.4 6 15.1-1.1 9.7-28 70.9-28.9 77.7-3.4 22.9 26.9 26.6 38.6 4 3.7-7.1 45.7-92.6 49.4-98.3 4.3-6.3 7.4-8.3 11.7-8 3.1 0 8.3.9 7.1 10.9-1.4 9.4-35.1 72.3-38.9 87.7-4.6 20.6 6.6 41.4 24.9 50.6 11.4 5.7 62.5 15.7 58.5-11.1zm5.7 92.3c-10.3 7.4-12.9 22-5.7 32.6 7.1 10.6 21.4 13.1 32 6 10.6-7.4 13.1-22 6-32.6-7.4-10.6-21.7-13.5-32.3-6z"],microsoft:[448,512,[],"f3ca","M0 32h214.6v214.6H0V32zm233.4 0H448v214.6H233.4V32zM0 265.4h214.6V480H0V265.4zm233.4 0H448V480H233.4V265.4z"],mix:[448,512,[],"f3cb","M0 64v348.9c0 56.2 88 58.1 88 0V174.3c7.9-52.9 88-50.4 88 6.5v175.3c0 57.9 96 58 96 0V240c5.3-54.7 88-52.5 88 4.3v23.8c0 59.9 88 56.6 88 0V64H0z"],mixcloud:[640,512,[],"f289","M424.43 219.729C416.124 134.727 344.135 68 256.919 68c-72.266 0-136.224 46.516-159.205 114.074-54.545 8.029-96.63 54.822-96.63 111.582 0 62.298 50.668 112.966 113.243 112.966h289.614c52.329 0 94.969-42.362 94.969-94.693 0-45.131-32.118-83.063-74.48-92.2zm-20.489 144.53H114.327c-39.04 0-70.881-31.564-70.881-70.604s31.841-70.604 70.881-70.604c18.827 0 36.548 7.475 49.838 20.766 19.963 19.963 50.133-10.227 30.18-30.18-14.675-14.398-32.672-24.365-52.053-29.349 19.935-44.3 64.79-73.926 114.628-73.926 69.496 0 125.979 56.483 125.979 125.702 0 13.568-2.215 26.857-6.369 39.594-8.943 27.517 32.133 38.939 40.147 13.29 2.769-8.306 4.984-16.889 6.369-25.472 19.381 7.476 33.502 26.303 33.502 48.453 0 28.795-23.535 52.33-52.607 52.33zm235.069-52.33c0 44.024-12.737 86.386-37.102 122.657-4.153 6.092-10.798 9.414-17.72 9.414-16.317 0-27.127-18.826-17.443-32.949 19.381-29.349 29.903-63.682 29.903-99.122s-10.521-69.773-29.903-98.845c-15.655-22.831 19.361-47.24 35.163-23.534 24.366 35.993 37.102 78.356 37.102 122.379zm-70.88 0c0 31.565-9.137 62.021-26.857 88.325-4.153 6.091-10.798 9.136-17.72 9.136-17.201 0-27.022-18.979-17.443-32.948 13.013-19.104 19.658-41.255 19.658-64.513 0-22.981-6.645-45.408-19.658-64.512-15.761-22.986 19.008-47.095 35.163-23.535 17.719 26.026 26.857 56.483 26.857 88.047z"],mizuni:[496,512,[],"f3cc","M248 8C111 8 0 119.1 0 256c0 137 111 248 248 248s248-111 248-248C496 119.1 385 8 248 8zm-80 351.9c-31.4 10.6-58.8 27.3-80 48.2V136c0-22.1 17.9-40 40-40s40 17.9 40 40v223.9zm120-9.9c-12.9-2-26.2-3.1-39.8-3.1-13.8 0-27.2 1.1-40.2 3.1V136c0-22.1 17.9-40 40-40s40 17.9 40 40v214zm120 57.7c-21.2-20.8-48.6-37.4-80-48V136c0-22.1 17.9-40 40-40s40 17.9 40 40v271.7z"],modx:[448,512,[],"f285","M356 241.8l36.7 23.7V480l-133-83.8L356 241.8zM440 75H226.3l-23 37.8 153.5 96.5L440 75zm-89 142.8L55.2 32v214.5l46 29L351 217.8zM97 294.2L8 437h213.7l125-200.5L97 294.2z"],monero:[496,512,[],"f3d0","M352 384h108.4C417 455.9 338.1 504 248 504S79 455.9 35.6 384H144V256.2L248 361l104-105v128zM88 336V128l159.4 159.4L408 128v208h74.8c8.5-25.1 13.2-52 13.2-80C496 119 385 8 248 8S0 119 0 256c0 28 4.6 54.9 13.2 80H88z"],napster:[496,512,[],"f3d2","M298.3 373.6c-14.2 13.6-31.3 24.1-50.4 30.5-19-6.4-36.2-16.9-50.3-30.5h100.7zm44-199.6c20-16.9 43.6-29.2 69.6-36.2V299c0 219.4-328 217.6-328 .3V137.7c25.9 6.9 49.6 19.6 69.5 36.4 56.8-40 132.5-39.9 188.9-.1zm-208.8-58.5c64.4-60 164.3-60.1 228.9-.2-7.1 3.5-13.9 7.3-20.6 11.5-58.7-30.5-129.2-30.4-187.9.1-6.3-4-13.9-8.2-20.4-11.4zM43.8 93.2v69.3c-58.4 36.5-58.4 121.1.1 158.3 26.4 245.1 381.7 240.3 407.6 1.5l.3-1.7c58.7-36.3 58.9-121.7.2-158.2V93.2c-17.3.5-34 3-50.1 7.4-82-91.5-225.5-91.5-307.5.1-16.3-4.4-33.1-7-50.6-7.5zM259.2 352s36-.3 61.3-1.5c10.2-.5 21.1-4 25.5-6.5 26.3-15.1 25.4-39.2 26.2-47.4-79.5-.6-99.9-3.9-113 55.4zm-135.5-55.3c.8 8.2-.1 32.3 26.2 47.4 4.4 2.5 15.2 6 25.5 6.5 25.3 1.1 61.3 1.5 61.3 1.5-13.2-59.4-33.7-56.1-113-55.4zm169.1 123.4c-3.2-5.3-6.9-7.3-6.9-7.3-24.8 7.3-52.2 6.9-75.9 0 0 0-2.9 1.5-6.4 6.6-2.8 4.1-3.7 9.6-3.7 9.6 29.1 17.6 67.1 17.6 96.2 0-.1-.1-.3-4-3.3-8.9z"],"nintendo-switch":[448,512,[],"f418","M95.9 33.5c-44.6 8-80.5 41-91.8 84.4C0 133.6-.3 142.8.2 264.4.4 376 .5 378.6 2.4 387.3c10.3 46.5 43.3 79.6 90.3 90.5 6.1 1.4 13.9 1.7 64.1 1.9 51.9.4 57.3.3 58.7-1.1 1.4-1.4 1.5-19.3 1.5-222.2 0-150.5-.3-221.3-.9-222.6-.9-1.7-2.5-1.8-56.9-1.7-44.2.1-57.5.4-63.3 1.4zm83.9 222.6V444l-37.8-.5c-34.8-.4-38.5-.6-45.5-2.3-29.9-7.7-52-30.7-58.3-60.7-2-9.4-2-240.1-.1-249.3 5.6-26.1 23.7-47.7 48-57.4 12.2-4.9 17.9-5.5 57.6-5.6l35.9-.1v188zm-75.9-131.2c-5.8 1.1-14.7 5.6-19.5 9.7-9.7 8.4-14.6 20.4-13.8 34.5.4 7.3.8 9.3 3.8 15.2 4.4 9 10.9 15.6 19.9 20 6.2 3.1 7.8 3.4 15.9 3.7 7.3.3 9.9 0 14.8-1.7 20.1-6.8 32.3-26.3 28.8-46.4-3.9-23.7-26.6-39.7-49.9-35zm158.2-92.3c-.4.3-.6 100.8-.6 223.5 0 202.3.1 222.8 1.5 223.4 2.5.9 74.5.6 83.4-.4 37.7-4.3 71-27.2 89-61.2 2.3-4.4 5.4-11.7 7-16.2 5.8-17.4 5.7-12.8 5.7-146.1 0-106.4-.2-122.3-1.5-129-9.2-48.3-46.1-84.8-94.5-93.1-6.5-1.1-16.5-1.4-48.8-1.4-22.4-.1-40.9.2-41.2.5zm99.1 202.1c14.5 3.8 26.3 14.8 31.2 28.9 3.1 8.7 3 21.5-.1 29.5-5.7 14.7-16.8 25-31.1 28.8-23.2 6-47.9-8-54.6-31-2-7-1.9-18.9.4-26.2 6.9-22.7 31-36.1 54.2-30z"],node:[640,512,[],"f419","M316.3 452c-2.1 0-4.2-.6-6.1-1.6L291 439c-2.9-1.6-1.5-2.2-.5-2.5 3.8-1.3 4.6-1.6 8.7-4 .4-.2 1-.1 1.4.1l14.8 8.8c.5.3 1.3.3 1.8 0L375 408c.5-.3.9-.9.9-1.6v-66.7c0-.7-.3-1.3-.9-1.6l-57.8-33.3c-.5-.3-1.2-.3-1.8 0l-57.8 33.3c-.6.3-.9 1-.9 1.6v66.7c0 .6.4 1.2.9 1.5l15.8 9.1c8.6 4.3 13.9-.8 13.9-5.8v-65.9c0-.9.7-1.7 1.7-1.7h7.3c.9 0 1.7.7 1.7 1.7v65.9c0 11.5-6.2 18-17.1 18-3.3 0-6 0-13.3-3.6l-15.2-8.7c-3.7-2.2-6.1-6.2-6.1-10.5v-66.7c0-4.3 2.3-8.4 6.1-10.5l57.8-33.4c3.7-2.1 8.5-2.1 12.1 0l57.8 33.4c3.7 2.2 6.1 6.2 6.1 10.5v66.7c0 4.3-2.3 8.4-6.1 10.5l-57.8 33.4c-1.7 1.1-3.8 1.7-6 1.7zm46.7-65.8c0-12.5-8.4-15.8-26.2-18.2-18-2.4-19.8-3.6-19.8-7.8 0-3.5 1.5-8.1 14.8-8.1 11.9 0 16.3 2.6 18.1 10.6.2.8.8 1.3 1.6 1.3h7.5c.5 0 .9-.2 1.2-.5.3-.4.5-.8.4-1.3-1.2-13.8-10.3-20.2-28.8-20.2-16.5 0-26.3 7-26.3 18.6 0 12.7 9.8 16.1 25.6 17.7 18.9 1.9 20.4 4.6 20.4 8.3 0 6.5-5.2 9.2-17.4 9.2-15.3 0-18.7-3.8-19.8-11.4-.1-.8-.8-1.4-1.7-1.4h-7.5c-.9 0-1.7.7-1.7 1.7 0 9.7 5.3 21.3 30.6 21.3 18.5 0 29-7.2 29-19.8zm54.5-50.1c0 6.1-5 11.1-11.1 11.1s-11.1-5-11.1-11.1c0-6.3 5.2-11.1 11.1-11.1 6-.1 11.1 4.8 11.1 11.1zm-1.8 0c0-5.2-4.2-9.3-9.4-9.3-5.1 0-9.3 4.1-9.3 9.3 0 5.2 4.2 9.4 9.3 9.4 5.2-.1 9.4-4.3 9.4-9.4zm-4.5 6.2h-2.6c-.1-.6-.5-3.8-.5-3.9-.2-.7-.4-1.1-1.3-1.1h-2.2v5h-2.4v-12.5h4.3c1.5 0 4.4 0 4.4 3.3 0 2.3-1.5 2.8-2.4 3.1 1.7.1 1.8 1.2 2.1 2.8.1 1 .3 2.7.6 3.3zm-2.8-8.8c0-1.7-1.2-1.7-1.8-1.7h-2v3.5h1.9c1.6 0 1.9-1.1 1.9-1.8zM137.3 191c0-2.7-1.4-5.1-3.7-6.4l-61.3-35.3c-1-.6-2.2-.9-3.4-1h-.6c-1.2 0-2.3.4-3.4 1L3.7 184.6C1.4 185.9 0 188.4 0 191l.1 95c0 1.3.7 2.5 1.8 3.2 1.1.7 2.5.7 3.7 0L42 268.3c2.3-1.4 3.7-3.8 3.7-6.4v-44.4c0-2.6 1.4-5.1 3.7-6.4l15.5-8.9c1.2-.7 2.4-1 3.7-1 1.3 0 2.6.3 3.7 1l15.5 8.9c2.3 1.3 3.7 3.8 3.7 6.4v44.4c0 2.6 1.4 5.1 3.7 6.4l36.4 20.9c1.1.7 2.6.7 3.7 0 1.1-.6 1.8-1.9 1.8-3.2l.2-95zM472.5 87.3v176.4c0 2.6-1.4 5.1-3.7 6.4l-61.3 35.4c-2.3 1.3-5.1 1.3-7.4 0l-61.3-35.4c-2.3-1.3-3.7-3.8-3.7-6.4v-70.8c0-2.6 1.4-5.1 3.7-6.4l61.3-35.4c2.3-1.3 5.1-1.3 7.4 0l15.3 8.8c1.7 1 3.9-.3 3.9-2.2v-94c0-2.8 3-4.6 5.5-3.2l36.5 20.4c2.3 1.2 3.8 3.7 3.8 6.4zm-46 128.9c0-.7-.4-1.3-.9-1.6l-21-12.2c-.6-.3-1.3-.3-1.9 0l-21 12.2c-.6.3-.9.9-.9 1.6v24.3c0 .7.4 1.3.9 1.6l21 12.1c.6.3 1.3.3 1.8 0l21-12.1c.6-.3.9-.9.9-1.6v-24.3zm209.8-.7c2.3-1.3 3.7-3.8 3.7-6.4V192c0-2.6-1.4-5.1-3.7-6.4l-60.9-35.4c-2.3-1.3-5.1-1.3-7.4 0l-61.3 35.4c-2.3 1.3-3.7 3.8-3.7 6.4v70.8c0 2.7 1.4 5.1 3.7 6.4l60.9 34.7c2.2 1.3 5 1.3 7.3 0l36.8-20.5c2.5-1.4 2.5-5 0-6.4L550 241.6c-1.2-.7-1.9-1.9-1.9-3.2v-22.2c0-1.3.7-2.5 1.9-3.2l19.2-11.1c1.1-.7 2.6-.7 3.7 0l19.2 11.1c1.1.7 1.9 1.9 1.9 3.2v17.4c0 2.8 3.1 4.6 5.6 3.2l36.7-21.3zM559 219c-.4.3-.7.7-.7 1.2v13.6c0 .5.3 1 .7 1.2l11.8 6.8c.4.3 1 .3 1.4 0L584 235c.4-.3.7-.7.7-1.2v-13.6c0-.5-.3-1-.7-1.2l-11.8-6.8c-.4-.3-1-.3-1.4 0L559 219zm-254.2 43.5v-70.4c0-2.6-1.6-5.1-3.9-6.4l-61.1-35.2c-2.1-1.2-5-1.4-7.4 0l-61.1 35.2c-2.3 1.3-3.9 3.7-3.9 6.4v70.4c0 2.8 1.9 5.2 4 6.4l61.2 35.2c2.4 1.4 5.2 1.3 7.4 0l61-35.2c1.8-1 3.1-2.7 3.6-4.7.1-.5.2-1.1.2-1.7zm-74.3-124.9l-.8.5h1.1l-.3-.5zm76.2 130.2l-.4-.7v.9l.4-.2z"],"node-js":[448,512,[],"f3d3","M224 508c-6.7 0-13.5-1.8-19.4-5.2l-61.7-36.5c-9.2-5.2-4.7-7-1.7-8 12.3-4.3 14.8-5.2 27.9-12.7 1.4-.8 3.2-.5 4.6.4l47.4 28.1c1.7 1 4.1 1 5.7 0l184.7-106.6c1.7-1 2.8-3 2.8-5V149.3c0-2.1-1.1-4-2.9-5.1L226.8 37.7c-1.7-1-4-1-5.7 0L36.6 144.3c-1.8 1-2.9 3-2.9 5.1v213.1c0 2 1.1 4 2.9 4.9l50.6 29.2c27.5 13.7 44.3-2.4 44.3-18.7V167.5c0-3 2.4-5.3 5.4-5.3h23.4c2.9 0 5.4 2.3 5.4 5.3V378c0 36.6-20 57.6-54.7 57.6-10.7 0-19.1 0-42.5-11.6l-48.4-27.9C8.1 389.2.7 376.3.7 362.4V149.3c0-13.8 7.4-26.8 19.4-33.7L204.6 9c11.7-6.6 27.2-6.6 38.8 0l184.7 106.7c12 6.9 19.4 19.8 19.4 33.7v213.1c0 13.8-7.4 26.7-19.4 33.7L243.4 502.8c-5.9 3.4-12.6 5.2-19.4 5.2zm149.1-210.1c0-39.9-27-50.5-83.7-58-57.4-7.6-63.2-11.5-63.2-24.9 0-11.1 4.9-25.9 47.4-25.9 37.9 0 51.9 8.2 57.7 33.8.5 2.4 2.7 4.2 5.2 4.2h24c1.5 0 2.9-.6 3.9-1.7s1.5-2.6 1.4-4.1c-3.7-44.1-33-64.6-92.2-64.6-52.7 0-84.1 22.2-84.1 59.5 0 40.4 31.3 51.6 81.8 56.6 60.5 5.9 65.2 14.8 65.2 26.7 0 20.6-16.6 29.4-55.5 29.4-48.9 0-59.6-12.3-63.2-36.6-.4-2.6-2.6-4.5-5.3-4.5h-23.9c-3 0-5.3 2.4-5.3 5.3 0 31.1 16.9 68.2 97.8 68.2 58.4-.1 92-23.2 92-63.4z"],npm:[576,512,[],"f3d4","M288 288h-32v-64h32v64zm288-128v192H288v32H160v-32H0V160h576zm-416 32H32v128h64v-96h32v96h32V192zm160 0H192v160h64v-32h64V192zm224 0H352v128h64v-96h32v96h32v-96h32v96h32V192z"],ns8:[640,512,[],"f3d5","M187.1 159.9l-34.2 113.7-54.5-113.7H49L0 320h44.9L76 213.5 126.6 320h56.9L232 159.9h-44.9zm452.5-.9c-2.9-18-23.9-28.1-42.1-31.3-44.6-7.8-101.9 16.3-88.5 58.8v.1c-43.8 8.7-74.3 26.8-94.2 48.2-3-9.8-13.6-16.6-34-16.6h-87.6c-9.3 0-12.9-2.3-11.5-7.4 1.6-5.5 1.9-6.8 3.7-12.2 2.1-6.4 7.8-7.1 13.3-7.1h133.5l9.7-31.5c-139.7 0-144.5-.5-160.1 1.2-12.3 1.3-23.5 4.8-30.6 15-6.8 9.9-14.4 35.6-17.6 47.1-5.4 19.4-.6 28.6 32.8 28.6h87.3c7.8 0 8.8 2.7 7.7 6.6-1.1 4.4-2.8 10-4.5 14.6-1.6 4.2-4.7 7.4-13.8 7.4H216.3L204.7 320c139.9 0 145.3-.6 160.9-2.3 6.6-.7 13-2.1 18.5-4.9.2 3.7.5 7.3 1.2 10.8 5.4 30.5 27.4 52.3 56.8 59.5 48.6 11.9 108.7-16.8 135.1-68 18.7-36.2 14.1-76.2-3.4-105.5h.1c29.6-5.9 70.3-22 65.7-50.6zM530.7 263.7c-5.9 29.5-36.6 47.8-61.6 43.9-30.9-4.8-38.5-39.5-14.1-64.8 16.2-16.8 45.2-24 68.5-26.9 6.7 14.1 10.3 32 7.2 47.8zm21.8-83.1c-4.2-6-9.8-18.5-2.5-26.3 6.7-7.2 20.9-10.1 31.8-7.7 15.3 3.4 19.7 15.9 4.9 24.4-10.7 6.1-23.6 8.1-34.2 9.6z"],nutritionix:[400,512,[],"f3d6","M88 8.1S221.4-.1 209 112.5c0 0 19.1-74.9 103-40.6 0 0-17.7 74-88 56 0 0 14.6-54.6 66.1-56.6 0 0-39.9-10.3-82.1 48.8 0 0-19.8-94.5-93.6-99.7 0 0 75.2 19.4 77.6 107.5 0 .1-106.4 7-104-119.8zm312 315.6c0 48.5-9.7 95.3-32 132.3-42.2 30.9-105 48-168 48-62.9 0-125.8-17.1-168-48C9.7 419 0 372.2 0 323.7 0 275.3 17.7 229 40 192c42.2-30.9 97.1-48.6 160-48.6 63 0 117.8 17.6 160 48.6 22.3 37 40 83.3 40 131.7zM120 428c0-15.5-12.5-28-28-28s-28 12.5-28 28 12.5 28 28 28 28-12.5 28-28zm0-66.2c0-15.5-12.5-28-28-28s-28 12.5-28 28 12.5 28 28 28 28-12.5 28-28zm0-66.2c0-15.5-12.5-28-28-28s-28 12.5-28 28 12.5 28 28 28 28-12.5 28-28zM192 428c0-15.5-12.5-28-28-28s-28 12.5-28 28 12.5 28 28 28 28-12.5 28-28zm0-66.2c0-15.5-12.5-28-28-28s-28 12.5-28 28 12.5 28 28 28 28-12.5 28-28zm0-66.2c0-15.5-12.5-28-28-28s-28 12.5-28 28 12.5 28 28 28 28-12.5 28-28zM264 428c0-15.5-12.5-28-28-28s-28 12.5-28 28 12.5 28 28 28 28-12.5 28-28zm0-66.2c0-15.5-12.5-28-28-28s-28 12.5-28 28 12.5 28 28 28 28-12.5 28-28zm0-66.2c0-15.5-12.5-28-28-28s-28 12.5-28 28 12.5 28 28 28 28-12.5 28-28zM336 428c0-15.5-12.5-28-28-28s-28 12.5-28 28 12.5 28 28 28 28-12.5 28-28zm0-66.2c0-15.5-12.5-28-28-28s-28 12.5-28 28 12.5 28 28 28 28-12.5 28-28zm0-66.2c0-15.5-12.5-28-28-28s-28 12.5-28 28 12.5 28 28 28 28-12.5 28-28zm24-39.6c-4.8-22.3-7.4-36.9-16-56-38.8-19.9-90.5-32-144-32S94.8 180.1 56 200c-8.8 19.5-11.2 33.9-16 56 42.2-7.9 98.7-14.8 160-14.8s117.8 6.9 160 14.8z"],odnoklassniki:[320,512,[],"f263","M275.1 334c-27.4 17.4-65.1 24.3-90 26.9l20.9 20.6 76.3 76.3c27.9 28.6-17.5 73.3-45.7 45.7-19.1-19.4-47.1-47.4-76.3-76.6L84 503.4c-28.2 27.5-73.6-17.6-45.4-45.7 19.4-19.4 47.1-47.4 76.3-76.3l20.6-20.6c-24.6-2.6-62.9-9.1-90.6-26.9-32.6-21-46.9-33.3-34.3-59 7.4-14.6 27.7-26.9 54.6-5.7 0 0 36.3 28.9 94.9 28.9s94.9-28.9 94.9-28.9c26.9-21.1 47.1-8.9 54.6 5.7 12.4 25.7-1.9 38-34.5 59.1zM30.3 129.7C30.3 58 88.6 0 160 0s129.7 58 129.7 129.7c0 71.4-58.3 129.4-129.7 129.4s-129.7-58-129.7-129.4zm66 0c0 35.1 28.6 63.7 63.7 63.7s63.7-28.6 63.7-63.7c0-35.4-28.6-64-63.7-64s-63.7 28.6-63.7 64z"],"odnoklassniki-square":[448,512,[],"f264","M184.2 177.1c0-22.1 17.9-40 39.8-40s39.8 17.9 39.8 40c0 22-17.9 39.8-39.8 39.8s-39.8-17.9-39.8-39.8zM448 80v352c0 26.5-21.5 48-48 48H48c-26.5 0-48-21.5-48-48V80c0-26.5 21.5-48 48-48h352c26.5 0 48 21.5 48 48zm-305.1 97.1c0 44.6 36.4 80.9 81.1 80.9s81.1-36.2 81.1-80.9c0-44.8-36.4-81.1-81.1-81.1s-81.1 36.2-81.1 81.1zm174.5 90.7c-4.6-9.1-17.3-16.8-34.1-3.6 0 0-22.7 18-59.3 18s-59.3-18-59.3-18c-16.8-13.2-29.5-5.5-34.1 3.6-7.9 16.1 1.1 23.7 21.4 37 17.3 11.1 41.2 15.2 56.6 16.8l-12.9 12.9c-18.2 18-35.5 35.5-47.7 47.7-17.6 17.6 10.7 45.8 28.4 28.6l47.7-47.9c18.2 18.2 35.7 35.7 47.7 47.9 17.6 17.2 46-10.7 28.6-28.6l-47.7-47.7-13-12.9c15.5-1.6 39.1-5.9 56.2-16.8 20.4-13.3 29.3-21 21.5-37z"],"old-republic":[496,512,[],"f510","M235.76 10.23c7.5-.31 15-.28 22.5-.09 3.61.14 7.2.4 10.79.73 4.92.27 9.79 1.03 14.67 1.62 2.93.43 5.83.98 8.75 1.46 7.9 1.33 15.67 3.28 23.39 5.4 12.24 3.47 24.19 7.92 35.76 13.21 26.56 12.24 50.94 29.21 71.63 49.88 20.03 20.09 36.72 43.55 48.89 69.19 1.13 2.59 2.44 5.1 3.47 7.74 2.81 6.43 5.39 12.97 7.58 19.63 4.14 12.33 7.34 24.99 9.42 37.83.57 3.14 1.04 6.3 1.4 9.47.55 3.83.94 7.69 1.18 11.56.83 8.34.84 16.73.77 25.1-.07 4.97-.26 9.94-.75 14.89-.24 3.38-.51 6.76-.98 10.12-.39 2.72-.63 5.46-1.11 8.17-.9 5.15-1.7 10.31-2.87 15.41-4.1 18.5-10.3 36.55-18.51 53.63-15.77 32.83-38.83 62.17-67.12 85.12a246.503 246.503 0 0 1-56.91 34.86c-6.21 2.68-12.46 5.25-18.87 7.41-3.51 1.16-7.01 2.38-10.57 3.39-6.62 1.88-13.29 3.64-20.04 5-4.66.91-9.34 1.73-14.03 2.48-5.25.66-10.5 1.44-15.79 1.74-6.69.66-13.41.84-20.12.81-6.82.03-13.65-.12-20.45-.79-3.29-.23-6.57-.5-9.83-.95-2.72-.39-5.46-.63-8.17-1.11-4.12-.72-8.25-1.37-12.35-2.22-4.25-.94-8.49-1.89-12.69-3.02-8.63-2.17-17.08-5.01-25.41-8.13-10.49-4.12-20.79-8.75-30.64-14.25-2.14-1.15-4.28-2.29-6.35-3.57-11.22-6.58-21.86-14.1-31.92-22.34-34.68-28.41-61.41-66.43-76.35-108.7-3.09-8.74-5.71-17.65-7.8-26.68-1.48-6.16-2.52-12.42-3.58-18.66-.4-2.35-.61-4.73-.95-7.09-.6-3.96-.75-7.96-1.17-11.94-.8-9.47-.71-18.99-.51-28.49.14-3.51.34-7.01.7-10.51.31-3.17.46-6.37.92-9.52.41-2.81.65-5.65 1.16-8.44.7-3.94 1.3-7.9 2.12-11.82 3.43-16.52 8.47-32.73 15.26-48.18 1.15-2.92 2.59-5.72 3.86-8.59 8.05-16.71 17.9-32.56 29.49-47.06 20-25.38 45.1-46.68 73.27-62.47 7.5-4.15 15.16-8.05 23.07-11.37 15.82-6.88 32.41-11.95 49.31-15.38 3.51-.67 7.04-1.24 10.56-1.85 2.62-.47 5.28-.7 7.91-1.08 3.53-.53 7.1-.68 10.65-1.04 2.46-.24 4.91-.36 7.36-.51m8.64 24.41c-9.23.1-18.43.99-27.57 2.23-7.3 1.08-14.53 2.6-21.71 4.3-13.91 3.5-27.48 8.34-40.46 14.42-10.46 4.99-20.59 10.7-30.18 17.22-4.18 2.92-8.4 5.8-12.34 9.03-5.08 3.97-9.98 8.17-14.68 12.59-2.51 2.24-4.81 4.7-7.22 7.06-28.22 28.79-48.44 65.39-57.5 104.69-2.04 8.44-3.54 17.02-4.44 25.65-1.1 8.89-1.44 17.85-1.41 26.8.11 7.14.38 14.28 1.22 21.37.62 7.12 1.87 14.16 3.2 21.18 1.07 4.65 2.03 9.32 3.33 13.91 6.29 23.38 16.5 45.7 30.07 65.75 8.64 12.98 18.78 24.93 29.98 35.77 16.28 15.82 35.05 29.04 55.34 39.22 7.28 3.52 14.66 6.87 22.27 9.63 5.04 1.76 10.06 3.57 15.22 4.98 11.26 3.23 22.77 5.6 34.39 7.06 2.91.29 5.81.61 8.72.9 13.82 1.08 27.74 1 41.54-.43 4.45-.6 8.92-.99 13.35-1.78 3.63-.67 7.28-1.25 10.87-2.1 4.13-.98 8.28-1.91 12.36-3.07 26.5-7.34 51.58-19.71 73.58-36.2 15.78-11.82 29.96-25.76 42.12-41.28 3.26-4.02 6.17-8.31 9.13-12.55 3.39-5.06 6.58-10.25 9.6-15.54 2.4-4.44 4.74-8.91 6.95-13.45 5.69-12.05 10.28-24.62 13.75-37.49 2.59-10.01 4.75-20.16 5.9-30.45 1.77-13.47 1.94-27.1 1.29-40.65-.29-3.89-.67-7.77-1-11.66-2.23-19.08-6.79-37.91-13.82-55.8-5.95-15.13-13.53-29.63-22.61-43.13-12.69-18.8-28.24-35.68-45.97-49.83-25.05-20-54.47-34.55-85.65-42.08-7.78-1.93-15.69-3.34-23.63-4.45-3.91-.59-7.85-.82-11.77-1.24-7.39-.57-14.81-.72-22.22-.58zM139.26 83.53c13.3-8.89 28.08-15.38 43.3-20.18-3.17 1.77-6.44 3.38-9.53 5.29-11.21 6.68-21.52 14.9-30.38 24.49-6.8 7.43-12.76 15.73-17.01 24.89-3.29 6.86-5.64 14.19-6.86 21.71-.93 4.85-1.3 9.81-1.17 14.75.13 13.66 4.44 27.08 11.29 38.82 5.92 10.22 13.63 19.33 22.36 27.26 4.85 4.36 10.24 8.09 14.95 12.6 2.26 2.19 4.49 4.42 6.43 6.91 2.62 3.31 4.89 6.99 5.99 11.1.9 3.02.66 6.2.69 9.31.02 4.1-.04 8.2.03 12.3.14 3.54-.02 7.09.11 10.63.08 2.38.02 4.76.05 7.14.16 5.77.06 11.53.15 17.3.11 2.91.02 5.82.13 8.74.03 1.63.13 3.28-.03 4.91-.91.12-1.82.18-2.73.16-10.99 0-21.88-2.63-31.95-6.93-6-2.7-11.81-5.89-17.09-9.83-5.75-4.19-11.09-8.96-15.79-14.31-6.53-7.24-11.98-15.39-16.62-23.95-1.07-2.03-2.24-4.02-3.18-6.12-1.16-2.64-2.62-5.14-3.67-7.82-4.05-9.68-6.57-19.94-8.08-30.31-.49-4.44-1.09-8.88-1.2-13.35-.7-15.73.84-31.55 4.67-46.82 2.12-8.15 4.77-16.18 8.31-23.83 6.32-14.2 15.34-27.18 26.3-38.19 6.28-6.2 13.13-11.84 20.53-16.67zm175.37-20.12c2.74.74 5.41 1.74 8.09 2.68 6.36 2.33 12.68 4.84 18.71 7.96 13.11 6.44 25.31 14.81 35.82 24.97 10.2 9.95 18.74 21.6 25.14 34.34 1.28 2.75 2.64 5.46 3.81 8.26 6.31 15.1 10 31.26 11.23 47.57.41 4.54.44 9.09.45 13.64.07 11.64-1.49 23.25-4.3 34.53-1.97 7.27-4.35 14.49-7.86 21.18-3.18 6.64-6.68 13.16-10.84 19.24-6.94 10.47-15.6 19.87-25.82 27.22-10.48 7.64-22.64 13.02-35.4 15.38-3.51.69-7.08 1.08-10.66 1.21-1.85.06-3.72.16-5.56-.1-.28-2.15 0-4.31-.01-6.46-.03-3.73.14-7.45.1-11.17.19-7.02.02-14.05.21-21.07.03-2.38-.03-4.76.03-7.14.17-5.07-.04-10.14.14-15.21.1-2.99-.24-6.04.51-8.96.66-2.5 1.78-4.86 3.09-7.08 4.46-7.31 11.06-12.96 17.68-18.26 5.38-4.18 10.47-8.77 15.02-13.84 7.68-8.37 14.17-17.88 18.78-28.27 2.5-5.93 4.52-12.1 5.55-18.46.86-4.37 1.06-8.83 1.01-13.27-.02-7.85-1.4-15.65-3.64-23.17-1.75-5.73-4.27-11.18-7.09-16.45-3.87-6.93-8.65-13.31-13.96-19.2-9.94-10.85-21.75-19.94-34.6-27.1-1.85-1.02-3.84-1.82-5.63-2.97zm-100.8 58.45c.98-1.18 1.99-2.33 3.12-3.38-.61.93-1.27 1.81-1.95 2.68-3.1 3.88-5.54 8.31-7.03 13.06-.87 3.27-1.68 6.6-1.73 10-.07 2.52-.08 5.07.32 7.57 1.13 7.63 4.33 14.85 8.77 21.12 2 2.7 4.25 5.27 6.92 7.33 1.62 1.27 3.53 2.09 5.34 3.05 3.11 1.68 6.32 3.23 9.07 5.48 2.67 2.09 4.55 5.33 4.4 8.79-.01 73.67 0 147.34-.01 221.02 0 1.35-.08 2.7.04 4.04.13 1.48.82 2.83 1.47 4.15.86 1.66 1.78 3.34 3.18 4.62.85.77 1.97 1.4 3.15 1.24 1.5-.2 2.66-1.35 3.45-2.57.96-1.51 1.68-3.16 2.28-4.85.76-2.13.44-4.42.54-6.63.14-4.03-.02-8.06.14-12.09.03-5.89.03-11.77.06-17.66.14-3.62.03-7.24.11-10.86.15-4.03-.02-8.06.14-12.09.03-5.99.03-11.98.07-17.97.14-3.62.02-7.24.11-10.86.14-3.93-.02-7.86.14-11.78.03-5.99.03-11.98.06-17.97.16-3.94-.01-7.88.19-11.82.29 1.44.13 2.92.22 4.38.19 3.61.42 7.23.76 10.84.32 3.44.44 6.89.86 10.32.37 3.1.51 6.22.95 9.31.57 4.09.87 8.21 1.54 12.29 1.46 9.04 2.83 18.11 5.09 26.99 1.13 4.82 2.4 9.61 4 14.3 2.54 7.9 5.72 15.67 10.31 22.62 1.73 2.64 3.87 4.98 6.1 7.21.27.25.55.51.88.71.6.25 1.31-.07 1.7-.57.71-.88 1.17-1.94 1.7-2.93 4.05-7.8 8.18-15.56 12.34-23.31.7-1.31 1.44-2.62 2.56-3.61 1.75-1.57 3.84-2.69 5.98-3.63 2.88-1.22 5.9-2.19 9.03-2.42 6.58-.62 13.11.75 19.56 1.85 3.69.58 7.4 1.17 11.13 1.41 3.74.1 7.48.05 11.21-.28 8.55-.92 16.99-2.96 24.94-6.25 5.3-2.24 10.46-4.83 15.31-7.93 11.46-7.21 21.46-16.57 30.04-27.01 1.17-1.42 2.25-2.9 3.46-4.28-1.2 3.24-2.67 6.37-4.16 9.48-1.25 2.9-2.84 5.61-4.27 8.42-5.16 9.63-11.02 18.91-17.75 27.52-4.03 5.21-8.53 10.05-13.33 14.57-6.64 6.05-14.07 11.37-22.43 14.76-8.21 3.37-17.31 4.63-26.09 3.29-3.56-.58-7.01-1.69-10.41-2.88-2.79-.97-5.39-2.38-8.03-3.69-3.43-1.71-6.64-3.81-9.71-6.08 2.71 3.06 5.69 5.86 8.7 8.61 4.27 3.76 8.74 7.31 13.63 10.23 3.98 2.45 8.29 4.4 12.84 5.51 1.46.37 2.96.46 4.45.6-1.25 1.1-2.63 2.04-3.99 2.98-9.61 6.54-20.01 11.86-30.69 16.43-20.86 8.7-43.17 13.97-65.74 15.34-4.66.24-9.32.36-13.98.36-4.98-.11-9.97-.13-14.92-.65-11.2-.76-22.29-2.73-33.17-5.43-10.35-2.71-20.55-6.12-30.3-10.55-8.71-3.86-17.12-8.42-24.99-13.79-1.83-1.31-3.74-2.53-5.37-4.08 6.6-1.19 13.03-3.39 18.99-6.48 5.74-2.86 10.99-6.66 15.63-11.07 2.24-2.19 4.29-4.59 6.19-7.09-3.43 2.13-6.93 4.15-10.62 5.78-4.41 2.16-9.07 3.77-13.81 5.02-5.73 1.52-11.74 1.73-17.61 1.14-8.13-.95-15.86-4.27-22.51-8.98-4.32-2.94-8.22-6.43-11.96-10.06-9.93-10.16-18.2-21.81-25.66-33.86-3.94-6.27-7.53-12.75-11.12-19.22-1.05-2.04-2.15-4.05-3.18-6.1 2.85 2.92 5.57 5.97 8.43 8.88 8.99 8.97 18.56 17.44 29.16 24.48 7.55 4.9 15.67 9.23 24.56 11.03 3.11.73 6.32.47 9.47.81 2.77.28 5.56.2 8.34.3 5.05.06 10.11.04 15.16-.16 3.65-.16 7.27-.66 10.89-1.09 2.07-.25 4.11-.71 6.14-1.2 3.88-.95 8.11-.96 11.83.61 4.76 1.85 8.44 5.64 11.38 9.71 2.16 3.02 4.06 6.22 5.66 9.58 1.16 2.43 2.46 4.79 3.55 7.26 1 2.24 2.15 4.42 3.42 6.52.67 1.02 1.4 2.15 2.62 2.55 1.06-.75 1.71-1.91 2.28-3.03 2.1-4.16 3.42-8.65 4.89-13.05 2.02-6.59 3.78-13.27 5.19-20.02 2.21-9.25 3.25-18.72 4.54-28.13.56-3.98.83-7.99 1.31-11.97.87-10.64 1.9-21.27 2.24-31.94.08-1.86.24-3.71.25-5.57.01-4.35.25-8.69.22-13.03-.01-2.38-.01-4.76 0-7.13.05-5.07-.2-10.14-.22-15.21-.2-6.61-.71-13.2-1.29-19.78-.73-5.88-1.55-11.78-3.12-17.51-2.05-7.75-5.59-15.03-9.8-21.82-3.16-5.07-6.79-9.88-11.09-14.03-3.88-3.86-8.58-7.08-13.94-8.45-1.5-.41-3.06-.45-4.59-.64.07-2.99.7-5.93 1.26-8.85 1.59-7.71 3.8-15.3 6.76-22.6 1.52-4.03 3.41-7.9 5.39-11.72 3.45-6.56 7.62-12.79 12.46-18.46zm31.27 1.7c.35-.06.71-.12 1.07-.19.19 1.79.09 3.58.1 5.37v38.13c-.01 1.74.13 3.49-.15 5.22-.36-.03-.71-.05-1.06-.05-.95-3.75-1.72-7.55-2.62-11.31-.38-1.53-.58-3.09-1.07-4.59-1.7-.24-3.43-.17-5.15-.2-5.06-.01-10.13 0-15.19-.01-1.66-.01-3.32.09-4.98-.03-.03-.39-.26-.91.16-1.18 1.28-.65 2.72-.88 4.06-1.35 3.43-1.14 6.88-2.16 10.31-3.31 1.39-.48 2.9-.72 4.16-1.54.04-.56.02-1.13-.05-1.68-1.23-.55-2.53-.87-3.81-1.28-3.13-1.03-6.29-1.96-9.41-3.02-1.79-.62-3.67-1-5.41-1.79-.03-.37-.07-.73-.11-1.09 5.09-.19 10.2.06 15.3-.12 3.36-.13 6.73.08 10.09-.07.12-.39.26-.77.37-1.16 1.08-4.94 2.33-9.83 3.39-14.75zm5.97-.2c.36.05.72.12 1.08.2.98 3.85 1.73 7.76 2.71 11.61.36 1.42.56 2.88 1.03 4.27 2.53.18 5.07-.01 7.61.05 5.16.12 10.33.12 15.49.07.76-.01 1.52.03 2.28.08-.04.36-.07.72-.1 1.08-1.82.83-3.78 1.25-5.67 1.89-3.73 1.23-7.48 2.39-11.22 3.57-.57.17-1.12.42-1.67.64-.15.55-.18 1.12-.12 1.69.87.48 1.82.81 2.77 1.09 4.88 1.52 9.73 3.14 14.63 4.6.38.13.78.27 1.13.49.4.27.23.79.15 1.18-1.66.13-3.31.03-4.97.04-5.17.01-10.33-.01-15.5.01-1.61.03-3.22-.02-4.82.21-.52 1.67-.72 3.42-1.17 5.11-.94 3.57-1.52 7.24-2.54 10.78-.36.01-.71.02-1.06.06-.29-1.73-.15-3.48-.15-5.22v-38.13c.02-1.78-.08-3.58.11-5.37zM65.05 168.33c1.12-2.15 2.08-4.4 3.37-6.46-1.82 7.56-2.91 15.27-3.62 23-.8 7.71-.85 15.49-.54 23.23 1.05 19.94 5.54 39.83 14.23 57.88 2.99 5.99 6.35 11.83 10.5 17.11 6.12 7.47 12.53 14.76 19.84 21.09 4.8 4.1 9.99 7.78 15.54 10.8 3.27 1.65 6.51 3.39 9.94 4.68 5.01 2.03 10.19 3.61 15.42 4.94 3.83.96 7.78 1.41 11.52 2.71 5 1.57 9.47 4.61 13.03 8.43 4.93 5.23 8.09 11.87 10.2 18.67.99 2.9 1.59 5.91 2.17 8.92.15.75.22 1.52.16 2.29-6.5 2.78-13.26 5.06-20.26 6.18-4.11.78-8.29.99-12.46 1.08-10.25.24-20.47-1.76-30.12-5.12-3.74-1.42-7.49-2.85-11.03-4.72-8.06-3.84-15.64-8.7-22.46-14.46-2.92-2.55-5.83-5.13-8.4-8.03-9.16-9.83-16.3-21.41-21.79-33.65-2.39-5.55-4.61-11.18-6.37-16.96-1.17-3.94-2.36-7.89-3.26-11.91-.75-2.94-1.22-5.95-1.87-8.92-.46-2.14-.69-4.32-1.03-6.48-.85-5.43-1.28-10.93-1.33-16.43.11-6.18.25-12.37 1.07-18.5.4-2.86.67-5.74 1.15-8.6.98-5.7 2.14-11.37 3.71-16.93 3.09-11.65 7.48-22.95 12.69-33.84zm363.73-6.44c1.1 1.66 1.91 3.48 2.78 5.26 2.1 4.45 4.24 8.9 6.02 13.49 7.61 18.76 12.3 38.79 13.04 59.05.02 1.76.07 3.52.11 5.29.13 9.57-1.27 19.09-3.18 28.45-.73 3.59-1.54 7.17-2.58 10.69-4.04 14.72-10 29-18.41 41.78-8.21 12.57-19.01 23.55-31.84 31.41-5.73 3.59-11.79 6.64-18.05 9.19-5.78 2.19-11.71 4.03-17.8 5.11-6.4 1.05-12.91 1.52-19.4 1.23-7.92-.48-15.78-2.07-23.21-4.85-1.94-.8-3.94-1.46-5.84-2.33-.21-1.51.25-2.99.53-4.46 1.16-5.74 3.03-11.36 5.7-16.58 2.37-4.51 5.52-8.65 9.46-11.9 2.43-2.05 5.24-3.61 8.16-4.83 3.58-1.5 7.47-1.97 11.24-2.83 7.23-1.71 14.37-3.93 21.15-7 10.35-4.65 19.71-11.38 27.65-19.46 1.59-1.61 3.23-3.18 4.74-4.87 3.37-3.76 6.71-7.57 9.85-11.53 7.48-10.07 12.82-21.59 16.71-33.48 1.58-5.3 3.21-10.6 4.21-16.05.63-2.87 1.04-5.78 1.52-8.68.87-6.09 1.59-12.22 1.68-18.38.12-6.65.14-13.32-.53-19.94-.73-7.99-1.87-15.96-3.71-23.78z"],opencart:[640,512,[],"f23d","M423.3 440.7c0 25.3-20.3 45.6-45.6 45.6s-45.8-20.3-45.8-45.6 20.6-45.8 45.8-45.8c25.4 0 45.6 20.5 45.6 45.8zm-253.9-45.8c-25.3 0-45.6 20.6-45.6 45.8s20.3 45.6 45.6 45.6 45.8-20.3 45.8-45.6-20.5-45.8-45.8-45.8zm291.7-270C158.9 124.9 81.9 112.1 0 25.7c34.4 51.7 53.3 148.9 373.1 144.2 333.3-5 130 86.1 70.8 188.9 186.7-166.7 319.4-233.9 17.2-233.9z"],openid:[448,512,[],"f19b","M271.5 432l-68 32C88.5 453.7 0 392.5 0 318.2c0-71.5 82.5-131 191.7-144.3v43c-71.5 12.5-124 53-124 101.3 0 51 58.5 93.3 135.7 103v-340l68-33.2v384zM448 291l-131.3-28.5 36.8-20.7c-19.5-11.5-43.5-20-70-24.8v-43c46.2 5.5 87.7 19.5 120.3 39.3l35-19.8L448 291z"],opera:[496,512,[],"f26a","M313.9 32.7c-170.2 0-252.6 223.8-147.5 355.1 36.5 45.4 88.6 75.6 147.5 75.6 36.3 0 70.3-11.1 99.4-30.4-43.8 39.2-101.9 63-165.3 63-3.9 0-8 0-11.9-.3C104.6 489.6 0 381.1 0 248 0 111 111 0 248 0h.8c63.1.3 120.7 24.1 164.4 63.1-29-19.4-63.1-30.4-99.3-30.4zm101.8 397.7c-40.9 24.7-90.7 23.6-132-5.8 56.2-20.5 97.7-91.6 97.7-176.6 0-84.7-41.2-155.8-97.4-176.6 41.8-29.2 91.2-30.3 132.9-5 105.9 98.7 105.5 265.7-1.2 364z"],"optin-monster":[576,512,[],"f23c","M550.671 450.303c0 11.62-15.673 19.457-32.158 14.863-12.16-3.243-31.346-17.565-36.211-27.294-5.674-11.62 4.054-32.698 18.916-30.806 15.674 1.621 49.453 25.401 49.453 43.237zM372.86 75.223c-3.783-72.151-100.796-79.718-125.928-23.51 44.588-24.321 90.257-15.673 125.928 23.51zM74.795 407.066c-15.673 1.621-49.452 25.401-49.452 43.237 0 11.62 15.673 19.457 32.157 14.863 12.16-3.243 31.076-17.565 35.94-27.294 5.946-11.62-3.782-32.698-18.645-30.806zm497.765 14.322c1.081 3.513 1.892 7.026 1.892 10.809.81 31.616-44.317 64.045-73.503 65.125-17.295.81-34.59-8.377-42.696-23.51-113.497 4.053-226.994 4.864-340.22 0-8.377 15.133-25.672 24.05-42.967 23.51-28.915-1.081-74.043-33.509-73.503-65.125.27-3.783.811-7.296 1.892-10.809-5.566-9.463-4.845-15.282 5.405-11.62 3.243-5.134 7.026-9.458 11.08-13.782-2.57-10.917 1.27-14.094 11.079-9.188 4.594-3.243 9.998-6.485 15.944-9.188 0-15.757 11.839-11.131 17.295-5.675 12.467-1.78 20.129.709 26.753 5.675v-19.726c-12.987 0-40.641-11.375-45.94-36.212-4.974-20.725 2.607-38.075 25.132-47.56.81-5.945 8.107-14.052 14.862-15.944 7.567-1.892 12.431 4.594 14.052 10.269 7.425 0 17.757 1.465 21.078 8.107 5.405-.541 11.079-1.352 16.484-1.892-2.432-1.892-5.134-3.513-8.107-4.594-5.134-8.917-13.782-11.079-24.591-11.62 0-.81 0-1.621.27-2.702-19.727-.541-44.048-5.675-54.857-17.835-21.321-23.638-15.935-83.577 12.16-103.498 8.377-5.675 21.618-.811 22.699 9.728 2.425 20.598.399 26.833 26.212 25.942 8.107-7.836 16.755-14.592 26.483-19.997-14.862-1.352-28.914 1.621-43.778 3.783 12.752-12.48 23.953-25.442 56.748-42.427 23.511-11.89 49.993-20.808 76.205-23.239-18.646-7.837-39.993-11.891-59.721-16.484 76.475-16.214 174.569-22.159 244.289 37.562 18.105 15.403 32.427 36.211 42.696 59.992 39.799 4.853 36.47-5.581 38.643-25.132 1.081-10.269 14.322-15.403 22.699-9.458 14.862 10.539 22.159 30.806 24.59 48.101 2.162 17.835.27 41.345-12.43 55.127-10.809 12.16-34.32 17.565-53.776 18.105v2.703c-11.08.27-20.268 2.432-25.673 11.62-2.972 1.081-5.674 2.703-8.377 4.594 5.675.54 11.35 1.351 16.755 1.891 1.869-5.619 12.535-8.377 21.077-8.377 1.621-5.405 6.756-11.89 14.052-10.269s14.052 9.998 14.863 15.944c10.809 4.324 22.159 12.16 25.131 25.672 1.892 8.107 1.621 15.133.27 21.888-5.726 25.262-33.361 36.212-45.939 36.212 0 6.756 0 13.241-.27 19.726 8.01-6.006 16.367-7.158 26.752-5.675 5.919-5.919 17.565-9.41 17.565 5.675 5.675 2.703 11.349 5.945 15.944 9.188 10.1-5.051 13.669-.539 10.809 9.188 4.053 4.323 8.107 8.917 11.079 13.782 10.136-3.62 11.021 2.078 5.409 11.62zm-73.773-254.016c17.295 6.756 26.212 22.159 30.265 35.67 1.081-10.539-2.702-39.453-13.782-51.073-7.296-7.296-14.052-5.134-14.052.81.001 6.216-1.35 11.62-2.431 14.593zm-18.646 12.43c12.971 15.673 17.024 41.615 12.7 62.963 10.809-2.162 20.537-6.215 26.212-12.16 1.892-2.162 3.783-4.864 4.864-7.566-1.081-21.348-10.269-42.697-29.725-48.912-3.242 3.243-9.187 4.864-14.051 5.675zm-21.889.811c7.567 20.537 12.431 42.696 14.322 64.585 3.513 0 7.567-.27 11.62-.811 5.945-24.321-.27-51.614-14.052-63.504-3.783 0-8.107 0-11.89-.27zM77.768 167.372c-1.081-2.973-2.432-8.377-2.432-14.593 0-5.945-7.026-8.107-14.052-.81-11.35 11.62-14.863 40.534-13.782 51.073 4.053-13.512 12.971-28.915 30.266-35.67zm5.675 75.394c-4.324-21.348-.27-47.291 12.701-62.963-4.865-.811-10.809-2.432-14.052-5.675-19.457 6.215-28.375 27.563-29.726 48.912 1.351 2.702 2.972 5.404 4.864 7.566 5.675 6.215 15.403 9.998 26.213 12.16zm41.345-61.073c-5.134 1.081-9.998 2.973-14.862 4.865l-12.16 5.134v-.27c-7.296 14.052-9.999 34.319-5.405 52.965 4.594.541 8.647.811 12.7.811 2.432-22.159 9.188-43.778 19.727-63.505zm88.095-23.239c0 42.155 34.319 76.205 76.205 76.205s76.205-34.05 76.205-76.205c0-41.886-34.319-75.935-76.205-75.935s-76.205 34.049-76.205 75.935zm152.41 97.283c9.969 50.608 3.299 64.692 16.484 58.099 15.944-8.107 22.699-39.183 22.97-57.019-12.971-.81-26.213-.81-39.454-1.08zm-71.611-.541v-.27c-.27 5.134.27 38.103 4.324 41.075 11.079 5.405 39.453 4.594 51.073 1.081 5.405-1.621 2.432-37.022 1.621-41.886-18.916-.27-38.102-.27-57.018 0zm-14.053 0v-.27c-19.456.27-38.642.27-57.829.811-1.892 9.187-4.594 48.911 1.892 51.614 12.971 5.675 41.616 5.134 54.586 1.621 4.595-2.432 2.433-45.399 1.351-53.776zm-85.662 57.56c5.405 2.432 8.647 2.432 9.728-4.324 1.892-8.647 2.432-36.752 4.865-52.155-12.16.27-24.591.811-36.752 1.621-5.405 19.727.27 45.129 22.159 54.858zm-65.666-11.08c43.778 47.02 92.689 85.663 155.923 106.47 67.558-19.186 115.659-59.991 163.219-107.011-11.095-4.315-7.715-10.363-7.296-11.62-8.918-.81-17.835-1.892-26.483-2.702-9.458 32.968-35.94 52.965-46.75 31.616-2.702-5.134-3.513-11.62-4.594-16.754-3.783 8.377-13.242 8.107-24.591 8.918-13.241 1.081-31.617 1.351-44.048-2.972-2.972 12.971-11.079 12.971-26.752 14.322-14.052 1.352-48.642 4.054-54.857-10.809-1.081 28.644-35.13 9.998-45.129-7.026-3.243-5.675-5.405-11.35-7.026-17.565-7.837.81-15.673 1.621-23.511 2.702 2.443 3.663 1.549 9.052-8.105 12.431zM115.6 453.545c-5.674-23.239-18.646-49.722-33.508-54.046-22.429-6.756-68.909 23.51-66.207 54.586 12.701 19.457 39.994 35.67 59.181 36.481 17.835.81 35.94-11.08 39.724-28.914.539-2.432.81-5.134.81-8.107zm7.296-5.944c33.509-19.457 69.179-35.671 105.931-47.02-38.643-20.537-68.098-47.831-97.283-77.016-2.162 1.352-5.134 2.432-7.836 3.513-1.637 4.91 8.718 5.33 5.405 12.431-2.162 4.054-8.648 7.567-15.133 9.188-2.161 2.702-5.134 4.864-7.836 6.485h-.27c-.27 13.511-.27 27.024.27 40.535 8.939 15.964 15.426 33.314 16.752 51.884zm320.764 12.7c-36.752-21.348-74.044-41.345-115.659-52.965-13.782 6.215-27.833 11.349-42.155 15.403-2.162.811-2.162.811-4.324 0-11.89-3.783-23.239-8.107-34.859-13.241-40.265 11.62-77.286 29.185-112.416 50.803h-.27v.27c.27 0 .27 0 .27-.27 103.227 4.054 206.455 3.513 309.413 0zm27.023-64.045l-.27.27c.541-13.782.811-27.563.811-41.345-2.973-1.621-5.675-4.054-8.107-6.756-6.485-1.351-12.971-5.134-15.133-8.918-1.892-4.053 1.351-7.566 5.945-10.269-.27-.541-.541-1.621-.541-2.432-2.972-.811-5.405-1.892-7.567-3.243-31.616 29.455-65.396 56.749-103.498 76.746 38.914 11.62 75.935 28.104 111.875 47.561 1.05-14.692 7.231-35.749 16.485-51.614zm23.24 3.244c-14.593 4.323-27.834 30.806-33.509 54.046 0 23.826 21.278 37.897 40.534 37.022 19.186-.811 46.48-17.024 59.181-36.481 2.973-31.077-43.507-61.344-66.206-54.587zM290.709 134.133c.045 0 .089.003.134.003.046 0 .09-.003.136-.003h-.27zm0 96.743c28.645 0 51.884-21.618 51.884-48.371 0-36.092-40.507-58.079-72.151-44.318 9.458 2.972 16.484 11.62 16.484 21.618 0 23.257-33.291 31.955-46.48 11.35-7.297 34.067 19.368 59.721 50.263 59.721zM68.039 474.083c.54 6.486 12.16 12.701 21.618 9.458 6.756-2.703 14.593-10.539 17.295-16.214 2.973-7.026-1.081-19.997-9.728-18.375-8.917 1.621-29.725 16.754-29.185 25.131zm410.75-25.131c-8.377-1.621-12.431 11.349-9.458 18.375 2.432 5.675 10.269 13.511 17.295 16.214 9.187 3.243 21.078-2.972 21.348-9.458.811-8.377-20.267-23.51-29.185-25.131z"],osi:[495,512,[],"f41a","M0 259.2C2.3 123.4 97.4 26.8 213.8 11.1c138.8-18.6 255.6 75.8 278 201.1 21.3 118.8-44 230-151.6 274-9.3 3.8-14.4 1.7-18-7.7-17.8-46.3-35.6-92.7-53.4-139-3.1-8.1-1-13.2 7-16.8 24.2-11 39.3-29.4 43.3-55.8 6.4-42.4-24.5-78.7-64.5-82.2-39-3.4-71.8 23.7-77.5 59.7-5.2 33 11.1 63.7 41.9 77.7 9.6 4.4 11.5 8.6 7.8 18.4-17.9 46.6-35.8 93.2-53.7 139.9-2.6 6.9-8.3 9.3-15.5 6.5-52.6-20.3-101.4-61-130.8-119C1.9 318.7 1.6 280.2 0 259.2zm20.9-1.9c.4 6.6.6 14.3 1.3 22.1 6.3 71.9 49.6 143.5 131 183.1 3.2 1.5 4.4.8 5.6-2.3 14.9-39.1 29.9-78.2 45-117.3 1.3-3.3.6-4.8-2.4-6.7-31.6-19.9-47.3-48.5-45.6-86 1-21.6 9.3-40.5 23.8-56.3 30-32.7 77-39.8 115.5-17.6 31.9 18.4 49.5 53.8 45.2 90.4-3.6 30.6-19.3 53.9-45.7 69.8-2.7 1.6-3.5 2.9-2.3 6 15.2 39.2 30.2 78.4 45.2 117.7 1.2 3.1 2.4 3.8 5.6 2.3 35.5-16.6 65.2-40.3 88.1-72 34.8-48.2 49.1-101.9 42.3-161C459.8 112 354.1 14.7 218 31.5 111.9 44.5 22.7 134 20.9 257.3z"],page4:[496,512,[],"f3d7","M248 504C111 504 0 393 0 256S111 8 248 8c20.9 0 41.3 2.6 60.7 7.5L42.3 392H248v112zm0-143.6V146.8L98.6 360.4H248zm96 31.6v92.7c45.7-19.2 84.5-51.7 111.4-92.7H344zm57.4-138.2l-21.2 8.4 21.2 8.3v-16.7zm-20.3 54.5c-6.7 0-8 6.3-8 12.9v7.7h16.2v-10c0-5.9-2.3-10.6-8.2-10.6zM496 256c0 37.3-8.2 72.7-23 104.4H344V27.3C433.3 64.8 496 153.1 496 256zM360.4 143.6h68.2V96h-13.9v32.6h-13.9V99h-13.9v29.6h-12.7V96h-13.9v47.6zm68.1 185.3H402v-11c0-15.4-5.6-25.2-20.9-25.2-15.4 0-20.7 10.6-20.7 25.9v25.3h68.2v-15zm0-103l-68.2 29.7V268l68.2 29.5v-16.6l-14.4-5.7v-26.5l14.4-5.9v-16.9zm-4.8-68.5h-35.6V184H402v-12.2h11c8.6 15.8 1.3 35.3-18.6 35.3-22.5 0-28.3-25.3-15.5-37.7l-11.6-10.6c-16.2 17.5-12.2 63.9 27.1 63.9 34 0 44.7-35.9 29.3-65.3z"],pagelines:[384,512,[],"f18c","M384 312.7c-55.1 136.7-187.1 54-187.1 54-40.5 81.8-107.4 134.4-184.6 134.7-16.1 0-16.6-24.4 0-24.4 64.4-.3 120.5-42.7 157.2-110.1-41.1 15.9-118.6 27.9-161.6-82.2 109-44.9 159.1 11.2 178.3 45.5 9.9-24.4 17-50.9 21.6-79.7 0 0-139.7 21.9-149.5-98.1 119.1-47.9 152.6 76.7 152.6 76.7 1.6-16.7 3.3-52.6 3.3-53.4 0 0-106.3-73.7-38.1-165.2 124.6 43 61.4 162.4 61.4 162.4.5 1.6.5 23.8 0 33.4 0 0 45.2-89 136.4-57.5-4.2 134-141.9 106.4-141.9 106.4-4.4 27.4-11.2 53.4-20 77.5 0 0 83-91.8 172-20z"],palfed:[576,512,[],"f3d8","M384.9 193.9c0-47.4-55.2-44.2-95.4-29.8-1.3 39.4-2.5 80.7-3 119.8.7 2.8 2.6 6.2 15.1 6.2 36.8 0 83.4-42.8 83.3-96.2zm-194.5 72.2c.2 0 6.5-2.7 11.2-2.7 26.6 0 20.7 44.1-14.4 44.1-21.5 0-37.1-18.1-37.1-43 0-42 42.9-95.6 100.7-126.5 1-12.4 3-22 10.5-28.2 11.2-9 26.6-3.5 29.5 11.1 72.2-22.2 135.2 1 135.2 72 0 77.9-79.3 152.6-140.1 138.2-.1 39.4.9 74.4 2.7 100v.2c.2 3.4.6 12.5-5.3 19.1-9.6 10.6-33.4 10-36.4-22.3-4.1-44.4.2-206.1 1.4-242.5-21.5 15-58.5 50.3-58.5 75.9.2 2.5.4 4 .6 4.6zM8 181.1s-.1 37.4 38.4 37.4h30l22.4 217.2s0 44.3 44.7 44.3h288.9s44.7-.4 44.7-44.3l22.4-217.2h30s38.4 1.2 38.4-37.4c0 0 .1-37.4-38.4-37.4h-30.1c-7.3-25.6-30.2-74.3-119.4-74.3h-28V50.3s-2.7-18.4-21.1-18.4h-85.8s-21.1 0-21.1 18.4v19.1h-28.1s-105 4.2-120.5 74.3h-29S8 142.5 8 181.1z"],patreon:[512,512,[],"f3d9","M512 194.8c0 101.3-82.4 183.8-183.8 183.8-101.7 0-184.4-82.4-184.4-183.8 0-101.6 82.7-184.3 184.4-184.3C429.6 10.5 512 93.2 512 194.8zM0 501.5h90v-491H0v491z"],paypal:[384,512,[],"f1ed","M111.4 295.9c-3.5 19.2-17.4 108.7-21.5 134-.3 1.8-1 2.5-3 2.5H12.3c-7.6 0-13.1-6.6-12.1-13.9L58.8 46.6c1.5-9.6 10.1-16.9 20-16.9 152.3 0 165.1-3.7 204 11.4 60.1 23.3 65.6 79.5 44 140.3-21.5 62.6-72.5 89.5-140.1 90.3-43.4.7-69.5-7-75.3 24.2zM357.1 152c-1.8-1.3-2.5-1.8-3 1.3-2 11.4-5.1 22.5-8.8 33.6-39.9 113.8-150.5 103.9-204.5 103.9-6.1 0-10.1 3.3-10.9 9.4-22.6 140.4-27.1 169.7-27.1 169.7-1 7.1 3.5 12.9 10.6 12.9h63.5c8.6 0 15.7-6.3 17.4-14.9.7-5.4-1.1 6.1 14.4-91.3 4.6-22 14.3-19.7 29.3-19.7 71 0 126.4-28.8 142.9-112.3 6.5-34.8 4.6-71.4-23.8-92.6z"],periscope:[448,512,[],"f3da","M370 63.6C331.4 22.6 280.5 0 226.6 0 111.9 0 18.5 96.2 18.5 214.4c0 75.1 57.8 159.8 82.7 192.7C137.8 455.5 192.6 512 226.6 512c41.6 0 112.9-94.2 120.9-105 24.6-33.1 82-118.3 82-192.6 0-56.5-21.1-110.1-59.5-150.8zM226.6 493.9c-42.5 0-190-167.3-190-279.4 0-107.4 83.9-196.3 190-196.3 100.8 0 184.7 89 184.7 196.3.1 112.1-147.4 279.4-184.7 279.4zM338 206.8c0 59.1-51.1 109.7-110.8 109.7-100.6 0-150.7-108.2-92.9-181.8v.4c0 24.5 20.1 44.4 44.8 44.4 24.7 0 44.8-19.9 44.8-44.4 0-18.2-11.1-33.8-26.9-40.7 76.6-19.2 141 39.3 141 112.4z"],phabricator:[496,512,[],"f3db","M323 262.1l-.1-13s21.7-19.8 21.1-21.2l-9.5-20c-.6-1.4-29.5-.5-29.5-.5l-9.4-9.3s.2-28.5-1.2-29.1l-20.1-9.2c-1.4-.6-20.7 21-20.7 21l-13.1-.2s-20.5-21.4-21.9-20.8l-20 8.3c-1.4.5.2 28.9.2 28.9l-9.1 9.1s-29.2-.9-29.7.4l-8.1 19.8c-.6 1.4 21 21 21 21l.1 12.9s-21.7 19.8-21.1 21.2l9.5 20c.6 1.4 29.5.5 29.5.5l9.4 9.3s-.2 31.8 1.2 32.3l20.1 8.3c1.4.6 20.7-23.5 20.7-23.5l13.1.2s20.5 23.8 21.8 23.3l20-7.5c1.4-.6-.2-32.1-.2-32.1l9.1-9.1s29.2.9 29.7-.5l8.1-19.8c.7-1.1-20.9-20.7-20.9-20.7zm-44.9-8.7c.7 17.1-12.8 31.6-30.1 32.4-17.3.8-32.1-12.5-32.8-29.6-.7-17.1 12.8-31.6 30.1-32.3 17.3-.8 32.1 12.5 32.8 29.5zm201.2-37.9l-97-97-.1.1c-75.1-73.3-195.4-72.8-269.8 1.6-50.9 51-27.8 27.9-95.7 95.3-22.3 22.3-22.3 58.7 0 81 69.9 69.4 46.4 46 97.4 97l.1-.1c75.1 73.3 195.4 72.9 269.8-1.6 51-50.9 27.9-27.9 95.3-95.3 22.3-22.3 22.3-58.7 0-81zM140.4 363.8c-59.6-59.5-59.6-156 0-215.5 59.5-59.6 156-59.5 215.6 0 59.5 59.5 59.6 156 0 215.6-59.6 59.5-156 59.4-215.6-.1z"],"phoenix-framework":[640,512,[],"f3dc","M212.9 344.3c3.8-.1 22.8-1.4 25.6-2.2-2.4-2.6-43.6-1-68-49.6-4.3-8.6-7.5-17.6-6.4-27.6 2.9-25.5 32.9-30 52-18.5 36 21.6 63.3 91.3 113.7 97.5 37 4.5 84.6-17 108.2-45.4-.6-.1-.8-.2-1-.1-.4.1-.8.2-1.1.3-33.3 12.1-94.3 9.7-134.7-14.8-37.6-22.8-53.1-58.7-51.8-74.6 1.8-21.3 22.9-23.2 35.9-19.6 14.4 3.9 24.4 17.6 38.9 27.4 15.6 10.4 32.9 13.7 51.3 10.3 14.9-2.7 34.4-12.3 36.5-14.5-1.1-.1-1.8-.1-2.5-.2-6.2-.6-12.4-.8-18.5-1.7C279.8 194.5 262.1 47.4 138.5 37.9 94.2 34.5 39.1 46 2.2 72.9c-.8.6-1.5 1.2-2.2 1.8.1.2.1.3.2.5.8 0 1.6-.1 2.4-.2 6.3-1 12.5-.8 18.7.3 23.8 4.3 47.7 23.1 55.9 76.5 5.3 34.3-.7 50.8 8 86.1 19 77.1 91 107.6 127.7 106.4zM75.3 64.9c-.9-1-.9-1.2-1.3-2 12.1-2.6 24.2-4.1 36.6-4.8-1.1 14.7-22.2 21.3-35.3 6.8zm196.9 350.5c-42.8 1.2-92-26.7-123.5-61.4-4.6-5-16.8-20.2-18.6-23.4l.4-.4c6.6 4.1 25.7 18.6 54.8 27 24.2 7 48.1 6.3 71.6-3.3 22.7-9.3 41-.5 43.1 2.9-18.5 3.8-20.1 4.4-24 7.9-5.1 4.4-4.6 11.7 7 17.2 26.2 12.4 63-2.8 97.2 25.4 2.4 2 8.1 7.8 10.1 10.7-.1.2-.3.3-.4.5-4.8-1.5-16.4-7.5-40.2-9.3-24.7-2-46.3 5.3-77.5 6.2zm174.8-252c16.4-5.2 41.3-13.4 66.5-3.3 16.1 6.5 26.2 18.7 32.1 34.6 3.5 9.4 5.1 19.7 5.1 28.7-.2 0-.4 0-.6.1-.2-.4-.4-.9-.5-1.3-5-22-29.9-43.8-67.6-29.9-50.2 18.6-130.4 9.7-176.9-48-.7-.9-2.4-1.7-1.3-3.2.1-.2 2.1.6 3 1.3 18.1 13.4 38.3 21.9 60.3 26.2 30.5 6.1 54.6 2.9 79.9-5.2zm102.7 117.5c-32.4.2-33.8 50.1-103.6 64.4-18.2 3.7-38.7 4.6-44.9 4.2v-.4c2.8-1.5 14.7-2.6 29.7-16.6 7.9-7.3 15.3-15.1 22.8-22.9 19.5-20.2 41.4-42.2 81.9-39 23.1 1.8 29.3 8.2 36.1 12.7.3.2.4.5.7.9-.5 0-.7.1-.9 0-7-2.7-14.3-3.3-21.8-3.3zm-12.3-24.1c-.1.2-.1.4-.2.6-28.9-4.4-48-7.9-68.5 4-17 9.9-31.4 20.5-62 24.4-27.1 3.4-45.1 2.4-66.1-8-.3-.2-.6-.4-1-.6 0-.2.1-.3.1-.5 24.9 3.8 36.4 5.1 55.5-5.8 22.3-12.9 40.1-26.6 71.3-31 29.6-4.1 51.3 2.5 70.9 16.9zM268.6 97.3c-.6-.6-1.1-1.2-2.1-2.3 7.6 0 29.7-1.2 53.4 8.4 19.7 8 32.2 21 50.2 32.9 11.1 7.3 23.4 9.3 36.4 8.1 4.3-.4 8.5-1.2 12.8-1.7.4-.1.9 0 1.5.3-.6.4-1.2.9-1.8 1.2-8.1 4-16.7 6.3-25.6 7.1-26.1 2.6-50.3-3.7-73.4-15.4-19.3-9.9-36.4-22.9-51.4-38.6zM640 335.7c-3.5 3.1-22.7 11.6-42.7 5.3-12.3-3.9-19.5-14.9-31.6-24.1-10-7.6-20.9-7.9-28.1-8.4.6-.8.9-1.2 1.2-1.4 14.8-9.2 30.5-12.2 47.3-6.5 12.5 4.2 19.2 13.5 30.4 24.2 10.8 10.4 21 9.9 23.1 10.5.1-.1.2 0 .4.4zm-212.5 137c2.2 1.2 1.6 1.5 1.5 2-18.5-1.4-33.9-7.6-46.8-22.2-21.8-24.7-41.7-27.9-48.6-29.7.5-.2.8-.4 1.1-.4 13.1.1 26.1.7 38.9 3.9 25.3 6.4 35 25.4 41.6 35.3 3.2 4.8 7.3 8.3 12.3 11.1z"],"phoenix-squadron":[513,512,[],"f511","M96.24 62.81C142.91 26.57 202.15 6.57 261.28 8.08c29.67-.38 59.29 5.38 87.17 15.37-24.2-4.64-49.18-6.35-73.6-2.45-43 5.35-83.26 27.23-112.16 59.35 5.69-.99 10.81-3.68 16.07-5.88 18.19-7.89 37.6-13.29 57.4-14.87 19.8-2.14 39.75-.43 59.45 1.93-14.46 2.79-29.2 4.58-43.11 9.61-34.53 11.11-65.46 33.26-86.55 62.82-13.84 19.77-23.7 42.99-24.74 67.33-.35 16.54 5.23 34.91 19.89 44.17 11.13 6.66 24.85 9.39 37.63 6.76 15.49-2.47 30.16-8.67 43.73-16.38 11.55-6.84 22.73-14.59 32.05-24.32 3.8-3.23 2.54-8.48 2.63-12.83-2.13-.34-4.4-1.11-6.32.3a203.975 203.975 0 0 1-35.93 15.42c-20.07 6.19-42.28 8.48-62.28.78 12.83 1.73 26.14.31 37.85-5.46 20.29-9.75 36.92-25.27 54.6-38.88 27.84-21.29 57.64-40.11 89.17-55.47 25.78-12.01 53.09-22.85 81.81-24.2-15.68 13.76-32.25 26.6-46.92 41.51-14.55 14.04-27.54 29.58-40.23 45.31-3.53 4.61-8.98 6.96-13.62 10.19-22.24 15.03-40.6 35.96-52.04 60.28-9.36 19.74-14.55 41.97-11.81 63.84 1.95 13.73 8.74 27.67 20.96 35.01 12.94 7.99 29.14 8.09 43.61 5.11 32.9-7.47 61.61-28.97 81.28-56 20.5-27.6 30.61-62.38 29.25-96.64-.52-7.52-1.58-15-1.67-22.55 8.02 19.54 14.87 39.83 16.7 61.01 2.01 14.32.75 28.84-1.62 43.02-1.92 11.02-5.69 21.58-7.81 32.53 20.36-22.73 34.17-51.24 39.46-81.31 5.72-35.37.58-72.36-14.25-104.95 20.84 32.12 32.43 69.79 35.81 107.8.5 12.77.5 25.58 0 38.34-2.91 34.26-12.97 67.95-29.76 98-26.2 47.48-68.2 85.89-117.54 108.32-78.52 36.34-175.2 31.41-248.72-14.72-38.84-23.78-71.06-58.32-91.68-98.96C10.72 337.43 2.04 305.5 0 273.13V241.7c3.94-69.97 40.99-136.32 96.24-178.89m222.65 80.57c5.51-.8 10.82-2.57 16.02-4.5 4.99-1.77 9.27-5.95 10.35-11.25-8.91 5-17.95 9.95-26.37 15.75z"],php:[640,512,[],"f457","M320 104.5c171.4 0 303.2 72.2 303.2 151.5S491.3 407.5 320 407.5c-171.4 0-303.2-72.2-303.2-151.5S148.7 104.5 320 104.5m0-16.8C143.3 87.7 0 163 0 256s143.3 168.3 320 168.3S640 349 640 256 496.7 87.7 320 87.7zM218.2 242.5c-7.9 40.5-35.8 36.3-70.1 36.3l13.7-70.6c38 0 63.8-4.1 56.4 34.3zM97.4 350.3h36.7l8.7-44.8c41.1 0 66.6 3 90.2-19.1 26.1-24 32.9-66.7 14.3-88.1-9.7-11.2-25.3-16.7-46.5-16.7h-70.7L97.4 350.3zm185.7-213.6h36.5l-8.7 44.8c31.5 0 60.7-2.3 74.8 10.7 14.8 13.6 7.7 31-8.3 113.1h-37c15.4-79.4 18.3-86 12.7-92-5.4-5.8-17.7-4.6-47.4-4.6l-18.8 96.6h-36.5l32.7-168.6zM505 242.5c-8 41.1-36.7 36.3-70.1 36.3l13.7-70.6c38.2 0 63.8-4.1 56.4 34.3zM384.2 350.3H421l8.7-44.8c43.2 0 67.1 2.5 90.2-19.1 26.1-24 32.9-66.7 14.3-88.1-9.7-11.2-25.3-16.7-46.5-16.7H417l-32.8 168.7z"],"pied-piper":[448,512,[],"f2ae","M32 419L0 479.2l.8-328C.8 85.3 54 32 120 32h327.2c-93 28.9-189.9 94.2-253.9 168.6C122.7 282 82.6 338 32 419M448 32S305.2 98.8 261.6 199.1c-23.2 53.6-28.9 118.1-71 158.6-28.9 27.8-69.8 38.2-105.3 56.3-23.2 12-66.4 40.5-84.9 66h328.4c66 0 119.3-53.3 119.3-119.2-.1 0-.1-328.8-.1-328.8z"],"pied-piper-alt":[576,512,[],"f1a8","M242 187c6.3-11.8 13.2-17 25.9-21.8 27.3-10.3 40.2-30.5 58.9-51.1 11.9 8.4 12 24.6 31.6 23v21.8l6.3.3c37.4-14.4 74.7-30.2 106.6-54.6 48.3-36.8 52.9-50 81.3-100l2-2.6c-.6 14.1-6.3 27.3-12.4 39.9-30.5 63.8-78.7 100.3-146.8 116.7-12.4 2.9-26.4 3.2-37.6 8.9 1.4 9.8 13.2 18.1 13.2 23 0 3.4-5.5 7.2-7.5 8.6-11.2-12.9-16.1-19.3-22.7-22.1-7.6-3.5-63.9-6.4-98.8 10zm137.9 256.9c-19 0-64.1 9.5-79.9 19.8l6.9 45.1c35.7 6.1 70.1 3.6 106-9.8-4.8-10-23.5-55.1-33-55.1zM244 246c-3.2-2-6.3-2.9-10.1-2.9-6.6 0-12.6 3.2-19.3 3.7l1.7 4.9L244 246zm-12.6 31.8l24.1 61.2 21-13.8-31.3-50.9-13.8 3.5zM555.5 0l-.6 1.1-.3.9.6-.6.3-1.4zm-59.2 382.1c-33.9-56.9-75.3-118.4-150-115.5l-.3-6c-1.1-13.5 32.8 3.2 35.1-31l-14.4 7.2c-19.8-45.7-8.6-54.3-65.5-54.3-14.7 0-26.7 1.7-41.4 4.6 2.9 18.6 2.2 36.7-10.9 50.3l19.5 5.5c-1.7 3.2-2.9 6.3-2.9 9.8 0 21 42.8 2.9 42.8 33.6 0 18.4-36.8 60.1-54.9 60.1-8 0-53.7-50-53.4-60.1l.3-4.6 52.3-11.5c13-2.6 12.3-22.7-2.9-22.7-3.7 0-43.1 9.2-49.4 10.6-2-5.2-7.5-14.1-13.8-14.1-3.2 0-6.3 3.2-9.5 4-9.2 2.6-31 2.9-21.5 20.1L15.9 298.5c-5.5 1.1-8.9 6.3-8.9 11.8 0 6 5.5 10.9 11.5 10.9 8 0 131.3-28.4 147.4-32.2 2.6 3.2 4.6 6.3 7.8 8.6 20.1 14.4 59.8 85.9 76.4 85.9 24.1 0 58-22.4 71.3-41.9 3.2-4.3 6.9-7.5 12.4-6.9.6 13.8-31.6 34.2-33 43.7-1.4 10.2-1 35.2-.3 41.1 26.7 8.1 52-3.6 77.9-2.9 4.3-21 10.6-41.9 9.8-63.5l-.3-9.5c-1.4-34.2-10.9-38.5-34.8-58.6-1.1-1.1-2.6-2.6-3.7-4 2.2-1.4 1.1-1 4.6-1.7 88.5 0 56.3 183.6 111.5 229.9 33.1-15 72.5-27.9 103.5-47.2-29-25.6-52.6-45.7-72.7-79.9zm-196.2 46v27.3l11.8-3.4-2.9-23.8h-8.9zm76.1 2.9c0-1.4-.6-3.2-.9-4.6-26.8 0-36.9 3.8-59.5 6.3l2 12.4c9-1.5 58.4-6.6 58.4-14.1z"],"pied-piper-hat":[640,512,[],"f4e5","M640 24.9c-80.8 53.6-89.4 92.5-96.4 104.4-6.7 12.2-11.7 60.3-23.3 83.6-11.7 23.6-54.2 42.2-66.1 50-11.7 7.8-28.3 38.1-41.9 64.2-108.1-4.4-167.4 38.8-259.2 93.6 29.4-9.7 43.3-16.7 43.3-16.7 94.2-36 139.3-68.3 281.1-49.2 1.1 0 1.9.6 2.8.8 3.9 2.2 5.3 6.9 3.1 10.8l-53.9 95.8c-2.5 4.7-7.8 7.2-13.1 6.1-126.8-23.8-226.9 17.3-318.9 18.6C24.1 488 0 453.4 0 451.8c0-1.1.6-1.7 1.7-1.7 0 0 38.3 0 103.1-15.3C178.4 294.5 244 245.4 315.4 245.4c0 0 71.7 0 90.6 61.9 22.8-39.7 28.3-49.2 28.3-49.2 5.3-9.4 35-77.2 86.4-141.4 51.5-64 90.4-79.9 119.3-91.8z"],"pied-piper-pp":[448,512,[],"f1a7","M205.3 174.6c0 21.1-14.2 38.1-31.7 38.1-7.1 0-12.8-1.2-17.2-3.7v-68c4.4-2.7 10.1-4.2 17.2-4.2 17.5 0 31.7 16.9 31.7 37.8zm52.6 67c-7.1 0-12.8 1.5-17.2 4.2v68c4.4 2.5 10.1 3.7 17.2 3.7 17.4 0 31.7-16.9 31.7-37.8 0-21.1-14.3-38.1-31.7-38.1zM448 80v352c0 26.5-21.5 48-48 48H48c-26.5 0-48-21.5-48-48V80c0-26.5 21.5-48 48-48h352c26.5 0 48 21.5 48 48zM185 255.1c41 0 74.2-35.6 74.2-79.6 0-44-33.2-79.6-74.2-79.6-12 0-24.1 3.2-34.6 8.8h-45.7V311l51.8-10.1v-50.6c8.6 3.1 18.1 4.8 28.5 4.8zm158.4 25.3c0-44-33.2-79.6-73.9-79.6-3.2 0-6.4.2-9.6.7-3.7 12.5-10.1 23.8-19.2 33.4-13.8 15-32.2 23.8-51.8 24.8V416l51.8-10.1v-50.6c8.6 3.2 18.2 4.7 28.7 4.7 40.8 0 74-35.6 74-79.6z"],pinterest:[496,512,[],"f0d2","M496 256c0 137-111 248-248 248-25.6 0-50.2-3.9-73.4-11.1 10.1-16.5 25.2-43.5 30.8-65 3-11.6 15.4-59 15.4-59 8.1 15.4 31.7 28.5 56.8 28.5 74.8 0 128.7-68.8 128.7-154.3 0-81.9-66.9-143.2-152.9-143.2-107 0-163.9 71.8-163.9 150.1 0 36.4 19.4 81.7 50.3 96.1 4.7 2.2 7.2 1.2 8.3-3.3.8-3.4 5-20.3 6.9-28.1.6-2.5.3-4.7-1.7-7.1-10.1-12.5-18.3-35.3-18.3-56.6 0-54.7 41.4-107.6 112-107.6 60.9 0 103.6 41.5 103.6 100.9 0 67.1-33.9 113.6-78 113.6-24.3 0-42.6-20.1-36.7-44.8 7-29.5 20.5-61.3 20.5-82.6 0-19-10.2-34.9-31.4-34.9-24.9 0-44.9 25.7-44.9 60.2 0 22 7.4 36.8 7.4 36.8s-24.5 103.8-29 123.2c-5 21.4-3 51.6-.9 71.2C65.4 450.9 0 361.1 0 256 0 119 111 8 248 8s248 111 248 248z"],"pinterest-p":[384,512,[],"f231","M204 6.5C101.4 6.5 0 74.9 0 185.6 0 256 39.6 296 63.6 296c9.9 0 15.6-27.6 15.6-35.4 0-9.3-23.7-29.1-23.7-67.8 0-80.4 61.2-137.4 140.4-137.4 68.1 0 118.5 38.7 118.5 109.8 0 53.1-21.3 152.7-90.3 152.7-24.9 0-46.2-18-46.2-43.8 0-37.8 26.4-74.4 26.4-113.4 0-66.2-93.9-54.2-93.9 25.8 0 16.8 2.1 35.4 9.6 50.7-13.8 59.4-42 147.9-42 209.1 0 18.9 2.7 37.5 4.5 56.4 3.4 3.8 1.7 3.4 6.9 1.5 50.4-69 48.6-82.5 71.4-172.8 12.3 23.4 44.1 36 69.3 36 106.2 0 153.9-103.5 153.9-196.8C384 71.3 298.2 6.5 204 6.5z"],"pinterest-square":[448,512,[],"f0d3","M448 80v352c0 26.5-21.5 48-48 48H154.4c9.8-16.4 22.4-40 27.4-59.3 3-11.5 15.3-58.4 15.3-58.4 8 15.3 31.4 28.2 56.3 28.2 74.1 0 127.4-68.1 127.4-152.7 0-81.1-66.2-141.8-151.4-141.8-106 0-162.2 71.1-162.2 148.6 0 36 19.2 80.8 49.8 95.1 4.7 2.2 7.1 1.2 8.2-3.3.8-3.4 5-20.1 6.8-27.8.6-2.5.3-4.6-1.7-7-10.1-12.3-18.3-34.9-18.3-56 0-54.2 41-106.6 110.9-106.6 60.3 0 102.6 41.1 102.6 99.9 0 66.4-33.5 112.4-77.2 112.4-24.1 0-42.1-19.9-36.4-44.4 6.9-29.2 20.3-60.7 20.3-81.8 0-53-75.5-45.7-75.5 25 0 21.7 7.3 36.5 7.3 36.5-31.4 132.8-36.1 134.5-29.6 192.6l2.2.8H48c-26.5 0-48-21.5-48-48V80c0-26.5 21.5-48 48-48h352c26.5 0 48 21.5 48 48z"],playstation:[576,512,[],"f3df","M570.9 372.3c-11.3 14.2-38.8 24.3-38.8 24.3L327 470.2v-54.3l150.9-53.8c17.1-6.1 19.8-14.8 5.8-19.4-13.9-4.6-39.1-3.3-56.2 2.9L327 381.1v-56.4c23.2-7.8 47.1-13.6 75.7-16.8 40.9-4.5 90.9.6 130.2 15.5 44.2 14 49.2 34.7 38 48.9zm-224.4-92.5v-139c0-16.3-3-31.3-18.3-35.6-11.7-3.8-19 7.1-19 23.4v347.9l-93.8-29.8V32c39.9 7.4 98 24.9 129.2 35.4C424.1 94.7 451 128.7 451 205.2c0 74.5-46 102.8-104.5 74.6zM43.2 410.2c-45.4-12.8-53-39.5-32.3-54.8 19.1-14.2 51.7-24.9 51.7-24.9l134.5-47.8v54.5l-96.8 34.6c-17.1 6.1-19.7 14.8-5.8 19.4 13.9 4.6 39.1 3.3 56.2-2.9l46.4-16.9v48.8c-51.6 9.3-101.4 7.3-153.9-10z"],"product-hunt":[512,512,[],"f288","M326.3 218.8c0 20.5-16.7 37.2-37.2 37.2h-70.3v-74.4h70.3c20.5 0 37.2 16.7 37.2 37.2zM504 256c0 137-111 248-248 248S8 393 8 256 119 8 256 8s248 111 248 248zm-128.1-37.2c0-47.9-38.9-86.8-86.8-86.8H169.2v248h49.6v-74.4h70.3c47.9 0 86.8-38.9 86.8-86.8z"],pushed:[432,512,[],"f3e1","M407 111.9l-98.5-9 14-33.4c10.4-23.5-10.8-40.4-28.7-37L22.5 76.9c-15.1 2.7-26 18.3-21.4 36.6l105.1 348.3c6.5 21.3 36.7 24.2 47.7 7l35.3-80.8 235.2-231.3c16.4-16.8 4.3-42.9-17.4-44.8zM297.6 53.6c5.1-.7 7.5 2.5 5.2 7.4L286 100.9 108.6 84.6l189-31zM22.7 107.9c-3.1-5.1 1-10 6.1-9.1l248.7 22.7-96.9 230.7L22.7 107.9zM136 456.4c-2.6 4-7.9 3.1-9.4-1.2L43.5 179.7l127.7 197.6c-7 15-35.2 79.1-35.2 79.1zm272.8-314.5L210.1 337.3l89.7-213.7 106.4 9.7c4 1.1 5.7 5.3 2.6 8.6z"],python:[448,512,[],"f3e2","M167.8 36.4c-45.2 8-53.4 24.7-53.4 55.6v40.7h106.9v13.6h-147c-31.1 0-58.3 18.7-66.8 54.2-9.8 40.7-10.2 66.1 0 108.6 7.6 31.6 25.7 54.2 56.8 54.2H101v-48.8c0-35.3 30.5-66.4 66.8-66.4h106.8c29.7 0 53.4-24.5 53.4-54.3V91.9c0-29-24.4-50.7-53.4-55.6-35.8-5.9-74.7-5.6-106.8.1zm-6.7 28.4c11 0 20.1 9.2 20.1 20.4s-9 20.3-20.1 20.3c-11.1 0-20.1-9.1-20.1-20.3.1-11.3 9-20.4 20.1-20.4zm185.2 81.4v47.5c0 36.8-31.2 67.8-66.8 67.8H172.7c-29.2 0-53.4 25-53.4 54.3v101.8c0 29 25.2 46 53.4 54.3 33.8 9.9 66.3 11.7 106.8 0 26.9-7.8 53.4-23.5 53.4-54.3v-40.7H226.2v-13.6h160.2c31.1 0 42.6-21.7 53.4-54.2 11.2-33.5 10.7-65.7 0-108.6-7.7-30.9-22.3-54.2-53.4-54.2h-40.1zM286.2 404c11.1 0 20.1 9.1 20.1 20.3 0 11.3-9 20.4-20.1 20.4-11 0-20.1-9.2-20.1-20.4.1-11.3 9.1-20.3 20.1-20.3z"],qq:[448,512,[],"f1d6","M433.754 420.445c-11.526 1.393-44.86-52.741-44.86-52.741 0 31.345-16.136 72.247-51.051 101.786 16.842 5.192 54.843 19.167 45.803 34.421-7.316 12.343-125.51 7.881-159.632 4.037-34.122 3.844-152.316 8.306-159.632-4.037-9.045-15.25 28.918-29.214 45.783-34.415-34.92-29.539-51.059-70.445-51.059-101.792 0 0-33.334 54.134-44.859 52.741-5.37-.65-12.424-29.644 9.347-99.704 10.261-33.024 21.995-60.478 40.144-105.779C60.683 98.063 108.982.006 224 0c113.737.006 163.156 96.133 160.264 214.963 18.118 45.223 29.912 72.85 40.144 105.778 21.768 70.06 14.716 99.053 9.346 99.704z"],quinscape:[489,512,[],"f459","M301.9 474.6h-1.3c-87.3 0-158.1-70.8-158.1-158.1s70.8-158.1 158.1-158.1c94.9 0 168.2 83.1 157 176.6 4 5.1 8.2 9.6 11.2 15.3 13.4-30.3 20.3-62.4 20.3-97.7C489.1 117.5 379.6 8 244.5 8 109.5 8 0 117.5 0 252.6s109.5 244.6 244.5 244.6c24.8 0 47.8-3.2 70.4-10.1-5.2-3.5-9-8.2-13.4-12.6l.4.1zm-21.2-69.8c0-54.8 44.4-99.2 99.2-99.2 54.8 0 99.2 44.4 99.2 99.2 0 54.8-44.4 99.2-99.2 99.2-54.8 0-99.2-44.4-99.2-99.2"],quora:[448,512,[],"f2c4","M440.5 386.7h-29.3c-1.5 13.5-10.5 30.8-33 30.8-20.5 0-35.3-14.2-49.5-35.8 44.2-34.2 74.7-87.5 74.7-153C403.5 111.2 306.8 32 205 32 105.3 32 7.3 111.7 7.3 228.7c0 134.1 131.3 221.6 249 189C276 451.3 302 480 351.5 480c81.8 0 90.8-75.3 89-93.3zM297 329.2C277.5 300 253.3 277 205.5 277c-30.5 0-54.3 10-69 22.8l12.2 24.3c6.2-3 13-4 19.8-4 35.5 0 53.7 30.8 69.2 61.3-10 3-20.7 4.2-32.7 4.2-75 0-107.5-53-107.5-156.7C97.5 124.5 130 71 205 71c76.2 0 108.7 53.5 108.7 157.7.1 41.8-5.4 75.6-16.7 100.5z"],"r-project":[581,512,[],"f4f7","M581 226.6C581 119.1 450.9 32 290.5 32S0 119.1 0 226.6C0 322.4 103.3 402 239.4 418.1V480h99.1v-61.5c24.3-2.7 47.6-7.4 69.4-13.9L448 480h112l-67.4-113.7c54.5-35.4 88.4-84.9 88.4-139.7zm-466.8 14.5c0-73.5 98.9-133 220.8-133s211.9 40.7 211.9 133c0 50.1-26.5 85-70.3 106.4-2.4-1.6-4.7-2.9-6.4-3.7-10.2-5.2-27.8-10.5-27.8-10.5s86.6-6.4 86.6-92.7-90.6-87.9-90.6-87.9h-199V361c-74.1-21.5-125.2-67.1-125.2-119.9zm225.1 38.3v-55.6c57.8 0 87.8-6.8 87.8 27.3 0 36.5-38.2 28.3-87.8 28.3zm-.9 72.5H365c10.8 0 18.9 11.7 24 19.2-16.1 1.9-33 2.8-50.6 2.9v-22.1z"],ravelry:[512,512,[],"f2d9","M407.4 61.5C331.6 22.1 257.8 31 182.9 66c-11.3 5.2-15.5 10.6-19.9 19-10.3 19.2-16.2 37.4-19.9 52.7-21.2 25.6-36.4 56.1-43.3 89.9-10.6 18-20.9 41.4-23.1 71.4 0 0-.7 7.6-.5 7.9-35.3-4.6-76.2-27-76.2-27 9.1 14.5 61.3 32.3 76.3 37.9 0 0 1.7 98 64.5 131.2-11.3-17.2-13.3-20.2-13.3-20.2S94.8 369 100.4 324.7c.7 0 1.5.2 2.2.2 23.9 87.4 103.2 151.4 196.9 151.4 6.2 0 12.1-.2 18-.7 14 1.5 27.6.5 40.1-3.9 6.9-2.2 13.8-6.4 20.2-10.8 70.2-39.1 100.9-82 123.1-147.7 5.4-16 8.1-35.5 9.8-52.2 8.7-82.3-30.6-161.6-103.3-199.5zM138.8 163.2s-1.2 12.3-.7 19.7c-3.4 2.5-10.1 8.1-18.2 16.7 5.2-12.8 11.3-25.1 18.9-36.4zm-31.2 121.9c4.4-17.2 13.3-39.1 29.8-55.1 0 0 1.7 48 15.8 90.1l-41.4-6.9c-2.2-9.2-3.5-18.5-4.2-28.1zm7.9 42.8c14.8 3.2 34 7.6 43.1 9.1 27.3 76.8 108.3 124.3 108.3 124.3 1 .5 1.7.7 2.7 1-73.1-11.6-132.7-64.7-154.1-134.4zM386 444.1c-14.5 4.7-36.2 8.4-64.7 3.7 0 0-91.1-23.1-127.5-107.8 38.2.7 52.4-.2 78-3.9 39.4-5.7 79-16.2 115-33 11.8-5.4 11.1-19.4 9.6-29.8-2-12.8-11.1-12.1-21.4-4.7 0 0-82 58.6-189.8 53.7-18.7-32-26.8-110.8-26.8-110.8 41.4-35.2 83.2-59.6 168.4-52.4.2-6.4 3-27.1-20.4-28.1 0 0-93.5-11.1-146 33.5 2.5-16.5 5.9-29.3 11.1-39.4 34.2-30.8 79-49.5 128.3-49.5 106.4 0 193 87.1 193 194.5-.2 76-43.8 142-106.8 174z"],react:[512,512,[],"f41b","M418.2 177.2c-5.4-1.8-10.8-3.5-16.2-5.1.9-3.7 1.7-7.4 2.5-11.1 12.3-59.6 4.2-107.5-23.1-123.3-26.3-15.1-69.2.6-112.6 38.4-4.3 3.7-8.5 7.6-12.5 11.5-2.7-2.6-5.5-5.2-8.3-7.7-45.5-40.4-91.1-57.4-118.4-41.5-26.2 15.2-34 60.3-23 116.7 1.1 5.6 2.3 11.1 3.7 16.7-6.4 1.8-12.7 3.8-18.6 5.9C38.3 196.2 0 225.4 0 255.6c0 31.2 40.8 62.5 96.3 81.5 4.5 1.5 9 3 13.6 4.3-1.5 6-2.8 11.9-4 18-10.5 55.5-2.3 99.5 23.9 114.6 27 15.6 72.4-.4 116.6-39.1 3.5-3.1 7-6.3 10.5-9.7 4.4 4.3 9 8.4 13.6 12.4 42.8 36.8 85.1 51.7 111.2 36.6 27-15.6 35.8-62.9 24.4-120.5-.9-4.4-1.9-8.9-3-13.5 3.2-.9 6.3-1.9 9.4-2.9 57.7-19.1 99.5-50 99.5-81.7 0-30.3-39.4-59.7-93.8-78.4zM282.9 92.3c37.2-32.4 71.9-45.1 87.7-36 16.9 9.7 23.4 48.9 12.8 100.4-.7 3.4-1.4 6.7-2.3 10-22.2-5-44.7-8.6-67.3-10.6-13-18.6-27.2-36.4-42.6-53.1 3.9-3.7 7.7-7.2 11.7-10.7zm-130 189.1c4.6 8.8 9.3 17.5 14.3 26.1 5.1 8.7 10.3 17.4 15.8 25.9-15.6-1.7-31.1-4.2-46.4-7.5 4.4-14.4 9.9-29.3 16.3-44.5zm0-50.6c-6.3-14.9-11.6-29.5-16-43.6 14.4-3.2 29.7-5.8 45.6-7.8-5.3 8.3-10.5 16.8-15.4 25.4-4.9 8.5-9.7 17.2-14.2 26zm11.4 25.3c6.6-13.8 13.8-27.3 21.4-40.6 7.6-13.3 15.8-26.2 24.4-38.9 15-1.1 30.3-1.7 45.9-1.7 15.6 0 31 .6 45.9 1.7 8.5 12.6 16.6 25.5 24.3 38.7 7.7 13.2 14.9 26.7 21.7 40.4-6.7 13.8-13.9 27.4-21.6 40.8-7.6 13.3-15.7 26.2-24.2 39-14.9 1.1-30.4 1.6-46.1 1.6-15.7 0-30.9-.5-45.6-1.4-8.7-12.7-16.9-25.7-24.6-39-7.7-13.3-14.8-26.8-21.5-40.6zm180.6 51.2c5.1-8.8 9.9-17.7 14.6-26.7 6.4 14.5 12 29.2 16.9 44.3-15.5 3.5-31.2 6.2-47 8 5.4-8.4 10.5-17 15.5-25.6zm14.4-76.5c-4.7-8.8-9.5-17.6-14.5-26.2-4.9-8.5-10-16.9-15.3-25.2 16.1 2 31.5 4.7 45.9 8-4.6 14.8-10 29.2-16.1 43.4zM256.2 118.3c10.5 11.4 20.4 23.4 29.6 35.8-19.8-.9-39.7-.9-59.5 0 9.8-12.9 19.9-24.9 29.9-35.8zM140.2 57c16.8-9.8 54.1 4.2 93.4 39 2.5 2.2 5 4.6 7.6 7-15.5 16.7-29.8 34.5-42.9 53.1-22.6 2-45 5.5-67.2 10.4-1.3-5.1-2.4-10.3-3.5-15.5-9.4-48.4-3.2-84.9 12.6-94zm-24.5 263.6c-4.2-1.2-8.3-2.5-12.4-3.9-21.3-6.7-45.5-17.3-63-31.2-10.1-7-16.9-17.8-18.8-29.9 0-18.3 31.6-41.7 77.2-57.6 5.7-2 11.5-3.8 17.3-5.5 6.8 21.7 15 43 24.5 63.6-9.6 20.9-17.9 42.5-24.8 64.5zm116.6 98c-16.5 15.1-35.6 27.1-56.4 35.3-11.1 5.3-23.9 5.8-35.3 1.3-15.9-9.2-22.5-44.5-13.5-92 1.1-5.6 2.3-11.2 3.7-16.7 22.4 4.8 45 8.1 67.9 9.8 13.2 18.7 27.7 36.6 43.2 53.4-3.2 3.1-6.4 6.1-9.6 8.9zm24.5-24.3c-10.2-11-20.4-23.2-30.3-36.3 9.6.4 19.5.6 29.5.6 10.3 0 20.4-.2 30.4-.7-9.2 12.7-19.1 24.8-29.6 36.4zm130.7 30c-.9 12.2-6.9 23.6-16.5 31.3-15.9 9.2-49.8-2.8-86.4-34.2-4.2-3.6-8.4-7.5-12.7-11.5 15.3-16.9 29.4-34.8 42.2-53.6 22.9-1.9 45.7-5.4 68.2-10.5 1 4.1 1.9 8.2 2.7 12.2 4.9 21.6 5.7 44.1 2.5 66.3zm18.2-107.5c-2.8.9-5.6 1.8-8.5 2.6-7-21.8-15.6-43.1-25.5-63.8 9.6-20.4 17.7-41.4 24.5-62.9 5.2 1.5 10.2 3.1 15 4.7 46.6 16 79.3 39.8 79.3 58 0 19.6-34.9 44.9-84.8 61.4zM256 210.2c25.3 0 45.8 20.5 45.8 45.8 0 25.3-20.5 45.8-45.8 45.8-25.3 0-45.8-20.5-45.8-45.8 0-25.3 20.5-45.8 45.8-45.8"],readme:[576,512,[],"f4d5","M528.3 46.5H388.5c-48.1 0-89.9 33.3-100.4 80.3-10.6-47-52.3-80.3-100.4-80.3H48c-26.5 0-48 21.5-48 48v245.8c0 26.5 21.5 48 48 48h89.7c102.2 0 132.7 24.4 147.3 75 .7 2.8 5.2 2.8 6 0 14.7-50.6 45.2-75 147.3-75H528c26.5 0 48-21.5 48-48V94.6c0-26.4-21.3-47.9-47.7-48.1zM242 311.9c0 1.9-1.5 3.5-3.5 3.5H78.2c-1.9 0-3.5-1.5-3.5-3.5V289c0-1.9 1.5-3.5 3.5-3.5h160.4c1.9 0 3.5 1.5 3.5 3.5v22.9zm0-60.9c0 1.9-1.5 3.5-3.5 3.5H78.2c-1.9 0-3.5-1.5-3.5-3.5v-22.9c0-1.9 1.5-3.5 3.5-3.5h160.4c1.9 0 3.5 1.5 3.5 3.5V251zm0-60.9c0 1.9-1.5 3.5-3.5 3.5H78.2c-1.9 0-3.5-1.5-3.5-3.5v-22.9c0-1.9 1.5-3.5 3.5-3.5h160.4c1.9 0 3.5 1.5 3.5 3.5v22.9zm259.3 121.7c0 1.9-1.5 3.5-3.5 3.5H337.5c-1.9 0-3.5-1.5-3.5-3.5v-22.9c0-1.9 1.5-3.5 3.5-3.5h160.4c1.9 0 3.5 1.5 3.5 3.5v22.9zm0-60.9c0 1.9-1.5 3.5-3.5 3.5H337.5c-1.9 0-3.5-1.5-3.5-3.5V228c0-1.9 1.5-3.5 3.5-3.5h160.4c1.9 0 3.5 1.5 3.5 3.5v22.9zm0-60.9c0 1.9-1.5 3.5-3.5 3.5H337.5c-1.9 0-3.5-1.5-3.5-3.5v-22.8c0-1.9 1.5-3.5 3.5-3.5h160.4c1.9 0 3.5 1.5 3.5 3.5V190z"],rebel:[512,512,[],"f1d0","M256.5 504C117.2 504 9 387.8 13.2 249.9 16 170.7 56.4 97.7 129.7 49.5c.3 0 1.9-.6 1.1.8-5.8 5.5-111.3 129.8-14.1 226.4 49.8 49.5 90 2.5 90 2.5 38.5-50.1-.6-125.9-.6-125.9-10-24.9-45.7-40.1-45.7-40.1l28.8-31.8c24.4 10.5 43.2 38.7 43.2 38.7.8-29.6-21.9-61.4-21.9-61.4L255.1 8l44.3 50.1c-20.5 28.8-21.9 62.6-21.9 62.6 13.8-23 43.5-39.3 43.5-39.3l28.5 31.8c-27.4 8.9-45.4 39.9-45.4 39.9-15.8 28.5-27.1 89.4.6 127.3 32.4 44.6 87.7-2.8 87.7-2.8 102.7-91.9-10.5-225-10.5-225-6.1-5.5.8-2.8.8-2.8 50.1 36.5 114.6 84.4 116.2 204.8C500.9 400.2 399 504 256.5 504z"],"red-river":[448,512,[],"f3e3","M353.2 32H94.8C42.4 32 0 74.4 0 126.8v258.4C0 437.6 42.4 480 94.8 480h258.4c52.4 0 94.8-42.4 94.8-94.8V126.8c0-52.4-42.4-94.8-94.8-94.8zM144.9 200.9v56.3c0 27-21.9 48.9-48.9 48.9V151.9c0-13.2 10.7-23.9 23.9-23.9h154.2c0 27-21.9 48.9-48.9 48.9h-56.3c-12.3-.6-24.6 11.6-24 24zm176.3 72h-56.3c-12.3-.6-24.6 11.6-24 24v56.3c0 27-21.9 48.9-48.9 48.9V247.9c0-13.2 10.7-23.9 23.9-23.9h154.2c0 27-21.9 48.9-48.9 48.9z"],reddit:[512,512,[],"f1a1","M201.5 305.5c-13.8 0-24.9-11.1-24.9-24.6 0-13.8 11.1-24.9 24.9-24.9 13.6 0 24.6 11.1 24.6 24.9 0 13.6-11.1 24.6-24.6 24.6zM504 256c0 137-111 248-248 248S8 393 8 256 119 8 256 8s248 111 248 248zm-132.3-41.2c-9.4 0-17.7 3.9-23.8 10-22.4-15.5-52.6-25.5-86.1-26.6l17.4-78.3 55.4 12.5c0 13.6 11.1 24.6 24.6 24.6 13.8 0 24.9-11.3 24.9-24.9s-11.1-24.9-24.9-24.9c-9.7 0-18 5.8-22.1 13.8l-61.2-13.6c-3-.8-6.1 1.4-6.9 4.4l-19.1 86.4c-33.2 1.4-63.1 11.3-85.5 26.8-6.1-6.4-14.7-10.2-24.1-10.2-34.9 0-46.3 46.9-14.4 62.8-1.1 5-1.7 10.2-1.7 15.5 0 52.6 59.2 95.2 132 95.2 73.1 0 132.3-42.6 132.3-95.2 0-5.3-.6-10.8-1.9-15.8 31.3-16 19.8-62.5-14.9-62.5zM302.8 331c-18.2 18.2-76.1 17.9-93.6 0-2.2-2.2-6.1-2.2-8.3 0-2.5 2.5-2.5 6.4 0 8.6 22.8 22.8 87.3 22.8 110.2 0 2.5-2.2 2.5-6.1 0-8.6-2.2-2.2-6.1-2.2-8.3 0zm7.7-75c-13.6 0-24.6 11.1-24.6 24.9 0 13.6 11.1 24.6 24.6 24.6 13.8 0 24.9-11.1 24.9-24.6 0-13.8-11-24.9-24.9-24.9z"],"reddit-alien":[512,512,[],"f281","M440.3 203.5c-15 0-28.2 6.2-37.9 15.9-35.7-24.7-83.8-40.6-137.1-42.3L293 52.3l88.2 19.8c0 21.6 17.6 39.2 39.2 39.2 22 0 39.7-18.1 39.7-39.7s-17.6-39.7-39.7-39.7c-15.4 0-28.7 9.3-35.3 22l-97.4-21.6c-4.9-1.3-9.7 2.2-11 7.1L246.3 177c-52.9 2.2-100.5 18.1-136.3 42.8-9.7-10.1-23.4-16.3-38.4-16.3-55.6 0-73.8 74.6-22.9 100.1-1.8 7.9-2.6 16.3-2.6 24.7 0 83.8 94.4 151.7 210.3 151.7 116.4 0 210.8-67.9 210.8-151.7 0-8.4-.9-17.2-3.1-25.1 49.9-25.6 31.5-99.7-23.8-99.7zM129.4 308.9c0-22 17.6-39.7 39.7-39.7 21.6 0 39.2 17.6 39.2 39.7 0 21.6-17.6 39.2-39.2 39.2-22 .1-39.7-17.6-39.7-39.2zm214.3 93.5c-36.4 36.4-139.1 36.4-175.5 0-4-3.5-4-9.7 0-13.7 3.5-3.5 9.7-3.5 13.2 0 27.8 28.5 120 29 149 0 3.5-3.5 9.7-3.5 13.2 0 4.1 4 4.1 10.2.1 13.7zm-.8-54.2c-21.6 0-39.2-17.6-39.2-39.2 0-22 17.6-39.7 39.2-39.7 22 0 39.7 17.6 39.7 39.7-.1 21.5-17.7 39.2-39.7 39.2z"],"reddit-square":[448,512,[],"f1a2","M283.2 345.5c2.7 2.7 2.7 6.8 0 9.2-24.5 24.5-93.8 24.6-118.4 0-2.7-2.4-2.7-6.5 0-9.2 2.4-2.4 6.5-2.4 8.9 0 18.7 19.2 81 19.6 100.5 0 2.4-2.3 6.6-2.3 9 0zm-91.3-53.8c0-14.9-11.9-26.8-26.5-26.8-14.9 0-26.8 11.9-26.8 26.8 0 14.6 11.9 26.5 26.8 26.5 14.6 0 26.5-11.9 26.5-26.5zm90.7-26.8c-14.6 0-26.5 11.9-26.5 26.8 0 14.6 11.9 26.5 26.5 26.5 14.9 0 26.8-11.9 26.8-26.5 0-14.9-11.9-26.8-26.8-26.8zM448 80v352c0 26.5-21.5 48-48 48H48c-26.5 0-48-21.5-48-48V80c0-26.5 21.5-48 48-48h352c26.5 0 48 21.5 48 48zm-99.7 140.6c-10.1 0-19 4.2-25.6 10.7-24.1-16.7-56.5-27.4-92.5-28.6l18.7-84.2 59.5 13.4c0 14.6 11.9 26.5 26.5 26.5 14.9 0 26.8-12.2 26.8-26.8 0-14.6-11.9-26.8-26.8-26.8-10.4 0-19.3 6.2-23.8 14.9l-65.7-14.6c-3.3-.9-6.5 1.5-7.4 4.8l-20.5 92.8c-35.7 1.5-67.8 12.2-91.9 28.9-6.5-6.8-15.8-11-25.9-11-37.5 0-49.8 50.4-15.5 67.5-1.2 5.4-1.8 11-1.8 16.7 0 56.5 63.7 102.3 141.9 102.3 78.5 0 142.2-45.8 142.2-102.3 0-5.7-.6-11.6-2.1-17 33.6-17.2 21.2-67.2-16.1-67.2z"],rendact:[496,512,[],"f3e4","M248 8C111 8 0 119 0 256s111 248 248 248c18.6 0 36.7-2.1 54.1-5.9-5.6-7.4-10.8-14.4-15.9-21.3-12.4 2.1-25.2 3.3-38.3 3.3C124.3 480 24 379.7 24 256S124.3 32 248 32s224 100.3 224 224c0 71-33 134.2-84.5 175.3-25.9 18.8-39.1 21.4-83.5-44.2-78.7-112.9-48-71.1-73.7-108.3 72.8 8.9 228.5-72 168.6-168.6C314-26.8 15 93.8 59.7 226.4c3.2 9.8 14.4 38.6 45.6 38.6 2 0 2.6-.6 2-1.7-4.4-8.7-20.1-9.8-20.1-37.4 0-40.5 40.5-89.6 100.3-120 66.1-32.3 131.9-30.2 158.2 5.4 27.2 38.3-20.9 119.2-120.4 136.9 7.5-9.4 57-75.2 62.8-84 22.7-34.6 23.6-49 14-59.2-15.5-16.9-29.5-10.3-50.7-11.7-10.8-.9-113.7 181.2-136.4 216.9-5.9 9-21.2 34.1-21.2 50.9 0 21.3 2.8 51.4 20.6 51.4 10.6 0 8-18.7 8-26.6 0-12.9 27.4-49.4 74.8-104.6 20.4 36.1 57.7 114.3 130.2 209.7 98-33.1 168.5-125.8 168.5-235C496 119 385 8 248 8z"],renren:[512,512,[],"f18b","M214 169.1c0 110.4-61 205.4-147.6 247.4C30 373.2 8 317.7 8 256.6 8 133.9 97.1 32.2 214 12.5v156.6zM255 504c-42.9 0-83.3-11-118.5-30.4C193.7 437.5 239.9 382.9 255 319c15.5 63.9 61.7 118.5 118.8 154.7C338.7 493 298.3 504 255 504zm190.6-87.5C359 374.5 298 279.6 298 169.1V12.5c116.9 19.7 206 121.4 206 244.1 0 61.1-22 116.6-58.4 159.9z"],replyd:[448,512,[],"f3e6","M320 480H128C57.6 480 0 422.4 0 352V160C0 89.6 57.6 32 128 32h192c70.4 0 128 57.6 128 128v192c0 70.4-57.6 128-128 128zM193.4 273.2c-6.1-2-11.6-3.1-16.4-3.1-7.2 0-13.5 1.9-18.9 5.6-5.4 3.7-9.6 9-12.8 15.8h-1.1l-4.2-18.3h-28v138.9h36.1v-89.7c1.5-5.4 4.4-9.8 8.7-13.2 4.3-3.4 9.8-5.1 16.2-5.1 4.6 0 9.8 1 15.6 3.1l4.8-34zm115.2 103.4c-3.2 2.4-7.7 4.8-13.7 7.1-6 2.3-12.8 3.5-20.4 3.5-12.2 0-21.1-3-26.5-8.9-5.5-5.9-8.5-14.7-9-26.4h83.3c.9-4.8 1.6-9.4 2.1-13.9.5-4.4.7-8.6.7-12.5 0-10.7-1.6-19.7-4.7-26.9-3.2-7.2-7.3-13-12.5-17.2-5.2-4.3-11.1-7.3-17.8-9.2-6.7-1.8-13.5-2.8-20.6-2.8-21.1 0-37.5 6.1-49.2 18.3s-17.5 30.5-17.5 55c0 22.8 5.2 40.7 15.6 53.7 10.4 13.1 26.8 19.6 49.2 19.6 10.7 0 20.9-1.5 30.4-4.6 9.5-3.1 17.1-6.8 22.6-11.2l-12-23.6zm-21.8-70.3c3.8 5.4 5.3 13.1 4.6 23.1h-51.7c.9-9.4 3.7-17 8.2-22.6 4.5-5.6 11.5-8.5 21-8.5 8.2-.1 14.1 2.6 17.9 8zm79.9 2.5c4.1 3.9 9.4 5.8 16.1 5.8 7 0 12.6-1.9 16.7-5.8s6.1-9.1 6.1-15.6-2-11.6-6.1-15.4c-4.1-3.8-9.6-5.7-16.7-5.7-6.7 0-12 1.9-16.1 5.7-4.1 3.8-6.1 8.9-6.1 15.4s2 11.7 6.1 15.6zm0 100.5c4.1 3.9 9.4 5.8 16.1 5.8 7 0 12.6-1.9 16.7-5.8s6.1-9.1 6.1-15.6-2-11.6-6.1-15.4c-4.1-3.8-9.6-5.7-16.7-5.7-6.7 0-12 1.9-16.1 5.7-4.1 3.8-6.1 8.9-6.1 15.4 0 6.6 2 11.7 6.1 15.6z"],researchgate:[448,512,[],"f4f8","M0 32v448h448V32H0zm262.2 334.4c-6.6 3-33.2 6-50-14.2-9.2-10.6-25.3-33.3-42.2-63.6-8.9 0-14.7 0-21.4-.6v46.4c0 23.5 6 21.2 25.8 23.9v8.1c-6.9-.3-23.1-.8-35.6-.8-13.1 0-26.1.6-33.6.8v-8.1c15.5-2.9 22-1.3 22-23.9V225c0-22.6-6.4-21-22-23.9V193c25.8 1 53.1-.6 70.9-.6 31.7 0 55.9 14.4 55.9 45.6 0 21.1-16.7 42.2-39.2 47.5 13.6 24.2 30 45.6 42.2 58.9 7.2 7.8 17.2 14.7 27.2 14.7v7.3zm22.9-135c-23.3 0-32.2-15.7-32.2-32.2V167c0-12.2 8.8-30.4 34-30.4s30.4 17.9 30.4 17.9l-10.7 7.2s-5.5-12.5-19.7-12.5c-7.9 0-19.7 7.3-19.7 19.7v26.8c0 13.4 6.6 23.3 17.9 23.3 14.1 0 21.5-10.9 21.5-26.8h-17.9v-10.7h30.4c0 20.5 4.7 49.9-34 49.9zm-116.5 44.7c-9.4 0-13.6-.3-20-.8v-69.7c6.4-.6 15-.6 22.5-.6 23.3 0 37.2 12.2 37.2 34.5 0 21.9-15 36.6-39.7 36.6z"],resolving:[496,512,[],"f3e7","M281.2 278.2c46-13.3 49.6-23.5 44-43.4L314 195.5c-6.1-20.9-18.4-28.1-71.1-12.8L54.7 236.8l28.6 98.6 197.9-57.2zM248.5 8C131.4 8 33.2 88.7 7.2 197.5l221.9-63.9c34.8-10.2 54.2-11.7 79.3-8.2 36.3 6.1 52.7 25 61.4 55.2l10.7 37.8c8.2 28.1 1 50.6-23.5 73.6-19.4 17.4-31.2 24.5-61.4 33.2L203 351.8l220.4 27.1 9.7 34.2-48.1 13.3-286.8-37.3 23 80.2c36.8 22 80.3 34.7 126.3 34.7 137 0 248.5-111.4 248.5-248.3C497 119.4 385.5 8 248.5 8zM38.3 388.6L0 256.8c0 48.5 14.3 93.4 38.3 131.8z"],rocketchat:[448,512,[],"f3e8","M448 256.2c0-87.2-99.6-153.3-219.8-153.3-18.8 0-37.3 1.6-55.3 4.8-11.1-10.5-24.2-20-38-27.4C61.2 44.2 0 79.4 0 79.4s56.9 47.1 47.6 88.3c-52.3 52.3-52.5 124.1 0 176.6C56.9 385.6 0 432.6 0 432.6s61.2 35.2 134.9-.8c13.8-7.5 26.9-16.9 38-27.4 18 3.2 36.5 4.8 55.3 4.8 120.3-.1 219.8-65.8 219.8-153zm-219.7 124c-23.7 0-46.3-2.8-67.3-7.8-21.3 25.8-68.1 61.7-113.6 50.1 14.8-16 36.7-43.1 32-87.6-27.3-21.4-43.6-48.7-43.6-78.5 0-68.4 86.2-123.9 192.5-123.9S420.8 188 420.8 256.4c0 68.3-86.2 123.8-192.5 123.8zm25.6-123.9c0 14.2-11.5 25.8-25.6 25.8-14.1 0-25.6-11.5-25.6-25.8 0-14.2 11.5-25.8 25.6-25.8 14.1 0 25.6 11.6 25.6 25.8zm88.9 0c0 14.2-11.4 25.8-25.6 25.8-14.1 0-25.6-11.5-25.6-25.8 0-14.2 11.4-25.8 25.6-25.8 14.1 0 25.6 11.6 25.6 25.8zm-177.9 0c0 14.2-11.4 25.8-25.6 25.8-14.1 0-25.6-11.5-25.6-25.8 0-14.2 11.4-25.8 25.6-25.8 14.2 0 25.6 11.6 25.6 25.8z"],rockrms:[496,512,[],"f3e9","M248 8C111 8 0 119 0 256s111 248 248 248 248-111 248-248S385 8 248 8zm157.4 419.5h-90l-112-131.3c-17.9-20.4-3.9-56.1 26.6-56.1h75.3l-84.6-99.3-84.3 98.9h-90L193.5 67.2c14.4-18.4 41.3-17.3 54.5 0l157.7 185.1c19 22.8 2 57.2-27.6 56.1-.6 0-74.2.2-74.2.2l101.5 118.9z"],safari:[512,512,[],"f267","M236.9 256.8c0-9.1 6.6-17.7 16.3-17.7 8.9 0 17.4 6.4 17.4 16.1 0 9.1-6.4 17.7-16.1 17.7-9 0-17.6-6.7-17.6-16.1zM504 256c0 137-111 248-248 248S8 393 8 256 119 8 256 8s248 111 248 248zm-26.6 0c0-122.3-99.1-221.4-221.4-221.4S34.6 133.7 34.6 256 133.7 477.4 256 477.4 477.4 378.3 477.4 256zm-72.5 96.6c0 3.6 13 10.2 16.3 12.2-27.4 41.5-69.8 71.4-117.9 83.3l-4.4-18.5c-.3-2.5-1.9-2.8-4.2-2.8-1.9 0-3 2.8-2.8 4.2l4.4 18.8c-13.3 2.8-26.8 4.2-40.4 4.2-36.3 0-72-10.2-103-29.1 1.7-2.8 12.2-18 12.2-20.2 0-1.9-1.7-3.6-3.6-3.6-3.9 0-12.2 16.6-14.7 19.9-41.8-27.7-72-70.6-83.6-119.6l19.1-4.2c2.2-.6 2.8-2.2 2.8-4.2 0-1.9-2.8-3-4.4-2.8L62 294.5c-2.5-12.7-3.9-25.5-3.9-38.5 0-37.1 10.5-73.6 30.2-104.9 2.8 1.7 16.1 10.8 18.3 10.8 1.9 0 3.6-1.4 3.6-3.3 0-3.9-14.7-11.3-18-13.6 28.2-41.2 71.1-70.9 119.8-81.9l4.2 18.5c.6 2.2 2.2 2.8 4.2 2.8s3-2.8 2.8-4.4L219 61.7c12.2-2.2 24.6-3.6 37.1-3.6 37.1 0 73.3 10.5 104.9 30.2-1.9 2.8-10.8 15.8-10.8 18 0 1.9 1.4 3.6 3.3 3.6 3.9 0 11.3-14.4 13.3-17.7 41 27.7 70.3 70 81.7 118.2l-15.5 3.3c-2.5.6-2.8 2.2-2.8 4.4 0 1.9 2.8 3 4.2 2.8l15.8-3.6c2.5 12.7 3.9 25.7 3.9 38.7 0 36.3-10 72-28.8 102.7-2.8-1.4-14.4-9.7-16.6-9.7-2.1 0-3.8 1.7-3.8 3.6zm-33.2-242.2c-13 12.2-134.2 123.7-137.6 129.5l-96.6 160.5c12.7-11.9 134.2-124 137.3-129.3l96.9-160.7z"],sass:[640,512,[],"f41e","M551.1 291.9c-22.4.1-41.8 5.5-58 13.5-5.9-11.9-12-22.3-13-30.1-1.2-9.1-2.5-14.5-1.1-25.3s7.7-26.1 7.6-27.2c-.1-1.1-1.4-6.6-14.3-6.7-12.9-.1-24 2.5-25.3 5.9-1.3 3.4-3.8 11.1-5.3 19.1-2.3 11.7-25.8 53.5-39.1 75.3-4.4-8.5-8.1-16-8.9-22-1.2-9.1-2.5-14.5-1.1-25.3s7.7-26.1 7.6-27.2c-.1-1.1-1.4-6.6-14.3-6.7-12.9-.1-24 2.5-25.3 5.9-1.3 3.4-2.7 11.4-5.3 19.1-2.6 7.7-33.9 77.3-42.1 95.4-4.2 9.2-7.8 16.6-10.4 21.6s-.2.3-.4.9c-2.2 4.3-3.5 6.7-3.5 6.7v.1c-1.7 3.2-3.6 6.1-4.5 6.1-.6 0-1.9-8.4.3-19.9 4.7-24.2 15.8-61.8 15.7-63.1-.1-.7 2.1-7.2-7.3-10.7-9.1-3.3-12.4 2.2-13.2 2.2-.8 0-1.4 2-1.4 2s10.1-42.4-19.4-42.4c-18.4 0-44 20.2-56.6 38.5-7.9 4.3-25 13.6-43 23.5-6.9 3.8-14 7.7-20.7 11.4-.5-.5-.9-1-1.4-1.5-35.8-38.2-101.9-65.2-99.1-116.5 1-18.7 7.5-67.8 127.1-127.4 98-48.8 176.4-35.4 189.9-5.6 19.4 42.5-41.9 121.6-143.7 133-38.8 4.3-59.2-10.7-64.3-16.3-5.3-5.9-6.1-6.2-8.1-5.1-3.3 1.8-1.2 7 0 10.1 3 7.9 15.5 21.9 36.8 28.9 18.7 6.1 64.2 9.5 119.2-11.8C367 196.5 415.1 130.2 401 74.7 386.6 18.3 293.1-.2 204.6 31.2 151.9 49.9 94.9 79.3 53.9 117.6 5.2 163.2-2.6 202.9.6 219.5c11.4 58.9 92.6 97.3 125.1 125.7-1.6.9-3.1 1.7-4.5 2.5-16.3 8.1-78.2 40.5-93.7 74.7-17.5 38.8 2.9 66.6 16.3 70.4 41.8 11.6 84.6-9.3 107.6-43.6s20.2-79.1 9.6-99.5c-.1-.3-.3-.5-.4-.8 4.2-2.5 8.5-5 12.8-7.5 8.3-4.9 16.4-9.4 23.5-13.3-4 10.8-6.9 23.8-8.4 42.6-1.8 22 7.3 50.5 19.1 61.7 5.2 4.9 11.5 5 15.4 5 13.8 0 20-11.4 26.9-25 8.5-16.6 16-35.9 16-35.9s-9.4 52.2 16.3 52.2c9.4 0 18.8-12.1 23-18.3v.1s.2-.4.7-1.2c1-1.5 1.5-2.4 1.5-2.4v-.3c3.8-6.5 12.1-21.4 24.6-46 16.2-31.8 31.7-71.5 31.7-71.5s1.4 9.7 6.2 25.8c2.8 9.5 8.7 19.9 13.4 30-3.8 5.2-6.1 8.2-6.1 8.2s0 .1.1.2c-3 4-6.4 8.3-9.9 12.5-12.8 15.2-28 32.6-30 37.6-2.4 5.9-1.8 10.3 2.8 13.7 3.4 2.6 9.4 3 15.7 2.5 11.5-.8 19.6-3.6 23.5-5.4 6.2-2.2 13.4-5.7 20.2-10.6 12.5-9.2 20.1-22.4 19.4-39.8-.4-9.6-3.5-19.2-7.3-28.2 1.1-1.6 2.3-3.3 3.4-5 19.8-28.9 35.1-60.6 35.1-60.6s1.4 9.7 6.2 25.8c2.4 8.1 7.1 17 11.4 25.7-18.6 15.1-30.1 32.6-34.1 44.1-7.4 21.3-1.6 30.9 9.3 33.1 4.9 1 11.9-1.3 17.1-3.5 6.5-2.2 14.3-5.7 21.6-11.1 12.5-9.2 24.6-22.1 23.8-39.6-.3-7.9-2.5-15.8-5.4-23.4 15.7-6.6 36.1-10.2 62.1-7.2 55.7 6.5 66.6 41.3 64.5 55.8-2.1 14.6-13.8 22.6-17.7 25-3.9 2.4-5.1 3.3-4.8 5.1.5 2.6 2.3 2.5 5.6 1.9 4.6-.8 29.2-11.8 30.3-38.7 1.6-34-31.1-71.4-89-71.1zM121.8 436.6c-18.4 20.1-44.2 27.7-55.3 21.3C54.6 451 59.3 421.4 82 400c13.8-13 31.6-25 43.4-32.4 2.7-1.6 6.6-4 11.4-6.9.8-.5 1.2-.7 1.2-.7.9-.6 1.9-1.1 2.9-1.7 8.3 30.4.3 57.2-19.1 78.3zm134.4-91.4c-6.4 15.7-19.9 55.7-28.1 53.6-7-1.8-11.3-32.3-1.4-62.3 5-15.1 15.6-33.1 21.9-40.1 10.1-11.3 21.2-14.9 23.8-10.4 3.5 5.9-12.2 49.4-16.2 59.2zm111 53c-2.7 1.4-5.2 2.3-6.4 1.6-.9-.5 1.1-2.4 1.1-2.4s13.9-14.9 19.4-21.7c3.2-4 6.9-8.7 10.9-13.9 0 .5.1 1 .1 1.6-.1 17.9-17.3 30-25.1 34.8zm85.6-19.5c-2-1.4-1.7-6.1 5-20.7 2.6-5.7 8.6-15.3 19-24.5 1.2 3.8 1.9 7.4 1.9 10.8-.1 22.5-16.2 30.9-25.9 34.4z"],schlix:[448,512,[],"f3ea","M350.5 157.7l-54.2-46.1 73.4-39 78.3 44.2-97.5 40.9zM192 122.1l45.7-28.2 34.7 34.6-55.4 29-25-35.4zm-65.1 6.6l31.9-22.1L176 135l-36.7 22.5-12.4-28.8zm-23.3 88.2l-8.8-34.8 29.6-18.3 13.1 35.3-33.9 17.8zm-21.2-83.7l23.9-18.1 8.9 24-26.7 18.3-6.1-24.2zM59 206.5l-3.6-28.4 22.3-15.5 6.1 28.7L59 206.5zm-30.6 16.6l20.8-12.8 3.3 33.4-22.9 12-1.2-32.6zM1.4 268l19.2-10.2.4 38.2-21 8.8L1.4 268zm59.1 59.3l-28.3 8.3-1.6-46.8 25.1-10.7 4.8 49.2zM99 263.2l-31.1 13-5.2-40.8L90.1 221l8.9 42.2zM123.2 377l-41.6 5.9-8.1-63.5 35.2-10.8 14.5 68.4zm28.5-139.9l21.2 57.1-46.2 13.6-13.7-54.1 38.7-16.6zm85.7 230.5l-70.9-3.3-24.3-95.8 55.2-8.6 40 107.7zm-84.9-279.7l42.2-22.4 28 45.9-50.8 21.3-19.4-44.8zm41 94.9l61.3-18.7 52.8 86.6-79.8 11.3-34.3-79.2zm51.4-85.6l67.3-28.8 65.5 65.4-88.6 26.2-44.2-62.8z"],scribd:[384,512,[],"f28a","M42.3 252.7c-16.1-19-24.7-45.9-24.8-79.9 0-100.4 75.2-153.1 167.2-153.1 98.6-1.6 156.8 49 184.3 70.6l-50.5 72.1-37.3-24.6 26.9-38.6c-36.5-24-79.4-36.5-123-35.8-50.7-.8-111.7 27.2-111.7 76.2 0 18.7 11.2 20.7 28.6 15.6 23.3-5.3 41.9.6 55.8 14 26.4 24.3 23.2 67.6-.7 91.9-29.2 29.5-85.2 27.3-114.8-8.4zm317.7 5.9c-15.5-18.8-38.9-29.4-63.2-28.6-38.1-2-71.1 28-70.5 67.2-.7 16.8 6 33 18.4 44.3 14.1 13.9 33 19.7 56.3 14.4 17.4-5.1 28.6-3.1 28.6 15.6 0 4.3-.5 8.5-1.4 12.7-16.7 40.9-59.5 64.4-121.4 64.4-51.9.2-102.4-16.4-144.1-47.3l33.7-39.4-35.6-27.4L0 406.3l15.4 13.8c52.5 46.8 120.4 72.5 190.7 72.2 51.4 0 94.4-10.5 133.6-44.1 57.1-51.4 54.2-149.2 20.3-189.6z"],searchengin:[460,512,[],"f3eb","M220.6 130.3l-67.2 28.2V43.2L98.7 233.5l54.7-24.2v130.3l67.2-209.3zm-83.2-96.7l-1.3 4.7-15.2 52.9C80.6 106.7 52 145.8 52 191.5c0 52.3 34.3 95.9 83.4 105.5v53.6C57.5 340.1 0 272.4 0 191.6c0-80.5 59.8-147.2 137.4-158zm311.4 447.2c-11.2 11.2-23.1 12.3-28.6 10.5-5.4-1.8-27.1-19.9-60.4-44.4-33.3-24.6-33.6-35.7-43-56.7-9.4-20.9-30.4-42.6-57.5-52.4l-9.7-14.7c-24.7 16.9-53 26.9-81.3 28.7l2.1-6.6 15.9-49.5c46.5-11.9 80.9-54 80.9-104.2 0-54.5-38.4-102.1-96-107.1V32.3C254.4 37.4 320 106.8 320 191.6c0 33.6-11.2 64.7-29 90.4l14.6 9.6c9.8 27.1 31.5 48 52.4 57.4s32.2 9.7 56.8 43c24.6 33.2 42.7 54.9 44.5 60.3s.7 17.3-10.5 28.5zm-9.9-17.9c0-4.4-3.6-8-8-8s-8 3.6-8 8 3.6 8 8 8 8-3.6 8-8z"],sellcast:[448,512,[],"f2da","M353.4 32H94.7C42.6 32 0 74.6 0 126.6v258.7C0 437.4 42.6 480 94.7 480h258.7c52.1 0 94.7-42.6 94.7-94.6V126.6c0-52-42.6-94.6-94.7-94.6zm-50 316.4c-27.9 48.2-89.9 64.9-138.2 37.2-22.9 39.8-54.9 8.6-42.3-13.2l15.7-27.2c5.9-10.3 19.2-13.9 29.5-7.9 18.6 10.8-.1-.1 18.5 10.7 27.6 15.9 63.4 6.3 79.4-21.3 15.9-27.6 6.3-63.4-21.3-79.4-17.8-10.2-.6-.4-18.6-10.6-24.6-14.2-3.4-51.9 21.6-37.5 18.6 10.8-.1-.1 18.5 10.7 48.4 28 65.1 90.3 37.2 138.5zm21.8-208.8c-17 29.5-16.3 28.8-19 31.5-6.5 6.5-16.3 8.7-26.5 3.6-18.6-10.8.1.1-18.5-10.7-27.6-15.9-63.4-6.3-79.4 21.3s-6.3 63.4 21.3 79.4c0 0 18.5 10.6 18.6 10.6 24.6 14.2 3.4 51.9-21.6 37.5-18.6-10.8.1.1-18.5-10.7-48.2-27.8-64.9-90.1-37.1-138.4 27.9-48.2 89.9-64.9 138.2-37.2l4.8-8.4c14.3-24.9 52-3.3 37.7 21.5z"],sellsy:[640,512,[],"f213","M539.71 237.308c3.064-12.257 4.29-24.821 4.29-37.384C544 107.382 468.618 32 376.076 32c-77.22 0-144.634 53.012-163.02 127.781-15.322-13.176-34.934-20.53-55.157-20.53-46.271 0-83.962 37.69-83.962 83.961 0 7.354.92 15.015 3.065 22.369-42.9 20.225-70.785 63.738-70.785 111.234C6.216 424.843 61.68 480 129.401 480h381.198c67.72 0 123.184-55.157 123.184-123.184.001-56.384-38.916-106.025-94.073-119.508zM199.88 401.554c0 8.274-7.048 15.321-15.321 15.321H153.61c-8.274 0-15.321-7.048-15.321-15.321V290.626c0-8.273 7.048-15.321 15.321-15.321h30.949c8.274 0 15.321 7.048 15.321 15.321v110.928zm89.477 0c0 8.274-7.048 15.321-15.322 15.321h-30.949c-8.274 0-15.321-7.048-15.321-15.321V270.096c0-8.274 7.048-15.321 15.321-15.321h30.949c8.274 0 15.322 7.048 15.322 15.321v131.458zm89.477 0c0 8.274-7.047 15.321-15.321 15.321h-30.949c-8.274 0-15.322-7.048-15.322-15.321V238.84c0-8.274 7.048-15.321 15.322-15.321h30.949c8.274 0 15.321 7.048 15.321 15.321v162.714zm87.027 0c0 8.274-7.048 15.321-15.322 15.321h-28.497c-8.274 0-15.321-7.048-15.321-15.321V176.941c0-8.579 7.047-15.628 15.321-15.628h28.497c8.274 0 15.322 7.048 15.322 15.628v224.613z"],servicestack:[496,512,[],"f3ec","M88 216c81.7 10.2 273.7 102.3 304 232H0c99.5-8.1 184.5-137 88-232zm32-152c32.3 35.6 47.7 83.9 46.4 133.6C249.3 231.3 373.7 321.3 400 448h96C455.3 231.9 222.8 79.5 120 64z"],shirtsinbulk:[448,512,[],"f214","M395.208 221.583H406v33.542h-10.792v-33.542zm0-9.625H406v-33.542h-10.792v33.542zm0 86.333H406V264.75h-10.792v33.541zM358.75 135.25h-33.542v10.5h33.542v-10.5zm36.458 206.208H406v-33.542h-10.792v33.542zM311.5 135.25h-33.542v10.5H311.5v-10.5zm-47.25 0H231v10.5h33.25v-10.5zm-47.25 0h-33.25v10.5H217v-10.5zm178.208 33.542H406V135.25h-33.542v10.5h22.75v23.042zm-255.792 259l30.625 13.417 4.375-9.917-30.625-13.417-4.375 9.917zM179.083 445l30.334 13.708 4.374-9.916-30.333-13.417-4.375 9.625zm216.125-60.375H406v-33.542h-10.792v33.542zm-334.833 8.167L91 406.208l4.375-9.624-30.625-13.709-4.375 9.917zm39.666 17.499l30.625 13.417 4.375-9.917-30.625-13.416-4.375 9.916zm132.417 38.501l4.375 9.916L267.459 445l-4.375-9.625-30.626 13.417zm118.417-52.208l4.375 9.624 30.624-13.416-4.374-9.917-30.625 13.709zM311.5 413.791l4.375 9.917 30.625-13.417-4.374-9.916-30.626 13.416zm-39.667 17.501l4.375 9.917 30.625-13.417-4.375-9.917-30.625 13.417zM311.5 46.583h-33.542v10.5H311.5v-10.5zm94.209 0h-33.251v10.5h33.251v-10.5zm-188.709 0h-33.25v10.5H217v-10.5zm141.75 0h-33.542v10.5h33.542v-10.5zm-94.5 0H231v10.5h33.25v-10.5zM448 3.708v406l-226.334 98.584L0 409.708v-406h448zm-29.166 116.958H29.166V390.75l192.792 85.75 196.875-85.75V120.666zm0-87.791H29.166V91.5h389.667V32.875zM75.542 46.583H42.291v10.5h33.251v-10.5zm94.5 0H136.5v10.5h33.542v-10.5zm-47.251 0H89.25v10.5h33.542v-10.5zm7.584 236.542c0-50.167 41.125-91.292 91.292-91.292 50.458 0 91.292 41.125 91.292 91.292 0 50.458-40.833 91.292-91.292 91.292-50.167-.001-91.292-40.834-91.292-91.292zm120.75 18.084c0 13.125-23.917 14.291-32.666 14.291-12.25 0-29.75-2.625-35.875-14.875h-.875L172.666 319c14.876 9.333 29.167 12.25 47.25 12.25 19.542 0 51.042-5.833 51.042-31.209 0-48.125-78.458-16.333-78.458-37.916 0-13.125 20.708-14.875 29.75-14.875 10.791 0 29.166 3.208 35.583 13.124h.875l8.751-16.916c-15.167-6.125-27.417-11.959-44.334-11.959-20.125 0-49.583 6.417-49.583 31.792 0 44.334 77.583 11.959 77.583 37.918zM122.791 135.25H89.25v10.5h33.542v-10.5zm-69.999 10.5h22.75v-10.5H42v33.542h10.792V145.75zm0 32.666H42v33.542h10.792v-33.542zm117.25-43.166H136.5v10.5h33.542v-10.5zm-117.25 86.333H42v33.542h10.792v-33.542zm0 86.334H42v33.542h10.792v-33.542zm0-43.167H42v33.542h10.792V264.75zm0 86.333H42v33.542h10.792v-33.542z"],simplybuilt:[512,512,[],"f215","M481.2 64h-106c-14.5 0-26.6 11.8-26.6 26.3v39.6H163.3V90.3c0-14.5-12-26.3-26.6-26.3h-106C16.1 64 4.3 75.8 4.3 90.3v331.4c0 14.5 11.8 26.3 26.6 26.3h450.4c14.8 0 26.6-11.8 26.6-26.3V90.3c-.2-14.5-12-26.3-26.7-26.3zM149.8 355.8c-36.6 0-66.4-29.7-66.4-66.4 0-36.9 29.7-66.6 66.4-66.6 36.9 0 66.6 29.7 66.6 66.6 0 36.7-29.7 66.4-66.6 66.4zm212.4 0c-36.9 0-66.6-29.7-66.6-66.6 0-36.6 29.7-66.4 66.6-66.4 36.6 0 66.4 29.7 66.4 66.4 0 36.9-29.8 66.6-66.4 66.6z"],sistrix:[448,512,[],"f3ee","M448 449L301.2 300.2c20-27.9 31.9-62.2 31.9-99.2 0-93.1-74.7-168.9-166.5-168.9C74.7 32 0 107.8 0 200.9s74.7 168.9 166.5 168.9c39.8 0 76.3-14.2 105-37.9l146 148.1 30.5-31zM166.5 330.8c-70.6 0-128.1-58.3-128.1-129.9S95.9 71 166.5 71s128.1 58.3 128.1 129.9-57.4 129.9-128.1 129.9z"],sith:[448,512,[],"f512","M0 32l69.71 118.75-58.86-11.52 69.84 91.03a146.741 146.741 0 0 0 0 51.45l-69.84 91.03 58.86-11.52L0 480l118.75-69.71-11.52 58.86 91.03-69.84c17.02 3.04 34.47 3.04 51.48 0l91.03 69.84-11.52-58.86L448 480l-69.71-118.78 58.86 11.52-69.84-91.03c3.03-17.01 3.04-34.44 0-51.45l69.84-91.03-58.86 11.52L448 32l-118.75 69.71 11.52-58.9-91.06 69.87c-8.5-1.52-17.1-2.29-25.71-2.29s-17.21.78-25.71 2.29l-91.06-69.87 11.52 58.9L0 32zm224 99.78c31.8 0 63.6 12.12 87.85 36.37 48.5 48.5 48.49 127.21 0 175.7s-127.2 48.46-175.7-.03c-48.5-48.5-48.49-127.21 0-175.7 24.24-24.25 56.05-36.34 87.85-36.34zm0 36.66c-22.42 0-44.83 8.52-61.92 25.61-34.18 34.18-34.19 89.68 0 123.87s89.65 34.18 123.84 0c34.18-34.18 34.19-89.68 0-123.87-17.09-17.09-39.5-25.61-61.92-25.61z"],skyatlas:[640,512,[],"f216","M640 329.3c0 65.9-52.5 114.4-117.5 114.4-165.9 0-196.6-249.7-359.7-249.7-146.9 0-147.1 212.2 5.6 212.2 42.5 0 90.9-17.8 125.3-42.5 5.6-4.1 16.9-16.3 22.8-16.3s10.9 5 10.9 10.9c0 7.8-13.1 19.1-18.7 24.1-40.9 35.6-100.3 61.2-154.7 61.2-83.4.1-154-59-154-144.9s67.5-149.1 152.8-149.1c185.3 0 222.5 245.9 361.9 245.9 99.9 0 94.8-139.7 3.4-139.7-17.5 0-35 11.6-46.9 11.6-8.4 0-15.9-7.2-15.9-15.6 0-11.6 5.3-23.7 5.3-36.3 0-66.6-50.9-114.7-116.9-114.7-53.1 0-80 36.9-88.8 36.9-6.2 0-11.2-5-11.2-11.2 0-5.6 4.1-10.3 7.8-14.4 25.3-28.8 64.7-43.7 102.8-43.7 79.4 0 139.1 58.4 139.1 137.8 0 6.9-.3 13.7-1.2 20.6 11.9-3.1 24.1-4.7 35.9-4.7 60.7 0 111.9 45.3 111.9 107.2z"],skype:[448,512,[],"f17e","M424.7 299.8c2.9-14 4.7-28.9 4.7-43.8 0-113.5-91.9-205.3-205.3-205.3-14.9 0-29.7 1.7-43.8 4.7C161.3 40.7 137.7 32 112 32 50.2 32 0 82.2 0 144c0 25.7 8.7 49.3 23.3 68.2-2.9 14-4.7 28.9-4.7 43.8 0 113.5 91.9 205.3 205.3 205.3 14.9 0 29.7-1.7 43.8-4.7 19 14.6 42.6 23.3 68.2 23.3 61.8 0 112-50.2 112-112 .1-25.6-8.6-49.2-23.2-68.1zm-194.6 91.5c-65.6 0-120.5-29.2-120.5-65 0-16 9-30.6 29.5-30.6 31.2 0 34.1 44.9 88.1 44.9 25.7 0 42.3-11.4 42.3-26.3 0-18.7-16-21.6-42-28-62.5-15.4-117.8-22-117.8-87.2 0-59.2 58.6-81.1 109.1-81.1 55.1 0 110.8 21.9 110.8 55.4 0 16.9-11.4 31.8-30.3 31.8-28.3 0-29.2-33.5-75-33.5-25.7 0-42 7-42 22.5 0 19.8 20.8 21.8 69.1 33 41.4 9.3 90.7 26.8 90.7 77.6 0 59.1-57.1 86.5-112 86.5z"],slack:[448,512,[],"f198","M244.2 217.5l19.3 57.7-59.8 20-19.3-57.7 59.8-20zm41.4 243.7C131.6 507.4 65 471.6 18.8 317.6S8.4 97 162.4 50.8C316.4 4.6 383 40.4 429.2 194.4c46.2 154 10.4 220.6-143.6 266.8zM366.2 265c-3.9-12.2-17.2-18.6-29.4-14.7l-29 9.7-19.3-57.7 29-9.7c12.2-3.9 18.6-17.2 14.7-29.4-3.9-12.2-17.2-18.6-29.4-14.7l-29 9.7-10-30.1c-3.9-12.2-17.2-18.6-29.4-14.7-12.2 3.9-18.6 17.2-14.7 29.4l10 30.1-59.8 20.1-10-30.1c-3.9-12.2-17.2-18.6-29.4-14.7-12.2 3.9-18.6 17.2-14.7 29.4l10 30.1-29 9.7c-12.2 3.9-18.6 17.2-14.7 29.4 3.2 9.3 12.2 15.4 21.5 15.8 4.3.6 7.7-1 36.9-10.7l19.3 57.7-29 9.7c-12.2 3.9-18.6 17.2-14.7 29.4 3.2 9.3 12.2 15.4 21.5 15.8 4.3.6 7.7-1 36.9-10.7l10 30.1c3.7 10.8 15.8 18.6 29.4 14.7 12.2-3.9 18.6-17.2 14.7-29.4l-10-30.1 59.8-20.1 10 30.1c3.7 10.8 15.8 18.6 29.4 14.7 12.2-3.9 18.6-17.2 14.7-29.4l-10-30.1 29-9.7c12.2-4.2 18.6-17.5 14.7-29.6z"],"slack-hash":[448,512,[],"f3ef","M446.2 270.4c-6.2-19-26.9-29.1-46-22.9l-45.4 15.1-30.3-90 45.4-15.1c19.1-6.2 29.1-26.8 23-45.9-6.2-19-26.9-29.1-46-22.9l-45.4 15.1-15.7-47c-6.2-19-26.9-29.1-46-22.9-19.1 6.2-29.1 26.8-23 45.9l15.7 47-93.4 31.2-15.7-47c-6.2-19-26.9-29.1-46-22.9-19.1 6.2-29.1 26.8-23 45.9l15.7 47-45.3 15c-19.1 6.2-29.1 26.8-23 45.9 5 14.5 19.1 24 33.6 24.6 6.8 1 12-1.6 57.7-16.8l30.3 90L78 354.8c-19 6.2-29.1 26.9-23 45.9 5 14.5 19.1 24 33.6 24.6 6.8 1 12-1.6 57.7-16.8l15.7 47c5.9 16.9 24.7 29 46 22.9 19.1-6.2 29.1-26.8 23-45.9l-15.7-47 93.6-31.3 15.7 47c5.9 16.9 24.7 29 46 22.9 19.1-6.2 29.1-26.8 23-45.9l-15.7-47 45.4-15.1c19-6 29.1-26.7 22.9-45.7zm-254.1 47.2l-30.3-90.2 93.5-31.3 30.3 90.2-93.5 31.3z"],slideshare:[512,512,[],"f1e7","M249.429 211.436c0 31.716-27.715 57.717-61.717 57.717-34.001 0-61.716-26.001-61.716-57.717 0-32.001 27.715-57.716 61.716-57.716 34.001 0 61.717 25.715 61.717 57.716zm254.294 50.002c-18.286 22.573-53.144 50.288-106.289 72.003C453.722 525.163 260 555.735 263.143 457.446c0 1.714-.286-52.859-.286-93.432-4.285-.858-8.571-2-13.714-3.143 0 40.858-.286 98.289-.286 96.575C252 555.735 58.278 525.163 114.566 333.441c-53.145-21.715-88.003-49.43-106.29-72.003-9.143-13.714.858-28.287 16.001-17.715 2 1.428 4.285 2.857 6.285 4.285V49.716C30.563 22.287 51.135 0 76.565 0h359.157c25.429 0 46.002 22.287 46.002 49.716v198.293l6-4.285c15.143-10.573 25.143 4 15.999 17.714zm-46.572-189.15c0-32.858-10.572-45.716-40.859-45.716H98.566c-31.716 0-40.573 10.858-40.573 45.716v192.293c67.717 35.43 125.72 29.144 157.435 28.001 13.429-.286 22.001 2.286 27.144 7.715 1.689 1.687 10.023 9.446 20.287 17.143 1.143-15.715 10.001-25.715 33.716-24.858 32.287 1.428 91.718 7.715 160.577-29.716V72.288zM331.146 153.72c-34.002 0-61.716 25.715-61.716 57.716 0 31.716 27.715 57.717 61.716 57.717 34.287 0 61.716-26.001 61.716-57.717 0-32.001-27.429-57.716-61.716-57.716z"],snapchat:[496,512,[],"f2ab","M248 8C111 8 0 119 0 256s111 248 248 248 248-111 248-248S385 8 248 8zm169.5 338.9c-3.5 8.1-18.1 14-44.8 18.2-1.4 1.9-2.5 9.8-4.3 15.9-1.1 3.7-3.7 5.9-8.1 5.9h-.2c-6.2 0-12.8-2.9-25.8-2.9-17.6 0-23.7 4-37.4 13.7-14.5 10.3-28.4 19.1-49.2 18.2-21 1.6-38.6-11.2-48.5-18.2-13.8-9.7-19.8-13.7-37.4-13.7-12.5 0-20.4 3.1-25.8 3.1-5.4 0-7.5-3.3-8.3-6-1.8-6.1-2.9-14.1-4.3-16-13.8-2.1-44.8-7.5-45.5-21.4-.2-3.6 2.3-6.8 5.9-7.4 46.3-7.6 67.1-55.1 68-57.1 0-.1.1-.2.2-.3 2.5-5 3-9.2 1.6-12.5-3.4-7.9-17.9-10.7-24-13.2-15.8-6.2-18-13.4-17-18.3 1.6-8.5 14.4-13.8 21.9-10.3 5.9 2.8 11.2 4.2 15.7 4.2 3.3 0 5.5-.8 6.6-1.4-1.4-23.9-4.7-58 3.8-77.1C183.1 100 230.7 96 244.7 96c.6 0 6.1-.1 6.7-.1 34.7 0 68 17.8 84.3 54.3 8.5 19.1 5.2 53.1 3.8 77.1 1.1.6 2.9 1.3 5.7 1.4 4.3-.2 9.2-1.6 14.7-4.2 4-1.9 9.6-1.6 13.6 0 6.3 2.3 10.3 6.8 10.4 11.9.1 6.5-5.7 12.1-17.2 16.6-1.4.6-3.1 1.1-4.9 1.7-6.5 2.1-16.4 5.2-19 11.5-1.4 3.3-.8 7.5 1.6 12.5.1.1.1.2.2.3.9 2 21.7 49.5 68 57.1 4 1 7.1 5.5 4.9 10.8z"],"snapchat-ghost":[512,512,[],"f2ac","M510.846 392.673c-5.211 12.157-27.239 21.089-67.36 27.318-2.064 2.786-3.775 14.686-6.507 23.956-1.625 5.566-5.623 8.869-12.128 8.869l-.297-.005c-9.395 0-19.203-4.323-38.852-4.323-26.521 0-35.662 6.043-56.254 20.588-21.832 15.438-42.771 28.764-74.027 27.399-31.646 2.334-58.025-16.908-72.871-27.404-20.714-14.643-29.828-20.582-56.241-20.582-18.864 0-30.736 4.72-38.852 4.72-8.073 0-11.213-4.922-12.422-9.04-2.703-9.189-4.404-21.263-6.523-24.13-20.679-3.209-67.31-11.344-68.498-32.15a10.627 10.627 0 0 1 8.877-11.069c69.583-11.455 100.924-82.901 102.227-85.934.074-.176.155-.344.237-.515 3.713-7.537 4.544-13.849 2.463-18.753-5.05-11.896-26.872-16.164-36.053-19.796-23.715-9.366-27.015-20.128-25.612-27.504 2.437-12.836 21.725-20.735 33.002-15.453 8.919 4.181 16.843 6.297 23.547 6.297 5.022 0 8.212-1.204 9.96-2.171-2.043-35.936-7.101-87.29 5.687-115.969C158.122 21.304 229.705 15.42 250.826 15.42c.944 0 9.141-.089 10.11-.089 52.148 0 102.254 26.78 126.723 81.643 12.777 28.65 7.749 79.792 5.695 116.009 1.582.872 4.357 1.942 8.599 2.139 6.397-.286 13.815-2.389 22.069-6.257 6.085-2.846 14.406-2.461 20.48.058l.029.01c9.476 3.385 15.439 10.215 15.589 17.87.184 9.747-8.522 18.165-25.878 25.018-2.118.835-4.694 1.655-7.434 2.525-9.797 3.106-24.6 7.805-28.616 17.271-2.079 4.904-1.256 11.211 2.46 18.748.087.168.166.342.239.515 1.301 3.03 32.615 74.46 102.23 85.934 6.427 1.058 11.163 7.877 7.725 15.859z"],"snapchat-square":[448,512,[],"f2ad","M400 32H48C21.5 32 0 53.5 0 80v352c0 26.5 21.5 48 48 48h352c26.5 0 48-21.5 48-48V80c0-26.5-21.5-48-48-48zm-6.5 314.9c-3.5 8.1-18.1 14-44.8 18.2-1.4 1.9-2.5 9.8-4.3 15.9-1.1 3.7-3.7 5.9-8.1 5.9h-.2c-6.2 0-12.8-2.9-25.8-2.9-17.6 0-23.7 4-37.4 13.7-14.5 10.3-28.4 19.1-49.2 18.2-21 1.6-38.6-11.2-48.5-18.2-13.8-9.7-19.8-13.7-37.4-13.7-12.5 0-20.4 3.1-25.8 3.1-5.4 0-7.5-3.3-8.3-6-1.8-6.1-2.9-14.1-4.3-16-13.8-2.1-44.8-7.5-45.5-21.4-.2-3.6 2.3-6.8 5.9-7.4 46.3-7.6 67.1-55.1 68-57.1 0-.1.1-.2.2-.3 2.5-5 3-9.2 1.6-12.5-3.4-7.9-17.9-10.7-24-13.2-15.8-6.2-18-13.4-17-18.3 1.6-8.5 14.4-13.8 21.9-10.3 5.9 2.8 11.2 4.2 15.7 4.2 3.3 0 5.5-.8 6.6-1.4-1.4-23.9-4.7-58 3.8-77.1C159.1 100 206.7 96 220.7 96c.6 0 6.1-.1 6.7-.1 34.7 0 68 17.8 84.3 54.3 8.5 19.1 5.2 53.1 3.8 77.1 1.1.6 2.9 1.3 5.7 1.4 4.3-.2 9.2-1.6 14.7-4.2 4-1.9 9.6-1.6 13.6 0 6.3 2.3 10.3 6.8 10.4 11.9.1 6.5-5.7 12.1-17.2 16.6-1.4.6-3.1 1.1-4.9 1.7-6.5 2.1-16.4 5.2-19 11.5-1.4 3.3-.8 7.5 1.6 12.5.1.1.1.2.2.3.9 2 21.7 49.5 68 57.1 4 1 7.1 5.5 4.9 10.8z"],soundcloud:[640,512,[],"f1be","M111.4 256.3l5.8 65-5.8 68.3c-.3 2.5-2.2 4.4-4.4 4.4s-4.2-1.9-4.2-4.4l-5.6-68.3 5.6-65c0-2.2 1.9-4.2 4.2-4.2 2.2 0 4.1 2 4.4 4.2zm21.4-45.6c-2.8 0-4.7 2.2-5 5l-5 105.6 5 68.3c.3 2.8 2.2 5 5 5 2.5 0 4.7-2.2 4.7-5l5.8-68.3-5.8-105.6c0-2.8-2.2-5-4.7-5zm25.5-24.1c-3.1 0-5.3 2.2-5.6 5.3l-4.4 130 4.4 67.8c.3 3.1 2.5 5.3 5.6 5.3 2.8 0 5.3-2.2 5.3-5.3l5.3-67.8-5.3-130c0-3.1-2.5-5.3-5.3-5.3zM7.2 283.2c-1.4 0-2.2 1.1-2.5 2.5L0 321.3l4.7 35c.3 1.4 1.1 2.5 2.5 2.5s2.2-1.1 2.5-2.5l5.6-35-5.6-35.6c-.3-1.4-1.1-2.5-2.5-2.5zm23.6-21.9c-1.4 0-2.5 1.1-2.5 2.5l-6.4 57.5 6.4 56.1c0 1.7 1.1 2.8 2.5 2.8s2.5-1.1 2.8-2.5l7.2-56.4-7.2-57.5c-.3-1.4-1.4-2.5-2.8-2.5zm25.3-11.4c-1.7 0-3.1 1.4-3.3 3.3L47 321.3l5.8 65.8c.3 1.7 1.7 3.1 3.3 3.1 1.7 0 3.1-1.4 3.1-3.1l6.9-65.8-6.9-68.1c0-1.9-1.4-3.3-3.1-3.3zm25.3-2.2c-1.9 0-3.6 1.4-3.6 3.6l-5.8 70 5.8 67.8c0 2.2 1.7 3.6 3.6 3.6s3.6-1.4 3.9-3.6l6.4-67.8-6.4-70c-.3-2.2-2-3.6-3.9-3.6zm241.4-110.9c-1.1-.8-2.8-1.4-4.2-1.4-2.2 0-4.2.8-5.6 1.9-1.9 1.7-3.1 4.2-3.3 6.7v.8l-3.3 176.7 1.7 32.5 1.7 31.7c.3 4.7 4.2 8.6 8.9 8.6s8.6-3.9 8.6-8.6l3.9-64.2-3.9-177.5c-.4-3-2-5.8-4.5-7.2zm-26.7 15.3c-1.4-.8-2.8-1.4-4.4-1.4s-3.1.6-4.4 1.4c-2.2 1.4-3.6 3.9-3.6 6.7l-.3 1.7-2.8 160.8s0 .3 3.1 65.6v.3c0 1.7.6 3.3 1.7 4.7 1.7 1.9 3.9 3.1 6.4 3.1 2.2 0 4.2-1.1 5.6-2.5 1.7-1.4 2.5-3.3 2.5-5.6l.3-6.7 3.1-58.6-3.3-162.8c-.3-2.8-1.7-5.3-3.9-6.7zm-111.4 22.5c-3.1 0-5.8 2.8-5.8 6.1l-4.4 140.6 4.4 67.2c.3 3.3 2.8 5.8 5.8 5.8 3.3 0 5.8-2.5 6.1-5.8l5-67.2-5-140.6c-.2-3.3-2.7-6.1-6.1-6.1zm376.7 62.8c-10.8 0-21.1 2.2-30.6 6.1-6.4-70.8-65.8-126.4-138.3-126.4-17.8 0-35 3.3-50.3 9.4-6.1 2.2-7.8 4.4-7.8 9.2v249.7c0 5 3.9 8.6 8.6 9.2h218.3c43.3 0 78.6-35 78.6-78.3.1-43.6-35.2-78.9-78.5-78.9zm-296.7-60.3c-4.2 0-7.5 3.3-7.8 7.8l-3.3 136.7 3.3 65.6c.3 4.2 3.6 7.5 7.8 7.5 4.2 0 7.5-3.3 7.5-7.5l3.9-65.6-3.9-136.7c-.3-4.5-3.3-7.8-7.5-7.8zm-53.6-7.8c-3.3 0-6.4 3.1-6.4 6.7l-3.9 145.3 3.9 66.9c.3 3.6 3.1 6.4 6.4 6.4 3.6 0 6.4-2.8 6.7-6.4l4.4-66.9-4.4-145.3c-.3-3.6-3.1-6.7-6.7-6.7zm26.7 3.4c-3.9 0-6.9 3.1-6.9 6.9L227 321.3l3.9 66.4c.3 3.9 3.1 6.9 6.9 6.9s6.9-3.1 6.9-6.9l4.2-66.4-4.2-141.7c0-3.9-3-6.9-6.9-6.9z"],speakap:[448,512,[],"f3f3","M352 32H96C43.2 32 0 75.2 0 128v256c0 52.8 43.2 96 96 96h256c52.8 0 96-43.2 96-96V128c0-52.8-43.2-96-96-96zM221 382.9c-39.6 0-81.9-17.8-81.9-53.7V302H179v17.8c0 15.1 19.5 24.5 41.9 24.5 24.2 0 41.3-10.4 41.3-29.5 0-23.8-27.2-31.9-54.7-42.6-31.9-12.4-63.1-26.2-63.1-69.1 0-48 38.6-66.4 79.9-66.4 37.6 0 75.5 14.1 75.5 41.9v31.2h-39.9v-16.1c0-12.1-17.8-18.5-35.6-18.5-19.5 0-35.6 8.1-35.6 26.2 0 22.1 22.5 29.2 47 38.9 35.9 12.4 71.1 27.2 71.1 71.5.1 48.6-40.8 71.1-85.8 71.1z"],spotify:[496,512,[],"f1bc","M248 8C111.1 8 0 119.1 0 256s111.1 248 248 248 248-111.1 248-248S384.9 8 248 8zm100.7 364.9c-4.2 0-6.8-1.3-10.7-3.6-62.4-37.6-135-39.2-206.7-24.5-3.9 1-9 2.6-11.9 2.6-9.7 0-15.8-7.7-15.8-15.8 0-10.3 6.1-15.2 13.6-16.8 81.9-18.1 165.6-16.5 237 26.2 6.1 3.9 9.7 7.4 9.7 16.5s-7.1 15.4-15.2 15.4zm26.9-65.6c-5.2 0-8.7-2.3-12.3-4.2-62.5-37-155.7-51.9-238.6-29.4-4.8 1.3-7.4 2.6-11.9 2.6-10.7 0-19.4-8.7-19.4-19.4s5.2-17.8 15.5-20.7c27.8-7.8 56.2-13.6 97.8-13.6 64.9 0 127.6 16.1 177 45.5 8.1 4.8 11.3 11 11.3 19.7-.1 10.8-8.5 19.5-19.4 19.5zm31-76.2c-5.2 0-8.4-1.3-12.9-3.9-71.2-42.5-198.5-52.7-280.9-29.7-3.6 1-8.1 2.6-12.9 2.6-13.2 0-23.3-10.3-23.3-23.6 0-13.6 8.4-21.3 17.4-23.9 35.2-10.3 74.6-15.2 117.5-15.2 73 0 149.5 15.2 205.4 47.8 7.8 4.5 12.9 10.7 12.9 22.6 0 13.6-11 23.3-23.2 23.3z"],"stack-exchange":[448,512,[],"f18d","M17.7 332.3h412.7v22c0 37.7-29.3 68-65.3 68h-19L259.3 512v-89.7H83c-36 0-65.3-30.3-65.3-68v-22zm0-23.6h412.7v-85H17.7v85zm0-109.4h412.7v-85H17.7v85zM365 0H83C47 0 17.7 30.3 17.7 67.7V90h412.7V67.7C430.3 30.3 401 0 365 0z"],"stack-overflow":[384,512,[],"f16c","M293.7 300l-181.2-84.5 16.7-36.5 181.3 84.7-16.8 36.3zm48-76L188.2 95.7l-25.5 30.8 153.5 128.3 25.5-30.8zm39.6-31.7L262 32l-32 24 119.3 160.3 32-24zM290.7 311L95 269.7 86.8 309l195.7 41 8.2-39zm31.6 129H42.7V320h-40v160h359.5V320h-40v120zm-39.8-80h-200v39.7h200V360z"],staylinked:[440,512,[],"f3f5","M201.6 127.4c4.1-3.2 10.3-3 13.8.5l170 167.3-2.7-2.7 44.3 41.3c3.7 3.5 3.3 9-.7 12.2l-198 163.9c-9.9 7.6-17.3.8-17.3.8L2.3 314.6c-3.5-3.5-3-9 1.2-12.2l45.8-34.9c4.2-3.2 10.4-3 13.9.5l151.9 147.5c3.7 3.5 10 3.7 14.2.4l93.2-74c4.1-3.2 4.5-8.7.9-12.2l-84-81.3c-3.6-3.5-9.9-3.7-14-.5l-.1.1c-4.1 3.2-10.4 3-14-.5l-68.1-64.3c-3.5-3.5-3.1-9 1.1-12.2l57.3-43.6m14.8 257.3c3.7 3.5 10.1 3.7 14.3.4l50.2-38.8-.3-.3 7.7-6c4.2-3.2 4.6-8.7.9-12.2l-57.1-54.4c-3.6-3.5-10-3.7-14.2-.5l-.1.1c-4.2 3.2-10.5 3.1-14.2-.4L109 180.8c-3.6-3.5-3.1-8.9 1.1-12.2l92.2-71.5c4.1-3.2 10.3-3 13.9.5l160.4 159c3.7 3.5 10 3.7 14.1.5l45.8-35.8c4.1-3.2 4.4-8.7.7-12.2L226.7 2.5c-1.5-1.2-8-5.5-16.3 1.1L3.6 165.7c-4.2 3.2-4.8 8.7-1.2 12.2l42.3 41.7"],steam:[496,512,[],"f1b6","M496 256c0 137-111.2 248-248.4 248-113.8 0-209.6-76.3-239-180.4l95.2 39.3c6.4 32.1 34.9 56.4 68.9 56.4 39.2 0 71.9-32.4 70.2-73.5l84.5-60.2c52.1 1.3 95.8-40.9 95.8-93.5 0-51.6-42-93.5-93.7-93.5s-93.7 42-93.7 93.5v1.2L176.6 279c-15.5-.9-30.7 3.4-43.5 12.1L0 236.1C10.2 108.4 117.1 8 247.6 8 384.8 8 496 119 496 256zM155.7 384.3l-30.5-12.6a52.79 52.79 0 0 0 27.2 25.8c26.9 11.2 57.8-1.6 69-28.4 5.4-13 5.5-27.3.1-40.3-5.4-13-15.5-23.2-28.5-28.6-12.9-5.4-26.7-5.2-38.9-.6l31.5 13c19.8 8.2 29.2 30.9 20.9 50.7-8.3 19.9-31 29.2-50.8 21zm173.8-129.9c-34.4 0-62.4-28-62.4-62.3s28-62.3 62.4-62.3 62.4 28 62.4 62.3-27.9 62.3-62.4 62.3zm.1-15.6c25.9 0 46.9-21 46.9-46.8 0-25.9-21-46.8-46.9-46.8s-46.9 21-46.9 46.8c.1 25.8 21.1 46.8 46.9 46.8z"],"steam-square":[448,512,[],"f1b7","M185.2 356.5c7.7-18.5-1-39.7-19.6-47.4l-29.5-12.2c11.4-4.3 24.3-4.5 36.4.5 12.2 5.1 21.6 14.6 26.7 26.7 5 12.2 5 25.6-.1 37.7-10.5 25.1-39.4 37-64.6 26.5-11.6-4.8-20.4-13.6-25.4-24.2l28.5 11.8c18.6 7.8 39.9-.9 47.6-19.4zM400 32H48C21.5 32 0 53.5 0 80v160.7l116.6 48.1c12-8.2 26.2-12.1 40.7-11.3l55.4-80.2v-1.1c0-48.2 39.3-87.5 87.6-87.5s87.6 39.3 87.6 87.5c0 49.2-40.9 88.7-89.6 87.5l-79 56.3c1.6 38.5-29.1 68.8-65.7 68.8-31.8 0-58.5-22.7-64.5-52.7L0 319.2V432c0 26.5 21.5 48 48 48h352c26.5 0 48-21.5 48-48V80c0-26.5-21.5-48-48-48zm-99.7 222.5c-32.2 0-58.4-26.1-58.4-58.3s26.2-58.3 58.4-58.3 58.4 26.2 58.4 58.3-26.2 58.3-58.4 58.3zm.1-14.6c24.2 0 43.9-19.6 43.9-43.8 0-24.2-19.6-43.8-43.9-43.8-24.2 0-43.9 19.6-43.9 43.8 0 24.2 19.7 43.8 43.9 43.8z"],"steam-symbol":[448,512,[],"f3f6","M395.5 177.5c0 33.8-27.5 61-61 61-33.8 0-61-27.3-61-61s27.3-61 61-61c33.5 0 61 27.2 61 61zm52.5.2c0 63-51 113.8-113.7 113.8L225 371.3c-4 43-40.5 76.8-84.5 76.8-40.5 0-74.7-28.8-83-67L0 358V250.7L97.2 290c15.1-9.2 32.2-13.3 52-11.5l71-101.7c.5-62.3 51.5-112.8 114-112.8C397 64 448 115 448 177.7zM203 363c0-34.7-27.8-62.5-62.5-62.5-4.5 0-9 .5-13.5 1.5l26 10.5c25.5 10.2 38 39 27.7 64.5-10.2 25.5-39.2 38-64.7 27.5-10.2-4-20.5-8.3-30.7-12.2 10.5 19.7 31.2 33.2 55.2 33.2 34.7 0 62.5-27.8 62.5-62.5zm207.5-185.3c0-42-34.3-76.2-76.2-76.2-42.3 0-76.5 34.2-76.5 76.2 0 42.2 34.3 76.2 76.5 76.2 41.9.1 76.2-33.9 76.2-76.2z"],"sticker-mule":[576,512,[],"f3f7","M353.1 509.8c-5.9 2.9-32.1 3.2-36.5-.5-4.1-3-2.2-11.9-1.5-15 2.2-15-2.5-7.9-9.8-11.5-3.1-1.5-4.1-5.5-4.6-10-.5-1.5-1-2.5-1.5-3.5-1.7-10.7 6.8-33.6 8.2-43.4 4.9-23.7-.7-37.2 1.5-46.9 3.7-16.2 4.1-3.5 4.1-29.9-1.4-25.9 3.3-36.9.5-38.9-14.8 0-64.3 10.7-112.2 2-46.1-8.9-59.4-29-65.4-30.9-10.3-4.5-23.2.5-27.3 7-.1.1-35 70.6-39.6 87.8-6.2 20.5-.5 47.4 4.1 66.8 0 .1 4.5 14.6 10.3 19.5 2.1 1.5 5.1 2.5 7.2 4.5 2.8 2.7 9.4 15.2 9.8 16 2.6 4.5 3.6 8-1.5 10.5-3.6 2-9.3 2.5-14.4 2.5-2.6.5-1.5 3.5-3.1 5-2.9 2.8-20.7 6.1-29.9 2.5-2.6-1-5.7-3-6.2-5-1.5-4 2.1-9-1-12.5-4.5-2.9-13.1-2-17-12-2.2-5.4-2.6-7.6-2.6-49.4 0-9.7-5.9-38.7-8.2-46.9-1.5-5.5-1.5-11.5 0-16 .3-.9 4.1-4.6 4.1-13-1-1.5-4.6-.5-5.1-1.5-10.4-80.6-5.9-79-7.7-98.3-1.5-16-10.9-43.9-6.7-64.3.5-2.4 3.4-21 24.2-38.9 31-26.7 48.4-38.3 159-11.5 1.1.4 66.3 21.1 110.7-9 15.5-11.3 28.8-11.3 35.5-16 .1-.1 61.7-52.1 87-65.3 47.2-29.4 69.9-16.7 75.1-18 4.7-1 13.4-25.8 17-25.8 5.5 0 1.6 20.2 3.6 25.9.5 2 3.6 5 6.2 5 2.3 0 1.7-.8 10.3-5 8.4-5.4 14.9-17.6 20.6-17 11.7 1.6-19 41.6-19 46.9 0 2 .2.8 4.6 9.5 2.6 5.5 4.6 13.5 6.2 20 8.3 29.7 5.7 14.6 13.4 36.9 20.2 50.1 20.6 45.2 20.6 52.9 0 7.5-4.1 11-7.2 16.5-1.5 3-4.6 7.5-7.2 8-2.7.7 7-1.5-13.4 2.5-7.2 1-13.4-4.5-14.9-9.5-1.6-4.7 2.8-10.1-11.8-22.9-10.3-10-21.1-11.3-31.9-17-9.8-5.7-11.9 1-18 8-18 22.9-34 46.9-52 69.8-11.8 15-24.2 30.4-33.5 47.4-3.9 6.8-9.5 28.1-10.3 29.9-6.2 17.7-5.5 25.8-16.5 68.3-3.1 10-5.7 21.4-8.7 32.4-2.2 6.8-7.4 49.3-.5 59.4 2.1 3.5 8.7 4.5 11.3 8 .1.1 9.6 18.2 9.3 20 0 6.1-9.4 5.6-11.3 6.5-4.8 2.9-3.8 5.9-6.4 7.4"],strava:[369,512,[],"f428","M301.6 292l-43.9 88.2-44.6-88.2h-67.6l112.2 220 111.5-220h-67.6zM151.4 0L0 292h89.2l62.2-116.1L213.1 292h88.5L151.4 0z"],stripe:[640,512,[],"f429","M640 261.6c0-45.5-22-81.4-64.2-81.4s-67.9 35.9-67.9 81.1c0 53.5 30.3 78.2 73.5 78.2 21.2 0 37.1-4.8 49.2-11.5v-33.4c-12.1 6.1-26 9.8-43.6 9.8-17.3 0-32.5-6.1-34.5-26.9h86.9c.2-2.3.6-11.6.6-15.9m-87.9-16.8c0-20 12.3-28.4 23.4-28.4 10.9 0 22.5 8.4 22.5 28.4h-45.9zm-112.9-64.6c-17.4 0-28.6 8.2-34.8 13.9l-2.3-11H363v204.8l44.4-9.4.1-50.2c6.4 4.7 15.9 11.2 31.4 11.2 31.8 0 60.8-23.2 60.8-79.6.1-51.6-29.3-79.7-60.5-79.7m-10.6 122.5c-10.4 0-16.6-3.8-20.9-8.4l-.3-66c4.6-5.1 11-8.8 21.2-8.8 16.2 0 27.4 18.2 27.4 41.4.1 23.9-10.9 41.8-27.4 41.8M346.4 124v36.2l-44.6 9.5v-36.2l44.6-9.5m-44.5 59.2h44.6v153.2h-44.6V183.2zm-47.8 13.1c10.4-19.1 31.1-15.2 37.1-13.1V224c-5.7-1.8-23.4-4.5-33.9 9.3v103.1H213V183.2h38.4l2.7 13.1m-89-13.1h33.7V221h-33.7v63.2c0 26.2 28 18 33.7 15.7v33.8c-5.9 3.2-16.6 5.9-31.2 5.9-26.3 0-46.1-17-46.1-43.3l.2-142.4 43.3-9.2.1 38.5zM44.9 228.3c0 20 67.9 10.5 67.9 63.4 0 32-25.4 47.8-62.3 47.8-15.3 0-32-3-48.5-10.1v-40c14.9 8.1 33.9 14.2 48.6 14.2 9.9 0 17-2.7 17-10.9 0-21.2-67.5-13.2-67.5-62.4 0-31.4 24-50.2 60-50.2 14.7 0 29.4 2.3 44.1 8.1V230c-13.5-7.3-30.7-11.4-44.2-11.4-9.3.1-15.1 2.8-15.1 9.7"],"stripe-s":[362,512,[],"f42a","M144.3 154.6c0-22.3 18.6-30.9 48.4-30.9 43.4 0 98.5 13.3 141.9 36.7V26.1C287.3 7.2 240.1 0 192.8 0 77.1 0 0 60.4 0 161.4c0 157.9 216.8 132.3 216.8 200.4 0 26.4-22.9 34.9-54.7 34.9-47.2 0-108.2-19.5-156.1-45.5v128.5c53 22.8 106.8 32.4 156 32.4 118.6 0 200.3-51 200.3-153.6 0-170.2-218-139.7-218-203.9"],studiovinari:[512,512,[],"f3f8","M480.3 187.7l4.2 28v28l-25.1 44.1-39.8 78.4-56.1 67.5-79.1 37.8-17.7 24.5-7.7 12-9.6 4s17.3-63.6 19.4-63.6c2.1 0 20.3.7 20.3.7l66.7-38.6-92.5 26.1-55.9 36.8-22.8 28-6.6 1.4 20.8-73.6 6.9-5.5 20.7 12.9 88.3-45.2 56.8-51.5 14.8-68.4-125.4 23.3 15.2-18.2-173.4-53.3 81.9-10.5-166-122.9L133.5 108 32.2 0l252.9 126.6-31.5-38L378 163 234.7 64l18.7 38.4-49.6-18.1L158.3 0l194.6 122L310 66.2l108 96.4 12-8.9-21-16.4 4.2-37.8L451 89.1l29.2 24.7 11.5 4.2-7 6.2 8.5 12-13.1 7.4-10.3 20.2 10.5 23.9z"],stumbleupon:[512,512,[],"f1a4","M502.9 266v69.7c0 62.1-50.3 112.4-112.4 112.4-61.8 0-112.4-49.8-112.4-111.3v-70.2l34.3 16 51.1-15.2V338c0 14.7 12 26.5 26.7 26.5S417 352.7 417 338v-72h85.9zm-224.7-58.2l34.3 16 51.1-15.2V173c0-60.5-51.1-109-112.1-109-60.8 0-112.1 48.2-112.1 108.2v162.4c0 14.9-12 26.7-26.7 26.7S86 349.5 86 334.6V266H0v69.7C0 397.7 50.3 448 112.4 448c61.6 0 112.4-49.5 112.4-110.8V176.9c0-14.7 12-26.7 26.7-26.7s26.7 12 26.7 26.7v30.9z"],"stumbleupon-circle":[496,512,[],"f1a3","M256 8C119 8 8 119 8 256s111 248 248 248 248-111 248-248S393 8 256 8zm0 177.5c-9.8 0-17.8 8-17.8 17.8v106.9c0 40.9-33.9 73.9-74.9 73.9-41.4 0-74.9-33.5-74.9-74.9v-46.5h57.3v45.8c0 10 8 17.8 17.8 17.8s17.8-7.9 17.8-17.8V200.1c0-40 34.2-72.1 74.7-72.1 40.7 0 74.7 32.3 74.7 72.6v23.7l-34.1 10.1-22.9-10.7v-20.6c.1-9.6-7.9-17.6-17.7-17.6zm167.6 123.6c0 41.4-33.5 74.9-74.9 74.9-41.2 0-74.9-33.2-74.9-74.2V263l22.9 10.7 34.1-10.1v47.1c0 9.8 8 17.6 17.8 17.6s17.8-7.9 17.8-17.6v-48h57.3c-.1 45.9-.1 46.4-.1 46.4z"],superpowers:[448,512,[],"f2dd","M448 32c-83.3 11-166.8 22-250 33-92 12.5-163.3 86.7-169 180-3.3 55.5 18 109.5 57.8 148.2L0 480c83.3-11 166.5-22 249.8-33 91.8-12.5 163.3-86.8 168.7-179.8 3.5-55.5-18-109.5-57.7-148.2L448 32zm-79.7 232.3c-4.2 79.5-74 139.2-152.8 134.5-79.5-4.7-140.7-71-136.3-151 4.5-79.2 74.3-139.3 153-134.5 79.3 4.7 140.5 71 136.1 151z"],supple:[640,512,[],"f3f9","M640 262.5c0 64.1-109 116.1-243.5 116.1-24.8 0-48.6-1.8-71.1-5 7.7.4 15.5.6 23.4.6 134.5 0 243.5-56.9 243.5-127.1 0-29.4-19.1-56.4-51.2-78 60 21.1 98.9 55.1 98.9 93.4zM47.7 227.9c-.1-70.2 108.8-127.3 243.3-127.6 7.9 0 15.6.2 23.3.5-22.5-3.2-46.3-4.9-71-4.9C108.8 96.3-.1 148.5 0 212.6c.1 38.3 39.1 72.3 99.3 93.3-32.3-21.5-51.5-48.6-51.6-78zm60.2 39.9s10.5 13.2 29.3 13.2c17.9 0 28.4-11.5 28.4-25.1 0-28-40.2-25.1-40.2-39.7 0-5.4 5.3-9.1 12.5-9.1 5.7 0 11.3 2.6 11.3 6.6v3.9h14.2v-7.9c0-12.1-15.4-16.8-25.4-16.8-16.5 0-28.5 10.2-28.5 24.1 0 26.6 40.2 25.4 40.2 39.9 0 6.6-5.8 10.1-12.3 10.1-11.9 0-20.7-10.1-20.7-10.1l-8.8 10.9zm120.8-73.6v54.4c0 11.3-7.1 17.8-17.8 17.8-10.7 0-17.8-6.5-17.8-17.7v-54.5h-15.8v55c0 18.9 13.4 31.9 33.7 31.9 20.1 0 33.4-13 33.4-31.9v-55h-15.7zm34.4 85.4h15.8v-29.5h15.5c16 0 27.2-11.5 27.2-28.1s-11.2-27.8-27.2-27.8h-39.1v13.4h7.8v72zm15.8-43v-29.1h12.9c8.7 0 13.7 5.7 13.7 14.4 0 8.9-5.1 14.7-14 14.7h-12.6zm57 43h15.8v-29.5h15.5c16 0 27.2-11.5 27.2-28.1s-11.2-27.8-27.2-27.8h-39.1v13.4h7.8v72zm15.7-43v-29.1h12.9c8.7 0 13.7 5.7 13.7 14.4 0 8.9-5 14.7-14 14.7h-12.6zm57.1 34.8c0 5.8 2.4 8.2 8.2 8.2h37.6c5.8 0 8.2-2.4 8.2-8.2v-13h-14.3v5.2c0 1.7-1 2.6-2.6 2.6h-18.6c-1.7 0-2.6-1-2.6-2.6v-61.2c0-5.7-2.4-8.2-8.2-8.2H401v13.4h5.2c1.7 0 2.6 1 2.6 2.6v61.2zm63.4 0c0 5.8 2.4 8.2 8.2 8.2H519c5.7 0 8.2-2.4 8.2-8.2v-13h-14.3v5.2c0 1.7-1 2.6-2.6 2.6h-19.7c-1.7 0-2.6-1-2.6-2.6v-20.3h27.7v-13.4H488v-22.4h19.2c1.7 0 2.6 1 2.6 2.6v5.2H524v-13c0-5.7-2.5-8.2-8.2-8.2h-51.6v13.4h7.8v63.9zm58.9-76v5.9h1.6v-5.9h2.7v-1.2h-7v1.2h2.7zm5.7-1.2v7.1h1.5v-5.7l2.3 5.7h1.3l2.3-5.7v5.7h1.5v-7.1h-2.3l-2.1 5.1-2.1-5.1h-2.4z"],teamspeak:[453,512,[],"f4f9","M203.4 65.8c-10.3-2-42.1-3.7-52.3-3.7-8.4 0-6.9 2-6.9 2 63.7 50.1 75.4 121.1 75.6 122.1 1.9 15.6-3.1 22.9-8.7 31.3-1.4 1.8-2.7 5 .3 8.9l28.9 39.4h-.3c9.4 14.1 28.1 32.2-5.4 39.9-7.7 1.9-18.7 6.3-11.2 15.8 2.2 2.9 8.8 9.5 1.8 16.5-7.5 4.8-2.7 7.9-1.8 10.4 1.8 6.6-2.6 11.2-4.3 12.7-4.2 3.2-3.7 7.8-2.6 10.1v.1c3.1 13.1-1.7 27.1-3.9 34.9l.1-.1c-1.4 3.2 22-2.6 22.1-2.6 87.5-23 150.6-88.5 150.6-165.7-.1-86.3-78.7-157.9-182-172m239.4 126.7c-12.9-35.4-34.5-62.3-58.9-85.4-75.8-71.7-175.6-76.6-214-74.8C133 34 79.5 43.7 67.1 59.4c-4.6 5.8-4 6.2-24.4 103.2C36.7 165 0 181.3 0 225.9c-.3 88.7 133.8 89.3 133.8-.3 0-24.6-13-46.1-32.5-58V49.2c12-3 119.7-28.8 223.6 37.3 29.8 19 104.3 76.9 96.9 182.8-1.5 21.2-5.7 40.8-13.6 56.4-57 113.5-170.7 133.8-212.8 137.9-68.3 6.6-122.5-14.1-124-14-2.7.1-12 7.1-11.7 7.4 36.6 12.1 74.3 23 121 23 48 0 84.3-8.2 118-21.6C401.8 417.3 453.6 340 453.6 253.7c0-22.9-4.3-43.2-10.8-61.2m-246.3 33.6c-8.5-15.6.5-15.9 4.8-23.4 5.5-9.6 1.7-22.1-1.7-32.2-9.6-28.5-33.8-84.5-83-103.6v100c4.5 4.5 54.1 33.5 58.2 116.8 4.1 82.7-52 121.6-51.7 123.3 22.8 4.5 66 10.7 76.7-8.6 5.7-10.2-1.5-28.1-.9-28.1-3.6-14.2 7.8-19.4 7.8-21.8 0-1.4-7.2-3.1-7.2-8.1 0-5.9 8.5-6.2 9.4-10.7.9-4.5-3.1-7.8-5.1-16.6-3.5-19 27.9-16 30.8-23.1 1.4-3.3-1.4-7.8-3.4-10.7"],telegram:[496,512,[],"f2c6","M248 8C111 8 0 119 0 256s111 248 248 248 248-111 248-248S385 8 248 8zm121.8 169.9l-40.7 191.8c-3 13.6-11.1 16.9-22.4 10.5l-62-45.7-29.9 28.8c-3.3 3.3-6.1 6.1-12.5 6.1l4.4-63.1 114.9-103.8c5-4.4-1.1-6.9-7.7-2.5l-142 89.4-61.2-19.1c-13.3-4.2-13.6-13.3 2.8-19.7l239.1-92.2c11.1-4 20.8 2.7 17.2 19.5z"],"telegram-plane":[448,512,[],"f3fe","M446.7 98.6l-67.6 318.8c-5.1 22.5-18.4 28.1-37.3 17.5l-103-75.9-49.7 47.8c-5.5 5.5-10.1 10.1-20.7 10.1l7.4-104.9 190.9-172.5c8.3-7.4-1.8-11.5-12.9-4.1L117.8 284 16.2 252.2c-22.1-6.9-22.5-22.1 4.6-32.7L418.2 66.4c18.4-6.9 34.5 4.1 28.5 32.2z"],"tencent-weibo":[384,512,[],"f1d5","M72.3 495.8c1.4 19.9-27.6 22.2-29.7 2.9C31 368.8 73.7 259.2 144 185.5c-15.6-34 9.2-77.1 50.6-77.1 30.3 0 55.1 24.6 55.1 55.1 0 44-49.5 70.8-86.9 45.1-65.7 71.3-101.4 169.8-90.5 287.2zM192 .1C66.1.1-12.3 134.3 43.7 242.4 52.4 259.8 79 246.9 70 229 23.7 136.4 91 29.8 192 29.8c75.4 0 136.9 61.4 136.9 136.9 0 90.8-86.9 153.9-167.7 133.1-19.1-4.1-25.6 24.4-6.6 29.1 110.7 23.2 204-60 204-162.3C358.6 74.7 284 .1 192 .1z"],themeisle:[512,512,[],"f2b2","M208 88.286c0-10 6.286-21.714 17.715-21.714 11.142 0 17.714 11.714 17.714 21.714 0 10.285-6.572 21.714-17.714 21.714C214.286 110 208 98.571 208 88.286zm304 160c0 36.001-11.429 102.286-36.286 129.714-22.858 24.858-87.428 61.143-120.857 70.572l-1.143.286v32.571c0 16.286-12.572 30.571-29.143 30.571-10 0-19.429-5.714-24.572-14.286-5.427 8.572-14.856 14.286-24.856 14.286-10 0-19.429-5.714-24.858-14.286-5.142 8.572-14.571 14.286-24.57 14.286-10.286 0-19.429-5.714-24.858-14.286-5.143 8.572-14.571 14.286-24.571 14.286-18.857 0-29.429-15.714-29.429-32.857-16.286 12.285-35.715 19.428-56.571 19.428-22 0-43.429-8.285-60.286-22.857 10.285-.286 20.571-2.286 30.285-5.714-20.857-5.714-39.428-18.857-52-36.286 21.37 4.645 46.209 1.673 67.143-11.143-22-22-56.571-58.857-68.572-87.428C1.143 321.714 0 303.714 0 289.429c0-49.714 20.286-160 86.286-160 10.571 0 18.857 4.858 23.143 14.857a158.792 158.792 0 0 1 12-15.428c2-2.572 5.714-5.429 7.143-8.286 7.999-12.571 11.714-21.142 21.714-34C182.571 45.428 232 17.143 285.143 17.143c6 0 12 .285 17.714 1.143C313.714 6.571 328.857 0 344.572 0c14.571 0 29.714 6 40 16.286.857.858 1.428 2.286 1.428 3.428 0 3.714-10.285 13.429-12.857 16.286 4.286 1.429 15.714 6.858 15.714 12 0 2.857-2.857 5.143-4.571 7.143 31.429 27.714 49.429 67.143 56.286 108 4.286-5.143 10.285-8.572 17.143-8.572 10.571 0 20.857 7.144 28.571 14.001C507.143 187.143 512 221.714 512 248.286zM188 89.428c0 18.286 12.571 37.143 32.286 37.143 19.714 0 32.285-18.857 32.285-37.143 0-18-12.571-36.857-32.285-36.857-19.715 0-32.286 18.858-32.286 36.857zM237.714 194c0-19.714 3.714-39.143 8.571-58.286-52.039 79.534-13.531 184.571 68.858 184.571 21.428 0 42.571-7.714 60-20 2-7.429 3.714-14.857 3.714-22.572 0-14.286-6.286-21.428-20.572-21.428-4.571 0-9.143.857-13.429 1.714-63.343 12.668-107.142 3.669-107.142-63.999zm-41.142 254.858c0-11.143-8.858-20.857-20.286-20.857-11.429 0-20 9.715-20 20.857v32.571c0 11.143 8.571 21.142 20 21.142 11.428 0 20.286-9.715 20.286-21.142v-32.571zm49.143 0c0-11.143-8.572-20.857-20-20.857-11.429 0-20.286 9.715-20.286 20.857v32.571c0 11.143 8.857 21.142 20.286 21.142 11.428 0 20-10 20-21.142v-32.571zm49.713 0c0-11.143-8.857-20.857-20.285-20.857-11.429 0-20.286 9.715-20.286 20.857v32.571c0 11.143 8.857 21.142 20.286 21.142 11.428 0 20.285-9.715 20.285-21.142v-32.571zm49.715 0c0-11.143-8.857-20.857-20.286-20.857-11.428 0-20.286 9.715-20.286 20.857v32.571c0 11.143 8.858 21.142 20.286 21.142 11.429 0 20.286-10 20.286-21.142v-32.571zM421.714 286c-30.857 59.142-90.285 102.572-158.571 102.572-96.571 0-160.571-84.572-160.571-176.572 0-16.857 2-33.429 6-49.714-20 33.715-29.714 72.572-29.714 111.429 0 60.286 24.857 121.715 71.429 160.857 5.143-9.714 14.857-16.286 26-16.286 10 0 19.428 5.714 24.571 14.286 5.429-8.571 14.571-14.286 24.858-14.286 10 0 19.428 5.714 24.571 14.286 5.429-8.571 14.857-14.286 24.858-14.286 10 0 19.428 5.714 24.857 14.286 5.143-8.571 14.571-14.286 24.572-14.286 10.857 0 20.857 6.572 25.714 16 43.427-36.286 68.569-92 71.426-148.286zm10.572-99.714c0-53.714-34.571-105.714-92.572-105.714-30.285 0-58.571 15.143-78.857 36.857C240.862 183.812 233.41 254 302.286 254c28.805 0 97.357-28.538 84.286 36.857 28.857-26 45.714-65.714 45.714-104.571z"],"trade-federation":[496,512,[],"f513","M202.45 108.42v81.61H38.94l48.22 61.91h114.73v196.75h64.61V252.96h83.27v-62.69h-83.18V171.1h145.62v-62.68H202.45zm4.86 6h197.77v50.68H259.44v30.93h83.18v50.93h-83.26v195.73h-52.73V245.94H89.86l-39.95-49.91h157.4v-81.61zM247.99 8.8C111.03 8.8 0 119.83 0 256.8s111.03 248 247.99 248S496 393.76 496 256.8 384.96 8.8 247.99 8.8zm.02 13.24c129.66 0 234.76 105.12 234.76 234.78s-105.1 234.76-234.76 234.76S13.23 386.47 13.23 256.81 118.35 22.04 248.01 22.04zm0 7.89c-125.3 0-226.89 101.57-226.89 226.87s101.59 226.89 226.89 226.89S474.88 382.1 474.88 256.8 373.31 29.93 248.01 29.93zm-.02 13.3c117.95 0 213.56 95.62 213.56 213.56s-95.62 213.56-213.56 213.56S34.43 374.75 34.43 256.8 130.04 43.23 247.99 43.23zm-73.32 104.8l15.66 18.05-22.16-9.45-12.33 20.47 2.15-23.99-23.28-5.4 23.48-5.37-2.06-23.81 12.37 20.67 22.01-9.32-15.84 18.15zm-47.14-46.54l8.01 12.4 12.69-5.86-9.32 11.45 9.5 10.25-13.77-5.32-6.82 12.2.81-14.74-13.71-2.71 14.27-3.78-1.66-13.89zm250.77 75.59l11.99 19.56 20.88-9.11-14.9 17.45 15.11 17.04-21.2-8.78-11.54 19.64 1.8-22.87-22.24-4.9 22.31-5.36-2.21-22.67zm-107.05 98.45l13.86-3.22-1.28-14.17 7.35 12.18 13.08-5.59-9.31 10.75 9.36 10.71-13.1-5.54-7.29 12.21 1.22-14.17-13.89-3.16zm-125.43 6.5l23.73-3.87-1.73-24.53 11.01 21.38 22.79-9.22-16.93 17.08 15.81 18.83-21.47-10.82-13.02 20.86 3.66-23.77-23.85-5.94zm63.44-165.66v81.61H54.16l36.72 46.01h117.7v196.75h48.82V245.01h83.27v-47.03H257.4v-34.83h145.73v-46.78H209.26zm10.86 11.2h171.8v24.36H246.88v56.23h82.93v23.94h-82.93v197.42h-26.76V232.1H96.31l-20.09-23.94h143.9v-80.59z"],trello:[448,512,[],"f181","M392 32H56C25.1 32 0 57.1 0 88v336c0 30.9 25.1 56 56 56h336c30.9 0 56-25.1 56-56V88c0-30.9-25.1-56-56-56zM194.9 371.4c0 14.8-12 26.9-26.9 26.9H85.1c-14.8 0-26.9-12-26.9-26.9V117.1c0-14.8 12-26.9 26.9-26.9H168c14.8 0 26.9 12 26.9 26.9v254.3zm194.9-112c0 14.8-12 26.9-26.9 26.9H280c-14.8 0-26.9-12-26.9-26.9V117.1c0-14.8 12-26.9 26.9-26.9h82.9c14.8 0 26.9 12 26.9 26.9v142.3z"],tripadvisor:[576,512,[],"f262","M166.4 280.521c0 13.236-10.73 23.966-23.966 23.966s-23.966-10.73-23.966-23.966 10.73-23.966 23.966-23.966 23.966 10.729 23.966 23.966zm264.962-23.956c-13.23 0-23.956 10.725-23.956 23.956 0 13.23 10.725 23.956 23.956 23.956 13.23 0 23.956-10.725 23.956-23.956-.001-13.231-10.726-23.956-23.956-23.956zm89.388 139.49c-62.667 49.104-153.276 38.109-202.379-24.559l-30.979 46.325-30.683-45.939c-48.277 60.39-135.622 71.891-197.885 26.055-64.058-47.158-77.759-137.316-30.601-201.374A186.762 186.762 0 0 0 0 139.416l90.286-.05a358.48 358.48 0 0 1 197.065-54.03 350.382 350.382 0 0 1 192.181 53.349l96.218.074a185.713 185.713 0 0 0-28.352 57.649c46.793 62.747 34.964 151.37-26.648 199.647zM259.366 281.761c-.007-63.557-51.535-115.075-115.092-115.068C80.717 166.7 29.2 218.228 29.206 281.785c.007 63.557 51.535 115.075 115.092 115.068 63.513-.075 114.984-51.539 115.068-115.052v-.04zm28.591-10.455c5.433-73.44 65.51-130.884 139.12-133.022a339.146 339.146 0 0 0-139.727-27.812 356.31 356.31 0 0 0-140.164 27.253c74.344 1.582 135.299 59.424 140.771 133.581zm251.706-28.767c-21.992-59.634-88.162-90.148-147.795-68.157-59.634 21.992-90.148 88.162-68.157 147.795v.032c22.038 59.607 88.198 90.091 147.827 68.113 59.615-22.004 90.113-88.162 68.125-147.783zm-326.039 37.975v.115c-.057 39.328-31.986 71.163-71.314 71.106-39.328-.057-71.163-31.986-71.106-71.314.057-39.328 31.986-71.163 71.314-71.106 39.259.116 71.042 31.94 71.106 71.199zm-24.512 0v-.084c-.051-25.784-20.994-46.645-46.778-46.594-25.784.051-46.645 20.994-46.594 46.777.051 25.784 20.994 46.645 46.777 46.594 25.726-.113 46.537-20.968 46.595-46.693zm313.423 0v.048c-.02 39.328-31.918 71.194-71.247 71.173s-71.194-31.918-71.173-71.247c.02-39.328 31.918-71.194 71.247-71.173 39.29.066 71.121 31.909 71.173 71.199zm-24.504-.008c-.009-25.784-20.918-46.679-46.702-46.67-25.784.009-46.679 20.918-46.67 46.702.009 25.784 20.918 46.678 46.702 46.67 25.765-.046 46.636-20.928 46.67-46.693v-.009z"],tumblr:[320,512,[],"f173","M309.8 480.3c-13.6 14.5-50 31.7-97.4 31.7-120.8 0-147-88.8-147-140.6v-144H17.9c-5.5 0-10-4.5-10-10v-68c0-7.2 4.5-13.6 11.3-16 62-21.8 81.5-76 84.3-117.1.8-11 6.5-16.3 16.1-16.3h70.9c5.5 0 10 4.5 10 10v115.2h83c5.5 0 10 4.4 10 9.9v81.7c0 5.5-4.5 10-10 10h-83.4V360c0 34.2 23.7 53.6 68 35.8 4.8-1.9 9-3.2 12.7-2.2 3.5.9 5.8 3.4 7.4 7.9l22 64.3c1.8 5 3.3 10.6-.4 14.5z"],"tumblr-square":[448,512,[],"f174","M400 32H48C21.5 32 0 53.5 0 80v352c0 26.5 21.5 48 48 48h352c26.5 0 48-21.5 48-48V80c0-26.5-21.5-48-48-48zm-82.3 364.2c-8.5 9.1-31.2 19.8-60.9 19.8-75.5 0-91.9-55.5-91.9-87.9v-90h-29.7c-3.4 0-6.2-2.8-6.2-6.2v-42.5c0-4.5 2.8-8.5 7.1-10 38.8-13.7 50.9-47.5 52.7-73.2.5-6.9 4.1-10.2 10-10.2h44.3c3.4 0 6.2 2.8 6.2 6.2v72h51.9c3.4 0 6.2 2.8 6.2 6.2v51.1c0 3.4-2.8 6.2-6.2 6.2h-52.1V321c0 21.4 14.8 33.5 42.5 22.4 3-1.2 5.6-2 8-1.4 2.2.5 3.6 2.1 4.6 4.9l13.8 40.2c1 3.2 2 6.7-.3 9.1z"],twitch:[448,512,[],"f1e8","M40.1 32L10 108.9v314.3h107V480h60.2l56.8-56.8h87l117-117V32H40.1zm357.8 254.1L331 353H224l-56.8 56.8V353H76.9V72.1h321v214zM331 149v116.9h-40.1V149H331zm-107 0v116.9h-40.1V149H224z"],twitter:[512,512,[],"f099","M459.37 151.716c.325 4.548.325 9.097.325 13.645 0 138.72-105.583 298.558-298.558 298.558-59.452 0-114.68-17.219-161.137-47.106 8.447.974 16.568 1.299 25.34 1.299 49.055 0 94.213-16.568 130.274-44.832-46.132-.975-84.792-31.188-98.112-72.772 6.498.974 12.995 1.624 19.818 1.624 9.421 0 18.843-1.3 27.614-3.573-48.081-9.747-84.143-51.98-84.143-102.985v-1.299c13.969 7.797 30.214 12.67 47.431 13.319-28.264-18.843-46.781-51.005-46.781-87.391 0-19.492 5.197-37.36 14.294-52.954 51.655 63.675 129.3 105.258 216.365 109.807-1.624-7.797-2.599-15.918-2.599-24.04 0-57.828 46.782-104.934 104.934-104.934 30.213 0 57.502 12.67 76.67 33.137 23.715-4.548 46.456-13.32 66.599-25.34-7.798 24.366-24.366 44.833-46.132 57.827 21.117-2.273 41.584-8.122 60.426-16.243-14.292 20.791-32.161 39.308-52.628 54.253z"],"twitter-square":[448,512,[],"f081","M400 32H48C21.5 32 0 53.5 0 80v352c0 26.5 21.5 48 48 48h352c26.5 0 48-21.5 48-48V80c0-26.5-21.5-48-48-48zm-48.9 158.8c.2 2.8.2 5.7.2 8.5 0 86.7-66 186.6-186.6 186.6-37.2 0-71.7-10.8-100.7-29.4 5.3.6 10.4.8 15.8.8 30.7 0 58.9-10.4 81.4-28-28.8-.6-53-19.5-61.3-45.5 10.1 1.5 19.2 1.5 29.6-1.2-30-6.1-52.5-32.5-52.5-64.4v-.8c8.7 4.9 18.9 7.9 29.6 8.3a65.447 65.447 0 0 1-29.2-54.6c0-12.2 3.2-23.4 8.9-33.1 32.3 39.8 80.8 65.8 135.2 68.6-9.3-44.5 24-80.6 64-80.6 18.9 0 35.9 7.9 47.9 20.7 14.8-2.8 29-8.3 41.6-15.8-4.9 15.2-15.2 28-28.8 36.1 13.2-1.4 26-5.1 37.8-10.2-8.9 13.1-20.1 24.7-32.9 34z"],typo3:[433,512,[],"f42b","M330.8 341c-7 2.3-11.6 2.3-18.5 2.3-57.2 0-140.6-198.5-140.6-264.9 0-24.7 5.4-32.4 13.9-39.4-69.5 8.5-149.3 34-176.3 66.4-5.4 7.7-9.3 20.8-9.3 37.1C0 246 106.8 480 184.1 480c36.3 0 97.3-59.5 146.7-139M294.5 32c71.8 0 138.8 11.6 138.8 52.5 0 82.6-52.5 182.3-78.8 182.3-47.9 0-101.7-132.1-101.7-198.5 0-30.9 11.6-36.3 41.7-36.3"],uber:[448,512,[],"f402","M414.1 32H33.9C15.2 32 0 47.2 0 65.9V446c0 18.8 15.2 34 33.9 34H414c18.7 0 33.9-15.2 33.9-33.9V65.9C448 47.2 432.8 32 414.1 32zM237.6 391.1C163 398.6 96.4 344.2 88.9 269.6h94.4V290c0 3.7 3 6.8 6.8 6.8H258c3.7 0 6.8-3 6.8-6.8v-67.9c0-3.7-3-6.8-6.8-6.8h-67.9c-3.7 0-6.8 3-6.8 6.8v20.4H88.9c7-69.4 65.4-122.2 135.1-122.2 69.7 0 128.1 52.8 135.1 122.2 7.5 74.5-46.9 141.1-121.5 148.6z"],uikit:[448,512,[],"f403","M443.9 128v256L218 512 0 384V169.7l87.6 45.1v117l133.5 75.5 135.8-75.5v-151l-101.1-57.6 87.6-53.1L443.9 128zM308.6 49.1L223.8 0l-88.6 54.8 86 47.3 87.4-53z"],uniregistry:[384,512,[],"f404","M281.1 220.1H384v-14.8H281.1v14.8zm0-37.1H384v-12.4H281.1V183zm0 74.2H384v-17.3H281.1v17.3zm-157.7 86.7H8.5c2.6 8.5 5.8 16.8 9.6 24.8h138.3c-12.9-5.7-24.1-14.2-33-24.8m145.7-12.4h109.7c1.8-7.3 3.1-14.7 3.9-22.3H278.3c-2.1 7.9-5.2 15.4-9.2 22.3m-41.5 37.1H367c3.7-8 5.8-16.2 8.5-24.8h-115c-8.8 10.7-20.1 19.2-32.9 24.8M384 32H281.1v2.5H384V32zM192 480c39.5 0 76.2-11.8 106.8-32.2H85.3C115.8 468.2 152.5 480 192 480m89.1-334.2H384V136H281.1v9.8zm0-37.1H384v-7.4H281.1v7.4zm0-37.1H384v-4.9H281.1v4.9zm-178.2 99H0V183h102.9v-12.4zM38.8 405.7h305.3c6.7-8.5 12.6-17.6 17.8-27.2H23c5.2 9.6 9.2 18.7 15.8 27.2m64.1-118.8v-12.4H0v12.4c0 2.5 0 5 .1 7.4h103.1c-.2-2.4-.3-4.9-.3-7.4m178.2 0c0 2.5-.1 5-.4 7.4h103.1c.1-2.5.2-4.9.2-7.4v-12.4H281.1v12.4zm-203 156h227.7c11.8-8.7 22.7-18.6 32.2-29.7H44.9c9.6 11 21.4 21 33.2 29.7m24.8-376.2H0v4.9h102.9v-4.9zm0-34.7H0v2.5h102.9V32zm0 173.3H0v14.8h102.9v-14.8zm0 34.6H0v17.3h102.9v-17.3zm0-103.9H0v9.9h102.9V136zm0-34.7H0v7.4h102.9v-7.4zm2.8 207.9H1.3c.9 7.6 2.2 15 3.9 22.3h109.7c-4-6.9-7.2-14.4-9.2-22.3"],untappd:[640,512,[],"f405","M401.3 49.9c-79.8 160.1-84.6 152.5-87.9 173.2l-5.2 32.8c-1.9 12-6.6 23.5-13.7 33.4L145.6 497.1c-7.6 10.6-20.4 16.2-33.4 14.6-40.3-5-77.8-32.2-95.3-68.5-5.7-11.8-4.5-25.8 3.1-36.4l148.9-207.9c7.1-9.9 16.4-18 27.2-23.7l29.3-15.5c18.5-9.8 9.7-11.9 135.6-138.9 1-4.8 1-7.3 3.6-8 3-.7 6.6-1 6.3-4.6l-.4-4.6c-.2-1.9 1.3-3.6 3.2-3.6 4.5-.1 13.2 1.2 25.6 10 12.3 8.9 16.4 16.8 17.7 21.1.6 1.8-.6 3.7-2.4 4.2l-4.5 1.1c-3.4.9-2.5 4.4-2.3 7.4.1 2.8-2.3 3.6-6.5 6.1zM230.1 36.4c3.4.9 2.5 4.4 2.3 7.4-.2 2.7 2.1 3.5 6.4 6 7.9 15.9 15.3 30.5 22.2 44 .7 1.3 2.3 1.5 3.3.5 11.2-12 24.6-26.2 40.5-42.6 1.3-1.4 1.4-3.5.1-4.9-8-8.2-16.5-16.9-25.6-26.1-1-4.7-1-7.3-3.6-8-3-.8-6.6-1-6.3-4.6.3-3.3 1.4-8.1-2.8-8.2-4.5-.1-13.2 1.1-25.6 10-12.3 8.9-16.4 16.8-17.7 21.1-1.4 4.2 3.6 4.6 6.8 5.4zM620 406.7L471.2 198.8c-13.2-18.5-26.6-23.4-56.4-39.1-11.2-5.9-14.2-10.9-30.5-28.9-1-1.1-2.9-.9-3.6.5-46.3 88.8-47.1 82.8-49 94.8-1.7 10.7-1.3 20 .3 29.8 1.9 12 6.6 23.5 13.7 33.4l148.9 207.9c7.6 10.6 20.2 16.2 33.1 14.7 40.3-4.9 78-32 95.7-68.6 5.4-11.9 4.3-25.9-3.4-36.6z"],usb:[640,512,[],"f287","M641.5 256c0 3.1-1.7 6.1-4.5 7.5L547.9 317c-1.4.8-2.8 1.4-4.5 1.4-1.4 0-3.1-.3-4.5-1.1-2.8-1.7-4.5-4.5-4.5-7.8v-35.6H295.7c25.3 39.6 40.5 106.9 69.6 106.9H392V354c0-5 3.9-8.9 8.9-8.9H490c5 0 8.9 3.9 8.9 8.9v89.1c0 5-3.9 8.9-8.9 8.9h-89.1c-5 0-8.9-3.9-8.9-8.9v-26.7h-26.7c-75.4 0-81.1-142.5-124.7-142.5H140.3c-8.1 30.6-35.9 53.5-69 53.5C32 327.3 0 295.3 0 256s32-71.3 71.3-71.3c33.1 0 61 22.8 69 53.5 39.1 0 43.9 9.5 74.6-60.4C255 88.7 273 95.7 323.8 95.7c7.5-20.9 27-35.6 50.4-35.6 29.5 0 53.5 23.9 53.5 53.5s-23.9 53.5-53.5 53.5c-23.4 0-42.9-14.8-50.4-35.6H294c-29.1 0-44.3 67.4-69.6 106.9h310.1v-35.6c0-3.3 1.7-6.1 4.5-7.8 2.8-1.7 6.4-1.4 8.9.3l89.1 53.5c2.8 1.1 4.5 4.1 4.5 7.2z"],ussunnah:[512,512,[],"f407","M156.8 285.1l5.7 14.4h-8.2c-1.3-3.2-3.1-7.7-3.8-9.5-2.5-6.3-1.1-8.4 0-10 1.9-2.7 3.2-4.4 3.6-5.2 0 2.2.8 5.7 2.7 10.3zm297.3 18.8c-2.1 13.8-5.7 27.1-10.5 39.7l43 23.4-44.8-18.8c-5.3 13.2-12 25.6-19.9 37.2l34.2 30.2-36.8-26.4c-8.4 11.8-18 22.6-28.7 32.3l24.9 34.7-28.1-31.8c-11 9.6-23.1 18-36.1 25.1l15.7 37.2-19.3-35.3c-13.1 6.8-27 12.1-41.6 15.9l6.7 38.4-10.5-37.4c-14.3 3.4-29.2 5.3-44.5 5.4L256 512l-1.9-38.4c-15.3-.1-30.2-2-44.5-5.3L199 505.6l6.7-38.2c-14.6-3.7-28.6-9.1-41.7-15.8l-19.2 35.1 15.6-37c-13-7-25.2-15.4-36.2-25.1l-27.9 31.6 24.7-34.4c-10.7-9.7-20.4-20.5-28.8-32.3l-36.5 26.2 33.9-29.9c-7.9-11.6-14.6-24.1-20-37.3l-44.4 18.7L67.8 344c-4.8-12.7-8.4-26.1-10.5-39.9l-51 9 50.3-14.2c-1.1-8.5-1.7-17.1-1.7-25.9 0-4.7.2-9.4.5-14.1L0 256l56-2.8c1.3-13.1 3.8-25.8 7.5-38.1L6.4 199l58.9 10.4c4-12 9.1-23.5 15.2-34.4l-55.1-30 58.3 24.6C90 159 97.2 149.2 105.3 140L55.8 96.4l53.9 38.7c8.1-8.6 17-16.5 26.6-23.6l-40-55.6 45.6 51.6c9.5-6.6 19.7-12.3 30.3-17.2l-27.3-64.9 33.8 62.1c10.5-4.4 21.4-7.9 32.7-10.4L199 6.4l19.5 69.2c11-2.1 22.3-3.2 33.8-3.4L256 0l3.6 72.2c11.5.2 22.8 1.4 33.8 3.5L313 6.4l-12.4 70.7c11.3 2.6 22.2 6.1 32.6 10.5l33.9-62.2-27.4 65.1c10.6 4.9 20.7 10.7 30.2 17.2l45.8-51.8-40.1 55.9c9.5 7.1 18.4 15 26.5 23.6l54.2-38.9-49.7 43.9c8 9.1 15.2 18.9 21.5 29.4l58.7-24.7-55.5 30.2c6.1 10.9 11.1 22.3 15.1 34.3l59.3-10.4-57.5 16.2c3.7 12.2 6.2 24.9 7.5 37.9L512 256l-56 2.8c.3 4.6.5 9.3.5 14.1 0 8.7-.6 17.3-1.6 25.8l50.7 14.3-51.5-9.1zm-21.8-31c0-97.5-79-176.5-176.5-176.5s-176.5 79-176.5 176.5 79 176.5 176.5 176.5 176.5-79 176.5-176.5zm-24 0c0 84.3-68.3 152.6-152.6 152.6s-152.6-68.3-152.6-152.6 68.3-152.6 152.6-152.6 152.6 68.3 152.6 152.6zM195 241c0 2.1 1.3 3.8 3.6 5.1 3.3 1.9 6.2 4.6 8.2 8.2 2.8-5.7 4.3-9.5 4.3-11.2 0-2.2-1.1-4.4-3.2-7-2.1-2.5-3.2-5.2-3.3-7.7-6.5 6.8-9.6 10.9-9.6 12.6zm-40.7-19c0 2.1 1.3 3.8 3.6 5.1 3.5 1.9 6.2 4.6 8.2 8.2 2.8-5.7 4.3-9.5 4.3-11.2 0-2.2-1.1-4.4-3.2-7-2.1-2.5-3.2-5.2-3.3-7.7-6.5 6.8-9.6 10.9-9.6 12.6zm-19 0c0 2.1 1.3 3.8 3.6 5.1 3.3 1.9 6.2 4.6 8.2 8.2 2.8-5.7 4.3-9.5 4.3-11.2 0-2.2-1.1-4.4-3.2-7-2.1-2.5-3.2-5.2-3.3-7.7-6.4 6.8-9.6 10.9-9.6 12.6zm204.9 87.9c-8.4-3-8.7-6.8-8.7-15.6V182c-8.2 12.5-14.2 18.6-18 18.6 6.3 14.4 9.5 23.9 9.5 28.3v64.3c0 2.2-2.2 6.5-4.7 6.5h-18c-2.8-7.5-10.2-26.9-15.3-40.3-2 2.5-7.2 9.2-10.7 13.7 2.4 1.6 4.1 3.6 5.2 6.3 2.6 6.7 6.4 16.5 7.9 20.2h-9.2c-3.9-10.4-9.6-25.4-11.8-31.1-2 2.5-7.2 9.2-10.7 13.7 2.4 1.6 4.1 3.6 5.2 6.3.8 2 2.8 7.3 4.3 10.9H256c-1.5-4.1-5.6-14.6-8.4-22-2 2.5-7.2 9.2-10.7 13.7 2.5 1.6 4.3 3.6 5.2 6.3.2.6.5 1.4.6 1.7H225c-4.6-13.9-11.4-27.7-11.4-34.1 0-2.2.3-5.1 1.1-8.2-8.8 10.8-14 15.9-14 25 0 7.5 10.4 28.3 10.4 33.3 0 1.7-.5 3.3-1.4 4.9-9.6-12.7-15.5-20.7-18.8-20.7h-12l-11.2-28c-3.8-9.6-5.7-16-5.7-18.8 0-3.8.5-7.7 1.7-12.2-1 1.3-3.7 4.7-5.5 7.1-.8-2.1-3.1-7.7-4.6-11.5-2.1 2.5-7.5 9.1-11.2 13.6.9 2.3 3.3 8.1 4.9 12.2-2.5 3.3-9.1 11.8-13.6 17.7-4 5.3-5.8 13.3-2.7 21.8 2.5 6.7 2 7.9-1.7 14.1H191c5.5 0 14.3 14 15.5 22 13.2-16 15.4-19.6 16.8-21.6h107c3.9 0 7.2-1.9 9.9-5.8zm20.1-26.6V181.7c-9 12.5-15.9 18.6-20.7 18.6 7.1 14.4 10.7 23.9 10.7 28.3v66.3c0 17.5 8.6 20.4 24 20.4 8.1 0 12.5-.8 13.7-2.7-4.3-1.6-7.6-2.5-9.9-3.3-8.1-3.2-17.8-7.4-17.8-26z"],vaadin:[448,512,[],"f408","M224.5 140.7c1.5-17.6 4.9-52.7 49.8-52.7h98.6c20.7 0 32.1-7.8 32.1-21.6V54.1c0-12.2 9.3-22.1 21.5-22.1S448 41.9 448 54.1v36.5c0 42.9-21.5 62-66.8 62H280.7c-30.1 0-33 14.7-33 27.1 0 1.3-.1 2.5-.2 3.7-.7 12.3-10.9 22.2-23.4 22.2s-22.7-9.8-23.4-22.2c-.1-1.2-.2-2.4-.2-3.7 0-12.3-3-27.1-33-27.1H66.8c-45.3 0-66.8-19.1-66.8-62V54.1C0 41.9 9.4 32 21.6 32s21.5 9.9 21.5 22.1v12.3C43.1 80.2 54.5 88 75.2 88h98.6c44.8 0 48.3 35.1 49.8 52.7h.9zM224 456c11.5 0 21.4-7 25.7-16.3 1.1-1.8 97.1-169.6 98.2-171.4 11.9-19.6-3.2-44.3-27.2-44.3-13.9 0-23.3 6.4-29.8 20.3L224 362l-66.9-117.7c-6.4-13.9-15.9-20.3-29.8-20.3-24 0-39.1 24.6-27.2 44.3 1.1 1.9 97.1 169.6 98.2 171.4 4.3 9.3 14.2 16.3 25.7 16.3z"],viacoin:[384,512,[],"f237","M384 32h-64l-80.7 192h-94.5L64 32H0l48 112H0v48h68.5l13.8 32H0v48h102.8L192 480l89.2-208H384v-48h-82.3l13.8-32H384v-48h-48l48-112zM192 336l-27-64h54l-27 64z"],viadeo:[448,512,[],"f2a9","M276.2 150.5v.7C258.3 98.6 233.6 47.8 205.4 0c43.3 29.2 67 100 70.8 150.5zm32.7 121.7c7.6 18.2 11 37.5 11 57 0 77.7-57.8 141-137.8 139.4l3.8-.3c74.2-46.7 109.3-118.6 109.3-205.1 0-38.1-6.5-75.9-18.9-112 1 11.7 1 23.7 1 35.4 0 91.8-18.1 241.6-116.6 280C95 455.2 49.4 398 49.4 329.2c0-75.6 57.4-142.3 135.4-142.3 16.8 0 33.7 3.1 49.1 9.6 1.7-15.1 6.5-29.9 13.4-43.3-19.9-7.2-41.2-10.7-62.5-10.7-161.5 0-238.7 195.9-129.9 313.7 67.9 74.6 192 73.9 259.8 0 56.6-61.3 60.9-142.4 36.4-201-12.7 8-27.1 13.9-42.2 17zM418.1 11.7c-31 66.5-81.3 47.2-115.8 80.1-12.4 12-20.6 34-20.6 50.5 0 14.1 4.5 27.1 12 38.8 47.4-11 98.3-46 118.2-90.7-.7 5.5-4.8 14.4-7.2 19.2-20.3 35.7-64.6 65.6-99.7 84.9 14.8 14.4 33.7 25.8 55 25.8 79 0 110.1-134.6 58.1-208.6z"],"viadeo-square":[448,512,[],"f2aa","M400 32H48C21.5 32 0 53.5 0 80v352c0 26.5 21.5 48 48 48h352c26.5 0 48-21.5 48-48V80c0-26.5-21.5-48-48-48zM280.7 381.2c-42.4 46.2-120 46.6-162.4 0-68-73.6-19.8-196.1 81.2-196.1 13.3 0 26.6 2.1 39.1 6.7-4.3 8.4-7.3 17.6-8.4 27.1-9.7-4.1-20.2-6-30.7-6-48.8 0-84.6 41.7-84.6 88.9 0 43 28.5 78.7 69.5 85.9 61.5-24 72.9-117.6 72.9-175 0-7.3 0-14.8-.6-22.1-11.2-32.9-26.6-64.6-44.2-94.5 27.1 18.3 41.9 62.5 44.2 94.1v.4c7.7 22.5 11.8 46.2 11.8 70 0 54.1-21.9 99-68.3 128.2l-2.4.2c50 1 86.2-38.6 86.2-87.2 0-12.2-2.1-24.3-6.9-35.7 9.5-1.9 18.5-5.6 26.4-10.5 15.3 36.6 12.6 87.3-22.8 125.6zM309 233.7c-13.3 0-25.1-7.1-34.4-16.1 21.9-12 49.6-30.7 62.3-53 1.5-3 4.1-8.6 4.5-12-12.5 27.9-44.2 49.8-73.9 56.7-4.7-7.3-7.5-15.5-7.5-24.3 0-10.3 5.2-24.1 12.9-31.6 21.6-20.5 53-8.5 72.4-50 32.5 46.2 13.1 130.3-36.3 130.3z"],viber:[512,512,[],"f409","M444 49.9C431.3 38.2 379.9.9 265.3.4c0 0-135.1-8.1-200.9 52.3C27.8 89.3 14.9 143 13.5 209.5c-1.4 66.5-3.1 191.1 117 224.9h.1l-.1 51.6s-.8 20.9 13 25.1c16.6 5.2 26.4-10.7 42.3-27.8 8.7-9.4 20.7-23.2 29.8-33.7 82.2 6.9 145.3-8.9 152.5-11.2 16.6-5.4 110.5-17.4 125.7-142 15.8-128.6-7.6-209.8-49.8-246.5zM457.9 287c-12.9 104-89 110.6-103 115.1-6 1.9-61.5 15.7-131.2 11.2 0 0-52 62.7-68.2 79-5.3 5.3-11.1 4.8-11-5.7 0-6.9.4-85.7.4-85.7-.1 0-.1 0 0 0-101.8-28.2-95.8-134.3-94.7-189.8 1.1-55.5 11.6-101 42.6-131.6 55.7-50.5 170.4-43 170.4-43 96.9.4 143.3 29.6 154.1 39.4 35.7 30.6 53.9 103.8 40.6 211.1zm-139-80.8c.4 8.6-12.5 9.2-12.9.6-1.1-22-11.4-32.7-32.6-33.9-8.6-.5-7.8-13.4.7-12.9 27.9 1.5 43.4 17.5 44.8 46.2zm20.3 11.3c1-42.4-25.5-75.6-75.8-79.3-8.5-.6-7.6-13.5.9-12.9 58 4.2 88.9 44.1 87.8 92.5-.1 8.6-13.1 8.2-12.9-.3zm47 13.4c.1 8.6-12.9 8.7-12.9.1-.6-81.5-54.9-125.9-120.8-126.4-8.5-.1-8.5-12.9 0-12.9 73.7.5 133 51.4 133.7 139.2zM374.9 329v.2c-10.8 19-31 40-51.8 33.3l-.2-.3c-21.1-5.9-70.8-31.5-102.2-56.5-16.2-12.8-31-27.9-42.4-42.4-10.3-12.9-20.7-28.2-30.8-46.6-21.3-38.5-26-55.7-26-55.7-6.7-20.8 14.2-41 33.3-51.8h.2c9.2-4.8 18-3.2 23.9 3.9 0 0 12.4 14.8 17.7 22.1 5 6.8 11.7 17.7 15.2 23.8 6.1 10.9 2.3 22-3.7 26.6l-12 9.6c-6.1 4.9-5.3 14-5.3 14s17.8 67.3 84.3 84.3c0 0 9.1.8 14-5.3l9.6-12c4.6-6 15.7-9.8 26.6-3.7 14.7 8.3 33.4 21.2 45.8 32.9 7 5.7 8.6 14.4 3.8 23.6z"],vimeo:[448,512,[],"f40a","M403.2 32H44.8C20.1 32 0 52.1 0 76.8v358.4C0 459.9 20.1 480 44.8 480h358.4c24.7 0 44.8-20.1 44.8-44.8V76.8c0-24.7-20.1-44.8-44.8-44.8zM377 180.8c-1.4 31.5-23.4 74.7-66 129.4-44 57.2-81.3 85.8-111.7 85.8-18.9 0-34.8-17.4-47.9-52.3-25.5-93.3-36.4-148-57.4-148-2.4 0-10.9 5.1-25.4 15.2l-15.2-19.6c37.3-32.8 72.9-69.2 95.2-71.2 25.2-2.4 40.7 14.8 46.5 51.7 20.7 131.2 29.9 151 67.6 91.6 13.5-21.4 20.8-37.7 21.8-48.9 3.5-33.2-25.9-30.9-45.8-22.4 15.9-52.1 46.3-77.4 91.2-76 33.3.9 49 22.5 47.1 64.7z"],"vimeo-square":[448,512,[],"f194","M400 32H48C21.5 32 0 53.5 0 80v352c0 26.5 21.5 48 48 48h352c26.5 0 48-21.5 48-48V80c0-26.5-21.5-48-48-48zm-16.2 149.6c-1.4 31.1-23.2 73.8-65.3 127.9-43.5 56.5-80.3 84.8-110.4 84.8-18.7 0-34.4-17.2-47.3-51.6-25.2-92.3-35.9-146.4-56.7-146.4-2.4 0-10.8 5-25.1 15.1L64 192c36.9-32.4 72.1-68.4 94.1-70.4 24.9-2.4 40.2 14.6 46 51.1 20.5 129.6 29.6 149.2 66.8 90.5 13.4-21.2 20.6-37.2 21.5-48.3 3.4-32.8-25.6-30.6-45.2-22.2 15.7-51.5 45.8-76.5 90.1-75.1 32.9 1 48.4 22.4 46.5 64z"],"vimeo-v":[448,512,[],"f27d","M447.8 153.6c-2 43.6-32.4 103.3-91.4 179.1-60.9 79.2-112.4 118.8-154.6 118.8-26.1 0-48.2-24.1-66.3-72.3C100.3 250 85.3 174.3 56.2 174.3c-3.4 0-15.1 7.1-35.2 21.1L0 168.2c51.6-45.3 100.9-95.7 131.8-98.5 34.9-3.4 56.3 20.5 64.4 71.5 28.7 181.5 41.4 208.9 93.6 126.7 18.7-29.6 28.8-52.1 30.2-67.6 4.8-45.9-35.8-42.8-63.3-31 22-72.1 64.1-107.1 126.2-105.1 45.8 1.2 67.5 31.1 64.9 89.4z"],vine:[384,512,[],"f1ca","M384 254.7v52.1c-18.4 4.2-36.9 6.1-52.1 6.1-36.9 77.4-103 143.8-125.1 156.2-14 7.9-27.1 8.4-42.7-.8C137 452 34.2 367.7 0 102.7h74.5C93.2 261.8 139 343.4 189.3 404.5c27.9-27.9 54.8-65.1 75.6-106.9-49.8-25.3-80.1-80.9-80.1-145.6 0-65.6 37.7-115.1 102.2-115.1 114.9 0 106.2 127.9 81.6 181.5 0 0-46.4 9.2-63.5-20.5 3.4-11.3 8.2-30.8 8.2-48.5 0-31.3-11.3-46.6-28.4-46.6-18.2 0-30.8 17.1-30.8 50 .1 79.2 59.4 118.7 129.9 101.9z"],vk:[576,512,[],"f189","M545 117.7c3.7-12.5 0-21.7-17.8-21.7h-58.9c-15 0-21.9 7.9-25.6 16.7 0 0-30 73.1-72.4 120.5-13.7 13.7-20 18.1-27.5 18.1-3.7 0-9.4-4.4-9.4-16.9V117.7c0-15-4.2-21.7-16.6-21.7h-92.6c-9.4 0-15 7-15 13.5 0 14.2 21.2 17.5 23.4 57.5v86.8c0 19-3.4 22.5-10.9 22.5-20 0-68.6-73.4-97.4-157.4-5.8-16.3-11.5-22.9-26.6-22.9H38.8c-16.8 0-20.2 7.9-20.2 16.7 0 15.6 20 93.1 93.1 195.5C160.4 378.1 229 416 291.4 416c37.5 0 42.1-8.4 42.1-22.9 0-66.8-3.4-73.1 15.4-73.1 8.7 0 23.7 4.4 58.7 38.1 40 40 46.6 57.9 69 57.9h58.9c16.8 0 25.3-8.4 20.4-25-11.2-34.9-86.9-106.7-90.3-111.5-8.7-11.2-6.2-16.2 0-26.2.1-.1 72-101.3 79.4-135.6z"],vnv:[640,512,[],"f40b","M104.9 352c-34.1 0-46.4-30.4-46.4-30.4L2.6 210.1S-7.8 192 13 192h32.8c10.4 0 13.2 8.7 18.8 18.1l36.7 74.5s5.2 13.1 21.1 13.1 21.1-13.1 21.1-13.1l36.7-74.5c5.6-9.5 8.4-18.1 18.8-18.1h32.8c20.8 0 10.4 18.1 10.4 18.1l-55.8 111.5S174.2 352 140 352h-35.1zm395 0c-34.1 0-46.4-30.4-46.4-30.4l-55.9-111.5S387.2 192 408 192h32.8c10.4 0 13.2 8.7 18.8 18.1l36.7 74.5s5.2 13.1 21.1 13.1 21.1-13.1 21.1-13.1l36.8-74.5c5.6-9.5 8.4-18.1 18.8-18.1H627c20.8 0 10.4 18.1 10.4 18.1l-55.9 111.5S569.3 352 535.1 352h-35.2zM337.6 192c34.1 0 46.4 30.4 46.4 30.4l55.9 111.5s10.4 18.1-10.4 18.1h-32.8c-10.4 0-13.2-8.7-18.8-18.1l-36.7-74.5s-5.2-13.1-21.1-13.1c-15.9 0-21.1 13.1-21.1 13.1l-36.7 74.5c-5.6 9.4-8.4 18.1-18.8 18.1h-32.9c-20.8 0-10.4-18.1-10.4-18.1l55.9-111.5s12.2-30.4 46.4-30.4h35.1z"],vuejs:[448,512,[],"f41f","M356.9 64.3H280l-56 88.6-48-88.6H0L224 448 448 64.3h-91.1zm-301.2 32h53.8L224 294.5 338.4 96.3h53.8L224 384.5 55.7 96.3z"],weibo:[512,512,[],"f18a","M407 177.6c7.6-24-13.4-46.8-37.4-41.7-22 4.8-28.8-28.1-7.1-32.8 50.1-10.9 92.3 37.1 76.5 84.8-6.8 21.2-38.8 10.8-32-10.3zM214.8 446.7C108.5 446.7 0 395.3 0 310.4c0-44.3 28-95.4 76.3-143.7C176 67 279.5 65.8 249.9 161c-4 13.1 12.3 5.7 12.3 6 79.5-33.6 140.5-16.8 114 51.4-3.7 9.4 1.1 10.9 8.3 13.1 135.7 42.3 34.8 215.2-169.7 215.2zm143.7-146.3c-5.4-55.7-78.5-94-163.4-85.7-84.8 8.6-148.8 60.3-143.4 116s78.5 94 163.4 85.7c84.8-8.6 148.8-60.3 143.4-116zM347.9 35.1c-25.9 5.6-16.8 43.7 8.3 38.3 72.3-15.2 134.8 52.8 111.7 124-7.4 24.2 29.1 37 37.4 12 31.9-99.8-55.1-195.9-157.4-174.3zm-78.5 311c-17.1 38.8-66.8 60-109.1 46.3-40.8-13.1-58-53.4-40.3-89.7 17.7-35.4 63.1-55.4 103.4-45.1 42 10.8 63.1 50.2 46 88.5zm-86.3-30c-12.9-5.4-30 .3-38 12.9-8.3 12.9-4.3 28 8.6 34 13.1 6 30.8.3 39.1-12.9 8-13.1 3.7-28.3-9.7-34zm32.6-13.4c-5.1-1.7-11.4.6-14.3 5.4-2.9 5.1-1.4 10.6 3.7 12.9 5.1 2 11.7-.3 14.6-5.4 2.8-5.2 1.1-10.9-4-12.9z"],weixin:[576,512,[],"f1d7","M385.2 167.6c6.4 0 12.6.3 18.8 1.1C387.4 90.3 303.3 32 207.7 32 100.5 32 13 104.8 13 197.4c0 53.4 29.3 97.5 77.9 131.6l-19.3 58.6 68-34.1c24.4 4.8 43.8 9.7 68.2 9.7 6.2 0 12.1-.3 18.3-.8-4-12.9-6.2-26.6-6.2-40.8-.1-84.9 72.9-154 165.3-154zm-104.5-52.9c14.5 0 24.2 9.7 24.2 24.4 0 14.5-9.7 24.2-24.2 24.2-14.8 0-29.3-9.7-29.3-24.2.1-14.7 14.6-24.4 29.3-24.4zm-136.4 48.6c-14.5 0-29.3-9.7-29.3-24.2 0-14.8 14.8-24.4 29.3-24.4 14.8 0 24.4 9.7 24.4 24.4 0 14.6-9.6 24.2-24.4 24.2zM563 319.4c0-77.9-77.9-141.3-165.4-141.3-92.7 0-165.4 63.4-165.4 141.3S305 460.7 397.6 460.7c19.3 0 38.9-5.1 58.6-9.9l53.4 29.3-14.8-48.6C534 402.1 563 363.2 563 319.4zm-219.1-24.5c-9.7 0-19.3-9.7-19.3-19.6 0-9.7 9.7-19.3 19.3-19.3 14.8 0 24.4 9.7 24.4 19.3 0 10-9.7 19.6-24.4 19.6zm107.1 0c-9.7 0-19.3-9.7-19.3-19.6 0-9.7 9.7-19.3 19.3-19.3 14.5 0 24.4 9.7 24.4 19.3.1 10-9.9 19.6-24.4 19.6z"],whatsapp:[448,512,[],"f232","M380.9 97.1C339 55.1 283.2 32 223.9 32c-122.4 0-222 99.6-222 222 0 39.1 10.2 77.3 29.6 111L0 480l117.7-30.9c32.4 17.7 68.9 27 106.1 27h.1c122.3 0 224.1-99.6 224.1-222 0-59.3-25.2-115-67.1-157zm-157 341.6c-33.2 0-65.7-8.9-94-25.7l-6.7-4-69.8 18.3L72 359.2l-4.4-7c-18.5-29.4-28.2-63.3-28.2-98.2 0-101.7 82.8-184.5 184.6-184.5 49.3 0 95.6 19.2 130.4 54.1 34.8 34.9 56.2 81.2 56.1 130.5 0 101.8-84.9 184.6-186.6 184.6zm101.2-138.2c-5.5-2.8-32.8-16.2-37.9-18-5.1-1.9-8.8-2.8-12.5 2.8-3.7 5.6-14.3 18-17.6 21.8-3.2 3.7-6.5 4.2-12 1.4-32.6-16.3-54-29.1-75.5-66-5.7-9.8 5.7-9.1 16.3-30.3 1.8-3.7.9-6.9-.5-9.7-1.4-2.8-12.5-30.1-17.1-41.2-4.5-10.8-9.1-9.3-12.5-9.5-3.2-.2-6.9-.2-10.6-.2-3.7 0-9.7 1.4-14.8 6.9-5.1 5.6-19.4 19-19.4 46.3 0 27.3 19.9 53.7 22.6 57.4 2.8 3.7 39.1 59.7 94.8 83.8 35.2 15.2 49 16.5 66.6 13.9 10.7-1.6 32.8-13.4 37.4-26.4 4.6-13 4.6-24.1 3.2-26.4-1.3-2.5-5-3.9-10.5-6.6z"],"whatsapp-square":[448,512,[],"f40c","M224 122.8c-72.7 0-131.8 59.1-131.9 131.8 0 24.9 7 49.2 20.2 70.1l3.1 5-13.3 48.6 49.9-13.1 4.8 2.9c20.2 12 43.4 18.4 67.1 18.4h.1c72.6 0 133.3-59.1 133.3-131.8 0-35.2-15.2-68.3-40.1-93.2-25-25-58-38.7-93.2-38.7zm77.5 188.4c-3.3 9.3-19.1 17.7-26.7 18.8-12.6 1.9-22.4.9-47.5-9.9-39.7-17.2-65.7-57.2-67.7-59.8-2-2.6-16.2-21.5-16.2-41s10.2-29.1 13.9-33.1c3.6-4 7.9-5 10.6-5 2.6 0 5.3 0 7.6.1 2.4.1 5.7-.9 8.9 6.8 3.3 7.9 11.2 27.4 12.2 29.4s1.7 4.3.3 6.9c-7.6 15.2-15.7 14.6-11.6 21.6 15.3 26.3 30.6 35.4 53.9 47.1 4 2 6.3 1.7 8.6-1 2.3-2.6 9.9-11.6 12.5-15.5 2.6-4 5.3-3.3 8.9-2 3.6 1.3 23.1 10.9 27.1 12.9s6.6 3 7.6 4.6c.9 1.9.9 9.9-2.4 19.1zM400 32H48C21.5 32 0 53.5 0 80v352c0 26.5 21.5 48 48 48h352c26.5 0 48-21.5 48-48V80c0-26.5-21.5-48-48-48zM223.9 413.2c-26.6 0-52.7-6.7-75.8-19.3L64 416l22.5-82.2c-13.9-24-21.2-51.3-21.2-79.3C65.4 167.1 136.5 96 223.9 96c42.4 0 82.2 16.5 112.2 46.5 29.9 30 47.9 69.8 47.9 112.2 0 87.4-72.7 158.5-160.1 158.5z"],whmcs:[448,512,[],"f40d","M448 161v-21.3l-28.5-8.8-2.2-10.4 20.1-20.7L427 80.4l-29 7.5-7.2-7.5 7.5-28.2-19.1-11.6-21.3 21-10.7-3.2-7-26.4h-22.6l-6.2 26.4-12.1 3.2-19.7-21-19.4 11 8.1 27.7-8.1 8.4-28.5-7.5-11 19.1 20.7 21-2.9 10.4-28.5 7.8-.3 21.7 28.8 7.5 2.4 12.1-20.1 19.9 10.4 18.5 29.6-7.5 7.2 8.6-8.1 26.9 19.9 11.6 19.4-20.4 11.6 2.9 6.7 28.5 22.6.3 6.7-28.8 11.6-3.5 20.7 21.6 20.4-12.1-8.8-28 7.8-8.1 28.8 8.8 10.3-20.1-20.9-18.8 2.2-12.1 29.1-7zm-119.2 45.2c-31.3 0-56.8-25.4-56.8-56.8s25.4-56.8 56.8-56.8 56.8 25.4 56.8 56.8c0 31.5-25.4 56.8-56.8 56.8zm72.3 16.4l46.9 14.5V277l-55.1 13.4-4.1 22.7 38.9 35.3-19.2 37.9-54-16.7-14.6 15.2 16.7 52.5-38.3 22.7-38.9-40.5-21.7 6.6-12.6 54-42.4-.5-12.6-53.6-21.7-5.6-36.4 38.4-37.4-21.7 15.2-50.5-13.7-16.1-55.5 14.1-19.7-34.8 37.9-37.4-4.8-22.8-54-14.1.5-40.9L54 219.9l5.7-19.7-38.9-39.4L41.5 125l53.6 14.1 15.2-15.7-15.2-52 36.4-20.7 36.8 39.4L191 84l11.6-52H245l11.6 45.9L234 72l-6.3-1.7-3.3 5.7-11 19.1-3.3 5.6 4.6 4.6 17.2 17.4-.3 1-23.8 6.5-6.2 1.7-.1 6.4-.2 12.9C153.8 161.6 118 204 118 254.7c0 58.3 47.3 105.7 105.7 105.7 50.5 0 92.7-35.4 103.2-82.8l13.2.2 6.9.1 1.6-6.7 5.6-24 1.9-.6 17.1 17.8 4.7 4.9 5.8-3.4 20.4-12.1 5.8-3.5-2-6.5-6.8-21.2z"],"wikipedia-w":[640,512,[],"f266","M640 51.2l-.3 12.2c-28.1.8-45 15.8-55.8 40.3-25 57.8-103.3 240-155.3 358.6H415l-81.9-193.1c-32.5 63.6-68.3 130-99.2 193.1-.3.3-15 0-15-.3C172 352.3 122.8 243.4 75.8 133.4 64.4 106.7 26.4 63.4.2 63.7c0-3.1-.3-10-.3-14.2h161.9v13.9c-19.2 1.1-52.8 13.3-43.3 34.2 21.9 49.7 103.6 240.3 125.6 288.6 15-29.7 57.8-109.2 75.3-142.8-13.9-28.3-58.6-133.9-72.8-160-9.7-17.8-36.1-19.4-55.8-19.7V49.8l142.5.3v13.1c-19.4.6-38.1 7.8-29.4 26.1 18.9 40 30.6 68.1 48.1 104.7 5.6-10.8 34.7-69.4 48.1-100.8 8.9-20.6-3.9-28.6-38.6-29.4.3-3.6 0-10.3.3-13.6 44.4-.3 111.1-.3 123.1-.6v13.6c-22.5.8-45.8 12.8-58.1 31.7l-59.2 122.8c6.4 16.1 63.3 142.8 69.2 156.7L559.2 91.8c-8.6-23.1-36.4-28.1-47.2-28.3V49.6l127.8 1.1.2.5z"],windows:[448,512,[],"f17a","M0 93.7l183.6-25.3v177.4H0V93.7zm0 324.6l183.6 25.3V268.4H0v149.9zm203.8 28L448 480V268.4H203.8v177.9zm0-380.6v180.1H448V32L203.8 65.7z"],"wolf-pack-battalion":[456,512,[],"f514","M239.73 471.53l10.56 15.84 5.28-12.32 5.28 7.04V512c21.06-7.92 21.11-66.86 25.51-97.21 4.62-31.89-.88-92.81 81.37-149.11-8.88-23.61-12-49.43-2.64-80.05 27.87 3.34 53.94 10.58 63.34 54.1l-30.35 8.36c11.15 23.04 17.01 46.76 13.2 72.14L384 313.18l-6.16 33.43-18.47-7.04-8.8 33.43-19.35-7.04 26.39 21.11 8.8-28.15 24.63 5.28 7.04-35.63 26.39 14.52c.25-20.02 6.97-58.06-8.8-84.45l26.39 5.28c3.99-22.07-2.38-39.21-7.92-56.74l22.43 9.68c-.44-25.07-29.94-56.79-61.58-58.5-20.22-1.09-56.74-25.17-54.1-51.9 1.96-19.87 17.45-42.62 43.11-49.7-43.99 36.51-9.68 67.3 5.28 73.46 4.4-11.44 17.54-69.08 0-130.2-40.47 22.87-89.73 65.1-93.25 147.79l-58.06 38.71-3.52 93.25 107.33-59.82 7.04 7.04-17.59 3.52-43.99 38.71-15.84-5.28-28.15 49.26-3.52 119.64 21.11 15.84-32.55 15.84-32.55-15.84 21.11-15.84-3.52-119.64-28.15-49.26-15.84 5.28-43.99-38.71-17.59-3.52 7.04-7.04 107.33 59.82-3.52-93.25-58.06-38.71C157.03 65.1 107.77 22.87 67.3 0c-17.54 61.12-4.4 118.76 0 130.2 14.96-6.16 49.26-36.95 5.28-73.46 25.66 7.08 41.15 29.83 43.11 49.7 2.63 26.74-33.88 50.81-54.1 51.9C29.94 160.06.44 191.78 0 216.85l22.43-9.68c-5.54 17.53-11.91 34.67-7.92 56.74l26.39-5.28c-15.76 26.39-9.05 64.43-8.8 84.45l26.39-14.52 7.04 35.63 24.63-5.28 8.8 28.15 26.39-21.11-19.34 7.05-8.8-33.43-18.47 7.04-6.16-33.43-27.27 7.04c-3.82-25.38 2.05-49.1 13.2-72.14l-30.35-8.36c9.4-43.52 35.47-50.77 63.34-54.1 9.36 30.62 6.24 56.45-2.64 80.05 82.25 56.3 76.75 117.23 81.37 149.11 4.4 30.35 4.45 89.29 25.51 97.21v-29.91l5.28-7.04 5.28 12.32 10.56-15.84 11.44 21.11 11.43-21.1zm79.17-95.01l-15.84-10.56c7.47-4.36 13.76-8.42 19.35-12.32-.6 7.26-.27 13.88-3.51 22.88zm28.15-49.26c-.4 10.94-.9 21.66-1.76 31.67-7.85-1.86-15.57-3.8-21.11-7.04 8.24-7.9 15.55-16.27 22.87-24.63zm24.63 5.28c-.02-13.43-2.05-24.21-5.28-33.43-5.38 9.09-11.23 18.18-18.47 27.27l23.75 6.16zm3.52-80.94c19.44 12.81 27.8 33.66 29.91 56.3-12.32-4.53-24.63-9.31-36.95-10.56 5.06-11.99 6.65-28.14 7.04-45.74zm-1.76-45.74c.81 14.3 1.84 28.82 1.76 42.23 19.22-8.11 29.78-9.72 43.99-14.08-10.6-18.95-27.22-25.52-45.75-28.15zM137.68 376.52l15.84-10.56c-7.47-4.36-13.76-8.42-19.35-12.32.6 7.26.27 13.88 3.51 22.88zm-28.15-49.26c.4 10.94.9 21.66 1.76 31.67 7.85-1.86 15.57-3.8 21.11-7.04-8.24-7.9-15.55-16.27-22.87-24.63zm-24.64 5.28c.02-13.43 2.05-24.21 5.28-33.43 5.38 9.09 11.23 18.18 18.47 27.27l-23.75 6.16zm-3.52-80.94c-19.44 12.81-27.8 33.66-29.91 56.3 12.32-4.53 24.63-9.31 36.95-10.56-5.05-11.99-6.65-28.14-7.04-45.74zm1.76-45.74c-.81 14.3-1.84 28.82-1.76 42.23-19.22-8.11-29.78-9.72-43.99-14.08 10.61-18.95 27.22-25.52 45.75-28.15z"],wordpress:[512,512,[],"f19a","M61.7 169.4l101.5 278C92.2 413 43.3 340.2 43.3 256c0-30.9 6.6-60.1 18.4-86.6zm337.9 75.9c0-26.3-9.4-44.5-17.5-58.7-10.8-17.5-20.9-32.4-20.9-49.9 0-19.6 14.8-37.8 35.7-37.8.9 0 1.8.1 2.8.2-37.9-34.7-88.3-55.9-143.7-55.9-74.3 0-139.7 38.1-177.8 95.9 5 .2 9.7.3 13.7.3 22.2 0 56.7-2.7 56.7-2.7 11.5-.7 12.8 16.2 1.4 17.5 0 0-11.5 1.3-24.3 2l77.5 230.4L249.8 247l-33.1-90.8c-11.5-.7-22.3-2-22.3-2-11.5-.7-10.1-18.2 1.3-17.5 0 0 35.1 2.7 56 2.7 22.2 0 56.7-2.7 56.7-2.7 11.5-.7 12.8 16.2 1.4 17.5 0 0-11.5 1.3-24.3 2l76.9 228.7 21.2-70.9c9-29.4 16-50.5 16-68.7zm-139.9 29.3l-63.8 185.5c19.1 5.6 39.2 8.7 60.1 8.7 24.8 0 48.5-4.3 70.6-12.1-.6-.9-1.1-1.9-1.5-2.9l-65.4-179.2zm183-120.7c.9 6.8 1.4 14 1.4 21.9 0 21.6-4 45.8-16.2 76.2l-65 187.9C426.2 403 468.7 334.5 468.7 256c0-37-9.4-71.8-26-102.1zM504 256c0 136.8-111.3 248-248 248C119.2 504 8 392.7 8 256 8 119.2 119.2 8 256 8c136.7 0 248 111.2 248 248zm-11.4 0c0-130.5-106.2-236.6-236.6-236.6C125.5 19.4 19.4 125.5 19.4 256S125.6 492.6 256 492.6c130.5 0 236.6-106.1 236.6-236.6z"],"wordpress-simple":[512,512,[],"f411","M256 8C119.3 8 8 119.2 8 256c0 136.7 111.3 248 248 248s248-111.3 248-248C504 119.2 392.7 8 256 8zM33 256c0-32.3 6.9-63 19.3-90.7l106.4 291.4C84.3 420.5 33 344.2 33 256zm223 223c-21.9 0-43-3.2-63-9.1l66.9-194.4 68.5 187.8c.5 1.1 1 2.1 1.6 3.1-23.1 8.1-48 12.6-74 12.6zm30.7-327.5c13.4-.7 25.5-2.1 25.5-2.1 12-1.4 10.6-19.1-1.4-18.4 0 0-36.1 2.8-59.4 2.8-21.9 0-58.7-2.8-58.7-2.8-12-.7-13.4 17.7-1.4 18.4 0 0 11.4 1.4 23.4 2.1l34.7 95.2L200.6 393l-81.2-241.5c13.4-.7 25.5-2.1 25.5-2.1 12-1.4 10.6-19.1-1.4-18.4 0 0-36.1 2.8-59.4 2.8-4.2 0-9.1-.1-14.4-.3C109.6 73 178.1 33 256 33c58 0 110.9 22.2 150.6 58.5-1-.1-1.9-.2-2.9-.2-21.9 0-37.4 19.1-37.4 39.6 0 18.4 10.6 33.9 21.9 52.3 8.5 14.8 18.4 33.9 18.4 61.5 0 19.1-7.3 41.2-17 72.1l-22.2 74.3-80.7-239.6zm81.4 297.2l68.1-196.9c12.7-31.8 17-57.2 17-79.9 0-8.2-.5-15.8-1.5-22.9 17.4 31.8 27.3 68.2 27.3 107 0 82.3-44.6 154.1-110.9 192.7z"],wpbeginner:[512,512,[],"f297","M462.799 322.374C519.01 386.682 466.961 480 370.944 480c-39.602 0-78.824-17.687-100.142-50.04-6.887.356-22.702.356-29.59 0C219.848 462.381 180.588 480 141.069 480c-95.49 0-148.348-92.996-91.855-157.626C-29.925 190.523 80.479 32 256.006 32c175.632 0 285.87 158.626 206.793 290.374zm-339.647-82.972h41.529v-58.075h-41.529v58.075zm217.18 86.072v-23.839c-60.506 20.915-132.355 9.198-187.589-33.971l.246 24.897c51.101 46.367 131.746 57.875 187.343 32.913zm-150.753-86.072h166.058v-58.075H189.579v58.075z"],wpexplorer:[512,512,[],"f2de","M512 256c0 141.2-114.7 256-256 256C114.8 512 0 397.3 0 256S114.7 0 256 0s256 114.7 256 256zm-32 0c0-123.2-100.3-224-224-224C132.5 32 32 132.5 32 256s100.5 224 224 224 224-100.5 224-224zM160.9 124.6l86.9 37.1-37.1 86.9-86.9-37.1 37.1-86.9zm110 169.1l46.6 94h-14.6l-50-100-48.9 100h-14l51.1-106.9-22.3-9.4 6-14 68.6 29.1-6 14.3-16.5-7.1zm-11.8-116.3l68.6 29.4-29.4 68.3L230 246l29.1-68.6zm80.3 42.9l54.6 23.1-23.4 54.3-54.3-23.1 23.1-54.3z"],wpforms:[448,512,[],"f298","M448 75.2v361.7c0 24.3-19 43.2-43.2 43.2H43.2C19.3 480 0 461.4 0 436.8V75.2C0 51.1 18.8 32 43.2 32h361.7c24 0 43.1 18.8 43.1 43.2zm-37.3 361.6V75.2c0-3-2.6-5.8-5.8-5.8h-9.3L285.3 144 224 94.1 162.8 144 52.5 69.3h-9.3c-3.2 0-5.8 2.8-5.8 5.8v361.7c0 3 2.6 5.8 5.8 5.8h361.7c3.2.1 5.8-2.7 5.8-5.8zM150.2 186v37H76.7v-37h73.5zm0 74.4v37.3H76.7v-37.3h73.5zm11.1-147.3l54-43.7H96.8l64.5 43.7zm210 72.9v37h-196v-37h196zm0 74.4v37.3h-196v-37.3h196zm-84.6-147.3l64.5-43.7H232.8l53.9 43.7zM371.3 335v37.3h-99.4V335h99.4z"],xbox:[512,512,[],"f412","M369.9 318.2c44.3 54.3 64.7 98.8 54.4 118.7-7.9 15.1-56.7 44.6-92.6 55.9-29.6 9.3-68.4 13.3-100.4 10.2-38.2-3.7-76.9-17.4-110.1-39C93.3 445.8 87 438.3 87 423.4c0-29.9 32.9-82.3 89.2-142.1 32-33.9 76.5-73.7 81.4-72.6 9.4 2.1 84.3 75.1 112.3 109.5zM188.6 143.8c-29.7-26.9-58.1-53.9-86.4-63.4-15.2-5.1-16.3-4.8-28.7 8.1-29.2 30.4-53.5 79.7-60.3 122.4-5.4 34.2-6.1 43.8-4.2 60.5 5.6 50.5 17.3 85.4 40.5 120.9 9.5 14.6 12.1 17.3 9.3 9.9-4.2-11-.3-37.5 9.5-64 14.3-39 53.9-112.9 120.3-194.4zm311.6 63.5C483.3 127.3 432.7 77 425.6 77c-7.3 0-24.2 6.5-36 13.9-23.3 14.5-41 31.4-64.3 52.8C367.7 197 427.5 283.1 448.2 346c6.8 20.7 9.7 41.1 7.4 52.3-1.7 8.5-1.7 8.5 1.4 4.6 6.1-7.7 19.9-31.3 25.4-43.5 7.4-16.2 15-40.2 18.6-58.7 4.3-22.5 3.9-70.8-.8-93.4zM141.3 43C189 40.5 251 77.5 255.6 78.4c.7.1 10.4-4.2 21.6-9.7 63.9-31.1 94-25.8 107.4-25.2-63.9-39.3-152.7-50-233.9-11.7-23.4 11.1-24 11.9-9.4 11.2z"],xing:[384,512,[],"f168","M162.7 210c-1.8 3.3-25.2 44.4-70.1 123.5-4.9 8.3-10.8 12.5-17.7 12.5H9.8c-7.7 0-12.1-7.5-8.5-14.4l69-121.3c.2 0 .2-.1 0-.3l-43.9-75.6c-4.3-7.8.3-14.1 8.5-14.1H100c7.3 0 13.3 4.1 18 12.2l44.7 77.5zM382.6 46.1l-144 253v.3L330.2 466c3.9 7.1.2 14.1-8.5 14.1h-65.2c-7.6 0-13.6-4-18-12.2l-92.4-168.5c3.3-5.8 51.5-90.8 144.8-255.2 4.6-8.1 10.4-12.2 17.5-12.2h65.7c8 0 12.3 6.7 8.5 14.1z"],"xing-square":[448,512,[],"f169","M400 32H48C21.5 32 0 53.5 0 80v352c0 26.5 21.5 48 48 48h352c26.5 0 48-21.5 48-48V80c0-26.5-21.5-48-48-48zM140.4 320.2H93.8c-5.5 0-8.7-5.3-6-10.3l49.3-86.7c.1 0 .1-.1 0-.2l-31.4-54c-3-5.6.2-10.1 6-10.1h46.6c5.2 0 9.5 2.9 12.9 8.7l31.9 55.3c-1.3 2.3-18 31.7-50.1 88.2-3.5 6.2-7.7 9.1-12.6 9.1zm219.7-214.1L257.3 286.8v.2l65.5 119c2.8 5.1.1 10.1-6 10.1h-46.6c-5.5 0-9.7-2.9-12.9-8.7l-66-120.3c2.3-4.1 36.8-64.9 103.4-182.3 3.3-5.8 7.4-8.7 12.5-8.7h46.9c5.7-.1 8.8 4.7 6 10z"],"y-combinator":[448,512,[],"f23b","M448 32v448H0V32h448zM236 287.5L313.5 142h-32.7L235 233c-4.7 9.3-9 18.3-12.8 26.8L210 233l-45.2-91h-35l76.7 143.8v94.5H236v-92.8z"],yahoo:[448,512,[],"f19e","M252 292l4 220c-12.7-2.2-23.5-3.9-32.3-3.9-8.4 0-19.2 1.7-32.3 3.9l4-220C140.4 197.2 85 95.2 21.4 0c11.9 3.1 23 3.9 33.2 3.9 9 0 20.4-.8 34.1-3.9 40.9 72.2 82.1 138.7 135 225.5C261 163.9 314.8 81.4 358.6 0c11.1 2.9 22 3.9 32.9 3.9 11.5 0 23.2-1 35-3.9C392.1 47.9 294.9 216.9 252 292z"],yandex:[256,512,[],"f413","M153.1 315.8L65.7 512H2l96-209.8c-45.1-22.9-75.2-64.4-75.2-141.1C22.7 53.7 90.8 0 171.7 0H254v512h-55.1V315.8h-45.8zm45.8-269.3h-29.4c-44.4 0-87.4 29.4-87.4 114.6 0 82.3 39.4 108.8 87.4 108.8h29.4V46.5z"],"yandex-international":[320,512,[],"f414","M129.5 512V345.9L18.5 48h55.8l81.8 229.7L250.2 0h51.3L180.8 347.8V512h-51.3z"],yelp:[384,512,[],"f1e9","M136.9 328c-1 .3-109.2 35.7-115.8 35.7-15.2-.9-18.5-16.2-19.9-31.2-1.5-14.2-1.4-29.8.3-46.8 1.9-18.8 5.5-45.1 24.2-44 4.8 0 67.1 25.9 112.7 44.4 17.1 6.8 18.6 35.8-1.5 41.9zm57.9-113.9c1.8 38.2-25.5 48.5-47.2 14.3L41.3 60.4c-1.5-6.6.3-12.4 5.3-17.4C62.2 26.5 146 3.2 168.1 8.9c7.5 1.9 12.1 6.1 13.8 12.6 1.3 8.3 11.5 167.4 12.9 192.6zm-1.4 164.8c0 4.6.2 116.4-1.7 121.5-2.3 6-7 9.7-14.3 11.2-10.1 1.7-27.1-1.9-51-10.7-22-8.1-56.7-21.5-49.3-42.5 2.8-6.9 51.4-62.8 77.3-93.6 12-15.2 39.8-5.5 39 14.1zm180.2-117.8c-5.6 3.7-110.8 28.2-118.1 30.6l.3-.6c-18.1 4.7-35.4-18.5-23.3-34.6 3.7-3.7 65.9-92.4 72.8-97 5.2-3.6 11.3-3.8 18.3-.6 18.4 8.8 55.1 63.1 57.4 84.6-.1 2.9 1.2 11.7-7.4 17.6zm10.1 130.7c-2.7 20.6-44.5 73.4-63.8 81-6.9 2.6-12.9 2-17.7-2-5-3.5-61.8-97.1-64.9-102.3-10.9-16.2 6.8-39.8 25.6-33.2 0 0 110.5 35.7 114.7 39.4 5.2 4.1 7.2 9.8 6.1 17.1z"],yoast:[448,512,[],"f2b1","M91.3 76h186l-7 18.9h-179c-39.7 0-71.9 31.6-71.9 70.3v205.4c0 35.4 24.9 70.3 84 70.3V460H91.3C41.2 460 0 419.8 0 370.5V165.2C0 115.9 40.7 76 91.3 76zm229.1-56h66.5C243.1 398.1 241.2 418.9 202.2 459.3c-20.8 21.6-49.3 31.7-78.3 32.7v-51.1c49.2-7.7 64.6-49.9 64.6-75.3 0-20.1.6-12.6-82.1-223.2h61.4L218.2 299 320.4 20zM448 161.5V460H234c6.6-9.6 10.7-16.3 12.1-19.4h182.5V161.5c0-32.5-17.1-51.9-48.2-62.9l6.7-17.6c41.7 13.6 60.9 43.1 60.9 80.5z"],youtube:[576,512,[],"f167","M549.655 124.083c-6.281-23.65-24.787-42.276-48.284-48.597C458.781 64 288 64 288 64S117.22 64 74.629 75.486c-23.497 6.322-42.003 24.947-48.284 48.597-11.412 42.867-11.412 132.305-11.412 132.305s0 89.438 11.412 132.305c6.281 23.65 24.787 41.5 48.284 47.821C117.22 448 288 448 288 448s170.78 0 213.371-11.486c23.497-6.321 42.003-24.171 48.284-47.821 11.412-42.867 11.412-132.305 11.412-132.305s0-89.438-11.412-132.305zm-317.51 213.508V175.185l142.739 81.205-142.739 81.201z"],"youtube-square":[448,512,[],"f431","M186.8 202.1l95.2 54.1-95.2 54.1V202.1zM448 80v352c0 26.5-21.5 48-48 48H48c-26.5 0-48-21.5-48-48V80c0-26.5 21.5-48 48-48h352c26.5 0 48 21.5 48 48zm-42 176.3s0-59.6-7.6-88.2c-4.2-15.8-16.5-28.2-32.2-32.4C337.9 128 224 128 224 128s-113.9 0-142.2 7.7c-15.7 4.2-28 16.6-32.2 32.4-7.6 28.5-7.6 88.2-7.6 88.2s0 59.6 7.6 88.2c4.2 15.8 16.5 27.7 32.2 31.9C110.1 384 224 384 224 384s113.9 0 142.2-7.7c15.7-4.2 28-16.1 32.2-31.9 7.6-28.5 7.6-88.1 7.6-88.1z"]};!function(c){try{c()}catch(c){if(!e)throw c}}(function(){!function c(l,v){var h=Object.keys(v).reduce(function(c,l){var h=v[l];return h.icon?c[h.iconName]=h.icon:c[l]=h,c},{});"function"==typeof t.hooks.addPack?t.hooks.addPack(l,h):t.styles[l]=f({},t.styles[l]||{},h),"fas"===l&&c("fa",v)}("fab",M)})}(),function(){"use strict";var c={};try{"undefined"!=typeof window&&(c=window)}catch(c){}var l=(c.navigator||{}).userAgent,h=void 0===l?"":l,v=c,z=(~h.indexOf("MSIE")||h.indexOf("Trident/"),"___FONT_AWESOME___"),e=function(){try{return!0}catch(c){return!1}}(),m=[1,2,3,4,5,6,7,8,9,10],a=m.concat([11,12,13,14,15,16,17,18,19,20]);["xs","sm","lg","fw","ul","li","border","pull-left","pull-right","spin","pulse","rotate-90","rotate-180","rotate-270","flip-horizontal","flip-vertical","stack","stack-1x","stack-2x","inverse","layers","layers-text","layers-counter"].concat(m.map(function(c){return c+"x"})).concat(a.map(function(c){return"w-"+c}));var s=v||{};s[z]||(s[z]={}),s[z].styles||(s[z].styles={}),s[z].hooks||(s[z].hooks={}),s[z].shims||(s[z].shims=[]);var t=s[z],f=Object.assign||function(c){for(var l=1;l<arguments.length;l++){var h=arguments[l];for(var v in h)Object.prototype.hasOwnProperty.call(h,v)&&(c[v]=h[v])}return c};var M={"address-book":[448,512,[],"f2b9","M436 160c6.6 0 12-5.4 12-12v-40c0-6.6-5.4-12-12-12h-20V48c0-26.5-21.5-48-48-48H48C21.5 0 0 21.5 0 48v416c0 26.5 21.5 48 48 48h320c26.5 0 48-21.5 48-48v-48h20c6.6 0 12-5.4 12-12v-40c0-6.6-5.4-12-12-12h-20v-64h20c6.6 0 12-5.4 12-12v-40c0-6.6-5.4-12-12-12h-20v-64h20zm-68 304H48V48h320v416zM208 256c35.3 0 64-28.7 64-64s-28.7-64-64-64-64 28.7-64 64 28.7 64 64 64zm-89.6 128h179.2c12.4 0 22.4-8.6 22.4-19.2v-19.2c0-31.8-30.1-57.6-67.2-57.6-10.8 0-18.7 8-44.8 8-26.9 0-33.4-8-44.8-8-37.1 0-67.2 25.8-67.2 57.6v19.2c0 10.6 10 19.2 22.4 19.2z"],"address-card":[576,512,[],"f2bb","M528 32H48C21.5 32 0 53.5 0 80v352c0 26.5 21.5 48 48 48h480c26.5 0 48-21.5 48-48V80c0-26.5-21.5-48-48-48zm0 400H48V80h480v352zM208 256c35.3 0 64-28.7 64-64s-28.7-64-64-64-64 28.7-64 64 28.7 64 64 64zm-89.6 128h179.2c12.4 0 22.4-8.6 22.4-19.2v-19.2c0-31.8-30.1-57.6-67.2-57.6-10.8 0-18.7 8-44.8 8-26.9 0-33.4-8-44.8-8-37.1 0-67.2 25.8-67.2 57.6v19.2c0 10.6 10 19.2 22.4 19.2zM360 320h112c4.4 0 8-3.6 8-8v-16c0-4.4-3.6-8-8-8H360c-4.4 0-8 3.6-8 8v16c0 4.4 3.6 8 8 8zm0-64h112c4.4 0 8-3.6 8-8v-16c0-4.4-3.6-8-8-8H360c-4.4 0-8 3.6-8 8v16c0 4.4 3.6 8 8 8zm0-64h112c4.4 0 8-3.6 8-8v-16c0-4.4-3.6-8-8-8H360c-4.4 0-8 3.6-8 8v16c0 4.4 3.6 8 8 8z"],"arrow-alt-circle-down":[512,512,[],"f358","M256 8C119 8 8 119 8 256s111 248 248 248 248-111 248-248S393 8 256 8zm0 448c-110.5 0-200-89.5-200-200S145.5 56 256 56s200 89.5 200 200-89.5 200-200 200zm-32-316v116h-67c-10.7 0-16 12.9-8.5 20.5l99 99c4.7 4.7 12.3 4.7 17 0l99-99c7.6-7.6 2.2-20.5-8.5-20.5h-67V140c0-6.6-5.4-12-12-12h-40c-6.6 0-12 5.4-12 12z"],"arrow-alt-circle-left":[512,512,[],"f359","M8 256c0 137 111 248 248 248s248-111 248-248S393 8 256 8 8 119 8 256zm448 0c0 110.5-89.5 200-200 200S56 366.5 56 256 145.5 56 256 56s200 89.5 200 200zm-72-20v40c0 6.6-5.4 12-12 12H256v67c0 10.7-12.9 16-20.5 8.5l-99-99c-4.7-4.7-4.7-12.3 0-17l99-99c7.6-7.6 20.5-2.2 20.5 8.5v67h116c6.6 0 12 5.4 12 12z"],"arrow-alt-circle-right":[512,512,[],"f35a","M504 256C504 119 393 8 256 8S8 119 8 256s111 248 248 248 248-111 248-248zm-448 0c0-110.5 89.5-200 200-200s200 89.5 200 200-89.5 200-200 200S56 366.5 56 256zm72 20v-40c0-6.6 5.4-12 12-12h116v-67c0-10.7 12.9-16 20.5-8.5l99 99c4.7 4.7 4.7 12.3 0 17l-99 99c-7.6 7.6-20.5 2.2-20.5-8.5v-67H140c-6.6 0-12-5.4-12-12z"],"arrow-alt-circle-up":[512,512,[],"f35b","M256 504c137 0 248-111 248-248S393 8 256 8 8 119 8 256s111 248 248 248zm0-448c110.5 0 200 89.5 200 200s-89.5 200-200 200S56 366.5 56 256 145.5 56 256 56zm20 328h-40c-6.6 0-12-5.4-12-12V256h-67c-10.7 0-16-12.9-8.5-20.5l99-99c4.7-4.7 12.3-4.7 17 0l99 99c7.6 7.6 2.2 20.5-8.5 20.5h-67v116c0 6.6-5.4 12-12 12z"],bell:[448,512,[],"f0f3","M425.403 330.939c-16.989-16.785-34.546-34.143-34.546-116.083 0-83.026-60.958-152.074-140.467-164.762A31.843 31.843 0 0 0 256 32c0-17.673-14.327-32-32-32s-32 14.327-32 32a31.848 31.848 0 0 0 5.609 18.095C118.101 62.783 57.143 131.831 57.143 214.857c0 81.933-17.551 99.292-34.543 116.078C-25.496 378.441 9.726 448 66.919 448H160c0 35.346 28.654 64 64 64 35.346 0 64-28.654 64-64h93.08c57.19 0 92.415-69.583 44.323-117.061zM224 472c-13.234 0-24-10.766-24-24h48c0 13.234-10.766 24-24 24zm157.092-72H66.9c-16.762 0-25.135-20.39-13.334-32.191 28.585-28.585 51.577-55.724 51.577-152.952C105.143 149.319 158.462 96 224 96s118.857 53.319 118.857 118.857c0 97.65 23.221 124.574 51.568 152.952C406.278 379.661 397.783 400 381.092 400z"],"bell-slash":[576,512,[],"f1f6","M130.9 400c-16.762 0-25.135-20.39-13.334-32.191 25.226-25.226 46.094-49.338 50.649-121.48l-46.777-41.274a168.48 168.48 0 0 0-.296 9.802c0 81.933-17.551 99.292-34.543 116.078C38.504 378.441 73.726 448 130.919 448H224c0 35.346 28.654 64 64 64s64-28.654 64-64h44.777l-54.4-48H130.9zM288 472c-13.234 0-24-10.766-24-24h48c0 13.234-10.766 24-24 24zm283.867.553l-67.931-59.571c13.104-24.118 11.524-56.318-14.532-82.042-16.989-16.785-34.546-34.143-34.546-116.083 0-83.026-60.958-152.074-140.467-164.762A31.848 31.848 0 0 0 320 32c0-17.673-14.327-32-32-32s-32 14.327-32 32a31.848 31.848 0 0 0 5.609 18.095c-41.471 6.618-77.891 28.571-103.249 59.841L36.459 3.037c-5.058-4.436-12.777-3.956-17.24 1.071L3.056 22.313C-1.407 27.34-.925 35.012 4.134 39.447l535.408 469.516c5.058 4.436 12.777 3.956 17.24-1.071l16.163-18.205c4.462-5.027 3.98-12.699-1.078-17.134zM288 96c65.538 0 118.857 53.319 118.857 118.857 0 97.65 23.221 124.574 51.568 152.952 2.908 2.908 4.573 6.328 5.209 9.832L194.482 141.612C216.258 113.867 250.075 96 288 96z"],bookmark:[384,512,[],"f02e","M336 0H48C21.49 0 0 21.49 0 48v464l192-112 192 112V48c0-26.51-21.49-48-48-48zm0 428.43l-144-84-144 84V54a6 6 0 0 1 6-6h276c3.314 0 6 2.683 6 5.996V428.43z"],building:[448,512,[],"f1ad","M128 148v-40c0-6.6 5.4-12 12-12h40c6.6 0 12 5.4 12 12v40c0 6.6-5.4 12-12 12h-40c-6.6 0-12-5.4-12-12zm140 12h40c6.6 0 12-5.4 12-12v-40c0-6.6-5.4-12-12-12h-40c-6.6 0-12 5.4-12 12v40c0 6.6 5.4 12 12 12zm-128 96h40c6.6 0 12-5.4 12-12v-40c0-6.6-5.4-12-12-12h-40c-6.6 0-12 5.4-12 12v40c0 6.6 5.4 12 12 12zm128 0h40c6.6 0 12-5.4 12-12v-40c0-6.6-5.4-12-12-12h-40c-6.6 0-12 5.4-12 12v40c0 6.6 5.4 12 12 12zm-76 84v-40c0-6.6-5.4-12-12-12h-40c-6.6 0-12 5.4-12 12v40c0 6.6 5.4 12 12 12h40c6.6 0 12-5.4 12-12zm76 12h40c6.6 0 12-5.4 12-12v-40c0-6.6-5.4-12-12-12h-40c-6.6 0-12 5.4-12 12v40c0 6.6 5.4 12 12 12zm180 124v36H0v-36c0-6.6 5.4-12 12-12h19.5V24c0-13.3 10.7-24 24-24h337c13.3 0 24 10.7 24 24v440H436c6.6 0 12 5.4 12 12zM79.5 463H192v-67c0-6.6 5.4-12 12-12h40c6.6 0 12 5.4 12 12v67h112.5V49L80 48l-.5 415z"],calendar:[448,512,[],"f133","M400 64h-48V12c0-6.6-5.4-12-12-12h-40c-6.6 0-12 5.4-12 12v52H160V12c0-6.6-5.4-12-12-12h-40c-6.6 0-12 5.4-12 12v52H48C21.5 64 0 85.5 0 112v352c0 26.5 21.5 48 48 48h352c26.5 0 48-21.5 48-48V112c0-26.5-21.5-48-48-48zm-6 400H54c-3.3 0-6-2.7-6-6V160h352v298c0 3.3-2.7 6-6 6z"],"calendar-alt":[448,512,[],"f073","M148 288h-40c-6.6 0-12-5.4-12-12v-40c0-6.6 5.4-12 12-12h40c6.6 0 12 5.4 12 12v40c0 6.6-5.4 12-12 12zm108-12v-40c0-6.6-5.4-12-12-12h-40c-6.6 0-12 5.4-12 12v40c0 6.6 5.4 12 12 12h40c6.6 0 12-5.4 12-12zm96 0v-40c0-6.6-5.4-12-12-12h-40c-6.6 0-12 5.4-12 12v40c0 6.6 5.4 12 12 12h40c6.6 0 12-5.4 12-12zm-96 96v-40c0-6.6-5.4-12-12-12h-40c-6.6 0-12 5.4-12 12v40c0 6.6 5.4 12 12 12h40c6.6 0 12-5.4 12-12zm-96 0v-40c0-6.6-5.4-12-12-12h-40c-6.6 0-12 5.4-12 12v40c0 6.6 5.4 12 12 12h40c6.6 0 12-5.4 12-12zm192 0v-40c0-6.6-5.4-12-12-12h-40c-6.6 0-12 5.4-12 12v40c0 6.6 5.4 12 12 12h40c6.6 0 12-5.4 12-12zm96-260v352c0 26.5-21.5 48-48 48H48c-26.5 0-48-21.5-48-48V112c0-26.5 21.5-48 48-48h48V12c0-6.6 5.4-12 12-12h40c6.6 0 12 5.4 12 12v52h128V12c0-6.6 5.4-12 12-12h40c6.6 0 12 5.4 12 12v52h48c26.5 0 48 21.5 48 48zm-48 346V160H48v298c0 3.3 2.7 6 6 6h340c3.3 0 6-2.7 6-6z"],"calendar-check":[448,512,[],"f274","M400 64h-48V12c0-6.627-5.373-12-12-12h-40c-6.627 0-12 5.373-12 12v52H160V12c0-6.627-5.373-12-12-12h-40c-6.627 0-12 5.373-12 12v52H48C21.49 64 0 85.49 0 112v352c0 26.51 21.49 48 48 48h352c26.51 0 48-21.49 48-48V112c0-26.51-21.49-48-48-48zm-6 400H54a6 6 0 0 1-6-6V160h352v298a6 6 0 0 1-6 6zm-52.849-200.65L198.842 404.519c-4.705 4.667-12.303 4.637-16.971-.068l-75.091-75.699c-4.667-4.705-4.637-12.303.068-16.971l22.719-22.536c4.705-4.667 12.303-4.637 16.97.069l44.104 44.461 111.072-110.181c4.705-4.667 12.303-4.637 16.971.068l22.536 22.718c4.667 4.705 4.636 12.303-.069 16.97z"],"calendar-minus":[448,512,[],"f272","M124 328c-6.6 0-12-5.4-12-12v-24c0-6.6 5.4-12 12-12h200c6.6 0 12 5.4 12 12v24c0 6.6-5.4 12-12 12H124zm324-216v352c0 26.5-21.5 48-48 48H48c-26.5 0-48-21.5-48-48V112c0-26.5 21.5-48 48-48h48V12c0-6.6 5.4-12 12-12h40c6.6 0 12 5.4 12 12v52h128V12c0-6.6 5.4-12 12-12h40c6.6 0 12 5.4 12 12v52h48c26.5 0 48 21.5 48 48zm-48 346V160H48v298c0 3.3 2.7 6 6 6h340c3.3 0 6-2.7 6-6z"],"calendar-plus":[448,512,[],"f271","M336 292v24c0 6.6-5.4 12-12 12h-76v76c0 6.6-5.4 12-12 12h-24c-6.6 0-12-5.4-12-12v-76h-76c-6.6 0-12-5.4-12-12v-24c0-6.6 5.4-12 12-12h76v-76c0-6.6 5.4-12 12-12h24c6.6 0 12 5.4 12 12v76h76c6.6 0 12 5.4 12 12zm112-180v352c0 26.5-21.5 48-48 48H48c-26.5 0-48-21.5-48-48V112c0-26.5 21.5-48 48-48h48V12c0-6.6 5.4-12 12-12h40c6.6 0 12 5.4 12 12v52h128V12c0-6.6 5.4-12 12-12h40c6.6 0 12 5.4 12 12v52h48c26.5 0 48 21.5 48 48zm-48 346V160H48v298c0 3.3 2.7 6 6 6h340c3.3 0 6-2.7 6-6z"],"calendar-times":[448,512,[],"f273","M311.7 374.7l-17 17c-4.7 4.7-12.3 4.7-17 0L224 337.9l-53.7 53.7c-4.7 4.7-12.3 4.7-17 0l-17-17c-4.7-4.7-4.7-12.3 0-17l53.7-53.7-53.7-53.7c-4.7-4.7-4.7-12.3 0-17l17-17c4.7-4.7 12.3-4.7 17 0l53.7 53.7 53.7-53.7c4.7-4.7 12.3-4.7 17 0l17 17c4.7 4.7 4.7 12.3 0 17L257.9 304l53.7 53.7c4.8 4.7 4.8 12.3.1 17zM448 112v352c0 26.5-21.5 48-48 48H48c-26.5 0-48-21.5-48-48V112c0-26.5 21.5-48 48-48h48V12c0-6.6 5.4-12 12-12h40c6.6 0 12 5.4 12 12v52h128V12c0-6.6 5.4-12 12-12h40c6.6 0 12 5.4 12 12v52h48c26.5 0 48 21.5 48 48zm-48 346V160H48v298c0 3.3 2.7 6 6 6h340c3.3 0 6-2.7 6-6z"],"caret-square-down":[448,512,[],"f150","M125.1 208h197.8c10.7 0 16.1 13 8.5 20.5l-98.9 98.3c-4.7 4.7-12.2 4.7-16.9 0l-98.9-98.3c-7.7-7.5-2.3-20.5 8.4-20.5zM448 80v352c0 26.5-21.5 48-48 48H48c-26.5 0-48-21.5-48-48V80c0-26.5 21.5-48 48-48h352c26.5 0 48 21.5 48 48zm-48 346V86c0-3.3-2.7-6-6-6H54c-3.3 0-6 2.7-6 6v340c0 3.3 2.7 6 6 6h340c3.3 0 6-2.7 6-6z"],"caret-square-left":[448,512,[],"f191","M272 157.1v197.8c0 10.7-13 16.1-20.5 8.5l-98.3-98.9c-4.7-4.7-4.7-12.2 0-16.9l98.3-98.9c7.5-7.7 20.5-2.3 20.5 8.4zM448 80v352c0 26.5-21.5 48-48 48H48c-26.5 0-48-21.5-48-48V80c0-26.5 21.5-48 48-48h352c26.5 0 48 21.5 48 48zm-48 346V86c0-3.3-2.7-6-6-6H54c-3.3 0-6 2.7-6 6v340c0 3.3 2.7 6 6 6h340c3.3 0 6-2.7 6-6z"],"caret-square-right":[448,512,[],"f152","M176 354.9V157.1c0-10.7 13-16.1 20.5-8.5l98.3 98.9c4.7 4.7 4.7 12.2 0 16.9l-98.3 98.9c-7.5 7.7-20.5 2.3-20.5-8.4zM448 80v352c0 26.5-21.5 48-48 48H48c-26.5 0-48-21.5-48-48V80c0-26.5 21.5-48 48-48h352c26.5 0 48 21.5 48 48zm-48 346V86c0-3.3-2.7-6-6-6H54c-3.3 0-6 2.7-6 6v340c0 3.3 2.7 6 6 6h340c3.3 0 6-2.7 6-6z"],"caret-square-up":[448,512,[],"f151","M322.9 304H125.1c-10.7 0-16.1-13-8.5-20.5l98.9-98.3c4.7-4.7 12.2-4.7 16.9 0l98.9 98.3c7.7 7.5 2.3 20.5-8.4 20.5zM448 80v352c0 26.5-21.5 48-48 48H48c-26.5 0-48-21.5-48-48V80c0-26.5 21.5-48 48-48h352c26.5 0 48 21.5 48 48zm-48 346V86c0-3.3-2.7-6-6-6H54c-3.3 0-6 2.7-6 6v340c0 3.3 2.7 6 6 6h340c3.3 0 6-2.7 6-6z"],"chart-bar":[512,512,[],"f080","M500 400c6.6 0 12 5.4 12 12v24c0 6.6-5.4 12-12 12H12c-6.6 0-12-5.4-12-12V76c0-6.6 5.4-12 12-12h24c6.6 0 12 5.4 12 12v324h452zm-356-60v-72c0-6.6-5.4-12-12-12h-24c-6.6 0-12 5.4-12 12v72c0 6.6 5.4 12 12 12h24c6.6 0 12-5.4 12-12zm96 0V140c0-6.6-5.4-12-12-12h-24c-6.6 0-12 5.4-12 12v200c0 6.6 5.4 12 12 12h24c6.6 0 12-5.4 12-12zm96 0V204c0-6.6-5.4-12-12-12h-24c-6.6 0-12 5.4-12 12v136c0 6.6 5.4 12 12 12h24c6.6 0 12-5.4 12-12zm96 0V108c0-6.6-5.4-12-12-12h-24c-6.6 0-12 5.4-12 12v232c0 6.6 5.4 12 12 12h24c6.6 0 12-5.4 12-12z"],"check-circle":[512,512,[],"f058","M256 8C119.033 8 8 119.033 8 256s111.033 248 248 248 248-111.033 248-248S392.967 8 256 8zm0 48c110.532 0 200 89.451 200 200 0 110.532-89.451 200-200 200-110.532 0-200-89.451-200-200 0-110.532 89.451-200 200-200m140.204 130.267l-22.536-22.718c-4.667-4.705-12.265-4.736-16.97-.068L215.346 303.697l-59.792-60.277c-4.667-4.705-12.265-4.736-16.97-.069l-22.719 22.536c-4.705 4.667-4.736 12.265-.068 16.971l90.781 91.516c4.667 4.705 12.265 4.736 16.97.068l172.589-171.204c4.704-4.668 4.734-12.266.067-16.971z"],"check-square":[448,512,[],"f14a","M400 32H48C21.49 32 0 53.49 0 80v352c0 26.51 21.49 48 48 48h352c26.51 0 48-21.49 48-48V80c0-26.51-21.49-48-48-48zm0 400H48V80h352v352zm-35.864-241.724L191.547 361.48c-4.705 4.667-12.303 4.637-16.97-.068l-90.781-91.516c-4.667-4.705-4.637-12.303.069-16.971l22.719-22.536c4.705-4.667 12.303-4.637 16.97.069l59.792 60.277 141.352-140.216c4.705-4.667 12.303-4.637 16.97.068l22.536 22.718c4.667 4.706 4.637 12.304-.068 16.971z"],circle:[512,512,[],"f111","M256 8C119 8 8 119 8 256s111 248 248 248 248-111 248-248S393 8 256 8zm0 448c-110.5 0-200-89.5-200-200S145.5 56 256 56s200 89.5 200 200-89.5 200-200 200z"],clipboard:[384,512,[],"f328","M336 64h-80c0-35.29-28.71-64-64-64s-64 28.71-64 64H48C21.49 64 0 85.49 0 112v352c0 26.51 21.49 48 48 48h288c26.51 0 48-21.49 48-48V112c0-26.51-21.49-48-48-48zm-6 400H54a6 6 0 0 1-6-6V118a6 6 0 0 1 6-6h42v36c0 6.627 5.373 12 12 12h168c6.627 0 12-5.373 12-12v-36h42a6 6 0 0 1 6 6v340a6 6 0 0 1-6 6zM192 40c13.255 0 24 10.745 24 24s-10.745 24-24 24-24-10.745-24-24 10.745-24 24-24"],clock:[512,512,[],"f017","M256 8C119 8 8 119 8 256s111 248 248 248 248-111 248-248S393 8 256 8zm0 448c-110.5 0-200-89.5-200-200S145.5 56 256 56s200 89.5 200 200-89.5 200-200 200zm61.8-104.4l-84.9-61.7c-3.1-2.3-4.9-5.9-4.9-9.7V116c0-6.6 5.4-12 12-12h32c6.6 0 12 5.4 12 12v141.7l66.8 48.6c5.4 3.9 6.5 11.4 2.6 16.8L334.6 349c-3.9 5.3-11.4 6.5-16.8 2.6z"],clone:[512,512,[],"f24d","M464 0H144c-26.51 0-48 21.49-48 48v48H48c-26.51 0-48 21.49-48 48v320c0 26.51 21.49 48 48 48h320c26.51 0 48-21.49 48-48v-48h48c26.51 0 48-21.49 48-48V48c0-26.51-21.49-48-48-48zM362 464H54a6 6 0 0 1-6-6V150a6 6 0 0 1 6-6h42v224c0 26.51 21.49 48 48 48h224v42a6 6 0 0 1-6 6zm96-96H150a6 6 0 0 1-6-6V54a6 6 0 0 1 6-6h308a6 6 0 0 1 6 6v308a6 6 0 0 1-6 6z"],"closed-captioning":[512,512,[],"f20a","M464 64H48C21.5 64 0 85.5 0 112v288c0 26.5 21.5 48 48 48h416c26.5 0 48-21.5 48-48V112c0-26.5-21.5-48-48-48zm-6 336H54c-3.3 0-6-2.7-6-6V118c0-3.3 2.7-6 6-6h404c3.3 0 6 2.7 6 6v276c0 3.3-2.7 6-6 6zm-211.1-85.7c1.7 2.4 1.5 5.6-.5 7.7-53.6 56.8-172.8 32.1-172.8-67.9 0-97.3 121.7-119.5 172.5-70.1 2.1 2 2.5 3.2 1 5.7l-17.5 30.5c-1.9 3.1-6.2 4-9.1 1.7-40.8-32-94.6-14.9-94.6 31.2 0 48 51 70.5 92.2 32.6 2.8-2.5 7.1-2.1 9.2.9l19.6 27.7zm190.4 0c1.7 2.4 1.5 5.6-.5 7.7-53.6 56.9-172.8 32.1-172.8-67.9 0-97.3 121.7-119.5 172.5-70.1 2.1 2 2.5 3.2 1 5.7L420 220.2c-1.9 3.1-6.2 4-9.1 1.7-40.8-32-94.6-14.9-94.6 31.2 0 48 51 70.5 92.2 32.6 2.8-2.5 7.1-2.1 9.2.9l19.6 27.7z"],comment:[512,512,[],"f075","M256 32C114.6 32 0 125.1 0 240c0 47.6 19.9 91.2 52.9 126.3C38 405.7 7 439.1 6.5 439.5c-6.6 7-8.4 17.2-4.6 26S14.4 480 24 480c61.5 0 110-25.7 139.1-46.3C192 442.8 223.2 448 256 448c141.4 0 256-93.1 256-208S397.4 32 256 32zm0 368c-26.7 0-53.1-4.1-78.4-12.1l-22.7-7.2-19.5 13.8c-14.3 10.1-33.9 21.4-57.5 29 7.3-12.1 14.4-25.7 19.9-40.2l10.6-28.1-20.6-21.8C69.7 314.1 48 282.2 48 240c0-88.2 93.3-160 208-160s208 71.8 208 160-93.3 160-208 160z"],"comment-alt":[512,512,[],"f27a","M448 0H64C28.7 0 0 28.7 0 64v288c0 35.3 28.7 64 64 64h96v84c0 7.1 5.8 12 12 12 2.4 0 4.9-.7 7.1-2.4L304 416h144c35.3 0 64-28.7 64-64V64c0-35.3-28.7-64-64-64zm16 352c0 8.8-7.2 16-16 16H288l-12.8 9.6L208 428v-60H64c-8.8 0-16-7.2-16-16V64c0-8.8 7.2-16 16-16h384c8.8 0 16 7.2 16 16v288z"],"comment-dots":[512,512,[],"f4ad","M144 208c-17.7 0-32 14.3-32 32s14.3 32 32 32 32-14.3 32-32-14.3-32-32-32zm112 0c-17.7 0-32 14.3-32 32s14.3 32 32 32 32-14.3 32-32-14.3-32-32-32zm112 0c-17.7 0-32 14.3-32 32s14.3 32 32 32 32-14.3 32-32-14.3-32-32-32zM256 32C114.6 32 0 125.1 0 240c0 47.6 19.9 91.2 52.9 126.3C38 405.7 7 439.1 6.5 439.5c-6.6 7-8.4 17.2-4.6 26S14.4 480 24 480c61.5 0 110-25.7 139.1-46.3C192 442.8 223.2 448 256 448c141.4 0 256-93.1 256-208S397.4 32 256 32zm0 368c-26.7 0-53.1-4.1-78.4-12.1l-22.7-7.2-19.5 13.8c-14.3 10.1-33.9 21.4-57.5 29 7.3-12.1 14.4-25.7 19.9-40.2l10.6-28.1-20.6-21.8C69.7 314.1 48 282.2 48 240c0-88.2 93.3-160 208-160s208 71.8 208 160-93.3 160-208 160z"],comments:[576,512,[],"f086","M532 386.2c27.5-27.1 44-61.1 44-98.2 0-80-76.5-146.1-176.2-157.9C368.3 72.5 294.3 32 208 32 93.1 32 0 103.6 0 192c0 37 16.5 71 44 98.2-15.3 30.7-37.3 54.5-37.7 54.9-6.3 6.7-8.1 16.5-4.4 25 3.6 8.5 12 14 21.2 14 53.5 0 96.7-20.2 125.2-38.8 9.2 2.1 18.7 3.7 28.4 4.9C208.1 407.6 281.8 448 368 448c20.8 0 40.8-2.4 59.8-6.8C456.3 459.7 499.4 480 553 480c9.2 0 17.5-5.5 21.2-14 3.6-8.5 1.9-18.3-4.4-25-.4-.3-22.5-24.1-37.8-54.8zm-392.8-92.3L122.1 305c-14.1 9.1-28.5 16.3-43.1 21.4 2.7-4.7 5.4-9.7 8-14.8l15.5-31.1L77.7 256C64.2 242.6 48 220.7 48 192c0-60.7 73.3-112 160-112s160 51.3 160 112-73.3 112-160 112c-16.5 0-33-1.9-49-5.6l-19.8-4.5zM498.3 352l-24.7 24.4 15.5 31.1c2.6 5.1 5.3 10.1 8 14.8-14.6-5.1-29-12.3-43.1-21.4l-17.1-11.1-19.9 4.6c-16 3.7-32.5 5.6-49 5.6-54 0-102.2-20.1-131.3-49.7C338 339.5 416 272.9 416 192c0-3.4-.4-6.7-.7-10C479.7 196.5 528 238.8 528 288c0 28.7-16.2 50.6-29.7 64z"],compass:[512,512,[],"f14e","M256 8C119.033 8 8 119.033 8 256s111.033 248 248 248 248-111.033 248-248S392.967 8 256 8zm0 448c-110.532 0-200-89.451-200-200 0-110.531 89.451-200 200-200 110.532 0 200 89.451 200 200 0 110.532-89.451 200-200 200zm91.326-312.131l-33.359 137.779a24.005 24.005 0 0 1-6.772 11.729l-102.64 97.779c-17.104 16.293-45.56.434-39.88-23.024l33.359-137.779a23.997 23.997 0 0 1 6.772-11.729l102.642-97.779c17.285-16.47 45.494-.175 39.878 23.024zM256 224c-17.673 0-32 14.327-32 32s14.327 32 32 32 32-14.327 32-32-14.327-32-32-32z"],copy:[448,512,[],"f0c5","M433.941 65.941l-51.882-51.882A48 48 0 0 0 348.118 0H176c-26.51 0-48 21.49-48 48v48H48c-26.51 0-48 21.49-48 48v320c0 26.51 21.49 48 48 48h224c26.51 0 48-21.49 48-48v-48h80c26.51 0 48-21.49 48-48V99.882a48 48 0 0 0-14.059-33.941zM266 464H54a6 6 0 0 1-6-6V150a6 6 0 0 1 6-6h74v224c0 26.51 21.49 48 48 48h96v42a6 6 0 0 1-6 6zm128-96H182a6 6 0 0 1-6-6V54a6 6 0 0 1 6-6h106v88c0 13.255 10.745 24 24 24h88v202a6 6 0 0 1-6 6zm6-256h-64V48h9.632c1.591 0 3.117.632 4.243 1.757l48.368 48.368a6 6 0 0 1 1.757 4.243V112z"],copyright:[512,512,[],"f1f9","M256 8C119.033 8 8 119.033 8 256s111.033 248 248 248 248-111.033 248-248S392.967 8 256 8zm0 448c-110.532 0-200-89.451-200-200 0-110.531 89.451-200 200-200 110.532 0 200 89.451 200 200 0 110.532-89.451 200-200 200zm107.351-101.064c-9.614 9.712-45.53 41.396-104.065 41.396-82.43 0-140.484-61.425-140.484-141.567 0-79.152 60.275-139.401 139.762-139.401 55.531 0 88.738 26.62 97.593 34.779a11.965 11.965 0 0 1 1.936 15.322l-18.155 28.113c-3.841 5.95-11.966 7.282-17.499 2.921-8.595-6.776-31.814-22.538-61.708-22.538-48.303 0-77.916 35.33-77.916 80.082 0 41.589 26.888 83.692 78.277 83.692 32.657 0 56.843-19.039 65.726-27.225 5.27-4.857 13.596-4.039 17.82 1.738l19.865 27.17a11.947 11.947 0 0 1-1.152 15.518z"],"credit-card":[576,512,[],"f09d","M527.9 32H48.1C21.5 32 0 53.5 0 80v352c0 26.5 21.5 48 48.1 48h479.8c26.6 0 48.1-21.5 48.1-48V80c0-26.5-21.5-48-48.1-48zM54.1 80h467.8c3.3 0 6 2.7 6 6v42H48.1V86c0-3.3 2.7-6 6-6zm467.8 352H54.1c-3.3 0-6-2.7-6-6V256h479.8v170c0 3.3-2.7 6-6 6zM192 332v40c0 6.6-5.4 12-12 12h-72c-6.6 0-12-5.4-12-12v-40c0-6.6 5.4-12 12-12h72c6.6 0 12 5.4 12 12zm192 0v40c0 6.6-5.4 12-12 12H236c-6.6 0-12-5.4-12-12v-40c0-6.6 5.4-12 12-12h136c6.6 0 12 5.4 12 12z"],"dot-circle":[512,512,[],"f192","M256 56c110.532 0 200 89.451 200 200 0 110.532-89.451 200-200 200-110.532 0-200-89.451-200-200 0-110.532 89.451-200 200-200m0-48C119.033 8 8 119.033 8 256s111.033 248 248 248 248-111.033 248-248S392.967 8 256 8zm0 168c-44.183 0-80 35.817-80 80s35.817 80 80 80 80-35.817 80-80-35.817-80-80-80z"],edit:[576,512,[],"f044","M402.3 344.9l32-32c5-5 13.7-1.5 13.7 5.7V464c0 26.5-21.5 48-48 48H48c-26.5 0-48-21.5-48-48V112c0-26.5 21.5-48 48-48h273.5c7.1 0 10.7 8.6 5.7 13.7l-32 32c-1.5 1.5-3.5 2.3-5.7 2.3H48v352h352V350.5c0-2.1.8-4.1 2.3-5.6zm156.6-201.8L296.3 405.7l-90.4 10c-26.2 2.9-48.5-19.2-45.6-45.6l10-90.4L432.9 17.1c22.9-22.9 59.9-22.9 82.7 0l43.2 43.2c22.9 22.9 22.9 60 .1 82.8zM460.1 174L402 115.9 216.2 301.8l-7.3 65.3 65.3-7.3L460.1 174zm64.8-79.7l-43.2-43.2c-4.1-4.1-10.8-4.1-14.8 0L436 82l58.1 58.1 30.9-30.9c4-4.2 4-10.8-.1-14.9z"],envelope:[512,512,[],"f0e0","M464 64H48C21.49 64 0 85.49 0 112v288c0 26.51 21.49 48 48 48h416c26.51 0 48-21.49 48-48V112c0-26.51-21.49-48-48-48zm0 48v40.805c-22.422 18.259-58.168 46.651-134.587 106.49-16.841 13.247-50.201 45.072-73.413 44.701-23.208.375-56.579-31.459-73.413-44.701C106.18 199.465 70.425 171.067 48 152.805V112h416zM48 400V214.398c22.914 18.251 55.409 43.862 104.938 82.646 21.857 17.205 60.134 55.186 103.062 54.955 42.717.231 80.509-37.199 103.053-54.947 49.528-38.783 82.032-64.401 104.947-82.653V400H48z"],"envelope-open":[512,512,[],"f2b6","M494.586 164.516c-4.697-3.883-111.723-89.95-135.251-108.657C337.231 38.191 299.437 0 256 0c-43.205 0-80.636 37.717-103.335 55.859-24.463 19.45-131.07 105.195-135.15 108.549A48.004 48.004 0 0 0 0 201.485V464c0 26.51 21.49 48 48 48h416c26.51 0 48-21.49 48-48V201.509a48 48 0 0 0-17.414-36.993zM464 458a6 6 0 0 1-6 6H54a6 6 0 0 1-6-6V204.347c0-1.813.816-3.526 2.226-4.665 15.87-12.814 108.793-87.554 132.364-106.293C200.755 78.88 232.398 48 256 48c23.693 0 55.857 31.369 73.41 45.389 23.573 18.741 116.503 93.493 132.366 106.316a5.99 5.99 0 0 1 2.224 4.663V458zm-31.991-187.704c4.249 5.159 3.465 12.795-1.745 16.981-28.975 23.283-59.274 47.597-70.929 56.863C336.636 362.283 299.205 400 256 400c-43.452 0-81.287-38.237-103.335-55.86-11.279-8.967-41.744-33.413-70.927-56.865-5.21-4.187-5.993-11.822-1.745-16.981l15.258-18.528c4.178-5.073 11.657-5.843 16.779-1.726 28.618 23.001 58.566 47.035 70.56 56.571C200.143 320.631 232.307 352 256 352c23.602 0 55.246-30.88 73.41-45.389 11.994-9.535 41.944-33.57 70.563-56.568 5.122-4.116 12.601-3.346 16.778 1.727l15.258 18.526z"],eye:[576,512,[],"f06e","M569.354 231.631C512.97 135.949 407.81 72 288 72 168.14 72 63.004 135.994 6.646 231.631a47.999 47.999 0 0 0 0 48.739C63.031 376.051 168.19 440 288 440c119.86 0 224.996-63.994 281.354-159.631a47.997 47.997 0 0 0 0-48.738zM288 392c-102.556 0-192.091-54.701-240-136 44.157-74.933 123.677-127.27 216.162-135.007C273.958 131.078 280 144.83 280 160c0 30.928-25.072 56-56 56s-56-25.072-56-56l.001-.042C157.794 179.043 152 200.844 152 224c0 75.111 60.889 136 136 136s136-60.889 136-136c0-31.031-10.4-59.629-27.895-82.515C451.704 164.638 498.009 205.106 528 256c-47.908 81.299-137.444 136-240 136z"],"eye-slash":[576,512,[],"f070","M272.702 359.139c-80.483-9.011-136.212-86.886-116.93-167.042l116.93 167.042zM288 392c-102.556 0-192.092-54.701-240-136 21.755-36.917 52.1-68.342 88.344-91.658l-27.541-39.343C67.001 152.234 31.921 188.741 6.646 231.631a47.999 47.999 0 0 0 0 48.739C63.004 376.006 168.14 440 288 440a332.89 332.89 0 0 0 39.648-2.367l-32.021-45.744A284.16 284.16 0 0 1 288 392zm281.354-111.631c-33.232 56.394-83.421 101.742-143.554 129.492l48.116 68.74c3.801 5.429 2.48 12.912-2.949 16.712L450.23 509.83c-5.429 3.801-12.912 2.48-16.712-2.949L102.084 33.399c-3.801-5.429-2.48-12.912 2.949-16.712L125.77 2.17c5.429-3.801 12.912-2.48 16.712 2.949l55.526 79.325C226.612 76.343 256.808 72 288 72c119.86 0 224.996 63.994 281.354 159.631a48.002 48.002 0 0 1 0 48.738zM528 256c-44.157-74.933-123.677-127.27-216.162-135.007C302.042 131.078 296 144.83 296 160c0 30.928 25.072 56 56 56s56-25.072 56-56l-.001-.042c30.632 57.277 16.739 130.26-36.928 171.719l26.695 38.135C452.626 346.551 498.308 306.386 528 256z"],file:[384,512,[],"f15b","M369.9 97.9L286 14C277 5 264.8-.1 252.1-.1H48C21.5 0 0 21.5 0 48v416c0 26.5 21.5 48 48 48h288c26.5 0 48-21.5 48-48V131.9c0-12.7-5.1-25-14.1-34zM332.1 128H256V51.9l76.1 76.1zM48 464V48h160v104c0 13.3 10.7 24 24 24h104v288H48z"],"file-alt":[384,512,[],"f15c","M288 248v28c0 6.6-5.4 12-12 12H108c-6.6 0-12-5.4-12-12v-28c0-6.6 5.4-12 12-12h168c6.6 0 12 5.4 12 12zm-12 72H108c-6.6 0-12 5.4-12 12v28c0 6.6 5.4 12 12 12h168c6.6 0 12-5.4 12-12v-28c0-6.6-5.4-12-12-12zm108-188.1V464c0 26.5-21.5 48-48 48H48c-26.5 0-48-21.5-48-48V48C0 21.5 21.5 0 48 0h204.1C264.8 0 277 5.1 286 14.1L369.9 98c9 8.9 14.1 21.2 14.1 33.9zm-128-80V128h76.1L256 51.9zM336 464V176H232c-13.3 0-24-10.7-24-24V48H48v416h288z"],"file-archive":[384,512,[],"f1c6","M369.941 97.941l-83.882-83.882A48 48 0 0 0 252.118 0H48C21.49 0 0 21.49 0 48v416c0 26.51 21.49 48 48 48h288c26.51 0 48-21.49 48-48V131.882a48 48 0 0 0-14.059-33.941zM256 51.882L332.118 128H256V51.882zM336 464H48V48h79.714v16h32V48H208v104c0 13.255 10.745 24 24 24h104v288zM192.27 96h-32V64h32v32zm-32 0v32h-32V96h32zm0 64v32h-32v-32h32zm32 0h-32v-32h32v32zm1.909 105.678A12 12 0 0 0 182.406 256H160.27v-32h-32v32l-19.69 97.106C101.989 385.611 126.834 416 160 416c33.052 0 57.871-30.192 51.476-62.62l-17.297-87.702zM160.27 390.073c-17.918 0-32.444-12.105-32.444-27.036 0-14.932 14.525-27.036 32.444-27.036s32.444 12.105 32.444 27.036c0 14.931-14.526 27.036-32.444 27.036zm32-166.073h-32v-32h32v32z"],"file-audio":[384,512,[],"f1c7","M369.941 97.941l-83.882-83.882A48 48 0 0 0 252.118 0H48C21.49 0 0 21.49 0 48v416c0 26.51 21.49 48 48 48h288c26.51 0 48-21.49 48-48V131.882a48 48 0 0 0-14.059-33.941zM332.118 128H256V51.882L332.118 128zM48 464V48h160v104c0 13.255 10.745 24 24 24h104v288H48zm144-76.024c0 10.691-12.926 16.045-20.485 8.485L136 360.486h-28c-6.627 0-12-5.373-12-12v-56c0-6.627 5.373-12 12-12h28l35.515-36.947c7.56-7.56 20.485-2.206 20.485 8.485v135.952zm41.201-47.13c9.051-9.297 9.06-24.133.001-33.439-22.149-22.752 12.235-56.246 34.395-33.481 27.198 27.94 27.212 72.444.001 100.401-21.793 22.386-56.947-10.315-34.397-33.481z"],"file-code":[384,512,[],"f1c9","M369.941 97.941l-83.882-83.882A48 48 0 0 0 252.118 0H48C21.49 0 0 21.49 0 48v416c0 26.51 21.49 48 48 48h288c26.51 0 48-21.49 48-48V131.882a48 48 0 0 0-14.059-33.941zM332.118 128H256V51.882L332.118 128zM48 464V48h160v104c0 13.255 10.745 24 24 24h104v288H48zm101.677-115.115L116.854 320l32.822-28.885a8.793 8.793 0 0 0 .605-12.624l-17.403-18.564c-3.384-3.613-8.964-3.662-12.438-.401L62.78 313.58c-3.703 3.474-3.704 9.367.001 12.84l57.659 54.055a8.738 8.738 0 0 0 6.012 2.381 8.746 8.746 0 0 0 6.427-2.782l17.403-18.563a8.795 8.795 0 0 0-.605-12.626zm84.284-127.85l-24.401-7.084a8.796 8.796 0 0 0-10.905 5.998L144.04 408.061c-1.353 4.66 1.338 9.552 5.998 10.905l24.403 7.084c4.68 1.355 9.557-1.354 10.905-5.998l54.612-188.112c1.354-4.66-1.337-9.552-5.997-10.905zm87.258 92.545l-57.658-54.055c-3.526-3.307-9.099-3.165-12.439.401l-17.403 18.563a8.795 8.795 0 0 0 .605 12.625L267.146 320l-32.822 28.885a8.793 8.793 0 0 0-.605 12.624l17.403 18.564a8.797 8.797 0 0 0 12.439.401h-.001l57.66-54.055c3.703-3.473 3.703-9.366-.001-12.839z"],"file-excel":[384,512,[],"f1c3","M369.9 97.9L286 14C277 5 264.8-.1 252.1-.1H48C21.5 0 0 21.5 0 48v416c0 26.5 21.5 48 48 48h288c26.5 0 48-21.5 48-48V131.9c0-12.7-5.1-25-14.1-34zM332.1 128H256V51.9l76.1 76.1zM48 464V48h160v104c0 13.3 10.7 24 24 24h104v288H48zm212-240h-28.8c-4.4 0-8.4 2.4-10.5 6.3-18 33.1-22.2 42.4-28.6 57.7-13.9-29.1-6.9-17.3-28.6-57.7-2.1-3.9-6.2-6.3-10.6-6.3H124c-9.3 0-15 10-10.4 18l46.3 78-46.3 78c-4.7 8 1.1 18 10.4 18h28.9c4.4 0 8.4-2.4 10.5-6.3 21.7-40 23-45 28.6-57.7 14.9 30.2 5.9 15.9 28.6 57.7 2.1 3.9 6.2 6.3 10.6 6.3H260c9.3 0 15-10 10.4-18L224 320c.7-1.1 30.3-50.5 46.3-78 4.7-8-1.1-18-10.3-18z"],"file-image":[384,512,[],"f1c5","M369.9 97.9L286 14C277 5 264.8-.1 252.1-.1H48C21.5 0 0 21.5 0 48v416c0 26.5 21.5 48 48 48h288c26.5 0 48-21.5 48-48V131.9c0-12.7-5.1-25-14.1-34zM332.1 128H256V51.9l76.1 76.1zM48 464V48h160v104c0 13.3 10.7 24 24 24h104v288H48zm32-48h224V288l-23.5-23.5c-4.7-4.7-12.3-4.7-17 0L176 352l-39.5-39.5c-4.7-4.7-12.3-4.7-17 0L80 352v64zm48-240c-26.5 0-48 21.5-48 48s21.5 48 48 48 48-21.5 48-48-21.5-48-48-48z"],"file-pdf":[384,512,[],"f1c1","M369.9 97.9L286 14C277 5 264.8-.1 252.1-.1H48C21.5 0 0 21.5 0 48v416c0 26.5 21.5 48 48 48h288c26.5 0 48-21.5 48-48V131.9c0-12.7-5.1-25-14.1-34zM332.1 128H256V51.9l76.1 76.1zM48 464V48h160v104c0 13.3 10.7 24 24 24h104v288H48zm250.2-143.7c-12.2-12-47-8.7-64.4-6.5-17.2-10.5-28.7-25-36.8-46.3 3.9-16.1 10.1-40.6 5.4-56-4.2-26.2-37.8-23.6-42.6-5.9-4.4 16.1-.4 38.5 7 67.1-10 23.9-24.9 56-35.4 74.4-20 10.3-47 26.2-51 46.2-3.3 15.8 26 55.2 76.1-31.2 22.4-7.4 46.8-16.5 68.4-20.1 18.9 10.2 41 17 55.8 17 25.5 0 28-28.2 17.5-38.7zm-198.1 77.8c5.1-13.7 24.5-29.5 30.4-35-19 30.3-30.4 35.7-30.4 35zm81.6-190.6c7.4 0 6.7 32.1 1.8 40.8-4.4-13.9-4.3-40.8-1.8-40.8zm-24.4 136.6c9.7-16.9 18-37 24.7-54.7 8.3 15.1 18.9 27.2 30.1 35.5-20.8 4.3-38.9 13.1-54.8 19.2zm131.6-5s-5 6-37.3-7.8c35.1-2.6 40.9 5.4 37.3 7.8z"],"file-powerpoint":[384,512,[],"f1c4","M369.9 97.9L286 14C277 5 264.8-.1 252.1-.1H48C21.5 0 0 21.5 0 48v416c0 26.5 21.5 48 48 48h288c26.5 0 48-21.5 48-48V131.9c0-12.7-5.1-25-14.1-34zM332.1 128H256V51.9l76.1 76.1zM48 464V48h160v104c0 13.3 10.7 24 24 24h104v288H48zm72-60V236c0-6.6 5.4-12 12-12h69.2c36.7 0 62.8 27 62.8 66.3 0 74.3-68.7 66.5-95.5 66.5V404c0 6.6-5.4 12-12 12H132c-6.6 0-12-5.4-12-12zm48.5-87.4h23c7.9 0 13.9-2.4 18.1-7.2 8.5-9.8 8.4-28.5.1-37.8-4.1-4.6-9.9-7-17.4-7h-23.9v52z"],"file-video":[384,512,[],"f1c8","M369.941 97.941l-83.882-83.882A48 48 0 0 0 252.118 0H48C21.49 0 0 21.49 0 48v416c0 26.51 21.49 48 48 48h288c26.51 0 48-21.49 48-48V131.882a48 48 0 0 0-14.059-33.941zM332.118 128H256V51.882L332.118 128zM48 464V48h160v104c0 13.255 10.745 24 24 24h104v288H48zm228.687-211.303L224 305.374V268c0-11.046-8.954-20-20-20H100c-11.046 0-20 8.954-20 20v104c0 11.046 8.954 20 20 20h104c11.046 0 20-8.954 20-20v-37.374l52.687 52.674C286.704 397.318 304 390.28 304 375.986V264.011c0-14.311-17.309-21.319-27.313-11.314z"],"file-word":[384,512,[],"f1c2","M369.9 97.9L286 14C277 5 264.8-.1 252.1-.1H48C21.5 0 0 21.5 0 48v416c0 26.5 21.5 48 48 48h288c26.5 0 48-21.5 48-48V131.9c0-12.7-5.1-25-14.1-34zM332.1 128H256V51.9l76.1 76.1zM48 464V48h160v104c0 13.3 10.7 24 24 24h104v288H48zm220.1-208c-5.7 0-10.6 4-11.7 9.5-20.6 97.7-20.4 95.4-21 103.5-.2-1.2-.4-2.6-.7-4.3-.8-5.1.3.2-23.6-99.5-1.3-5.4-6.1-9.2-11.7-9.2h-13.3c-5.5 0-10.3 3.8-11.7 9.1-24.4 99-24 96.2-24.8 103.7-.1-1.1-.2-2.5-.5-4.2-.7-5.2-14.1-73.3-19.1-99-1.1-5.6-6-9.7-11.8-9.7h-16.8c-7.8 0-13.5 7.3-11.7 14.8 8 32.6 26.7 109.5 33.2 136 1.3 5.4 6.1 9.1 11.7 9.1h25.2c5.5 0 10.3-3.7 11.6-9.1l17.9-71.4c1.5-6.2 2.5-12 3-17.3l2.9 17.3c.1.4 12.6 50.5 17.9 71.4 1.3 5.3 6.1 9.1 11.6 9.1h24.7c5.5 0 10.3-3.7 11.6-9.1 20.8-81.9 30.2-119 34.5-136 1.9-7.6-3.8-14.9-11.6-14.9h-15.8z"],flag:[512,512,[],"f024","M336.174 80c-49.132 0-93.305-32-161.913-32-31.301 0-58.303 6.482-80.721 15.168a48.04 48.04 0 0 0 2.142-20.727C93.067 19.575 74.167 1.594 51.201.104 23.242-1.71 0 20.431 0 48c0 17.764 9.657 33.262 24 41.562V496c0 8.837 7.163 16 16 16h16c8.837 0 16-7.163 16-16v-83.443C109.869 395.28 143.259 384 199.826 384c49.132 0 93.305 32 161.913 32 58.479 0 101.972-22.617 128.548-39.981C503.846 367.161 512 352.051 512 335.855V95.937c0-34.459-35.264-57.768-66.904-44.117C409.193 67.309 371.641 80 336.174 80zM464 336c-21.783 15.412-60.824 32-102.261 32-59.945 0-102.002-32-161.913-32-43.361 0-96.379 9.403-127.826 24V128c21.784-15.412 60.824-32 102.261-32 59.945 0 102.002 32 161.913 32 43.271 0 96.32-17.366 127.826-32v240z"],folder:[512,512,[],"f07b","M464 128H272l-64-64H48C21.49 64 0 85.49 0 112v288c0 26.51 21.49 48 48 48h416c26.51 0 48-21.49 48-48V176c0-26.51-21.49-48-48-48zm-6 272H54c-3.314 0-6-2.678-6-5.992V117.992A5.993 5.993 0 0 1 54 112h134.118l64 64H458a6 6 0 0 1 6 6v212a6 6 0 0 1-6 6z"],"folder-open":[576,512,[],"f07c","M527.943 224H480v-48c0-26.51-21.49-48-48-48H272l-64-64H48C21.49 64 0 85.49 0 112v288c0 26.51 21.49 48 48 48h400a48.001 48.001 0 0 0 40.704-22.56l79.942-128c19.948-31.917-3.038-73.44-40.703-73.44zM54 112h134.118l64 64H426a6 6 0 0 1 6 6v42H152a48 48 0 0 0-41.098 23.202L48 351.449V117.993A5.993 5.993 0 0 1 54 112zm394 288H72l77.234-128H528l-80 128z"],"font-awesome-logo-full":[3992,512,["Font Awesome"],"f4e6","M454.6 0H57.4C25.9 0 0 25.9 0 57.4v397.3C0 486.1 25.9 512 57.4 512h397.3c31.4 0 57.4-25.9 57.4-57.4V57.4C512 25.9 486.1 0 454.6 0zm-58.9 324.9c0 4.8-4.1 6.9-8.9 8.9-19.2 8.1-39.7 15.7-61.5 15.7-40.5 0-68.7-44.8-163.2 2.5v51.8c0 30.3-45.7 30.2-45.7 0v-250c-9-7-15-17.9-15-30.3 0-21 17.1-38.2 38.2-38.2 21 0 38.2 17.1 38.2 38.2 0 12.2-5.8 23.2-14.9 30.2v21c37.1-12 65.5-34.4 146.1-3.4 26.6 11.4 68.7-15.7 76.5-15.7 5.5 0 10.3 4.1 10.3 8.9v160.4zm432.9-174.2h-137v70.1H825c39.8 0 40.4 62.2 0 62.2H691.6v105.6c0 45.5-70.7 46.4-70.7 0V128.3c0-22 18-39.8 39.8-39.8h167.8c39.6 0 40.5 62.2.1 62.2zm191.1 23.4c-169.3 0-169.1 252.4 0 252.4 169.9 0 169.9-252.4 0-252.4zm0 196.1c-81.6 0-82.1-139.8 0-139.8 82.5 0 82.4 139.8 0 139.8zm372.4 53.4c-17.5 0-31.4-13.9-31.4-31.4v-117c0-62.4-72.6-52.5-99.1-16.4v133.4c0 41.5-63.3 41.8-63.3 0V208c0-40 63.1-41.6 63.1 0v3.4c43.3-51.6 162.4-60.4 162.4 39.3v141.5c.3 30.4-31.5 31.4-31.7 31.4zm179.7 2.9c-44.3 0-68.3-22.9-68.3-65.8V235.2H1488c-35.6 0-36.7-55.3 0-55.3h15.5v-37.3c0-41.3 63.8-42.1 63.8 0v37.5h24.9c35.4 0 35.7 55.3 0 55.3h-24.9v108.5c0 29.6 26.1 26.3 27.4 26.3 31.4 0 52.6 56.3-22.9 56.3zM1992 123c-19.5-50.2-95.5-50-114.5 0-107.3 275.7-99.5 252.7-99.5 262.8 0 42.8 58.3 51.2 72.1 14.4l13.5-35.9H2006l13 35.9c14.2 37.7 72.1 27.2 72.1-14.4 0-10.1 5.3 6.8-99.1-262.8zm-108.9 179.1l51.7-142.9 51.8 142.9h-103.5zm591.3-85.6l-53.7 176.3c-12.4 41.2-72 41-84 0l-42.3-135.9-42.3 135.9c-12.4 40.9-72 41.2-84.5 0l-54.2-176.3c-12.5-39.4 49.8-56.1 60.2-16.9L2213 342l45.3-139.5c10.9-32.7 59.6-34.7 71.2 0l45.3 139.5 39.3-142.4c10.3-38.3 72.6-23.8 60.3 16.9zm275.4 75.1c0-42.4-33.9-117.5-119.5-117.5-73.2 0-124.4 56.3-124.4 126 0 77.2 55.3 126.4 128.5 126.4 31.7 0 93-11.5 93-39.8 0-18.3-21.1-31.5-39.3-22.4-49.4 26.2-109 8.4-115.9-43.8h148.3c16.3 0 29.3-13.4 29.3-28.9zM2571 277.7c9.5-73.4 113.9-68.6 118.6 0H2571zm316.7 148.8c-31.4 0-81.6-10.5-96.6-31.9-12.4-17 2.5-39.8 21.8-39.8 16.3 0 36.8 22.9 77.7 22.9 27.4 0 40.4-11 40.4-25.8 0-39.8-142.9-7.4-142.9-102 0-40.4 35.3-75.7 98.6-75.7 31.4 0 74.1 9.9 87.6 29.4 10.8 14.8-1.4 36.2-20.9 36.2-15.1 0-26.7-17.3-66.2-17.3-22.9 0-37.8 10.5-37.8 23.8 0 35.9 142.4 6 142.4 103.1-.1 43.7-37.4 77.1-104.1 77.1zm266.8-252.4c-169.3 0-169.1 252.4 0 252.4 170.1 0 169.6-252.4 0-252.4zm0 196.1c-81.8 0-82-139.8 0-139.8 82.5 0 82.4 139.8 0 139.8zm476.9 22V268.7c0-53.8-61.4-45.8-85.7-10.5v134c0 41.3-63.8 42.1-63.8 0V268.7c0-52.1-59.5-47.4-85.7-10.1v133.6c0 41.5-63.3 41.8-63.3 0V208c0-40 63.1-41.6 63.1 0v3.4c9.9-14.4 41.8-37.3 78.6-37.3 35.3 0 57.7 16.4 66.7 43.8 13.9-21.8 45.8-43.8 82.6-43.8 44.3 0 70.7 23.4 70.7 72.7v145.3c.5 17.3-13.5 31.4-31.9 31.4 3.5.1-31.3 1.1-31.3-31.3zM3992 291.6c0-42.4-32.4-117.5-117.9-117.5-73.2 0-127.5 56.3-127.5 126 0 77.2 58.3 126.4 131.6 126.4 31.7 0 91.5-11.5 91.5-39.8 0-18.3-21.1-31.5-39.3-22.4-49.4 26.2-110.5 8.4-117.5-43.8h149.8c16.3 0 29.1-13.4 29.3-28.9zm-180.5-13.9c9.7-74.4 115.9-68.3 120.1 0h-120.1z"],frown:[496,512,[],"f119","M248 8C111 8 0 119 0 256s111 248 248 248 248-111 248-248S385 8 248 8zm0 448c-110.3 0-200-89.7-200-200S137.7 56 248 56s200 89.7 200 200-89.7 200-200 200zm-80-216c17.7 0 32-14.3 32-32s-14.3-32-32-32-32 14.3-32 32 14.3 32 32 32zm160-64c-17.7 0-32 14.3-32 32s14.3 32 32 32 32-14.3 32-32-14.3-32-32-32zm-80 128c-40.2 0-78 17.7-103.8 48.6-8.5 10.2-7.1 25.3 3.1 33.8 10.2 8.5 25.3 7.1 33.8-3.1 16.6-19.9 41-31.4 66.9-31.4s50.3 11.4 66.9 31.4c4.8 5.7 11.6 8.6 18.5 8.6 5.4 0 10.9-1.8 15.4-5.6 10.2-8.5 11.5-23.6 3.1-33.8C326 321.7 288.2 304 248 304z"],futbol:[496,512,[],"f1e3","M483.8 179.4C449.8 74.6 352.6 8 248.1 8c-25.4 0-51.2 3.9-76.7 12.2C41.2 62.5-30.1 202.4 12.2 332.6 46.2 437.4 143.4 504 247.9 504c25.4 0 51.2-3.9 76.7-12.2 130.2-42.3 201.5-182.2 159.2-312.4zm-74.5 193.7l-52.2 6.4-43.7-60.9 24.4-75.2 71.1-22.1 38.9 36.4c-.2 30.7-7.4 61.1-21.7 89.2-4.7 9.3-10.7 17.8-16.8 26.2zm0-235.4l-10.4 53.1-70.7 22-64.2-46.5V92.5l47.4-26.2c39.2 13 73.4 38 97.9 71.4zM184.9 66.4L232 92.5v73.8l-64.2 46.5-70.6-22-10.1-52.5c24.3-33.4 57.9-58.6 97.8-71.9zM139 379.5L85.9 373c-14.4-20.1-37.3-59.6-37.8-115.3l39-36.4 71.1 22.2 24.3 74.3-43.5 61.7zm48.2 67l-22.4-48.1 43.6-61.7H287l44.3 61.7-22.4 48.1c-6.2 1.8-57.6 20.4-121.7 0z"],gem:[576,512,[],"f3a5","M464 0H112c-4 0-7.8 2-10 5.4L2 152.6c-2.9 4.4-2.6 10.2.7 14.2l276 340.8c4.8 5.9 13.8 5.9 18.6 0l276-340.8c3.3-4.1 3.6-9.8.7-14.2L474.1 5.4C471.8 2 468.1 0 464 0zm-19.3 48l63.3 96h-68.4l-51.7-96h56.8zm-202.1 0h90.7l51.7 96H191l51.6-96zm-111.3 0h56.8l-51.7 96H68l63.3-96zm-43 144h51.4L208 352 88.3 192zm102.9 0h193.6L288 435.3 191.2 192zM368 352l68.2-160h51.4L368 352z"],"hand-lizard":[576,512,[],"f258","M556.686 290.542L410.328 64.829C397.001 44.272 374.417 32 349.917 32H56C25.121 32 0 57.122 0 88v8c0 44.112 35.888 80 80 80h196.042l-18.333 48H144c-48.523 0-88 39.477-88 88 0 30.879 25.121 56 56 56h131.552c2.987 0 5.914.549 8.697 1.631L352 408.418V480h224V355.829c0-23.225-6.679-45.801-19.314-65.287zM528 432H400v-23.582c0-19.948-12.014-37.508-30.604-44.736l-99.751-38.788A71.733 71.733 0 0 0 243.552 320H112c-4.411 0-8-3.589-8-8 0-22.056 17.944-40 40-40h113.709c19.767 0 37.786-12.407 44.84-30.873l24.552-64.281c8.996-23.553-8.428-48.846-33.63-48.846H80c-17.645 0-32-14.355-32-32v-8c0-4.411 3.589-8 8-8h293.917c8.166 0 15.693 4.09 20.137 10.942l146.358 225.715A71.84 71.84 0 0 1 528 355.829V432z"],"hand-paper":[448,512,[],"f256","M372.57 112.641v-10.825c0-43.612-40.52-76.691-83.039-65.546-25.629-49.5-94.09-47.45-117.982.747C130.269 26.456 89.144 57.945 89.144 102v126.13c-19.953-7.427-43.308-5.068-62.083 8.871-29.355 21.796-35.794 63.333-14.55 93.153L132.48 498.569a32 32 0 0 0 26.062 13.432h222.897c14.904 0 27.835-10.289 31.182-24.813l30.184-130.958A203.637 203.637 0 0 0 448 310.564V179c0-40.62-35.523-71.992-75.43-66.359zm27.427 197.922c0 11.731-1.334 23.469-3.965 34.886L368.707 464h-201.92L51.591 302.303c-14.439-20.27 15.023-42.776 29.394-22.605l27.128 38.079c8.995 12.626 29.031 6.287 29.031-9.283V102c0-25.645 36.571-24.81 36.571.691V256c0 8.837 7.163 16 16 16h6.856c8.837 0 16-7.163 16-16V67c0-25.663 36.571-24.81 36.571.691V256c0 8.837 7.163 16 16 16h6.856c8.837 0 16-7.163 16-16V101.125c0-25.672 36.57-24.81 36.57.691V256c0 8.837 7.163 16 16 16h6.857c8.837 0 16-7.163 16-16v-76.309c0-26.242 36.57-25.64 36.57-.691v131.563z"],"hand-peace":[448,512,[],"f25b","M362.146 191.976c-13.71-21.649-38.761-34.016-65.006-30.341V74c0-40.804-32.811-74-73.141-74-40.33 0-73.14 33.196-73.14 74L160 168l-18.679-78.85C126.578 50.843 83.85 32.11 46.209 47.208 8.735 62.238-9.571 104.963 5.008 142.85l55.757 144.927c-30.557 24.956-43.994 57.809-24.733 92.218l54.853 97.999C102.625 498.97 124.73 512 148.575 512h205.702c30.744 0 57.558-21.44 64.555-51.797l27.427-118.999a67.801 67.801 0 0 0 1.729-15.203L448 256c0-44.956-43.263-77.343-85.854-64.024zM399.987 326c0 1.488-.169 2.977-.502 4.423l-27.427 119.001c-1.978 8.582-9.29 14.576-17.782 14.576H148.575c-6.486 0-12.542-3.621-15.805-9.449l-54.854-98c-4.557-8.141-2.619-18.668 4.508-24.488l26.647-21.764a16 16 0 0 0 4.812-18.139l-64.09-166.549C37.226 92.956 84.37 74.837 96.51 106.389l59.784 155.357A16 16 0 0 0 171.227 272h11.632c8.837 0 16-7.163 16-16V74c0-34.375 50.281-34.43 50.281 0v182c0 8.837 7.163 16 16 16h6.856c8.837 0 16-7.163 16-16v-28c0-25.122 36.567-25.159 36.567 0v28c0 8.837 7.163 16 16 16h6.856c8.837 0 16-7.163 16-16 0-25.12 36.567-25.16 36.567 0v70z"],"hand-point-down":[448,512,[],"f0a7","M188.8 512c45.616 0 83.2-37.765 83.2-83.2v-35.647a93.148 93.148 0 0 0 22.064-7.929c22.006 2.507 44.978-3.503 62.791-15.985C409.342 368.1 448 331.841 448 269.299V248c0-60.063-40-98.512-40-127.2v-2.679c4.952-5.747 8-13.536 8-22.12V32c0-17.673-12.894-32-28.8-32H156.8C140.894 0 128 14.327 128 32v64c0 8.584 3.048 16.373 8 22.12v2.679c0 6.964-6.193 14.862-23.668 30.183l-.148.129-.146.131c-9.937 8.856-20.841 18.116-33.253 25.851C48.537 195.798 0 207.486 0 252.8c0 56.928 35.286 92 83.2 92 8.026 0 15.489-.814 22.4-2.176V428.8c0 45.099 38.101 83.2 83.2 83.2zm0-48c-18.7 0-35.2-16.775-35.2-35.2V270.4c-17.325 0-35.2 26.4-70.4 26.4-26.4 0-35.2-20.625-35.2-44 0-8.794 32.712-20.445 56.1-34.926 14.575-9.074 27.225-19.524 39.875-30.799 18.374-16.109 36.633-33.836 39.596-59.075h176.752C364.087 170.79 400 202.509 400 248v21.299c0 40.524-22.197 57.124-61.325 50.601-8.001 14.612-33.979 24.151-53.625 12.925-18.225 19.365-46.381 17.787-61.05 4.95V428.8c0 18.975-16.225 35.2-35.2 35.2zM328 64c0-13.255 10.745-24 24-24s24 10.745 24 24-10.745 24-24 24-24-10.745-24-24z"],"hand-point-left":[512,512,[],"f0a5","M0 220.8C0 266.416 37.765 304 83.2 304h35.647a93.148 93.148 0 0 0 7.929 22.064c-2.507 22.006 3.503 44.978 15.985 62.791C143.9 441.342 180.159 480 242.701 480H264c60.063 0 98.512-40 127.2-40h2.679c5.747 4.952 13.536 8 22.12 8h64c17.673 0 32-12.894 32-28.8V188.8c0-15.906-14.327-28.8-32-28.8h-64c-8.584 0-16.373 3.048-22.12 8H391.2c-6.964 0-14.862-6.193-30.183-23.668l-.129-.148-.131-.146c-8.856-9.937-18.116-20.841-25.851-33.253C316.202 80.537 304.514 32 259.2 32c-56.928 0-92 35.286-92 83.2 0 8.026.814 15.489 2.176 22.4H83.2C38.101 137.6 0 175.701 0 220.8zm48 0c0-18.7 16.775-35.2 35.2-35.2h158.4c0-17.325-26.4-35.2-26.4-70.4 0-26.4 20.625-35.2 44-35.2 8.794 0 20.445 32.712 34.926 56.1 9.074 14.575 19.524 27.225 30.799 39.875 16.109 18.374 33.836 36.633 59.075 39.596v176.752C341.21 396.087 309.491 432 264 432h-21.299c-40.524 0-57.124-22.197-50.601-61.325-14.612-8.001-24.151-33.979-12.925-53.625-19.365-18.225-17.787-46.381-4.95-61.05H83.2C64.225 256 48 239.775 48 220.8zM448 360c13.255 0 24 10.745 24 24s-10.745 24-24 24-24-10.745-24-24 10.745-24 24-24z"],"hand-point-right":[512,512,[],"f0a4","M428.8 137.6h-86.177a115.52 115.52 0 0 0 2.176-22.4c0-47.914-35.072-83.2-92-83.2-45.314 0-57.002 48.537-75.707 78.784-7.735 12.413-16.994 23.317-25.851 33.253l-.131.146-.129.148C135.662 161.807 127.764 168 120.8 168h-2.679c-5.747-4.952-13.536-8-22.12-8H32c-17.673 0-32 12.894-32 28.8v230.4C0 435.106 14.327 448 32 448h64c8.584 0 16.373-3.048 22.12-8h2.679c28.688 0 67.137 40 127.2 40h21.299c62.542 0 98.8-38.658 99.94-91.145 12.482-17.813 18.491-40.785 15.985-62.791A93.148 93.148 0 0 0 393.152 304H428.8c45.435 0 83.2-37.584 83.2-83.2 0-45.099-38.101-83.2-83.2-83.2zm0 118.4h-91.026c12.837 14.669 14.415 42.825-4.95 61.05 11.227 19.646 1.687 45.624-12.925 53.625 6.524 39.128-10.076 61.325-50.6 61.325H248c-45.491 0-77.21-35.913-120-39.676V215.571c25.239-2.964 42.966-21.222 59.075-39.596 11.275-12.65 21.725-25.3 30.799-39.875C232.355 112.712 244.006 80 252.8 80c23.375 0 44 8.8 44 35.2 0 35.2-26.4 53.075-26.4 70.4h158.4c18.425 0 35.2 16.5 35.2 35.2 0 18.975-16.225 35.2-35.2 35.2zM88 384c0 13.255-10.745 24-24 24s-24-10.745-24-24 10.745-24 24-24 24 10.745 24 24z"],"hand-point-up":[448,512,[],"f0a6","M105.6 83.2v86.177a115.52 115.52 0 0 0-22.4-2.176c-47.914 0-83.2 35.072-83.2 92 0 45.314 48.537 57.002 78.784 75.707 12.413 7.735 23.317 16.994 33.253 25.851l.146.131.148.129C129.807 376.338 136 384.236 136 391.2v2.679c-4.952 5.747-8 13.536-8 22.12v64c0 17.673 12.894 32 28.8 32h230.4c15.906 0 28.8-14.327 28.8-32v-64c0-8.584-3.048-16.373-8-22.12V391.2c0-28.688 40-67.137 40-127.2v-21.299c0-62.542-38.658-98.8-91.145-99.94-17.813-12.482-40.785-18.491-62.791-15.985A93.148 93.148 0 0 0 272 118.847V83.2C272 37.765 234.416 0 188.8 0c-45.099 0-83.2 38.101-83.2 83.2zm118.4 0v91.026c14.669-12.837 42.825-14.415 61.05 4.95 19.646-11.227 45.624-1.687 53.625 12.925 39.128-6.524 61.325 10.076 61.325 50.6V264c0 45.491-35.913 77.21-39.676 120H183.571c-2.964-25.239-21.222-42.966-39.596-59.075-12.65-11.275-25.3-21.725-39.875-30.799C80.712 279.645 48 267.994 48 259.2c0-23.375 8.8-44 35.2-44 35.2 0 53.075 26.4 70.4 26.4V83.2c0-18.425 16.5-35.2 35.2-35.2 18.975 0 35.2 16.225 35.2 35.2zM352 424c13.255 0 24 10.745 24 24s-10.745 24-24 24-24-10.745-24-24 10.745-24 24-24z"],"hand-pointer":[448,512,[],"f25a","M358.182 179.361c-19.493-24.768-52.679-31.945-79.872-19.098-15.127-15.687-36.182-22.487-56.595-19.629V67c0-36.944-29.736-67-66.286-67S89.143 30.056 89.143 67v161.129c-19.909-7.41-43.272-5.094-62.083 8.872-29.355 21.795-35.793 63.333-14.55 93.152l109.699 154.001C134.632 501.59 154.741 512 176 512h178.286c30.802 0 57.574-21.5 64.557-51.797l27.429-118.999A67.873 67.873 0 0 0 448 326v-84c0-46.844-46.625-79.273-89.818-62.639zM80.985 279.697l27.126 38.079c8.995 12.626 29.031 6.287 29.031-9.283V67c0-25.12 36.571-25.16 36.571 0v175c0 8.836 7.163 16 16 16h6.857c8.837 0 16-7.164 16-16v-35c0-25.12 36.571-25.16 36.571 0v35c0 8.836 7.163 16 16 16H272c8.837 0 16-7.164 16-16v-21c0-25.12 36.571-25.16 36.571 0v21c0 8.836 7.163 16 16 16h6.857c8.837 0 16-7.164 16-16 0-25.121 36.571-25.16 36.571 0v84c0 1.488-.169 2.977-.502 4.423l-27.43 119.001c-1.978 8.582-9.29 14.576-17.782 14.576H176c-5.769 0-11.263-2.878-14.697-7.697l-109.712-154c-14.406-20.223 14.994-42.818 29.394-22.606zM176.143 400v-96c0-8.837 6.268-16 14-16h6c7.732 0 14 7.163 14 16v96c0 8.837-6.268 16-14 16h-6c-7.733 0-14-7.163-14-16zm75.428 0v-96c0-8.837 6.268-16 14-16h6c7.732 0 14 7.163 14 16v96c0 8.837-6.268 16-14 16h-6c-7.732 0-14-7.163-14-16zM327 400v-96c0-8.837 6.268-16 14-16h6c7.732 0 14 7.163 14 16v96c0 8.837-6.268 16-14 16h-6c-7.732 0-14-7.163-14-16z"],"hand-rock":[512,512,[],"f255","M408.864 79.052c-22.401-33.898-66.108-42.273-98.813-23.588-29.474-31.469-79.145-31.093-108.334-.022-47.16-27.02-108.71 5.055-110.671 60.806C44.846 105.407 0 140.001 0 187.429v56.953c0 32.741 14.28 63.954 39.18 85.634l97.71 85.081c4.252 3.702 3.11 5.573 3.11 32.903 0 17.673 14.327 32 32 32h252c17.673 0 32-14.327 32-32 0-23.513-1.015-30.745 3.982-42.37l42.835-99.656c6.094-14.177 9.183-29.172 9.183-44.568V146.963c0-52.839-54.314-88.662-103.136-67.911zM464 261.406a64.505 64.505 0 0 1-5.282 25.613l-42.835 99.655c-5.23 12.171-7.883 25.04-7.883 38.25V432H188v-10.286c0-16.37-7.14-31.977-19.59-42.817l-97.71-85.08C56.274 281.255 48 263.236 48 244.381v-56.953c0-33.208 52-33.537 52 .677v41.228a16 16 0 0 0 5.493 12.067l7 6.095A16 16 0 0 0 139 235.429V118.857c0-33.097 52-33.725 52 .677v26.751c0 8.836 7.164 16 16 16h7c8.836 0 16-7.164 16-16v-41.143c0-33.134 52-33.675 52 .677v40.466c0 8.836 7.163 16 16 16h7c8.837 0 16-7.164 16-16v-27.429c0-33.03 52-33.78 52 .677v26.751c0 8.836 7.163 16 16 16h7c8.837 0 16-7.164 16-16 0-33.146 52-33.613 52 .677v114.445z"],"hand-scissors":[512,512,[],"f257","M256 480l70-.013c5.114 0 10.231-.583 15.203-1.729l118.999-27.427C490.56 443.835 512 417.02 512 386.277V180.575c0-23.845-13.03-45.951-34.005-57.69l-97.999-54.853c-34.409-19.261-67.263-5.824-92.218 24.733L142.85 37.008c-37.887-14.579-80.612 3.727-95.642 41.201-15.098 37.642 3.635 80.37 41.942 95.112L168 192l-94-9.141c-40.804 0-74 32.811-74 73.14 0 40.33 33.196 73.141 74 73.141h87.635c-3.675 26.245 8.692 51.297 30.341 65.006C178.657 436.737 211.044 480 256 480zm0-48.013c-25.16 0-25.12-36.567 0-36.567 8.837 0 16-7.163 16-16v-6.856c0-8.837-7.163-16-16-16h-28c-25.159 0-25.122-36.567 0-36.567h28c8.837 0 16-7.163 16-16v-6.856c0-8.837-7.163-16-16-16H74c-34.43 0-34.375-50.281 0-50.281h182c8.837 0 16-7.163 16-16v-11.632a16 16 0 0 0-10.254-14.933L106.389 128.51c-31.552-12.14-13.432-59.283 19.222-46.717l166.549 64.091a16.001 16.001 0 0 0 18.139-4.812l21.764-26.647c5.82-7.127 16.348-9.064 24.488-4.508l98 54.854c5.828 3.263 9.449 9.318 9.449 15.805v205.701c0 8.491-5.994 15.804-14.576 17.782l-119.001 27.427a19.743 19.743 0 0 1-4.423.502h-70z"],"hand-spock":[512,512,[],"f259","M21.096 381.79l129.092 121.513a32 32 0 0 0 21.932 8.698h237.6c14.17 0 26.653-9.319 30.68-22.904l31.815-107.313A115.955 115.955 0 0 0 477 348.811v-36.839c0-4.051.476-8.104 1.414-12.045l31.73-133.41c10.099-42.412-22.316-82.738-65.544-82.525-4.144-24.856-22.543-47.165-49.85-53.992-35.803-8.952-72.227 12.655-81.25 48.75L296.599 184 274.924 52.01c-8.286-36.07-44.303-58.572-80.304-50.296-29.616 6.804-50.138 32.389-51.882 61.295-42.637.831-73.455 40.563-64.071 81.844l31.04 136.508c-27.194-22.515-67.284-19.992-91.482 5.722-25.376 26.961-24.098 69.325 2.871 94.707zm32.068-61.811l.002-.001c7.219-7.672 19.241-7.98 26.856-.813l53.012 49.894C143.225 378.649 160 371.4 160 357.406v-69.479c0-1.193-.134-2.383-.397-3.546l-34.13-150.172c-5.596-24.617 31.502-32.86 37.054-8.421l30.399 133.757a16 16 0 0 0 15.603 12.454h8.604c10.276 0 17.894-9.567 15.594-19.583l-41.62-181.153c-5.623-24.469 31.39-33.076 37.035-8.508l45.22 196.828A16 16 0 0 0 288.956 272h13.217a16 16 0 0 0 15.522-12.119l42.372-169.49c6.104-24.422 42.962-15.159 36.865 9.217L358.805 252.12c-2.521 10.088 5.115 19.88 15.522 19.88h9.694a16 16 0 0 0 15.565-12.295L426.509 146.6c5.821-24.448 42.797-15.687 36.966 8.802L431.72 288.81a100.094 100.094 0 0 0-2.72 23.162v36.839c0 6.548-.943 13.051-2.805 19.328L397.775 464h-219.31L53.978 346.836c-7.629-7.18-7.994-19.229-.814-26.857z"],handshake:[640,512,[],"f2b5","M519.2 127.9l-47.6-47.6A56.252 56.252 0 0 0 432 64H205.2c-14.8 0-29.1 5.9-39.6 16.3L118 127.9H0v255.7h64c17.6 0 31.8-14.2 31.9-31.7h9.1l84.6 76.4c30.9 25.1 73.8 25.7 105.6 3.8 12.5 10.8 26 15.9 41.1 15.9 18.2 0 35.3-7.4 48.8-24 22.1 8.7 48.2 2.6 64-16.8l26.2-32.3c5.6-6.9 9.1-14.8 10.9-23h57.9c.1 17.5 14.4 31.7 31.9 31.7h64V127.9H519.2zM48 351.6c-8.8 0-16-7.2-16-16s7.2-16 16-16 16 7.2 16 16c0 8.9-7.2 16-16 16zm390-6.9l-26.1 32.2c-2.8 3.4-7.8 4-11.3 1.2l-23.9-19.4-30 36.5c-6 7.3-15 4.8-18 2.4l-36.8-31.5-15.6 19.2c-13.9 17.1-39.2 19.7-55.3 6.6l-97.3-88H96V175.8h41.9l61.7-61.6c2-.8 3.7-1.5 5.7-2.3H262l-38.7 35.5c-29.4 26.9-31.1 72.3-4.4 101.3 14.8 16.2 61.2 41.2 101.5 4.4l8.2-7.5 108.2 87.8c3.4 2.8 3.9 7.9 1.2 11.3zm106-40.8h-69.2c-2.3-2.8-4.9-5.4-7.7-7.7l-102.7-83.4 12.5-11.4c6.5-6 7-16.1 1-22.6L367 167.1c-6-6.5-16.1-6.9-22.6-1l-55.2 50.6c-9.5 8.7-25.7 9.4-34.6 0-9.3-9.9-8.5-25.1 1.2-33.9l65.6-60.1c7.4-6.8 17-10.5 27-10.5l83.7-.2c2.1 0 4.1.8 5.5 2.3l61.7 61.6H544v128zm48 47.7c-8.8 0-16-7.2-16-16s7.2-16 16-16 16 7.2 16 16c0 8.9-7.2 16-16 16z"],hdd:[576,512,[],"f0a0","M567.403 235.642L462.323 84.589A48 48 0 0 0 422.919 64H153.081a48 48 0 0 0-39.404 20.589L8.597 235.642A48.001 48.001 0 0 0 0 263.054V400c0 26.51 21.49 48 48 48h480c26.51 0 48-21.49 48-48V263.054c0-9.801-3-19.366-8.597-27.412zM153.081 112h269.838l77.913 112H75.168l77.913-112zM528 400H48V272h480v128zm-32-64c0 17.673-14.327 32-32 32s-32-14.327-32-32 14.327-32 32-32 32 14.327 32 32zm-96 0c0 17.673-14.327 32-32 32s-32-14.327-32-32 14.327-32 32-32 32 14.327 32 32z"],heart:[512,512,[],"f004","M458.4 64.3C400.6 15.7 311.3 23 256 79.3 200.7 23 111.4 15.6 53.6 64.3-21.6 127.6-10.6 230.8 43 285.5l175.4 178.7c10 10.2 23.4 15.9 37.6 15.9 14.3 0 27.6-5.6 37.6-15.8L469 285.6c53.5-54.7 64.7-157.9-10.6-221.3zm-23.6 187.5L259.4 430.5c-2.4 2.4-4.4 2.4-6.8 0L77.2 251.8c-36.5-37.2-43.9-107.6 7.3-150.7 38.9-32.7 98.9-27.8 136.5 10.5l35 35.7 35-35.7c37.8-38.5 97.8-43.2 136.5-10.6 51.1 43.1 43.5 113.9 7.3 150.8z"],hospital:[448,512,[],"f0f8","M128 244v-40c0-6.627 5.373-12 12-12h40c6.627 0 12 5.373 12 12v40c0 6.627-5.373 12-12 12h-40c-6.627 0-12-5.373-12-12zm140 12h40c6.627 0 12-5.373 12-12v-40c0-6.627-5.373-12-12-12h-40c-6.627 0-12 5.373-12 12v40c0 6.627 5.373 12 12 12zm-76 84v-40c0-6.627-5.373-12-12-12h-40c-6.627 0-12 5.373-12 12v40c0 6.627 5.373 12 12 12h40c6.627 0 12-5.373 12-12zm76 12h40c6.627 0 12-5.373 12-12v-40c0-6.627-5.373-12-12-12h-40c-6.627 0-12 5.373-12 12v40c0 6.627 5.373 12 12 12zm180 124v36H0v-36c0-6.627 5.373-12 12-12h19.5V85.035C31.5 73.418 42.245 64 55.5 64H144V24c0-13.255 10.745-24 24-24h112c13.255 0 24 10.745 24 24v40h88.5c13.255 0 24 9.418 24 21.035V464H436c6.627 0 12 5.373 12 12zM79.5 463H192v-67c0-6.627 5.373-12 12-12h40c6.627 0 12 5.373 12 12v67h112.5V112H304v24c0 13.255-10.745 24-24 24H168c-13.255 0-24-10.745-24-24v-24H79.5v351zM266 64h-26V38a6 6 0 0 0-6-6h-20a6 6 0 0 0-6 6v26h-26a6 6 0 0 0-6 6v20a6 6 0 0 0 6 6h26v26a6 6 0 0 0 6 6h20a6 6 0 0 0 6-6V96h26a6 6 0 0 0 6-6V70a6 6 0 0 0-6-6z"],hourglass:[384,512,[],"f254","M368 48h4c6.627 0 12-5.373 12-12V12c0-6.627-5.373-12-12-12H12C5.373 0 0 5.373 0 12v24c0 6.627 5.373 12 12 12h4c0 80.564 32.188 165.807 97.18 208C47.899 298.381 16 383.9 16 464h-4c-6.627 0-12 5.373-12 12v24c0 6.627 5.373 12 12 12h360c6.627 0 12-5.373 12-12v-24c0-6.627-5.373-12-12-12h-4c0-80.564-32.188-165.807-97.18-208C336.102 213.619 368 128.1 368 48zM64 48h256c0 101.62-57.307 184-128 184S64 149.621 64 48zm256 416H64c0-101.62 57.308-184 128-184s128 82.38 128 184z"],"id-badge":[384,512,[],"f2c1","M336 0H48C21.5 0 0 21.5 0 48v416c0 26.5 21.5 48 48 48h288c26.5 0 48-21.5 48-48V48c0-26.5-21.5-48-48-48zm0 464H48V48h288v416zM144 112h96c8.8 0 16-7.2 16-16s-7.2-16-16-16h-96c-8.8 0-16 7.2-16 16s7.2 16 16 16zm48 176c35.3 0 64-28.7 64-64s-28.7-64-64-64-64 28.7-64 64 28.7 64 64 64zm-89.6 128h179.2c12.4 0 22.4-8.6 22.4-19.2v-19.2c0-31.8-30.1-57.6-67.2-57.6-10.8 0-18.7 8-44.8 8-26.9 0-33.4-8-44.8-8-37.1 0-67.2 25.8-67.2 57.6v19.2c0 10.6 10 19.2 22.4 19.2z"],"id-card":[576,512,[],"f2c2","M528 32H48C21.5 32 0 53.5 0 80v352c0 26.5 21.5 48 48 48h480c26.5 0 48-21.5 48-48V80c0-26.5-21.5-48-48-48zm0 400H303.2c.9-4.5.8 3.6.8-22.4 0-31.8-30.1-57.6-67.2-57.6-10.8 0-18.7 8-44.8 8-26.9 0-33.4-8-44.8-8-37.1 0-67.2 25.8-67.2 57.6 0 26-.2 17.9.8 22.4H48V144h480v288zm-168-80h112c4.4 0 8-3.6 8-8v-16c0-4.4-3.6-8-8-8H360c-4.4 0-8 3.6-8 8v16c0 4.4 3.6 8 8 8zm0-64h112c4.4 0 8-3.6 8-8v-16c0-4.4-3.6-8-8-8H360c-4.4 0-8 3.6-8 8v16c0 4.4 3.6 8 8 8zm0-64h112c4.4 0 8-3.6 8-8v-16c0-4.4-3.6-8-8-8H360c-4.4 0-8 3.6-8 8v16c0 4.4 3.6 8 8 8zm-168 96c35.3 0 64-28.7 64-64s-28.7-64-64-64-64 28.7-64 64 28.7 64 64 64z"],image:[512,512,[],"f03e","M464 64H48C21.49 64 0 85.49 0 112v288c0 26.51 21.49 48 48 48h416c26.51 0 48-21.49 48-48V112c0-26.51-21.49-48-48-48zm-6 336H54a6 6 0 0 1-6-6V118a6 6 0 0 1 6-6h404a6 6 0 0 1 6 6v276a6 6 0 0 1-6 6zM128 152c-22.091 0-40 17.909-40 40s17.909 40 40 40 40-17.909 40-40-17.909-40-40-40zM96 352h320v-80l-87.515-87.515c-4.686-4.686-12.284-4.686-16.971 0L192 304l-39.515-39.515c-4.686-4.686-12.284-4.686-16.971 0L96 304v48z"],images:[576,512,[],"f302","M480 416v16c0 26.51-21.49 48-48 48H48c-26.51 0-48-21.49-48-48V176c0-26.51 21.49-48 48-48h16v48H54a6 6 0 0 0-6 6v244a6 6 0 0 0 6 6h372a6 6 0 0 0 6-6v-10h48zm42-336H150a6 6 0 0 0-6 6v244a6 6 0 0 0 6 6h372a6 6 0 0 0 6-6V86a6 6 0 0 0-6-6zm6-48c26.51 0 48 21.49 48 48v256c0 26.51-21.49 48-48 48H144c-26.51 0-48-21.49-48-48V80c0-26.51 21.49-48 48-48h384zM264 144c0 22.091-17.909 40-40 40s-40-17.909-40-40 17.909-40 40-40 40 17.909 40 40zm-72 96l39.515-39.515c4.686-4.686 12.284-4.686 16.971 0L288 240l103.515-103.515c4.686-4.686 12.284-4.686 16.971 0L480 208v80H192v-48z"],keyboard:[576,512,[],"f11c","M528 64H48C21.49 64 0 85.49 0 112v288c0 26.51 21.49 48 48 48h480c26.51 0 48-21.49 48-48V112c0-26.51-21.49-48-48-48zm8 336c0 4.411-3.589 8-8 8H48c-4.411 0-8-3.589-8-8V112c0-4.411 3.589-8 8-8h480c4.411 0 8 3.589 8 8v288zM170 270v-28c0-6.627-5.373-12-12-12h-28c-6.627 0-12 5.373-12 12v28c0 6.627 5.373 12 12 12h28c6.627 0 12-5.373 12-12zm96 0v-28c0-6.627-5.373-12-12-12h-28c-6.627 0-12 5.373-12 12v28c0 6.627 5.373 12 12 12h28c6.627 0 12-5.373 12-12zm96 0v-28c0-6.627-5.373-12-12-12h-28c-6.627 0-12 5.373-12 12v28c0 6.627 5.373 12 12 12h28c6.627 0 12-5.373 12-12zm96 0v-28c0-6.627-5.373-12-12-12h-28c-6.627 0-12 5.373-12 12v28c0 6.627 5.373 12 12 12h28c6.627 0 12-5.373 12-12zm-336 82v-28c0-6.627-5.373-12-12-12H82c-6.627 0-12 5.373-12 12v28c0 6.627 5.373 12 12 12h28c6.627 0 12-5.373 12-12zm384 0v-28c0-6.627-5.373-12-12-12h-28c-6.627 0-12 5.373-12 12v28c0 6.627 5.373 12 12 12h28c6.627 0 12-5.373 12-12zM122 188v-28c0-6.627-5.373-12-12-12H82c-6.627 0-12 5.373-12 12v28c0 6.627 5.373 12 12 12h28c6.627 0 12-5.373 12-12zm96 0v-28c0-6.627-5.373-12-12-12h-28c-6.627 0-12 5.373-12 12v28c0 6.627 5.373 12 12 12h28c6.627 0 12-5.373 12-12zm96 0v-28c0-6.627-5.373-12-12-12h-28c-6.627 0-12 5.373-12 12v28c0 6.627 5.373 12 12 12h28c6.627 0 12-5.373 12-12zm96 0v-28c0-6.627-5.373-12-12-12h-28c-6.627 0-12 5.373-12 12v28c0 6.627 5.373 12 12 12h28c6.627 0 12-5.373 12-12zm96 0v-28c0-6.627-5.373-12-12-12h-28c-6.627 0-12 5.373-12 12v28c0 6.627 5.373 12 12 12h28c6.627 0 12-5.373 12-12zm-98 158v-16c0-6.627-5.373-12-12-12H180c-6.627 0-12 5.373-12 12v16c0 6.627 5.373 12 12 12h216c6.627 0 12-5.373 12-12z"],lemon:[512,512,[],"f094","M484.112 27.889C455.989-.233 416.108-8.057 387.059 8.865 347.604 31.848 223.504-41.111 91.196 91.197-41.277 223.672 31.923 347.472 8.866 387.058c-16.922 29.051-9.1 68.932 19.022 97.054 28.135 28.135 68.011 35.938 97.057 19.021 39.423-22.97 163.557 49.969 295.858-82.329 132.474-132.477 59.273-256.277 82.331-295.861 16.922-29.05 9.1-68.931-19.022-97.054zm-22.405 72.894c-38.8 66.609 45.6 165.635-74.845 286.08-120.44 120.443-219.475 36.048-286.076 74.843-22.679 13.207-64.035-27.241-50.493-50.488 38.8-66.609-45.6-165.635 74.845-286.08C245.573 4.702 344.616 89.086 411.219 50.292c22.73-13.24 64.005 27.288 50.488 50.491zm-169.861 8.736c1.37 10.96-6.404 20.957-17.365 22.327-54.846 6.855-135.779 87.787-142.635 142.635-1.373 10.989-11.399 18.734-22.326 17.365-10.961-1.37-18.735-11.366-17.365-22.326 9.162-73.286 104.167-168.215 177.365-177.365 10.953-1.368 20.956 6.403 22.326 17.364z"],"life-ring":[512,512,[],"f1cd","M256 504c136.967 0 248-111.033 248-248S392.967 8 256 8 8 119.033 8 256s111.033 248 248 248zm-103.398-76.72l53.411-53.411c31.806 13.506 68.128 13.522 99.974 0l53.411 53.411c-63.217 38.319-143.579 38.319-206.796 0zM336 256c0 44.112-35.888 80-80 80s-80-35.888-80-80 35.888-80 80-80 80 35.888 80 80zm91.28 103.398l-53.411-53.411c13.505-31.806 13.522-68.128 0-99.974l53.411-53.411c38.319 63.217 38.319 143.579 0 206.796zM359.397 84.72l-53.411 53.411c-31.806-13.505-68.128-13.522-99.973 0L152.602 84.72c63.217-38.319 143.579-38.319 206.795 0zM84.72 152.602l53.411 53.411c-13.506 31.806-13.522 68.128 0 99.974L84.72 359.398c-38.319-63.217-38.319-143.579 0-206.796z"],lightbulb:[384,512,[],"f0eb","M272 428v28c0 10.449-6.68 19.334-16 22.629V488c0 13.255-10.745 24-24 24h-80c-13.255 0-24-10.745-24-24v-9.371c-9.32-3.295-16-12.18-16-22.629v-28c0-6.627 5.373-12 12-12h136c6.627 0 12 5.373 12 12zM128 176c0-35.29 28.71-64 64-64 8.837 0 16-7.164 16-16s-7.163-16-16-16c-52.935 0-96 43.065-96 96 0 8.836 7.164 16 16 16s16-7.164 16-16zm64-128c70.734 0 128 57.254 128 128 0 77.602-37.383 60.477-80.98 160h-94.04C101.318 236.33 64 253.869 64 176c0-70.735 57.254-128 128-128m0-48C94.805 0 16 78.803 16 176c0 101.731 51.697 91.541 90.516 192.674 3.55 9.249 12.47 15.326 22.376 15.326h126.215c9.906 0 18.826-6.078 22.376-15.326C316.303 267.541 368 277.731 368 176 368 78.803 289.195 0 192 0z"],"list-alt":[512,512,[],"f022","M464 32H48C21.49 32 0 53.49 0 80v352c0 26.51 21.49 48 48 48h416c26.51 0 48-21.49 48-48V80c0-26.51-21.49-48-48-48zm-6 400H54a6 6 0 0 1-6-6V86a6 6 0 0 1 6-6h404a6 6 0 0 1 6 6v340a6 6 0 0 1-6 6zm-42-92v24c0 6.627-5.373 12-12 12H204c-6.627 0-12-5.373-12-12v-24c0-6.627 5.373-12 12-12h200c6.627 0 12 5.373 12 12zm0-96v24c0 6.627-5.373 12-12 12H204c-6.627 0-12-5.373-12-12v-24c0-6.627 5.373-12 12-12h200c6.627 0 12 5.373 12 12zm0-96v24c0 6.627-5.373 12-12 12H204c-6.627 0-12-5.373-12-12v-24c0-6.627 5.373-12 12-12h200c6.627 0 12 5.373 12 12zm-252 12c0 19.882-16.118 36-36 36s-36-16.118-36-36 16.118-36 36-36 36 16.118 36 36zm0 96c0 19.882-16.118 36-36 36s-36-16.118-36-36 16.118-36 36-36 36 16.118 36 36zm0 96c0 19.882-16.118 36-36 36s-36-16.118-36-36 16.118-36 36-36 36 16.118 36 36z"],map:[576,512,[],"f279","M508.505 36.17L381.517 92.576 207.179 34.463a47.992 47.992 0 0 0-34.674 1.674l-144 64A48 48 0 0 0 0 144v287.967c0 34.938 35.991 57.864 67.4
```
