## Supplementary material for "Characterizing and inferring quantitative cell cycle phase in single-cell RNA-seq data analysis": Source code of our analysis, including scripts and data necessary to reproduce the work.: jquery.min.js

```
/*! jQuery v1.12.4 | (c) jQuery Foundation | jquery.org/license */
!function(a,b){"object"==typeof module&&"object"==typeof module.exports?module.exports=a.document?b(a,!0):function(a){if(!a.document)throw new Error("jQuery requires a window with a document");return b(a)}:b(a)}("undefined"!=typeof window?window:this,function(a,b){var c=[],d=a.document,e=c.slice,f=c.concat,g=c.push,h=c.indexOf,i={},j=i.toString,k=i.hasOwnProperty,l={},m="1.12.4",n=function(a,b){return new n.fn.init(a,b)},o=/^[\s\uFEFF\xA0]+|[\s\uFEFF\xA0]+$/g,p=/^-ms-/,q=/-([\da-z])/gi,r=function(a,b){return b.toUpperCase()};n.fn=n.prototype={jquery:m,constructor:n,selector:"",length:0,toArray:function(){return e.call(this)},get:function(a){return null!=a?0>a?this[a+this.length]:this[a]:e.call(this)},pushStack:function(a){var b=n.merge(this.constructor(),a);return b.prevObject=this,b.context=this.context,b},each:function(a){return n.each(this,a)},map:function(a){return this.pushStack(n.map(this,function(b,c){return a.call(b,c,b)}))},slice:function(){return this.pushStack(e.apply(this,arguments))},first:function(){return this.eq(0)},last:function(){return this.eq(-1)},eq:function(a){var b=this.length,c=+a+(0>a?b:0);return this.pushStack(c>=0&&b>c?[this[c]]:[])},end:function(){return this.prevObject||this.constructor()},push:g,sort:c.sort,splice:c.splice},n.extend=n.fn.extend=function(){var a,b,c,d,e,f,g=arguments[0]||{},h=1,i=arguments.length,j=!1;for("boolean"==typeof g&&(j=g,g=arguments[h]||{},h++),"object"==typeof g||n.isFunction(g)||(g={}),h===i&&(g=this,h--);i>h;h++)if(null!=(e=arguments[h]))for(d in e)a=g[d],c=e[d],g!==c&&(j&&c&&(n.isPlainObject(c)||(b=n.isArray(c)))?(b?(b=!1,f=a&&n.isArray(a)?a:[]):f=a&&n.isPlainObject(a)?a:{},g[d]=n.extend(j,f,c)):void 0!==c&&(g[d]=c));return g},n.extend({expando:"jQuery"+(m+Math.random()).replace(/\D/g,""),isReady:!0,error:function(a){throw new Error(a)},noop:function(){},isFunction:function(a){return"function"===n.type(a)},isArray:Array.isArray||function(a){return"array"===n.type(a)},isWindow:function(a){return null!=a&&a==a.window},isNumeric:function(a){var b=a&&a.toString();return!n.isArray(a)&&b-parseFloat(b)+1>=0},isEmptyObject:function(a){var b;for(b in a)return!1;return!0},isPlainObject:function(a){var b;if(!a||"object"!==n.type(a)||a.nodeType||n.isWindow(a))return!1;try{if(a.constructor&&!k.call(a,"constructor")&&!k.call(a.constructor.prototype,"isPrototypeOf"))return!1}catch(c){return!1}if(!l.ownFirst)for(b in a)return k.call(a,b);for(b in a);return void 0===b||k.call(a,b)},type:function(a){return null==a?a+"":"object"==typeof a||"function"==typeof a?i[j.call(a)]||"object":typeof a},globalEval:function(b){b&&n.trim(b)&&(a.execScript||function(b){a.eval.call(a,b)})(b)},camelCase:function(a){return a.replace(p,"ms-").replace(q,r)},nodeName:function(a,b){return a.nodeName&&a.nodeName.toLowerCase()===b.toLowerCase()},each:function(a,b){var c,d=0;if(s(a)){for(c=a.length;c>d;d++)if(b.call(a[d],d,a[d])===!1)break}else for(d in a)if(b.call(a[d],d,a[d])===!1)break;return a},trim:function(a){return null==a?"":(a+"").replace(o,"")},makeArray:function(a,b){var c=b||[];return null!=a&&(s(Object(a))?n.merge(c,"string"==typeof a?[a]:a):g.call(c,a)),c},inArray:function(a,b,c){var d;if(b){if(h)return h.call(b,a,c);for(d=b.length,c=c?0>c?Math.max(0,d+c):c:0;d>c;c++)if(c in b&&b[c]===a)return c}return-1},merge:function(a,b){var c=+b.length,d=0,e=a.length;while(c>d)a[e++]=b[d++];if(c!==c)while(void 0!==b[d])a[e++]=b[d++];return a.length=e,a},grep:function(a,b,c){for(var d,e=[],f=0,g=a.length,h=!c;g>f;f++)d=!b(a[f],f),d!==h&&e.push(a[f]);return e},map:function(a,b,c){var d,e,g=0,h=[];if(s(a))for(d=a.length;d>g;g++)e=b(a[g],g,c),null!=e&&h.push(e);else for(g in a)e=b(a[g],g,c),null!=e&&h.push(e);return f.apply([],h)},guid:1,proxy:function(a,b){var c,d,f;return"string"==typeof b&&(f=a[b],b=a,a=f),n.isFunction(a)?(c=e.call(arguments,2),d=function(){return a.apply(b||this,c.concat(e.call(arguments)))},d.guid=a.guid=a.guid||n.guid++,d):void 0},now:function(){return+new Date},support:l}),"function"==typeof Symbol&&(n.fn[Symbol.iterator]=c[Symbol.iterator]),n.each("Boolean Number String Function Array Date RegExp Object Error Symbol".split(" "),function(a,b){i["[object "+b+"]"]=b.toLowerCase()});function s(a){var b=!!a&&"length"in a&&a.length,c=n.type(a);return"function"===c||n.isWindow(a)?!1:"array"===c||0===b||"number"==typeof b&&b>0&&b-1 in a}var t=function(a){var b,c,d,e,f,g,h,i,j,k,l,m,n,o,p,q,r,s,t,u="sizzle"+1*new Date,v=a.document,w=0,x=0,y=ga(),z=ga(),A=ga(),B=function(a,b){return a===b&&(l=!0),0},C=1<<31,D={}.hasOwnProperty,E=[],F=E.pop,G=E.push,H=E.push,I=E.slice,J=function(a,b){for(var c=0,d=a.length;d>c;c++)if(a[c]===b)return c;return-1},K="checked|selected|async|autofocus|autoplay|controls|defer|disabled|hidden|ismap|loop|multiple|open|readonly|required|scoped",L="[\\x20\\t\\r\\n\\f]",M="(?:\\\\.|[\\w-]|[^\\x00-\\xa0])+",N="\\["+L+"*("+M+")(?:"+L+"*([*^$|!~]?=)"+L+"*(?:'((?:\\\\.|[^\\\\'])*)'|\"((?:\\\\.|[^\\\\\"])*)\"|("+M+"))|)"+L+"*\\]",O=":("+M+")(?:\\((('((?:\\\\.|[^\\\\'])*)'|\"((?:\\\\.|[^\\\\\"])*)\")|((?:\\\\.|[^\\\\()[\\]]|"+N+")*)|.*)\\)|)",P=new RegExp(L+"+","g"),Q=new RegExp("^"+L+"+|((?:^|[^\\\\])(?:\\\\.)*)"+L+"+$","g"),R=new RegExp("^"+L+"*,"+L+"*"),S=new RegExp("^"+L+"*([>+~]|"+L+")"+L+"*"),T=new RegExp("="+L+"*([^\\]'\"]*?)"+L+"*\\]","g"),U=new RegExp(O),V=new RegExp("^"+M+"$"),W={ID:new RegExp("^#("+M+")"),CLASS:new RegExp("^\\.("+M+")"),TAG:new RegExp("^("+M+"|[*])"),ATTR:new RegExp("^"+N),PSEUDO:new RegExp("^"+O),CHILD:new RegExp("^:(only|first|last|nth|nth-last)-(child|of-type)(?:\\("+L+"*(even|odd|(([+-]|)(\\d*)n|)"+L+"*(?:([+-]|)"+L+"*(\\d+)|))"+L+"*\\)|)","i"),bool:new RegExp("^(?:"+K+")$","i"),needsContext:new RegExp("^"+L+"*[>+~]|:(even|odd|eq|gt|lt|nth|first|last)(?:\\("+L+"*((?:-\\d)?\\d*)"+L+"*\\)|)(?=[^-]|$)","i")},X=/^(?:input|select|textarea|button)$/i,Y=/^h\d$/i,Z=/^[^{]+\{\s*\[native \w/,$=/^(?:#([\w-]+)|(\w+)|\.([\w-]+))$/,_=/[+~]/,aa=/'|\\/g,ba=new RegExp("\\\\([\\da-f]{1,6}"+L+"?|("+L+")|.)","ig"),ca=function(a,b,c){var d="0x"+b-65536;return d!==d||c?b:0>d?String.fromCharCode(d+65536):String.fromCharCode(d>>10|55296,1023&d|56320)},da=function(){m()};try{H.apply(E=I.call(v.childNodes),v.childNodes),E[v.childNodes.length].nodeType}catch(ea){H={apply:E.length?function(a,b){G.apply(a,I.call(b))}:function(a,b){var c=a.length,d=0;while(a[c++]=b[d++]);a.length=c-1}}}function fa(a,b,d,e){var f,h,j,k,l,o,r,s,w=b&&b.ownerDocument,x=b?b.nodeType:9;if(d=d||[],"string"!=typeof a||!a||1!==x&&9!==x&&11!==x)return d;if(!e&&((b?b.ownerDocument||b:v)!==n&&m(b),b=b||n,p)){if(11!==x&&(o=$.exec(a)))if(f=o[1]){if(9===x){if(!(j=b.getElementById(f)))return d;if(j.id===f)return d.push(j),d}else if(w&&(j=w.getElementById(f))&&t(b,j)&&j.id===f)return d.push(j),d}else{if(o[2])return H.apply(d,b.getElementsByTagName(a)),d;if((f=o[3])&&c.getElementsByClassName&&b.getElementsByClassName)return H.apply(d,b.getElementsByClassName(f)),d}if(c.qsa&&!A[a+" "]&&(!q||!q.test(a))){if(1!==x)w=b,s=a;else if("object"!==b.nodeName.toLowerCase()){(k=b.getAttribute("id"))?k=k.replace(aa,"\\$&"):b.setAttribute("id",k=u),r=g(a),h=r.length,l=V.test(k)?"#"+k:"[id='"+k+"']";while(h--)r[h]=l+" "+qa(r[h]);s=r.join(","),w=_.test(a)&&oa(b.parentNode)||b}if(s)try{return H.apply(d,w.querySelectorAll(s)),d}catch(y){}finally{k===u&&b.removeAttribute("id")}}}return i(a.replace(Q,"$1"),b,d,e)}function ga(){var a=[];function b(c,e){return a.push(c+" ")>d.cacheLength&&delete b[a.shift()],b[c+" "]=e}return b}function ha(a){return a[u]=!0,a}function ia(a){var b=n.createElement("div");try{return!!a(b)}catch(c){return!1}finally{b.parentNode&&b.parentNode.removeChild(b),b=null}}function ja(a,b){var c=a.split("|"),e=c.length;while(e--)d.attrHandle[c[e]]=b}function ka(a,b){var c=b&&a,d=c&&1===a.nodeType&&1===b.nodeType&&(~b.sourceIndex||C)-(~a.sourceIndex||C);if(d)return d;if(c)while(c=c.nextSibling)if(c===b)return-1;return a?1:-1}function la(a){return function(b){var c=b.nodeName.toLowerCase();return"input"===c&&b.type===a}}function ma(a){return function(b){var c=b.nodeName.toLowerCase();return("input"===c||"button"===c)&&b.type===a}}function na(a){return ha(function(b){return b=+b,ha(function(c,d){var e,f=a([],c.length,b),g=f.length;while(g--)c[e=f[g]]&&(c[e]=!(d[e]=c[e]))})})}function oa(a){return a&&"undefined"!=typeof a.getElementsByTagName&&a}c=fa.support={},f=fa.isXML=function(a){var b=a&&(a.ownerDocument||a).documentElement;return b?"HTML"!==b.nodeName:!1},m=fa.setDocument=function(a){var b,e,g=a?a.ownerDocument||a:v;return g!==n&&9===g.nodeType&&g.documentElement?(n=g,o=n.documentElement,p=!f(n),(e=n.defaultView)&&e.top!==e&&(e.addEventListener?e.addEventListener("unload",da,!1):e.attachEvent&&e.attachEvent("onunload",da)),c.attributes=ia(function(a){return a.className="i",!a.getAttribute("className")}),c.getElementsByTagName=ia(function(a){return a.appendChild(n.createComment("")),!a.getElementsByTagName("*").length}),c.getElementsByClassName=Z.test(n.getElementsByClassName),c.getById=ia(function(a){return o.appendChild(a).id=u,!n.getElementsByName||!n.getElementsByName(u).length}),c.getById?(d.find.ID=function(a,b){if("undefined"!=typeof b.getElementById&&p){var c=b.getElementById(a);return c?[c]:[]}},d.filter.ID=function(a){var b=a.replace(ba,ca);return function(a){return a.getAttribute("id")===b}}):(delete d.find.ID,d.filter.ID=function(a){var b=a.replace(ba,ca);return function(a){var c="undefined"!=typeof a.getAttributeNode&&a.getAttributeNode("id");return c&&c.value===b}}),d.find.TAG=c.getElementsByTagName?function(a,b){return"undefined"!=typeof b.getElementsByTagName?b.getElementsByTagName(a):c.qsa?b.querySelectorAll(a):void 0}:function(a,b){var c,d=[],e=0,f=b.getElementsByTagName(a);if("*"===a){while(c=f[e++])1===c.nodeType&&d.push(c);return d}return f},d.find.CLASS=c.getElementsByClassName&&function(a,b){return"undefined"!=typeof b.getElementsByClassName&&p?b.getElementsByClassName(a):void 0},r=[],q=[],(c.qsa=Z.test(n.querySelectorAll))&&(ia(function(a){o.appendChild(a).innerHTML="<a id='"+u+"'></a><select id='"+u+"-\r\\' msallowcapture=''><option selected=''></option></select>",a.querySelectorAll("[msallowcapture^='']").length&&q.push("[*^$]="+L+"*(?:''|\"\")"),a.querySelectorAll("[selected]").length||q.push("\\["+L+"*(?:value|"+K+")"),a.querySelectorAll("[id~="+u+"-]").length||q.push("~="),a.querySelectorAll(":checked").length||q.push(":checked"),a.querySelectorAll("a#"+u+"+*").length||q.push(".#.+[+~]")}),ia(function(a){var b=n.createElement("input");b.setAttribute("type","hidden"),a.appendChild(b).setAttribute("name","D"),a.querySelectorAll("[name=d]").length&&q.push("name"+L+"*[*^$|!~]?="),a.querySelectorAll(":enabled").length||q.push(":enabled",":disabled"),a.querySelectorAll("*,:x"),q.push(",.*:")})),(c.matchesSelector=Z.test(s=o.matches||o.webkitMatchesSelector||o.mozMatchesSelector||o.oMatchesSelector||o.msMatchesSelector))&&ia(function(a){c.disconnectedMatch=s.call(a,"div"),s.call(a,"[s!='']:x"),r.push("!=",O)}),q=q.length&&new RegExp(q.join("|")),r=r.length&&new RegExp(r.join("|")),b=Z.test(o.compareDocumentPosition),t=b||Z.test(o.contains)?function(a,b){var c=9===a.nodeType?a.documentElement:a,d=b&&b.parentNode;return a===d||!(!d||1!==d.nodeType||!(c.contains?c.contains(d):a.compareDocumentPosition&&16&a.compareDocumentPosition(d)))}:function(a,b){if(b)while(b=b.parentNode)if(b===a)return!0;return!1},B=b?function(a,b){if(a===b)return l=!0,0;var d=!a.compareDocumentPosition-!b.compareDocumentPosition;return d?d:(d=(a.ownerDocument||a)===(b.ownerDocument||b)?a.compareDocumentPosition(b):1,1&d||!c.sortDetached&&b.compareDocumentPosition(a)===d?a===n||a.ownerDocument===v&&t(v,a)?-1:b===n||b.ownerDocument===v&&t(v,b)?1:k?J(k,a)-J(k,b):0:4&d?-1:1)}:function(a,b){if(a===b)return l=!0,0;var c,d=0,e=a.parentNode,f=b.parentNode,g=[a],h=[b];if(!e||!f)return a===n?-1:b===n?1:e?-1:f?1:k?J(k,a)-J(k,b):0;if(e===f)return ka(a,b);c=a;while(c=c.parentNode)g.unshift(c);c=b;while(c=c.parentNode)h.unshift(c);while(g[d]===h[d])d++;return d?ka(g[d],h[d]):g[d]===v?-1:h[d]===v?1:0},n):n},fa.matches=function(a,b){return fa(a,null,null,b)},fa.matchesSelector=function(a,b){if((a.ownerDocument||a)!==n&&m(a),b=b.replace(T,"='$1']"),c.matchesSelector&&p&&!A[b+" "]&&(!r||!r.test(b))&&(!q||!q.test(b)))try{var d=s.call(a,b);if(d||c.disconnectedMatch||a.document&&11!==a.document.nodeType)return d}catch(e){}return fa(b,n,null,[a]).length>0},fa.contains=function(a,b){return(a.ownerDocument||a)!==n&&m(a),t(a,b)},fa.attr=function(a,b){(a.ownerDocument||a)!==n&&m(a);var e=d.attrHandle[b.toLowerCase()],f=e&&D.call(d.attrHandle,b.toLowerCase())?e(a,b,!p):void 0;return void 0!==f?f:c.attributes||!p?a.getAttribute(b):(f=a.getAttributeNode(b))&&f.specified?f.value:null},fa.error=function(a){throw new Error("Syntax error, unrecognized expression: "+a)},fa.uniqueSort=function(a){var b,d=[],e=0,f=0;if(l=!c.detectDuplicates,k=!c.sortStable&&a.slice(0),a.sort(B),l){while(b=a[f++])b===a[f]&&(e=d.push(f));while(e--)a.splice(d[e],1)}return k=null,a},e=fa.getText=function(a){var b,c="",d=0,f=a.nodeType;if(f){if(1===f||9===f||11===f){if("string"==typeof a.textContent)return a.textContent;for(a=a.firstChild;a;a=a.nextSibling)c+=e(a)}else if(3===f||4===f)return a.nodeValue}else while(b=a[d++])c+=e(b);return c},d=fa.selectors={cacheLength:50,createPseudo:ha,match:W,attrHandle:{},find:{},relative:{">":{dir:"parentNode",first:!0}," ":{dir:"parentNode"},"+":{dir:"previousSibling",first:!0},"~":{dir:"previousSibling"}},preFilter:{ATTR:function(a){return a[1]=a[1].replace(ba,ca),a[3]=(a[3]||a[4]||a[5]||"").replace(ba,ca),"~="===a[2]&&(a[3]=" "+a[3]+" "),a.slice(0,4)},CHILD:function(a){return a[1]=a[1].toLowerCase(),"nth"===a[1].slice(0,3)?(a[3]||fa.error(a[0]),a[4]=+(a[4]?a[5]+(a[6]||1):2*("even"===a[3]||"odd"===a[3])),a[5]=+(a[7]+a[8]||"odd"===a[3])):a[3]&&fa.error(a[0]),a},PSEUDO:function(a){var b,c=!a[6]&&a[2];return W.CHILD.test(a[0])?null:(a[3]?a[2]=a[4]||a[5]||"":c&&U.test(c)&&(b=g(c,!0))&&(b=c.indexOf(")",c.length-b)-c.length)&&(a[0]=a[0].slice(0,b),a[2]=c.slice(0,b)),a.slice(0,3))}},filter:{TAG:function(a){var b=a.replace(ba,ca).toLowerCase();return"*"===a?function(){return!0}:function(a){return a.nodeName&&a.nodeName.toLowerCase()===b}},CLASS:function(a){var b=y[a+" "];return b||(b=new RegExp("(^|"+L+")"+a+"("+L+"|$)"))&&y(a,function(a){return b.test("string"==typeof a.className&&a.className||"undefined"!=typeof a.getAttribute&&a.getAttribute("class")||"")})},ATTR:function(a,b,c){return function(d){var e=fa.attr(d,a);return null==e?"!="===b:b?(e+="","="===b?e===c:"!="===b?e!==c:"^="===b?c&&0===e.indexOf(c):"*="===b?c&&e.indexOf(c)>-1:"$="===b?c&&e.slice(-c.length)===c:"~="===b?(" "+e.replace(P," ")+" ").indexOf(c)>-1:"|="===b?e===c||e.slice(0,c.length+1)===c+"-":!1):!0}},CHILD:function(a,b,c,d,e){var f="nth"!==a.slice(0,3),g="last"!==a.slice(-4),h="of-type"===b;return 1===d&&0===e?function(a){return!!a.parentNode}:function(b,c,i){var j,k,l,m,n,o,p=f!==g?"nextSibling":"previousSibling",q=b.parentNode,r=h&&b.nodeName.toLowerCase(),s=!i&&!h,t=!1;if(q){if(f){while(p){m=b;while(m=m[p])if(h?m.nodeName.toLowerCase()===r:1===m.nodeType)return!1;o=p="only"===a&&!o&&"nextSibling"}return!0}if(o=[g?q.firstChild:q.lastChild],g&&s){m=q,l=m[u]||(m[u]={}),k=l[m.uniqueID]||(l[m.uniqueID]={}),j=k[a]||[],n=j[0]===w&&j[1],t=n&&j[2],m=n&&q.childNodes[n];while(m=++n&&m&&m[p]||(t=n=0)||o.pop())if(1===m.nodeType&&++t&&m===b){k[a]=[w,n,t];break}}else if(s&&(m=b,l=m[u]||(m[u]={}),k=l[m.uniqueID]||(l[m.uniqueID]={}),j=k[a]||[],n=j[0]===w&&j[1],t=n),t===!1)while(m=++n&&m&&m[p]||(t=n=0)||o.pop())if((h?m.nodeName.toLowerCase()===r:1===m.nodeType)&&++t&&(s&&(l=m[u]||(m[u]={}),k=l[m.uniqueID]||(l[m.uniqueID]={}),k[a]=[w,t]),m===b))break;return t-=e,t===d||t%d===0&&t/d>=0}}},PSEUDO:function(a,b){var c,e=d.pseudos[a]||d.setFilters[a.toLowerCase()]||fa.error("unsupported pseudo: "+a);return e[u]?e(b):e.length>1?(c=[a,a,"",b],d.setFilters.hasOwnProperty(a.toLowerCase())?ha(function(a,c){var d,f=e(a,b),g=f.length;while(g--)d=J(a,f[g]),a[d]=!(c[d]=f[g])}):function(a){return e(a,0,c)}):e}},pseudos:{not:ha(function(a){var b=[],c=[],d=h(a.replace(Q,"$1"));return d[u]?ha(function(a,b,c,e){var f,g=d(a,null,e,[]),h=a.length;while(h--)(f=g[h])&&(a[h]=!(b[h]=f))}):function(a,e,f){return b[0]=a,d(b,null,f,c),b[0]=null,!c.pop()}}),has:ha(function(a){return function(b){return fa(a,b).length>0}}),contains:ha(function(a){return a=a.replace(ba,ca),function(b){return(b.textContent||b.innerText||e(b)).indexOf(a)>-1}}),lang:ha(function(a){return V.test(a||"")||fa.error("unsupported lang: "+a),a=a.replace(ba,ca).toLowerCase(),function(b){var c;do if(c=p?b.lang:b.getAttribute("xml:lang")||b.getAttribute("lang"))return c=c.toLowerCase(),c===a||0===c.indexOf(a+"-");while((b=b.parentNode)&&1===b.nodeType);return!1}}),target:function(b){var c=a.location&&a.location.hash;return c&&c.slice(1)===b.id},root:function(a){return a===o},focus:function(a){return a===n.activeElement&&(!n.hasFocus||n.hasFocus())&&!!(a.type||a.href||~a.tabIndex)},enabled:function(a){return a.disabled===!1},disabled:function(a){return a.disabled===!0},checked:function(a){var b=a.nodeName.toLowerCase();return"input"===b&&!!a.checked||"option"===b&&!!a.selected},selected:function(a){return a.parentNode&&a.parentNode.selectedIndex,a.selected===!0},empty:function(a){for(a=a.firstChild;a;a=a.nextSibling)if(a.nodeType<6)return!1;return!0},parent:function(a){return!d.pseudos.empty(a)},header:function(a){return Y.test(a.nodeName)},input:function(a){return X.test(a.nodeName)},button:function(a){var b=a.nodeName.toLowerCase();return"input"===b&&"button"===a.type||"button"===b},text:function(a){var b;return"input"===a.nodeName.toLowerCase()&&"text"===a.type&&(null==(b=a.getAttribute("type"))||"text"===b.toLowerCase())},first:na(function(){return[0]}),last:na(function(a,b){return[b-1]}),eq:na(function(a,b,c){return[0>c?c+b:c]}),even:na(function(a,b){for(var c=0;b>c;c+=2)a.push(c);return a}),odd:na(function(a,b){for(var c=1;b>c;c+=2)a.push(c);return a}),lt:na(function(a,b,c){for(var d=0>c?c+b:c;--d>=0;)a.push(d);return a}),gt:na(function(a,b,c){for(var d=0>c?c+b:c;++d<b;)a.push(d);return a})}},d.pseudos.nth=d.pseudos.eq;for(b in{radio:!0,checkbox:!0,file:!0,password:!0,image:!0})d.pseudos[b]=la(b);for(b in{submit:!0,reset:!0})d.pseudos[b]=ma(b);function pa(){}pa.prototype=d.filters=d.pseudos,d.setFilters=new pa,g=fa.tokenize=function(a,b){var c,e,f,g,h,i,j,k=z[a+" "];if(k)return b?0:k.slice(0);h=a,i=[],j=d.preFilter;while(h){c&&!(e=R.exec(h))||(e&&(h=h.slice(e[0].length)||h),i.push(f=[])),c=!1,(e=S.exec(h))&&(c=e.shift(),f.push({value:c,type:e[0].replace(Q," ")}),h=h.slice(c.length));for(g in d.filter)!(e=W[g].exec(h))||j[g]&&!(e=j[g](e))||(c=e.shift(),f.push({value:c,type:g,matches:e}),h=h.slice(c.length));if(!c)break}return b?h.length:h?fa.error(a):z(a,i).slice(0)};function qa(a){for(var b=0,c=a.length,d="";c>b;b++)d+=a[b].value;return d}function ra(a,b,c){var d=b.dir,e=c&&"parentNode"===d,f=x++;return b.first?function(b,c,f){while(b=b[d])if(1===b.nodeType||e)return a(b,c,f)}:function(b,c,g){var h,i,j,k=[w,f];if(g){while(b=b[d])if((1===b.nodeType||e)&&a(b,c,g))return!0}else while(b=b[d])if(1===b.nodeType||e){if(j=b[u]||(b[u]={}),i=j[b.uniqueID]||(j[b.uniqueID]={}),(h=i[d])&&h[0]===w&&h[1]===f)return k[2]=h[2];if(i[d]=k,k[2]=a(b,c,g))return!0}}}function sa(a){return a.length>1?function(b,c,d){var e=a.length;while(e--)if(!a[e](b,c,d))return!1;return!0}:a[0]}function ta(a,b,c){for(var d=0,e=b.length;e>d;d++)fa(a,b[d],c);return c}function ua(a,b,c,d,e){for(var f,g=[],h=0,i=a.length,j=null!=b;i>h;h++)(f=a[h])&&(c&&!c(f,d,e)||(g.push(f),j&&b.push(h)));return g}function va(a,b,c,d,e,f){return d&&!d[u]&&(d=va(d)),e&&!e[u]&&(e=va(e,f)),ha(function(f,g,h,i){var j,k,l,m=[],n=[],o=g.length,p=f||ta(b||"*",h.nodeType?[h]:h,[]),q=!a||!f&&b?p:ua(p,m,a,h,i),r=c?e||(f?a:o||d)?[]:g:q;if(c&&c(q,r,h,i),d){j=ua(r,n),d(j,[],h,i),k=j.length;while(k--)(l=j[k])&&(r[n[k]]=!(q[n[k]]=l))}if(f){if(e||a){if(e){j=[],k=r.length;while(k--)(l=r[k])&&j.push(q[k]=l);e(null,r=[],j,i)}k=r.length;while(k--)(l=r[k])&&(j=e?J(f,l):m[k])>-1&&(f[j]=!(g[j]=l))}}else r=ua(r===g?r.splice(o,r.length):r),e?e(null,g,r,i):H.apply(g,r)})}function wa(a){for(var b,c,e,f=a.length,g=d.relative[a[0].type],h=g||d.relative[" "],i=g?1:0,k=ra(function(a){return a===b},h,!0),l=ra(function(a){return J(b,a)>-1},h,!0),m=[function(a,c,d){var e=!g&&(d||c!==j)||((b=c).nodeType?k(a,c,d):l(a,c,d));return b=null,e}];f>i;i++)if(c=d.relative[a[i].type])m=[ra(sa(m),c)];else{if(c=d.filter[a[i].type].apply(null,a[i].matches),c[u]){for(e=++i;f>e;e++)if(d.relative[a[e].type])break;return va(i>1&&sa(m),i>1&&qa(a.slice(0,i-1).concat({value:" "===a[i-2].type?"*":""})).replace(Q,"$1"),c,e>i&&wa(a.slice(i,e)),f>e&&wa(a=a.slice(e)),f>e&&qa(a))}m.push(c)}return sa(m)}function xa(a,b){var c=b.length>0,e=a.length>0,f=function(f,g,h,i,k){var l,o,q,r=0,s="0",t=f&&[],u=[],v=j,x=f||e&&d.find.TAG("*",k),y=w+=null==v?1:Math.random()||.1,z=x.length;for(k&&(j=g===n||g||k);s!==z&&null!=(l=x[s]);s++){if(e&&l){o=0,g||l.ownerDocument===n||(m(l),h=!p);while(q=a[o++])if(q(l,g||n,h)){i.push(l);break}k&&(w=y)}c&&((l=!q&&l)&&r--,f&&t.push(l))}if(r+=s,c&&s!==r){o=0;while(q=b[o++])q(t,u,g,h);if(f){if(r>0)while(s--)t[s]||u[s]||(u[s]=F.call(i));u=ua(u)}H.apply(i,u),k&&!f&&u.length>0&&r+b.length>1&&fa.uniqueSort(i)}return k&&(w=y,j=v),t};return c?ha(f):f}return h=fa.compile=function(a,b){var c,d=[],e=[],f=A[a+" "];if(!f){b||(b=g(a)),c=b.length;while(c--)f=wa(b[c]),f[u]?d.push(f):e.push(f);f=A(a,xa(e,d)),f.selector=a}return f},i=fa.select=function(a,b,e,f){var i,j,k,l,m,n="function"==typeof a&&a,o=!f&&g(a=n.selector||a);if(e=e||[],1===o.length){if(j=o[0]=o[0].slice(0),j.length>2&&"ID"===(k=j[0]).type&&c.getById&&9===b.nodeType&&p&&d.relative[j[1].type]){if(b=(d.find.ID(k.matches[0].replace(ba,ca),b)||[])[0],!b)return e;n&&(b=b.parentNode),a=a.slice(j.shift().value.length)}i=W.needsContext.test(a)?0:j.length;while(i--){if(k=j[i],d.relative[l=k.type])break;if((m=d.find[l])&&(f=m(k.matches[0].replace(ba,ca),_.test(j[0].type)&&oa(b.parentNode)||b))){if(j.splice(i,1),a=f.length&&qa(j),!a)return H.apply(e,f),e;break}}}return(n||h(a,o))(f,b,!p,e,!b||_.test(a)&&oa(b.parentNode)||b),e},c.sortStable=u.split("").sort(B).join("")===u,c.detectDuplicates=!!l,m(),c.sortDetached=ia(function(a){return 1&a.compareDocumentPosition(n.createElement("div"))}),ia(function(a){return a.innerHTML="<a href='#'></a>","#"===a.firstChild.getAttribute("href")})||ja("type|href|height|width",function(a,b,c){return c?void 0:a.getAttribute(b,"type"===b.toLowerCase()?1:2)}),c.attributes&&ia(function(a){return a.innerHTML="<input/>",a.firstChild.setAttribute("value",""),""===a.firstChild.getAttribute("value")})||ja("value",function(a,b,c){return c||"input"!==a.nodeName.toLowerCase()?void 0:a.defaultValue}),ia(function(a){return null==a.getAttribute("disabled")})||ja(K,function(a,b,c){var d;return c?void 0:a[b]===!0?b.toLowerCase():(d=a.getAttributeNode(b))&&d.specified?d.value:null}),fa}(a);n.find=t,n.expr=t.selectors,n.expr[":"]=n.expr.pseudos,n.uniqueSort=n.unique=t.uniqueSort,n.text=t.getText,n.isXMLDoc=t.isXML,n.contains=t.contains;var u=function(a,b,c){var d=[],e=void 0!==c;while((a=a[b])&&9!==a.nodeType)if(1===a.nodeType){if(e&&n(a).is(c))break;d.push(a)}return d},v=function(a,b){for(var c=[];a;a=a.nextSibling)1===a.nodeType&&a!==b&&c.push(a);return c},w=n.expr.match.needsContext,x=/^<([\w-]+)\s*\/?>(?:<\/\1>|)$/,y=/^.[^:#\[\.,]*$/;function z(a,b,c){if(n.isFunction(b))return n.grep(a,function(a,d){return!!b.call(a,d,a)!==c});if(b.nodeType)return n.grep(a,function(a){return a===b!==c});if("string"==typeof b){if(y.test(b))return n.filter(b,a,c);b=n.filter(b,a)}return n.grep(a,function(a){return n.inArray(a,b)>-1!==c})}n.filter=function(a,b,c){var d=b[0];return c&&(a=":not("+a+")"),1===b.length&&1===d.nodeType?n.find.matchesSelector(d,a)?[d]:[]:n.find.matches(a,n.grep(b,function(a){return 1===a.nodeType}))},n.fn.extend({find:function(a){var b,c=[],d=this,e=d.length;if("string"!=typeof a)return this.pushStack(n(a).filter(function(){for(b=0;e>b;b++)if(n.contains(d[b],this))return!0}));for(b=0;e>b;b++)n.find(a,d[b],c);return c=this.pushStack(e>1?n.unique(c):c),c.selector=this.selector?this.selector+" "+a:a,c},filter:function(a){return this.pushStack(z(this,a||[],!1))},not:function(a){return this.pushStack(z(this,a||[],!0))},is:function(a){return!!z(this,"string"==typeof a&&w.test(a)?n(a):a||[],!1).length}});var A,B=/^(?:\s*(<[\w\W]+>)[^>]*|#([\w-]*))$/,C=n.fn.init=function(a,b,c){var e,f;if(!a)return this;if(c=c||A,"string"==typeof a){if(e="<"===a.charAt(0)&&">"===a.charAt(a.length-1)&&a.length>=3?[null,a,null]:B.exec(a),!e||!e[1]&&b)return!b||b.jquery?(b||c).find(a):this.constructor(b).find(a);if(e[1]){if(b=b instanceof n?b[0]:b,n.merge(this,n.parseHTML(e[1],b&&b.nodeType?b.ownerDocument||b:d,!0)),x.test(e[1])&&n.isPlainObject(b))for(e in b)n.isFunction(this[e])?this[e](b[e]):this.attr(e,b[e]);return this}if(f=d.getElementById(e[2]),f&&f.parentNode){if(f.id!==e[2])return A.find(a);this.length=1,this[0]=f}return this.context=d,this.selector=a,this}return a.nodeType?(this.context=this[0]=a,this.length=1,this):n.isFunction(a)?"undefined"!=typeof c.ready?c.ready(a):a(n):(void 0!==a.selector&&(this.selector=a.selector,this.context=a.context),n.makeArray(a,this))};C.prototype=n.fn,A=n(d);var D=/^(?:parents|prev(?:Until|All))/,E={children:!0,contents:!0,next:!0,prev:!0};n.fn.extend({has:function(a){var b,c=n(a,this),d=c.length;return this.filter(function(){for(b=0;d>b;b++)if(n.contains(this,c[b]))return!0})},closest:function(a,b){for(var c,d=0,e=this.length,f=[],g=w.test(a)||"string"!=typeof a?n(a,b||this.context):0;e>d;d++)for(c=this[d];c&&c!==b;c=c.parentNode)if(c.nodeType<11&&(g?g.index(c)>-1:1===c.nodeType&&n.find.matchesSelector(c,a))){f.push(c);break}return this.pushStack(f.length>1?n.uniqueSort(f):f)},index:function(a){return a?"string"==typeof a?n.inArray(this[0],n(a)):n.inArray(a.jquery?a[0]:a,this):this[0]&&this[0].parentNode?this.first().prevAll().length:-1},add:function(a,b){return this.pushStack(n.uniqueSort(n.merge(this.get(),n(a,b))))},addBack:function(a){return this.add(null==a?this.prevObject:this.prevObject.filter(a))}});function F(a,b){do a=a[b];while(a&&1!==a.nodeType);return a}n.each({parent:function(a){var b=a.parentNode;return b&&11!==b.nodeType?b:null},parents:function(a){return u(a,"parentNode")},parentsUntil:function(a,b,c){return u(a,"parentNode",c)},next:function(a){return F(a,"nextSibling")},prev:function(a){return F(a,"previousSibling")},nextAll:function(a){return u(a,"nextSibling")},prevAll:function(a){return u(a,"previousSibling")},nextUntil:function(a,b,c){return u(a,"nextSibling",c)},prevUntil:function(a,b,c){return u(a,"previousSibling",c)},siblings:function(a){return v((a.parentNode||{}).firstChild,a)},children:function(a){return v(a.firstChild)},contents:function(a){return n.nodeName(a,"iframe")?a.contentDocument||a.contentWindow.document:n.merge([],a.childNodes)}},function(a,b){n.fn[a]=function(c,d){var e=n.map(this,b,c);return"Until"!==a.slice(-5)&&(d=c),d&&"string"==typeof d&&(e=n.filter(d,e)),this.length>1&&(E[a]||(e=n.uniqueSort(e)),D.test(a)&&(e=e.reverse())),this.pushStack(e)}});var G=/\S+/g;function H(a){var b={};return n.each(a.match(G)||[],function(a,c){b[c]=!0}),b}n.Callbacks=function(a){a="string"==typeof a?H(a):n.extend({},a);var b,c,d,e,f=[],g=[],h=-1,i=function(){for(e=a.once,d=b=!0;g.length;h=-1){c=g.shift();while(++h<f.length)f[h].apply(c[0],c[1])===!1&&a.stopOnFalse&&(h=f.length,c=!1)}a.memory||(c=!1),b=!1,e&&(f=c?[]:"")},j={add:function(){return f&&(c&&!b&&(h=f.length-1,g.push(c)),function d(b){n.each(b,function(b,c){n.isFunction(c)?a.unique&&j.has(c)||f.push(c):c&&c.length&&"string"!==n.type(c)&&d(c)})}(arguments),c&&!b&&i()),this},remove:function(){return n.each(arguments,function(a,b){var c;while((c=n.inArray(b,f,c))>-1)f.splice(c,1),h>=c&&h--}),this},has:function(a){return a?n.inArray(a,f)>-1:f.length>0},empty:function(){return f&&(f=[]),this},disable:function(){return e=g=[],f=c="",this},disabled:function(){return!f},lock:function(){return e=!0,c||j.disable(),this},locked:function(){return!!e},fireWith:function(a,c){return e||(c=c||[],c=[a,c.slice?c.slice():c],g.push(c),b||i()),this},fire:function(){return j.fireWith(this,arguments),this},fired:function(){return!!d}};return j},n.extend({Deferred:function(a){var b=[["resolve","done",n.Callbacks("once memory"),"resolved"],["reject","fail",n.Callbacks("once memory"),"rejected"],["notify","progress",n.Callbacks("memory")]],c="pending",d={state:function(){return c},always:function(){return e.done(arguments).fail(arguments),this},then:function(){var a=arguments;return n.Deferred(function(c){n.each(b,function(b,f){var g=n.isFunction(a[b])&&a[b];e[f[1]](function(){var a=g&&g.apply(this,arguments);a&&n.isFunction(a.promise)?a.promise().progress(c.notify).done(c.resolve).fail(c.reject):c[f[0]+"With"](this===d?c.promise():this,g?[a]:arguments)})}),a=null}).promise()},promise:function(a){return null!=a?n.extend(a,d):d}},e={};return d.pipe=d.then,n.each(b,function(a,f){var g=f[2],h=f[3];d[f[1]]=g.add,h&&g.add(function(){c=h},b[1^a][2].disable,b[2][2].lock),e[f[0]]=function(){return e[f[0]+"With"](this===e?d:this,arguments),this},e[f[0]+"With"]=g.fireWith}),d.promise(e),a&&a.call(e,e),e},when:function(a){var b=0,c=e.call(arguments),d=c.length,f=1!==d||a&&n.isFunction(a.promise)?d:0,g=1===f?a:n.Deferred(),h=function(a,b,c){return function(d){b[a]=this,c[a]=arguments.length>1?e.call(arguments):d,c===i?g.notifyWith(b,c):--f||g.resolveWith(b,c)}},i,j,k;if(d>1)for(i=new Array(d),j=new Array(d),k=new Array(d);d>b;b++)c[b]&&n.isFunction(c[b].promise)?c[b].promise().progress(h(b,j,i)).done(h(b,k,c)).fail(g.reject):--f;return f||g.resolveWith(k,c),g.promise()}});var I;n.fn.ready=function(a){return n.ready.promise().done(a),this},n.extend({isReady:!1,readyWait:1,holdReady:function(a){a?n.readyWait++:n.ready(!0)},ready:function(a){(a===!0?--n.readyWait:n.isReady)||(n.isReady=!0,a!==!0&&--n.readyWait>0||(I.resolveWith(d,[n]),n.fn.triggerHandler&&(n(d).triggerHandler("ready"),n(d).off("ready"))))}});function J(){d.addEventListener?(d.removeEventListener("DOMContentLoaded",K),a.removeEventListener("load",K)):(d.detachEvent("onreadystatechange",K),a.detachEvent("onload",K))}function K(){(d.addEventListener||"load"===a.event.type||"complete"===d.readyState)&&(J(),n.ready())}n.ready.promise=function(b){if(!I)if(I=n.Deferred(),"complete"===d.readyState||"loading"!==d.readyState&&!d.documentElement.doScroll)a.setTimeout(n.ready);else if(d.addEventListener)d.addEventListener("DOMContentLoaded",K),a.addEventListener("load",K);else{d.attachEvent("onreadystatechange",K),a.attachEvent("onload",K);var c=!1;try{c=null==a.frameElement&&d.documentElement}catch(e){}c&&c.doScroll&&!function f(){if(!n.isReady){try{c.doScroll("left")}catch(b){return a.setTimeout(f,50)}J(),n.ready()}}()}return I.promise(b)},n.ready.promise();var L;for(L in n(l))break;l.ownFirst="0"===L,l.inlineBlockNeedsLayout=!1,n(function(){var a,b,c,e;c=d.getElementsByTagName("body")[0],c&&c.style&&(b=d.createElement("div"),e=d.createElement("div"),e.style.cssText="position:absolute;border:0;width:0;height:0;top:0;left:-9999px",c.appendChild(e).appendChild(b),"undefined"!=typeof b.style.zoom&&(b.style.cssText="display:inline;margin:0;border:0;padding:1px;width:1px;zoom:1",l.inlineBlockNeedsLayout=a=3===b.offsetWidth,a&&(c.style.zoom=1)),c.removeChild(e))}),function(){var a=d.createElement("div");l.deleteExpando=!0;try{delete a.test}catch(b){l.deleteExpando=!1}a=null}();var M=function(a){var b=n.noData[(a.nodeName+" ").toLowerCase()],c=+a.nodeType||1;return 1!==c&&9!==c?!1:!b||b!==!0&&a.getAttribute("classid")===b},N=/^(?:\{[\w\W]*\}|\[[\w\W]*\])$/,O=/([A-Z])/g;function P(a,b,c){if(void 0===c&&1===a.nodeType){var d="data-"+b.replace(O,"-$1").toLowerCase();if(c=a.getAttribute(d),"string"==typeof c){try{c="true"===c?!0:"false"===c?!1:"null"===c?null:+c+""===c?+c:N.test(c)?n.parseJSON(c):c}catch(e){}n.data(a,b,c)}else c=void 0;
}return c}function Q(a){var b;for(b in a)if(("data"!==b||!n.isEmptyObject(a[b]))&&"toJSON"!==b)return!1;return!0}function R(a,b,d,e){if(M(a)){var f,g,h=n.expando,i=a.nodeType,j=i?n.cache:a,k=i?a[h]:a[h]&&h;if(k&&j[k]&&(e||j[k].data)||void 0!==d||"string"!=typeof b)return k||(k=i?a[h]=c.pop()||n.guid++:h),j[k]||(j[k]=i?{}:{toJSON:n.noop}),"object"!=typeof b&&"function"!=typeof b||(e?j[k]=n.extend(j[k],b):j[k].data=n.extend(j[k].data,b)),g=j[k],e||(g.data||(g.data={}),g=g.data),void 0!==d&&(g[n.camelCase(b)]=d),"string"==typeof b?(f=g[b],null==f&&(f=g[n.camelCase(b)])):f=g,f}}function S(a,b,c){if(M(a)){var d,e,f=a.nodeType,g=f?n.cache:a,h=f?a[n.expando]:n.expando;if(g[h]){if(b&&(d=c?g[h]:g[h].data)){n.isArray(b)?b=b.concat(n.map(b,n.camelCase)):b in d?b=[b]:(b=n.camelCase(b),b=b in d?[b]:b.split(" ")),e=b.length;while(e--)delete d[b[e]];if(c?!Q(d):!n.isEmptyObject(d))return}(c||(delete g[h].data,Q(g[h])))&&(f?n.cleanData([a],!0):l.deleteExpando||g!=g.window?delete g[h]:g[h]=void 0)}}}n.extend({cache:{},noData:{"applet ":!0,"embed ":!0,"object ":"clsid:D27CDB6E-AE6D-11cf-96B8-444553540000"},hasData:function(a){return a=a.nodeType?n.cache[a[n.expando]]:a[n.expando],!!a&&!Q(a)},data:function(a,b,c){return R(a,b,c)},removeData:function(a,b){return S(a,b)},_data:function(a,b,c){return R(a,b,c,!0)},_removeData:function(a,b){return S(a,b,!0)}}),n.fn.extend({data:function(a,b){var c,d,e,f=this[0],g=f&&f.attributes;if(void 0===a){if(this.length&&(e=n.data(f),1===f.nodeType&&!n._data(f,"parsedAttrs"))){c=g.length;while(c--)g[c]&&(d=g[c].name,0===d.indexOf("data-")&&(d=n.camelCase(d.slice(5)),P(f,d,e[d])));n._data(f,"parsedAttrs",!0)}return e}return"object"==typeof a?this.each(function(){n.data(this,a)}):arguments.length>1?this.each(function(){n.data(this,a,b)}):f?P(f,a,n.data(f,a)):void 0},removeData:function(a){return this.each(function(){n.removeData(this,a)})}}),n.extend({queue:function(a,b,c){var d;return a?(b=(b||"fx")+"queue",d=n._data(a,b),c&&(!d||n.isArray(c)?d=n._data(a,b,n.makeArray(c)):d.push(c)),d||[]):void 0},dequeue:function(a,b){b=b||"fx";var c=n.queue(a,b),d=c.length,e=c.shift(),f=n._queueHooks(a,b),g=function(){n.dequeue(a,b)};"inprogress"===e&&(e=c.shift(),d--),e&&("fx"===b&&c.unshift("inprogress"),delete f.stop,e.call(a,g,f)),!d&&f&&f.empty.fire()},_queueHooks:function(a,b){var c=b+"queueHooks";return n._data(a,c)||n._data(a,c,{empty:n.Callbacks("once memory").add(function(){n._removeData(a,b+"queue"),n._removeData(a,c)})})}}),n.fn.extend({queue:function(a,b){var c=2;return"string"!=typeof a&&(b=a,a="fx",c--),arguments.length<c?n.queue(this[0],a):void 0===b?this:this.each(function(){var c=n.queue(this,a,b);n._queueHooks(this,a),"fx"===a&&"inprogress"!==c[0]&&n.dequeue(this,a)})},dequeue:function(a){return this.each(function(){n.dequeue(this,a)})},clearQueue:function(a){return this.queue(a||"fx",[])},promise:function(a,b){var c,d=1,e=n.Deferred(),f=this,g=this.length,h=function(){--d||e.resolveWith(f,[f])};"string"!=typeof a&&(b=a,a=void 0),a=a||"fx";while(g--)c=n._data(f[g],a+"queueHooks"),c&&c.empty&&(d++,c.empty.add(h));return h(),e.promise(b)}}),function(){var a;l.shrinkWrapBlocks=function(){if(null!=a)return a;a=!1;var b,c,e;return c=d.getElementsByTagName("body")[0],c&&c.style?(b=d.createElement("div"),e=d.createElement("div"),e.style.cssText="position:absolute;border:0;width:0;height:0;top:0;left:-9999px",c.appendChild(e).appendChild(b),"undefined"!=typeof b.style.zoom&&(b.style.cssText="-webkit-box-sizing:content-box;-moz-box-sizing:content-box;box-sizing:content-box;display:block;margin:0;border:0;padding:1px;width:1px;zoom:1",b.appendChild(d.createElement("div")).style.width="5px",a=3!==b.offsetWidth),c.removeChild(e),a):void 0}}();var T=/[+-]?(?:\d*\.|)\d+(?:[eE][+-]?\d+|)/.source,U=new RegExp("^(?:([+-])=|)("+T+")([a-z%]*)$","i"),V=["Top","Right","Bottom","Left"],W=function(a,b){return a=b||a,"none"===n.css(a,"display")||!n.contains(a.ownerDocument,a)};function X(a,b,c,d){var e,f=1,g=20,h=d?function(){return d.cur()}:function(){return n.css(a,b,"")},i=h(),j=c&&c[3]||(n.cssNumber[b]?"":"px"),k=(n.cssNumber[b]||"px"!==j&&+i)&&U.exec(n.css(a,b));if(k&&k[3]!==j){j=j||k[3],c=c||[],k=+i||1;do f=f||".5",k/=f,n.style(a,b,k+j);while(f!==(f=h()/i)&&1!==f&&--g)}return c&&(k=+k||+i||0,e=c[1]?k+(c[1]+1)*c[2]:+c[2],d&&(d.unit=j,d.start=k,d.end=e)),e}var Y=function(a,b,c,d,e,f,g){var h=0,i=a.length,j=null==c;if("object"===n.type(c)){e=!0;for(h in c)Y(a,b,h,c[h],!0,f,g)}else if(void 0!==d&&(e=!0,n.isFunction(d)||(g=!0),j&&(g?(b.call(a,d),b=null):(j=b,b=function(a,b,c){return j.call(n(a),c)})),b))for(;i>h;h++)b(a[h],c,g?d:d.call(a[h],h,b(a[h],c)));return e?a:j?b.call(a):i?b(a[0],c):f},Z=/^(?:checkbox|radio)$/i,$=/<([\w:-]+)/,_=/^$|\/(?:java|ecma)script/i,aa=/^\s+/,ba="abbr|article|aside|audio|bdi|canvas|data|datalist|details|dialog|figcaption|figure|footer|header|hgroup|main|mark|meter|nav|output|picture|progress|section|summary|template|time|video";function ca(a){var b=ba.split("|"),c=a.createDocumentFragment();if(c.createElement)while(b.length)c.createElement(b.pop());return c}!function(){var a=d.createElement("div"),b=d.createDocumentFragment(),c=d.createElement("input");a.innerHTML="  <link/><table></table><a href='/a'>a</a><input type='checkbox'/>",l.leadingWhitespace=3===a.firstChild.nodeType,l.tbody=!a.getElementsByTagName("tbody").length,l.htmlSerialize=!!a.getElementsByTagName("link").length,l.html5Clone="<:nav></:nav>"!==d.createElement("nav").cloneNode(!0).outerHTML,c.type="checkbox",c.checked=!0,b.appendChild(c),l.appendChecked=c.checked,a.innerHTML="<textarea>x</textarea>",l.noCloneChecked=!!a.cloneNode(!0).lastChild.defaultValue,b.appendChild(a),c=d.createElement("input"),c.setAttribute("type","radio"),c.setAttribute("checked","checked"),c.setAttribute("name","t"),a.appendChild(c),l.checkClone=a.cloneNode(!0).cloneNode(!0).lastChild.checked,l.noCloneEvent=!!a.addEventListener,a[n.expando]=1,l.attributes=!a.getAttribute(n.expando)}();var da={option:[1,"<select multiple='multiple'>","</select>"],legend:[1,"<fieldset>","</fieldset>"],area:[1,"<map>","</map>"],param:[1,"<object>","</object>"],thead:[1,"<table>","</table>"],tr:[2,"<table><tbody>","</tbody></table>"],col:[2,"<table><tbody></tbody><colgroup>","</colgroup></table>"],td:[3,"<table><tbody><tr>","</tr></tbody></table>"],_default:l.htmlSerialize?[0,"",""]:[1,"X<div>","</div>"]};da.optgroup=da.option,da.tbody=da.tfoot=da.colgroup=da.caption=da.thead,da.th=da.td;function ea(a,b){var c,d,e=0,f="undefined"!=typeof a.getElementsByTagName?a.getElementsByTagName(b||"*"):"undefined"!=typeof a.querySelectorAll?a.querySelectorAll(b||"*"):void 0;if(!f)for(f=[],c=a.childNodes||a;null!=(d=c[e]);e++)!b||n.nodeName(d,b)?f.push(d):n.merge(f,ea(d,b));return void 0===b||b&&n.nodeName(a,b)?n.merge([a],f):f}function fa(a,b){for(var c,d=0;null!=(c=a[d]);d++)n._data(c,"globalEval",!b||n._data(b[d],"globalEval"))}var ga=/<|&#?\w+;/,ha=/<tbody/i;function ia(a){Z.test(a.type)&&(a.defaultChecked=a.checked)}function ja(a,b,c,d,e){for(var f,g,h,i,j,k,m,o=a.length,p=ca(b),q=[],r=0;o>r;r++)if(g=a[r],g||0===g)if("object"===n.type(g))n.merge(q,g.nodeType?[g]:g);else if(ga.test(g)){i=i||p.appendChild(b.createElement("div")),j=($.exec(g)||["",""])[1].toLowerCase(),m=da[j]||da._default,i.innerHTML=m[1]+n.htmlPrefilter(g)+m[2],f=m[0];while(f--)i=i.lastChild;if(!l.leadingWhitespace&&aa.test(g)&&q.push(b.createTextNode(aa.exec(g)[0])),!l.tbody){g="table"!==j||ha.test(g)?"<table>"!==m[1]||ha.test(g)?0:i:i.firstChild,f=g&&g.childNodes.length;while(f--)n.nodeName(k=g.childNodes[f],"tbody")&&!k.childNodes.length&&g.removeChild(k)}n.merge(q,i.childNodes),i.textContent="";while(i.firstChild)i.removeChild(i.firstChild);i=p.lastChild}else q.push(b.createTextNode(g));i&&p.removeChild(i),l.appendChecked||n.grep(ea(q,"input"),ia),r=0;while(g=q[r++])if(d&&n.inArray(g,d)>-1)e&&e.push(g);else if(h=n.contains(g.ownerDocument,g),i=ea(p.appendChild(g),"script"),h&&fa(i),c){f=0;while(g=i[f++])_.test(g.type||"")&&c.push(g)}return i=null,p}!function(){var b,c,e=d.createElement("div");for(b in{submit:!0,change:!0,focusin:!0})c="on"+b,(l[b]=c in a)||(e.setAttribute(c,"t"),l[b]=e.attributes[c].expando===!1);e=null}();var ka=/^(?:input|select|textarea)$/i,la=/^key/,ma=/^(?:mouse|pointer|contextmenu|drag|drop)|click/,na=/^(?:focusinfocus|focusoutblur)$/,oa=/^([^.]*)(?:\.(.+)|)/;function pa(){return!0}function qa(){return!1}function ra(){try{return d.activeElement}catch(a){}}function sa(a,b,c,d,e,f){var g,h;if("object"==typeof b){"string"!=typeof c&&(d=d||c,c=void 0);for(h in b)sa(a,h,c,d,b[h],f);return a}if(null==d&&null==e?(e=c,d=c=void 0):null==e&&("string"==typeof c?(e=d,d=void 0):(e=d,d=c,c=void 0)),e===!1)e=qa;else if(!e)return a;return 1===f&&(g=e,e=function(a){return n().off(a),g.apply(this,arguments)},e.guid=g.guid||(g.guid=n.guid++)),a.each(function(){n.event.add(this,b,e,d,c)})}n.event={global:{},add:function(a,b,c,d,e){var f,g,h,i,j,k,l,m,o,p,q,r=n._data(a);if(r){c.handler&&(i=c,c=i.handler,e=i.selector),c.guid||(c.guid=n.guid++),(g=r.events)||(g=r.events={}),(k=r.handle)||(k=r.handle=function(a){return"undefined"==typeof n||a&&n.event.triggered===a.type?void 0:n.event.dispatch.apply(k.elem,arguments)},k.elem=a),b=(b||"").match(G)||[""],h=b.length;while(h--)f=oa.exec(b[h])||[],o=q=f[1],p=(f[2]||"").split(".").sort(),o&&(j=n.event.special[o]||{},o=(e?j.delegateType:j.bindType)||o,j=n.event.special[o]||{},l=n.extend({type:o,origType:q,data:d,handler:c,guid:c.guid,selector:e,needsContext:e&&n.expr.match.needsContext.test(e),namespace:p.join(".")},i),(m=g[o])||(m=g[o]=[],m.delegateCount=0,j.setup&&j.setup.call(a,d,p,k)!==!1||(a.addEventListener?a.addEventListener(o,k,!1):a.attachEvent&&a.attachEvent("on"+o,k))),j.add&&(j.add.call(a,l),l.handler.guid||(l.handler.guid=c.guid)),e?m.splice(m.delegateCount++,0,l):m.push(l),n.event.global[o]=!0);a=null}},remove:function(a,b,c,d,e){var f,g,h,i,j,k,l,m,o,p,q,r=n.hasData(a)&&n._data(a);if(r&&(k=r.events)){b=(b||"").match(G)||[""],j=b.length;while(j--)if(h=oa.exec(b[j])||[],o=q=h[1],p=(h[2]||"").split(".").sort(),o){l=n.event.special[o]||{},o=(d?l.delegateType:l.bindType)||o,m=k[o]||[],h=h[2]&&new RegExp("(^|\\.)"+p.join("\\.(?:.*\\.|)")+"(\\.|$)"),i=f=m.length;while(f--)g=m[f],!e&&q!==g.origType||c&&c.guid!==g.guid||h&&!h.test(g.namespace)||d&&d!==g.selector&&("**"!==d||!g.selector)||(m.splice(f,1),g.selector&&m.delegateCount--,l.remove&&l.remove.call(a,g));i&&!m.length&&(l.teardown&&l.teardown.call(a,p,r.handle)!==!1||n.removeEvent(a,o,r.handle),delete k[o])}else for(o in k)n.event.remove(a,o+b[j],c,d,!0);n.isEmptyObject(k)&&(delete r.handle,n._removeData(a,"events"))}},trigger:function(b,c,e,f){var g,h,i,j,l,m,o,p=[e||d],q=k.call(b,"type")?b.type:b,r=k.call(b,"namespace")?b.namespace.split("."):[];if(i=m=e=e||d,3!==e.nodeType&&8!==e.nodeType&&!na.test(q+n.event.triggered)&&(q.indexOf(".")>-1&&(r=q.split("."),q=r.shift(),r.sort()),h=q.indexOf(":")<0&&"on"+q,b=b[n.expando]?b:new n.Event(q,"object"==typeof b&&b),b.isTrigger=f?2:3,b.namespace=r.join("."),b.rnamespace=b.namespace?new RegExp("(^|\\.)"+r.join("\\.(?:.*\\.|)")+"(\\.|$)"):null,b.result=void 0,b.target||(b.target=e),c=null==c?[b]:n.makeArray(c,[b]),l=n.event.special[q]||{},f||!l.trigger||l.trigger.apply(e,c)!==!1)){if(!f&&!l.noBubble&&!n.isWindow(e)){for(j=l.delegateType||q,na.test(j+q)||(i=i.parentNode);i;i=i.parentNode)p.push(i),m=i;m===(e.ownerDocument||d)&&p.push(m.defaultView||m.parentWindow||a)}o=0;while((i=p[o++])&&!b.isPropagationStopped())b.type=o>1?j:l.bindType||q,g=(n._data(i,"events")||{})[b.type]&&n._data(i,"handle"),g&&g.apply(i,c),g=h&&i[h],g&&g.apply&&M(i)&&(b.result=g.apply(i,c),b.result===!1&&b.preventDefault());if(b.type=q,!f&&!b.isDefaultPrevented()&&(!l._default||l._default.apply(p.pop(),c)===!1)&&M(e)&&h&&e[q]&&!n.isWindow(e)){m=e[h],m&&(e[h]=null),n.event.triggered=q;try{e[q]()}catch(s){}n.event.triggered=void 0,m&&(e[h]=m)}return b.result}},dispatch:function(a){a=n.event.fix(a);var b,c,d,f,g,h=[],i=e.call(arguments),j=(n._data(this,"events")||{})[a.type]||[],k=n.event.special[a.type]||{};if(i[0]=a,a.delegateTarget=this,!k.preDispatch||k.preDispatch.call(this,a)!==!1){h=n.event.handlers.call(this,a,j),b=0;while((f=h[b++])&&!a.isPropagationStopped()){a.currentTarget=f.elem,c=0;while((g=f.handlers[c++])&&!a.isImmediatePropagationStopped())a.rnamespace&&!a.rnamespace.test(g.namespace)||(a.handleObj=g,a.data=g.data,d=((n.event.special[g.origType]||{}).handle||g.handler).apply(f.elem,i),void 0!==d&&(a.result=d)===!1&&(a.preventDefault(),a.stopPropagation()))}return k.postDispatch&&k.postDispatch.call(this,a),a.result}},handlers:function(a,b){var c,d,e,f,g=[],h=b.delegateCount,i=a.target;if(h&&i.nodeType&&("click"!==a.type||isNaN(a.button)||a.button<1))for(;i!=this;i=i.parentNode||this)if(1===i.nodeType&&(i.disabled!==!0||"click"!==a.type)){for(d=[],c=0;h>c;c++)f=b[c],e=f.selector+" ",void 0===d[e]&&(d[e]=f.needsContext?n(e,this).index(i)>-1:n.find(e,this,null,[i]).length),d[e]&&d.push(f);d.length&&g.push({elem:i,handlers:d})}return h<b.length&&g.push({elem:this,handlers:b.slice(h)}),g},fix:function(a){if(a[n.expando])return a;var b,c,e,f=a.type,g=a,h=this.fixHooks[f];h||(this.fixHooks[f]=h=ma.test(f)?this.mouseHooks:la.test(f)?this.keyHooks:{}),e=h.props?this.props.concat(h.props):this.props,a=new n.Event(g),b=e.length;while(b--)c=e[b],a[c]=g[c];return a.target||(a.target=g.srcElement||d),3===a.target.nodeType&&(a.target=a.target.parentNode),a.metaKey=!!a.metaKey,h.filter?h.filter(a,g):a},props:"altKey bubbles cancelable ctrlKey currentTarget detail eventPhase metaKey relatedTarget shiftKey target timeStamp view which".split(" "),fixHooks:{},keyHooks:{props:"char charCode key keyCode".split(" "),filter:function(a,b){return null==a.which&&(a.which=null!=b.charCode?b.charCode:b.keyCode),a}},mouseHooks:{props:"button buttons clientX clientY fromElement offsetX offsetY pageX pageY screenX screenY toElement".split(" "),filter:function(a,b){var c,e,f,g=b.button,h=b.fromElement;return null==a.pageX&&null!=b.clientX&&(e=a.target.ownerDocument||d,f=e.documentElement,c=e.body,a.pageX=b.clientX+(f&&f.scrollLeft||c&&c.scrollLeft||0)-(f&&f.clientLeft||c&&c.clientLeft||0),a.pageY=b.clientY+(f&&f.scrollTop||c&&c.scrollTop||0)-(f&&f.clientTop||c&&c.clientTop||0)),!a.relatedTarget&&h&&(a.relatedTarget=h===a.target?b.toElement:h),a.which||void 0===g||(a.which=1&g?1:2&g?3:4&g?2:0),a}},special:{load:{noBubble:!0},focus:{trigger:function(){if(this!==ra()&&this.focus)try{return this.focus(),!1}catch(a){}},delegateType:"focusin"},blur:{trigger:function(){return this===ra()&&this.blur?(this.blur(),!1):void 0},delegateType:"focusout"},click:{trigger:function(){return n.nodeName(this,"input")&&"checkbox"===this.type&&this.click?(this.click(),!1):void 0},_default:function(a){return n.nodeName(a.target,"a")}},beforeunload:{postDispatch:function(a){void 0!==a.result&&a.originalEvent&&(a.originalEvent.returnValue=a.result)}}},simulate:function(a,b,c){var d=n.extend(new n.Event,c,{type:a,isSimulated:!0});n.event.trigger(d,null,b),d.isDefaultPrevented()&&c.preventDefault()}},n.removeEvent=d.removeEventListener?function(a,b,c){a.removeEventListener&&a.removeEventListener(b,c)}:function(a,b,c){var d="on"+b;a.detachEvent&&("undefined"==typeof a[d]&&(a[d]=null),a.detachEvent(d,c))},n.Event=function(a,b){return this instanceof n.Event?(a&&a.type?(this.originalEvent=a,this.type=a.type,this.isDefaultPrevented=a.defaultPrevented||void 0===a.defaultPrevented&&a.returnValue===!1?pa:qa):this.type=a,b&&n.extend(this,b),this.timeStamp=a&&a.timeStamp||n.now(),void(this[n.expando]=!0)):new n.Event(a,b)},n.Event.prototype={constructor:n.Event,isDefaultPrevented:qa,isPropagationStopped:qa,isImmediatePropagationStopped:qa,preventDefault:function(){var a=this.originalEvent;this.isDefaultPrevented=pa,a&&(a.preventDefault?a.preventDefault():a.returnValue=!1)},stopPropagation:function(){var a=this.originalEvent;this.isPropagationStopped=pa,a&&!this.isSimulated&&(a.stopPropagation&&a.stopPropagation(),a.cancelBubble=!0)},stopImmediatePropagation:function(){var a=this.originalEvent;this.isImmediatePropagationStopped=pa,a&&a.stopImmediatePropagation&&a.stopImmediatePropagation(),this.stopPropagation()}},n.each({mouseenter:"mouseover",mouseleave:"mouseout",pointerenter:"pointerover",pointerleave:"pointerout"},function(a,b){n.event.special[a]={delegateType:b,bindType:b,handle:function(a){var c,d=this,e=a.relatedTarget,f=a.handleObj;return e&&(e===d||n.contains(d,e))||(a.type=f.origType,c=f.handler.apply(this,arguments),a.type=b),c}}}),l.submit||(n.event.special.submit={setup:function(){return n.nodeName(this,"form")?!1:void n.event.add(this,"click._submit keypress._submit",function(a){var b=a.target,c=n.nodeName(b,"input")||n.nodeName(b,"button")?n.prop(b,"form"):void 0;c&&!n._data(c,"submit")&&(n.event.add(c,"submit._submit",function(a){a._submitBubble=!0}),n._data(c,"submit",!0))})},postDispatch:function(a){a._submitBubble&&(delete a._submitBubble,this.parentNode&&!a.isTrigger&&n.event.simulate("submit",this.parentNode,a))},teardown:function(){return n.nodeName(this,"form")?!1:void n.event.remove(this,"._submit")}}),l.change||(n.event.special.change={setup:function(){return ka.test(this.nodeName)?("checkbox"!==this.type&&"radio"!==this.type||(n.event.add(this,"propertychange._change",function(a){"checked"===a.originalEvent.propertyName&&(this._justChanged=!0)}),n.event.add(this,"click._change",function(a){this._justChanged&&!a.isTrigger&&(this._justChanged=!1),n.event.simulate("change",this,a)})),!1):void n.event.add(this,"beforeactivate._change",function(a){var b=a.target;ka.test(b.nodeName)&&!n._data(b,"change")&&(n.event.add(b,"change._change",function(a){!this.parentNode||a.isSimulated||a.isTrigger||n.event.simulate("change",this.parentNode,a)}),n._data(b,"change",!0))})},handle:function(a){var b=a.target;return this!==b||a.isSimulated||a.isTrigger||"radio"!==b.type&&"checkbox"!==b.type?a.handleObj.handler.apply(this,arguments):void 0},teardown:function(){return n.event.remove(this,"._change"),!ka.test(this.nodeName)}}),l.focusin||n.each({focus:"focusin",blur:"focusout"},function(a,b){var c=function(a){n.event.simulate(b,a.target,n.event.fix(a))};n.event.special[b]={setup:function(){var d=this.ownerDocument||this,e=n._data(d,b);e||d.addEventListener(a,c,!0),n._data(d,b,(e||0)+1)},teardown:function(){var d=this.ownerDocument||this,e=n._data(d,b)-1;e?n._data(d,b,e):(d.removeEventListener(a,c,!0),n._removeData(d,b))}}}),n.fn.extend({on:function(a,b,c,d){return sa(this,a,b,c,d)},one:function(a,b,c,d){return sa(this,a,b,c,d,1)},off:function(a,b,c){var d,e;if(a&&a.preventDefault&&a.handleObj)return d=a.handleObj,n(a.delegateTarget).off(d.namespace?d.origType+"."+d.namespace:d.origType,d.selector,d.handler),this;if("object"==typeof a){for(e in a)this.off(e,b,a[e]);return this}return b!==!1&&"function"!=typeof b||(c=b,b=void 0),c===!1&&(c=qa),this.each(function(){n.event.remove(this,a,c,b)})},trigger:function(a,b){return this.each(function(){n.event.trigger(a,b,this)})},triggerHandler:function(a,b){var c=this[0];return c?n.event.trigger(a,b,c,!0):void 0}});var ta=/ jQuery\d+="(?:null|\d+)"/g,ua=new RegExp("<(?:"+ba+")[\\s/>]","i"),va=/<(?!area|br|col|embed|hr|img|input|link|meta|param)(([\w:-]+)[^>]*)\/>/gi,wa=/<script|<style|<link/i,xa=/checked\s*(?:[^=]|=\s*.checked.)/i,ya=/^true\/(.*)/,za=/^\s*<!(?:\[CDATA\[|--)|(?:\]\]|--)>\s*$/g,Aa=ca(d),Ba=Aa.appendChild(d.createElement("div"));function Ca(a,b){return n.nodeName(a,"table")&&n.nodeName(11!==b.nodeType?b:b.firstChild,"tr")?a.getElementsByTagName("tbody")[0]||a.appendChild(a.ownerDocument.createElement("tbody")):a}function Da(a){return a.type=(null!==n.find.attr(a,"type"))+"/"+a.type,a}function Ea(a){var b=ya.exec(a.type);return b?a.type=b[1]:a.removeAttribute("type"),a}function Fa(a,b){if(1===b.nodeType&&n.hasData(a)){var c,d,e,f=n._data(a),g=n._data(b,f),h=f.events;if(h){delete g.handle,g.events={};for(c in h)for(d=0,e=h[c].length;e>d;d++)n.event.add(b,c,h[c][d])}g.data&&(g.data=n.extend({},g.data))}}function Ga(a,b){var c,d,e;if(1===b.nodeType){if(c=b.nodeName.toLowerCase(),!l.noCloneEvent&&b[n.expando]){e=n._data(b);for(d in e.events)n.removeEvent(b,d,e.handle);b.removeAttribute(n.expando)}"script"===c&&b.text!==a.text?(Da(b).text=a.text,Ea(b)):"object"===c?(b.parentNode&&(b.outerHTML=a.outerHTML),l.html5Clone&&a.innerHTML&&!n.trim(b.innerHTML)&&(b.innerHTML=a.innerHTML)):"input"===c&&Z.test(a.type)?(b.defaultChecked=b.checked=a.checked,b.value!==a.value&&(b.value=a.value)):"option"===c?b.defaultSelected=b.selected=a.defaultSelected:"input"!==c&&"textarea"!==c||(b.defaultValue=a.defaultValue)}}function Ha(a,b,c,d){b=f.apply([],b);var e,g,h,i,j,k,m=0,o=a.length,p=o-1,q=b[0],r=n.isFunction(q);if(r||o>1&&"string"==typeof q&&!l.checkClone&&xa.test(q))return a.each(function(e){var f=a.eq(e);r&&(b[0]=q.call(this,e,f.html())),Ha(f,b,c,d)});if(o&&(k=ja(b,a[0].ownerDocument,!1,a,d),e=k.firstChild,1===k.childNodes.length&&(k=e),e||d)){for(i=n.map(ea(k,"script"),Da),h=i.length;o>m;m++)g=k,m!==p&&(g=n.clone(g,!0,!0),h&&n.merge(i,ea(g,"script"))),c.call(a[m],g,m);if(h)for(j=i[i.length-1].ownerDocument,n.map(i,Ea),m=0;h>m;m++)g=i[m],_.test(g.type||"")&&!n._data(g,"globalEval")&&n.contains(j,g)&&(g.src?n._evalUrl&&n._evalUrl(g.src):n.globalEval((g.text||g.textContent||g.innerHTML||"").replace(za,"")));k=e=null}return a}function Ia(a,b,c){for(var d,e=b?n.filter(b,a):a,f=0;null!=(d=e[f]);f++)c||1!==d.nodeType||n.cleanData(ea(d)),d.parentNode&&(c&&n.contains(d.ownerDocument,d)&&fa(ea(d,"script")),d.parentNode.removeChild(d));return a}n.extend({htmlPrefilter:function(a){return a.replace(va,"<$1></$2>")},clone:function(a,b,c){var d,e,f,g,h,i=n.contains(a.ownerDocument,a);if(l.html5Clone||n.isXMLDoc(a)||!ua.test("<"+a.nodeName+">")?f=a.cloneNode(!0):(Ba.innerHTML=a.outerHTML,Ba.removeChild(f=Ba.firstChild)),!(l.noCloneEvent&&l.noCloneChecked||1!==a.nodeType&&11!==a.nodeType||n.isXMLDoc(a)))for(d=ea(f),h=ea(a),g=0;null!=(e=h[g]);++g)d[g]&&Ga(e,d[g]);if(b)if(c)for(h=h||ea(a),d=d||ea(f),g=0;null!=(e=h[g]);g++)Fa(e,d[g]);else Fa(a,f);return d=ea(f,"script"),d.length>0&&fa(d,!i&&ea(a,"script")),d=h=e=null,f},cleanData:function(a,b){for(var d,e,f,g,h=0,i=n.expando,j=n.cache,k=l.attributes,m=n.event.special;null!=(d=a[h]);h++)if((b||M(d))&&(f=d[i],g=f&&j[f])){if(g.events)for(e in g.events)m[e]?n.event.remove(d,e):n.removeEvent(d,e,g.handle);j[f]&&(delete j[f],k||"undefined"==typeof d.removeAttribute?d[i]=void 0:d.removeAttribute(i),c.push(f))}}}),n.fn.extend({domManip:Ha,detach:function(a){return Ia(this,a,!0)},remove:function(a){return Ia(this,a)},text:function(a){return Y(this,function(a){return void 0===a?n.text(this):this.empty().append((this[0]&&this[0].ownerDocument||d).createTextNode(a))},null,a,arguments.length)},append:function(){return Ha(this,arguments,function(a){if(1===this.nodeType||11===this.nodeType||9===this.nodeType){var b=Ca(this,a);b.appendChild(a)}})},prepend:function(){return Ha(this,arguments,function(a){if(1===this.nodeType||11===this.nodeType||9===this.nodeType){var b=Ca(this,a);b.insertBefore(a,b.firstChild)}})},before:function(){return Ha(this,arguments,function(a){this.parentNode&&this.parentNode.insertBefore(a,this)})},after:function(){return Ha(this,arguments,function(a){this.parentNode&&this.parentNode.insertBefore(a,this.nextSibling)})},empty:function(){for(var a,b=0;null!=(a=this[b]);b++){1===a.nodeType&&n.cleanData(ea(a,!1));while(a.firstChild)a.removeChild(a.firstChild);a.options&&n.nodeName(a,"select")&&(a.options.length=0)}return this},clone:function(a,b){return a=null==a?!1:a,b=null==b?a:b,this.map(function(){return n.clone(this,a,b)})},html:function(a){return Y(this,function(a){var b=this[0]||{},c=0,d=this.length;if(void 0===a)return 1===b.nodeType?b.innerHTML.replace(ta,""):void 0;if("string"==typeof a&&!wa.test(a)&&(l.htmlSerialize||!ua.test(a))&&(l.leadingWhitespace||!aa.test(a))&&!da[($.exec(a)||["",""])[1].toLowerCase()]){a=n.htmlPrefilter(a);try{for(;d>c;c++)b=this[c]||{},1===b.nodeType&&(n.cleanData(ea(b,!1)),b.innerHTML=a);b=0}catch(e){}}b&&this.empty().append(a)},null,a,arguments.length)},replaceWith:function(){var a=[];return Ha(this,arguments,function(b){var c=this.parentNode;n.inArray(this,a)<0&&(n.cleanData(ea(this)),c&&c.replaceChild(b,this))},a)}}),n.each({appendTo:"append",prependTo:"prepend",insertBefore:"before",insertAfter:"after",replaceAll:"replaceWith"},function(a,b){n.fn[a]=function(a){for(var c,d=0,e=[],f=n(a),h=f.length-1;h>=d;d++)c=d===h?this:this.clone(!0),n(f[d])[b](c),g.apply(e,c.get());return this.pushStack(e)}});var Ja,Ka={HTML:"block",BODY:"block"};function La(a,b){var c=n(b.createElement(a)).appendTo(b.body),d=n.css(c[0],"display");return c.detach(),d}function Ma(a){var b=d,c=Ka[a];return c||(c=La(a,b),"none"!==c&&c||(Ja=(Ja||n("<iframe frameborder='0' width='0' height='0'/>")).appendTo(b.documentElement),b=(Ja[0].contentWindow||Ja[0].contentDocument).document,b.write(),b.close(),c=La(a,b),Ja.detach()),Ka[a]=c),c}var Na=/^margin/,Oa=new RegExp("^("+T+")(?!px)[a-z%]+$","i"),Pa=function(a,b,c,d){var e,f,g={};for(f in b)g[f]=a.style[f],a.style[f]=b[f];e=c.apply(a,d||[]);for(f in b)a.style[f]=g[f];return e},Qa=d.documentElement;!function(){var b,c,e,f,g,h,i=d.createElement("div"),j=d.createElement("div");if(j.style){j.style.cssText="float:left;opacity:.5",l.opacity="0.5"===j.style.opacity,l.cssFloat=!!j.style.cssFloat,j.style.backgroundClip="content-box",j.cloneNode(!0).style.backgroundClip="",l.clearCloneStyle="content-box"===j.style.backgroundClip,i=d.createElement("div"),i.style.cssText="border:0;width:8px;height:0;top:0;left:-9999px;padding:0;margin-top:1px;position:absolute",j.innerHTML="",i.appendChild(j),l.boxSizing=""===j.style.boxSizing||""===j.style.MozBoxSizing||""===j.style.WebkitBoxSizing,n.extend(l,{reliableHiddenOffsets:function(){return null==b&&k(),f},boxSizingReliable:function(){return null==b&&k(),e},pixelMarginRight:function(){return null==b&&k(),c},pixelPosition:function(){return null==b&&k(),b},reliableMarginRight:function(){return null==b&&k(),g},reliableMarginLeft:function(){return null==b&&k(),h}});function k(){var k,l,m=d.documentElement;m.appendChild(i),j.style.cssText="-webkit-box-sizing:border-box;box-sizing:border-box;position:relative;display:block;margin:auto;border:1px;padding:1px;top:1%;width:50%",b=e=h=!1,c=g=!0,a.getComputedStyle&&(l=a.getComputedStyle(j),b="1%"!==(l||{}).top,h="2px"===(l||{}).marginLeft,e="4px"===(l||{width:"4px"}).width,j.style.marginRight="50%",c="4px"===(l||{marginRight:"4px"}).marginRight,k=j.appendChild(d.createElement("div")),k.style.cssText=j.style.cssText="-webkit-box-sizing:content-box;-moz-box-sizing:content-box;box-sizing:content-box;display:block;margin:0;border:0;padding:0",k.style.marginRight=k.style.width="0",j.style.width="1px",g=!parseFloat((a.getComputedStyle(k)||{}).marginRight),j.removeChild(k)),j.style.display="none",f=0===j.getClientRects().length,f&&(j.style.display="",j.innerHTML="<table><tr><td></td><td>t</td></tr></table>",j.childNodes[0].style.borderCollapse="separate",k=j.getElementsByTagName("td"),k[0].style.cssText="margin:0;border:0;padding:0;display:none",f=0===k[0].offsetHeight,f&&(k[0].style.display="",k[1].style.display="none",f=0===k[0].offsetHeight)),m.removeChild(i)}}}();var Ra,Sa,Ta=/^(top|right|bottom|left)$/;a.getComputedStyle?(Ra=function(b){var c=b.ownerDocument.defaultView;return c&&c.opener||(c=a),c.getComputedStyle(b)},Sa=function(a,b,c){var d,e,f,g,h=a.style;return c=c||Ra(a),g=c?c.getPropertyValue(b)||c[b]:void 0,""!==g&&void 0!==g||n.contains(a.ownerDocument,a)||(g=n.style(a,b)),c&&!l.pixelMarginRight()&&Oa.test(g)&&Na.test(b)&&(d=h.width,e=h.minWidth,f=h.maxWidth,h.minWidth=h.maxWidth=h.width=g,g=c.width,h.width=d,h.minWidth=e,h.maxWidth=f),void 0===g?g:g+""}):Qa.currentStyle&&(Ra=function(a){return a.currentStyle},Sa=function(a,b,c){var d,e,f,g,h=a.style;return c=c||Ra(a),g=c?c[b]:void 0,null==g&&h&&h[b]&&(g=h[b]),Oa.test(g)&&!Ta.test(b)&&(d=h.left,e=a.runtimeStyle,f=e&&e.left,f&&(e.left=a.currentStyle.left),h.left="fontSize"===b?"1em":g,g=h.pixelLeft+"px",h.left=d,f&&(e.left=f)),void 0===g?g:g+""||"auto"});function Ua(a,b){return{get:function(){return a()?void delete this.get:(this.get=b).apply(this,arguments)}}}var Va=/alpha\([^)]*\)/i,Wa=/opacity\s*=\s*([^)]*)/i,Xa=/^(none|table(?!-c[ea]).+)/,Ya=new RegExp("^("+T+")(.*)$","i"),Za={position:"absolute",visibility:"hidden",display:"block"},$a={letterSpacing:"0",fontWeight:"400"},_a=["Webkit","O","Moz","ms"],ab=d.createElement("div").style;function bb(a){if(a in ab)return a;var b=a.charAt(0).toUpperCase()+a.slice(1),c=_a.length;while(c--)if(a=_a[c]+b,a in ab)return a}function cb(a,b){for(var c,d,e,f=[],g=0,h=a.length;h>g;g++)d=a[g],d.style&&(f[g]=n._data(d,"olddisplay"),c=d.style.display,b?(f[g]||"none"!==c||(d.style.display=""),""===d.style.display&&W(d)&&(f[g]=n._data(d,"olddisplay",Ma(d.nodeName)))):(e=W(d),(c&&"none"!==c||!e)&&n._data(d,"olddisplay",e?c:n.css(d,"display"))));for(g=0;h>g;g++)d=a[g],d.style&&(b&&"none"!==d.style.display&&""!==d.style.display||(d.style.display=b?f[g]||"":"none"));return a}function db(a,b,c){var d=Ya.exec(b);return d?Math.max(0,d[1]-(c||0))+(d[2]||"px"):b}function eb(a,b,c,d,e){for(var f=c===(d?"border":"content")?4:"width"===b?1:0,g=0;4>f;f+=2)"margin"===c&&(g+=n.css(a,c+V[f],!0,e)),d?("content"===c&&(g-=n.css(a,"padding"+V[f],!0,e)),"margin"!==c&&(g-=n.css(a,"border"+V[f]+"Width",!0,e))):(g+=n.css(a,"padding"+V[f],!0,e),"padding"!==c&&(g+=n.css(a,"border"+V[f]+"Width",!0,e)));return g}function fb(a,b,c){var d=!0,e="width"===b?a.offsetWidth:a.offsetHeight,f=Ra(a),g=l.boxSizing&&"border-box"===n.css(a,"boxSizing",!1,f);if(0>=e||null==e){if(e=Sa(a,b,f),(0>e||null==e)&&(e=a.style[b]),Oa.test(e))return e;d=g&&(l.boxSizingReliable()||e===a.style[b]),e=parseFloat(e)||0}return e+eb(a,b,c||(g?"border":"content"),d,f)+"px"}n.extend({cssHooks:{opacity:{get:function(a,b){if(b){var c=Sa(a,"opacity");return""===c?"1":c}}}},cssNumber:{animationIterationCount:!0,columnCount:!0,fillOpacity:!0,flexGrow:!0,flexShrink:!0,fontWeight:!0,lineHeight:!0,opacity:!0,order:!0,orphans:!0,widows:!0,zIndex:!0,zoom:!0},cssProps:{"float":l.cssFloat?"cssFloat":"styleFloat"},style:function(a,b,c,d){if(a&&3!==a.nodeType&&8!==a.nodeType&&a.style){var e,f,g,h=n.camelCase(b),i=a.style;if(b=n.cssProps[h]||(n.cssProps[h]=bb(h)||h),g=n.cssHooks[b]||n.cssHooks[h],void 0===c)return g&&"get"in g&&void 0!==(e=g.get(a,!1,d))?e:i[b];if(f=typeof c,"string"===f&&(e=U.exec(c))&&e[1]&&(c=X(a,b,e),f="number"),null!=c&&c===c&&("number"===f&&(c+=e&&e[3]||(n.cssNumber[h]?"":"px")),l.clearCloneStyle||""!==c||0!==b.indexOf("background")||(i[b]="inherit"),!(g&&"set"in g&&void 0===(c=g.set(a,c,d)))))try{i[b]=c}catch(j){}}},css:function(a,b,c,d){var e,f,g,h=n.camelCase(b);return b=n.cssProps[h]||(n.cssProps[h]=bb(h)||h),g=n.cssHooks[b]||n.cssHooks[h],g&&"get"in g&&(f=g.get(a,!0,c)),void 0===f&&(f=Sa(a,b,d)),"normal"===f&&b in $a&&(f=$a[b]),""===c||c?(e=parseFloat(f),c===!0||isFinite(e)?e||0:f):f}}),n.each(["height","width"],function(a,b){n.cssHooks[b]={get:function(a,c,d){return c?Xa.test(n.css(a,"display"))&&0===a.offsetWidth?Pa(a,Za,function(){return fb(a,b,d)}):fb(a,b,d):void 0},set:function(a,c,d){var e=d&&Ra(a);return db(a,c,d?eb(a,b,d,l.boxSizing&&"border-box"===n.css(a,"boxSizing",!1,e),e):0)}}}),l.opacity||(n.cssHooks.opacity={get:function(a,b){return Wa.test((b&&a.currentStyle?a.currentStyle.filter:a.style.filter)||"")?.01*parseFloat(RegExp.$1)+"":b?"1":""},set:function(a,b){var c=a.style,d=a.currentStyle,e=n.isNumeric(b)?"alpha(opacity="+100*b+")":"",f=d&&d.filter||c.filter||"";c.zoom=1,(b>=1||""===b)&&""===n.trim(f.replace(Va,""))&&c.removeAttribute&&(c.removeAttribute("filter"),""===b||d&&!d.filter)||(c.filter=Va.test(f)?f.replace(Va,e):f+" "+e)}}),n.cssHooks.marginRight=Ua(l.reliableMarginRight,function(a,b){return b?Pa(a,{display:"inline-block"},Sa,[a,"marginRight"]):void 0}),n.cssHooks.marginLeft=Ua(l.reliableMarginLeft,function(a,b){return b?(parseFloat(Sa(a,"marginLeft"))||(n.contains(a.ownerDocument,a)?a.getBoundingClientRect().left-Pa(a,{
marginLeft:0},function(){return a.getBoundingClientRect().left}):0))+"px":void 0}),n.each({margin:"",padding:"",border:"Width"},function(a,b){n.cssHooks[a+b]={expand:function(c){for(var d=0,e={},f="string"==typeof c?c.split(" "):[c];4>d;d++)e[a+V[d]+b]=f[d]||f[d-2]||f[0];return e}},Na.test(a)||(n.cssHooks[a+b].set=db)}),n.fn.extend({css:function(a,b){return Y(this,function(a,b,c){var d,e,f={},g=0;if(n.isArray(b)){for(d=Ra(a),e=b.length;e>g;g++)f[b[g]]=n.css(a,b[g],!1,d);return f}return void 0!==c?n.style(a,b,c):n.css(a,b)},a,b,arguments.length>1)},show:function(){return cb(this,!0)},hide:function(){return cb(this)},toggle:function(a){return"boolean"==typeof a?a?this.show():this.hide():this.each(function(){W(this)?n(this).show():n(this).hide()})}});function gb(a,b,c,d,e){return new gb.prototype.init(a,b,c,d,e)}n.Tween=gb,gb.prototype={constructor:gb,init:function(a,b,c,d,e,f){this.elem=a,this.prop=c,this.easing=e||n.easing._default,this.options=b,this.start=this.now=this.cur(),this.end=d,this.unit=f||(n.cssNumber[c]?"":"px")},cur:function(){var a=gb.propHooks[this.prop];return a&&a.get?a.get(this):gb.propHooks._default.get(this)},run:function(a){var b,c=gb.propHooks[this.prop];return this.options.duration?this.pos=b=n.easing[this.easing](a,this.options.duration*a,0,1,this.options.duration):this.pos=b=a,this.now=(this.end-this.start)*b+this.start,this.options.step&&this.options.step.call(this.elem,this.now,this),c&&c.set?c.set(this):gb.propHooks._default.set(this),this}},gb.prototype.init.prototype=gb.prototype,gb.propHooks={_default:{get:function(a){var b;return 1!==a.elem.nodeType||null!=a.elem[a.prop]&&null==a.elem.style[a.prop]?a.elem[a.prop]:(b=n.css(a.elem,a.prop,""),b&&"auto"!==b?b:0)},set:function(a){n.fx.step[a.prop]?n.fx.step[a.prop](a):1!==a.elem.nodeType||null==a.elem.style[n.cssProps[a.prop]]&&!n.cssHooks[a.prop]?a.elem[a.prop]=a.now:n.style(a.elem,a.prop,a.now+a.unit)}}},gb.propHooks.scrollTop=gb.propHooks.scrollLeft={set:function(a){a.elem.nodeType&&a.elem.parentNode&&(a.elem[a.prop]=a.now)}},n.easing={linear:function(a){return a},swing:function(a){return.5-Math.cos(a*Math.PI)/2},_default:"swing"},n.fx=gb.prototype.init,n.fx.step={};var hb,ib,jb=/^(?:toggle|show|hide)$/,kb=/queueHooks$/;function lb(){return a.setTimeout(function(){hb=void 0}),hb=n.now()}function mb(a,b){var c,d={height:a},e=0;for(b=b?1:0;4>e;e+=2-b)c=V[e],d["margin"+c]=d["padding"+c]=a;return b&&(d.opacity=d.width=a),d}function nb(a,b,c){for(var d,e=(qb.tweeners[b]||[]).concat(qb.tweeners["*"]),f=0,g=e.length;g>f;f++)if(d=e[f].call(c,b,a))return d}function ob(a,b,c){var d,e,f,g,h,i,j,k,m=this,o={},p=a.style,q=a.nodeType&&W(a),r=n._data(a,"fxshow");c.queue||(h=n._queueHooks(a,"fx"),null==h.unqueued&&(h.unqueued=0,i=h.empty.fire,h.empty.fire=function(){h.unqueued||i()}),h.unqueued++,m.always(function(){m.always(function(){h.unqueued--,n.queue(a,"fx").length||h.empty.fire()})})),1===a.nodeType&&("height"in b||"width"in b)&&(c.overflow=[p.overflow,p.overflowX,p.overflowY],j=n.css(a,"display"),k="none"===j?n._data(a,"olddisplay")||Ma(a.nodeName):j,"inline"===k&&"none"===n.css(a,"float")&&(l.inlineBlockNeedsLayout&&"inline"!==Ma(a.nodeName)?p.zoom=1:p.display="inline-block")),c.overflow&&(p.overflow="hidden",l.shrinkWrapBlocks()||m.always(function(){p.overflow=c.overflow[0],p.overflowX=c.overflow[1],p.overflowY=c.overflow[2]}));for(d in b)if(e=b[d],jb.exec(e)){if(delete b[d],f=f||"toggle"===e,e===(q?"hide":"show")){if("show"!==e||!r||void 0===r[d])continue;q=!0}o[d]=r&&r[d]||n.style(a,d)}else j=void 0;if(n.isEmptyObject(o))"inline"===("none"===j?Ma(a.nodeName):j)&&(p.display=j);else{r?"hidden"in r&&(q=r.hidden):r=n._data(a,"fxshow",{}),f&&(r.hidden=!q),q?n(a).show():m.done(function(){n(a).hide()}),m.done(function(){var b;n._removeData(a,"fxshow");for(b in o)n.style(a,b,o[b])});for(d in o)g=nb(q?r[d]:0,d,m),d in r||(r[d]=g.start,q&&(g.end=g.start,g.start="width"===d||"height"===d?1:0))}}function pb(a,b){var c,d,e,f,g;for(c in a)if(d=n.camelCase(c),e=b[d],f=a[c],n.isArray(f)&&(e=f[1],f=a[c]=f[0]),c!==d&&(a[d]=f,delete a[c]),g=n.cssHooks[d],g&&"expand"in g){f=g.expand(f),delete a[d];for(c in f)c in a||(a[c]=f[c],b[c]=e)}else b[d]=e}function qb(a,b,c){var d,e,f=0,g=qb.prefilters.length,h=n.Deferred().always(function(){delete i.elem}),i=function(){if(e)return!1;for(var b=hb||lb(),c=Math.max(0,j.startTime+j.duration-b),d=c/j.duration||0,f=1-d,g=0,i=j.tweens.length;i>g;g++)j.tweens[g].run(f);return h.notifyWith(a,[j,f,c]),1>f&&i?c:(h.resolveWith(a,[j]),!1)},j=h.promise({elem:a,props:n.extend({},b),opts:n.extend(!0,{specialEasing:{},easing:n.easing._default},c),originalProperties:b,originalOptions:c,startTime:hb||lb(),duration:c.duration,tweens:[],createTween:function(b,c){var d=n.Tween(a,j.opts,b,c,j.opts.specialEasing[b]||j.opts.easing);return j.tweens.push(d),d},stop:function(b){var c=0,d=b?j.tweens.length:0;if(e)return this;for(e=!0;d>c;c++)j.tweens[c].run(1);return b?(h.notifyWith(a,[j,1,0]),h.resolveWith(a,[j,b])):h.rejectWith(a,[j,b]),this}}),k=j.props;for(pb(k,j.opts.specialEasing);g>f;f++)if(d=qb.prefilters[f].call(j,a,k,j.opts))return n.isFunction(d.stop)&&(n._queueHooks(j.elem,j.opts.queue).stop=n.proxy(d.stop,d)),d;return n.map(k,nb,j),n.isFunction(j.opts.start)&&j.opts.start.call(a,j),n.fx.timer(n.extend(i,{elem:a,anim:j,queue:j.opts.queue})),j.progress(j.opts.progress).done(j.opts.done,j.opts.complete).fail(j.opts.fail).always(j.opts.always)}n.Animation=n.extend(qb,{tweeners:{"*":[function(a,b){var c=this.createTween(a,b);return X(c.elem,a,U.exec(b),c),c}]},tweener:function(a,b){n.isFunction(a)?(b=a,a=["*"]):a=a.match(G);for(var c,d=0,e=a.length;e>d;d++)c=a[d],qb.tweeners[c]=qb.tweeners[c]||[],qb.tweeners[c].unshift(b)},prefilters:[ob],prefilter:function(a,b){b?qb.prefilters.unshift(a):qb.prefilters.push(a)}}),n.speed=function(a,b,c){var d=a&&"object"==typeof a?n.extend({},a):{complete:c||!c&&b||n.isFunction(a)&&a,duration:a,easing:c&&b||b&&!n.isFunction(b)&&b};return d.duration=n.fx.off?0:"number"==typeof d.duration?d.duration:d.duration in n.fx.speeds?n.fx.speeds[d.duration]:n.fx.speeds._default,null!=d.queue&&d.queue!==!0||(d.queue="fx"),d.old=d.complete,d.complete=function(){n.isFunction(d.old)&&d.old.call(this),d.queue&&n.dequeue(this,d.queue)},d},n.fn.extend({fadeTo:function(a,b,c,d){return this.filter(W).css("opacity",0).show().end().animate({opacity:b},a,c,d)},animate:function(a,b,c,d){var e=n.isEmptyObject(a),f=n.speed(b,c,d),g=function(){var b=qb(this,n.extend({},a),f);(e||n._data(this,"finish"))&&b.stop(!0)};return g.finish=g,e||f.queue===!1?this.each(g):this.queue(f.queue,g)},stop:function(a,b,c){var d=function(a){var b=a.stop;delete a.stop,b(c)};return"string"!=typeof a&&(c=b,b=a,a=void 0),b&&a!==!1&&this.queue(a||"fx",[]),this.each(function(){var b=!0,e=null!=a&&a+"queueHooks",f=n.timers,g=n._data(this);if(e)g[e]&&g[e].stop&&d(g[e]);else for(e in g)g[e]&&g[e].stop&&kb.test(e)&&d(g[e]);for(e=f.length;e--;)f[e].elem!==this||null!=a&&f[e].queue!==a||(f[e].anim.stop(c),b=!1,f.splice(e,1));!b&&c||n.dequeue(this,a)})},finish:function(a){return a!==!1&&(a=a||"fx"),this.each(function(){var b,c=n._data(this),d=c[a+"queue"],e=c[a+"queueHooks"],f=n.timers,g=d?d.length:0;for(c.finish=!0,n.queue(this,a,[]),e&&e.stop&&e.stop.call(this,!0),b=f.length;b--;)f[b].elem===this&&f[b].queue===a&&(f[b].anim.stop(!0),f.splice(b,1));for(b=0;g>b;b++)d[b]&&d[b].finish&&d[b].finish.call(this);delete c.finish})}}),n.each(["toggle","show","hide"],function(a,b){var c=n.fn[b];n.fn[b]=function(a,d,e){return null==a||"boolean"==typeof a?c.apply(this,arguments):this.animate(mb(b,!0),a,d,e)}}),n.each({slideDown:mb("show"),slideUp:mb("hide"),slideToggle:mb("toggle"),fadeIn:{opacity:"show"},fadeOut:{opacity:"hide"},fadeToggle:{opacity:"toggle"}},function(a,b){n.fn[a]=function(a,c,d){return this.animate(b,a,c,d)}}),n.timers=[],n.fx.tick=function(){var a,b=n.timers,c=0;for(hb=n.now();c<b.length;c++)a=b[c],a()||b[c]!==a||b.splice(c--,1);b.length||n.fx.stop(),hb=void 0},n.fx.timer=function(a){n.timers.push(a),a()?n.fx.start():n.timers.pop()},n.fx.interval=13,n.fx.start=function(){ib||(ib=a.setInterval(n.fx.tick,n.fx.interval))},n.fx.stop=function(){a.clearInterval(ib),ib=null},n.fx.speeds={slow:600,fast:200,_default:400},n.fn.delay=function(b,c){return b=n.fx?n.fx.speeds[b]||b:b,c=c||"fx",this.queue(c,function(c,d){var e=a.setTimeout(c,b);d.stop=function(){a.clearTimeout(e)}})},function(){var a,b=d.createElement("input"),c=d.createElement("div"),e=d.createElement("select"),f=e.appendChild(d.createElement("option"));c=d.createElement("div"),c.setAttribute("className","t"),c.innerHTML="  <link/><table></table><a href='/a'>a</a><input type='checkbox'/>",a=c.getElementsByTagName("a")[0],b.setAttribute("type","checkbox"),c.appendChild(b),a=c.getElementsByTagName("a")[0],a.style.cssText="top:1px",l.getSetAttribute="t"!==c.className,l.style=/top/.test(a.getAttribute("style")),l.hrefNormalized="/a"===a.getAttribute("href"),l.checkOn=!!b.value,l.optSelected=f.selected,l.enctype=!!d.createElement("form").enctype,e.disabled=!0,l.optDisabled=!f.disabled,b=d.createElement("input"),b.setAttribute("value",""),l.input=""===b.getAttribute("value"),b.value="t",b.setAttribute("type","radio"),l.radioValue="t"===b.value}();var rb=/\r/g,sb=/[\x20\t\r\n\f]+/g;n.fn.extend({val:function(a){var b,c,d,e=this[0];{if(arguments.length)return d=n.isFunction(a),this.each(function(c){var e;1===this.nodeType&&(e=d?a.call(this,c,n(this).val()):a,null==e?e="":"number"==typeof e?e+="":n.isArray(e)&&(e=n.map(e,function(a){return null==a?"":a+""})),b=n.valHooks[this.type]||n.valHooks[this.nodeName.toLowerCase()],b&&"set"in b&&void 0!==b.set(this,e,"value")||(this.value=e))});if(e)return b=n.valHooks[e.type]||n.valHooks[e.nodeName.toLowerCase()],b&&"get"in b&&void 0!==(c=b.get(e,"value"))?c:(c=e.value,"string"==typeof c?c.replace(rb,""):null==c?"":c)}}}),n.extend({valHooks:{option:{get:function(a){var b=n.find.attr(a,"value");return null!=b?b:n.trim(n.text(a)).replace(sb," ")}},select:{get:function(a){for(var b,c,d=a.options,e=a.selectedIndex,f="select-one"===a.type||0>e,g=f?null:[],h=f?e+1:d.length,i=0>e?h:f?e:0;h>i;i++)if(c=d[i],(c.selected||i===e)&&(l.optDisabled?!c.disabled:null===c.getAttribute("disabled"))&&(!c.parentNode.disabled||!n.nodeName(c.parentNode,"optgroup"))){if(b=n(c).val(),f)return b;g.push(b)}return g},set:function(a,b){var c,d,e=a.options,f=n.makeArray(b),g=e.length;while(g--)if(d=e[g],n.inArray(n.valHooks.option.get(d),f)>-1)try{d.selected=c=!0}catch(h){d.scrollHeight}else d.selected=!1;return c||(a.selectedIndex=-1),e}}}}),n.each(["radio","checkbox"],function(){n.valHooks[this]={set:function(a,b){return n.isArray(b)?a.checked=n.inArray(n(a).val(),b)>-1:void 0}},l.checkOn||(n.valHooks[this].get=function(a){return null===a.getAttribute("value")?"on":a.value})});var tb,ub,vb=n.expr.attrHandle,wb=/^(?:checked|selected)$/i,xb=l.getSetAttribute,yb=l.input;n.fn.extend({attr:function(a,b){return Y(this,n.attr,a,b,arguments.length>1)},removeAttr:function(a){return this.each(function(){n.removeAttr(this,a)})}}),n.extend({attr:function(a,b,c){var d,e,f=a.nodeType;if(3!==f&&8!==f&&2!==f)return"undefined"==typeof a.getAttribute?n.prop(a,b,c):(1===f&&n.isXMLDoc(a)||(b=b.toLowerCase(),e=n.attrHooks[b]||(n.expr.match.bool.test(b)?ub:tb)),void 0!==c?null===c?void n.removeAttr(a,b):e&&"set"in e&&void 0!==(d=e.set(a,c,b))?d:(a.setAttribute(b,c+""),c):e&&"get"in e&&null!==(d=e.get(a,b))?d:(d=n.find.attr(a,b),null==d?void 0:d))},attrHooks:{type:{set:function(a,b){if(!l.radioValue&&"radio"===b&&n.nodeName(a,"input")){var c=a.value;return a.setAttribute("type",b),c&&(a.value=c),b}}}},removeAttr:function(a,b){var c,d,e=0,f=b&&b.match(G);if(f&&1===a.nodeType)while(c=f[e++])d=n.propFix[c]||c,n.expr.match.bool.test(c)?yb&&xb||!wb.test(c)?a[d]=!1:a[n.camelCase("default-"+c)]=a[d]=!1:n.attr(a,c,""),a.removeAttribute(xb?c:d)}}),ub={set:function(a,b,c){return b===!1?n.removeAttr(a,c):yb&&xb||!wb.test(c)?a.setAttribute(!xb&&n.propFix[c]||c,c):a[n.camelCase("default-"+c)]=a[c]=!0,c}},n.each(n.expr.match.bool.source.match(/\w+/g),function(a,b){var c=vb[b]||n.find.attr;yb&&xb||!wb.test(b)?vb[b]=function(a,b,d){var e,f;return d||(f=vb[b],vb[b]=e,e=null!=c(a,b,d)?b.toLowerCase():null,vb[b]=f),e}:vb[b]=function(a,b,c){return c?void 0:a[n.camelCase("default-"+b)]?b.toLowerCase():null}}),yb&&xb||(n.attrHooks.value={set:function(a,b,c){return n.nodeName(a,"input")?void(a.defaultValue=b):tb&&tb.set(a,b,c)}}),xb||(tb={set:function(a,b,c){var d=a.getAttributeNode(c);return d||a.setAttributeNode(d=a.ownerDocument.createAttribute(c)),d.value=b+="","value"===c||b===a.getAttribute(c)?b:void 0}},vb.id=vb.name=vb.coords=function(a,b,c){var d;return c?void 0:(d=a.getAttributeNode(b))&&""!==d.value?d.value:null},n.valHooks.button={get:function(a,b){var c=a.getAttributeNode(b);return c&&c.specified?c.value:void 0},set:tb.set},n.attrHooks.contenteditable={set:function(a,b,c){tb.set(a,""===b?!1:b,c)}},n.each(["width","height"],function(a,b){n.attrHooks[b]={set:function(a,c){return""===c?(a.setAttribute(b,"auto"),c):void 0}}})),l.style||(n.attrHooks.style={get:function(a){return a.style.cssText||void 0},set:function(a,b){return a.style.cssText=b+""}});var zb=/^(?:input|select|textarea|button|object)$/i,Ab=/^(?:a|area)$/i;n.fn.extend({prop:function(a,b){return Y(this,n.prop,a,b,arguments.length>1)},removeProp:function(a){return a=n.propFix[a]||a,this.each(function(){try{this[a]=void 0,delete this[a]}catch(b){}})}}),n.extend({prop:function(a,b,c){var d,e,f=a.nodeType;if(3!==f&&8!==f&&2!==f)return 1===f&&n.isXMLDoc(a)||(b=n.propFix[b]||b,e=n.propHooks[b]),void 0!==c?e&&"set"in e&&void 0!==(d=e.set(a,c,b))?d:a[b]=c:e&&"get"in e&&null!==(d=e.get(a,b))?d:a[b]},propHooks:{tabIndex:{get:function(a){var b=n.find.attr(a,"tabindex");return b?parseInt(b,10):zb.test(a.nodeName)||Ab.test(a.nodeName)&&a.href?0:-1}}},propFix:{"for":"htmlFor","class":"className"}}),l.hrefNormalized||n.each(["href","src"],function(a,b){n.propHooks[b]={get:function(a){return a.getAttribute(b,4)}}}),l.optSelected||(n.propHooks.selected={get:function(a){var b=a.parentNode;return b&&(b.selectedIndex,b.parentNode&&b.parentNode.selectedIndex),null},set:function(a){var b=a.parentNode;b&&(b.selectedIndex,b.parentNode&&b.parentNode.selectedIndex)}}),n.each(["tabIndex","readOnly","maxLength","cellSpacing","cellPadding","rowSpan","colSpan","useMap","frameBorder","contentEditable"],function(){n.propFix[this.toLowerCase()]=this}),l.enctype||(n.propFix.enctype="encoding");var Bb=/[\t\r\n\f]/g;function Cb(a){return n.attr(a,"class")||""}n.fn.extend({addClass:function(a){var b,c,d,e,f,g,h,i=0;if(n.isFunction(a))return this.each(function(b){n(this).addClass(a.call(this,b,Cb(this)))});if("string"==typeof a&&a){b=a.match(G)||[];while(c=this[i++])if(e=Cb(c),d=1===c.nodeType&&(" "+e+" ").replace(Bb," ")){g=0;while(f=b[g++])d.indexOf(" "+f+" ")<0&&(d+=f+" ");h=n.trim(d),e!==h&&n.attr(c,"class",h)}}return this},removeClass:function(a){var b,c,d,e,f,g,h,i=0;if(n.isFunction(a))return this.each(function(b){n(this).removeClass(a.call(this,b,Cb(this)))});if(!arguments.length)return this.attr("class","");if("string"==typeof a&&a){b=a.match(G)||[];while(c=this[i++])if(e=Cb(c),d=1===c.nodeType&&(" "+e+" ").replace(Bb," ")){g=0;while(f=b[g++])while(d.indexOf(" "+f+" ")>-1)d=d.replace(" "+f+" "," ");h=n.trim(d),e!==h&&n.attr(c,"class",h)}}return this},toggleClass:function(a,b){var c=typeof a;return"boolean"==typeof b&&"string"===c?b?this.addClass(a):this.removeClass(a):n.isFunction(a)?this.each(function(c){n(this).toggleClass(a.call(this,c,Cb(this),b),b)}):this.each(function(){var b,d,e,f;if("string"===c){d=0,e=n(this),f=a.match(G)||[];while(b=f[d++])e.hasClass(b)?e.removeClass(b):e.addClass(b)}else void 0!==a&&"boolean"!==c||(b=Cb(this),b&&n._data(this,"__className__",b),n.attr(this,"class",b||a===!1?"":n._data(this,"__className__")||""))})},hasClass:function(a){var b,c,d=0;b=" "+a+" ";while(c=this[d++])if(1===c.nodeType&&(" "+Cb(c)+" ").replace(Bb," ").indexOf(b)>-1)return!0;return!1}}),n.each("blur focus focusin focusout load resize scroll unload click dblclick mousedown mouseup mousemove mouseover mouseout mouseenter mouseleave change select submit keydown keypress keyup error contextmenu".split(" "),function(a,b){n.fn[b]=function(a,c){return arguments.length>0?this.on(b,null,a,c):this.trigger(b)}}),n.fn.extend({hover:function(a,b){return this.mouseenter(a).mouseleave(b||a)}});var Db=a.location,Eb=n.now(),Fb=/\?/,Gb=/(,)|(\[|{)|(}|])|"(?:[^"\\\r\n]|\\["\\\/bfnrt]|\\u[\da-fA-F]{4})*"\s*:?|true|false|null|-?(?!0\d)\d+(?:\.\d+|)(?:[eE][+-]?\d+|)/g;n.parseJSON=function(b){if(a.JSON&&a.JSON.parse)return a.JSON.parse(b+"");var c,d=null,e=n.trim(b+"");return e&&!n.trim(e.replace(Gb,function(a,b,e,f){return c&&b&&(d=0),0===d?a:(c=e||b,d+=!f-!e,"")}))?Function("return "+e)():n.error("Invalid JSON: "+b)},n.parseXML=function(b){var c,d;if(!b||"string"!=typeof b)return null;try{a.DOMParser?(d=new a.DOMParser,c=d.parseFromString(b,"text/xml")):(c=new a.ActiveXObject("Microsoft.XMLDOM"),c.async="false",c.loadXML(b))}catch(e){c=void 0}return c&&c.documentElement&&!c.getElementsByTagName("parsererror").length||n.error("Invalid XML: "+b),c};var Hb=/#.*$/,Ib=/([?&])_=[^&]*/,Jb=/^(.*?):[ \t]*([^\r\n]*)\r?$/gm,Kb=/^(?:about|app|app-storage|.+-extension|file|res|widget):$/,Lb=/^(?:GET|HEAD)$/,Mb=/^\/\//,Nb=/^([\w.+-]+:)(?:\/\/(?:[^\/?#]*@|)([^\/?#:]*)(?::(\d+)|)|)/,Ob={},Pb={},Qb="*/".concat("*"),Rb=Db.href,Sb=Nb.exec(Rb.toLowerCase())||[];function Tb(a){return function(b,c){"string"!=typeof b&&(c=b,b="*");var d,e=0,f=b.toLowerCase().match(G)||[];if(n.isFunction(c))while(d=f[e++])"+"===d.charAt(0)?(d=d.slice(1)||"*",(a[d]=a[d]||[]).unshift(c)):(a[d]=a[d]||[]).push(c)}}function Ub(a,b,c,d){var e={},f=a===Pb;function g(h){var i;return e[h]=!0,n.each(a[h]||[],function(a,h){var j=h(b,c,d);return"string"!=typeof j||f||e[j]?f?!(i=j):void 0:(b.dataTypes.unshift(j),g(j),!1)}),i}return g(b.dataTypes[0])||!e["*"]&&g("*")}function Vb(a,b){var c,d,e=n.ajaxSettings.flatOptions||{};for(d in b)void 0!==b[d]&&((e[d]?a:c||(c={}))[d]=b[d]);return c&&n.extend(!0,a,c),a}function Wb(a,b,c){var d,e,f,g,h=a.contents,i=a.dataTypes;while("*"===i[0])i.shift(),void 0===e&&(e=a.mimeType||b.getResponseHeader("Content-Type"));if(e)for(g in h)if(h[g]&&h[g].test(e)){i.unshift(g);break}if(i[0]in c)f=i[0];else{for(g in c){if(!i[0]||a.converters[g+" "+i[0]]){f=g;break}d||(d=g)}f=f||d}return f?(f!==i[0]&&i.unshift(f),c[f]):void 0}function Xb(a,b,c,d){var e,f,g,h,i,j={},k=a.dataTypes.slice();if(k[1])for(g in a.converters)j[g.toLowerCase()]=a.converters[g];f=k.shift();while(f)if(a.responseFields[f]&&(c[a.responseFields[f]]=b),!i&&d&&a.dataFilter&&(b=a.dataFilter(b,a.dataType)),i=f,f=k.shift())if("*"===f)f=i;else if("*"!==i&&i!==f){if(g=j[i+" "+f]||j["* "+f],!g)for(e in j)if(h=e.split(" "),h[1]===f&&(g=j[i+" "+h[0]]||j["* "+h[0]])){g===!0?g=j[e]:j[e]!==!0&&(f=h[0],k.unshift(h[1]));break}if(g!==!0)if(g&&a["throws"])b=g(b);else try{b=g(b)}catch(l){return{state:"parsererror",error:g?l:"No conversion from "+i+" to "+f}}}return{state:"success",data:b}}n.extend({active:0,lastModified:{},etag:{},ajaxSettings:{url:Rb,type:"GET",isLocal:Kb.test(Sb[1]),global:!0,processData:!0,async:!0,contentType:"application/x-www-form-urlencoded; charset=UTF-8",accepts:{"*":Qb,text:"text/plain",html:"text/html",xml:"application/xml, text/xml",json:"application/json, text/javascript"},contents:{xml:/\bxml\b/,html:/\bhtml/,json:/\bjson\b/},responseFields:{xml:"responseXML",text:"responseText",json:"responseJSON"},converters:{"* text":String,"text html":!0,"text json":n.parseJSON,"text xml":n.parseXML},flatOptions:{url:!0,context:!0}},ajaxSetup:function(a,b){return b?Vb(Vb(a,n.ajaxSettings),b):Vb(n.ajaxSettings,a)},ajaxPrefilter:Tb(Ob),ajaxTransport:Tb(Pb),ajax:function(b,c){"object"==typeof b&&(c=b,b=void 0),c=c||{};var d,e,f,g,h,i,j,k,l=n.ajaxSetup({},c),m=l.context||l,o=l.context&&(m.nodeType||m.jquery)?n(m):n.event,p=n.Deferred(),q=n.Callbacks("once memory"),r=l.statusCode||{},s={},t={},u=0,v="canceled",w={readyState:0,getResponseHeader:function(a){var b;if(2===u){if(!k){k={};while(b=Jb.exec(g))k[b[1].toLowerCase()]=b[2]}b=k[a.toLowerCase()]}return null==b?null:b},getAllResponseHeaders:function(){return 2===u?g:null},setRequestHeader:function(a,b){var c=a.toLowerCase();return u||(a=t[c]=t[c]||a,s[a]=b),this},overrideMimeType:function(a){return u||(l.mimeType=a),this},statusCode:function(a){var b;if(a)if(2>u)for(b in a)r[b]=[r[b],a[b]];else w.always(a[w.status]);return this},abort:function(a){var b=a||v;return j&&j.abort(b),y(0,b),this}};if(p.promise(w).complete=q.add,w.success=w.done,w.error=w.fail,l.url=((b||l.url||Rb)+"").replace(Hb,"").replace(Mb,Sb[1]+"//"),l.type=c.method||c.type||l.method||l.type,l.dataTypes=n.trim(l.dataType||"*").toLowerCase().match(G)||[""],null==l.crossDomain&&(d=Nb.exec(l.url.toLowerCase()),l.crossDomain=!(!d||d[1]===Sb[1]&&d[2]===Sb[2]&&(d[3]||("http:"===d[1]?"80":"443"))===(Sb[3]||("http:"===Sb[1]?"80":"443")))),l.data&&l.processData&&"string"!=typeof l.data&&(l.data=n.param(l.data,l.traditional)),Ub(Ob,l,c,w),2===u)return w;i=n.event&&l.global,i&&0===n.active++&&n.event.trigger("ajaxStart"),l.type=l.type.toUpperCase(),l.hasContent=!Lb.test(l.type),f=l.url,l.hasContent||(l.data&&(f=l.url+=(Fb.test(f)?"&":"?")+l.data,delete l.data),l.cache===!1&&(l.url=Ib.test(f)?f.replace(Ib,"$1_="+Eb++):f+(Fb.test(f)?"&":"?")+"_="+Eb++)),l.ifModified&&(n.lastModified[f]&&w.setRequestHeader("If-Modified-Since",n.lastModified[f]),n.etag[f]&&w.setRequestHeader("If-None-Match",n.etag[f])),(l.data&&l.hasContent&&l.contentType!==!1||c.contentType)&&w.setRequestHeader("Content-Type",l.contentType),w.setRequestHeader("Accept",l.dataTypes[0]&&l.accepts[l.dataTypes[0]]?l.accepts[l.dataTypes[0]]+("*"!==l.dataTypes[0]?", "+Qb+"; q=0.01":""):l.accepts["*"]);for(e in l.headers)w.setRequestHeader(e,l.headers[e]);if(l.beforeSend&&(l.beforeSend.call(m,w,l)===!1||2===u))return w.abort();v="abort";for(e in{success:1,error:1,complete:1})w[e](l[e]);if(j=Ub(Pb,l,c,w)){if(w.readyState=1,i&&o.trigger("ajaxSend",[w,l]),2===u)return w;l.async&&l.timeout>0&&(h=a.setTimeout(function(){w.abort("timeout")},l.timeout));try{u=1,j.send(s,y)}catch(x){if(!(2>u))throw x;y(-1,x)}}else y(-1,"No Transport");function y(b,c,d,e){var k,s,t,v,x,y=c;2!==u&&(u=2,h&&a.clearTimeout(h),j=void 0,g=e||"",w.readyState=b>0?4:0,k=b>=200&&300>b||304===b,d&&(v=Wb(l,w,d)),v=Xb(l,v,w,k),k?(l.ifModified&&(x=w.getResponseHeader("Last-Modified"),x&&(n.lastModified[f]=x),x=w.getResponseHeader("etag"),x&&(n.etag[f]=x)),204===b||"HEAD"===l.type?y="nocontent":304===b?y="notmodified":(y=v.state,s=v.data,t=v.error,k=!t)):(t=y,!b&&y||(y="error",0>b&&(b=0))),w.status=b,w.statusText=(c||y)+"",k?p.resolveWith(m,[s,y,w]):p.rejectWith(m,[w,y,t]),w.statusCode(r),r=void 0,i&&o.trigger(k?"ajaxSuccess":"ajaxError",[w,l,k?s:t]),q.fireWith(m,[w,y]),i&&(o.trigger("ajaxComplete",[w,l]),--n.active||n.event.trigger("ajaxStop")))}return w},getJSON:function(a,b,c){return n.get(a,b,c,"json")},getScript:function(a,b){return n.get(a,void 0,b,"script")}}),n.each(["get","post"],function(a,b){n[b]=function(a,c,d,e){return n.isFunction(c)&&(e=e||d,d=c,c=void 0),n.ajax(n.extend({url:a,type:b,dataType:e,data:c,success:d},n.isPlainObject(a)&&a))}}),n._evalUrl=function(a){return n.ajax({url:a,type:"GET",dataType:"script",cache:!0,async:!1,global:!1,"throws":!0})},n.fn.extend({wrapAll:function(a){if(n.isFunction(a))return this.each(function(b){n(this).wrapAll(a.call(this,b))});if(this[0]){var b=n(a,this[0].ownerDocument).eq(0).clone(!0);this[0].parentNode&&b.insertBefore(this[0]),b.map(function(){var a=this;while(a.firstChild&&1===a.firstChild.nodeType)a=a.firstChild;return a}).append(this)}return this},wrapInner:function(a){return n.isFunction(a)?this.each(function(b){n(this).wrapInner(a.call(this,b))}):this.each(function(){var b=n(this),c=b.contents();c.length?c.wrapAll(a):b.append(a)})},wrap:function(a){var b=n.isFunction(a);return this.each(function(c){n(this).wrapAll(b?a.call(this,c):a)})},unwrap:function(){return this.parent().each(function(){n.nodeName(this,"body")||n(this).replaceWith(this.childNodes)}).end()}});function Yb(a){return a.style&&a.style.display||n.css(a,"display")}function Zb(a){if(!n.contains(a.ownerDocument||d,a))return!0;while(a&&1===a.nodeType){if("none"===Yb(a)||"hidden"===a.type)return!0;a=a.parentNode}return!1}n.expr.filters.hidden=function(a){return l.reliableHiddenOffsets()?a.offsetWidth<=0&&a.offsetHeight<=0&&!a.getClientRects().length:Zb(a)},n.expr.filters.visible=function(a){return!n.expr.filters.hidden(a)};var $b=/%20/g,_b=/\[\]$/,ac=/\r?\n/g,bc=/^(?:submit|button|image|reset|file)$/i,cc=/^(?:input|select|textarea|keygen)/i;function dc(a,b,c,d){var e;if(n.isArray(b))n.each(b,function(b,e){c||_b.test(a)?d(a,e):dc(a+"["+("object"==typeof e&&null!=e?b:"")+"]",e,c,d)});else if(c||"object"!==n.type(b))d(a,b);else for(e in b)dc(a+"["+e+"]",b[e],c,d)}n.param=function(a,b){var c,d=[],e=function(a,b){b=n.isFunction(b)?b():null==b?"":b,d[d.length]=encodeURIComponent(a)+"="+encodeURIComponent(b)};if(void 0===b&&(b=n.ajaxSettings&&n.ajaxSettings.traditional),n.isArray(a)||a.jquery&&!n.isPlainObject(a))n.each(a,function(){e(this.name,this.value)});else for(c in a)dc(c,a[c],b,e);return d.join("&").replace($b,"+")},n.fn.extend({serialize:function(){return n.param(this.serializeArray())},serializeArray:function(){return this.map(function(){var a=n.prop(this,"elements");return a?n.makeArray(a):this}).filter(function(){var a=this.type;return this.name&&!n(this).is(":disabled")&&cc.test(this.nodeName)&&!bc.test(a)&&(this.checked||!Z.test(a))}).map(function(a,b){var c=n(this).val();return null==c?null:n.isArray(c)?n.map(c,function(a){return{name:b.name,value:a.replace(ac,"\r\n")}}):{name:b.name,value:c.replace(ac,"\r\n")}}).get()}}),n.ajaxSettings.xhr=void 0!==a.ActiveXObject?function(){return this.isLocal?ic():d.documentMode>8?hc():/^(get|post|head|put|delete|options)$/i.test(this.type)&&hc()||ic()}:hc;var ec=0,fc={},gc=n.ajaxSettings.xhr();a.attachEvent&&a.attachEvent("onunload",function(){for(var a in fc)fc[a](void 0,!0)}),l.cors=!!gc&&"withCredentials"in gc,gc=l.ajax=!!gc,gc&&n.ajaxTransport(function(b){if(!b.crossDomain||l.cors){var c;return{send:function(d,e){var f,g=b.xhr(),h=++ec;if(g.open(b.type,b.url,b.async,b.username,b.password),b.xhrFields)for(f in b.xhrFields)g[f]=b.xhrFields[f];b.mimeType&&g.overrideMimeType&&g.overrideMimeType(b.mimeType),b.crossDomain||d["X-Requested-With"]||(d["X-Requested-With"]="XMLHttpRequest");for(f in d)void 0!==d[f]&&g.setRequestHeader(f,d[f]+"");g.send(b.hasContent&&b.data||null),c=function(a,d){var f,i,j;if(c&&(d||4===g.readyState))if(delete fc[h],c=void 0,g.onreadystatechange=n.noop,d)4!==g.readyState&&g.abort();else{j={},f=g.status,"string"==typeof g.responseText&&(j.text=g.responseText);try{i=g.statusText}catch(k){i=""}f||!b.isLocal||b.crossDomain?1223===f&&(f=204):f=j.text?200:404}j&&e(f,i,j,g.getAllResponseHeaders())},b.async?4===g.readyState?a.setTimeout(c):g.onreadystatechange=fc[h]=c:c()},abort:function(){c&&c(void 0,!0)}}}});function hc(){try{return new a.XMLHttpRequest}catch(b){}}function ic(){try{return new a.ActiveXObject("Microsoft.XMLHTTP")}catch(b){}}n.ajaxSetup({accepts:{script:"text/javascript, application/javascript, application/ecmascript, application/x-ecmascript"},contents:{script:/\b(?:java|ecma)script\b/},converters:{"text script":function(a){return n.globalEval(a),a}}}),n.ajaxPrefilter("script",function(a){void 0===a.cache&&(a.cache=!1),a.crossDomain&&(a.type="GET",a.global=!1)}),n.ajaxTransport("script",function(a){if(a.crossDomain){var b,c=d.head||n("head")[0]||d.documentElement;return{send:function(e,f){b=d.createElement("script"),b.async=!0,a.scriptCharset&&(b.charset=a.scriptCharset),b.src=a.url,b.onload=b.onreadystatechange=function(a,c){(c||!b.readyState||/loaded|complete/.test(b.readyState))&&(b.onload=b.onreadystatechange=null,b.parentNode&&b.parentNode.removeChild(b),b=null,c||f(200,"success"))},c.insertBefore(b,c.firstChild)},abort:function(){b&&b.onload(void 0,!0)}}}});var jc=[],kc=/(=)\?(?=&|$)|\?\?/;n.ajaxSetup({jsonp:"callback",jsonpCallback:function(){var a=jc.pop()||n.expando+"_"+Eb++;return this[a]=!0,a}}),n.ajaxPrefilter("json jsonp",function(b,c,d){var e,f,g,h=b.jsonp!==!1&&(kc.test(b.url)?"url":"string"==typeof b.data&&0===(b.contentType||"").indexOf("application/x-www-form-urlencoded")&&kc.test(b.data)&&"data");return h||"jsonp"===b.dataTypes[0]?(e=b.jsonpCallback=n.isFunction(b.jsonpCallback)?b.jsonpCallback():b.jsonpCallback,h?b[h]=b[h].replace(kc,"$1"+e):b.jsonp!==!1&&(b.url+=(Fb.test(b.url)?"&":"?")+b.jsonp+"="+e),b.converters["script json"]=function(){return g||n.error(e+" was not called"),g[0]},b.dataTypes[0]="json",f=a[e],a[e]=function(){g=arguments},d.always(function(){void 0===f?n(a).removeProp(e):a[e]=f,b[e]&&(b.jsonpCallback=c.jsonpCallback,jc.push(e)),g&&n.isFunction(f)&&f(g[0]),g=f=void 0}),"script"):void 0}),n.parseHTML=function(a,b,c){if(!a||"string"!=typeof a)return null;"boolean"==typeof b&&(c=b,b=!1),b=b||d;var e=x.exec(a),f=!c&&[];return e?[b.createElement(e[1])]:(e=ja([a],b,f),f&&f.length&&n(f).remove(),n.merge([],e.childNodes))};var lc=n.fn.load;n.fn.load=function(a,b,c){if("string"!=typeof a&&lc)return lc.apply(this,arguments);var d,e,f,g=this,h=a.indexOf(" ");return h>-1&&(d=n.trim(a.slice(h,a.length)),a=a.slice(0,h)),n.isFunction(b)?(c=b,b=void 0):b&&"object"==typeof b&&(e="POST"),g.length>0&&n.ajax({url:a,type:e||"GET",dataType:"html",data:b}).done(function(a){f=arguments,g.html(d?n("<div>").append(n.parseHTML(a)).find(d):a)}).always(c&&function(a,b){g.each(function(){c.apply(this,f||[a.responseText,b,a])})}),this},n.each(["ajaxStart","ajaxStop","ajaxComplete","ajaxError","ajaxSuccess","ajaxSend"],function(a,b){n.fn[b]=function(a){return this.on(b,a)}}),n.expr.filters.animated=function(a){return n.grep(n.timers,function(b){return a===b.elem}).length};function mc(a){return n.isWindow(a)?a:9===a.nodeType?a.defaultView||a.parentWindow:!1}n.offset={setOffset:function(a,b,c){var d,e,f,g,h,i,j,k=n.css(a,"position"),l=n(a),m={};"static"===k&&(a.style.position="relative"),h=l.offset(),f=n.css(a,"top"),i=n.css(a,"left"),j=("absolute"===k||"fixed"===k)&&n.inArray("auto",[f,i])>-1,j?(d=l.position(),g=d.top,e=d.left):(g=parseFloat(f)||0,e=parseFloat(i)||0),n.isFunction(b)&&(b=b.call(a,c,n.extend({},h))),null!=b.top&&(m.top=b.top-h.top+g),null!=b.left&&(m.left=b.left-h.left+e),"using"in b?b.using.call(a,m):l.css(m)}},n.fn.extend({offset:function(a){if(arguments.length)return void 0===a?this:this.each(function(b){n.offset.setOffset(this,a,b)});var b,c,d={top:0,left:0},e=this[0],f=e&&e.ownerDocument;if(f)return b=f.documentElement,n.contains(b,e)?("undefined"!=typeof e.getBoundingClientRect&&(d=e.getBoundingClientRect()),c=mc(f),{top:d.top+(c.pageYOffset||b.scrollTop)-(b.clientTop||0),left:d.left+(c.pageXOffset||b.scrollLeft)-(b.clientLeft||0)}):d},position:function(){if(this[0]){var a,b,c={top:0,left:0},d=this[0];return"fixed"===n.css(d,"position")?b=d.getBoundingClientRect():(a=this.offsetParent(),b=this.offset(),n.nodeName(a[0],"html")||(c=a.offset()),c.top+=n.css(a[0],"borderTopWidth",!0),c.left+=n.css(a[0],"borderLeftWidth",!0)),{top:b.top-c.top-n.css(d,"marginTop",!0),left:b.left-c.left-n.css(d,"marginLeft",!0)}}},offsetParent:function(){return this.map(function(){var a=this.offsetParent;while(a&&!n.nodeName(a,"html")&&"static"===n.css(a,"position"))a=a.offsetParent;return a||Qa})}}),n.each({scrollLeft:"pageXOffset",scrollTop:"pageYOffset"},function(a,b){var c=/Y/.test(b);n.fn[a]=function(d){return Y(this,function(a,d,e){var f=mc(a);return void 0===e?f?b in f?f[b]:f.document.documentElement[d]:a[d]:void(f?f.scrollTo(c?n(f).scrollLeft():e,c?e:n(f).scrollTop()):a[d]=e)},a,d,arguments.length,null)}}),n.each(["top","left"],function(a,b){n.cssHooks[b]=Ua(l.pixelPosition,function(a,c){return c?(c=Sa(a,b),Oa.test(c)?n(a).position()[b]+"px":c):void 0})}),n.each({Height:"height",Width:"width"},function(a,b){n.each({
padding:"inner"+a,content:b,"":"outer"+a},function(c,d){n.fn[d]=function(d,e){var f=arguments.length&&(c||"boolean"!=typeof d),g=c||(d===!0||e===!0?"margin":"border");return Y(this,function(b,c,d){var e;return n.isWindow(b)?b.document.documentElement["client"+a]:9===b.nodeType?(e=b.documentElement,Math.max(b.body["scroll"+a],e["scroll"+a],b.body["offset"+a],e["offset"+a],e["client"+a])):void 0===d?n.css(b,c,g):n.style(b,c,d,g)},b,f?d:void 0,f,null)}})}),n.fn.extend({bind:function(a,b,c){return this.on(a,null,b,c)},unbind:function(a,b){return this.off(a,null,b)},delegate:function(a,b,c,d){return this.on(b,a,c,d)},undelegate:function(a,b,c){return 1===arguments.length?this.off(a,"**"):this.off(b,a||"**",c)}}),n.fn.size=function(){return this.length},n.fn.andSelf=n.fn.addBack,"function"==typeof define&&define.amd&&define("jquery",[],function(){return n});var nc=a.jQuery,oc=a.$;return n.noConflict=function(b){return a.$===n&&(a.$=oc),b&&a.jQuery===n&&(a.jQuery=nc),n},b||(a.jQuery=a.$=n),n});
```
