## Supplementary material for "Characterizing and inferring quantitative cell cycle phase in single-cell RNA-seq data analysis": Source code of our analysis, including scripts and data necessary to reproduce the work.: LICENSE.txt

Copyright 2005, 2014 jQuery Foundation and other contributors,
https://jquery.org/

This software consists of voluntary contributions made by many
individuals. For exact contribution history, see the revision history
available at https://github.com/jquery/jquery

The following license applies to all parts of this software except as
documented below:

====

Permission is hereby granted, free of charge, to any person obtaining
a copy of this software and associated documentation files (the
"Software"), to deal in the Software without restriction, including
without limitation the rights to use, copy, modify, merge, publish,
distribute, sublicense, and/or sell copies of the Software, and to
permit persons to whom the Software is furnished to do so, subject to
the following conditions:

The above copyright notice and this permission notice shall be
included in all copies or substantial portions of the Software.

THE SOFTWARE IS PROVIDED "AS IS", WITHOUT WARRANTY OF ANY KIND,
EXPRESS OR IMPLIED, INCLUDING BUT NOT LIMITED TO THE WARRANTIES OF
MERCHANTABILITY, FITNESS FOR A PARTICULAR PURPOSE AND
NONINFRINGEMENT. IN NO EVENT SHALL THE AUTHORS OR COPYRIGHT HOLDERS BE
LIABLE FOR ANY CLAIM, DAMAGES OR OTHER LIABILITY, WHETHER IN AN ACTION
OF CONTRACT, TORT OR OTHERWISE, ARISING FROM, OUT OF OR IN CONNECTION
WITH THE SOFTWARE OR THE USE OR OTHER DEALINGS IN THE SOFTWARE.

====

All files located in the node_modules and external directories are
externally maintained libraries used by this software which have their
own licenses; we recommend you read them, as their terms may differ from
the terms above.
