## Supplementary material for "Characterizing and inferring quantitative cell cycle phase in single-cell RNA-seq data analysis": Source code of our analysis, including scripts and data necessary to reproduce the work.: index.html

jQuery UI Example Page


### Welcome to jQuery UI!

This page demonstrates the widgets and theme you selected in Download Builder. Please make sure you are using them with a compatible jQuery version.

### YOUR COMPONENTS:

#### Accordion

##### First

Lorem ipsum dolor sit amet. Lorem ipsum dolor sit amet. Lorem ipsum dolor sit amet.

##### Second

Phasellus mattis tincidunt nibh.

##### Third

Nam dui erat, auctor a, dignissim quis.

#### Autocomplete

#### Button

A button element


Choice 1
Choice 2
Choice 3


#### Tabs

- First
- Second
- Third

Lorem ipsum dolor sit amet, consectetur adipisicing elit, sed do eiusmod tempor incididunt ut labore et dolore magna aliqua. Ut enim ad minim veniam, quis nostrud exercitation ullamco laboris nisi ut aliquip ex ea commodo consequat.

Phasellus mattis tincidunt nibh. Cras orci urna, blandit id, pretium vel, aliquet ornare, felis. Maecenas scelerisque sem non nisl. Fusce sed lorem in enim dictum bibendum.

Nam dui erat, auctor a, dignissim quis, sollicitudin eu, felis. Pellentesque nisi urna, interdum eget, sagittis et, consequat vestibulum, lacus. Mauris porttitor ullamcorper augue.

#### Dialog

Open Dialog

#### Overlay and Shadow Classes *(not currently used in UI widgets)*

Lorem ipsum dolor sit amet, Nulla nec tortor. Donec id elit quis purus consectetur consequat.

Nam congue semper tellus. Sed erat dolor, dapibus sit amet, venenatis ornare, ultrices ut, nisi. Aliquam ante. Suspendisse scelerisque dui nec velit. Duis augue augue, gravida euismod, vulputate ac, facilisis id, sem. Morbi in orci.

Nulla purus lacus, pulvinar vel, malesuada ac, mattis nec, quam. Nam molestie scelerisque quam. Nullam feugiat cursus lacus.orem ipsum dolor sit amet, consectetur adipiscing elit. Donec libero risus, commodo vitae, pharetra mollis, posuere eu, pede. Nulla nec tortor. Donec id elit quis purus consectetur consequat.

Nam congue semper tellus. Sed erat dolor, dapibus sit amet, venenatis ornare, ultrices ut, nisi. Aliquam ante. Suspendisse scelerisque dui nec velit. Duis augue augue, gravida euismod, vulputate ac, facilisis id, sem. Morbi in orci. Nulla purus lacus, pulvinar vel, malesuada ac, mattis nec, quam. Nam molestie scelerisque quam.

Nullam feugiat cursus lacus.orem ipsum dolor sit amet, consectetur adipiscing elit. Donec libero risus, commodo vitae, pharetra mollis, posuere eu, pede. Nulla nec tortor. Donec id elit quis purus consectetur consequat. Nam congue semper tellus. Sed erat dolor, dapibus sit amet, venenatis ornare, ultrices ut, nisi. Aliquam ante.

Suspendisse scelerisque dui nec velit. Duis augue augue, gravida euismod, vulputate ac, facilisis id, sem. Morbi in orci. Nulla purus lacus, pulvinar vel, malesuada ac, mattis nec, quam. Nam molestie scelerisque quam. Nullam feugiat cursus lacus.orem ipsum dolor sit amet, consectetur adipiscing elit. Donec libero risus, commodo vitae, pharetra mollis, posuere eu, pede. Nulla nec tortor. Donec id elit quis purus consectetur consequat. Nam congue semper tellus. Sed erat dolor, dapibus sit amet, venenatis ornare, ultrices ut, nisi.

Lorem ipsum dolor sit amet, consectetur adipisicing elit, sed do eiusmod tempor incididunt ut labore et dolore magna aliqua. Ut enim ad minim veniam, quis nostrud exercitation ullamco laboris nisi ut aliquip ex ea commodo consequat.

Lorem ipsum dolor sit amet, consectetur adipisicing elit, sed do eiusmod tempor incididunt ut labore et dolore magna aliqua. Ut enim ad minim veniam, quis nostrud exercitation ullamco laboris nisi ut aliquip ex ea commodo consequat.

#### Framework Icons (content color preview)


#### Slider

#### Progressbar

#### Selectmenu

Slower
Slow
Medium
Fast
Faster


#### Spinner


#### Menu

- Item 1
- Item 2
- Item 3
  - Item 3-1
  - Item 3-2
  - Item 3-3
  - Item 3-4
  - Item 3-5
- Item 4
- Item 5

#### Tooltip

Tooltips can be attached to any element. When you hover
the element with your mouse, the title attribute is displayed in a little box next to the element, just like a native tooltip.

#### Highlight / Error

**Hey!** Sample ui-state-highlight style.

  

**Alert:** Sample ui-state-error style.
