## Supplementary material for "Characterizing and inferring quantitative cell cycle phase in single-cell RNA-seq data analysis": Source code of our analysis, including scripts and data necessary to reproduce the work.: jquery-ui.js

```
/*! jQuery UI - v1.11.4 - 2016-01-05
* http://jqueryui.com
* Includes: core.js, widget.js, mouse.js, position.js, draggable.js, droppable.js, resizable.js, selectable.js, sortable.js, accordion.js, autocomplete.js, button.js, dialog.js, menu.js, progressbar.js, selectmenu.js, slider.js, spinner.js, tabs.js, tooltip.js, effect.js, effect-blind.js, effect-bounce.js, effect-clip.js, effect-drop.js, effect-explode.js, effect-fade.js, effect-fold.js, effect-highlight.js, effect-puff.js, effect-pulsate.js, effect-scale.js, effect-shake.js, effect-size.js, effect-slide.js, effect-transfer.js
* Copyright jQuery Foundation and other contributors; Licensed MIT */

(function( factory ) {
	if ( typeof define === "function" && define.amd ) {

		// AMD. Register as an anonymous module.
		define([ "jquery" ], factory );
	} else {

		// Browser globals
		factory( jQuery );
	}
}(function( $ ) {
/*!
 * jQuery UI Core 1.11.4
 * http://jqueryui.com
 *
 * Copyright jQuery Foundation and other contributors
 * Released under the MIT license.
 * http://jquery.org/license
 *
 * http://api.jqueryui.com/category/ui-core/
 */


// $.ui might exist from components with no dependencies, e.g., $.ui.position
$.ui = $.ui || {};

$.extend( $.ui, {
	version: "1.11.4",

	keyCode: {
		BACKSPACE: 8,
		COMMA: 188,
		DELETE: 46,
		DOWN: 40,
		END: 35,
		ENTER: 13,
		ESCAPE: 27,
		HOME: 36,
		LEFT: 37,
		PAGE_DOWN: 34,
		PAGE_UP: 33,
		PERIOD: 190,
		RIGHT: 39,
		SPACE: 32,
		TAB: 9,
		UP: 38
	}
});

// plugins
$.fn.extend({
	scrollParent: function( includeHidden ) {
		var position = this.css( "position" ),
			excludeStaticParent = position === "absolute",
			overflowRegex = includeHidden ? /(auto|scroll|hidden)/ : /(auto|scroll)/,
			scrollParent = this.parents().filter( function() {
				var parent = $( this );
				if ( excludeStaticParent && parent.css( "position" ) === "static" ) {
					return false;
				}
				return overflowRegex.test( parent.css( "overflow" ) + parent.css( "overflow-y" ) + parent.css( "overflow-x" ) );
			}).eq( 0 );

		return position === "fixed" || !scrollParent.length ? $( this[ 0 ].ownerDocument || document ) : scrollParent;
	},

	uniqueId: (function() {
		var uuid = 0;

		return function() {
			return this.each(function() {
				if ( !this.id ) {
					this.id = "ui-id-" + ( ++uuid );
				}
			});
		};
	})(),

	removeUniqueId: function() {
		return this.each(function() {
			if ( /^ui-id-\d+$/.test( this.id ) ) {
				$( this ).removeAttr( "id" );
			}
		});
	}
});

// selectors
function focusable( element, isTabIndexNotNaN ) {
	var map, mapName, img,
		nodeName = element.nodeName.toLowerCase();
	if ( "area" === nodeName ) {
		map = element.parentNode;
		mapName = map.name;
		if ( !element.href || !mapName || map.nodeName.toLowerCase() !== "map" ) {
			return false;
		}
		img = $( "img[usemap='#" + mapName + "']" )[ 0 ];
		return !!img && visible( img );
	}
	return ( /^(input|select|textarea|button|object)$/.test( nodeName ) ?
		!element.disabled :
		"a" === nodeName ?
			element.href || isTabIndexNotNaN :
			isTabIndexNotNaN) &&
		// the element and all of its ancestors must be visible
		visible( element );
}

function visible( element ) {
	return $.expr.filters.visible( element ) &&
		!$( element ).parents().addBack().filter(function() {
			return $.css( this, "visibility" ) === "hidden";
		}).length;
}

$.extend( $.expr[ ":" ], {
	data: $.expr.createPseudo ?
		$.expr.createPseudo(function( dataName ) {
			return function( elem ) {
				return !!$.data( elem, dataName );
			};
		}) :
		// support: jQuery <1.8
		function( elem, i, match ) {
			return !!$.data( elem, match[ 3 ] );
		},

	focusable: function( element ) {
		return focusable( element, !isNaN( $.attr( element, "tabindex" ) ) );
	},

	tabbable: function( element ) {
		var tabIndex = $.attr( element, "tabindex" ),
			isTabIndexNaN = isNaN( tabIndex );
		return ( isTabIndexNaN || tabIndex >= 0 ) && focusable( element, !isTabIndexNaN );
	}
});

// support: jQuery <1.8
if ( !$( "<a>" ).outerWidth( 1 ).jquery ) {
	$.each( [ "Width", "Height" ], function( i, name ) {
		var side = name === "Width" ? [ "Left", "Right" ] : [ "Top", "Bottom" ],
			type = name.toLowerCase(),
			orig = {
				innerWidth: $.fn.innerWidth,
				innerHeight: $.fn.innerHeight,
				outerWidth: $.fn.outerWidth,
				outerHeight: $.fn.outerHeight
			};

		function reduce( elem, size, border, margin ) {
			$.each( side, function() {
				size -= parseFloat( $.css( elem, "padding" + this ) ) || 0;
				if ( border ) {
					size -= parseFloat( $.css( elem, "border" + this + "Width" ) ) || 0;
				}
				if ( margin ) {
					size -= parseFloat( $.css( elem, "margin" + this ) ) || 0;
				}
			});
			return size;
		}

		$.fn[ "inner" + name ] = function( size ) {
			if ( size === undefined ) {
				return orig[ "inner" + name ].call( this );
			}

			return this.each(function() {
				$( this ).css( type, reduce( this, size ) + "px" );
			});
		};

		$.fn[ "outer" + name] = function( size, margin ) {
			if ( typeof size !== "number" ) {
				return orig[ "outer" + name ].call( this, size );
			}

			return this.each(function() {
				$( this).css( type, reduce( this, size, true, margin ) + "px" );
			});
		};
	});
}

// support: jQuery <1.8
if ( !$.fn.addBack ) {
	$.fn.addBack = function( selector ) {
		return this.add( selector == null ?
			this.prevObject : this.prevObject.filter( selector )
		);
	};
}

// support: jQuery 1.6.1, 1.6.2 (http://bugs.jquery.com/ticket/9413)
if ( $( "<a>" ).data( "a-b", "a" ).removeData( "a-b" ).data( "a-b" ) ) {
	$.fn.removeData = (function( removeData ) {
		return function( key ) {
			if ( arguments.length ) {
				return removeData.call( this, $.camelCase( key ) );
			} else {
				return removeData.call( this );
			}
		};
	})( $.fn.removeData );
}

// deprecated
$.ui.ie = !!/msie [\w.]+/.exec( navigator.userAgent.toLowerCase() );

$.fn.extend({
	focus: (function( orig ) {
		return function( delay, fn ) {
			return typeof delay === "number" ?
				this.each(function() {
					var elem = this;
					setTimeout(function() {
						$( elem ).focus();
						if ( fn ) {
							fn.call( elem );
						}
					}, delay );
				}) :
				orig.apply( this, arguments );
		};
	})( $.fn.focus ),

	disableSelection: (function() {
		var eventType = "onselectstart" in document.createElement( "div" ) ?
			"selectstart" :
			"mousedown";

		return function() {
			return this.bind( eventType + ".ui-disableSelection", function( event ) {
				event.preventDefault();
			});
		};
	})(),

	enableSelection: function() {
		return this.unbind( ".ui-disableSelection" );
	},

	zIndex: function( zIndex ) {
		if ( zIndex !== undefined ) {
			return this.css( "zIndex", zIndex );
		}

		if ( this.length ) {
			var elem = $( this[ 0 ] ), position, value;
			while ( elem.length && elem[ 0 ] !== document ) {
				// Ignore z-index if position is set to a value where z-index is ignored by the browser
				// This makes behavior of this function consistent across browsers
				// WebKit always returns auto if the element is positioned
				position = elem.css( "position" );
				if ( position === "absolute" || position === "relative" || position === "fixed" ) {
					// IE returns 0 when zIndex is not specified
					// other browsers return a string
					// we ignore the case of nested elements with an explicit value of 0
					// <div style="z-index: -10;"><div style="z-index: 0;"></div></div>
					value = parseInt( elem.css( "zIndex" ), 10 );
					if ( !isNaN( value ) && value !== 0 ) {
						return value;
					}
				}
				elem = elem.parent();
			}
		}

		return 0;
	}
});

// $.ui.plugin is deprecated. Use $.widget() extensions instead.
$.ui.plugin = {
	add: function( module, option, set ) {
		var i,
			proto = $.ui[ module ].prototype;
		for ( i in set ) {
			proto.plugins[ i ] = proto.plugins[ i ] || [];
			proto.plugins[ i ].push( [ option, set[ i ] ] );
		}
	},
	call: function( instance, name, args, allowDisconnected ) {
		var i,
			set = instance.plugins[ name ];

		if ( !set ) {
			return;
		}

		if ( !allowDisconnected && ( !instance.element[ 0 ].parentNode || instance.element[ 0 ].parentNode.nodeType === 11 ) ) {
			return;
		}

		for ( i = 0; i < set.length; i++ ) {
			if ( instance.options[ set[ i ][ 0 ] ] ) {
				set[ i ][ 1 ].apply( instance.element, args );
			}
		}
	}
};


/*!
 * jQuery UI Widget 1.11.4
 * http://jqueryui.com
 *
 * Copyright jQuery Foundation and other contributors
 * Released under the MIT license.
 * http://jquery.org/license
 *
 * http://api.jqueryui.com/jQuery.widget/
 */


var widget_uuid = 0,
	widget_slice = Array.prototype.slice;

$.cleanData = (function( orig ) {
	return function( elems ) {
		var events, elem, i;
		for ( i = 0; (elem = elems[i]) != null; i++ ) {
			try {

				// Only trigger remove when necessary to save time
				events = $._data( elem, "events" );
				if ( events && events.remove ) {
					$( elem ).triggerHandler( "remove" );
				}

			// http://bugs.jquery.com/ticket/8235
			} catch ( e ) {}
		}
		orig( elems );
	};
})( $.cleanData );

$.widget = function( name, base, prototype ) {
	var fullName, existingConstructor, constructor, basePrototype,
		// proxiedPrototype allows the provided prototype to remain unmodified
		// so that it can be used as a mixin for multiple widgets (#8876)
		proxiedPrototype = {},
		namespace = name.split( "." )[ 0 ];

	name = name.split( "." )[ 1 ];
	fullName = namespace + "-" + name;

	if ( !prototype ) {
		prototype = base;
		base = $.Widget;
	}

	// create selector for plugin
	$.expr[ ":" ][ fullName.toLowerCase() ] = function( elem ) {
		return !!$.data( elem, fullName );
	};

	$[ namespace ] = $[ namespace ] || {};
	existingConstructor = $[ namespace ][ name ];
	constructor = $[ namespace ][ name ] = function( options, element ) {
		// allow instantiation without "new" keyword
		if ( !this._createWidget ) {
			return new constructor( options, element );
		}

		// allow instantiation without initializing for simple inheritance
		// must use "new" keyword (the code above always passes args)
		if ( arguments.length ) {
			this._createWidget( options, element );
		}
	};
	// extend with the existing constructor to carry over any static properties
	$.extend( constructor, existingConstructor, {
		version: prototype.version,
		// copy the object used to create the prototype in case we need to
		// redefine the widget later
		_proto: $.extend( {}, prototype ),
		// track widgets that inherit from this widget in case this widget is
		// redefined after a widget inherits from it
		_childConstructors: []
	});

	basePrototype = new base();
	// we need to make the options hash a property directly on the new instance
	// otherwise we'll modify the options hash on the prototype that we're
	// inheriting from
	basePrototype.options = $.widget.extend( {}, basePrototype.options );
	$.each( prototype, function( prop, value ) {
		if ( !$.isFunction( value ) ) {
			proxiedPrototype[ prop ] = value;
			return;
		}
		proxiedPrototype[ prop ] = (function() {
			var _super = function() {
					return base.prototype[ prop ].apply( this, arguments );
				},
				_superApply = function( args ) {
					return base.prototype[ prop ].apply( this, args );
				};
			return function() {
				var __super = this._super,
					__superApply = this._superApply,
					returnValue;

				this._super = _super;
				this._superApply = _superApply;

				returnValue = value.apply( this, arguments );

				this._super = __super;
				this._superApply = __superApply;

				return returnValue;
			};
		})();
	});
	constructor.prototype = $.widget.extend( basePrototype, {
		// TODO: remove support for widgetEventPrefix
		// always use the name + a colon as the prefix, e.g., draggable:start
		// don't prefix for widgets that aren't DOM-based
		widgetEventPrefix: existingConstructor ? (basePrototype.widgetEventPrefix || name) : name
	}, proxiedPrototype, {
		constructor: constructor,
		namespace: namespace,
		widgetName: name,
		widgetFullName: fullName
	});

	// If this widget is being redefined then we need to find all widgets that
	// are inheriting from it and redefine all of them so that they inherit from
	// the new version of this widget. We're essentially trying to replace one
	// level in the prototype chain.
	if ( existingConstructor ) {
		$.each( existingConstructor._childConstructors, function( i, child ) {
			var childPrototype = child.prototype;

			// redefine the child widget using the same prototype that was
			// originally used, but inherit from the new version of the base
			$.widget( childPrototype.namespace + "." + childPrototype.widgetName, constructor, child._proto );
		});
		// remove the list of existing child constructors from the old constructor
		// so the old child constructors can be garbage collected
		delete existingConstructor._childConstructors;
	} else {
		base._childConstructors.push( constructor );
	}

	$.widget.bridge( name, constructor );

	return constructor;
};

$.widget.extend = function( target ) {
	var input = widget_slice.call( arguments, 1 ),
		inputIndex = 0,
		inputLength = input.length,
		key,
		value;
	for ( ; inputIndex < inputLength; inputIndex++ ) {
		for ( key in input[ inputIndex ] ) {
			value = input[ inputIndex ][ key ];
			if ( input[ inputIndex ].hasOwnProperty( key ) && value !== undefined ) {
				// Clone objects
				if ( $.isPlainObject( value ) ) {
					target[ key ] = $.isPlainObject( target[ key ] ) ?
						$.widget.extend( {}, target[ key ], value ) :
						// Don't extend strings, arrays, etc. with objects
						$.widget.extend( {}, value );
				// Copy everything else by reference
				} else {
					target[ key ] = value;
				}
			}
		}
	}
	return target;
};

$.widget.bridge = function( name, object ) {
	var fullName = object.prototype.widgetFullName || name;
	$.fn[ name ] = function( options ) {
		var isMethodCall = typeof options === "string",
			args = widget_slice.call( arguments, 1 ),
			returnValue = this;

		if ( isMethodCall ) {
			this.each(function() {
				var methodValue,
					instance = $.data( this, fullName );
				if ( options === "instance" ) {
					returnValue = instance;
					return false;
				}
				if ( !instance ) {
					return $.error( "cannot call methods on " + name + " prior to initialization; " +
						"attempted to call method '" + options + "'" );
				}
				if ( !$.isFunction( instance[options] ) || options.charAt( 0 ) === "_" ) {
					return $.error( "no such method '" + options + "' for " + name + " widget instance" );
				}
				methodValue = instance[ options ].apply( instance, args );
				if ( methodValue !== instance && methodValue !== undefined ) {
					returnValue = methodValue && methodValue.jquery ?
						returnValue.pushStack( methodValue.get() ) :
						methodValue;
					return false;
				}
			});
		} else {

			// Allow multiple hashes to be passed on init
			if ( args.length ) {
				options = $.widget.extend.apply( null, [ options ].concat(args) );
			}

			this.each(function() {
				var instance = $.data( this, fullName );
				if ( instance ) {
					instance.option( options || {} );
					if ( instance._init ) {
						instance._init();
					}
				} else {
					$.data( this, fullName, new object( options, this ) );
				}
			});
		}

		return returnValue;
	};
};

$.Widget = function( /* options, element */ ) {};
$.Widget._childConstructors = [];

$.Widget.prototype = {
	widgetName: "widget",
	widgetEventPrefix: "",
	defaultElement: "<div>",
	options: {
		disabled: false,

		// callbacks
		create: null
	},
	_createWidget: function( options, element ) {
		element = $( element || this.defaultElement || this )[ 0 ];
		this.element = $( element );
		this.uuid = widget_uuid++;
		this.eventNamespace = "." + this.widgetName + this.uuid;

		this.bindings = $();
		this.hoverable = $();
		this.focusable = $();

		if ( element !== this ) {
			$.data( element, this.widgetFullName, this );
			this._on( true, this.element, {
				remove: function( event ) {
					if ( event.target === element ) {
						this.destroy();
					}
				}
			});
			this.document = $( element.style ?
				// element within the document
				element.ownerDocument :
				// element is window or document
				element.document || element );
			this.window = $( this.document[0].defaultView || this.document[0].parentWindow );
		}

		this.options = $.widget.extend( {},
			this.options,
			this._getCreateOptions(),
			options );

		this._create();
		this._trigger( "create", null, this._getCreateEventData() );
		this._init();
	},
	_getCreateOptions: $.noop,
	_getCreateEventData: $.noop,
	_create: $.noop,
	_init: $.noop,

	destroy: function() {
		this._destroy();
		// we can probably remove the unbind calls in 2.0
		// all event bindings should go through this._on()
		this.element
			.unbind( this.eventNamespace )
			.removeData( this.widgetFullName )
			// support: jquery <1.6.3
			// http://bugs.jquery.com/ticket/9413
			.removeData( $.camelCase( this.widgetFullName ) );
		this.widget()
			.unbind( this.eventNamespace )
			.removeAttr( "aria-disabled" )
			.removeClass(
				this.widgetFullName + "-disabled " +
				"ui-state-disabled" );

		// clean up events and states
		this.bindings.unbind( this.eventNamespace );
		this.hoverable.removeClass( "ui-state-hover" );
		this.focusable.removeClass( "ui-state-focus" );
	},
	_destroy: $.noop,

	widget: function() {
		return this.element;
	},

	option: function( key, value ) {
		var options = key,
			parts,
			curOption,
			i;

		if ( arguments.length === 0 ) {
			// don't return a reference to the internal hash
			return $.widget.extend( {}, this.options );
		}

		if ( typeof key === "string" ) {
			// handle nested keys, e.g., "foo.bar" => { foo: { bar: ___ } }
			options = {};
			parts = key.split( "." );
			key = parts.shift();
			if ( parts.length ) {
				curOption = options[ key ] = $.widget.extend( {}, this.options[ key ] );
				for ( i = 0; i < parts.length - 1; i++ ) {
					curOption[ parts[ i ] ] = curOption[ parts[ i ] ] || {};
					curOption = curOption[ parts[ i ] ];
				}
				key = parts.pop();
				if ( arguments.length === 1 ) {
					return curOption[ key ] === undefined ? null : curOption[ key ];
				}
				curOption[ key ] = value;
			} else {
				if ( arguments.length === 1 ) {
					return this.options[ key ] === undefined ? null : this.options[ key ];
				}
				options[ key ] = value;
			}
		}

		this._setOptions( options );

		return this;
	},
	_setOptions: function( options ) {
		var key;

		for ( key in options ) {
			this._setOption( key, options[ key ] );
		}

		return this;
	},
	_setOption: function( key, value ) {
		this.options[ key ] = value;

		if ( key === "disabled" ) {
			this.widget()
				.toggleClass( this.widgetFullName + "-disabled", !!value );

			// If the widget is becoming disabled, then nothing is interactive
			if ( value ) {
				this.hoverable.removeClass( "ui-state-hover" );
				this.focusable.removeClass( "ui-state-focus" );
			}
		}

		return this;
	},

	enable: function() {
		return this._setOptions({ disabled: false });
	},
	disable: function() {
		return this._setOptions({ disabled: true });
	},

	_on: function( suppressDisabledCheck, element, handlers ) {
		var delegateElement,
			instance = this;

		// no suppressDisabledCheck flag, shuffle arguments
		if ( typeof suppressDisabledCheck !== "boolean" ) {
			handlers = element;
			element = suppressDisabledCheck;
			suppressDisabledCheck = false;
		}

		// no element argument, shuffle and use this.element
		if ( !handlers ) {
			handlers = element;
			element = this.element;
			delegateElement = this.widget();
		} else {
			element = delegateElement = $( element );
			this.bindings = this.bindings.add( element );
		}

		$.each( handlers, function( event, handler ) {
			function handlerProxy() {
				// allow widgets to customize the disabled handling
				// - disabled as an array instead of boolean
				// - disabled class as method for disabling individual parts
				if ( !suppressDisabledCheck &&
						( instance.options.disabled === true ||
							$( this ).hasClass( "ui-state-disabled" ) ) ) {
					return;
				}
				return ( typeof handler === "string" ? instance[ handler ] : handler )
					.apply( instance, arguments );
			}

			// copy the guid so direct unbinding works
			if ( typeof handler !== "string" ) {
				handlerProxy.guid = handler.guid =
					handler.guid || handlerProxy.guid || $.guid++;
			}

			var match = event.match( /^([\w:-]*)\s*(.*)$/ ),
				eventName = match[1] + instance.eventNamespace,
				selector = match[2];
			if ( selector ) {
				delegateElement.delegate( selector, eventName, handlerProxy );
			} else {
				element.bind( eventName, handlerProxy );
			}
		});
	},

	_off: function( element, eventName ) {
		eventName = (eventName || "").split( " " ).join( this.eventNamespace + " " ) +
			this.eventNamespace;
		element.unbind( eventName ).undelegate( eventName );

		// Clear the stack to avoid memory leaks (#10056)
		this.bindings = $( this.bindings.not( element ).get() );
		this.focusable = $( this.focusable.not( element ).get() );
		this.hoverable = $( this.hoverable.not( element ).get() );
	},

	_delay: function( handler, delay ) {
		function handlerProxy() {
			return ( typeof handler === "string" ? instance[ handler ] : handler )
				.apply( instance, arguments );
		}
		var instance = this;
		return setTimeout( handlerProxy, delay || 0 );
	},

	_hoverable: function( element ) {
		this.hoverable = this.hoverable.add( element );
		this._on( element, {
			mouseenter: function( event ) {
				$( event.currentTarget ).addClass( "ui-state-hover" );
			},
			mouseleave: function( event ) {
				$( event.currentTarget ).removeClass( "ui-state-hover" );
			}
		});
	},

	_focusable: function( element ) {
		this.focusable = this.focusable.add( element );
		this._on( element, {
			focusin: function( event ) {
				$( event.currentTarget ).addClass( "ui-state-focus" );
			},
			focusout: function( event ) {
				$( event.currentTarget ).removeClass( "ui-state-focus" );
			}
		});
	},

	_trigger: function( type, event, data ) {
		var prop, orig,
			callback = this.options[ type ];

		data = data || {};
		event = $.Event( event );
		event.type = ( type === this.widgetEventPrefix ?
			type :
			this.widgetEventPrefix + type ).toLowerCase();
		// the original event may come from any element
		// so we need to reset the target on the new event
		event.target = this.element[ 0 ];

		// copy original event properties over to the new event
		orig = event.originalEvent;
		if ( orig ) {
			for ( prop in orig ) {
				if ( !( prop in event ) ) {
					event[ prop ] = orig[ prop ];
				}
			}
		}

		this.element.trigger( event, data );
		return !( $.isFunction( callback ) &&
			callback.apply( this.element[0], [ event ].concat( data ) ) === false ||
			event.isDefaultPrevented() );
	}
};

$.each( { show: "fadeIn", hide: "fadeOut" }, function( method, defaultEffect ) {
	$.Widget.prototype[ "_" + method ] = function( element, options, callback ) {
		if ( typeof options === "string" ) {
			options = { effect: options };
		}
		var hasOptions,
			effectName = !options ?
				method :
				options === true || typeof options === "number" ?
					defaultEffect :
					options.effect || defaultEffect;
		options = options || {};
		if ( typeof options === "number" ) {
			options = { duration: options };
		}
		hasOptions = !$.isEmptyObject( options );
		options.complete = callback;
		if ( options.delay ) {
			element.delay( options.delay );
		}
		if ( hasOptions && $.effects && $.effects.effect[ effectName ] ) {
			element[ method ]( options );
		} else if ( effectName !== method && element[ effectName ] ) {
			element[ effectName ]( options.duration, options.easing, callback );
		} else {
			element.queue(function( next ) {
				$( this )[ method ]();
				if ( callback ) {
					callback.call( element[ 0 ] );
				}
				next();
			});
		}
	};
});

var widget = $.widget;


/*!
 * jQuery UI Mouse 1.11.4
 * http://jqueryui.com
 *
 * Copyright jQuery Foundation and other contributors
 * Released under the MIT license.
 * http://jquery.org/license
 *
 * http://api.jqueryui.com/mouse/
 */


var mouseHandled = false;
$( document ).mouseup( function() {
	mouseHandled = false;
});

var mouse = $.widget("ui.mouse", {
	version: "1.11.4",
	options: {
		cancel: "input,textarea,button,select,option",
		distance: 1,
		delay: 0
	},
	_mouseInit: function() {
		var that = this;

		this.element
			.bind("mousedown." + this.widgetName, function(event) {
				return that._mouseDown(event);
			})
			.bind("click." + this.widgetName, function(event) {
				if (true === $.data(event.target, that.widgetName + ".preventClickEvent")) {
					$.removeData(event.target, that.widgetName + ".preventClickEvent");
					event.stopImmediatePropagation();
					return false;
				}
			});

		this.started = false;
	},

	// TODO: make sure destroying one instance of mouse doesn't mess with
	// other instances of mouse
	_mouseDestroy: function() {
		this.element.unbind("." + this.widgetName);
		if ( this._mouseMoveDelegate ) {
			this.document
				.unbind("mousemove." + this.widgetName, this._mouseMoveDelegate)
				.unbind("mouseup." + this.widgetName, this._mouseUpDelegate);
		}
	},

	_mouseDown: function(event) {
		// don't let more than one widget handle mouseStart
		if ( mouseHandled ) {
			return;
		}

		this._mouseMoved = false;

		// we may have missed mouseup (out of window)
		(this._mouseStarted && this._mouseUp(event));

		this._mouseDownEvent = event;

		var that = this,
			btnIsLeft = (event.which === 1),
			// event.target.nodeName works around a bug in IE 8 with
			// disabled inputs (#7620)
			elIsCancel = (typeof this.options.cancel === "string" && event.target.nodeName ? $(event.target).closest(this.options.cancel).length : false);
		if (!btnIsLeft || elIsCancel || !this._mouseCapture(event)) {
			return true;
		}

		this.mouseDelayMet = !this.options.delay;
		if (!this.mouseDelayMet) {
			this._mouseDelayTimer = setTimeout(function() {
				that.mouseDelayMet = true;
			}, this.options.delay);
		}

		if (this._mouseDistanceMet(event) && this._mouseDelayMet(event)) {
			this._mouseStarted = (this._mouseStart(event) !== false);
			if (!this._mouseStarted) {
				event.preventDefault();
				return true;
			}
		}

		// Click event may never have fired (Gecko & Opera)
		if (true === $.data(event.target, this.widgetName + ".preventClickEvent")) {
			$.removeData(event.target, this.widgetName + ".preventClickEvent");
		}

		// these delegates are required to keep context
		this._mouseMoveDelegate = function(event) {
			return that._mouseMove(event);
		};
		this._mouseUpDelegate = function(event) {
			return that._mouseUp(event);
		};

		this.document
			.bind( "mousemove." + this.widgetName, this._mouseMoveDelegate )
			.bind( "mouseup." + this.widgetName, this._mouseUpDelegate );

		event.preventDefault();

		mouseHandled = true;
		return true;
	},

	_mouseMove: function(event) {
		// Only check for mouseups outside the document if you've moved inside the document
		// at least once. This prevents the firing of mouseup in the case of IE<9, which will
		// fire a mousemove event if content is placed under the cursor. See #7778
		// Support: IE <9
		if ( this._mouseMoved ) {
			// IE mouseup check - mouseup happened when mouse was out of window
			if ($.ui.ie && ( !document.documentMode || document.documentMode < 9 ) && !event.button) {
				return this._mouseUp(event);

			// Iframe mouseup check - mouseup occurred in another document
			} else if ( !event.which ) {
				return this._mouseUp( event );
			}
		}

		if ( event.which || event.button ) {
			this._mouseMoved = true;
		}

		if (this._mouseStarted) {
			this._mouseDrag(event);
			return event.preventDefault();
		}

		if (this._mouseDistanceMet(event) && this._mouseDelayMet(event)) {
			this._mouseStarted =
				(this._mouseStart(this._mouseDownEvent, event) !== false);
			(this._mouseStarted ? this._mouseDrag(event) : this._mouseUp(event));
		}

		return !this._mouseStarted;
	},

	_mouseUp: function(event) {
		this.document
			.unbind( "mousemove." + this.widgetName, this._mouseMoveDelegate )
			.unbind( "mouseup." + this.widgetName, this._mouseUpDelegate );

		if (this._mouseStarted) {
			this._mouseStarted = false;

			if (event.target === this._mouseDownEvent.target) {
				$.data(event.target, this.widgetName + ".preventClickEvent", true);
			}

			this._mouseStop(event);
		}

		mouseHandled = false;
		return false;
	},

	_mouseDistanceMet: function(event) {
		return (Math.max(
				Math.abs(this._mouseDownEvent.pageX - event.pageX),
				Math.abs(this._mouseDownEvent.pageY - event.pageY)
			) >= this.options.distance
		);
	},

	_mouseDelayMet: function(/* event */) {
		return this.mouseDelayMet;
	},

	// These are placeholder methods, to be overriden by extending plugin
	_mouseStart: function(/* event */) {},
	_mouseDrag: function(/* event */) {},
	_mouseStop: function(/* event */) {},
	_mouseCapture: function(/* event */) { return true; }
});


/*!
 * jQuery UI Position 1.11.4
 * http://jqueryui.com
 *
 * Copyright jQuery Foundation and other contributors
 * Released under the MIT license.
 * http://jquery.org/license
 *
 * http://api.jqueryui.com/position/
 */

(function() {

$.ui = $.ui || {};

var cachedScrollbarWidth, supportsOffsetFractions,
	max = Math.max,
	abs = Math.abs,
	round = Math.round,
	rhorizontal = /left|center|right/,
	rvertical = /top|center|bottom/,
	roffset = /[\+\-]\d+(\.[\d]+)?%?/,
	rposition = /^\w+/,
	rpercent = /%$/,
	_position = $.fn.position;

function getOffsets( offsets, width, height ) {
	return [
		parseFloat( offsets[ 0 ] ) * ( rpercent.test( offsets[ 0 ] ) ? width / 100 : 1 ),
		parseFloat( offsets[ 1 ] ) * ( rpercent.test( offsets[ 1 ] ) ? height / 100 : 1 )
	];
}

function parseCss( element, property ) {
	return parseInt( $.css( element, property ), 10 ) || 0;
}

function getDimensions( elem ) {
	var raw = elem[0];
	if ( raw.nodeType === 9 ) {
		return {
			width: elem.width(),
			height: elem.height(),
			offset: { top: 0, left: 0 }
		};
	}
	if ( $.isWindow( raw ) ) {
		return {
			width: elem.width(),
			height: elem.height(),
			offset: { top: elem.scrollTop(), left: elem.scrollLeft() }
		};
	}
	if ( raw.preventDefault ) {
		return {
			width: 0,
			height: 0,
			offset: { top: raw.pageY, left: raw.pageX }
		};
	}
	return {
		width: elem.outerWidth(),
		height: elem.outerHeight(),
		offset: elem.offset()
	};
}

$.position = {
	scrollbarWidth: function() {
		if ( cachedScrollbarWidth !== undefined ) {
			return cachedScrollbarWidth;
		}
		var w1, w2,
			div = $( "<div style='display:block;position:absolute;width:50px;height:50px;overflow:hidden;'><div style='height:100px;width:auto;'></div></div>" ),
			innerDiv = div.children()[0];

		$( "body" ).append( div );
		w1 = innerDiv.offsetWidth;
		div.css( "overflow", "scroll" );

		w2 = innerDiv.offsetWidth;

		if ( w1 === w2 ) {
			w2 = div[0].clientWidth;
		}

		div.remove();

		return (cachedScrollbarWidth = w1 - w2);
	},
	getScrollInfo: function( within ) {
		var overflowX = within.isWindow || within.isDocument ? "" :
				within.element.css( "overflow-x" ),
			overflowY = within.isWindow || within.isDocument ? "" :
				within.element.css( "overflow-y" ),
			hasOverflowX = overflowX === "scroll" ||
				( overflowX === "auto" && within.width < within.element[0].scrollWidth ),
			hasOverflowY = overflowY === "scroll" ||
				( overflowY === "auto" && within.height < within.element[0].scrollHeight );
		return {
			width: hasOverflowY ? $.position.scrollbarWidth() : 0,
			height: hasOverflowX ? $.position.scrollbarWidth() : 0
		};
	},
	getWithinInfo: function( element ) {
		var withinElement = $( element || window ),
			isWindow = $.isWindow( withinElement[0] ),
			isDocument = !!withinElement[ 0 ] && withinElement[ 0 ].nodeType === 9;
		return {
			element: withinElement,
			isWindow: isWindow,
			isDocument: isDocument,
			offset: withinElement.offset() || { left: 0, top: 0 },
			scrollLeft: withinElement.scrollLeft(),
			scrollTop: withinElement.scrollTop(),

			// support: jQuery 1.6.x
			// jQuery 1.6 doesn't support .outerWidth/Height() on documents or windows
			width: isWindow || isDocument ? withinElement.width() : withinElement.outerWidth(),
			height: isWindow || isDocument ? withinElement.height() : withinElement.outerHeight()
		};
	}
};

$.fn.position = function( options ) {
	if ( !options || !options.of ) {
		return _position.apply( this, arguments );
	}

	// make a copy, we don't want to modify arguments
	options = $.extend( {}, options );

	var atOffset, targetWidth, targetHeight, targetOffset, basePosition, dimensions,
		target = $( options.of ),
		within = $.position.getWithinInfo( options.within ),
		scrollInfo = $.position.getScrollInfo( within ),
		collision = ( options.collision || "flip" ).split( " " ),
		offsets = {};

	dimensions = getDimensions( target );
	if ( target[0].preventDefault ) {
		// force left top to allow flipping
		options.at = "left top";
	}
	targetWidth = dimensions.width;
	targetHeight = dimensions.height;
	targetOffset = dimensions.offset;
	// clone to reuse original targetOffset later
	basePosition = $.extend( {}, targetOffset );

	// force my and at to have valid horizontal and vertical positions
	// if a value is missing or invalid, it will be converted to center
	$.each( [ "my", "at" ], function() {
		var pos = ( options[ this ] || "" ).split( " " ),
			horizontalOffset,
			verticalOffset;

		if ( pos.length === 1) {
			pos = rhorizontal.test( pos[ 0 ] ) ?
				pos.concat( [ "center" ] ) :
				rvertical.test( pos[ 0 ] ) ?
					[ "center" ].concat( pos ) :
					[ "center", "center" ];
		}
		pos[ 0 ] = rhorizontal.test( pos[ 0 ] ) ? pos[ 0 ] : "center";
		pos[ 1 ] = rvertical.test( pos[ 1 ] ) ? pos[ 1 ] : "center";

		// calculate offsets
		horizontalOffset = roffset.exec( pos[ 0 ] );
		verticalOffset = roffset.exec( pos[ 1 ] );
		offsets[ this ] = [
			horizontalOffset ? horizontalOffset[ 0 ] : 0,
			verticalOffset ? verticalOffset[ 0 ] : 0
		];

		// reduce to just the positions without the offsets
		options[ this ] = [
			rposition.exec( pos[ 0 ] )[ 0 ],
			rposition.exec( pos[ 1 ] )[ 0 ]
		];
	});

	// normalize collision option
	if ( collision.length === 1 ) {
		collision[ 1 ] = collision[ 0 ];
	}

	if ( options.at[ 0 ] === "right" ) {
		basePosition.left += targetWidth;
	} else if ( options.at[ 0 ] === "center" ) {
		basePosition.left += targetWidth / 2;
	}

	if ( options.at[ 1 ] === "bottom" ) {
		basePosition.top += targetHeight;
	} else if ( options.at[ 1 ] === "center" ) {
		basePosition.top += targetHeight / 2;
	}

	atOffset = getOffsets( offsets.at, targetWidth, targetHeight );
	basePosition.left += atOffset[ 0 ];
	basePosition.top += atOffset[ 1 ];

	return this.each(function() {
		var collisionPosition, using,
			elem = $( this ),
			elemWidth = elem.outerWidth(),
			elemHeight = elem.outerHeight(),
			marginLeft = parseCss( this, "marginLeft" ),
			marginTop = parseCss( this, "marginTop" ),
			collisionWidth = elemWidth + marginLeft + parseCss( this, "marginRight" ) + scrollInfo.width,
			collisionHeight = elemHeight + marginTop + parseCss( this, "marginBottom" ) + scrollInfo.height,
			position = $.extend( {}, basePosition ),
			myOffset = getOffsets( offsets.my, elem.outerWidth(), elem.outerHeight() );

		if ( options.my[ 0 ] === "right" ) {
			position.left -= elemWidth;
		} else if ( options.my[ 0 ] === "center" ) {
			position.left -= elemWidth / 2;
		}

		if ( options.my[ 1 ] === "bottom" ) {
			position.top -= elemHeight;
		} else if ( options.my[ 1 ] === "center" ) {
			position.top -= elemHeight / 2;
		}

		position.left += myOffset[ 0 ];
		position.top += myOffset[ 1 ];

		// if the browser doesn't support fractions, then round for consistent results
		if ( !supportsOffsetFractions ) {
			position.left = round( position.left );
			position.top = round( position.top );
		}

		collisionPosition = {
			marginLeft: marginLeft,
			marginTop: marginTop
		};

		$.each( [ "left", "top" ], function( i, dir ) {
			if ( $.ui.position[ collision[ i ] ] ) {
				$.ui.position[ collision[ i ] ][ dir ]( position, {
					targetWidth: targetWidth,
					targetHeight: targetHeight,
					elemWidth: elemWidth,
					elemHeight: elemHeight,
					collisionPosition: collisionPosition,
					collisionWidth: collisionWidth,
					collisionHeight: collisionHeight,
					offset: [ atOffset[ 0 ] + myOffset[ 0 ], atOffset [ 1 ] + myOffset[ 1 ] ],
					my: options.my,
					at: options.at,
					within: within,
					elem: elem
				});
			}
		});

		if ( options.using ) {
			// adds feedback as second argument to using callback, if present
			using = function( props ) {
				var left = targetOffset.left - position.left,
					right = left + targetWidth - elemWidth,
					top = targetOffset.top - position.top,
					bottom = top + targetHeight - elemHeight,
					feedback = {
						target: {
							element: target,
							left: targetOffset.left,
							top: targetOffset.top,
							width: targetWidth,
							height: targetHeight
						},
						element: {
							element: elem,
							left: position.left,
							top: position.top,
							width: elemWidth,
							height: elemHeight
						},
						horizontal: right < 0 ? "left" : left > 0 ? "right" : "center",
						vertical: bottom < 0 ? "top" : top > 0 ? "bottom" : "middle"
					};
				if ( targetWidth < elemWidth && abs( left + right ) < targetWidth ) {
					feedback.horizontal = "center";
				}
				if ( targetHeight < elemHeight && abs( top + bottom ) < targetHeight ) {
					feedback.vertical = "middle";
				}
				if ( max( abs( left ), abs( right ) ) > max( abs( top ), abs( bottom ) ) ) {
					feedback.important = "horizontal";
				} else {
					feedback.important = "vertical";
				}
				options.using.call( this, props, feedback );
			};
		}

		elem.offset( $.extend( position, { using: using } ) );
	});
};

$.ui.position = {
	fit: {
		left: function( position, data ) {
			var within = data.within,
				withinOffset = within.isWindow ? within.scrollLeft : within.offset.left,
				outerWidth = within.width,
				collisionPosLeft = position.left - data.collisionPosition.marginLeft,
				overLeft = withinOffset - collisionPosLeft,
				overRight = collisionPosLeft + data.collisionWidth - outerWidth - withinOffset,
				newOverRight;

			// element is wider than within
			if ( data.collisionWidth > outerWidth ) {
				// element is initially over the left side of within
				if ( overLeft > 0 && overRight <= 0 ) {
					newOverRight = position.left + overLeft + data.collisionWidth - outerWidth - withinOffset;
					position.left += overLeft - newOverRight;
				// element is initially over right side of within
				} else if ( overRight > 0 && overLeft <= 0 ) {
					position.left = withinOffset;
				// element is initially over both left and right sides of within
				} else {
					if ( overLeft > overRight ) {
						position.left = withinOffset + outerWidth - data.collisionWidth;
					} else {
						position.left = withinOffset;
					}
				}
			// too far left -> align with left edge
			} else if ( overLeft > 0 ) {
				position.left += overLeft;
			// too far right -> align with right edge
			} else if ( overRight > 0 ) {
				position.left -= overRight;
			// adjust based on position and margin
			} else {
				position.left = max( position.left - collisionPosLeft, position.left );
			}
		},
		top: function( position, data ) {
			var within = data.within,
				withinOffset = within.isWindow ? within.scrollTop : within.offset.top,
				outerHeight = data.within.height,
				collisionPosTop = position.top - data.collisionPosition.marginTop,
				overTop = withinOffset - collisionPosTop,
				overBottom = collisionPosTop + data.collisionHeight - outerHeight - withinOffset,
				newOverBottom;

			// element is taller than within
			if ( data.collisionHeight > outerHeight ) {
				// element is initially over the top of within
				if ( overTop > 0 && overBottom <= 0 ) {
					newOverBottom = position.top + overTop + data.collisionHeight - outerHeight - withinOffset;
					position.top += overTop - newOverBottom;
				// element is initially over bottom of within
				} else if ( overBottom > 0 && overTop <= 0 ) {
					position.top = withinOffset;
				// element is initially over both top and bottom of within
				} else {
					if ( overTop > overBottom ) {
						position.top = withinOffset + outerHeight - data.collisionHeight;
					} else {
						position.top = withinOffset;
					}
				}
			// too far up -> align with top
			} else if ( overTop > 0 ) {
				position.top += overTop;
			// too far down -> align with bottom edge
			} else if ( overBottom > 0 ) {
				position.top -= overBottom;
			// adjust based on position and margin
			} else {
				position.top = max( position.top - collisionPosTop, position.top );
			}
		}
	},
	flip: {
		left: function( position, data ) {
			var within = data.within,
				withinOffset = within.offset.left + within.scrollLeft,
				outerWidth = within.width,
				offsetLeft = within.isWindow ? within.scrollLeft : within.offset.left,
				collisionPosLeft = position.left - data.collisionPosition.marginLeft,
				overLeft = collisionPosLeft - offsetLeft,
				overRight = collisionPosLeft + data.collisionWidth - outerWidth - offsetLeft,
				myOffset = data.my[ 0 ] === "left" ?
					-data.elemWidth :
					data.my[ 0 ] === "right" ?
						data.elemWidth :
						0,
				atOffset = data.at[ 0 ] === "left" ?
					data.targetWidth :
					data.at[ 0 ] === "right" ?
						-data.targetWidth :
						0,
				offset = -2 * data.offset[ 0 ],
				newOverRight,
				newOverLeft;

			if ( overLeft < 0 ) {
				newOverRight = position.left + myOffset + atOffset + offset + data.collisionWidth - outerWidth - withinOffset;
				if ( newOverRight < 0 || newOverRight < abs( overLeft ) ) {
					position.left += myOffset + atOffset + offset;
				}
			} else if ( overRight > 0 ) {
				newOverLeft = position.left - data.collisionPosition.marginLeft + myOffset + atOffset + offset - offsetLeft;
				if ( newOverLeft > 0 || abs( newOverLeft ) < overRight ) {
					position.left += myOffset + atOffset + offset;
				}
			}
		},
		top: function( position, data ) {
			var within = data.within,
				withinOffset = within.offset.top + within.scrollTop,
				outerHeight = within.height,
				offsetTop = within.isWindow ? within.scrollTop : within.offset.top,
				collisionPosTop = position.top - data.collisionPosition.marginTop,
				overTop = collisionPosTop - offsetTop,
				overBottom = collisionPosTop + data.collisionHeight - outerHeight - offsetTop,
				top = data.my[ 1 ] === "top",
				myOffset = top ?
					-data.elemHeight :
					data.my[ 1 ] === "bottom" ?
						data.elemHeight :
						0,
				atOffset = data.at[ 1 ] === "top" ?
					data.targetHeight :
					data.at[ 1 ] === "bottom" ?
						-data.targetHeight :
						0,
				offset = -2 * data.offset[ 1 ],
				newOverTop,
				newOverBottom;
			if ( overTop < 0 ) {
				newOverBottom = position.top + myOffset + atOffset + offset + data.collisionHeight - outerHeight - withinOffset;
				if ( newOverBottom < 0 || newOverBottom < abs( overTop ) ) {
					position.top += myOffset + atOffset + offset;
				}
			} else if ( overBottom > 0 ) {
				newOverTop = position.top - data.collisionPosition.marginTop + myOffset + atOffset + offset - offsetTop;
				if ( newOverTop > 0 || abs( newOverTop ) < overBottom ) {
					position.top += myOffset + atOffset + offset;
				}
			}
		}
	},
	flipfit: {
		left: function() {
			$.ui.position.flip.left.apply( this, arguments );
			$.ui.position.fit.left.apply( this, arguments );
		},
		top: function() {
			$.ui.position.flip.top.apply( this, arguments );
			$.ui.position.fit.top.apply( this, arguments );
		}
	}
};

// fraction support test
(function() {
	var testElement, testElementParent, testElementStyle, offsetLeft, i,
		body = document.getElementsByTagName( "body" )[ 0 ],
		div = document.createElement( "div" );

	//Create a "fake body" for testing based on method used in jQuery.support
	testElement = document.createElement( body ? "div" : "body" );
	testElementStyle = {
		visibility: "hidden",
		width: 0,
		height: 0,
		border: 0,
		margin: 0,
		background: "none"
	};
	if ( body ) {
		$.extend( testElementStyle, {
			position: "absolute",
			left: "-1000px",
			top: "-1000px"
		});
	}
	for ( i in testElementStyle ) {
		testElement.style[ i ] = testElementStyle[ i ];
	}
	testElement.appendChild( div );
	testElementParent = body || document.documentElement;
	testElementParent.insertBefore( testElement, testElementParent.firstChild );

	div.style.cssText = "position: absolute; left: 10.7432222px;";

	offsetLeft = $( div ).offset().left;
	supportsOffsetFractions = offsetLeft > 10 && offsetLeft < 11;

	testElement.innerHTML = "";
	testElementParent.removeChild( testElement );
})();

})();

var position = $.ui.position;


/*!
 * jQuery UI Draggable 1.11.4
 * http://jqueryui.com
 *
 * Copyright jQuery Foundation and other contributors
 * Released under the MIT license.
 * http://jquery.org/license
 *
 * http://api.jqueryui.com/draggable/
 */


$.widget("ui.draggable", $.ui.mouse, {
	version: "1.11.4",
	widgetEventPrefix: "drag",
	options: {
		addClasses: true,
		appendTo: "parent",
		axis: false,
		connectToSortable: false,
		containment: false,
		cursor: "auto",
		cursorAt: false,
		grid: false,
		handle: false,
		helper: "original",
		iframeFix: false,
		opacity: false,
		refreshPositions: false,
		revert: false,
		revertDuration: 500,
		scope: "default",
		scroll: true,
		scrollSensitivity: 20,
		scrollSpeed: 20,
		snap: false,
		snapMode: "both",
		snapTolerance: 20,
		stack: false,
		zIndex: false,

		// callbacks
		drag: null,
		start: null,
		stop: null
	},
	_create: function() {

		if ( this.options.helper === "original" ) {
			this._setPositionRelative();
		}
		if (this.options.addClasses){
			this.element.addClass("ui-draggable");
		}
		if (this.options.disabled){
			this.element.addClass("ui-draggable-disabled");
		}
		this._setHandleClassName();

		this._mouseInit();
	},

	_setOption: function( key, value ) {
		this._super( key, value );
		if ( key === "handle" ) {
			this._removeHandleClassName();
			this._setHandleClassName();
		}
	},

	_destroy: function() {
		if ( ( this.helper || this.element ).is( ".ui-draggable-dragging" ) ) {
			this.destroyOnClear = true;
			return;
		}
		this.element.removeClass( "ui-draggable ui-draggable-dragging ui-draggable-disabled" );
		this._removeHandleClassName();
		this._mouseDestroy();
	},

	_mouseCapture: function(event) {
		var o = this.options;

		this._blurActiveElement( event );

		// among others, prevent a drag on a resizable-handle
		if (this.helper || o.disabled || $(event.target).closest(".ui-resizable-handle").length > 0) {
			return false;
		}

		//Quit if we're not on a valid handle
		this.handle = this._getHandle(event);
		if (!this.handle) {
			return false;
		}

		this._blockFrames( o.iframeFix === true ? "iframe" : o.iframeFix );

		return true;

	},

	_blockFrames: function( selector ) {
		this.iframeBlocks = this.document.find( selector ).map(function() {
			var iframe = $( this );

			return $( "<div>" )
				.css( "position", "absolute" )
				.appendTo( iframe.parent() )
				.outerWidth( iframe.outerWidth() )
				.outerHeight( iframe.outerHeight() )
				.offset( iframe.offset() )[ 0 ];
		});
	},

	_unblockFrames: function() {
		if ( this.iframeBlocks ) {
			this.iframeBlocks.remove();
			delete this.iframeBlocks;
		}
	},

	_blurActiveElement: function( event ) {
		var document = this.document[ 0 ];

		// Only need to blur if the event occurred on the draggable itself, see #10527
		if ( !this.handleElement.is( event.target ) ) {
			return;
		}

		// support: IE9
		// IE9 throws an "Unspecified error" accessing document.activeElement from an <iframe>
		try {

			// Support: IE9, IE10
			// If the <body> is blurred, IE will switch windows, see #9520
			if ( document.activeElement && document.activeElement.nodeName.toLowerCase() !== "body" ) {

				// Blur any element that currently has focus, see #4261
				$( document.activeElement ).blur();
			}
		} catch ( error ) {}
	},

	_mouseStart: function(event) {

		var o = this.options;

		//Create and append the visible helper
		this.helper = this._createHelper(event);

		this.helper.addClass("ui-draggable-dragging");

		//Cache the helper size
		this._cacheHelperProportions();

		//If ddmanager is used for droppables, set the global draggable
		if ($.ui.ddmanager) {
			$.ui.ddmanager.current = this;
		}

		/*
		 * - Position generation -
		 * This block generates everything position related - it's the core of draggables.
		 */

		//Cache the margins of the original element
		this._cacheMargins();

		//Store the helper's css position
		this.cssPosition = this.helper.css( "position" );
		this.scrollParent = this.helper.scrollParent( true );
		this.offsetParent = this.helper.offsetParent();
		this.hasFixedAncestor = this.helper.parents().filter(function() {
				return $( this ).css( "position" ) === "fixed";
			}).length > 0;

		//The element's absolute position on the page minus margins
		this.positionAbs = this.element.offset();
		this._refreshOffsets( event );

		//Generate the original position
		this.originalPosition = this.position = this._generatePosition( event, false );
		this.originalPageX = event.pageX;
		this.originalPageY = event.pageY;

		//Adjust the mouse offset relative to the helper if "cursorAt" is supplied
		(o.cursorAt && this._adjustOffsetFromHelper(o.cursorAt));

		//Set a containment if given in the options
		this._setContainment();

		//Trigger event + callbacks
		if (this._trigger("start", event) === false) {
			this._clear();
			return false;
		}

		//Recache the helper size
		this._cacheHelperProportions();

		//Prepare the droppable offsets
		if ($.ui.ddmanager && !o.dropBehaviour) {
			$.ui.ddmanager.prepareOffsets(this, event);
		}

		// Reset helper's right/bottom css if they're set and set explicit width/height instead
		// as this prevents resizing of elements with right/bottom set (see #7772)
		this._normalizeRightBottom();

		this._mouseDrag(event, true); //Execute the drag once - this causes the helper not to be visible before getting its correct position

		//If the ddmanager is used for droppables, inform the manager that dragging has started (see #5003)
		if ( $.ui.ddmanager ) {
			$.ui.ddmanager.dragStart(this, event);
		}

		return true;
	},

	_refreshOffsets: function( event ) {
		this.offset = {
			top: this.positionAbs.top - this.margins.top,
			left: this.positionAbs.left - this.margins.left,
			scroll: false,
			parent: this._getParentOffset(),
			relative: this._getRelativeOffset()
		};

		this.offset.click = {
			left: event.pageX - this.offset.left,
			top: event.pageY - this.offset.top
		};
	},

	_mouseDrag: function(event, noPropagation) {
		// reset any necessary cached properties (see #5009)
		if ( this.hasFixedAncestor ) {
			this.offset.parent = this._getParentOffset();
		}

		//Compute the helpers position
		this.position = this._generatePosition( event, true );
		this.positionAbs = this._convertPositionTo("absolute");

		//Call plugins and callbacks and use the resulting position if something is returned
		if (!noPropagation) {
			var ui = this._uiHash();
			if (this._trigger("drag", event, ui) === false) {
				this._mouseUp({});
				return false;
			}
			this.position = ui.position;
		}

		this.helper[ 0 ].style.left = this.position.left + "px";
		this.helper[ 0 ].style.top = this.position.top + "px";

		if ($.ui.ddmanager) {
			$.ui.ddmanager.drag(this, event);
		}

		return false;
	},

	_mouseStop: function(event) {

		//If we are using droppables, inform the manager about the drop
		var that = this,
			dropped = false;
		if ($.ui.ddmanager && !this.options.dropBehaviour) {
			dropped = $.ui.ddmanager.drop(this, event);
		}

		//if a drop comes from outside (a sortable)
		if (this.dropped) {
			dropped = this.dropped;
			this.dropped = false;
		}

		if ((this.options.revert === "invalid" && !dropped) || (this.options.revert === "valid" && dropped) || this.options.revert === true || ($.isFunction(this.options.revert) && this.options.revert.call(this.element, dropped))) {
			$(this.helper).animate(this.originalPosition, parseInt(this.options.revertDuration, 10), function() {
				if (that._trigger("stop", event) !== false) {
					that._clear();
				}
			});
		} else {
			if (this._trigger("stop", event) !== false) {
				this._clear();
			}
		}

		return false;
	},

	_mouseUp: function( event ) {
		this._unblockFrames();

		//If the ddmanager is used for droppables, inform the manager that dragging has stopped (see #5003)
		if ( $.ui.ddmanager ) {
			$.ui.ddmanager.dragStop(this, event);
		}

		// Only need to focus if the event occurred on the draggable itself, see #10527
		if ( this.handleElement.is( event.target ) ) {
			// The interaction is over; whether or not the click resulted in a drag, focus the element
			this.element.focus();
		}

		return $.ui.mouse.prototype._mouseUp.call(this, event);
	},

	cancel: function() {

		if (this.helper.is(".ui-draggable-dragging")) {
			this._mouseUp({});
		} else {
			this._clear();
		}

		return this;

	},

	_getHandle: function(event) {
		return this.options.handle ?
			!!$( event.target ).closest( this.element.find( this.options.handle ) ).length :
			true;
	},

	_setHandleClassName: function() {
		this.handleElement = this.options.handle ?
			this.element.find( this.options.handle ) : this.element;
		this.handleElement.addClass( "ui-draggable-handle" );
	},

	_removeHandleClassName: function() {
		this.handleElement.removeClass( "ui-draggable-handle" );
	},

	_createHelper: function(event) {

		var o = this.options,
			helperIsFunction = $.isFunction( o.helper ),
			helper = helperIsFunction ?
				$( o.helper.apply( this.element[ 0 ], [ event ] ) ) :
				( o.helper === "clone" ?
					this.element.clone().removeAttr( "id" ) :
					this.element );

		if (!helper.parents("body").length) {
			helper.appendTo((o.appendTo === "parent" ? this.element[0].parentNode : o.appendTo));
		}

		// http://bugs.jqueryui.com/ticket/9446
		// a helper function can return the original element
		// which wouldn't have been set to relative in _create
		if ( helperIsFunction && helper[ 0 ] === this.element[ 0 ] ) {
			this._setPositionRelative();
		}

		if (helper[0] !== this.element[0] && !(/(fixed|absolute)/).test(helper.css("position"))) {
			helper.css("position", "absolute");
		}

		return helper;

	},

	_setPositionRelative: function() {
		if ( !( /^(?:r|a|f)/ ).test( this.element.css( "position" ) ) ) {
			this.element[ 0 ].style.position = "relative";
		}
	},

	_adjustOffsetFromHelper: function(obj) {
		if (typeof obj === "string") {
			obj = obj.split(" ");
		}
		if ($.isArray(obj)) {
			obj = { left: +obj[0], top: +obj[1] || 0 };
		}
		if ("left" in obj) {
			this.offset.click.left = obj.left + this.margins.left;
		}
		if ("right" in obj) {
			this.offset.click.left = this.helperProportions.width - obj.right + this.margins.left;
		}
		if ("top" in obj) {
			this.offset.click.top = obj.top + this.margins.top;
		}
		if ("bottom" in obj) {
			this.offset.click.top = this.helperProportions.height - obj.bottom + this.margins.top;
		}
	},

	_isRootNode: function( element ) {
		return ( /(html|body)/i ).test( element.tagName ) || element === this.document[ 0 ];
	},

	_getParentOffset: function() {

		//Get the offsetParent and cache its position
		var po = this.offsetParent.offset(),
			document = this.document[ 0 ];

		// This is a special case where we need to modify a offset calculated on start, since the following happened:
		// 1. The position of the helper is absolute, so it's position is calculated based on the next positioned parent
		// 2. The actual offset parent is a child of the scroll parent, and the scroll parent isn't the document, which means that
		//    the scroll is included in the initial calculation of the offset of the parent, and never recalculated upon drag
		if (this.cssPosition === "absolute" && this.scrollParent[0] !== document && $.contains(this.scrollParent[0], this.offsetParent[0])) {
			po.left += this.scrollParent.scrollLeft();
			po.top += this.scrollParent.scrollTop();
		}

		if ( this._isRootNode( this.offsetParent[ 0 ] ) ) {
			po = { top: 0, left: 0 };
		}

		return {
			top: po.top + (parseInt(this.offsetParent.css("borderTopWidth"), 10) || 0),
			left: po.left + (parseInt(this.offsetParent.css("borderLeftWidth"), 10) || 0)
		};

	},

	_getRelativeOffset: function() {
		if ( this.cssPosition !== "relative" ) {
			return { top: 0, left: 0 };
		}

		var p = this.element.position(),
			scrollIsRootNode = this._isRootNode( this.scrollParent[ 0 ] );

		return {
			top: p.top - ( parseInt(this.helper.css( "top" ), 10) || 0 ) + ( !scrollIsRootNode ? this.scrollParent.scrollTop() : 0 ),
			left: p.left - ( parseInt(this.helper.css( "left" ), 10) || 0 ) + ( !scrollIsRootNode ? this.scrollParent.scrollLeft() : 0 )
		};

	},

	_cacheMargins: function() {
		this.margins = {
			left: (parseInt(this.element.css("marginLeft"), 10) || 0),
			top: (parseInt(this.element.css("marginTop"), 10) || 0),
			right: (parseInt(this.element.css("marginRight"), 10) || 0),
			bottom: (parseInt(this.element.css("marginBottom"), 10) || 0)
		};
	},

	_cacheHelperProportions: function() {
		this.helperProportions = {
			width: this.helper.outerWidth(),
			height: this.helper.outerHeight()
		};
	},

	_setContainment: function() {

		var isUserScrollable, c, ce,
			o = this.options,
			document = this.document[ 0 ];

		this.relativeContainer = null;

		if ( !o.containment ) {
			this.containment = null;
			return;
		}

		if ( o.containment === "window" ) {
			this.containment = [
				$( window ).scrollLeft() - this.offset.relative.left - this.offset.parent.left,
				$( window ).scrollTop() - this.offset.relative.top - this.offset.parent.top,
				$( window ).scrollLeft() + $( window ).width() - this.helperProportions.width - this.margins.left,
				$( window ).scrollTop() + ( $( window ).height() || document.body.parentNode.scrollHeight ) - this.helperProportions.height - this.margins.top
			];
			return;
		}

		if ( o.containment === "document") {
			this.containment = [
				0,
				0,
				$( document ).width() - this.helperProportions.width - this.margins.left,
				( $( document ).height() || document.body.parentNode.scrollHeight ) - this.helperProportions.height - this.margins.top
			];
			return;
		}

		if ( o.containment.constructor === Array ) {
			this.containment = o.containment;
			return;
		}

		if ( o.containment === "parent" ) {
			o.containment = this.helper[ 0 ].parentNode;
		}

		c = $( o.containment );
		ce = c[ 0 ];

		if ( !ce ) {
			return;
		}

		isUserScrollable = /(scroll|auto)/.test( c.css( "overflow" ) );

		this.containment = [
			( parseInt( c.css( "borderLeftWidth" ), 10 ) || 0 ) + ( parseInt( c.css( "paddingLeft" ), 10 ) || 0 ),
			( parseInt( c.css( "borderTopWidth" ), 10 ) || 0 ) + ( parseInt( c.css( "paddingTop" ), 10 ) || 0 ),
			( isUserScrollable ? Math.max( ce.scrollWidth, ce.offsetWidth ) : ce.offsetWidth ) -
				( parseInt( c.css( "borderRightWidth" ), 10 ) || 0 ) -
				( parseInt( c.css( "paddingRight" ), 10 ) || 0 ) -
				this.helperProportions.width -
				this.margins.left -
				this.margins.right,
			( isUserScrollable ? Math.max( ce.scrollHeight, ce.offsetHeight ) : ce.offsetHeight ) -
				( parseInt( c.css( "borderBottomWidth" ), 10 ) || 0 ) -
				( parseInt( c.css( "paddingBottom" ), 10 ) || 0 ) -
				this.helperProportions.height -
				this.margins.top -
				this.margins.bottom
		];
		this.relativeContainer = c;
	},

	_convertPositionTo: function(d, pos) {

		if (!pos) {
			pos = this.position;
		}

		var mod = d === "absolute" ? 1 : -1,
			scrollIsRootNode = this._isRootNode( this.scrollParent[ 0 ] );

		return {
			top: (
				pos.top	+																// The absolute mouse position
				this.offset.relative.top * mod +										// Only for relative positioned nodes: Relative offset from element to offset parent
				this.offset.parent.top * mod -										// The offsetParent's offset without borders (offset + border)
				( ( this.cssPosition === "fixed" ? -this.offset.scroll.top : ( scrollIsRootNode ? 0 : this.offset.scroll.top ) ) * mod)
			),
			left: (
				pos.left +																// The absolute mouse position
				this.offset.relative.left * mod +										// Only for relative positioned nodes: Relative offset from element to offset parent
				this.offset.parent.left * mod	-										// The offsetParent's offset without borders (offset + border)
				( ( this.cssPosition === "fixed" ? -this.offset.scroll.left : ( scrollIsRootNode ? 0 : this.offset.scroll.left ) ) * mod)
			)
		};

	},

	_generatePosition: function( event, constrainPosition ) {

		var containment, co, top, left,
			o = this.options,
			scrollIsRootNode = this._isRootNode( this.scrollParent[ 0 ] ),
			pageX = event.pageX,
			pageY = event.pageY;

		// Cache the scroll
		if ( !scrollIsRootNode || !this.offset.scroll ) {
			this.offset.scroll = {
				top: this.scrollParent.scrollTop(),
				left: this.scrollParent.scrollLeft()
			};
		}

		/*
		 * - Position constraining -
		 * Constrain the position to a mix of grid, containment.
		 */

		// If we are not dragging yet, we won't check for options
		if ( constrainPosition ) {
			if ( this.containment ) {
				if ( this.relativeContainer ){
					co = this.relativeContainer.offset();
					containment = [
						this.containment[ 0 ] + co.left,
						this.containment[ 1 ] + co.top,
						this.containment[ 2 ] + co.left,
						this.containment[ 3 ] + co.top
					];
				} else {
					containment = this.containment;
				}

				if (event.pageX - this.offset.click.left < containment[0]) {
					pageX = containment[0] + this.offset.click.left;
				}
				if (event.pageY - this.offset.click.top < containment[1]) {
					pageY = containment[1] + this.offset.click.top;
				}
				if (event.pageX - this.offset.click.left > containment[2]) {
					pageX = containment[2] + this.offset.click.left;
				}
				if (event.pageY - this.offset.click.top > containment[3]) {
					pageY = containment[3] + this.offset.click.top;
				}
			}

			if (o.grid) {
				//Check for grid elements set to 0 to prevent divide by 0 error causing invalid argument errors in IE (see ticket #6950)
				top = o.grid[1] ? this.originalPageY + Math.round((pageY - this.originalPageY) / o.grid[1]) * o.grid[1] : this.originalPageY;
				pageY = containment ? ((top - this.offset.click.top >= containment[1] || top - this.offset.click.top > containment[3]) ? top : ((top - this.offset.click.top >= containment[1]) ? top - o.grid[1] : top + o.grid[1])) : top;

				left = o.grid[0] ? this.originalPageX + Math.round((pageX - this.originalPageX) / o.grid[0]) * o.grid[0] : this.originalPageX;
				pageX = containment ? ((left - this.offset.click.left >= containment[0] || left - this.offset.click.left > containment[2]) ? left : ((left - this.offset.click.left >= containment[0]) ? left - o.grid[0] : left + o.grid[0])) : left;
			}

			if ( o.axis === "y" ) {
				pageX = this.originalPageX;
			}

			if ( o.axis === "x" ) {
				pageY = this.originalPageY;
			}
		}

		return {
			top: (
				pageY -																	// The absolute mouse position
				this.offset.click.top	-												// Click offset (relative to the element)
				this.offset.relative.top -												// Only for relative positioned nodes: Relative offset from element to offset parent
				this.offset.parent.top +												// The offsetParent's offset without borders (offset + border)
				( this.cssPosition === "fixed" ? -this.offset.scroll.top : ( scrollIsRootNode ? 0 : this.offset.scroll.top ) )
			),
			left: (
				pageX -																	// The absolute mouse position
				this.offset.click.left -												// Click offset (relative to the element)
				this.offset.relative.left -												// Only for relative positioned nodes: Relative offset from element to offset parent
				this.offset.parent.left +												// The offsetParent's offset without borders (offset + border)
				( this.cssPosition === "fixed" ? -this.offset.scroll.left : ( scrollIsRootNode ? 0 : this.offset.scroll.left ) )
			)
		};

	},

	_clear: function() {
		this.helper.removeClass("ui-draggable-dragging");
		if (this.helper[0] !== this.element[0] && !this.cancelHelperRemoval) {
			this.helper.remove();
		}
		this.helper = null;
		this.cancelHelperRemoval = false;
		if ( this.destroyOnClear ) {
			this.destroy();
		}
	},

	_normalizeRightBottom: function() {
		if ( this.options.axis !== "y" && this.helper.css( "right" ) !== "auto" ) {
			this.helper.width( this.helper.width() );
			this.helper.css( "right", "auto" );
		}
		if ( this.options.axis !== "x" && this.helper.css( "bottom" ) !== "auto" ) {
			this.helper.height( this.helper.height() );
			this.helper.css( "bottom", "auto" );
		}
	},

	// From now on bulk stuff - mainly helpers

	_trigger: function( type, event, ui ) {
		ui = ui || this._uiHash();
		$.ui.plugin.call( this, type, [ event, ui, this ], true );

		// Absolute position and offset (see #6884 ) have to be recalculated after plugins
		if ( /^(drag|start|stop)/.test( type ) ) {
			this.positionAbs = this._convertPositionTo( "absolute" );
			ui.offset = this.positionAbs;
		}
		return $.Widget.prototype._trigger.call( this, type, event, ui );
	},

	plugins: {},

	_uiHash: function() {
		return {
			helper: this.helper,
			position: this.position,
			originalPosition: this.originalPosition,
			offset: this.positionAbs
		};
	}

});

$.ui.plugin.add( "draggable", "connectToSortable", {
	start: function( event, ui, draggable ) {
		var uiSortable = $.extend( {}, ui, {
			item: draggable.element
		});

		draggable.sortables = [];
		$( draggable.options.connectToSortable ).each(function() {
			var sortable = $( this ).sortable( "instance" );

			if ( sortable && !sortable.options.disabled ) {
				draggable.sortables.push( sortable );

				// refreshPositions is called at drag start to refresh the containerCache
				// which is used in drag. This ensures it's initialized and synchronized
				// with any changes that might have happened on the page since initialization.
				sortable.refreshPositions();
				sortable._trigger("activate", event, uiSortable);
			}
		});
	},
	stop: function( event, ui, draggable ) {
		var uiSortable = $.extend( {}, ui, {
			item: draggable.element
		});

		draggable.cancelHelperRemoval = false;

		$.each( draggable.sortables, function() {
			var sortable = this;

			if ( sortable.isOver ) {
				sortable.isOver = 0;

				// Allow this sortable to handle removing the helper
				draggable.cancelHelperRemoval = true;
				sortable.cancelHelperRemoval = false;

				// Use _storedCSS To restore properties in the sortable,
				// as this also handles revert (#9675) since the draggable
				// may have modified them in unexpected ways (#8809)
				sortable._storedCSS = {
					position: sortable.placeholder.css( "position" ),
					top: sortable.placeholder.css( "top" ),
					left: sortable.placeholder.css( "left" )
				};

				sortable._mouseStop(event);

				// Once drag has ended, the sortable should return to using
				// its original helper, not the shared helper from draggable
				sortable.options.helper = sortable.options._helper;
			} else {
				// Prevent this Sortable from removing the helper.
				// However, don't set the draggable to remove the helper
				// either as another connected Sortable may yet handle the removal.
				sortable.cancelHelperRemoval = true;

				sortable._trigger( "deactivate", event, uiSortable );
			}
		});
	},
	drag: function( event, ui, draggable ) {
		$.each( draggable.sortables, function() {
			var innermostIntersecting = false,
				sortable = this;

			// Copy over variables that sortable's _intersectsWith uses
			sortable.positionAbs = draggable.positionAbs;
			sortable.helperProportions = draggable.helperProportions;
			sortable.offset.click = draggable.offset.click;

			if ( sortable._intersectsWith( sortable.containerCache ) ) {
				innermostIntersecting = true;

				$.each( draggable.sortables, function() {
					// Copy over variables that sortable's _intersectsWith uses
					this.positionAbs = draggable.positionAbs;
					this.helperProportions = draggable.helperProportions;
					this.offset.click = draggable.offset.click;

					if ( this !== sortable &&
							this._intersectsWith( this.containerCache ) &&
							$.contains( sortable.element[ 0 ], this.element[ 0 ] ) ) {
						innermostIntersecting = false;
					}

					return innermostIntersecting;
				});
			}

			if ( innermostIntersecting ) {
				// If it intersects, we use a little isOver variable and set it once,
				// so that the move-in stuff gets fired only once.
				if ( !sortable.isOver ) {
					sortable.isOver = 1;

					// Store draggable's parent in case we need to reappend to it later.
					draggable._parent = ui.helper.parent();

					sortable.currentItem = ui.helper
						.appendTo( sortable.element )
						.data( "ui-sortable-item", true );

					// Store helper option to later restore it
					sortable.options._helper = sortable.options.helper;

					sortable.options.helper = function() {
						return ui.helper[ 0 ];
					};

					// Fire the start events of the sortable with our passed browser event,
					// and our own helper (so it doesn't create a new one)
					event.target = sortable.currentItem[ 0 ];
					sortable._mouseCapture( event, true );
					sortable._mouseStart( event, true, true );

					// Because the browser event is way off the new appended portlet,
					// modify necessary variables to reflect the changes
					sortable.offset.click.top = draggable.offset.click.top;
					sortable.offset.click.left = draggable.offset.click.left;
					sortable.offset.parent.left -= draggable.offset.parent.left -
						sortable.offset.parent.left;
					sortable.offset.parent.top -= draggable.offset.parent.top -
						sortable.offset.parent.top;

					draggable._trigger( "toSortable", event );

					// Inform draggable that the helper is in a valid drop zone,
					// used solely in the revert option to handle "valid/invalid".
					draggable.dropped = sortable.element;

					// Need to refreshPositions of all sortables in the case that
					// adding to one sortable changes the location of the other sortables (#9675)
					$.each( draggable.sortables, function() {
						this.refreshPositions();
					});

					// hack so receive/update callbacks work (mostly)
					draggable.currentItem = draggable.element;
					sortable.fromOutside = draggable;
				}

				if ( sortable.currentItem ) {
					sortable._mouseDrag( event );
					// Copy the sortable's position because the draggable's can potentially reflect
					// a relative position, while sortable is always absolute, which the dragged
					// element has now become. (#8809)
					ui.position = sortable.position;
				}
			} else {
				// If it doesn't intersect with the sortable, and it intersected before,
				// we fake the drag stop of the sortable, but make sure it doesn't remove
				// the helper by using cancelHelperRemoval.
				if ( sortable.isOver ) {

					sortable.isOver = 0;
					sortable.cancelHelperRemoval = true;

					// Calling sortable's mouseStop would trigger a revert,
					// so revert must be temporarily false until after mouseStop is called.
					sortable.options._revert = sortable.options.revert;
					sortable.options.revert = false;

					sortable._trigger( "out", event, sortable._uiHash( sortable ) );
					sortable._mouseStop( event, true );

					// restore sortable behaviors that were modfied
					// when the draggable entered the sortable area (#9481)
					sortable.options.revert = sortable.options._revert;
					sortable.options.helper = sortable.options._helper;

					if ( sortable.placeholder ) {
						sortable.placeholder.remove();
					}

					// Restore and recalculate the draggable's offset considering the sortable
					// may have modified them in unexpected ways. (#8809, #10669)
					ui.helper.appendTo( draggable._parent );
					draggable._refreshOffsets( event );
					ui.position = draggable._generatePosition( event, true );

					draggable._trigger( "fromSortable", event );

					// Inform draggable that the helper is no longer in a valid drop zone
					draggable.dropped = false;

					// Need to refreshPositions of all sortables just in case removing
					// from one sortable changes the location of other sortables (#9675)
					$.each( draggable.sortables, function() {
						this.refreshPositions();
					});
				}
			}
		});
	}
});

$.ui.plugin.add("draggable", "cursor", {
	start: function( event, ui, instance ) {
		var t = $( "body" ),
			o = instance.options;

		if (t.css("cursor")) {
			o._cursor = t.css("cursor");
		}
		t.css("cursor", o.cursor);
	},
	stop: function( event, ui, instance ) {
		var o = instance.options;
		if (o._cursor) {
			$("body").css("cursor", o._cursor);
		}
	}
});

$.ui.plugin.add("draggable", "opacity", {
	start: function( event, ui, instance ) {
		var t = $( ui.helper ),
			o = instance.options;
		if (t.css("opacity")) {
			o._opacity = t.css("opacity");
		}
		t.css("opacity", o.opacity);
	},
	stop: function( event, ui, instance ) {
		var o = instance.options;
		if (o._opacity) {
			$(ui.helper).css("opacity", o._opacity);
		}
	}
});

$.ui.plugin.add("draggable", "scroll", {
	start: function( event, ui, i ) {
		if ( !i.scrollParentNotHidden ) {
			i.scrollParentNotHidden = i.helper.scrollParent( false );
		}

		if ( i.scrollParentNotHidden[ 0 ] !== i.document[ 0 ] && i.scrollParentNotHidden[ 0 ].tagName !== "HTML" ) {
			i.overflowOffset = i.scrollParentNotHidden.offset();
		}
	},
	drag: function( event, ui, i  ) {

		var o = i.options,
			scrolled = false,
			scrollParent = i.scrollParentNotHidden[ 0 ],
			document = i.document[ 0 ];

		if ( scrollParent !== document && scrollParent.tagName !== "HTML" ) {
			if ( !o.axis || o.axis !== "x" ) {
				if ( ( i.overflowOffset.top + scrollParent.offsetHeight ) - event.pageY < o.scrollSensitivity ) {
					scrollParent.scrollTop = scrolled = scrollParent.scrollTop + o.scrollSpeed;
				} else if ( event.pageY - i.overflowOffset.top < o.scrollSensitivity ) {
					scrollParent.scrollTop = scrolled = scrollParent.scrollTop - o.scrollSpeed;
				}
			}

			if ( !o.axis || o.axis !== "y" ) {
				if ( ( i.overflowOffset.left + scrollParent.offsetWidth ) - event.pageX < o.scrollSensitivity ) {
					scrollParent.scrollLeft = scrolled = scrollParent.scrollLeft + o.scrollSpeed;
				} else if ( event.pageX - i.overflowOffset.left < o.scrollSensitivity ) {
					scrollParent.scrollLeft = scrolled = scrollParent.scrollLeft - o.scrollSpeed;
				}
			}

		} else {

			if (!o.axis || o.axis !== "x") {
				if (event.pageY - $(document).scrollTop() < o.scrollSensitivity) {
					scrolled = $(document).scrollTop($(document).scrollTop() - o.scrollSpeed);
				} else if ($(window).height() - (event.pageY - $(document).scrollTop()) < o.scrollSensitivity) {
					scrolled = $(document).scrollTop($(document).scrollTop() + o.scrollSpeed);
				}
			}

			if (!o.axis || o.axis !== "y") {
				if (event.pageX - $(document).scrollLeft() < o.scrollSensitivity) {
					scrolled = $(document).scrollLeft($(document).scrollLeft() - o.scrollSpeed);
				} else if ($(window).width() - (event.pageX - $(document).scrollLeft()) < o.scrollSensitivity) {
					scrolled = $(document).scrollLeft($(document).scrollLeft() + o.scrollSpeed);
				}
			}

		}

		if (scrolled !== false && $.ui.ddmanager && !o.dropBehaviour) {
			$.ui.ddmanager.prepareOffsets(i, event);
		}

	}
});

$.ui.plugin.add("draggable", "snap", {
	start: function( event, ui, i ) {

		var o = i.options;

		i.snapElements = [];

		$(o.snap.constructor !== String ? ( o.snap.items || ":data(ui-draggable)" ) : o.snap).each(function() {
			var $t = $(this),
				$o = $t.offset();
			if (this !== i.element[0]) {
				i.snapElements.push({
					item: this,
					width: $t.outerWidth(), height: $t.outerHeight(),
					top: $o.top, left: $o.left
				});
			}
		});

	},
	drag: function( event, ui, inst ) {

		var ts, bs, ls, rs, l, r, t, b, i, first,
			o = inst.options,
			d = o.snapTolerance,
			x1 = ui.offset.left, x2 = x1 + inst.helperProportions.width,
			y1 = ui.offset.top, y2 = y1 + inst.helperProportions.height;

		for (i = inst.snapElements.length - 1; i >= 0; i--){

			l = inst.snapElements[i].left - inst.margins.left;
			r = l + inst.snapElements[i].width;
			t = inst.snapElements[i].top - inst.margins.top;
			b = t + inst.snapElements[i].height;

			if ( x2 < l - d || x1 > r + d || y2 < t - d || y1 > b + d || !$.contains( inst.snapElements[ i ].item.ownerDocument, inst.snapElements[ i ].item ) ) {
				if (inst.snapElements[i].snapping) {
					(inst.options.snap.release && inst.options.snap.release.call(inst.element, event, $.extend(inst._uiHash(), { snapItem: inst.snapElements[i].item })));
				}
				inst.snapElements[i].snapping = false;
				continue;
			}

			if (o.snapMode !== "inner") {
				ts = Math.abs(t - y2) <= d;
				bs = Math.abs(b - y1) <= d;
				ls = Math.abs(l - x2) <= d;
				rs = Math.abs(r - x1) <= d;
				if (ts) {
					ui.position.top = inst._convertPositionTo("relative", { top: t - inst.helperProportions.height, left: 0 }).top;
				}
				if (bs) {
					ui.position.top = inst._convertPositionTo("relative", { top: b, left: 0 }).top;
				}
				if (ls) {
					ui.position.left = inst._convertPositionTo("relative", { top: 0, left: l - inst.helperProportions.width }).left;
				}
				if (rs) {
					ui.position.left = inst._convertPositionTo("relative", { top: 0, left: r }).left;
				}
			}

			first = (ts || bs || ls || rs);

			if (o.snapMode !== "outer") {
				ts = Math.abs(t - y1) <= d;
				bs = Math.abs(b - y2) <= d;
				ls = Math.abs(l - x1) <= d;
				rs = Math.abs(r - x2) <= d;
				if (ts) {
					ui.position.top = inst._convertPositionTo("relative", { top: t, left: 0 }).top;
				}
				if (bs) {
					ui.position.top = inst._convertPositionTo("relative", { top: b - inst.helperProportions.height, left: 0 }).top;
				}
				if (ls) {
					ui.position.left = inst._convertPositionTo("relative", { top: 0, left: l }).left;
				}
				if (rs) {
					ui.position.left = inst._convertPositionTo("relative", { top: 0, left: r - inst.helperProportions.width }).left;
				}
			}

			if (!inst.snapElements[i].snapping && (ts || bs || ls || rs || first)) {
				(inst.options.snap.snap && inst.options.snap.snap.call(inst.element, event, $.extend(inst._uiHash(), { snapItem: inst.snapElements[i].item })));
			}
			inst.snapElements[i].snapping = (ts || bs || ls || rs || first);

		}

	}
});

$.ui.plugin.add("draggable", "stack", {
	start: function( event, ui, instance ) {
		var min,
			o = instance.options,
			group = $.makeArray($(o.stack)).sort(function(a, b) {
				return (parseInt($(a).css("zIndex"), 10) || 0) - (parseInt($(b).css("zIndex"), 10) || 0);
			});

		if (!group.length) { return; }

		min = parseInt($(group[0]).css("zIndex"), 10) || 0;
		$(group).each(function(i) {
			$(this).css("zIndex", min + i);
		});
		this.css("zIndex", (min + group.length));
	}
});

$.ui.plugin.add("draggable", "zIndex", {
	start: function( event, ui, instance ) {
		var t = $( ui.helper ),
			o = instance.options;

		if (t.css("zIndex")) {
			o._zIndex = t.css("zIndex");
		}
		t.css("zIndex", o.zIndex);
	},
	stop: function( event, ui, instance ) {
		var o = instance.options;

		if (o._zIndex) {
			$(ui.helper).css("zIndex", o._zIndex);
		}
	}
});

var draggable = $.ui.draggable;


/*!
 * jQuery UI Droppable 1.11.4
 * http://jqueryui.com
 *
 * Copyright jQuery Foundation and other contributors
 * Released under the MIT license.
 * http://jquery.org/license
 *
 * http://api.jqueryui.com/droppable/
 */


$.widget( "ui.droppable", {
	version: "1.11.4",
	widgetEventPrefix: "drop",
	options: {
		accept: "*",
		activeClass: false,
		addClasses: true,
		greedy: false,
		hoverClass: false,
		scope: "default",
		tolerance: "intersect",

		// callbacks
		activate: null,
		deactivate: null,
		drop: null,
		out: null,
		over: null
	},
	_create: function() {

		var proportions,
			o = this.options,
			accept = o.accept;

		this.isover = false;
		this.isout = true;

		this.accept = $.isFunction( accept ) ? accept : function( d ) {
			return d.is( accept );
		};

		this.proportions = function( /* valueToWrite */ ) {
			if ( arguments.length ) {
				// Store the droppable's proportions
				proportions = arguments[ 0 ];
			} else {
				// Retrieve or derive the droppable's proportions
				return proportions ?
					proportions :
					proportions = {
						width: this.element[ 0 ].offsetWidth,
						height: this.element[ 0 ].offsetHeight
					};
			}
		};

		this._addToManager( o.scope );

		o.addClasses && this.element.addClass( "ui-droppable" );

	},

	_addToManager: function( scope ) {
		// Add the reference and positions to the manager
		$.ui.ddmanager.droppables[ scope ] = $.ui.ddmanager.droppables[ scope ] || [];
		$.ui.ddmanager.droppables[ scope ].push( this );
	},

	_splice: function( drop ) {
		var i = 0;
		for ( ; i < drop.length; i++ ) {
			if ( drop[ i ] === this ) {
				drop.splice( i, 1 );
			}
		}
	},

	_destroy: function() {
		var drop = $.ui.ddmanager.droppables[ this.options.scope ];

		this._splice( drop );

		this.element.removeClass( "ui-droppable ui-droppable-disabled" );
	},

	_setOption: function( key, value ) {

		if ( key === "accept" ) {
			this.accept = $.isFunction( value ) ? value : function( d ) {
				return d.is( value );
			};
		} else if ( key === "scope" ) {
			var drop = $.ui.ddmanager.droppables[ this.options.scope ];

			this._splice( drop );
			this._addToManager( value );
		}

		this._super( key, value );
	},

	_activate: function( event ) {
		var draggable = $.ui.ddmanager.current;
		if ( this.options.activeClass ) {
			this.element.addClass( this.options.activeClass );
		}
		if ( draggable ){
			this._trigger( "activate", event, this.ui( draggable ) );
		}
	},

	_deactivate: function( event ) {
		var draggable = $.ui.ddmanager.current;
		if ( this.options.activeClass ) {
			this.element.removeClass( this.options.activeClass );
		}
		if ( draggable ){
			this._trigger( "deactivate", event, this.ui( draggable ) );
		}
	},

	_over: function( event ) {

		var draggable = $.ui.ddmanager.current;

		// Bail if draggable and droppable are same element
		if ( !draggable || ( draggable.currentItem || draggable.element )[ 0 ] === this.element[ 0 ] ) {
			return;
		}

		if ( this.accept.call( this.element[ 0 ], ( draggable.currentItem || draggable.element ) ) ) {
			if ( this.options.hoverClass ) {
				this.element.addClass( this.options.hoverClass );
			}
			this._trigger( "over", event, this.ui( draggable ) );
		}

	},

	_out: function( event ) {

		var draggable = $.ui.ddmanager.current;

		// Bail if draggable and droppable are same element
		if ( !draggable || ( draggable.currentItem || draggable.element )[ 0 ] === this.element[ 0 ] ) {
			return;
		}

		if ( this.accept.call( this.element[ 0 ], ( draggable.currentItem || draggable.element ) ) ) {
			if ( this.options.hoverClass ) {
				this.element.removeClass( this.options.hoverClass );
			}
			this._trigger( "out", event, this.ui( draggable ) );
		}

	},

	_drop: function( event, custom ) {

		var draggable = custom || $.ui.ddmanager.current,
			childrenIntersection = false;

		// Bail if draggable and droppable are same element
		if ( !draggable || ( draggable.currentItem || draggable.element )[ 0 ] === this.element[ 0 ] ) {
			return false;
		}

		this.element.find( ":data(ui-droppable)" ).not( ".ui-draggable-dragging" ).each(function() {
			var inst = $( this ).droppable( "instance" );
			if (
				inst.options.greedy &&
				!inst.options.disabled &&
				inst.options.scope === draggable.options.scope &&
				inst.accept.call( inst.element[ 0 ], ( draggable.currentItem || draggable.element ) ) &&
				$.ui.intersect( draggable, $.extend( inst, { offset: inst.element.offset() } ), inst.options.tolerance, event )
			) { childrenIntersection = true; return false; }
		});
		if ( childrenIntersection ) {
			return false;
		}

		if ( this.accept.call( this.element[ 0 ], ( draggable.currentItem || draggable.element ) ) ) {
			if ( this.options.activeClass ) {
				this.element.removeClass( this.options.activeClass );
			}
			if ( this.options.hoverClass ) {
				this.element.removeClass( this.options.hoverClass );
			}
			this._trigger( "drop", event, this.ui( draggable ) );
			return this.element;
		}

		return false;

	},

	ui: function( c ) {
		return {
			draggable: ( c.currentItem || c.element ),
			helper: c.helper,
			position: c.position,
			offset: c.positionAbs
		};
	}

});

$.ui.intersect = (function() {
	function isOverAxis( x, reference, size ) {
		return ( x >= reference ) && ( x < ( reference + size ) );
	}

	return function( draggable, droppable, toleranceMode, event ) {

		if ( !droppable.offset ) {
			return false;
		}

		var x1 = ( draggable.positionAbs || draggable.position.absolute ).left + draggable.margins.left,
			y1 = ( draggable.positionAbs || draggable.position.absolute ).top + draggable.margins.top,
			x2 = x1 + draggable.helperProportions.width,
			y2 = y1 + draggable.helperProportions.height,
			l = droppable.offset.left,
			t = droppable.offset.top,
			r = l + droppable.proportions().width,
			b = t + droppable.proportions().height;

		switch ( toleranceMode ) {
		case "fit":
			return ( l <= x1 && x2 <= r && t <= y1 && y2 <= b );
		case "intersect":
			return ( l < x1 + ( draggable.helperProportions.width / 2 ) && // Right Half
				x2 - ( draggable.helperProportions.width / 2 ) < r && // Left Half
				t < y1 + ( draggable.helperProportions.height / 2 ) && // Bottom Half
				y2 - ( draggable.helperProportions.height / 2 ) < b ); // Top Half
		case "pointer":
			return isOverAxis( event.pageY, t, droppable.proportions().height ) && isOverAxis( event.pageX, l, droppable.proportions().width );
		case "touch":
			return (
				( y1 >= t && y1 <= b ) || // Top edge touching
				( y2 >= t && y2 <= b ) || // Bottom edge touching
				( y1 < t && y2 > b ) // Surrounded vertically
			) && (
				( x1 >= l && x1 <= r ) || // Left edge touching
				( x2 >= l && x2 <= r ) || // Right edge touching
				( x1 < l && x2 > r ) // Surrounded horizontally
			);
		default:
			return false;
		}
	};
})();

/*
	This manager tracks offsets of draggables and droppables
*/
$.ui.ddmanager = {
	current: null,
	droppables: { "default": [] },
	prepareOffsets: function( t, event ) {

		var i, j,
			m = $.ui.ddmanager.droppables[ t.options.scope ] || [],
			type = event ? event.type : null, // workaround for #2317
			list = ( t.currentItem || t.element ).find( ":data(ui-droppable)" ).addBack();

		droppablesLoop: for ( i = 0; i < m.length; i++ ) {

			// No disabled and non-accepted
			if ( m[ i ].options.disabled || ( t && !m[ i ].accept.call( m[ i ].element[ 0 ], ( t.currentItem || t.element ) ) ) ) {
				continue;
			}

			// Filter out elements in the current dragged item
			for ( j = 0; j < list.length; j++ ) {
				if ( list[ j ] === m[ i ].element[ 0 ] ) {
					m[ i ].proportions().height = 0;
					continue droppablesLoop;
				}
			}

			m[ i ].visible = m[ i ].element.css( "display" ) !== "none";
			if ( !m[ i ].visible ) {
				continue;
			}

			// Activate the droppable if used directly from draggables
			if ( type === "mousedown" ) {
				m[ i ]._activate.call( m[ i ], event );
			}

			m[ i ].offset = m[ i ].element.offset();
			m[ i ].proportions({ width: m[ i ].element[ 0 ].offsetWidth, height: m[ i ].element[ 0 ].offsetHeight });

		}

	},
	drop: function( draggable, event ) {

		var dropped = false;
		// Create a copy of the droppables in case the list changes during the drop (#9116)
		$.each( ( $.ui.ddmanager.droppables[ draggable.options.scope ] || [] ).slice(), function() {

			if ( !this.options ) {
				return;
			}
			if ( !this.options.disabled && this.visible && $.ui.intersect( draggable, this, this.options.tolerance, event ) ) {
				dropped = this._drop.call( this, event ) || dropped;
			}

			if ( !this.options.disabled && this.visible && this.accept.call( this.element[ 0 ], ( draggable.currentItem || draggable.element ) ) ) {
				this.isout = true;
				this.isover = false;
				this._deactivate.call( this, event );
			}

		});
		return dropped;

	},
	dragStart: function( draggable, event ) {
		// Listen for scrolling so that if the dragging causes scrolling the position of the droppables can be recalculated (see #5003)
		draggable.element.parentsUntil( "body" ).bind( "scroll.droppable", function() {
			if ( !draggable.options.refreshPositions ) {
				$.ui.ddmanager.prepareOffsets( draggable, event );
			}
		});
	},
	drag: function( draggable, event ) {

		// If you have a highly dynamic page, you might try this option. It renders positions every time you move the mouse.
		if ( draggable.options.refreshPositions ) {
			$.ui.ddmanager.prepareOffsets( draggable, event );
		}

		// Run through all droppables and check their positions based on specific tolerance options
		$.each( $.ui.ddmanager.droppables[ draggable.options.scope ] || [], function() {

			if ( this.options.disabled || this.greedyChild || !this.visible ) {
				return;
			}

			var parentInstance, scope, parent,
				intersects = $.ui.intersect( draggable, this, this.options.tolerance, event ),
				c = !intersects && this.isover ? "isout" : ( intersects && !this.isover ? "isover" : null );
			if ( !c ) {
				return;
			}

			if ( this.options.greedy ) {
				// find droppable parents with same scope
				scope = this.options.scope;
				parent = this.element.parents( ":data(ui-droppable)" ).filter(function() {
					return $( this ).droppable( "instance" ).options.scope === scope;
				});

				if ( parent.length ) {
					parentInstance = $( parent[ 0 ] ).droppable( "instance" );
					parentInstance.greedyChild = ( c === "isover" );
				}
			}

			// we just moved into a greedy child
			if ( parentInstance && c === "isover" ) {
				parentInstance.isover = false;
				parentInstance.isout = true;
				parentInstance._out.call( parentInstance, event );
			}

			this[ c ] = true;
			this[c === "isout" ? "isover" : "isout"] = false;
			this[c === "isover" ? "_over" : "_out"].call( this, event );

			// we just moved out of a greedy child
			if ( parentInstance && c === "isout" ) {
				parentInstance.isout = false;
				parentInstance.isover = true;
				parentInstance._over.call( parentInstance, event );
			}
		});

	},
	dragStop: function( draggable, event ) {
		draggable.element.parentsUntil( "body" ).unbind( "scroll.droppable" );
		// Call prepareOffsets one final time since IE does not fire return scroll events when overflow was caused by drag (see #5003)
		if ( !draggable.options.refreshPositions ) {
			$.ui.ddmanager.prepareOffsets( draggable, event );
		}
	}
};

var droppable = $.ui.droppable;


/*!
 * jQuery UI Resizable 1.11.4
 * http://jqueryui.com
 *
 * Copyright jQuery Foundation and other contributors
 * Released under the MIT license.
 * http://jquery.org/license
 *
 * http://api.jqueryui.com/resizable/
 */


$.widget("ui.resizable", $.ui.mouse, {
	version: "1.11.4",
	widgetEventPrefix: "resize",
	options: {
		alsoResize: false,
		animate: false,
		animateDuration: "slow",
		animateEasing: "swing",
		aspectRatio: false,
		autoHide: false,
		containment: false,
		ghost: false,
		grid: false,
		handles: "e,s,se",
		helper: false,
		maxHeight: null,
		maxWidth: null,
		minHeight: 10,
		minWidth: 10,
		// See #7960
		zIndex: 90,

		// callbacks
		resize: null,
		start: null,
		stop: null
	},

	_num: function( value ) {
		return parseInt( value, 10 ) || 0;
	},

	_isNumber: function( value ) {
		return !isNaN( parseInt( value, 10 ) );
	},

	_hasScroll: function( el, a ) {

		if ( $( el ).css( "overflow" ) === "hidden") {
			return false;
		}

		var scroll = ( a && a === "left" ) ? "scrollLeft" : "scrollTop",
			has = false;

		if ( el[ scroll ] > 0 ) {
			return true;
		}

		// TODO: determine which cases actually cause this to happen
		// if the element doesn't have the scroll set, see if it's possible to
		// set the scroll
		el[ scroll ] = 1;
		has = ( el[ scroll ] > 0 );
		el[ scroll ] = 0;
		return has;
	},

	_create: function() {

		var n, i, handle, axis, hname,
			that = this,
			o = this.options;
		this.element.addClass("ui-resizable");

		$.extend(this, {
			_aspectRatio: !!(o.aspectRatio),
			aspectRatio: o.aspectRatio,
			originalElement: this.element,
			_proportionallyResizeElements: [],
			_helper: o.helper || o.ghost || o.animate ? o.helper || "ui-resizable-helper" : null
		});

		// Wrap the element if it cannot hold child nodes
		if (this.element[0].nodeName.match(/^(canvas|textarea|input|select|button|img)$/i)) {

			this.element.wrap(
				$("<div class='ui-wrapper' style='overflow: hidden;'></div>").css({
					position: this.element.css("position"),
					width: this.element.outerWidth(),
					height: this.element.outerHeight(),
					top: this.element.css("top"),
					left: this.element.css("left")
				})
			);

			this.element = this.element.parent().data(
				"ui-resizable", this.element.resizable( "instance" )
			);

			this.elementIsWrapper = true;

			this.element.css({
				marginLeft: this.originalElement.css("marginLeft"),
				marginTop: this.originalElement.css("marginTop"),
				marginRight: this.originalElement.css("marginRight"),
				marginBottom: this.originalElement.css("marginBottom")
			});
			this.originalElement.css({
				marginLeft: 0,
				marginTop: 0,
				marginRight: 0,
				marginBottom: 0
			});
			// support: Safari
			// Prevent Safari textarea resize
			this.originalResizeStyle = this.originalElement.css("resize");
			this.originalElement.css("resize", "none");

			this._proportionallyResizeElements.push( this.originalElement.css({
				position: "static",
				zoom: 1,
				display: "block"
			}) );

			// support: IE9
			// avoid IE jump (hard set the margin)
			this.originalElement.css({ margin: this.originalElement.css("margin") });

			this._proportionallyResize();
		}

		this.handles = o.handles ||
			( !$(".ui-resizable-handle", this.element).length ?
				"e,s,se" : {
					n: ".ui-resizable-n",
					e: ".ui-resizable-e",
					s: ".ui-resizable-s",
					w: ".ui-resizable-w",
					se: ".ui-resizable-se",
					sw: ".ui-resizable-sw",
					ne: ".ui-resizable-ne",
					nw: ".ui-resizable-nw"
				} );

		this._handles = $();
		if ( this.handles.constructor === String ) {

			if ( this.handles === "all") {
				this.handles = "n,e,s,w,se,sw,ne,nw";
			}

			n = this.handles.split(",");
			this.handles = {};

			for (i = 0; i < n.length; i++) {

				handle = $.trim(n[i]);
				hname = "ui-resizable-" + handle;
				axis = $("<div class='ui-resizable-handle " + hname + "'></div>");

				axis.css({ zIndex: o.zIndex });

				// TODO : What's going on here?
				if ("se" === handle) {
					axis.addClass("ui-icon ui-icon-gripsmall-diagonal-se");
				}

				this.handles[handle] = ".ui-resizable-" + handle;
				this.element.append(axis);
			}

		}

		this._renderAxis = function(target) {

			var i, axis, padPos, padWrapper;

			target = target || this.element;

			for (i in this.handles) {

				if (this.handles[i].constructor === String) {
					this.handles[i] = this.element.children( this.handles[ i ] ).first().show();
				} else if ( this.handles[ i ].jquery || this.handles[ i ].nodeType ) {
					this.handles[ i ] = $( this.handles[ i ] );
					this._on( this.handles[ i ], { "mousedown": that._mouseDown });
				}

				if (this.elementIsWrapper && this.originalElement[0].nodeName.match(/^(textarea|input|select|button)$/i)) {

					axis = $(this.handles[i], this.element);

					padWrapper = /sw|ne|nw|se|n|s/.test(i) ? axis.outerHeight() : axis.outerWidth();

					padPos = [ "padding",
						/ne|nw|n/.test(i) ? "Top" :
						/se|sw|s/.test(i) ? "Bottom" :
						/^e$/.test(i) ? "Right" : "Left" ].join("");

					target.css(padPos, padWrapper);

					this._proportionallyResize();
				}

				this._handles = this._handles.add( this.handles[ i ] );
			}
		};

		// TODO: make renderAxis a prototype function
		this._renderAxis(this.element);

		this._handles = this._handles.add( this.element.find( ".ui-resizable-handle" ) );
		this._handles.disableSelection();

		this._handles.mouseover(function() {
			if (!that.resizing) {
				if (this.className) {
					axis = this.className.match(/ui-resizable-(se|sw|ne|nw|n|e|s|w)/i);
				}
				that.axis = axis && axis[1] ? axis[1] : "se";
			}
		});

		if (o.autoHide) {
			this._handles.hide();
			$(this.element)
				.addClass("ui-resizable-autohide")
				.mouseenter(function() {
					if (o.disabled) {
						return;
					}
					$(this).removeClass("ui-resizable-autohide");
					that._handles.show();
				})
				.mouseleave(function() {
					if (o.disabled) {
						return;
					}
					if (!that.resizing) {
						$(this).addClass("ui-resizable-autohide");
						that._handles.hide();
					}
				});
		}

		this._mouseInit();
	},

	_destroy: function() {

		this._mouseDestroy();

		var wrapper,
			_destroy = function(exp) {
				$(exp)
					.removeClass("ui-resizable ui-resizable-disabled ui-resizable-resizing")
					.removeData("resizable")
					.removeData("ui-resizable")
					.unbind(".resizable")
					.find(".ui-resizable-handle")
						.remove();
			};

		// TODO: Unwrap at same DOM position
		if (this.elementIsWrapper) {
			_destroy(this.element);
			wrapper = this.element;
			this.originalElement.css({
				position: wrapper.css("position"),
				width: wrapper.outerWidth(),
				height: wrapper.outerHeight(),
				top: wrapper.css("top"),
				left: wrapper.css("left")
			}).insertAfter( wrapper );
			wrapper.remove();
		}

		this.originalElement.css("resize", this.originalResizeStyle);
		_destroy(this.originalElement);

		return this;
	},

	_mouseCapture: function(event) {
		var i, handle,
			capture = false;

		for (i in this.handles) {
			handle = $(this.handles[i])[0];
			if (handle === event.target || $.contains(handle, event.target)) {
				capture = true;
			}
		}

		return !this.options.disabled && capture;
	},

	_mouseStart: function(event) {

		var curleft, curtop, cursor,
			o = this.options,
			el = this.element;

		this.resizing = true;

		this._renderProxy();

		curleft = this._num(this.helper.css("left"));
		curtop = this._num(this.helper.css("top"));

		if (o.containment) {
			curleft += $(o.containment).scrollLeft() || 0;
			curtop += $(o.containment).scrollTop() || 0;
		}

		this.offset = this.helper.offset();
		this.position = { left: curleft, top: curtop };

		this.size = this._helper ? {
				width: this.helper.width(),
				height: this.helper.height()
			} : {
				width: el.width(),
				height: el.height()
			};

		this.originalSize = this._helper ? {
				width: el.outerWidth(),
				height: el.outerHeight()
			} : {
				width: el.width(),
				height: el.height()
			};

		this.sizeDiff = {
			width: el.outerWidth() - el.width(),
			height: el.outerHeight() - el.height()
		};

		this.originalPosition = { left: curleft, top: curtop };
		this.originalMousePosition = { left: event.pageX, top: event.pageY };

		this.aspectRatio = (typeof o.aspectRatio === "number") ?
			o.aspectRatio :
			((this.originalSize.width / this.originalSize.height) || 1);

		cursor = $(".ui-resizable-" + this.axis).css("cursor");
		$("body").css("cursor", cursor === "auto" ? this.axis + "-resize" : cursor);

		el.addClass("ui-resizable-resizing");
		this._propagate("start", event);
		return true;
	},

	_mouseDrag: function(event) {

		var data, props,
			smp = this.originalMousePosition,
			a = this.axis,
			dx = (event.pageX - smp.left) || 0,
			dy = (event.pageY - smp.top) || 0,
			trigger = this._change[a];

		this._updatePrevProperties();

		if (!trigger) {
			return false;
		}

		data = trigger.apply(this, [ event, dx, dy ]);

		this._updateVirtualBoundaries(event.shiftKey);
		if (this._aspectRatio || event.shiftKey) {
			data = this._updateRatio(data, event);
		}

		data = this._respectSize(data, event);

		this._updateCache(data);

		this._propagate("resize", event);

		props = this._applyChanges();

		if ( !this._helper && this._proportionallyResizeElements.length ) {
			this._proportionallyResize();
		}

		if ( !$.isEmptyObject( props ) ) {
			this._updatePrevProperties();
			this._trigger( "resize", event, this.ui() );
			this._applyChanges();
		}

		return false;
	},

	_mouseStop: function(event) {

		this.resizing = false;
		var pr, ista, soffseth, soffsetw, s, left, top,
			o = this.options, that = this;

		if (this._helper) {

			pr = this._proportionallyResizeElements;
			ista = pr.length && (/textarea/i).test(pr[0].nodeName);
			soffseth = ista && this._hasScroll(pr[0], "left") ? 0 : that.sizeDiff.height;
			soffsetw = ista ? 0 : that.sizeDiff.width;

			s = {
				width: (that.helper.width()  - soffsetw),
				height: (that.helper.height() - soffseth)
			};
			left = (parseInt(that.element.css("left"), 10) +
				(that.position.left - that.originalPosition.left)) || null;
			top = (parseInt(that.element.css("top"), 10) +
				(that.position.top - that.originalPosition.top)) || null;

			if (!o.animate) {
				this.element.css($.extend(s, { top: top, left: left }));
			}

			that.helper.height(that.size.height);
			that.helper.width(that.size.width);

			if (this._helper && !o.animate) {
				this._proportionallyResize();
			}
		}

		$("body").css("cursor", "auto");

		this.element.removeClass("ui-resizable-resizing");

		this._propagate("stop", event);

		if (this._helper) {
			this.helper.remove();
		}

		return false;

	},

	_updatePrevProperties: function() {
		this.prevPosition = {
			top: this.position.top,
			left: this.position.left
		};
		this.prevSize = {
			width: this.size.width,
			height: this.size.height
		};
	},

	_applyChanges: function() {
		var props = {};

		if ( this.position.top !== this.prevPosition.top ) {
			props.top = this.position.top + "px";
		}
		if ( this.position.left !== this.prevPosition.left ) {
			props.left = this.position.left + "px";
		}
		if ( this.size.width !== this.prevSize.width ) {
			props.width = this.size.width + "px";
		}
		if ( this.size.height !== this.prevSize.height ) {
			props.height = this.size.height + "px";
		}

		this.helper.css( props );

		return props;
	},

	_updateVirtualBoundaries: function(forceAspectRatio) {
		var pMinWidth, pMaxWidth, pMinHeight, pMaxHeight, b,
			o = this.options;

		b = {
			minWidth: this._isNumber(o.minWidth) ? o.minWidth : 0,
			maxWidth: this._isNumber(o.maxWidth) ? o.maxWidth : Infinity,
			minHeight: this._isNumber(o.minHeight) ? o.minHeight : 0,
			maxHeight: this._isNumber(o.maxHeight) ? o.maxHeight : Infinity
		};

		if (this._aspectRatio || forceAspectRatio) {
			pMinWidth = b.minHeight * this.aspectRatio;
			pMinHeight = b.minWidth / this.aspectRatio;
			pMaxWidth = b.maxHeight * this.aspectRatio;
			pMaxHeight = b.maxWidth / this.aspectRatio;

			if (pMinWidth > b.minWidth) {
				b.minWidth = pMinWidth;
			}
			if (pMinHeight > b.minHeight) {
				b.minHeight = pMinHeight;
			}
			if (pMaxWidth < b.maxWidth) {
				b.maxWidth = pMaxWidth;
			}
			if (pMaxHeight < b.maxHeight) {
				b.maxHeight = pMaxHeight;
			}
		}
		this._vBoundaries = b;
	},

	_updateCache: function(data) {
		this.offset = this.helper.offset();
		if (this._isNumber(data.left)) {
			this.position.left = data.left;
		}
		if (this._isNumber(data.top)) {
			this.position.top = data.top;
		}
		if (this._isNumber(data.height)) {
			this.size.height = data.height;
		}
		if (this._isNumber(data.width)) {
			this.size.width = data.width;
		}
	},

	_updateRatio: function( data ) {

		var cpos = this.position,
			csize = this.size,
			a = this.axis;

		if (this._isNumber(data.height)) {
			data.width = (data.height * this.aspectRatio);
		} else if (this._isNumber(data.width)) {
			data.height = (data.width / this.aspectRatio);
		}

		if (a === "sw") {
			data.left = cpos.left + (csize.width - data.width);
			data.top = null;
		}
		if (a === "nw") {
			data.top = cpos.top + (csize.height - data.height);
			data.left = cpos.left + (csize.width - data.width);
		}

		return data;
	},

	_respectSize: function( data ) {

		var o = this._vBoundaries,
			a = this.axis,
			ismaxw = this._isNumber(data.width) && o.maxWidth && (o.maxWidth < data.width),
			ismaxh = this._isNumber(data.height) && o.maxHeight && (o.maxHeight < data.height),
			isminw = this._isNumber(data.width) && o.minWidth && (o.minWidth > data.width),
			isminh = this._isNumber(data.height) && o.minHeight && (o.minHeight > data.height),
			dw = this.originalPosition.left + this.originalSize.width,
			dh = this.position.top + this.size.height,
			cw = /sw|nw|w/.test(a), ch = /nw|ne|n/.test(a);
		if (isminw) {
			data.width = o.minWidth;
		}
		if (isminh) {
			data.height = o.minHeight;
		}
		if (ismaxw) {
			data.width = o.maxWidth;
		}
		if (ismaxh) {
			data.height = o.maxHeight;
		}

		if (isminw && cw) {
			data.left = dw - o.minWidth;
		}
		if (ismaxw && cw) {
			data.left = dw - o.maxWidth;
		}
		if (isminh && ch) {
			data.top = dh - o.minHeight;
		}
		if (ismaxh && ch) {
			data.top = dh - o.maxHeight;
		}

		// Fixing jump error on top/left - bug #2330
		if (!data.width && !data.height && !data.left && data.top) {
			data.top = null;
		} else if (!data.width && !data.height && !data.top && data.left) {
			data.left = null;
		}

		return data;
	},

	_getPaddingPlusBorderDimensions: function( element ) {
		var i = 0,
			widths = [],
			borders = [
				element.css( "borderTopWidth" ),
				element.css( "borderRightWidth" ),
				element.css( "borderBottomWidth" ),
				element.css( "borderLeftWidth" )
			],
			paddings = [
				element.css( "paddingTop" ),
				element.css( "paddingRight" ),
				element.css( "paddingBottom" ),
				element.css( "paddingLeft" )
			];

		for ( ; i < 4; i++ ) {
			widths[ i ] = ( parseInt( borders[ i ], 10 ) || 0 );
			widths[ i ] += ( parseInt( paddings[ i ], 10 ) || 0 );
		}

		return {
			height: widths[ 0 ] + widths[ 2 ],
			width: widths[ 1 ] + widths[ 3 ]
		};
	},

	_proportionallyResize: function() {

		if (!this._proportionallyResizeElements.length) {
			return;
		}

		var prel,
			i = 0,
			element = this.helper || this.element;

		for ( ; i < this._proportionallyResizeElements.length; i++) {

			prel = this._proportionallyResizeElements[i];

			// TODO: Seems like a bug to cache this.outerDimensions
			// considering that we are in a loop.
			if (!this.outerDimensions) {
				this.outerDimensions = this._getPaddingPlusBorderDimensions( prel );
			}

			prel.css({
				height: (element.height() - this.outerDimensions.height) || 0,
				width: (element.width() - this.outerDimensions.width) || 0
			});

		}

	},

	_renderProxy: function() {

		var el = this.element, o = this.options;
		this.elementOffset = el.offset();

		if (this._helper) {

			this.helper = this.helper || $("<div style='overflow:hidden;'></div>");

			this.helper.addClass(this._helper).css({
				width: this.element.outerWidth() - 1,
				height: this.element.outerHeight() - 1,
				position: "absolute",
				left: this.elementOffset.left + "px",
				top: this.elementOffset.top + "px",
				zIndex: ++o.zIndex //TODO: Don't modify option
			});

			this.helper
				.appendTo("body")
				.disableSelection();

		} else {
			this.helper = this.element;
		}

	},

	_change: {
		e: function(event, dx) {
			return { width: this.originalSize.width + dx };
		},
		w: function(event, dx) {
			var cs = this.originalSize, sp = this.originalPosition;
			return { left: sp.left + dx, width: cs.width - dx };
		},
		n: function(event, dx, dy) {
			var cs = this.originalSize, sp = this.originalPosition;
			return { top: sp.top + dy, height: cs.height - dy };
		},
		s: function(event, dx, dy) {
			return { height: this.originalSize.height + dy };
		},
		se: function(event, dx, dy) {
			return $.extend(this._change.s.apply(this, arguments),
				this._change.e.apply(this, [ event, dx, dy ]));
		},
		sw: function(event, dx, dy) {
			return $.extend(this._change.s.apply(this, arguments),
				this._change.w.apply(this, [ event, dx, dy ]));
		},
		ne: function(event, dx, dy) {
			return $.extend(this._change.n.apply(this, arguments),
				this._change.e.apply(this, [ event, dx, dy ]));
		},
		nw: function(event, dx, dy) {
			return $.extend(this._change.n.apply(this, arguments),
				this._change.w.apply(this, [ event, dx, dy ]));
		}
	},

	_propagate: function(n, event) {
		$.ui.plugin.call(this, n, [ event, this.ui() ]);
		(n !== "resize" && this._trigger(n, event, this.ui()));
	},

	plugins: {},

	ui: function() {
		return {
			originalElement: this.originalElement,
			element: this.element,
			helper: this.helper,
			position: this.position,
			size: this.size,
			originalSize: this.originalSize,
			originalPosition: this.originalPosition
		};
	}

});

/*
 * Resizable Extensions
 */

$.ui.plugin.add("resizable", "animate", {

	stop: function( event ) {
		var that = $(this).resizable( "instance" ),
			o = that.options,
			pr = that._proportionallyResizeElements,
			ista = pr.length && (/textarea/i).test(pr[0].nodeName),
			soffseth = ista && that._hasScroll(pr[0], "left") ? 0 : that.sizeDiff.height,
			soffsetw = ista ? 0 : that.sizeDiff.width,
			style = { width: (that.size.width - soffsetw), height: (that.size.height - soffseth) },
			left = (parseInt(that.element.css("left"), 10) +
				(that.position.left - that.originalPosition.left)) || null,
			top = (parseInt(that.element.css("top"), 10) +
				(that.position.top - that.originalPosition.top)) || null;

		that.element.animate(
			$.extend(style, top && left ? { top: top, left: left } : {}), {
				duration: o.animateDuration,
				easing: o.animateEasing,
				step: function() {

					var data = {
						width: parseInt(that.element.css("width"), 10),
						height: parseInt(that.element.css("height"), 10),
						top: parseInt(that.element.css("top"), 10),
						left: parseInt(that.element.css("left"), 10)
					};

					if (pr && pr.length) {
						$(pr[0]).css({ width: data.width, height: data.height });
					}

					// propagating resize, and updating values for each animation step
					that._updateCache(data);
					that._propagate("resize", event);

				}
			}
		);
	}

});

$.ui.plugin.add( "resizable", "containment", {

	start: function() {
		var element, p, co, ch, cw, width, height,
			that = $( this ).resizable( "instance" ),
			o = that.options,
			el = that.element,
			oc = o.containment,
			ce = ( oc instanceof $ ) ? oc.get( 0 ) : ( /parent/.test( oc ) ) ? el.parent().get( 0 ) : oc;

		if ( !ce ) {
			return;
		}

		that.containerElement = $( ce );

		if ( /document/.test( oc ) || oc === document ) {
			that.containerOffset = {
				left: 0,
				top: 0
			};
			that.containerPosition = {
				left: 0,
				top: 0
			};

			that.parentData = {
				element: $( document ),
				left: 0,
				top: 0,
				width: $( document ).width(),
				height: $( document ).height() || document.body.parentNode.scrollHeight
			};
		} else {
			element = $( ce );
			p = [];
			$([ "Top", "Right", "Left", "Bottom" ]).each(function( i, name ) {
				p[ i ] = that._num( element.css( "padding" + name ) );
			});

			that.containerOffset = element.offset();
			that.containerPosition = element.position();
			that.containerSize = {
				height: ( element.innerHeight() - p[ 3 ] ),
				width: ( element.innerWidth() - p[ 1 ] )
			};

			co = that.containerOffset;
			ch = that.containerSize.height;
			cw = that.containerSize.width;
			width = ( that._hasScroll ( ce, "left" ) ? ce.scrollWidth : cw );
			height = ( that._hasScroll ( ce ) ? ce.scrollHeight : ch ) ;

			that.parentData = {
				element: ce,
				left: co.left,
				top: co.top,
				width: width,
				height: height
			};
		}
	},

	resize: function( event ) {
		var woset, hoset, isParent, isOffsetRelative,
			that = $( this ).resizable( "instance" ),
			o = that.options,
			co = that.containerOffset,
			cp = that.position,
			pRatio = that._aspectRatio || event.shiftKey,
			cop = {
				top: 0,
				left: 0
			},
			ce = that.containerElement,
			continueResize = true;

		if ( ce[ 0 ] !== document && ( /static/ ).test( ce.css( "position" ) ) ) {
			cop = co;
		}

		if ( cp.left < ( that._helper ? co.left : 0 ) ) {
			that.size.width = that.size.width +
				( that._helper ?
					( that.position.left - co.left ) :
					( that.position.left - cop.left ) );

			if ( pRatio ) {
				that.size.height = that.size.width / that.aspectRatio;
				continueResize = false;
			}
			that.position.left = o.helper ? co.left : 0;
		}

		if ( cp.top < ( that._helper ? co.top : 0 ) ) {
			that.size.height = that.size.height +
				( that._helper ?
					( that.position.top - co.top ) :
					that.position.top );

			if ( pRatio ) {
				that.size.width = that.size.height * that.aspectRatio;
				continueResize = false;
			}
			that.position.top = that._helper ? co.top : 0;
		}

		isParent = that.containerElement.get( 0 ) === that.element.parent().get( 0 );
		isOffsetRelative = /relative|absolute/.test( that.containerElement.css( "position" ) );

		if ( isParent && isOffsetRelative ) {
			that.offset.left = that.parentData.left + that.position.left;
			that.offset.top = that.parentData.top + that.position.top;
		} else {
			that.offset.left = that.element.offset().left;
			that.offset.top = that.element.offset().top;
		}

		woset = Math.abs( that.sizeDiff.width +
			(that._helper ?
				that.offset.left - cop.left :
				(that.offset.left - co.left)) );

		hoset = Math.abs( that.sizeDiff.height +
			(that._helper ?
				that.offset.top - cop.top :
				(that.offset.top - co.top)) );

		if ( woset + that.size.width >= that.parentData.width ) {
			that.size.width = that.parentData.width - woset;
			if ( pRatio ) {
				that.size.height = that.size.width / that.aspectRatio;
				continueResize = false;
			}
		}

		if ( hoset + that.size.height >= that.parentData.height ) {
			that.size.height = that.parentData.height - hoset;
			if ( pRatio ) {
				that.size.width = that.size.height * that.aspectRatio;
				continueResize = false;
			}
		}

		if ( !continueResize ) {
			that.position.left = that.prevPosition.left;
			that.position.top = that.prevPosition.top;
			that.size.width = that.prevSize.width;
			that.size.height = that.prevSize.height;
		}
	},

	stop: function() {
		var that = $( this ).resizable( "instance" ),
			o = that.options,
			co = that.containerOffset,
			cop = that.containerPosition,
			ce = that.containerElement,
			helper = $( that.helper ),
			ho = helper.offset(),
			w = helper.outerWidth() - that.sizeDiff.width,
			h = helper.outerHeight() - that.sizeDiff.height;

		if ( that._helper && !o.animate && ( /relative/ ).test( ce.css( "position" ) ) ) {
			$( this ).css({
				left: ho.left - cop.left - co.left,
				width: w,
				height: h
			});
		}

		if ( that._helper && !o.animate && ( /static/ ).test( ce.css( "position" ) ) ) {
			$( this ).css({
				left: ho.left - cop.left - co.left,
				width: w,
				height: h
			});
		}
	}
});

$.ui.plugin.add("resizable", "alsoResize", {

	start: function() {
		var that = $(this).resizable( "instance" ),
			o = that.options;

		$(o.alsoResize).each(function() {
			var el = $(this);
			el.data("ui-resizable-alsoresize", {
				width: parseInt(el.width(), 10), height: parseInt(el.height(), 10),
				left: parseInt(el.css("left"), 10), top: parseInt(el.css("top"), 10)
			});
		});
	},

	resize: function(event, ui) {
		var that = $(this).resizable( "instance" ),
			o = that.options,
			os = that.originalSize,
			op = that.originalPosition,
			delta = {
				height: (that.size.height - os.height) || 0,
				width: (that.size.width - os.width) || 0,
				top: (that.position.top - op.top) || 0,
				left: (that.position.left - op.left) || 0
			};

			$(o.alsoResize).each(function() {
				var el = $(this), start = $(this).data("ui-resizable-alsoresize"), style = {},
					css = el.parents(ui.originalElement[0]).length ?
							[ "width", "height" ] :
							[ "width", "height", "top", "left" ];

				$.each(css, function(i, prop) {
					var sum = (start[prop] || 0) + (delta[prop] || 0);
					if (sum && sum >= 0) {
						style[prop] = sum || null;
					}
				});

				el.css(style);
			});
	},

	stop: function() {
		$(this).removeData("resizable-alsoresize");
	}
});

$.ui.plugin.add("resizable", "ghost", {

	start: function() {

		var that = $(this).resizable( "instance" ), o = that.options, cs = that.size;

		that.ghost = that.originalElement.clone();
		that.ghost
			.css({
				opacity: 0.25,
				display: "block",
				position: "relative",
				height: cs.height,
				width: cs.width,
				margin: 0,
				left: 0,
				top: 0
			})
			.addClass("ui-resizable-ghost")
			.addClass(typeof o.ghost === "string" ? o.ghost : "");

		that.ghost.appendTo(that.helper);

	},

	resize: function() {
		var that = $(this).resizable( "instance" );
		if (that.ghost) {
			that.ghost.css({
				position: "relative",
				height: that.size.height,
				width: that.size.width
			});
		}
	},

	stop: function() {
		var that = $(this).resizable( "instance" );
		if (that.ghost && that.helper) {
			that.helper.get(0).removeChild(that.ghost.get(0));
		}
	}

});

$.ui.plugin.add("resizable", "grid", {

	resize: function() {
		var outerDimensions,
			that = $(this).resizable( "instance" ),
			o = that.options,
			cs = that.size,
			os = that.originalSize,
			op = that.originalPosition,
			a = that.axis,
			grid = typeof o.grid === "number" ? [ o.grid, o.grid ] : o.grid,
			gridX = (grid[0] || 1),
			gridY = (grid[1] || 1),
			ox = Math.round((cs.width - os.width) / gridX) * gridX,
			oy = Math.round((cs.height - os.height) / gridY) * gridY,
			newWidth = os.width + ox,
			newHeight = os.height + oy,
			isMaxWidth = o.maxWidth && (o.maxWidth < newWidth),
			isMaxHeight = o.maxHeight && (o.maxHeight < newHeight),
			isMinWidth = o.minWidth && (o.minWidth > newWidth),
			isMinHeight = o.minHeight && (o.minHeight > newHeight);

		o.grid = grid;

		if (isMinWidth) {
			newWidth += gridX;
		}
		if (isMinHeight) {
			newHeight += gridY;
		}
		if (isMaxWidth) {
			newWidth -= gridX;
		}
		if (isMaxHeight) {
			newHeight -= gridY;
		}

		if (/^(se|s|e)$/.test(a)) {
			that.size.width = newWidth;
			that.size.height = newHeight;
		} else if (/^(ne)$/.test(a)) {
			that.size.width = newWidth;
			that.size.height = newHeight;
			that.position.top = op.top - oy;
		} else if (/^(sw)$/.test(a)) {
			that.size.width = newWidth;
			that.size.height = newHeight;
			that.position.left = op.left - ox;
		} else {
			if ( newHeight - gridY <= 0 || newWidth - gridX <= 0) {
				outerDimensions = that._getPaddingPlusBorderDimensions( this );
			}

			if ( newHeight - gridY > 0 ) {
				that.size.height = newHeight;
				that.position.top = op.top - oy;
			} else {
				newHeight = gridY - outerDimensions.height;
				that.size.height = newHeight;
				that.position.top = op.top + os.height - newHeight;
			}
			if ( newWidth - gridX > 0 ) {
				that.size.width = newWidth;
				that.position.left = op.left - ox;
			} else {
				newWidth = gridX - outerDimensions.width;
				that.size.width = newWidth;
				that.position.left = op.left + os.width - newWidth;
			}
		}
	}

});

var resizable = $.ui.resizable;


/*!
 * jQuery UI Selectable 1.11.4
 * http://jqueryui.com
 *
 * Copyright jQuery Foundation and other contributors
 * Released under the MIT license.
 * http://jquery.org/license
 *
 * http://api.jqueryui.com/selectable/
 */


var selectable = $.widget("ui.selectable", $.ui.mouse, {
	version: "1.11.4",
	options: {
		appendTo: "body",
		autoRefresh: true,
		distance: 0,
		filter: "*",
		tolerance: "touch",

		// callbacks
		selected: null,
		selecting: null,
		start: null,
		stop: null,
		unselected: null,
		unselecting: null
	},
	_create: function() {
		var selectees,
			that = this;

		this.element.addClass("ui-selectable");

		this.dragged = false;

		// cache selectee children based on filter
		this.refresh = function() {
			selectees = $(that.options.filter, that.element[0]);
			selectees.addClass("ui-selectee");
			selectees.each(function() {
				var $this = $(this),
					pos = $this.offset();
				$.data(this, "selectable-item", {
					element: this,
					$element: $this,
					left: pos.left,
					top: pos.top,
					right: pos.left + $this.outerWidth(),
					bottom: pos.top + $this.outerHeight(),
					startselected: false,
					selected: $this.hasClass("ui-selected"),
					selecting: $this.hasClass("ui-selecting"),
					unselecting: $this.hasClass("ui-unselecting")
				});
			});
		};
		this.refresh();

		this.selectees = selectees.addClass("ui-selectee");

		this._mouseInit();

		this.helper = $("<div class='ui-selectable-helper'></div>");
	},

	_destroy: function() {
		this.selectees
			.removeClass("ui-selectee")
			.removeData("selectable-item");
		this.element
			.removeClass("ui-selectable ui-selectable-disabled");
		this._mouseDestroy();
	},

	_mouseStart: function(event) {
		var that = this,
			options = this.options;

		this.opos = [ event.pageX, event.pageY ];

		if (this.options.disabled) {
			return;
		}

		this.selectees = $(options.filter, this.element[0]);

		this._trigger("start", event);

		$(options.appendTo).append(this.helper);
		// position helper (lasso)
		this.helper.css({
			"left": event.pageX,
			"top": event.pageY,
			"width": 0,
			"height": 0
		});

		if (options.autoRefresh) {
			this.refresh();
		}

		this.selectees.filter(".ui-selected").each(function() {
			var selectee = $.data(this, "selectable-item");
			selectee.startselected = true;
			if (!event.metaKey && !event.ctrlKey) {
				selectee.$element.removeClass("ui-selected");
				selectee.selected = false;
				selectee.$element.addClass("ui-unselecting");
				selectee.unselecting = true;
				// selectable UNSELECTING callback
				that._trigger("unselecting", event, {
					unselecting: selectee.element
				});
			}
		});

		$(event.target).parents().addBack().each(function() {
			var doSelect,
				selectee = $.data(this, "selectable-item");
			if (selectee) {
				doSelect = (!event.metaKey && !event.ctrlKey) || !selectee.$element.hasClass("ui-selected");
				selectee.$element
					.removeClass(doSelect ? "ui-unselecting" : "ui-selected")
					.addClass(doSelect ? "ui-selecting" : "ui-unselecting");
				selectee.unselecting = !doSelect;
				selectee.selecting = doSelect;
				selectee.selected = doSelect;
				// selectable (UN)SELECTING callback
				if (doSelect) {
					that._trigger("selecting", event, {
						selecting: selectee.element
					});
				} else {
					that._trigger("unselecting", event, {
						unselecting: selectee.element
					});
				}
				return false;
			}
		});

	},

	_mouseDrag: function(event) {

		this.dragged = true;

		if (this.options.disabled) {
			return;
		}

		var tmp,
			that = this,
			options = this.options,
			x1 = this.opos[0],
			y1 = this.opos[1],
			x2 = event.pageX,
			y2 = event.pageY;

		if (x1 > x2) { tmp = x2; x2 = x1; x1 = tmp; }
		if (y1 > y2) { tmp = y2; y2 = y1; y1 = tmp; }
		this.helper.css({ left: x1, top: y1, width: x2 - x1, height: y2 - y1 });

		this.selectees.each(function() {
			var selectee = $.data(this, "selectable-item"),
				hit = false;

			//prevent helper from being selected if appendTo: selectable
			if (!selectee || selectee.element === that.element[0]) {
				return;
			}

			if (options.tolerance === "touch") {
				hit = ( !(selectee.left > x2 || selectee.right < x1 || selectee.top > y2 || selectee.bottom < y1) );
			} else if (options.tolerance === "fit") {
				hit = (selectee.left > x1 && selectee.right < x2 && selectee.top > y1 && selectee.bottom < y2);
			}

			if (hit) {
				// SELECT
				if (selectee.selected) {
					selectee.$element.removeClass("ui-selected");
					selectee.selected = false;
				}
				if (selectee.unselecting) {
					selectee.$element.removeClass("ui-unselecting");
					selectee.unselecting = false;
				}
				if (!selectee.selecting) {
					selectee.$element.addClass("ui-selecting");
					selectee.selecting = true;
					// selectable SELECTING callback
					that._trigger("selecting", event, {
						selecting: selectee.element
					});
				}
			} else {
				// UNSELECT
				if (selectee.selecting) {
					if ((event.metaKey || event.ctrlKey) && selectee.startselected) {
						selectee.$element.removeClass("ui-selecting");
						selectee.selecting = false;
						selectee.$element.addClass("ui-selected");
						selectee.selected = true;
					} else {
						selectee.$element.removeClass("ui-selecting");
						selectee.selecting = false;
						if (selectee.startselected) {
							selectee.$element.addClass("ui-unselecting");
							selectee.unselecting = true;
						}
						// selectable UNSELECTING callback
						that._trigger("unselecting", event, {
							unselecting: selectee.element
						});
					}
				}
				if (selectee.selected) {
					if (!event.metaKey && !event.ctrlKey && !selectee.startselected) {
						selectee.$element.removeClass("ui-selected");
						selectee.selected = false;

						selectee.$element.addClass("ui-unselecting");
						selectee.unselecting = true;
						// selectable UNSELECTING callback
						that._trigger("unselecting", event, {
							unselecting: selectee.element
						});
					}
				}
			}
		});

		return false;
	},

	_mouseStop: function(event) {
		var that = this;

		this.dragged = false;

		$(".ui-unselecting", this.element[0]).each(function() {
			var selectee = $.data(this, "selectable-item");
			selectee.$element.removeClass("ui-unselecting");
			selectee.unselecting = false;
			selectee.startselected = false;
			that._trigger("unselected", event, {
				unselected: selectee.element
			});
		});
		$(".ui-selecting", this.element[0]).each(function() {
			var selectee = $.data(this, "selectable-item");
			selectee.$element.removeClass("ui-selecting").addClass("ui-selected");
			selectee.selecting = false;
			selectee.selected = true;
			selectee.startselected = true;
			that._trigger("selected", event, {
				selected: selectee.element
			});
		});
		this._trigger("stop", event);

		this.helper.remove();

		return false;
	}

});


/*!
 * jQuery UI Sortable 1.11.4
 * http://jqueryui.com
 *
 * Copyright jQuery Foundation and other contributors
 * Released under the MIT license.
 * http://jquery.org/license
 *
 * http://api.jqueryui.com/sortable/
 */


var sortable = $.widget("ui.sortable", $.ui.mouse, {
	version: "1.11.4",
	widgetEventPrefix: "sort",
	ready: false,
	options: {
		appendTo: "parent",
		axis: false,
		connectWith: false,
		containment: false,
		cursor: "auto",
		cursorAt: false,
		dropOnEmpty: true,
		forcePlaceholderSize: false,
		forceHelperSize: false,
		grid: false,
		handle: false,
		helper: "original",
		items: "> *",
		opacity: false,
		placeholder: false,
		revert: false,
		scroll: true,
		scrollSensitivity: 20,
		scrollSpeed: 20,
		scope: "default",
		tolerance: "intersect",
		zIndex: 1000,

		// callbacks
		activate: null,
		beforeStop: null,
		change: null,
		deactivate: null,
		out: null,
		over: null,
		receive: null,
		remove: null,
		sort: null,
		start: null,
		stop: null,
		update: null
	},

	_isOverAxis: function( x, reference, size ) {
		return ( x >= reference ) && ( x < ( reference + size ) );
	},

	_isFloating: function( item ) {
		return (/left|right/).test(item.css("float")) || (/inline|table-cell/).test(item.css("display"));
	},

	_create: function() {
		this.containerCache = {};
		this.element.addClass("ui-sortable");

		//Get the items
		this.refresh();

		//Let's determine the parent's offset
		this.offset = this.element.offset();

		//Initialize mouse events for interaction
		this._mouseInit();

		this._setHandleClassName();

		//We're ready to go
		this.ready = true;

	},

	_setOption: function( key, value ) {
		this._super( key, value );

		if ( key === "handle" ) {
			this._setHandleClassName();
		}
	},

	_setHandleClassName: function() {
		this.element.find( ".ui-sortable-handle" ).removeClass( "ui-sortable-handle" );
		$.each( this.items, function() {
			( this.instance.options.handle ?
				this.item.find( this.instance.options.handle ) : this.item )
				.addClass( "ui-sortable-handle" );
		});
	},

	_destroy: function() {
		this.element
			.removeClass( "ui-sortable ui-sortable-disabled" )
			.find( ".ui-sortable-handle" )
				.removeClass( "ui-sortable-handle" );
		this._mouseDestroy();

		for ( var i = this.items.length - 1; i >= 0; i-- ) {
			this.items[i].item.removeData(this.widgetName + "-item");
		}

		return this;
	},

	_mouseCapture: function(event, overrideHandle) {
		var currentItem = null,
			validHandle = false,
			that = this;

		if (this.reverting) {
			return false;
		}

		if(this.options.disabled || this.options.type === "static") {
			return false;
		}

		//We have to refresh the items data once first
		this._refreshItems(event);

		//Find out if the clicked node (or one of its parents) is a actual item in this.items
		$(event.target).parents().each(function() {
			if($.data(this, that.widgetName + "-item") === that) {
				currentItem = $(this);
				return false;
			}
		});
		if($.data(event.target, that.widgetName + "-item") === that) {
			currentItem = $(event.target);
		}

		if(!currentItem) {
			return false;
		}
		if(this.options.handle && !overrideHandle) {
			$(this.options.handle, currentItem).find("*").addBack().each(function() {
				if(this === event.target) {
					validHandle = true;
				}
			});
			if(!validHandle) {
				return false;
			}
		}

		this.currentItem = currentItem;
		this._removeCurrentsFromItems();
		return true;

	},

	_mouseStart: function(event, overrideHandle, noActivation) {

		var i, body,
			o = this.options;

		this.currentContainer = this;

		//We only need to call refreshPositions, because the refreshItems call has been moved to mouseCapture
		this.refreshPositions();

		//Create and append the visible helper
		this.helper = this._createHelper(event);

		//Cache the helper size
		this._cacheHelperProportions();

		/*
		 * - Position generation -
		 * This block generates everything position related - it's the core of draggables.
		 */

		//Cache the margins of the original element
		this._cacheMargins();

		//Get the next scrolling parent
		this.scrollParent = this.helper.scrollParent();

		//The element's absolute position on the page minus margins
		this.offset = this.currentItem.offset();
		this.offset = {
			top: this.offset.top - this.margins.top,
			left: this.offset.left - this.margins.left
		};

		$.extend(this.offset, {
			click: { //Where the click happened, relative to the element
				left: event.pageX - this.offset.left,
				top: event.pageY - this.offset.top
			},
			parent: this._getParentOffset(),
			relative: this._getRelativeOffset() //This is a relative to absolute position minus the actual position calculation - only used for relative positioned helper
		});

		// Only after we got the offset, we can change the helper's position to absolute
		// TODO: Still need to figure out a way to make relative sorting possible
		this.helper.css("position", "absolute");
		this.cssPosition = this.helper.css("position");

		//Generate the original position
		this.originalPosition = this._generatePosition(event);
		this.originalPageX = event.pageX;
		this.originalPageY = event.pageY;

		//Adjust the mouse offset relative to the helper if "cursorAt" is supplied
		(o.cursorAt && this._adjustOffsetFromHelper(o.cursorAt));

		//Cache the former DOM position
		this.domPosition = { prev: this.currentItem.prev()[0], parent: this.currentItem.parent()[0] };

		//If the helper is not the original, hide the original so it's not playing any role during the drag, won't cause anything bad this way
		if(this.helper[0] !== this.currentItem[0]) {
			this.currentItem.hide();
		}

		//Create the placeholder
		this._createPlaceholder();

		//Set a containment if given in the options
		if(o.containment) {
			this._setContainment();
		}

		if( o.cursor && o.cursor !== "auto" ) { // cursor option
			body = this.document.find( "body" );

			// support: IE
			this.storedCursor = body.css( "cursor" );
			body.css( "cursor", o.cursor );

			this.storedStylesheet = $( "<style>*{ cursor: "+o.cursor+" !important; }</style>" ).appendTo( body );
		}

		if(o.opacity) { // opacity option
			if (this.helper.css("opacity")) {
				this._storedOpacity = this.helper.css("opacity");
			}
			this.helper.css("opacity", o.opacity);
		}

		if(o.zIndex) { // zIndex option
			if (this.helper.css("zIndex")) {
				this._storedZIndex = this.helper.css("zIndex");
			}
			this.helper.css("zIndex", o.zIndex);
		}

		//Prepare scrolling
		if(this.scrollParent[0] !== this.document[0] && this.scrollParent[0].tagName !== "HTML") {
			this.overflowOffset = this.scrollParent.offset();
		}

		//Call callbacks
		this._trigger("start", event, this._uiHash());

		//Recache the helper size
		if(!this._preserveHelperProportions) {
			this._cacheHelperProportions();
		}


		//Post "activate" events to possible containers
		if( !noActivation ) {
			for ( i = this.containers.length - 1; i >= 0; i-- ) {
				this.containers[ i ]._trigger( "activate", event, this._uiHash( this ) );
			}
		}

		//Prepare possible droppables
		if($.ui.ddmanager) {
			$.ui.ddmanager.current = this;
		}

		if ($.ui.ddmanager && !o.dropBehaviour) {
			$.ui.ddmanager.prepareOffsets(this, event);
		}

		this.dragging = true;

		this.helper.addClass("ui-sortable-helper");
		this._mouseDrag(event); //Execute the drag once - this causes the helper not to be visible before getting its correct position
		return true;

	},

	_mouseDrag: function(event) {
		var i, item, itemElement, intersection,
			o = this.options,
			scrolled = false;

		//Compute the helpers position
		this.position = this._generatePosition(event);
		this.positionAbs = this._convertPositionTo("absolute");

		if (!this.lastPositionAbs) {
			this.lastPositionAbs = this.positionAbs;
		}

		//Do scrolling
		if(this.options.scroll) {
			if(this.scrollParent[0] !== this.document[0] && this.scrollParent[0].tagName !== "HTML") {

				if((this.overflowOffset.top + this.scrollParent[0].offsetHeight) - event.pageY < o.scrollSensitivity) {
					this.scrollParent[0].scrollTop = scrolled = this.scrollParent[0].scrollTop + o.scrollSpeed;
				} else if(event.pageY - this.overflowOffset.top < o.scrollSensitivity) {
					this.scrollParent[0].scrollTop = scrolled = this.scrollParent[0].scrollTop - o.scrollSpeed;
				}

				if((this.overflowOffset.left + this.scrollParent[0].offsetWidth) - event.pageX < o.scrollSensitivity) {
					this.scrollParent[0].scrollLeft = scrolled = this.scrollParent[0].scrollLeft + o.scrollSpeed;
				} else if(event.pageX - this.overflowOffset.left < o.scrollSensitivity) {
					this.scrollParent[0].scrollLeft = scrolled = this.scrollParent[0].scrollLeft - o.scrollSpeed;
				}

			} else {

				if(event.pageY - this.document.scrollTop() < o.scrollSensitivity) {
					scrolled = this.document.scrollTop(this.document.scrollTop() - o.scrollSpeed);
				} else if(this.window.height() - (event.pageY - this.document.scrollTop()) < o.scrollSensitivity) {
					scrolled = this.document.scrollTop(this.document.scrollTop() + o.scrollSpeed);
				}

				if(event.pageX - this.document.scrollLeft() < o.scrollSensitivity) {
					scrolled = this.document.scrollLeft(this.document.scrollLeft() - o.scrollSpeed);
				} else if(this.window.width() - (event.pageX - this.document.scrollLeft()) < o.scrollSensitivity) {
					scrolled = this.document.scrollLeft(this.document.scrollLeft() + o.scrollSpeed);
				}

			}

			if(scrolled !== false && $.ui.ddmanager && !o.dropBehaviour) {
				$.ui.ddmanager.prepareOffsets(this, event);
			}
		}

		//Regenerate the absolute position used for position checks
		this.positionAbs = this._convertPositionTo("absolute");

		//Set the helper position
		if(!this.options.axis || this.options.axis !== "y") {
			this.helper[0].style.left = this.position.left+"px";
		}
		if(!this.options.axis || this.options.axis !== "x") {
			this.helper[0].style.top = this.position.top+"px";
		}

		//Rearrange
		for (i = this.items.length - 1; i >= 0; i--) {

			//Cache variables and intersection, continue if no intersection
			item = this.items[i];
			itemElement = item.item[0];
			intersection = this._intersectsWithPointer(item);
			if (!intersection) {
				continue;
			}

			// Only put the placeholder inside the current Container, skip all
			// items from other containers. This works because when moving
			// an item from one container to another the
			// currentContainer is switched before the placeholder is moved.
			//
			// Without this, moving items in "sub-sortables" can cause
			// the placeholder to jitter between the outer and inner container.
			if (item.instance !== this.currentContainer) {
				continue;
			}

			// cannot intersect with itself
			// no useless actions that have been done before
			// no action if the item moved is the parent of the item checked
			if (itemElement !== this.currentItem[0] &&
				this.placeholder[intersection === 1 ? "next" : "prev"]()[0] !== itemElement &&
				!$.contains(this.placeholder[0], itemElement) &&
				(this.options.type === "semi-dynamic" ? !$.contains(this.element[0], itemElement) : true)
			) {

				this.direction = intersection === 1 ? "down" : "up";

				if (this.options.tolerance === "pointer" || this._intersectsWithSides(item)) {
					this._rearrange(event, item);
				} else {
					break;
				}

				this._trigger("change", event, this._uiHash());
				break;
			}
		}

		//Post events to containers
		this._contactContainers(event);

		//Interconnect with droppables
		if($.ui.ddmanager) {
			$.ui.ddmanager.drag(this, event);
		}

		//Call callbacks
		this._trigger("sort", event, this._uiHash());

		this.lastPositionAbs = this.positionAbs;
		return false;

	},

	_mouseStop: function(event, noPropagation) {

		if(!event) {
			return;
		}

		//If we are using droppables, inform the manager about the drop
		if ($.ui.ddmanager && !this.options.dropBehaviour) {
			$.ui.ddmanager.drop(this, event);
		}

		if(this.options.revert) {
			var that = this,
				cur = this.placeholder.offset(),
				axis = this.options.axis,
				animation = {};

			if ( !axis || axis === "x" ) {
				animation.left = cur.left - this.offset.parent.left - this.margins.left + (this.offsetParent[0] === this.document[0].body ? 0 : this.offsetParent[0].scrollLeft);
			}
			if ( !axis || axis === "y" ) {
				animation.top = cur.top - this.offset.parent.top - this.margins.top + (this.offsetParent[0] === this.document[0].body ? 0 : this.offsetParent[0].scrollTop);
			}
			this.reverting = true;
			$(this.helper).animate( animation, parseInt(this.options.revert, 10) || 500, function() {
				that._clear(event);
			});
		} else {
			this._clear(event, noPropagation);
		}

		return false;

	},

	cancel: function() {

		if(this.dragging) {

			this._mouseUp({ target: null });

			if(this.options.helper === "original") {
				this.currentItem.css(this._storedCSS).removeClass("ui-sortable-helper");
			} else {
				this.currentItem.show();
			}

			//Post deactivating events to containers
			for (var i = this.containers.length - 1; i >= 0; i--){
				this.containers[i]._trigger("deactivate", null, this._uiHash(this));
				if(this.containers[i].containerCache.over) {
					this.containers[i]._trigger("out", null, this._uiHash(this));
					this.containers[i].containerCache.over = 0;
				}
			}

		}

		if (this.placeholder) {
			//$(this.placeholder[0]).remove(); would have been the jQuery way - unfortunately, it unbinds ALL events from the original node!
			if(this.placeholder[0].parentNode) {
				this.placeholder[0].parentNode.removeChild(this.placeholder[0]);
			}
			if(this.options.helper !== "original" && this.helper && this.helper[0].parentNode) {
				this.helper.remove();
			}

			$.extend(this, {
				helper: null,
				dragging: false,
				reverting: false,
				_noFinalSort: null
			});

			if(this.domPosition.prev) {
				$(this.domPosition.prev).after(this.currentItem);
			} else {
				$(this.domPosition.parent).prepend(this.currentItem);
			}
		}

		return this;

	},

	serialize: function(o) {

		var items = this._getItemsAsjQuery(o && o.connected),
			str = [];
		o = o || {};

		$(items).each(function() {
			var res = ($(o.item || this).attr(o.attribute || "id") || "").match(o.expression || (/(.+)[\-=_](.+)/));
			if (res) {
				str.push((o.key || res[1]+"[]")+"="+(o.key && o.expression ? res[1] : res[2]));
			}
		});

		if(!str.length && o.key) {
			str.push(o.key + "=");
		}

		return str.join("&");

	},

	toArray: function(o) {

		var items = this._getItemsAsjQuery(o && o.connected),
			ret = [];

		o = o || {};

		items.each(function() { ret.push($(o.item || this).attr(o.attribute || "id") || ""); });
		return ret;

	},

	/* Be careful with the following core functions */
	_intersectsWith: function(item) {

		var x1 = this.positionAbs.left,
			x2 = x1 + this.helperProportions.width,
			y1 = this.positionAbs.top,
			y2 = y1 + this.helperProportions.height,
			l = item.left,
			r = l + item.width,
			t = item.top,
			b = t + item.height,
			dyClick = this.offset.click.top,
			dxClick = this.offset.click.left,
			isOverElementHeight = ( this.options.axis === "x" ) || ( ( y1 + dyClick ) > t && ( y1 + dyClick ) < b ),
			isOverElementWidth = ( this.options.axis === "y" ) || ( ( x1 + dxClick ) > l && ( x1 + dxClick ) < r ),
			isOverElement = isOverElementHeight && isOverElementWidth;

		if ( this.options.tolerance === "pointer" ||
			this.options.forcePointerForContainers ||
			(this.options.tolerance !== "pointer" && this.helperProportions[this.floating ? "width" : "height"] > item[this.floating ? "width" : "height"])
		) {
			return isOverElement;
		} else {

			return (l < x1 + (this.helperProportions.width / 2) && // Right Half
				x2 - (this.helperProportions.width / 2) < r && // Left Half
				t < y1 + (this.helperProportions.height / 2) && // Bottom Half
				y2 - (this.helperProportions.height / 2) < b ); // Top Half

		}
	},

	_intersectsWithPointer: function(item) {

		var isOverElementHeight = (this.options.axis === "x") || this._isOverAxis(this.positionAbs.top + this.offset.click.top, item.top, item.height),
			isOverElementWidth = (this.options.axis === "y") || this._isOverAxis(this.positionAbs.left + this.offset.click.left, item.left, item.width),
			isOverElement = isOverElementHeight && isOverElementWidth,
			verticalDirection = this._getDragVerticalDirection(),
			horizontalDirection = this._getDragHorizontalDirection();

		if (!isOverElement) {
			return false;
		}

		return this.floating ?
			( ((horizontalDirection && horizontalDirection === "right") || verticalDirection === "down") ? 2 : 1 )
			: ( verticalDirection && (verticalDirection === "down" ? 2 : 1) );

	},

	_intersectsWithSides: function(item) {

		var isOverBottomHalf = this._isOverAxis(this.positionAbs.top + this.offset.click.top, item.top + (item.height/2), item.height),
			isOverRightHalf = this._isOverAxis(this.positionAbs.left + this.offset.click.left, item.left + (item.width/2), item.width),
			verticalDirection = this._getDragVerticalDirection(),
			horizontalDirection = this._getDragHorizontalDirection();

		if (this.floating && horizontalDirection) {
			return ((horizontalDirection === "right" && isOverRightHalf) || (horizontalDirection === "left" && !isOverRightHalf));
		} else {
			return verticalDirection && ((verticalDirection === "down" && isOverBottomHalf) || (verticalDirection === "up" && !isOverBottomHalf));
		}

	},

	_getDragVerticalDirection: function() {
		var delta = this.positionAbs.top - this.lastPositionAbs.top;
		return delta !== 0 && (delta > 0 ? "down" : "up");
	},

	_getDragHorizontalDirection: function() {
		var delta = this.positionAbs.left - this.lastPositionAbs.left;
		return delta !== 0 && (delta > 0 ? "right" : "left");
	},

	refresh: function(event) {
		this._refreshItems(event);
		this._setHandleClassName();
		this.refreshPositions();
		return this;
	},

	_connectWith: function() {
		var options = this.options;
		return options.connectWith.constructor === String ? [options.connectWith] : options.connectWith;
	},

	_getItemsAsjQuery: function(connected) {

		var i, j, cur, inst,
			items = [],
			queries = [],
			connectWith = this._connectWith();

		if(connectWith && connected) {
			for (i = connectWith.length - 1; i >= 0; i--){
				cur = $(connectWith[i], this.document[0]);
				for ( j = cur.length - 1; j >= 0; j--){
					inst = $.data(cur[j], this.widgetFullName);
					if(inst && inst !== this && !inst.options.disabled) {
						queries.push([$.isFunction(inst.options.items) ? inst.options.items.call(inst.element) : $(inst.options.items, inst.element).not(".ui-sortable-helper").not(".ui-sortable-placeholder"), inst]);
					}
				}
			}
		}

		queries.push([$.isFunction(this.options.items) ? this.options.items.call(this.element, null, { options: this.options, item: this.currentItem }) : $(this.options.items, this.element).not(".ui-sortable-helper").not(".ui-sortable-placeholder"), this]);

		function addItems() {
			items.push( this );
		}
		for (i = queries.length - 1; i >= 0; i--){
			queries[i][0].each( addItems );
		}

		return $(items);

	},

	_removeCurrentsFromItems: function() {

		var list = this.currentItem.find(":data(" + this.widgetName + "-item)");

		this.items = $.grep(this.items, function (item) {
			for (var j=0; j < list.length; j++) {
				if(list[j] === item.item[0]) {
					return false;
				}
			}
			return true;
		});

	},

	_refreshItems: function(event) {

		this.items = [];
		this.containers = [this];

		var i, j, cur, inst, targetData, _queries, item, queriesLength,
			items = this.items,
			queries = [[$.isFunction(this.options.items) ? this.options.items.call(this.element[0], event, { item: this.currentItem }) : $(this.options.items, this.element), this]],
			connectWith = this._connectWith();

		if(connectWith && this.ready) { //Shouldn't be run the first time through due to massive slow-down
			for (i = connectWith.length - 1; i >= 0; i--){
				cur = $(connectWith[i], this.document[0]);
				for (j = cur.length - 1; j >= 0; j--){
					inst = $.data(cur[j], this.widgetFullName);
					if(inst && inst !== this && !inst.options.disabled) {
						queries.push([$.isFunction(inst.options.items) ? inst.options.items.call(inst.element[0], event, { item: this.currentItem }) : $(inst.options.items, inst.element), inst]);
						this.containers.push(inst);
					}
				}
			}
		}

		for (i = queries.length - 1; i >= 0; i--) {
			targetData = queries[i][1];
			_queries = queries[i][0];

			for (j=0, queriesLength = _queries.length; j < queriesLength; j++) {
				item = $(_queries[j]);

				item.data(this.widgetName + "-item", targetData); // Data for target checking (mouse manager)

				items.push({
					item: item,
					instance: targetData,
					width: 0, height: 0,
					left: 0, top: 0
				});
			}
		}

	},

	refreshPositions: function(fast) {

		// Determine whether items are being displayed horizontally
		this.floating = this.items.length ?
			this.options.axis === "x" || this._isFloating( this.items[ 0 ].item ) :
			false;

		//This has to be redone because due to the item being moved out/into the offsetParent, the offsetParent's position will change
		if(this.offsetParent && this.helper) {
			this.offset.parent = this._getParentOffset();
		}

		var i, item, t, p;

		for (i = this.items.length - 1; i >= 0; i--){
			item = this.items[i];

			//We ignore calculating positions of all connected containers when we're not over them
			if(item.instance !== this.currentContainer && this.currentContainer && item.item[0] !== this.currentItem[0]) {
				continue;
			}

			t = this.options.toleranceElement ? $(this.options.toleranceElement, item.item) : item.item;

			if (!fast) {
				item.width = t.outerWidth();
				item.height = t.outerHeight();
			}

			p = t.offset();
			item.left = p.left;
			item.top = p.top;
		}

		if(this.options.custom && this.options.custom.refreshContainers) {
			this.options.custom.refreshContainers.call(this);
		} else {
			for (i = this.containers.length - 1; i >= 0; i--){
				p = this.containers[i].element.offset();
				this.containers[i].containerCache.left = p.left;
				this.containers[i].containerCache.top = p.top;
				this.containers[i].containerCache.width = this.containers[i].element.outerWidth();
				this.containers[i].containerCache.height = this.containers[i].element.outerHeight();
			}
		}

		return this;
	},

	_createPlaceholder: function(that) {
		that = that || this;
		var className,
			o = that.options;

		if(!o.placeholder || o.placeholder.constructor === String) {
			className = o.placeholder;
			o.placeholder = {
				element: function() {

					var nodeName = that.currentItem[0].nodeName.toLowerCase(),
						element = $( "<" + nodeName + ">", that.document[0] )
							.addClass(className || that.currentItem[0].className+" ui-sortable-placeholder")
							.removeClass("ui-sortable-helper");

					if ( nodeName === "tbody" ) {
						that._createTrPlaceholder(
							that.currentItem.find( "tr" ).eq( 0 ),
							$( "<tr>", that.document[ 0 ] ).appendTo( element )
						);
					} else if ( nodeName === "tr" ) {
						that._createTrPlaceholder( that.currentItem, element );
					} else if ( nodeName === "img" ) {
						element.attr( "src", that.currentItem.attr( "src" ) );
					}

					if ( !className ) {
						element.css( "visibility", "hidden" );
					}

					return element;
				},
				update: function(container, p) {

					// 1. If a className is set as 'placeholder option, we don't force sizes - the class is responsible for that
					// 2. The option 'forcePlaceholderSize can be enabled to force it even if a class name is specified
					if(className && !o.forcePlaceholderSize) {
						return;
					}

					//If the element doesn't have a actual height by itself (without styles coming from a stylesheet), it receives the inline height from the dragged item
					if(!p.height()) { p.height(that.currentItem.innerHeight() - parseInt(that.currentItem.css("paddingTop")||0, 10) - parseInt(that.currentItem.css("paddingBottom")||0, 10)); }
					if(!p.width()) { p.width(that.currentItem.innerWidth() - parseInt(that.currentItem.css("paddingLeft")||0, 10) - parseInt(that.currentItem.css("paddingRight")||0, 10)); }
				}
			};
		}

		//Create the placeholder
		that.placeholder = $(o.placeholder.element.call(that.element, that.currentItem));

		//Append it after the actual current item
		that.currentItem.after(that.placeholder);

		//Update the size of the placeholder (TODO: Logic to fuzzy, see line 316/317)
		o.placeholder.update(that, that.placeholder);

	},

	_createTrPlaceholder: function( sourceTr, targetTr ) {
		var that = this;

		sourceTr.children().each(function() {
			$( "<td>&#160;</td>", that.document[ 0 ] )
				.attr( "colspan", $( this ).attr( "colspan" ) || 1 )
				.appendTo( targetTr );
		});
	},

	_contactContainers: function(event) {
		var i, j, dist, itemWithLeastDistance, posProperty, sizeProperty, cur, nearBottom, floating, axis,
			innermostContainer = null,
			innermostIndex = null;

		// get innermost container that intersects with item
		for (i = this.containers.length - 1; i >= 0; i--) {

			// never consider a container that's located within the item itself
			if($.contains(this.currentItem[0], this.containers[i].element[0])) {
				continue;
			}

			if(this._intersectsWith(this.containers[i].containerCache)) {

				// if we've already found a container and it's more "inner" than this, then continue
				if(innermostContainer && $.contains(this.containers[i].element[0], innermostContainer.element[0])) {
					continue;
				}

				innermostContainer = this.containers[i];
				innermostIndex = i;

			} else {
				// container doesn't intersect. trigger "out" event if necessary
				if(this.containers[i].containerCache.over) {
					this.containers[i]._trigger("out", event, this._uiHash(this));
					this.containers[i].containerCache.over = 0;
				}
			}

		}

		// if no intersecting containers found, return
		if(!innermostContainer) {
			return;
		}

		// move the item into the container if it's not there already
		if(this.containers.length === 1) {
			if (!this.containers[innermostIndex].containerCache.over) {
				this.containers[innermostIndex]._trigger("over", event, this._uiHash(this));
				this.containers[innermostIndex].containerCache.over = 1;
			}
		} else {

			//When entering a new container, we will find the item with the least distance and append our item near it
			dist = 10000;
			itemWithLeastDistance = null;
			floating = innermostContainer.floating || this._isFloating(this.currentItem);
			posProperty = floating ? "left" : "top";
			sizeProperty = floating ? "width" : "height";
			axis = floating ? "clientX" : "clientY";

			for (j = this.items.length - 1; j >= 0; j--) {
				if(!$.contains(this.containers[innermostIndex].element[0], this.items[j].item[0])) {
					continue;
				}
				if(this.items[j].item[0] === this.currentItem[0]) {
					continue;
				}

				cur = this.items[j].item.offset()[posProperty];
				nearBottom = false;
				if ( event[ axis ] - cur > this.items[ j ][ sizeProperty ] / 2 ) {
					nearBottom = true;
				}

				if ( Math.abs( event[ axis ] - cur ) < dist ) {
					dist = Math.abs( event[ axis ] - cur );
					itemWithLeastDistance = this.items[ j ];
					this.direction = nearBottom ? "up": "down";
				}
			}

			//Check if dropOnEmpty is enabled
			if(!itemWithLeastDistance && !this.options.dropOnEmpty) {
				return;
			}

			if(this.currentContainer === this.containers[innermostIndex]) {
				if ( !this.currentContainer.containerCache.over ) {
					this.containers[ innermostIndex ]._trigger( "over", event, this._uiHash() );
					this.currentContainer.containerCache.over = 1;
				}
				return;
			}

			itemWithLeastDistance ? this._rearrange(event, itemWithLeastDistance, null, true) : this._rearrange(event, null, this.containers[innermostIndex].element, true);
			this._trigger("change", event, this._uiHash());
			this.containers[innermostIndex]._trigger("change", event, this._uiHash(this));
			this.currentContainer = this.containers[innermostIndex];

			//Update the placeholder
			this.options.placeholder.update(this.currentContainer, this.placeholder);

			this.containers[innermostIndex]._trigger("over", event, this._uiHash(this));
			this.containers[innermostIndex].containerCache.over = 1;
		}


	},

	_createHelper: function(event) {

		var o = this.options,
			helper = $.isFunction(o.helper) ? $(o.helper.apply(this.element[0], [event, this.currentItem])) : (o.helper === "clone" ? this.currentItem.clone() : this.currentItem);

		//Add the helper to the DOM if that didn't happen already
		if(!helper.parents("body").length) {
			$(o.appendTo !== "parent" ? o.appendTo : this.currentItem[0].parentNode)[0].appendChild(helper[0]);
		}

		if(helper[0] === this.currentItem[0]) {
			this._storedCSS = { width: this.currentItem[0].style.width, height: this.currentItem[0].style.height, position: this.currentItem.css("position"), top: this.currentItem.css("top"), left: this.currentItem.css("left") };
		}

		if(!helper[0].style.width || o.forceHelperSize) {
			helper.width(this.currentItem.width());
		}
		if(!helper[0].style.height || o.forceHelperSize) {
			helper.height(this.currentItem.height());
		}

		return helper;

	},

	_adjustOffsetFromHelper: function(obj) {
		if (typeof obj === "string") {
			obj = obj.split(" ");
		}
		if ($.isArray(obj)) {
			obj = {left: +obj[0], top: +obj[1] || 0};
		}
		if ("left" in obj) {
			this.offset.click.left = obj.left + this.margins.left;
		}
		if ("right" in obj) {
			this.offset.click.left = this.helperProportions.width - obj.right + this.margins.left;
		}
		if ("top" in obj) {
			this.offset.click.top = obj.top + this.margins.top;
		}
		if ("bottom" in obj) {
			this.offset.click.top = this.helperProportions.height - obj.bottom + this.margins.top;
		}
	},

	_getParentOffset: function() {


		//Get the offsetParent and cache its position
		this.offsetParent = this.helper.offsetParent();
		var po = this.offsetParent.offset();

		// This is a special case where we need to modify a offset calculated on start, since the following happened:
		// 1. The position of the helper is absolute, so it's position is calculated based on the next positioned parent
		// 2. The actual offset parent is a child of the scroll parent, and the scroll parent isn't the document, which means that
		//    the scroll is included in the initial calculation of the offset of the parent, and never recalculated upon drag
		if(this.cssPosition === "absolute" && this.scrollParent[0] !== this.document[0] && $.contains(this.scrollParent[0], this.offsetParent[0])) {
			po.left += this.scrollParent.scrollLeft();
			po.top += this.scrollParent.scrollTop();
		}

		// This needs to be actually done for all browsers, since pageX/pageY includes this information
		// with an ugly IE fix
		if( this.offsetParent[0] === this.document[0].body || (this.offsetParent[0].tagName && this.offsetParent[0].tagName.toLowerCase() === "html" && $.ui.ie)) {
			po = { top: 0, left: 0 };
		}

		return {
			top: po.top + (parseInt(this.offsetParent.css("borderTopWidth"),10) || 0),
			left: po.left + (parseInt(this.offsetParent.css("borderLeftWidth"),10) || 0)
		};

	},

	_getRelativeOffset: function() {

		if(this.cssPosition === "relative") {
			var p = this.currentItem.position();
			return {
				top: p.top - (parseInt(this.helper.css("top"),10) || 0) + this.scrollParent.scrollTop(),
				left: p.left - (parseInt(this.helper.css("left"),10) || 0) + this.scrollParent.scrollLeft()
			};
		} else {
			return { top: 0, left: 0 };
		}

	},

	_cacheMargins: function() {
		this.margins = {
			left: (parseInt(this.currentItem.css("marginLeft"),10) || 0),
			top: (parseInt(this.currentItem.css("marginTop"),10) || 0)
		};
	},

	_cacheHelperProportions: function() {
		this.helperProportions = {
			width: this.helper.outerWidth(),
			height: this.helper.outerHeight()
		};
	},

	_setContainment: function() {

		var ce, co, over,
			o = this.options;
		if(o.containment === "parent") {
			o.containment = this.helper[0].parentNode;
		}
		if(o.containment === "document" || o.containment === "window") {
			this.containment = [
				0 - this.offset.relative.left - this.offset.parent.left,
				0 - this.offset.relative.top - this.offset.parent.top,
				o.containment === "document" ? this.document.width() : this.window.width() - this.helperProportions.width - this.margins.left,
				(o.containment === "document" ? this.document.width() : this.window.height() || this.document[0].body.parentNode.scrollHeight) - this.helperProportions.height - this.margins.top
			];
		}

		if(!(/^(document|window|parent)$/).test(o.containment)) {
			ce = $(o.containment)[0];
			co = $(o.containment).offset();
			over = ($(ce).css("overflow") !== "hidden");

			this.containment = [
				co.left + (parseInt($(ce).css("borderLeftWidth"),10) || 0) + (parseInt($(ce).css("paddingLeft"),10) || 0) - this.margins.left,
				co.top + (parseInt($(ce).css("borderTopWidth"),10) || 0) + (parseInt($(ce).css("paddingTop"),10) || 0) - this.margins.top,
				co.left+(over ? Math.max(ce.scrollWidth,ce.offsetWidth) : ce.offsetWidth) - (parseInt($(ce).css("borderLeftWidth"),10) || 0) - (parseInt($(ce).css("paddingRight"),10) || 0) - this.helperProportions.width - this.margins.left,
				co.top+(over ? Math.max(ce.scrollHeight,ce.offsetHeight) : ce.offsetHeight) - (parseInt($(ce).css("borderTopWidth"),10) || 0) - (parseInt($(ce).css("paddingBottom"),10) || 0) - this.helperProportions.height - this.margins.top
			];
		}

	},

	_convertPositionTo: function(d, pos) {

		if(!pos) {
			pos = this.position;
		}
		var mod = d === "absolute" ? 1 : -1,
			scroll = this.cssPosition === "absolute" && !(this.scrollParent[0] !== this.document[0] && $.contains(this.scrollParent[0], this.offsetParent[0])) ? this.offsetParent : this.scrollParent,
			scrollIsRootNode = (/(html|body)/i).test(scroll[0].tagName);

		return {
			top: (
				pos.top	+																// The absolute mouse position
				this.offset.relative.top * mod +										// Only for relative positioned nodes: Relative offset from element to offset parent
				this.offset.parent.top * mod -											// The offsetParent's offset without borders (offset + border)
				( ( this.cssPosition === "fixed" ? -this.scrollParent.scrollTop() : ( scrollIsRootNode ? 0 : scroll.scrollTop() ) ) * mod)
			),
			left: (
				pos.left +																// The absolute mouse position
				this.offset.relative.left * mod +										// Only for relative positioned nodes: Relative offset from element to offset parent
				this.offset.parent.left * mod	-										// The offsetParent's offset without borders (offset + border)
				( ( this.cssPosition === "fixed" ? -this.scrollParent.scrollLeft() : scrollIsRootNode ? 0 : scroll.scrollLeft() ) * mod)
			)
		};

	},

	_generatePosition: function(event) {

		var top, left,
			o = this.options,
			pageX = event.pageX,
			pageY = event.pageY,
			scroll = this.cssPosition === "absolute" && !(this.scrollParent[0] !== this.document[0] && $.contains(this.scrollParent[0], this.offsetParent[0])) ? this.offsetParent : this.scrollParent, scrollIsRootNode = (/(html|body)/i).test(scroll[0].tagName);

		// This is another very weird special case that only happens for relative elements:
		// 1. If the css position is relative
		// 2. and the scroll parent is the document or similar to the offset parent
		// we have to refresh the relative offset during the scroll so there are no jumps
		if(this.cssPosition === "relative" && !(this.scrollParent[0] !== this.document[0] && this.scrollParent[0] !== this.offsetParent[0])) {
			this.offset.relative = this._getRelativeOffset();
		}

		/*
		 * - Position constraining -
		 * Constrain the position to a mix of grid, containment.
		 */

		if(this.originalPosition) { //If we are not dragging yet, we won't check for options

			if(this.containment) {
				if(event.pageX - this.offset.click.left < this.containment[0]) {
					pageX = this.containment[0] + this.offset.click.left;
				}
				if(event.pageY - this.offset.click.top < this.containment[1]) {
					pageY = this.containment[1] + this.offset.click.top;
				}
				if(event.pageX - this.offset.click.left > this.containment[2]) {
					pageX = this.containment[2] + this.offset.click.left;
				}
				if(event.pageY - this.offset.click.top > this.containment[3]) {
					pageY = this.containment[3] + this.offset.click.top;
				}
			}

			if(o.grid) {
				top = this.originalPageY + Math.round((pageY - this.originalPageY) / o.grid[1]) * o.grid[1];
				pageY = this.containment ? ( (top - this.offset.click.top >= this.containment[1] && top - this.offset.click.top <= this.containment[3]) ? top : ((top - this.offset.click.top >= this.containment[1]) ? top - o.grid[1] : top + o.grid[1])) : top;

				left = this.originalPageX + Math.round((pageX - this.originalPageX) / o.grid[0]) * o.grid[0];
				pageX = this.containment ? ( (left - this.offset.click.left >= this.containment[0] && left - this.offset.click.left <= this.containment[2]) ? left : ((left - this.offset.click.left >= this.containment[0]) ? left - o.grid[0] : left + o.grid[0])) : left;
			}

		}

		return {
			top: (
				pageY -																// The absolute mouse position
				this.offset.click.top -													// Click offset (relative to the element)
				this.offset.relative.top	-											// Only for relative positioned nodes: Relative offset from element to offset parent
				this.offset.parent.top +												// The offsetParent's offset without borders (offset + border)
				( ( this.cssPosition === "fixed" ? -this.scrollParent.scrollTop() : ( scrollIsRootNode ? 0 : scroll.scrollTop() ) ))
			),
			left: (
				pageX -																// The absolute mouse position
				this.offset.click.left -												// Click offset (relative to the element)
				this.offset.relative.left	-											// Only for relative positioned nodes: Relative offset from element to offset parent
				this.offset.parent.left +												// The offsetParent's offset without borders (offset + border)
				( ( this.cssPosition === "fixed" ? -this.scrollParent.scrollLeft() : scrollIsRootNode ? 0 : scroll.scrollLeft() ))
			)
		};

	},

	_rearrange: function(event, i, a, hardRefresh) {

		a ? a[0].appendChild(this.placeholder[0]) : i.item[0].parentNode.insertBefore(this.placeholder[0], (this.direction === "down" ? i.item[0] : i.item[0].nextSibling));

		//Various things done here to improve the performance:
		// 1. we create a setTimeout, that calls refreshPositions
		// 2. on the instance, we have a counter variable, that get's higher after every append
		// 3. on the local scope, we copy the counter variable, and check in the timeout, if it's still the same
		// 4. this lets only the last addition to the timeout stack through
		this.counter = this.counter ? ++this.counter : 1;
		var counter = this.counter;

		this._delay(function() {
			if(counter === this.counter) {
				this.refreshPositions(!hardRefresh); //Precompute after each DOM insertion, NOT on mousemove
			}
		});

	},

	_clear: function(event, noPropagation) {

		this.reverting = false;
		// We delay all events that have to be triggered to after the point where the placeholder has been removed and
		// everything else normalized again
		var i,
			delayedTriggers = [];

		// We first have to update the dom position of the actual currentItem
		// Note: don't do it if the current item is already removed (by a user), or it gets reappended (see #4088)
		if(!this._noFinalSort && this.currentItem.parent().length) {
			this.placeholder.before(this.currentItem);
		}
		this._noFinalSort = null;

		if(this.helper[0] === this.currentItem[0]) {
			for(i in this._storedCSS) {
				if(this._storedCSS[i] === "auto" || this._storedCSS[i] === "static") {
					this._storedCSS[i] = "";
				}
			}
			this.currentItem.css(this._storedCSS).removeClass("ui-sortable-helper");
		} else {
			this.currentItem.show();
		}

		if(this.fromOutside && !noPropagation) {
			delayedTriggers.push(function(event) { this._trigger("receive", event, this._uiHash(this.fromOutside)); });
		}
		if((this.fromOutside || this.domPosition.prev !== this.currentItem.prev().not(".ui-sortable-helper")[0] || this.domPosition.parent !== this.currentItem.parent()[0]) && !noPropagation) {
			delayedTriggers.push(function(event) { this._trigger("update", event, this._uiHash()); }); //Trigger update callback if the DOM position has changed
		}

		// Check if the items Container has Changed and trigger appropriate
		// events.
		if (this !== this.currentContainer) {
			if(!noPropagation) {
				delayedTriggers.push(function(event) { this._trigger("remove", event, this._uiHash()); });
				delayedTriggers.push((function(c) { return function(event) { c._trigger("receive", event, this._uiHash(this)); };  }).call(this, this.currentContainer));
				delayedTriggers.push((function(c) { return function(event) { c._trigger("update", event, this._uiHash(this));  }; }).call(this, this.currentContainer));
			}
		}


		//Post events to containers
		function delayEvent( type, instance, container ) {
			return function( event ) {
				container._trigger( type, event, instance._uiHash( instance ) );
			};
		}
		for (i = this.containers.length - 1; i >= 0; i--){
			if (!noPropagation) {
				delayedTriggers.push( delayEvent( "deactivate", this, this.containers[ i ] ) );
			}
			if(this.containers[i].containerCache.over) {
				delayedTriggers.push( delayEvent( "out", this, this.containers[ i ] ) );
				this.containers[i].containerCache.over = 0;
			}
		}

		//Do what was originally in plugins
		if ( this.storedCursor ) {
			this.document.find( "body" ).css( "cursor", this.storedCursor );
			this.storedStylesheet.remove();
		}
		if(this._storedOpacity) {
			this.helper.css("opacity", this._storedOpacity);
		}
		if(this._storedZIndex) {
			this.helper.css("zIndex", this._storedZIndex === "auto" ? "" : this._storedZIndex);
		}

		this.dragging = false;

		if(!noPropagation) {
			this._trigger("beforeStop", event, this._uiHash());
		}

		//$(this.placeholder[0]).remove(); would have been the jQuery way - unfortunately, it unbinds ALL events from the original node!
		this.placeholder[0].parentNode.removeChild(this.placeholder[0]);

		if ( !this.cancelHelperRemoval ) {
			if ( this.helper[ 0 ] !== this.currentItem[ 0 ] ) {
				this.helper.remove();
			}
			this.helper = null;
		}

		if(!noPropagation) {
			for (i=0; i < delayedTriggers.length; i++) {
				delayedTriggers[i].call(this, event);
			} //Trigger all delayed events
			this._trigger("stop", event, this._uiHash());
		}

		this.fromOutside = false;
		return !this.cancelHelperRemoval;

	},

	_trigger: function() {
		if ($.Widget.prototype._trigger.apply(this, arguments) === false) {
			this.cancel();
		}
	},

	_uiHash: function(_inst) {
		var inst = _inst || this;
		return {
			helper: inst.helper,
			placeholder: inst.placeholder || $([]),
			position: inst.position,
			originalPosition: inst.originalPosition,
			offset: inst.positionAbs,
			item: inst.currentItem,
			sender: _inst ? _inst.element : null
		};
	}

});


/*!
 * jQuery UI Accordion 1.11.4
 * http://jqueryui.com
 *
 * Copyright jQuery Foundation and other contributors
 * Released under the MIT license.
 * http://jquery.org/license
 *
 * http://api.jqueryui.com/accordion/
 */


var accordion = $.widget( "ui.accordion", {
	version: "1.11.4",
	options: {
		active: 0,
		animate: {},
		collapsible: false,
		event: "click",
		header: "> li > :first-child,> :not(li):even",
		heightStyle: "auto",
		icons: {
			activeHeader: "ui-icon-triangle-1-s",
			header: "ui-icon-triangle-1-e"
		},

		// callbacks
		activate: null,
		beforeActivate: null
	},

	hideProps: {
		borderTopWidth: "hide",
		borderBottomWidth: "hide",
		paddingTop: "hide",
		paddingBottom: "hide",
		height: "hide"
	},

	showProps: {
		borderTopWidth: "show",
		borderBottomWidth: "show",
		paddingTop: "show",
		paddingBottom: "show",
		height: "show"
	},

	_create: function() {
		var options = this.options;
		this.prevShow = this.prevHide = $();
		this.element.addClass( "ui-accordion ui-widget ui-helper-reset" )
			// ARIA
			.attr( "role", "tablist" );

		// don't allow collapsible: false and active: false / null
		if ( !options.collapsible && (options.active === false || options.active == null) ) {
			options.active = 0;
		}

		this._processPanels();
		// handle negative values
		if ( options.active < 0 ) {
			options.active += this.headers.length;
		}
		this._refresh();
	},

	_getCreateEventData: function() {
		return {
			header: this.active,
			panel: !this.active.length ? $() : this.active.next()
		};
	},

	_createIcons: function() {
		var icons = this.options.icons;
		if ( icons ) {
			$( "<span>" )
				.addClass( "ui-accordion-header-icon ui-icon " + icons.header )
				.prependTo( this.headers );
			this.active.children( ".ui-accordion-header-icon" )
				.removeClass( icons.header )
				.addClass( icons.activeHeader );
			this.headers.addClass( "ui-accordion-icons" );
		}
	},

	_destroyIcons: function() {
		this.headers
			.removeClass( "ui-accordion-icons" )
			.children( ".ui-accordion-header-icon" )
				.remove();
	},

	_destroy: function() {
		var contents;

		// clean up main element
		this.element
			.removeClass( "ui-accordion ui-widget ui-helper-reset" )
			.removeAttr( "role" );

		// clean up headers
		this.headers
			.removeClass( "ui-accordion-header ui-accordion-header-active ui-state-default " +
				"ui-corner-all ui-state-active ui-state-disabled ui-corner-top" )
			.removeAttr( "role" )
			.removeAttr( "aria-expanded" )
			.removeAttr( "aria-selected" )
			.removeAttr( "aria-controls" )
			.removeAttr( "tabIndex" )
			.removeUniqueId();

		this._destroyIcons();

		// clean up content panels
		contents = this.headers.next()
			.removeClass( "ui-helper-reset ui-widget-content ui-corner-bottom " +
				"ui-accordion-content ui-accordion-content-active ui-state-disabled" )
			.css( "display", "" )
			.removeAttr( "role" )
			.removeAttr( "aria-hidden" )
			.removeAttr( "aria-labelledby" )
			.removeUniqueId();

		if ( this.options.heightStyle !== "content" ) {
			contents.css( "height", "" );
		}
	},

	_setOption: function( key, value ) {
		if ( key === "active" ) {
			// _activate() will handle invalid values and update this.options
			this._activate( value );
			return;
		}

		if ( key === "event" ) {
			if ( this.options.event ) {
				this._off( this.headers, this.options.event );
			}
			this._setupEvents( value );
		}

		this._super( key, value );

		// setting collapsible: false while collapsed; open first panel
		if ( key === "collapsible" && !value && this.options.active === false ) {
			this._activate( 0 );
		}

		if ( key === "icons" ) {
			this._destroyIcons();
			if ( value ) {
				this._createIcons();
			}
		}

		// #5332 - opacity doesn't cascade to positioned elements in IE
		// so we need to add the disabled class to the headers and panels
		if ( key === "disabled" ) {
			this.element
				.toggleClass( "ui-state-disabled", !!value )
				.attr( "aria-disabled", value );
			this.headers.add( this.headers.next() )
				.toggleClass( "ui-state-disabled", !!value );
		}
	},

	_keydown: function( event ) {
		if ( event.altKey || event.ctrlKey ) {
			return;
		}

		var keyCode = $.ui.keyCode,
			length = this.headers.length,
			currentIndex = this.headers.index( event.target ),
			toFocus = false;

		switch ( event.keyCode ) {
			case keyCode.RIGHT:
			case keyCode.DOWN:
				toFocus = this.headers[ ( currentIndex + 1 ) % length ];
				break;
			case keyCode.LEFT:
			case keyCode.UP:
				toFocus = this.headers[ ( currentIndex - 1 + length ) % length ];
				break;
			case keyCode.SPACE:
			case keyCode.ENTER:
				this._eventHandler( event );
				break;
			case keyCode.HOME:
				toFocus = this.headers[ 0 ];
				break;
			case keyCode.END:
				toFocus = this.headers[ length - 1 ];
				break;
		}

		if ( toFocus ) {
			$( event.target ).attr( "tabIndex", -1 );
			$( toFocus ).attr( "tabIndex", 0 );
			toFocus.focus();
			event.preventDefault();
		}
	},

	_panelKeyDown: function( event ) {
		if ( event.keyCode === $.ui.keyCode.UP && event.ctrlKey ) {
			$( event.currentTarget ).prev().focus();
		}
	},

	refresh: function() {
		var options = this.options;
		this._processPanels();

		// was collapsed or no panel
		if ( ( options.active === false && options.collapsible === true ) || !this.headers.length ) {
			options.active = false;
			this.active = $();
		// active false only when collapsible is true
		} else if ( options.active === false ) {
			this._activate( 0 );
		// was active, but active panel is gone
		} else if ( this.active.length && !$.contains( this.element[ 0 ], this.active[ 0 ] ) ) {
			// all remaining panel are disabled
			if ( this.headers.length === this.headers.find(".ui-state-disabled").length ) {
				options.active = false;
				this.active = $();
			// activate previous panel
			} else {
				this._activate( Math.max( 0, options.active - 1 ) );
			}
		// was active, active panel still exists
		} else {
			// make sure active index is correct
			options.active = this.headers.index( this.active );
		}

		this._destroyIcons();

		this._refresh();
	},

	_processPanels: function() {
		var prevHeaders = this.headers,
			prevPanels = this.panels;

		this.headers = this.element.find( this.options.header )
			.addClass( "ui-accordion-header ui-state-default ui-corner-all" );

		this.panels = this.headers.next()
			.addClass( "ui-accordion-content ui-helper-reset ui-widget-content ui-corner-bottom" )
			.filter( ":not(.ui-accordion-content-active)" )
			.hide();

		// Avoid memory leaks (#10056)
		if ( prevPanels ) {
			this._off( prevHeaders.not( this.headers ) );
			this._off( prevPanels.not( this.panels ) );
		}
	},

	_refresh: function() {
		var maxHeight,
			options = this.options,
			heightStyle = options.heightStyle,
			parent = this.element.parent();

		this.active = this._findActive( options.active )
			.addClass( "ui-accordion-header-active ui-state-active ui-corner-top" )
			.removeClass( "ui-corner-all" );
		this.active.next()
			.addClass( "ui-accordion-content-active" )
			.show();

		this.headers
			.attr( "role", "tab" )
			.each(function() {
				var header = $( this ),
					headerId = header.uniqueId().attr( "id" ),
					panel = header.next(),
					panelId = panel.uniqueId().attr( "id" );
				header.attr( "aria-controls", panelId );
				panel.attr( "aria-labelledby", headerId );
			})
			.next()
				.attr( "role", "tabpanel" );

		this.headers
			.not( this.active )
			.attr({
				"aria-selected": "false",
				"aria-expanded": "false",
				tabIndex: -1
			})
			.next()
				.attr({
					"aria-hidden": "true"
				})
				.hide();

		// make sure at least one header is in the tab order
		if ( !this.active.length ) {
			this.headers.eq( 0 ).attr( "tabIndex", 0 );
		} else {
			this.active.attr({
				"aria-selected": "true",
				"aria-expanded": "true",
				tabIndex: 0
			})
			.next()
				.attr({
					"aria-hidden": "false"
				});
		}

		this._createIcons();

		this._setupEvents( options.event );

		if ( heightStyle === "fill" ) {
			maxHeight = parent.height();
			this.element.siblings( ":visible" ).each(function() {
				var elem = $( this ),
					position = elem.css( "position" );

				if ( position === "absolute" || position === "fixed" ) {
					return;
				}
				maxHeight -= elem.outerHeight( true );
			});

			this.headers.each(function() {
				maxHeight -= $( this ).outerHeight( true );
			});

			this.headers.next()
				.each(function() {
					$( this ).height( Math.max( 0, maxHeight -
						$( this ).innerHeight() + $( this ).height() ) );
				})
				.css( "overflow", "auto" );
		} else if ( heightStyle === "auto" ) {
			maxHeight = 0;
			this.headers.next()
				.each(function() {
					maxHeight = Math.max( maxHeight, $( this ).css( "height", "" ).height() );
				})
				.height( maxHeight );
		}
	},

	_activate: function( index ) {
		var active = this._findActive( index )[ 0 ];

		// trying to activate the already active panel
		if ( active === this.active[ 0 ] ) {
			return;
		}

		// trying to collapse, simulate a click on the currently active header
		active = active || this.active[ 0 ];

		this._eventHandler({
			target: active,
			currentTarget: active,
			preventDefault: $.noop
		});
	},

	_findActive: function( selector ) {
		return typeof selector === "number" ? this.headers.eq( selector ) : $();
	},

	_setupEvents: function( event ) {
		var events = {
			keydown: "_keydown"
		};
		if ( event ) {
			$.each( event.split( " " ), function( index, eventName ) {
				events[ eventName ] = "_eventHandler";
			});
		}

		this._off( this.headers.add( this.headers.next() ) );
		this._on( this.headers, events );
		this._on( this.headers.next(), { keydown: "_panelKeyDown" });
		this._hoverable( this.headers );
		this._focusable( this.headers );
	},

	_eventHandler: function( event ) {
		var options = this.options,
			active = this.active,
			clicked = $( event.currentTarget ),
			clickedIsActive = clicked[ 0 ] === active[ 0 ],
			collapsing = clickedIsActive && options.collapsible,
			toShow = collapsing ? $() : clicked.next(),
			toHide = active.next(),
			eventData = {
				oldHeader: active,
				oldPanel: toHide,
				newHeader: collapsing ? $() : clicked,
				newPanel: toShow
			};

		event.preventDefault();

		if (
				// click on active header, but not collapsible
				( clickedIsActive && !options.collapsible ) ||
				// allow canceling activation
				( this._trigger( "beforeActivate", event, eventData ) === false ) ) {
			return;
		}

		options.active = collapsing ? false : this.headers.index( clicked );

		// when the call to ._toggle() comes after the class changes
		// it causes a very odd bug in IE 8 (see #6720)
		this.active = clickedIsActive ? $() : clicked;
		this._toggle( eventData );

		// switch classes
		// corner classes on the previously active header stay after the animation
		active.removeClass( "ui-accordion-header-active ui-state-active" );
		if ( options.icons ) {
			active.children( ".ui-accordion-header-icon" )
				.removeClass( options.icons.activeHeader )
				.addClass( options.icons.header );
		}

		if ( !clickedIsActive ) {
			clicked
				.removeClass( "ui-corner-all" )
				.addClass( "ui-accordion-header-active ui-state-active ui-corner-top" );
			if ( options.icons ) {
				clicked.children( ".ui-accordion-header-icon" )
					.removeClass( options.icons.header )
					.addClass( options.icons.activeHeader );
			}

			clicked
				.next()
				.addClass( "ui-accordion-content-active" );
		}
	},

	_toggle: function( data ) {
		var toShow = data.newPanel,
			toHide = this.prevShow.length ? this.prevShow : data.oldPanel;

		// handle activating a panel during the animation for another activation
		this.prevShow.add( this.prevHide ).stop( true, true );
		this.prevShow = toShow;
		this.prevHide = toHide;

		if ( this.options.animate ) {
			this._animate( toShow, toHide, data );
		} else {
			toHide.hide();
			toShow.show();
			this._toggleComplete( data );
		}

		toHide.attr({
			"aria-hidden": "true"
		});
		toHide.prev().attr({
			"aria-selected": "false",
			"aria-expanded": "false"
		});
		// if we're switching panels, remove the old header from the tab order
		// if we're opening from collapsed state, remove the previous header from the tab order
		// if we're collapsing, then keep the collapsing header in the tab order
		if ( toShow.length && toHide.length ) {
			toHide.prev().attr({
				"tabIndex": -1,
				"aria-expanded": "false"
			});
		} else if ( toShow.length ) {
			this.headers.filter(function() {
				return parseInt( $( this ).attr( "tabIndex" ), 10 ) === 0;
			})
			.attr( "tabIndex", -1 );
		}

		toShow
			.attr( "aria-hidden", "false" )
			.prev()
				.attr({
					"aria-selected": "true",
					"aria-expanded": "true",
					tabIndex: 0
				});
	},

	_animate: function( toShow, toHide, data ) {
		var total, easing, duration,
			that = this,
			adjust = 0,
			boxSizing = toShow.css( "box-sizing" ),
			down = toShow.length &&
				( !toHide.length || ( toShow.index() < toHide.index() ) ),
			animate = this.options.animate || {},
			options = down && animate.down || animate,
			complete = function() {
				that._toggleComplete( data );
			};

		if ( typeof options === "number" ) {
			duration = options;
		}
		if ( typeof options === "string" ) {
			easing = options;
		}
		// fall back from options to animation in case of partial down settings
		easing = easing || options.easing || animate.easing;
		duration = duration || options.duration || animate.duration;

		if ( !toHide.length ) {
			return toShow.animate( this.showProps, duration, easing, complete );
		}
		if ( !toShow.length ) {
			return toHide.animate( this.hideProps, duration, easing, complete );
		}

		total = toShow.show().outerHeight();
		toHide.animate( this.hideProps, {
			duration: duration,
			easing: easing,
			step: function( now, fx ) {
				fx.now = Math.round( now );
			}
		});
		toShow
			.hide()
			.animate( this.showProps, {
				duration: duration,
				easing: easing,
				complete: complete,
				step: function( now, fx ) {
					fx.now = Math.round( now );
					if ( fx.prop !== "height" ) {
						if ( boxSizing === "content-box" ) {
							adjust += fx.now;
						}
					} else if ( that.options.heightStyle !== "content" ) {
						fx.now = Math.round( total - toHide.outerHeight() - adjust );
						adjust = 0;
					}
				}
			});
	},

	_toggleComplete: function( data ) {
		var toHide = data.oldPanel;

		toHide
			.removeClass( "ui-accordion-content-active" )
			.prev()
				.removeClass( "ui-corner-top" )
				.addClass( "ui-corner-all" );

		// Work around for rendering bug in IE (#5421)
		if ( toHide.length ) {
			toHide.parent()[ 0 ].className = toHide.parent()[ 0 ].className;
		}
		this._trigger( "activate", null, data );
	}
});


/*!
 * jQuery UI Menu 1.11.4
 * http://jqueryui.com
 *
 * Copyright jQuery Foundation and other contributors
 * Released under the MIT license.
 * http://jquery.org/license
 *
 * http://api.jqueryui.com/menu/
 */


var menu = $.widget( "ui.menu", {
	version: "1.11.4",
	defaultElement: "<ul>",
	delay: 300,
	options: {
		icons: {
			submenu: "ui-icon-carat-1-e"
		},
		items: "> *",
		menus: "ul",
		position: {
			my: "left-1 top",
			at: "right top"
		},
		role: "menu",

		// callbacks
		blur: null,
		focus: null,
		select: null
	},

	_create: function() {
		this.activeMenu = this.element;

		// Flag used to prevent firing of the click handler
		// as the event bubbles up through nested menus
		this.mouseHandled = false;
		this.element
			.uniqueId()
			.addClass( "ui-menu ui-widget ui-widget-content" )
			.toggleClass( "ui-menu-icons", !!this.element.find( ".ui-icon" ).length )
			.attr({
				role: this.options.role,
				tabIndex: 0
			});

		if ( this.options.disabled ) {
			this.element
				.addClass( "ui-state-disabled" )
				.attr( "aria-disabled", "true" );
		}

		this._on({
			// Prevent focus from sticking to links inside menu after clicking
			// them (focus should always stay on UL during navigation).
			"mousedown .ui-menu-item": function( event ) {
				event.preventDefault();
			},
			"click .ui-menu-item": function( event ) {
				var target = $( event.target );
				if ( !this.mouseHandled && target.not( ".ui-state-disabled" ).length ) {
					this.select( event );

					// Only set the mouseHandled flag if the event will bubble, see #9469.
					if ( !event.isPropagationStopped() ) {
						this.mouseHandled = true;
					}

					// Open submenu on click
					if ( target.has( ".ui-menu" ).length ) {
						this.expand( event );
					} else if ( !this.element.is( ":focus" ) && $( this.document[ 0 ].activeElement ).closest( ".ui-menu" ).length ) {

						// Redirect focus to the menu
						this.element.trigger( "focus", [ true ] );

						// If the active item is on the top level, let it stay active.
						// Otherwise, blur the active item since it is no longer visible.
						if ( this.active && this.active.parents( ".ui-menu" ).length === 1 ) {
							clearTimeout( this.timer );
						}
					}
				}
			},
			"mouseenter .ui-menu-item": function( event ) {
				// Ignore mouse events while typeahead is active, see #10458.
				// Prevents focusing the wrong item when typeahead causes a scroll while the mouse
				// is over an item in the menu
				if ( this.previousFilter ) {
					return;
				}
				var target = $( event.currentTarget );
				// Remove ui-state-active class from siblings of the newly focused menu item
				// to avoid a jump caused by adjacent elements both having a class with a border
				target.siblings( ".ui-state-active" ).removeClass( "ui-state-active" );
				this.focus( event, target );
			},
			mouseleave: "collapseAll",
			"mouseleave .ui-menu": "collapseAll",
			focus: function( event, keepActiveItem ) {
				// If there's already an active item, keep it active
				// If not, activate the first item
				var item = this.active || this.element.find( this.options.items ).eq( 0 );

				if ( !keepActiveItem ) {
					this.focus( event, item );
				}
			},
			blur: function( event ) {
				this._delay(function() {
					if ( !$.contains( this.element[0], this.document[0].activeElement ) ) {
						this.collapseAll( event );
					}
				});
			},
			keydown: "_keydown"
		});

		this.refresh();

		// Clicks outside of a menu collapse any open menus
		this._on( this.document, {
			click: function( event ) {
				if ( this._closeOnDocumentClick( event ) ) {
					this.collapseAll( event );
				}

				// Reset the mouseHandled flag
				this.mouseHandled = false;
			}
		});
	},

	_destroy: function() {
		// Destroy (sub)menus
		this.element
			.removeAttr( "aria-activedescendant" )
			.find( ".ui-menu" ).addBack()
				.removeClass( "ui-menu ui-widget ui-widget-content ui-menu-icons ui-front" )
				.removeAttr( "role" )
				.removeAttr( "tabIndex" )
				.removeAttr( "aria-labelledby" )
				.removeAttr( "aria-expanded" )
				.removeAttr( "aria-hidden" )
				.removeAttr( "aria-disabled" )
				.removeUniqueId()
				.show();

		// Destroy menu items
		this.element.find( ".ui-menu-item" )
			.removeClass( "ui-menu-item" )
			.removeAttr( "role" )
			.removeAttr( "aria-disabled" )
			.removeUniqueId()
			.removeClass( "ui-state-hover" )
			.removeAttr( "tabIndex" )
			.removeAttr( "role" )
			.removeAttr( "aria-haspopup" )
			.children().each( function() {
				var elem = $( this );
				if ( elem.data( "ui-menu-submenu-carat" ) ) {
					elem.remove();
				}
			});

		// Destroy menu dividers
		this.element.find( ".ui-menu-divider" ).removeClass( "ui-menu-divider ui-widget-content" );
	},

	_keydown: function( event ) {
		var match, prev, character, skip,
			preventDefault = true;

		switch ( event.keyCode ) {
		case $.ui.keyCode.PAGE_UP:
			this.previousPage( event );
			break;
		case $.ui.keyCode.PAGE_DOWN:
			this.nextPage( event );
			break;
		case $.ui.keyCode.HOME:
			this._move( "first", "first", event );
			break;
		case $.ui.keyCode.END:
			this._move( "last", "last", event );
			break;
		case $.ui.keyCode.UP:
			this.previous( event );
			break;
		case $.ui.keyCode.DOWN:
			this.next( event );
			break;
		case $.ui.keyCode.LEFT:
			this.collapse( event );
			break;
		case $.ui.keyCode.RIGHT:
			if ( this.active && !this.active.is( ".ui-state-disabled" ) ) {
				this.expand( event );
			}
			break;
		case $.ui.keyCode.ENTER:
		case $.ui.keyCode.SPACE:
			this._activate( event );
			break;
		case $.ui.keyCode.ESCAPE:
			this.collapse( event );
			break;
		default:
			preventDefault = false;
			prev = this.previousFilter || "";
			character = String.fromCharCode( event.keyCode );
			skip = false;

			clearTimeout( this.filterTimer );

			if ( character === prev ) {
				skip = true;
			} else {
				character = prev + character;
			}

			match = this._filterMenuItems( character );
			match = skip && match.index( this.active.next() ) !== -1 ?
				this.active.nextAll( ".ui-menu-item" ) :
				match;

			// If no matches on the current filter, reset to the last character pressed
			// to move down the menu to the first item that starts with that character
			if ( !match.length ) {
				character = String.fromCharCode( event.keyCode );
				match = this._filterMenuItems( character );
			}

			if ( match.length ) {
				this.focus( event, match );
				this.previousFilter = character;
				this.filterTimer = this._delay(function() {
					delete this.previousFilter;
				}, 1000 );
			} else {
				delete this.previousFilter;
			}
		}

		if ( preventDefault ) {
			event.preventDefault();
		}
	},

	_activate: function( event ) {
		if ( !this.active.is( ".ui-state-disabled" ) ) {
			if ( this.active.is( "[aria-haspopup='true']" ) ) {
				this.expand( event );
			} else {
				this.select( event );
			}
		}
	},

	refresh: function() {
		var menus, items,
			that = this,
			icon = this.options.icons.submenu,
			submenus = this.element.find( this.options.menus );

		this.element.toggleClass( "ui-menu-icons", !!this.element.find( ".ui-icon" ).length );

		// Initialize nested menus
		submenus.filter( ":not(.ui-menu)" )
			.addClass( "ui-menu ui-widget ui-widget-content ui-front" )
			.hide()
			.attr({
				role: this.options.role,
				"aria-hidden": "true",
				"aria-expanded": "false"
			})
			.each(function() {
				var menu = $( this ),
					item = menu.parent(),
					submenuCarat = $( "<span>" )
						.addClass( "ui-menu-icon ui-icon " + icon )
						.data( "ui-menu-submenu-carat", true );

				item
					.attr( "aria-haspopup", "true" )
					.prepend( submenuCarat );
				menu.attr( "aria-labelledby", item.attr( "id" ) );
			});

		menus = submenus.add( this.element );
		items = menus.find( this.options.items );

		// Initialize menu-items containing spaces and/or dashes only as dividers
		items.not( ".ui-menu-item" ).each(function() {
			var item = $( this );
			if ( that._isDivider( item ) ) {
				item.addClass( "ui-widget-content ui-menu-divider" );
			}
		});

		// Don't refresh list items that are already adapted
		items.not( ".ui-menu-item, .ui-menu-divider" )
			.addClass( "ui-menu-item" )
			.uniqueId()
			.attr({
				tabIndex: -1,
				role: this._itemRole()
			});

		// Add aria-disabled attribute to any disabled menu item
		items.filter( ".ui-state-disabled" ).attr( "aria-disabled", "true" );

		// If the active item has been removed, blur the menu
		if ( this.active && !$.contains( this.element[ 0 ], this.active[ 0 ] ) ) {
			this.blur();
		}
	},

	_itemRole: function() {
		return {
			menu: "menuitem",
			listbox: "option"
		}[ this.options.role ];
	},

	_setOption: function( key, value ) {
		if ( key === "icons" ) {
			this.element.find( ".ui-menu-icon" )
				.removeClass( this.options.icons.submenu )
				.addClass( value.submenu );
		}
		if ( key === "disabled" ) {
			this.element
				.toggleClass( "ui-state-disabled", !!value )
				.attr( "aria-disabled", value );
		}
		this._super( key, value );
	},

	focus: function( event, item ) {
		var nested, focused;
		this.blur( event, event && event.type === "focus" );

		this._scrollIntoView( item );

		this.active = item.first();
		focused = this.active.addClass( "ui-state-focus" ).removeClass( "ui-state-active" );
		// Only update aria-activedescendant if there's a role
		// otherwise we assume focus is managed elsewhere
		if ( this.options.role ) {
			this.element.attr( "aria-activedescendant", focused.attr( "id" ) );
		}

		// Highlight active parent menu item, if any
		this.active
			.parent()
			.closest( ".ui-menu-item" )
			.addClass( "ui-state-active" );

		if ( event && event.type === "keydown" ) {
			this._close();
		} else {
			this.timer = this._delay(function() {
				this._close();
			}, this.delay );
		}

		nested = item.children( ".ui-menu" );
		if ( nested.length && event && ( /^mouse/.test( event.type ) ) ) {
			this._startOpening(nested);
		}
		this.activeMenu = item.parent();

		this._trigger( "focus", event, { item: item } );
	},

	_scrollIntoView: function( item ) {
		var borderTop, paddingTop, offset, scroll, elementHeight, itemHeight;
		if ( this._hasScroll() ) {
			borderTop = parseFloat( $.css( this.activeMenu[0], "borderTopWidth" ) ) || 0;
			paddingTop = parseFloat( $.css( this.activeMenu[0], "paddingTop" ) ) || 0;
			offset = item.offset().top - this.activeMenu.offset().top - borderTop - paddingTop;
			scroll = this.activeMenu.scrollTop();
			elementHeight = this.activeMenu.height();
			itemHeight = item.outerHeight();

			if ( offset < 0 ) {
				this.activeMenu.scrollTop( scroll + offset );
			} else if ( offset + itemHeight > elementHeight ) {
				this.activeMenu.scrollTop( scroll + offset - elementHeight + itemHeight );
			}
		}
	},

	blur: function( event, fromFocus ) {
		if ( !fromFocus ) {
			clearTimeout( this.timer );
		}

		if ( !this.active ) {
			return;
		}

		this.active.removeClass( "ui-state-focus" );
		this.active = null;

		this._trigger( "blur", event, { item: this.active } );
	},

	_startOpening: function( submenu ) {
		clearTimeout( this.timer );

		// Don't open if already open fixes a Firefox bug that caused a .5 pixel
		// shift in the submenu position when mousing over the carat icon
		if ( submenu.attr( "aria-hidden" ) !== "true" ) {
			return;
		}

		this.timer = this._delay(function() {
			this._close();
			this._open( submenu );
		}, this.delay );
	},

	_open: function( submenu ) {
		var position = $.extend({
			of: this.active
		}, this.options.position );

		clearTimeout( this.timer );
		this.element.find( ".ui-menu" ).not( submenu.parents( ".ui-menu" ) )
			.hide()
			.attr( "aria-hidden", "true" );

		submenu
			.show()
			.removeAttr( "aria-hidden" )
			.attr( "aria-expanded", "true" )
			.position( position );
	},

	collapseAll: function( event, all ) {
		clearTimeout( this.timer );
		this.timer = this._delay(function() {
			// If we were passed an event, look for the submenu that contains the event
			var currentMenu = all ? this.element :
				$( event && event.target ).closest( this.element.find( ".ui-menu" ) );

			// If we found no valid submenu ancestor, use the main menu to close all sub menus anyway
			if ( !currentMenu.length ) {
				currentMenu = this.element;
			}

			this._close( currentMenu );

			this.blur( event );
			this.activeMenu = currentMenu;
		}, this.delay );
	},

	// With no arguments, closes the currently active menu - if nothing is active
	// it closes all menus.  If passed an argument, it will search for menus BELOW
	_close: function( startMenu ) {
		if ( !startMenu ) {
			startMenu = this.active ? this.active.parent() : this.element;
		}

		startMenu
			.find( ".ui-menu" )
				.hide()
				.attr( "aria-hidden", "true" )
				.attr( "aria-expanded", "false" )
			.end()
			.find( ".ui-state-active" ).not( ".ui-state-focus" )
				.removeClass( "ui-state-active" );
	},

	_closeOnDocumentClick: function( event ) {
		return !$( event.target ).closest( ".ui-menu" ).length;
	},

	_isDivider: function( item ) {

		// Match hyphen, em dash, en dash
		return !/[^\-\u2014\u2013\s]/.test( item.text() );
	},

	collapse: function( event ) {
		var newItem = this.active &&
			this.active.parent().closest( ".ui-menu-item", this.element );
		if ( newItem && newItem.length ) {
			this._close();
			this.focus( event, newItem );
		}
	},

	expand: function( event ) {
		var newItem = this.active &&
			this.active
				.children( ".ui-menu " )
				.find( this.options.items )
				.first();

		if ( newItem && newItem.length ) {
			this._open( newItem.parent() );

			// Delay so Firefox will not hide activedescendant change in expanding submenu from AT
			this._delay(function() {
				this.focus( event, newItem );
			});
		}
	},

	next: function( event ) {
		this._move( "next", "first", event );
	},

	previous: function( event ) {
		this._move( "prev", "last", event );
	},

	isFirstItem: function() {
		return this.active && !this.active.prevAll( ".ui-menu-item" ).length;
	},

	isLastItem: function() {
		return this.active && !this.active.nextAll( ".ui-menu-item" ).length;
	},

	_move: function( direction, filter, event ) {
		var next;
		if ( this.active ) {
			if ( direction === "first" || direction === "last" ) {
				next = this.active
					[ direction === "first" ? "prevAll" : "nextAll" ]( ".ui-menu-item" )
					.eq( -1 );
			} else {
				next = this.active
					[ direction + "All" ]( ".ui-menu-item" )
					.eq( 0 );
			}
		}
		if ( !next || !next.length || !this.active ) {
			next = this.activeMenu.find( this.options.items )[ filter ]();
		}

		this.focus( event, next );
	},

	nextPage: function( event ) {
		var item, base, height;

		if ( !this.active ) {
			this.next( event );
			return;
		}
		if ( this.isLastItem() ) {
			return;
		}
		if ( this._hasScroll() ) {
			base = this.active.offset().top;
			height = this.element.height();
			this.active.nextAll( ".ui-menu-item" ).each(function() {
				item = $( this );
				return item.offset().top - base - height < 0;
			});

			this.focus( event, item );
		} else {
			this.focus( event, this.activeMenu.find( this.options.items )
				[ !this.active ? "first" : "last" ]() );
		}
	},

	previousPage: function( event ) {
		var item, base, height;
		if ( !this.active ) {
			this.next( event );
			return;
		}
		if ( this.isFirstItem() ) {
			return;
		}
		if ( this._hasScroll() ) {
			base = this.active.offset().top;
			height = this.element.height();
			this.active.prevAll( ".ui-menu-item" ).each(function() {
				item = $( this );
				return item.offset().top - base + height > 0;
			});

			this.focus( event, item );
		} else {
			this.focus( event, this.activeMenu.find( this.options.items ).first() );
		}
	},

	_hasScroll: function() {
		return this.element.outerHeight() < this.element.prop( "scrollHeight" );
	},

	select: function( event ) {
		// TODO: It should never be possible to not have an active item at this
		// point, but the tests don't trigger mouseenter before click.
		this.active = this.active || $( event.target ).closest( ".ui-menu-item" );
		var ui = { item: this.active };
		if ( !this.active.has( ".ui-menu" ).length ) {
			this.collapseAll( event, true );
		}
		this._trigger( "select", event, ui );
	},

	_filterMenuItems: function(character) {
		var escapedCharacter = character.replace( /[\-\[\]{}()*+?.,\\\^$|#\s]/g, "\\$&" ),
			regex = new RegExp( "^" + escapedCharacter, "i" );

		return this.activeMenu
			.find( this.options.items )

			// Only match on items, not dividers or other content (#10571)
			.filter( ".ui-menu-item" )
			.filter(function() {
				return regex.test( $.trim( $( this ).text() ) );
			});
	}
});


/*!
 * jQuery UI Autocomplete 1.11.4
 * http://jqueryui.com
 *
 * Copyright jQuery Foundation and other contributors
 * Released under the MIT license.
 * http://jquery.org/license
 *
 * http://api.jqueryui.com/autocomplete/
 */


$.widget( "ui.autocomplete", {
	version: "1.11.4",
	defaultElement: "<input>",
	options: {
		appendTo: null,
		autoFocus: false,
		delay: 300,
		minLength: 1,
		position: {
			my: "left top",
			at: "left bottom",
			collision: "none"
		},
		source: null,

		// callbacks
		change: null,
		close: null,
		focus: null,
		open: null,
		response: null,
		search: null,
		select: null
	},

	requestIndex: 0,
	pending: 0,

	_create: function() {
		// Some browsers only repeat keydown events, not keypress events,
		// so we use the suppressKeyPress flag to determine if we've already
		// handled the keydown event. #7269
		// Unfortunately the code for & in keypress is the same as the up arrow,
		// so we use the suppressKeyPressRepeat flag to avoid handling keypress
		// events when we know the keydown event was used to modify the
		// search term. #7799
		var suppressKeyPress, suppressKeyPressRepeat, suppressInput,
			nodeName = this.element[ 0 ].nodeName.toLowerCase(),
			isTextarea = nodeName === "textarea",
			isInput = nodeName === "input";

		this.isMultiLine =
			// Textareas are always multi-line
			isTextarea ? true :
			// Inputs are always single-line, even if inside a contentEditable element
			// IE also treats inputs as contentEditable
			isInput ? false :
			// All other element types are determined by whether or not they're contentEditable
			this.element.prop( "isContentEditable" );

		this.valueMethod = this.element[ isTextarea || isInput ? "val" : "text" ];
		this.isNewMenu = true;

		this.element
			.addClass( "ui-autocomplete-input" )
			.attr( "autocomplete", "off" );

		this._on( this.element, {
			keydown: function( event ) {
				if ( this.element.prop( "readOnly" ) ) {
					suppressKeyPress = true;
					suppressInput = true;
					suppressKeyPressRepeat = true;
					return;
				}

				suppressKeyPress = false;
				suppressInput = false;
				suppressKeyPressRepeat = false;
				var keyCode = $.ui.keyCode;
				switch ( event.keyCode ) {
				case keyCode.PAGE_UP:
					suppressKeyPress = true;
					this._move( "previousPage", event );
					break;
				case keyCode.PAGE_DOWN:
					suppressKeyPress = true;
					this._move( "nextPage", event );
					break;
				case keyCode.UP:
					suppressKeyPress = true;
					this._keyEvent( "previous", event );
					break;
				case keyCode.DOWN:
					suppressKeyPress = true;
					this._keyEvent( "next", event );
					break;
				case keyCode.ENTER:
					// when menu is open and has focus
					if ( this.menu.active ) {
						// #6055 - Opera still allows the keypress to occur
						// which causes forms to submit
						suppressKeyPress = true;
						event.preventDefault();
						this.menu.select( event );
					}
					break;
				case keyCode.TAB:
					if ( this.menu.active ) {
						this.menu.select( event );
					}
					break;
				case keyCode.ESCAPE:
					if ( this.menu.element.is( ":visible" ) ) {
						if ( !this.isMultiLine ) {
							this._value( this.term );
						}
						this.close( event );
						// Different browsers have different default behavior for escape
						// Single press can mean undo or clear
						// Double press in IE means clear the whole form
						event.preventDefault();
					}
					break;
				default:
					suppressKeyPressRepeat = true;
					// search timeout should be triggered before the input value is changed
					this._searchTimeout( event );
					break;
				}
			},
			keypress: function( event ) {
				if ( suppressKeyPress ) {
					suppressKeyPress = false;
					if ( !this.isMultiLine || this.menu.element.is( ":visible" ) ) {
						event.preventDefault();
					}
					return;
				}
				if ( suppressKeyPressRepeat ) {
					return;
				}

				// replicate some key handlers to allow them to repeat in Firefox and Opera
				var keyCode = $.ui.keyCode;
				switch ( event.keyCode ) {
				case keyCode.PAGE_UP:
					this._move( "previousPage", event );
					break;
				case keyCode.PAGE_DOWN:
					this._move( "nextPage", event );
					break;
				case keyCode.UP:
					this._keyEvent( "previous", event );
					break;
				case keyCode.DOWN:
					this._keyEvent( "next", event );
					break;
				}
			},
			input: function( event ) {
				if ( suppressInput ) {
					suppressInput = false;
					event.preventDefault();
					return;
				}
				this._searchTimeout( event );
			},
			focus: function() {
				this.selectedItem = null;
				this.previous = this._value();
			},
			blur: function( event ) {
				if ( this.cancelBlur ) {
					delete this.cancelBlur;
					return;
				}

				clearTimeout( this.searching );
				this.close( event );
				this._change( event );
			}
		});

		this._initSource();
		this.menu = $( "<ul>" )
			.addClass( "ui-autocomplete ui-front" )
			.appendTo( this._appendTo() )
			.menu({
				// disable ARIA support, the live region takes care of that
				role: null
			})
			.hide()
			.menu( "instance" );

		this._on( this.menu.element, {
			mousedown: function( event ) {
				// prevent moving focus out of the text field
				event.preventDefault();

				// IE doesn't prevent moving focus even with event.preventDefault()
				// so we set a flag to know when we should ignore the blur event
				this.cancelBlur = true;
				this._delay(function() {
					delete this.cancelBlur;
				});

				// clicking on the scrollbar causes focus to shift to the body
				// but we can't detect a mouseup or a click immediately afterward
				// so we have to track the next mousedown and close the menu if
				// the user clicks somewhere outside of the autocomplete
				var menuElement = this.menu.element[ 0 ];
				if ( !$( event.target ).closest( ".ui-menu-item" ).length ) {
					this._delay(function() {
						var that = this;
						this.document.one( "mousedown", function( event ) {
							if ( event.target !== that.element[ 0 ] &&
									event.target !== menuElement &&
									!$.contains( menuElement, event.target ) ) {
								that.close();
							}
						});
					});
				}
			},
			menufocus: function( event, ui ) {
				var label, item;
				// support: Firefox
				// Prevent accidental activation of menu items in Firefox (#7024 #9118)
				if ( this.isNewMenu ) {
					this.isNewMenu = false;
					if ( event.originalEvent && /^mouse/.test( event.originalEvent.type ) ) {
						this.menu.blur();

						this.document.one( "mousemove", function() {
							$( event.target ).trigger( event.originalEvent );
						});

						return;
					}
				}

				item = ui.item.data( "ui-autocomplete-item" );
				if ( false !== this._trigger( "focus", event, { item: item } ) ) {
					// use value to match what will end up in the input, if it was a key event
					if ( event.originalEvent && /^key/.test( event.originalEvent.type ) ) {
						this._value( item.value );
					}
				}

				// Announce the value in the liveRegion
				label = ui.item.attr( "aria-label" ) || item.value;
				if ( label && $.trim( label ).length ) {
					this.liveRegion.children().hide();
					$( "<div>" ).text( label ).appendTo( this.liveRegion );
				}
			},
			menuselect: function( event, ui ) {
				var item = ui.item.data( "ui-autocomplete-item" ),
					previous = this.previous;

				// only trigger when focus was lost (click on menu)
				if ( this.element[ 0 ] !== this.document[ 0 ].activeElement ) {
					this.element.focus();
					this.previous = previous;
					// #6109 - IE triggers two focus events and the second
					// is asynchronous, so we need to reset the previous
					// term synchronously and asynchronously :-(
					this._delay(function() {
						this.previous = previous;
						this.selectedItem = item;
					});
				}

				if ( false !== this._trigger( "select", event, { item: item } ) ) {
					this._value( item.value );
				}
				// reset the term after the select event
				// this allows custom select handling to work properly
				this.term = this._value();

				this.close( event );
				this.selectedItem = item;
			}
		});

		this.liveRegion = $( "<span>", {
				role: "status",
				"aria-live": "assertive",
				"aria-relevant": "additions"
			})
			.addClass( "ui-helper-hidden-accessible" )
			.appendTo( this.document[ 0 ].body );

		// turning off autocomplete prevents the browser from remembering the
		// value when navigating through history, so we re-enable autocomplete
		// if the page is unloaded before the widget is destroyed. #7790
		this._on( this.window, {
			beforeunload: function() {
				this.element.removeAttr( "autocomplete" );
			}
		});
	},

	_destroy: function() {
		clearTimeout( this.searching );
		this.element
			.removeClass( "ui-autocomplete-input" )
			.removeAttr( "autocomplete" );
		this.menu.element.remove();
		this.liveRegion.remove();
	},

	_setOption: function( key, value ) {
		this._super( key, value );
		if ( key === "source" ) {
			this._initSource();
		}
		if ( key === "appendTo" ) {
			this.menu.element.appendTo( this._appendTo() );
		}
		if ( key === "disabled" && value && this.xhr ) {
			this.xhr.abort();
		}
	},

	_appendTo: function() {
		var element = this.options.appendTo;

		if ( element ) {
			element = element.jquery || element.nodeType ?
				$( element ) :
				this.document.find( element ).eq( 0 );
		}

		if ( !element || !element[ 0 ] ) {
			element = this.element.closest( ".ui-front" );
		}

		if ( !element.length ) {
			element = this.document[ 0 ].body;
		}

		return element;
	},

	_initSource: function() {
		var array, url,
			that = this;
		if ( $.isArray( this.options.source ) ) {
			array = this.options.source;
			this.source = function( request, response ) {
				response( $.ui.autocomplete.filter( array, request.term ) );
			};
		} else if ( typeof this.options.source === "string" ) {
			url = this.options.source;
			this.source = function( request, response ) {
				if ( that.xhr ) {
					that.xhr.abort();
				}
				that.xhr = $.ajax({
					url: url,
					data: request,
					dataType: "json",
					success: function( data ) {
						response( data );
					},
					error: function() {
						response([]);
					}
				});
			};
		} else {
			this.source = this.options.source;
		}
	},

	_searchTimeout: function( event ) {
		clearTimeout( this.searching );
		this.searching = this._delay(function() {

			// Search if the value has changed, or if the user retypes the same value (see #7434)
			var equalValues = this.term === this._value(),
				menuVisible = this.menu.element.is( ":visible" ),
				modifierKey = event.altKey || event.ctrlKey || event.metaKey || event.shiftKey;

			if ( !equalValues || ( equalValues && !menuVisible && !modifierKey ) ) {
				this.selectedItem = null;
				this.search( null, event );
			}
		}, this.options.delay );
	},

	search: function( value, event ) {
		value = value != null ? value : this._value();

		// always save the actual value, not the one passed as an argument
		this.term = this._value();

		if ( value.length < this.options.minLength ) {
			return this.close( event );
		}

		if ( this._trigger( "search", event ) === false ) {
			return;
		}

		return this._search( value );
	},

	_search: function( value ) {
		this.pending++;
		this.element.addClass( "ui-autocomplete-loading" );
		this.cancelSearch = false;

		this.source( { term: value }, this._response() );
	},

	_response: function() {
		var index = ++this.requestIndex;

		return $.proxy(function( content ) {
			if ( index === this.requestIndex ) {
				this.__response( content );
			}

			this.pending--;
			if ( !this.pending ) {
				this.element.removeClass( "ui-autocomplete-loading" );
			}
		}, this );
	},

	__response: function( content ) {
		if ( content ) {
			content = this._normalize( content );
		}
		this._trigger( "response", null, { content: content } );
		if ( !this.options.disabled && content && content.length && !this.cancelSearch ) {
			this._suggest( content );
			this._trigger( "open" );
		} else {
			// use ._close() instead of .close() so we don't cancel future searches
			this._close();
		}
	},

	close: function( event ) {
		this.cancelSearch = true;
		this._close( event );
	},

	_close: function( event ) {
		if ( this.menu.element.is( ":visible" ) ) {
			this.menu.element.hide();
			this.menu.blur();
			this.isNewMenu = true;
			this._trigger( "close", event );
		}
	},

	_change: function( event ) {
		if ( this.previous !== this._value() ) {
			this._trigger( "change", event, { item: this.selectedItem } );
		}
	},

	_normalize: function( items ) {
		// assume all items have the right format when the first item is complete
		if ( items.length && items[ 0 ].label && items[ 0 ].value ) {
			return items;
		}
		return $.map( items, function( item ) {
			if ( typeof item === "string" ) {
				return {
					label: item,
					value: item
				};
			}
			return $.extend( {}, item, {
				label: item.label || item.value,
				value: item.value || item.label
			});
		});
	},

	_suggest: function( items ) {
		var ul = this.menu.element.empty();
		this._renderMenu( ul, items );
		this.isNewMenu = true;
		this.menu.refresh();

		// size and position menu
		ul.show();
		this._resizeMenu();
		ul.position( $.extend({
			of: this.element
		}, this.options.position ) );

		if ( this.options.autoFocus ) {
			this.menu.next();
		}
	},

	_resizeMenu: function() {
		var ul = this.menu.element;
		ul.outerWidth( Math.max(
			// Firefox wraps long text (possibly a rounding bug)
			// so we add 1px to avoid the wrapping (#7513)
			ul.width( "" ).outerWidth() + 1,
			this.element.outerWidth()
		) );
	},

	_renderMenu: function( ul, items ) {
		var that = this;
		$.each( items, function( index, item ) {
			that._renderItemData( ul, item );
		});
	},

	_renderItemData: function( ul, item ) {
		return this._renderItem( ul, item ).data( "ui-autocomplete-item", item );
	},

	_renderItem: function( ul, item ) {
		return $( "<li>" ).text( item.label ).appendTo( ul );
	},

	_move: function( direction, event ) {
		if ( !this.menu.element.is( ":visible" ) ) {
			this.search( null, event );
			return;
		}
		if ( this.menu.isFirstItem() && /^previous/.test( direction ) ||
				this.menu.isLastItem() && /^next/.test( direction ) ) {

			if ( !this.isMultiLine ) {
				this._value( this.term );
			}

			this.menu.blur();
			return;
		}
		this.menu[ direction ]( event );
	},

	widget: function() {
		return this.menu.element;
	},

	_value: function() {
		return this.valueMethod.apply( this.element, arguments );
	},

	_keyEvent: function( keyEvent, event ) {
		if ( !this.isMultiLine || this.menu.element.is( ":visible" ) ) {
			this._move( keyEvent, event );

			// prevents moving cursor to beginning/end of the text field in some browsers
			event.preventDefault();
		}
	}
});

$.extend( $.ui.autocomplete, {
	escapeRegex: function( value ) {
		return value.replace( /[\-\[\]{}()*+?.,\\\^$|#\s]/g, "\\$&" );
	},
	filter: function( array, term ) {
		var matcher = new RegExp( $.ui.autocomplete.escapeRegex( term ), "i" );
		return $.grep( array, function( value ) {
			return matcher.test( value.label || value.value || value );
		});
	}
});

// live region extension, adding a `messages` option
// NOTE: This is an experimental API. We are still investigating
// a full solution for string manipulation and internationalization.
$.widget( "ui.autocomplete", $.ui.autocomplete, {
	options: {
		messages: {
			noResults: "No search results.",
			results: function( amount ) {
				return amount + ( amount > 1 ? " results are" : " result is" ) +
					" available, use up and down arrow keys to navigate.";
			}
		}
	},

	__response: function( content ) {
		var message;
		this._superApply( arguments );
		if ( this.options.disabled || this.cancelSearch ) {
			return;
		}
		if ( content && content.length ) {
			message = this.options.messages.results( content.length );
		} else {
			message = this.options.messages.noResults;
		}
		this.liveRegion.children().hide();
		$( "<div>" ).text( message ).appendTo( this.liveRegion );
	}
});

var autocomplete = $.ui.autocomplete;


/*!
 * jQuery UI Button 1.11.4
 * http://jqueryui.com
 *
 * Copyright jQuery Foundation and other contributors
 * Released under the MIT license.
 * http://jquery.org/license
 *
 * http://api.jqueryui.com/button/
 */


var lastActive,
	baseClasses = "ui-button ui-widget ui-state-default ui-corner-all",
	typeClasses = "ui-button-icons-only ui-button-icon-only ui-button-text-icons ui-button-text-icon-primary ui-button-text-icon-secondary ui-button-text-only",
	formResetHandler = function() {
		var form = $( this );
		setTimeout(function() {
			form.find( ":ui-button" ).button( "refresh" );
		}, 1 );
	},
	radioGroup = function( radio ) {
		var name = radio.name,
			form = radio.form,
			radios = $( [] );
		if ( name ) {
			name = name.replace( /'/g, "\\'" );
			if ( form ) {
				radios = $( form ).find( "[name='" + name + "'][type=radio]" );
			} else {
				radios = $( "[name='" + name + "'][type=radio]", radio.ownerDocument )
					.filter(function() {
						return !this.form;
					});
			}
		}
		return radios;
	};

$.widget( "ui.button", {
	version: "1.11.4",
	defaultElement: "<button>",
	options: {
		disabled: null,
		text: true,
		label: null,
		icons: {
			primary: null,
			secondary: null
		}
	},
	_create: function() {
		this.element.closest( "form" )
			.unbind( "reset" + this.eventNamespace )
			.bind( "reset" + this.eventNamespace, formResetHandler );

		if ( typeof this.options.disabled !== "boolean" ) {
			this.options.disabled = !!this.element.prop( "disabled" );
		} else {
			this.element.prop( "disabled", this.options.disabled );
		}

		this._determineButtonType();
		this.hasTitle = !!this.buttonElement.attr( "title" );

		var that = this,
			options = this.options,
			toggleButton = this.type === "checkbox" || this.type === "radio",
			activeClass = !toggleButton ? "ui-state-active" : "";

		if ( options.label === null ) {
			options.label = (this.type === "input" ? this.buttonElement.val() : this.buttonElement.html());
		}

		this._hoverable( this.buttonElement );

		this.buttonElement
			.addClass( baseClasses )
			.attr( "role", "button" )
			.bind( "mouseenter" + this.eventNamespace, function() {
				if ( options.disabled ) {
					return;
				}
				if ( this === lastActive ) {
					$( this ).addClass( "ui-state-active" );
				}
			})
			.bind( "mouseleave" + this.eventNamespace, function() {
				if ( options.disabled ) {
					return;
				}
				$( this ).removeClass( activeClass );
			})
			.bind( "click" + this.eventNamespace, function( event ) {
				if ( options.disabled ) {
					event.preventDefault();
					event.stopImmediatePropagation();
				}
			});

		// Can't use _focusable() because the element that receives focus
		// and the element that gets the ui-state-focus class are different
		this._on({
			focus: function() {
				this.buttonElement.addClass( "ui-state-focus" );
			},
			blur: function() {
				this.buttonElement.removeClass( "ui-state-focus" );
			}
		});

		if ( toggleButton ) {
			this.element.bind( "change" + this.eventNamespace, function() {
				that.refresh();
			});
		}

		if ( this.type === "checkbox" ) {
			this.buttonElement.bind( "click" + this.eventNamespace, function() {
				if ( options.disabled ) {
					return false;
				}
			});
		} else if ( this.type === "radio" ) {
			this.buttonElement.bind( "click" + this.eventNamespace, function() {
				if ( options.disabled ) {
					return false;
				}
				$( this ).addClass( "ui-state-active" );
				that.buttonElement.attr( "aria-pressed", "true" );

				var radio = that.element[ 0 ];
				radioGroup( radio )
					.not( radio )
					.map(function() {
						return $( this ).button( "widget" )[ 0 ];
					})
					.removeClass( "ui-state-active" )
					.attr( "aria-pressed", "false" );
			});
		} else {
			this.buttonElement
				.bind( "mousedown" + this.eventNamespace, function() {
					if ( options.disabled ) {
						return false;
					}
					$( this ).addClass( "ui-state-active" );
					lastActive = this;
					that.document.one( "mouseup", function() {
						lastActive = null;
					});
				})
				.bind( "mouseup" + this.eventNamespace, function() {
					if ( options.disabled ) {
						return false;
					}
					$( this ).removeClass( "ui-state-active" );
				})
				.bind( "keydown" + this.eventNamespace, function(event) {
					if ( options.disabled ) {
						return false;
					}
					if ( event.keyCode === $.ui.keyCode.SPACE || event.keyCode === $.ui.keyCode.ENTER ) {
						$( this ).addClass( "ui-state-active" );
					}
				})
				// see #8559, we bind to blur here in case the button element loses
				// focus between keydown and keyup, it would be left in an "active" state
				.bind( "keyup" + this.eventNamespace + " blur" + this.eventNamespace, function() {
					$( this ).removeClass( "ui-state-active" );
				});

			if ( this.buttonElement.is("a") ) {
				this.buttonElement.keyup(function(event) {
					if ( event.keyCode === $.ui.keyCode.SPACE ) {
						// TODO pass through original event correctly (just as 2nd argument doesn't work)
						$( this ).click();
					}
				});
			}
		}

		this._setOption( "disabled", options.disabled );
		this._resetButton();
	},

	_determineButtonType: function() {
		var ancestor, labelSelector, checked;

		if ( this.element.is("[type=checkbox]") ) {
			this.type = "checkbox";
		} else if ( this.element.is("[type=radio]") ) {
			this.type = "radio";
		} else if ( this.element.is("input") ) {
			this.type = "input";
		} else {
			this.type = "button";
		}

		if ( this.type === "checkbox" || this.type === "radio" ) {
			// we don't search against the document in case the element
			// is disconnected from the DOM
			ancestor = this.element.parents().last();
			labelSelector = "label[for='" + this.element.attr("id") + "']";
			this.buttonElement = ancestor.find( labelSelector );
			if ( !this.buttonElement.length ) {
				ancestor = ancestor.length ? ancestor.siblings() : this.element.siblings();
				this.buttonElement = ancestor.filter( labelSelector );
				if ( !this.buttonElement.length ) {
					this.buttonElement = ancestor.find( labelSelector );
				}
			}
			this.element.addClass( "ui-helper-hidden-accessible" );

			checked = this.element.is( ":checked" );
			if ( checked ) {
				this.buttonElement.addClass( "ui-state-active" );
			}
			this.buttonElement.prop( "aria-pressed", checked );
		} else {
			this.buttonElement = this.element;
		}
	},

	widget: function() {
		return this.buttonElement;
	},

	_destroy: function() {
		this.element
			.removeClass( "ui-helper-hidden-accessible" );
		this.buttonElement
			.removeClass( baseClasses + " ui-state-active " + typeClasses )
			.removeAttr( "role" )
			.removeAttr( "aria-pressed" )
			.html( this.buttonElement.find(".ui-button-text").html() );

		if ( !this.hasTitle ) {
			this.buttonElement.removeAttr( "title" );
		}
	},

	_setOption: function( key, value ) {
		this._super( key, value );
		if ( key === "disabled" ) {
			this.widget().toggleClass( "ui-state-disabled", !!value );
			this.element.prop( "disabled", !!value );
			if ( value ) {
				if ( this.type === "checkbox" || this.type === "radio" ) {
					this.buttonElement.removeClass( "ui-state-focus" );
				} else {
					this.buttonElement.removeClass( "ui-state-focus ui-state-active" );
				}
			}
			return;
		}
		this._resetButton();
	},

	refresh: function() {
		//See #8237 & #8828
		var isDisabled = this.element.is( "input, button" ) ? this.element.is( ":disabled" ) : this.element.hasClass( "ui-button-disabled" );

		if ( isDisabled !== this.options.disabled ) {
			this._setOption( "disabled", isDisabled );
		}
		if ( this.type === "radio" ) {
			radioGroup( this.element[0] ).each(function() {
				if ( $( this ).is( ":checked" ) ) {
					$( this ).button( "widget" )
						.addClass( "ui-state-active" )
						.attr( "aria-pressed", "true" );
				} else {
					$( this ).button( "widget" )
						.removeClass( "ui-state-active" )
						.attr( "aria-pressed", "false" );
				}
			});
		} else if ( this.type === "checkbox" ) {
			if ( this.element.is( ":checked" ) ) {
				this.buttonElement
					.addClass( "ui-state-active" )
					.attr( "aria-pressed", "true" );
			} else {
				this.buttonElement
					.removeClass( "ui-state-active" )
					.attr( "aria-pressed", "false" );
			}
		}
	},

	_resetButton: function() {
		if ( this.type === "input" ) {
			if ( this.options.label ) {
				this.element.val( this.options.label );
			}
			return;
		}
		var buttonElement = this.buttonElement.removeClass( typeClasses ),
			buttonText = $( "<span></span>", this.document[0] )
				.addClass( "ui-button-text" )
				.html( this.options.label )
				.appendTo( buttonElement.empty() )
				.text(),
			icons = this.options.icons,
			multipleIcons = icons.primary && icons.secondary,
			buttonClasses = [];

		if ( icons.primary || icons.secondary ) {
			if ( this.options.text ) {
				buttonClasses.push( "ui-button-text-icon" + ( multipleIcons ? "s" : ( icons.primary ? "-primary" : "-secondary" ) ) );
			}

			if ( icons.primary ) {
				buttonElement.prepend( "<span class='ui-button-icon-primary ui-icon " + icons.primary + "'></span>" );
			}

			if ( icons.secondary ) {
				buttonElement.append( "<span class='ui-button-icon-secondary ui-icon " + icons.secondary + "'></span>" );
			}

			if ( !this.options.text ) {
				buttonClasses.push( multipleIcons ? "ui-button-icons-only" : "ui-button-icon-only" );

				if ( !this.hasTitle ) {
					buttonElement.attr( "title", $.trim( buttonText ) );
				}
			}
		} else {
			buttonClasses.push( "ui-button-text-only" );
		}
		buttonElement.addClass( buttonClasses.join( " " ) );
	}
});

$.widget( "ui.buttonset", {
	version: "1.11.4",
	options: {
		items: "button, input[type=button], input[type=submit], input[type=reset], input[type=checkbox], input[type=radio], a, :data(ui-button)"
	},

	_create: function() {
		this.element.addClass( "ui-buttonset" );
	},

	_init: function() {
		this.refresh();
	},

	_setOption: function( key, value ) {
		if ( key === "disabled" ) {
			this.buttons.button( "option", key, value );
		}

		this._super( key, value );
	},

	refresh: function() {
		var rtl = this.element.css( "direction" ) === "rtl",
			allButtons = this.element.find( this.options.items ),
			existingButtons = allButtons.filter( ":ui-button" );

		// Initialize new buttons
		allButtons.not( ":ui-button" ).button();

		// Refresh existing buttons
		existingButtons.button( "refresh" );

		this.buttons = allButtons
			.map(function() {
				return $( this ).button( "widget" )[ 0 ];
			})
				.removeClass( "ui-corner-all ui-corner-left ui-corner-right" )
				.filter( ":first" )
					.addClass( rtl ? "ui-corner-right" : "ui-corner-left" )
				.end()
				.filter( ":last" )
					.addClass( rtl ? "ui-corner-left" : "ui-corner-right" )
				.end()
			.end();
	},

	_destroy: function() {
		this.element.removeClass( "ui-buttonset" );
		this.buttons
			.map(function() {
				return $( this ).button( "widget" )[ 0 ];
			})
				.removeClass( "ui-corner-left ui-corner-right" )
			.end()
			.button( "destroy" );
	}
});

var button = $.ui.button;


/*!
 * jQuery UI Dialog 1.11.4
 * http://jqueryui.com
 *
 * Copyright jQuery Foundation and other contributors
 * Released under the MIT license.
 * http://jquery.org/license
 *
 * http://api.jqueryui.com/dialog/
 */


var dialog = $.widget( "ui.dialog", {
	version: "1.11.4",
	options: {
		appendTo: "body",
		autoOpen: true,
		buttons: [],
		closeOnEscape: true,
		closeText: "Close",
		dialogClass: "",
		draggable: true,
		hide: null,
		height: "auto",
		maxHeight: null,
		maxWidth: null,
		minHeight: 150,
		minWidth: 150,
		modal: false,
		position: {
			my: "center",
			at: "center",
			of: window,
			collision: "fit",
			// Ensure the titlebar is always visible
			using: function( pos ) {
				var topOffset = $( this ).css( pos ).offset().top;
				if ( topOffset < 0 ) {
					$( this ).css( "top", pos.top - topOffset );
				}
			}
		},
		resizable: true,
		show: null,
		title: null,
		width: 300,

		// callbacks
		beforeClose: null,
		close: null,
		drag: null,
		dragStart: null,
		dragStop: null,
		focus: null,
		open: null,
		resize: null,
		resizeStart: null,
		resizeStop: null
	},

	sizeRelatedOptions: {
		buttons: true,
		height: true,
		maxHeight: true,
		maxWidth: true,
		minHeight: true,
		minWidth: true,
		width: true
	},

	resizableRelatedOptions: {
		maxHeight: true,
		maxWidth: true,
		minHeight: true,
		minWidth: true
	},

	_create: function() {
		this.originalCss = {
			display: this.element[ 0 ].style.display,
			width: this.element[ 0 ].style.width,
			minHeight: this.element[ 0 ].style.minHeight,
			maxHeight: this.element[ 0 ].style.maxHeight,
			height: this.element[ 0 ].style.height
		};
		this.originalPosition = {
			parent: this.element.parent(),
			index: this.element.parent().children().index( this.element )
		};
		this.originalTitle = this.element.attr( "title" );
		this.options.title = this.options.title || this.originalTitle;

		this._createWrapper();

		this.element
			.show()
			.removeAttr( "title" )
			.addClass( "ui-dialog-content ui-widget-content" )
			.appendTo( this.uiDialog );

		this._createTitlebar();
		this._createButtonPane();

		if ( this.options.draggable && $.fn.draggable ) {
			this._makeDraggable();
		}
		if ( this.options.resizable && $.fn.resizable ) {
			this._makeResizable();
		}

		this._isOpen = false;

		this._trackFocus();
	},

	_init: function() {
		if ( this.options.autoOpen ) {
			this.open();
		}
	},

	_appendTo: function() {
		var element = this.options.appendTo;
		if ( element && (element.jquery || element.nodeType) ) {
			return $( element );
		}
		return this.document.find( element || "body" ).eq( 0 );
	},

	_destroy: function() {
		var next,
			originalPosition = this.originalPosition;

		this._untrackInstance();
		this._destroyOverlay();

		this.element
			.removeUniqueId()
			.removeClass( "ui-dialog-content ui-widget-content" )
			.css( this.originalCss )
			// Without detaching first, the following becomes really slow
			.detach();

		this.uiDialog.stop( true, true ).remove();

		if ( this.originalTitle ) {
			this.element.attr( "title", this.originalTitle );
		}

		next = originalPosition.parent.children().eq( originalPosition.index );
		// Don't try to place the dialog next to itself (#8613)
		if ( next.length && next[ 0 ] !== this.element[ 0 ] ) {
			next.before( this.element );
		} else {
			originalPosition.parent.append( this.element );
		}
	},

	widget: function() {
		return this.uiDialog;
	},

	disable: $.noop,
	enable: $.noop,

	close: function( event ) {
		var activeElement,
			that = this;

		if ( !this._isOpen || this._trigger( "beforeClose", event ) === false ) {
			return;
		}

		this._isOpen = false;
		this._focusedElement = null;
		this._destroyOverlay();
		this._untrackInstance();

		if ( !this.opener.filter( ":focusable" ).focus().length ) {

			// support: IE9
			// IE9 throws an "Unspecified error" accessing document.activeElement from an <iframe>
			try {
				activeElement = this.document[ 0 ].activeElement;

				// Support: IE9, IE10
				// If the <body> is blurred, IE will switch windows, see #4520
				if ( activeElement && activeElement.nodeName.toLowerCase() !== "body" ) {

					// Hiding a focused element doesn't trigger blur in WebKit
					// so in case we have nothing to focus on, explicitly blur the active element
					// https://bugs.webkit.org/show_bug.cgi?id=47182
					$( activeElement ).blur();
				}
			} catch ( error ) {}
		}

		this._hide( this.uiDialog, this.options.hide, function() {
			that._trigger( "close", event );
		});
	},

	isOpen: function() {
		return this._isOpen;
	},

	moveToTop: function() {
		this._moveToTop();
	},

	_moveToTop: function( event, silent ) {
		var moved = false,
			zIndices = this.uiDialog.siblings( ".ui-front:visible" ).map(function() {
				return +$( this ).css( "z-index" );
			}).get(),
			zIndexMax = Math.max.apply( null, zIndices );

		if ( zIndexMax >= +this.uiDialog.css( "z-index" ) ) {
			this.uiDialog.css( "z-index", zIndexMax + 1 );
			moved = true;
		}

		if ( moved && !silent ) {
			this._trigger( "focus", event );
		}
		return moved;
	},

	open: function() {
		var that = this;
		if ( this._isOpen ) {
			if ( this._moveToTop() ) {
				this._focusTabbable();
			}
			return;
		}

		this._isOpen = true;
		this.opener = $( this.document[ 0 ].activeElement );

		this._size();
		this._position();
		this._createOverlay();
		this._moveToTop( null, true );

		// Ensure the overlay is moved to the top with the dialog, but only when
		// opening. The overlay shouldn't move after the dialog is open so that
		// modeless dialogs opened after the modal dialog stack properly.
		if ( this.overlay ) {
			this.overlay.css( "z-index", this.uiDialog.css( "z-index" ) - 1 );
		}

		this._show( this.uiDialog, this.options.show, function() {
			that._focusTabbable();
			that._trigger( "focus" );
		});

		// Track the dialog immediately upon openening in case a focus event
		// somehow occurs outside of the dialog before an element inside the
		// dialog is focused (#10152)
		this._makeFocusTarget();

		this._trigger( "open" );
	},

	_focusTabbable: function() {
		// Set focus to the first match:
		// 1. An element that was focused previously
		// 2. First element inside the dialog matching [autofocus]
		// 3. Tabbable element inside the content element
		// 4. Tabbable element inside the buttonpane
		// 5. The close button
		// 6. The dialog itself
		var hasFocus = this._focusedElement;
		if ( !hasFocus ) {
			hasFocus = this.element.find( "[autofocus]" );
		}
		if ( !hasFocus.length ) {
			hasFocus = this.element.find( ":tabbable" );
		}
		if ( !hasFocus.length ) {
			hasFocus = this.uiDialogButtonPane.find( ":tabbable" );
		}
		if ( !hasFocus.length ) {
			hasFocus = this.uiDialogTitlebarClose.filter( ":tabbable" );
		}
		if ( !hasFocus.length ) {
			hasFocus = this.uiDialog;
		}
		hasFocus.eq( 0 ).focus();
	},

	_keepFocus: function( event ) {
		function checkFocus() {
			var activeElement = this.document[0].activeElement,
				isActive = this.uiDialog[0] === activeElement ||
					$.contains( this.uiDialog[0], activeElement );
			if ( !isActive ) {
				this._focusTabbable();
			}
		}
		event.preventDefault();
		checkFocus.call( this );
		// support: IE
		// IE <= 8 doesn't prevent moving focus even with event.preventDefault()
		// so we check again later
		this._delay( checkFocus );
	},

	_createWrapper: function() {
		this.uiDialog = $("<div>")
			.addClass( "ui-dialog ui-widget ui-widget-content ui-corner-all ui-front " +
				this.options.dialogClass )
			.hide()
			.attr({
				// Setting tabIndex makes the div focusable
				tabIndex: -1,
				role: "dialog"
			})
			.appendTo( this._appendTo() );

		this._on( this.uiDialog, {
			keydown: function( event ) {
				if ( this.options.closeOnEscape && !event.isDefaultPrevented() && event.keyCode &&
						event.keyCode === $.ui.keyCode.ESCAPE ) {
					event.preventDefault();
					this.close( event );
					return;
				}

				// prevent tabbing out of dialogs
				if ( event.keyCode !== $.ui.keyCode.TAB || event.isDefaultPrevented() ) {
					return;
				}
				var tabbables = this.uiDialog.find( ":tabbable" ),
					first = tabbables.filter( ":first" ),
					last = tabbables.filter( ":last" );

				if ( ( event.target === last[0] || event.target === this.uiDialog[0] ) && !event.shiftKey ) {
					this._delay(function() {
						first.focus();
					});
					event.preventDefault();
				} else if ( ( event.target === first[0] || event.target === this.uiDialog[0] ) && event.shiftKey ) {
					this._delay(function() {
						last.focus();
					});
					event.preventDefault();
				}
			},
			mousedown: function( event ) {
				if ( this._moveToTop( event ) ) {
					this._focusTabbable();
				}
			}
		});

		// We assume that any existing aria-describedby attribute means
		// that the dialog content is marked up properly
		// otherwise we brute force the content as the description
		if ( !this.element.find( "[aria-describedby]" ).length ) {
			this.uiDialog.attr({
				"aria-describedby": this.element.uniqueId().attr( "id" )
			});
		}
	},

	_createTitlebar: function() {
		var uiDialogTitle;

		this.uiDialogTitlebar = $( "<div>" )
			.addClass( "ui-dialog-titlebar ui-widget-header ui-corner-all ui-helper-clearfix" )
			.prependTo( this.uiDialog );
		this._on( this.uiDialogTitlebar, {
			mousedown: function( event ) {
				// Don't prevent click on close button (#8838)
				// Focusing a dialog that is partially scrolled out of view
				// causes the browser to scroll it into view, preventing the click event
				if ( !$( event.target ).closest( ".ui-dialog-titlebar-close" ) ) {
					// Dialog isn't getting focus when dragging (#8063)
					this.uiDialog.focus();
				}
			}
		});

		// support: IE
		// Use type="button" to prevent enter keypresses in textboxes from closing the
		// dialog in IE (#9312)
		this.uiDialogTitlebarClose = $( "<button type='button'></button>" )
			.button({
				label: this.options.closeText,
				icons: {
					primary: "ui-icon-closethick"
				},
				text: false
			})
			.addClass( "ui-dialog-titlebar-close" )
			.appendTo( this.uiDialogTitlebar );
		this._on( this.uiDialogTitlebarClose, {
			click: function( event ) {
				event.preventDefault();
				this.close( event );
			}
		});

		uiDialogTitle = $( "<span>" )
			.uniqueId()
			.addClass( "ui-dialog-title" )
			.prependTo( this.uiDialogTitlebar );
		this._title( uiDialogTitle );

		this.uiDialog.attr({
			"aria-labelledby": uiDialogTitle.attr( "id" )
		});
	},

	_title: function( title ) {
		if ( !this.options.title ) {
			title.html( "&#160;" );
		}
		title.text( this.options.title );
	},

	_createButtonPane: function() {
		this.uiDialogButtonPane = $( "<div>" )
			.addClass( "ui-dialog-buttonpane ui-widget-content ui-helper-clearfix" );

		this.uiButtonSet = $( "<div>" )
			.addClass( "ui-dialog-buttonset" )
			.appendTo( this.uiDialogButtonPane );

		this._createButtons();
	},

	_createButtons: function() {
		var that = this,
			buttons = this.options.buttons;

		// if we already have a button pane, remove it
		this.uiDialogButtonPane.remove();
		this.uiButtonSet.empty();

		if ( $.isEmptyObject( buttons ) || ($.isArray( buttons ) && !buttons.length) ) {
			this.uiDialog.removeClass( "ui-dialog-buttons" );
			return;
		}

		$.each( buttons, function( name, props ) {
			var click, buttonOptions;
			props = $.isFunction( props ) ?
				{ click: props, text: name } :
				props;
			// Default to a non-submitting button
			props = $.extend( { type: "button" }, props );
			// Change the context for the click callback to be the main element
			click = props.click;
			props.click = function() {
				click.apply( that.element[ 0 ], arguments );
			};
			buttonOptions = {
				icons: props.icons,
				text: props.showText
			};
			delete props.icons;
			delete props.showText;
			$( "<button></button>", props )
				.button( buttonOptions )
				.appendTo( that.uiButtonSet );
		});
		this.uiDialog.addClass( "ui-dialog-buttons" );
		this.uiDialogButtonPane.appendTo( this.uiDialog );
	},

	_makeDraggable: function() {
		var that = this,
			options = this.options;

		function filteredUi( ui ) {
			return {
				position: ui.position,
				offset: ui.offset
			};
		}

		this.uiDialog.draggable({
			cancel: ".ui-dialog-content, .ui-dialog-titlebar-close",
			handle: ".ui-dialog-titlebar",
			containment: "document",
			start: function( event, ui ) {
				$( this ).addClass( "ui-dialog-dragging" );
				that._blockFrames();
				that._trigger( "dragStart", event, filteredUi( ui ) );
			},
			drag: function( event, ui ) {
				that._trigger( "drag", event, filteredUi( ui ) );
			},
			stop: function( event, ui ) {
				var left = ui.offset.left - that.document.scrollLeft(),
					top = ui.offset.top - that.document.scrollTop();

				options.position = {
					my: "left top",
					at: "left" + (left >= 0 ? "+" : "") + left + " " +
						"top" + (top >= 0 ? "+" : "") + top,
					of: that.window
				};
				$( this ).removeClass( "ui-dialog-dragging" );
				that._unblockFrames();
				that._trigger( "dragStop", event, filteredUi( ui ) );
			}
		});
	},

	_makeResizable: function() {
		var that = this,
			options = this.options,
			handles = options.resizable,
			// .ui-resizable has position: relative defined in the stylesheet
			// but dialogs have to use absolute or fixed positioning
			position = this.uiDialog.css("position"),
			resizeHandles = typeof handles === "string" ?
				handles	:
				"n,e,s,w,se,sw,ne,nw";

		function filteredUi( ui ) {
			return {
				originalPosition: ui.originalPosition,
				originalSize: ui.originalSize,
				position: ui.position,
				size: ui.size
			};
		}

		this.uiDialog.resizable({
			cancel: ".ui-dialog-content",
			containment: "document",
			alsoResize: this.element,
			maxWidth: options.maxWidth,
			maxHeight: options.maxHeight,
			minWidth: options.minWidth,
			minHeight: this._minHeight(),
			handles: resizeHandles,
			start: function( event, ui ) {
				$( this ).addClass( "ui-dialog-resizing" );
				that._blockFrames();
				that._trigger( "resizeStart", event, filteredUi( ui ) );
			},
			resize: function( event, ui ) {
				that._trigger( "resize", event, filteredUi( ui ) );
			},
			stop: function( event, ui ) {
				var offset = that.uiDialog.offset(),
					left = offset.left - that.document.scrollLeft(),
					top = offset.top - that.document.scrollTop();

				options.height = that.uiDialog.height();
				options.width = that.uiDialog.width();
				options.position = {
					my: "left top",
					at: "left" + (left >= 0 ? "+" : "") + left + " " +
						"top" + (top >= 0 ? "+" : "") + top,
					of: that.window
				};
				$( this ).removeClass( "ui-dialog-resizing" );
				that._unblockFrames();
				that._trigger( "resizeStop", event, filteredUi( ui ) );
			}
		})
		.css( "position", position );
	},

	_trackFocus: function() {
		this._on( this.widget(), {
			focusin: function( event ) {
				this._makeFocusTarget();
				this._focusedElement = $( event.target );
			}
		});
	},

	_makeFocusTarget: function() {
		this._untrackInstance();
		this._trackingInstances().unshift( this );
	},

	_untrackInstance: function() {
		var instances = this._trackingInstances(),
			exists = $.inArray( this, instances );
		if ( exists !== -1 ) {
			instances.splice( exists, 1 );
		}
	},

	_trackingInstances: function() {
		var instances = this.document.data( "ui-dialog-instances" );
		if ( !instances ) {
			instances = [];
			this.document.data( "ui-dialog-instances", instances );
		}
		return instances;
	},

	_minHeight: function() {
		var options = this.options;

		return options.height === "auto" ?
			options.minHeight :
			Math.min( options.minHeight, options.height );
	},

	_position: function() {
		// Need to show the dialog to get the actual offset in the position plugin
		var isVisible = this.uiDialog.is( ":visible" );
		if ( !isVisible ) {
			this.uiDialog.show();
		}
		this.uiDialog.position( this.options.position );
		if ( !isVisible ) {
			this.uiDialog.hide();
		}
	},

	_setOptions: function( options ) {
		var that = this,
			resize = false,
			resizableOptions = {};

		$.each( options, function( key, value ) {
			that._setOption( key, value );

			if ( key in that.sizeRelatedOptions ) {
				resize = true;
			}
			if ( key in that.resizableRelatedOptions ) {
				resizableOptions[ key ] = value;
			}
		});

		if ( resize ) {
			this._size();
			this._position();
		}
		if ( this.uiDialog.is( ":data(ui-resizable)" ) ) {
			this.uiDialog.resizable( "option", resizableOptions );
		}
	},

	_setOption: function( key, value ) {
		var isDraggable, isResizable,
			uiDialog = this.uiDialog;

		if ( key === "dialogClass" ) {
			uiDialog
				.removeClass( this.options.dialogClass )
				.addClass( value );
		}

		if ( key === "disabled" ) {
			return;
		}

		this._super( key, value );

		if ( key === "appendTo" ) {
			this.uiDialog.appendTo( this._appendTo() );
		}

		if ( key === "buttons" ) {
			this._createButtons();
		}

		if ( key === "closeText" ) {
			this.uiDialogTitlebarClose.button({
				// Ensure that we always pass a string
				label: "" + value
			});
		}

		if ( key === "draggable" ) {
			isDraggable = uiDialog.is( ":data(ui-draggable)" );
			if ( isDraggable && !value ) {
				uiDialog.draggable( "destroy" );
			}

			if ( !isDraggable && value ) {
				this._makeDraggable();
			}
		}

		if ( key === "position" ) {
			this._position();
		}

		if ( key === "resizable" ) {
			// currently resizable, becoming non-resizable
			isResizable = uiDialog.is( ":data(ui-resizable)" );
			if ( isResizable && !value ) {
				uiDialog.resizable( "destroy" );
			}

			// currently resizable, changing handles
			if ( isResizable && typeof value === "string" ) {
				uiDialog.resizable( "option", "handles", value );
			}

			// currently non-resizable, becoming resizable
			if ( !isResizable && value !== false ) {
				this._makeResizable();
			}
		}

		if ( key === "title" ) {
			this._title( this.uiDialogTitlebar.find( ".ui-dialog-title" ) );
		}
	},

	_size: function() {
		// If the user has resized the dialog, the .ui-dialog and .ui-dialog-content
		// divs will both have width and height set, so we need to reset them
		var nonContentHeight, minContentHeight, maxContentHeight,
			options = this.options;

		// Reset content sizing
		this.element.show().css({
			width: "auto",
			minHeight: 0,
			maxHeight: "none",
			height: 0
		});

		if ( options.minWidth > options.width ) {
			options.width = options.minWidth;
		}

		// reset wrapper sizing
		// determine the height of all the non-content elements
		nonContentHeight = this.uiDialog.css({
				height: "auto",
				width: options.width
			})
			.outerHeight();
		minContentHeight = Math.max( 0, options.minHeight - nonContentHeight );
		maxContentHeight = typeof options.maxHeight === "number" ?
			Math.max( 0, options.maxHeight - nonContentHeight ) :
			"none";

		if ( options.height === "auto" ) {
			this.element.css({
				minHeight: minContentHeight,
				maxHeight: maxContentHeight,
				height: "auto"
			});
		} else {
			this.element.height( Math.max( 0, options.height - nonContentHeight ) );
		}

		if ( this.uiDialog.is( ":data(ui-resizable)" ) ) {
			this.uiDialog.resizable( "option", "minHeight", this._minHeight() );
		}
	},

	_blockFrames: function() {
		this.iframeBlocks = this.document.find( "iframe" ).map(function() {
			var iframe = $( this );

			return $( "<div>" )
				.css({
					position: "absolute",
					width: iframe.outerWidth(),
					height: iframe.outerHeight()
				})
				.appendTo( iframe.parent() )
				.offset( iframe.offset() )[0];
		});
	},

	_unblockFrames: function() {
		if ( this.iframeBlocks ) {
			this.iframeBlocks.remove();
			delete this.iframeBlocks;
		}
	},

	_allowInteraction: function( event ) {
		if ( $( event.target ).closest( ".ui-dialog" ).length ) {
			return true;
		}

		// TODO: Remove hack when datepicker implements
		// the .ui-front logic (#8989)
		return !!$( event.target ).closest( ".ui-datepicker" ).length;
	},

	_createOverlay: function() {
		if ( !this.options.modal ) {
			return;
		}

		// We use a delay in case the overlay is created from an
		// event that we're going to be cancelling (#2804)
		var isOpening = true;
		this._delay(function() {
			isOpening = false;
		});

		if ( !this.document.data( "ui-dialog-overlays" ) ) {

			// Prevent use of anchors and inputs
			// Using _on() for an event handler shared across many instances is
			// safe because the dialogs stack and must be closed in reverse order
			this._on( this.document, {
				focusin: function( event ) {
					if ( isOpening ) {
						return;
					}

					if ( !this._allowInteraction( event ) ) {
						event.preventDefault();
						this._trackingInstances()[ 0 ]._focusTabbable();
					}
				}
			});
		}

		this.overlay = $( "<div>" )
			.addClass( "ui-widget-overlay ui-front" )
			.appendTo( this._appendTo() );
		this._on( this.overlay, {
			mousedown: "_keepFocus"
		});
		this.document.data( "ui-dialog-overlays",
			(this.document.data( "ui-dialog-overlays" ) || 0) + 1 );
	},

	_destroyOverlay: function() {
		if ( !this.options.modal ) {
			return;
		}

		if ( this.overlay ) {
			var overlays = this.document.data( "ui-dialog-overlays" ) - 1;

			if ( !overlays ) {
				this.document
					.unbind( "focusin" )
					.removeData( "ui-dialog-overlays" );
			} else {
				this.document.data( "ui-dialog-overlays", overlays );
			}

			this.overlay.remove();
			this.overlay = null;
		}
	}
});


/*!
 * jQuery UI Progressbar 1.11.4
 * http://jqueryui.com
 *
 * Copyright jQuery Foundation and other contributors
 * Released under the MIT license.
 * http://jquery.org/license
 *
 * http://api.jqueryui.com/progressbar/
 */


var progressbar = $.widget( "ui.progressbar", {
	version: "1.11.4",
	options: {
		max: 100,
		value: 0,

		change: null,
		complete: null
	},

	min: 0,

	_create: function() {
		// Constrain initial value
		this.oldValue = this.options.value = this._constrainedValue();

		this.element
			.addClass( "ui-progressbar ui-widget ui-widget-content ui-corner-all" )
			.attr({
				// Only set static values, aria-valuenow and aria-valuemax are
				// set inside _refreshValue()
				role: "progressbar",
				"aria-valuemin": this.min
			});

		this.valueDiv = $( "<div class='ui-progressbar-value ui-widget-header ui-corner-left'></div>" )
			.appendTo( this.element );

		this._refreshValue();
	},

	_destroy: function() {
		this.element
			.removeClass( "ui-progressbar ui-widget ui-widget-content ui-corner-all" )
			.removeAttr( "role" )
			.removeAttr( "aria-valuemin" )
			.removeAttr( "aria-valuemax" )
			.removeAttr( "aria-valuenow" );

		this.valueDiv.remove();
	},

	value: function( newValue ) {
		if ( newValue === undefined ) {
			return this.options.value;
		}

		this.options.value = this._constrainedValue( newValue );
		this._refreshValue();
	},

	_constrainedValue: function( newValue ) {
		if ( newValue === undefined ) {
			newValue = this.options.value;
		}

		this.indeterminate = newValue === false;

		// sanitize value
		if ( typeof newValue !== "number" ) {
			newValue = 0;
		}

		return this.indeterminate ? false :
			Math.min( this.options.max, Math.max( this.min, newValue ) );
	},

	_setOptions: function( options ) {
		// Ensure "value" option is set after other values (like max)
		var value = options.value;
		delete options.value;

		this._super( options );

		this.options.value = this._constrainedValue( value );
		this._refreshValue();
	},

	_setOption: function( key, value ) {
		if ( key === "max" ) {
			// Don't allow a max less than min
			value = Math.max( this.min, value );
		}
		if ( key === "disabled" ) {
			this.element
				.toggleClass( "ui-state-disabled", !!value )
				.attr( "aria-disabled", value );
		}
		this._super( key, value );
	},

	_percentage: function() {
		return this.indeterminate ? 100 : 100 * ( this.options.value - this.min ) / ( this.options.max - this.min );
	},

	_refreshValue: function() {
		var value = this.options.value,
			percentage = this._percentage();

		this.valueDiv
			.toggle( this.indeterminate || value > this.min )
			.toggleClass( "ui-corner-right", value === this.options.max )
			.width( percentage.toFixed(0) + "%" );

		this.element.toggleClass( "ui-progressbar-indeterminate", this.indeterminate );

		if ( this.indeterminate ) {
			this.element.removeAttr( "aria-valuenow" );
			if ( !this.overlayDiv ) {
				this.overlayDiv = $( "<div class='ui-progressbar-overlay'></div>" ).appendTo( this.valueDiv );
			}
		} else {
			this.element.attr({
				"aria-valuemax": this.options.max,
				"aria-valuenow": value
			});
			if ( this.overlayDiv ) {
				this.overlayDiv.remove();
				this.overlayDiv = null;
			}
		}

		if ( this.oldValue !== value ) {
			this.oldValue = value;
			this._trigger( "change" );
		}
		if ( value === this.options.max ) {
			this._trigger( "complete" );
		}
	}
});


/*!
 * jQuery UI Selectmenu 1.11.4
 * http://jqueryui.com
 *
 * Copyright jQuery Foundation and other contributors
 * Released under the MIT license.
 * http://jquery.org/license
 *
 * http://api.jqueryui.com/selectmenu
 */


var selectmenu = $.widget( "ui.selectmenu", {
	version: "1.11.4",
	defaultElement: "<select>",
	options: {
		appendTo: null,
		disabled: null,
		icons: {
			button: "ui-icon-triangle-1-s"
		},
		position: {
			my: "left top",
			at: "left bottom",
			collision: "none"
		},
		width: null,

		// callbacks
		change: null,
		close: null,
		focus: null,
		open: null,
		select: null
	},

	_create: function() {
		var selectmenuId = this.element.uniqueId().attr( "id" );
		this.ids = {
			element: selectmenuId,
			button: selectmenuId + "-button",
			menu: selectmenuId + "-menu"
		};

		this._drawButton();
		this._drawMenu();

		if ( this.options.disabled ) {
			this.disable();
		}
	},

	_drawButton: function() {
		var that = this;

		// Associate existing label with the new button
		this.label = $( "label[for='" + this.ids.element + "']" ).attr( "for", this.ids.button );
		this._on( this.label, {
			click: function( event ) {
				this.button.focus();
				event.preventDefault();
			}
		});

		// Hide original select element
		this.element.hide();

		// Create button
		this.button = $( "<span>", {
			"class": "ui-selectmenu-button ui-widget ui-state-default ui-corner-all",
			tabindex: this.options.disabled ? -1 : 0,
			id: this.ids.button,
			role: "combobox",
			"aria-expanded": "false",
			"aria-autocomplete": "list",
			"aria-owns": this.ids.menu,
			"aria-haspopup": "true"
		})
			.insertAfter( this.element );

		$( "<span>", {
			"class": "ui-icon " + this.options.icons.button
		})
			.prependTo( this.button );

		this.buttonText = $( "<span>", {
			"class": "ui-selectmenu-text"
		})
			.appendTo( this.button );

		this._setText( this.buttonText, this.element.find( "option:selected" ).text() );
		this._resizeButton();

		this._on( this.button, this._buttonEvents );
		this.button.one( "focusin", function() {

			// Delay rendering the menu items until the button receives focus.
			// The menu may have already been rendered via a programmatic open.
			if ( !that.menuItems ) {
				that._refreshMenu();
			}
		});
		this._hoverable( this.button );
		this._focusable( this.button );
	},

	_drawMenu: function() {
		var that = this;

		// Create menu
		this.menu = $( "<ul>", {
			"aria-hidden": "true",
			"aria-labelledby": this.ids.button,
			id: this.ids.menu
		});

		// Wrap menu
		this.menuWrap = $( "<div>", {
			"class": "ui-selectmenu-menu ui-front"
		})
			.append( this.menu )
			.appendTo( this._appendTo() );

		// Initialize menu widget
		this.menuInstance = this.menu
			.menu({
				role: "listbox",
				select: function( event, ui ) {
					event.preventDefault();

					// support: IE8
					// If the item was selected via a click, the text selection
					// will be destroyed in IE
					that._setSelection();

					that._select( ui.item.data( "ui-selectmenu-item" ), event );
				},
				focus: function( event, ui ) {
					var item = ui.item.data( "ui-selectmenu-item" );

					// Prevent inital focus from firing and check if its a newly focused item
					if ( that.focusIndex != null && item.index !== that.focusIndex ) {
						that._trigger( "focus", event, { item: item } );
						if ( !that.isOpen ) {
							that._select( item, event );
						}
					}
					that.focusIndex = item.index;

					that.button.attr( "aria-activedescendant",
						that.menuItems.eq( item.index ).attr( "id" ) );
				}
			})
			.menu( "instance" );

		// Adjust menu styles to dropdown
		this.menu
			.addClass( "ui-corner-bottom" )
			.removeClass( "ui-corner-all" );

		// Don't close the menu on mouseleave
		this.menuInstance._off( this.menu, "mouseleave" );

		// Cancel the menu's collapseAll on document click
		this.menuInstance._closeOnDocumentClick = function() {
			return false;
		};

		// Selects often contain empty items, but never contain dividers
		this.menuInstance._isDivider = function() {
			return false;
		};
	},

	refresh: function() {
		this._refreshMenu();
		this._setText( this.buttonText, this._getSelectedItem().text() );
		if ( !this.options.width ) {
			this._resizeButton();
		}
	},

	_refreshMenu: function() {
		this.menu.empty();

		var item,
			options = this.element.find( "option" );

		if ( !options.length ) {
			return;
		}

		this._parseOptions( options );
		this._renderMenu( this.menu, this.items );

		this.menuInstance.refresh();
		this.menuItems = this.menu.find( "li" ).not( ".ui-selectmenu-optgroup" );

		item = this._getSelectedItem();

		// Update the menu to have the correct item focused
		this.menuInstance.focus( null, item );
		this._setAria( item.data( "ui-selectmenu-item" ) );

		// Set disabled state
		this._setOption( "disabled", this.element.prop( "disabled" ) );
	},

	open: function( event ) {
		if ( this.options.disabled ) {
			return;
		}

		// If this is the first time the menu is being opened, render the items
		if ( !this.menuItems ) {
			this._refreshMenu();
		} else {

			// Menu clears focus on close, reset focus to selected item
			this.menu.find( ".ui-state-focus" ).removeClass( "ui-state-focus" );
			this.menuInstance.focus( null, this._getSelectedItem() );
		}

		this.isOpen = true;
		this._toggleAttr();
		this._resizeMenu();
		this._position();

		this._on( this.document, this._documentClick );

		this._trigger( "open", event );
	},

	_position: function() {
		this.menuWrap.position( $.extend( { of: this.button }, this.options.position ) );
	},

	close: function( event ) {
		if ( !this.isOpen ) {
			return;
		}

		this.isOpen = false;
		this._toggleAttr();

		this.range = null;
		this._off( this.document );

		this._trigger( "close", event );
	},

	widget: function() {
		return this.button;
	},

	menuWidget: function() {
		return this.menu;
	},

	_renderMenu: function( ul, items ) {
		var that = this,
			currentOptgroup = "";

		$.each( items, function( index, item ) {
			if ( item.optgroup !== currentOptgroup ) {
				$( "<li>", {
					"class": "ui-selectmenu-optgroup ui-menu-divider" +
						( item.element.parent( "optgroup" ).prop( "disabled" ) ?
							" ui-state-disabled" :
							"" ),
					text: item.optgroup
				})
					.appendTo( ul );

				currentOptgroup = item.optgroup;
			}

			that._renderItemData( ul, item );
		});
	},

	_renderItemData: function( ul, item ) {
		return this._renderItem( ul, item ).data( "ui-selectmenu-item", item );
	},

	_renderItem: function( ul, item ) {
		var li = $( "<li>" );

		if ( item.disabled ) {
			li.addClass( "ui-state-disabled" );
		}
		this._setText( li, item.label );

		return li.appendTo( ul );
	},

	_setText: function( element, value ) {
		if ( value ) {
			element.text( value );
		} else {
			element.html( "&#160;" );
		}
	},

	_move: function( direction, event ) {
		var item, next,
			filter = ".ui-menu-item";

		if ( this.isOpen ) {
			item = this.menuItems.eq( this.focusIndex );
		} else {
			item = this.menuItems.eq( this.element[ 0 ].selectedIndex );
			filter += ":not(.ui-state-disabled)";
		}

		if ( direction === "first" || direction === "last" ) {
			next = item[ direction === "first" ? "prevAll" : "nextAll" ]( filter ).eq( -1 );
		} else {
			next = item[ direction + "All" ]( filter ).eq( 0 );
		}

		if ( next.length ) {
			this.menuInstance.focus( event, next );
		}
	},

	_getSelectedItem: function() {
		return this.menuItems.eq( this.element[ 0 ].selectedIndex );
	},

	_toggle: function( event ) {
		this[ this.isOpen ? "close" : "open" ]( event );
	},

	_setSelection: function() {
		var selection;

		if ( !this.range ) {
			return;
		}

		if ( window.getSelection ) {
			selection = window.getSelection();
			selection.removeAllRanges();
			selection.addRange( this.range );

		// support: IE8
		} else {
			this.range.select();
		}

		// support: IE
		// Setting the text selection kills the button focus in IE, but
		// restoring the focus doesn't kill the selection.
		this.button.focus();
	},

	_documentClick: {
		mousedown: function( event ) {
			if ( !this.isOpen ) {
				return;
			}

			if ( !$( event.target ).closest( ".ui-selectmenu-menu, #" + this.ids.button ).length ) {
				this.close( event );
			}
		}
	},

	_buttonEvents: {

		// Prevent text selection from being reset when interacting with the selectmenu (#10144)
		mousedown: function() {
			var selection;

			if ( window.getSelection ) {
				selection = window.getSelection();
				if ( selection.rangeCount ) {
					this.range = selection.getRangeAt( 0 );
				}

			// support: IE8
			} else {
				this.range = document.selection.createRange();
			}
		},

		click: function( event ) {
			this._setSelection();
			this._toggle( event );
		},

		keydown: function( event ) {
			var preventDefault = true;
			switch ( event.keyCode ) {
				case $.ui.keyCode.TAB:
				case $.ui.keyCode.ESCAPE:
					this.close( event );
					preventDefault = false;
					break;
				case $.ui.keyCode.ENTER:
					if ( this.isOpen ) {
						this._selectFocusedItem( event );
					}
					break;
				case $.ui.keyCode.UP:
					if ( event.altKey ) {
						this._toggle( event );
					} else {
						this._move( "prev", event );
					}
					break;
				case $.ui.keyCode.DOWN:
					if ( event.altKey ) {
						this._toggle( event );
					} else {
						this._move( "next", event );
					}
					break;
				case $.ui.keyCode.SPACE:
					if ( this.isOpen ) {
						this._selectFocusedItem( event );
					} else {
						this._toggle( event );
					}
					break;
				case $.ui.keyCode.LEFT:
					this._move( "prev", event );
					break;
				case $.ui.keyCode.RIGHT:
					this._move( "next", event );
					break;
				case $.ui.keyCode.HOME:
				case $.ui.keyCode.PAGE_UP:
					this._move( "first", event );
					break;
				case $.ui.keyCode.END:
				case $.ui.keyCode.PAGE_DOWN:
					this._move( "last", event );
					break;
				default:
					this.menu.trigger( event );
					preventDefault = false;
			}

			if ( preventDefault ) {
				event.preventDefault();
			}
		}
	},

	_selectFocusedItem: function( event ) {
		var item = this.menuItems.eq( this.focusIndex );
		if ( !item.hasClass( "ui-state-disabled" ) ) {
			this._select( item.data( "ui-selectmenu-item" ), event );
		}
	},

	_select: function( item, event ) {
		var oldIndex = this.element[ 0 ].selectedIndex;

		// Change native select element
		this.element[ 0 ].selectedIndex = item.index;
		this._setText( this.buttonText, item.label );
		this._setAria( item );
		this._trigger( "select", event, { item: item } );

		if ( item.index !== oldIndex ) {
			this._trigger( "change", event, { item: item } );
		}

		this.close( event );
	},

	_setAria: function( item ) {
		var id = this.menuItems.eq( item.index ).attr( "id" );

		this.button.attr({
			"aria-labelledby": id,
			"aria-activedescendant": id
		});
		this.menu.attr( "aria-activedescendant", id );
	},

	_setOption: function( key, value ) {
		if ( key === "icons" ) {
			this.button.find( "span.ui-icon" )
				.removeClass( this.options.icons.button )
				.addClass( value.button );
		}

		this._super( key, value );

		if ( key === "appendTo" ) {
			this.menuWrap.appendTo( this._appendTo() );
		}

		if ( key === "disabled" ) {
			this.menuInstance.option( "disabled", value );
			this.button
				.toggleClass( "ui-state-disabled", value )
				.attr( "aria-disabled", value );

			this.element.prop( "disabled", value );
			if ( value ) {
				this.button.attr( "tabindex", -1 );
				this.close();
			} else {
				this.button.attr( "tabindex", 0 );
			}
		}

		if ( key === "width" ) {
			this._resizeButton();
		}
	},

	_appendTo: function() {
		var element = this.options.appendTo;

		if ( element ) {
			element = element.jquery || element.nodeType ?
				$( element ) :
				this.document.find( element ).eq( 0 );
		}

		if ( !element || !element[ 0 ] ) {
			element = this.element.closest( ".ui-front" );
		}

		if ( !element.length ) {
			element = this.document[ 0 ].body;
		}

		return element;
	},

	_toggleAttr: function() {
		this.button
			.toggleClass( "ui-corner-top", this.isOpen )
			.toggleClass( "ui-corner-all", !this.isOpen )
			.attr( "aria-expanded", this.isOpen );
		this.menuWrap.toggleClass( "ui-selectmenu-open", this.isOpen );
		this.menu.attr( "aria-hidden", !this.isOpen );
	},

	_resizeButton: function() {
		var width = this.options.width;

		if ( !width ) {
			width = this.element.show().outerWidth();
			this.element.hide();
		}

		this.button.outerWidth( width );
	},

	_resizeMenu: function() {
		this.menu.outerWidth( Math.max(
			this.button.outerWidth(),

			// support: IE10
			// IE10 wraps long text (possibly a rounding bug)
			// so we add 1px to avoid the wrapping
			this.menu.width( "" ).outerWidth() + 1
		) );
	},

	_getCreateOptions: function() {
		return { disabled: this.element.prop( "disabled" ) };
	},

	_parseOptions: function( options ) {
		var data = [];
		options.each(function( index, item ) {
			var option = $( item ),
				optgroup = option.parent( "optgroup" );
			data.push({
				element: option,
				index: index,
				value: option.val(),
				label: option.text(),
				optgroup: optgroup.attr( "label" ) || "",
				disabled: optgroup.prop( "disabled" ) || option.prop( "disabled" )
			});
		});
		this.items = data;
	},

	_destroy: function() {
		this.menuWrap.remove();
		this.button.remove();
		this.element.show();
		this.element.removeUniqueId();
		this.label.attr( "for", this.ids.element );
	}
});


/*!
 * jQuery UI Slider 1.11.4
 * http://jqueryui.com
 *
 * Copyright jQuery Foundation and other contributors
 * Released under the MIT license.
 * http://jquery.org/license
 *
 * http://api.jqueryui.com/slider/
 */


var slider = $.widget( "ui.slider", $.ui.mouse, {
	version: "1.11.4",
	widgetEventPrefix: "slide",

	options: {
		animate: false,
		distance: 0,
		max: 100,
		min: 0,
		orientation: "horizontal",
		range: false,
		step: 1,
		value: 0,
		values: null,

		// callbacks
		change: null,
		slide: null,
		start: null,
		stop: null
	},

	// number of pages in a slider
	// (how many times can you page up/down to go through the whole range)
	numPages: 5,

	_create: function() {
		this._keySliding = false;
		this._mouseSliding = false;
		this._animateOff = true;
		this._handleIndex = null;
		this._detectOrientation();
		this._mouseInit();
		this._calculateNewMax();

		this.element
			.addClass( "ui-slider" +
				" ui-slider-" + this.orientation +
				" ui-widget" +
				" ui-widget-content" +
				" ui-corner-all");

		this._refresh();
		this._setOption( "disabled", this.options.disabled );

		this._animateOff = false;
	},

	_refresh: function() {
		this._createRange();
		this._createHandles();
		this._setupEvents();
		this._refreshValue();
	},

	_createHandles: function() {
		var i, handleCount,
			options = this.options,
			existingHandles = this.element.find( ".ui-slider-handle" ).addClass( "ui-state-default ui-corner-all" ),
			handle = "<span class='ui-slider-handle ui-state-default ui-corner-all' tabindex='0'></span>",
			handles = [];

		handleCount = ( options.values && options.values.length ) || 1;

		if ( existingHandles.length > handleCount ) {
			existingHandles.slice( handleCount ).remove();
			existingHandles = existingHandles.slice( 0, handleCount );
		}

		for ( i = existingHandles.length; i < handleCount; i++ ) {
			handles.push( handle );
		}

		this.handles = existingHandles.add( $( handles.join( "" ) ).appendTo( this.element ) );

		this.handle = this.handles.eq( 0 );

		this.handles.each(function( i ) {
			$( this ).data( "ui-slider-handle-index", i );
		});
	},

	_createRange: function() {
		var options = this.options,
			classes = "";

		if ( options.range ) {
			if ( options.range === true ) {
				if ( !options.values ) {
					options.values = [ this._valueMin(), this._valueMin() ];
				} else if ( options.values.length && options.values.length !== 2 ) {
					options.values = [ options.values[0], options.values[0] ];
				} else if ( $.isArray( options.values ) ) {
					options.values = options.values.slice(0);
				}
			}

			if ( !this.range || !this.range.length ) {
				this.range = $( "<div></div>" )
					.appendTo( this.element );

				classes = "ui-slider-range" +
				// note: this isn't the most fittingly semantic framework class for this element,
				// but worked best visually with a variety of themes
				" ui-widget-header ui-corner-all";
			} else {
				this.range.removeClass( "ui-slider-range-min ui-slider-range-max" )
					// Handle range switching from true to min/max
					.css({
						"left": "",
						"bottom": ""
					});
			}

			this.range.addClass( classes +
				( ( options.range === "min" || options.range === "max" ) ? " ui-slider-range-" + options.range : "" ) );
		} else {
			if ( this.range ) {
				this.range.remove();
			}
			this.range = null;
		}
	},

	_setupEvents: function() {
		this._off( this.handles );
		this._on( this.handles, this._handleEvents );
		this._hoverable( this.handles );
		this._focusable( this.handles );
	},

	_destroy: function() {
		this.handles.remove();
		if ( this.range ) {
			this.range.remove();
		}

		this.element
			.removeClass( "ui-slider" +
				" ui-slider-horizontal" +
				" ui-slider-vertical" +
				" ui-widget" +
				" ui-widget-content" +
				" ui-corner-all" );

		this._mouseDestroy();
	},

	_mouseCapture: function( event ) {
		var position, normValue, distance, closestHandle, index, allowed, offset, mouseOverHandle,
			that = this,
			o = this.options;

		if ( o.disabled ) {
			return false;
		}

		this.elementSize = {
			width: this.element.outerWidth(),
			height: this.element.outerHeight()
		};
		this.elementOffset = this.element.offset();

		position = { x: event.pageX, y: event.pageY };
		normValue = this._normValueFromMouse( position );
		distance = this._valueMax() - this._valueMin() + 1;
		this.handles.each(function( i ) {
			var thisDistance = Math.abs( normValue - that.values(i) );
			if (( distance > thisDistance ) ||
				( distance === thisDistance &&
					(i === that._lastChangedValue || that.values(i) === o.min ))) {
				distance = thisDistance;
				closestHandle = $( this );
				index = i;
			}
		});

		allowed = this._start( event, index );
		if ( allowed === false ) {
			return false;
		}
		this._mouseSliding = true;

		this._handleIndex = index;

		closestHandle
			.addClass( "ui-state-active" )
			.focus();

		offset = closestHandle.offset();
		mouseOverHandle = !$( event.target ).parents().addBack().is( ".ui-slider-handle" );
		this._clickOffset = mouseOverHandle ? { left: 0, top: 0 } : {
			left: event.pageX - offset.left - ( closestHandle.width() / 2 ),
			top: event.pageY - offset.top -
				( closestHandle.height() / 2 ) -
				( parseInt( closestHandle.css("borderTopWidth"), 10 ) || 0 ) -
				( parseInt( closestHandle.css("borderBottomWidth"), 10 ) || 0) +
				( parseInt( closestHandle.css("marginTop"), 10 ) || 0)
		};

		if ( !this.handles.hasClass( "ui-state-hover" ) ) {
			this._slide( event, index, normValue );
		}
		this._animateOff = true;
		return true;
	},

	_mouseStart: function() {
		return true;
	},

	_mouseDrag: function( event ) {
		var position = { x: event.pageX, y: event.pageY },
			normValue = this._normValueFromMouse( position );

		this._slide( event, this._handleIndex, normValue );

		return false;
	},

	_mouseStop: function( event ) {
		this.handles.removeClass( "ui-state-active" );
		this._mouseSliding = false;

		this._stop( event, this._handleIndex );
		this._change( event, this._handleIndex );

		this._handleIndex = null;
		this._clickOffset = null;
		this._animateOff = false;

		return false;
	},

	_detectOrientation: function() {
		this.orientation = ( this.options.orientation === "vertical" ) ? "vertical" : "horizontal";
	},

	_normValueFromMouse: function( position ) {
		var pixelTotal,
			pixelMouse,
			percentMouse,
			valueTotal,
			valueMouse;

		if ( this.orientation === "horizontal" ) {
			pixelTotal = this.elementSize.width;
			pixelMouse = position.x - this.elementOffset.left - ( this._clickOffset ? this._clickOffset.left : 0 );
		} else {
			pixelTotal = this.elementSize.height;
			pixelMouse = position.y - this.elementOffset.top - ( this._clickOffset ? this._clickOffset.top : 0 );
		}

		percentMouse = ( pixelMouse / pixelTotal );
		if ( percentMouse > 1 ) {
			percentMouse = 1;
		}
		if ( percentMouse < 0 ) {
			percentMouse = 0;
		}
		if ( this.orientation === "vertical" ) {
			percentMouse = 1 - percentMouse;
		}

		valueTotal = this._valueMax() - this._valueMin();
		valueMouse = this._valueMin() + percentMouse * valueTotal;

		return this._trimAlignValue( valueMouse );
	},

	_start: function( event, index ) {
		var uiHash = {
			handle: this.handles[ index ],
			value: this.value()
		};
		if ( this.options.values && this.options.values.length ) {
			uiHash.value = this.values( index );
			uiHash.values = this.values();
		}
		return this._trigger( "start", event, uiHash );
	},

	_slide: function( event, index, newVal ) {
		var otherVal,
			newValues,
			allowed;

		if ( this.options.values && this.options.values.length ) {
			otherVal = this.values( index ? 0 : 1 );

			if ( ( this.options.values.length === 2 && this.options.range === true ) &&
					( ( index === 0 && newVal > otherVal) || ( index === 1 && newVal < otherVal ) )
				) {
				newVal = otherVal;
			}

			if ( newVal !== this.values( index ) ) {
				newValues = this.values();
				newValues[ index ] = newVal;
				// A slide can be canceled by returning false from the slide callback
				allowed = this._trigger( "slide", event, {
					handle: this.handles[ index ],
					value: newVal,
					values: newValues
				} );
				otherVal = this.values( index ? 0 : 1 );
				if ( allowed !== false ) {
					this.values( index, newVal );
				}
			}
		} else {
			if ( newVal !== this.value() ) {
				// A slide can be canceled by returning false from the slide callback
				allowed = this._trigger( "slide", event, {
					handle: this.handles[ index ],
					value: newVal
				} );
				if ( allowed !== false ) {
					this.value( newVal );
				}
			}
		}
	},

	_stop: function( event, index ) {
		var uiHash = {
			handle: this.handles[ index ],
			value: this.value()
		};
		if ( this.options.values && this.options.values.length ) {
			uiHash.value = this.values( index );
			uiHash.values = this.values();
		}

		this._trigger( "stop", event, uiHash );
	},

	_change: function( event, index ) {
		if ( !this._keySliding && !this._mouseSliding ) {
			var uiHash = {
				handle: this.handles[ index ],
				value: this.value()
			};
			if ( this.options.values && this.options.values.length ) {
				uiHash.value = this.values( index );
				uiHash.values = this.values();
			}

			//store the last changed value index for reference when handles overlap
			this._lastChangedValue = index;

			this._trigger( "change", event, uiHash );
		}
	},

	value: function( newValue ) {
		if ( arguments.length ) {
			this.options.value = this._trimAlignValue( newValue );
			this._refreshValue();
			this._change( null, 0 );
			return;
		}

		return this._value();
	},

	values: function( index, newValue ) {
		var vals,
			newValues,
			i;

		if ( arguments.length > 1 ) {
			this.options.values[ index ] = this._trimAlignValue( newValue );
			this._refreshValue();
			this._change( null, index );
			return;
		}

		if ( arguments.length ) {
			if ( $.isArray( arguments[ 0 ] ) ) {
				vals = this.options.values;
				newValues = arguments[ 0 ];
				for ( i = 0; i < vals.length; i += 1 ) {
					vals[ i ] = this._trimAlignValue( newValues[ i ] );
					this._change( null, i );
				}
				this._refreshValue();
			} else {
				if ( this.options.values && this.options.values.length ) {
					return this._values( index );
				} else {
					return this.value();
				}
			}
		} else {
			return this._values();
		}
	},

	_setOption: function( key, value ) {
		var i,
			valsLength = 0;

		if ( key === "range" && this.options.range === true ) {
			if ( value === "min" ) {
				this.options.value = this._values( 0 );
				this.options.values = null;
			} else if ( value === "max" ) {
				this.options.value = this._values( this.options.values.length - 1 );
				this.options.values = null;
			}
		}

		if ( $.isArray( this.options.values ) ) {
			valsLength = this.options.values.length;
		}

		if ( key === "disabled" ) {
			this.element.toggleClass( "ui-state-disabled", !!value );
		}

		this._super( key, value );

		switch ( key ) {
			case "orientation":
				this._detectOrientation();
				this.element
					.removeClass( "ui-slider-horizontal ui-slider-vertical" )
					.addClass( "ui-slider-" + this.orientation );
				this._refreshValue();

				// Reset positioning from previous orientation
				this.handles.css( value === "horizontal" ? "bottom" : "left", "" );
				break;
			case "value":
				this._animateOff = true;
				this._refreshValue();
				this._change( null, 0 );
				this._animateOff = false;
				break;
			case "values":
				this._animateOff = true;
				this._refreshValue();
				for ( i = 0; i < valsLength; i += 1 ) {
					this._change( null, i );
				}
				this._animateOff = false;
				break;
			case "step":
			case "min":
			case "max":
				this._animateOff = true;
				this._calculateNewMax();
				this._refreshValue();
				this._animateOff = false;
				break;
			case "range":
				this._animateOff = true;
				this._refresh();
				this._animateOff = false;
				break;
		}
	},

	//internal value getter
	// _value() returns value trimmed by min and max, aligned by step
	_value: function() {
		var val = this.options.value;
		val = this._trimAlignValue( val );

		return val;
	},

	//internal values getter
	// _values() returns array of values trimmed by min and max, aligned by step
	// _values( index ) returns single value trimmed by min and max, aligned by step
	_values: function( index ) {
		var val,
			vals,
			i;

		if ( arguments.length ) {
			val = this.options.values[ index ];
			val = this._trimAlignValue( val );

			return val;
		} else if ( this.options.values && this.options.values.length ) {
			// .slice() creates a copy of the array
			// this copy gets trimmed by min and max and then returned
			vals = this.options.values.slice();
			for ( i = 0; i < vals.length; i += 1) {
				vals[ i ] = this._trimAlignValue( vals[ i ] );
			}

			return vals;
		} else {
			return [];
		}
	},

	// returns the step-aligned value that val is closest to, between (inclusive) min and max
	_trimAlignValue: function( val ) {
		if ( val <= this._valueMin() ) {
			return this._valueMin();
		}
		if ( val >= this._valueMax() ) {
			return this._valueMax();
		}
		var step = ( this.options.step > 0 ) ? this.options.step : 1,
			valModStep = (val - this._valueMin()) % step,
			alignValue = val - valModStep;

		if ( Math.abs(valModStep) * 2 >= step ) {
			alignValue += ( valModStep > 0 ) ? step : ( -step );
		}

		// Since JavaScript has problems with large floats, round
		// the final value to 5 digits after the decimal point (see #4124)
		return parseFloat( alignValue.toFixed(5) );
	},

	_calculateNewMax: function() {
		var max = this.options.max,
			min = this._valueMin(),
			step = this.options.step,
			aboveMin = Math.floor( ( +( max - min ).toFixed( this._precision() ) ) / step ) * step;
		max = aboveMin + min;
		this.max = parseFloat( max.toFixed( this._precision() ) );
	},

	_precision: function() {
		var precision = this._precisionOf( this.options.step );
		if ( this.options.min !== null ) {
			precision = Math.max( precision, this._precisionOf( this.options.min ) );
		}
		return precision;
	},

	_precisionOf: function( num ) {
		var str = num.toString(),
			decimal = str.indexOf( "." );
		return decimal === -1 ? 0 : str.length - decimal - 1;
	},

	_valueMin: function() {
		return this.options.min;
	},

	_valueMax: function() {
		return this.max;
	},

	_refreshValue: function() {
		var lastValPercent, valPercent, value, valueMin, valueMax,
			oRange = this.options.range,
			o = this.options,
			that = this,
			animate = ( !this._animateOff ) ? o.animate : false,
			_set = {};

		if ( this.options.values && this.options.values.length ) {
			this.handles.each(function( i ) {
				valPercent = ( that.values(i) - that._valueMin() ) / ( that._valueMax() - that._valueMin() ) * 100;
				_set[ that.orientation === "horizontal" ? "left" : "bottom" ] = valPercent + "%";
				$( this ).stop( 1, 1 )[ animate ? "animate" : "css" ]( _set, o.animate );
				if ( that.options.range === true ) {
					if ( that.orientation === "horizontal" ) {
						if ( i === 0 ) {
							that.range.stop( 1, 1 )[ animate ? "animate" : "css" ]( { left: valPercent + "%" }, o.animate );
						}
						if ( i === 1 ) {
							that.range[ animate ? "animate" : "css" ]( { width: ( valPercent - lastValPercent ) + "%" }, { queue: false, duration: o.animate } );
						}
					} else {
						if ( i === 0 ) {
							that.range.stop( 1, 1 )[ animate ? "animate" : "css" ]( { bottom: ( valPercent ) + "%" }, o.animate );
						}
						if ( i === 1 ) {
							that.range[ animate ? "animate" : "css" ]( { height: ( valPercent - lastValPercent ) + "%" }, { queue: false, duration: o.animate } );
						}
					}
				}
				lastValPercent = valPercent;
			});
		} else {
			value = this.value();
			valueMin = this._valueMin();
			valueMax = this._valueMax();
			valPercent = ( valueMax !== valueMin ) ?
					( value - valueMin ) / ( valueMax - valueMin ) * 100 :
					0;
			_set[ this.orientation === "horizontal" ? "left" : "bottom" ] = valPercent + "%";
			this.handle.stop( 1, 1 )[ animate ? "animate" : "css" ]( _set, o.animate );

			if ( oRange === "min" && this.orientation === "horizontal" ) {
				this.range.stop( 1, 1 )[ animate ? "animate" : "css" ]( { width: valPercent + "%" }, o.animate );
			}
			if ( oRange === "max" && this.orientation === "horizontal" ) {
				this.range[ animate ? "animate" : "css" ]( { width: ( 100 - valPercent ) + "%" }, { queue: false, duration: o.animate } );
			}
			if ( oRange === "min" && this.orientation === "vertical" ) {
				this.range.stop( 1, 1 )[ animate ? "animate" : "css" ]( { height: valPercent + "%" }, o.animate );
			}
			if ( oRange === "max" && this.orientation === "vertical" ) {
				this.range[ animate ? "animate" : "css" ]( { height: ( 100 - valPercent ) + "%" }, { queue: false, duration: o.animate } );
			}
		}
	},

	_handleEvents: {
		keydown: function( event ) {
			var allowed, curVal, newVal, step,
				index = $( event.target ).data( "ui-slider-handle-index" );

			switch ( event.keyCode ) {
				case $.ui.keyCode.HOME:
				case $.ui.keyCode.END:
				case $.ui.keyCode.PAGE_UP:
				case $.ui.keyCode.PAGE_DOWN:
				case $.ui.keyCode.UP:
				case $.ui.keyCode.RIGHT:
				case $.ui.keyCode.DOWN:
				case $.ui.keyCode.LEFT:
					event.preventDefault();
					if ( !this._keySliding ) {
						this._keySliding = true;
						$( event.target ).addClass( "ui-state-active" );
						allowed = this._start( event, index );
						if ( allowed === false ) {
							return;
						}
					}
					break;
			}

			step = this.options.step;
			if ( this.options.values && this.options.values.length ) {
				curVal = newVal = this.values( index );
			} else {
				curVal = newVal = this.value();
			}

			switch ( event.keyCode ) {
				case $.ui.keyCode.HOME:
					newVal = this._valueMin();
					break;
				case $.ui.keyCode.END:
					newVal = this._valueMax();
					break;
				case $.ui.keyCode.PAGE_UP:
					newVal = this._trimAlignValue(
						curVal + ( ( this._valueMax() - this._valueMin() ) / this.numPages )
					);
					break;
				case $.ui.keyCode.PAGE_DOWN:
					newVal = this._trimAlignValue(
						curVal - ( (this._valueMax() - this._valueMin()) / this.numPages ) );
					break;
				case $.ui.keyCode.UP:
				case $.ui.keyCode.RIGHT:
					if ( curVal === this._valueMax() ) {
						return;
					}
					newVal = this._trimAlignValue( curVal + step );
					break;
				case $.ui.keyCode.DOWN:
				case $.ui.keyCode.LEFT:
					if ( curVal === this._valueMin() ) {
						return;
					}
					newVal = this._trimAlignValue( curVal - step );
					break;
			}

			this._slide( event, index, newVal );
		},
		keyup: function( event ) {
			var index = $( event.target ).data( "ui-slider-handle-index" );

			if ( this._keySliding ) {
				this._keySliding = false;
				this._stop( event, index );
				this._change( event, index );
				$( event.target ).removeClass( "ui-state-active" );
			}
		}
	}
});


/*!
 * jQuery UI Spinner 1.11.4
 * http://jqueryui.com
 *
 * Copyright jQuery Foundation and other contributors
 * Released under the MIT license.
 * http://jquery.org/license
 *
 * http://api.jqueryui.com/spinner/
 */


function spinner_modifier( fn ) {
	return function() {
		var previous = this.element.val();
		fn.apply( this, arguments );
		this._refresh();
		if ( previous !== this.element.val() ) {
			this._trigger( "change" );
		}
	};
}

var spinner = $.widget( "ui.spinner", {
	version: "1.11.4",
	defaultElement: "<input>",
	widgetEventPrefix: "spin",
	options: {
		culture: null,
		icons: {
			down: "ui-icon-triangle-1-s",
			up: "ui-icon-triangle-1-n"
		},
		incremental: true,
		max: null,
		min: null,
		numberFormat: null,
		page: 10,
		step: 1,

		change: null,
		spin: null,
		start: null,
		stop: null
	},

	_create: function() {
		// handle string values that need to be parsed
		this._setOption( "max", this.options.max );
		this._setOption( "min", this.options.min );
		this._setOption( "step", this.options.step );

		// Only format if there is a value, prevents the field from being marked
		// as invalid in Firefox, see #9573.
		if ( this.value() !== "" ) {
			// Format the value, but don't constrain.
			this._value( this.element.val(), true );
		}

		this._draw();
		this._on( this._events );
		this._refresh();

		// turning off autocomplete prevents the browser from remembering the
		// value when navigating through history, so we re-enable autocomplete
		// if the page is unloaded before the widget is destroyed. #7790
		this._on( this.window, {
			beforeunload: function() {
				this.element.removeAttr( "autocomplete" );
			}
		});
	},

	_getCreateOptions: function() {
		var options = {},
			element = this.element;

		$.each( [ "min", "max", "step" ], function( i, option ) {
			var value = element.attr( option );
			if ( value !== undefined && value.length ) {
				options[ option ] = value;
			}
		});

		return options;
	},

	_events: {
		keydown: function( event ) {
			if ( this._start( event ) && this._keydown( event ) ) {
				event.preventDefault();
			}
		},
		keyup: "_stop",
		focus: function() {
			this.previous = this.element.val();
		},
		blur: function( event ) {
			if ( this.cancelBlur ) {
				delete this.cancelBlur;
				return;
			}

			this._stop();
			this._refresh();
			if ( this.previous !== this.element.val() ) {
				this._trigger( "change", event );
			}
		},
		mousewheel: function( event, delta ) {
			if ( !delta ) {
				return;
			}
			if ( !this.spinning && !this._start( event ) ) {
				return false;
			}

			this._spin( (delta > 0 ? 1 : -1) * this.options.step, event );
			clearTimeout( this.mousewheelTimer );
			this.mousewheelTimer = this._delay(function() {
				if ( this.spinning ) {
					this._stop( event );
				}
			}, 100 );
			event.preventDefault();
		},
		"mousedown .ui-spinner-button": function( event ) {
			var previous;

			// We never want the buttons to have focus; whenever the user is
			// interacting with the spinner, the focus should be on the input.
			// If the input is focused then this.previous is properly set from
			// when the input first received focus. If the input is not focused
			// then we need to set this.previous based on the value before spinning.
			previous = this.element[0] === this.document[0].activeElement ?
				this.previous : this.element.val();
			function checkFocus() {
				var isActive = this.element[0] === this.document[0].activeElement;
				if ( !isActive ) {
					this.element.focus();
					this.previous = previous;
					// support: IE
					// IE sets focus asynchronously, so we need to check if focus
					// moved off of the input because the user clicked on the button.
					this._delay(function() {
						this.previous = previous;
					});
				}
			}

			// ensure focus is on (or stays on) the text field
			event.preventDefault();
			checkFocus.call( this );

			// support: IE
			// IE doesn't prevent moving focus even with event.preventDefault()
			// so we set a flag to know when we should ignore the blur event
			// and check (again) if focus moved off of the input.
			this.cancelBlur = true;
			this._delay(function() {
				delete this.cancelBlur;
				checkFocus.call( this );
			});

			if ( this._start( event ) === false ) {
				return;
			}

			this._repeat( null, $( event.currentTarget ).hasClass( "ui-spinner-up" ) ? 1 : -1, event );
		},
		"mouseup .ui-spinner-button": "_stop",
		"mouseenter .ui-spinner-button": function( event ) {
			// button will add ui-state-active if mouse was down while mouseleave and kept down
			if ( !$( event.currentTarget ).hasClass( "ui-state-active" ) ) {
				return;
			}

			if ( this._start( event ) === false ) {
				return false;
			}
			this._repeat( null, $( event.currentTarget ).hasClass( "ui-spinner-up" ) ? 1 : -1, event );
		},
		// TODO: do we really want to consider this a stop?
		// shouldn't we just stop the repeater and wait until mouseup before
		// we trigger the stop event?
		"mouseleave .ui-spinner-button": "_stop"
	},

	_draw: function() {
		var uiSpinner = this.uiSpinner = this.element
			.addClass( "ui-spinner-input" )
			.attr( "autocomplete", "off" )
			.wrap( this._uiSpinnerHtml() )
			.parent()
				// add buttons
				.append( this._buttonHtml() );

		this.element.attr( "role", "spinbutton" );

		// button bindings
		this.buttons = uiSpinner.find( ".ui-spinner-button" )
			.attr( "tabIndex", -1 )
			.button()
			.removeClass( "ui-corner-all" );

		// IE 6 doesn't understand height: 50% for the buttons
		// unless the wrapper has an explicit height
		if ( this.buttons.height() > Math.ceil( uiSpinner.height() * 0.5 ) &&
				uiSpinner.height() > 0 ) {
			uiSpinner.height( uiSpinner.height() );
		}

		// disable spinner if element was already disabled
		if ( this.options.disabled ) {
			this.disable();
		}
	},

	_keydown: function( event ) {
		var options = this.options,
			keyCode = $.ui.keyCode;

		switch ( event.keyCode ) {
		case keyCode.UP:
			this._repeat( null, 1, event );
			return true;
		case keyCode.DOWN:
			this._repeat( null, -1, event );
			return true;
		case keyCode.PAGE_UP:
			this._repeat( null, options.page, event );
			return true;
		case keyCode.PAGE_DOWN:
			this._repeat( null, -options.page, event );
			return true;
		}

		return false;
	},

	_uiSpinnerHtml: function() {
		return "<span class='ui-spinner ui-widget ui-widget-content ui-corner-all'></span>";
	},

	_buttonHtml: function() {
		return "" +
			"<a class='ui-spinner-button ui-spinner-up ui-corner-tr'>" +
				"<span class='ui-icon " + this.options.icons.up + "'>&#9650;</span>" +
			"</a>" +
			"<a class='ui-spinner-button ui-spinner-down ui-corner-br'>" +
				"<span class='ui-icon " + this.options.icons.down + "'>&#9660;</span>" +
			"</a>";
	},

	_start: function( event ) {
		if ( !this.spinning && this._trigger( "start", event ) === false ) {
			return false;
		}

		if ( !this.counter ) {
			this.counter = 1;
		}
		this.spinning = true;
		return true;
	},

	_repeat: function( i, steps, event ) {
		i = i || 500;

		clearTimeout( this.timer );
		this.timer = this._delay(function() {
			this._repeat( 40, steps, event );
		}, i );

		this._spin( steps * this.options.step, event );
	},

	_spin: function( step, event ) {
		var value = this.value() || 0;

		if ( !this.counter ) {
			this.counter = 1;
		}

		value = this._adjustValue( value + step * this._increment( this.counter ) );

		if ( !this.spinning || this._trigger( "spin", event, { value: value } ) !== false) {
			this._value( value );
			this.counter++;
		}
	},

	_increment: function( i ) {
		var incremental = this.options.incremental;

		if ( incremental ) {
			return $.isFunction( incremental ) ?
				incremental( i ) :
				Math.floor( i * i * i / 50000 - i * i / 500 + 17 * i / 200 + 1 );
		}

		return 1;
	},

	_precision: function() {
		var precision = this._precisionOf( this.options.step );
		if ( this.options.min !== null ) {
			precision = Math.max( precision, this._precisionOf( this.options.min ) );
		}
		return precision;
	},

	_precisionOf: function( num ) {
		var str = num.toString(),
			decimal = str.indexOf( "." );
		return decimal === -1 ? 0 : str.length - decimal - 1;
	},

	_adjustValue: function( value ) {
		var base, aboveMin,
			options = this.options;

		// make sure we're at a valid step
		// - find out where we are relative to the base (min or 0)
		base = options.min !== null ? options.min : 0;
		aboveMin = value - base;
		// - round to the nearest step
		aboveMin = Math.round(aboveMin / options.step) * options.step;
		// - rounding is based on 0, so adjust back to our base
		value = base + aboveMin;

		// fix precision from bad JS floating point math
		value = parseFloat( value.toFixed( this._precision() ) );

		// clamp the value
		if ( options.max !== null && value > options.max) {
			return options.max;
		}
		if ( options.min !== null && value < options.min ) {
			return options.min;
		}

		return value;
	},

	_stop: function( event ) {
		if ( !this.spinning ) {
			return;
		}

		clearTimeout( this.timer );
		clearTimeout( this.mousewheelTimer );
		this.counter = 0;
		this.spinning = false;
		this._trigger( "stop", event );
	},

	_setOption: function( key, value ) {
		if ( key === "culture" || key === "numberFormat" ) {
			var prevValue = this._parse( this.element.val() );
			this.options[ key ] = value;
			this.element.val( this._format( prevValue ) );
			return;
		}

		if ( key === "max" || key === "min" || key === "step" ) {
			if ( typeof value === "string" ) {
				value = this._parse( value );
			}
		}
		if ( key === "icons" ) {
			this.buttons.first().find( ".ui-icon" )
				.removeClass( this.options.icons.up )
				.addClass( value.up );
			this.buttons.last().find( ".ui-icon" )
				.removeClass( this.options.icons.down )
				.addClass( value.down );
		}

		this._super( key, value );

		if ( key === "disabled" ) {
			this.widget().toggleClass( "ui-state-disabled", !!value );
			this.element.prop( "disabled", !!value );
			this.buttons.button( value ? "disable" : "enable" );
		}
	},

	_setOptions: spinner_modifier(function( options ) {
		this._super( options );
	}),

	_parse: function( val ) {
		if ( typeof val === "string" && val !== "" ) {
			val = window.Globalize && this.options.numberFormat ?
				Globalize.parseFloat( val, 10, this.options.culture ) : +val;
		}
		return val === "" || isNaN( val ) ? null : val;
	},

	_format: function( value ) {
		if ( value === "" ) {
			return "";
		}
		return window.Globalize && this.options.numberFormat ?
			Globalize.format( value, this.options.numberFormat, this.options.culture ) :
			value;
	},

	_refresh: function() {
		this.element.attr({
			"aria-valuemin": this.options.min,
			"aria-valuemax": this.options.max,
			// TODO: what should we do with values that can't be parsed?
			"aria-valuenow": this._parse( this.element.val() )
		});
	},

	isValid: function() {
		var value = this.value();

		// null is invalid
		if ( value === null ) {
			return false;
		}

		// if value gets adjusted, it's invalid
		return value === this._adjustValue( value );
	},

	// update the value without triggering change
	_value: function( value, allowAny ) {
		var parsed;
		if ( value !== "" ) {
			parsed = this._parse( value );
			if ( parsed !== null ) {
				if ( !allowAny ) {
					parsed = this._adjustValue( parsed );
				}
				value = this._format( parsed );
			}
		}
		this.element.val( value );
		this._refresh();
	},

	_destroy: function() {
		this.element
			.removeClass( "ui-spinner-input" )
			.prop( "disabled", false )
			.removeAttr( "autocomplete" )
			.removeAttr( "role" )
			.removeAttr( "aria-valuemin" )
			.removeAttr( "aria-valuemax" )
			.removeAttr( "aria-valuenow" );
		this.uiSpinner.replaceWith( this.element );
	},

	stepUp: spinner_modifier(function( steps ) {
		this._stepUp( steps );
	}),
	_stepUp: function( steps ) {
		if ( this._start() ) {
			this._spin( (steps || 1) * this.options.step );
			this._stop();
		}
	},

	stepDown: spinner_modifier(function( steps ) {
		this._stepDown( steps );
	}),
	_stepDown: function( steps ) {
		if ( this._start() ) {
			this._spin( (steps || 1) * -this.options.step );
			this._stop();
		}
	},

	pageUp: spinner_modifier(function( pages ) {
		this._stepUp( (pages || 1) * this.options.page );
	}),

	pageDown: spinner_modifier(function( pages ) {
		this._stepDown( (pages || 1) * this.options.page );
	}),

	value: function( newVal ) {
		if ( !arguments.length ) {
			return this._parse( this.element.val() );
		}
		spinner_modifier( this._value ).call( this, newVal );
	},

	widget: function() {
		return this.uiSpinner;
	}
});


/*!
 * jQuery UI Tabs 1.11.4
 * http://jqueryui.com
 *
 * Copyright jQuery Foundation and other contributors
 * Released under the MIT license.
 * http://jquery.org/license
 *
 * http://api.jqueryui.com/tabs/
 */


var tabs = $.widget( "ui.tabs", {
	version: "1.11.4",
	delay: 300,
	options: {
		active: null,
		collapsible: false,
		event: "click",
		heightStyle: "content",
		hide: null,
		show: null,

		// callbacks
		activate: null,
		beforeActivate: null,
		beforeLoad: null,
		load: null
	},

	_isLocal: (function() {
		var rhash = /#.*$/;

		return function( anchor ) {
			var anchorUrl, locationUrl;

			// support: IE7
			// IE7 doesn't normalize the href property when set via script (#9317)
			anchor = anchor.cloneNode( false );

			anchorUrl = anchor.href.replace( rhash, "" );
			locationUrl = location.href.replace( rhash, "" );

			// decoding may throw an error if the URL isn't UTF-8 (#9518)
			try {
				anchorUrl = decodeURIComponent( anchorUrl );
			} catch ( error ) {}
			try {
				locationUrl = decodeURIComponent( locationUrl );
			} catch ( error ) {}

			return anchor.hash.length > 1 && anchorUrl === locationUrl;
		};
	})(),

	_create: function() {
		var that = this,
			options = this.options;

		this.running = false;

		this.element
			.addClass( "ui-tabs ui-widget ui-widget-content ui-corner-all" )
			.toggleClass( "ui-tabs-collapsible", options.collapsible );

		this._processTabs();
		options.active = this._initialActive();

		// Take disabling tabs via class attribute from HTML
		// into account and update option properly.
		if ( $.isArray( options.disabled ) ) {
			options.disabled = $.unique( options.disabled.concat(
				$.map( this.tabs.filter( ".ui-state-disabled" ), function( li ) {
					return that.tabs.index( li );
				})
			) ).sort();
		}

		// check for length avoids error when initializing empty list
		if ( this.options.active !== false && this.anchors.length ) {
			this.active = this._findActive( options.active );
		} else {
			this.active = $();
		}

		this._refresh();

		if ( this.active.length ) {
			this.load( options.active );
		}
	},

	_initialActive: function() {
		var active = this.options.active,
			collapsible = this.options.collapsible,
			locationHash = location.hash.substring( 1 );

		if ( active === null ) {
			// check the fragment identifier in the URL
			if ( locationHash ) {
				this.tabs.each(function( i, tab ) {
					if ( $( tab ).attr( "aria-controls" ) === locationHash ) {
						active = i;
						return false;
					}
				});
			}

			// check for a tab marked active via a class
			if ( active === null ) {
				active = this.tabs.index( this.tabs.filter( ".ui-tabs-active" ) );
			}

			// no active tab, set to false
			if ( active === null || active === -1 ) {
				active = this.tabs.length ? 0 : false;
			}
		}

		// handle numbers: negative, out of range
		if ( active !== false ) {
			active = this.tabs.index( this.tabs.eq( active ) );
			if ( active === -1 ) {
				active = collapsible ? false : 0;
			}
		}

		// don't allow collapsible: false and active: false
		if ( !collapsible && active === false && this.anchors.length ) {
			active = 0;
		}

		return active;
	},

	_getCreateEventData: function() {
		return {
			tab: this.active,
			panel: !this.active.length ? $() : this._getPanelForTab( this.active )
		};
	},

	_tabKeydown: function( event ) {
		var focusedTab = $( this.document[0].activeElement ).closest( "li" ),
			selectedIndex = this.tabs.index( focusedTab ),
			goingForward = true;

		if ( this._handlePageNav( event ) ) {
			return;
		}

		switch ( event.keyCode ) {
			case $.ui.keyCode.RIGHT:
			case $.ui.keyCode.DOWN:
				selectedIndex++;
				break;
			case $.ui.keyCode.UP:
			case $.ui.keyCode.LEFT:
				goingForward = false;
				selectedIndex--;
				break;
			case $.ui.keyCode.END:
				selectedIndex = this.anchors.length - 1;
				break;
			case $.ui.keyCode.HOME:
				selectedIndex = 0;
				break;
			case $.ui.keyCode.SPACE:
				// Activate only, no collapsing
				event.preventDefault();
				clearTimeout( this.activating );
				this._activate( selectedIndex );
				return;
			case $.ui.keyCode.ENTER:
				// Toggle (cancel delayed activation, allow collapsing)
				event.preventDefault();
				clearTimeout( this.activating );
				// Determine if we should collapse or activate
				this._activate( selectedIndex === this.options.active ? false : selectedIndex );
				return;
			default:
				return;
		}

		// Focus the appropriate tab, based on which key was pressed
		event.preventDefault();
		clearTimeout( this.activating );
		selectedIndex = this._focusNextTab( selectedIndex, goingForward );

		// Navigating with control/command key will prevent automatic activation
		if ( !event.ctrlKey && !event.metaKey ) {

			// Update aria-selected immediately so that AT think the tab is already selected.
			// Otherwise AT may confuse the user by stating that they need to activate the tab,
			// but the tab will already be activated by the time the announcement finishes.
			focusedTab.attr( "aria-selected", "false" );
			this.tabs.eq( selectedIndex ).attr( "aria-selected", "true" );

			this.activating = this._delay(function() {
				this.option( "active", selectedIndex );
			}, this.delay );
		}
	},

	_panelKeydown: function( event ) {
		if ( this._handlePageNav( event ) ) {
			return;
		}

		// Ctrl+up moves focus to the current tab
		if ( event.ctrlKey && event.keyCode === $.ui.keyCode.UP ) {
			event.preventDefault();
			this.active.focus();
		}
	},

	// Alt+page up/down moves focus to the previous/next tab (and activates)
	_handlePageNav: function( event ) {
		if ( event.altKey && event.keyCode === $.ui.keyCode.PAGE_UP ) {
			this._activate( this._focusNextTab( this.options.active - 1, false ) );
			return true;
		}
		if ( event.altKey && event.keyCode === $.ui.keyCode.PAGE_DOWN ) {
			this._activate( this._focusNextTab( this.options.active + 1, true ) );
			return true;
		}
	},

	_findNextTab: function( index, goingForward ) {
		var lastTabIndex = this.tabs.length - 1;

		function constrain() {
			if ( index > lastTabIndex ) {
				index = 0;
			}
			if ( index < 0 ) {
				index = lastTabIndex;
			}
			return index;
		}

		while ( $.inArray( constrain(), this.options.disabled ) !== -1 ) {
			index = goingForward ? index + 1 : index - 1;
		}

		return index;
	},

	_focusNextTab: function( index, goingForward ) {
		index = this._findNextTab( index, goingForward );
		this.tabs.eq( index ).focus();
		return index;
	},

	_setOption: function( key, value ) {
		if ( key === "active" ) {
			// _activate() will handle invalid values and update this.options
			this._activate( value );
			return;
		}

		if ( key === "disabled" ) {
			// don't use the widget factory's disabled handling
			this._setupDisabled( value );
			return;
		}

		this._super( key, value);

		if ( key === "collapsible" ) {
			this.element.toggleClass( "ui-tabs-collapsible", value );
			// Setting collapsible: false while collapsed; open first panel
			if ( !value && this.options.active === false ) {
				this._activate( 0 );
			}
		}

		if ( key === "event" ) {
			this._setupEvents( value );
		}

		if ( key === "heightStyle" ) {
			this._setupHeightStyle( value );
		}
	},

	_sanitizeSelector: function( hash ) {
		return hash ? hash.replace( /[!"$%&'()*+,.\/:;<=>?@\[\]\^`{|}~]/g, "\\$&" ) : "";
	},

	refresh: function() {
		var options = this.options,
			lis = this.tablist.children( ":has(a[href])" );

		// get disabled tabs from class attribute from HTML
		// this will get converted to a boolean if needed in _refresh()
		options.disabled = $.map( lis.filter( ".ui-state-disabled" ), function( tab ) {
			return lis.index( tab );
		});

		this._processTabs();

		// was collapsed or no tabs
		if ( options.active === false || !this.anchors.length ) {
			options.active = false;
			this.active = $();
		// was active, but active tab is gone
		} else if ( this.active.length && !$.contains( this.tablist[ 0 ], this.active[ 0 ] ) ) {
			// all remaining tabs are disabled
			if ( this.tabs.length === options.disabled.length ) {
				options.active = false;
				this.active = $();
			// activate previous tab
			} else {
				this._activate( this._findNextTab( Math.max( 0, options.active - 1 ), false ) );
			}
		// was active, active tab still exists
		} else {
			// make sure active index is correct
			options.active = this.tabs.index( this.active );
		}

		this._refresh();
	},

	_refresh: function() {
		this._setupDisabled( this.options.disabled );
		this._setupEvents( this.options.event );
		this._setupHeightStyle( this.options.heightStyle );

		this.tabs.not( this.active ).attr({
			"aria-selected": "false",
			"aria-expanded": "false",
			tabIndex: -1
		});
		this.panels.not( this._getPanelForTab( this.active ) )
			.hide()
			.attr({
				"aria-hidden": "true"
			});

		// Make sure one tab is in the tab order
		if ( !this.active.length ) {
			this.tabs.eq( 0 ).attr( "tabIndex", 0 );
		} else {
			this.active
				.addClass( "ui-tabs-active ui-state-active" )
				.attr({
					"aria-selected": "true",
					"aria-expanded": "true",
					tabIndex: 0
				});
			this._getPanelForTab( this.active )
				.show()
				.attr({
					"aria-hidden": "false"
				});
		}
	},

	_processTabs: function() {
		var that = this,
			prevTabs = this.tabs,
			prevAnchors = this.anchors,
			prevPanels = this.panels;

		this.tablist = this._getList()
			.addClass( "ui-tabs-nav ui-helper-reset ui-helper-clearfix ui-widget-header ui-corner-all" )
			.attr( "role", "tablist" )

			// Prevent users from focusing disabled tabs via click
			.delegate( "> li", "mousedown" + this.eventNamespace, function( event ) {
				if ( $( this ).is( ".ui-state-disabled" ) ) {
					event.preventDefault();
				}
			})

			// support: IE <9
			// Preventing the default action in mousedown doesn't prevent IE
			// from focusing the element, so if the anchor gets focused, blur.
			// We don't have to worry about focusing the previously focused
			// element since clicking on a non-focusable element should focus
			// the body anyway.
			.delegate( ".ui-tabs-anchor", "focus" + this.eventNamespace, function() {
				if ( $( this ).closest( "li" ).is( ".ui-state-disabled" ) ) {
					this.blur();
				}
			});

		this.tabs = this.tablist.find( "> li:has(a[href])" )
			.addClass( "ui-state-default ui-corner-top" )
			.attr({
				role: "tab",
				tabIndex: -1
			});

		this.anchors = this.tabs.map(function() {
				return $( "a", this )[ 0 ];
			})
			.addClass( "ui-tabs-anchor" )
			.attr({
				role: "presentation",
				tabIndex: -1
			});

		this.panels = $();

		this.anchors.each(function( i, anchor ) {
			var selector, panel, panelId,
				anchorId = $( anchor ).uniqueId().attr( "id" ),
				tab = $( anchor ).closest( "li" ),
				originalAriaControls = tab.attr( "aria-controls" );

			// inline tab
			if ( that._isLocal( anchor ) ) {
				selector = anchor.hash;
				panelId = selector.substring( 1 );
				panel = that.element.find( that._sanitizeSelector( selector ) );
			// remote tab
			} else {
				// If the tab doesn't already have aria-controls,
				// generate an id by using a throw-away element
				panelId = tab.attr( "aria-controls" ) || $( {} ).uniqueId()[ 0 ].id;
				selector = "#" + panelId;
				panel = that.element.find( selector );
				if ( !panel.length ) {
					panel = that._createPanel( panelId );
					panel.insertAfter( that.panels[ i - 1 ] || that.tablist );
				}
				panel.attr( "aria-live", "polite" );
			}

			if ( panel.length) {
				that.panels = that.panels.add( panel );
			}
			if ( originalAriaControls ) {
				tab.data( "ui-tabs-aria-controls", originalAriaControls );
			}
			tab.attr({
				"aria-controls": panelId,
				"aria-labelledby": anchorId
			});
			panel.attr( "aria-labelledby", anchorId );
		});

		this.panels
			.addClass( "ui-tabs-panel ui-widget-content ui-corner-bottom" )
			.attr( "role", "tabpanel" );

		// Avoid memory leaks (#10056)
		if ( prevTabs ) {
			this._off( prevTabs.not( this.tabs ) );
			this._off( prevAnchors.not( this.anchors ) );
			this._off( prevPanels.not( this.panels ) );
		}
	},

	// allow overriding how to find the list for rare usage scenarios (#7715)
	_getList: function() {
		return this.tablist || this.element.find( "ol,ul" ).eq( 0 );
	},

	_createPanel: function( id ) {
		return $( "<div>" )
			.attr( "id", id )
			.addClass( "ui-tabs-panel ui-widget-content ui-corner-bottom" )
			.data( "ui-tabs-destroy", true );
	},

	_setupDisabled: function( disabled ) {
		if ( $.isArray( disabled ) ) {
			if ( !disabled.length ) {
				disabled = false;
			} else if ( disabled.length === this.anchors.length ) {
				disabled = true;
			}
		}

		// disable tabs
		for ( var i = 0, li; ( li = this.tabs[ i ] ); i++ ) {
			if ( disabled === true || $.inArray( i, disabled ) !== -1 ) {
				$( li )
					.addClass( "ui-state-disabled" )
					.attr( "aria-disabled", "true" );
			} else {
				$( li )
					.removeClass( "ui-state-disabled" )
					.removeAttr( "aria-disabled" );
			}
		}

		this.options.disabled = disabled;
	},

	_setupEvents: function( event ) {
		var events = {};
		if ( event ) {
			$.each( event.split(" "), function( index, eventName ) {
				events[ eventName ] = "_eventHandler";
			});
		}

		this._off( this.anchors.add( this.tabs ).add( this.panels ) );
		// Always prevent the default action, even when disabled
		this._on( true, this.anchors, {
			click: function( event ) {
				event.preventDefault();
			}
		});
		this._on( this.anchors, events );
		this._on( this.tabs, { keydown: "_tabKeydown" } );
		this._on( this.panels, { keydown: "_panelKeydown" } );

		this._focusable( this.tabs );
		this._hoverable( this.tabs );
	},

	_setupHeightStyle: function( heightStyle ) {
		var maxHeight,
			parent = this.element.parent();

		if ( heightStyle === "fill" ) {
			maxHeight = parent.height();
			maxHeight -= this.element.outerHeight() - this.element.height();

			this.element.siblings( ":visible" ).each(function() {
				var elem = $( this ),
					position = elem.css( "position" );

				if ( position === "absolute" || position === "fixed" ) {
					return;
				}
				maxHeight -= elem.outerHeight( true );
			});

			this.element.children().not( this.panels ).each(function() {
				maxHeight -= $( this ).outerHeight( true );
			});

			this.panels.each(function() {
				$( this ).height( Math.max( 0, maxHeight -
					$( this ).innerHeight() + $( this ).height() ) );
			})
			.css( "overflow", "auto" );
		} else if ( heightStyle === "auto" ) {
			maxHeight = 0;
			this.panels.each(function() {
				maxHeight = Math.max( maxHeight, $( this ).height( "" ).height() );
			}).height( maxHeight );
		}
	},

	_eventHandler: function( event ) {
		var options = this.options,
			active = this.active,
			anchor = $( event.currentTarget ),
			tab = anchor.closest( "li" ),
			clickedIsActive = tab[ 0 ] === active[ 0 ],
			collapsing = clickedIsActive && options.collapsible,
			toShow = collapsing ? $() : this._getPanelForTab( tab ),
			toHide = !active.length ? $() : this._getPanelForTab( active ),
			eventData = {
				oldTab: active,
				oldPanel: toHide,
				newTab: collapsing ? $() : tab,
				newPanel: toShow
			};

		event.preventDefault();

		if ( tab.hasClass( "ui-state-disabled" ) ||
				// tab is already loading
				tab.hasClass( "ui-tabs-loading" ) ||
				// can't switch durning an animation
				this.running ||
				// click on active header, but not collapsible
				( clickedIsActive && !options.collapsible ) ||
				// allow canceling activation
				( this._trigger( "beforeActivate", event, eventData ) === false ) ) {
			return;
		}

		options.active = collapsing ? false : this.tabs.index( tab );

		this.active = clickedIsActive ? $() : tab;
		if ( this.xhr ) {
			this.xhr.abort();
		}

		if ( !toHide.length && !toShow.length ) {
			$.error( "jQuery UI Tabs: Mismatching fragment identifier." );
		}

		if ( toShow.length ) {
			this.load( this.tabs.index( tab ), event );
		}
		this._toggle( event, eventData );
	},

	// handles show/hide for selecting tabs
	_toggle: function( event, eventData ) {
		var that = this,
			toShow = eventData.newPanel,
			toHide = eventData.oldPanel;

		this.running = true;

		function complete() {
			that.running = false;
			that._trigger( "activate", event, eventData );
		}

		function show() {
			eventData.newTab.closest( "li" ).addClass( "ui-tabs-active ui-state-active" );

			if ( toShow.length && that.options.show ) {
				that._show( toShow, that.options.show, complete );
			} else {
				toShow.show();
				complete();
			}
		}

		// start out by hiding, then showing, then completing
		if ( toHide.length && this.options.hide ) {
			this._hide( toHide, this.options.hide, function() {
				eventData.oldTab.closest( "li" ).removeClass( "ui-tabs-active ui-state-active" );
				show();
			});
		} else {
			eventData.oldTab.closest( "li" ).removeClass( "ui-tabs-active ui-state-active" );
			toHide.hide();
			show();
		}

		toHide.attr( "aria-hidden", "true" );
		eventData.oldTab.attr({
			"aria-selected": "false",
			"aria-expanded": "false"
		});
		// If we're switching tabs, remove the old tab from the tab order.
		// If we're opening from collapsed state, remove the previous tab from the tab order.
		// If we're collapsing, then keep the collapsing tab in the tab order.
		if ( toShow.length && toHide.length ) {
			eventData.oldTab.attr( "tabIndex", -1 );
		} else if ( toShow.length ) {
			this.tabs.filter(function() {
				return $( this ).attr( "tabIndex" ) === 0;
			})
			.attr( "tabIndex", -1 );
		}

		toShow.attr( "aria-hidden", "false" );
		eventData.newTab.attr({
			"aria-selected": "true",
			"aria-expanded": "true",
			tabIndex: 0
		});
	},

	_activate: function( index ) {
		var anchor,
			active = this._findActive( index );

		// trying to activate the already active panel
		if ( active[ 0 ] === this.active[ 0 ] ) {
			return;
		}

		// trying to collapse, simulate a click on the current active header
		if ( !active.length ) {
			active = this.active;
		}

		anchor = active.find( ".ui-tabs-anchor" )[ 0 ];
		this._eventHandler({
			target: anchor,
			currentTarget: anchor,
			preventDefault: $.noop
		});
	},

	_findActive: function( index ) {
		return index === false ? $() : this.tabs.eq( index );
	},

	_getIndex: function( index ) {
		// meta-function to give users option to provide a href string instead of a numerical index.
		if ( typeof index === "string" ) {
			index = this.anchors.index( this.anchors.filter( "[href$='" + index + "']" ) );
		}

		return index;
	},

	_destroy: function() {
		if ( this.xhr ) {
			this.xhr.abort();
		}

		this.element.removeClass( "ui-tabs ui-widget ui-widget-content ui-corner-all ui-tabs-collapsible" );

		this.tablist
			.removeClass( "ui-tabs-nav ui-helper-reset ui-helper-clearfix ui-widget-header ui-corner-all" )
			.removeAttr( "role" );

		this.anchors
			.removeClass( "ui-tabs-anchor" )
			.removeAttr( "role" )
			.removeAttr( "tabIndex" )
			.removeUniqueId();

		this.tablist.unbind( this.eventNamespace );

		this.tabs.add( this.panels ).each(function() {
			if ( $.data( this, "ui-tabs-destroy" ) ) {
				$( this ).remove();
			} else {
				$( this )
					.removeClass( "ui-state-default ui-state-active ui-state-disabled " +
						"ui-corner-top ui-corner-bottom ui-widget-content ui-tabs-active ui-tabs-panel" )
					.removeAttr( "tabIndex" )
					.removeAttr( "aria-live" )
					.removeAttr( "aria-busy" )
					.removeAttr( "aria-selected" )
					.removeAttr( "aria-labelledby" )
					.removeAttr( "aria-hidden" )
					.removeAttr( "aria-expanded" )
					.removeAttr( "role" );
			}
		});

		this.tabs.each(function() {
			var li = $( this ),
				prev = li.data( "ui-tabs-aria-controls" );
			if ( prev ) {
				li
					.attr( "aria-controls", prev )
					.removeData( "ui-tabs-aria-controls" );
			} else {
				li.removeAttr( "aria-controls" );
			}
		});

		this.panels.show();

		if ( this.options.heightStyle !== "content" ) {
			this.panels.css( "height", "" );
		}
	},

	enable: function( index ) {
		var disabled = this.options.disabled;
		if ( disabled === false ) {
			return;
		}

		if ( index === undefined ) {
			disabled = false;
		} else {
			index = this._getIndex( index );
			if ( $.isArray( disabled ) ) {
				disabled = $.map( disabled, function( num ) {
					return num !== index ? num : null;
				});
			} else {
				disabled = $.map( this.tabs, function( li, num ) {
					return num !== index ? num : null;
				});
			}
		}
		this._setupDisabled( disabled );
	},

	disable: function( index ) {
		var disabled = this.options.disabled;
		if ( disabled === true ) {
			return;
		}

		if ( index === undefined ) {
			disabled = true;
		} else {
			index = this._getIndex( index );
			if ( $.inArray( index, disabled ) !== -1 ) {
				return;
			}
			if ( $.isArray( disabled ) ) {
				disabled = $.merge( [ index ], disabled ).sort();
			} else {
				disabled = [ index ];
			}
		}
		this._setupDisabled( disabled );
	},

	load: function( index, event ) {
		index = this._getIndex( index );
		var that = this,
			tab = this.tabs.eq( index ),
			anchor = tab.find( ".ui-tabs-anchor" ),
			panel = this._getPanelForTab( tab ),
			eventData = {
				tab: tab,
				panel: panel
			},
			complete = function( jqXHR, status ) {
				if ( status === "abort" ) {
					that.panels.stop( false, true );
				}

				tab.removeClass( "ui-tabs-loading" );
				panel.removeAttr( "aria-busy" );

				if ( jqXHR === that.xhr ) {
					delete that.xhr;
				}
			};

		// not remote
		if ( this._isLocal( anchor[ 0 ] ) ) {
			return;
		}

		this.xhr = $.ajax( this._ajaxSettings( anchor, event, eventData ) );

		// support: jQuery <1.8
		// jQuery <1.8 returns false if the request is canceled in beforeSend,
		// but as of 1.8, $.ajax() always returns a jqXHR object.
		if ( this.xhr && this.xhr.statusText !== "canceled" ) {
			tab.addClass( "ui-tabs-loading" );
			panel.attr( "aria-busy", "true" );

			this.xhr
				.done(function( response, status, jqXHR ) {
					// support: jQuery <1.8
					// http://bugs.jquery.com/ticket/11778
					setTimeout(function() {
						panel.html( response );
						that._trigger( "load", event, eventData );

						complete( jqXHR, status );
					}, 1 );
				})
				.fail(function( jqXHR, status ) {
					// support: jQuery <1.8
					// http://bugs.jquery.com/ticket/11778
					setTimeout(function() {
						complete( jqXHR, status );
					}, 1 );
				});
		}
	},

	_ajaxSettings: function( anchor, event, eventData ) {
		var that = this;
		return {
			url: anchor.attr( "href" ),
			beforeSend: function( jqXHR, settings ) {
				return that._trigger( "beforeLoad", event,
					$.extend( { jqXHR: jqXHR, ajaxSettings: settings }, eventData ) );
			}
		};
	},

	_getPanelForTab: function( tab ) {
		var id = $( tab ).attr( "aria-controls" );
		return this.element.find( this._sanitizeSelector( "#" + id ) );
	}
});


/*!
 * jQuery UI Tooltip 1.11.4
 * http://jqueryui.com
 *
 * Copyright jQuery Foundation and other contributors
 * Released under the MIT license.
 * http://jquery.org/license
 *
 * http://api.jqueryui.com/tooltip/
 */


var tooltip = $.widget( "ui.tooltip", {
	version: "1.11.4",
	options: {
		content: function() {
			// support: IE<9, Opera in jQuery <1.7
			// .text() can't accept undefined, so coerce to a string
			var title = $( this ).attr( "title" ) || "";
			// Escape title, since we're going from an attribute to raw HTML
			return $( "<a>" ).text( title ).html();
		},
		hide: true,
		// Disabled elements have inconsistent behavior across browsers (#8661)
		items: "[title]:not([disabled])",
		position: {
			my: "left top+15",
			at: "left bottom",
			collision: "flipfit flip"
		},
		show: true,
		tooltipClass: null,
		track: false,

		// callbacks
		close: null,
		open: null
	},

	_addDescribedBy: function( elem, id ) {
		var describedby = (elem.attr( "aria-describedby" ) || "").split( /\s+/ );
		describedby.push( id );
		elem
			.data( "ui-tooltip-id", id )
			.attr( "aria-describedby", $.trim( describedby.join( " " ) ) );
	},

	_removeDescribedBy: function( elem ) {
		var id = elem.data( "ui-tooltip-id" ),
			describedby = (elem.attr( "aria-describedby" ) || "").split( /\s+/ ),
			index = $.inArray( id, describedby );

		if ( index !== -1 ) {
			describedby.splice( index, 1 );
		}

		elem.removeData( "ui-tooltip-id" );
		describedby = $.trim( describedby.join( " " ) );
		if ( describedby ) {
			elem.attr( "aria-describedby", describedby );
		} else {
			elem.removeAttr( "aria-describedby" );
		}
	},

	_create: function() {
		this._on({
			mouseover: "open",
			focusin: "open"
		});

		// IDs of generated tooltips, needed for destroy
		this.tooltips = {};

		// IDs of parent tooltips where we removed the title attribute
		this.parents = {};

		if ( this.options.disabled ) {
			this._disable();
		}

		// Append the aria-live region so tooltips announce correctly
		this.liveRegion = $( "<div>" )
			.attr({
				role: "log",
				"aria-live": "assertive",
				"aria-relevant": "additions"
			})
			.addClass( "ui-helper-hidden-accessible" )
			.appendTo( this.document[ 0 ].body );
	},

	_setOption: function( key, value ) {
		var that = this;

		if ( key === "disabled" ) {
			this[ value ? "_disable" : "_enable" ]();
			this.options[ key ] = value;
			// disable element style changes
			return;
		}

		this._super( key, value );

		if ( key === "content" ) {
			$.each( this.tooltips, function( id, tooltipData ) {
				that._updateContent( tooltipData.element );
			});
		}
	},

	_disable: function() {
		var that = this;

		// close open tooltips
		$.each( this.tooltips, function( id, tooltipData ) {
			var event = $.Event( "blur" );
			event.target = event.currentTarget = tooltipData.element[ 0 ];
			that.close( event, true );
		});

		// remove title attributes to prevent native tooltips
		this.element.find( this.options.items ).addBack().each(function() {
			var element = $( this );
			if ( element.is( "[title]" ) ) {
				element
					.data( "ui-tooltip-title", element.attr( "title" ) )
					.removeAttr( "title" );
			}
		});
	},

	_enable: function() {
		// restore title attributes
		this.element.find( this.options.items ).addBack().each(function() {
			var element = $( this );
			if ( element.data( "ui-tooltip-title" ) ) {
				element.attr( "title", element.data( "ui-tooltip-title" ) );
			}
		});
	},

	open: function( event ) {
		var that = this,
			target = $( event ? event.target : this.element )
				// we need closest here due to mouseover bubbling,
				// but always pointing at the same event target
				.closest( this.options.items );

		// No element to show a tooltip for or the tooltip is already open
		if ( !target.length || target.data( "ui-tooltip-id" ) ) {
			return;
		}

		if ( target.attr( "title" ) ) {
			target.data( "ui-tooltip-title", target.attr( "title" ) );
		}

		target.data( "ui-tooltip-open", true );

		// kill parent tooltips, custom or native, for hover
		if ( event && event.type === "mouseover" ) {
			target.parents().each(function() {
				var parent = $( this ),
					blurEvent;
				if ( parent.data( "ui-tooltip-open" ) ) {
					blurEvent = $.Event( "blur" );
					blurEvent.target = blurEvent.currentTarget = this;
					that.close( blurEvent, true );
				}
				if ( parent.attr( "title" ) ) {
					parent.uniqueId();
					that.parents[ this.id ] = {
						element: this,
						title: parent.attr( "title" )
					};
					parent.attr( "title", "" );
				}
			});
		}

		this._registerCloseHandlers( event, target );
		this._updateContent( target, event );
	},

	_updateContent: function( target, event ) {
		var content,
			contentOption = this.options.content,
			that = this,
			eventType = event ? event.type : null;

		if ( typeof contentOption === "string" ) {
			return this._open( event, target, contentOption );
		}

		content = contentOption.call( target[0], function( response ) {

			// IE may instantly serve a cached response for ajax requests
			// delay this call to _open so the other call to _open runs first
			that._delay(function() {

				// Ignore async response if tooltip was closed already
				if ( !target.data( "ui-tooltip-open" ) ) {
					return;
				}

				// jQuery creates a special event for focusin when it doesn't
				// exist natively. To improve performance, the native event
				// object is reused and the type is changed. Therefore, we can't
				// rely on the type being correct after the event finished
				// bubbling, so we set it back to the previous value. (#8740)
				if ( event ) {
					event.type = eventType;
				}
				this._open( event, target, response );
			});
		});
		if ( content ) {
			this._open( event, target, content );
		}
	},

	_open: function( event, target, content ) {
		var tooltipData, tooltip, delayedShow, a11yContent,
			positionOption = $.extend( {}, this.options.position );

		if ( !content ) {
			return;
		}

		// Content can be updated multiple times. If the tooltip already
		// exists, then just update the content and bail.
		tooltipData = this._find( target );
		if ( tooltipData ) {
			tooltipData.tooltip.find( ".ui-tooltip-content" ).html( content );
			return;
		}

		// if we have a title, clear it to prevent the native tooltip
		// we have to check first to avoid defining a title if none exists
		// (we don't want to cause an element to start matching [title])
		//
		// We use removeAttr only for key events, to allow IE to export the correct
		// accessible attributes. For mouse events, set to empty string to avoid
		// native tooltip showing up (happens only when removing inside mouseover).
		if ( target.is( "[title]" ) ) {
			if ( event && event.type === "mouseover" ) {
				target.attr( "title", "" );
			} else {
				target.removeAttr( "title" );
			}
		}

		tooltipData = this._tooltip( target );
		tooltip = tooltipData.tooltip;
		this._addDescribedBy( target, tooltip.attr( "id" ) );
		tooltip.find( ".ui-tooltip-content" ).html( content );

		// Support: Voiceover on OS X, JAWS on IE <= 9
		// JAWS announces deletions even when aria-relevant="additions"
		// Voiceover will sometimes re-read the entire log region's contents from the beginning
		this.liveRegion.children().hide();
		if ( content.clone ) {
			a11yContent = content.clone();
			a11yContent.removeAttr( "id" ).find( "[id]" ).removeAttr( "id" );
		} else {
			a11yContent = content;
		}
		$( "<div>" ).html( a11yContent ).appendTo( this.liveRegion );

		function position( event ) {
			positionOption.of = event;
			if ( tooltip.is( ":hidden" ) ) {
				return;
			}
			tooltip.position( positionOption );
		}
		if ( this.options.track && event && /^mouse/.test( event.type ) ) {
			this._on( this.document, {
				mousemove: position
			});
			// trigger once to override element-relative positioning
			position( event );
		} else {
			tooltip.position( $.extend({
				of: target
			}, this.options.position ) );
		}

		tooltip.hide();

		this._show( tooltip, this.options.show );
		// Handle tracking tooltips that are shown with a delay (#8644). As soon
		// as the tooltip is visible, position the tooltip using the most recent
		// event.
		if ( this.options.show && this.options.show.delay ) {
			delayedShow = this.delayedShow = setInterval(function() {
				if ( tooltip.is( ":visible" ) ) {
					position( positionOption.of );
					clearInterval( delayedShow );
				}
			}, $.fx.interval );
		}

		this._trigger( "open", event, { tooltip: tooltip } );
	},

	_registerCloseHandlers: function( event, target ) {
		var events = {
			keyup: function( event ) {
				if ( event.keyCode === $.ui.keyCode.ESCAPE ) {
					var fakeEvent = $.Event(event);
					fakeEvent.currentTarget = target[0];
					this.close( fakeEvent, true );
				}
			}
		};

		// Only bind remove handler for delegated targets. Non-delegated
		// tooltips will handle this in destroy.
		if ( target[ 0 ] !== this.element[ 0 ] ) {
			events.remove = function() {
				this._removeTooltip( this._find( target ).tooltip );
			};
		}

		if ( !event || event.type === "mouseover" ) {
			events.mouseleave = "close";
		}
		if ( !event || event.type === "focusin" ) {
			events.focusout = "close";
		}
		this._on( true, target, events );
	},

	close: function( event ) {
		var tooltip,
			that = this,
			target = $( event ? event.currentTarget : this.element ),
			tooltipData = this._find( target );

		// The tooltip may already be closed
		if ( !tooltipData ) {

			// We set ui-tooltip-open immediately upon open (in open()), but only set the
			// additional data once there's actually content to show (in _open()). So even if the
			// tooltip doesn't have full data, we always remove ui-tooltip-open in case we're in
			// the period between open() and _open().
			target.removeData( "ui-tooltip-open" );
			return;
		}

		tooltip = tooltipData.tooltip;

		// disabling closes the tooltip, so we need to track when we're closing
		// to avoid an infinite loop in case the tooltip becomes disabled on close
		if ( tooltipData.closing ) {
			return;
		}

		// Clear the interval for delayed tracking tooltips
		clearInterval( this.delayedShow );

		// only set title if we had one before (see comment in _open())
		// If the title attribute has changed since open(), don't restore
		if ( target.data( "ui-tooltip-title" ) && !target.attr( "title" ) ) {
			target.attr( "title", target.data( "ui-tooltip-title" ) );
		}

		this._removeDescribedBy( target );

		tooltipData.hiding = true;
		tooltip.stop( true );
		this._hide( tooltip, this.options.hide, function() {
			that._removeTooltip( $( this ) );
		});

		target.removeData( "ui-tooltip-open" );
		this._off( target, "mouseleave focusout keyup" );

		// Remove 'remove' binding only on delegated targets
		if ( target[ 0 ] !== this.element[ 0 ] ) {
			this._off( target, "remove" );
		}
		this._off( this.document, "mousemove" );

		if ( event && event.type === "mouseleave" ) {
			$.each( this.parents, function( id, parent ) {
				$( parent.element ).attr( "title", parent.title );
				delete that.parents[ id ];
			});
		}

		tooltipData.closing = true;
		this._trigger( "close", event, { tooltip: tooltip } );
		if ( !tooltipData.hiding ) {
			tooltipData.closing = false;
		}
	},

	_tooltip: function( element ) {
		var tooltip = $( "<div>" )
				.attr( "role", "tooltip" )
				.addClass( "ui-tooltip ui-widget ui-corner-all ui-widget-content " +
					( this.options.tooltipClass || "" ) ),
			id = tooltip.uniqueId().attr( "id" );

		$( "<div>" )
			.addClass( "ui-tooltip-content" )
			.appendTo( tooltip );

		tooltip.appendTo( this.document[0].body );

		return this.tooltips[ id ] = {
			element: element,
			tooltip: tooltip
		};
	},

	_find: function( target ) {
		var id = target.data( "ui-tooltip-id" );
		return id ? this.tooltips[ id ] : null;
	},

	_removeTooltip: function( tooltip ) {
		tooltip.remove();
		delete this.tooltips[ tooltip.attr( "id" ) ];
	},

	_destroy: function() {
		var that = this;

		// close open tooltips
		$.each( this.tooltips, function( id, tooltipData ) {
			// Delegate to close method to handle common cleanup
			var event = $.Event( "blur" ),
				element = tooltipData.element;
			event.target = event.currentTarget = element[ 0 ];
			that.close( event, true );

			// Remove immediately; destroying an open tooltip doesn't use the
			// hide animation
			$( "#" + id ).remove();

			// Restore the title
			if ( element.data( "ui-tooltip-title" ) ) {
				// If the title attribute has changed since open(), don't restore
				if ( !element.attr( "title" ) ) {
					element.attr( "title", element.data( "ui-tooltip-title" ) );
				}
				element.removeData( "ui-tooltip-title" );
			}
		});
		this.liveRegion.remove();
	}
});


/*!
 * jQuery UI Effects 1.11.4
 * http://jqueryui.com
 *
 * Copyright jQuery Foundation and other contributors
 * Released under the MIT license.
 * http://jquery.org/license
 *
 * http://api.jqueryui.com/category/effects-core/
 */


var dataSpace = "ui-effects-",

	// Create a local jQuery because jQuery Color relies on it and the
	// global may not exist with AMD and a custom build (#10199)
	jQuery = $;

$.effects = {
	effect: {}
};

/*!
 * jQuery Color Animations v2.1.2
 * https://github.com/jquery/jquery-color
 *
 * Copyright 2014 jQuery Foundation and other contributors
 * Released under the MIT license.
 * http://jquery.org/license
 *
 * Date: Wed Jan 16 08:47:09 2013 -0600
 */
(function( jQuery, undefined ) {

	var stepHooks = "backgroundColor borderBottomColor borderLeftColor borderRightColor borderTopColor color columnRuleColor outlineColor textDecorationColor textEmphasisColor",

	// plusequals test for += 100 -= 100
	rplusequals = /^([\-+])=\s*(\d+\.?\d*)/,
	// a set of RE's that can match strings and generate color tuples.
	stringParsers = [ {
			re: /rgba?\(\s*(\d{1,3})\s*,\s*(\d{1,3})\s*,\s*(\d{1,3})\s*(?:,\s*(\d?(?:\.\d+)?)\s*)?\)/,
			parse: function( execResult ) {
				return [
					execResult[ 1 ],
					execResult[ 2 ],
					execResult[ 3 ],
					execResult[ 4 ]
				];
			}
		}, {
			re: /rgba?\(\s*(\d+(?:\.\d+)?)\%\s*,\s*(\d+(?:\.\d+)?)\%\s*,\s*(\d+(?:\.\d+)?)\%\s*(?:,\s*(\d?(?:\.\d+)?)\s*)?\)/,
			parse: function( execResult ) {
				return [
					execResult[ 1 ] * 2.55,
					execResult[ 2 ] * 2.55,
					execResult[ 3 ] * 2.55,
					execResult[ 4 ]
				];
			}
		}, {
			// this regex ignores A-F because it's compared against an already lowercased string
			re: /#([a-f0-9]{2})([a-f0-9]{2})([a-f0-9]{2})/,
			parse: function( execResult ) {
				return [
					parseInt( execResult[ 1 ], 16 ),
					parseInt( execResult[ 2 ], 16 ),
					parseInt( execResult[ 3 ], 16 )
				];
			}
		}, {
			// this regex ignores A-F because it's compared against an already lowercased string
			re: /#([a-f0-9])([a-f0-9])([a-f0-9])/,
			parse: function( execResult ) {
				return [
					parseInt( execResult[ 1 ] + execResult[ 1 ], 16 ),
					parseInt( execResult[ 2 ] + execResult[ 2 ], 16 ),
					parseInt( execResult[ 3 ] + execResult[ 3 ], 16 )
				];
			}
		}, {
			re: /hsla?\(\s*(\d+(?:\.\d+)?)\s*,\s*(\d+(?:\.\d+)?)\%\s*,\s*(\d+(?:\.\d+)?)\%\s*(?:,\s*(\d?(?:\.\d+)?)\s*)?\)/,
			space: "hsla",
			parse: function( execResult ) {
				return [
					execResult[ 1 ],
					execResult[ 2 ] / 100,
					execResult[ 3 ] / 100,
					execResult[ 4 ]
				];
			}
		} ],

	// jQuery.Color( )
	color = jQuery.Color = function( color, green, blue, alpha ) {
		return new jQuery.Color.fn.parse( color, green, blue, alpha );
	},
	spaces = {
		rgba: {
			props: {
				red: {
					idx: 0,
					type: "byte"
				},
				green: {
					idx: 1,
					type: "byte"
				},
				blue: {
					idx: 2,
					type: "byte"
				}
			}
		},

		hsla: {
			props: {
				hue: {
					idx: 0,
					type: "degrees"
				},
				saturation: {
					idx: 1,
					type: "percent"
				},
				lightness: {
					idx: 2,
					type: "percent"
				}
			}
		}
	},
	propTypes = {
		"byte": {
			floor: true,
			max: 255
		},
		"percent": {
			max: 1
		},
		"degrees": {
			mod: 360,
			floor: true
		}
	},
	support = color.support = {},

	// element for support tests
	supportElem = jQuery( "<p>" )[ 0 ],

	// colors = jQuery.Color.names
	colors,

	// local aliases of functions called often
	each = jQuery.each;

// determine rgba support immediately
supportElem.style.cssText = "background-color:rgba(1,1,1,.5)";
support.rgba = supportElem.style.backgroundColor.indexOf( "rgba" ) > -1;

// define cache name and alpha properties
// for rgba and hsla spaces
each( spaces, function( spaceName, space ) {
	space.cache = "_" + spaceName;
	space.props.alpha = {
		idx: 3,
		type: "percent",
		def: 1
	};
});

function clamp( value, prop, allowEmpty ) {
	var type = propTypes[ prop.type ] || {};

	if ( value == null ) {
		return (allowEmpty || !prop.def) ? null : prop.def;
	}

	// ~~ is an short way of doing floor for positive numbers
	value = type.floor ? ~~value : parseFloat( value );

	// IE will pass in empty strings as value for alpha,
	// which will hit this case
	if ( isNaN( value ) ) {
		return prop.def;
	}

	if ( type.mod ) {
		// we add mod before modding to make sure that negatives values
		// get converted properly: -10 -> 350
		return (value + type.mod) % type.mod;
	}

	// for now all property types without mod have min and max
	return 0 > value ? 0 : type.max < value ? type.max : value;
}

function stringParse( string ) {
	var inst = color(),
		rgba = inst._rgba = [];

	string = string.toLowerCase();

	each( stringParsers, function( i, parser ) {
		var parsed,
			match = parser.re.exec( string ),
			values = match && parser.parse( match ),
			spaceName = parser.space || "rgba";

		if ( values ) {
			parsed = inst[ spaceName ]( values );

			// if this was an rgba parse the assignment might happen twice
			// oh well....
			inst[ spaces[ spaceName ].cache ] = parsed[ spaces[ spaceName ].cache ];
			rgba = inst._rgba = parsed._rgba;

			// exit each( stringParsers ) here because we matched
			return false;
		}
	});

	// Found a stringParser that handled it
	if ( rgba.length ) {

		// if this came from a parsed string, force "transparent" when alpha is 0
		// chrome, (and maybe others) return "transparent" as rgba(0,0,0,0)
		if ( rgba.join() === "0,0,0,0" ) {
			jQuery.extend( rgba, colors.transparent );
		}
		return inst;
	}

	// named colors
	return colors[ string ];
}

color.fn = jQuery.extend( color.prototype, {
	parse: function( red, green, blue, alpha ) {
		if ( red === undefined ) {
			this._rgba = [ null, null, null, null ];
			return this;
		}
		if ( red.jquery || red.nodeType ) {
			red = jQuery( red ).css( green );
			green = undefined;
		}

		var inst = this,
			type = jQuery.type( red ),
			rgba = this._rgba = [];

		// more than 1 argument specified - assume ( red, green, blue, alpha )
		if ( green !== undefined ) {
			red = [ red, green, blue, alpha ];
			type = "array";
		}

		if ( type === "string" ) {
			return this.parse( stringParse( red ) || colors._default );
		}

		if ( type === "array" ) {
			each( spaces.rgba.props, function( key, prop ) {
				rgba[ prop.idx ] = clamp( red[ prop.idx ], prop );
			});
			return this;
		}

		if ( type === "object" ) {
			if ( red instanceof color ) {
				each( spaces, function( spaceName, space ) {
					if ( red[ space.cache ] ) {
						inst[ space.cache ] = red[ space.cache ].slice();
					}
				});
			} else {
				each( spaces, function( spaceName, space ) {
					var cache = space.cache;
					each( space.props, function( key, prop ) {

						// if the cache doesn't exist, and we know how to convert
						if ( !inst[ cache ] && space.to ) {

							// if the value was null, we don't need to copy it
							// if the key was alpha, we don't need to copy it either
							if ( key === "alpha" || red[ key ] == null ) {
								return;
							}
							inst[ cache ] = space.to( inst._rgba );
						}

						// this is the only case where we allow nulls for ALL properties.
						// call clamp with alwaysAllowEmpty
						inst[ cache ][ prop.idx ] = clamp( red[ key ], prop, true );
					});

					// everything defined but alpha?
					if ( inst[ cache ] && jQuery.inArray( null, inst[ cache ].slice( 0, 3 ) ) < 0 ) {
						// use the default of 1
						inst[ cache ][ 3 ] = 1;
						if ( space.from ) {
							inst._rgba = space.from( inst[ cache ] );
						}
					}
				});
			}
			return this;
		}
	},
	is: function( compare ) {
		var is = color( compare ),
			same = true,
			inst = this;

		each( spaces, function( _, space ) {
			var localCache,
				isCache = is[ space.cache ];
			if (isCache) {
				localCache = inst[ space.cache ] || space.to && space.to( inst._rgba ) || [];
				each( space.props, function( _, prop ) {
					if ( isCache[ prop.idx ] != null ) {
						same = ( isCache[ prop.idx ] === localCache[ prop.idx ] );
						return same;
					}
				});
			}
			return same;
		});
		return same;
	},
	_space: function() {
		var used = [],
			inst = this;
		each( spaces, function( spaceName, space ) {
			if ( inst[ space.cache ] ) {
				used.push( spaceName );
			}
		});
		return used.pop();
	},
	transition: function( other, distance ) {
		var end = color( other ),
			spaceName = end._space(),
			space = spaces[ spaceName ],
			startColor = this.alpha() === 0 ? color( "transparent" ) : this,
			start = startColor[ space.cache ] || space.to( startColor._rgba ),
			result = start.slice();

		end = end[ space.cache ];
		each( space.props, function( key, prop ) {
			var index = prop.idx,
				startValue = start[ index ],
				endValue = end[ index ],
				type = propTypes[ prop.type ] || {};

			// if null, don't override start value
			if ( endValue === null ) {
				return;
			}
			// if null - use end
			if ( startValue === null ) {
				result[ index ] = endValue;
			} else {
				if ( type.mod ) {
					if ( endValue - startValue > type.mod / 2 ) {
						startValue += type.mod;
					} else if ( startValue - endValue > type.mod / 2 ) {
						startValue -= type.mod;
					}
				}
				result[ index ] = clamp( ( endValue - startValue ) * distance + startValue, prop );
			}
		});
		return this[ spaceName ]( result );
	},
	blend: function( opaque ) {
		// if we are already opaque - return ourself
		if ( this._rgba[ 3 ] === 1 ) {
			return this;
		}

		var rgb = this._rgba.slice(),
			a = rgb.pop(),
			blend = color( opaque )._rgba;

		return color( jQuery.map( rgb, function( v, i ) {
			return ( 1 - a ) * blend[ i ] + a * v;
		}));
	},
	toRgbaString: function() {
		var prefix = "rgba(",
			rgba = jQuery.map( this._rgba, function( v, i ) {
				return v == null ? ( i > 2 ? 1 : 0 ) : v;
			});

		if ( rgba[ 3 ] === 1 ) {
			rgba.pop();
			prefix = "rgb(";
		}

		return prefix + rgba.join() + ")";
	},
	toHslaString: function() {
		var prefix = "hsla(",
			hsla = jQuery.map( this.hsla(), function( v, i ) {
				if ( v == null ) {
					v = i > 2 ? 1 : 0;
				}

				// catch 1 and 2
				if ( i && i < 3 ) {
					v = Math.round( v * 100 ) + "%";
				}
				return v;
			});

		if ( hsla[ 3 ] === 1 ) {
			hsla.pop();
			prefix = "hsl(";
		}
		return prefix + hsla.join() + ")";
	},
	toHexString: function( includeAlpha ) {
		var rgba = this._rgba.slice(),
			alpha = rgba.pop();

		if ( includeAlpha ) {
			rgba.push( ~~( alpha * 255 ) );
		}

		return "#" + jQuery.map( rgba, function( v ) {

			// default to 0 when nulls exist
			v = ( v || 0 ).toString( 16 );
			return v.length === 1 ? "0" + v : v;
		}).join("");
	},
	toString: function() {
		return this._rgba[ 3 ] === 0 ? "transparent" : this.toRgbaString();
	}
});
color.fn.parse.prototype = color.fn;

// hsla conversions adapted from:
// https://code.google.com/p/maashaack/source/browse/packages/graphics/trunk/src/graphics/colors/HUE2RGB.as?r=5021

function hue2rgb( p, q, h ) {
	h = ( h + 1 ) % 1;
	if ( h * 6 < 1 ) {
		return p + ( q - p ) * h * 6;
	}
	if ( h * 2 < 1) {
		return q;
	}
	if ( h * 3 < 2 ) {
		return p + ( q - p ) * ( ( 2 / 3 ) - h ) * 6;
	}
	return p;
}

spaces.hsla.to = function( rgba ) {
	if ( rgba[ 0 ] == null || rgba[ 1 ] == null || rgba[ 2 ] == null ) {
		return [ null, null, null, rgba[ 3 ] ];
	}
	var r = rgba[ 0 ] / 255,
		g = rgba[ 1 ] / 255,
		b = rgba[ 2 ] / 255,
		a = rgba[ 3 ],
		max = Math.max( r, g, b ),
		min = Math.min( r, g, b ),
		diff = max - min,
		add = max + min,
		l = add * 0.5,
		h, s;

	if ( min === max ) {
		h = 0;
	} else if ( r === max ) {
		h = ( 60 * ( g - b ) / diff ) + 360;
	} else if ( g === max ) {
		h = ( 60 * ( b - r ) / diff ) + 120;
	} else {
		h = ( 60 * ( r - g ) / diff ) + 240;
	}

	// chroma (diff) == 0 means greyscale which, by definition, saturation = 0%
	// otherwise, saturation is based on the ratio of chroma (diff) to lightness (add)
	if ( diff === 0 ) {
		s = 0;
	} else if ( l <= 0.5 ) {
		s = diff / add;
	} else {
		s = diff / ( 2 - add );
	}
	return [ Math.round(h) % 360, s, l, a == null ? 1 : a ];
};

spaces.hsla.from = function( hsla ) {
	if ( hsla[ 0 ] == null || hsla[ 1 ] == null || hsla[ 2 ] == null ) {
		return [ null, null, null, hsla[ 3 ] ];
	}
	var h = hsla[ 0 ] / 360,
		s = hsla[ 1 ],
		l = hsla[ 2 ],
		a = hsla[ 3 ],
		q = l <= 0.5 ? l * ( 1 + s ) : l + s - l * s,
		p = 2 * l - q;

	return [
		Math.round( hue2rgb( p, q, h + ( 1 / 3 ) ) * 255 ),
		Math.round( hue2rgb( p, q, h ) * 255 ),
		Math.round( hue2rgb( p, q, h - ( 1 / 3 ) ) * 255 ),
		a
	];
};

each( spaces, function( spaceName, space ) {
	var props = space.props,
		cache = space.cache,
		to = space.to,
		from = space.from;

	// makes rgba() and hsla()
	color.fn[ spaceName ] = function( value ) {

		// generate a cache for this space if it doesn't exist
		if ( to && !this[ cache ] ) {
			this[ cache ] = to( this._rgba );
		}
		if ( value === undefined ) {
			return this[ cache ].slice();
		}

		var ret,
			type = jQuery.type( value ),
			arr = ( type === "array" || type === "object" ) ? value : arguments,
			local = this[ cache ].slice();

		each( props, function( key, prop ) {
			var val = arr[ type === "object" ? key : prop.idx ];
			if ( val == null ) {
				val = local[ prop.idx ];
			}
			local[ prop.idx ] = clamp( val, prop );
		});

		if ( from ) {
			ret = color( from( local ) );
			ret[ cache ] = local;
			return ret;
		} else {
			return color( local );
		}
	};

	// makes red() green() blue() alpha() hue() saturation() lightness()
	each( props, function( key, prop ) {
		// alpha is included in more than one space
		if ( color.fn[ key ] ) {
			return;
		}
		color.fn[ key ] = function( value ) {
			var vtype = jQuery.type( value ),
				fn = ( key === "alpha" ? ( this._hsla ? "hsla" : "rgba" ) : spaceName ),
				local = this[ fn ](),
				cur = local[ prop.idx ],
				match;

			if ( vtype === "undefined" ) {
				return cur;
			}

			if ( vtype === "function" ) {
				value = value.call( this, cur );
				vtype = jQuery.type( value );
			}
			if ( value == null && prop.empty ) {
				return this;
			}
			if ( vtype === "string" ) {
				match = rplusequals.exec( value );
				if ( match ) {
					value = cur + parseFloat( match[ 2 ] ) * ( match[ 1 ] === "+" ? 1 : -1 );
				}
			}
			local[ prop.idx ] = value;
			return this[ fn ]( local );
		};
	});
});

// add cssHook and .fx.step function for each named hook.
// accept a space separated string of properties
color.hook = function( hook ) {
	var hooks = hook.split( " " );
	each( hooks, function( i, hook ) {
		jQuery.cssHooks[ hook ] = {
			set: function( elem, value ) {
				var parsed, curElem,
					backgroundColor = "";

				if ( value !== "transparent" && ( jQuery.type( value ) !== "string" || ( parsed = stringParse( value ) ) ) ) {
					value = color( parsed || value );
					if ( !support.rgba && value._rgba[ 3 ] !== 1 ) {
						curElem = hook === "backgroundColor" ? elem.parentNode : elem;
						while (
							(backgroundColor === "" || backgroundColor === "transparent") &&
							curElem && curElem.style
						) {
							try {
								backgroundColor = jQuery.css( curElem, "backgroundColor" );
								curElem = curElem.parentNode;
							} catch ( e ) {
							}
						}

						value = value.blend( backgroundColor && backgroundColor !== "transparent" ?
							backgroundColor :
							"_default" );
					}

					value = value.toRgbaString();
				}
				try {
					elem.style[ hook ] = value;
				} catch ( e ) {
					// wrapped to prevent IE from throwing errors on "invalid" values like 'auto' or 'inherit'
				}
			}
		};
		jQuery.fx.step[ hook ] = function( fx ) {
			if ( !fx.colorInit ) {
				fx.start = color( fx.elem, hook );
				fx.end = color( fx.end );
				fx.colorInit = true;
			}
			jQuery.cssHooks[ hook ].set( fx.elem, fx.start.transition( fx.end, fx.pos ) );
		};
	});

};

color.hook( stepHooks );

jQuery.cssHooks.borderColor = {
	expand: function( value ) {
		var expanded = {};

		each( [ "Top", "Right", "Bottom", "Left" ], function( i, part ) {
			expanded[ "border" + part + "Color" ] = value;
		});
		return expanded;
	}
};

// Basic color names only.
// Usage of any of the other color names requires adding yourself or including
// jquery.color.svg-names.js.
colors = jQuery.Color.names = {
	// 4.1. Basic color keywords
	aqua: "#00ffff",
	black: "#000000",
	blue: "#0000ff",
	fuchsia: "#ff00ff",
	gray: "#808080",
	green: "#008000",
	lime: "#00ff00",
	maroon: "#800000",
	navy: "#000080",
	olive: "#808000",
	purple: "#800080",
	red: "#ff0000",
	silver: "#c0c0c0",
	teal: "#008080",
	white: "#ffffff",
	yellow: "#ffff00",

	// 4.2.3. "transparent" color keyword
	transparent: [ null, null, null, 0 ],

	_default: "#ffffff"
};

})( jQuery );

/******************************************************************************/
/****************************** CLASS ANIMATIONS ******************************/
/******************************************************************************/
(function() {

var classAnimationActions = [ "add", "remove", "toggle" ],
	shorthandStyles = {
		border: 1,
		borderBottom: 1,
		borderColor: 1,
		borderLeft: 1,
		borderRight: 1,
		borderTop: 1,
		borderWidth: 1,
		margin: 1,
		padding: 1
	};

$.each([ "borderLeftStyle", "borderRightStyle", "borderBottomStyle", "borderTopStyle" ], function( _, prop ) {
	$.fx.step[ prop ] = function( fx ) {
		if ( fx.end !== "none" && !fx.setAttr || fx.pos === 1 && !fx.setAttr ) {
			jQuery.style( fx.elem, prop, fx.end );
			fx.setAttr = true;
		}
	};
});

function getElementStyles( elem ) {
	var key, len,
		style = elem.ownerDocument.defaultView ?
			elem.ownerDocument.defaultView.getComputedStyle( elem, null ) :
			elem.currentStyle,
		styles = {};

	if ( style && style.length && style[ 0 ] && style[ style[ 0 ] ] ) {
		len = style.length;
		while ( len-- ) {
			key = style[ len ];
			if ( typeof style[ key ] === "string" ) {
				styles[ $.camelCase( key ) ] = style[ key ];
			}
		}
	// support: Opera, IE <9
	} else {
		for ( key in style ) {
			if ( typeof style[ key ] === "string" ) {
				styles[ key ] = style[ key ];
			}
		}
	}

	return styles;
}

function styleDifference( oldStyle, newStyle ) {
	var diff = {},
		name, value;

	for ( name in newStyle ) {
		value = newStyle[ name ];
		if ( oldStyle[ name ] !== value ) {
			if ( !shorthandStyles[ name ] ) {
				if ( $.fx.step[ name ] || !isNaN( parseFloat( value ) ) ) {
					diff[ name ] = value;
				}
			}
		}
	}

	return diff;
}

// support: jQuery <1.8
if ( !$.fn.addBack ) {
	$.fn.addBack = function( selector ) {
		return this.add( selector == null ?
			this.prevObject : this.prevObject.filter( selector )
		);
	};
}

$.effects.animateClass = function( value, duration, easing, callback ) {
	var o = $.speed( duration, easing, callback );

	return this.queue( function() {
		var animated = $( this ),
			baseClass = animated.attr( "class" ) || "",
			applyClassChange,
			allAnimations = o.children ? animated.find( "*" ).addBack() : animated;

		// map the animated objects to store the original styles.
		allAnimations = allAnimations.map(function() {
			var el = $( this );
			return {
				el: el,
				start: getElementStyles( this )
			};
		});

		// apply class change
		applyClassChange = function() {
			$.each( classAnimationActions, function(i, action) {
				if ( value[ action ] ) {
					animated[ action + "Class" ]( value[ action ] );
				}
			});
		};
		applyClassChange();

		// map all animated objects again - calculate new styles and diff
		allAnimations = allAnimations.map(function() {
			this.end = getElementStyles( this.el[ 0 ] );
			this.diff = styleDifference( this.start, this.end );
			return this;
		});

		// apply original class
		animated.attr( "class", baseClass );

		// map all animated objects again - this time collecting a promise
		allAnimations = allAnimations.map(function() {
			var styleInfo = this,
				dfd = $.Deferred(),
				opts = $.extend({}, o, {
					queue: false,
					complete: function() {
						dfd.resolve( styleInfo );
					}
				});

			this.el.animate( this.diff, opts );
			return dfd.promise();
		});

		// once all animations have completed:
		$.when.apply( $, allAnimations.get() ).done(function() {

			// set the final class
			applyClassChange();

			// for each animated element,
			// clear all css properties that were animated
			$.each( arguments, function() {
				var el = this.el;
				$.each( this.diff, function(key) {
					el.css( key, "" );
				});
			});

			// this is guarnteed to be there if you use jQuery.speed()
			// it also handles dequeuing the next anim...
			o.complete.call( animated[ 0 ] );
		});
	});
};

$.fn.extend({
	addClass: (function( orig ) {
		return function( classNames, speed, easing, callback ) {
			return speed ?
				$.effects.animateClass.call( this,
					{ add: classNames }, speed, easing, callback ) :
				orig.apply( this, arguments );
		};
	})( $.fn.addClass ),

	removeClass: (function( orig ) {
		return function( classNames, speed, easing, callback ) {
			return arguments.length > 1 ?
				$.effects.animateClass.call( this,
					{ remove: classNames }, speed, easing, callback ) :
				orig.apply( this, arguments );
		};
	})( $.fn.removeClass ),

	toggleClass: (function( orig ) {
		return function( classNames, force, speed, easing, callback ) {
			if ( typeof force === "boolean" || force === undefined ) {
				if ( !speed ) {
					// without speed parameter
					return orig.apply( this, arguments );
				} else {
					return $.effects.animateClass.call( this,
						(force ? { add: classNames } : { remove: classNames }),
						speed, easing, callback );
				}
			} else {
				// without force parameter
				return $.effects.animateClass.call( this,
					{ toggle: classNames }, force, speed, easing );
			}
		};
	})( $.fn.toggleClass ),

	switchClass: function( remove, add, speed, easing, callback) {
		return $.effects.animateClass.call( this, {
			add: add,
			remove: remove
		}, speed, easing, callback );
	}
});

})();

/******************************************************************************/
/*********************************** EFFECTS **********************************/
/******************************************************************************/

(function() {

$.extend( $.effects, {
	version: "1.11.4",

	// Saves a set of properties in a data storage
	save: function( element, set ) {
		for ( var i = 0; i < set.length; i++ ) {
			if ( set[ i ] !== null ) {
				element.data( dataSpace + set[ i ], element[ 0 ].style[ set[ i ] ] );
			}
		}
	},

	// Restores a set of previously saved properties from a data storage
	restore: function( element, set ) {
		var val, i;
		for ( i = 0; i < set.length; i++ ) {
			if ( set[ i ] !== null ) {
				val = element.data( dataSpace + set[ i ] );
				// support: jQuery 1.6.2
				// http://bugs.jquery.com/ticket/9917
				// jQuery 1.6.2 incorrectly returns undefined for any falsy value.
				// We can't differentiate between "" and 0 here, so we just assume
				// empty string since it's likely to be a more common value...
				if ( val === undefined ) {
					val = "";
				}
				element.css( set[ i ], val );
			}
		}
	},

	setMode: function( el, mode ) {
		if (mode === "toggle") {
			mode = el.is( ":hidden" ) ? "show" : "hide";
		}
		return mode;
	},

	// Translates a [top,left] array into a baseline value
	// this should be a little more flexible in the future to handle a string & hash
	getBaseline: function( origin, original ) {
		var y, x;
		switch ( origin[ 0 ] ) {
			case "top": y = 0; break;
			case "middle": y = 0.5; break;
			case "bottom": y = 1; break;
			default: y = origin[ 0 ] / original.height;
		}
		switch ( origin[ 1 ] ) {
			case "left": x = 0; break;
			case "center": x = 0.5; break;
			case "right": x = 1; break;
			default: x = origin[ 1 ] / original.width;
		}
		return {
			x: x,
			y: y
		};
	},

	// Wraps the element around a wrapper that copies position properties
	createWrapper: function( element ) {

		// if the element is already wrapped, return it
		if ( element.parent().is( ".ui-effects-wrapper" )) {
			return element.parent();
		}

		// wrap the element
		var props = {
				width: element.outerWidth(true),
				height: element.outerHeight(true),
				"float": element.css( "float" )
			},
			wrapper = $( "<div></div>" )
				.addClass( "ui-effects-wrapper" )
				.css({
					fontSize: "100%",
					background: "transparent",
					border: "none",
					margin: 0,
					padding: 0
				}),
			// Store the size in case width/height are defined in % - Fixes #5245
			size = {
				width: element.width(),
				height: element.height()
			},
			active = document.activeElement;

		// support: Firefox
		// Firefox incorrectly exposes anonymous content
		// https://bugzilla.mozilla.org/show_bug.cgi?id=561664
		try {
			active.id;
		} catch ( e ) {
			active = document.body;
		}

		element.wrap( wrapper );

		// Fixes #7595 - Elements lose focus when wrapped.
		if ( element[ 0 ] === active || $.contains( element[ 0 ], active ) ) {
			$( active ).focus();
		}

		wrapper = element.parent(); //Hotfix for jQuery 1.4 since some change in wrap() seems to actually lose the reference to the wrapped element

		// transfer positioning properties to the wrapper
		if ( element.css( "position" ) === "static" ) {
			wrapper.css({ position: "relative" });
			element.css({ position: "relative" });
		} else {
			$.extend( props, {
				position: element.css( "position" ),
				zIndex: element.css( "z-index" )
			});
			$.each([ "top", "left", "bottom", "right" ], function(i, pos) {
				props[ pos ] = element.css( pos );
				if ( isNaN( parseInt( props[ pos ], 10 ) ) ) {
					props[ pos ] = "auto";
				}
			});
			element.css({
				position: "relative",
				top: 0,
				left: 0,
				right: "auto",
				bottom: "auto"
			});
		}
		element.css(size);

		return wrapper.css( props ).show();
	},

	removeWrapper: function( element ) {
		var active = document.activeElement;

		if ( element.parent().is( ".ui-effects-wrapper" ) ) {
			element.parent().replaceWith( element );

			// Fixes #7595 - Elements lose focus when wrapped.
			if ( element[ 0 ] === active || $.contains( element[ 0 ], active ) ) {
				$( active ).focus();
			}
		}

		return element;
	},

	setTransition: function( element, list, factor, value ) {
		value = value || {};
		$.each( list, function( i, x ) {
			var unit = element.cssUnit( x );
			if ( unit[ 0 ] > 0 ) {
				value[ x ] = unit[ 0 ] * factor + unit[ 1 ];
			}
		});
		return value;
	}
});

// return an effect options object for the given parameters:
function _normalizeArguments( effect, options, speed, callback ) {

	// allow passing all options as the first parameter
	if ( $.isPlainObject( effect ) ) {
		options = effect;
		effect = effect.effect;
	}

	// convert to an object
	effect = { effect: effect };

	// catch (effect, null, ...)
	if ( options == null ) {
		options = {};
	}

	// catch (effect, callback)
	if ( $.isFunction( options ) ) {
		callback = options;
		speed = null;
		options = {};
	}

	// catch (effect, speed, ?)
	if ( typeof options === "number" || $.fx.speeds[ options ] ) {
		callback = speed;
		speed = options;
		options = {};
	}

	// catch (effect, options, callback)
	if ( $.isFunction( speed ) ) {
		callback = speed;
		speed = null;
	}

	// add options to effect
	if ( options ) {
		$.extend( effect, options );
	}

	speed = speed || options.duration;
	effect.duration = $.fx.off ? 0 :
		typeof speed === "number" ? speed :
		speed in $.fx.speeds ? $.fx.speeds[ speed ] :
		$.fx.speeds._default;

	effect.complete = callback || options.complete;

	return effect;
}

function standardAnimationOption( option ) {
	// Valid standard speeds (nothing, number, named speed)
	if ( !option || typeof option === "number" || $.fx.speeds[ option ] ) {
		return true;
	}

	// Invalid strings - treat as "normal" speed
	if ( typeof option === "string" && !$.effects.effect[ option ] ) {
		return true;
	}

	// Complete callback
	if ( $.isFunction( option ) ) {
		return true;
	}

	// Options hash (but not naming an effect)
	if ( typeof option === "object" && !option.effect ) {
		return true;
	}

	// Didn't match any standard API
	return false;
}

$.fn.extend({
	effect: function( /* effect, options, speed, callback */ ) {
		var args = _normalizeArguments.apply( this, arguments ),
			mode = args.mode,
			queue = args.queue,
			effectMethod = $.effects.effect[ args.effect ];

		if ( $.fx.off || !effectMethod ) {
			// delegate to the original method (e.g., .show()) if possible
			if ( mode ) {
				return this[ mode ]( args.duration, args.complete );
			} else {
				return this.each( function() {
					if ( args.complete ) {
						args.complete.call( this );
					}
				});
			}
		}

		function run( next ) {
			var elem = $( this ),
				complete = args.complete,
				mode = args.mode;

			function done() {
				if ( $.isFunction( complete ) ) {
					complete.call( elem[0] );
				}
				if ( $.isFunction( next ) ) {
					next();
				}
			}

			// If the element already has the correct final state, delegate to
			// the core methods so the internal tracking of "olddisplay" works.
			if ( elem.is( ":hidden" ) ? mode === "hide" : mode === "show" ) {
				elem[ mode ]();
				done();
			} else {
				effectMethod.call( elem[0], args, done );
			}
		}

		return queue === false ? this.each( run ) : this.queue( queue || "fx", run );
	},

	show: (function( orig ) {
		return function( option ) {
			if ( standardAnimationOption( option ) ) {
				return orig.apply( this, arguments );
			} else {
				var args = _normalizeArguments.apply( this, arguments );
				args.mode = "show";
				return this.effect.call( this, args );
			}
		};
	})( $.fn.show ),

	hide: (function( orig ) {
		return function( option ) {
			if ( standardAnimationOption( option ) ) {
				return orig.apply( this, arguments );
			} else {
				var args = _normalizeArguments.apply( this, arguments );
				args.mode = "hide";
				return this.effect.call( this, args );
			}
		};
	})( $.fn.hide ),

	toggle: (function( orig ) {
		return function( option ) {
			if ( standardAnimationOption( option ) || typeof option === "boolean" ) {
				return orig.apply( this, arguments );
			} else {
				var args = _normalizeArguments.apply( this, arguments );
				args.mode = "toggle";
				return this.effect.call( this, args );
			}
		};
	})( $.fn.toggle ),

	// helper functions
	cssUnit: function(key) {
		var style = this.css( key ),
			val = [];

		$.each( [ "em", "px", "%", "pt" ], function( i, unit ) {
			if ( style.indexOf( unit ) > 0 ) {
				val = [ parseFloat( style ), unit ];
			}
		});
		return val;
	}
});

})();

/******************************************************************************/
/*********************************** EASING ***********************************/
/******************************************************************************/

(function() {

// based on easing equations from Robert Penner (http://www.robertpenner.com/easing)

var baseEasings = {};

$.each( [ "Quad", "Cubic", "Quart", "Quint", "Expo" ], function( i, name ) {
	baseEasings[ name ] = function( p ) {
		return Math.pow( p, i + 2 );
	};
});

$.extend( baseEasings, {
	Sine: function( p ) {
		return 1 - Math.cos( p * Math.PI / 2 );
	},
	Circ: function( p ) {
		return 1 - Math.sqrt( 1 - p * p );
	},
	Elastic: function( p ) {
		return p === 0 || p === 1 ? p :
			-Math.pow( 2, 8 * (p - 1) ) * Math.sin( ( (p - 1) * 80 - 7.5 ) * Math.PI / 15 );
	},
	Back: function( p ) {
		return p * p * ( 3 * p - 2 );
	},
	Bounce: function( p ) {
		var pow2,
			bounce = 4;

		while ( p < ( ( pow2 = Math.pow( 2, --bounce ) ) - 1 ) / 11 ) {}
		return 1 / Math.pow( 4, 3 - bounce ) - 7.5625 * Math.pow( ( pow2 * 3 - 2 ) / 22 - p, 2 );
	}
});

$.each( baseEasings, function( name, easeIn ) {
	$.easing[ "easeIn" + name ] = easeIn;
	$.easing[ "easeOut" + name ] = function( p ) {
		return 1 - easeIn( 1 - p );
	};
	$.easing[ "easeInOut" + name ] = function( p ) {
		return p < 0.5 ?
			easeIn( p * 2 ) / 2 :
			1 - easeIn( p * -2 + 2 ) / 2;
	};
});

})();

var effect = $.effects;


/*!
 * jQuery UI Effects Blind 1.11.4
 * http://jqueryui.com
 *
 * Copyright jQuery Foundation and other contributors
 * Released under the MIT license.
 * http://jquery.org/license
 *
 * http://api.jqueryui.com/blind-effect/
 */


var effectBlind = $.effects.effect.blind = function( o, done ) {
	// Create element
	var el = $( this ),
		rvertical = /up|down|vertical/,
		rpositivemotion = /up|left|vertical|horizontal/,
		props = [ "position", "top", "bottom", "left", "right", "height", "width" ],
		mode = $.effects.setMode( el, o.mode || "hide" ),
		direction = o.direction || "up",
		vertical = rvertical.test( direction ),
		ref = vertical ? "height" : "width",
		ref2 = vertical ? "top" : "left",
		motion = rpositivemotion.test( direction ),
		animation = {},
		show = mode === "show",
		wrapper, distance, margin;

	// if already wrapped, the wrapper's properties are my property. #6245
	if ( el.parent().is( ".ui-effects-wrapper" ) ) {
		$.effects.save( el.parent(), props );
	} else {
		$.effects.save( el, props );
	}
	el.show();
	wrapper = $.effects.createWrapper( el ).css({
		overflow: "hidden"
	});

	distance = wrapper[ ref ]();
	margin = parseFloat( wrapper.css( ref2 ) ) || 0;

	animation[ ref ] = show ? distance : 0;
	if ( !motion ) {
		el
			.css( vertical ? "bottom" : "right", 0 )
			.css( vertical ? "top" : "left", "auto" )
			.css({ position: "absolute" });

		animation[ ref2 ] = show ? margin : distance + margin;
	}

	// start at 0 if we are showing
	if ( show ) {
		wrapper.css( ref, 0 );
		if ( !motion ) {
			wrapper.css( ref2, margin + distance );
		}
	}

	// Animate
	wrapper.animate( animation, {
		duration: o.duration,
		easing: o.easing,
		queue: false,
		complete: function() {
			if ( mode === "hide" ) {
				el.hide();
			}
			$.effects.restore( el, props );
			$.effects.removeWrapper( el );
			done();
		}
	});
};


/*!
 * jQuery UI Effects Bounce 1.11.4
 * http://jqueryui.com
 *
 * Copyright jQuery Foundation and other contributors
 * Released under the MIT license.
 * http://jquery.org/license
 *
 * http://api.jqueryui.com/bounce-effect/
 */


var effectBounce = $.effects.effect.bounce = function( o, done ) {
	var el = $( this ),
		props = [ "position", "top", "bottom", "left", "right", "height", "width" ],

		// defaults:
		mode = $.effects.setMode( el, o.mode || "effect" ),
		hide = mode === "hide",
		show = mode === "show",
		direction = o.direction || "up",
		distance = o.distance,
		times = o.times || 5,

		// number of internal animations
		anims = times * 2 + ( show || hide ? 1 : 0 ),
		speed = o.duration / anims,
		easing = o.easing,

		// utility:
		ref = ( direction === "up" || direction === "down" ) ? "top" : "left",
		motion = ( direction === "up" || direction === "left" ),
		i,
		upAnim,
		downAnim,

		// we will need to re-assemble the queue to stack our animations in place
		queue = el.queue(),
		queuelen = queue.length;

	// Avoid touching opacity to prevent clearType and PNG issues in IE
	if ( show || hide ) {
		props.push( "opacity" );
	}

	$.effects.save( el, props );
	el.show();
	$.effects.createWrapper( el ); // Create Wrapper

	// default distance for the BIGGEST bounce is the outer Distance / 3
	if ( !distance ) {
		distance = el[ ref === "top" ? "outerHeight" : "outerWidth" ]() / 3;
	}

	if ( show ) {
		downAnim = { opacity: 1 };
		downAnim[ ref ] = 0;

		// if we are showing, force opacity 0 and set the initial position
		// then do the "first" animation
		el.css( "opacity", 0 )
			.css( ref, motion ? -distance * 2 : distance * 2 )
			.animate( downAnim, speed, easing );
	}

	// start at the smallest distance if we are hiding
	if ( hide ) {
		distance = distance / Math.pow( 2, times - 1 );
	}

	downAnim = {};
	downAnim[ ref ] = 0;
	// Bounces up/down/left/right then back to 0 -- times * 2 animations happen here
	for ( i = 0; i < times; i++ ) {
		upAnim = {};
		upAnim[ ref ] = ( motion ? "-=" : "+=" ) + distance;

		el.animate( upAnim, speed, easing )
			.animate( downAnim, speed, easing );

		distance = hide ? distance * 2 : distance / 2;
	}

	// Last Bounce when Hiding
	if ( hide ) {
		upAnim = { opacity: 0 };
		upAnim[ ref ] = ( motion ? "-=" : "+=" ) + distance;

		el.animate( upAnim, speed, easing );
	}

	el.queue(function() {
		if ( hide ) {
			el.hide();
		}
		$.effects.restore( el, props );
		$.effects.removeWrapper( el );
		done();
	});

	// inject all the animations we just queued to be first in line (after "inprogress")
	if ( queuelen > 1) {
		queue.splice.apply( queue,
			[ 1, 0 ].concat( queue.splice( queuelen, anims + 1 ) ) );
	}
	el.dequeue();

};


/*!
 * jQuery UI Effects Clip 1.11.4
 * http://jqueryui.com
 *
 * Copyright jQuery Foundation and other contributors
 * Released under the MIT license.
 * http://jquery.org/license
 *
 * http://api.jqueryui.com/clip-effect/
 */


var effectClip = $.effects.effect.clip = function( o, done ) {
	// Create element
	var el = $( this ),
		props = [ "position", "top", "bottom", "left", "right", "height", "width" ],
		mode = $.effects.setMode( el, o.mode || "hide" ),
		show = mode === "show",
		direction = o.direction || "vertical",
		vert = direction === "vertical",
		size = vert ? "height" : "width",
		position = vert ? "top" : "left",
		animation = {},
		wrapper, animate, distance;

	// Save & Show
	$.effects.save( el, props );
	el.show();

	// Create Wrapper
	wrapper = $.effects.createWrapper( el ).css({
		overflow: "hidden"
	});
	animate = ( el[0].tagName === "IMG" ) ? wrapper : el;
	distance = animate[ size ]();

	// Shift
	if ( show ) {
		animate.css( size, 0 );
		animate.css( position, distance / 2 );
	}

	// Create Animation Object:
	animation[ size ] = show ? distance : 0;
	animation[ position ] = show ? 0 : distance / 2;

	// Animate
	animate.animate( animation, {
		queue: false,
		duration: o.duration,
		easing: o.easing,
		complete: function() {
			if ( !show ) {
				el.hide();
			}
			$.effects.restore( el, props );
			$.effects.removeWrapper( el );
			done();
		}
	});

};


/*!
 * jQuery UI Effects Drop 1.11.4
 * http://jqueryui.com
 *
 * Copyright jQuery Foundation and other contributors
 * Released under the MIT license.
 * http://jquery.org/license
 *
 * http://api.jqueryui.com/drop-effect/
 */


var effectDrop = $.effects.effect.drop = function( o, done ) {

	var el = $( this ),
		props = [ "position", "top", "bottom", "left", "right", "opacity", "height", "width" ],
		mode = $.effects.setMode( el, o.mode || "hide" ),
		show = mode === "show",
		direction = o.direction || "left",
		ref = ( direction === "up" || direction === "down" ) ? "top" : "left",
		motion = ( direction === "up" || direction === "left" ) ? "pos" : "neg",
		animation = {
			opacity: show ? 1 : 0
		},
		distance;

	// Adjust
	$.effects.save( el, props );
	el.show();
	$.effects.createWrapper( el );

	distance = o.distance || el[ ref === "top" ? "outerHeight" : "outerWidth" ]( true ) / 2;

	if ( show ) {
		el
			.css( "opacity", 0 )
			.css( ref, motion === "pos" ? -distance : distance );
	}

	// Animation
	animation[ ref ] = ( show ?
		( motion === "pos" ? "+=" : "-=" ) :
		( motion === "pos" ? "-=" : "+=" ) ) +
		distance;

	// Animate
	el.animate( animation, {
		queue: false,
		duration: o.duration,
		easing: o.easing,
		complete: function() {
			if ( mode === "hide" ) {
				el.hide();
			}
			$.effects.restore( el, props );
			$.effects.removeWrapper( el );
			done();
		}
	});
};


/*!
 * jQuery UI Effects Explode 1.11.4
 * http://jqueryui.com
 *
 * Copyright jQuery Foundation and other contributors
 * Released under the MIT license.
 * http://jquery.org/license
 *
 * http://api.jqueryui.com/explode-effect/
 */


var effectExplode = $.effects.effect.explode = function( o, done ) {

	var rows = o.pieces ? Math.round( Math.sqrt( o.pieces ) ) : 3,
		cells = rows,
		el = $( this ),
		mode = $.effects.setMode( el, o.mode || "hide" ),
		show = mode === "show",

		// show and then visibility:hidden the element before calculating offset
		offset = el.show().css( "visibility", "hidden" ).offset(),

		// width and height of a piece
		width = Math.ceil( el.outerWidth() / cells ),
		height = Math.ceil( el.outerHeight() / rows ),
		pieces = [],

		// loop
		i, j, left, top, mx, my;

	// children animate complete:
	function childComplete() {
		pieces.push( this );
		if ( pieces.length === rows * cells ) {
			animComplete();
		}
	}

	// clone the element for each row and cell.
	for ( i = 0; i < rows ; i++ ) { // ===>
		top = offset.top + i * height;
		my = i - ( rows - 1 ) / 2 ;

		for ( j = 0; j < cells ; j++ ) { // |||
			left = offset.left + j * width;
			mx = j - ( cells - 1 ) / 2 ;

			// Create a clone of the now hidden main element that will be absolute positioned
			// within a wrapper div off the -left and -top equal to size of our pieces
			el
				.clone()
				.appendTo( "body" )
				.wrap( "<div></div>" )
				.css({
					position: "absolute",
					visibility: "visible",
					left: -j * width,
					top: -i * height
				})

			// select the wrapper - make it overflow: hidden and absolute positioned based on
			// where the original was located +left and +top equal to the size of pieces
				.parent()
				.addClass( "ui-effects-explode" )
				.css({
					position: "absolute",
					overflow: "hidden",
					width: width,
					height: height,
					left: left + ( show ? mx * width : 0 ),
					top: top + ( show ? my * height : 0 ),
					opacity: show ? 0 : 1
				}).animate({
					left: left + ( show ? 0 : mx * width ),
					top: top + ( show ? 0 : my * height ),
					opacity: show ? 1 : 0
				}, o.duration || 500, o.easing, childComplete );
		}
	}

	function animComplete() {
		el.css({
			visibility: "visible"
		});
		$( pieces ).remove();
		if ( !show ) {
			el.hide();
		}
		done();
	}
};


/*!
 * jQuery UI Effects Fade 1.11.4
 * http://jqueryui.com
 *
 * Copyright jQuery Foundation and other contributors
 * Released under the MIT license.
 * http://jquery.org/license
 *
 * http://api.jqueryui.com/fade-effect/
 */


var effectFade = $.effects.effect.fade = function( o, done ) {
	var el = $( this ),
		mode = $.effects.setMode( el, o.mode || "toggle" );

	el.animate({
		opacity: mode
	}, {
		queue: false,
		duration: o.duration,
		easing: o.easing,
		complete: done
	});
};


/*!
 * jQuery UI Effects Fold 1.11.4
 * http://jqueryui.com
 *
 * Copyright jQuery Foundation and other contributors
 * Released under the MIT license.
 * http://jquery.org/license
 *
 * http://api.jqueryui.com/fold-effect/
 */


var effectFold = $.effects.effect.fold = function( o, done ) {

	// Create element
	var el = $( this ),
		props = [ "position", "top", "bottom", "left", "right", "height", "width" ],
		mode = $.effects.setMode( el, o.mode || "hide" ),
		show = mode === "show",
		hide = mode === "hide",
		size = o.size || 15,
		percent = /([0-9]+)%/.exec( size ),
		horizFirst = !!o.horizFirst,
		widthFirst = show !== horizFirst,
		ref = widthFirst ? [ "width", "height" ] : [ "height", "width" ],
		duration = o.duration / 2,
		wrapper, distance,
		animation1 = {},
		animation2 = {};

	$.effects.save( el, props );
	el.show();

	// Create Wrapper
	wrapper = $.effects.createWrapper( el ).css({
		overflow: "hidden"
	});
	distance = widthFirst ?
		[ wrapper.width(), wrapper.height() ] :
		[ wrapper.height(), wrapper.width() ];

	if ( percent ) {
		size = parseInt( percent[ 1 ], 10 ) / 100 * distance[ hide ? 0 : 1 ];
	}
	if ( show ) {
		wrapper.css( horizFirst ? {
			height: 0,
			width: size
		} : {
			height: size,
			width: 0
		});
	}

	// Animation
	animation1[ ref[ 0 ] ] = show ? distance[ 0 ] : size;
	animation2[ ref[ 1 ] ] = show ? distance[ 1 ] : 0;

	// Animate
	wrapper
		.animate( animation1, duration, o.easing )
		.animate( animation2, duration, o.easing, function() {
			if ( hide ) {
				el.hide();
			}
			$.effects.restore( el, props );
			$.effects.removeWrapper( el );
			done();
		});

};


/*!
 * jQuery UI Effects Highlight 1.11.4
 * http://jqueryui.com
 *
 * Copyright jQuery Foundation and other contributors
 * Released under the MIT license.
 * http://jquery.org/license
 *
 * http://api.jqueryui.com/highlight-effect/
 */


var effectHighlight = $.effects.effect.highlight = function( o, done ) {
	var elem = $( this ),
		props = [ "backgroundImage", "backgroundColor", "opacity" ],
		mode = $.effects.setMode( elem, o.mode || "show" ),
		animation = {
			backgroundColor: elem.css( "backgroundColor" )
		};

	if (mode === "hide") {
		animation.opacity = 0;
	}

	$.effects.save( elem, props );

	elem
		.show()
		.css({
			backgroundImage: "none",
			backgroundColor: o.color || "#ffff99"
		})
		.animate( animation, {
			queue: false,
			duration: o.duration,
			easing: o.easing,
			complete: function() {
				if ( mode === "hide" ) {
					elem.hide();
				}
				$.effects.restore( elem, props );
				done();
			}
		});
};


/*!
 * jQuery UI Effects Size 1.11.4
 * http://jqueryui.com
 *
 * Copyright jQuery Foundation and other contributors
 * Released under the MIT license.
 * http://jquery.org/license
 *
 * http://api.jqueryui.com/size-effect/
 */


var effectSize = $.effects.effect.size = function( o, done ) {

	// Create element
	var original, baseline, factor,
		el = $( this ),
		props0 = [ "position", "top", "bottom", "left", "right", "width", "height", "overflow", "opacity" ],

		// Always restore
		props1 = [ "position", "top", "bottom", "left", "right", "overflow", "opacity" ],

		// Copy for children
		props2 = [ "width", "height", "overflow" ],
		cProps = [ "fontSize" ],
		vProps = [ "borderTopWidth", "borderBottomWidth", "paddingTop", "paddingBottom" ],
		hProps = [ "borderLeftWidth", "borderRightWidth", "paddingLeft", "paddingRight" ],

		// Set options
		mode = $.effects.setMode( el, o.mode || "effect" ),
		restore = o.restore || mode !== "effect",
		scale = o.scale || "both",
		origin = o.origin || [ "middle", "center" ],
		position = el.css( "position" ),
		props = restore ? props0 : props1,
		zero = {
			height: 0,
			width: 0,
			outerHeight: 0,
			outerWidth: 0
		};

	if ( mode === "show" ) {
		el.show();
	}
	original = {
		height: el.height(),
		width: el.width(),
		outerHeight: el.outerHeight(),
		outerWidth: el.outerWidth()
	};

	if ( o.mode === "toggle" && mode === "show" ) {
		el.from = o.to || zero;
		el.to = o.from || original;
	} else {
		el.from = o.from || ( mode === "show" ? zero : original );
		el.to = o.to || ( mode === "hide" ? zero : original );
	}

	// Set scaling factor
	factor = {
		from: {
			y: el.from.height / original.height,
			x: el.from.width / original.width
		},
		to: {
			y: el.to.height / original.height,
			x: el.to.width / original.width
		}
	};

	// Scale the css box
	if ( scale === "box" || scale === "both" ) {

		// Vertical props scaling
		if ( factor.from.y !== factor.to.y ) {
			props = props.concat( vProps );
			el.from = $.effects.setTransition( el, vProps, factor.from.y, el.from );
			el.to = $.effects.setTransition( el, vProps, factor.to.y, el.to );
		}

		// Horizontal props scaling
		if ( factor.from.x !== factor.to.x ) {
			props = props.concat( hProps );
			el.from = $.effects.setTransition( el, hProps, factor.from.x, el.from );
			el.to = $.effects.setTransition( el, hProps, factor.to.x, el.to );
		}
	}

	// Scale the content
	if ( scale === "content" || scale === "both" ) {

		// Vertical props scaling
		if ( factor.from.y !== factor.to.y ) {
			props = props.concat( cProps ).concat( props2 );
			el.from = $.effects.setTransition( el, cProps, factor.from.y, el.from );
			el.to = $.effects.setTransition( el, cProps, factor.to.y, el.to );
		}
	}

	$.effects.save( el, props );
	el.show();
	$.effects.createWrapper( el );
	el.css( "overflow", "hidden" ).css( el.from );

	// Adjust
	if (origin) { // Calculate baseline shifts
		baseline = $.effects.getBaseline( origin, original );
		el.from.top = ( original.outerHeight - el.outerHeight() ) * baseline.y;
		el.from.left = ( original.outerWidth - el.outerWidth() ) * baseline.x;
		el.to.top = ( original.outerHeight - el.to.outerHeight ) * baseline.y;
		el.to.left = ( original.outerWidth - el.to.outerWidth ) * baseline.x;
	}
	el.css( el.from ); // set top & left

	// Animate
	if ( scale === "content" || scale === "both" ) { // Scale the children

		// Add margins/font-size
		vProps = vProps.concat([ "marginTop", "marginBottom" ]).concat(cProps);
		hProps = hProps.concat([ "marginLeft", "marginRight" ]);
		props2 = props0.concat(vProps).concat(hProps);

		el.find( "*[width]" ).each( function() {
			var child = $( this ),
				c_original = {
					height: child.height(),
					width: child.width(),
					outerHeight: child.outerHeight(),
					outerWidth: child.outerWidth()
				};
			if (restore) {
				$.effects.save(child, props2);
			}

			child.from = {
				height: c_original.height * factor.from.y,
				width: c_original.width * factor.from.x,
				outerHeight: c_original.outerHeight * factor.from.y,
				outerWidth: c_original.outerWidth * factor.from.x
			};
			child.to = {
				height: c_original.height * factor.to.y,
				width: c_original.width * factor.to.x,
				outerHeight: c_original.height * factor.to.y,
				outerWidth: c_original.width * factor.to.x
			};

			// Vertical props scaling
			if ( factor.from.y !== factor.to.y ) {
				child.from = $.effects.setTransition( child, vProps, factor.from.y, child.from );
				child.to = $.effects.setTransition( child, vProps, factor.to.y, child.to );
			}

			// Horizontal props scaling
			if ( factor.from.x !== factor.to.x ) {
				child.from = $.effects.setTransition( child, hProps, factor.from.x, child.from );
				child.to = $.effects.setTransition( child, hProps, factor.to.x, child.to );
			}

			// Animate children
			child.css( child.from );
			child.animate( child.to, o.duration, o.easing, function() {

				// Restore children
				if ( restore ) {
					$.effects.restore( child, props2 );
				}
			});
		});
	}

	// Animate
	el.animate( el.to, {
		queue: false,
		duration: o.duration,
		easing: o.easing,
		complete: function() {
			if ( el.to.opacity === 0 ) {
				el.css( "opacity", el.from.opacity );
			}
			if ( mode === "hide" ) {
				el.hide();
			}
			$.effects.restore( el, props );
			if ( !restore ) {

				// we need to calculate our new positioning based on the scaling
				if ( position === "static" ) {
					el.css({
						position: "relative",
						top: el.to.top,
						left: el.to.left
					});
				} else {
					$.each([ "top", "left" ], function( idx, pos ) {
						el.css( pos, function( _, str ) {
							var val = parseInt( str, 10 ),
								toRef = idx ? el.to.left : el.to.top;

							// if original was "auto", recalculate the new value from wrapper
							if ( str === "auto" ) {
								return toRef + "px";
							}

							return val + toRef + "px";
						});
					});
				}
			}

			$.effects.removeWrapper( el );
			done();
		}
	});

};


/*!
 * jQuery UI Effects Scale 1.11.4
 * http://jqueryui.com
 *
 * Copyright jQuery Foundation and other contributors
 * Released under the MIT license.
 * http://jquery.org/license
 *
 * http://api.jqueryui.com/scale-effect/
 */


var effectScale = $.effects.effect.scale = function( o, done ) {

	// Create element
	var el = $( this ),
		options = $.extend( true, {}, o ),
		mode = $.effects.setMode( el, o.mode || "effect" ),
		percent = parseInt( o.percent, 10 ) ||
			( parseInt( o.percent, 10 ) === 0 ? 0 : ( mode === "hide" ? 0 : 100 ) ),
		direction = o.direction || "both",
		origin = o.origin,
		original = {
			height: el.height(),
			width: el.width(),
			outerHeight: el.outerHeight(),
			outerWidth: el.outerWidth()
		},
		factor = {
			y: direction !== "horizontal" ? (percent / 100) : 1,
			x: direction !== "vertical" ? (percent / 100) : 1
		};

	// We are going to pass this effect to the size effect:
	options.effect = "size";
	options.queue = false;
	options.complete = done;

	// Set default origin and restore for show/hide
	if ( mode !== "effect" ) {
		options.origin = origin || [ "middle", "center" ];
		options.restore = true;
	}

	options.from = o.from || ( mode === "show" ? {
		height: 0,
		width: 0,
		outerHeight: 0,
		outerWidth: 0
	} : original );
	options.to = {
		height: original.height * factor.y,
		width: original.width * factor.x,
		outerHeight: original.outerHeight * factor.y,
		outerWidth: original.outerWidth * factor.x
	};

	// Fade option to support puff
	if ( options.fade ) {
		if ( mode === "show" ) {
			options.from.opacity = 0;
			options.to.opacity = 1;
		}
		if ( mode === "hide" ) {
			options.from.opacity = 1;
			options.to.opacity = 0;
		}
	}

	// Animate
	el.effect( options );

};


/*!
 * jQuery UI Effects Puff 1.11.4
 * http://jqueryui.com
 *
 * Copyright jQuery Foundation and other contributors
 * Released under the MIT license.
 * http://jquery.org/license
 *
 * http://api.jqueryui.com/puff-effect/
 */


var effectPuff = $.effects.effect.puff = function( o, done ) {
	var elem = $( this ),
		mode = $.effects.setMode( elem, o.mode || "hide" ),
		hide = mode === "hide",
		percent = parseInt( o.percent, 10 ) || 150,
		factor = percent / 100,
		original = {
			height: elem.height(),
			width: elem.width(),
			outerHeight: elem.outerHeight(),
			outerWidth: elem.outerWidth()
		};

	$.extend( o, {
		effect: "scale",
		queue: false,
		fade: true,
		mode: mode,
		complete: done,
		percent: hide ? percent : 100,
		from: hide ?
			original :
			{
				height: original.height * factor,
				width: original.width * factor,
				outerHeight: original.outerHeight * factor,
				outerWidth: original.outerWidth * factor
			}
	});

	elem.effect( o );
};


/*!
 * jQuery UI Effects Pulsate 1.11.4
 * http://jqueryui.com
 *
 * Copyright jQuery Foundation and other contributors
 * Released under the MIT license.
 * http://jquery.org/license
 *
 * http://api.jqueryui.com/pulsate-effect/
 */


var effectPulsate = $.effects.effect.pulsate = function( o, done ) {
	var elem = $( this ),
		mode = $.effects.setMode( elem, o.mode || "show" ),
		show = mode === "show",
		hide = mode === "hide",
		showhide = ( show || mode === "hide" ),

		// showing or hiding leaves of the "last" animation
		anims = ( ( o.times || 5 ) * 2 ) + ( showhide ? 1 : 0 ),
		duration = o.duration / anims,
		animateTo = 0,
		queue = elem.queue(),
		queuelen = queue.length,
		i;

	if ( show || !elem.is(":visible")) {
		elem.css( "opacity", 0 ).show();
		animateTo = 1;
	}

	// anims - 1 opacity "toggles"
	for ( i = 1; i < anims; i++ ) {
		elem.animate({
			opacity: animateTo
		}, duration, o.easing );
		animateTo = 1 - animateTo;
	}

	elem.animate({
		opacity: animateTo
	}, duration, o.easing);

	elem.queue(function() {
		if ( hide ) {
			elem.hide();
		}
		done();
	});

	// We just queued up "anims" animations, we need to put them next in the queue
	if ( queuelen > 1 ) {
		queue.splice.apply( queue,
			[ 1, 0 ].concat( queue.splice( queuelen, anims + 1 ) ) );
	}
	elem.dequeue();
};


/*!
 * jQuery UI Effects Shake 1.11.4
 * http://jqueryui.com
 *
 * Copyright jQuery Foundation and other contributors
 * Released under the MIT license.
 * http://jquery.org/license
 *
 * http://api.jqueryui.com/shake-effect/
 */


var effectShake = $.effects.effect.shake = function( o, done ) {

	var el = $( this ),
		props = [ "position", "top", "bottom", "left", "right", "height", "width" ],
		mode = $.effects.setMode( el, o.mode || "effect" ),
		direction = o.direction || "left",
		distance = o.distance || 20,
		times = o.times || 3,
		anims = times * 2 + 1,
		speed = Math.round( o.duration / anims ),
		ref = (direction === "up" || direction === "down") ? "top" : "left",
		positiveMotion = (direction === "up" || direction === "left"),
		animation = {},
		animation1 = {},
		animation2 = {},
		i,

		// we will need to re-assemble the queue to stack our animations in place
		queue = el.queue(),
		queuelen = queue.length;

	$.effects.save( el, props );
	el.show();
	$.effects.createWrapper( el );

	// Animation
	animation[ ref ] = ( positiveMotion ? "-=" : "+=" ) + distance;
	animation1[ ref ] = ( positiveMotion ? "+=" : "-=" ) + distance * 2;
	animation2[ ref ] = ( positiveMotion ? "-=" : "+=" ) + distance * 2;

	// Animate
	el.animate( animation, speed, o.easing );

	// Shakes
	for ( i = 1; i < times; i++ ) {
		el.animate( animation1, speed, o.easing ).animate( animation2, speed, o.easing );
	}
	el
		.animate( animation1, speed, o.easing )
		.animate( animation, speed / 2, o.easing )
		.queue(function() {
			if ( mode === "hide" ) {
				el.hide();
			}
			$.effects.restore( el, props );
			$.effects.removeWrapper( el );
			done();
		});

	// inject all the animations we just queued to be first in line (after "inprogress")
	if ( queuelen > 1) {
		queue.splice.apply( queue,
			[ 1, 0 ].concat( queue.splice( queuelen, anims + 1 ) ) );
	}
	el.dequeue();

};


/*!
 * jQuery UI Effects Slide 1.11.4
 * http://jqueryui.com
 *
 * Copyright jQuery Foundation and other contributors
 * Released under the MIT license.
 * http://jquery.org/license
 *
 * http://api.jqueryui.com/slide-effect/
 */


var effectSlide = $.effects.effect.slide = function( o, done ) {

	// Create element
	var el = $( this ),
		props = [ "position", "top", "bottom", "left", "right", "width", "height" ],
		mode = $.effects.setMode( el, o.mode || "show" ),
		show = mode === "show",
		direction = o.direction || "left",
		ref = (direction === "up" || direction === "down") ? "top" : "left",
		positiveMotion = (direction === "up" || direction === "left"),
		distance,
		animation = {};

	// Adjust
	$.effects.save( el, props );
	el.show();
	distance = o.distance || el[ ref === "top" ? "outerHeight" : "outerWidth" ]( true );

	$.effects.createWrapper( el ).css({
		overflow: "hidden"
	});

	if ( show ) {
		el.css( ref, positiveMotion ? (isNaN(distance) ? "-" + distance : -distance) : distance );
	}

	// Animation
	animation[ ref ] = ( show ?
		( positiveMotion ? "+=" : "-=") :
		( positiveMotion ? "-=" : "+=")) +
		distance;

	// Animate
	el.animate( animation, {
		queue: false,
		duration: o.duration,
		easing: o.easing,
		complete: function() {
			if ( mode === "hide" ) {
				el.hide();
			}
			$.effects.restore( el, props );
			$.effects.removeWrapper( el );
			done();
		}
	});
};


/*!
 * jQuery UI Effects Transfer 1.11.4
 * http://jqueryui.com
 *
 * Copyright jQuery Foundation and other contributors
 * Released under the MIT license.
 * http://jquery.org/license
 *
 * http://api.jqueryui.com/transfer-effect/
 */


var effectTransfer = $.effects.effect.transfer = function( o, done ) {
	var elem = $( this ),
		target = $( o.to ),
		targetFixed = target.css( "position" ) === "fixed",
		body = $("body"),
		fixTop = targetFixed ? body.scrollTop() : 0,
		fixLeft = targetFixed ? body.scrollLeft() : 0,
		endPosition = target.offset(),
		animation = {
			top: endPosition.top - fixTop,
			left: endPosition.left - fixLeft,
			height: target.innerHeight(),
			width: target.innerWidth()
		},
		startPosition = elem.offset(),
		transfer = $( "<div class='ui-effects-transfer'></div>" )
			.appendTo( document.body )
			.addClass( o.className )
			.css({
				top: startPosition.top - fixTop,
				left: startPosition.left - fixLeft,
				height: elem.innerHeight(),
				width: elem.innerWidth(),
				position: targetFixed ? "fixed" : "absolute"
			})
			.animate( animation, o.duration, o.easing, function() {
				transfer.remove();
				done();
			});
};


}));
```
