## Supplementary material for "Characterizing and inferring quantitative cell cycle phase in single-cell RNA-seq data analysis": Source code of our analysis, including scripts and data necessary to reproduce the work.: jquery-ui.min.js

```
/*! jQuery UI - v1.11.4 - 2016-01-05
* http://jqueryui.com
* Includes: core.js, widget.js, mouse.js, position.js, draggable.js, droppable.js, resizable.js, selectable.js, sortable.js, accordion.js, autocomplete.js, button.js, dialog.js, menu.js, progressbar.js, selectmenu.js, slider.js, spinner.js, tabs.js, tooltip.js, effect.js, effect-blind.js, effect-bounce.js, effect-clip.js, effect-drop.js, effect-explode.js, effect-fade.js, effect-fold.js, effect-highlight.js, effect-puff.js, effect-pulsate.js, effect-scale.js, effect-shake.js, effect-size.js, effect-slide.js, effect-transfer.js
* Copyright jQuery Foundation and other contributors; Licensed MIT */

(function(e){"function"==typeof define&&define.amd?define(["jquery"],e):e(jQuery)})(function(e){function t(t,s){var n,a,o,r=t.nodeName.toLowerCase();return"area"===r?(n=t.parentNode,a=n.name,t.href&&a&&"map"===n.nodeName.toLowerCase()?(o=e("img[usemap='#"+a+"']")[0],!!o&&i(o)):!1):(/^(input|select|textarea|button|object)$/.test(r)?!t.disabled:"a"===r?t.href||s:s)&&i(t)}function i(t){return e.expr.filters.visible(t)&&!e(t).parents().addBack().filter(function(){return"hidden"===e.css(this,"visibility")}).length}function s(e){return function(){var t=this.element.val();e.apply(this,arguments),this._refresh(),t!==this.element.val()&&this._trigger("change")}}e.ui=e.ui||{},e.extend(e.ui,{version:"1.11.4",keyCode:{BACKSPACE:8,COMMA:188,DELETE:46,DOWN:40,END:35,ENTER:13,ESCAPE:27,HOME:36,LEFT:37,PAGE_DOWN:34,PAGE_UP:33,PERIOD:190,RIGHT:39,SPACE:32,TAB:9,UP:38}}),e.fn.extend({scrollParent:function(t){var i=this.css("position"),s="absolute"===i,n=t?/(auto|scroll|hidden)/:/(auto|scroll)/,a=this.parents().filter(function(){var t=e(this);return s&&"static"===t.css("position")?!1:n.test(t.css("overflow")+t.css("overflow-y")+t.css("overflow-x"))}).eq(0);return"fixed"!==i&&a.length?a:e(this[0].ownerDocument||document)},uniqueId:function(){var e=0;return function(){return this.each(function(){this.id||(this.id="ui-id-"+ ++e)})}}(),removeUniqueId:function(){return this.each(function(){/^ui-id-\d+$/.test(this.id)&&e(this).removeAttr("id")})}}),e.extend(e.expr[":"],{data:e.expr.createPseudo?e.expr.createPseudo(function(t){return function(i){return!!e.data(i,t)}}):function(t,i,s){return!!e.data(t,s[3])},focusable:function(i){return t(i,!isNaN(e.attr(i,"tabindex")))},tabbable:function(i){var s=e.attr(i,"tabindex"),n=isNaN(s);return(n||s>=0)&&t(i,!n)}}),e("<a>").outerWidth(1).jquery||e.each(["Width","Height"],function(t,i){function s(t,i,s,a){return e.each(n,function(){i-=parseFloat(e.css(t,"padding"+this))||0,s&&(i-=parseFloat(e.css(t,"border"+this+"Width"))||0),a&&(i-=parseFloat(e.css(t,"margin"+this))||0)}),i}var n="Width"===i?["Left","Right"]:["Top","Bottom"],a=i.toLowerCase(),o={innerWidth:e.fn.innerWidth,innerHeight:e.fn.innerHeight,outerWidth:e.fn.outerWidth,outerHeight:e.fn.outerHeight};e.fn["inner"+i]=function(t){return void 0===t?o["inner"+i].call(this):this.each(function(){e(this).css(a,s(this,t)+"px")})},e.fn["outer"+i]=function(t,n){return"number"!=typeof t?o["outer"+i].call(this,t):this.each(function(){e(this).css(a,s(this,t,!0,n)+"px")})}}),e.fn.addBack||(e.fn.addBack=function(e){return this.add(null==e?this.prevObject:this.prevObject.filter(e))}),e("<a>").data("a-b","a").removeData("a-b").data("a-b")&&(e.fn.removeData=function(t){return function(i){return arguments.length?t.call(this,e.camelCase(i)):t.call(this)}}(e.fn.removeData)),e.ui.ie=!!/msie [\w.]+/.exec(navigator.userAgent.toLowerCase()),e.fn.extend({focus:function(t){return function(i,s){return"number"==typeof i?this.each(function(){var t=this;setTimeout(function(){e(t).focus(),s&&s.call(t)},i)}):t.apply(this,arguments)}}(e.fn.focus),disableSelection:function(){var e="onselectstart"in document.createElement("div")?"selectstart":"mousedown";return function(){return this.bind(e+".ui-disableSelection",function(e){e.preventDefault()})}}(),enableSelection:function(){return this.unbind(".ui-disableSelection")},zIndex:function(t){if(void 0!==t)return this.css("zIndex",t);if(this.length)for(var i,s,n=e(this[0]);n.length&&n[0]!==document;){if(i=n.css("position"),("absolute"===i||"relative"===i||"fixed"===i)&&(s=parseInt(n.css("zIndex"),10),!isNaN(s)&&0!==s))return s;n=n.parent()}return 0}}),e.ui.plugin={add:function(t,i,s){var n,a=e.ui[t].prototype;for(n in s)a.plugins[n]=a.plugins[n]||[],a.plugins[n].push([i,s[n]])},call:function(e,t,i,s){var n,a=e.plugins[t];if(a&&(s||e.element[0].parentNode&&11!==e.element[0].parentNode.nodeType))for(n=0;a.length>n;n++)e.options[a[n][0]]&&a[n][1].apply(e.element,i)}};var n=0,a=Array.prototype.slice;e.cleanData=function(t){return function(i){var s,n,a;for(a=0;null!=(n=i[a]);a++)try{s=e._data(n,"events"),s&&s.remove&&e(n).triggerHandler("remove")}catch(o){}t(i)}}(e.cleanData),e.widget=function(t,i,s){var n,a,o,r,h={},l=t.split(".")[0];return t=t.split(".")[1],n=l+"-"+t,s||(s=i,i=e.Widget),e.expr[":"][n.toLowerCase()]=function(t){return!!e.data(t,n)},e[l]=e[l]||{},a=e[l][t],o=e[l][t]=function(e,t){return this._createWidget?(arguments.length&&this._createWidget(e,t),void 0):new o(e,t)},e.extend(o,a,{version:s.version,_proto:e.extend({},s),_childConstructors:[]}),r=new i,r.options=e.widget.extend({},r.options),e.each(s,function(t,s){return e.isFunction(s)?(h[t]=function(){var e=function(){return i.prototype[t].apply(this,arguments)},n=function(e){return i.prototype[t].apply(this,e)};return function(){var t,i=this._super,a=this._superApply;return this._super=e,this._superApply=n,t=s.apply(this,arguments),this._super=i,this._superApply=a,t}}(),void 0):(h[t]=s,void 0)}),o.prototype=e.widget.extend(r,{widgetEventPrefix:a?r.widgetEventPrefix||t:t},h,{constructor:o,namespace:l,widgetName:t,widgetFullName:n}),a?(e.each(a._childConstructors,function(t,i){var s=i.prototype;e.widget(s.namespace+"."+s.widgetName,o,i._proto)}),delete a._childConstructors):i._childConstructors.push(o),e.widget.bridge(t,o),o},e.widget.extend=function(t){for(var i,s,n=a.call(arguments,1),o=0,r=n.length;r>o;o++)for(i in n[o])s=n[o][i],n[o].hasOwnProperty(i)&&void 0!==s&&(t[i]=e.isPlainObject(s)?e.isPlainObject(t[i])?e.widget.extend({},t[i],s):e.widget.extend({},s):s);return t},e.widget.bridge=function(t,i){var s=i.prototype.widgetFullName||t;e.fn[t]=function(n){var o="string"==typeof n,r=a.call(arguments,1),h=this;return o?this.each(function(){var i,a=e.data(this,s);return"instance"===n?(h=a,!1):a?e.isFunction(a[n])&&"_"!==n.charAt(0)?(i=a[n].apply(a,r),i!==a&&void 0!==i?(h=i&&i.jquery?h.pushStack(i.get()):i,!1):void 0):e.error("no such method '"+n+"' for "+t+" widget instance"):e.error("cannot call methods on "+t+" prior to initialization; "+"attempted to call method '"+n+"'")}):(r.length&&(n=e.widget.extend.apply(null,[n].concat(r))),this.each(function(){var t=e.data(this,s);t?(t.option(n||{}),t._init&&t._init()):e.data(this,s,new i(n,this))})),h}},e.Widget=function(){},e.Widget._childConstructors=[],e.Widget.prototype={widgetName:"widget",widgetEventPrefix:"",defaultElement:"<div>",options:{disabled:!1,create:null},_createWidget:function(t,i){i=e(i||this.defaultElement||this)[0],this.element=e(i),this.uuid=n++,this.eventNamespace="."+this.widgetName+this.uuid,this.bindings=e(),this.hoverable=e(),this.focusable=e(),i!==this&&(e.data(i,this.widgetFullName,this),this._on(!0,this.element,{remove:function(e){e.target===i&&this.destroy()}}),this.document=e(i.style?i.ownerDocument:i.document||i),this.window=e(this.document[0].defaultView||this.document[0].parentWindow)),this.options=e.widget.extend({},this.options,this._getCreateOptions(),t),this._create(),this._trigger("create",null,this._getCreateEventData()),this._init()},_getCreateOptions:e.noop,_getCreateEventData:e.noop,_create:e.noop,_init:e.noop,destroy:function(){this._destroy(),this.element.unbind(this.eventNamespace).removeData(this.widgetFullName).removeData(e.camelCase(this.widgetFullName)),this.widget().unbind(this.eventNamespace).removeAttr("aria-disabled").removeClass(this.widgetFullName+"-disabled "+"ui-state-disabled"),this.bindings.unbind(this.eventNamespace),this.hoverable.removeClass("ui-state-hover"),this.focusable.removeClass("ui-state-focus")},_destroy:e.noop,widget:function(){return this.element},option:function(t,i){var s,n,a,o=t;if(0===arguments.length)return e.widget.extend({},this.options);if("string"==typeof t)if(o={},s=t.split("."),t=s.shift(),s.length){for(n=o[t]=e.widget.extend({},this.options[t]),a=0;s.length-1>a;a++)n[s[a]]=n[s[a]]||{},n=n[s[a]];if(t=s.pop(),1===arguments.length)return void 0===n[t]?null:n[t];n[t]=i}else{if(1===arguments.length)return void 0===this.options[t]?null:this.options[t];o[t]=i}return this._setOptions(o),this},_setOptions:function(e){var t;for(t in e)this._setOption(t,e[t]);return this},_setOption:function(e,t){return this.options[e]=t,"disabled"===e&&(this.widget().toggleClass(this.widgetFullName+"-disabled",!!t),t&&(this.hoverable.removeClass("ui-state-hover"),this.focusable.removeClass("ui-state-focus"))),this},enable:function(){return this._setOptions({disabled:!1})},disable:function(){return this._setOptions({disabled:!0})},_on:function(t,i,s){var n,a=this;"boolean"!=typeof t&&(s=i,i=t,t=!1),s?(i=n=e(i),this.bindings=this.bindings.add(i)):(s=i,i=this.element,n=this.widget()),e.each(s,function(s,o){function r(){return t||a.options.disabled!==!0&&!e(this).hasClass("ui-state-disabled")?("string"==typeof o?a[o]:o).apply(a,arguments):void 0}"string"!=typeof o&&(r.guid=o.guid=o.guid||r.guid||e.guid++);var h=s.match(/^([\w:-]*)\s*(.*)$/),l=h[1]+a.eventNamespace,u=h[2];u?n.delegate(u,l,r):i.bind(l,r)})},_off:function(t,i){i=(i||"").split(" ").join(this.eventNamespace+" ")+this.eventNamespace,t.unbind(i).undelegate(i),this.bindings=e(this.bindings.not(t).get()),this.focusable=e(this.focusable.not(t).get()),this.hoverable=e(this.hoverable.not(t).get())},_delay:function(e,t){function i(){return("string"==typeof e?s[e]:e).apply(s,arguments)}var s=this;return setTimeout(i,t||0)},_hoverable:function(t){this.hoverable=this.hoverable.add(t),this._on(t,{mouseenter:function(t){e(t.currentTarget).addClass("ui-state-hover")},mouseleave:function(t){e(t.currentTarget).removeClass("ui-state-hover")}})},_focusable:function(t){this.focusable=this.focusable.add(t),this._on(t,{focusin:function(t){e(t.currentTarget).addClass("ui-state-focus")},focusout:function(t){e(t.currentTarget).removeClass("ui-state-focus")}})},_trigger:function(t,i,s){var n,a,o=this.options[t];if(s=s||{},i=e.Event(i),i.type=(t===this.widgetEventPrefix?t:this.widgetEventPrefix+t).toLowerCase(),i.target=this.element[0],a=i.originalEvent)for(n in a)n in i||(i[n]=a[n]);return this.element.trigger(i,s),!(e.isFunction(o)&&o.apply(this.element[0],[i].concat(s))===!1||i.isDefaultPrevented())}},e.each({show:"fadeIn",hide:"fadeOut"},function(t,i){e.Widget.prototype["_"+t]=function(s,n,a){"string"==typeof n&&(n={effect:n});var o,r=n?n===!0||"number"==typeof n?i:n.effect||i:t;n=n||{},"number"==typeof n&&(n={duration:n}),o=!e.isEmptyObject(n),n.complete=a,n.delay&&s.delay(n.delay),o&&e.effects&&e.effects.effect[r]?s[t](n):r!==t&&s[r]?s[r](n.duration,n.easing,a):s.queue(function(i){e(this)[t](),a&&a.call(s[0]),i()})}}),e.widget;var o=!1;e(document).mouseup(function(){o=!1}),e.widget("ui.mouse",{version:"1.11.4",options:{cancel:"input,textarea,button,select,option",distance:1,delay:0},_mouseInit:function(){var t=this;this.element.bind("mousedown."+this.widgetName,function(e){return t._mouseDown(e)}).bind("click."+this.widgetName,function(i){return!0===e.data(i.target,t.widgetName+".preventClickEvent")?(e.removeData(i.target,t.widgetName+".preventClickEvent"),i.stopImmediatePropagation(),!1):void 0}),this.started=!1},_mouseDestroy:function(){this.element.unbind("."+this.widgetName),this._mouseMoveDelegate&&this.document.unbind("mousemove."+this.widgetName,this._mouseMoveDelegate).unbind("mouseup."+this.widgetName,this._mouseUpDelegate)},_mouseDown:function(t){if(!o){this._mouseMoved=!1,this._mouseStarted&&this._mouseUp(t),this._mouseDownEvent=t;var i=this,s=1===t.which,n="string"==typeof this.options.cancel&&t.target.nodeName?e(t.target).closest(this.options.cancel).length:!1;return s&&!n&&this._mouseCapture(t)?(this.mouseDelayMet=!this.options.delay,this.mouseDelayMet||(this._mouseDelayTimer=setTimeout(function(){i.mouseDelayMet=!0},this.options.delay)),this._mouseDistanceMet(t)&&this._mouseDelayMet(t)&&(this._mouseStarted=this._mouseStart(t)!==!1,!this._mouseStarted)?(t.preventDefault(),!0):(!0===e.data(t.target,this.widgetName+".preventClickEvent")&&e.removeData(t.target,this.widgetName+".preventClickEvent"),this._mouseMoveDelegate=function(e){return i._mouseMove(e)},this._mouseUpDelegate=function(e){return i._mouseUp(e)},this.document.bind("mousemove."+this.widgetName,this._mouseMoveDelegate).bind("mouseup."+this.widgetName,this._mouseUpDelegate),t.preventDefault(),o=!0,!0)):!0}},_mouseMove:function(t){if(this._mouseMoved){if(e.ui.ie&&(!document.documentMode||9>document.documentMode)&&!t.button)return this._mouseUp(t);if(!t.which)return this._mouseUp(t)}return(t.which||t.button)&&(this._mouseMoved=!0),this._mouseStarted?(this._mouseDrag(t),t.preventDefault()):(this._mouseDistanceMet(t)&&this._mouseDelayMet(t)&&(this._mouseStarted=this._mouseStart(this._mouseDownEvent,t)!==!1,this._mouseStarted?this._mouseDrag(t):this._mouseUp(t)),!this._mouseStarted)},_mouseUp:function(t){return this.document.unbind("mousemove."+this.widgetName,this._mouseMoveDelegate).unbind("mouseup."+this.widgetName,this._mouseUpDelegate),this._mouseStarted&&(this._mouseStarted=!1,t.target===this._mouseDownEvent.target&&e.data(t.target,this.widgetName+".preventClickEvent",!0),this._mouseStop(t)),o=!1,!1},_mouseDistanceMet:function(e){return Math.max(Math.abs(this._mouseDownEvent.pageX-e.pageX),Math.abs(this._mouseDownEvent.pageY-e.pageY))>=this.options.distance},_mouseDelayMet:function(){return this.mouseDelayMet},_mouseStart:function(){},_mouseDrag:function(){},_mouseStop:function(){},_mouseCapture:function(){return!0}}),function(){function t(e,t,i){return[parseFloat(e[0])*(p.test(e[0])?t/100:1),parseFloat(e[1])*(p.test(e[1])?i/100:1)]}function i(t,i){return parseInt(e.css(t,i),10)||0}function s(t){var i=t[0];return 9===i.nodeType?{width:t.width(),height:t.height(),offset:{top:0,left:0}}:e.isWindow(i)?{width:t.width(),height:t.height(),offset:{top:t.scrollTop(),left:t.scrollLeft()}}:i.preventDefault?{width:0,height:0,offset:{top:i.pageY,left:i.pageX}}:{width:t.outerWidth(),height:t.outerHeight(),offset:t.offset()}}e.ui=e.ui||{};var n,a,o=Math.max,r=Math.abs,h=Math.round,l=/left|center|right/,u=/top|center|bottom/,d=/[\+\-]\d+(\.[\d]+)?%?/,c=/^\w+/,p=/%$/,f=e.fn.position;e.position={scrollbarWidth:function(){if(void 0!==n)return n;var t,i,s=e("<div style='display:block;position:absolute;width:50px;height:50px;overflow:hidden;'><div style='height:100px;width:auto;'></div></div>"),a=s.children()[0];return e("body").append(s),t=a.offsetWidth,s.css("overflow","scroll"),i=a.offsetWidth,t===i&&(i=s[0].clientWidth),s.remove(),n=t-i},getScrollInfo:function(t){var i=t.isWindow||t.isDocument?"":t.element.css("overflow-x"),s=t.isWindow||t.isDocument?"":t.element.css("overflow-y"),n="scroll"===i||"auto"===i&&t.width<t.element[0].scrollWidth,a="scroll"===s||"auto"===s&&t.height<t.element[0].scrollHeight;return{width:a?e.position.scrollbarWidth():0,height:n?e.position.scrollbarWidth():0}},getWithinInfo:function(t){var i=e(t||window),s=e.isWindow(i[0]),n=!!i[0]&&9===i[0].nodeType;return{element:i,isWindow:s,isDocument:n,offset:i.offset()||{left:0,top:0},scrollLeft:i.scrollLeft(),scrollTop:i.scrollTop(),width:s||n?i.width():i.outerWidth(),height:s||n?i.height():i.outerHeight()}}},e.fn.position=function(n){if(!n||!n.of)return f.apply(this,arguments);n=e.extend({},n);var p,m,g,v,y,b,_=e(n.of),x=e.position.getWithinInfo(n.within),w=e.position.getScrollInfo(x),k=(n.collision||"flip").split(" "),T={};return b=s(_),_[0].preventDefault&&(n.at="left top"),m=b.width,g=b.height,v=b.offset,y=e.extend({},v),e.each(["my","at"],function(){var e,t,i=(n[this]||"").split(" ");1===i.length&&(i=l.test(i[0])?i.concat(["center"]):u.test(i[0])?["center"].concat(i):["center","center"]),i[0]=l.test(i[0])?i[0]:"center",i[1]=u.test(i[1])?i[1]:"center",e=d.exec(i[0]),t=d.exec(i[1]),T[this]=[e?e[0]:0,t?t[0]:0],n[this]=[c.exec(i[0])[0],c.exec(i[1])[0]]}),1===k.length&&(k[1]=k[0]),"right"===n.at[0]?y.left+=m:"center"===n.at[0]&&(y.left+=m/2),"bottom"===n.at[1]?y.top+=g:"center"===n.at[1]&&(y.top+=g/2),p=t(T.at,m,g),y.left+=p[0],y.top+=p[1],this.each(function(){var s,l,u=e(this),d=u.outerWidth(),c=u.outerHeight(),f=i(this,"marginLeft"),b=i(this,"marginTop"),D=d+f+i(this,"marginRight")+w.width,S=c+b+i(this,"marginBottom")+w.height,N=e.extend({},y),M=t(T.my,u.outerWidth(),u.outerHeight());"right"===n.my[0]?N.left-=d:"center"===n.my[0]&&(N.left-=d/2),"bottom"===n.my[1]?N.top-=c:"center"===n.my[1]&&(N.top-=c/2),N.left+=M[0],N.top+=M[1],a||(N.left=h(N.left),N.top=h(N.top)),s={marginLeft:f,marginTop:b},e.each(["left","top"],function(t,i){e.ui.position[k[t]]&&e.ui.position[k[t]][i](N,{targetWidth:m,targetHeight:g,elemWidth:d,elemHeight:c,collisionPosition:s,collisionWidth:D,collisionHeight:S,offset:[p[0]+M[0],p[1]+M[1]],my:n.my,at:n.at,within:x,elem:u})}),n.using&&(l=function(e){var t=v.left-N.left,i=t+m-d,s=v.top-N.top,a=s+g-c,h={target:{element:_,left:v.left,top:v.top,width:m,height:g},element:{element:u,left:N.left,top:N.top,width:d,height:c},horizontal:0>i?"left":t>0?"right":"center",vertical:0>a?"top":s>0?"bottom":"middle"};d>m&&m>r(t+i)&&(h.horizontal="center"),c>g&&g>r(s+a)&&(h.vertical="middle"),h.important=o(r(t),r(i))>o(r(s),r(a))?"horizontal":"vertical",n.using.call(this,e,h)}),u.offset(e.extend(N,{using:l}))})},e.ui.position={fit:{left:function(e,t){var i,s=t.within,n=s.isWindow?s.scrollLeft:s.offset.left,a=s.width,r=e.left-t.collisionPosition.marginLeft,h=n-r,l=r+t.collisionWidth-a-n;t.collisionWidth>a?h>0&&0>=l?(i=e.left+h+t.collisionWidth-a-n,e.left+=h-i):e.left=l>0&&0>=h?n:h>l?n+a-t.collisionWidth:n:h>0?e.left+=h:l>0?e.left-=l:e.left=o(e.left-r,e.left)},top:function(e,t){var i,s=t.within,n=s.isWindow?s.scrollTop:s.offset.top,a=t.within.height,r=e.top-t.collisionPosition.marginTop,h=n-r,l=r+t.collisionHeight-a-n;t.collisionHeight>a?h>0&&0>=l?(i=e.top+h+t.collisionHeight-a-n,e.top+=h-i):e.top=l>0&&0>=h?n:h>l?n+a-t.collisionHeight:n:h>0?e.top+=h:l>0?e.top-=l:e.top=o(e.top-r,e.top)}},flip:{left:function(e,t){var i,s,n=t.within,a=n.offset.left+n.scrollLeft,o=n.width,h=n.isWindow?n.scrollLeft:n.offset.left,l=e.left-t.collisionPosition.marginLeft,u=l-h,d=l+t.collisionWidth-o-h,c="left"===t.my[0]?-t.elemWidth:"right"===t.my[0]?t.elemWidth:0,p="left"===t.at[0]?t.targetWidth:"right"===t.at[0]?-t.targetWidth:0,f=-2*t.offset[0];0>u?(i=e.left+c+p+f+t.collisionWidth-o-a,(0>i||r(u)>i)&&(e.left+=c+p+f)):d>0&&(s=e.left-t.collisionPosition.marginLeft+c+p+f-h,(s>0||d>r(s))&&(e.left+=c+p+f))},top:function(e,t){var i,s,n=t.within,a=n.offset.top+n.scrollTop,o=n.height,h=n.isWindow?n.scrollTop:n.offset.top,l=e.top-t.collisionPosition.marginTop,u=l-h,d=l+t.collisionHeight-o-h,c="top"===t.my[1],p=c?-t.elemHeight:"bottom"===t.my[1]?t.elemHeight:0,f="top"===t.at[1]?t.targetHeight:"bottom"===t.at[1]?-t.targetHeight:0,m=-2*t.offset[1];0>u?(s=e.top+p+f+m+t.collisionHeight-o-a,(0>s||r(u)>s)&&(e.top+=p+f+m)):d>0&&(i=e.top-t.collisionPosition.marginTop+p+f+m-h,(i>0||d>r(i))&&(e.top+=p+f+m))}},flipfit:{left:function(){e.ui.position.flip.left.apply(this,arguments),e.ui.position.fit.left.apply(this,arguments)},top:function(){e.ui.position.flip.top.apply(this,arguments),e.ui.position.fit.top.apply(this,arguments)}}},function(){var t,i,s,n,o,r=document.getElementsByTagName("body")[0],h=document.createElement("div");t=document.createElement(r?"div":"body"),s={visibility:"hidden",width:0,height:0,border:0,margin:0,background:"none"},r&&e.extend(s,{position:"absolute",left:"-1000px",top:"-1000px"});for(o in s)t.style[o]=s[o];t.appendChild(h),i=r||document.documentElement,i.insertBefore(t,i.firstChild),h.style.cssText="position: absolute; left: 10.7432222px;",n=e(h).offset().left,a=n>10&&11>n,t.innerHTML="",i.removeChild(t)}()}(),e.ui.position,e.widget("ui.draggable",e.ui.mouse,{version:"1.11.4",widgetEventPrefix:"drag",options:{addClasses:!0,appendTo:"parent",axis:!1,connectToSortable:!1,containment:!1,cursor:"auto",cursorAt:!1,grid:!1,handle:!1,helper:"original",iframeFix:!1,opacity:!1,refreshPositions:!1,revert:!1,revertDuration:500,scope:"default",scroll:!0,scrollSensitivity:20,scrollSpeed:20,snap:!1,snapMode:"both",snapTolerance:20,stack:!1,zIndex:!1,drag:null,start:null,stop:null},_create:function(){"original"===this.options.helper&&this._setPositionRelative(),this.options.addClasses&&this.element.addClass("ui-draggable"),this.options.disabled&&this.element.addClass("ui-draggable-disabled"),this._setHandleClassName(),this._mouseInit()},_setOption:function(e,t){this._super(e,t),"handle"===e&&(this._removeHandleClassName(),this._setHandleClassName())},_destroy:function(){return(this.helper||this.element).is(".ui-draggable-dragging")?(this.destroyOnClear=!0,void 0):(this.element.removeClass("ui-draggable ui-draggable-dragging ui-draggable-disabled"),this._removeHandleClassName(),this._mouseDestroy(),void 0)},_mouseCapture:function(t){var i=this.options;return this._blurActiveElement(t),this.helper||i.disabled||e(t.target).closest(".ui-resizable-handle").length>0?!1:(this.handle=this._getHandle(t),this.handle?(this._blockFrames(i.iframeFix===!0?"iframe":i.iframeFix),!0):!1)},_blockFrames:function(t){this.iframeBlocks=this.document.find(t).map(function(){var t=e(this);return e("<div>").css("position","absolute").appendTo(t.parent()).outerWidth(t.outerWidth()).outerHeight(t.outerHeight()).offset(t.offset())[0]})},_unblockFrames:function(){this.iframeBlocks&&(this.iframeBlocks.remove(),delete this.iframeBlocks)},_blurActiveElement:function(t){var i=this.document[0];if(this.handleElement.is(t.target))try{i.activeElement&&"body"!==i.activeElement.nodeName.toLowerCase()&&e(i.activeElement).blur()}catch(s){}},_mouseStart:function(t){var i=this.options;return this.helper=this._createHelper(t),this.helper.addClass("ui-draggable-dragging"),this._cacheHelperProportions(),e.ui.ddmanager&&(e.ui.ddmanager.current=this),this._cacheMargins(),this.cssPosition=this.helper.css("position"),this.scrollParent=this.helper.scrollParent(!0),this.offsetParent=this.helper.offsetParent(),this.hasFixedAncestor=this.helper.parents().filter(function(){return"fixed"===e(this).css("position")}).length>0,this.positionAbs=this.element.offset(),this._refreshOffsets(t),this.originalPosition=this.position=this._generatePosition(t,!1),this.originalPageX=t.pageX,this.originalPageY=t.pageY,i.cursorAt&&this._adjustOffsetFromHelper(i.cursorAt),this._setContainment(),this._trigger("start",t)===!1?(this._clear(),!1):(this._cacheHelperProportions(),e.ui.ddmanager&&!i.dropBehaviour&&e.ui.ddmanager.prepareOffsets(this,t),this._normalizeRightBottom(),this._mouseDrag(t,!0),e.ui.ddmanager&&e.ui.ddmanager.dragStart(this,t),!0)},_refreshOffsets:function(e){this.offset={top:this.positionAbs.top-this.margins.top,left:this.positionAbs.left-this.margins.left,scroll:!1,parent:this._getParentOffset(),relative:this._getRelativeOffset()},this.offset.click={left:e.pageX-this.offset.left,top:e.pageY-this.offset.top}},_mouseDrag:function(t,i){if(this.hasFixedAncestor&&(this.offset.parent=this._getParentOffset()),this.position=this._generatePosition(t,!0),this.positionAbs=this._convertPositionTo("absolute"),!i){var s=this._uiHash();if(this._trigger("drag",t,s)===!1)return this._mouseUp({}),!1;this.position=s.position}return this.helper[0].style.left=this.position.left+"px",this.helper[0].style.top=this.position.top+"px",e.ui.ddmanager&&e.ui.ddmanager.drag(this,t),!1},_mouseStop:function(t){var i=this,s=!1;return e.ui.ddmanager&&!this.options.dropBehaviour&&(s=e.ui.ddmanager.drop(this,t)),this.dropped&&(s=this.dropped,this.dropped=!1),"invalid"===this.options.revert&&!s||"valid"===this.options.revert&&s||this.options.revert===!0||e.isFunction(this.options.revert)&&this.options.revert.call(this.element,s)?e(this.helper).animate(this.originalPosition,parseInt(this.options.revertDuration,10),function(){i._trigger("stop",t)!==!1&&i._clear()}):this._trigger("stop",t)!==!1&&this._clear(),!1},_mouseUp:function(t){return this._unblockFrames(),e.ui.ddmanager&&e.ui.ddmanager.dragStop(this,t),this.handleElement.is(t.target)&&this.element.focus(),e.ui.mouse.prototype._mouseUp.call(this,t)},cancel:function(){return this.helper.is(".ui-draggable-dragging")?this._mouseUp({}):this._clear(),this},_getHandle:function(t){return this.options.handle?!!e(t.target).closest(this.element.find(this.options.handle)).length:!0},_setHandleClassName:function(){this.handleElement=this.options.handle?this.element.find(this.options.handle):this.element,this.handleElement.addClass("ui-draggable-handle")},_removeHandleClassName:function(){this.handleElement.removeClass("ui-draggable-handle")},_createHelper:function(t){var i=this.options,s=e.isFunction(i.helper),n=s?e(i.helper.apply(this.element[0],[t])):"clone"===i.helper?this.element.clone().removeAttr("id"):this.element;return n.parents("body").length||n.appendTo("parent"===i.appendTo?this.element[0].parentNode:i.appendTo),s&&n[0]===this.element[0]&&this._setPositionRelative(),n[0]===this.element[0]||/(fixed|absolute)/.test(n.css("position"))||n.css("position","absolute"),n},_setPositionRelative:function(){/^(?:r|a|f)/.test(this.element.css("position"))||(this.element[0].style.position="relative")},_adjustOffsetFromHelper:function(t){"string"==typeof t&&(t=t.split(" ")),e.isArray(t)&&(t={left:+t[0],top:+t[1]||0}),"left"in t&&(this.offset.click.left=t.left+this.margins.left),"right"in t&&(this.offset.click.left=this.helperProportions.width-t.right+this.margins.left),"top"in t&&(this.offset.click.top=t.top+this.margins.top),"bottom"in t&&(this.offset.click.top=this.helperProportions.height-t.bottom+this.margins.top)},_isRootNode:function(e){return/(html|body)/i.test(e.tagName)||e===this.document[0]},_getParentOffset:function(){var t=this.offsetParent.offset(),i=this.document[0];return"absolute"===this.cssPosition&&this.scrollParent[0]!==i&&e.contains(this.scrollParent[0],this.offsetParent[0])&&(t.left+=this.scrollParent.scrollLeft(),t.top+=this.scrollParent.scrollTop()),this._isRootNode(this.offsetParent[0])&&(t={top:0,left:0}),{top:t.top+(parseInt(this.offsetParent.css("borderTopWidth"),10)||0),left:t.left+(parseInt(this.offsetParent.css("borderLeftWidth"),10)||0)}},_getRelativeOffset:function(){if("relative"!==this.cssPosition)return{top:0,left:0};var e=this.element.position(),t=this._isRootNode(this.scrollParent[0]);return{top:e.top-(parseInt(this.helper.css("top"),10)||0)+(t?0:this.scrollParent.scrollTop()),left:e.left-(parseInt(this.helper.css("left"),10)||0)+(t?0:this.scrollParent.scrollLeft())}},_cacheMargins:function(){this.margins={left:parseInt(this.element.css("marginLeft"),10)||0,top:parseInt(this.element.css("marginTop"),10)||0,right:parseInt(this.element.css("marginRight"),10)||0,bottom:parseInt(this.element.css("marginBottom"),10)||0}},_cacheHelperProportions:function(){this.helperProportions={width:this.helper.outerWidth(),height:this.helper.outerHeight()}},_setContainment:function(){var t,i,s,n=this.options,a=this.document[0];return this.relativeContainer=null,n.containment?"window"===n.containment?(this.containment=[e(window).scrollLeft()-this.offset.relative.left-this.offset.parent.left,e(window).scrollTop()-this.offset.relative.top-this.offset.parent.top,e(window).scrollLeft()+e(window).width()-this.helperProportions.width-this.margins.left,e(window).scrollTop()+(e(window).height()||a.body.parentNode.scrollHeight)-this.helperProportions.height-this.margins.top],void 0):"document"===n.containment?(this.containment=[0,0,e(a).width()-this.helperProportions.width-this.margins.left,(e(a).height()||a.body.parentNode.scrollHeight)-this.helperProportions.height-this.margins.top],void 0):n.containment.constructor===Array?(this.containment=n.containment,void 0):("parent"===n.containment&&(n.containment=this.helper[0].parentNode),i=e(n.containment),s=i[0],s&&(t=/(scroll|auto)/.test(i.css("overflow")),this.containment=[(parseInt(i.css("borderLeftWidth"),10)||0)+(parseInt(i.css("paddingLeft"),10)||0),(parseInt(i.css("borderTopWidth"),10)||0)+(parseInt(i.css("paddingTop"),10)||0),(t?Math.max(s.scrollWidth,s.offsetWidth):s.offsetWidth)-(parseInt(i.css("borderRightWidth"),10)||0)-(parseInt(i.css("paddingRight"),10)||0)-this.helperProportions.width-this.margins.left-this.margins.right,(t?Math.max(s.scrollHeight,s.offsetHeight):s.offsetHeight)-(parseInt(i.css("borderBottomWidth"),10)||0)-(parseInt(i.css("paddingBottom"),10)||0)-this.helperProportions.height-this.margins.top-this.margins.bottom],this.relativeContainer=i),void 0):(this.containment=null,void 0)},_convertPositionTo:function(e,t){t||(t=this.position);var i="absolute"===e?1:-1,s=this._isRootNode(this.scrollParent[0]);return{top:t.top+this.offset.relative.top*i+this.offset.parent.top*i-("fixed"===this.cssPosition?-this.offset.scroll.top:s?0:this.offset.scroll.top)*i,left:t.left+this.offset.relative.left*i+this.offset.parent.left*i-("fixed"===this.cssPosition?-this.offset.scroll.left:s?0:this.offset.scroll.left)*i}},_generatePosition:function(e,t){var i,s,n,a,o=this.options,r=this._isRootNode(this.scrollParent[0]),h=e.pageX,l=e.pageY;return r&&this.offset.scroll||(this.offset.scroll={top:this.scrollParent.scrollTop(),left:this.scrollParent.scrollLeft()}),t&&(this.containment&&(this.relativeContainer?(s=this.relativeContainer.offset(),i=[this.containment[0]+s.left,this.containment[1]+s.top,this.containment[2]+s.left,this.containment[3]+s.top]):i=this.containment,e.pageX-this.offset.click.left<i[0]&&(h=i[0]+this.offset.click.left),e.pageY-this.offset.click.top<i[1]&&(l=i[1]+this.offset.click.top),e.pageX-this.offset.click.left>i[2]&&(h=i[2]+this.offset.click.left),e.pageY-this.offset.click.top>i[3]&&(l=i[3]+this.offset.click.top)),o.grid&&(n=o.grid[1]?this.originalPageY+Math.round((l-this.originalPageY)/o.grid[1])*o.grid[1]:this.originalPageY,l=i?n-this.offset.click.top>=i[1]||n-this.offset.click.top>i[3]?n:n-this.offset.click.top>=i[1]?n-o.grid[1]:n+o.grid[1]:n,a=o.grid[0]?this.originalPageX+Math.round((h-this.originalPageX)/o.grid[0])*o.grid[0]:this.originalPageX,h=i?a-this.offset.click.left>=i[0]||a-this.offset.click.left>i[2]?a:a-this.offset.click.left>=i[0]?a-o.grid[0]:a+o.grid[0]:a),"y"===o.axis&&(h=this.originalPageX),"x"===o.axis&&(l=this.originalPageY)),{top:l-this.offset.click.top-this.offset.relative.top-this.offset.parent.top+("fixed"===this.cssPosition?-this.offset.scroll.top:r?0:this.offset.scroll.top),left:h-this.offset.click.left-this.offset.relative.left-this.offset.parent.left+("fixed"===this.cssPosition?-this.offset.scroll.left:r?0:this.offset.scroll.left)}},_clear:function(){this.helper.removeClass("ui-draggable-dragging"),this.helper[0]===this.element[0]||this.cancelHelperRemoval||this.helper.remove(),this.helper=null,this.cancelHelperRemoval=!1,this.destroyOnClear&&this.destroy()},_normalizeRightBottom:function(){"y"!==this.options.axis&&"auto"!==this.helper.css("right")&&(this.helper.width(this.helper.width()),this.helper.css("right","auto")),"x"!==this.options.axis&&"auto"!==this.helper.css("bottom")&&(this.helper.height(this.helper.height()),this.helper.css("bottom","auto"))},_trigger:function(t,i,s){return s=s||this._uiHash(),e.ui.plugin.call(this,t,[i,s,this],!0),/^(drag|start|stop)/.test(t)&&(this.positionAbs=this._convertPositionTo("absolute"),s.offset=this.positionAbs),e.Widget.prototype._trigger.call(this,t,i,s)},plugins:{},_uiHash:function(){return{helper:this.helper,position:this.position,originalPosition:this.originalPosition,offset:this.positionAbs}}}),e.ui.plugin.add("draggable","connectToSortable",{start:function(t,i,s){var n=e.extend({},i,{item:s.element});s.sortables=[],e(s.options.connectToSortable).each(function(){var i=e(this).sortable("instance");i&&!i.options.disabled&&(s.sortables.push(i),i.refreshPositions(),i._trigger("activate",t,n))})},stop:function(t,i,s){var n=e.extend({},i,{item:s.element});s.cancelHelperRemoval=!1,e.each(s.sortables,function(){var e=this;e.isOver?(e.isOver=0,s.cancelHelperRemoval=!0,e.cancelHelperRemoval=!1,e._storedCSS={position:e.placeholder.css("position"),top:e.placeholder.css("top"),left:e.placeholder.css("left")},e._mouseStop(t),e.options.helper=e.options._helper):(e.cancelHelperRemoval=!0,e._trigger("deactivate",t,n))})},drag:function(t,i,s){e.each(s.sortables,function(){var n=!1,a=this;a.positionAbs=s.positionAbs,a.helperProportions=s.helperProportions,a.offset.click=s.offset.click,a._intersectsWith(a.containerCache)&&(n=!0,e.each(s.sortables,function(){return this.positionAbs=s.positionAbs,this.helperProportions=s.helperProportions,this.offset.click=s.offset.click,this!==a&&this._intersectsWith(this.containerCache)&&e.contains(a.element[0],this.element[0])&&(n=!1),n
})),n?(a.isOver||(a.isOver=1,s._parent=i.helper.parent(),a.currentItem=i.helper.appendTo(a.element).data("ui-sortable-item",!0),a.options._helper=a.options.helper,a.options.helper=function(){return i.helper[0]},t.target=a.currentItem[0],a._mouseCapture(t,!0),a._mouseStart(t,!0,!0),a.offset.click.top=s.offset.click.top,a.offset.click.left=s.offset.click.left,a.offset.parent.left-=s.offset.parent.left-a.offset.parent.left,a.offset.parent.top-=s.offset.parent.top-a.offset.parent.top,s._trigger("toSortable",t),s.dropped=a.element,e.each(s.sortables,function(){this.refreshPositions()}),s.currentItem=s.element,a.fromOutside=s),a.currentItem&&(a._mouseDrag(t),i.position=a.position)):a.isOver&&(a.isOver=0,a.cancelHelperRemoval=!0,a.options._revert=a.options.revert,a.options.revert=!1,a._trigger("out",t,a._uiHash(a)),a._mouseStop(t,!0),a.options.revert=a.options._revert,a.options.helper=a.options._helper,a.placeholder&&a.placeholder.remove(),i.helper.appendTo(s._parent),s._refreshOffsets(t),i.position=s._generatePosition(t,!0),s._trigger("fromSortable",t),s.dropped=!1,e.each(s.sortables,function(){this.refreshPositions()}))})}}),e.ui.plugin.add("draggable","cursor",{start:function(t,i,s){var n=e("body"),a=s.options;n.css("cursor")&&(a._cursor=n.css("cursor")),n.css("cursor",a.cursor)},stop:function(t,i,s){var n=s.options;n._cursor&&e("body").css("cursor",n._cursor)}}),e.ui.plugin.add("draggable","opacity",{start:function(t,i,s){var n=e(i.helper),a=s.options;n.css("opacity")&&(a._opacity=n.css("opacity")),n.css("opacity",a.opacity)},stop:function(t,i,s){var n=s.options;n._opacity&&e(i.helper).css("opacity",n._opacity)}}),e.ui.plugin.add("draggable","scroll",{start:function(e,t,i){i.scrollParentNotHidden||(i.scrollParentNotHidden=i.helper.scrollParent(!1)),i.scrollParentNotHidden[0]!==i.document[0]&&"HTML"!==i.scrollParentNotHidden[0].tagName&&(i.overflowOffset=i.scrollParentNotHidden.offset())},drag:function(t,i,s){var n=s.options,a=!1,o=s.scrollParentNotHidden[0],r=s.document[0];o!==r&&"HTML"!==o.tagName?(n.axis&&"x"===n.axis||(s.overflowOffset.top+o.offsetHeight-t.pageY<n.scrollSensitivity?o.scrollTop=a=o.scrollTop+n.scrollSpeed:t.pageY-s.overflowOffset.top<n.scrollSensitivity&&(o.scrollTop=a=o.scrollTop-n.scrollSpeed)),n.axis&&"y"===n.axis||(s.overflowOffset.left+o.offsetWidth-t.pageX<n.scrollSensitivity?o.scrollLeft=a=o.scrollLeft+n.scrollSpeed:t.pageX-s.overflowOffset.left<n.scrollSensitivity&&(o.scrollLeft=a=o.scrollLeft-n.scrollSpeed))):(n.axis&&"x"===n.axis||(t.pageY-e(r).scrollTop()<n.scrollSensitivity?a=e(r).scrollTop(e(r).scrollTop()-n.scrollSpeed):e(window).height()-(t.pageY-e(r).scrollTop())<n.scrollSensitivity&&(a=e(r).scrollTop(e(r).scrollTop()+n.scrollSpeed))),n.axis&&"y"===n.axis||(t.pageX-e(r).scrollLeft()<n.scrollSensitivity?a=e(r).scrollLeft(e(r).scrollLeft()-n.scrollSpeed):e(window).width()-(t.pageX-e(r).scrollLeft())<n.scrollSensitivity&&(a=e(r).scrollLeft(e(r).scrollLeft()+n.scrollSpeed)))),a!==!1&&e.ui.ddmanager&&!n.dropBehaviour&&e.ui.ddmanager.prepareOffsets(s,t)}}),e.ui.plugin.add("draggable","snap",{start:function(t,i,s){var n=s.options;s.snapElements=[],e(n.snap.constructor!==String?n.snap.items||":data(ui-draggable)":n.snap).each(function(){var t=e(this),i=t.offset();this!==s.element[0]&&s.snapElements.push({item:this,width:t.outerWidth(),height:t.outerHeight(),top:i.top,left:i.left})})},drag:function(t,i,s){var n,a,o,r,h,l,u,d,c,p,f=s.options,m=f.snapTolerance,g=i.offset.left,v=g+s.helperProportions.width,y=i.offset.top,b=y+s.helperProportions.height;for(c=s.snapElements.length-1;c>=0;c--)h=s.snapElements[c].left-s.margins.left,l=h+s.snapElements[c].width,u=s.snapElements[c].top-s.margins.top,d=u+s.snapElements[c].height,h-m>v||g>l+m||u-m>b||y>d+m||!e.contains(s.snapElements[c].item.ownerDocument,s.snapElements[c].item)?(s.snapElements[c].snapping&&s.options.snap.release&&s.options.snap.release.call(s.element,t,e.extend(s._uiHash(),{snapItem:s.snapElements[c].item})),s.snapElements[c].snapping=!1):("inner"!==f.snapMode&&(n=m>=Math.abs(u-b),a=m>=Math.abs(d-y),o=m>=Math.abs(h-v),r=m>=Math.abs(l-g),n&&(i.position.top=s._convertPositionTo("relative",{top:u-s.helperProportions.height,left:0}).top),a&&(i.position.top=s._convertPositionTo("relative",{top:d,left:0}).top),o&&(i.position.left=s._convertPositionTo("relative",{top:0,left:h-s.helperProportions.width}).left),r&&(i.position.left=s._convertPositionTo("relative",{top:0,left:l}).left)),p=n||a||o||r,"outer"!==f.snapMode&&(n=m>=Math.abs(u-y),a=m>=Math.abs(d-b),o=m>=Math.abs(h-g),r=m>=Math.abs(l-v),n&&(i.position.top=s._convertPositionTo("relative",{top:u,left:0}).top),a&&(i.position.top=s._convertPositionTo("relative",{top:d-s.helperProportions.height,left:0}).top),o&&(i.position.left=s._convertPositionTo("relative",{top:0,left:h}).left),r&&(i.position.left=s._convertPositionTo("relative",{top:0,left:l-s.helperProportions.width}).left)),!s.snapElements[c].snapping&&(n||a||o||r||p)&&s.options.snap.snap&&s.options.snap.snap.call(s.element,t,e.extend(s._uiHash(),{snapItem:s.snapElements[c].item})),s.snapElements[c].snapping=n||a||o||r||p)}}),e.ui.plugin.add("draggable","stack",{start:function(t,i,s){var n,a=s.options,o=e.makeArray(e(a.stack)).sort(function(t,i){return(parseInt(e(t).css("zIndex"),10)||0)-(parseInt(e(i).css("zIndex"),10)||0)});o.length&&(n=parseInt(e(o[0]).css("zIndex"),10)||0,e(o).each(function(t){e(this).css("zIndex",n+t)}),this.css("zIndex",n+o.length))}}),e.ui.plugin.add("draggable","zIndex",{start:function(t,i,s){var n=e(i.helper),a=s.options;n.css("zIndex")&&(a._zIndex=n.css("zIndex")),n.css("zIndex",a.zIndex)},stop:function(t,i,s){var n=s.options;n._zIndex&&e(i.helper).css("zIndex",n._zIndex)}}),e.ui.draggable,e.widget("ui.droppable",{version:"1.11.4",widgetEventPrefix:"drop",options:{accept:"*",activeClass:!1,addClasses:!0,greedy:!1,hoverClass:!1,scope:"default",tolerance:"intersect",activate:null,deactivate:null,drop:null,out:null,over:null},_create:function(){var t,i=this.options,s=i.accept;this.isover=!1,this.isout=!0,this.accept=e.isFunction(s)?s:function(e){return e.is(s)},this.proportions=function(){return arguments.length?(t=arguments[0],void 0):t?t:t={width:this.element[0].offsetWidth,height:this.element[0].offsetHeight}},this._addToManager(i.scope),i.addClasses&&this.element.addClass("ui-droppable")},_addToManager:function(t){e.ui.ddmanager.droppables[t]=e.ui.ddmanager.droppables[t]||[],e.ui.ddmanager.droppables[t].push(this)},_splice:function(e){for(var t=0;e.length>t;t++)e[t]===this&&e.splice(t,1)},_destroy:function(){var t=e.ui.ddmanager.droppables[this.options.scope];this._splice(t),this.element.removeClass("ui-droppable ui-droppable-disabled")},_setOption:function(t,i){if("accept"===t)this.accept=e.isFunction(i)?i:function(e){return e.is(i)};else if("scope"===t){var s=e.ui.ddmanager.droppables[this.options.scope];this._splice(s),this._addToManager(i)}this._super(t,i)},_activate:function(t){var i=e.ui.ddmanager.current;this.options.activeClass&&this.element.addClass(this.options.activeClass),i&&this._trigger("activate",t,this.ui(i))},_deactivate:function(t){var i=e.ui.ddmanager.current;this.options.activeClass&&this.element.removeClass(this.options.activeClass),i&&this._trigger("deactivate",t,this.ui(i))},_over:function(t){var i=e.ui.ddmanager.current;i&&(i.currentItem||i.element)[0]!==this.element[0]&&this.accept.call(this.element[0],i.currentItem||i.element)&&(this.options.hoverClass&&this.element.addClass(this.options.hoverClass),this._trigger("over",t,this.ui(i)))},_out:function(t){var i=e.ui.ddmanager.current;i&&(i.currentItem||i.element)[0]!==this.element[0]&&this.accept.call(this.element[0],i.currentItem||i.element)&&(this.options.hoverClass&&this.element.removeClass(this.options.hoverClass),this._trigger("out",t,this.ui(i)))},_drop:function(t,i){var s=i||e.ui.ddmanager.current,n=!1;return s&&(s.currentItem||s.element)[0]!==this.element[0]?(this.element.find(":data(ui-droppable)").not(".ui-draggable-dragging").each(function(){var i=e(this).droppable("instance");return i.options.greedy&&!i.options.disabled&&i.options.scope===s.options.scope&&i.accept.call(i.element[0],s.currentItem||s.element)&&e.ui.intersect(s,e.extend(i,{offset:i.element.offset()}),i.options.tolerance,t)?(n=!0,!1):void 0}),n?!1:this.accept.call(this.element[0],s.currentItem||s.element)?(this.options.activeClass&&this.element.removeClass(this.options.activeClass),this.options.hoverClass&&this.element.removeClass(this.options.hoverClass),this._trigger("drop",t,this.ui(s)),this.element):!1):!1},ui:function(e){return{draggable:e.currentItem||e.element,helper:e.helper,position:e.position,offset:e.positionAbs}}}),e.ui.intersect=function(){function e(e,t,i){return e>=t&&t+i>e}return function(t,i,s,n){if(!i.offset)return!1;var a=(t.positionAbs||t.position.absolute).left+t.margins.left,o=(t.positionAbs||t.position.absolute).top+t.margins.top,r=a+t.helperProportions.width,h=o+t.helperProportions.height,l=i.offset.left,u=i.offset.top,d=l+i.proportions().width,c=u+i.proportions().height;switch(s){case"fit":return a>=l&&d>=r&&o>=u&&c>=h;case"intersect":return a+t.helperProportions.width/2>l&&d>r-t.helperProportions.width/2&&o+t.helperProportions.height/2>u&&c>h-t.helperProportions.height/2;case"pointer":return e(n.pageY,u,i.proportions().height)&&e(n.pageX,l,i.proportions().width);case"touch":return(o>=u&&c>=o||h>=u&&c>=h||u>o&&h>c)&&(a>=l&&d>=a||r>=l&&d>=r||l>a&&r>d);default:return!1}}}(),e.ui.ddmanager={current:null,droppables:{"default":[]},prepareOffsets:function(t,i){var s,n,a=e.ui.ddmanager.droppables[t.options.scope]||[],o=i?i.type:null,r=(t.currentItem||t.element).find(":data(ui-droppable)").addBack();e:for(s=0;a.length>s;s++)if(!(a[s].options.disabled||t&&!a[s].accept.call(a[s].element[0],t.currentItem||t.element))){for(n=0;r.length>n;n++)if(r[n]===a[s].element[0]){a[s].proportions().height=0;continue e}a[s].visible="none"!==a[s].element.css("display"),a[s].visible&&("mousedown"===o&&a[s]._activate.call(a[s],i),a[s].offset=a[s].element.offset(),a[s].proportions({width:a[s].element[0].offsetWidth,height:a[s].element[0].offsetHeight}))}},drop:function(t,i){var s=!1;return e.each((e.ui.ddmanager.droppables[t.options.scope]||[]).slice(),function(){this.options&&(!this.options.disabled&&this.visible&&e.ui.intersect(t,this,this.options.tolerance,i)&&(s=this._drop.call(this,i)||s),!this.options.disabled&&this.visible&&this.accept.call(this.element[0],t.currentItem||t.element)&&(this.isout=!0,this.isover=!1,this._deactivate.call(this,i)))}),s},dragStart:function(t,i){t.element.parentsUntil("body").bind("scroll.droppable",function(){t.options.refreshPositions||e.ui.ddmanager.prepareOffsets(t,i)})},drag:function(t,i){t.options.refreshPositions&&e.ui.ddmanager.prepareOffsets(t,i),e.each(e.ui.ddmanager.droppables[t.options.scope]||[],function(){if(!this.options.disabled&&!this.greedyChild&&this.visible){var s,n,a,o=e.ui.intersect(t,this,this.options.tolerance,i),r=!o&&this.isover?"isout":o&&!this.isover?"isover":null;r&&(this.options.greedy&&(n=this.options.scope,a=this.element.parents(":data(ui-droppable)").filter(function(){return e(this).droppable("instance").options.scope===n}),a.length&&(s=e(a[0]).droppable("instance"),s.greedyChild="isover"===r)),s&&"isover"===r&&(s.isover=!1,s.isout=!0,s._out.call(s,i)),this[r]=!0,this["isout"===r?"isover":"isout"]=!1,this["isover"===r?"_over":"_out"].call(this,i),s&&"isout"===r&&(s.isout=!1,s.isover=!0,s._over.call(s,i)))}})},dragStop:function(t,i){t.element.parentsUntil("body").unbind("scroll.droppable"),t.options.refreshPositions||e.ui.ddmanager.prepareOffsets(t,i)}},e.ui.droppable,e.widget("ui.resizable",e.ui.mouse,{version:"1.11.4",widgetEventPrefix:"resize",options:{alsoResize:!1,animate:!1,animateDuration:"slow",animateEasing:"swing",aspectRatio:!1,autoHide:!1,containment:!1,ghost:!1,grid:!1,handles:"e,s,se",helper:!1,maxHeight:null,maxWidth:null,minHeight:10,minWidth:10,zIndex:90,resize:null,start:null,stop:null},_num:function(e){return parseInt(e,10)||0},_isNumber:function(e){return!isNaN(parseInt(e,10))},_hasScroll:function(t,i){if("hidden"===e(t).css("overflow"))return!1;var s=i&&"left"===i?"scrollLeft":"scrollTop",n=!1;return t[s]>0?!0:(t[s]=1,n=t[s]>0,t[s]=0,n)},_create:function(){var t,i,s,n,a,o=this,r=this.options;if(this.element.addClass("ui-resizable"),e.extend(this,{_aspectRatio:!!r.aspectRatio,aspectRatio:r.aspectRatio,originalElement:this.element,_proportionallyResizeElements:[],_helper:r.helper||r.ghost||r.animate?r.helper||"ui-resizable-helper":null}),this.element[0].nodeName.match(/^(canvas|textarea|input|select|button|img)$/i)&&(this.element.wrap(e("<div class='ui-wrapper' style='overflow: hidden;'></div>").css({position:this.element.css("position"),width:this.element.outerWidth(),height:this.element.outerHeight(),top:this.element.css("top"),left:this.element.css("left")})),this.element=this.element.parent().data("ui-resizable",this.element.resizable("instance")),this.elementIsWrapper=!0,this.element.css({marginLeft:this.originalElement.css("marginLeft"),marginTop:this.originalElement.css("marginTop"),marginRight:this.originalElement.css("marginRight"),marginBottom:this.originalElement.css("marginBottom")}),this.originalElement.css({marginLeft:0,marginTop:0,marginRight:0,marginBottom:0}),this.originalResizeStyle=this.originalElement.css("resize"),this.originalElement.css("resize","none"),this._proportionallyResizeElements.push(this.originalElement.css({position:"static",zoom:1,display:"block"})),this.originalElement.css({margin:this.originalElement.css("margin")}),this._proportionallyResize()),this.handles=r.handles||(e(".ui-resizable-handle",this.element).length?{n:".ui-resizable-n",e:".ui-resizable-e",s:".ui-resizable-s",w:".ui-resizable-w",se:".ui-resizable-se",sw:".ui-resizable-sw",ne:".ui-resizable-ne",nw:".ui-resizable-nw"}:"e,s,se"),this._handles=e(),this.handles.constructor===String)for("all"===this.handles&&(this.handles="n,e,s,w,se,sw,ne,nw"),t=this.handles.split(","),this.handles={},i=0;t.length>i;i++)s=e.trim(t[i]),a="ui-resizable-"+s,n=e("<div class='ui-resizable-handle "+a+"'></div>"),n.css({zIndex:r.zIndex}),"se"===s&&n.addClass("ui-icon ui-icon-gripsmall-diagonal-se"),this.handles[s]=".ui-resizable-"+s,this.element.append(n);this._renderAxis=function(t){var i,s,n,a;t=t||this.element;for(i in this.handles)this.handles[i].constructor===String?this.handles[i]=this.element.children(this.handles[i]).first().show():(this.handles[i].jquery||this.handles[i].nodeType)&&(this.handles[i]=e(this.handles[i]),this._on(this.handles[i],{mousedown:o._mouseDown})),this.elementIsWrapper&&this.originalElement[0].nodeName.match(/^(textarea|input|select|button)$/i)&&(s=e(this.handles[i],this.element),a=/sw|ne|nw|se|n|s/.test(i)?s.outerHeight():s.outerWidth(),n=["padding",/ne|nw|n/.test(i)?"Top":/se|sw|s/.test(i)?"Bottom":/^e$/.test(i)?"Right":"Left"].join(""),t.css(n,a),this._proportionallyResize()),this._handles=this._handles.add(this.handles[i])},this._renderAxis(this.element),this._handles=this._handles.add(this.element.find(".ui-resizable-handle")),this._handles.disableSelection(),this._handles.mouseover(function(){o.resizing||(this.className&&(n=this.className.match(/ui-resizable-(se|sw|ne|nw|n|e|s|w)/i)),o.axis=n&&n[1]?n[1]:"se")}),r.autoHide&&(this._handles.hide(),e(this.element).addClass("ui-resizable-autohide").mouseenter(function(){r.disabled||(e(this).removeClass("ui-resizable-autohide"),o._handles.show())}).mouseleave(function(){r.disabled||o.resizing||(e(this).addClass("ui-resizable-autohide"),o._handles.hide())})),this._mouseInit()},_destroy:function(){this._mouseDestroy();var t,i=function(t){e(t).removeClass("ui-resizable ui-resizable-disabled ui-resizable-resizing").removeData("resizable").removeData("ui-resizable").unbind(".resizable").find(".ui-resizable-handle").remove()};return this.elementIsWrapper&&(i(this.element),t=this.element,this.originalElement.css({position:t.css("position"),width:t.outerWidth(),height:t.outerHeight(),top:t.css("top"),left:t.css("left")}).insertAfter(t),t.remove()),this.originalElement.css("resize",this.originalResizeStyle),i(this.originalElement),this},_mouseCapture:function(t){var i,s,n=!1;for(i in this.handles)s=e(this.handles[i])[0],(s===t.target||e.contains(s,t.target))&&(n=!0);return!this.options.disabled&&n},_mouseStart:function(t){var i,s,n,a=this.options,o=this.element;return this.resizing=!0,this._renderProxy(),i=this._num(this.helper.css("left")),s=this._num(this.helper.css("top")),a.containment&&(i+=e(a.containment).scrollLeft()||0,s+=e(a.containment).scrollTop()||0),this.offset=this.helper.offset(),this.position={left:i,top:s},this.size=this._helper?{width:this.helper.width(),height:this.helper.height()}:{width:o.width(),height:o.height()},this.originalSize=this._helper?{width:o.outerWidth(),height:o.outerHeight()}:{width:o.width(),height:o.height()},this.sizeDiff={width:o.outerWidth()-o.width(),height:o.outerHeight()-o.height()},this.originalPosition={left:i,top:s},this.originalMousePosition={left:t.pageX,top:t.pageY},this.aspectRatio="number"==typeof a.aspectRatio?a.aspectRatio:this.originalSize.width/this.originalSize.height||1,n=e(".ui-resizable-"+this.axis).css("cursor"),e("body").css("cursor","auto"===n?this.axis+"-resize":n),o.addClass("ui-resizable-resizing"),this._propagate("start",t),!0},_mouseDrag:function(t){var i,s,n=this.originalMousePosition,a=this.axis,o=t.pageX-n.left||0,r=t.pageY-n.top||0,h=this._change[a];return this._updatePrevProperties(),h?(i=h.apply(this,[t,o,r]),this._updateVirtualBoundaries(t.shiftKey),(this._aspectRatio||t.shiftKey)&&(i=this._updateRatio(i,t)),i=this._respectSize(i,t),this._updateCache(i),this._propagate("resize",t),s=this._applyChanges(),!this._helper&&this._proportionallyResizeElements.length&&this._proportionallyResize(),e.isEmptyObject(s)||(this._updatePrevProperties(),this._trigger("resize",t,this.ui()),this._applyChanges()),!1):!1},_mouseStop:function(t){this.resizing=!1;var i,s,n,a,o,r,h,l=this.options,u=this;return this._helper&&(i=this._proportionallyResizeElements,s=i.length&&/textarea/i.test(i[0].nodeName),n=s&&this._hasScroll(i[0],"left")?0:u.sizeDiff.height,a=s?0:u.sizeDiff.width,o={width:u.helper.width()-a,height:u.helper.height()-n},r=parseInt(u.element.css("left"),10)+(u.position.left-u.originalPosition.left)||null,h=parseInt(u.element.css("top"),10)+(u.position.top-u.originalPosition.top)||null,l.animate||this.element.css(e.extend(o,{top:h,left:r})),u.helper.height(u.size.height),u.helper.width(u.size.width),this._helper&&!l.animate&&this._proportionallyResize()),e("body").css("cursor","auto"),this.element.removeClass("ui-resizable-resizing"),this._propagate("stop",t),this._helper&&this.helper.remove(),!1},_updatePrevProperties:function(){this.prevPosition={top:this.position.top,left:this.position.left},this.prevSize={width:this.size.width,height:this.size.height}},_applyChanges:function(){var e={};return this.position.top!==this.prevPosition.top&&(e.top=this.position.top+"px"),this.position.left!==this.prevPosition.left&&(e.left=this.position.left+"px"),this.size.width!==this.prevSize.width&&(e.width=this.size.width+"px"),this.size.height!==this.prevSize.height&&(e.height=this.size.height+"px"),this.helper.css(e),e},_updateVirtualBoundaries:function(e){var t,i,s,n,a,o=this.options;a={minWidth:this._isNumber(o.minWidth)?o.minWidth:0,maxWidth:this._isNumber(o.maxWidth)?o.maxWidth:1/0,minHeight:this._isNumber(o.minHeight)?o.minHeight:0,maxHeight:this._isNumber(o.maxHeight)?o.maxHeight:1/0},(this._aspectRatio||e)&&(t=a.minHeight*this.aspectRatio,s=a.minWidth/this.aspectRatio,i=a.maxHeight*this.aspectRatio,n=a.maxWidth/this.aspectRatio,t>a.minWidth&&(a.minWidth=t),s>a.minHeight&&(a.minHeight=s),a.maxWidth>i&&(a.maxWidth=i),a.maxHeight>n&&(a.maxHeight=n)),this._vBoundaries=a},_updateCache:function(e){this.offset=this.helper.offset(),this._isNumber(e.left)&&(this.position.left=e.left),this._isNumber(e.top)&&(this.position.top=e.top),this._isNumber(e.height)&&(this.size.height=e.height),this._isNumber(e.width)&&(this.size.width=e.width)},_updateRatio:function(e){var t=this.position,i=this.size,s=this.axis;return this._isNumber(e.height)?e.width=e.height*this.aspectRatio:this._isNumber(e.width)&&(e.height=e.width/this.aspectRatio),"sw"===s&&(e.left=t.left+(i.width-e.width),e.top=null),"nw"===s&&(e.top=t.top+(i.height-e.height),e.left=t.left+(i.width-e.width)),e},_respectSize:function(e){var t=this._vBoundaries,i=this.axis,s=this._isNumber(e.width)&&t.maxWidth&&t.maxWidth<e.width,n=this._isNumber(e.height)&&t.maxHeight&&t.maxHeight<e.height,a=this._isNumber(e.width)&&t.minWidth&&t.minWidth>e.width,o=this._isNumber(e.height)&&t.minHeight&&t.minHeight>e.height,r=this.originalPosition.left+this.originalSize.width,h=this.position.top+this.size.height,l=/sw|nw|w/.test(i),u=/nw|ne|n/.test(i);return a&&(e.width=t.minWidth),o&&(e.height=t.minHeight),s&&(e.width=t.maxWidth),n&&(e.height=t.maxHeight),a&&l&&(e.left=r-t.minWidth),s&&l&&(e.left=r-t.maxWidth),o&&u&&(e.top=h-t.minHeight),n&&u&&(e.top=h-t.maxHeight),e.width||e.height||e.left||!e.top?e.width||e.height||e.top||!e.left||(e.left=null):e.top=null,e},_getPaddingPlusBorderDimensions:function(e){for(var t=0,i=[],s=[e.css("borderTopWidth"),e.css("borderRightWidth"),e.css("borderBottomWidth"),e.css("borderLeftWidth")],n=[e.css("paddingTop"),e.css("paddingRight"),e.css("paddingBottom"),e.css("paddingLeft")];4>t;t++)i[t]=parseInt(s[t],10)||0,i[t]+=parseInt(n[t],10)||0;return{height:i[0]+i[2],width:i[1]+i[3]}},_proportionallyResize:function(){if(this._proportionallyResizeElements.length)for(var e,t=0,i=this.helper||this.element;this._proportionallyResizeElements.length>t;t++)e=this._proportionallyResizeElements[t],this.outerDimensions||(this.outerDimensions=this._getPaddingPlusBorderDimensions(e)),e.css({height:i.height()-this.outerDimensions.height||0,width:i.width()-this.outerDimensions.width||0})},_renderProxy:function(){var t=this.element,i=this.options;this.elementOffset=t.offset(),this._helper?(this.helper=this.helper||e("<div style='overflow:hidden;'></div>"),this.helper.addClass(this._helper).css({width:this.element.outerWidth()-1,height:this.element.outerHeight()-1,position:"absolute",left:this.elementOffset.left+"px",top:this.elementOffset.top+"px",zIndex:++i.zIndex}),this.helper.appendTo("body").disableSelection()):this.helper=this.element},_change:{e:function(e,t){return{width:this.originalSize.width+t}},w:function(e,t){var i=this.originalSize,s=this.originalPosition;return{left:s.left+t,width:i.width-t}},n:function(e,t,i){var s=this.originalSize,n=this.originalPosition;return{top:n.top+i,height:s.height-i}},s:function(e,t,i){return{height:this.originalSize.height+i}},se:function(t,i,s){return e.extend(this._change.s.apply(this,arguments),this._change.e.apply(this,[t,i,s]))},sw:function(t,i,s){return e.extend(this._change.s.apply(this,arguments),this._change.w.apply(this,[t,i,s]))},ne:function(t,i,s){return e.extend(this._change.n.apply(this,arguments),this._change.e.apply(this,[t,i,s]))},nw:function(t,i,s){return e.extend(this._change.n.apply(this,arguments),this._change.w.apply(this,[t,i,s]))}},_propagate:function(t,i){e.ui.plugin.call(this,t,[i,this.ui()]),"resize"!==t&&this._trigger(t,i,this.ui())},plugins:{},ui:function(){return{originalElement:this.originalElement,element:this.element,helper:this.helper,position:this.position,size:this.size,originalSize:this.originalSize,originalPosition:this.originalPosition}}}),e.ui.plugin.add("resizable","animate",{stop:function(t){var i=e(this).resizable("instance"),s=i.options,n=i._proportionallyResizeElements,a=n.length&&/textarea/i.test(n[0].nodeName),o=a&&i._hasScroll(n[0],"left")?0:i.sizeDiff.height,r=a?0:i.sizeDiff.width,h={width:i.size.width-r,height:i.size.height-o},l=parseInt(i.element.css("left"),10)+(i.position.left-i.originalPosition.left)||null,u=parseInt(i.element.css("top"),10)+(i.position.top-i.originalPosition.top)||null;i.element.animate(e.extend(h,u&&l?{top:u,left:l}:{}),{duration:s.animateDuration,easing:s.animateEasing,step:function(){var s={width:parseInt(i.element.css("width"),10),height:parseInt(i.element.css("height"),10),top:parseInt(i.element.css("top"),10),left:parseInt(i.element.css("left"),10)};n&&n.length&&e(n[0]).css({width:s.width,height:s.height}),i._updateCache(s),i._propagate("resize",t)}})}}),e.ui.plugin.add("resizable","containment",{start:function(){var t,i,s,n,a,o,r,h=e(this).resizable("instance"),l=h.options,u=h.element,d=l.containment,c=d instanceof e?d.get(0):/parent/.test(d)?u.parent().get(0):d;c&&(h.containerElement=e(c),/document/.test(d)||d===document?(h.containerOffset={left:0,top:0},h.containerPosition={left:0,top:0},h.parentData={element:e(document),left:0,top:0,width:e(document).width(),height:e(document).height()||document.body.parentNode.scrollHeight}):(t=e(c),i=[],e(["Top","Right","Left","Bottom"]).each(function(e,s){i[e]=h._num(t.css("padding"+s))}),h.containerOffset=t.offset(),h.containerPosition=t.position(),h.containerSize={height:t.innerHeight()-i[3],width:t.innerWidth()-i[1]},s=h.containerOffset,n=h.containerSize.height,a=h.containerSize.width,o=h._hasScroll(c,"left")?c.scrollWidth:a,r=h._hasScroll(c)?c.scrollHeight:n,h.parentData={element:c,left:s.left,top:s.top,width:o,height:r}))},resize:function(t){var i,s,n,a,o=e(this).resizable("instance"),r=o.options,h=o.containerOffset,l=o.position,u=o._aspectRatio||t.shiftKey,d={top:0,left:0},c=o.containerElement,p=!0;c[0]!==document&&/static/.test(c.css("position"))&&(d=h),l.left<(o._helper?h.left:0)&&(o.size.width=o.size.width+(o._helper?o.position.left-h.left:o.position.left-d.left),u&&(o.size.height=o.size.width/o.aspectRatio,p=!1),o.position.left=r.helper?h.left:0),l.top<(o._helper?h.top:0)&&(o.size.height=o.size.height+(o._helper?o.position.top-h.top:o.position.top),u&&(o.size.width=o.size.height*o.aspectRatio,p=!1),o.position.top=o._helper?h.top:0),n=o.containerElement.get(0)===o.element.parent().get(0),a=/relative|absolute/.test(o.containerElement.css("position")),n&&a?(o.offset.left=o.parentData.left+o.position.left,o.offset.top=o.parentData.top+o.position.top):(o.offset.left=o.element.offset().left,o.offset.top=o.element.offset().top),i=Math.abs(o.sizeDiff.width+(o._helper?o.offset.left-d.left:o.offset.left-h.left)),s=Math.abs(o.sizeDiff.height+(o._helper?o.offset.top-d.top:o.offset.top-h.top)),i+o.size.width>=o.parentData.width&&(o.size.width=o.parentData.width-i,u&&(o.size.height=o.size.width/o.aspectRatio,p=!1)),s+o.size.height>=o.parentData.height&&(o.size.height=o.parentData.height-s,u&&(o.size.width=o.size.height*o.aspectRatio,p=!1)),p||(o.position.left=o.prevPosition.left,o.position.top=o.prevPosition.top,o.size.width=o.prevSize.width,o.size.height=o.prevSize.height)},stop:function(){var t=e(this).resizable("instance"),i=t.options,s=t.containerOffset,n=t.containerPosition,a=t.containerElement,o=e(t.helper),r=o.offset(),h=o.outerWidth()-t.sizeDiff.width,l=o.outerHeight()-t.sizeDiff.height;t._helper&&!i.animate&&/relative/.test(a.css("position"))&&e(this).css({left:r.left-n.left-s.left,width:h,height:l}),t._helper&&!i.animate&&/static/.test(a.css("position"))&&e(this).css({left:r.left-n.left-s.left,width:h,height:l})}}),e.ui.plugin.add("resizable","alsoResize",{start:function(){var t=e(this).resizable("instance"),i=t.options;e(i.alsoResize).each(function(){var t=e(this);t.data("ui-resizable-alsoresize",{width:parseInt(t.width(),10),height:parseInt(t.height(),10),left:parseInt(t.css("left"),10),top:parseInt(t.css("top"),10)})})},resize:function(t,i){var s=e(this).resizable("instance"),n=s.options,a=s.originalSize,o=s.originalPosition,r={height:s.size.height-a.height||0,width:s.size.width-a.width||0,top:s.position.top-o.top||0,left:s.position.left-o.left||0};e(n.alsoResize).each(function(){var t=e(this),s=e(this).data("ui-resizable-alsoresize"),n={},a=t.parents(i.originalElement[0]).length?["width","height"]:["width","height","top","left"];e.each(a,function(e,t){var i=(s[t]||0)+(r[t]||0);i&&i>=0&&(n[t]=i||null)}),t.css(n)})},stop:function(){e(this).removeData("resizable-alsoresize")}}),e.ui.plugin.add("resizable","ghost",{start:function(){var t=e(this).resizable("instance"),i=t.options,s=t.size;t.ghost=t.originalElement.clone(),t.ghost.css({opacity:.25,display:"block",position:"relative",height:s.height,width:s.width,margin:0,left:0,top:0}).addClass("ui-resizable-ghost").addClass("string"==typeof i.ghost?i.ghost:""),t.ghost.appendTo(t.helper)},resize:function(){var t=e(this).resizable("instance");t.ghost&&t.ghost.css({position:"relative",height:t.size.height,width:t.size.width})},stop:function(){var t=e(this).resizable("instance");t.ghost&&t.helper&&t.helper.get(0).removeChild(t.ghost.get(0))}}),e.ui.plugin.add("resizable","grid",{resize:function(){var t,i=e(this).resizable("instance"),s=i.options,n=i.size,a=i.originalSize,o=i.originalPosition,r=i.axis,h="number"==typeof s.grid?[s.grid,s.grid]:s.grid,l=h[0]||1,u=h[1]||1,d=Math.round((n.width-a.width)/l)*l,c=Math.round((n.height-a.height)/u)*u,p=a.width+d,f=a.height+c,m=s.maxWidth&&p>s.maxWidth,g=s.maxHeight&&f>s.maxHeight,v=s.minWidth&&s.minWidth>p,y=s.minHeight&&s.minHeight>f;s.grid=h,v&&(p+=l),y&&(f+=u),m&&(p-=l),g&&(f-=u),/^(se|s|e)$/.test(r)?(i.size.width=p,i.size.height=f):/^(ne)$/.test(r)?(i.size.width=p,i.size.height=f,i.position.top=o.top-c):/^(sw)$/.test(r)?(i.size.width=p,i.size.height=f,i.position.left=o.left-d):((0>=f-u||0>=p-l)&&(t=i._getPaddingPlusBorderDimensions(this)),f-u>0?(i.size.height=f,i.position.top=o.top-c):(f=u-t.height,i.size.height=f,i.position.top=o.top+a.height-f),p-l>0?(i.size.width=p,i.position.left=o.left-d):(p=l-t.width,i.size.width=p,i.position.left=o.left+a.width-p))}}),e.ui.resizable,e.widget("ui.selectable",e.ui.mouse,{version:"1.11.4",options:{appendTo:"body",autoRefresh:!0,distance:0,filter:"*",tolerance:"touch",selected:null,selecting:null,start:null,stop:null,unselected:null,unselecting:null},_create:function(){var t,i=this;this.element.addClass("ui-selectable"),this.dragged=!1,this.refresh=function(){t=e(i.options.filter,i.element[0]),t.addClass("ui-selectee"),t.each(function(){var t=e(this),i=t.offset();e.data(this,"selectable-item",{element:this,$element:t,left:i.left,top:i.top,right:i.left+t.outerWidth(),bottom:i.top+t.outerHeight(),startselected:!1,selected:t.hasClass("ui-selected"),selecting:t.hasClass("ui-selecting"),unselecting:t.hasClass("ui-unselecting")})})},this.refresh(),this.selectees=t.addClass("ui-selectee"),this._mouseInit(),this.helper=e("<div class='ui-selectable-helper'></div>")},_destroy:function(){this.selectees.removeClass("ui-selectee").removeData("selectable-item"),this.element.removeClass("ui-selectable ui-selectable-disabled"),this._mouseDestroy()},_mouseStart:function(t){var i=this,s=this.options;this.opos=[t.pageX,t.pageY],this.options.disabled||(this.selectees=e(s.filter,this.element[0]),this._trigger("start",t),e(s.appendTo).append(this.helper),this.helper.css({left:t.pageX,top:t.pageY,width:0,height:0}),s.autoRefresh&&this.refresh(),this.selectees.filter(".ui-selected").each(function(){var s=e.data(this,"selectable-item");s.startselected=!0,t.metaKey||t.ctrlKey||(s.$element.removeClass("ui-selected"),s.selected=!1,s.$element.addClass("ui-unselecting"),s.unselecting=!0,i._trigger("unselecting",t,{unselecting:s.element}))}),e(t.target).parents().addBack().each(function(){var s,n=e.data(this,"selectable-item");return n?(s=!t.metaKey&&!t.ctrlKey||!n.$element.hasClass("ui-selected"),n.$element.removeClass(s?"ui-unselecting":"ui-selected").addClass(s?"ui-selecting":"ui-unselecting"),n.unselecting=!s,n.selecting=s,n.selected=s,s?i._trigger("selecting",t,{selecting:n.element}):i._trigger("unselecting",t,{unselecting:n.element}),!1):void 0}))},_mouseDrag:function(t){if(this.dragged=!0,!this.options.disabled){var i,s=this,n=this.options,a=this.opos[0],o=this.opos[1],r=t.pageX,h=t.pageY;return a>r&&(i=r,r=a,a=i),o>h&&(i=h,h=o,o=i),this.helper.css({left:a,top:o,width:r-a,height:h-o}),this.selectees.each(function(){var i=e.data(this,"selectable-item"),l=!1;
i&&i.element!==s.element[0]&&("touch"===n.tolerance?l=!(i.left>r||a>i.right||i.top>h||o>i.bottom):"fit"===n.tolerance&&(l=i.left>a&&r>i.right&&i.top>o&&h>i.bottom),l?(i.selected&&(i.$element.removeClass("ui-selected"),i.selected=!1),i.unselecting&&(i.$element.removeClass("ui-unselecting"),i.unselecting=!1),i.selecting||(i.$element.addClass("ui-selecting"),i.selecting=!0,s._trigger("selecting",t,{selecting:i.element}))):(i.selecting&&((t.metaKey||t.ctrlKey)&&i.startselected?(i.$element.removeClass("ui-selecting"),i.selecting=!1,i.$element.addClass("ui-selected"),i.selected=!0):(i.$element.removeClass("ui-selecting"),i.selecting=!1,i.startselected&&(i.$element.addClass("ui-unselecting"),i.unselecting=!0),s._trigger("unselecting",t,{unselecting:i.element}))),i.selected&&(t.metaKey||t.ctrlKey||i.startselected||(i.$element.removeClass("ui-selected"),i.selected=!1,i.$element.addClass("ui-unselecting"),i.unselecting=!0,s._trigger("unselecting",t,{unselecting:i.element})))))}),!1}},_mouseStop:function(t){var i=this;return this.dragged=!1,e(".ui-unselecting",this.element[0]).each(function(){var s=e.data(this,"selectable-item");s.$element.removeClass("ui-unselecting"),s.unselecting=!1,s.startselected=!1,i._trigger("unselected",t,{unselected:s.element})}),e(".ui-selecting",this.element[0]).each(function(){var s=e.data(this,"selectable-item");s.$element.removeClass("ui-selecting").addClass("ui-selected"),s.selecting=!1,s.selected=!0,s.startselected=!0,i._trigger("selected",t,{selected:s.element})}),this._trigger("stop",t),this.helper.remove(),!1}}),e.widget("ui.sortable",e.ui.mouse,{version:"1.11.4",widgetEventPrefix:"sort",ready:!1,options:{appendTo:"parent",axis:!1,connectWith:!1,containment:!1,cursor:"auto",cursorAt:!1,dropOnEmpty:!0,forcePlaceholderSize:!1,forceHelperSize:!1,grid:!1,handle:!1,helper:"original",items:"> *",opacity:!1,placeholder:!1,revert:!1,scroll:!0,scrollSensitivity:20,scrollSpeed:20,scope:"default",tolerance:"intersect",zIndex:1e3,activate:null,beforeStop:null,change:null,deactivate:null,out:null,over:null,receive:null,remove:null,sort:null,start:null,stop:null,update:null},_isOverAxis:function(e,t,i){return e>=t&&t+i>e},_isFloating:function(e){return/left|right/.test(e.css("float"))||/inline|table-cell/.test(e.css("display"))},_create:function(){this.containerCache={},this.element.addClass("ui-sortable"),this.refresh(),this.offset=this.element.offset(),this._mouseInit(),this._setHandleClassName(),this.ready=!0},_setOption:function(e,t){this._super(e,t),"handle"===e&&this._setHandleClassName()},_setHandleClassName:function(){this.element.find(".ui-sortable-handle").removeClass("ui-sortable-handle"),e.each(this.items,function(){(this.instance.options.handle?this.item.find(this.instance.options.handle):this.item).addClass("ui-sortable-handle")})},_destroy:function(){this.element.removeClass("ui-sortable ui-sortable-disabled").find(".ui-sortable-handle").removeClass("ui-sortable-handle"),this._mouseDestroy();for(var e=this.items.length-1;e>=0;e--)this.items[e].item.removeData(this.widgetName+"-item");return this},_mouseCapture:function(t,i){var s=null,n=!1,a=this;return this.reverting?!1:this.options.disabled||"static"===this.options.type?!1:(this._refreshItems(t),e(t.target).parents().each(function(){return e.data(this,a.widgetName+"-item")===a?(s=e(this),!1):void 0}),e.data(t.target,a.widgetName+"-item")===a&&(s=e(t.target)),s?!this.options.handle||i||(e(this.options.handle,s).find("*").addBack().each(function(){this===t.target&&(n=!0)}),n)?(this.currentItem=s,this._removeCurrentsFromItems(),!0):!1:!1)},_mouseStart:function(t,i,s){var n,a,o=this.options;if(this.currentContainer=this,this.refreshPositions(),this.helper=this._createHelper(t),this._cacheHelperProportions(),this._cacheMargins(),this.scrollParent=this.helper.scrollParent(),this.offset=this.currentItem.offset(),this.offset={top:this.offset.top-this.margins.top,left:this.offset.left-this.margins.left},e.extend(this.offset,{click:{left:t.pageX-this.offset.left,top:t.pageY-this.offset.top},parent:this._getParentOffset(),relative:this._getRelativeOffset()}),this.helper.css("position","absolute"),this.cssPosition=this.helper.css("position"),this.originalPosition=this._generatePosition(t),this.originalPageX=t.pageX,this.originalPageY=t.pageY,o.cursorAt&&this._adjustOffsetFromHelper(o.cursorAt),this.domPosition={prev:this.currentItem.prev()[0],parent:this.currentItem.parent()[0]},this.helper[0]!==this.currentItem[0]&&this.currentItem.hide(),this._createPlaceholder(),o.containment&&this._setContainment(),o.cursor&&"auto"!==o.cursor&&(a=this.document.find("body"),this.storedCursor=a.css("cursor"),a.css("cursor",o.cursor),this.storedStylesheet=e("<style>*{ cursor: "+o.cursor+" !important; }</style>").appendTo(a)),o.opacity&&(this.helper.css("opacity")&&(this._storedOpacity=this.helper.css("opacity")),this.helper.css("opacity",o.opacity)),o.zIndex&&(this.helper.css("zIndex")&&(this._storedZIndex=this.helper.css("zIndex")),this.helper.css("zIndex",o.zIndex)),this.scrollParent[0]!==this.document[0]&&"HTML"!==this.scrollParent[0].tagName&&(this.overflowOffset=this.scrollParent.offset()),this._trigger("start",t,this._uiHash()),this._preserveHelperProportions||this._cacheHelperProportions(),!s)for(n=this.containers.length-1;n>=0;n--)this.containers[n]._trigger("activate",t,this._uiHash(this));return e.ui.ddmanager&&(e.ui.ddmanager.current=this),e.ui.ddmanager&&!o.dropBehaviour&&e.ui.ddmanager.prepareOffsets(this,t),this.dragging=!0,this.helper.addClass("ui-sortable-helper"),this._mouseDrag(t),!0},_mouseDrag:function(t){var i,s,n,a,o=this.options,r=!1;for(this.position=this._generatePosition(t),this.positionAbs=this._convertPositionTo("absolute"),this.lastPositionAbs||(this.lastPositionAbs=this.positionAbs),this.options.scroll&&(this.scrollParent[0]!==this.document[0]&&"HTML"!==this.scrollParent[0].tagName?(this.overflowOffset.top+this.scrollParent[0].offsetHeight-t.pageY<o.scrollSensitivity?this.scrollParent[0].scrollTop=r=this.scrollParent[0].scrollTop+o.scrollSpeed:t.pageY-this.overflowOffset.top<o.scrollSensitivity&&(this.scrollParent[0].scrollTop=r=this.scrollParent[0].scrollTop-o.scrollSpeed),this.overflowOffset.left+this.scrollParent[0].offsetWidth-t.pageX<o.scrollSensitivity?this.scrollParent[0].scrollLeft=r=this.scrollParent[0].scrollLeft+o.scrollSpeed:t.pageX-this.overflowOffset.left<o.scrollSensitivity&&(this.scrollParent[0].scrollLeft=r=this.scrollParent[0].scrollLeft-o.scrollSpeed)):(t.pageY-this.document.scrollTop()<o.scrollSensitivity?r=this.document.scrollTop(this.document.scrollTop()-o.scrollSpeed):this.window.height()-(t.pageY-this.document.scrollTop())<o.scrollSensitivity&&(r=this.document.scrollTop(this.document.scrollTop()+o.scrollSpeed)),t.pageX-this.document.scrollLeft()<o.scrollSensitivity?r=this.document.scrollLeft(this.document.scrollLeft()-o.scrollSpeed):this.window.width()-(t.pageX-this.document.scrollLeft())<o.scrollSensitivity&&(r=this.document.scrollLeft(this.document.scrollLeft()+o.scrollSpeed))),r!==!1&&e.ui.ddmanager&&!o.dropBehaviour&&e.ui.ddmanager.prepareOffsets(this,t)),this.positionAbs=this._convertPositionTo("absolute"),this.options.axis&&"y"===this.options.axis||(this.helper[0].style.left=this.position.left+"px"),this.options.axis&&"x"===this.options.axis||(this.helper[0].style.top=this.position.top+"px"),i=this.items.length-1;i>=0;i--)if(s=this.items[i],n=s.item[0],a=this._intersectsWithPointer(s),a&&s.instance===this.currentContainer&&n!==this.currentItem[0]&&this.placeholder[1===a?"next":"prev"]()[0]!==n&&!e.contains(this.placeholder[0],n)&&("semi-dynamic"===this.options.type?!e.contains(this.element[0],n):!0)){if(this.direction=1===a?"down":"up","pointer"!==this.options.tolerance&&!this._intersectsWithSides(s))break;this._rearrange(t,s),this._trigger("change",t,this._uiHash());break}return this._contactContainers(t),e.ui.ddmanager&&e.ui.ddmanager.drag(this,t),this._trigger("sort",t,this._uiHash()),this.lastPositionAbs=this.positionAbs,!1},_mouseStop:function(t,i){if(t){if(e.ui.ddmanager&&!this.options.dropBehaviour&&e.ui.ddmanager.drop(this,t),this.options.revert){var s=this,n=this.placeholder.offset(),a=this.options.axis,o={};a&&"x"!==a||(o.left=n.left-this.offset.parent.left-this.margins.left+(this.offsetParent[0]===this.document[0].body?0:this.offsetParent[0].scrollLeft)),a&&"y"!==a||(o.top=n.top-this.offset.parent.top-this.margins.top+(this.offsetParent[0]===this.document[0].body?0:this.offsetParent[0].scrollTop)),this.reverting=!0,e(this.helper).animate(o,parseInt(this.options.revert,10)||500,function(){s._clear(t)})}else this._clear(t,i);return!1}},cancel:function(){if(this.dragging){this._mouseUp({target:null}),"original"===this.options.helper?this.currentItem.css(this._storedCSS).removeClass("ui-sortable-helper"):this.currentItem.show();for(var t=this.containers.length-1;t>=0;t--)this.containers[t]._trigger("deactivate",null,this._uiHash(this)),this.containers[t].containerCache.over&&(this.containers[t]._trigger("out",null,this._uiHash(this)),this.containers[t].containerCache.over=0)}return this.placeholder&&(this.placeholder[0].parentNode&&this.placeholder[0].parentNode.removeChild(this.placeholder[0]),"original"!==this.options.helper&&this.helper&&this.helper[0].parentNode&&this.helper.remove(),e.extend(this,{helper:null,dragging:!1,reverting:!1,_noFinalSort:null}),this.domPosition.prev?e(this.domPosition.prev).after(this.currentItem):e(this.domPosition.parent).prepend(this.currentItem)),this},serialize:function(t){var i=this._getItemsAsjQuery(t&&t.connected),s=[];return t=t||{},e(i).each(function(){var i=(e(t.item||this).attr(t.attribute||"id")||"").match(t.expression||/(.+)[\-=_](.+)/);i&&s.push((t.key||i[1]+"[]")+"="+(t.key&&t.expression?i[1]:i[2]))}),!s.length&&t.key&&s.push(t.key+"="),s.join("&")},toArray:function(t){var i=this._getItemsAsjQuery(t&&t.connected),s=[];return t=t||{},i.each(function(){s.push(e(t.item||this).attr(t.attribute||"id")||"")}),s},_intersectsWith:function(e){var t=this.positionAbs.left,i=t+this.helperProportions.width,s=this.positionAbs.top,n=s+this.helperProportions.height,a=e.left,o=a+e.width,r=e.top,h=r+e.height,l=this.offset.click.top,u=this.offset.click.left,d="x"===this.options.axis||s+l>r&&h>s+l,c="y"===this.options.axis||t+u>a&&o>t+u,p=d&&c;return"pointer"===this.options.tolerance||this.options.forcePointerForContainers||"pointer"!==this.options.tolerance&&this.helperProportions[this.floating?"width":"height"]>e[this.floating?"width":"height"]?p:t+this.helperProportions.width/2>a&&o>i-this.helperProportions.width/2&&s+this.helperProportions.height/2>r&&h>n-this.helperProportions.height/2},_intersectsWithPointer:function(e){var t="x"===this.options.axis||this._isOverAxis(this.positionAbs.top+this.offset.click.top,e.top,e.height),i="y"===this.options.axis||this._isOverAxis(this.positionAbs.left+this.offset.click.left,e.left,e.width),s=t&&i,n=this._getDragVerticalDirection(),a=this._getDragHorizontalDirection();return s?this.floating?a&&"right"===a||"down"===n?2:1:n&&("down"===n?2:1):!1},_intersectsWithSides:function(e){var t=this._isOverAxis(this.positionAbs.top+this.offset.click.top,e.top+e.height/2,e.height),i=this._isOverAxis(this.positionAbs.left+this.offset.click.left,e.left+e.width/2,e.width),s=this._getDragVerticalDirection(),n=this._getDragHorizontalDirection();return this.floating&&n?"right"===n&&i||"left"===n&&!i:s&&("down"===s&&t||"up"===s&&!t)},_getDragVerticalDirection:function(){var e=this.positionAbs.top-this.lastPositionAbs.top;return 0!==e&&(e>0?"down":"up")},_getDragHorizontalDirection:function(){var e=this.positionAbs.left-this.lastPositionAbs.left;return 0!==e&&(e>0?"right":"left")},refresh:function(e){return this._refreshItems(e),this._setHandleClassName(),this.refreshPositions(),this},_connectWith:function(){var e=this.options;return e.connectWith.constructor===String?[e.connectWith]:e.connectWith},_getItemsAsjQuery:function(t){function i(){r.push(this)}var s,n,a,o,r=[],h=[],l=this._connectWith();if(l&&t)for(s=l.length-1;s>=0;s--)for(a=e(l[s],this.document[0]),n=a.length-1;n>=0;n--)o=e.data(a[n],this.widgetFullName),o&&o!==this&&!o.options.disabled&&h.push([e.isFunction(o.options.items)?o.options.items.call(o.element):e(o.options.items,o.element).not(".ui-sortable-helper").not(".ui-sortable-placeholder"),o]);for(h.push([e.isFunction(this.options.items)?this.options.items.call(this.element,null,{options:this.options,item:this.currentItem}):e(this.options.items,this.element).not(".ui-sortable-helper").not(".ui-sortable-placeholder"),this]),s=h.length-1;s>=0;s--)h[s][0].each(i);return e(r)},_removeCurrentsFromItems:function(){var t=this.currentItem.find(":data("+this.widgetName+"-item)");this.items=e.grep(this.items,function(e){for(var i=0;t.length>i;i++)if(t[i]===e.item[0])return!1;return!0})},_refreshItems:function(t){this.items=[],this.containers=[this];var i,s,n,a,o,r,h,l,u=this.items,d=[[e.isFunction(this.options.items)?this.options.items.call(this.element[0],t,{item:this.currentItem}):e(this.options.items,this.element),this]],c=this._connectWith();if(c&&this.ready)for(i=c.length-1;i>=0;i--)for(n=e(c[i],this.document[0]),s=n.length-1;s>=0;s--)a=e.data(n[s],this.widgetFullName),a&&a!==this&&!a.options.disabled&&(d.push([e.isFunction(a.options.items)?a.options.items.call(a.element[0],t,{item:this.currentItem}):e(a.options.items,a.element),a]),this.containers.push(a));for(i=d.length-1;i>=0;i--)for(o=d[i][1],r=d[i][0],s=0,l=r.length;l>s;s++)h=e(r[s]),h.data(this.widgetName+"-item",o),u.push({item:h,instance:o,width:0,height:0,left:0,top:0})},refreshPositions:function(t){this.floating=this.items.length?"x"===this.options.axis||this._isFloating(this.items[0].item):!1,this.offsetParent&&this.helper&&(this.offset.parent=this._getParentOffset());var i,s,n,a;for(i=this.items.length-1;i>=0;i--)s=this.items[i],s.instance!==this.currentContainer&&this.currentContainer&&s.item[0]!==this.currentItem[0]||(n=this.options.toleranceElement?e(this.options.toleranceElement,s.item):s.item,t||(s.width=n.outerWidth(),s.height=n.outerHeight()),a=n.offset(),s.left=a.left,s.top=a.top);if(this.options.custom&&this.options.custom.refreshContainers)this.options.custom.refreshContainers.call(this);else for(i=this.containers.length-1;i>=0;i--)a=this.containers[i].element.offset(),this.containers[i].containerCache.left=a.left,this.containers[i].containerCache.top=a.top,this.containers[i].containerCache.width=this.containers[i].element.outerWidth(),this.containers[i].containerCache.height=this.containers[i].element.outerHeight();return this},_createPlaceholder:function(t){t=t||this;var i,s=t.options;s.placeholder&&s.placeholder.constructor!==String||(i=s.placeholder,s.placeholder={element:function(){var s=t.currentItem[0].nodeName.toLowerCase(),n=e("<"+s+">",t.document[0]).addClass(i||t.currentItem[0].className+" ui-sortable-placeholder").removeClass("ui-sortable-helper");return"tbody"===s?t._createTrPlaceholder(t.currentItem.find("tr").eq(0),e("<tr>",t.document[0]).appendTo(n)):"tr"===s?t._createTrPlaceholder(t.currentItem,n):"img"===s&&n.attr("src",t.currentItem.attr("src")),i||n.css("visibility","hidden"),n},update:function(e,n){(!i||s.forcePlaceholderSize)&&(n.height()||n.height(t.currentItem.innerHeight()-parseInt(t.currentItem.css("paddingTop")||0,10)-parseInt(t.currentItem.css("paddingBottom")||0,10)),n.width()||n.width(t.currentItem.innerWidth()-parseInt(t.currentItem.css("paddingLeft")||0,10)-parseInt(t.currentItem.css("paddingRight")||0,10)))}}),t.placeholder=e(s.placeholder.element.call(t.element,t.currentItem)),t.currentItem.after(t.placeholder),s.placeholder.update(t,t.placeholder)},_createTrPlaceholder:function(t,i){var s=this;t.children().each(function(){e("<td>&#160;</td>",s.document[0]).attr("colspan",e(this).attr("colspan")||1).appendTo(i)})},_contactContainers:function(t){var i,s,n,a,o,r,h,l,u,d,c=null,p=null;for(i=this.containers.length-1;i>=0;i--)if(!e.contains(this.currentItem[0],this.containers[i].element[0]))if(this._intersectsWith(this.containers[i].containerCache)){if(c&&e.contains(this.containers[i].element[0],c.element[0]))continue;c=this.containers[i],p=i}else this.containers[i].containerCache.over&&(this.containers[i]._trigger("out",t,this._uiHash(this)),this.containers[i].containerCache.over=0);if(c)if(1===this.containers.length)this.containers[p].containerCache.over||(this.containers[p]._trigger("over",t,this._uiHash(this)),this.containers[p].containerCache.over=1);else{for(n=1e4,a=null,u=c.floating||this._isFloating(this.currentItem),o=u?"left":"top",r=u?"width":"height",d=u?"clientX":"clientY",s=this.items.length-1;s>=0;s--)e.contains(this.containers[p].element[0],this.items[s].item[0])&&this.items[s].item[0]!==this.currentItem[0]&&(h=this.items[s].item.offset()[o],l=!1,t[d]-h>this.items[s][r]/2&&(l=!0),n>Math.abs(t[d]-h)&&(n=Math.abs(t[d]-h),a=this.items[s],this.direction=l?"up":"down"));if(!a&&!this.options.dropOnEmpty)return;if(this.currentContainer===this.containers[p])return this.currentContainer.containerCache.over||(this.containers[p]._trigger("over",t,this._uiHash()),this.currentContainer.containerCache.over=1),void 0;a?this._rearrange(t,a,null,!0):this._rearrange(t,null,this.containers[p].element,!0),this._trigger("change",t,this._uiHash()),this.containers[p]._trigger("change",t,this._uiHash(this)),this.currentContainer=this.containers[p],this.options.placeholder.update(this.currentContainer,this.placeholder),this.containers[p]._trigger("over",t,this._uiHash(this)),this.containers[p].containerCache.over=1}},_createHelper:function(t){var i=this.options,s=e.isFunction(i.helper)?e(i.helper.apply(this.element[0],[t,this.currentItem])):"clone"===i.helper?this.currentItem.clone():this.currentItem;return s.parents("body").length||e("parent"!==i.appendTo?i.appendTo:this.currentItem[0].parentNode)[0].appendChild(s[0]),s[0]===this.currentItem[0]&&(this._storedCSS={width:this.currentItem[0].style.width,height:this.currentItem[0].style.height,position:this.currentItem.css("position"),top:this.currentItem.css("top"),left:this.currentItem.css("left")}),(!s[0].style.width||i.forceHelperSize)&&s.width(this.currentItem.width()),(!s[0].style.height||i.forceHelperSize)&&s.height(this.currentItem.height()),s},_adjustOffsetFromHelper:function(t){"string"==typeof t&&(t=t.split(" ")),e.isArray(t)&&(t={left:+t[0],top:+t[1]||0}),"left"in t&&(this.offset.click.left=t.left+this.margins.left),"right"in t&&(this.offset.click.left=this.helperProportions.width-t.right+this.margins.left),"top"in t&&(this.offset.click.top=t.top+this.margins.top),"bottom"in t&&(this.offset.click.top=this.helperProportions.height-t.bottom+this.margins.top)},_getParentOffset:function(){this.offsetParent=this.helper.offsetParent();var t=this.offsetParent.offset();return"absolute"===this.cssPosition&&this.scrollParent[0]!==this.document[0]&&e.contains(this.scrollParent[0],this.offsetParent[0])&&(t.left+=this.scrollParent.scrollLeft(),t.top+=this.scrollParent.scrollTop()),(this.offsetParent[0]===this.document[0].body||this.offsetParent[0].tagName&&"html"===this.offsetParent[0].tagName.toLowerCase()&&e.ui.ie)&&(t={top:0,left:0}),{top:t.top+(parseInt(this.offsetParent.css("borderTopWidth"),10)||0),left:t.left+(parseInt(this.offsetParent.css("borderLeftWidth"),10)||0)}},_getRelativeOffset:function(){if("relative"===this.cssPosition){var e=this.currentItem.position();return{top:e.top-(parseInt(this.helper.css("top"),10)||0)+this.scrollParent.scrollTop(),left:e.left-(parseInt(this.helper.css("left"),10)||0)+this.scrollParent.scrollLeft()}}return{top:0,left:0}},_cacheMargins:function(){this.margins={left:parseInt(this.currentItem.css("marginLeft"),10)||0,top:parseInt(this.currentItem.css("marginTop"),10)||0}},_cacheHelperProportions:function(){this.helperProportions={width:this.helper.outerWidth(),height:this.helper.outerHeight()}},_setContainment:function(){var t,i,s,n=this.options;"parent"===n.containment&&(n.containment=this.helper[0].parentNode),("document"===n.containment||"window"===n.containment)&&(this.containment=[0-this.offset.relative.left-this.offset.parent.left,0-this.offset.relative.top-this.offset.parent.top,"document"===n.containment?this.document.width():this.window.width()-this.helperProportions.width-this.margins.left,("document"===n.containment?this.document.width():this.window.height()||this.document[0].body.parentNode.scrollHeight)-this.helperProportions.height-this.margins.top]),/^(document|window|parent)$/.test(n.containment)||(t=e(n.containment)[0],i=e(n.containment).offset(),s="hidden"!==e(t).css("overflow"),this.containment=[i.left+(parseInt(e(t).css("borderLeftWidth"),10)||0)+(parseInt(e(t).css("paddingLeft"),10)||0)-this.margins.left,i.top+(parseInt(e(t).css("borderTopWidth"),10)||0)+(parseInt(e(t).css("paddingTop"),10)||0)-this.margins.top,i.left+(s?Math.max(t.scrollWidth,t.offsetWidth):t.offsetWidth)-(parseInt(e(t).css("borderLeftWidth"),10)||0)-(parseInt(e(t).css("paddingRight"),10)||0)-this.helperProportions.width-this.margins.left,i.top+(s?Math.max(t.scrollHeight,t.offsetHeight):t.offsetHeight)-(parseInt(e(t).css("borderTopWidth"),10)||0)-(parseInt(e(t).css("paddingBottom"),10)||0)-this.helperProportions.height-this.margins.top])},_convertPositionTo:function(t,i){i||(i=this.position);var s="absolute"===t?1:-1,n="absolute"!==this.cssPosition||this.scrollParent[0]!==this.document[0]&&e.contains(this.scrollParent[0],this.offsetParent[0])?this.scrollParent:this.offsetParent,a=/(html|body)/i.test(n[0].tagName);return{top:i.top+this.offset.relative.top*s+this.offset.parent.top*s-("fixed"===this.cssPosition?-this.scrollParent.scrollTop():a?0:n.scrollTop())*s,left:i.left+this.offset.relative.left*s+this.offset.parent.left*s-("fixed"===this.cssPosition?-this.scrollParent.scrollLeft():a?0:n.scrollLeft())*s}},_generatePosition:function(t){var i,s,n=this.options,a=t.pageX,o=t.pageY,r="absolute"!==this.cssPosition||this.scrollParent[0]!==this.document[0]&&e.contains(this.scrollParent[0],this.offsetParent[0])?this.scrollParent:this.offsetParent,h=/(html|body)/i.test(r[0].tagName);return"relative"!==this.cssPosition||this.scrollParent[0]!==this.document[0]&&this.scrollParent[0]!==this.offsetParent[0]||(this.offset.relative=this._getRelativeOffset()),this.originalPosition&&(this.containment&&(t.pageX-this.offset.click.left<this.containment[0]&&(a=this.containment[0]+this.offset.click.left),t.pageY-this.offset.click.top<this.containment[1]&&(o=this.containment[1]+this.offset.click.top),t.pageX-this.offset.click.left>this.containment[2]&&(a=this.containment[2]+this.offset.click.left),t.pageY-this.offset.click.top>this.containment[3]&&(o=this.containment[3]+this.offset.click.top)),n.grid&&(i=this.originalPageY+Math.round((o-this.originalPageY)/n.grid[1])*n.grid[1],o=this.containment?i-this.offset.click.top>=this.containment[1]&&i-this.offset.click.top<=this.containment[3]?i:i-this.offset.click.top>=this.containment[1]?i-n.grid[1]:i+n.grid[1]:i,s=this.originalPageX+Math.round((a-this.originalPageX)/n.grid[0])*n.grid[0],a=this.containment?s-this.offset.click.left>=this.containment[0]&&s-this.offset.click.left<=this.containment[2]?s:s-this.offset.click.left>=this.containment[0]?s-n.grid[0]:s+n.grid[0]:s)),{top:o-this.offset.click.top-this.offset.relative.top-this.offset.parent.top+("fixed"===this.cssPosition?-this.scrollParent.scrollTop():h?0:r.scrollTop()),left:a-this.offset.click.left-this.offset.relative.left-this.offset.parent.left+("fixed"===this.cssPosition?-this.scrollParent.scrollLeft():h?0:r.scrollLeft())}},_rearrange:function(e,t,i,s){i?i[0].appendChild(this.placeholder[0]):t.item[0].parentNode.insertBefore(this.placeholder[0],"down"===this.direction?t.item[0]:t.item[0].nextSibling),this.counter=this.counter?++this.counter:1;var n=this.counter;this._delay(function(){n===this.counter&&this.refreshPositions(!s)})},_clear:function(e,t){function i(e,t,i){return function(s){i._trigger(e,s,t._uiHash(t))}}this.reverting=!1;var s,n=[];if(!this._noFinalSort&&this.currentItem.parent().length&&this.placeholder.before(this.currentItem),this._noFinalSort=null,this.helper[0]===this.currentItem[0]){for(s in this._storedCSS)("auto"===this._storedCSS[s]||"static"===this._storedCSS[s])&&(this._storedCSS[s]="");this.currentItem.css(this._storedCSS).removeClass("ui-sortable-helper")}else this.currentItem.show();for(this.fromOutside&&!t&&n.push(function(e){this._trigger("receive",e,this._uiHash(this.fromOutside))}),!this.fromOutside&&this.domPosition.prev===this.currentItem.prev().not(".ui-sortable-helper")[0]&&this.domPosition.parent===this.currentItem.parent()[0]||t||n.push(function(e){this._trigger("update",e,this._uiHash())}),this!==this.currentContainer&&(t||(n.push(function(e){this._trigger("remove",e,this._uiHash())}),n.push(function(e){return function(t){e._trigger("receive",t,this._uiHash(this))}}.call(this,this.currentContainer)),n.push(function(e){return function(t){e._trigger("update",t,this._uiHash(this))}}.call(this,this.currentContainer)))),s=this.containers.length-1;s>=0;s--)t||n.push(i("deactivate",this,this.containers[s])),this.containers[s].containerCache.over&&(n.push(i("out",this,this.containers[s])),this.containers[s].containerCache.over=0);if(this.storedCursor&&(this.document.find("body").css("cursor",this.storedCursor),this.storedStylesheet.remove()),this._storedOpacity&&this.helper.css("opacity",this._storedOpacity),this._storedZIndex&&this.helper.css("zIndex","auto"===this._storedZIndex?"":this._storedZIndex),this.dragging=!1,t||this._trigger("beforeStop",e,this._uiHash()),this.placeholder[0].parentNode.removeChild(this.placeholder[0]),this.cancelHelperRemoval||(this.helper[0]!==this.currentItem[0]&&this.helper.remove(),this.helper=null),!t){for(s=0;n.length>s;s++)n[s].call(this,e);this._trigger("stop",e,this._uiHash())}return this.fromOutside=!1,!this.cancelHelperRemoval},_trigger:function(){e.Widget.prototype._trigger.apply(this,arguments)===!1&&this.cancel()},_uiHash:function(t){var i=t||this;return{helper:i.helper,placeholder:i.placeholder||e([]),position:i.position,originalPosition:i.originalPosition,offset:i.positionAbs,item:i.currentItem,sender:t?t.element:null}}}),e.widget("ui.accordion",{version:"1.11.4",options:{active:0,animate:{},collapsible:!1,event:"click",header:"> li > :first-child,> :not(li):even",heightStyle:"auto",icons:{activeHeader:"ui-icon-triangle-1-s",header:"ui-icon-triangle-1-e"},activate:null,beforeActivate:null},hideProps:{borderTopWidth:"hide",borderBottomWidth:"hide",paddingTop:"hide",paddingBottom:"hide",height:"hide"},showProps:{borderTopWidth:"show",borderBottomWidth:"show",paddingTop:"show",paddingBottom:"show",height:"show"},_create:function(){var t=this.options;this.prevShow=this.prevHide=e(),this.element.addClass("ui-accordion ui-widget ui-helper-reset").attr("role","tablist"),t.collapsible||t.active!==!1&&null!=t.active||(t.active=0),this._processPanels(),0>t.active&&(t.active+=this.headers.length),this._refresh()},_getCreateEventData:function(){return{header:this.active,panel:this.active.length?this.active.next():e()}},_createIcons:function(){var t=this.options.icons;t&&(e("<span>").addClass("ui-accordion-header-icon ui-icon "+t.header).prependTo(this.headers),this.active.children(".ui-accordion-header-icon").removeClass(t.header).addClass(t.activeHeader),this.headers.addClass("ui-accordion-icons"))},_destroyIcons:function(){this.headers.removeClass("ui-accordion-icons").children(".ui-accordion-header-icon").remove()},_destroy:function(){var e;this.element.removeClass("ui-accordion ui-widget ui-helper-reset").removeAttr("role"),this.headers.removeClass("ui-accordion-header ui-accordion-header-active ui-state-default ui-corner-all ui-state-active ui-state-disabled ui-corner-top").removeAttr("role").removeAttr("aria-expanded").removeAttr("aria-selected").removeAttr("aria-controls").removeAttr("tabIndex").removeUniqueId(),this._destroyIcons(),e=this.headers.next().removeClass("ui-helper-reset ui-widget-content ui-corner-bottom ui-accordion-content ui-accordion-content-active ui-state-disabled").css("display","").removeAttr("role").removeAttr("aria-hidden").removeAttr("aria-labelledby").removeUniqueId(),"content"!==this.options.heightStyle&&e.css("height","")},_setOption:function(e,t){return"active"===e?(this._activate(t),void 0):("event"===e&&(this.options.event&&this._off(this.headers,this.options.event),this._setupEvents(t)),this._super(e,t),"collapsible"!==e||t||this.options.active!==!1||this._activate(0),"icons"===e&&(this._destroyIcons(),t&&this._createIcons()),"disabled"===e&&(this.element.toggleClass("ui-state-disabled",!!t).attr("aria-disabled",t),this.headers.add(this.headers.next()).toggleClass("ui-state-disabled",!!t)),void 0)},_keydown:function(t){if(!t.altKey&&!t.ctrlKey){var i=e.ui.keyCode,s=this.headers.length,n=this.headers.index(t.target),a=!1;switch(t.keyCode){case i.RIGHT:case i.DOWN:a=this.headers[(n+1)%s];break;case i.LEFT:case i.UP:a=this.headers[(n-1+s)%s];break;case i.SPACE:case i.ENTER:this._eventHandler(t);break;case i.HOME:a=this.headers[0];break;case i.END:a=this.headers[s-1]}a&&(e(t.target).attr("tabIndex",-1),e(a).attr("tabIndex",0),a.focus(),t.preventDefault())}},_panelKeyDown:function(t){t.keyCode===e.ui.keyCode.UP&&t.ctrlKey&&e(t.currentTarget).prev().focus()},refresh:function(){var t=this.options;this._processPanels(),t.active===!1&&t.collapsible===!0||!this.headers.length?(t.active=!1,this.active=e()):t.active===!1?this._activate(0):this.active.length&&!e.contains(this.element[0],this.active[0])?this.headers.length===this.headers.find(".ui-state-disabled").length?(t.active=!1,this.active=e()):this._activate(Math.max(0,t.active-1)):t.active=this.headers.index(this.active),this._destroyIcons(),this._refresh()},_processPanels:function(){var e=this.headers,t=this.panels;this.headers=this.element.find(this.options.header).addClass("ui-accordion-header ui-state-default ui-corner-all"),this.panels=this.headers.next().addClass("ui-accordion-content ui-helper-reset ui-widget-content ui-corner-bottom").filter(":not(.ui-accordion-content-active)").hide(),t&&(this._off(e.not(this.headers)),this._off(t.not(this.panels)))},_refresh:function(){var t,i=this.options,s=i.heightStyle,n=this.element.parent();this.active=this._findActive(i.active).addClass("ui-accordion-header-active ui-state-active ui-corner-top").removeClass("ui-corner-all"),this.active.next().addClass("ui-accordion-content-active").show(),this.headers.attr("role","tab").each(function(){var t=e(this),i=t.uniqueId().attr("id"),s=t.next(),n=s.uniqueId().attr("id");t.attr("aria-controls",n),s.attr("aria-labelledby",i)}).next().attr("role","tabpanel"),this.headers.not(this.active).attr({"aria-selected":"false","aria-expanded":"false",tabIndex:-1}).next().attr({"aria-hidden":"true"}).hide(),this.active.length?this.active.attr({"aria-selected":"true","aria-expanded":"true",tabIndex:0}).next().attr({"aria-hidden":"false"}):this.headers.eq(0).attr("tabIndex",0),this._createIcons(),this._setupEvents(i.event),"fill"===s?(t=n.height(),this.element.siblings(":visible").each(function(){var i=e(this),s=i.css("position");"absolute"!==s&&"fixed"!==s&&(t-=i.outerHeight(!0))}),this.headers.each(function(){t-=e(this).outerHeight(!0)}),this.headers.next().each(function(){e(this).height(Math.max(0,t-e(this).innerHeight()+e(this).height()))}).css("overflow","auto")):"auto"===s&&(t=0,this.headers.next().each(function(){t=Math.max(t,e(this).css("height","").height())}).height(t))},_activate:function(t){var i=this._findActive(t)[0];i!==this.active[0]&&(i=i||this.active[0],this._eventHandler({target:i,currentTarget:i,preventDefault:e.noop}))},_findActive:function(t){return"number"==typeof t?this.headers.eq(t):e()},_setupEvents:function(t){var i={keydown:"_keydown"};t&&e.each(t.split(" "),function(e,t){i[t]="_eventHandler"}),this._off(this.headers.add(this.headers.next())),this._on(this.headers,i),this._on(this.headers.next(),{keydown:"_panelKeyDown"}),this._hoverable(this.headers),this._focusable(this.headers)},_eventHandler:function(t){var i=this.options,s=this.active,n=e(t.currentTarget),a=n[0]===s[0],o=a&&i.collapsible,r=o?e():n.next(),h=s.next(),l={oldHeader:s,oldPanel:h,newHeader:o?e():n,newPanel:r};
t.preventDefault(),a&&!i.collapsible||this._trigger("beforeActivate",t,l)===!1||(i.active=o?!1:this.headers.index(n),this.active=a?e():n,this._toggle(l),s.removeClass("ui-accordion-header-active ui-state-active"),i.icons&&s.children(".ui-accordion-header-icon").removeClass(i.icons.activeHeader).addClass(i.icons.header),a||(n.removeClass("ui-corner-all").addClass("ui-accordion-header-active ui-state-active ui-corner-top"),i.icons&&n.children(".ui-accordion-header-icon").removeClass(i.icons.header).addClass(i.icons.activeHeader),n.next().addClass("ui-accordion-content-active")))},_toggle:function(t){var i=t.newPanel,s=this.prevShow.length?this.prevShow:t.oldPanel;this.prevShow.add(this.prevHide).stop(!0,!0),this.prevShow=i,this.prevHide=s,this.options.animate?this._animate(i,s,t):(s.hide(),i.show(),this._toggleComplete(t)),s.attr({"aria-hidden":"true"}),s.prev().attr({"aria-selected":"false","aria-expanded":"false"}),i.length&&s.length?s.prev().attr({tabIndex:-1,"aria-expanded":"false"}):i.length&&this.headers.filter(function(){return 0===parseInt(e(this).attr("tabIndex"),10)}).attr("tabIndex",-1),i.attr("aria-hidden","false").prev().attr({"aria-selected":"true","aria-expanded":"true",tabIndex:0})},_animate:function(e,t,i){var s,n,a,o=this,r=0,h=e.css("box-sizing"),l=e.length&&(!t.length||e.index()<t.index()),u=this.options.animate||{},d=l&&u.down||u,c=function(){o._toggleComplete(i)};return"number"==typeof d&&(a=d),"string"==typeof d&&(n=d),n=n||d.easing||u.easing,a=a||d.duration||u.duration,t.length?e.length?(s=e.show().outerHeight(),t.animate(this.hideProps,{duration:a,easing:n,step:function(e,t){t.now=Math.round(e)}}),e.hide().animate(this.showProps,{duration:a,easing:n,complete:c,step:function(e,i){i.now=Math.round(e),"height"!==i.prop?"content-box"===h&&(r+=i.now):"content"!==o.options.heightStyle&&(i.now=Math.round(s-t.outerHeight()-r),r=0)}}),void 0):t.animate(this.hideProps,a,n,c):e.animate(this.showProps,a,n,c)},_toggleComplete:function(e){var t=e.oldPanel;t.removeClass("ui-accordion-content-active").prev().removeClass("ui-corner-top").addClass("ui-corner-all"),t.length&&(t.parent()[0].className=t.parent()[0].className),this._trigger("activate",null,e)}}),e.widget("ui.menu",{version:"1.11.4",defaultElement:"<ul>",delay:300,options:{icons:{submenu:"ui-icon-carat-1-e"},items:"> *",menus:"ul",position:{my:"left-1 top",at:"right top"},role:"menu",blur:null,focus:null,select:null},_create:function(){this.activeMenu=this.element,this.mouseHandled=!1,this.element.uniqueId().addClass("ui-menu ui-widget ui-widget-content").toggleClass("ui-menu-icons",!!this.element.find(".ui-icon").length).attr({role:this.options.role,tabIndex:0}),this.options.disabled&&this.element.addClass("ui-state-disabled").attr("aria-disabled","true"),this._on({"mousedown .ui-menu-item":function(e){e.preventDefault()},"click .ui-menu-item":function(t){var i=e(t.target);!this.mouseHandled&&i.not(".ui-state-disabled").length&&(this.select(t),t.isPropagationStopped()||(this.mouseHandled=!0),i.has(".ui-menu").length?this.expand(t):!this.element.is(":focus")&&e(this.document[0].activeElement).closest(".ui-menu").length&&(this.element.trigger("focus",[!0]),this.active&&1===this.active.parents(".ui-menu").length&&clearTimeout(this.timer)))},"mouseenter .ui-menu-item":function(t){if(!this.previousFilter){var i=e(t.currentTarget);i.siblings(".ui-state-active").removeClass("ui-state-active"),this.focus(t,i)}},mouseleave:"collapseAll","mouseleave .ui-menu":"collapseAll",focus:function(e,t){var i=this.active||this.element.find(this.options.items).eq(0);t||this.focus(e,i)},blur:function(t){this._delay(function(){e.contains(this.element[0],this.document[0].activeElement)||this.collapseAll(t)})},keydown:"_keydown"}),this.refresh(),this._on(this.document,{click:function(e){this._closeOnDocumentClick(e)&&this.collapseAll(e),this.mouseHandled=!1}})},_destroy:function(){this.element.removeAttr("aria-activedescendant").find(".ui-menu").addBack().removeClass("ui-menu ui-widget ui-widget-content ui-menu-icons ui-front").removeAttr("role").removeAttr("tabIndex").removeAttr("aria-labelledby").removeAttr("aria-expanded").removeAttr("aria-hidden").removeAttr("aria-disabled").removeUniqueId().show(),this.element.find(".ui-menu-item").removeClass("ui-menu-item").removeAttr("role").removeAttr("aria-disabled").removeUniqueId().removeClass("ui-state-hover").removeAttr("tabIndex").removeAttr("role").removeAttr("aria-haspopup").children().each(function(){var t=e(this);t.data("ui-menu-submenu-carat")&&t.remove()}),this.element.find(".ui-menu-divider").removeClass("ui-menu-divider ui-widget-content")},_keydown:function(t){var i,s,n,a,o=!0;switch(t.keyCode){case e.ui.keyCode.PAGE_UP:this.previousPage(t);break;case e.ui.keyCode.PAGE_DOWN:this.nextPage(t);break;case e.ui.keyCode.HOME:this._move("first","first",t);break;case e.ui.keyCode.END:this._move("last","last",t);break;case e.ui.keyCode.UP:this.previous(t);break;case e.ui.keyCode.DOWN:this.next(t);break;case e.ui.keyCode.LEFT:this.collapse(t);break;case e.ui.keyCode.RIGHT:this.active&&!this.active.is(".ui-state-disabled")&&this.expand(t);break;case e.ui.keyCode.ENTER:case e.ui.keyCode.SPACE:this._activate(t);break;case e.ui.keyCode.ESCAPE:this.collapse(t);break;default:o=!1,s=this.previousFilter||"",n=String.fromCharCode(t.keyCode),a=!1,clearTimeout(this.filterTimer),n===s?a=!0:n=s+n,i=this._filterMenuItems(n),i=a&&-1!==i.index(this.active.next())?this.active.nextAll(".ui-menu-item"):i,i.length||(n=String.fromCharCode(t.keyCode),i=this._filterMenuItems(n)),i.length?(this.focus(t,i),this.previousFilter=n,this.filterTimer=this._delay(function(){delete this.previousFilter},1e3)):delete this.previousFilter}o&&t.preventDefault()},_activate:function(e){this.active.is(".ui-state-disabled")||(this.active.is("[aria-haspopup='true']")?this.expand(e):this.select(e))},refresh:function(){var t,i,s=this,n=this.options.icons.submenu,a=this.element.find(this.options.menus);this.element.toggleClass("ui-menu-icons",!!this.element.find(".ui-icon").length),a.filter(":not(.ui-menu)").addClass("ui-menu ui-widget ui-widget-content ui-front").hide().attr({role:this.options.role,"aria-hidden":"true","aria-expanded":"false"}).each(function(){var t=e(this),i=t.parent(),s=e("<span>").addClass("ui-menu-icon ui-icon "+n).data("ui-menu-submenu-carat",!0);i.attr("aria-haspopup","true").prepend(s),t.attr("aria-labelledby",i.attr("id"))}),t=a.add(this.element),i=t.find(this.options.items),i.not(".ui-menu-item").each(function(){var t=e(this);s._isDivider(t)&&t.addClass("ui-widget-content ui-menu-divider")}),i.not(".ui-menu-item, .ui-menu-divider").addClass("ui-menu-item").uniqueId().attr({tabIndex:-1,role:this._itemRole()}),i.filter(".ui-state-disabled").attr("aria-disabled","true"),this.active&&!e.contains(this.element[0],this.active[0])&&this.blur()},_itemRole:function(){return{menu:"menuitem",listbox:"option"}[this.options.role]},_setOption:function(e,t){"icons"===e&&this.element.find(".ui-menu-icon").removeClass(this.options.icons.submenu).addClass(t.submenu),"disabled"===e&&this.element.toggleClass("ui-state-disabled",!!t).attr("aria-disabled",t),this._super(e,t)},focus:function(e,t){var i,s;this.blur(e,e&&"focus"===e.type),this._scrollIntoView(t),this.active=t.first(),s=this.active.addClass("ui-state-focus").removeClass("ui-state-active"),this.options.role&&this.element.attr("aria-activedescendant",s.attr("id")),this.active.parent().closest(".ui-menu-item").addClass("ui-state-active"),e&&"keydown"===e.type?this._close():this.timer=this._delay(function(){this._close()},this.delay),i=t.children(".ui-menu"),i.length&&e&&/^mouse/.test(e.type)&&this._startOpening(i),this.activeMenu=t.parent(),this._trigger("focus",e,{item:t})},_scrollIntoView:function(t){var i,s,n,a,o,r;this._hasScroll()&&(i=parseFloat(e.css(this.activeMenu[0],"borderTopWidth"))||0,s=parseFloat(e.css(this.activeMenu[0],"paddingTop"))||0,n=t.offset().top-this.activeMenu.offset().top-i-s,a=this.activeMenu.scrollTop(),o=this.activeMenu.height(),r=t.outerHeight(),0>n?this.activeMenu.scrollTop(a+n):n+r>o&&this.activeMenu.scrollTop(a+n-o+r))},blur:function(e,t){t||clearTimeout(this.timer),this.active&&(this.active.removeClass("ui-state-focus"),this.active=null,this._trigger("blur",e,{item:this.active}))},_startOpening:function(e){clearTimeout(this.timer),"true"===e.attr("aria-hidden")&&(this.timer=this._delay(function(){this._close(),this._open(e)},this.delay))},_open:function(t){var i=e.extend({of:this.active},this.options.position);clearTimeout(this.timer),this.element.find(".ui-menu").not(t.parents(".ui-menu")).hide().attr("aria-hidden","true"),t.show().removeAttr("aria-hidden").attr("aria-expanded","true").position(i)},collapseAll:function(t,i){clearTimeout(this.timer),this.timer=this._delay(function(){var s=i?this.element:e(t&&t.target).closest(this.element.find(".ui-menu"));s.length||(s=this.element),this._close(s),this.blur(t),this.activeMenu=s},this.delay)},_close:function(e){e||(e=this.active?this.active.parent():this.element),e.find(".ui-menu").hide().attr("aria-hidden","true").attr("aria-expanded","false").end().find(".ui-state-active").not(".ui-state-focus").removeClass("ui-state-active")},_closeOnDocumentClick:function(t){return!e(t.target).closest(".ui-menu").length},_isDivider:function(e){return!/[^\-\u2014\u2013\s]/.test(e.text())},collapse:function(e){var t=this.active&&this.active.parent().closest(".ui-menu-item",this.element);t&&t.length&&(this._close(),this.focus(e,t))},expand:function(e){var t=this.active&&this.active.children(".ui-menu ").find(this.options.items).first();t&&t.length&&(this._open(t.parent()),this._delay(function(){this.focus(e,t)}))},next:function(e){this._move("next","first",e)},previous:function(e){this._move("prev","last",e)},isFirstItem:function(){return this.active&&!this.active.prevAll(".ui-menu-item").length},isLastItem:function(){return this.active&&!this.active.nextAll(".ui-menu-item").length},_move:function(e,t,i){var s;this.active&&(s="first"===e||"last"===e?this.active["first"===e?"prevAll":"nextAll"](".ui-menu-item").eq(-1):this.active[e+"All"](".ui-menu-item").eq(0)),s&&s.length&&this.active||(s=this.activeMenu.find(this.options.items)[t]()),this.focus(i,s)},nextPage:function(t){var i,s,n;return this.active?(this.isLastItem()||(this._hasScroll()?(s=this.active.offset().top,n=this.element.height(),this.active.nextAll(".ui-menu-item").each(function(){return i=e(this),0>i.offset().top-s-n}),this.focus(t,i)):this.focus(t,this.activeMenu.find(this.options.items)[this.active?"last":"first"]())),void 0):(this.next(t),void 0)},previousPage:function(t){var i,s,n;return this.active?(this.isFirstItem()||(this._hasScroll()?(s=this.active.offset().top,n=this.element.height(),this.active.prevAll(".ui-menu-item").each(function(){return i=e(this),i.offset().top-s+n>0}),this.focus(t,i)):this.focus(t,this.activeMenu.find(this.options.items).first())),void 0):(this.next(t),void 0)},_hasScroll:function(){return this.element.outerHeight()<this.element.prop("scrollHeight")},select:function(t){this.active=this.active||e(t.target).closest(".ui-menu-item");var i={item:this.active};this.active.has(".ui-menu").length||this.collapseAll(t,!0),this._trigger("select",t,i)},_filterMenuItems:function(t){var i=t.replace(/[\-\[\]{}()*+?.,\\\^$|#\s]/g,"\\$&"),s=RegExp("^"+i,"i");return this.activeMenu.find(this.options.items).filter(".ui-menu-item").filter(function(){return s.test(e.trim(e(this).text()))})}}),e.widget("ui.autocomplete",{version:"1.11.4",defaultElement:"<input>",options:{appendTo:null,autoFocus:!1,delay:300,minLength:1,position:{my:"left top",at:"left bottom",collision:"none"},source:null,change:null,close:null,focus:null,open:null,response:null,search:null,select:null},requestIndex:0,pending:0,_create:function(){var t,i,s,n=this.element[0].nodeName.toLowerCase(),a="textarea"===n,o="input"===n;this.isMultiLine=a?!0:o?!1:this.element.prop("isContentEditable"),this.valueMethod=this.element[a||o?"val":"text"],this.isNewMenu=!0,this.element.addClass("ui-autocomplete-input").attr("autocomplete","off"),this._on(this.element,{keydown:function(n){if(this.element.prop("readOnly"))return t=!0,s=!0,i=!0,void 0;t=!1,s=!1,i=!1;var a=e.ui.keyCode;switch(n.keyCode){case a.PAGE_UP:t=!0,this._move("previousPage",n);break;case a.PAGE_DOWN:t=!0,this._move("nextPage",n);break;case a.UP:t=!0,this._keyEvent("previous",n);break;case a.DOWN:t=!0,this._keyEvent("next",n);break;case a.ENTER:this.menu.active&&(t=!0,n.preventDefault(),this.menu.select(n));break;case a.TAB:this.menu.active&&this.menu.select(n);break;case a.ESCAPE:this.menu.element.is(":visible")&&(this.isMultiLine||this._value(this.term),this.close(n),n.preventDefault());break;default:i=!0,this._searchTimeout(n)}},keypress:function(s){if(t)return t=!1,(!this.isMultiLine||this.menu.element.is(":visible"))&&s.preventDefault(),void 0;if(!i){var n=e.ui.keyCode;switch(s.keyCode){case n.PAGE_UP:this._move("previousPage",s);break;case n.PAGE_DOWN:this._move("nextPage",s);break;case n.UP:this._keyEvent("previous",s);break;case n.DOWN:this._keyEvent("next",s)}}},input:function(e){return s?(s=!1,e.preventDefault(),void 0):(this._searchTimeout(e),void 0)},focus:function(){this.selectedItem=null,this.previous=this._value()},blur:function(e){return this.cancelBlur?(delete this.cancelBlur,void 0):(clearTimeout(this.searching),this.close(e),this._change(e),void 0)}}),this._initSource(),this.menu=e("<ul>").addClass("ui-autocomplete ui-front").appendTo(this._appendTo()).menu({role:null}).hide().menu("instance"),this._on(this.menu.element,{mousedown:function(t){t.preventDefault(),this.cancelBlur=!0,this._delay(function(){delete this.cancelBlur});var i=this.menu.element[0];e(t.target).closest(".ui-menu-item").length||this._delay(function(){var t=this;this.document.one("mousedown",function(s){s.target===t.element[0]||s.target===i||e.contains(i,s.target)||t.close()})})},menufocus:function(t,i){var s,n;return this.isNewMenu&&(this.isNewMenu=!1,t.originalEvent&&/^mouse/.test(t.originalEvent.type))?(this.menu.blur(),this.document.one("mousemove",function(){e(t.target).trigger(t.originalEvent)}),void 0):(n=i.item.data("ui-autocomplete-item"),!1!==this._trigger("focus",t,{item:n})&&t.originalEvent&&/^key/.test(t.originalEvent.type)&&this._value(n.value),s=i.item.attr("aria-label")||n.value,s&&e.trim(s).length&&(this.liveRegion.children().hide(),e("<div>").text(s).appendTo(this.liveRegion)),void 0)},menuselect:function(e,t){var i=t.item.data("ui-autocomplete-item"),s=this.previous;this.element[0]!==this.document[0].activeElement&&(this.element.focus(),this.previous=s,this._delay(function(){this.previous=s,this.selectedItem=i})),!1!==this._trigger("select",e,{item:i})&&this._value(i.value),this.term=this._value(),this.close(e),this.selectedItem=i}}),this.liveRegion=e("<span>",{role:"status","aria-live":"assertive","aria-relevant":"additions"}).addClass("ui-helper-hidden-accessible").appendTo(this.document[0].body),this._on(this.window,{beforeunload:function(){this.element.removeAttr("autocomplete")}})},_destroy:function(){clearTimeout(this.searching),this.element.removeClass("ui-autocomplete-input").removeAttr("autocomplete"),this.menu.element.remove(),this.liveRegion.remove()},_setOption:function(e,t){this._super(e,t),"source"===e&&this._initSource(),"appendTo"===e&&this.menu.element.appendTo(this._appendTo()),"disabled"===e&&t&&this.xhr&&this.xhr.abort()},_appendTo:function(){var t=this.options.appendTo;return t&&(t=t.jquery||t.nodeType?e(t):this.document.find(t).eq(0)),t&&t[0]||(t=this.element.closest(".ui-front")),t.length||(t=this.document[0].body),t},_initSource:function(){var t,i,s=this;e.isArray(this.options.source)?(t=this.options.source,this.source=function(i,s){s(e.ui.autocomplete.filter(t,i.term))}):"string"==typeof this.options.source?(i=this.options.source,this.source=function(t,n){s.xhr&&s.xhr.abort(),s.xhr=e.ajax({url:i,data:t,dataType:"json",success:function(e){n(e)},error:function(){n([])}})}):this.source=this.options.source},_searchTimeout:function(e){clearTimeout(this.searching),this.searching=this._delay(function(){var t=this.term===this._value(),i=this.menu.element.is(":visible"),s=e.altKey||e.ctrlKey||e.metaKey||e.shiftKey;(!t||t&&!i&&!s)&&(this.selectedItem=null,this.search(null,e))},this.options.delay)},search:function(e,t){return e=null!=e?e:this._value(),this.term=this._value(),e.length<this.options.minLength?this.close(t):this._trigger("search",t)!==!1?this._search(e):void 0},_search:function(e){this.pending++,this.element.addClass("ui-autocomplete-loading"),this.cancelSearch=!1,this.source({term:e},this._response())},_response:function(){var t=++this.requestIndex;return e.proxy(function(e){t===this.requestIndex&&this.__response(e),this.pending--,this.pending||this.element.removeClass("ui-autocomplete-loading")},this)},__response:function(e){e&&(e=this._normalize(e)),this._trigger("response",null,{content:e}),!this.options.disabled&&e&&e.length&&!this.cancelSearch?(this._suggest(e),this._trigger("open")):this._close()},close:function(e){this.cancelSearch=!0,this._close(e)},_close:function(e){this.menu.element.is(":visible")&&(this.menu.element.hide(),this.menu.blur(),this.isNewMenu=!0,this._trigger("close",e))},_change:function(e){this.previous!==this._value()&&this._trigger("change",e,{item:this.selectedItem})},_normalize:function(t){return t.length&&t[0].label&&t[0].value?t:e.map(t,function(t){return"string"==typeof t?{label:t,value:t}:e.extend({},t,{label:t.label||t.value,value:t.value||t.label})})},_suggest:function(t){var i=this.menu.element.empty();this._renderMenu(i,t),this.isNewMenu=!0,this.menu.refresh(),i.show(),this._resizeMenu(),i.position(e.extend({of:this.element},this.options.position)),this.options.autoFocus&&this.menu.next()},_resizeMenu:function(){var e=this.menu.element;e.outerWidth(Math.max(e.width("").outerWidth()+1,this.element.outerWidth()))},_renderMenu:function(t,i){var s=this;e.each(i,function(e,i){s._renderItemData(t,i)})},_renderItemData:function(e,t){return this._renderItem(e,t).data("ui-autocomplete-item",t)},_renderItem:function(t,i){return e("<li>").text(i.label).appendTo(t)},_move:function(e,t){return this.menu.element.is(":visible")?this.menu.isFirstItem()&&/^previous/.test(e)||this.menu.isLastItem()&&/^next/.test(e)?(this.isMultiLine||this._value(this.term),this.menu.blur(),void 0):(this.menu[e](t),void 0):(this.search(null,t),void 0)},widget:function(){return this.menu.element},_value:function(){return this.valueMethod.apply(this.element,arguments)},_keyEvent:function(e,t){(!this.isMultiLine||this.menu.element.is(":visible"))&&(this._move(e,t),t.preventDefault())}}),e.extend(e.ui.autocomplete,{escapeRegex:function(e){return e.replace(/[\-\[\]{}()*+?.,\\\^$|#\s]/g,"\\$&")},filter:function(t,i){var s=RegExp(e.ui.autocomplete.escapeRegex(i),"i");return e.grep(t,function(e){return s.test(e.label||e.value||e)})}}),e.widget("ui.autocomplete",e.ui.autocomplete,{options:{messages:{noResults:"No search results.",results:function(e){return e+(e>1?" results are":" result is")+" available, use up and down arrow keys to navigate."}}},__response:function(t){var i;this._superApply(arguments),this.options.disabled||this.cancelSearch||(i=t&&t.length?this.options.messages.results(t.length):this.options.messages.noResults,this.liveRegion.children().hide(),e("<div>").text(i).appendTo(this.liveRegion))}}),e.ui.autocomplete;var r,h="ui-button ui-widget ui-state-default ui-corner-all",l="ui-button-icons-only ui-button-icon-only ui-button-text-icons ui-button-text-icon-primary ui-button-text-icon-secondary ui-button-text-only",u=function(){var t=e(this);setTimeout(function(){t.find(":ui-button").button("refresh")},1)},d=function(t){var i=t.name,s=t.form,n=e([]);return i&&(i=i.replace(/'/g,"\\'"),n=s?e(s).find("[name='"+i+"'][type=radio]"):e("[name='"+i+"'][type=radio]",t.ownerDocument).filter(function(){return!this.form})),n};e.widget("ui.button",{version:"1.11.4",defaultElement:"<button>",options:{disabled:null,text:!0,label:null,icons:{primary:null,secondary:null}},_create:function(){this.element.closest("form").unbind("reset"+this.eventNamespace).bind("reset"+this.eventNamespace,u),"boolean"!=typeof this.options.disabled?this.options.disabled=!!this.element.prop("disabled"):this.element.prop("disabled",this.options.disabled),this._determineButtonType(),this.hasTitle=!!this.buttonElement.attr("title");var t=this,i=this.options,s="checkbox"===this.type||"radio"===this.type,n=s?"":"ui-state-active";null===i.label&&(i.label="input"===this.type?this.buttonElement.val():this.buttonElement.html()),this._hoverable(this.buttonElement),this.buttonElement.addClass(h).attr("role","button").bind("mouseenter"+this.eventNamespace,function(){i.disabled||this===r&&e(this).addClass("ui-state-active")}).bind("mouseleave"+this.eventNamespace,function(){i.disabled||e(this).removeClass(n)}).bind("click"+this.eventNamespace,function(e){i.disabled&&(e.preventDefault(),e.stopImmediatePropagation())}),this._on({focus:function(){this.buttonElement.addClass("ui-state-focus")},blur:function(){this.buttonElement.removeClass("ui-state-focus")}}),s&&this.element.bind("change"+this.eventNamespace,function(){t.refresh()}),"checkbox"===this.type?this.buttonElement.bind("click"+this.eventNamespace,function(){return i.disabled?!1:void 0}):"radio"===this.type?this.buttonElement.bind("click"+this.eventNamespace,function(){if(i.disabled)return!1;e(this).addClass("ui-state-active"),t.buttonElement.attr("aria-pressed","true");var s=t.element[0];d(s).not(s).map(function(){return e(this).button("widget")[0]}).removeClass("ui-state-active").attr("aria-pressed","false")}):(this.buttonElement.bind("mousedown"+this.eventNamespace,function(){return i.disabled?!1:(e(this).addClass("ui-state-active"),r=this,t.document.one("mouseup",function(){r=null}),void 0)}).bind("mouseup"+this.eventNamespace,function(){return i.disabled?!1:(e(this).removeClass("ui-state-active"),void 0)}).bind("keydown"+this.eventNamespace,function(t){return i.disabled?!1:((t.keyCode===e.ui.keyCode.SPACE||t.keyCode===e.ui.keyCode.ENTER)&&e(this).addClass("ui-state-active"),void 0)}).bind("keyup"+this.eventNamespace+" blur"+this.eventNamespace,function(){e(this).removeClass("ui-state-active")}),this.buttonElement.is("a")&&this.buttonElement.keyup(function(t){t.keyCode===e.ui.keyCode.SPACE&&e(this).click()})),this._setOption("disabled",i.disabled),this._resetButton()},_determineButtonType:function(){var e,t,i;this.type=this.element.is("[type=checkbox]")?"checkbox":this.element.is("[type=radio]")?"radio":this.element.is("input")?"input":"button","checkbox"===this.type||"radio"===this.type?(e=this.element.parents().last(),t="label[for='"+this.element.attr("id")+"']",this.buttonElement=e.find(t),this.buttonElement.length||(e=e.length?e.siblings():this.element.siblings(),this.buttonElement=e.filter(t),this.buttonElement.length||(this.buttonElement=e.find(t))),this.element.addClass("ui-helper-hidden-accessible"),i=this.element.is(":checked"),i&&this.buttonElement.addClass("ui-state-active"),this.buttonElement.prop("aria-pressed",i)):this.buttonElement=this.element},widget:function(){return this.buttonElement},_destroy:function(){this.element.removeClass("ui-helper-hidden-accessible"),this.buttonElement.removeClass(h+" ui-state-active "+l).removeAttr("role").removeAttr("aria-pressed").html(this.buttonElement.find(".ui-button-text").html()),this.hasTitle||this.buttonElement.removeAttr("title")},_setOption:function(e,t){return this._super(e,t),"disabled"===e?(this.widget().toggleClass("ui-state-disabled",!!t),this.element.prop("disabled",!!t),t&&("checkbox"===this.type||"radio"===this.type?this.buttonElement.removeClass("ui-state-focus"):this.buttonElement.removeClass("ui-state-focus ui-state-active")),void 0):(this._resetButton(),void 0)},refresh:function(){var t=this.element.is("input, button")?this.element.is(":disabled"):this.element.hasClass("ui-button-disabled");t!==this.options.disabled&&this._setOption("disabled",t),"radio"===this.type?d(this.element[0]).each(function(){e(this).is(":checked")?e(this).button("widget").addClass("ui-state-active").attr("aria-pressed","true"):e(this).button("widget").removeClass("ui-state-active").attr("aria-pressed","false")}):"checkbox"===this.type&&(this.element.is(":checked")?this.buttonElement.addClass("ui-state-active").attr("aria-pressed","true"):this.buttonElement.removeClass("ui-state-active").attr("aria-pressed","false"))},_resetButton:function(){if("input"===this.type)return this.options.label&&this.element.val(this.options.label),void 0;var t=this.buttonElement.removeClass(l),i=e("<span></span>",this.document[0]).addClass("ui-button-text").html(this.options.label).appendTo(t.empty()).text(),s=this.options.icons,n=s.primary&&s.secondary,a=[];s.primary||s.secondary?(this.options.text&&a.push("ui-button-text-icon"+(n?"s":s.primary?"-primary":"-secondary")),s.primary&&t.prepend("<span class='ui-button-icon-primary ui-icon "+s.primary+"'></span>"),s.secondary&&t.append("<span class='ui-button-icon-secondary ui-icon "+s.secondary+"'></span>"),this.options.text||(a.push(n?"ui-button-icons-only":"ui-button-icon-only"),this.hasTitle||t.attr("title",e.trim(i)))):a.push("ui-button-text-only"),t.addClass(a.join(" "))}}),e.widget("ui.buttonset",{version:"1.11.4",options:{items:"button, input[type=button], input[type=submit], input[type=reset], input[type=checkbox], input[type=radio], a, :data(ui-button)"},_create:function(){this.element.addClass("ui-buttonset")},_init:function(){this.refresh()},_setOption:function(e,t){"disabled"===e&&this.buttons.button("option",e,t),this._super(e,t)},refresh:function(){var t="rtl"===this.element.css("direction"),i=this.element.find(this.options.items),s=i.filter(":ui-button");i.not(":ui-button").button(),s.button("refresh"),this.buttons=i.map(function(){return e(this).button("widget")[0]}).removeClass("ui-corner-all ui-corner-left ui-corner-right").filter(":first").addClass(t?"ui-corner-right":"ui-corner-left").end().filter(":last").addClass(t?"ui-corner-left":"ui-corner-right").end().end()},_destroy:function(){this.element.removeClass("ui-buttonset"),this.buttons.map(function(){return e(this).button("widget")[0]}).removeClass("ui-corner-left ui-corner-right").end().button("destroy")}}),e.ui.button,e.widget("ui.dialog",{version:"1.11.4",options:{appendTo:"body",autoOpen:!0,buttons:[],closeOnEscape:!0,closeText:"Close",dialogClass:"",draggable:!0,hide:null,height:"auto",maxHeight:null,maxWidth:null,minHeight:150,minWidth:150,modal:!1,position:{my:"center",at:"center",of:window,collision:"fit",using:function(t){var i=e(this).css(t).offset().top;0>i&&e(this).css("top",t.top-i)}},resizable:!0,show:null,title:null,width:300,beforeClose:null,close:null,drag:null,dragStart:null,dragStop:null,focus:null,open:null,resize:null,resizeStart:null,resizeStop:null},sizeRelatedOptions:{buttons:!0,height:!0,maxHeight:!0,maxWidth:!0,minHeight:!0,minWidth:!0,width:!0},resizableRelatedOptions:{maxHeight:!0,maxWidth:!0,minHeight:!0,minWidth:!0},_create:function(){this.originalCss={display:this.element[0].style.display,width:this.element[0].style.width,minHeight:this.element[0].style.minHeight,maxHeight:this.element[0].style.maxHeight,height:this.element[0].style.height},this.originalPosition={parent:this.element.parent(),index:this.element.parent().children().index(this.element)},this.originalTitle=this.element.attr("title"),this.options.title=this.options.title||this.originalTitle,this._createWrapper(),this.element.show().removeAttr("title").addClass("ui-dialog-content ui-widget-content").appendTo(this.uiDialog),this._createTitlebar(),this._createButtonPane(),this.options.draggable&&e.fn.draggable&&this._makeDraggable(),this.options.resizable&&e.fn.resizable&&this._makeResizable(),this._isOpen=!1,this._trackFocus()},_init:function(){this.options.autoOpen&&this.open()},_appendTo:function(){var t=this.options.appendTo;return t&&(t.jquery||t.nodeType)?e(t):this.document.find(t||"body").eq(0)},_destroy:function(){var e,t=this.originalPosition;this._untrackInstance(),this._destroyOverlay(),this.element.removeUniqueId().removeClass("ui-dialog-content ui-widget-content").css(this.originalCss).detach(),this.uiDialog.stop(!0,!0).remove(),this.originalTitle&&this.element.attr("title",this.originalTitle),e=t.parent.children().eq(t.index),e.length&&e[0]!==this.element[0]?e.before(this.element):t.parent.append(this.element)},widget:function(){return this.uiDialog},disable:e.noop,enable:e.noop,close:function(t){var i,s=this;if(this._isOpen&&this._trigger("beforeClose",t)!==!1){if(this._isOpen=!1,this._focusedElement=null,this._destroyOverlay(),this._untrackInstance(),!this.opener.filter(":focusable").focus().length)try{i=this.document[0].activeElement,i&&"body"!==i.nodeName.toLowerCase()&&e(i).blur()}catch(n){}this._hide(this.uiDialog,this.options.hide,function(){s._trigger("close",t)})}},isOpen:function(){return this._isOpen},moveToTop:function(){this._moveToTop()},_moveToTop:function(t,i){var s=!1,n=this.uiDialog.siblings(".ui-front:visible").map(function(){return+e(this).css("z-index")}).get(),a=Math.max.apply(null,n);return a>=+this.uiDialog.css("z-index")&&(this.uiDialog.css("z-index",a+1),s=!0),s&&!i&&this._trigger("focus",t),s},open:function(){var t=this;return this._isOpen?(this._moveToTop()&&this._focusTabbable(),void 0):(this._isOpen=!0,this.opener=e(this.document[0].activeElement),this._size(),this._position(),this._createOverlay(),this._moveToTop(null,!0),this.overlay&&this.overlay.css("z-index",this.uiDialog.css("z-index")-1),this._show(this.uiDialog,this.options.show,function(){t._focusTabbable(),t._trigger("focus")}),this._makeFocusTarget(),this._trigger("open"),void 0)},_focusTabbable:function(){var e=this._focusedElement;e||(e=this.element.find("[autofocus]")),e.length||(e=this.element.find(":tabbable")),e.length||(e=this.uiDialogButtonPane.find(":tabbable")),e.length||(e=this.uiDialogTitlebarClose.filter(":tabbable")),e.length||(e=this.uiDialog),e.eq(0).focus()},_keepFocus:function(t){function i(){var t=this.document[0].activeElement,i=this.uiDialog[0]===t||e.contains(this.uiDialog[0],t);i||this._focusTabbable()}t.preventDefault(),i.call(this),this._delay(i)},_createWrapper:function(){this.uiDialog=e("<div>").addClass("ui-dialog ui-widget ui-widget-content ui-corner-all ui-front "+this.options.dialogClass).hide().attr({tabIndex:-1,role:"dialog"}).appendTo(this._appendTo()),this._on(this.uiDialog,{keydown:function(t){if(this.options.closeOnEscape&&!t.isDefaultPrevented()&&t.keyCode&&t.keyCode===e.ui.keyCode.ESCAPE)return t.preventDefault(),this.close(t),void 0;if(t.keyCode===e.ui.keyCode.TAB&&!t.isDefaultPrevented()){var i=this.uiDialog.find(":tabbable"),s=i.filter(":first"),n=i.filter(":last");t.target!==n[0]&&t.target!==this.uiDialog[0]||t.shiftKey?t.target!==s[0]&&t.target!==this.uiDialog[0]||!t.shiftKey||(this._delay(function(){n.focus()}),t.preventDefault()):(this._delay(function(){s.focus()}),t.preventDefault())}},mousedown:function(e){this._moveToTop(e)&&this._focusTabbable()}}),this.element.find("[aria-describedby]").length||this.uiDialog.attr({"aria-describedby":this.element.uniqueId().attr("id")})},_createTitlebar:function(){var t;this.uiDialogTitlebar=e("<div>").addClass("ui-dialog-titlebar ui-widget-header ui-corner-all ui-helper-clearfix").prependTo(this.uiDialog),this._on(this.uiDialogTitlebar,{mousedown:function(t){e(t.target).closest(".ui-dialog-titlebar-close")||this.uiDialog.focus()}}),this.uiDialogTitlebarClose=e("<button type='button'></button>").button({label:this.options.closeText,icons:{primary:"ui-icon-closethick"},text:!1}).addClass("ui-dialog-titlebar-close").appendTo(this.uiDialogTitlebar),this._on(this.uiDialogTitlebarClose,{click:function(e){e.preventDefault(),this.close(e)}}),t=e("<span>").uniqueId().addClass("ui-dialog-title").prependTo(this.uiDialogTitlebar),this._title(t),this.uiDialog.attr({"aria-labelledby":t.attr("id")})},_title:function(e){this.options.title||e.html("&#160;"),e.text(this.options.title)
},_createButtonPane:function(){this.uiDialogButtonPane=e("<div>").addClass("ui-dialog-buttonpane ui-widget-content ui-helper-clearfix"),this.uiButtonSet=e("<div>").addClass("ui-dialog-buttonset").appendTo(this.uiDialogButtonPane),this._createButtons()},_createButtons:function(){var t=this,i=this.options.buttons;return this.uiDialogButtonPane.remove(),this.uiButtonSet.empty(),e.isEmptyObject(i)||e.isArray(i)&&!i.length?(this.uiDialog.removeClass("ui-dialog-buttons"),void 0):(e.each(i,function(i,s){var n,a;s=e.isFunction(s)?{click:s,text:i}:s,s=e.extend({type:"button"},s),n=s.click,s.click=function(){n.apply(t.element[0],arguments)},a={icons:s.icons,text:s.showText},delete s.icons,delete s.showText,e("<button></button>",s).button(a).appendTo(t.uiButtonSet)}),this.uiDialog.addClass("ui-dialog-buttons"),this.uiDialogButtonPane.appendTo(this.uiDialog),void 0)},_makeDraggable:function(){function t(e){return{position:e.position,offset:e.offset}}var i=this,s=this.options;this.uiDialog.draggable({cancel:".ui-dialog-content, .ui-dialog-titlebar-close",handle:".ui-dialog-titlebar",containment:"document",start:function(s,n){e(this).addClass("ui-dialog-dragging"),i._blockFrames(),i._trigger("dragStart",s,t(n))},drag:function(e,s){i._trigger("drag",e,t(s))},stop:function(n,a){var o=a.offset.left-i.document.scrollLeft(),r=a.offset.top-i.document.scrollTop();s.position={my:"left top",at:"left"+(o>=0?"+":"")+o+" "+"top"+(r>=0?"+":"")+r,of:i.window},e(this).removeClass("ui-dialog-dragging"),i._unblockFrames(),i._trigger("dragStop",n,t(a))}})},_makeResizable:function(){function t(e){return{originalPosition:e.originalPosition,originalSize:e.originalSize,position:e.position,size:e.size}}var i=this,s=this.options,n=s.resizable,a=this.uiDialog.css("position"),o="string"==typeof n?n:"n,e,s,w,se,sw,ne,nw";this.uiDialog.resizable({cancel:".ui-dialog-content",containment:"document",alsoResize:this.element,maxWidth:s.maxWidth,maxHeight:s.maxHeight,minWidth:s.minWidth,minHeight:this._minHeight(),handles:o,start:function(s,n){e(this).addClass("ui-dialog-resizing"),i._blockFrames(),i._trigger("resizeStart",s,t(n))},resize:function(e,s){i._trigger("resize",e,t(s))},stop:function(n,a){var o=i.uiDialog.offset(),r=o.left-i.document.scrollLeft(),h=o.top-i.document.scrollTop();s.height=i.uiDialog.height(),s.width=i.uiDialog.width(),s.position={my:"left top",at:"left"+(r>=0?"+":"")+r+" "+"top"+(h>=0?"+":"")+h,of:i.window},e(this).removeClass("ui-dialog-resizing"),i._unblockFrames(),i._trigger("resizeStop",n,t(a))}}).css("position",a)},_trackFocus:function(){this._on(this.widget(),{focusin:function(t){this._makeFocusTarget(),this._focusedElement=e(t.target)}})},_makeFocusTarget:function(){this._untrackInstance(),this._trackingInstances().unshift(this)},_untrackInstance:function(){var t=this._trackingInstances(),i=e.inArray(this,t);-1!==i&&t.splice(i,1)},_trackingInstances:function(){var e=this.document.data("ui-dialog-instances");return e||(e=[],this.document.data("ui-dialog-instances",e)),e},_minHeight:function(){var e=this.options;return"auto"===e.height?e.minHeight:Math.min(e.minHeight,e.height)},_position:function(){var e=this.uiDialog.is(":visible");e||this.uiDialog.show(),this.uiDialog.position(this.options.position),e||this.uiDialog.hide()},_setOptions:function(t){var i=this,s=!1,n={};e.each(t,function(e,t){i._setOption(e,t),e in i.sizeRelatedOptions&&(s=!0),e in i.resizableRelatedOptions&&(n[e]=t)}),s&&(this._size(),this._position()),this.uiDialog.is(":data(ui-resizable)")&&this.uiDialog.resizable("option",n)},_setOption:function(e,t){var i,s,n=this.uiDialog;"dialogClass"===e&&n.removeClass(this.options.dialogClass).addClass(t),"disabled"!==e&&(this._super(e,t),"appendTo"===e&&this.uiDialog.appendTo(this._appendTo()),"buttons"===e&&this._createButtons(),"closeText"===e&&this.uiDialogTitlebarClose.button({label:""+t}),"draggable"===e&&(i=n.is(":data(ui-draggable)"),i&&!t&&n.draggable("destroy"),!i&&t&&this._makeDraggable()),"position"===e&&this._position(),"resizable"===e&&(s=n.is(":data(ui-resizable)"),s&&!t&&n.resizable("destroy"),s&&"string"==typeof t&&n.resizable("option","handles",t),s||t===!1||this._makeResizable()),"title"===e&&this._title(this.uiDialogTitlebar.find(".ui-dialog-title")))},_size:function(){var e,t,i,s=this.options;this.element.show().css({width:"auto",minHeight:0,maxHeight:"none",height:0}),s.minWidth>s.width&&(s.width=s.minWidth),e=this.uiDialog.css({height:"auto",width:s.width}).outerHeight(),t=Math.max(0,s.minHeight-e),i="number"==typeof s.maxHeight?Math.max(0,s.maxHeight-e):"none","auto"===s.height?this.element.css({minHeight:t,maxHeight:i,height:"auto"}):this.element.height(Math.max(0,s.height-e)),this.uiDialog.is(":data(ui-resizable)")&&this.uiDialog.resizable("option","minHeight",this._minHeight())},_blockFrames:function(){this.iframeBlocks=this.document.find("iframe").map(function(){var t=e(this);return e("<div>").css({position:"absolute",width:t.outerWidth(),height:t.outerHeight()}).appendTo(t.parent()).offset(t.offset())[0]})},_unblockFrames:function(){this.iframeBlocks&&(this.iframeBlocks.remove(),delete this.iframeBlocks)},_allowInteraction:function(t){return e(t.target).closest(".ui-dialog").length?!0:!!e(t.target).closest(".ui-datepicker").length},_createOverlay:function(){if(this.options.modal){var t=!0;this._delay(function(){t=!1}),this.document.data("ui-dialog-overlays")||this._on(this.document,{focusin:function(e){t||this._allowInteraction(e)||(e.preventDefault(),this._trackingInstances()[0]._focusTabbable())}}),this.overlay=e("<div>").addClass("ui-widget-overlay ui-front").appendTo(this._appendTo()),this._on(this.overlay,{mousedown:"_keepFocus"}),this.document.data("ui-dialog-overlays",(this.document.data("ui-dialog-overlays")||0)+1)}},_destroyOverlay:function(){if(this.options.modal&&this.overlay){var e=this.document.data("ui-dialog-overlays")-1;e?this.document.data("ui-dialog-overlays",e):this.document.unbind("focusin").removeData("ui-dialog-overlays"),this.overlay.remove(),this.overlay=null}}}),e.widget("ui.progressbar",{version:"1.11.4",options:{max:100,value:0,change:null,complete:null},min:0,_create:function(){this.oldValue=this.options.value=this._constrainedValue(),this.element.addClass("ui-progressbar ui-widget ui-widget-content ui-corner-all").attr({role:"progressbar","aria-valuemin":this.min}),this.valueDiv=e("<div class='ui-progressbar-value ui-widget-header ui-corner-left'></div>").appendTo(this.element),this._refreshValue()},_destroy:function(){this.element.removeClass("ui-progressbar ui-widget ui-widget-content ui-corner-all").removeAttr("role").removeAttr("aria-valuemin").removeAttr("aria-valuemax").removeAttr("aria-valuenow"),this.valueDiv.remove()},value:function(e){return void 0===e?this.options.value:(this.options.value=this._constrainedValue(e),this._refreshValue(),void 0)},_constrainedValue:function(e){return void 0===e&&(e=this.options.value),this.indeterminate=e===!1,"number"!=typeof e&&(e=0),this.indeterminate?!1:Math.min(this.options.max,Math.max(this.min,e))},_setOptions:function(e){var t=e.value;delete e.value,this._super(e),this.options.value=this._constrainedValue(t),this._refreshValue()},_setOption:function(e,t){"max"===e&&(t=Math.max(this.min,t)),"disabled"===e&&this.element.toggleClass("ui-state-disabled",!!t).attr("aria-disabled",t),this._super(e,t)},_percentage:function(){return this.indeterminate?100:100*(this.options.value-this.min)/(this.options.max-this.min)},_refreshValue:function(){var t=this.options.value,i=this._percentage();this.valueDiv.toggle(this.indeterminate||t>this.min).toggleClass("ui-corner-right",t===this.options.max).width(i.toFixed(0)+"%"),this.element.toggleClass("ui-progressbar-indeterminate",this.indeterminate),this.indeterminate?(this.element.removeAttr("aria-valuenow"),this.overlayDiv||(this.overlayDiv=e("<div class='ui-progressbar-overlay'></div>").appendTo(this.valueDiv))):(this.element.attr({"aria-valuemax":this.options.max,"aria-valuenow":t}),this.overlayDiv&&(this.overlayDiv.remove(),this.overlayDiv=null)),this.oldValue!==t&&(this.oldValue=t,this._trigger("change")),t===this.options.max&&this._trigger("complete")}}),e.widget("ui.selectmenu",{version:"1.11.4",defaultElement:"<select>",options:{appendTo:null,disabled:null,icons:{button:"ui-icon-triangle-1-s"},position:{my:"left top",at:"left bottom",collision:"none"},width:null,change:null,close:null,focus:null,open:null,select:null},_create:function(){var e=this.element.uniqueId().attr("id");this.ids={element:e,button:e+"-button",menu:e+"-menu"},this._drawButton(),this._drawMenu(),this.options.disabled&&this.disable()},_drawButton:function(){var t=this;this.label=e("label[for='"+this.ids.element+"']").attr("for",this.ids.button),this._on(this.label,{click:function(e){this.button.focus(),e.preventDefault()}}),this.element.hide(),this.button=e("<span>",{"class":"ui-selectmenu-button ui-widget ui-state-default ui-corner-all",tabindex:this.options.disabled?-1:0,id:this.ids.button,role:"combobox","aria-expanded":"false","aria-autocomplete":"list","aria-owns":this.ids.menu,"aria-haspopup":"true"}).insertAfter(this.element),e("<span>",{"class":"ui-icon "+this.options.icons.button}).prependTo(this.button),this.buttonText=e("<span>",{"class":"ui-selectmenu-text"}).appendTo(this.button),this._setText(this.buttonText,this.element.find("option:selected").text()),this._resizeButton(),this._on(this.button,this._buttonEvents),this.button.one("focusin",function(){t.menuItems||t._refreshMenu()}),this._hoverable(this.button),this._focusable(this.button)},_drawMenu:function(){var t=this;this.menu=e("<ul>",{"aria-hidden":"true","aria-labelledby":this.ids.button,id:this.ids.menu}),this.menuWrap=e("<div>",{"class":"ui-selectmenu-menu ui-front"}).append(this.menu).appendTo(this._appendTo()),this.menuInstance=this.menu.menu({role:"listbox",select:function(e,i){e.preventDefault(),t._setSelection(),t._select(i.item.data("ui-selectmenu-item"),e)},focus:function(e,i){var s=i.item.data("ui-selectmenu-item");null!=t.focusIndex&&s.index!==t.focusIndex&&(t._trigger("focus",e,{item:s}),t.isOpen||t._select(s,e)),t.focusIndex=s.index,t.button.attr("aria-activedescendant",t.menuItems.eq(s.index).attr("id"))}}).menu("instance"),this.menu.addClass("ui-corner-bottom").removeClass("ui-corner-all"),this.menuInstance._off(this.menu,"mouseleave"),this.menuInstance._closeOnDocumentClick=function(){return!1},this.menuInstance._isDivider=function(){return!1}},refresh:function(){this._refreshMenu(),this._setText(this.buttonText,this._getSelectedItem().text()),this.options.width||this._resizeButton()},_refreshMenu:function(){this.menu.empty();var e,t=this.element.find("option");t.length&&(this._parseOptions(t),this._renderMenu(this.menu,this.items),this.menuInstance.refresh(),this.menuItems=this.menu.find("li").not(".ui-selectmenu-optgroup"),e=this._getSelectedItem(),this.menuInstance.focus(null,e),this._setAria(e.data("ui-selectmenu-item")),this._setOption("disabled",this.element.prop("disabled")))},open:function(e){this.options.disabled||(this.menuItems?(this.menu.find(".ui-state-focus").removeClass("ui-state-focus"),this.menuInstance.focus(null,this._getSelectedItem())):this._refreshMenu(),this.isOpen=!0,this._toggleAttr(),this._resizeMenu(),this._position(),this._on(this.document,this._documentClick),this._trigger("open",e))},_position:function(){this.menuWrap.position(e.extend({of:this.button},this.options.position))},close:function(e){this.isOpen&&(this.isOpen=!1,this._toggleAttr(),this.range=null,this._off(this.document),this._trigger("close",e))},widget:function(){return this.button},menuWidget:function(){return this.menu},_renderMenu:function(t,i){var s=this,n="";e.each(i,function(i,a){a.optgroup!==n&&(e("<li>",{"class":"ui-selectmenu-optgroup ui-menu-divider"+(a.element.parent("optgroup").prop("disabled")?" ui-state-disabled":""),text:a.optgroup}).appendTo(t),n=a.optgroup),s._renderItemData(t,a)})},_renderItemData:function(e,t){return this._renderItem(e,t).data("ui-selectmenu-item",t)},_renderItem:function(t,i){var s=e("<li>");return i.disabled&&s.addClass("ui-state-disabled"),this._setText(s,i.label),s.appendTo(t)},_setText:function(e,t){t?e.text(t):e.html("&#160;")},_move:function(e,t){var i,s,n=".ui-menu-item";this.isOpen?i=this.menuItems.eq(this.focusIndex):(i=this.menuItems.eq(this.element[0].selectedIndex),n+=":not(.ui-state-disabled)"),s="first"===e||"last"===e?i["first"===e?"prevAll":"nextAll"](n).eq(-1):i[e+"All"](n).eq(0),s.length&&this.menuInstance.focus(t,s)},_getSelectedItem:function(){return this.menuItems.eq(this.element[0].selectedIndex)},_toggle:function(e){this[this.isOpen?"close":"open"](e)},_setSelection:function(){var e;this.range&&(window.getSelection?(e=window.getSelection(),e.removeAllRanges(),e.addRange(this.range)):this.range.select(),this.button.focus())},_documentClick:{mousedown:function(t){this.isOpen&&(e(t.target).closest(".ui-selectmenu-menu, #"+this.ids.button).length||this.close(t))}},_buttonEvents:{mousedown:function(){var e;window.getSelection?(e=window.getSelection(),e.rangeCount&&(this.range=e.getRangeAt(0))):this.range=document.selection.createRange()},click:function(e){this._setSelection(),this._toggle(e)},keydown:function(t){var i=!0;switch(t.keyCode){case e.ui.keyCode.TAB:case e.ui.keyCode.ESCAPE:this.close(t),i=!1;break;case e.ui.keyCode.ENTER:this.isOpen&&this._selectFocusedItem(t);break;case e.ui.keyCode.UP:t.altKey?this._toggle(t):this._move("prev",t);break;case e.ui.keyCode.DOWN:t.altKey?this._toggle(t):this._move("next",t);break;case e.ui.keyCode.SPACE:this.isOpen?this._selectFocusedItem(t):this._toggle(t);break;case e.ui.keyCode.LEFT:this._move("prev",t);break;case e.ui.keyCode.RIGHT:this._move("next",t);break;case e.ui.keyCode.HOME:case e.ui.keyCode.PAGE_UP:this._move("first",t);break;case e.ui.keyCode.END:case e.ui.keyCode.PAGE_DOWN:this._move("last",t);break;default:this.menu.trigger(t),i=!1}i&&t.preventDefault()}},_selectFocusedItem:function(e){var t=this.menuItems.eq(this.focusIndex);t.hasClass("ui-state-disabled")||this._select(t.data("ui-selectmenu-item"),e)},_select:function(e,t){var i=this.element[0].selectedIndex;this.element[0].selectedIndex=e.index,this._setText(this.buttonText,e.label),this._setAria(e),this._trigger("select",t,{item:e}),e.index!==i&&this._trigger("change",t,{item:e}),this.close(t)},_setAria:function(e){var t=this.menuItems.eq(e.index).attr("id");this.button.attr({"aria-labelledby":t,"aria-activedescendant":t}),this.menu.attr("aria-activedescendant",t)},_setOption:function(e,t){"icons"===e&&this.button.find("span.ui-icon").removeClass(this.options.icons.button).addClass(t.button),this._super(e,t),"appendTo"===e&&this.menuWrap.appendTo(this._appendTo()),"disabled"===e&&(this.menuInstance.option("disabled",t),this.button.toggleClass("ui-state-disabled",t).attr("aria-disabled",t),this.element.prop("disabled",t),t?(this.button.attr("tabindex",-1),this.close()):this.button.attr("tabindex",0)),"width"===e&&this._resizeButton()},_appendTo:function(){var t=this.options.appendTo;return t&&(t=t.jquery||t.nodeType?e(t):this.document.find(t).eq(0)),t&&t[0]||(t=this.element.closest(".ui-front")),t.length||(t=this.document[0].body),t},_toggleAttr:function(){this.button.toggleClass("ui-corner-top",this.isOpen).toggleClass("ui-corner-all",!this.isOpen).attr("aria-expanded",this.isOpen),this.menuWrap.toggleClass("ui-selectmenu-open",this.isOpen),this.menu.attr("aria-hidden",!this.isOpen)},_resizeButton:function(){var e=this.options.width;e||(e=this.element.show().outerWidth(),this.element.hide()),this.button.outerWidth(e)},_resizeMenu:function(){this.menu.outerWidth(Math.max(this.button.outerWidth(),this.menu.width("").outerWidth()+1))},_getCreateOptions:function(){return{disabled:this.element.prop("disabled")}},_parseOptions:function(t){var i=[];t.each(function(t,s){var n=e(s),a=n.parent("optgroup");i.push({element:n,index:t,value:n.val(),label:n.text(),optgroup:a.attr("label")||"",disabled:a.prop("disabled")||n.prop("disabled")})}),this.items=i},_destroy:function(){this.menuWrap.remove(),this.button.remove(),this.element.show(),this.element.removeUniqueId(),this.label.attr("for",this.ids.element)}}),e.widget("ui.slider",e.ui.mouse,{version:"1.11.4",widgetEventPrefix:"slide",options:{animate:!1,distance:0,max:100,min:0,orientation:"horizontal",range:!1,step:1,value:0,values:null,change:null,slide:null,start:null,stop:null},numPages:5,_create:function(){this._keySliding=!1,this._mouseSliding=!1,this._animateOff=!0,this._handleIndex=null,this._detectOrientation(),this._mouseInit(),this._calculateNewMax(),this.element.addClass("ui-slider ui-slider-"+this.orientation+" ui-widget"+" ui-widget-content"+" ui-corner-all"),this._refresh(),this._setOption("disabled",this.options.disabled),this._animateOff=!1},_refresh:function(){this._createRange(),this._createHandles(),this._setupEvents(),this._refreshValue()},_createHandles:function(){var t,i,s=this.options,n=this.element.find(".ui-slider-handle").addClass("ui-state-default ui-corner-all"),a="<span class='ui-slider-handle ui-state-default ui-corner-all' tabindex='0'></span>",o=[];for(i=s.values&&s.values.length||1,n.length>i&&(n.slice(i).remove(),n=n.slice(0,i)),t=n.length;i>t;t++)o.push(a);this.handles=n.add(e(o.join("")).appendTo(this.element)),this.handle=this.handles.eq(0),this.handles.each(function(t){e(this).data("ui-slider-handle-index",t)})},_createRange:function(){var t=this.options,i="";t.range?(t.range===!0&&(t.values?t.values.length&&2!==t.values.length?t.values=[t.values[0],t.values[0]]:e.isArray(t.values)&&(t.values=t.values.slice(0)):t.values=[this._valueMin(),this._valueMin()]),this.range&&this.range.length?this.range.removeClass("ui-slider-range-min ui-slider-range-max").css({left:"",bottom:""}):(this.range=e("<div></div>").appendTo(this.element),i="ui-slider-range ui-widget-header ui-corner-all"),this.range.addClass(i+("min"===t.range||"max"===t.range?" ui-slider-range-"+t.range:""))):(this.range&&this.range.remove(),this.range=null)},_setupEvents:function(){this._off(this.handles),this._on(this.handles,this._handleEvents),this._hoverable(this.handles),this._focusable(this.handles)},_destroy:function(){this.handles.remove(),this.range&&this.range.remove(),this.element.removeClass("ui-slider ui-slider-horizontal ui-slider-vertical ui-widget ui-widget-content ui-corner-all"),this._mouseDestroy()},_mouseCapture:function(t){var i,s,n,a,o,r,h,l,u=this,d=this.options;return d.disabled?!1:(this.elementSize={width:this.element.outerWidth(),height:this.element.outerHeight()},this.elementOffset=this.element.offset(),i={x:t.pageX,y:t.pageY},s=this._normValueFromMouse(i),n=this._valueMax()-this._valueMin()+1,this.handles.each(function(t){var i=Math.abs(s-u.values(t));(n>i||n===i&&(t===u._lastChangedValue||u.values(t)===d.min))&&(n=i,a=e(this),o=t)}),r=this._start(t,o),r===!1?!1:(this._mouseSliding=!0,this._handleIndex=o,a.addClass("ui-state-active").focus(),h=a.offset(),l=!e(t.target).parents().addBack().is(".ui-slider-handle"),this._clickOffset=l?{left:0,top:0}:{left:t.pageX-h.left-a.width()/2,top:t.pageY-h.top-a.height()/2-(parseInt(a.css("borderTopWidth"),10)||0)-(parseInt(a.css("borderBottomWidth"),10)||0)+(parseInt(a.css("marginTop"),10)||0)},this.handles.hasClass("ui-state-hover")||this._slide(t,o,s),this._animateOff=!0,!0))},_mouseStart:function(){return!0},_mouseDrag:function(e){var t={x:e.pageX,y:e.pageY},i=this._normValueFromMouse(t);return this._slide(e,this._handleIndex,i),!1},_mouseStop:function(e){return this.handles.removeClass("ui-state-active"),this._mouseSliding=!1,this._stop(e,this._handleIndex),this._change(e,this._handleIndex),this._handleIndex=null,this._clickOffset=null,this._animateOff=!1,!1},_detectOrientation:function(){this.orientation="vertical"===this.options.orientation?"vertical":"horizontal"},_normValueFromMouse:function(e){var t,i,s,n,a;return"horizontal"===this.orientation?(t=this.elementSize.width,i=e.x-this.elementOffset.left-(this._clickOffset?this._clickOffset.left:0)):(t=this.elementSize.height,i=e.y-this.elementOffset.top-(this._clickOffset?this._clickOffset.top:0)),s=i/t,s>1&&(s=1),0>s&&(s=0),"vertical"===this.orientation&&(s=1-s),n=this._valueMax()-this._valueMin(),a=this._valueMin()+s*n,this._trimAlignValue(a)},_start:function(e,t){var i={handle:this.handles[t],value:this.value()};return this.options.values&&this.options.values.length&&(i.value=this.values(t),i.values=this.values()),this._trigger("start",e,i)},_slide:function(e,t,i){var s,n,a;this.options.values&&this.options.values.length?(s=this.values(t?0:1),2===this.options.values.length&&this.options.range===!0&&(0===t&&i>s||1===t&&s>i)&&(i=s),i!==this.values(t)&&(n=this.values(),n[t]=i,a=this._trigger("slide",e,{handle:this.handles[t],value:i,values:n}),s=this.values(t?0:1),a!==!1&&this.values(t,i))):i!==this.value()&&(a=this._trigger("slide",e,{handle:this.handles[t],value:i}),a!==!1&&this.value(i))},_stop:function(e,t){var i={handle:this.handles[t],value:this.value()};this.options.values&&this.options.values.length&&(i.value=this.values(t),i.values=this.values()),this._trigger("stop",e,i)},_change:function(e,t){if(!this._keySliding&&!this._mouseSliding){var i={handle:this.handles[t],value:this.value()};this.options.values&&this.options.values.length&&(i.value=this.values(t),i.values=this.values()),this._lastChangedValue=t,this._trigger("change",e,i)}},value:function(e){return arguments.length?(this.options.value=this._trimAlignValue(e),this._refreshValue(),this._change(null,0),void 0):this._value()},values:function(t,i){var s,n,a;if(arguments.length>1)return this.options.values[t]=this._trimAlignValue(i),this._refreshValue(),this._change(null,t),void 0;if(!arguments.length)return this._values();if(!e.isArray(arguments[0]))return this.options.values&&this.options.values.length?this._values(t):this.value();for(s=this.options.values,n=arguments[0],a=0;s.length>a;a+=1)s[a]=this._trimAlignValue(n[a]),this._change(null,a);this._refreshValue()},_setOption:function(t,i){var s,n=0;switch("range"===t&&this.options.range===!0&&("min"===i?(this.options.value=this._values(0),this.options.values=null):"max"===i&&(this.options.value=this._values(this.options.values.length-1),this.options.values=null)),e.isArray(this.options.values)&&(n=this.options.values.length),"disabled"===t&&this.element.toggleClass("ui-state-disabled",!!i),this._super(t,i),t){case"orientation":this._detectOrientation(),this.element.removeClass("ui-slider-horizontal ui-slider-vertical").addClass("ui-slider-"+this.orientation),this._refreshValue(),this.handles.css("horizontal"===i?"bottom":"left","");break;case"value":this._animateOff=!0,this._refreshValue(),this._change(null,0),this._animateOff=!1;break;case"values":for(this._animateOff=!0,this._refreshValue(),s=0;n>s;s+=1)this._change(null,s);this._animateOff=!1;break;case"step":case"min":case"max":this._animateOff=!0,this._calculateNewMax(),this._refreshValue(),this._animateOff=!1;break;case"range":this._animateOff=!0,this._refresh(),this._animateOff=!1}},_value:function(){var e=this.options.value;return e=this._trimAlignValue(e)},_values:function(e){var t,i,s;if(arguments.length)return t=this.options.values[e],t=this._trimAlignValue(t);if(this.options.values&&this.options.values.length){for(i=this.options.values.slice(),s=0;i.length>s;s+=1)i[s]=this._trimAlignValue(i[s]);return i}return[]},_trimAlignValue:function(e){if(this._valueMin()>=e)return this._valueMin();if(e>=this._valueMax())return this._valueMax();var t=this.options.step>0?this.options.step:1,i=(e-this._valueMin())%t,s=e-i;return 2*Math.abs(i)>=t&&(s+=i>0?t:-t),parseFloat(s.toFixed(5))},_calculateNewMax:function(){var e=this.options.max,t=this._valueMin(),i=this.options.step,s=Math.floor(+(e-t).toFixed(this._precision())/i)*i;e=s+t,this.max=parseFloat(e.toFixed(this._precision()))},_precision:function(){var e=this._precisionOf(this.options.step);return null!==this.options.min&&(e=Math.max(e,this._precisionOf(this.options.min))),e},_precisionOf:function(e){var t=""+e,i=t.indexOf(".");return-1===i?0:t.length-i-1},_valueMin:function(){return this.options.min},_valueMax:function(){return this.max},_refreshValue:function(){var t,i,s,n,a,o=this.options.range,r=this.options,h=this,l=this._animateOff?!1:r.animate,u={};this.options.values&&this.options.values.length?this.handles.each(function(s){i=100*((h.values(s)-h._valueMin())/(h._valueMax()-h._valueMin())),u["horizontal"===h.orientation?"left":"bottom"]=i+"%",e(this).stop(1,1)[l?"animate":"css"](u,r.animate),h.options.range===!0&&("horizontal"===h.orientation?(0===s&&h.range.stop(1,1)[l?"animate":"css"]({left:i+"%"},r.animate),1===s&&h.range[l?"animate":"css"]({width:i-t+"%"},{queue:!1,duration:r.animate})):(0===s&&h.range.stop(1,1)[l?"animate":"css"]({bottom:i+"%"},r.animate),1===s&&h.range[l?"animate":"css"]({height:i-t+"%"},{queue:!1,duration:r.animate}))),t=i}):(s=this.value(),n=this._valueMin(),a=this._valueMax(),i=a!==n?100*((s-n)/(a-n)):0,u["horizontal"===this.orientation?"left":"bottom"]=i+"%",this.handle.stop(1,1)[l?"animate":"css"](u,r.animate),"min"===o&&"horizontal"===this.orientation&&this.range.stop(1,1)[l?"animate":"css"]({width:i+"%"},r.animate),"max"===o&&"horizontal"===this.orientation&&this.range[l?"animate":"css"]({width:100-i+"%"},{queue:!1,duration:r.animate}),"min"===o&&"vertical"===this.orientation&&this.range.stop(1,1)[l?"animate":"css"]({height:i+"%"},r.animate),"max"===o&&"vertical"===this.orientation&&this.range[l?"animate":"css"]({height:100-i+"%"},{queue:!1,duration:r.animate}))},_handleEvents:{keydown:function(t){var i,s,n,a,o=e(t.target).data("ui-slider-handle-index");switch(t.keyCode){case e.ui.keyCode.HOME:case e.ui.keyCode.END:case e.ui.keyCode.PAGE_UP:case e.ui.keyCode.PAGE_DOWN:case e.ui.keyCode.UP:case e.ui.keyCode.RIGHT:case e.ui.keyCode.DOWN:case e.ui.keyCode.LEFT:if(t.preventDefault(),!this._keySliding&&(this._keySliding=!0,e(t.target).addClass("ui-state-active"),i=this._start(t,o),i===!1))return}switch(a=this.options.step,s=n=this.options.values&&this.options.values.length?this.values(o):this.value(),t.keyCode){case e.ui.keyCode.HOME:n=this._valueMin();break;case e.ui.keyCode.END:n=this._valueMax();break;case e.ui.keyCode.PAGE_UP:n=this._trimAlignValue(s+(this._valueMax()-this._valueMin())/this.numPages);break;case e.ui.keyCode.PAGE_DOWN:n=this._trimAlignValue(s-(this._valueMax()-this._valueMin())/this.numPages);break;case e.ui.keyCode.UP:case e.ui.keyCode.RIGHT:if(s===this._valueMax())return;n=this._trimAlignValue(s+a);break;case e.ui.keyCode.DOWN:case e.ui.keyCode.LEFT:if(s===this._valueMin())return;n=this._trimAlignValue(s-a)}this._slide(t,o,n)},keyup:function(t){var i=e(t.target).data("ui-slider-handle-index");this._keySliding&&(this._keySliding=!1,this._stop(t,i),this._change(t,i),e(t.target).removeClass("ui-state-active"))}}}),e.widget("ui.spinner",{version:"1.11.4",defaultElement:"<input>",widgetEventPrefix:"spin",options:{culture:null,icons:{down:"ui-icon-triangle-1-s",up:"ui-icon-triangle-1-n"},incremental:!0,max:null,min:null,numberFormat:null,page:10,step:1,change:null,spin:null,start:null,stop:null},_create:function(){this._setOption("max",this.options.max),this._setOption("min",this.options.min),this._setOption("step",this.options.step),""!==this.value()&&this._value(this.element.val(),!0),this._draw(),this._on(this._events),this._refresh(),this._on(this.window,{beforeunload:function(){this.element.removeAttr("autocomplete")}})},_getCreateOptions:function(){var t={},i=this.element;return e.each(["min","max","step"],function(e,s){var n=i.attr(s);void 0!==n&&n.length&&(t[s]=n)}),t},_events:{keydown:function(e){this._start(e)&&this._keydown(e)&&e.preventDefault()},keyup:"_stop",focus:function(){this.previous=this.element.val()},blur:function(e){return this.cancelBlur?(delete this.cancelBlur,void 0):(this._stop(),this._refresh(),this.previous!==this.element.val()&&this._trigger("change",e),void 0)},mousewheel:function(e,t){if(t){if(!this.spinning&&!this._start(e))return!1;this._spin((t>0?1:-1)*this.options.step,e),clearTimeout(this.mousewheelTimer),this.mousewheelTimer=this._delay(function(){this.spinning&&this._stop(e)},100),e.preventDefault()}},"mousedown .ui-spinner-button":function(t){function i(){var e=this.element[0]===this.document[0].activeElement;e||(this.element.focus(),this.previous=s,this._delay(function(){this.previous=s}))}var s;s=this.element[0]===this.document[0].activeElement?this.previous:this.element.val(),t.preventDefault(),i.call(this),this.cancelBlur=!0,this._delay(function(){delete this.cancelBlur,i.call(this)}),this._start(t)!==!1&&this._repeat(null,e(t.currentTarget).hasClass("ui-spinner-up")?1:-1,t)},"mouseup .ui-spinner-button":"_stop","mouseenter .ui-spinner-button":function(t){return e(t.currentTarget).hasClass("ui-state-active")?this._start(t)===!1?!1:(this._repeat(null,e(t.currentTarget).hasClass("ui-spinner-up")?1:-1,t),void 0):void 0},"mouseleave .ui-spinner-button":"_stop"},_draw:function(){var e=this.uiSpinner=this.element.addClass("ui-spinner-input").attr("autocomplete","off").wrap(this._uiSpinnerHtml()).parent().append(this._buttonHtml());this.element.attr("role","spinbutton"),this.buttons=e.find(".ui-spinner-button").attr("tabIndex",-1).button().removeClass("ui-corner-all"),this.buttons.height()>Math.ceil(.5*e.height())&&e.height()>0&&e.height(e.height()),this.options.disabled&&this.disable()},_keydown:function(t){var i=this.options,s=e.ui.keyCode;switch(t.keyCode){case s.UP:return this._repeat(null,1,t),!0;case s.DOWN:return this._repeat(null,-1,t),!0;case s.PAGE_UP:return this._repeat(null,i.page,t),!0;case s.PAGE_DOWN:return this._repeat(null,-i.page,t),!0}return!1},_uiSpinnerHtml:function(){return"<span class='ui-spinner ui-widget ui-widget-content ui-corner-all'></span>"},_buttonHtml:function(){return"<a class='ui-spinner-button ui-spinner-up ui-corner-tr'><span class='ui-icon "+this.options.icons.up+"'>&#9650;</span>"+"</a>"+"<a class='ui-spinner-button ui-spinner-down ui-corner-br'>"+"<span class='ui-icon "+this.options.icons.down+"'>&#9660;</span>"+"</a>"},_start:function(e){return this.spinning||this._trigger("start",e)!==!1?(this.counter||(this.counter=1),this.spinning=!0,!0):!1},_repeat:function(e,t,i){e=e||500,clearTimeout(this.timer),this.timer=this._delay(function(){this._repeat(40,t,i)},e),this._spin(t*this.options.step,i)},_spin:function(e,t){var i=this.value()||0;this.counter||(this.counter=1),i=this._adjustValue(i+e*this._increment(this.counter)),this.spinning&&this._trigger("spin",t,{value:i})===!1||(this._value(i),this.counter++)},_increment:function(t){var i=this.options.incremental;return i?e.isFunction(i)?i(t):Math.floor(t*t*t/5e4-t*t/500+17*t/200+1):1},_precision:function(){var e=this._precisionOf(this.options.step);return null!==this.options.min&&(e=Math.max(e,this._precisionOf(this.options.min))),e},_precisionOf:function(e){var t=""+e,i=t.indexOf(".");return-1===i?0:t.length-i-1},_adjustValue:function(e){var t,i,s=this.options;return t=null!==s.min?s.min:0,i=e-t,i=Math.round(i/s.step)*s.step,e=t+i,e=parseFloat(e.toFixed(this._precision())),null!==s.max&&e>s.max?s.max:null!==s.min&&s.min>e?s.min:e},_stop:function(e){this.spinning&&(clearTimeout(this.timer),clearTimeout(this.mousewheelTimer),this.counter=0,this.spinning=!1,this._trigger("stop",e))},_setOption:function(e,t){if("culture"===e||"numberFormat"===e){var i=this._parse(this.element.val());return this.options[e]=t,this.element.val(this._format(i)),void 0}("max"===e||"min"===e||"step"===e)&&"string"==typeof t&&(t=this._parse(t)),"icons"===e&&(this.buttons.first().find(".ui-icon").removeClass(this.options.icons.up).addClass(t.up),this.buttons.last().find(".ui-icon").removeClass(this.options.icons.down).addClass(t.down)),this._super(e,t),"disabled"===e&&(this.widget().toggleClass("ui-state-disabled",!!t),this.element.prop("disabled",!!t),this.buttons.button(t?"disable":"enable"))},_setOptions:s(function(e){this._super(e)}),_parse:function(e){return"string"==typeof e&&""!==e&&(e=window.Globalize&&this.options.numberFormat?Globalize.parseFloat(e,10,this.options.culture):+e),""===e||isNaN(e)?null:e
},_format:function(e){return""===e?"":window.Globalize&&this.options.numberFormat?Globalize.format(e,this.options.numberFormat,this.options.culture):e},_refresh:function(){this.element.attr({"aria-valuemin":this.options.min,"aria-valuemax":this.options.max,"aria-valuenow":this._parse(this.element.val())})},isValid:function(){var e=this.value();return null===e?!1:e===this._adjustValue(e)},_value:function(e,t){var i;""!==e&&(i=this._parse(e),null!==i&&(t||(i=this._adjustValue(i)),e=this._format(i))),this.element.val(e),this._refresh()},_destroy:function(){this.element.removeClass("ui-spinner-input").prop("disabled",!1).removeAttr("autocomplete").removeAttr("role").removeAttr("aria-valuemin").removeAttr("aria-valuemax").removeAttr("aria-valuenow"),this.uiSpinner.replaceWith(this.element)},stepUp:s(function(e){this._stepUp(e)}),_stepUp:function(e){this._start()&&(this._spin((e||1)*this.options.step),this._stop())},stepDown:s(function(e){this._stepDown(e)}),_stepDown:function(e){this._start()&&(this._spin((e||1)*-this.options.step),this._stop())},pageUp:s(function(e){this._stepUp((e||1)*this.options.page)}),pageDown:s(function(e){this._stepDown((e||1)*this.options.page)}),value:function(e){return arguments.length?(s(this._value).call(this,e),void 0):this._parse(this.element.val())},widget:function(){return this.uiSpinner}}),e.widget("ui.tabs",{version:"1.11.4",delay:300,options:{active:null,collapsible:!1,event:"click",heightStyle:"content",hide:null,show:null,activate:null,beforeActivate:null,beforeLoad:null,load:null},_isLocal:function(){var e=/#.*$/;return function(t){var i,s;t=t.cloneNode(!1),i=t.href.replace(e,""),s=location.href.replace(e,"");try{i=decodeURIComponent(i)}catch(n){}try{s=decodeURIComponent(s)}catch(n){}return t.hash.length>1&&i===s}}(),_create:function(){var t=this,i=this.options;this.running=!1,this.element.addClass("ui-tabs ui-widget ui-widget-content ui-corner-all").toggleClass("ui-tabs-collapsible",i.collapsible),this._processTabs(),i.active=this._initialActive(),e.isArray(i.disabled)&&(i.disabled=e.unique(i.disabled.concat(e.map(this.tabs.filter(".ui-state-disabled"),function(e){return t.tabs.index(e)}))).sort()),this.active=this.options.active!==!1&&this.anchors.length?this._findActive(i.active):e(),this._refresh(),this.active.length&&this.load(i.active)},_initialActive:function(){var t=this.options.active,i=this.options.collapsible,s=location.hash.substring(1);return null===t&&(s&&this.tabs.each(function(i,n){return e(n).attr("aria-controls")===s?(t=i,!1):void 0}),null===t&&(t=this.tabs.index(this.tabs.filter(".ui-tabs-active"))),(null===t||-1===t)&&(t=this.tabs.length?0:!1)),t!==!1&&(t=this.tabs.index(this.tabs.eq(t)),-1===t&&(t=i?!1:0)),!i&&t===!1&&this.anchors.length&&(t=0),t},_getCreateEventData:function(){return{tab:this.active,panel:this.active.length?this._getPanelForTab(this.active):e()}},_tabKeydown:function(t){var i=e(this.document[0].activeElement).closest("li"),s=this.tabs.index(i),n=!0;if(!this._handlePageNav(t)){switch(t.keyCode){case e.ui.keyCode.RIGHT:case e.ui.keyCode.DOWN:s++;break;case e.ui.keyCode.UP:case e.ui.keyCode.LEFT:n=!1,s--;break;case e.ui.keyCode.END:s=this.anchors.length-1;break;case e.ui.keyCode.HOME:s=0;break;case e.ui.keyCode.SPACE:return t.preventDefault(),clearTimeout(this.activating),this._activate(s),void 0;case e.ui.keyCode.ENTER:return t.preventDefault(),clearTimeout(this.activating),this._activate(s===this.options.active?!1:s),void 0;default:return}t.preventDefault(),clearTimeout(this.activating),s=this._focusNextTab(s,n),t.ctrlKey||t.metaKey||(i.attr("aria-selected","false"),this.tabs.eq(s).attr("aria-selected","true"),this.activating=this._delay(function(){this.option("active",s)},this.delay))}},_panelKeydown:function(t){this._handlePageNav(t)||t.ctrlKey&&t.keyCode===e.ui.keyCode.UP&&(t.preventDefault(),this.active.focus())},_handlePageNav:function(t){return t.altKey&&t.keyCode===e.ui.keyCode.PAGE_UP?(this._activate(this._focusNextTab(this.options.active-1,!1)),!0):t.altKey&&t.keyCode===e.ui.keyCode.PAGE_DOWN?(this._activate(this._focusNextTab(this.options.active+1,!0)),!0):void 0},_findNextTab:function(t,i){function s(){return t>n&&(t=0),0>t&&(t=n),t}for(var n=this.tabs.length-1;-1!==e.inArray(s(),this.options.disabled);)t=i?t+1:t-1;return t},_focusNextTab:function(e,t){return e=this._findNextTab(e,t),this.tabs.eq(e).focus(),e},_setOption:function(e,t){return"active"===e?(this._activate(t),void 0):"disabled"===e?(this._setupDisabled(t),void 0):(this._super(e,t),"collapsible"===e&&(this.element.toggleClass("ui-tabs-collapsible",t),t||this.options.active!==!1||this._activate(0)),"event"===e&&this._setupEvents(t),"heightStyle"===e&&this._setupHeightStyle(t),void 0)},_sanitizeSelector:function(e){return e?e.replace(/[!"$%&'()*+,.\/:;<=>?@\[\]\^`{|}~]/g,"\\$&"):""},refresh:function(){var t=this.options,i=this.tablist.children(":has(a[href])");t.disabled=e.map(i.filter(".ui-state-disabled"),function(e){return i.index(e)}),this._processTabs(),t.active!==!1&&this.anchors.length?this.active.length&&!e.contains(this.tablist[0],this.active[0])?this.tabs.length===t.disabled.length?(t.active=!1,this.active=e()):this._activate(this._findNextTab(Math.max(0,t.active-1),!1)):t.active=this.tabs.index(this.active):(t.active=!1,this.active=e()),this._refresh()},_refresh:function(){this._setupDisabled(this.options.disabled),this._setupEvents(this.options.event),this._setupHeightStyle(this.options.heightStyle),this.tabs.not(this.active).attr({"aria-selected":"false","aria-expanded":"false",tabIndex:-1}),this.panels.not(this._getPanelForTab(this.active)).hide().attr({"aria-hidden":"true"}),this.active.length?(this.active.addClass("ui-tabs-active ui-state-active").attr({"aria-selected":"true","aria-expanded":"true",tabIndex:0}),this._getPanelForTab(this.active).show().attr({"aria-hidden":"false"})):this.tabs.eq(0).attr("tabIndex",0)},_processTabs:function(){var t=this,i=this.tabs,s=this.anchors,n=this.panels;this.tablist=this._getList().addClass("ui-tabs-nav ui-helper-reset ui-helper-clearfix ui-widget-header ui-corner-all").attr("role","tablist").delegate("> li","mousedown"+this.eventNamespace,function(t){e(this).is(".ui-state-disabled")&&t.preventDefault()}).delegate(".ui-tabs-anchor","focus"+this.eventNamespace,function(){e(this).closest("li").is(".ui-state-disabled")&&this.blur()}),this.tabs=this.tablist.find("> li:has(a[href])").addClass("ui-state-default ui-corner-top").attr({role:"tab",tabIndex:-1}),this.anchors=this.tabs.map(function(){return e("a",this)[0]}).addClass("ui-tabs-anchor").attr({role:"presentation",tabIndex:-1}),this.panels=e(),this.anchors.each(function(i,s){var n,a,o,r=e(s).uniqueId().attr("id"),h=e(s).closest("li"),l=h.attr("aria-controls");t._isLocal(s)?(n=s.hash,o=n.substring(1),a=t.element.find(t._sanitizeSelector(n))):(o=h.attr("aria-controls")||e({}).uniqueId()[0].id,n="#"+o,a=t.element.find(n),a.length||(a=t._createPanel(o),a.insertAfter(t.panels[i-1]||t.tablist)),a.attr("aria-live","polite")),a.length&&(t.panels=t.panels.add(a)),l&&h.data("ui-tabs-aria-controls",l),h.attr({"aria-controls":o,"aria-labelledby":r}),a.attr("aria-labelledby",r)}),this.panels.addClass("ui-tabs-panel ui-widget-content ui-corner-bottom").attr("role","tabpanel"),i&&(this._off(i.not(this.tabs)),this._off(s.not(this.anchors)),this._off(n.not(this.panels)))},_getList:function(){return this.tablist||this.element.find("ol,ul").eq(0)},_createPanel:function(t){return e("<div>").attr("id",t).addClass("ui-tabs-panel ui-widget-content ui-corner-bottom").data("ui-tabs-destroy",!0)},_setupDisabled:function(t){e.isArray(t)&&(t.length?t.length===this.anchors.length&&(t=!0):t=!1);for(var i,s=0;i=this.tabs[s];s++)t===!0||-1!==e.inArray(s,t)?e(i).addClass("ui-state-disabled").attr("aria-disabled","true"):e(i).removeClass("ui-state-disabled").removeAttr("aria-disabled");this.options.disabled=t},_setupEvents:function(t){var i={};t&&e.each(t.split(" "),function(e,t){i[t]="_eventHandler"}),this._off(this.anchors.add(this.tabs).add(this.panels)),this._on(!0,this.anchors,{click:function(e){e.preventDefault()}}),this._on(this.anchors,i),this._on(this.tabs,{keydown:"_tabKeydown"}),this._on(this.panels,{keydown:"_panelKeydown"}),this._focusable(this.tabs),this._hoverable(this.tabs)},_setupHeightStyle:function(t){var i,s=this.element.parent();"fill"===t?(i=s.height(),i-=this.element.outerHeight()-this.element.height(),this.element.siblings(":visible").each(function(){var t=e(this),s=t.css("position");"absolute"!==s&&"fixed"!==s&&(i-=t.outerHeight(!0))}),this.element.children().not(this.panels).each(function(){i-=e(this).outerHeight(!0)}),this.panels.each(function(){e(this).height(Math.max(0,i-e(this).innerHeight()+e(this).height()))}).css("overflow","auto")):"auto"===t&&(i=0,this.panels.each(function(){i=Math.max(i,e(this).height("").height())}).height(i))},_eventHandler:function(t){var i=this.options,s=this.active,n=e(t.currentTarget),a=n.closest("li"),o=a[0]===s[0],r=o&&i.collapsible,h=r?e():this._getPanelForTab(a),l=s.length?this._getPanelForTab(s):e(),u={oldTab:s,oldPanel:l,newTab:r?e():a,newPanel:h};t.preventDefault(),a.hasClass("ui-state-disabled")||a.hasClass("ui-tabs-loading")||this.running||o&&!i.collapsible||this._trigger("beforeActivate",t,u)===!1||(i.active=r?!1:this.tabs.index(a),this.active=o?e():a,this.xhr&&this.xhr.abort(),l.length||h.length||e.error("jQuery UI Tabs: Mismatching fragment identifier."),h.length&&this.load(this.tabs.index(a),t),this._toggle(t,u))},_toggle:function(t,i){function s(){a.running=!1,a._trigger("activate",t,i)}function n(){i.newTab.closest("li").addClass("ui-tabs-active ui-state-active"),o.length&&a.options.show?a._show(o,a.options.show,s):(o.show(),s())}var a=this,o=i.newPanel,r=i.oldPanel;this.running=!0,r.length&&this.options.hide?this._hide(r,this.options.hide,function(){i.oldTab.closest("li").removeClass("ui-tabs-active ui-state-active"),n()}):(i.oldTab.closest("li").removeClass("ui-tabs-active ui-state-active"),r.hide(),n()),r.attr("aria-hidden","true"),i.oldTab.attr({"aria-selected":"false","aria-expanded":"false"}),o.length&&r.length?i.oldTab.attr("tabIndex",-1):o.length&&this.tabs.filter(function(){return 0===e(this).attr("tabIndex")}).attr("tabIndex",-1),o.attr("aria-hidden","false"),i.newTab.attr({"aria-selected":"true","aria-expanded":"true",tabIndex:0})},_activate:function(t){var i,s=this._findActive(t);s[0]!==this.active[0]&&(s.length||(s=this.active),i=s.find(".ui-tabs-anchor")[0],this._eventHandler({target:i,currentTarget:i,preventDefault:e.noop}))},_findActive:function(t){return t===!1?e():this.tabs.eq(t)},_getIndex:function(e){return"string"==typeof e&&(e=this.anchors.index(this.anchors.filter("[href$='"+e+"']"))),e},_destroy:function(){this.xhr&&this.xhr.abort(),this.element.removeClass("ui-tabs ui-widget ui-widget-content ui-corner-all ui-tabs-collapsible"),this.tablist.removeClass("ui-tabs-nav ui-helper-reset ui-helper-clearfix ui-widget-header ui-corner-all").removeAttr("role"),this.anchors.removeClass("ui-tabs-anchor").removeAttr("role").removeAttr("tabIndex").removeUniqueId(),this.tablist.unbind(this.eventNamespace),this.tabs.add(this.panels).each(function(){e.data(this,"ui-tabs-destroy")?e(this).remove():e(this).removeClass("ui-state-default ui-state-active ui-state-disabled ui-corner-top ui-corner-bottom ui-widget-content ui-tabs-active ui-tabs-panel").removeAttr("tabIndex").removeAttr("aria-live").removeAttr("aria-busy").removeAttr("aria-selected").removeAttr("aria-labelledby").removeAttr("aria-hidden").removeAttr("aria-expanded").removeAttr("role")}),this.tabs.each(function(){var t=e(this),i=t.data("ui-tabs-aria-controls");i?t.attr("aria-controls",i).removeData("ui-tabs-aria-controls"):t.removeAttr("aria-controls")}),this.panels.show(),"content"!==this.options.heightStyle&&this.panels.css("height","")},enable:function(t){var i=this.options.disabled;i!==!1&&(void 0===t?i=!1:(t=this._getIndex(t),i=e.isArray(i)?e.map(i,function(e){return e!==t?e:null}):e.map(this.tabs,function(e,i){return i!==t?i:null})),this._setupDisabled(i))},disable:function(t){var i=this.options.disabled;if(i!==!0){if(void 0===t)i=!0;else{if(t=this._getIndex(t),-1!==e.inArray(t,i))return;i=e.isArray(i)?e.merge([t],i).sort():[t]}this._setupDisabled(i)}},load:function(t,i){t=this._getIndex(t);var s=this,n=this.tabs.eq(t),a=n.find(".ui-tabs-anchor"),o=this._getPanelForTab(n),r={tab:n,panel:o},h=function(e,t){"abort"===t&&s.panels.stop(!1,!0),n.removeClass("ui-tabs-loading"),o.removeAttr("aria-busy"),e===s.xhr&&delete s.xhr};this._isLocal(a[0])||(this.xhr=e.ajax(this._ajaxSettings(a,i,r)),this.xhr&&"canceled"!==this.xhr.statusText&&(n.addClass("ui-tabs-loading"),o.attr("aria-busy","true"),this.xhr.done(function(e,t,n){setTimeout(function(){o.html(e),s._trigger("load",i,r),h(n,t)},1)}).fail(function(e,t){setTimeout(function(){h(e,t)},1)})))},_ajaxSettings:function(t,i,s){var n=this;return{url:t.attr("href"),beforeSend:function(t,a){return n._trigger("beforeLoad",i,e.extend({jqXHR:t,ajaxSettings:a},s))}}},_getPanelForTab:function(t){var i=e(t).attr("aria-controls");return this.element.find(this._sanitizeSelector("#"+i))}}),e.widget("ui.tooltip",{version:"1.11.4",options:{content:function(){var t=e(this).attr("title")||"";return e("<a>").text(t).html()},hide:!0,items:"[title]:not([disabled])",position:{my:"left top+15",at:"left bottom",collision:"flipfit flip"},show:!0,tooltipClass:null,track:!1,close:null,open:null},_addDescribedBy:function(t,i){var s=(t.attr("aria-describedby")||"").split(/\s+/);s.push(i),t.data("ui-tooltip-id",i).attr("aria-describedby",e.trim(s.join(" ")))},_removeDescribedBy:function(t){var i=t.data("ui-tooltip-id"),s=(t.attr("aria-describedby")||"").split(/\s+/),n=e.inArray(i,s);-1!==n&&s.splice(n,1),t.removeData("ui-tooltip-id"),s=e.trim(s.join(" ")),s?t.attr("aria-describedby",s):t.removeAttr("aria-describedby")},_create:function(){this._on({mouseover:"open",focusin:"open"}),this.tooltips={},this.parents={},this.options.disabled&&this._disable(),this.liveRegion=e("<div>").attr({role:"log","aria-live":"assertive","aria-relevant":"additions"}).addClass("ui-helper-hidden-accessible").appendTo(this.document[0].body)},_setOption:function(t,i){var s=this;return"disabled"===t?(this[i?"_disable":"_enable"](),this.options[t]=i,void 0):(this._super(t,i),"content"===t&&e.each(this.tooltips,function(e,t){s._updateContent(t.element)}),void 0)},_disable:function(){var t=this;e.each(this.tooltips,function(i,s){var n=e.Event("blur");n.target=n.currentTarget=s.element[0],t.close(n,!0)}),this.element.find(this.options.items).addBack().each(function(){var t=e(this);t.is("[title]")&&t.data("ui-tooltip-title",t.attr("title")).removeAttr("title")})},_enable:function(){this.element.find(this.options.items).addBack().each(function(){var t=e(this);t.data("ui-tooltip-title")&&t.attr("title",t.data("ui-tooltip-title"))})},open:function(t){var i=this,s=e(t?t.target:this.element).closest(this.options.items);s.length&&!s.data("ui-tooltip-id")&&(s.attr("title")&&s.data("ui-tooltip-title",s.attr("title")),s.data("ui-tooltip-open",!0),t&&"mouseover"===t.type&&s.parents().each(function(){var t,s=e(this);s.data("ui-tooltip-open")&&(t=e.Event("blur"),t.target=t.currentTarget=this,i.close(t,!0)),s.attr("title")&&(s.uniqueId(),i.parents[this.id]={element:this,title:s.attr("title")},s.attr("title",""))}),this._registerCloseHandlers(t,s),this._updateContent(s,t))},_updateContent:function(e,t){var i,s=this.options.content,n=this,a=t?t.type:null;return"string"==typeof s?this._open(t,e,s):(i=s.call(e[0],function(i){n._delay(function(){e.data("ui-tooltip-open")&&(t&&(t.type=a),this._open(t,e,i))})}),i&&this._open(t,e,i),void 0)},_open:function(t,i,s){function n(e){l.of=e,o.is(":hidden")||o.position(l)}var a,o,r,h,l=e.extend({},this.options.position);if(s){if(a=this._find(i))return a.tooltip.find(".ui-tooltip-content").html(s),void 0;i.is("[title]")&&(t&&"mouseover"===t.type?i.attr("title",""):i.removeAttr("title")),a=this._tooltip(i),o=a.tooltip,this._addDescribedBy(i,o.attr("id")),o.find(".ui-tooltip-content").html(s),this.liveRegion.children().hide(),s.clone?(h=s.clone(),h.removeAttr("id").find("[id]").removeAttr("id")):h=s,e("<div>").html(h).appendTo(this.liveRegion),this.options.track&&t&&/^mouse/.test(t.type)?(this._on(this.document,{mousemove:n}),n(t)):o.position(e.extend({of:i},this.options.position)),o.hide(),this._show(o,this.options.show),this.options.show&&this.options.show.delay&&(r=this.delayedShow=setInterval(function(){o.is(":visible")&&(n(l.of),clearInterval(r))},e.fx.interval)),this._trigger("open",t,{tooltip:o})}},_registerCloseHandlers:function(t,i){var s={keyup:function(t){if(t.keyCode===e.ui.keyCode.ESCAPE){var s=e.Event(t);s.currentTarget=i[0],this.close(s,!0)}}};i[0]!==this.element[0]&&(s.remove=function(){this._removeTooltip(this._find(i).tooltip)}),t&&"mouseover"!==t.type||(s.mouseleave="close"),t&&"focusin"!==t.type||(s.focusout="close"),this._on(!0,i,s)},close:function(t){var i,s=this,n=e(t?t.currentTarget:this.element),a=this._find(n);return a?(i=a.tooltip,a.closing||(clearInterval(this.delayedShow),n.data("ui-tooltip-title")&&!n.attr("title")&&n.attr("title",n.data("ui-tooltip-title")),this._removeDescribedBy(n),a.hiding=!0,i.stop(!0),this._hide(i,this.options.hide,function(){s._removeTooltip(e(this))}),n.removeData("ui-tooltip-open"),this._off(n,"mouseleave focusout keyup"),n[0]!==this.element[0]&&this._off(n,"remove"),this._off(this.document,"mousemove"),t&&"mouseleave"===t.type&&e.each(this.parents,function(t,i){e(i.element).attr("title",i.title),delete s.parents[t]}),a.closing=!0,this._trigger("close",t,{tooltip:i}),a.hiding||(a.closing=!1)),void 0):(n.removeData("ui-tooltip-open"),void 0)},_tooltip:function(t){var i=e("<div>").attr("role","tooltip").addClass("ui-tooltip ui-widget ui-corner-all ui-widget-content "+(this.options.tooltipClass||"")),s=i.uniqueId().attr("id");return e("<div>").addClass("ui-tooltip-content").appendTo(i),i.appendTo(this.document[0].body),this.tooltips[s]={element:t,tooltip:i}},_find:function(e){var t=e.data("ui-tooltip-id");return t?this.tooltips[t]:null},_removeTooltip:function(e){e.remove(),delete this.tooltips[e.attr("id")]},_destroy:function(){var t=this;e.each(this.tooltips,function(i,s){var n=e.Event("blur"),a=s.element;n.target=n.currentTarget=a[0],t.close(n,!0),e("#"+i).remove(),a.data("ui-tooltip-title")&&(a.attr("title")||a.attr("title",a.data("ui-tooltip-title")),a.removeData("ui-tooltip-title"))}),this.liveRegion.remove()}});var c="ui-effects-",p=e;e.effects={effect:{}},function(e,t){function i(e,t,i){var s=d[t.type]||{};return null==e?i||!t.def?null:t.def:(e=s.floor?~~e:parseFloat(e),isNaN(e)?t.def:s.mod?(e+s.mod)%s.mod:0>e?0:e>s.max?s.max:e)}function s(i){var s=l(),n=s._rgba=[];return i=i.toLowerCase(),f(h,function(e,a){var o,r=a.re.exec(i),h=r&&a.parse(r),l=a.space||"rgba";return h?(o=s[l](h),s[u[l].cache]=o[u[l].cache],n=s._rgba=o._rgba,!1):t}),n.length?("0,0,0,0"===n.join()&&e.extend(n,a.transparent),s):a[i]}function n(e,t,i){return i=(i+1)%1,1>6*i?e+6*(t-e)*i:1>2*i?t:2>3*i?e+6*(t-e)*(2/3-i):e}var a,o="backgroundColor borderBottomColor borderLeftColor borderRightColor borderTopColor color columnRuleColor outlineColor textDecorationColor textEmphasisColor",r=/^([\-+])=\s*(\d+\.?\d*)/,h=[{re:/rgba?\(\s*(\d{1,3})\s*,\s*(\d{1,3})\s*,\s*(\d{1,3})\s*(?:,\s*(\d?(?:\.\d+)?)\s*)?\)/,parse:function(e){return[e[1],e[2],e[3],e[4]]}},{re:/rgba?\(\s*(\d+(?:\.\d+)?)\%\s*,\s*(\d+(?:\.\d+)?)\%\s*,\s*(\d+(?:\.\d+)?)\%\s*(?:,\s*(\d?(?:\.\d+)?)\s*)?\)/,parse:function(e){return[2.55*e[1],2.55*e[2],2.55*e[3],e[4]]}},{re:/#([a-f0-9]{2})([a-f0-9]{2})([a-f0-9]{2})/,parse:function(e){return[parseInt(e[1],16),parseInt(e[2],16),parseInt(e[3],16)]}},{re:/#([a-f0-9])([a-f0-9])([a-f0-9])/,parse:function(e){return[parseInt(e[1]+e[1],16),parseInt(e[2]+e[2],16),parseInt(e[3]+e[3],16)]}},{re:/hsla?\(\s*(\d+(?:\.\d+)?)\s*,\s*(\d+(?:\.\d+)?)\%\s*,\s*(\d+(?:\.\d+)?)\%\s*(?:,\s*(\d?(?:\.\d+)?)\s*)?\)/,space:"hsla",parse:function(e){return[e[1],e[2]/100,e[3]/100,e[4]]}}],l=e.Color=function(t,i,s,n){return new e.Color.fn.parse(t,i,s,n)},u={rgba:{props:{red:{idx:0,type:"byte"},green:{idx:1,type:"byte"},blue:{idx:2,type:"byte"}}},hsla:{props:{hue:{idx:0,type:"degrees"},saturation:{idx:1,type:"percent"},lightness:{idx:2,type:"percent"}}}},d={"byte":{floor:!0,max:255},percent:{max:1},degrees:{mod:360,floor:!0}},c=l.support={},p=e("<p>")[0],f=e.each;p.style.cssText="background-color:rgba(1,1,1,.5)",c.rgba=p.style.backgroundColor.indexOf("rgba")>-1,f(u,function(e,t){t.cache="_"+e,t.props.alpha={idx:3,type:"percent",def:1}}),l.fn=e.extend(l.prototype,{parse:function(n,o,r,h){if(n===t)return this._rgba=[null,null,null,null],this;(n.jquery||n.nodeType)&&(n=e(n).css(o),o=t);var d=this,c=e.type(n),p=this._rgba=[];return o!==t&&(n=[n,o,r,h],c="array"),"string"===c?this.parse(s(n)||a._default):"array"===c?(f(u.rgba.props,function(e,t){p[t.idx]=i(n[t.idx],t)}),this):"object"===c?(n instanceof l?f(u,function(e,t){n[t.cache]&&(d[t.cache]=n[t.cache].slice())}):f(u,function(t,s){var a=s.cache;f(s.props,function(e,t){if(!d[a]&&s.to){if("alpha"===e||null==n[e])return;d[a]=s.to(d._rgba)}d[a][t.idx]=i(n[e],t,!0)}),d[a]&&0>e.inArray(null,d[a].slice(0,3))&&(d[a][3]=1,s.from&&(d._rgba=s.from(d[a])))}),this):t},is:function(e){var i=l(e),s=!0,n=this;return f(u,function(e,a){var o,r=i[a.cache];return r&&(o=n[a.cache]||a.to&&a.to(n._rgba)||[],f(a.props,function(e,i){return null!=r[i.idx]?s=r[i.idx]===o[i.idx]:t})),s}),s},_space:function(){var e=[],t=this;return f(u,function(i,s){t[s.cache]&&e.push(i)}),e.pop()},transition:function(e,t){var s=l(e),n=s._space(),a=u[n],o=0===this.alpha()?l("transparent"):this,r=o[a.cache]||a.to(o._rgba),h=r.slice();return s=s[a.cache],f(a.props,function(e,n){var a=n.idx,o=r[a],l=s[a],u=d[n.type]||{};null!==l&&(null===o?h[a]=l:(u.mod&&(l-o>u.mod/2?o+=u.mod:o-l>u.mod/2&&(o-=u.mod)),h[a]=i((l-o)*t+o,n)))}),this[n](h)},blend:function(t){if(1===this._rgba[3])return this;var i=this._rgba.slice(),s=i.pop(),n=l(t)._rgba;return l(e.map(i,function(e,t){return(1-s)*n[t]+s*e}))},toRgbaString:function(){var t="rgba(",i=e.map(this._rgba,function(e,t){return null==e?t>2?1:0:e});return 1===i[3]&&(i.pop(),t="rgb("),t+i.join()+")"},toHslaString:function(){var t="hsla(",i=e.map(this.hsla(),function(e,t){return null==e&&(e=t>2?1:0),t&&3>t&&(e=Math.round(100*e)+"%"),e});return 1===i[3]&&(i.pop(),t="hsl("),t+i.join()+")"},toHexString:function(t){var i=this._rgba.slice(),s=i.pop();return t&&i.push(~~(255*s)),"#"+e.map(i,function(e){return e=(e||0).toString(16),1===e.length?"0"+e:e}).join("")},toString:function(){return 0===this._rgba[3]?"transparent":this.toRgbaString()}}),l.fn.parse.prototype=l.fn,u.hsla.to=function(e){if(null==e[0]||null==e[1]||null==e[2])return[null,null,null,e[3]];var t,i,s=e[0]/255,n=e[1]/255,a=e[2]/255,o=e[3],r=Math.max(s,n,a),h=Math.min(s,n,a),l=r-h,u=r+h,d=.5*u;return t=h===r?0:s===r?60*(n-a)/l+360:n===r?60*(a-s)/l+120:60*(s-n)/l+240,i=0===l?0:.5>=d?l/u:l/(2-u),[Math.round(t)%360,i,d,null==o?1:o]},u.hsla.from=function(e){if(null==e[0]||null==e[1]||null==e[2])return[null,null,null,e[3]];var t=e[0]/360,i=e[1],s=e[2],a=e[3],o=.5>=s?s*(1+i):s+i-s*i,r=2*s-o;return[Math.round(255*n(r,o,t+1/3)),Math.round(255*n(r,o,t)),Math.round(255*n(r,o,t-1/3)),a]},f(u,function(s,n){var a=n.props,o=n.cache,h=n.to,u=n.from;l.fn[s]=function(s){if(h&&!this[o]&&(this[o]=h(this._rgba)),s===t)return this[o].slice();var n,r=e.type(s),d="array"===r||"object"===r?s:arguments,c=this[o].slice();return f(a,function(e,t){var s=d["object"===r?e:t.idx];null==s&&(s=c[t.idx]),c[t.idx]=i(s,t)}),u?(n=l(u(c)),n[o]=c,n):l(c)},f(a,function(t,i){l.fn[t]||(l.fn[t]=function(n){var a,o=e.type(n),h="alpha"===t?this._hsla?"hsla":"rgba":s,l=this[h](),u=l[i.idx];return"undefined"===o?u:("function"===o&&(n=n.call(this,u),o=e.type(n)),null==n&&i.empty?this:("string"===o&&(a=r.exec(n),a&&(n=u+parseFloat(a[2])*("+"===a[1]?1:-1))),l[i.idx]=n,this[h](l)))})})}),l.hook=function(t){var i=t.split(" ");f(i,function(t,i){e.cssHooks[i]={set:function(t,n){var a,o,r="";if("transparent"!==n&&("string"!==e.type(n)||(a=s(n)))){if(n=l(a||n),!c.rgba&&1!==n._rgba[3]){for(o="backgroundColor"===i?t.parentNode:t;(""===r||"transparent"===r)&&o&&o.style;)try{r=e.css(o,"backgroundColor"),o=o.parentNode}catch(h){}n=n.blend(r&&"transparent"!==r?r:"_default")}n=n.toRgbaString()}try{t.style[i]=n}catch(h){}}},e.fx.step[i]=function(t){t.colorInit||(t.start=l(t.elem,i),t.end=l(t.end),t.colorInit=!0),e.cssHooks[i].set(t.elem,t.start.transition(t.end,t.pos))}})},l.hook(o),e.cssHooks.borderColor={expand:function(e){var t={};return f(["Top","Right","Bottom","Left"],function(i,s){t["border"+s+"Color"]=e}),t}},a=e.Color.names={aqua:"#00ffff",black:"#000000",blue:"#0000ff",fuchsia:"#ff00ff",gray:"#808080",green:"#008000",lime:"#00ff00",maroon:"#800000",navy:"#000080",olive:"#808000",purple:"#800080",red:"#ff0000",silver:"#c0c0c0",teal:"#008080",white:"#ffffff",yellow:"#ffff00",transparent:[null,null,null,0],_default:"#ffffff"}}(p),function(){function t(t){var i,s,n=t.ownerDocument.defaultView?t.ownerDocument.defaultView.getComputedStyle(t,null):t.currentStyle,a={};if(n&&n.length&&n[0]&&n[n[0]])for(s=n.length;s--;)i=n[s],"string"==typeof n[i]&&(a[e.camelCase(i)]=n[i]);else for(i in n)"string"==typeof n[i]&&(a[i]=n[i]);return a}function i(t,i){var s,a,o={};for(s in i)a=i[s],t[s]!==a&&(n[s]||(e.fx.step[s]||!isNaN(parseFloat(a)))&&(o[s]=a));return o}var s=["add","remove","toggle"],n={border:1,borderBottom:1,borderColor:1,borderLeft:1,borderRight:1,borderTop:1,borderWidth:1,margin:1,padding:1};e.each(["borderLeftStyle","borderRightStyle","borderBottomStyle","borderTopStyle"],function(t,i){e.fx.step[i]=function(e){("none"!==e.end&&!e.setAttr||1===e.pos&&!e.setAttr)&&(p.style(e.elem,i,e.end),e.setAttr=!0)}}),e.fn.addBack||(e.fn.addBack=function(e){return this.add(null==e?this.prevObject:this.prevObject.filter(e))}),e.effects.animateClass=function(n,a,o,r){var h=e.speed(a,o,r);return this.queue(function(){var a,o=e(this),r=o.attr("class")||"",l=h.children?o.find("*").addBack():o;l=l.map(function(){var i=e(this);return{el:i,start:t(this)}}),a=function(){e.each(s,function(e,t){n[t]&&o[t+"Class"](n[t])})},a(),l=l.map(function(){return this.end=t(this.el[0]),this.diff=i(this.start,this.end),this}),o.attr("class",r),l=l.map(function(){var t=this,i=e.Deferred(),s=e.extend({},h,{queue:!1,complete:function(){i.resolve(t)}});return this.el.animate(this.diff,s),i.promise()}),e.when.apply(e,l.get()).done(function(){a(),e.each(arguments,function(){var t=this.el;e.each(this.diff,function(e){t.css(e,"")})}),h.complete.call(o[0])})})},e.fn.extend({addClass:function(t){return function(i,s,n,a){return s?e.effects.animateClass.call(this,{add:i},s,n,a):t.apply(this,arguments)}}(e.fn.addClass),removeClass:function(t){return function(i,s,n,a){return arguments.length>1?e.effects.animateClass.call(this,{remove:i},s,n,a):t.apply(this,arguments)}}(e.fn.removeClass),toggleClass:function(t){return function(i,s,n,a,o){return"boolean"==typeof s||void 0===s?n?e.effects.animateClass.call(this,s?{add:i}:{remove:i},n,a,o):t.apply(this,arguments):e.effects.animateClass.call(this,{toggle:i},s,n,a)}}(e.fn.toggleClass),switchClass:function(t,i,s,n,a){return e.effects.animateClass.call(this,{add:i,remove:t},s,n,a)}})}(),function(){function t(t,i,s,n){return e.isPlainObject(t)&&(i=t,t=t.effect),t={effect:t},null==i&&(i={}),e.isFunction(i)&&(n=i,s=null,i={}),("number"==typeof i||e.fx.speeds[i])&&(n=s,s=i,i={}),e.isFunction(s)&&(n=s,s=null),i&&e.extend(t,i),s=s||i.duration,t.duration=e.fx.off?0:"number"==typeof s?s:s in e.fx.speeds?e.fx.speeds[s]:e.fx.speeds._default,t.complete=n||i.complete,t}function i(t){return!t||"number"==typeof t||e.fx.speeds[t]?!0:"string"!=typeof t||e.effects.effect[t]?e.isFunction(t)?!0:"object"!=typeof t||t.effect?!1:!0:!0}e.extend(e.effects,{version:"1.11.4",save:function(e,t){for(var i=0;t.length>i;i++)null!==t[i]&&e.data(c+t[i],e[0].style[t[i]])},restore:function(e,t){var i,s;for(s=0;t.length>s;s++)null!==t[s]&&(i=e.data(c+t[s]),void 0===i&&(i=""),e.css(t[s],i))},setMode:function(e,t){return"toggle"===t&&(t=e.is(":hidden")?"show":"hide"),t},getBaseline:function(e,t){var i,s;switch(e[0]){case"top":i=0;break;case"middle":i=.5;break;case"bottom":i=1;break;default:i=e[0]/t.height}switch(e[1]){case"left":s=0;break;case"center":s=.5;break;case"right":s=1;break;default:s=e[1]/t.width}return{x:s,y:i}},createWrapper:function(t){if(t.parent().is(".ui-effects-wrapper"))return t.parent();var i={width:t.outerWidth(!0),height:t.outerHeight(!0),"float":t.css("float")},s=e("<div></div>").addClass("ui-effects-wrapper").css({fontSize:"100%",background:"transparent",border:"none",margin:0,padding:0}),n={width:t.width(),height:t.height()},a=document.activeElement;try{a.id}catch(o){a=document.body}return t.wrap(s),(t[0]===a||e.contains(t[0],a))&&e(a).focus(),s=t.parent(),"static"===t.css("position")?(s.css({position:"relative"}),t.css({position:"relative"})):(e.extend(i,{position:t.css("position"),zIndex:t.css("z-index")}),e.each(["top","left","bottom","right"],function(e,s){i[s]=t.css(s),isNaN(parseInt(i[s],10))&&(i[s]="auto")}),t.css({position:"relative",top:0,left:0,right:"auto",bottom:"auto"})),t.css(n),s.css(i).show()},removeWrapper:function(t){var i=document.activeElement;return t.parent().is(".ui-effects-wrapper")&&(t.parent().replaceWith(t),(t[0]===i||e.contains(t[0],i))&&e(i).focus()),t},setTransition:function(t,i,s,n){return n=n||{},e.each(i,function(e,i){var a=t.cssUnit(i);a[0]>0&&(n[i]=a[0]*s+a[1])}),n}}),e.fn.extend({effect:function(){function i(t){function i(){e.isFunction(a)&&a.call(n[0]),e.isFunction(t)&&t()}var n=e(this),a=s.complete,r=s.mode;(n.is(":hidden")?"hide"===r:"show"===r)?(n[r](),i()):o.call(n[0],s,i)}var s=t.apply(this,arguments),n=s.mode,a=s.queue,o=e.effects.effect[s.effect];return e.fx.off||!o?n?this[n](s.duration,s.complete):this.each(function(){s.complete&&s.complete.call(this)}):a===!1?this.each(i):this.queue(a||"fx",i)},show:function(e){return function(s){if(i(s))return e.apply(this,arguments);var n=t.apply(this,arguments);return n.mode="show",this.effect.call(this,n)}}(e.fn.show),hide:function(e){return function(s){if(i(s))return e.apply(this,arguments);var n=t.apply(this,arguments);return n.mode="hide",this.effect.call(this,n)}}(e.fn.hide),toggle:function(e){return function(s){if(i(s)||"boolean"==typeof s)return e.apply(this,arguments);var n=t.apply(this,arguments);return n.mode="toggle",this.effect.call(this,n)}}(e.fn.toggle),cssUnit:function(t){var i=this.css(t),s=[];return e.each(["em","px","%","pt"],function(e,t){i.indexOf(t)>0&&(s=[parseFloat(i),t])}),s}})}(),function(){var t={};e.each(["Quad","Cubic","Quart","Quint","Expo"],function(e,i){t[i]=function(t){return Math.pow(t,e+2)}}),e.extend(t,{Sine:function(e){return 1-Math.cos(e*Math.PI/2)},Circ:function(e){return 1-Math.sqrt(1-e*e)},Elastic:function(e){return 0===e||1===e?e:-Math.pow(2,8*(e-1))*Math.sin((80*(e-1)-7.5)*Math.PI/15)},Back:function(e){return e*e*(3*e-2)},Bounce:function(e){for(var t,i=4;((t=Math.pow(2,--i))-1)/11>e;);return 1/Math.pow(4,3-i)-7.5625*Math.pow((3*t-2)/22-e,2)}}),e.each(t,function(t,i){e.easing["easeIn"+t]=i,e.easing["easeOut"+t]=function(e){return 1-i(1-e)},e.easing["easeInOut"+t]=function(e){return.5>e?i(2*e)/2:1-i(-2*e+2)/2}})}(),e.effects,e.effects.effect.blind=function(t,i){var s,n,a,o=e(this),r=/up|down|vertical/,h=/up|left|vertical|horizontal/,l=["position","top","bottom","left","right","height","width"],u=e.effects.setMode(o,t.mode||"hide"),d=t.direction||"up",c=r.test(d),p=c?"height":"width",f=c?"top":"left",m=h.test(d),g={},v="show"===u;o.parent().is(".ui-effects-wrapper")?e.effects.save(o.parent(),l):e.effects.save(o,l),o.show(),s=e.effects.createWrapper(o).css({overflow:"hidden"}),n=s[p](),a=parseFloat(s.css(f))||0,g[p]=v?n:0,m||(o.css(c?"bottom":"right",0).css(c?"top":"left","auto").css({position:"absolute"}),g[f]=v?a:n+a),v&&(s.css(p,0),m||s.css(f,a+n)),s.animate(g,{duration:t.duration,easing:t.easing,queue:!1,complete:function(){"hide"===u&&o.hide(),e.effects.restore(o,l),e.effects.removeWrapper(o),i()
}})},e.effects.effect.bounce=function(t,i){var s,n,a,o=e(this),r=["position","top","bottom","left","right","height","width"],h=e.effects.setMode(o,t.mode||"effect"),l="hide"===h,u="show"===h,d=t.direction||"up",c=t.distance,p=t.times||5,f=2*p+(u||l?1:0),m=t.duration/f,g=t.easing,v="up"===d||"down"===d?"top":"left",y="up"===d||"left"===d,b=o.queue(),_=b.length;for((u||l)&&r.push("opacity"),e.effects.save(o,r),o.show(),e.effects.createWrapper(o),c||(c=o["top"===v?"outerHeight":"outerWidth"]()/3),u&&(a={opacity:1},a[v]=0,o.css("opacity",0).css(v,y?2*-c:2*c).animate(a,m,g)),l&&(c/=Math.pow(2,p-1)),a={},a[v]=0,s=0;p>s;s++)n={},n[v]=(y?"-=":"+=")+c,o.animate(n,m,g).animate(a,m,g),c=l?2*c:c/2;l&&(n={opacity:0},n[v]=(y?"-=":"+=")+c,o.animate(n,m,g)),o.queue(function(){l&&o.hide(),e.effects.restore(o,r),e.effects.removeWrapper(o),i()}),_>1&&b.splice.apply(b,[1,0].concat(b.splice(_,f+1))),o.dequeue()},e.effects.effect.clip=function(t,i){var s,n,a,o=e(this),r=["position","top","bottom","left","right","height","width"],h=e.effects.setMode(o,t.mode||"hide"),l="show"===h,u=t.direction||"vertical",d="vertical"===u,c=d?"height":"width",p=d?"top":"left",f={};e.effects.save(o,r),o.show(),s=e.effects.createWrapper(o).css({overflow:"hidden"}),n="IMG"===o[0].tagName?s:o,a=n[c](),l&&(n.css(c,0),n.css(p,a/2)),f[c]=l?a:0,f[p]=l?0:a/2,n.animate(f,{queue:!1,duration:t.duration,easing:t.easing,complete:function(){l||o.hide(),e.effects.restore(o,r),e.effects.removeWrapper(o),i()}})},e.effects.effect.drop=function(t,i){var s,n=e(this),a=["position","top","bottom","left","right","opacity","height","width"],o=e.effects.setMode(n,t.mode||"hide"),r="show"===o,h=t.direction||"left",l="up"===h||"down"===h?"top":"left",u="up"===h||"left"===h?"pos":"neg",d={opacity:r?1:0};e.effects.save(n,a),n.show(),e.effects.createWrapper(n),s=t.distance||n["top"===l?"outerHeight":"outerWidth"](!0)/2,r&&n.css("opacity",0).css(l,"pos"===u?-s:s),d[l]=(r?"pos"===u?"+=":"-=":"pos"===u?"-=":"+=")+s,n.animate(d,{queue:!1,duration:t.duration,easing:t.easing,complete:function(){"hide"===o&&n.hide(),e.effects.restore(n,a),e.effects.removeWrapper(n),i()}})},e.effects.effect.explode=function(t,i){function s(){b.push(this),b.length===d*c&&n()}function n(){p.css({visibility:"visible"}),e(b).remove(),m||p.hide(),i()}var a,o,r,h,l,u,d=t.pieces?Math.round(Math.sqrt(t.pieces)):3,c=d,p=e(this),f=e.effects.setMode(p,t.mode||"hide"),m="show"===f,g=p.show().css("visibility","hidden").offset(),v=Math.ceil(p.outerWidth()/c),y=Math.ceil(p.outerHeight()/d),b=[];for(a=0;d>a;a++)for(h=g.top+a*y,u=a-(d-1)/2,o=0;c>o;o++)r=g.left+o*v,l=o-(c-1)/2,p.clone().appendTo("body").wrap("<div></div>").css({position:"absolute",visibility:"visible",left:-o*v,top:-a*y}).parent().addClass("ui-effects-explode").css({position:"absolute",overflow:"hidden",width:v,height:y,left:r+(m?l*v:0),top:h+(m?u*y:0),opacity:m?0:1}).animate({left:r+(m?0:l*v),top:h+(m?0:u*y),opacity:m?1:0},t.duration||500,t.easing,s)},e.effects.effect.fade=function(t,i){var s=e(this),n=e.effects.setMode(s,t.mode||"toggle");s.animate({opacity:n},{queue:!1,duration:t.duration,easing:t.easing,complete:i})},e.effects.effect.fold=function(t,i){var s,n,a=e(this),o=["position","top","bottom","left","right","height","width"],r=e.effects.setMode(a,t.mode||"hide"),h="show"===r,l="hide"===r,u=t.size||15,d=/([0-9]+)%/.exec(u),c=!!t.horizFirst,p=h!==c,f=p?["width","height"]:["height","width"],m=t.duration/2,g={},v={};e.effects.save(a,o),a.show(),s=e.effects.createWrapper(a).css({overflow:"hidden"}),n=p?[s.width(),s.height()]:[s.height(),s.width()],d&&(u=parseInt(d[1],10)/100*n[l?0:1]),h&&s.css(c?{height:0,width:u}:{height:u,width:0}),g[f[0]]=h?n[0]:u,v[f[1]]=h?n[1]:0,s.animate(g,m,t.easing).animate(v,m,t.easing,function(){l&&a.hide(),e.effects.restore(a,o),e.effects.removeWrapper(a),i()})},e.effects.effect.highlight=function(t,i){var s=e(this),n=["backgroundImage","backgroundColor","opacity"],a=e.effects.setMode(s,t.mode||"show"),o={backgroundColor:s.css("backgroundColor")};"hide"===a&&(o.opacity=0),e.effects.save(s,n),s.show().css({backgroundImage:"none",backgroundColor:t.color||"#ffff99"}).animate(o,{queue:!1,duration:t.duration,easing:t.easing,complete:function(){"hide"===a&&s.hide(),e.effects.restore(s,n),i()}})},e.effects.effect.size=function(t,i){var s,n,a,o=e(this),r=["position","top","bottom","left","right","width","height","overflow","opacity"],h=["position","top","bottom","left","right","overflow","opacity"],l=["width","height","overflow"],u=["fontSize"],d=["borderTopWidth","borderBottomWidth","paddingTop","paddingBottom"],c=["borderLeftWidth","borderRightWidth","paddingLeft","paddingRight"],p=e.effects.setMode(o,t.mode||"effect"),f=t.restore||"effect"!==p,m=t.scale||"both",g=t.origin||["middle","center"],v=o.css("position"),y=f?r:h,b={height:0,width:0,outerHeight:0,outerWidth:0};"show"===p&&o.show(),s={height:o.height(),width:o.width(),outerHeight:o.outerHeight(),outerWidth:o.outerWidth()},"toggle"===t.mode&&"show"===p?(o.from=t.to||b,o.to=t.from||s):(o.from=t.from||("show"===p?b:s),o.to=t.to||("hide"===p?b:s)),a={from:{y:o.from.height/s.height,x:o.from.width/s.width},to:{y:o.to.height/s.height,x:o.to.width/s.width}},("box"===m||"both"===m)&&(a.from.y!==a.to.y&&(y=y.concat(d),o.from=e.effects.setTransition(o,d,a.from.y,o.from),o.to=e.effects.setTransition(o,d,a.to.y,o.to)),a.from.x!==a.to.x&&(y=y.concat(c),o.from=e.effects.setTransition(o,c,a.from.x,o.from),o.to=e.effects.setTransition(o,c,a.to.x,o.to))),("content"===m||"both"===m)&&a.from.y!==a.to.y&&(y=y.concat(u).concat(l),o.from=e.effects.setTransition(o,u,a.from.y,o.from),o.to=e.effects.setTransition(o,u,a.to.y,o.to)),e.effects.save(o,y),o.show(),e.effects.createWrapper(o),o.css("overflow","hidden").css(o.from),g&&(n=e.effects.getBaseline(g,s),o.from.top=(s.outerHeight-o.outerHeight())*n.y,o.from.left=(s.outerWidth-o.outerWidth())*n.x,o.to.top=(s.outerHeight-o.to.outerHeight)*n.y,o.to.left=(s.outerWidth-o.to.outerWidth)*n.x),o.css(o.from),("content"===m||"both"===m)&&(d=d.concat(["marginTop","marginBottom"]).concat(u),c=c.concat(["marginLeft","marginRight"]),l=r.concat(d).concat(c),o.find("*[width]").each(function(){var i=e(this),s={height:i.height(),width:i.width(),outerHeight:i.outerHeight(),outerWidth:i.outerWidth()};f&&e.effects.save(i,l),i.from={height:s.height*a.from.y,width:s.width*a.from.x,outerHeight:s.outerHeight*a.from.y,outerWidth:s.outerWidth*a.from.x},i.to={height:s.height*a.to.y,width:s.width*a.to.x,outerHeight:s.height*a.to.y,outerWidth:s.width*a.to.x},a.from.y!==a.to.y&&(i.from=e.effects.setTransition(i,d,a.from.y,i.from),i.to=e.effects.setTransition(i,d,a.to.y,i.to)),a.from.x!==a.to.x&&(i.from=e.effects.setTransition(i,c,a.from.x,i.from),i.to=e.effects.setTransition(i,c,a.to.x,i.to)),i.css(i.from),i.animate(i.to,t.duration,t.easing,function(){f&&e.effects.restore(i,l)})})),o.animate(o.to,{queue:!1,duration:t.duration,easing:t.easing,complete:function(){0===o.to.opacity&&o.css("opacity",o.from.opacity),"hide"===p&&o.hide(),e.effects.restore(o,y),f||("static"===v?o.css({position:"relative",top:o.to.top,left:o.to.left}):e.each(["top","left"],function(e,t){o.css(t,function(t,i){var s=parseInt(i,10),n=e?o.to.left:o.to.top;return"auto"===i?n+"px":s+n+"px"})})),e.effects.removeWrapper(o),i()}})},e.effects.effect.scale=function(t,i){var s=e(this),n=e.extend(!0,{},t),a=e.effects.setMode(s,t.mode||"effect"),o=parseInt(t.percent,10)||(0===parseInt(t.percent,10)?0:"hide"===a?0:100),r=t.direction||"both",h=t.origin,l={height:s.height(),width:s.width(),outerHeight:s.outerHeight(),outerWidth:s.outerWidth()},u={y:"horizontal"!==r?o/100:1,x:"vertical"!==r?o/100:1};n.effect="size",n.queue=!1,n.complete=i,"effect"!==a&&(n.origin=h||["middle","center"],n.restore=!0),n.from=t.from||("show"===a?{height:0,width:0,outerHeight:0,outerWidth:0}:l),n.to={height:l.height*u.y,width:l.width*u.x,outerHeight:l.outerHeight*u.y,outerWidth:l.outerWidth*u.x},n.fade&&("show"===a&&(n.from.opacity=0,n.to.opacity=1),"hide"===a&&(n.from.opacity=1,n.to.opacity=0)),s.effect(n)},e.effects.effect.puff=function(t,i){var s=e(this),n=e.effects.setMode(s,t.mode||"hide"),a="hide"===n,o=parseInt(t.percent,10)||150,r=o/100,h={height:s.height(),width:s.width(),outerHeight:s.outerHeight(),outerWidth:s.outerWidth()};e.extend(t,{effect:"scale",queue:!1,fade:!0,mode:n,complete:i,percent:a?o:100,from:a?h:{height:h.height*r,width:h.width*r,outerHeight:h.outerHeight*r,outerWidth:h.outerWidth*r}}),s.effect(t)},e.effects.effect.pulsate=function(t,i){var s,n=e(this),a=e.effects.setMode(n,t.mode||"show"),o="show"===a,r="hide"===a,h=o||"hide"===a,l=2*(t.times||5)+(h?1:0),u=t.duration/l,d=0,c=n.queue(),p=c.length;for((o||!n.is(":visible"))&&(n.css("opacity",0).show(),d=1),s=1;l>s;s++)n.animate({opacity:d},u,t.easing),d=1-d;n.animate({opacity:d},u,t.easing),n.queue(function(){r&&n.hide(),i()}),p>1&&c.splice.apply(c,[1,0].concat(c.splice(p,l+1))),n.dequeue()},e.effects.effect.shake=function(t,i){var s,n=e(this),a=["position","top","bottom","left","right","height","width"],o=e.effects.setMode(n,t.mode||"effect"),r=t.direction||"left",h=t.distance||20,l=t.times||3,u=2*l+1,d=Math.round(t.duration/u),c="up"===r||"down"===r?"top":"left",p="up"===r||"left"===r,f={},m={},g={},v=n.queue(),y=v.length;for(e.effects.save(n,a),n.show(),e.effects.createWrapper(n),f[c]=(p?"-=":"+=")+h,m[c]=(p?"+=":"-=")+2*h,g[c]=(p?"-=":"+=")+2*h,n.animate(f,d,t.easing),s=1;l>s;s++)n.animate(m,d,t.easing).animate(g,d,t.easing);n.animate(m,d,t.easing).animate(f,d/2,t.easing).queue(function(){"hide"===o&&n.hide(),e.effects.restore(n,a),e.effects.removeWrapper(n),i()}),y>1&&v.splice.apply(v,[1,0].concat(v.splice(y,u+1))),n.dequeue()},e.effects.effect.slide=function(t,i){var s,n=e(this),a=["position","top","bottom","left","right","width","height"],o=e.effects.setMode(n,t.mode||"show"),r="show"===o,h=t.direction||"left",l="up"===h||"down"===h?"top":"left",u="up"===h||"left"===h,d={};e.effects.save(n,a),n.show(),s=t.distance||n["top"===l?"outerHeight":"outerWidth"](!0),e.effects.createWrapper(n).css({overflow:"hidden"}),r&&n.css(l,u?isNaN(s)?"-"+s:-s:s),d[l]=(r?u?"+=":"-=":u?"-=":"+=")+s,n.animate(d,{queue:!1,duration:t.duration,easing:t.easing,complete:function(){"hide"===o&&n.hide(),e.effects.restore(n,a),e.effects.removeWrapper(n),i()}})},e.effects.effect.transfer=function(t,i){var s=e(this),n=e(t.to),a="fixed"===n.css("position"),o=e("body"),r=a?o.scrollTop():0,h=a?o.scrollLeft():0,l=n.offset(),u={top:l.top-r,left:l.left-h,height:n.innerHeight(),width:n.innerWidth()},d=s.offset(),c=e("<div class='ui-effects-transfer'></div>").appendTo(document.body).addClass(t.className).css({top:d.top-r,left:d.left-h,height:s.innerHeight(),width:s.innerWidth(),position:a?"fixed":"absolute"}).animate(u,t.duration,t.easing,function(){c.remove(),i()})}});
```
