## Supplementary material for "Characterizing and inferring quantitative cell cycle phase in single-cell RNA-seq data analysis": Source code of our analysis, including scripts and data necessary to reproduce the work.: README

This a jQuery UI custom build, downloaded from:
http://jqueryui.com/download/#!version=1.11.4&components=1111111111110111111111111111111111111

It includes all components except the datepicker, because it conflicts with
bootstrap-datepicker that is packaged with Shiny.

The copy of jQuery that is bundled with the download, under external/, is not
included because Shiny already has its own copy of jQuery.
