## Supplementary material for "Characterizing and inferring quantitative cell cycle phase in single-cell RNA-seq data analysis": Source code of our analysis, including scripts and data necessary to reproduce the work.: codefolding.js

```
window.initializeCodeFolding = function(show) {

  // handlers for show-all and hide all
  $("#rmd-show-all-code").click(function() {
    $('div.r-code-collapse').each(function() {
      $(this).collapse('show');
    });
  });
  $("#rmd-hide-all-code").click(function() {
    $('div.r-code-collapse').each(function() {
      $(this).collapse('hide');
    });
  });

  // index for unique code element ids
  var currentIndex = 1;

  // select all R code blocks
  var rCodeBlocks = $('pre.r, pre.python, pre.bash, pre.sql, pre.cpp, pre.stan');
  rCodeBlocks.each(function() {

    // create a collapsable div to wrap the code in
    var div = $('<div class="collapse r-code-collapse"></div>');
    if (show)
      div.addClass('in');
    var id = 'rcode-643E0F36' + currentIndex++;
    div.attr('id', id);
    $(this).before(div);
    $(this).detach().appendTo(div);

    // add a show code button right above
    var showCodeText = $('<span>' + (show ? 'Hide' : 'Code') + '</span>');
    var showCodeButton = $('<button type="button" class="btn btn-default btn-xs code-folding-btn pull-right"></button>');
    showCodeButton.append(showCodeText);
    showCodeButton
        .attr('data-toggle', 'collapse')
        .attr('data-target', '#' + id)
        .attr('aria-expanded', show)
        .attr('aria-controls', id);

    var buttonRow = $('<div class="row"></div>');
    var buttonCol = $('<div class="col-md-12"></div>');

    buttonCol.append(showCodeButton);
    buttonRow.append(buttonCol);

    div.before(buttonRow);

    // update state of button on show/hide
    div.on('hidden.bs.collapse', function () {
      showCodeText.text('Code');
    });
    div.on('show.bs.collapse', function () {
      showCodeText.text('Hide');
    });
  });

}
```
