## Supplementary material for "Characterizing and inferring quantitative cell cycle phase in single-cell RNA-seq data analysis": Source code of our analysis, including scripts and data necessary to reproduce the work.: sourceembed.js

```
window.initializeSourceEmbed = function(filename) {
  $("#rmd-download-source").click(function() {
    var src = $("#rmd-source-code").html();
    var a = document.createElement('a');
    a.href = "data:text/x-r-markdown;base64," + src;
    a.download = filename;
    document.body.appendChild(a);
    a.click();
    document.body.removeChild(a);
  });
};
```
