## Supplementary material for "Characterizing and inferring quantitative cell cycle phase in single-cell RNA-seq data analysis": Source code of our analysis, including scripts and data necessary to reproduce the work.: tabsets.js

```
/**
 * jQuery Plugin: Sticky Tabs
 *
 * @author Aidan Lister <>
 * adapted by Ruben Arslan to activate parent tabs too
 * http://www.aidanlister.com/2014/03/persisting-the-tab-state-in-bootstrap/
 */
(function($) {
  "use strict";
  $.fn.rmarkdownStickyTabs = function() {
    var context = this;
    // Show the tab corresponding with the hash in the URL, or the first tab
    var showStuffFromHash = function() {
      var hash = window.location.hash;
      var selector = hash ? 'a[href="' + hash + '"]' : 'li.active > a';
      var $selector = $(selector, context);
      if($selector.data('toggle') === "tab") {
        $selector.tab('show');
        // walk up the ancestors of this element, show any hidden tabs
        $selector.parents('.section.tabset').each(function(i, elm) {
          var link = $('a[href="#' + $(elm).attr('id') + '"]');
          if(link.data('toggle') === "tab") {
            link.tab("show");
          }
        });
      }
    };


    // Set the correct tab when the page loads
    showStuffFromHash(context);

    // Set the correct tab when a user uses their back/forward button
    $(window).on('hashchange', function() {
      showStuffFromHash(context);
    });

    // Change the URL when tabs are clicked
    $('a', context).on('click', function(e) {
      history.pushState(null, null, this.href);
      showStuffFromHash(context);
    });

    return this;
  };
}(jQuery));

window.buildTabsets = function(tocID) {

  // build a tabset from a section div with the .tabset class
  function buildTabset(tabset) {

    // check for fade and pills options
    var fade = tabset.hasClass("tabset-fade");
    var pills = tabset.hasClass("tabset-pills");
    var navClass = pills ? "nav-pills" : "nav-tabs";

    // determine the heading level of the tabset and tabs
    var match = tabset.attr('class').match(/level(\d) /);
    if (match === null)
      return;
    var tabsetLevel = Number(match[1]);
    var tabLevel = tabsetLevel + 1;

    // find all subheadings immediately below
    var tabs = tabset.find("div.section.level" + tabLevel);
    if (!tabs.length)
      return;

    // create tablist and tab-content elements
    var tabList = $('<ul class="nav ' + navClass + '" role="tablist"></ul>');
    $(tabs[0]).before(tabList);
    var tabContent = $('<div class="tab-content"></div>');
    $(tabs[0]).before(tabContent);

    // build the tabset
    var activeTab = 0;
    tabs.each(function(i) {

      // get the tab div
      var tab = $(tabs[i]);

      // get the id then sanitize it for use with bootstrap tabs
      var id = tab.attr('id');

      // see if this is marked as the active tab
      if (tab.hasClass('active'))
        activeTab = i;

      // remove any table of contents entries associated with
      // this ID (since we'll be removing the heading element)
      $("div#" + tocID + " li a[href='#" + id + "']").parent().remove();

      // sanitize the id for use with bootstrap tabs
      id = id.replace(/[.\/?&!#<>]/g, '').replace(/\s/g, '_');
      tab.attr('id', id);

      // get the heading element within it, grab it's text, then remove it
      var heading = tab.find('h' + tabLevel + ':first');
      var headingText = heading.html();
      heading.remove();

      // build and append the tab list item
      var a = $('<a role="tab" data-toggle="tab">' + headingText + '</a>');
      a.attr('href', '#' + id);
      a.attr('aria-controls', id);
      var li = $('<li role="presentation"></li>');
      li.append(a);
      tabList.append(li);

      // set it's attributes
      tab.attr('role', 'tabpanel');
      tab.addClass('tab-pane');
      tab.addClass('tabbed-pane');
      if (fade)
        tab.addClass('fade');

      // move it into the tab content div
      tab.detach().appendTo(tabContent);
    });

    // set active tab
    $(tabList.children('li')[activeTab]).addClass('active');
    var active = $(tabContent.children('div.section')[activeTab]);
    active.addClass('active');
    if (fade)
      active.addClass('in');

    if (tabset.hasClass("tabset-sticky"))
      tabset.rmarkdownStickyTabs();
  }

  // convert section divs with the .tabset class to tabsets
  var tabsets = $("div.section.tabset");
  tabsets.each(function(i) {
    buildTabset($(tabsets[i]));
  });
};
```
