## Supplementary material for "Characterizing and inferring quantitative cell cycle phase in single-cell RNA-seq data analysis": Source code of our analysis, including scripts and data necessary to reproduce the work.: jquery.tocify.js

```
/* jquery Tocify - v1.9.1 - 2013-10-22
 * http://www.gregfranko.com/jquery.tocify.js/
 * Copyright (c) 2013 Greg Franko; Licensed MIT */

// Immediately-Invoked Function Expression (IIFE) [Ben Alman Blog Post](http://benalman.com/news/2010/11/immediately-invoked-function-expression/) that calls another IIFE that contains all of the plugin logic.  I used this pattern so that anyone viewing this code would not have to scroll to the bottom of the page to view the local parameters that were passed to the main IIFE.
(function(tocify) {

    // ECMAScript 5 Strict Mode: [John Resig Blog Post](http://ejohn.org/blog/ecmascript-5-strict-mode-json-and-more/)
    "use strict";

    // Calls the second IIFE and locally passes in the global jQuery, window, and document objects
    tocify(window.jQuery, window, document);

  }

  // Locally passes in `jQuery`, the `window` object, the `document` object, and an `undefined` variable.  The `jQuery`, `window` and `document` objects are passed in locally, to improve performance, since javascript first searches for a variable match within the local variables set before searching the global variables set.  All of the global variables are also passed in locally to be minifier friendly. `undefined` can be passed in locally, because it is not a reserved word in JavaScript.
  (function($, window, document, undefined) {

    // ECMAScript 5 Strict Mode: [John Resig Blog Post](http://ejohn.org/blog/ecmascript-5-strict-mode-json-and-more/)
    "use strict";

    var tocClassName = "tocify",
      tocClass = "." + tocClassName,
      tocFocusClassName = "tocify-focus",
      tocHoverClassName = "tocify-hover",
      hideTocClassName = "tocify-hide",
      hideTocClass = "." + hideTocClassName,
      headerClassName = "tocify-header",
      headerClass = "." + headerClassName,
      subheaderClassName = "tocify-subheader",
      subheaderClass = "." + subheaderClassName,
      itemClassName = "tocify-item",
      itemClass = "." + itemClassName,
      extendPageClassName = "tocify-extend-page",
      extendPageClass = "." + extendPageClassName;

    // Calling the jQueryUI Widget Factory Method
    $.widget("toc.tocify", {

      //Plugin version
      version: "1.9.1",

      // These options will be used as defaults
      options: {

        // **context**: Accepts String: Any jQuery selector
        // The container element that holds all of the elements used to generate the table of contents
        context: "body",

        // **ignoreSelector**: Accepts String: Any jQuery selector
        // A selector to any element that would be matched by selectors that you wish to be ignored
        ignoreSelector: null,

        // **selectors**: Accepts an Array of Strings: Any jQuery selectors
        // The element's used to generate the table of contents.  The order is very important since it will determine the table of content's nesting structure
        selectors: "h1, h2, h3",

        // **showAndHide**: Accepts a boolean: true or false
        // Used to determine if elements should be shown and hidden
        showAndHide: true,

        // **showEffect**: Accepts String: "none", "fadeIn", "show", or "slideDown"
        // Used to display any of the table of contents nested items
        showEffect: "slideDown",

        // **showEffectSpeed**: Accepts Number (milliseconds) or String: "slow", "medium", or "fast"
        // The time duration of the show animation
        showEffectSpeed: "medium",

        // **hideEffect**: Accepts String: "none", "fadeOut", "hide", or "slideUp"
        // Used to hide any of the table of contents nested items
        hideEffect: "slideUp",

        // **hideEffectSpeed**: Accepts Number (milliseconds) or String: "slow", "medium", or "fast"
        // The time duration of the hide animation
        hideEffectSpeed: "medium",

        // **smoothScroll**: Accepts a boolean: true or false
        // Determines if a jQuery animation should be used to scroll to specific table of contents items on the page
        smoothScroll: true,

        // **smoothScrollSpeed**: Accepts Number (milliseconds) or String: "slow", "medium", or "fast"
        // The time duration of the smoothScroll animation
        smoothScrollSpeed: "medium",

        // **scrollTo**: Accepts Number (pixels)
        // The amount of space between the top of page and the selected table of contents item after the page has been scrolled
        scrollTo: 0,

        // **showAndHideOnScroll**: Accepts a boolean: true or false
        // Determines if table of contents nested items should be shown and hidden while scrolling
        showAndHideOnScroll: true,

        // **highlightOnScroll**: Accepts a boolean: true or false
        // Determines if table of contents nested items should be highlighted (set to a different color) while scrolling
        highlightOnScroll: true,

        // **highlightOffset**: Accepts a number
        // The offset distance in pixels to trigger the next active table of contents item
        highlightOffset: 40,

        // **theme**: Accepts a string: "bootstrap", "jqueryui", or "none"
        // Determines if Twitter Bootstrap, jQueryUI, or Tocify classes should be added to the table of contents
        theme: "bootstrap",

        // **extendPage**: Accepts a boolean: true or false
        // If a user scrolls to the bottom of the page and the page is not tall enough to scroll to the last table of contents item, then the page height is increased
        extendPage: true,

        // **extendPageOffset**: Accepts a number: pixels
        // How close to the bottom of the page a user must scroll before the page is extended
        extendPageOffset: 100,

        // **history**: Accepts a boolean: true or false
        // Adds a hash to the page url to maintain history
        history: true,

        // **scrollHistory**: Accepts a boolean: true or false
        // Adds a hash to the page url, to maintain history, when scrolling to a TOC item
        scrollHistory: false,

        // **hashGenerator**: How the hash value (the anchor segment of the URL, following the
        // # character) will be generated.
        //
        // "compact" (default) - #CompressesEverythingTogether
        // "pretty" - #looks-like-a-nice-url-and-is-easily-readable
        // function(text, element){} - Your own hash generation function that accepts the text as an
        // argument, and returns the hash value.
        hashGenerator: "compact",

        // **highlightDefault**: Accepts a boolean: true or false
        // Set's the first TOC item as active if no other TOC item is active.
        highlightDefault: true

      },

      // _Create
      // -------
      //      Constructs the plugin.  Only called once.
      _create: function() {

        var self = this;

        self.extendPageScroll = true;

        // Internal array that keeps track of all TOC items (Helps to recognize if there are duplicate TOC item strings)
        self.items = [];

        // Generates the HTML for the dynamic table of contents
        self._generateToc();

        // Adds CSS classes to the newly generated table of contents HTML
        self._addCSSClasses();

        self.webkit = (function() {

          for (var prop in window) {

            if (prop) {

              if (prop.toLowerCase().indexOf("webkit") !== -1) {

                return true;

              }

            }

          }

          return false;

        }());

        // Adds jQuery event handlers to the newly generated table of contents
        self._setEventHandlers();

        // Binding to the Window load event to make sure the correct scrollTop is calculated
        $(window).load(function() {

          // Sets the active TOC item
          self._setActiveElement(true);

          // Once all animations on the page are complete, this callback function will be called
          $("html, body").promise().done(function() {

            setTimeout(function() {

              self.extendPageScroll = false;

            }, 0);

          });

        });

      },

      // _generateToc
      // ------------
      //      Generates the HTML for the dynamic table of contents
      _generateToc: function() {

        // _Local variables_

        // Stores the plugin context in the self variable
        var self = this,

          // All of the HTML tags found within the context provided (i.e. body) that match the top level jQuery selector above
          firstElem,

          // Instantiated variable that will store the top level newly created unordered list DOM element
          ul,
          ignoreSelector = self.options.ignoreSelector;


        // Determine the element to start the toc with
        // get all the top level selectors
        firstElem = [];
        var selectors = this.options.selectors.replace(/ /g, "").split(",");
        // find the first set that have at least one non-ignored element
        for(var i = 0; i < selectors.length; i++) {
          var foundSelectors = $(this.options.context).find(selectors[i]);
          for (var s = 0; s < foundSelectors.length; s++) {
            if (!$(foundSelectors[s]).is(ignoreSelector)) {
              firstElem = foundSelectors;
              break;
            }
          }
          if (firstElem.length> 0)
            break;
        }

        if (!firstElem.length) {

          self.element.addClass(hideTocClassName);

          return;

        }

        self.element.addClass(tocClassName);

        // Loops through each top level selector
        firstElem.each(function(index) {

          //If the element matches the ignoreSelector then we skip it
          if ($(this).is(ignoreSelector)) {
            return;
          }

          // Creates an unordered list HTML element and adds a dynamic ID and standard class name
          ul = $("<ul/>", {
            "id": headerClassName + index,
            "class": headerClassName
          }).

          // Appends a top level list item HTML element to the previously created HTML header
          append(self._nestElements($(this), index));

          // Add the created unordered list element to the HTML element calling the plugin
          self.element.append(ul);

          // Finds all of the HTML tags between the header and subheader elements
          $(this).nextUntil(this.nodeName.toLowerCase()).each(function() {

            // If there are no nested subheader elemements
            if ($(this).find(self.options.selectors).length === 0) {

              // Loops through all of the subheader elements
              $(this).filter(self.options.selectors).each(function() {

                //If the element matches the ignoreSelector then we skip it
                if ($(this).is(ignoreSelector)) {
                  return;
                }

                self._appendSubheaders.call(this, self, ul);

              });

            }

            // If there are nested subheader elements
            else {

              // Loops through all of the subheader elements
              $(this).find(self.options.selectors).each(function() {

                //If the element matches the ignoreSelector then we skip it
                if ($(this).is(ignoreSelector)) {
                  return;
                }

                self._appendSubheaders.call(this, self, ul);

              });

            }

          });

        });

      },

      _setActiveElement: function(pageload) {

        var self = this,

          hash = window.location.hash.substring(1),

          elem = self.element.find('li[data-unique="' + hash + '"]');

        if (hash.length) {

          // Removes highlighting from all of the list item's
          self.element.find("." + self.focusClass).removeClass(self.focusClass);

          // Highlights the current list item that was clicked
          elem.addClass(self.focusClass);

          // Triggers the click event on the currently focused TOC item
          elem.click();

        } else {

          // Removes highlighting from all of the list item's
          self.element.find("." + self.focusClass).removeClass(self.focusClass);

          if (!hash.length && pageload && self.options.highlightDefault) {

            // Highlights the first TOC item if no other items are highlighted
            self.element.find(itemClass).first().addClass(self.focusClass);

          }

        }

        return self;

      },

      // _nestElements
      // -------------
      //      Helps create the table of contents list by appending nested list items
      _nestElements: function(self, index) {

        var arr, item, hashValue;

        arr = $.grep(this.items, function(item) {

          return item === self.text();

        });

        // If there is already a duplicate TOC item
        if (arr.length) {

          // Adds the current TOC item text and index (for slight randomization) to the internal array
          this.items.push(self.text() + index);

        }

        // If there not a duplicate TOC item
        else {

          // Adds the current TOC item text to the internal array
          this.items.push(self.text());

        }

        hashValue = this._generateHashValue(arr, self, index);

        // Appends a list item HTML element to the last unordered list HTML element found within the HTML element calling the plugin
        item = $("<li/>", {

          // Sets a common class name to the list item
          "class": itemClassName,

          "data-unique": hashValue

        });

        if (this.options.theme !== "bootstrap3") {

          item.append($("<a/>", {

            "text": self.text()

          }));

        } else {

          item.text(self.text());

        }

        // Adds an HTML anchor tag before the currently traversed HTML element
        self.before($("<div/>", {

          // Sets a name attribute on the anchor tag to the text of the currently traversed HTML element (also making sure that all whitespace is replaced with an underscore)
          "name": hashValue,

          "data-unique": hashValue

        }));

        return item;

      },

      // _generateHashValue
      // ------------------
      //      Generates the hash value that will be used to refer to each item.
      _generateHashValue: function(arr, self, index) {

        var hashValue = "",
          hashGeneratorOption = this.options.hashGenerator;

        if (hashGeneratorOption === "pretty") {

          // prettify the text
          hashValue = self.text().toLowerCase().replace(/\s/g, "-");

          // fix double hyphens
          while (hashValue.indexOf("--") > -1) {
            hashValue = hashValue.replace(/--/g, "-");
          }

          // fix colon-space instances
          while (hashValue.indexOf(":-") > -1) {
            hashValue = hashValue.replace(/:-/g, "-");
          }

        } else if (typeof hashGeneratorOption === "function") {

          // call the function
          hashValue = hashGeneratorOption(self.text(), self);

        } else {

          // compact - the default
          hashValue = self.text().replace(/\s/g, "");

        }

        // add the index if we need to
        if (arr.length) {
          hashValue += "" + index;
        }

        // return the value
        return hashValue;

      },

      // _appendElements
      // ---------------
      //      Helps create the table of contents list by appending subheader elements

      _appendSubheaders: function(self, ul) {

        // The current element index
        var index = $(this).index(self.options.selectors),

          // Finds the previous header DOM element
          previousHeader = $(self.options.selectors).eq(index - 1),

          currentTagName = +$(this).prop("tagName").charAt(1),

          previousTagName = +previousHeader.prop("tagName").charAt(1),

          lastSubheader;

        // If the current header DOM element is smaller than the previous header DOM element or the first subheader
        if (currentTagName < previousTagName) {

          // Selects the last unordered list HTML found within the HTML element calling the plugin
          self.element.find(subheaderClass + "[data-tag=" + currentTagName + "]").last().append(self._nestElements($(this), index));

        }

        // If the current header DOM element is the same type of header(eg. h4) as the previous header DOM element
        else if (currentTagName === previousTagName) {

          ul.find(itemClass).last().after(self._nestElements($(this), index));

        } else {

          // Selects the last unordered list HTML found within the HTML element calling the plugin
          ul.find(itemClass).last().

          // Appends an unorderedList HTML element to the dynamic `unorderedList` variable and sets a common class name
          after($("<ul/>", {

            "class": subheaderClassName,

            "data-tag": currentTagName

          })).next(subheaderClass).

          // Appends a list item HTML element to the last unordered list HTML element found within the HTML element calling the plugin
          append(self._nestElements($(this), index));
        }

      },

      // _setEventHandlers
      // ----------------
      //      Adds jQuery event handlers to the newly generated table of contents
      _setEventHandlers: function() {

        // _Local variables_

        // Stores the plugin context in the self variable
        var self = this,

          // Instantiates a new variable that will be used to hold a specific element's context
          $self,

          // Instantiates a new variable that will be used to determine the smoothScroll animation time duration
          duration;

        // Event delegation that looks for any clicks on list item elements inside of the HTML element calling the plugin
        this.element.on("click.tocify", "li", function(event) {

          if (self.options.history) {

            window.location.hash = $(this).attr("data-unique");

          }

          // Removes highlighting from all of the list item's
          self.element.find("." + self.focusClass).removeClass(self.focusClass);

          // Highlights the current list item that was clicked
          $(this).addClass(self.focusClass);

          // If the showAndHide option is true
          if (self.options.showAndHide) {

            var elem = $('li[data-unique="' + $(this).attr("data-unique") + '"]');

            self._triggerShow(elem);

          }

          self._scrollTo($(this));

        });

        // Mouseenter and Mouseleave event handlers for the list item's within the HTML element calling the plugin
        this.element.find("li").on({

          // Mouseenter event handler
          "mouseenter.tocify": function() {

            // Adds a hover CSS class to the current list item
            $(this).addClass(self.hoverClass);

            // Makes sure the cursor is set to the pointer icon
            $(this).css("cursor", "pointer");

          },

          // Mouseleave event handler
          "mouseleave.tocify": function() {

            if (self.options.theme !== "bootstrap") {

              // Removes the hover CSS class from the current list item
              $(this).removeClass(self.hoverClass);

            }

          }
        });

        // only attach handler if needed (expensive in IE)
        if (self.options.extendPage || self.options.highlightOnScroll || self.options.scrollHistory || self.options.showAndHideOnScroll) {
          // Window scroll event handler
          $(window).on("scroll.tocify", function() {

            // Once all animations on the page are complete, this callback function will be called
            $("html, body").promise().done(function() {

              // Local variables

              // Stores how far the user has scrolled
              var winScrollTop = $(window).scrollTop(),

                // Stores the height of the window
                winHeight = $(window).height(),

                // Stores the height of the document
                docHeight = $(document).height(),

                scrollHeight = $("body")[0].scrollHeight,

                // Instantiates a variable that will be used to hold a selected HTML element
                elem,

                lastElem,

                lastElemOffset,

                currentElem;

              if (self.options.extendPage) {

                // If the user has scrolled to the bottom of the page and the last toc item is not focused
                if ((self.webkit && winScrollTop >= scrollHeight - winHeight - self.options.extendPageOffset) || (!self.webkit && winHeight + winScrollTop > docHeight - self.options.extendPageOffset)) {

                  if (!$(extendPageClass).length) {

                    lastElem = $('div[data-unique="' + $(itemClass).last().attr("data-unique") + '"]');

                    if (!lastElem.length) return;

                    // Gets the top offset of the page header that is linked to the last toc item
                    lastElemOffset = lastElem.offset().top;

                    // Appends a div to the bottom of the page and sets the height to the difference of the window scrollTop and the last element's position top offset
                    $(self.options.context).append($("<div/>", {

                      "class": extendPageClassName,

                      "height": Math.abs(lastElemOffset - winScrollTop) + "px",

                      "data-unique": extendPageClassName

                    }));

                    if (self.extendPageScroll) {

                      currentElem = self.element.find('li.' + self.focusClass);

                      self._scrollTo($('div[data-unique="' + currentElem.attr("data-unique") + '"]'));

                    }

                  }

                }

              }

              // The zero timeout ensures the following code is run after the scroll events
              setTimeout(function() {

                // _Local variables_

                // Stores the distance to the closest anchor
                var closestAnchorDistance = null,

                  // Stores the index of the closest anchor
                  closestAnchorIdx = null,

                  // Keeps a reference to all anchors
                  anchors = $(self.options.context).find("div[data-unique]"),

                  anchorText;

                // Determines the index of the closest anchor
                anchors.each(function(idx) {
                  var distance = Math.abs(($(this).next().length ? $(this).next() : $(this)).offset().top - winScrollTop - self.options.highlightOffset);
                  if (closestAnchorDistance == null || distance < closestAnchorDistance) {
                    closestAnchorDistance = distance;
                    closestAnchorIdx = idx;
                  } else {
                    return false;
                  }
                });

                anchorText = $(anchors[closestAnchorIdx]).attr("data-unique");

                // Stores the list item HTML element that corresponds to the currently traversed anchor tag
                elem = $('li[data-unique="' + anchorText + '"]');

                // If the `highlightOnScroll` option is true and a next element is found
                if (self.options.highlightOnScroll && elem.length) {

                  // Removes highlighting from all of the list item's
                  self.element.find("." + self.focusClass).removeClass(self.focusClass);

                  // Highlights the corresponding list item
                  elem.addClass(self.focusClass);

                }

                if (self.options.scrollHistory) {

                  if (window.location.hash !== "#" + anchorText) {

                    window.location.replace("#" + anchorText);

                  }
                }

                // If the `showAndHideOnScroll` option is true
                if (self.options.showAndHideOnScroll && self.options.showAndHide) {

                  self._triggerShow(elem, true);

                }

              }, 0);

            });

          });
        }

      },

      // Show
      // ----
      //      Opens the current sub-header
      show: function(elem, scroll) {

        // Stores the plugin context in the `self` variable
        var self = this,
          element = elem;

        // If the sub-header is not already visible
        if (!elem.is(":visible")) {

          // If the current element does not have any nested subheaders, is not a header, and its parent is not visible
          if (!elem.find(subheaderClass).length && !elem.parent().is(headerClass) && !elem.parent().is(":visible")) {

            // Sets the current element to all of the subheaders within the current header
            elem = elem.parents(subheaderClass).add(elem);

          }

          // If the current element does not have any nested subheaders and is not a header
          else if (!elem.children(subheaderClass).length && !elem.parent().is(headerClass)) {

            // Sets the current element to the closest subheader
            elem = elem.closest(subheaderClass);

          }

          //Determines what jQuery effect to use
          switch (self.options.showEffect) {

            //Uses `no effect`
            case "none":

              elem.show();

              break;

              //Uses the jQuery `show` special effect
            case "show":

              elem.show(self.options.showEffectSpeed);

              break;

              //Uses the jQuery `slideDown` special effect
            case "slideDown":

              elem.slideDown(self.options.showEffectSpeed);

              break;

              //Uses the jQuery `fadeIn` special effect
            case "fadeIn":

              elem.fadeIn(self.options.showEffectSpeed);

              break;

              //If none of the above options were passed, then a `jQueryUI show effect` is expected
            default:

              elem.show();

              break;

          }

        }

        // If the current subheader parent element is a header
        if (elem.parent().is(headerClass)) {

          // Hides all non-active sub-headers
          self.hide($(subheaderClass).not(elem));

        }

        // If the current subheader parent element is not a header
        else {

          // Hides all non-active sub-headers
          self.hide($(subheaderClass).not(elem.closest(headerClass).find(subheaderClass).not(elem.siblings())));

        }

        // Maintains chainablity
        return self;

      },

      // Hide
      // ----
      //      Closes the current sub-header
      hide: function(elem) {

        // Stores the plugin context in the `self` variable
        var self = this;

        //Determines what jQuery effect to use
        switch (self.options.hideEffect) {

          // Uses `no effect`
          case "none":

            elem.hide();

            break;

            // Uses the jQuery `hide` special effect
          case "hide":

            elem.hide(self.options.hideEffectSpeed);

            break;

            // Uses the jQuery `slideUp` special effect
          case "slideUp":

            elem.slideUp(self.options.hideEffectSpeed);

            break;

            // Uses the jQuery `fadeOut` special effect
          case "fadeOut":

            elem.fadeOut(self.options.hideEffectSpeed);

            break;

            // If none of the above options were passed, then a `jqueryUI hide effect` is expected
          default:

            elem.hide();

            break;

        }

        // Maintains chainablity
        return self;
      },

      // _triggerShow
      // ------------
      //      Determines what elements get shown on scroll and click
      _triggerShow: function(elem, scroll) {

        var self = this;

        // If the current element's parent is a header element or the next element is a nested subheader element
        if (elem.parent().is(headerClass) || elem.next().is(subheaderClass)) {

          // Shows the next sub-header element
          self.show(elem.next(subheaderClass), scroll);

        }

        // If the current element's parent is a subheader element
        else if (elem.parent().is(subheaderClass)) {

          // Shows the parent sub-header element
          self.show(elem.parent(), scroll);

        }

        // Maintains chainability
        return self;

      },

      // _addCSSClasses
      // --------------
      //      Adds CSS classes to the newly generated table of contents HTML
      _addCSSClasses: function() {

        // If the user wants a jqueryUI theme
        if (this.options.theme === "jqueryui") {

          this.focusClass = "ui-state-default";

          this.hoverClass = "ui-state-hover";

          //Adds the default styling to the dropdown list
          this.element.addClass("ui-widget").find(".toc-title").addClass("ui-widget-header").end().find("li").addClass("ui-widget-content");

        }

        // If the user wants a twitterBootstrap theme
        else if (this.options.theme === "bootstrap") {

          this.element.find(headerClass + "," + subheaderClass).addClass("nav nav-list");

          this.focusClass = "active";

        }

        // If the user wants a twitterBootstrap theme
        else if (this.options.theme === "bootstrap3") {

          this.element.find(headerClass + "," + subheaderClass).addClass("list-group");

          this.element.find(itemClass).addClass("list-group-item");

          this.focusClass = "active";

        }

        // If a user does not want a prebuilt theme
        else {

          // Adds more neutral classes (instead of jqueryui)

          this.focusClass = tocFocusClassName;

          this.hoverClass = tocHoverClassName;

        }

        //Maintains chainability
        return this;

      },

      // setOption
      // ---------
      //      Sets a single Tocify option after the plugin is invoked
      setOption: function() {

        // Calls the jQueryUI Widget Factory setOption method
        $.Widget.prototype._setOption.apply(this, arguments);

      },

      // setOptions
      // ----------
      //      Sets a single or multiple Tocify options after the plugin is invoked
      setOptions: function() {

        // Calls the jQueryUI Widget Factory setOptions method
        $.Widget.prototype._setOptions.apply(this, arguments);

      },

      // _scrollTo
      // ---------
      //      Scrolls to a specific element
      _scrollTo: function(elem) {

        var self = this,
          duration = self.options.smoothScroll || 0,
          scrollTo = self.options.scrollTo,
          currentDiv = $('div[data-unique="' + elem.attr("data-unique") + '"]');

        if (!currentDiv.length) {

          return self;

        }

        // Once all animations on the page are complete, this callback function will be called
        $("html, body").promise().done(function() {

          // Animates the html and body element scrolltops
          $("html, body").animate({

            // Sets the jQuery `scrollTop` to the top offset of the HTML div tag that matches the current list item's `data-unique` tag
            "scrollTop": currentDiv.offset().top - ($.isFunction(scrollTo) ? scrollTo.call() : scrollTo) + "px"

          }, {

            // Sets the smoothScroll animation time duration to the smoothScrollSpeed option
            "duration": duration

          });

        });

        // Maintains chainability
        return self;

      }

    });

  })); //end of plugin
```
