## Supplementary material for "Characterizing and inferring quantitative cell cycle phase in single-cell RNA-seq data analysis": Source code of our analysis, including scripts and data necessary to reproduce the work.: totals.html

Sequencing depth per C1 chip


peco-paper

- Home
- About
- License

- Source code


### Sequencing depth per C1 chip

###### *John Blischak*

#### *2017-11-28*

workflowr

- Summary
- Checks
- Past versions

**Last updated:** 2020-01-23

**Checks:**  7  0

**Knit directory:** `peco-paper/`

| File | Version | Author | Date | Message |
| --- | --- | --- | --- | --- |
| html | d77bc47 | jhsiao999 | 2020-01-23 | Build site. |
| Rmd | 7c203e3 | jhsiao999 | 2020-01-23 | move totals.Rmd and change eset to sce |

---

#### Setup

```
library("dplyr")
library("DT")
library("ggplot2")
library("reshape2")
library("SingleCellExperiment")
theme_set(cowplot::theme_cowplot())
```

```
sce_raw = readRDS("data/sce-raw.rds")
anno = data.frame(colData(sce_raw))
anno$experiment = factor(anno$experiment)
```

#### Total sequencing depth

```
ggplot(anno, aes(x = raw, color = experiment)) +
  geom_density() +
  labs(x = "Number of raw sequences per single cell", y = "Number of cells",
       title = "Distribution of total raw sequences per single cell") +
  scale_color_discrete(name = "C1 chip")
```

Past versions of distribution-sequencing-depth-1.png

| Version | Author | Date |
| --- | --- | --- |
| d77bc47 | jhsiao999 | 2020-01-23 |

Mapped reads per cell

```
ggplot(anno, aes(x = experiment, y = mapped, color = experiment)) +
  geom_violin() + 
  geom_boxplot(alpha = .01, width = .2, position = position_dodge(width = .9)) +
  labs(x = "C1 chip", y = "Number of reads",
       title = "Number of mapped sequences per single cell") +
  theme(legend.title = element_blank(),
        axis.text.x = element_text(angle = 45, hjust = 1, vjust = 1))
```

Past versions of mapped-1.png

| Version | Author | Date |
| --- | --- | --- |
| d77bc47 | jhsiao999 | 2020-01-23 |

Sum of sequences across the 96 single cells per C1 chip.

```
total_per_experiment <- anno %>%
  group_by(experiment) %>%
  summarize(raw = sum(raw) / 10^6,
            mapped = sum(mapped) / 10^6,
            molecules = sum(molecules) / 10^6)
datatable(total_per_experiment,
          options = list(pageLength = nrow(total_per_experiment)),
          colnames = c("C1 chip", "Number of raw sequences",
                       "Number of mapped",
                       "Number of molecules"))
```

```
ggplot(melt(total_per_experiment, id.vars = "experiment",
            variable.name = "type", value.name = "count"),
       aes(x = experiment, y = count, color = type)) +
  geom_point() +
  labs(title = "Sequencing depth per C1 chip",
       x = "C1 chip", y = "Number of sequences") +
  theme(legend.title = element_blank(),
        axis.text.x = element_text(angle = 45, hjust = 1, vjust = 1))
```

Past versions of unnamed-chunk-1-1.png

| Version | Author | Date |
| --- | --- | --- |
| d77bc47 | jhsiao999 | 2020-01-23 |

  

Session information

```
sessionInfo()
```

```
R version 3.5.1 (2018-07-02)
Platform: x86_64-pc-linux-gnu (64-bit)
Running under: Scientific Linux 7.4 (Nitrogen)

Matrix products: default
BLAS/LAPACK: /software/openblas-0.2.19-el7-x86_64/lib/libopenblas_haswellp-r0.2.19.so

locale:
 [1] LC_CTYPE=en_US.UTF-8       LC_NUMERIC=C              
 [3] LC_TIME=en_US.UTF-8        LC_COLLATE=en_US.UTF-8    
 [5] LC_MONETARY=en_US.UTF-8    LC_MESSAGES=en_US.UTF-8   
 [7] LC_PAPER=en_US.UTF-8       LC_NAME=C                 
 [9] LC_ADDRESS=C               LC_TELEPHONE=C            
[11] LC_MEASUREMENT=en_US.UTF-8 LC_IDENTIFICATION=C       

attached base packages:
[1] parallel  stats4    stats     graphics  grDevices utils     datasets 
[8] methods   base     

other attached packages:
 [1] SingleCellExperiment_1.4.1  SummarizedExperiment_1.12.0
 [3] DelayedArray_0.8.0          BiocParallel_1.16.0        
 [5] matrixStats_0.55.0          Biobase_2.42.0             
 [7] GenomicRanges_1.34.0        GenomeInfoDb_1.18.1        
 [9] IRanges_2.16.0              S4Vectors_0.20.1           
[11] BiocGenerics_0.28.0         reshape2_1.4.3             
[13] ggplot2_3.2.1               DT_0.5                     
[15] dplyr_0.8.0.1              

loaded via a namespace (and not attached):
 [1] tidyselect_0.2.5       purrr_0.3.2            lattice_0.20-38       
 [4] colorspace_1.3-2       htmltools_0.3.6        yaml_2.2.0            
 [7] rlang_0.4.0            later_0.7.5            pillar_1.3.1          
[10] glue_1.3.0             withr_2.1.2            GenomeInfoDbData_1.2.0
[13] plyr_1.8.4             stringr_1.3.1          zlibbioc_1.28.0       
[16] munsell_0.5.0          gtable_0.2.0           workflowr_1.6.0       
[19] htmlwidgets_1.3        evaluate_0.12          labeling_0.3          
[22] knitr_1.20             crosstalk_1.0.0        httpuv_1.4.5          
[25] Rcpp_1.0.3             xtable_1.8-4           promises_1.0.1        
[28] scales_1.0.0           backports_1.1.2        jsonlite_1.6          
[31] XVector_0.22.0         mime_0.6               fs_1.3.1              
[34] digest_0.6.20          stringi_1.2.4          shiny_1.2.0           
[37] cowplot_0.9.4          grid_3.5.1             rprojroot_1.3-2       
[40] tools_3.5.1            bitops_1.0-6           magrittr_1.5          
[43] lazyeval_0.2.1         RCurl_1.95-4.11        tibble_2.1.1          
[46] crayon_1.3.4           whisker_0.3-2          pkgconfig_2.0.3       
[49] Matrix_1.2-17          assertthat_0.2.1       rmarkdown_1.10        
[52] R6_2.4.0               git2r_0.26.1           compiler_3.5.1
```
