## Supplementary material for "Characterizing and inferring quantitative cell cycle phase in single-cell RNA-seq data analysis": Source code of our analysis, including scripts and data necessary to reproduce the work.: README.md

# Peco

__peco__ is a supervised approach for predicting _continuous_ cell cycle phase in single-cell RNA-seq (scRNA-seq) data analysis. 

We trained __peco__ using cyclic gene expression signatures learned from Fluorescence Ubiquitin Cell Cycle Indicator (FUCCI) reporters. The data was collected from  fluorescence imaging with scRNA-seq to measure cell cycle phase and gene expression levels in human induced pluripotent stem cells (iPSCs). 

* Our paper: [Characterizing and inferring quantitative cell-cycle phase in single-cell RNA-seq data analysis](doi.org/10.1101/526848).


## Software

* [peco 0.99.4](https://github.com/jhsiao999/peco) is submitted and pending review on Bioconductor.

The development version can be downloaded from GitHub

```{r, eval=F}
devtools::install_github("jhsiao999/peco")
library(peco)
```


## Data access

* GEO record [GSE121265](http://www.ncbi.nlm.nih.gov/ geo/query/acc.cgi?acc=GSE121265) 
for all raw and processed sequencing data

* The processed data data sets are also available in as a gzip compressed 
tarball on the [Gilad lab website](https://giladlab.uchicago.edu/data/): https://giladlab.uchicago.edu/wp-content/uploads/2019/02/Hsiao_et_al_2019.tar.gz.

* All data sets used in our analysis are listed and downloadable at https://jdblischak.github.io/fucci-seq/data-overview.html.


## Code availability

Find out how we 

* [processed scRNA-seq and imaging data](access_data.html)

* [infer an angel for each cell based on FUCCI fluorescence intensities](https://jdblischak.github.io/fucci-seq/images-circle-ordering-eval.html)

* [estimated the cyclic trends of gene expression levels](https://jdblischak.github.io/fucci-seq/npreg-trendfilter-quantile.html)

* [analyzed our data and generated figures in our paper](https://github.com/jhsiao999/peco-paper/tree/master/code)


## Contact

Please contact me at []
for questions on the package or the methods.

## How to cite

> Hsiao, C. J., Tung, P., Blischak, J. D., Burnett, J., Dey, K. K.,
> Barr, A. K., Stephens, M., and Gilad, Y. (2018). [Characterizing and
> inferring quantitative cell-cycle phase in single-cell RNA-seq data
> analysis.](https://doi.org/10.1101/526848) bioRxiv doi:10.1101/526848

## License

Copyright (c) 2018-2019, Chiaowen Joyce Hsiao.

All source code and software in this repository are made available
under the terms of the [GNU General Public
License](https://www.gnu.org/licenses/gpl-3.0.en.html). See
file [LICENSE](LICENSE) for the full text of the license.
