## Supplementary figures and images for "Characterizing and inferring quantitative cell cycle phase in single-cell RNA-seq data analysis"

### after-filter-1.png

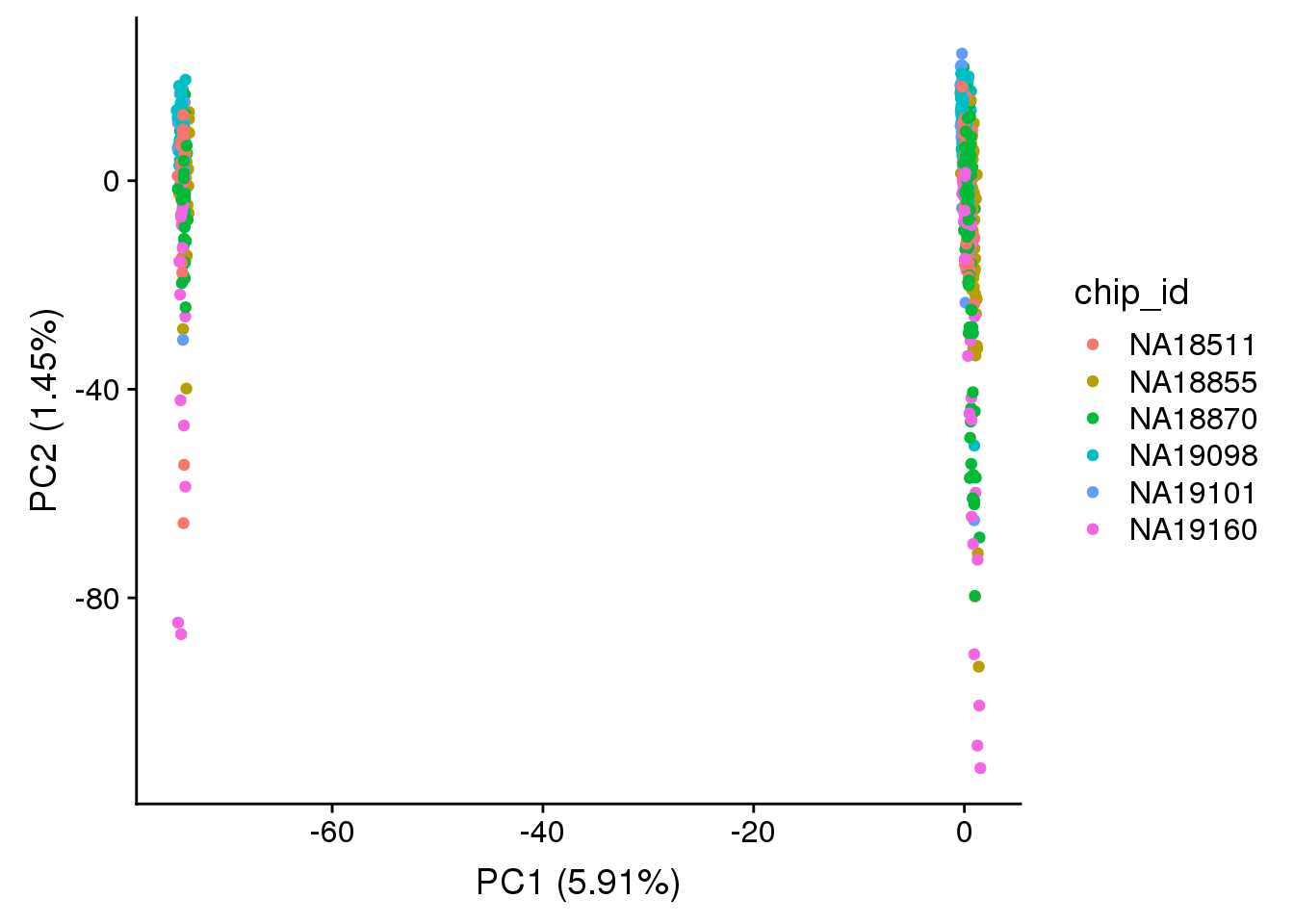

### after-filter-2.png

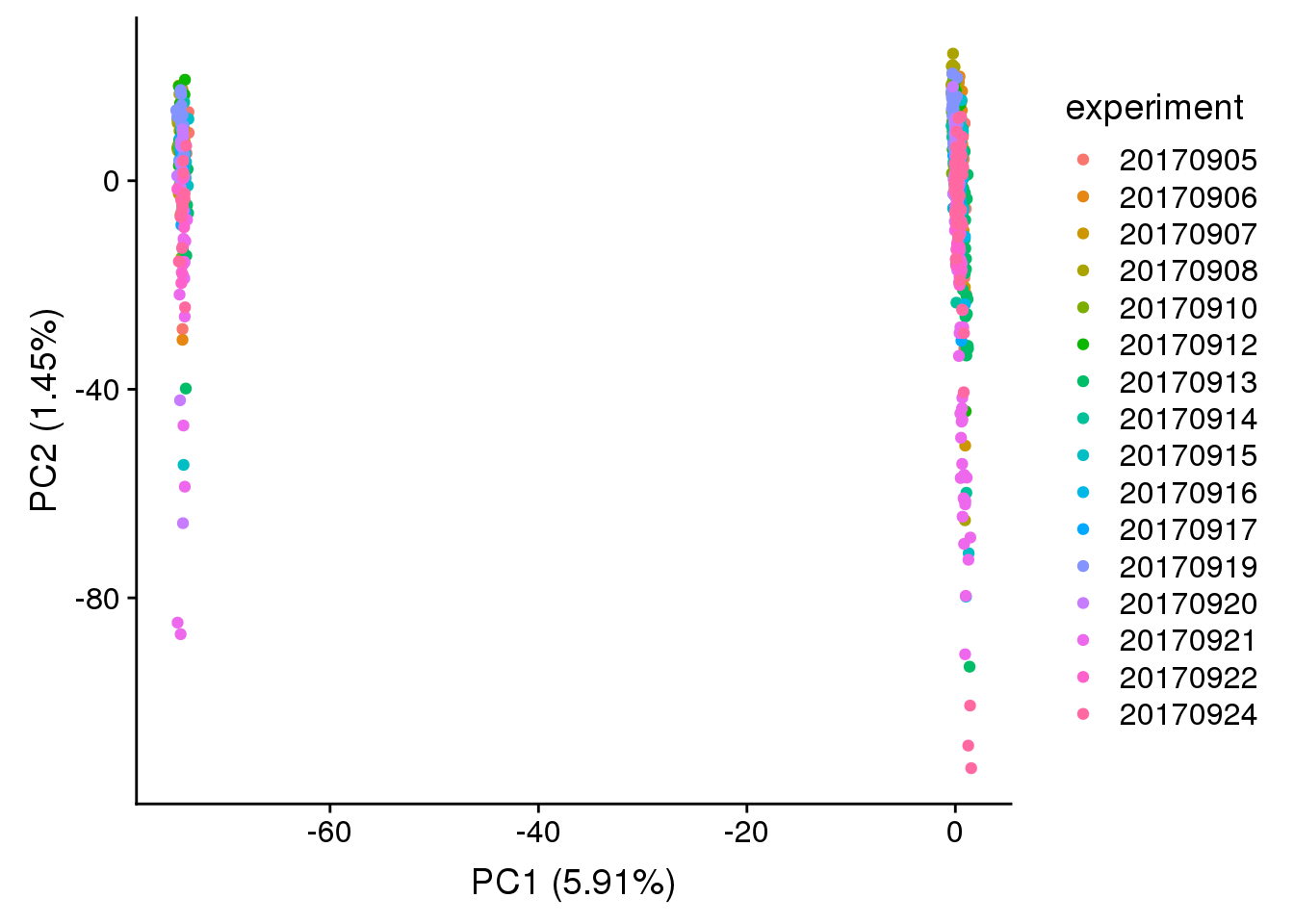

### after-filter-tf-1.png

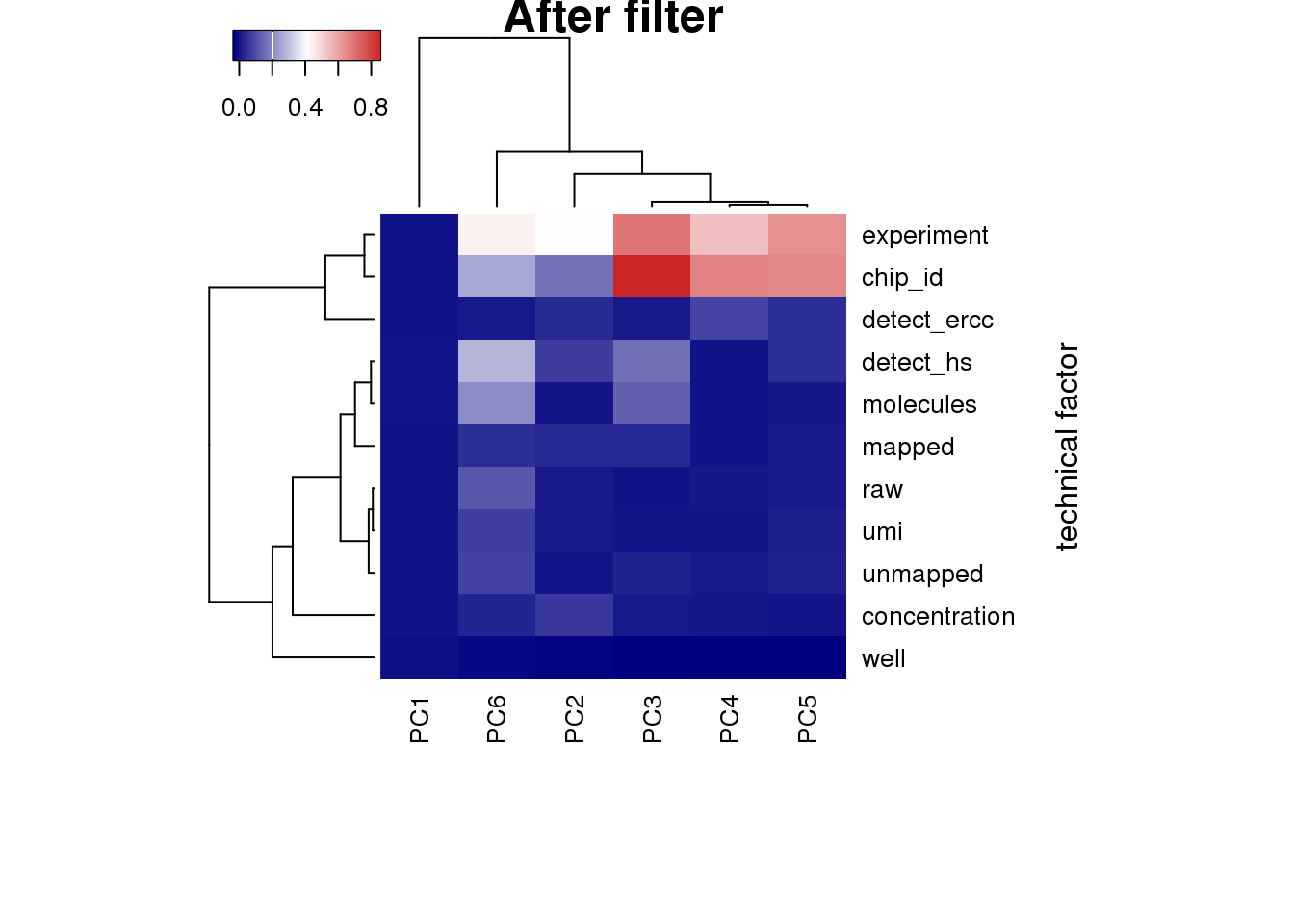

### before-filter-1.png

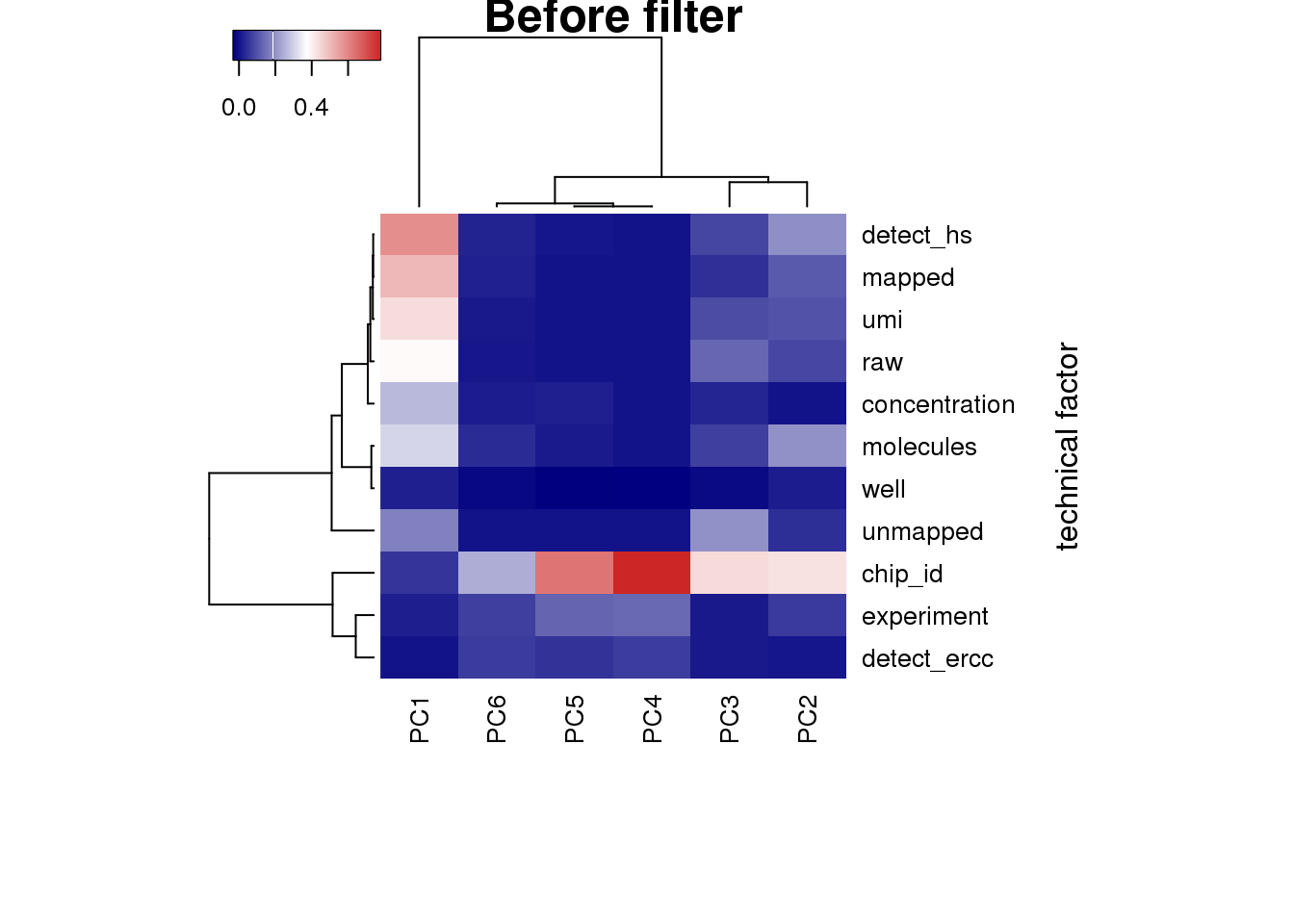

### before-filter-2.png

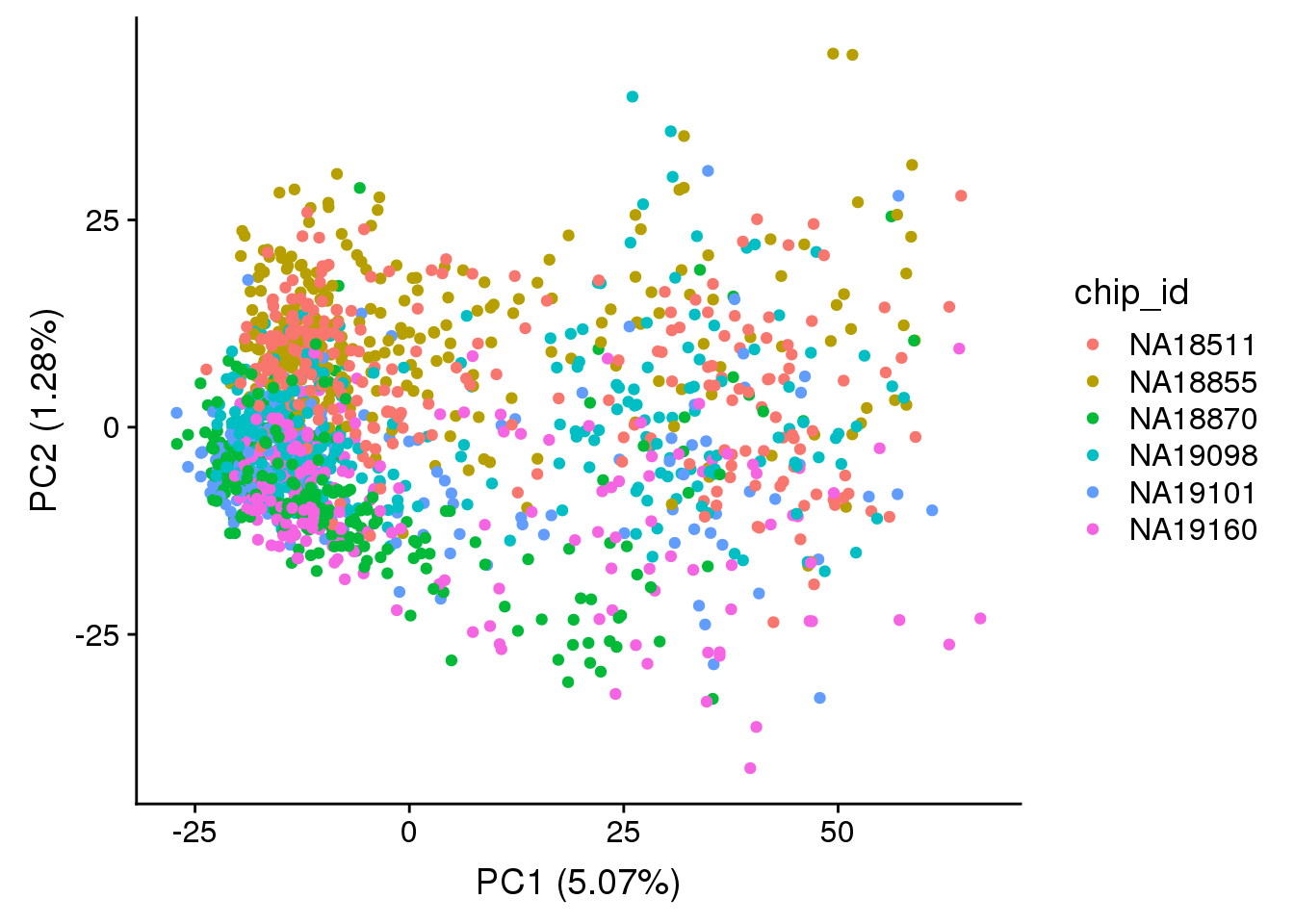

### boxplot-adjusted-1.png

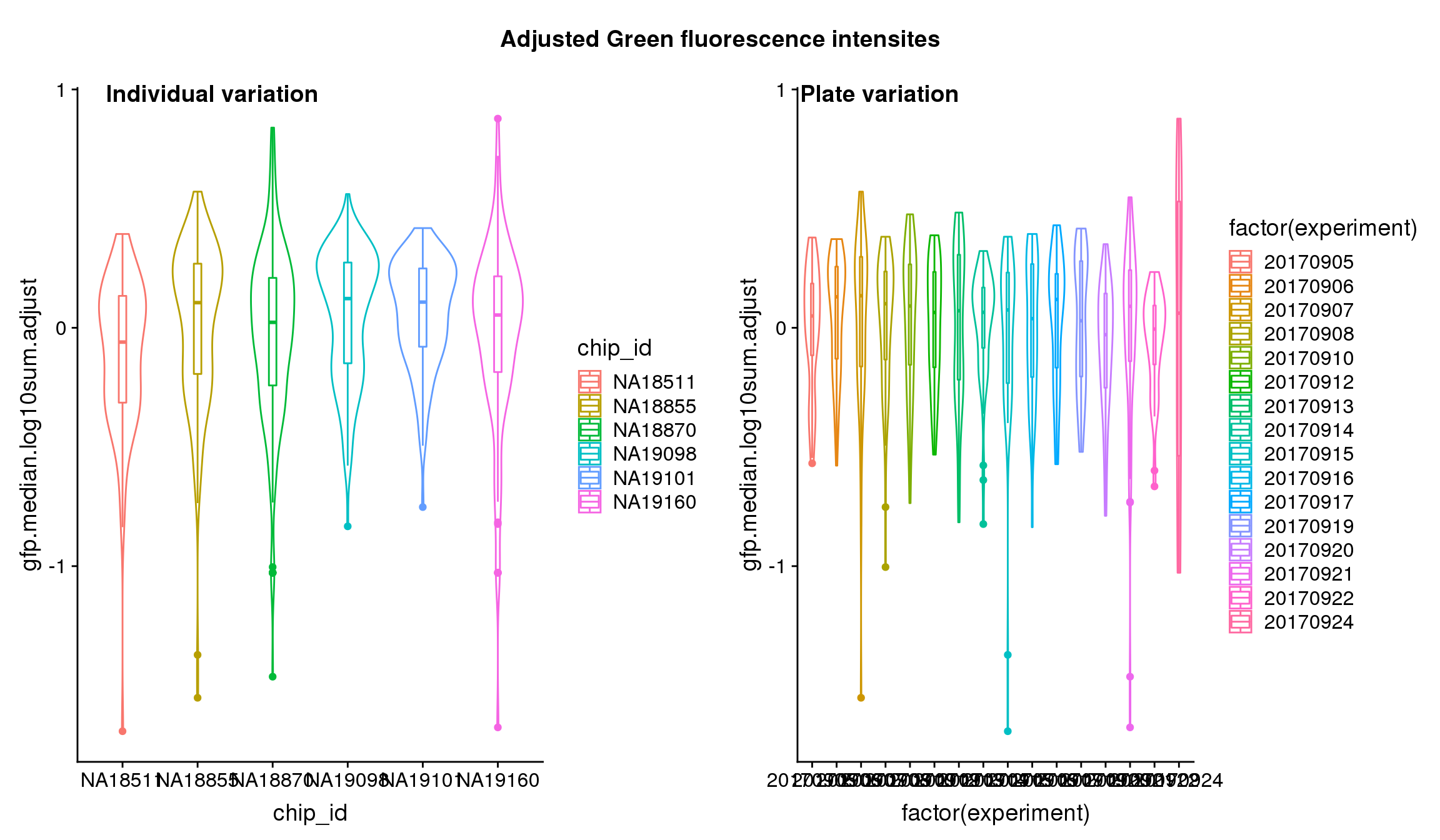

### boxplot-adjusted-2.png

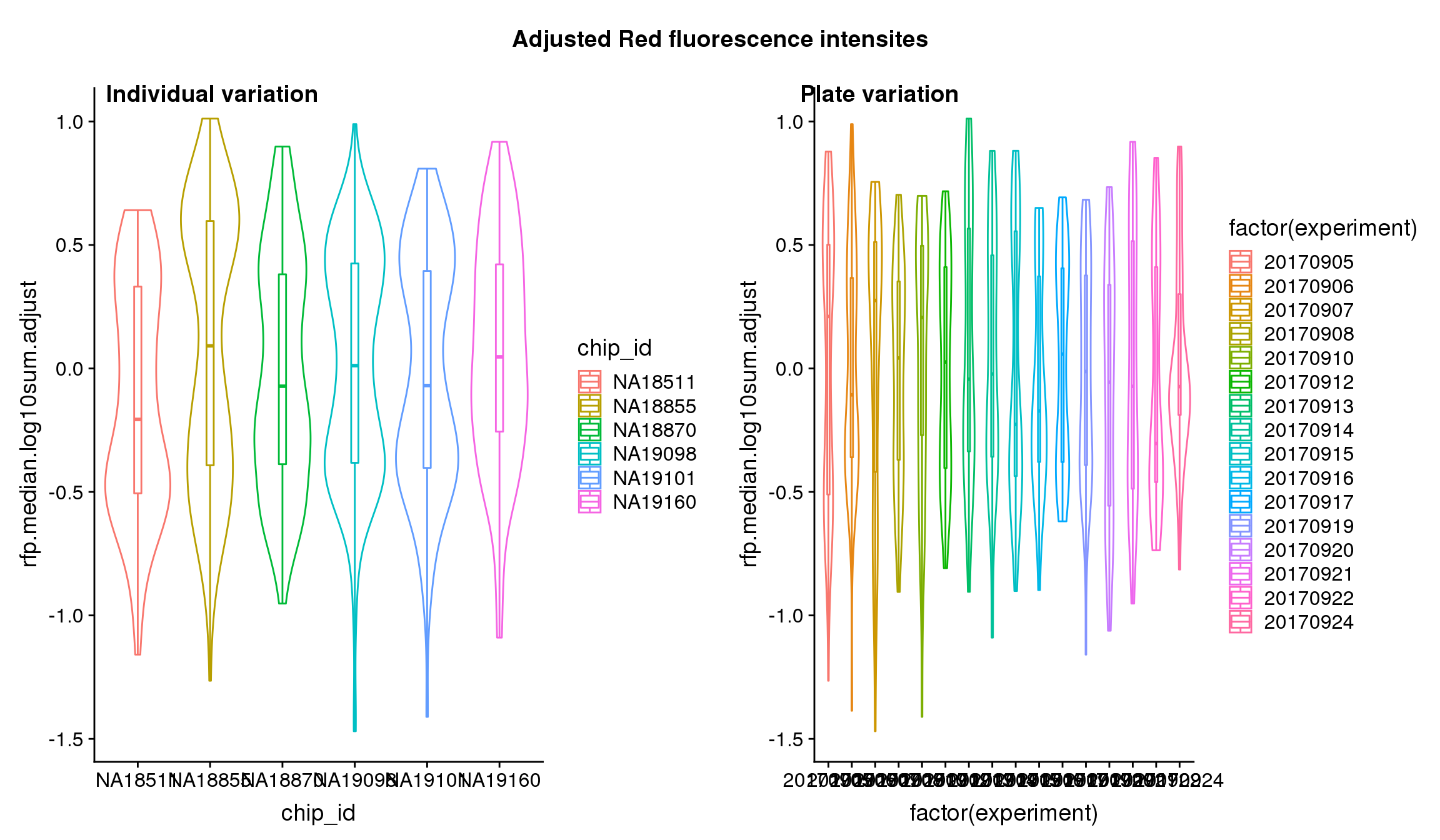

### boxplot-adjusted-3.png

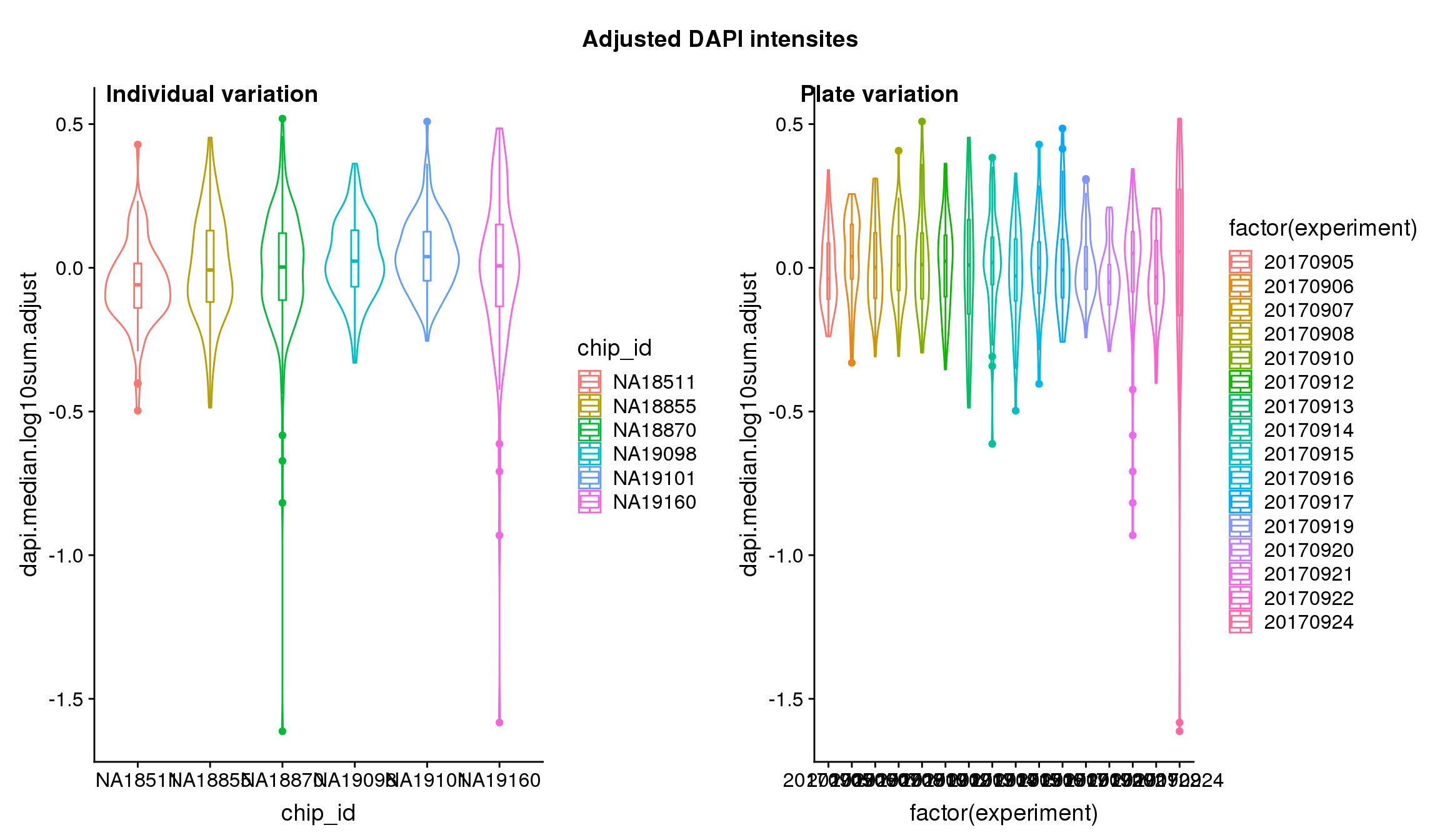

### boxplot-adjusted-4.png

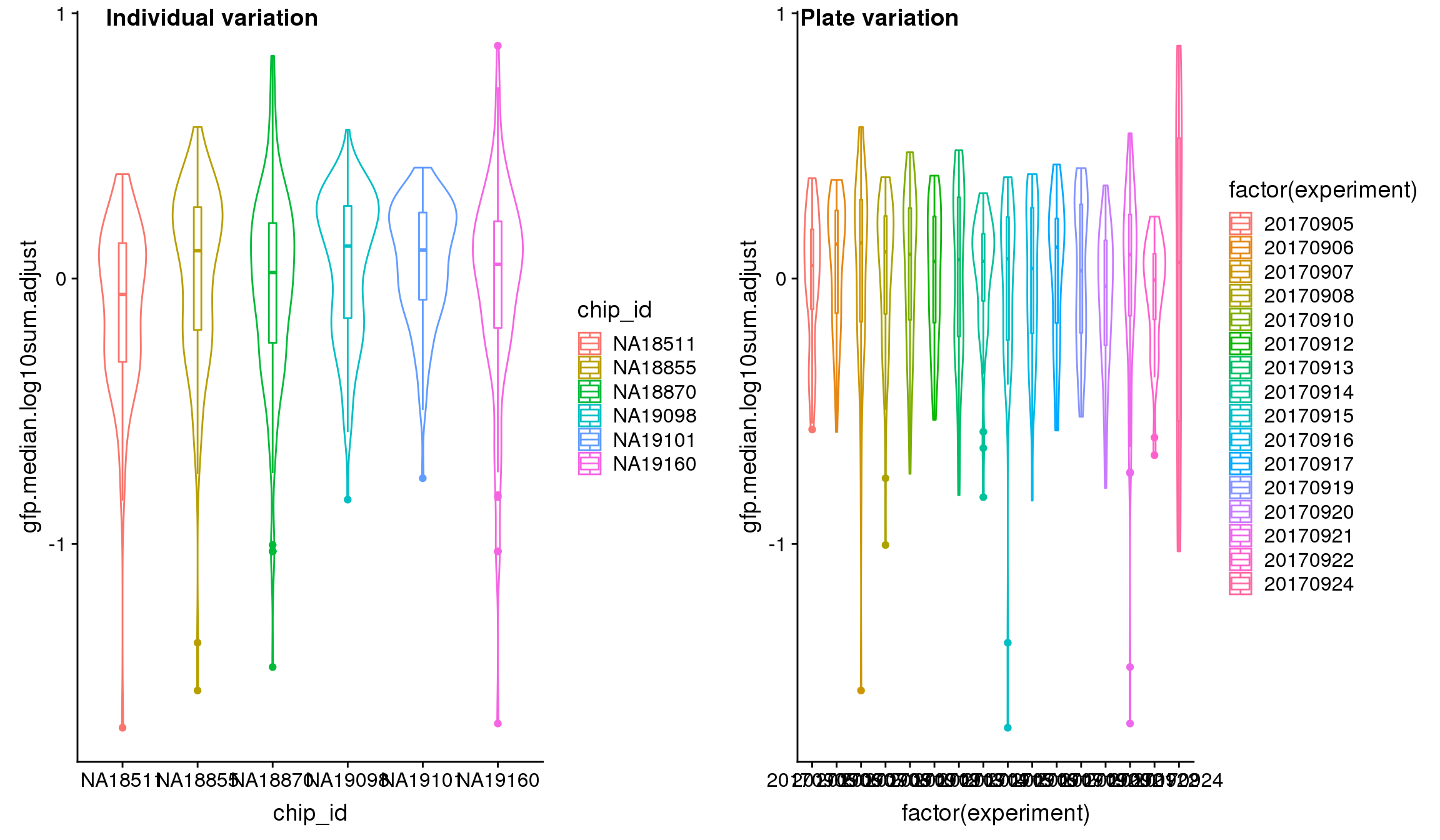

### boxplot-adjusted-5.png

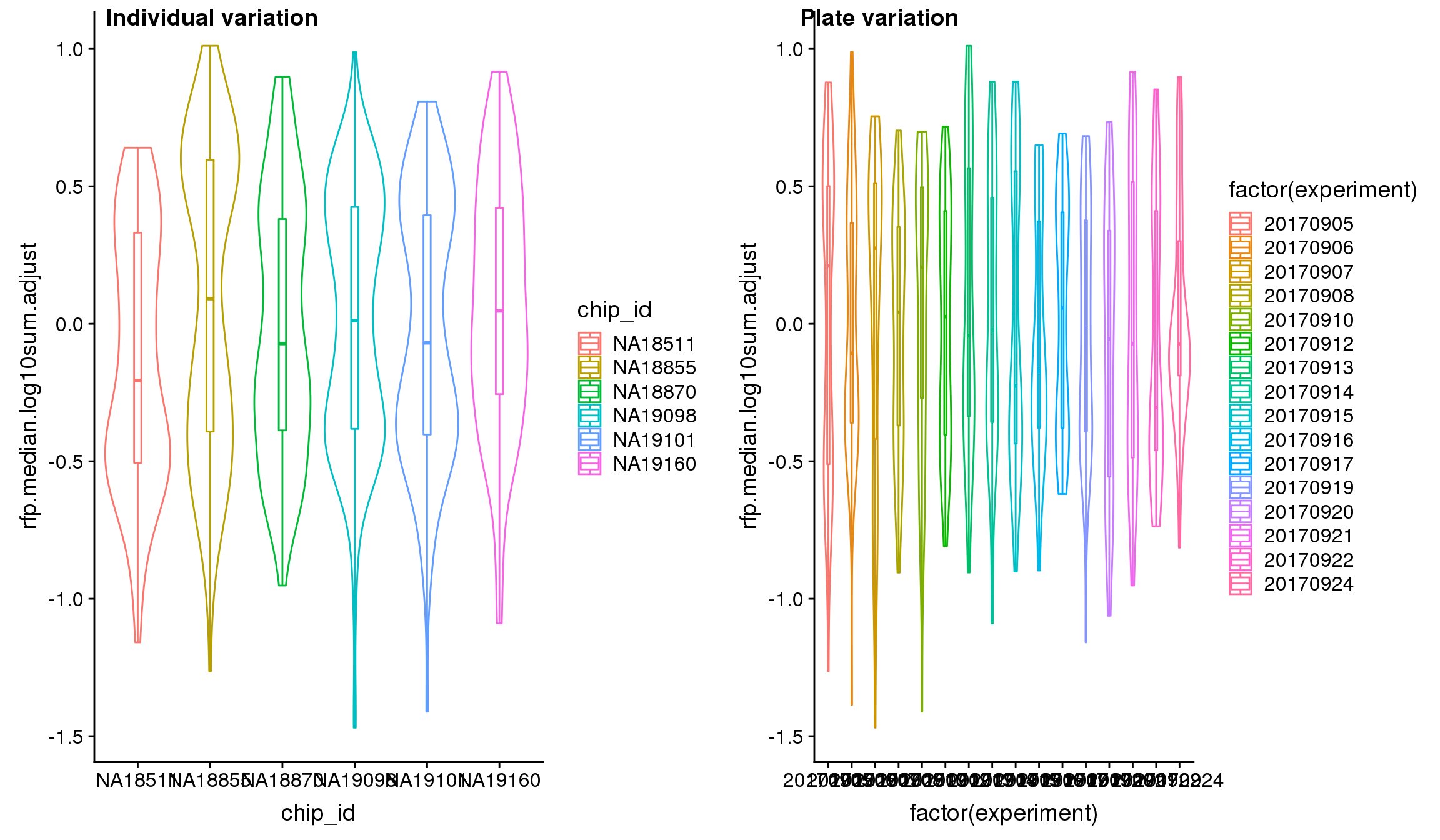

### boxplot-adjusted-6.png

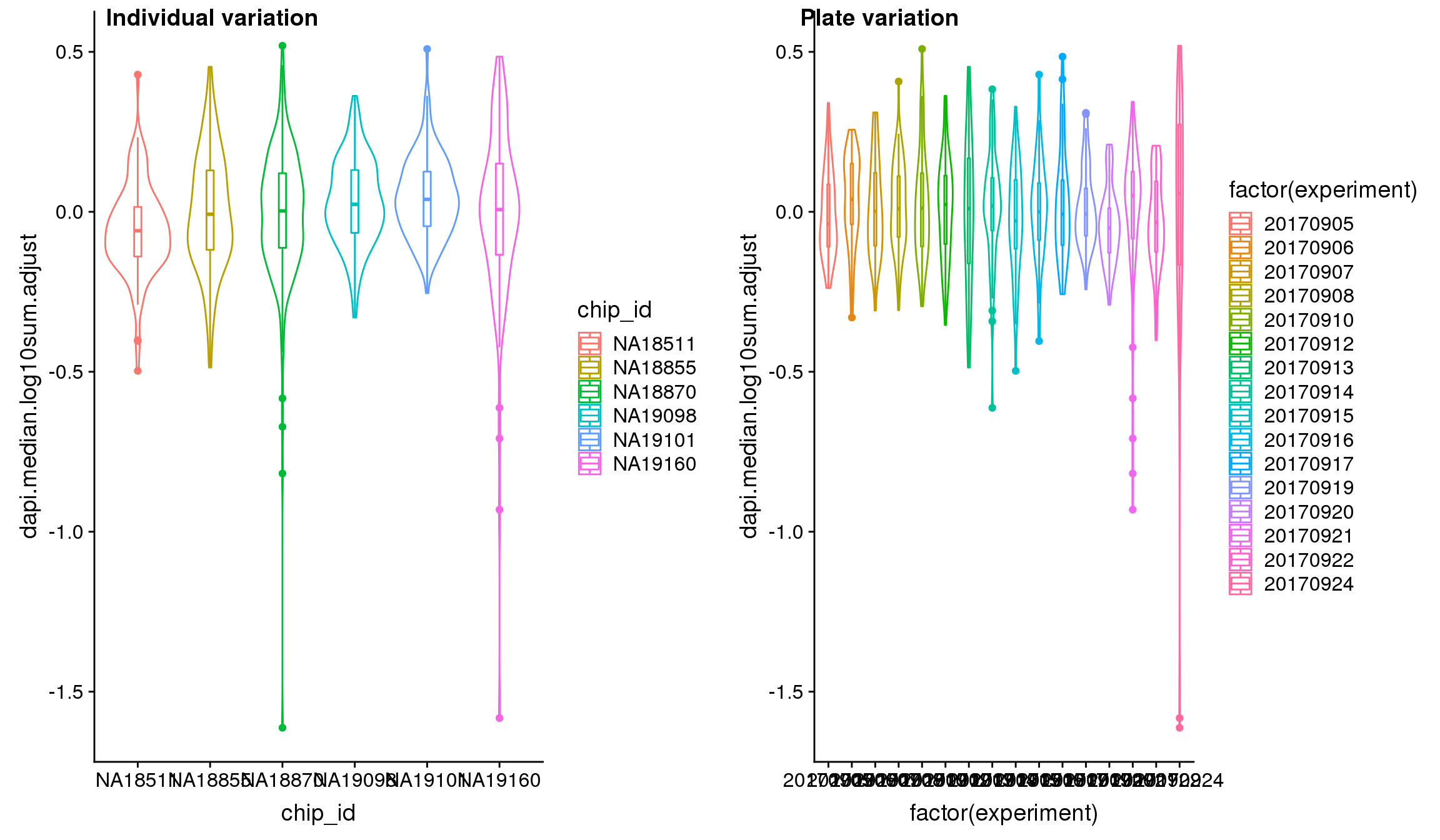

### check-cyclic-1.png

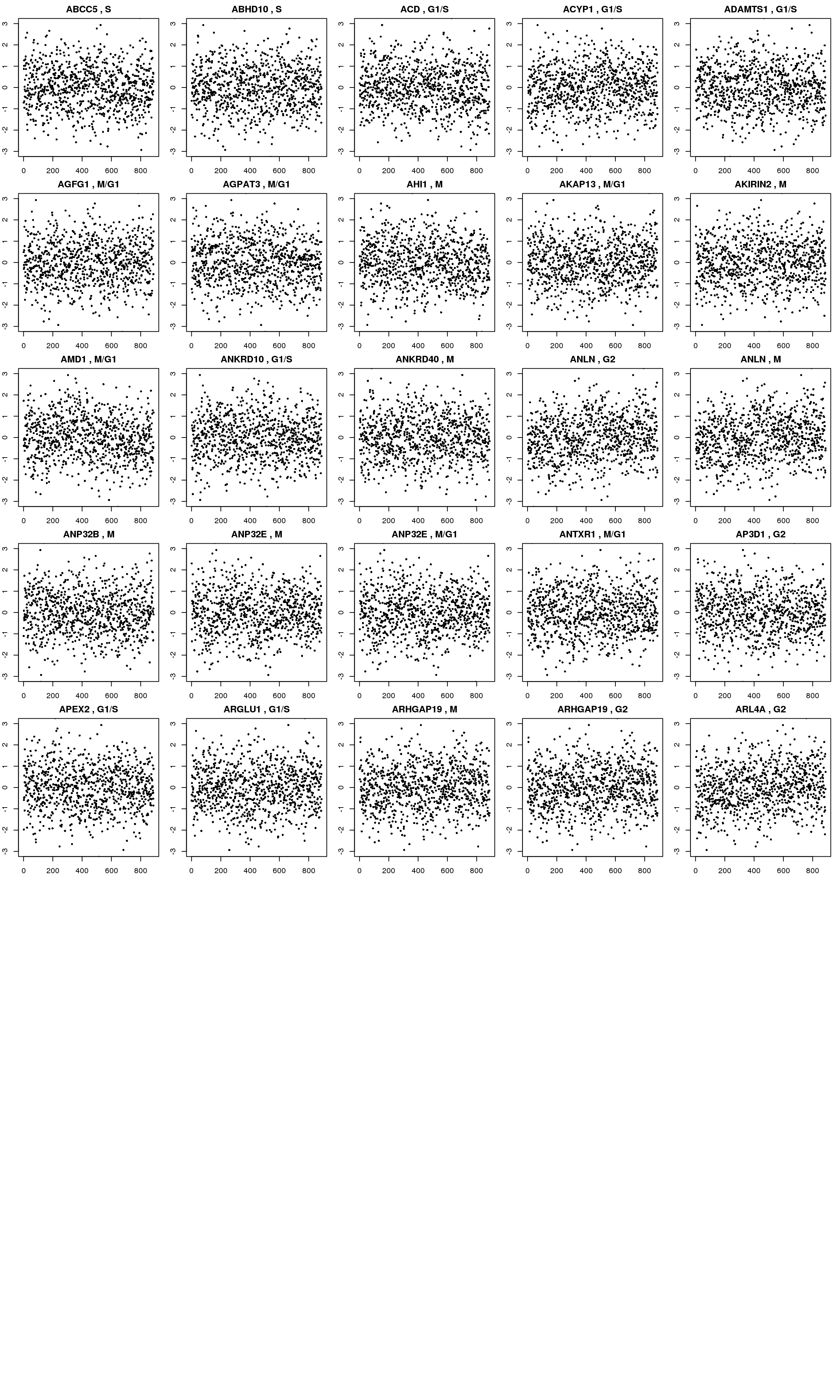

### check-high-low-undetected-cells-1.png

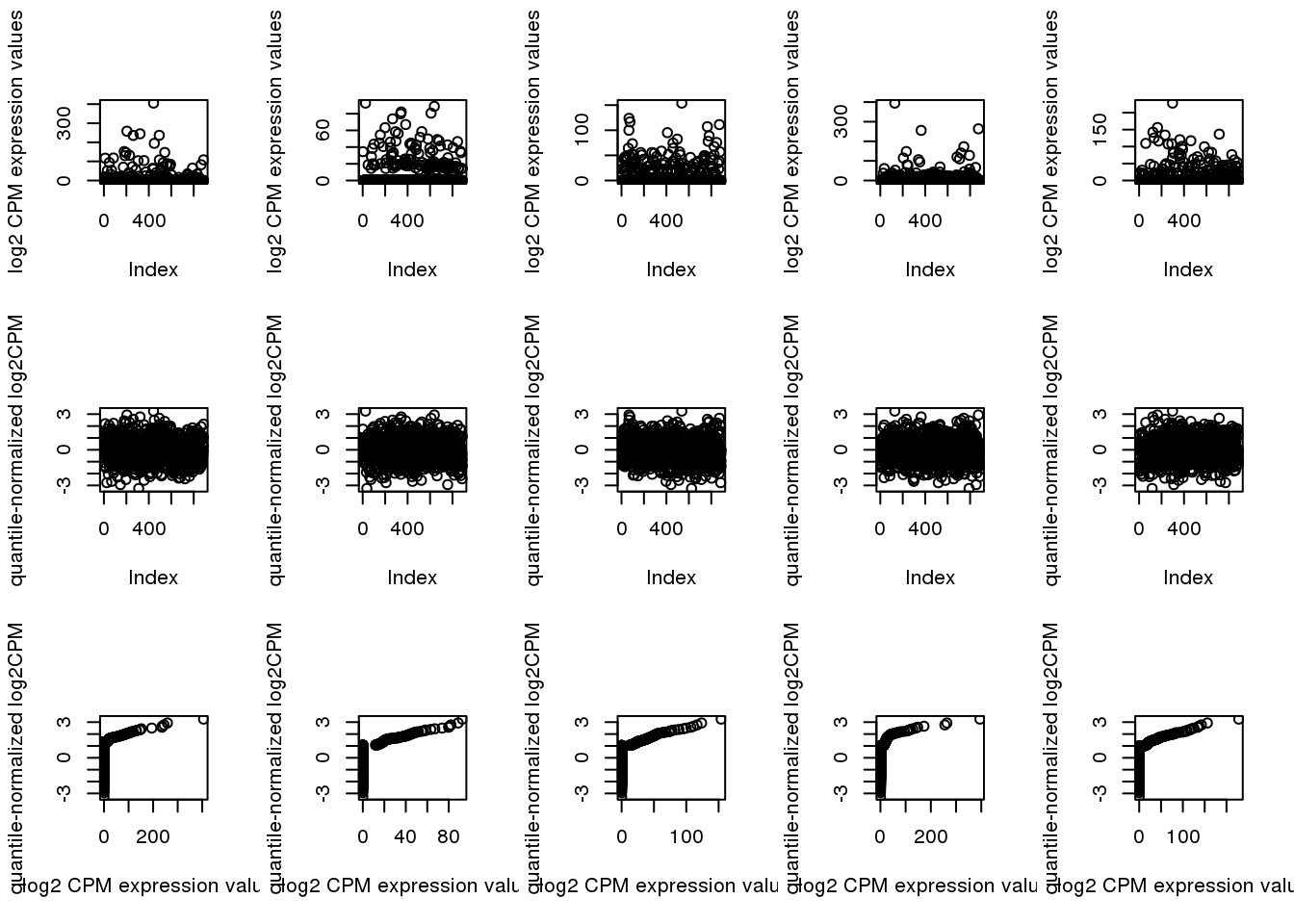

### check-high-low-undetected-cells-2.png

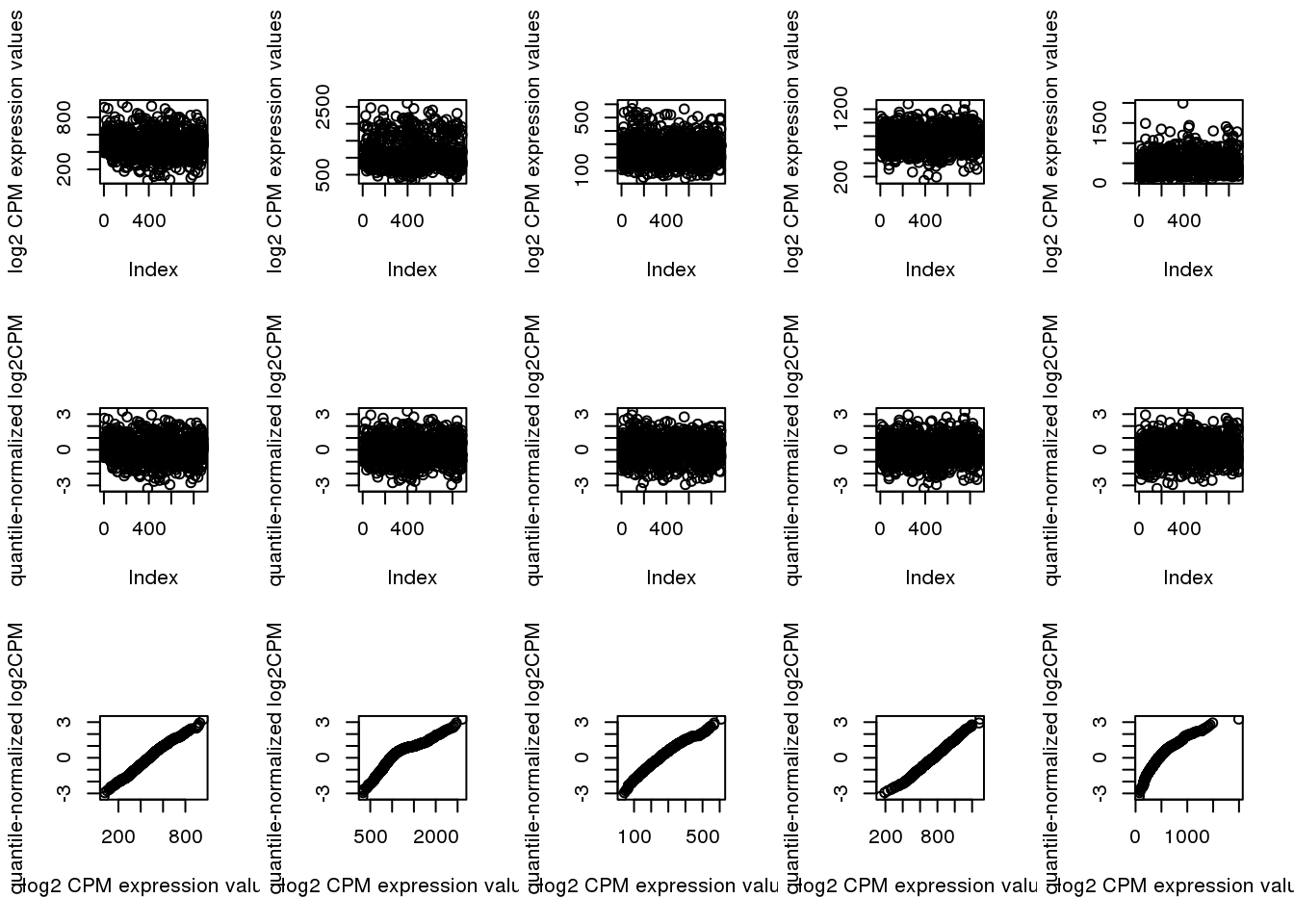

### conversion-efficiency-1.png

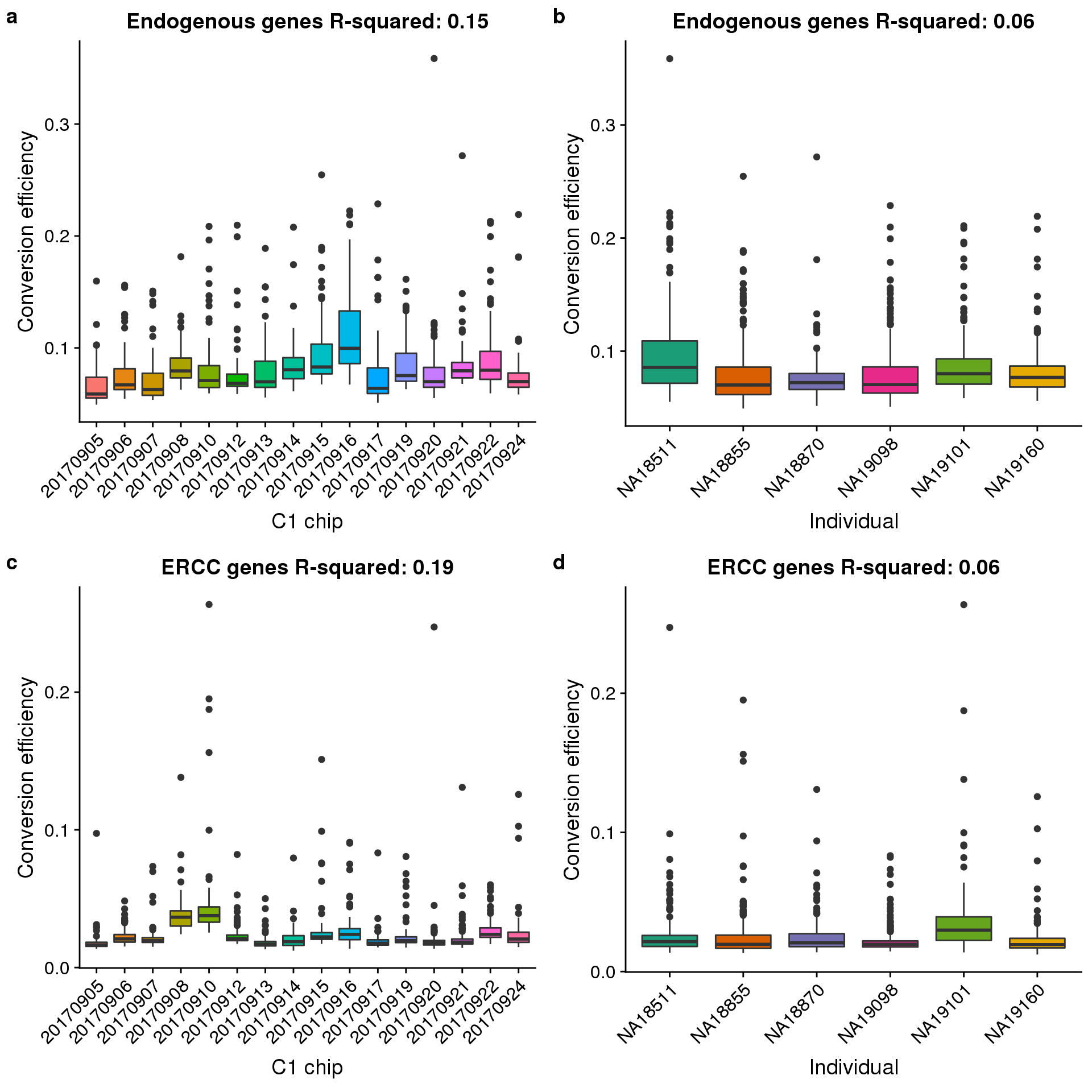

### convertion-1.png

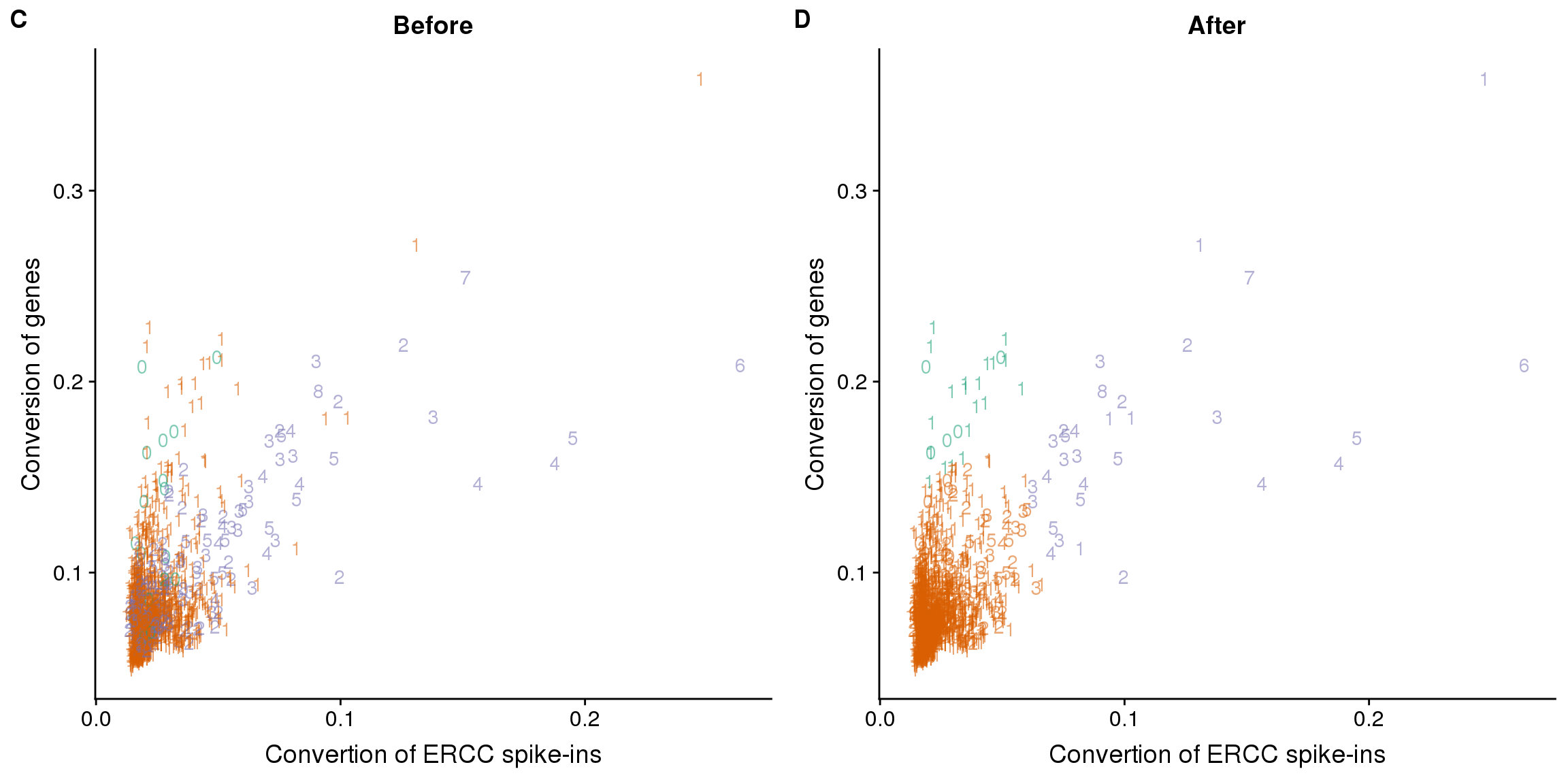

### cor-1.png

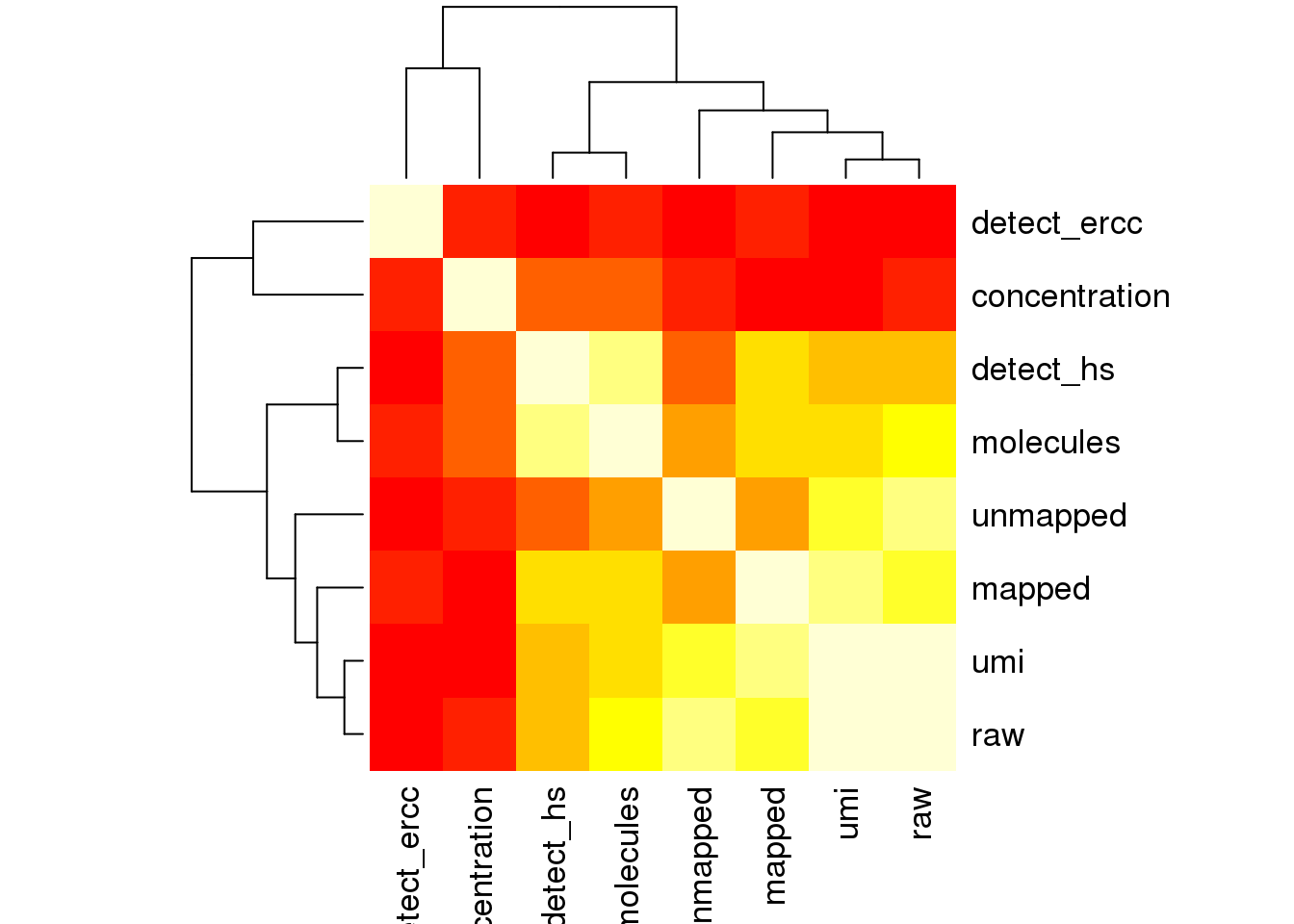

### distribution-sequencing-depth-1.png

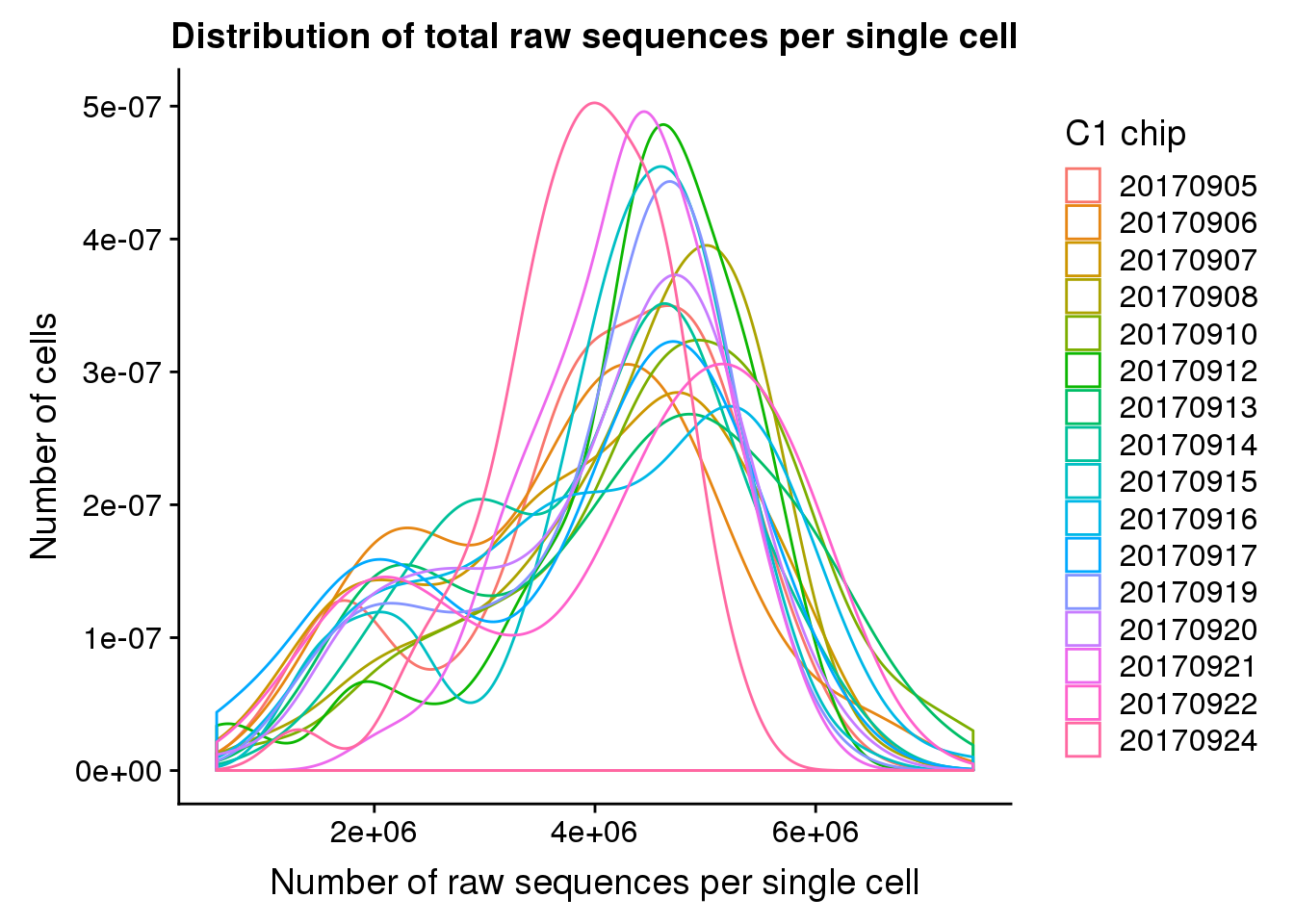

### ercc-percentage-1.png

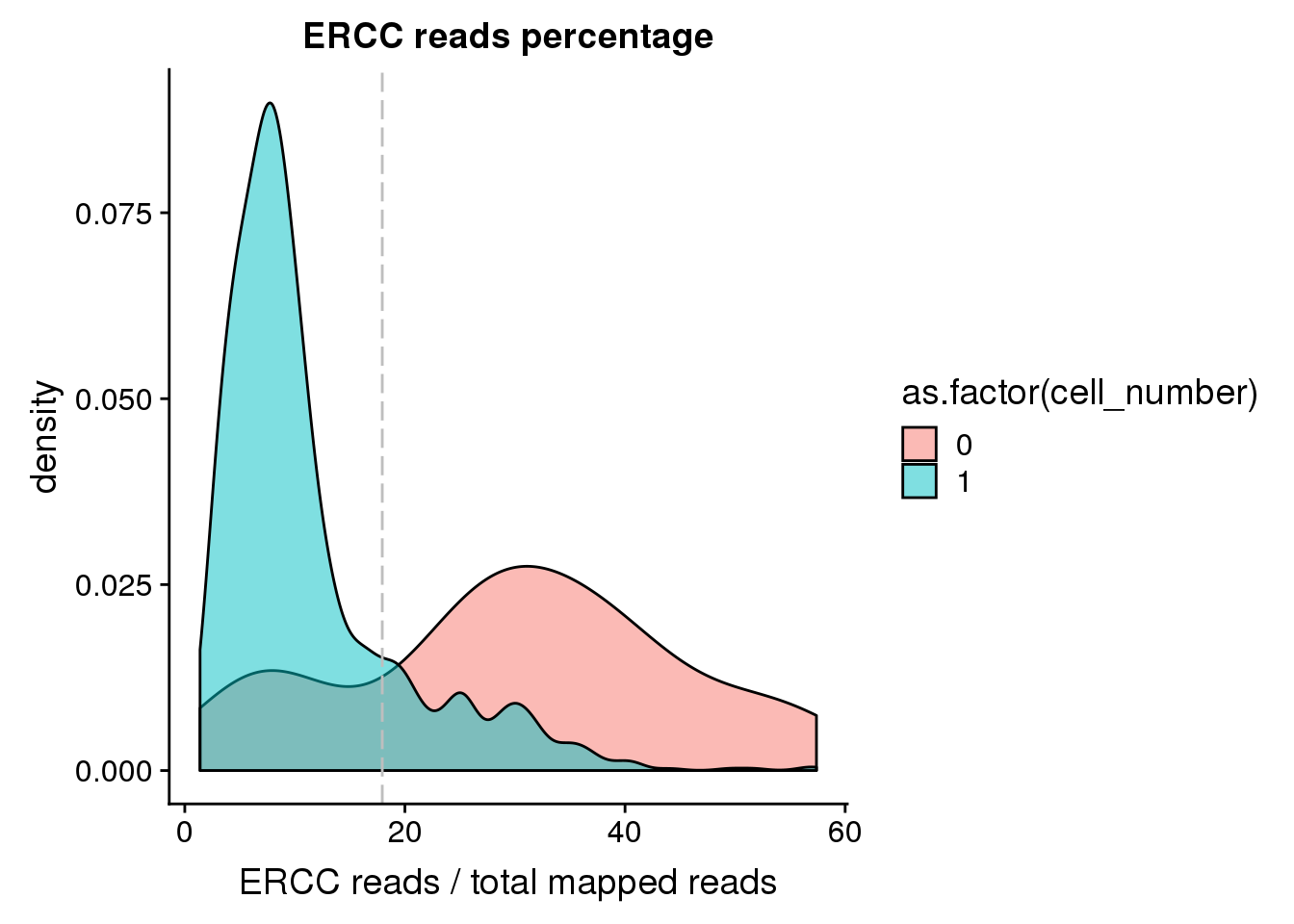

### ercc_exp-1.png

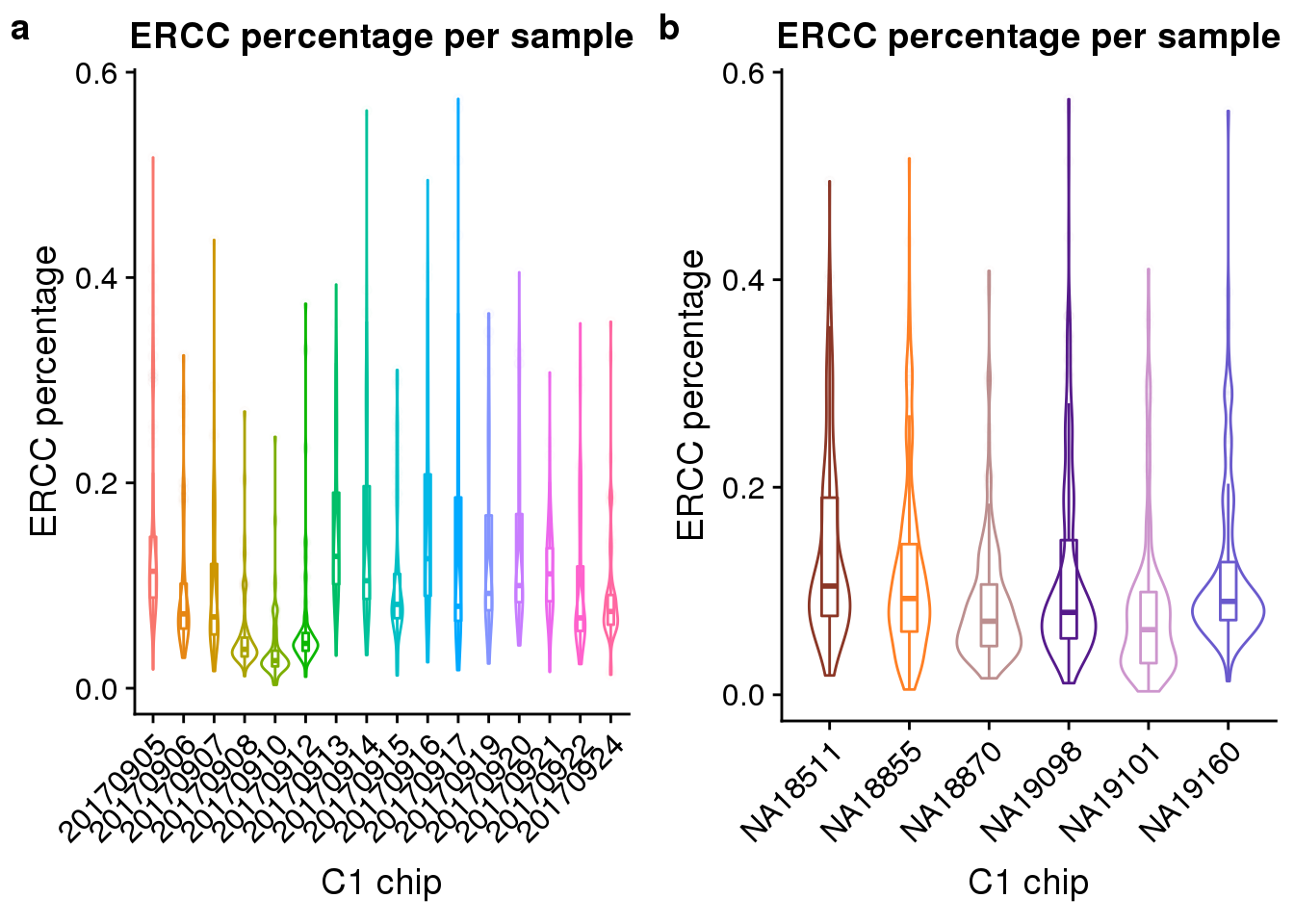

### fucci-1.png

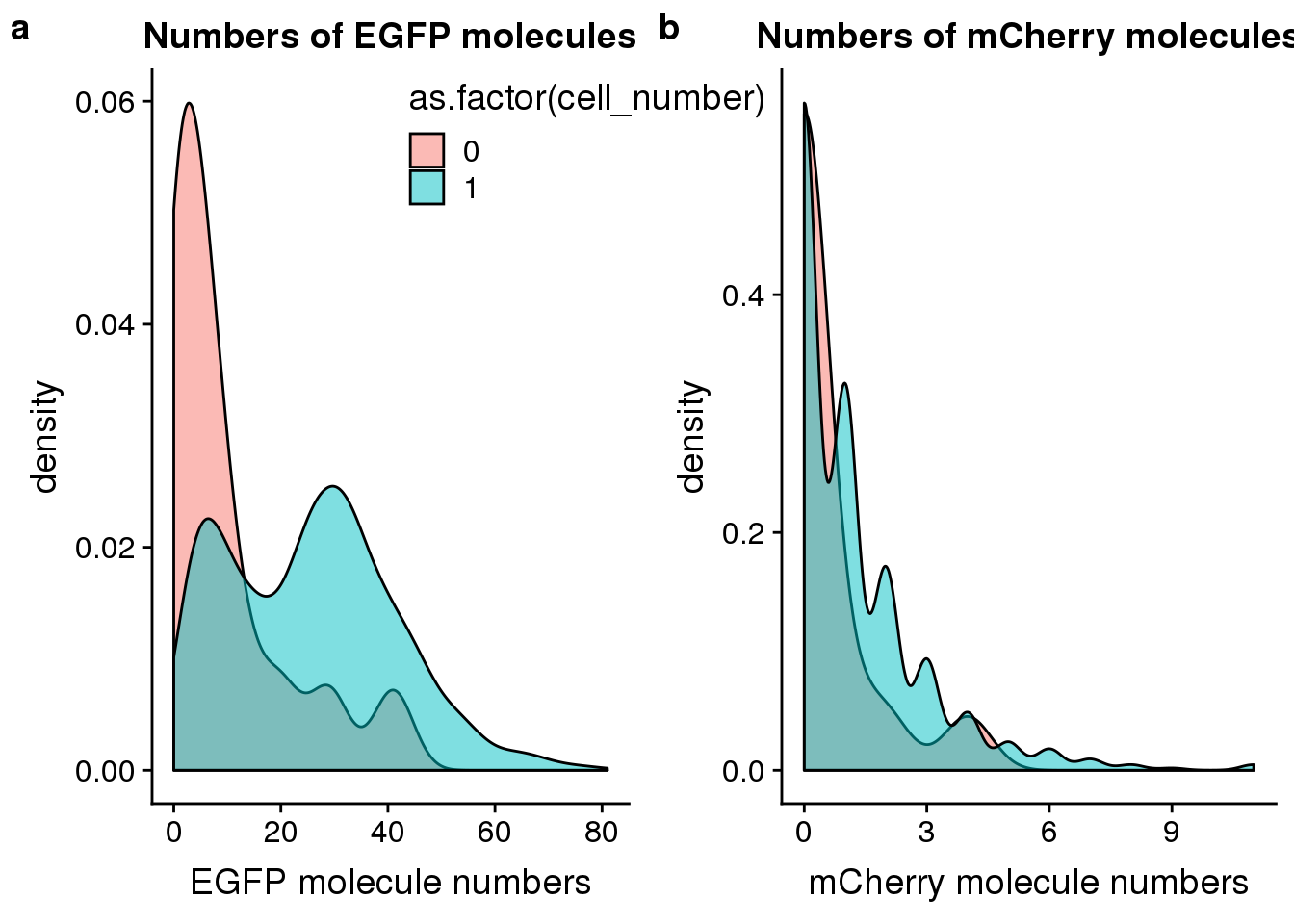

### gene-number-1.png

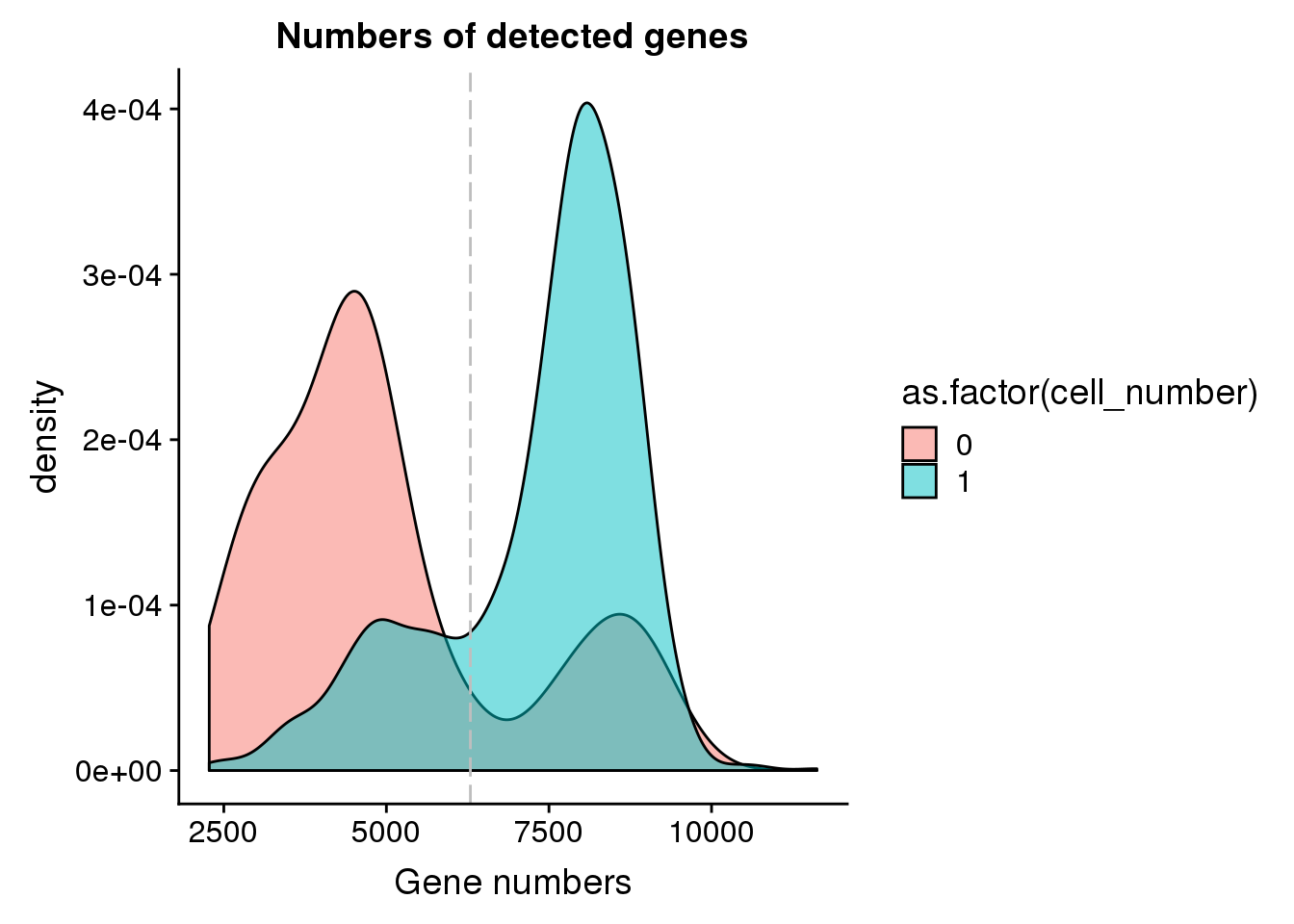

### gene-number-exp-1.png

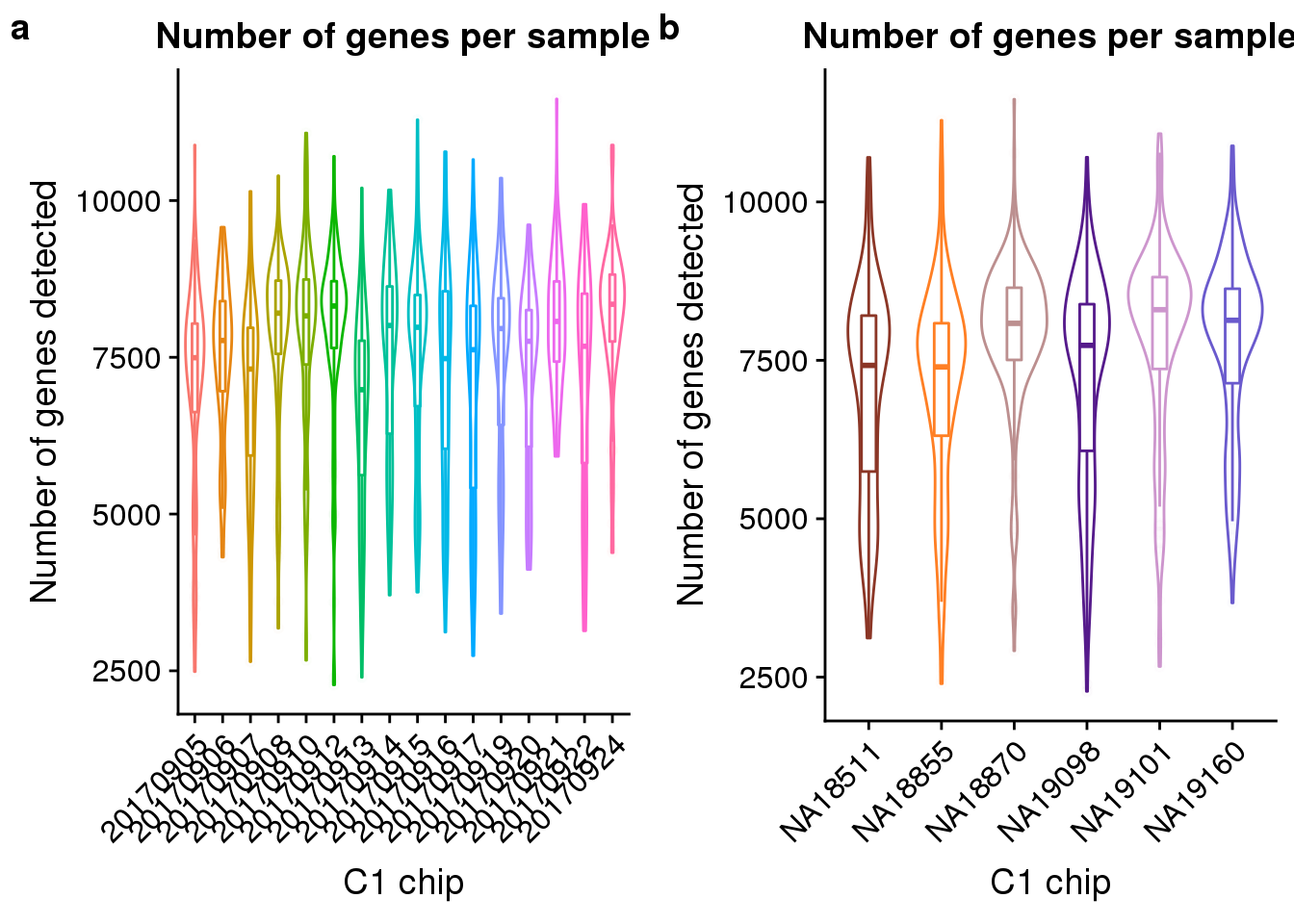

### gene-v-ercc-1.png

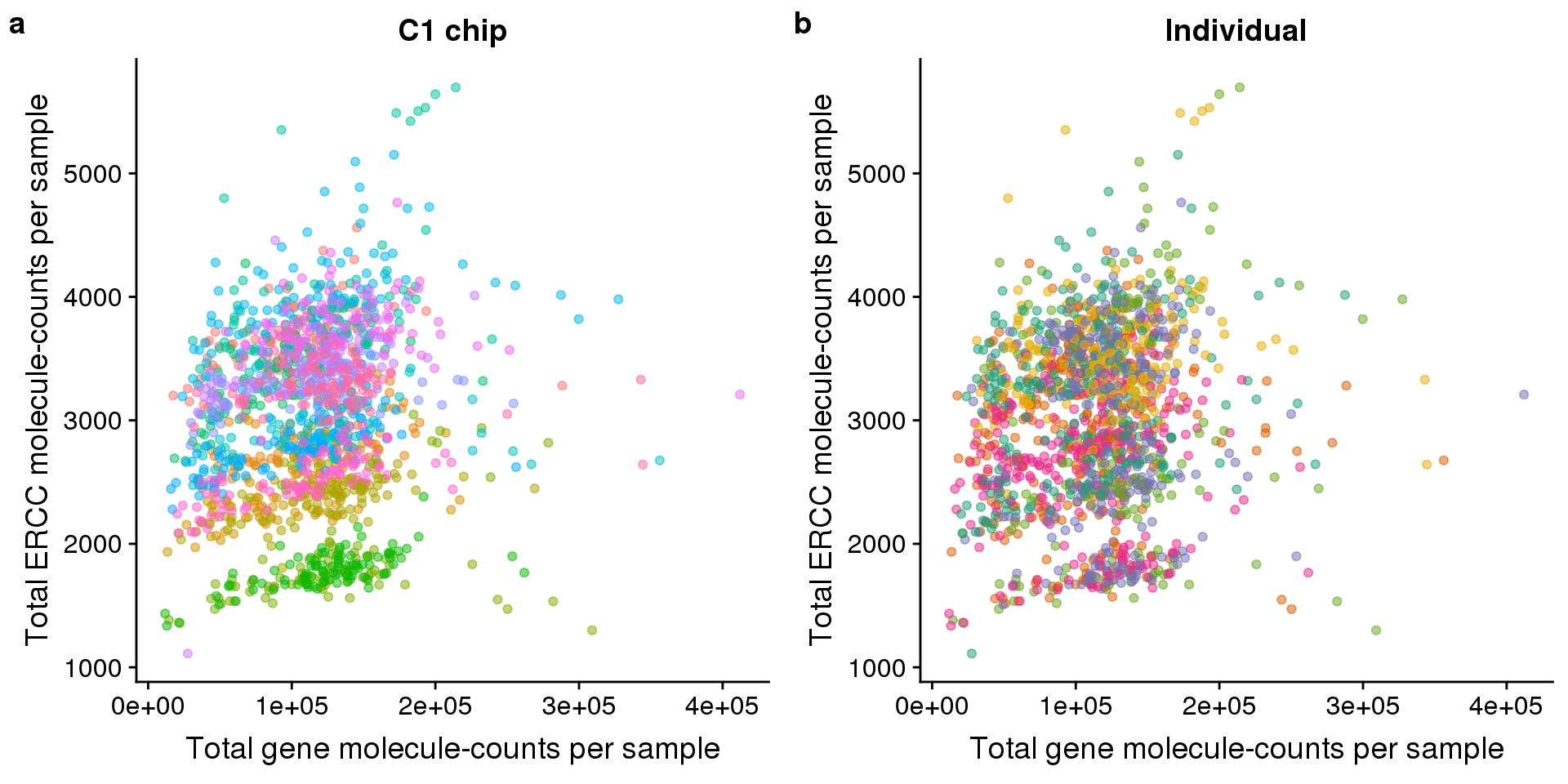

### get-permute-pvals-1.png

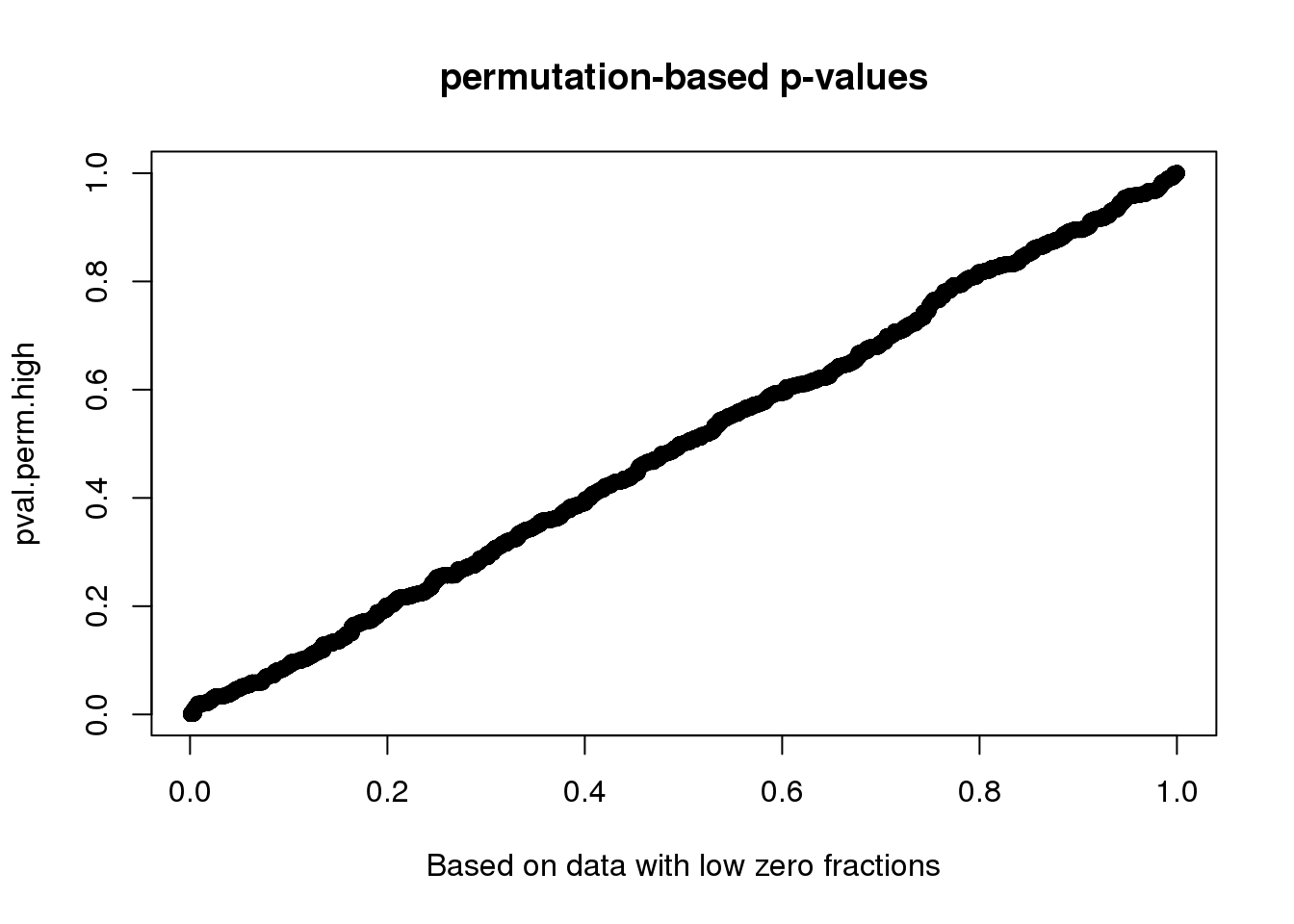

### lda-1.png

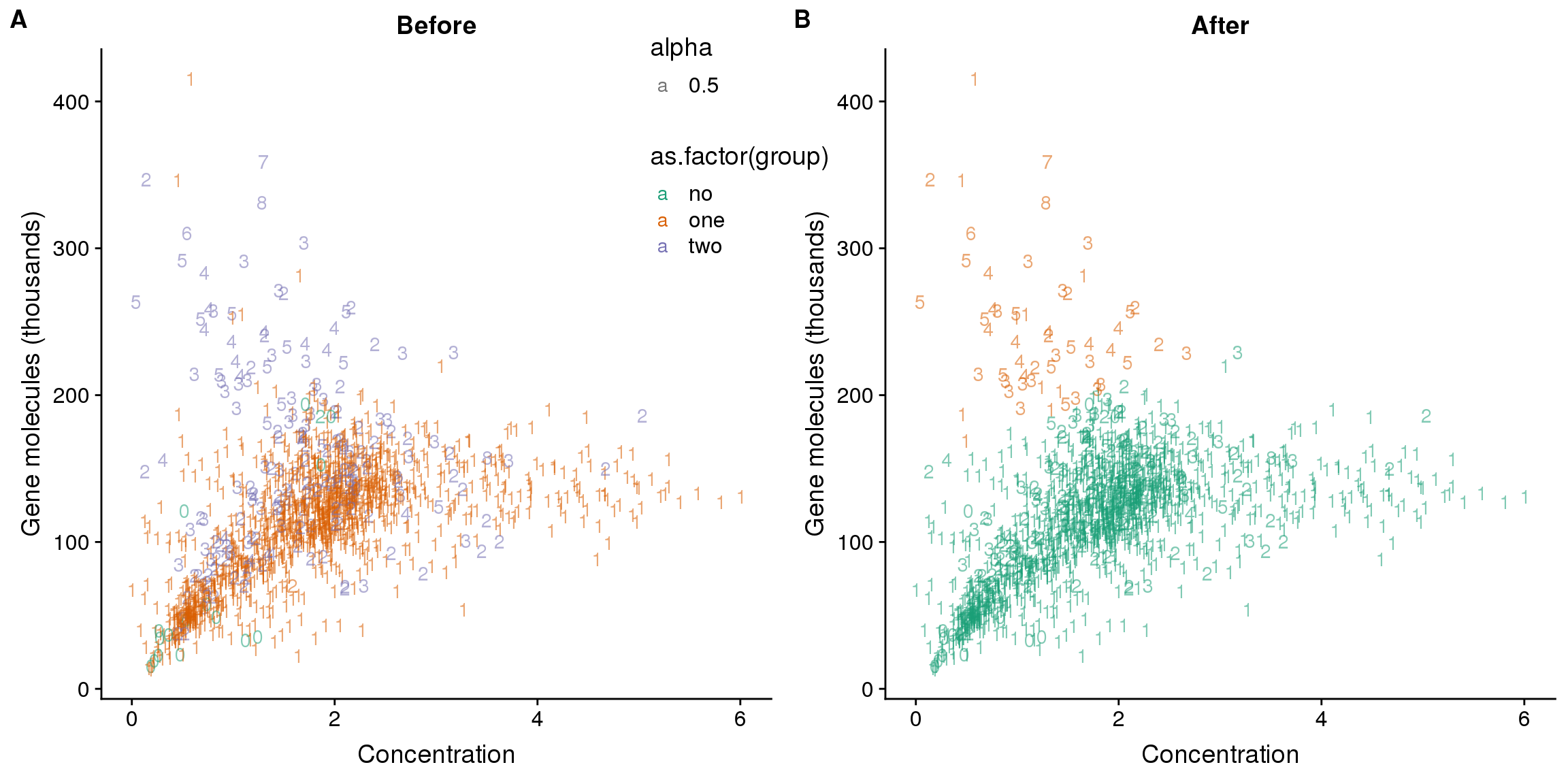

### load-data-1.png

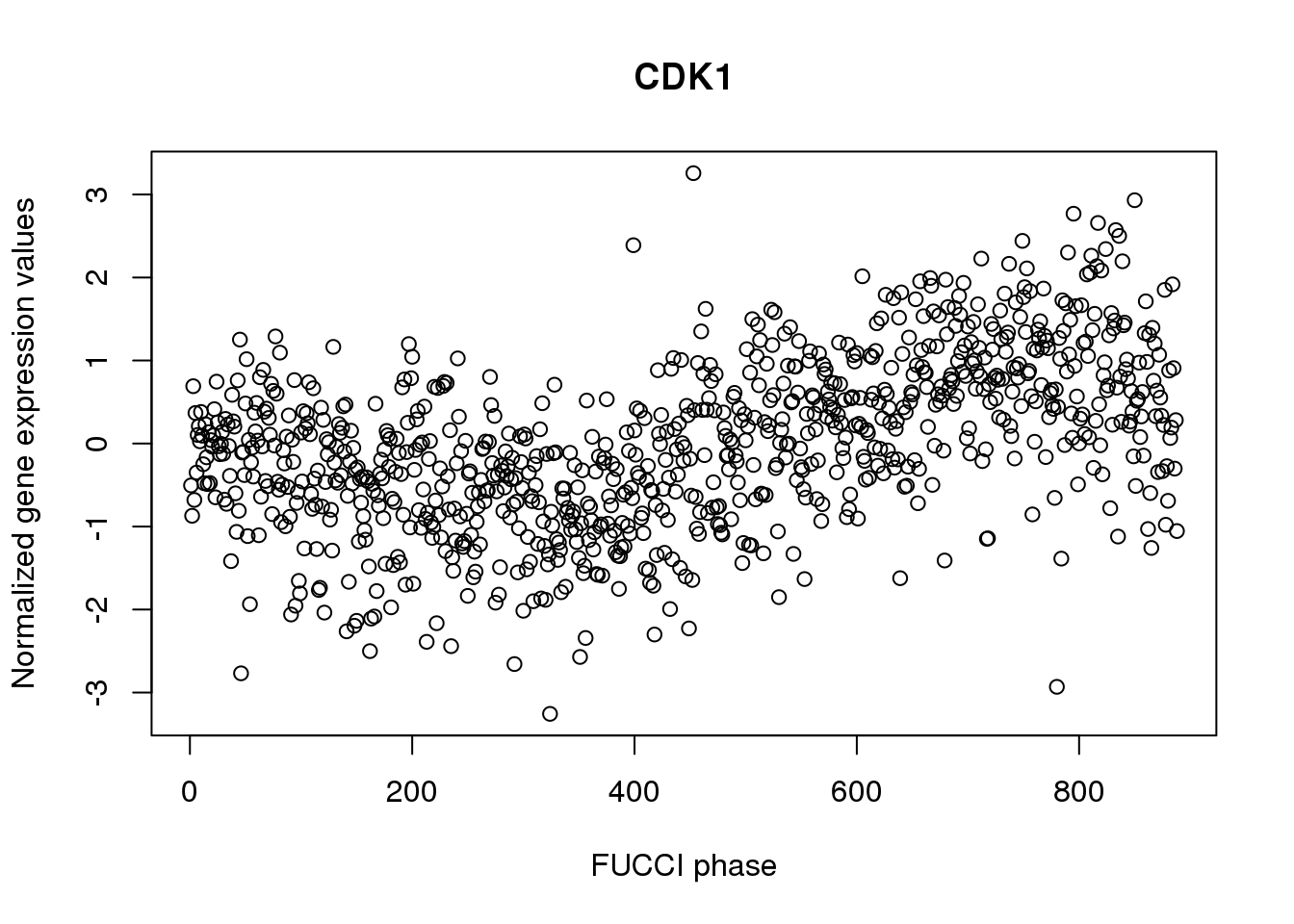

### load-permdist-results-1.png

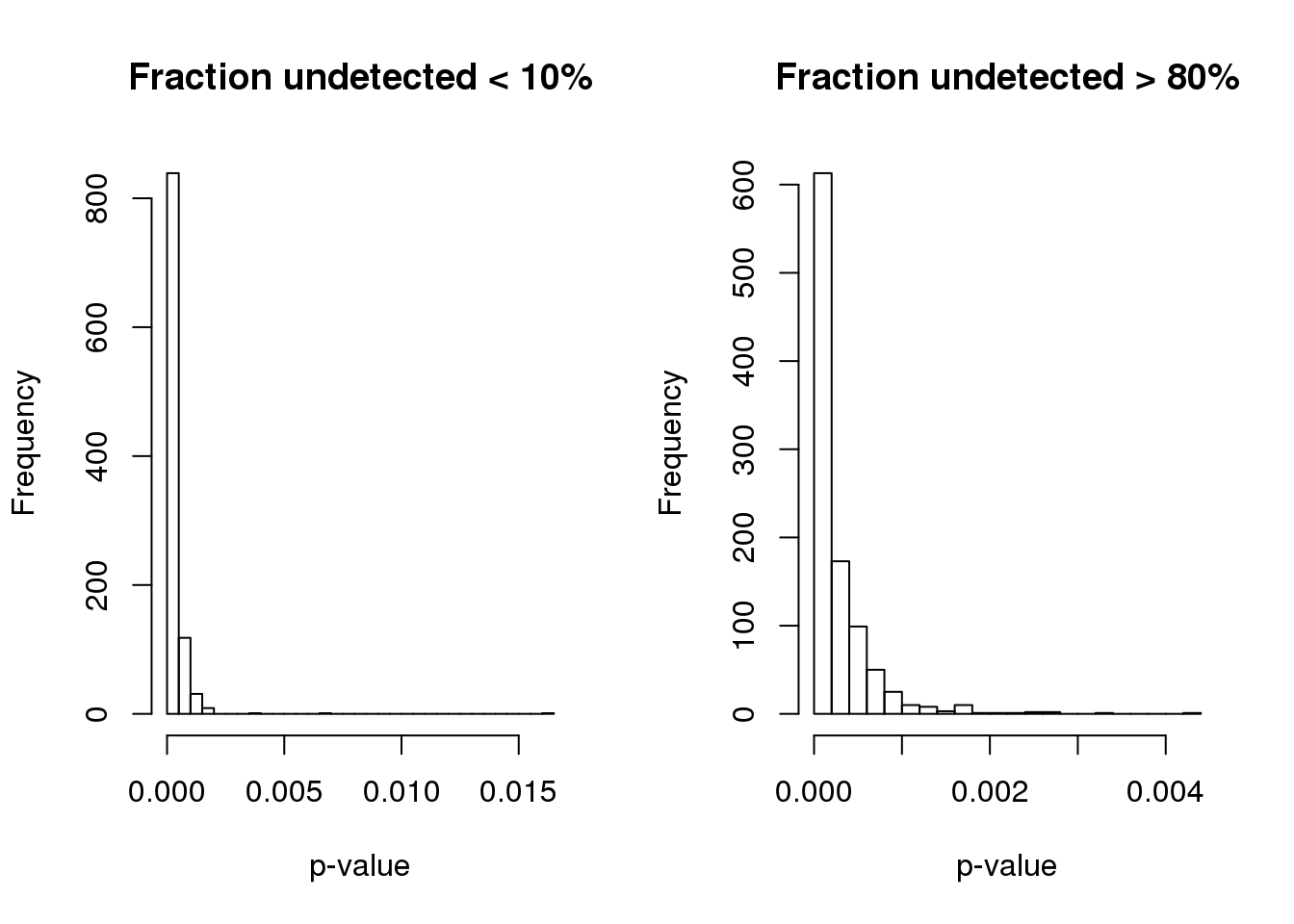

### mapped-1.png

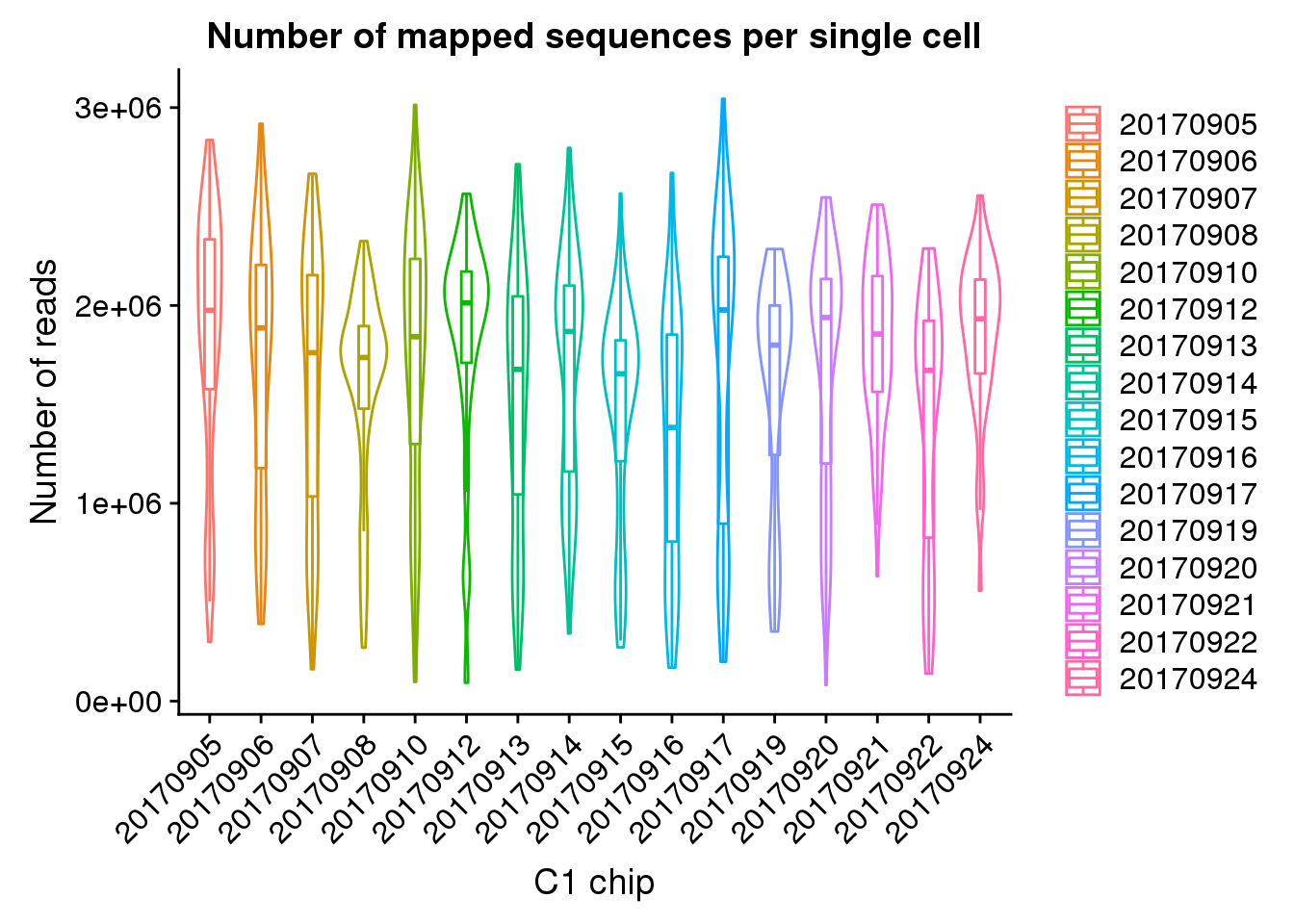

### peco-1.png

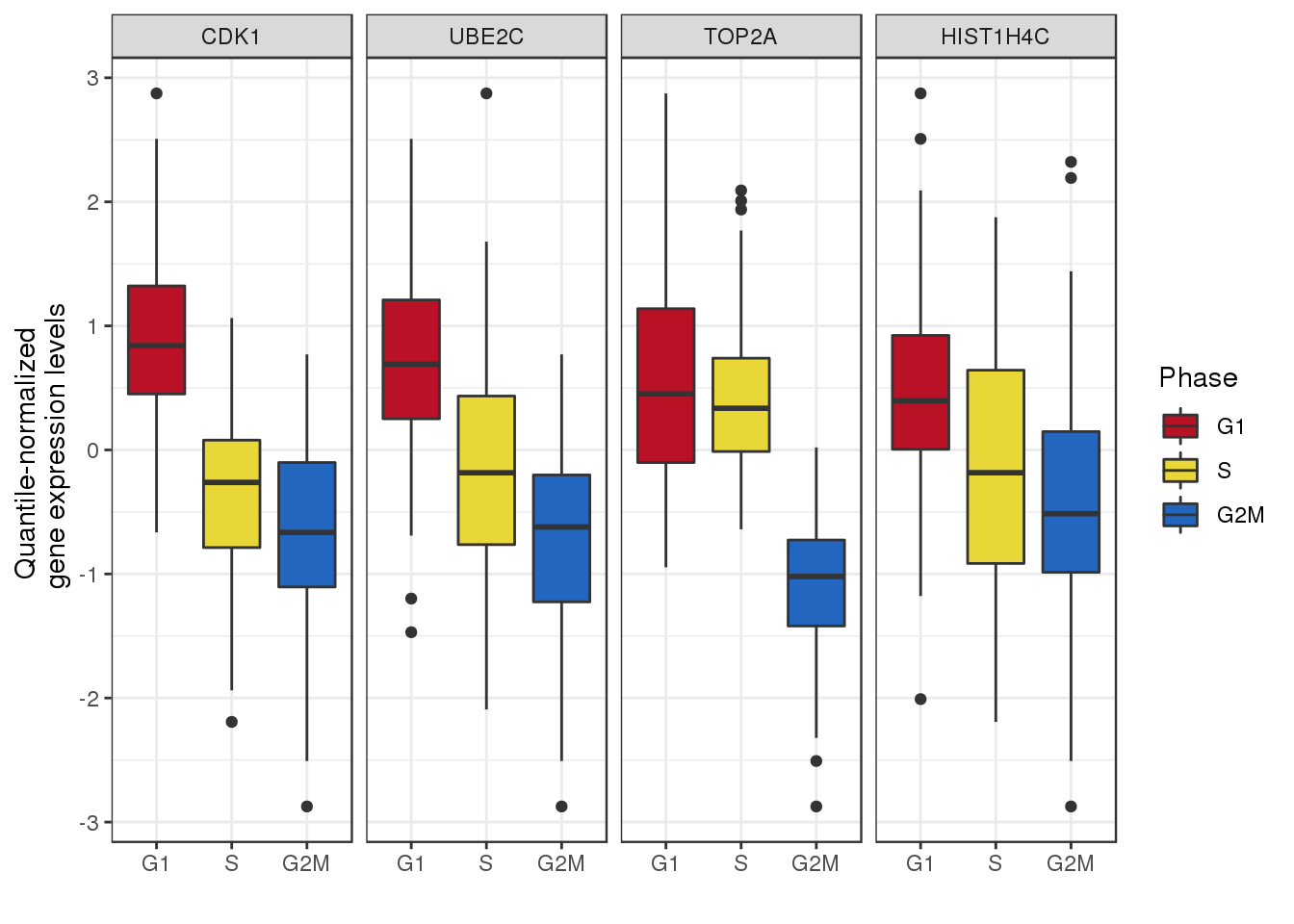

### peco-2.png

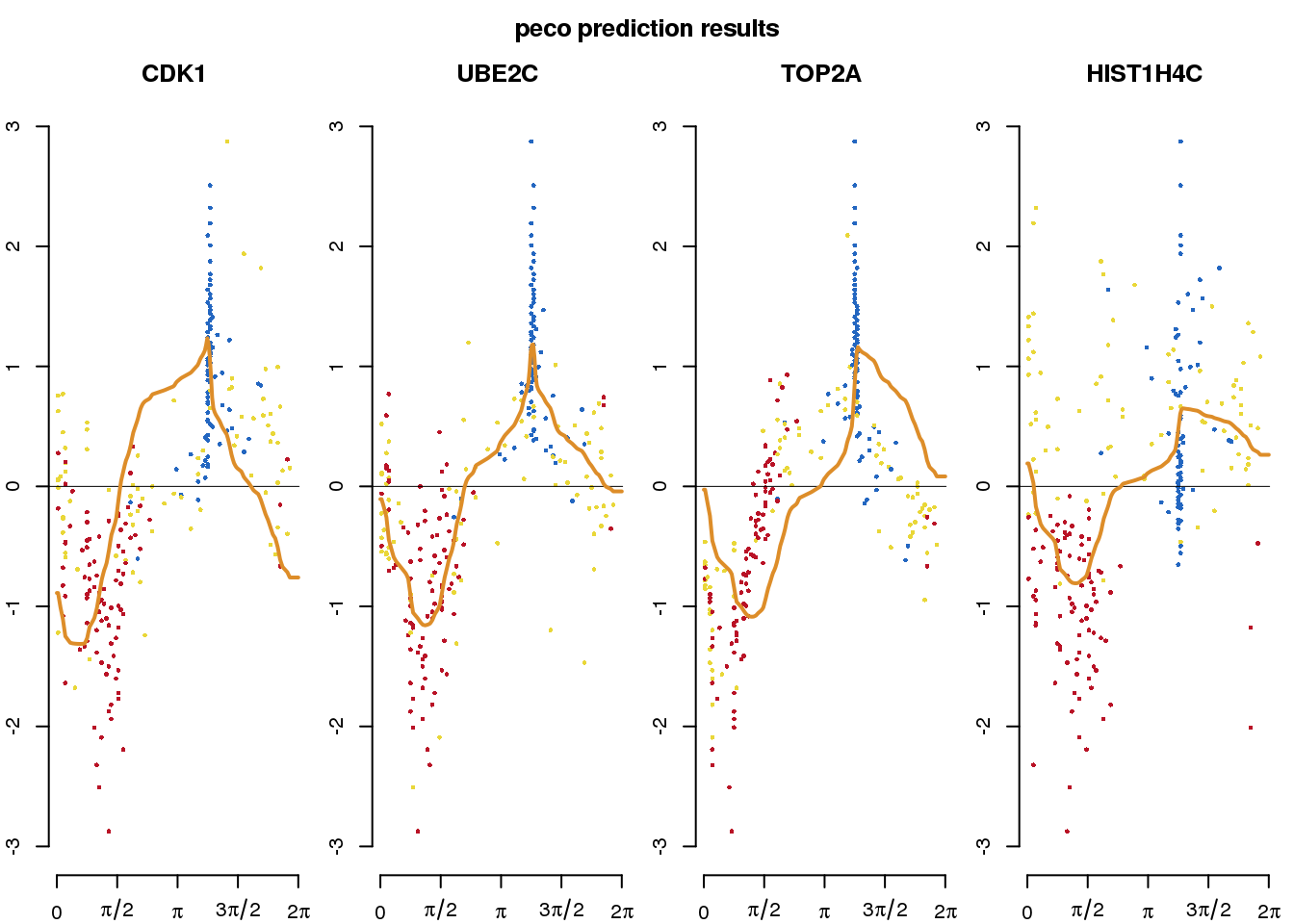

### plot-oscope-1.png

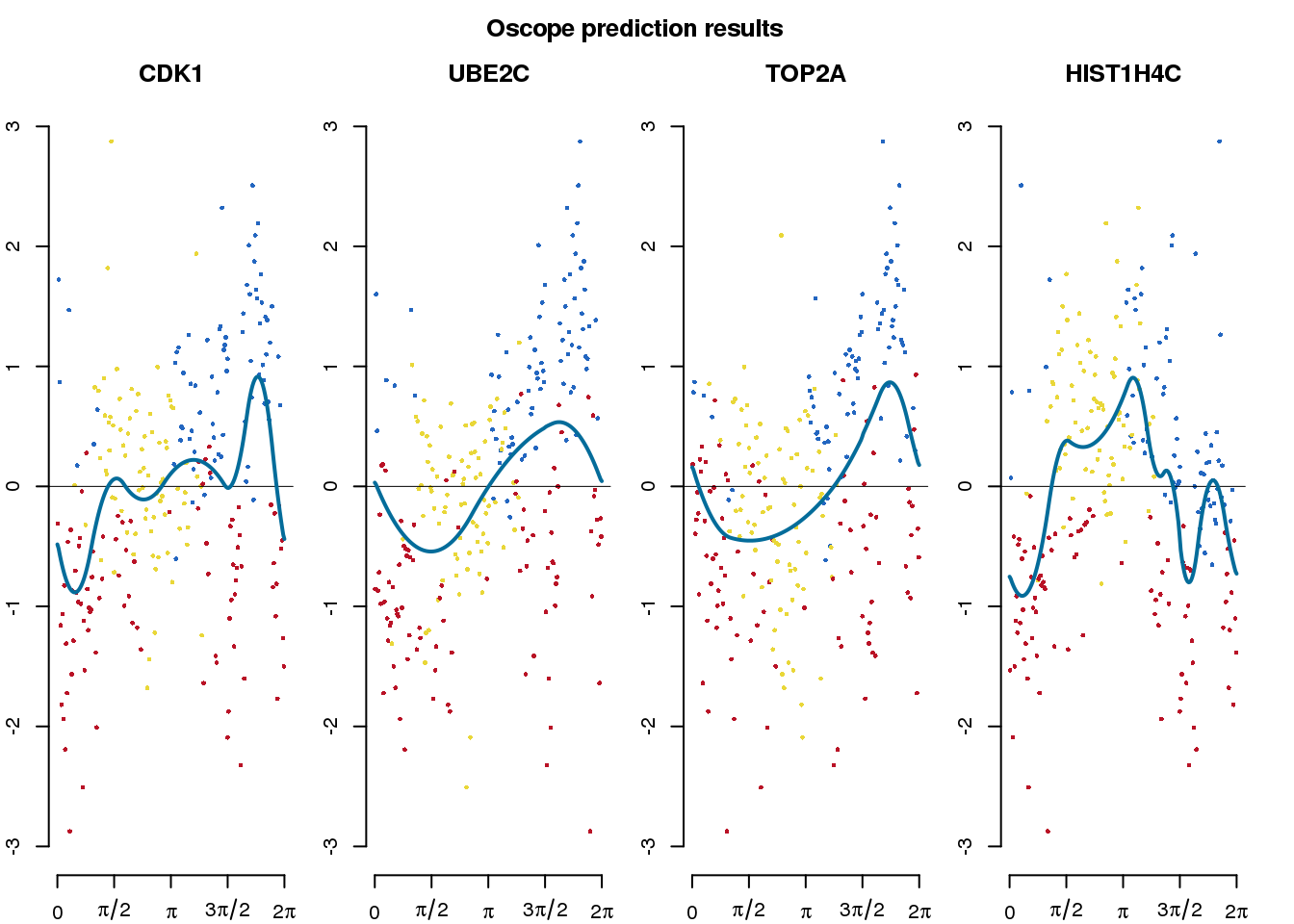

### plot-recar-1.png

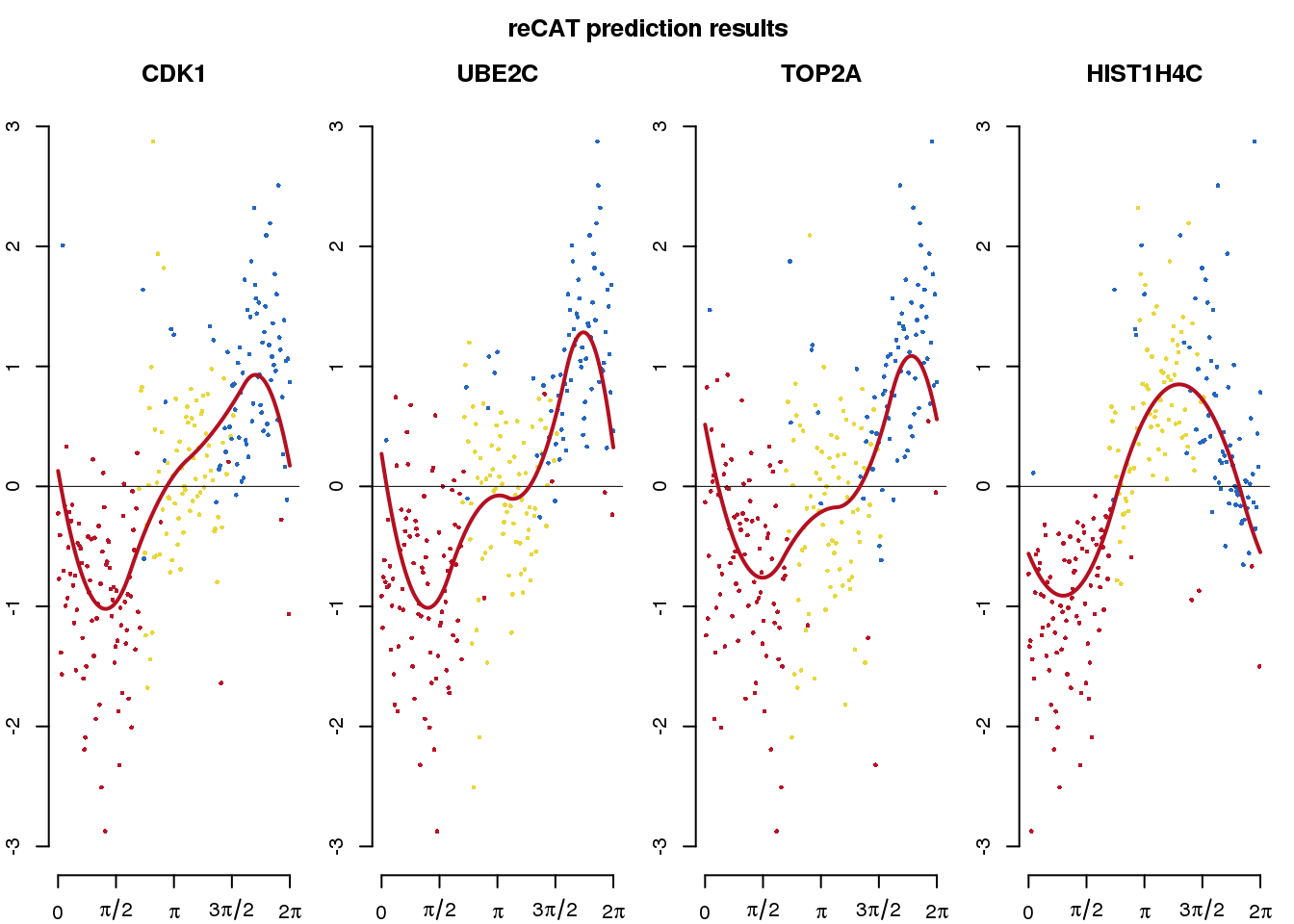

### plots-1.png

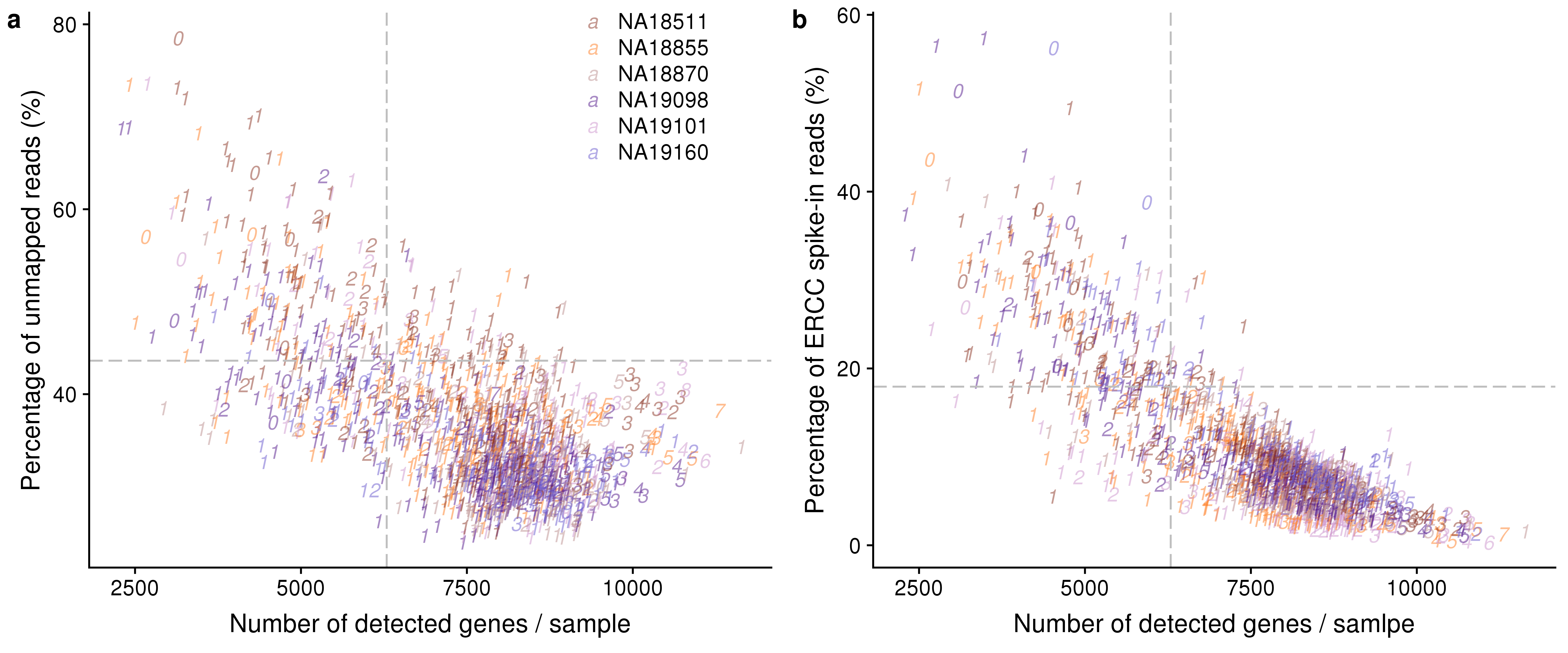

### plots-2.png

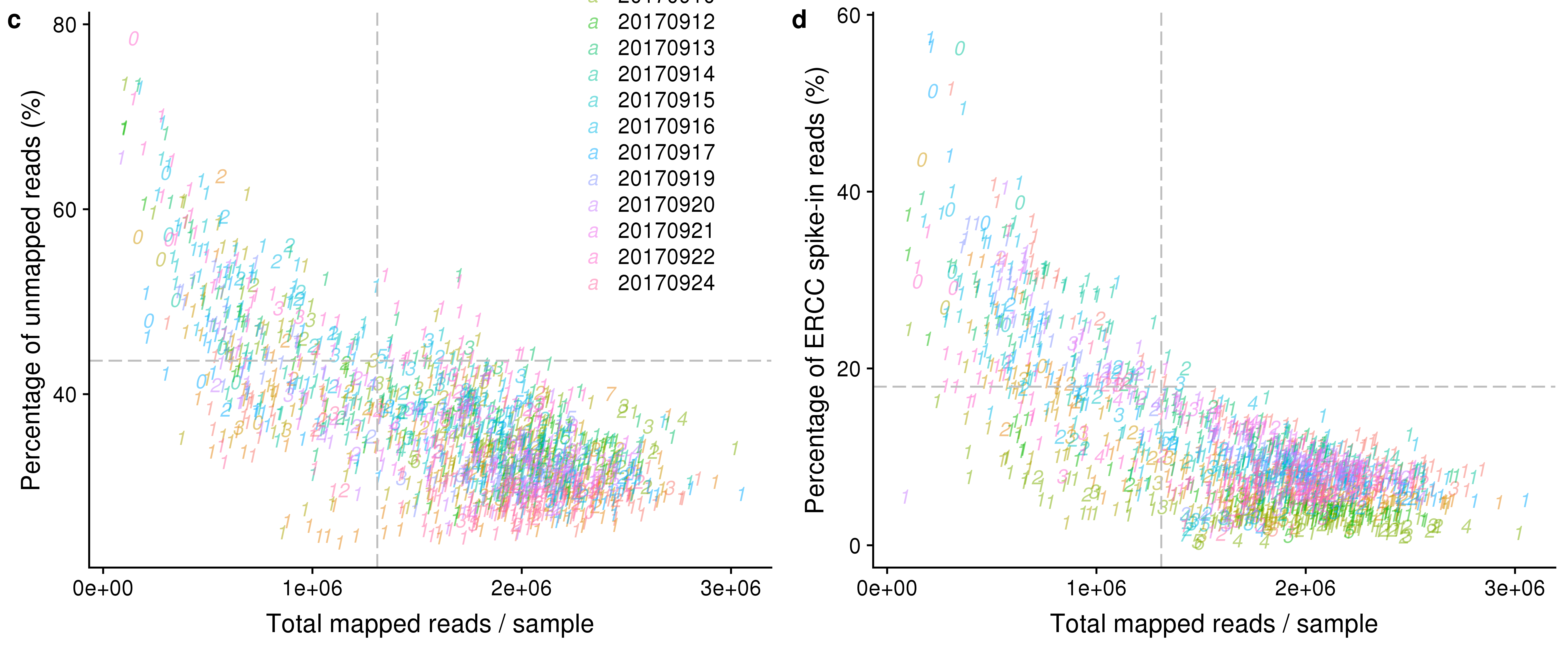

### pve-top10-genes-1.png

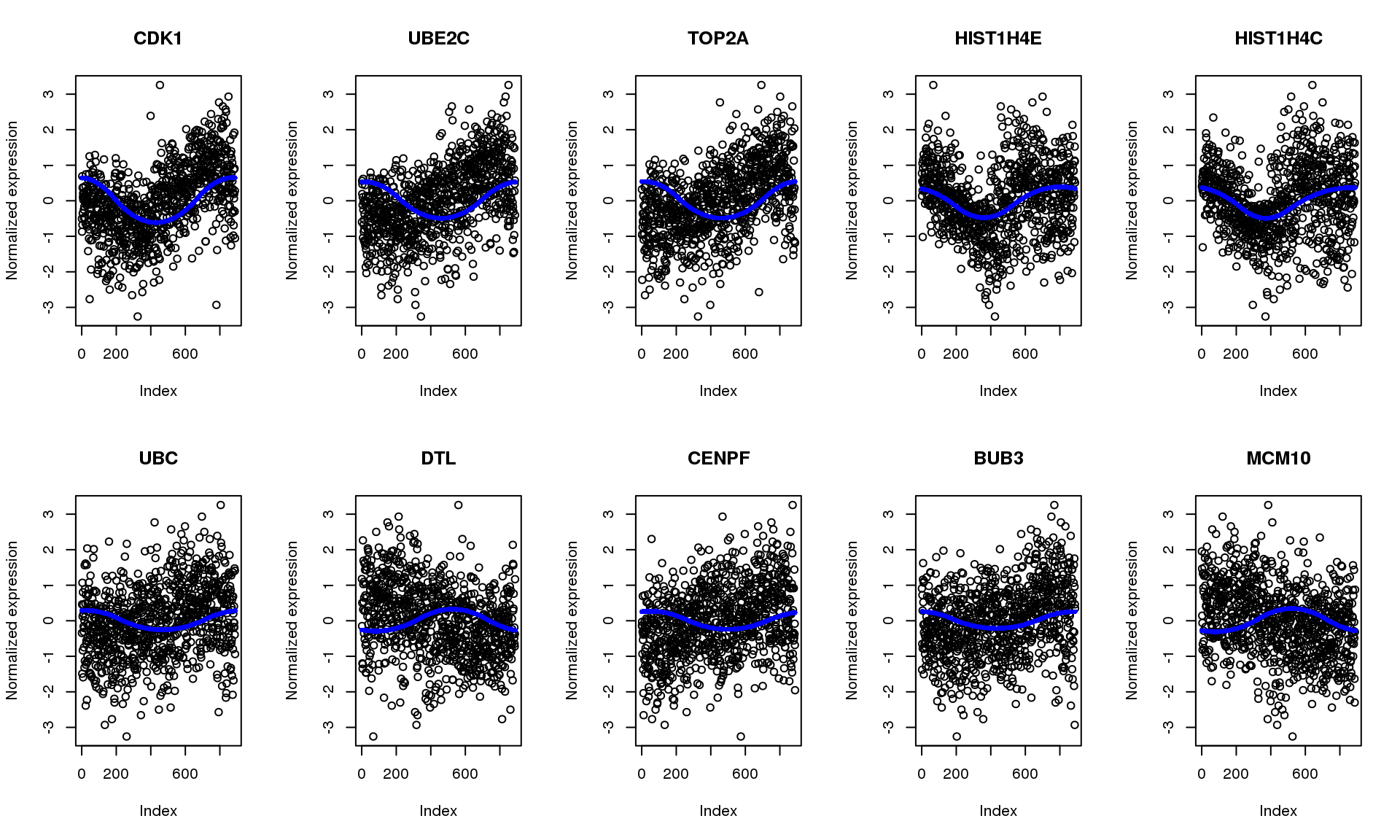

### reads-v-molecules-1.png

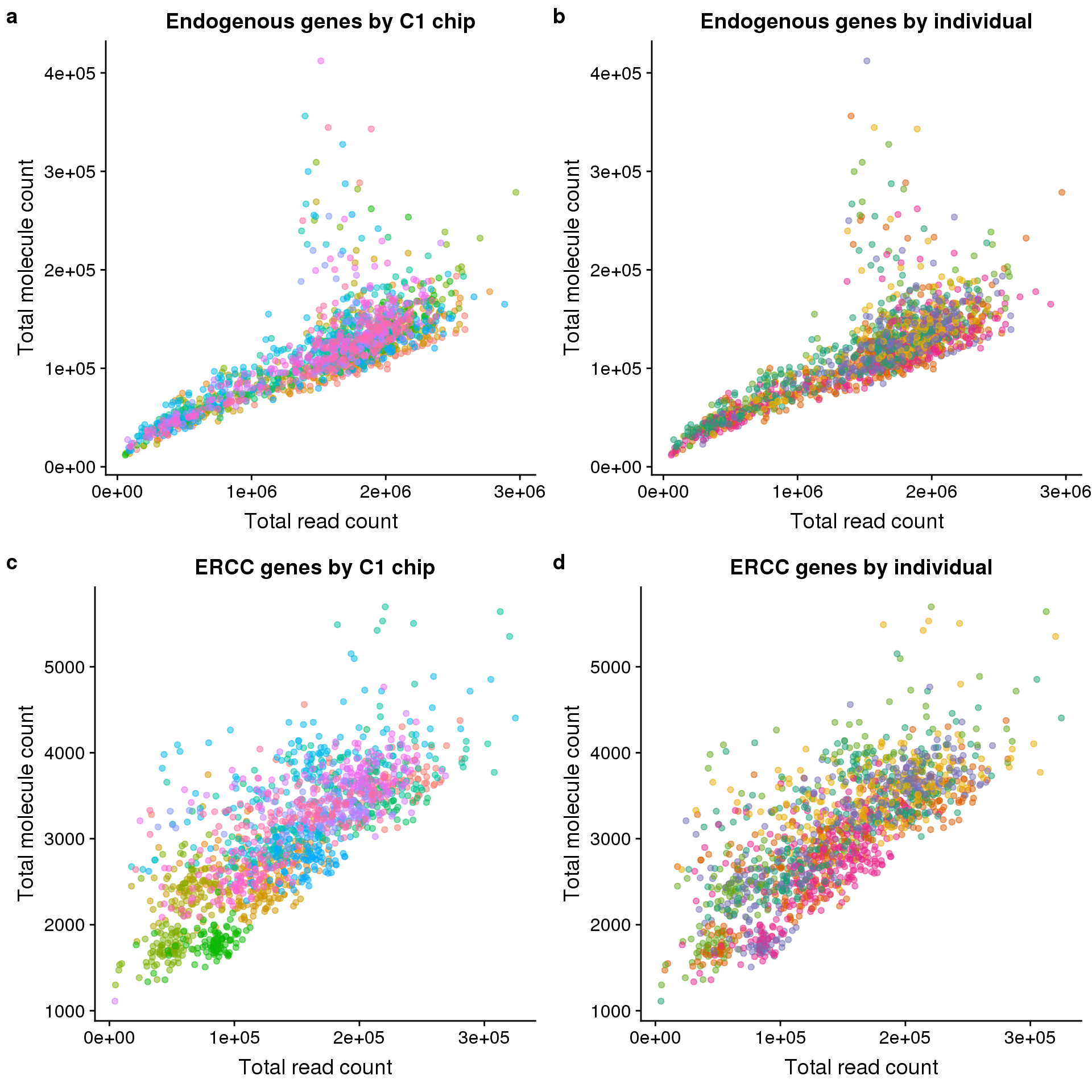

### top-1.png

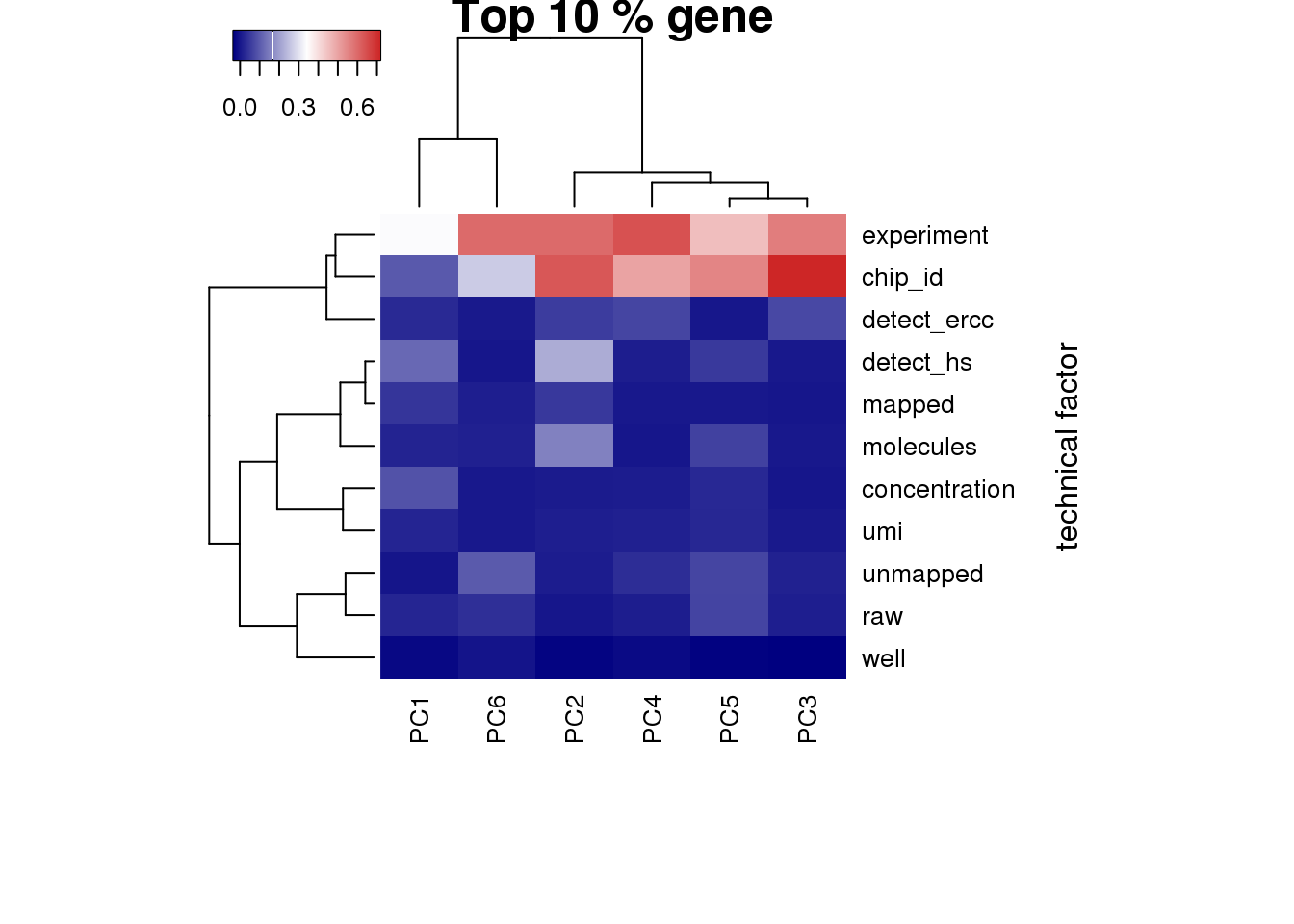

### total-reads-1.png

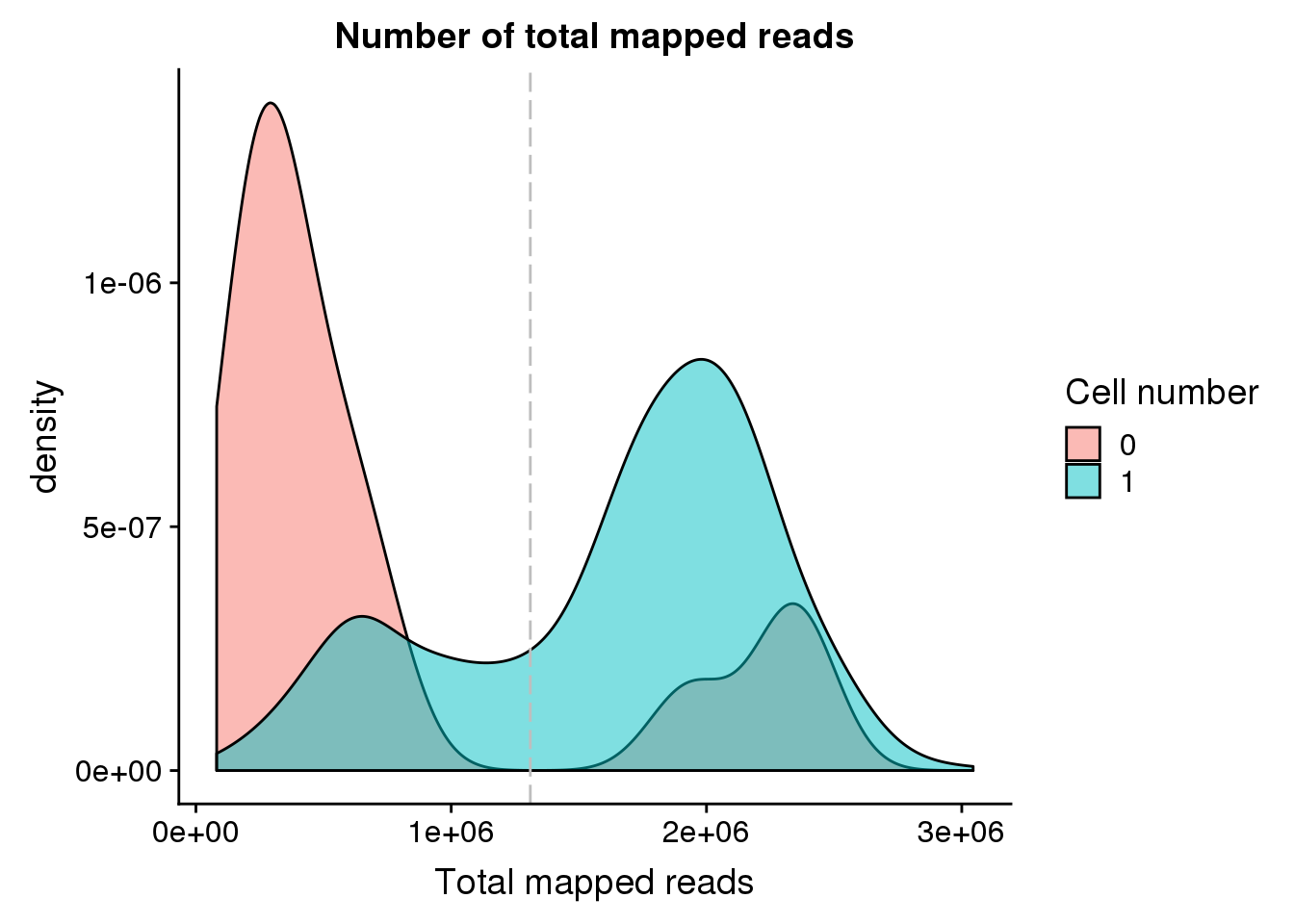

### ui-icons_444444_256x240.png

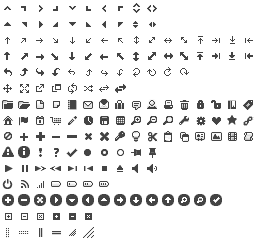

### ui-icons_555555_256x240.png

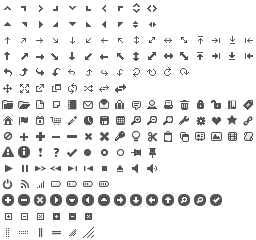

### ui-icons_777620_256x240.png

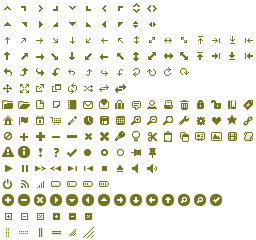

### ui-icons_777777_256x240.png

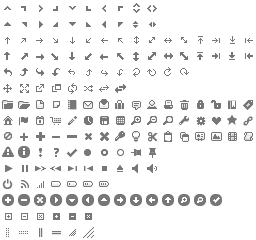

### ui-icons_cc0000_256x240.png

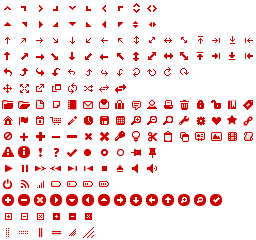

### ui-icons_ffffff_256x240.png

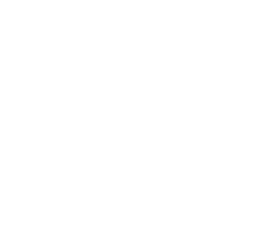

### unmapped-ratios-1.png

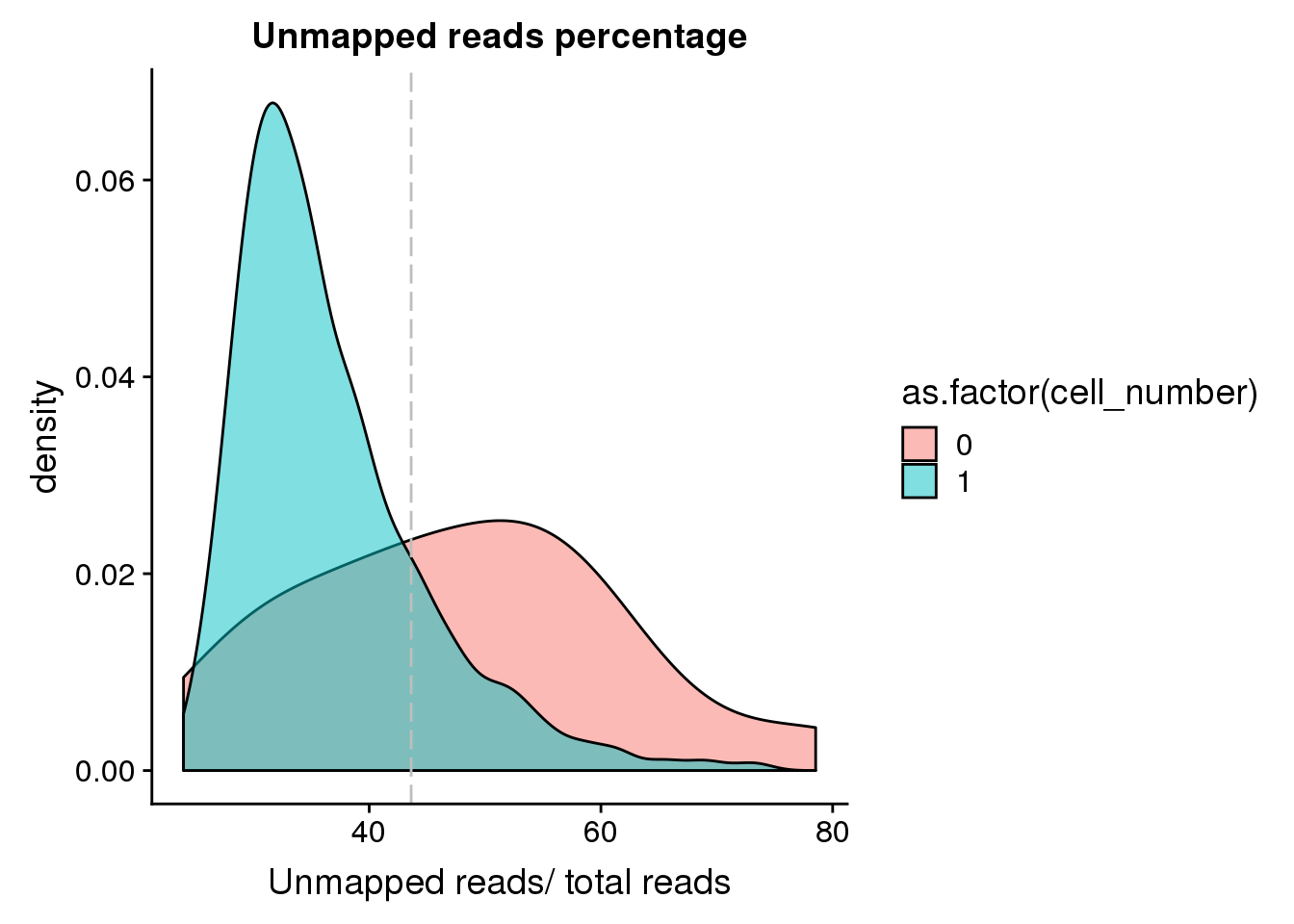

### unmapped_exp-1.png

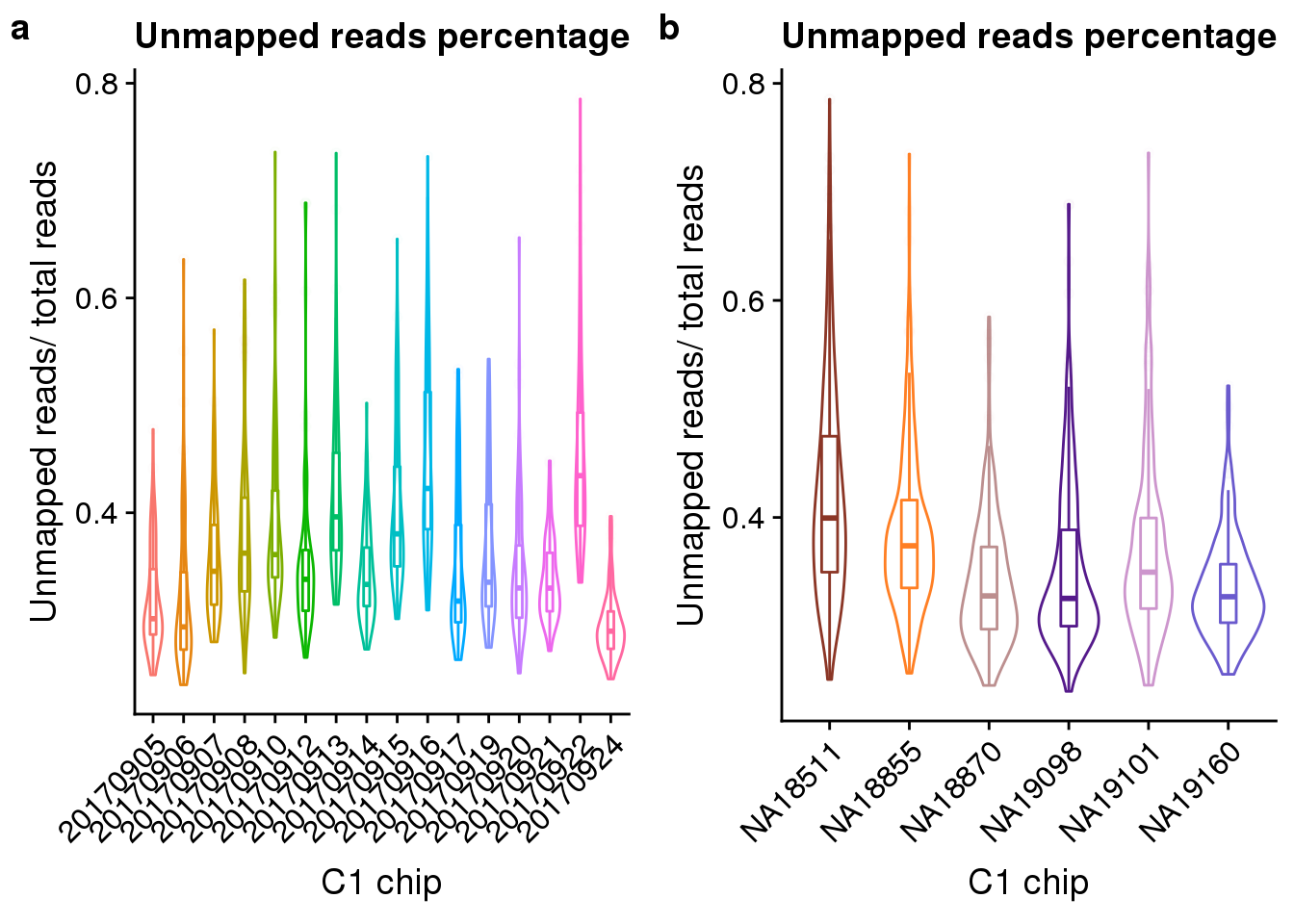

### unnamed-chunk-1-1.png

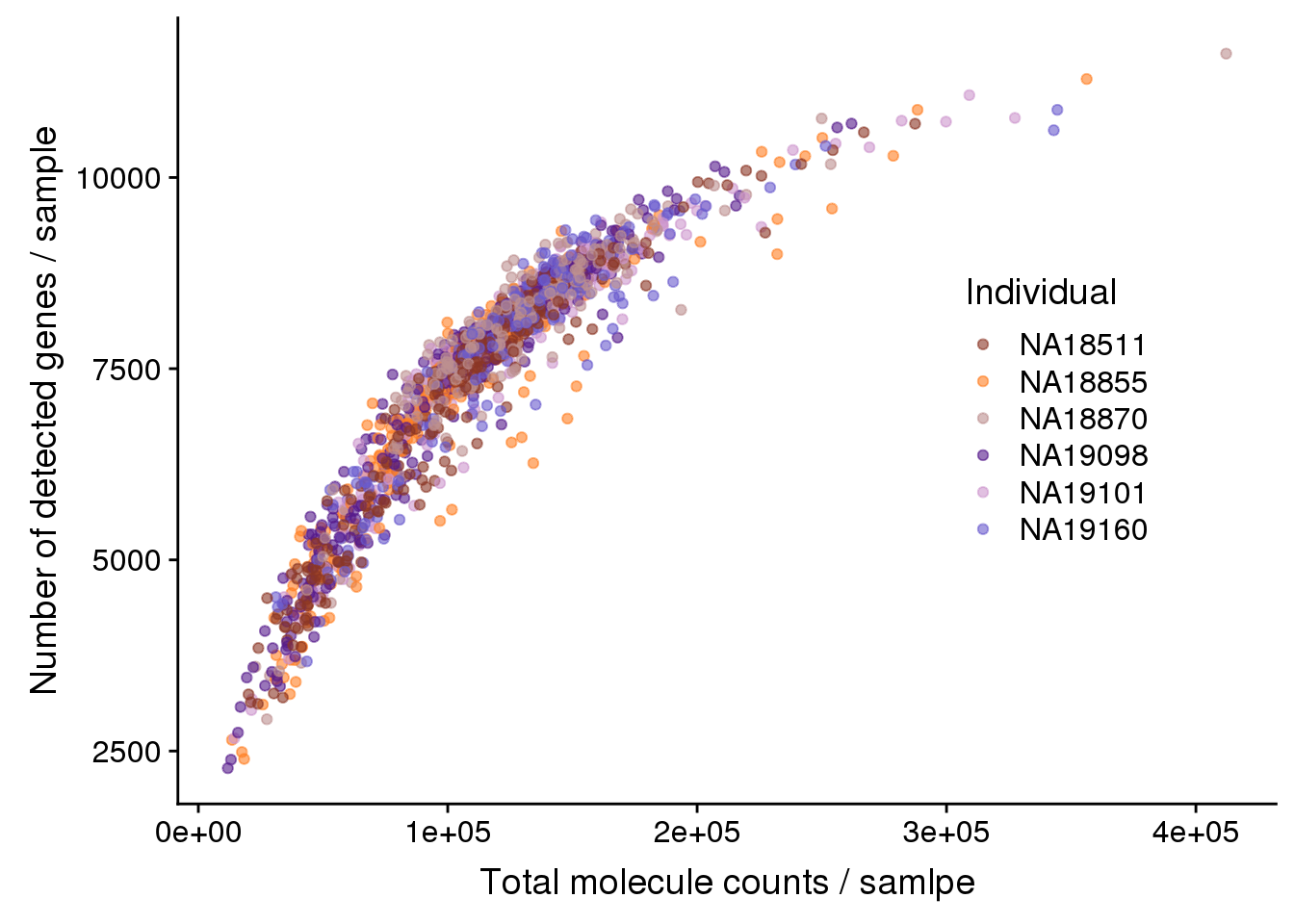

### unnamed-chunk-1-1.png

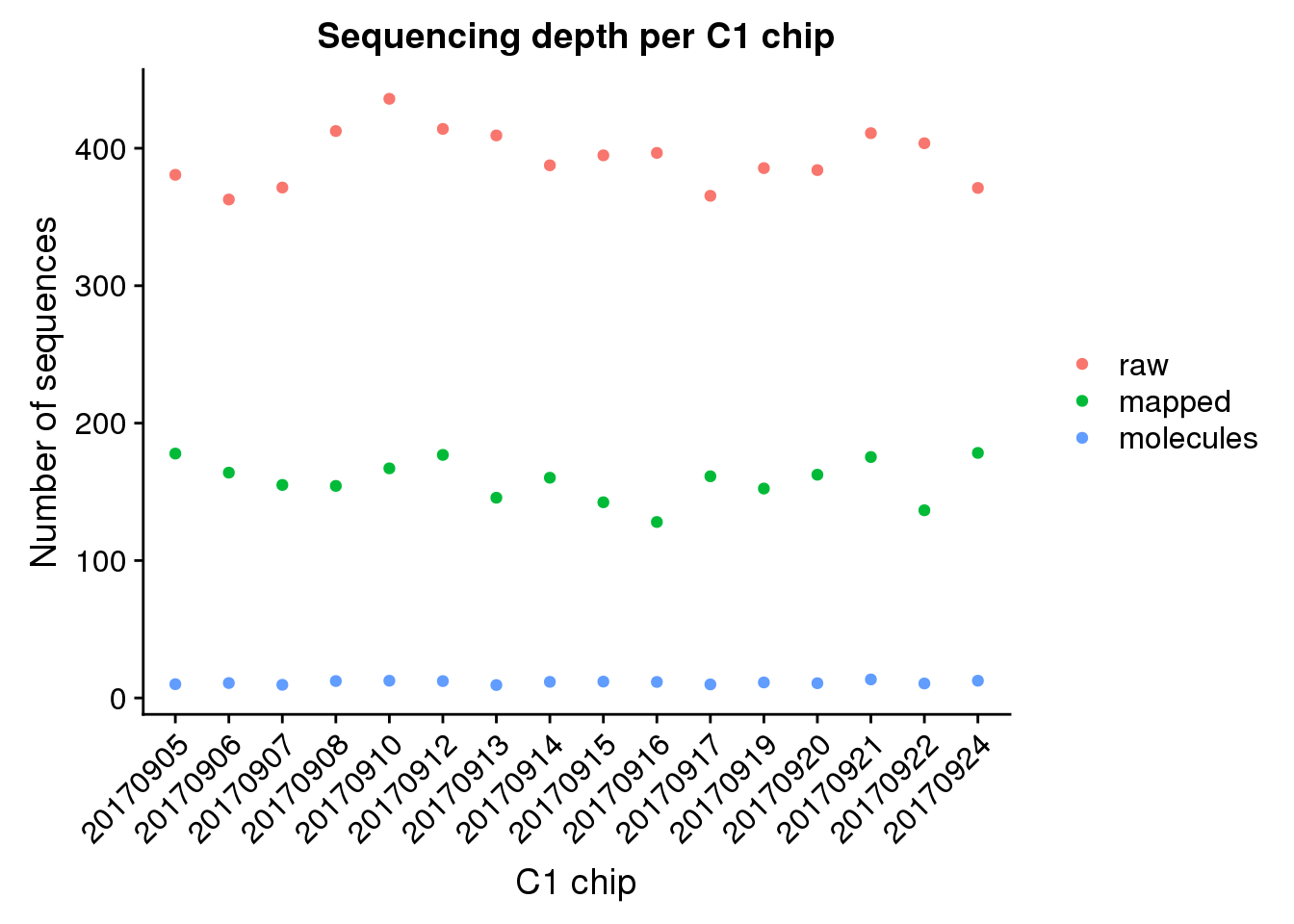

### unnamed-chunk-4-1.png

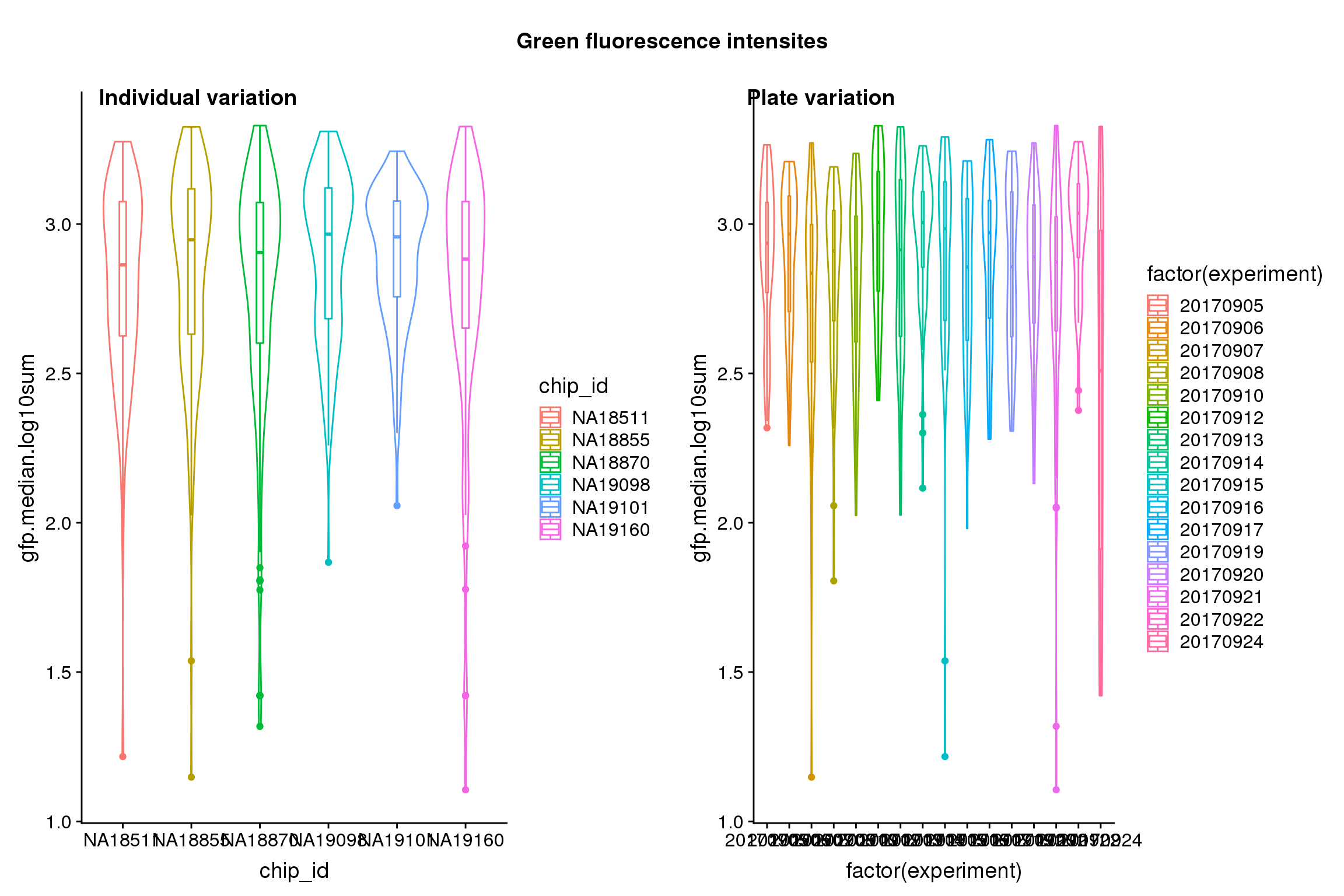

### unnamed-chunk-4-2.png

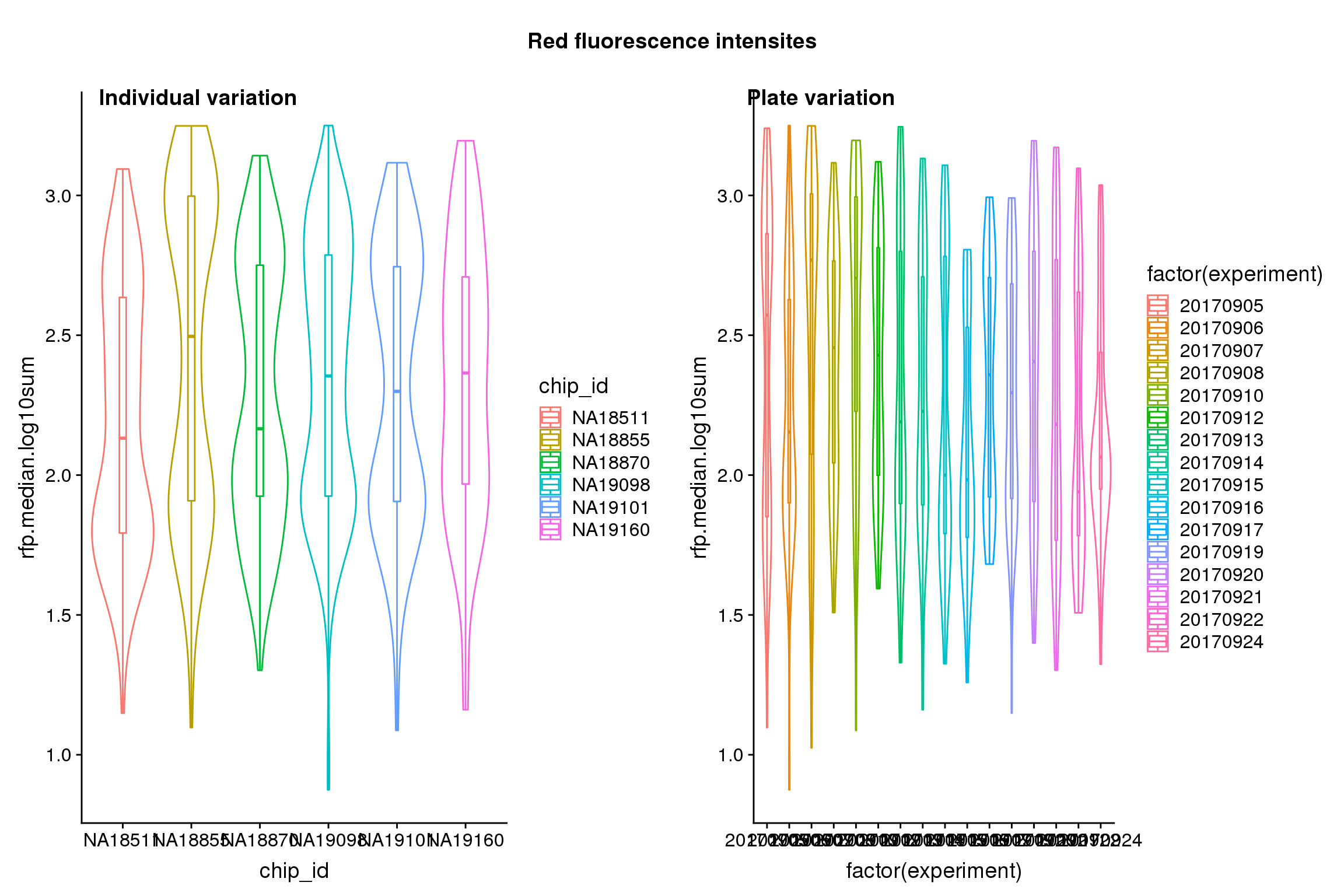

### unnamed-chunk-4-3.png

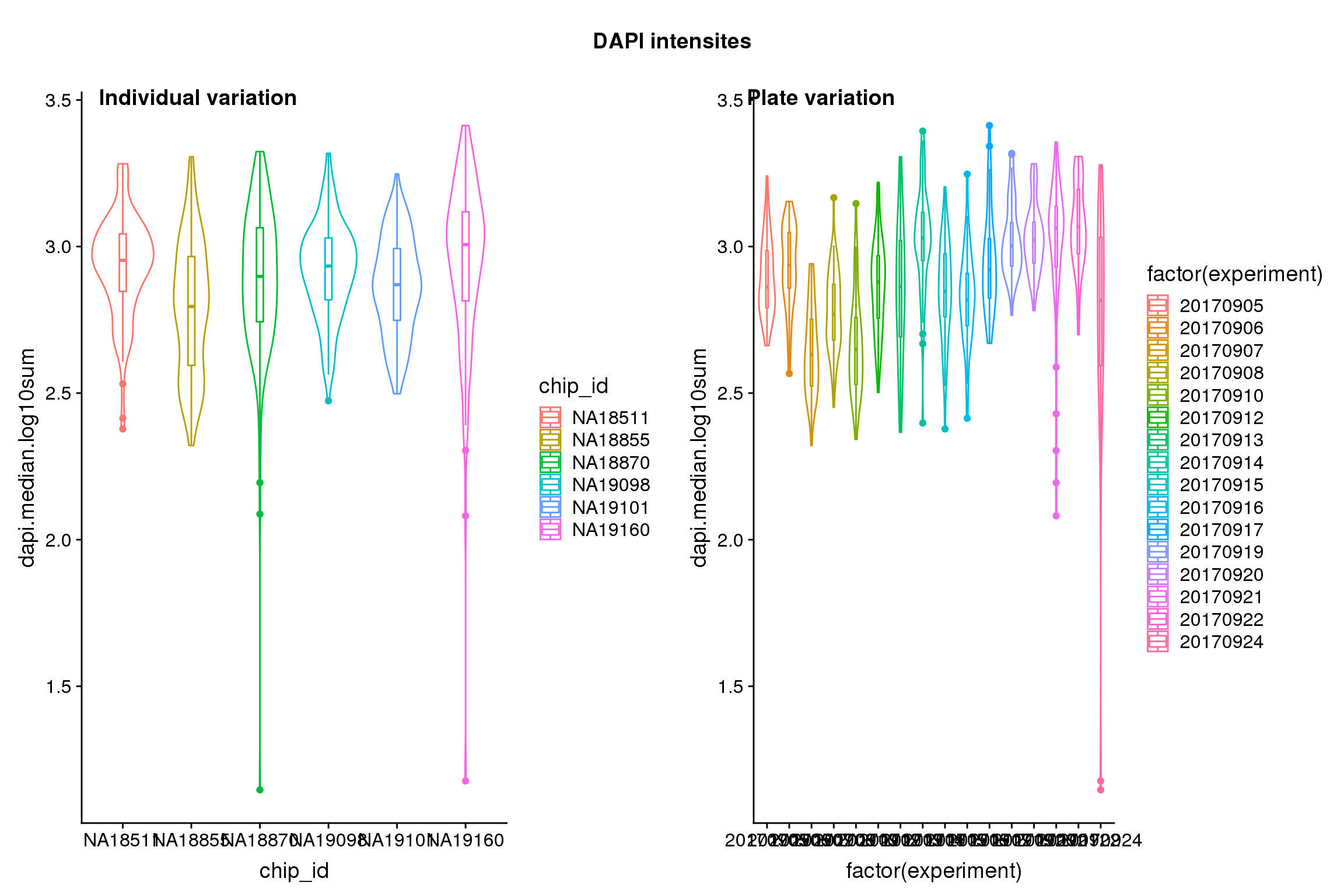

### unnamed-chunk-4-4.png

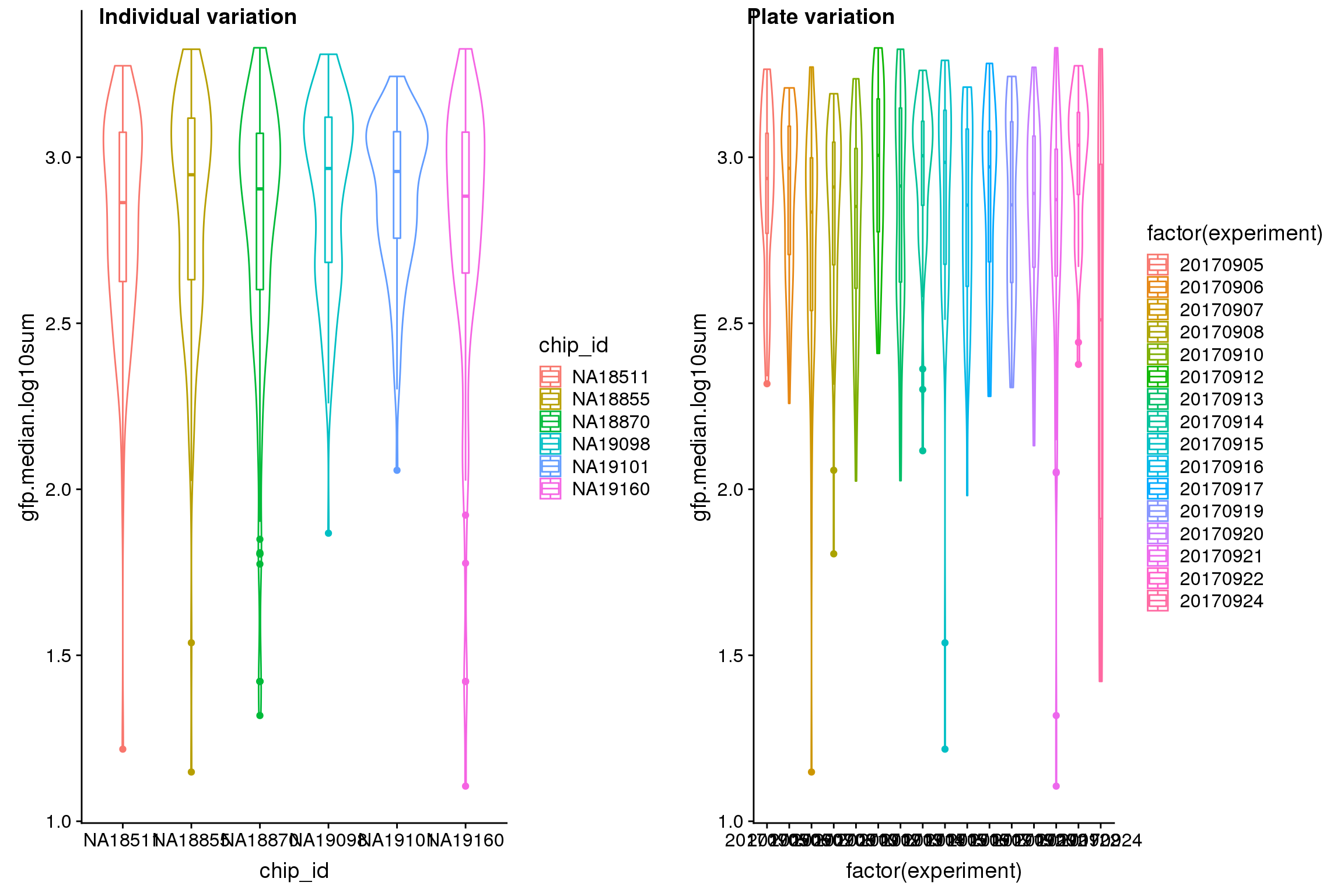

### unnamed-chunk-4-5.png

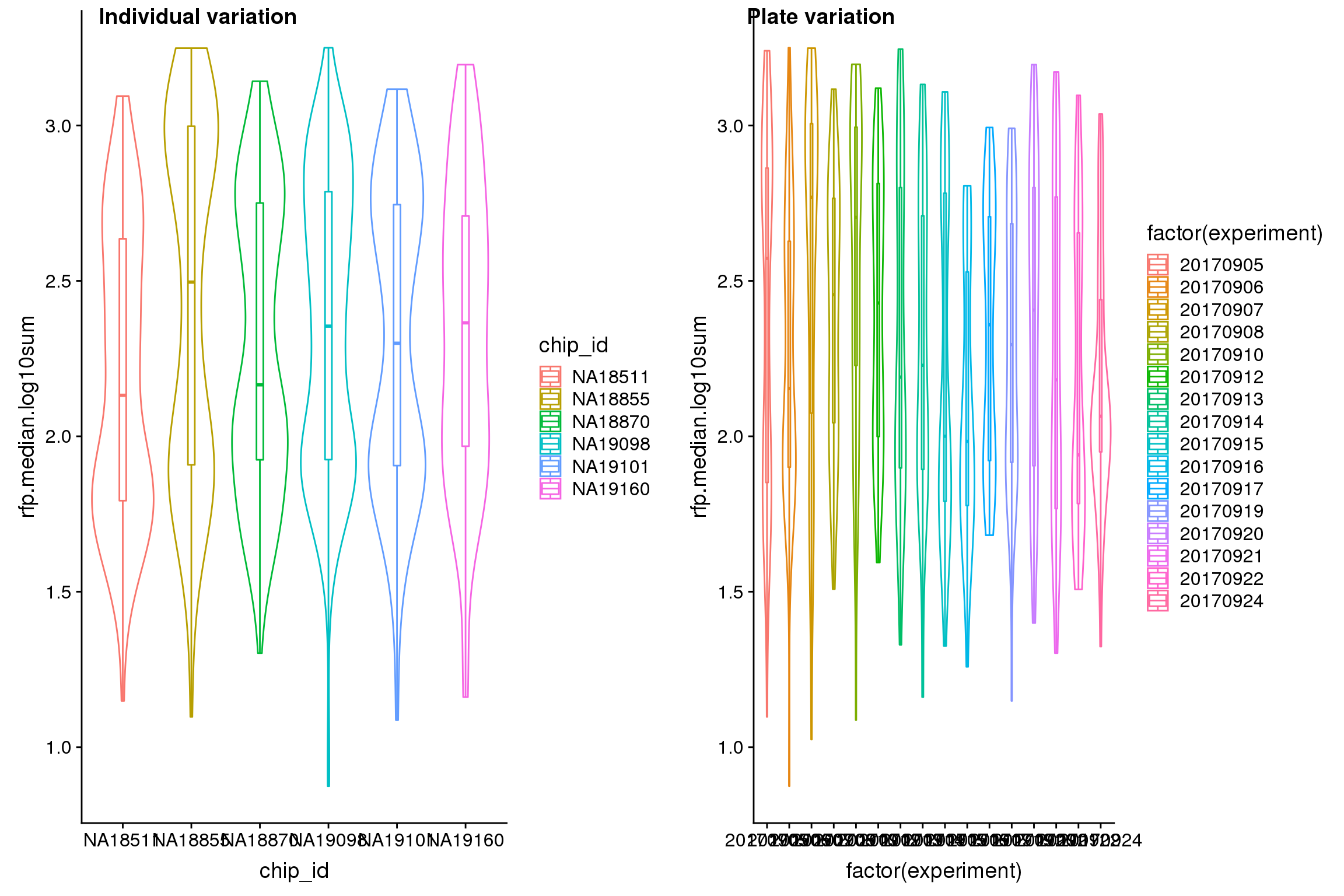

### unnamed-chunk-4-6.png

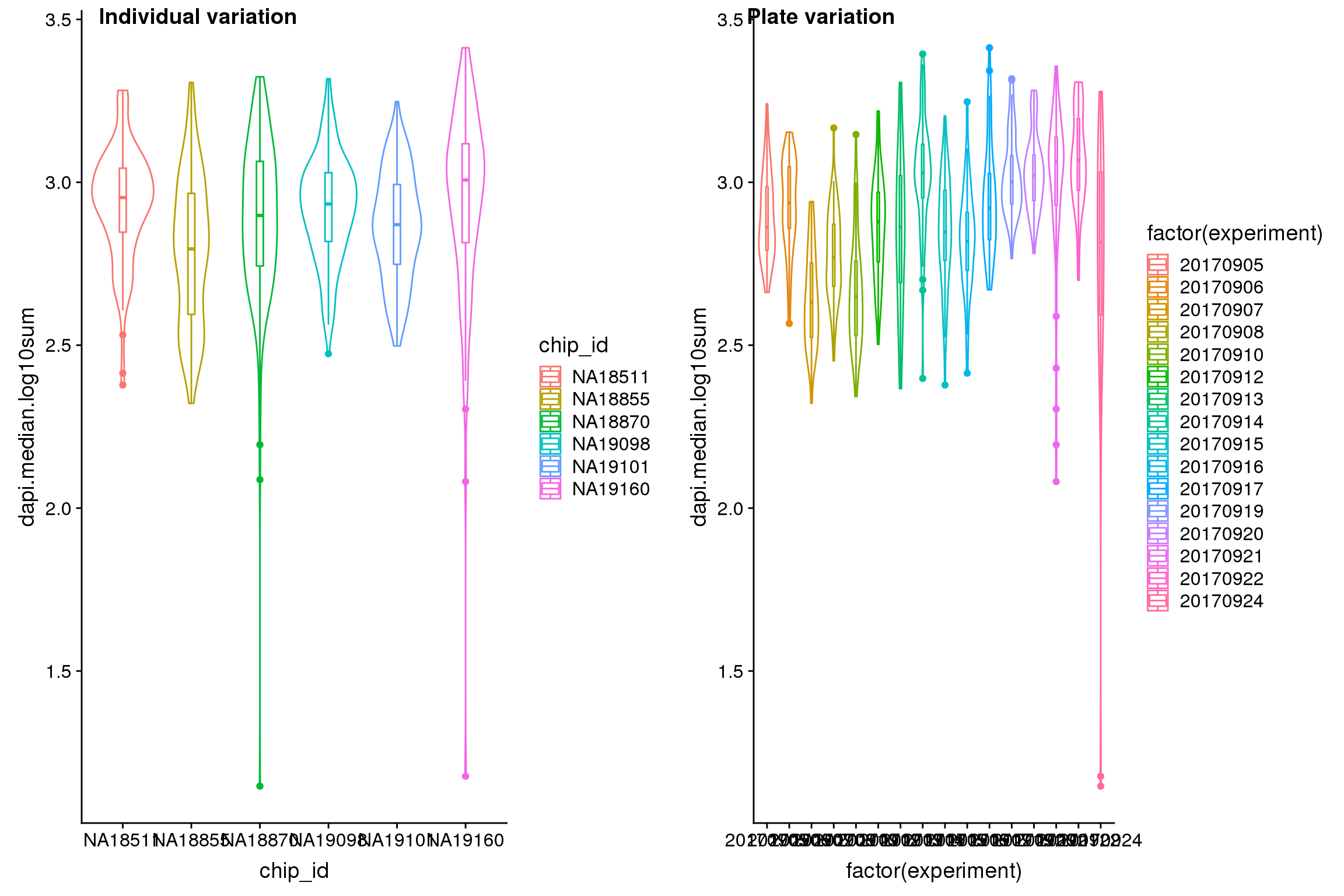
