## Supplemental figures for "Characterizing and inferring quantitative cell cycle phase in single-cell RNA-seq data analysis"

#### Supplemental Figure S1

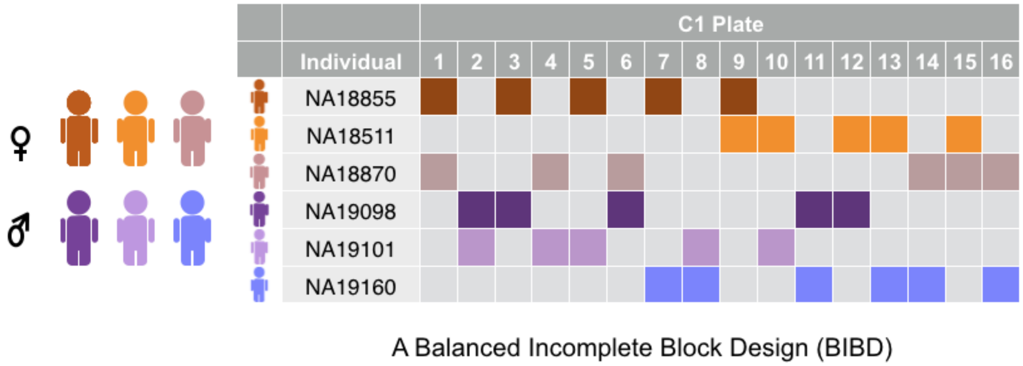


Supplemental Fig. S1: C1 study design. The table displays the distribution of cells from six individual cell lines across sixteen C1 96-well plates, with rows corresponding to cell lines and columns corresponding to C1 plates. Specifically, we used a balanced incomplete block design (BIBD) in which cells from unique pairs of individuals were distributed across fifteen 96-well C1 plates on the C1 platform. We also included data from one additional plate (containing individuals NA18855 and NA18511), which we collected as part of a pilot study. In total, we collected data from 1,536 scRNA-seq samples distributed across sixteen C1 plates.

#### Supplemental Figure S2

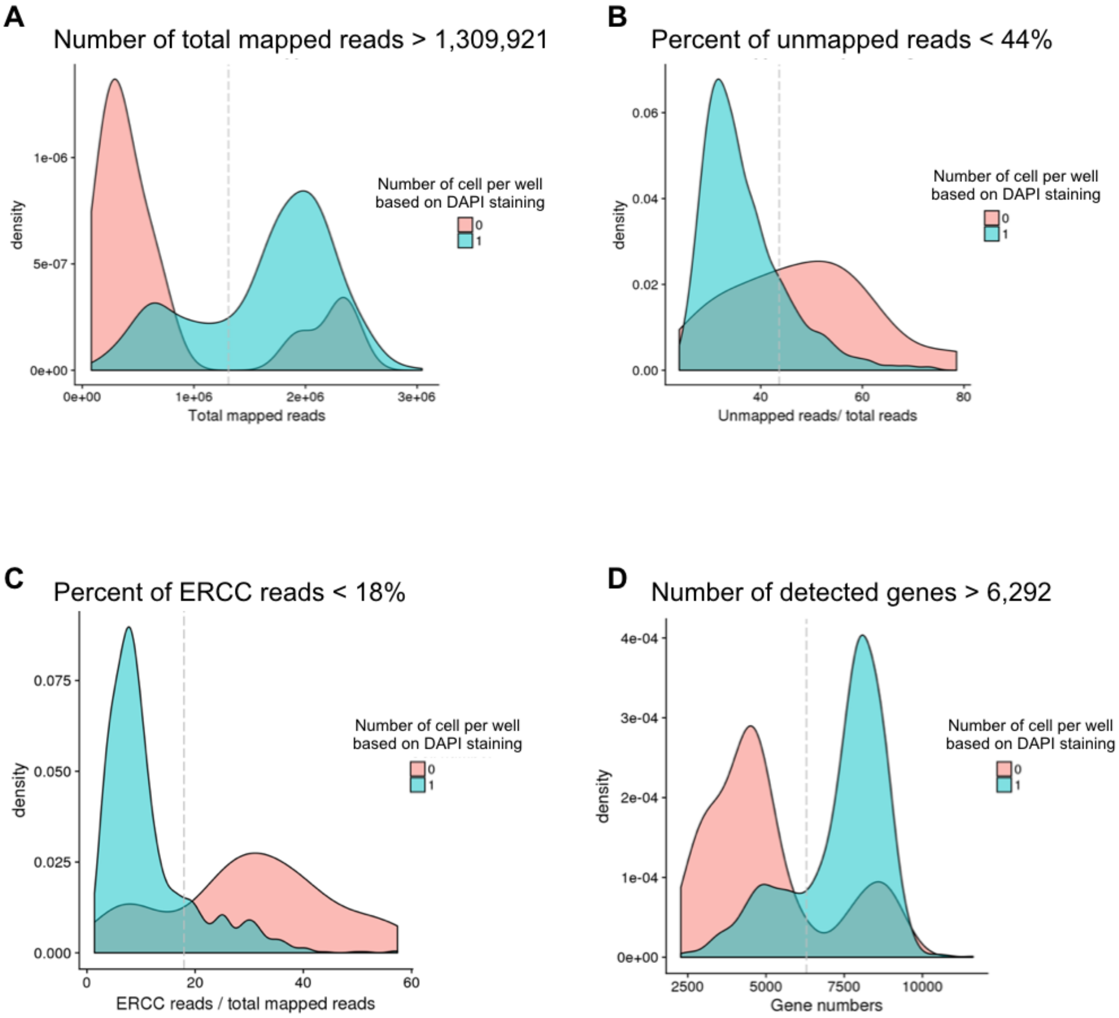


Supplemental Fig. S2: Filtering criteria for including single-cell samples. We used DAPI to determine the number of cells captured in each C1 well and compared common scRNA-seq data metrics between empty wells and single-cell wells in order to determine filtering criteria for single-cell samples. Using this approach, we determined filtering criteria for (A) the number of total mapped reads (>= 1,309,921), (B) the percentage of unmapped reads (< 44%), (C) the percentage of ERCC reads (< 18%), and (D) the number of detected genes (>= 6,292 genes with least one read).

#### Supplemental Figure S3

| | Mapped Reads (million) | Unmapped Reads Proportion (%) | Number of singleton samples | ERCC reads proportion (%) | Molecule count (million) |
|---|---|---|---|---|---|
| Overall | 2.0 (0.30) | 33.0 (4.12) | 888 | 7.6 (3.15) | .13 (.024) |
| NA18511 | 1.9 (0.25) | 35.0 (4.31) | 103 | 9.0 (2.43) | .13 (.019) |
| NA18855 | 1.9 (0.31) | 34.9 (4.17) | 163 | 8.2 (3.58) | .12 (.022) |
| NA18870 | 2.0 (0.30) | 32.5 (4.30) | 206 | 7.4 (3.47) | .13 (.023) |
| NA19098 | 2.1 (0.32) | 31.3 (2.95) | 182 | 6.8 (2.25) | .13 (.024) |
| NA19101 | 2.0 (0.29) | 33.3 (4.27) | 122 | 5.9 (3.27) | .15 (.022) |
| NA19160 | 2.0 (0.27) | 32.0 (3.14) | 112 | 8.8 (2.33) | .14 (.022) |

Supplemental Fig. S3: Summary table of scRNA-seq quality metrics. We computed means and standard deviations (in parentheses) of these metrics across single-cell samples for each of the six cell lines and also for the entire dataset.

#### Supplemental Figure S4

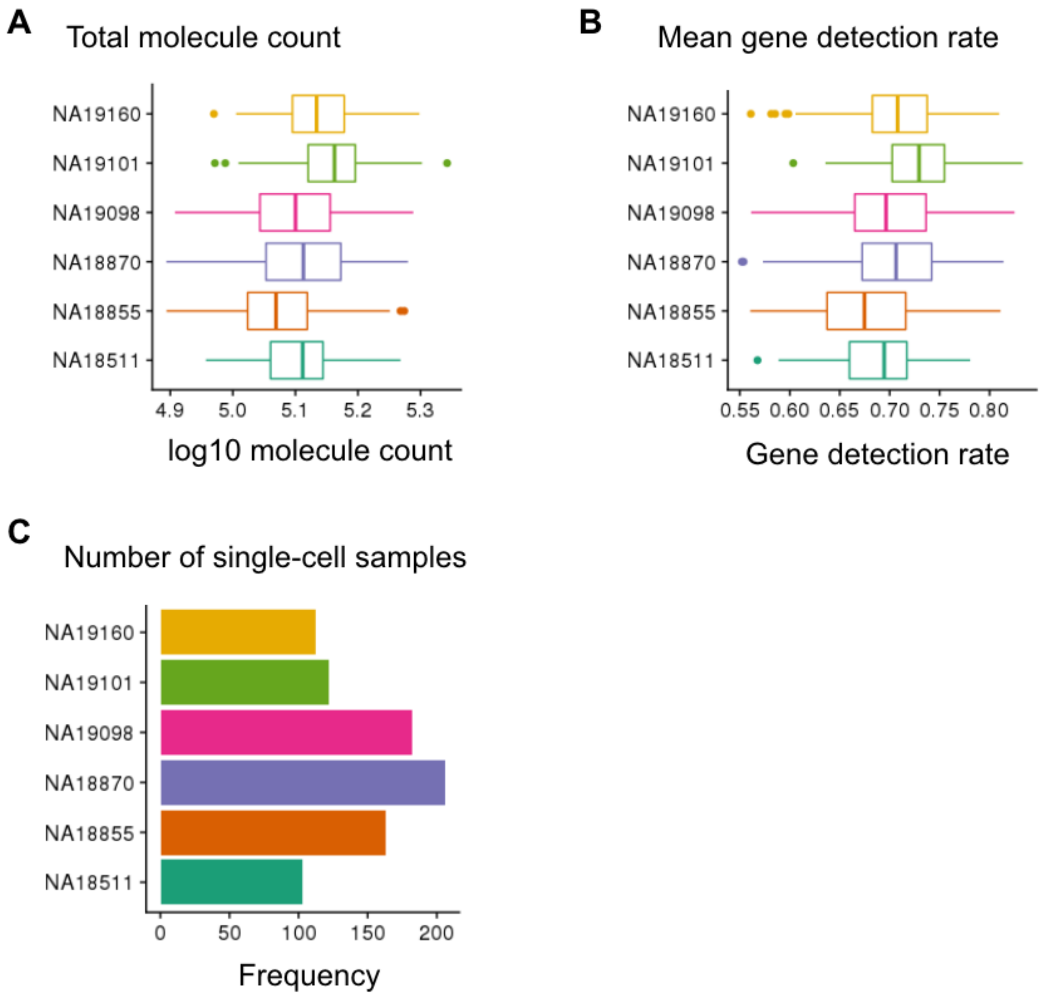


Supplemental Fig. S4: Distribution of scRNA-seq quality metrics for the six cell lines. We show the distribution of single-cell samples in (A) the total molecule count, (B) the mean gene detection rate (i.e., fraction of genes with at least one read), (C) the number of single-cell samples.

#### Supplemental Figure S5

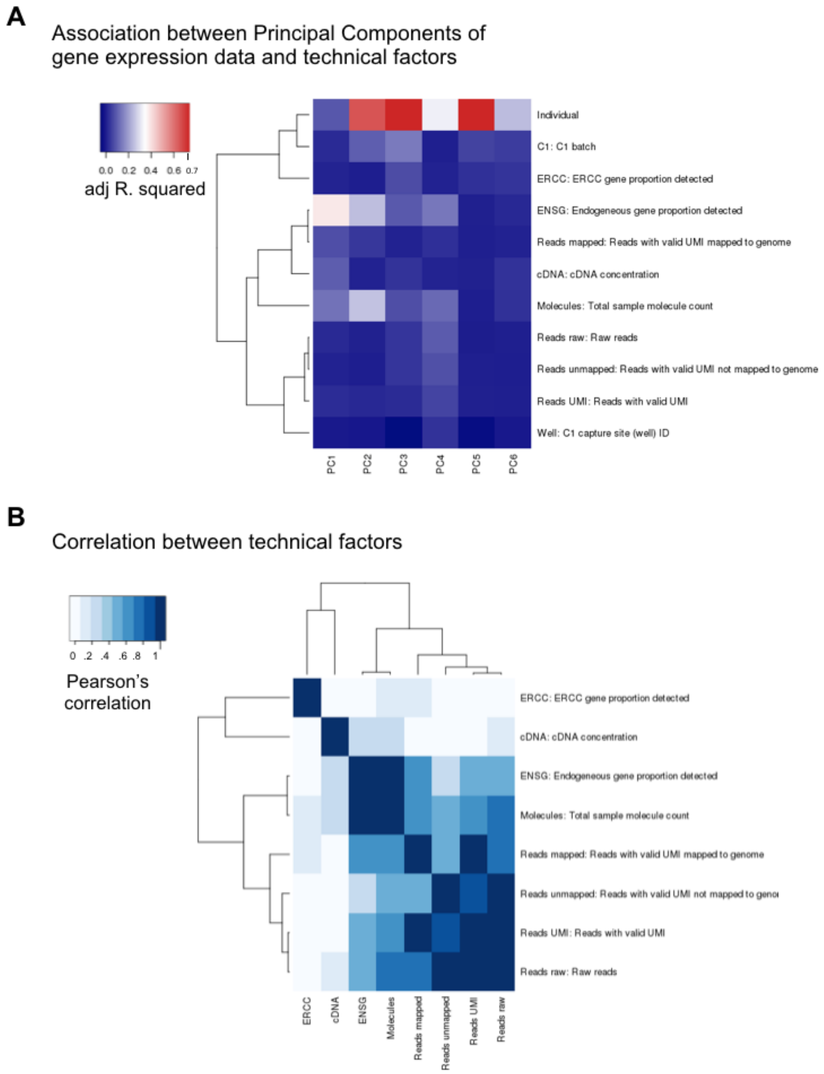


Supplemental Fig. S5: Major sources of variation in our gene expression data of 888 quality samples and 11,040 genes. (A) Principal Component Analysis (PCA) was applied to the $\log_2$ CPM of the gene expression data. We computed the proportion of variance explained (i.e., adjusted R-squared) in each of the principal components by: individual identity of the single-cell sample (Individual), C1 processing batch (C1), capture site or well (Well), fraction of ERCC genes detected (ERCC), fraction of endogenous genes detected (ENSG), cDNA concentration (cDNA), sample total molecule count (Molecules), number of raw reads (Reads raw), number of raw reads with valid UMI (Reads UMI), number of reads with valid UMI mapped to the genome (Reads mapped), and number of reads with valid UMI not mapped to the genome (Reads unmapped). (B) Pearson's correlation between technical factors that are known to influence sample variation in gene expression data.

#### Supplemental Figure S6

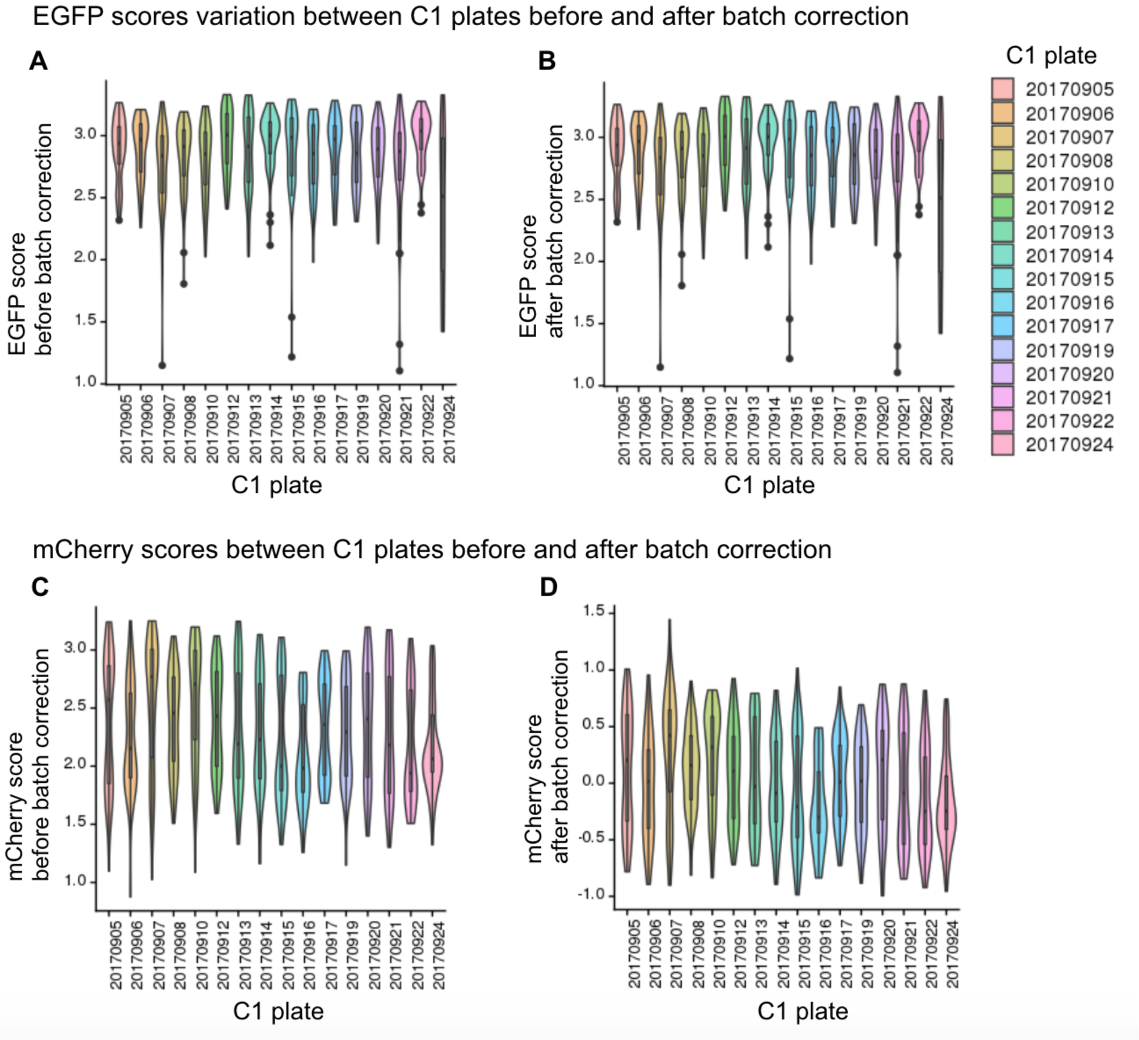


Supplemental Fig. S6: FUCCI scores for the sixteen C1 plates before and after correcting for C1 plate effect. EGFP and mCherry scores are computed by taking $\log_{10}$ sum of fluorescence intensity in the predefined cell area after background noise correction. (A) and (B) show EGFP scores variation between C1 plates before and after correcting for C1 plate effect. (C) and (D) show mCherry scores before and after correcting for C1 plate effect. We applied a linear model to account for plate effects on these scores without removing individual effects.

#### Supplemental Figure S7

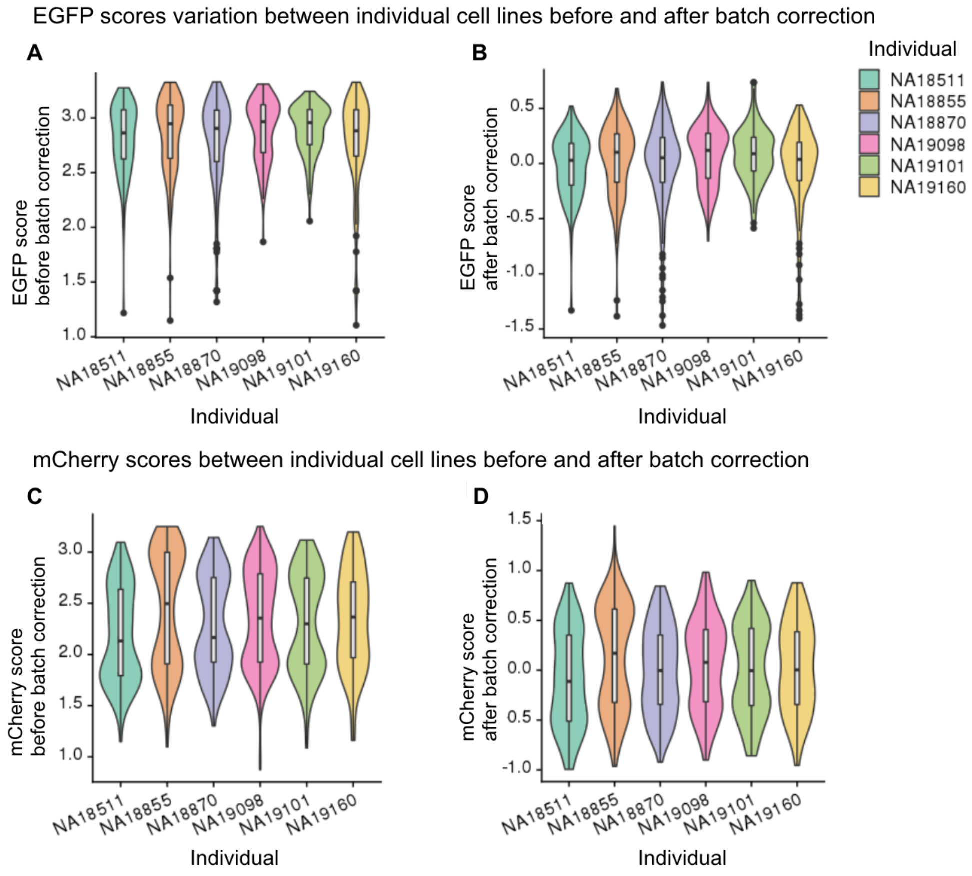


Supplemental Fig. S7: FUCCI scores for the six cell lines before and after correcting for C1 plate effect. We computed EGFP and mCherry scores by taking $\log_{10}$ sum of fluorescence intensity in the predefined cell area after background noise correction. (A) and (B) show EGFP scores variation between individual cell lines before and after correcting for C1 plate effect. (C) and (D) show mCherry scores variation before and after correcting for C1 plate effect. We applied a linear model to account for plate effects on these scores without removing individual effects.

#### Supplemental Figure S8

**A** PAM-based classification

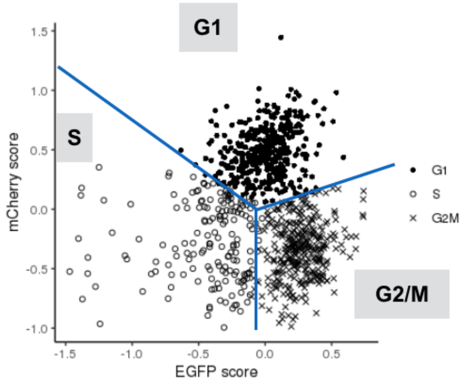


**B** PAM-based classification vs Whitfield et al. 2002 classification

Whitfield et al. cell-cycle classification

| PAM-based classification | G1/S | S | G2 | M | M/G1 | Misclassfication rate |
|---|---|---|---|---|---|---|
| G1 | 53 | 39 | 13 | 27 | 22 | 51% |
| S | 8 | 14 | 14 | 24 | 25 | 74% |
| G2/M | 26 | 51 | 99 | 91 | 53 | 24% |

*Note*. Misclassfication rate of G1 genes = fraction of G1 genes in PAM-based classification that were classifed as S, G2 or M genes. Misclassfication rate of S genes in PAM-basd classification that were classifed as G2, M, or M/G1 genes. Misclassfication rate of G2/M genes = fraction of G2/M genes in PAM-based classfication that were classifed as G1/S or S genes in Whitfield et al. 2002.

Supplemental Fig. S8: Classification obtained from the PAM-based method. We applied Partition Around Medoids (PAM) to EGFP and mCherry scores to cluster the 888 single-cell samples into G1, S, or G2/M phase (384, 172 332 cells in each phase, respectively), using *pam* function in the R package *clust* (Maechler et al. 2019). (A) shows a scatter plot of mCherry scores (Y-axis) versus EGFP scores (X-axis) overlaid with phase boundaries obtained from the PAM method (blue lines). (B) compares discrete cell cycle assignment using PAM-based classification versus using Whitfield et al. (2002) cell cycle classifications.

#### Supplemental Figure S9A-C

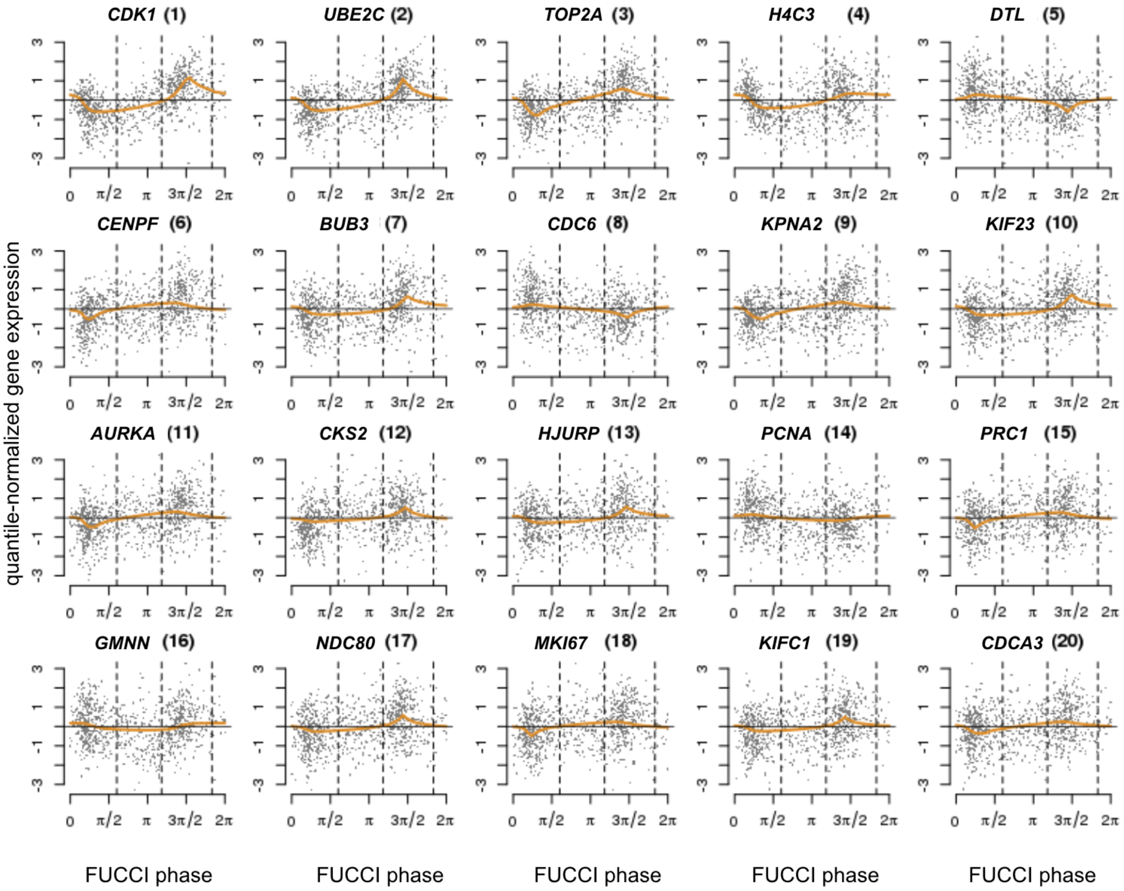


Supplemental Fig. S9A: Cyclic trends of gene expression based on FUCCI phase in 54 genes (a subset of the top 101 cyclic genes also known to be cell cycle genes in Whitfield et al. 2002). For each gene, we ordered 888 single-cell samples by FUCCI phase and applied trend filtering to estimate the cyclic trend of gene expression. The orange line corresponds to the fitted cyclic trend based on FUCCI phase. We ordered the 54 genes by their proportion of variance explained (PVE) by the fitted cyclic trend and showed their order in the figure title (in parenthesis) from large to small PVE. This figure shows top 20 of the 54 genes.

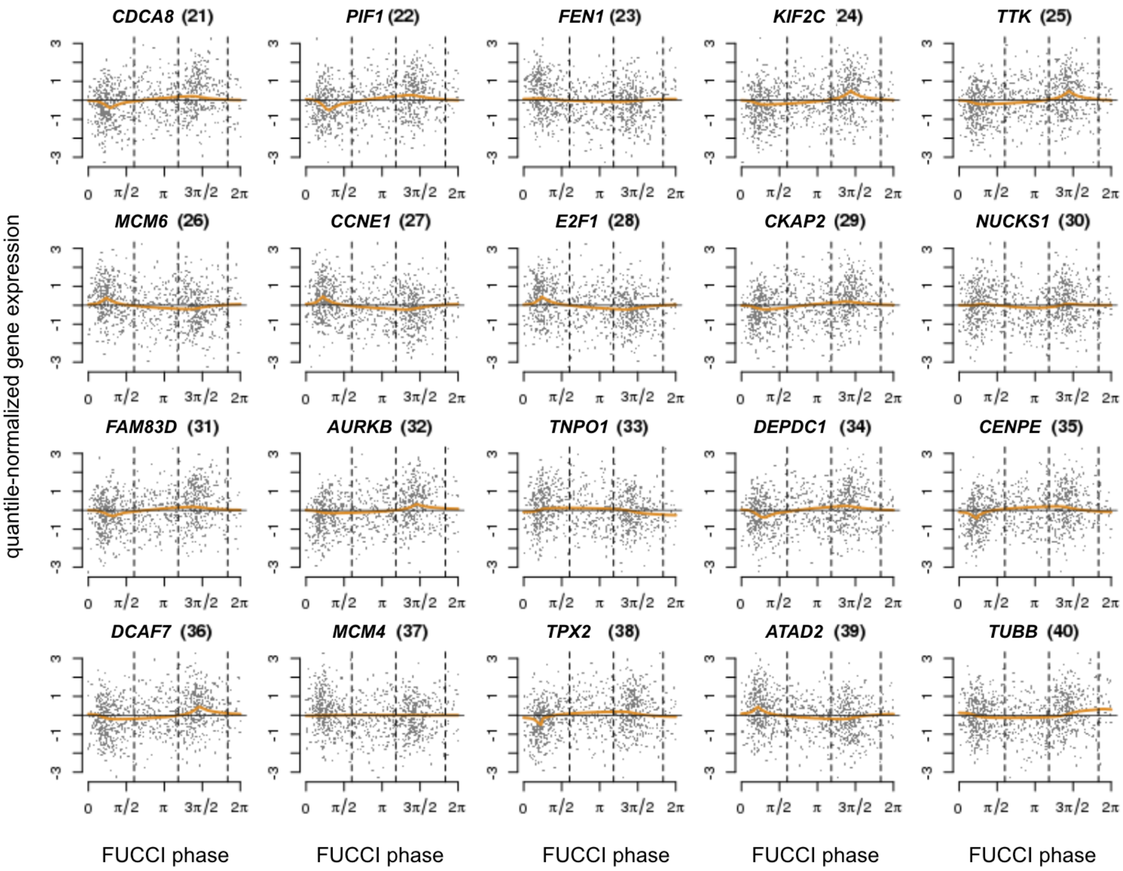


Supplemental Fig. S9B: Cyclic trends of gene expression based on FUCCI phase in 54 genes (a subset of the top 101 cyclic genes also known to be cell cycle genes in Whitfield et al. 2002). For each gene, we ordered 888 single-cell samples by FUCCI phase and applied trend filtering to estimate the cyclic trend of gene expression. The orange line corresponds to the fitted cyclic trend based on FUCCI phase. We ordered the 54 genes by their proportion of variance explained (PVE) by the fitted cyclic trend and showed their order in the figure title (in parenthesis) from large to small PVE. This figure shows genes in top 21 to 40.

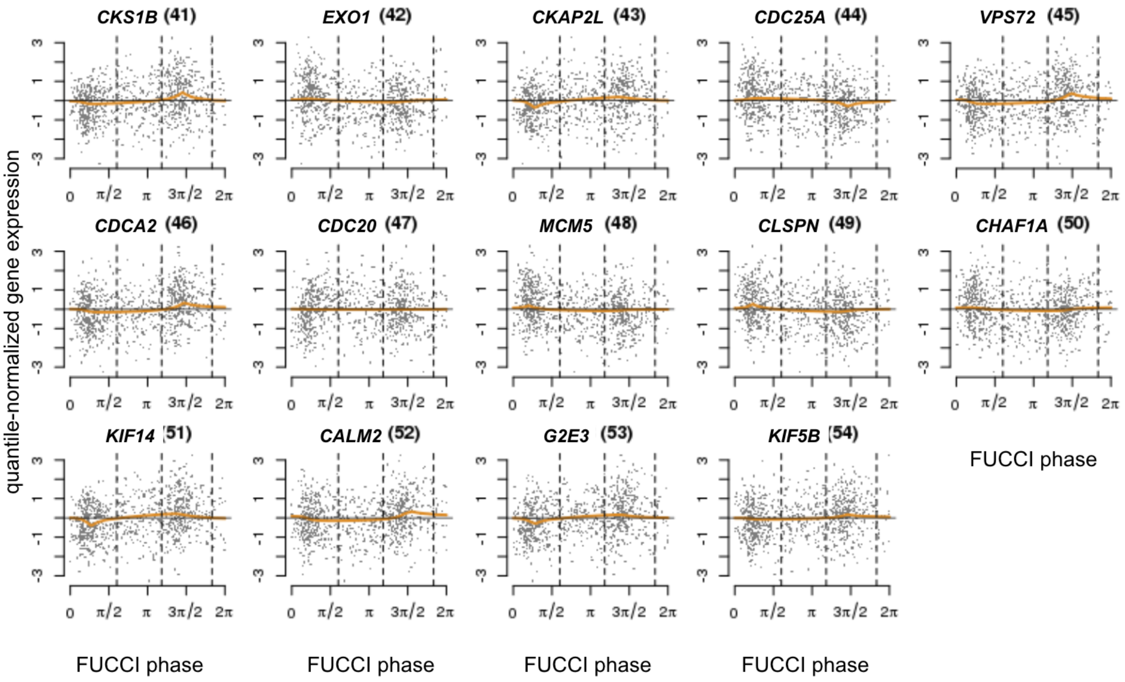


Supplemental Fig. S9C: Cyclic trends of gene expression based on FUCCI phase in 54 genes (a subset of the top 101 cyclic genes also known to be cell cycle genes in Whitfield et al. 2002). For each gene, we ordered 888 single-cell samples by FUCCI phase and applied trend filtering to estimate the cyclic trend of gene expression. The orange line corresponds to the fitted cyclic trend based on FUCCI phase. We ordered the 54 genes by their proportion of variance explained (PVE) by the fitted cyclic trend and showed their order in the figure title (in parenthesis) from large to small PVE. This figure shows genes in top 40 to 54.

#### Supplemental Figure S10

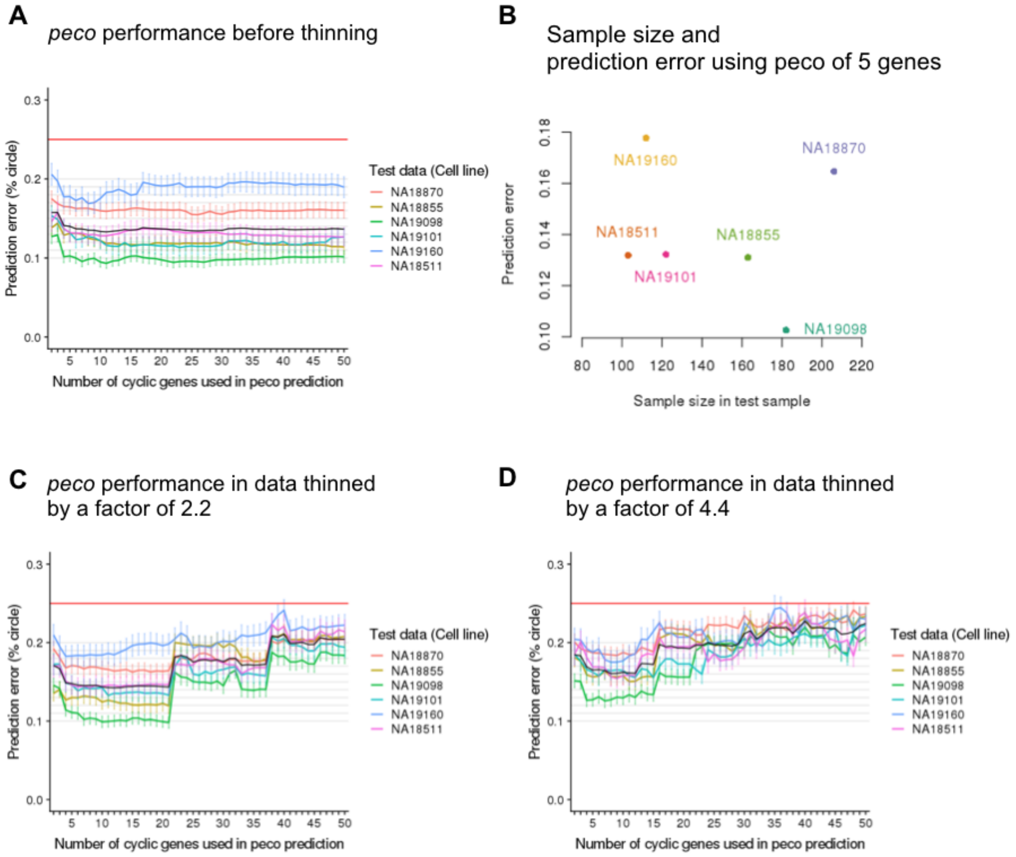


Supplemental Fig. S10: Performance of peco in unthinned and thinned data. We applied six-fold cross-validation. In each fold, we trained our predictor on cells from five individuals and tested its performance on cells from the remaining individual. In panel (A), (C), (D), Y-axis corresponds to prediction error (between 0 to 25%, or $\pi/4$), and X-axis corresponds to the number of top cyclic genes used in the predictor. The six lines correspond to performances in the six folds, specifically average prediction error among cells in the test samples, and error bars correspond to standard errors. (A) The performance of our predictor built between 5 to 50 genes in unthinned data. In (C) and (D), we repeated the analysis in (A) after thinning the test data (total sample molecule count in the un-thinned data was 56,724 $\pm$ 12,762) by a factor of 2.2 (total sample molecule count 25,581 $\pm$ 15,220) and 4.4 (total sample molecule count 13,651$\pm$13,577. (C) and (D) show the performance of our predictor in data thinned by a factor of 2.2 and 4.4, respectively. (B) shows that number of cells was not correlated with prediction error of FUCCI phase using our predictor of 5 genes.

#### Supplemental Figure S11

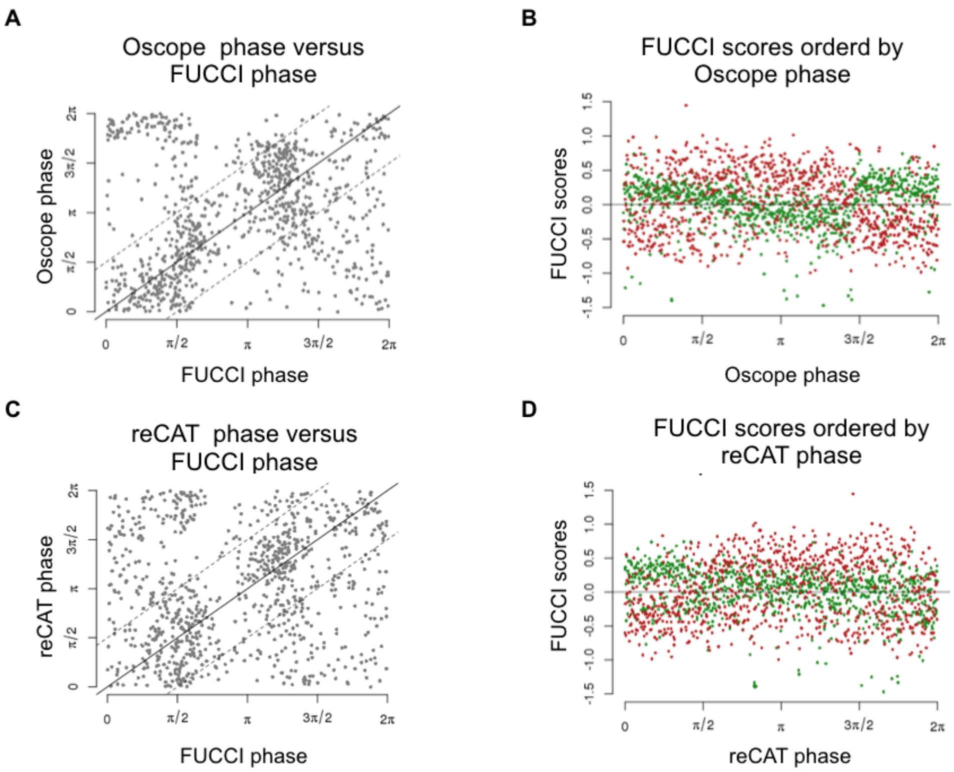


Supplemental Fig. S11: Comparison of FUCCI phase with phase assignment by Oscope and reCAT. In (A) and (C), we plot FUCCI phase (X-axis) against continuous phase (Y-axis) based on Oscope and reCAT, respectively. In (B) and (D), we order EGFP (green dots) and mCherry (red dots) scores by Oscope/Cyclone based phase, respectively.

#### Supplemental Figure S12

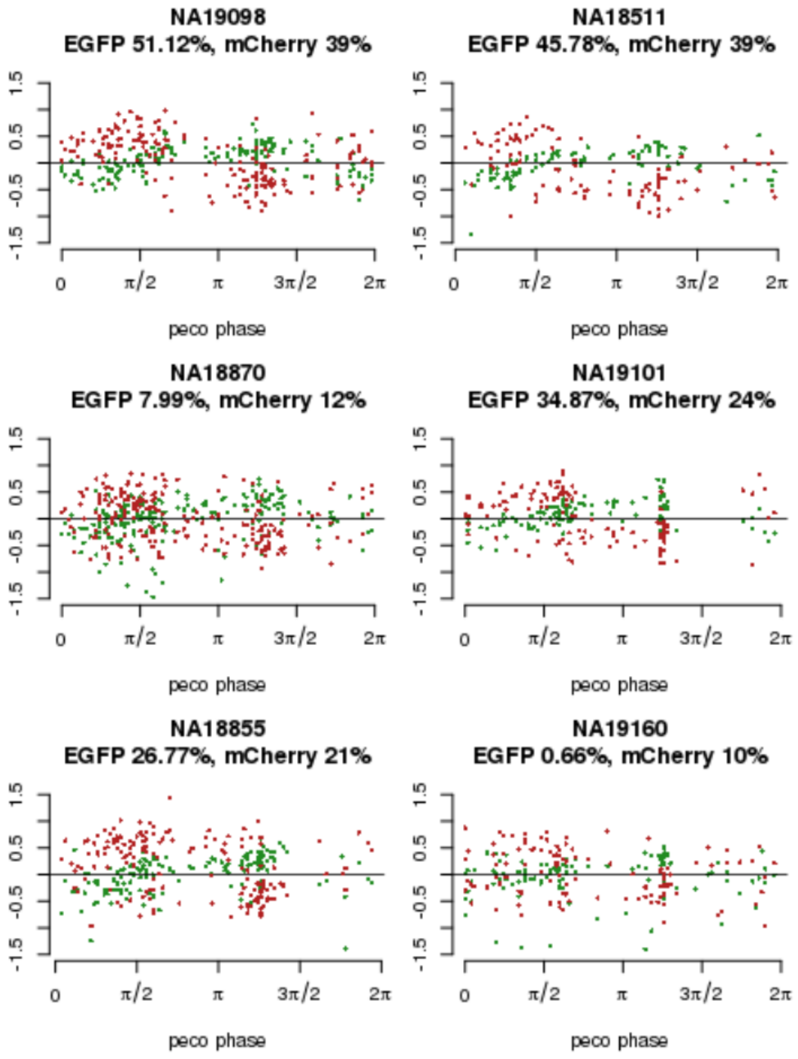


Supplemental Fig. S12: FUCCI scores and inferred phase from peco in individual cell lines. In (A) to (F), we order EGFP (green dots) and mCherry (red dots) scores by inferred phase from peco in each individual cell line (based on our cross-validation results). We also estimated proportion of variance explained by inferred phase from peco in EGFP and mCherry scores (as shown on top of each plot).

#### Supplemental Figure S13

**A**

Seurat discrete phase (rows) vs. PAM based discrete classification (columns)

| | G1 | S | G2M |
|---|---|---|---|
| G1 | 86 | 36 | 68 |
| S | 238 | 44 | 76 |
| G2M | 60 | 92 | 188 |
| % Misclass-ified | 78% | 74% | 43% |

**B**

Cyclone discrete phase (rows) vs. PAM based discrete classification (columns)

| | G1 | S | G2M |
|---|---|---|---|
| G1 | 250 | 76 | 44 |
| S | 84 | 21 | 58 |
| G2M | 50 | 75 | 230 |
| % Misclass-ified | 34% | 88% | 31% |

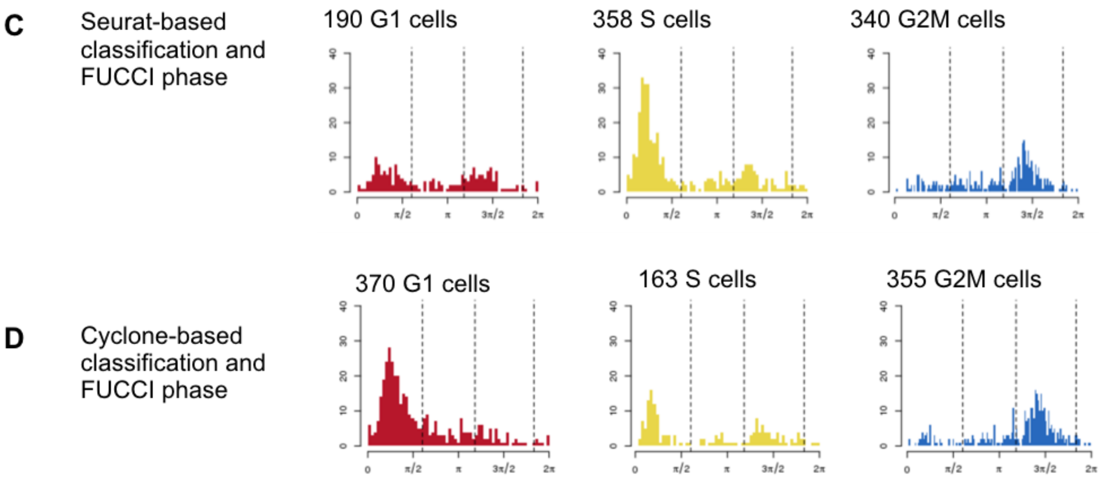


Supplemental Fig. S13: Comparison of FUCCI phase with phase assignment by Seurat and Cyclone. In (A) and (B), classification obtained from PAM is compared with Seurat/Cyclone-based classification. (C) and (D) show the FUCCI phase distribution among single-cell samples in each discrete class by Seurat and Cyclone-based classification, respectively.

#### Supplemental Figure S14

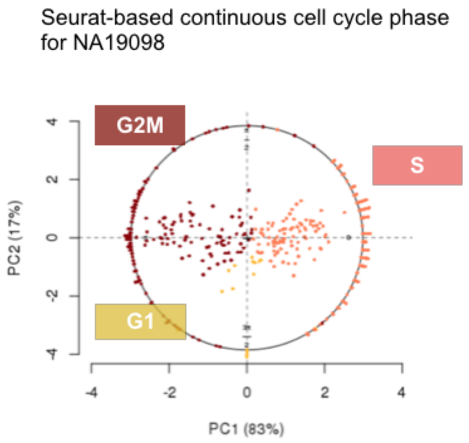


Supplemental Fig. S14: Continuous cell cycle phase assignment based on the two Seurat phase-specific scores for samples from cell line NA19098. Seurat uses the mean expression levels of 43 S-phase marker genes and 54 G2/M phase genes to compute two phase-specific scores for each cell. We applied PCA to transform the two phase-specific scores to PC scores. X- and Y-axis correspond to PC1 and PC2 score. The dots inside the circle correspond to the cells and their PC scores. We transformed these scores to angles on the unit circle (see Methods for details). The colors correspond to Seurat-based G1, S, G2/M phase assignment.

#### Supplemental Figure S15

Comparison of prediction error on our data.

| | peco | Cyclone | Seurat | Oscope | reCAT |
|---|---|---|---|---|---|
| NA18511 | 13.2(1.21) | 15.2(1.27) | 19.6(1.51)* | 20.4(1.39)* | 21.4(1.39)* |
| NA18855 | 13.1(.97) | 18.4(1.02)* | 18.6(.01)* | 15.9(.99)* | 21.5(1.08)* |
| NA18870 | 16.5(.96) | 18.6(.98) | 21.7(.99)* | 22.9(1.03)* | 23.9(1.01)* |
| NA19098 | 10.3(.71) | 14.3(.79)* | 19.4(.96)* | 24.5(1.04)* | 20.9(.96)* |
| NA19101 | 13.2(1.09) | 15.1(1.03) | 20.2(1.28)* | 18.8(1.17)* | 23.3(1.34)* |
| NA19160 | 17.8(1.44) | 22.9(1.41)* | 21.9(1.35)* | 24.1(1.35)* | 24.3(1.37)* |

Note. Reported in each cell are percent of a unit circle with standard errors in parentheses

*We compared prediction error between methods and peco using Wilcoxon test and determined signficance at P-value < .05

Supplemental Fig. S15: Comparison of prediction error on our data in cross-validation. We performed Wilcoxon test to compare prediction error between peco and each method in each test data set (samples from an individual cell line). We report prediction error as percentage of the unit circle, along with standard error associated with mean prediction error in parentheses.

#### Supplemental Figure S16

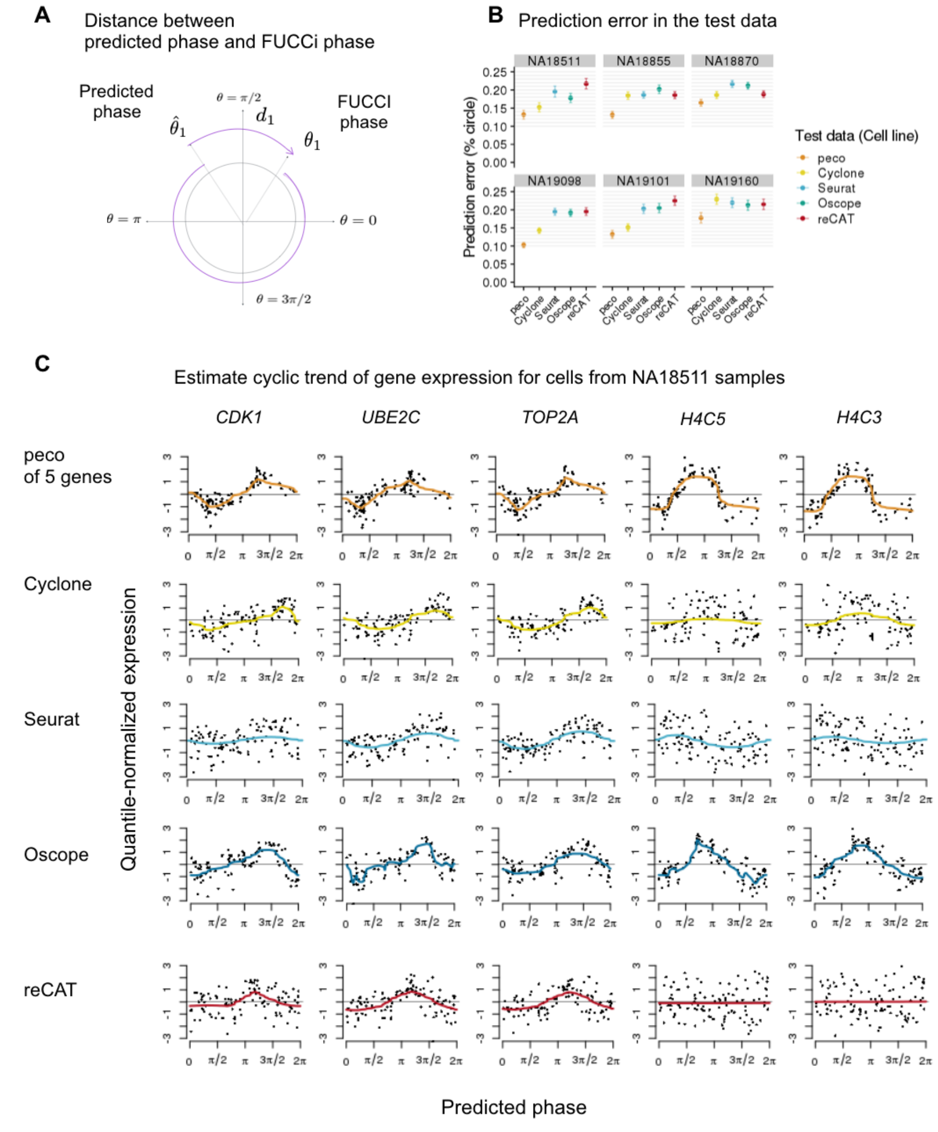


Supplemental Fig. S16: Comparison with existing methods on our data. (A) We defined prediction error as the distance between predicted phase and FUCCI phase (as a percentage of a unit circle). (B) We applied six-fold cross-validation to test the performance of predictors. The six panels correspond to performances in the six folds. Each panel compares the mean prediction error of FUCCI phase in the test data (error bars correspond to standard error) using our method and existing tools (Seurat, Cyclone, reCAT and Oscope). (C) Estimated cyclic trend of top 5 cyclic genes in samples from cell line NA18511. Rows correspond to the results of the five methods. Specifically, we ordered the samples according to the predicted phase of each method and used trendfilter to estimate cyclic trend of gene expression. The colored line corresponds to the estimated cyclic expression level along the predicted phase.

#### Supplemental Figure S17A-E

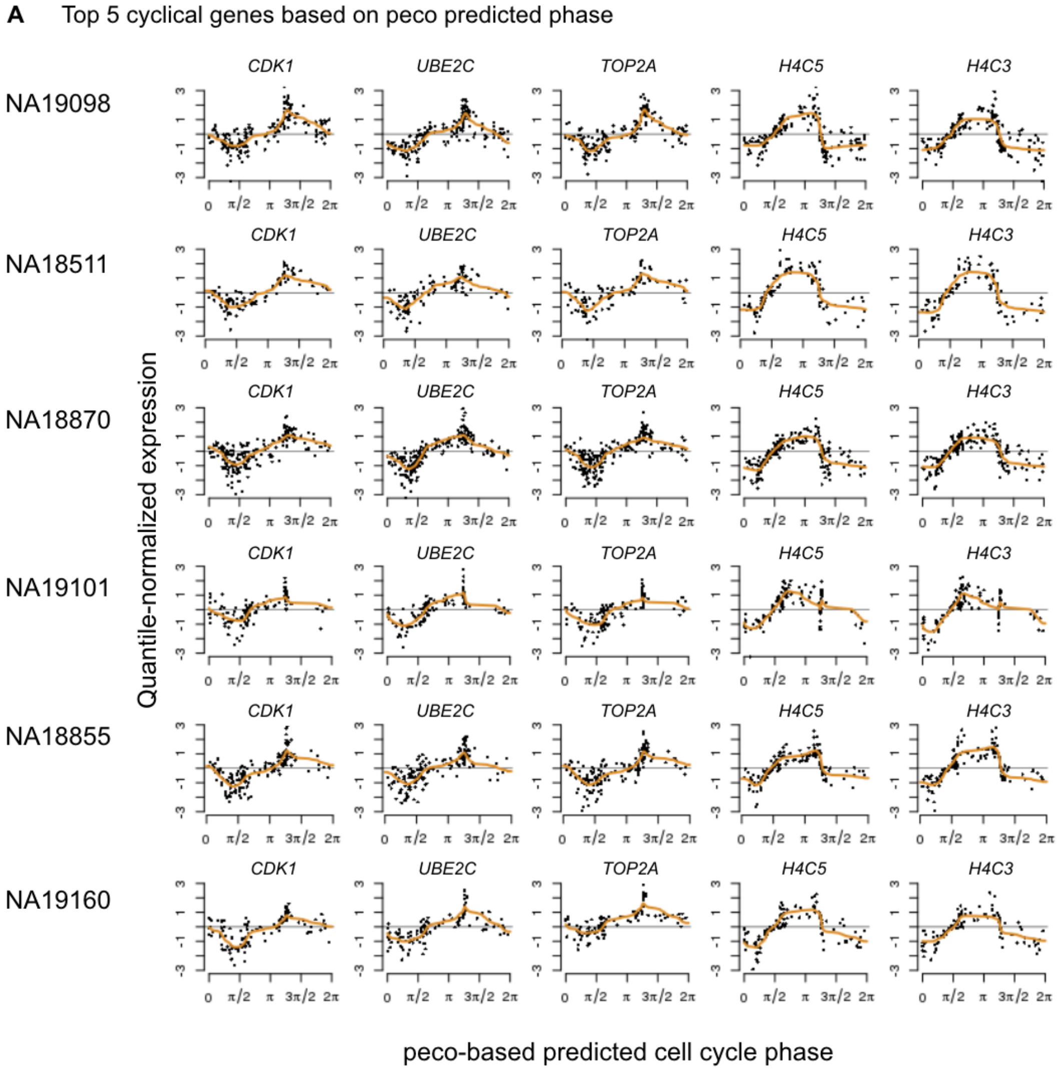


Supplemental Fig. S17A: peco prediction results for the six cell lines, using the simple predictor of 5 genes (*CDK1*, *UBE2C*, *TOP2A*, *H4C5*, *H4C3*). Rows correspond to results for individual cell lines. For example, for cell line NA19098, we ordered samples by FUCCI phase and used *trendfilter* to estimate the cyclic trend of gene expression in the top 5 cyclic genes. The colored line represents the predicted cyclic trend.

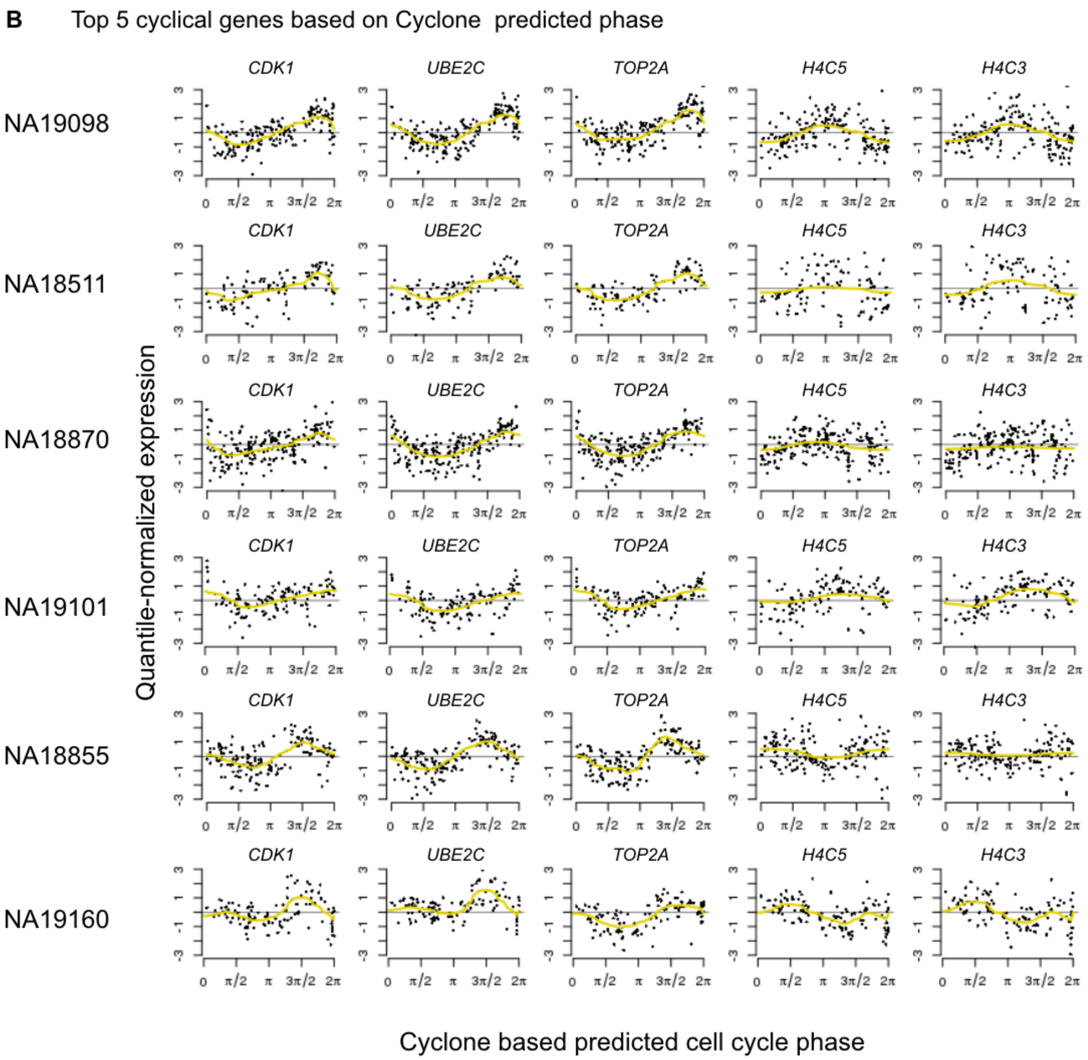


Supplemental Fig. S17B: Cyclone prediction results for the six cell lines. As described in the Results, we transform the three phase-specific Cyclone scores to angles on the unit cirlce, using the same approach for deriving FUCCI phase from FUCCI scores. Rows correspond to results for individual cell lines. For example, for cell line NA19098, we ordered samples by Cyclone-based predicted phase and used *trendfilter* to estimate the cyclic trend of gene expression in the top 5 cyclic genes.

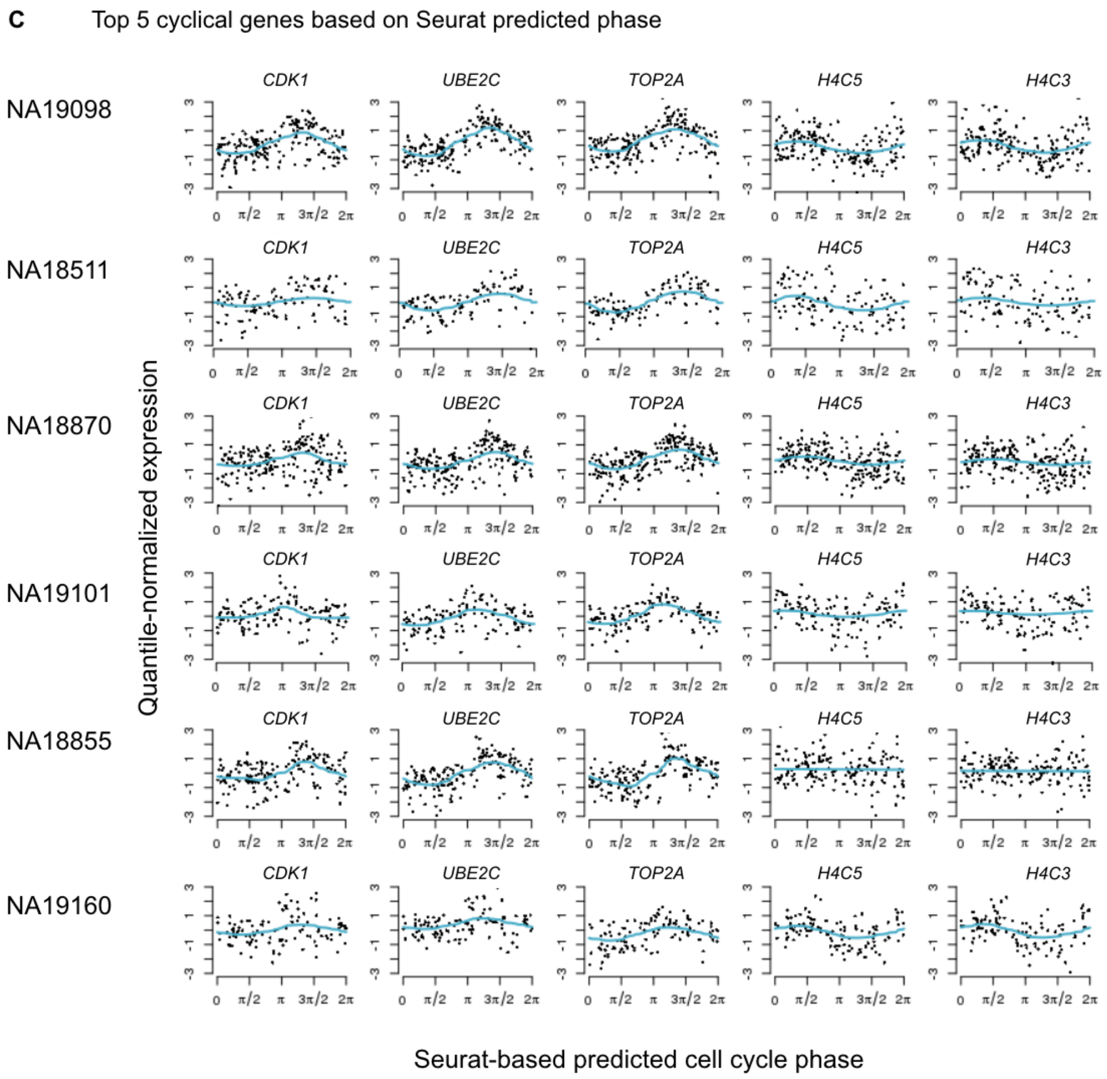


Supplemental Fig. S17C: Seurat prediction results for the six cell lines. As described in the Results, we transform the two phase-specific Seurat scores to angles on the unit cirlce, using the same approach for deriving FUCCI phase from FUCCI scores. Rows correspond to results for individual cell lines. For example, for cell line NA19098, we ordered samples by Seurat-based predicted phase and used *trendfilter* to estimate the cyclic trend of gene expression in the top 5 cyclic genes.

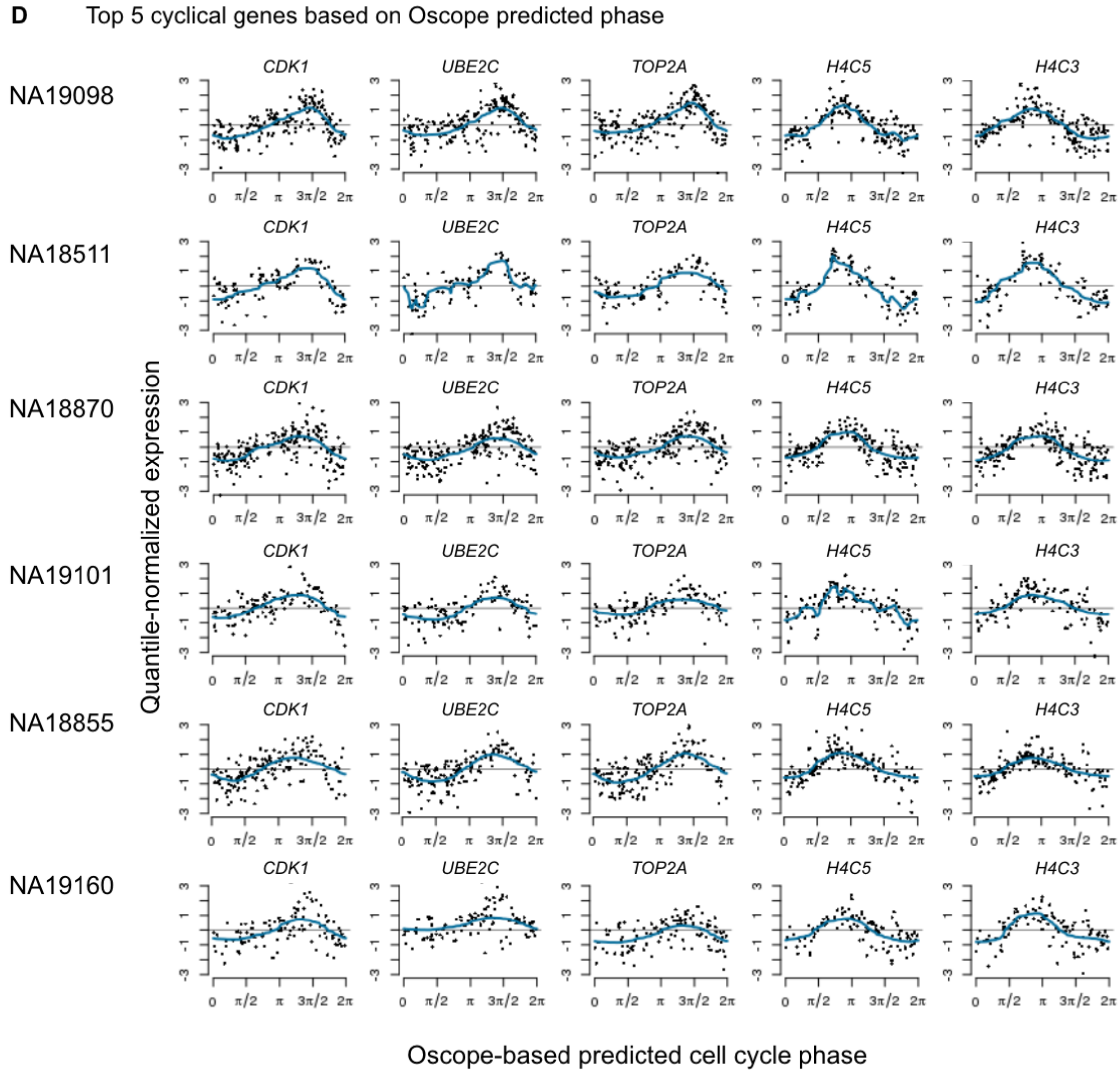


Supplemental Fig. S17D: Oscope prediction results for the six cell lines. As described in the Results, we estimated the cyclic ordering of cells across the 888 high-quality single-cell samples in the data. We then assigned each cell an angle on the unit circle based on the ordering per individual cell line. Rows correspond to results for individual cell lines. For example, for cell line NA19098, we ordered samples by Oscope predicted phase and used *trendfilter* to estimate the cyclic trend of gene expression in the top 5 cyclic genes.

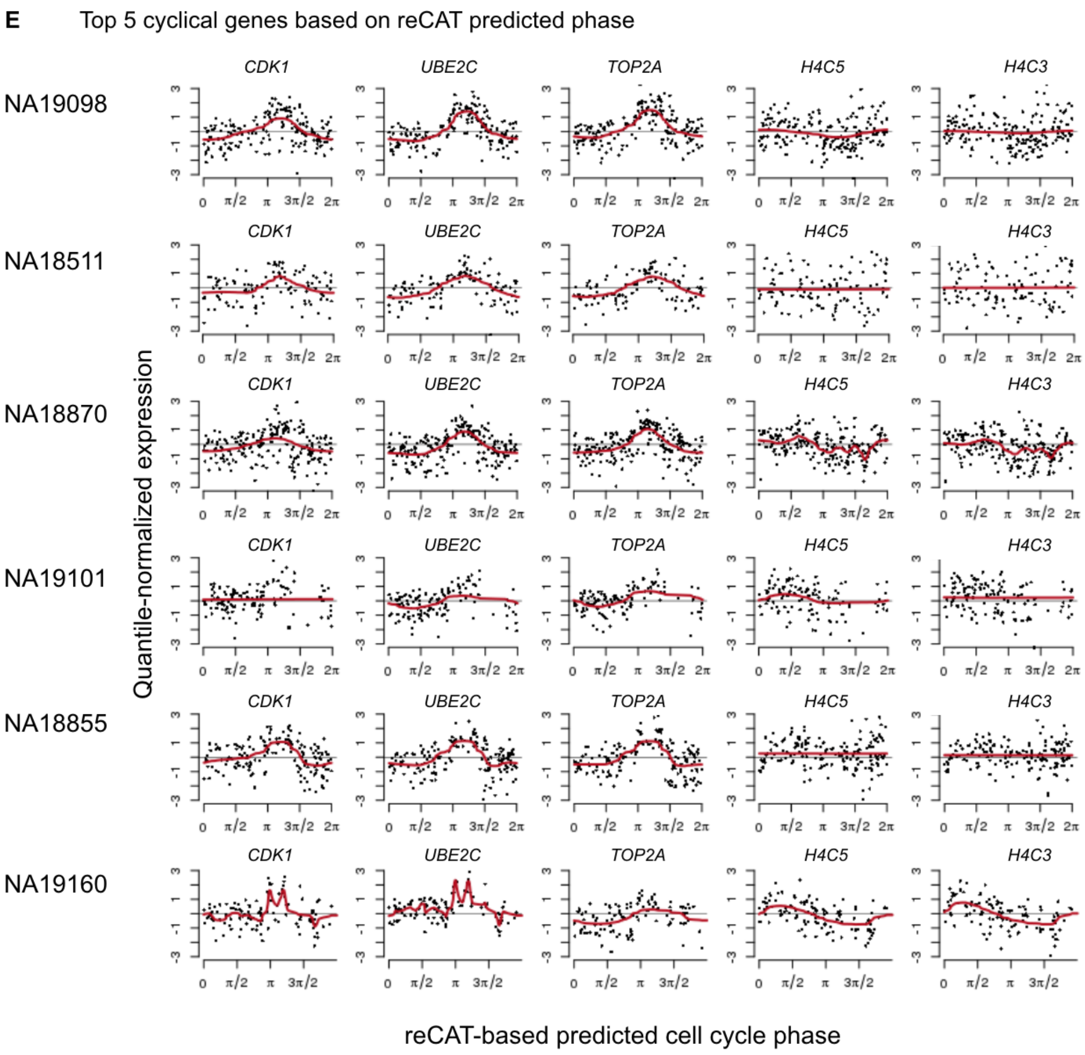


Supplemental Fig. S17E: reCAT prediction results for the six cell lines. As described in the Results, we estimated the cyclic ordering of cells across the 888 high-quality single-cell samples in the data. We then assigned each cell an angle on the unit circle based on the ordering per individual cell line. Rows correspond to results for individual cell lines. For example, for cell line NA19098, we ordered samples by reCAT predicted phase and used *trendfilter* to estimate the cyclic trend of gene expression in the top 5 cyclic genes.

#### Supplemental Figure 18

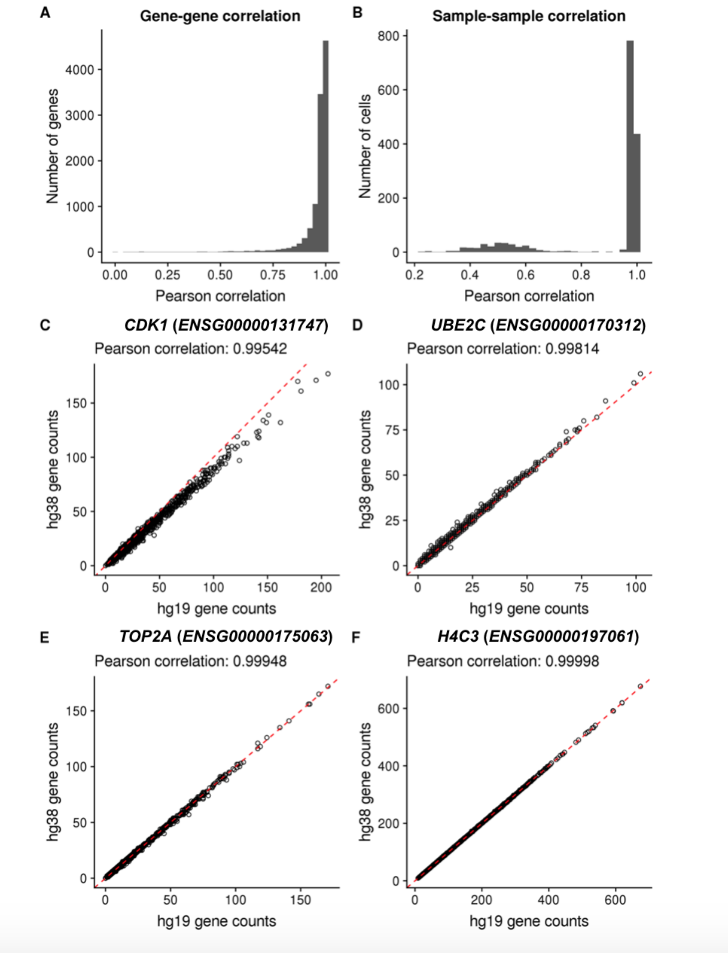


Supplemental Fig. S18: Comparison of global gene expression profiles obtain from genome build hg19 vs hg38 (A) Histogram of Pearson correlation for each gene comparing its counts across single cells when mapping to genome build hg19 vs hg38. The median correlation was 0.983919, and 1,070 genes (9.8%) had a correlation less than 0.9. (B) Histogram of Pearson correlation for each single-cell sample comparing its gene counts when mapped to genome build hg19 vs hg38. The median correlation was 0.9849, and 306 single cells (19.9%) had a correlation less than 0.9. (C) A scatterplot of cell cycle gene *CDK1* (ENSG00000131747) gene counts for hg19 (x-axis) vs hg38 (y-axis). The Pearson correlation across single cells was 0.99542. (D) A scatterplot of cell cycle gene *UBE2C* (ENSG00000170312) gene counts for hg19 (x-axis) versus hg38 (y-axis). The Pearson correlation across single cells was 0.99814. (E) A scatterplot of cell cycle gene *TOP2A* (ENSG00000175063) gene counts for hg19 (x-axis) versus hg38 (y-axis). The Pearson correlation across single cells was 0.99948. (F) A scatterplot of cell cycle gene *H4C5* (ENSG00000197061) gene counts for hg19 (x-axis) versus hg38 (y-axis). The Pearson correlation across single cells was 0.99998. The dashed red line represents the 1-1 line. All plots used data for the 10,297 protein-coding genes that were shared between genome builds hg19 and hg38. Note that cell cycle gene *H4C3* (ENSG0000198518) was deprecated in the annotation for genome build hg38.

#### Supplemental Figure S19

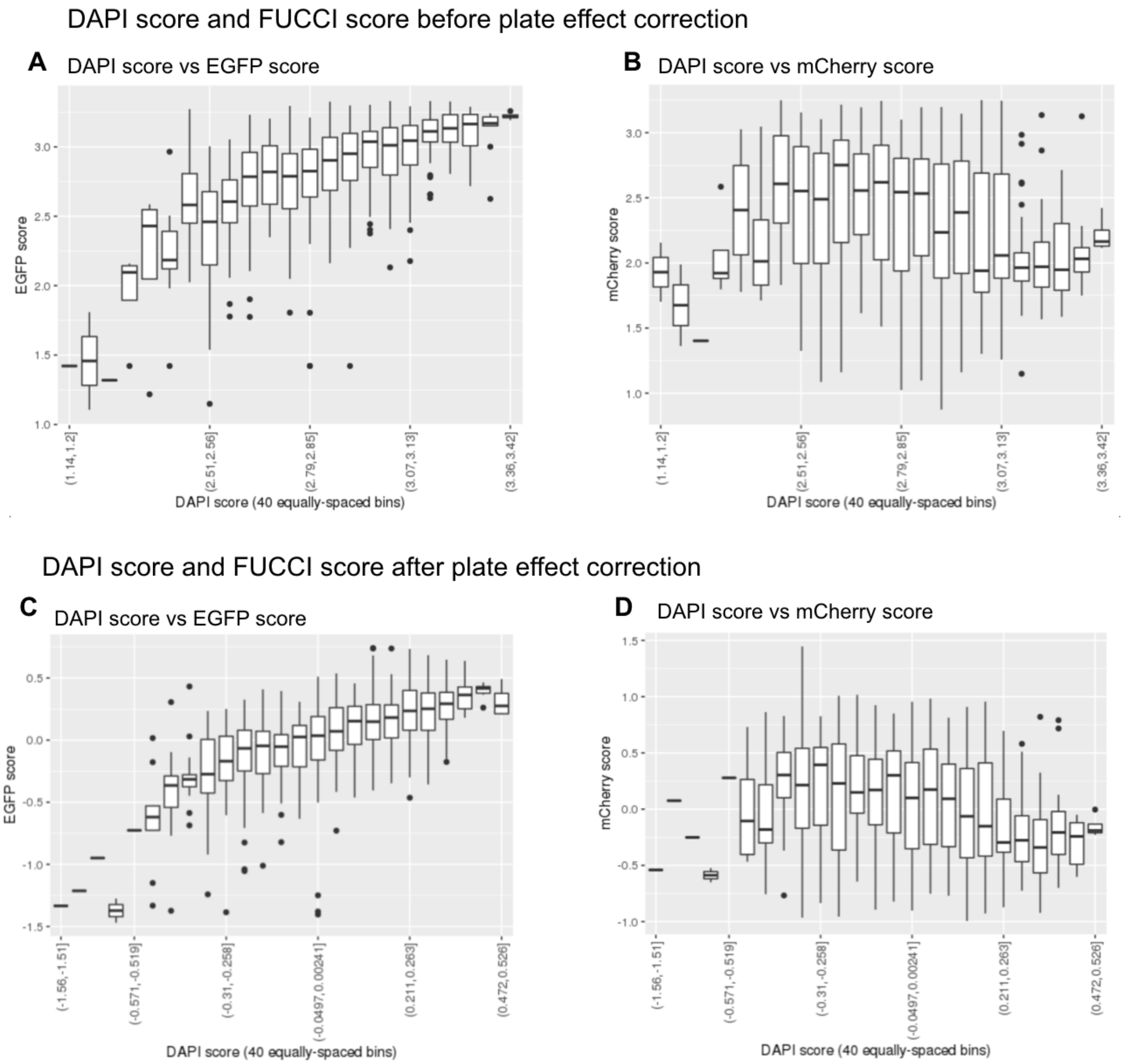


Supplemental Fig. S19: FUCCI score and DAPI score of the 888 single-cell samples before and after C1 plate effect correction. (A) and (B) show the relationship between EGFP and mCherry scores with DAPI score before plate effects correction. (C) and (D) show the relationship between EGFP and mCherry scores with DAPI score after plate effects correction. The 40 boxplots in each plot correspond to the distribution of EGFP/mCherry scores in 40 equally-spaced intervals of DAPI scores.

#### Supplemental Figure 20

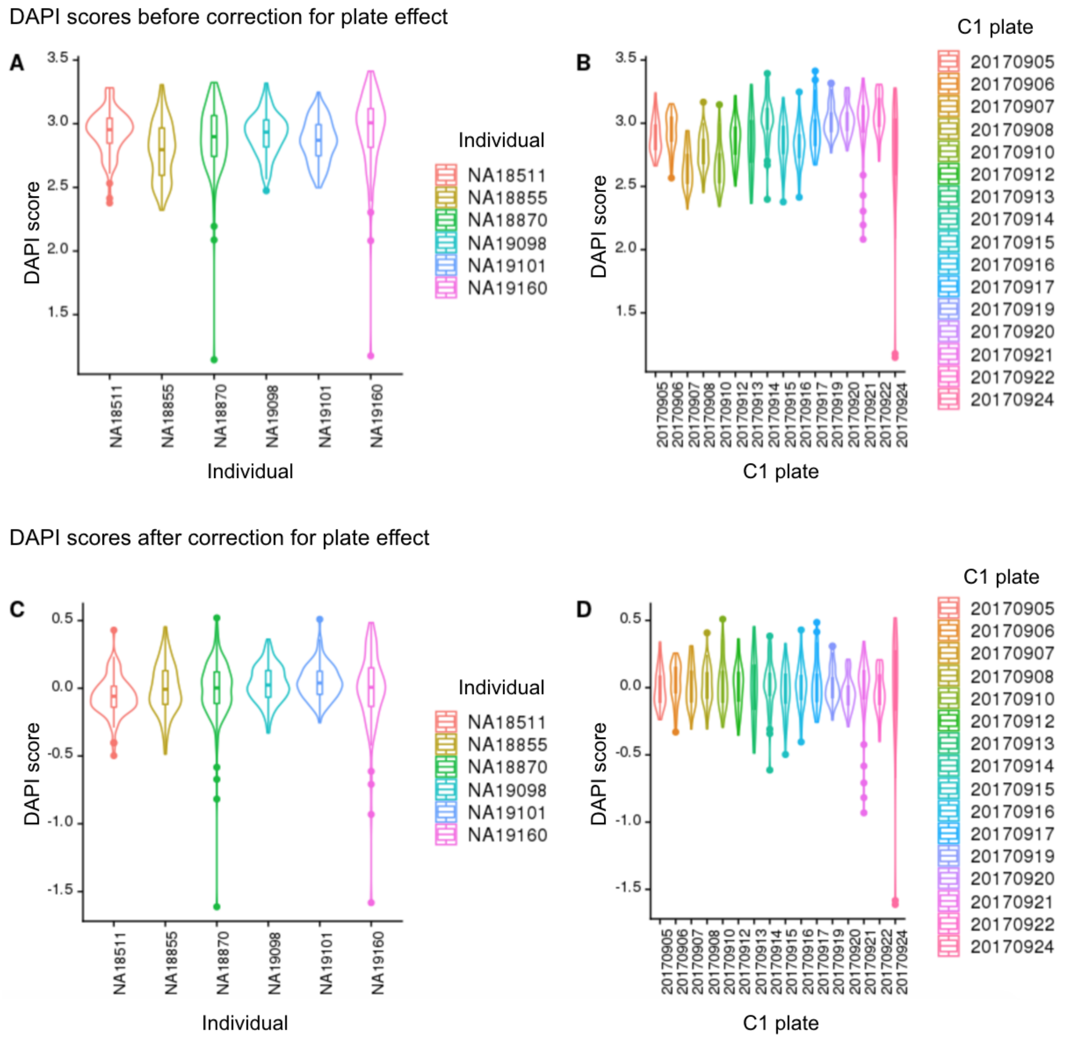


Supplemental Fig. S20: DAPI scores variation before and after correcting for C1 plate effect. To adjust for C1 plate effect, we fitted an analysis of variance model (score $\sim$ plate + individual) and subtracted the marginal means of plate effect from and DAPI scores controlling for individual effect. (A) and (B) show DAPI scores variation before correcting for C1 plate effect between individual cell lines. (C) and (D) show DAPI scores variation after correcting for C1 plate effect between C1 plates.

### Supplemental Figure 21

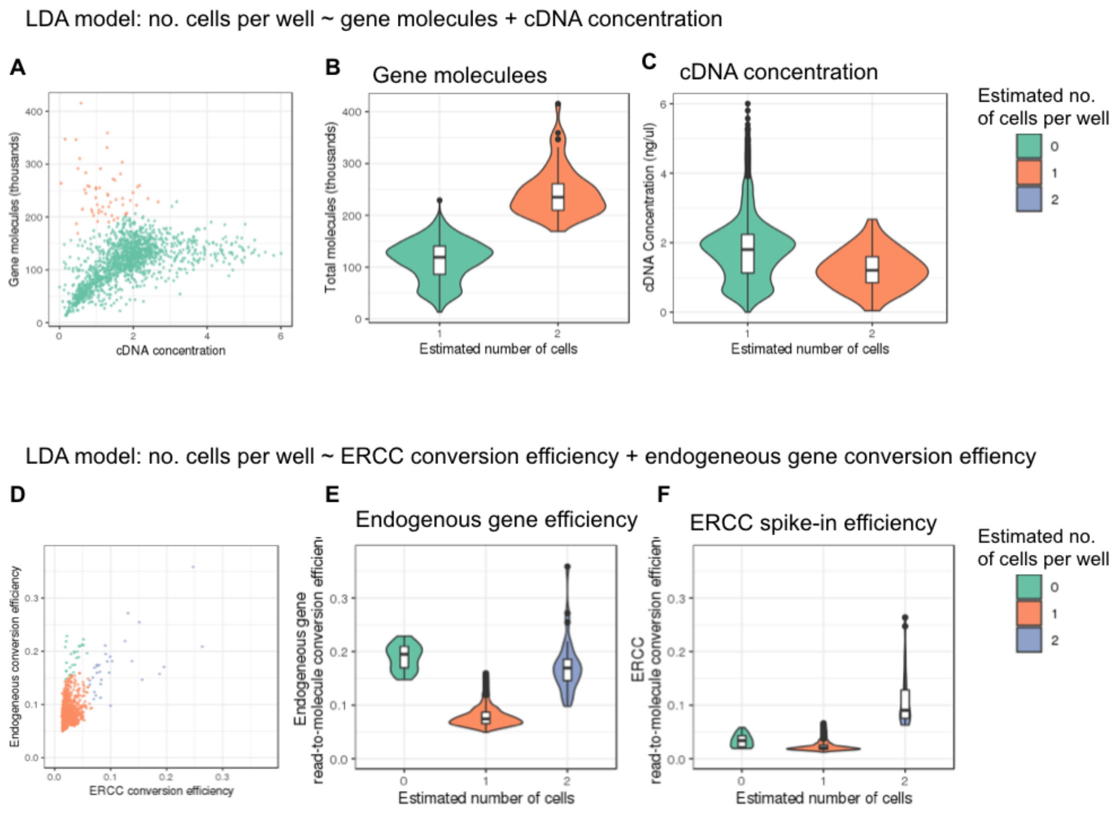


Supplemental Fig. S21: As a part of the quality control analysis, we used LDA to determine the number of cells captured in each well. Specifically, we fitted two LDA models: 1) number of cells $\sim$ gene molecule count + cDNA amplicons concentration, and 2) number of cells $\sim$ ERCC spike-in control read-to-molecule conversion efficiency + endogenous gene read-to-molecule conversion efficiency. We determined the observed number of cells captured in each C1 well based on DAPI staining results. (A) plots the relationship between cDNA concentration and gene molecule, with sample points colored by the predicted number of cells per well in LDA analysis. (B) and (C) show the distribution of gene molecule and cDNA concentration in wells predicted to have 1 cell and wells predicted to have 2 cells. (D) plots the relationship between the read-to-molecule conversion efficiency of ERCC controls and endogenous genes, with sample points colored by the predicted number of cells per well in LDA analysis. (E) and (F) show the distribution of ERCC and endogenous gene read-to-molecule conversion efficiency in wells predicted to have 0, 1, and 2 cells in LDA analysis.
